# Identification of novel myeloid-derived cell states with implication in cancer outcome

**DOI:** 10.1101/2023.01.04.522727

**Authors:** Gabriela Rapozo Guimarães, Giovanna Resk Maklouf, Cristiane Esteves Teixeira, Leandro de Oliveira Santos, Nayara Gusmão Tessarollo, Marco Antônio Pretti, Nayara Evelin Toledo, Jéssica Gonçalves Vieira da Cruz, Marcelo Falchetti, Mylla M. Dimas, Alessandra Freitas Serain, Fabiane Carvalho de Macedo, Fabiana Resende Rodrigues, Nina Carrossini Bastos, Jesse Lopes da Silva, Edroaldo Lummertz da Rocha, Cláudia Bessa Pereira Chaves, Andreia Cristina de Melo, Pedro Manoel Mendes Moraes-Vieira, Marcelo A. Mori, Mariana Boroni

**Author notes:** These authors share the co-first authorship (Gabriela Rapozo Guimarães, Giovanna Resk Maklouf). Corresponding author: Mariana Boroni.

## Abstract

Tumor-associated myeloid-derived cells (MDCs) significantly impact cancer prognosis and treatment response due to their remarkable plasticity and tumorigenic behaviors. We integrated single-cell RNA-Sequencing datasets from seven different cancers, resulting in a comprehensive collection of 29 MDC subpopulations in the tumor microenvironment (TME). Distinguishing resident-tissue from monocyte-derived macrophages, we discovered a resident-tissue-like subpopulation within monocyte-derived macrophages. Additionally, hypoxia-driven macrophages emerged as a prominent TME component. Deconvolution of these profiles revealed five subpopulations as independent prognostic markers across various cancer types. Validation in large cohorts confirmed the FOLR2-expressing macrophage association with poor clinical outcomes in ovarian and triple-negative breast cancer. Moreover, the marker TREM2, commonly used to define immunosuppressive tumor-associated macrophages, cannot solely predict cancer prognosis, as different polarization states of macrophages express this marker in a context-dependent manner. This comprehensive MDC atlas offers valuable insights and a foundation for novel analyses, advancing strategies for treating solid cancers.

## Introduction

The tumor microenvironment (TME) represents a dynamic network consisting of diverse cell types and acellular components that intricately interact with malignant cells. The understanding of immune cells within the TME, particularly lymphocytes, has paved the way for the development of immunotherapies capable of inducing long-lasting responses across a wide spectrum of cancers^1^. However, the current immune-checkpoint inhibitor therapies only confer benefits to a subset of individuals^2^. Among the crucial players in the TME are myeloid-derived cells (MDCs), comprising a heterogeneous array of populations including monocytes, macrophages, conventional dendritic cells (cDC), and polymorphonuclear granulocytes. These MDCs exhibit remarkable plasticity, allowing them to adopt various cellular states and perform a wide range of functions when exposed to different niches^3^. Intriguingly, while MDCs possess the capacity to enhance cancer cell phagocytosis and induce cytotoxic tumor death when properly activated^4,5^, they can also foster tumor growth by facilitating and sustaining cancer hallmarks^6^.

Regarding ontogeny, most MDCs originate from bone marrow progenitor cells and subsequently migrate into the TME. Macrophages, however, can derive from two primary sources: 1) erythro-myeloid progenitors in the yolk sac and fetal liver before birth, which exhibit enriched expression of *FOLR2*, *PLTP*, and *LYVE1* markers^7,8^; and 2) circulating monocytes originating from the bone marrow and recruited into the tumor^9^. Recruited macrophages have a transient lifespan and necessitate constant replenishment from circulating monocytes^10^. Their ontogeny can be indicated by the expression of monocyte-related markers such as *FN1*, *SELL*, and *VCAN^11^*.

Macrophages, regardless of their origin, can exhibit distinct phenotypes. Traditionally, they have been classified into two main polarization states: classically activated M1, characterized by a proinflammatory phenotype, and alternatively activated M2, associated with tissue remodeling and/or anti-inflammatory properties^9^. In the context of the TME, M2 macrophages are commonly referred to as tumor-associated macrophages (TAMs), and their presence has been correlated with poor prognosis in several tumor types^12–14^. However, it is important to note that not all TAMs display a clear M2-like phenotype, as they often express markers associated with both activation states^15,16^. This underscores the need to move beyond the simplistic M1/M2 dichotomy and delineate distinct TAM states.

TAMs play a multifaceted role within the TME, influencing tumor growth, epithelial-mesenchymal plasticity, extracellular matrix remodeling, cell invasion, and angiogenesis, subsequently impacting tumor progression, metastasis, and therapy resistance. Furthermore, the interaction between TAMs and malignant cells can facilitate immune evasion through TME shaping^17^. Given the pivotal role of TAMs and other MDCs in promoting tumor growth and metastasis, they have emerged as promising targets for cancer therapy. Numerous strategies aimed at depleting or modulating the functional/phenotypic reprogramming, infiltration, or activation of TAMs are being explored^18^. However, the lack of precise markers to distinguish between MDC subpopulations and states poses a challenge to the effectiveness of these treatments. Therefore, a thorough characterization of MDC populations within the TME is urgently required to overcome this limitation.

While recent studies have utilized high-resolution technologies such as single-cell RNA sequencing (scRNA-Seq) and spatial transcriptomics to investigate the immune landscape of different cancer types, a comprehensive pan-cancer integrated analysis encompassing MDCs in the TME has been lacking. To address this critical gap, we employed data from three distinct scRNA-Seq technologies, integrating them to construct the most comprehensive pan-cancer catalog of MDC subpopulations to date. Our analysis incorporated data from seven solid tumor types, enabling a thorough characterization of tumor-associated cell types. Consequently, we identified novel MDC populations/states within these tumors and uncovered abnormally expanded MDC subpopulations associated with a poor prognosis across various tumor origins. These findings pave the way for the development of effective targeted immunotherapy strategies aimed at specific MDC subpopulations.

## Results

### A thorough integration strategy recovers an in-depth pan-cancer repertoire of MDCs in the tumor microenvironment

To define the landscape of cell subtypes in the TME, we integrated scRNA-Seq data comprising 392,204 high-quality cells from 13 public datasets (Supplementary Table 1 and Supplementary Fig. 1a). The integrated datasets comprised three different technologies (10x Genomics, InDrop, and Smart-Seq2) (Supplementary Fig. 1b) and samples from different anatomical sites of healthy donors and patients diagnosed with different cancer types, totaling 142 individuals (Supplementary Table 2). Our pipeline allowed for the integration of a heterogeneous group of datasets, and even though we applied stringent quality control, it retained crucial biological variation while efficiently removing typical contaminants such as doublets and ambient RNA, as well as noise due to batch effect (Fig. 1a). The final integrated dataset encompassed seven tumors, including breast, colorectal, liver, lung, ovary, skin, and uveal melanomas, as well as their adjacent tissue counterparts, metastatic samples, blood samples, and adjacent lymph nodes (Fig. 1b).

**Figure 1.**
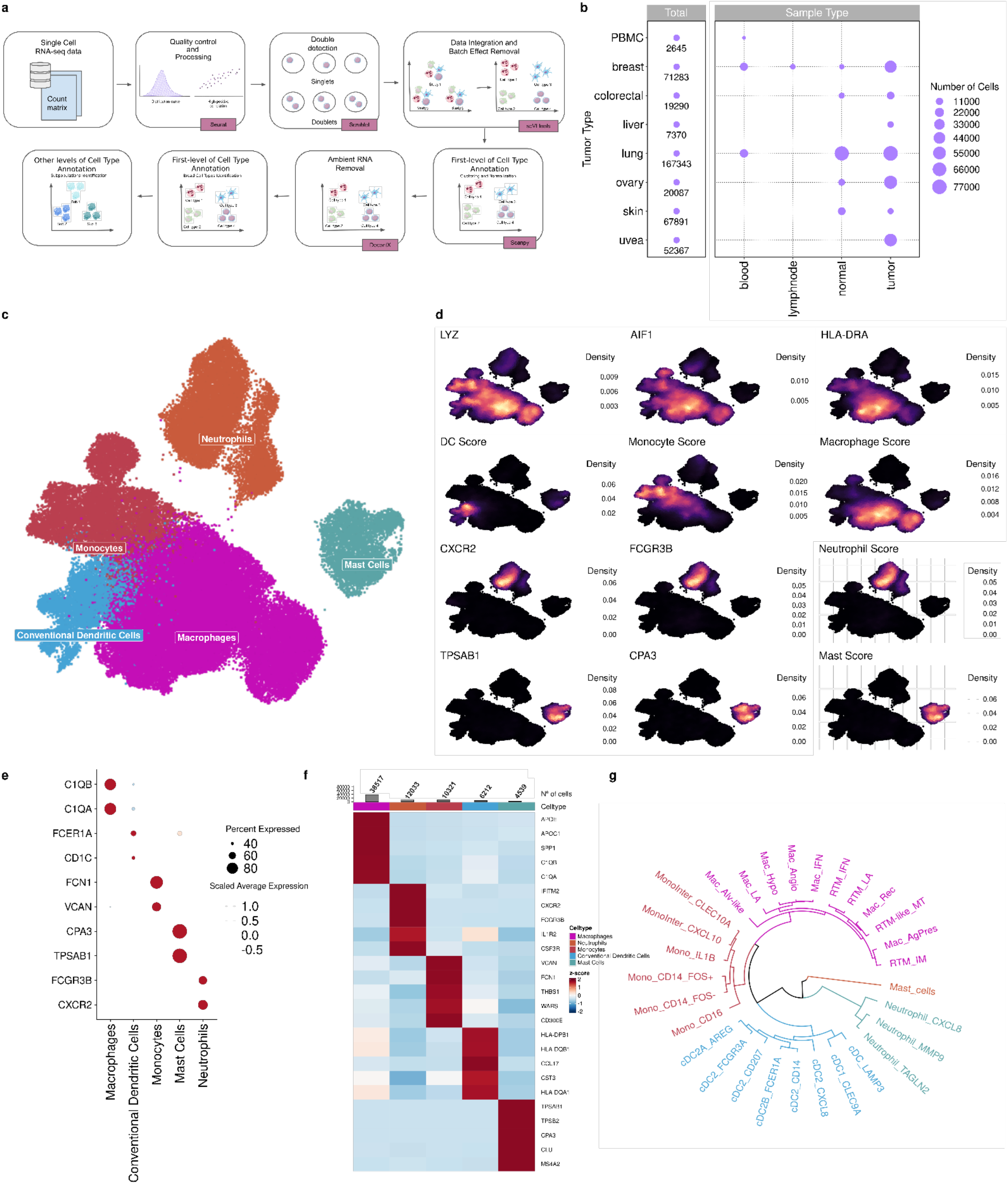
Multi-tissue single-cell atlas of tumor-infiltrating myeloid cells reveals novel states and gene programs. **a** Schematic overview of our integration approach. Data from 13 datasets were pre- processed for quality control to remove either empty droplets or doublets and then integrated using technical co-variations to remove batch effects. Broad cell types were identified and contamination by ambient RNA was removed. MDCs were then sub-clustered and annotated based on gene markers to identify cell states. **b** Dotplot showing the distribution of sample types across the eight tissues analyzed in this study. Dot size indicates the number of cells by sample type. **c** Uniform Manifold Approximation and Projection (UMAP) of 51,687 MDCs, color-coded according to cell types. **d** Density plots highlighting the expression of gene markers and scores (DC - *CD1C*, *FCER1A*, *LAMP3*, and *CLEC9A*); Monocyte - *VCAN* and *FCN1*; Macrophage - *C1QA* and *CD68*; Neutrophil - *CXCR2* and *FCGR3B*; Mast - *TPSAB1* and *CPA3*) for each cell type. **e** Dotplot showing the expression of specific markers by each MDCs subpopulation. The dot size indicates the percent of expressing cells, and the dot color represents the scaled average expression. **f** Heatmap showing expression levels of specific markers from each MDCs subpopulation. The color scale represents the scaled expression of each gene. Cells type proportions related to total MDCs were added to the top. **g** Dendrogram illustrates the relationship among the MDCs populations.

Using our approach, we identified homogeneous cell groups distributed in clusters (Supplementary Fig. 1c). The clusters were annotated based on canonical gene markers (Supplementary Table 3) of major cell types and differentially expressed genes (DEGs) (Supplementary Fig. 1d, e), yielding 10 broadly recognized cell types. Low-quality cells were annotated as non-identifiable (Supplementary Fig. 1c). MDCs were the second-largest group of cells in the TME (Supplementary Fig. 1d), and their proportion in tumor samples was 1.74 times greater than that in normal samples (Supplementary Fig. 1f). The percentage of these cells varied across sample types from 1.7% to 25%, except for metastatic lung samples, where over 60% of TME cells matched mononuclear phagocytes (Supplementary Fig. 1g).

Among the MDC subpopulations (Fig. 1c-e), mononuclear phagocytes (n = 51,687), positive for *LYZ*, *AIF1,* and *HLA-DRA,* were the majority. Other populations, such as neutrophils, marked by *CXCR2* and *FCGR3B*, and mast cells, identified by *TPSAB1* and *CPA3* expression, were also largely detected. Megakaryocytes were the smallest group among MDCs (n = 185) and were not further investigated in this study (Supplementary Fig. 1d).

To characterize the MDC subpopulations, we conducted unsupervised clustering and identified five major lineages (Fig. 1e, f): mast cells (n = 4,539), neutrophils (n = 12,033), cDCs (n = 6,212), monocytes (n = 10,321), and macrophages (n = 38,517). Further separation of the MDCs resulted in 29 distinct subpopulations (Fig. 1g). To avoid subjective and arbitrary definitions of cell clusters, we employed entropy-based statistics to accurately quantify the purity of each cluster. Our approach improved the ability to detect and define robust signatures, ranging from common cell states to less frequent ones, performing better than previous strategies that evaluated small and isolated datasets. The purity of each cluster was displayed with scores above 0.84 of ROGUE values (Supplementary Fig. 1h, i). Our findings provide new insights into the MDC subpopulations and contribute to a better characterization of their distribution in the TME.

### MDCs are heterogeneous and phenotypically diverse across tumor samples

Mast cells, one of the less frequent MDC subpopulations, were identified as a single state and were found to be equally distributed in tumor and normal samples (Supplementary Fig. 1f). Neutrophils were further subdivided into three functionally distinct subpopulations (Supplementary Fig. 2a–d): the Neutrophil_TAGLN2 population (n = 6,570) expressed numerous genes involved in the interferon signaling pathway, reflecting the probable pro-inflammatory role of these cells; the Neutrophil_MMP9 population (n = 4,024) was enriched in pathways related to signaling processes by VEGF; and the Neutrophil_CXCL8 population (n = 1,439) was linked to anti-inflammatory pathways such as IL-4 and -13 signaling (Supplementary Fig. 2e and Supplementary Table 4). This last population was found to be proportionally more prevalent in tumors than in blood or normal tissues, particularly in ovarian and colorectal tumors, although more of these cells were identified in lung and breast samples (data generated by the InDrop technique) (Supplementary Fig. 2f-i).

cDCs were the fourth most prevalent subpopulation of MDCs, accounting for 9% of these cells (Fig. 1f). Within the cDC population, three main groups were identified (Fig. 2a, b): cDC type 1 (cDC1), characterized by the expression of *CLEC9A* and *CADM1,* denoted as cDC1_CLEC9A (n = 580); cDC type 2 (cDC2) (n = 5347), marked by *CLEC10A* and *CD1C* expression, and further subclassified into six phenotypes (Fig. 2c); and migratory mature cDC (cDC_Mig), expressing *LAMP3* and *CCR7*, named cDC_LAMP3 (n = 285) (Fig. 2a,b).

**Figure 2.**
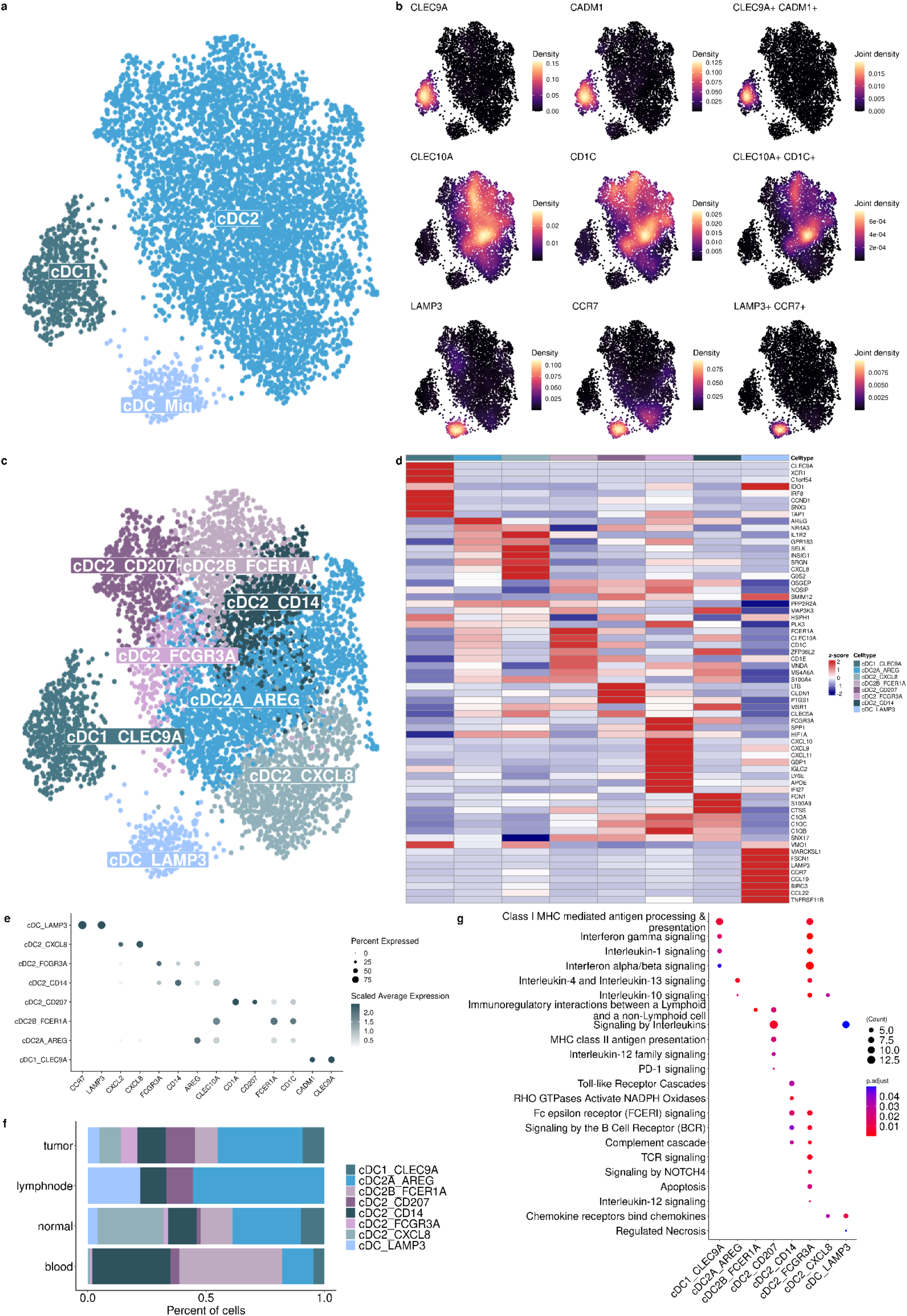
Characterization of conventional Dendritic Cells subpopulations across the conditions. **a** UMAP of cDC subpopulations colored by the three cDC states: cDC1, cDC2, and cDC_Mig. **b** Density plots highlight the gene marker expression and co-expression for each cell type. **c** UMAP of cDC subpopulations colored by the eight subpopulations/states. **d** Heatmap showing the DEGs per cluster. The color scale represents the scaled expression of each gene. **e** Dotplot showing the mean expression of gene markers for each cDC subpopulation. Dot size indicates the fraction of expressing cells, colored based on the normalized expression levels scale. **f** Barplot showing the relative distribution of cDC subpopulations across sample types. **g** Enrichment pathways analysis of cDC subpopulations using Reactome database. The size of each circle represents the number of genes belonging to a given pathway and such circles are colored by p-adjust.

The cDC2 subpopulations were categorized into previously described types A, resembling an anti-inflammatory profile, and B, a proinflammatory profile^19^, named cDC2A_AREG (n = 2,044) and cDC2B_FCER1A (n = 726), respectively (Fig. 2c-e). cDC2A_AREG was enriched in tumors and lymph nodes, whereas cDC2B_FCER1A was found mostly in the blood (Fig. 2f and Supplementary Fig. 3a). We also found two other subpopulations that were present predominantly in blood and tumor samples, respectively (Supplementary Fig. 3a): cDC2_CD14 (n = 765) and cDC2_FCGR3A (n = 314), likely to be monocyte-derived.

Another cDC2 subpopulation was identified by the expression of a chemokine set (*CXCL2* and *CXCL8*), named cDC2_CXCL8 (n = 993) (Fig. 2c,d), mainly found in normal skin samples (Supplementary Fig. 3b,c). We also identified a cDC2 subpopulation expressing *CD207*, a langerin protein encoder (cDC_CD207, n = 505), which was abundant in tumor samples, particularly in lung and ovarian tumors and lymph nodes (Fig. 2f and Supplementary Fig. 3b,c). Although langerin is primarily expressed by epidermal macrophages known as Langerhans cells^20,21^, cDC_CD207 is a type 2 DC subpopulation (expressing *CD1A*) distinct from the aforementioned cell (Fig. 2d,e).

The cDC_LAMP3 mature subpopulation was identified based on the expression of *LAMP3* and *CCR7* genes (Fig. 2b-d). Interestingly, these cells showed increased expression of genes coding for PD-1 and PD-2 ligands, suggesting a potential immunosuppressive role (Supplementary Fig. 3d). Additionally, functional enrichment analysis further supported the cell’s profile (Fig. 2g). DC-activating signals, such as pro-inflammatory cytokines (interferon-alpha/beta and gamma) and antigen presentation, were observed in cDC1-CLEC9A and cDC_CD207. On the other hand, cDC2A_AREG appeared to have regulatory properties, including IL-10, -4, and -13 signaling (Fig. 2g and Supplementary Table 5).

Monocytes are precursors of tumor-infiltrating myeloid cells, and we identified four major groups of these cells (Fig. 3a, b), reflecting six subpopulations/states with distinct gene programs (Fig. 3c-e). One of these subpopulations, Mono_FCG3RA (n = 2,018), exhibited a non-classical phenotype characterized by the co-expression of *FCGR3A* and *FAM110A* within a single cluster (Fig. 3b-d). The classical phenotype was marked by the expression of common monocyte markers such as *CD14* and *SELL* (Fig. 3b, d), and further differentiated into two subpopulations based on *FOS* expression: Mono_CD14_FOS^-^ (n = 2,150) and Mono_CD14_FOS^+^ (n = 1,497) (Fig. 3c-e). Mono_CD14_FOS^+^ was enriched in lymph nodes and normal tissues, such as the ovary (Supplementary Fig. 4a-c), while Mono_CD14_FOS^-^ (n = 2,150) was more prevalent in blood samples (Fig. 3f and Supplementary Fig. 4a-c). These two states have not been previously described, and their functions remain unclear, although both are enriched in Toll-like receptor activation pathways, which may facilitate their differentiation into macrophages or dendritic cells (Fig. 3g and Supplementary Table 5). Additionally, Mono_IL1B (n = 1,739) was identified as displaying a pro-inflammatory profile with high expression levels of *IL1B*, *CXCL2,* and *EREG* (Fig. 3c-e), as well as other inflammatory genes (Supplementary Fig. 4d and Table 5). This subpopulation also showed increased expression of genes involved in Toll-like receptor cascades and death receptor ligands that activate caspase cascades (Fig. 3g). At this resolution, we were also able to identify two monocyte subpopulations displaying an intermediate phenotype between monocytes and antigen-presenting cells, expressing both genes coding for CD16 and CD14. We called this population Mono_Inter, from which *CXCL10* and *CLEC10A* expressions distinguished the subpopulations into MonoInter_CXCL10 (n = 1,238) and MonoInter_CLEC10A (n = 1,559), respectively (Fig. 3c-e). These subpopulations expressed markers related to antigen presentation, such as *HLA-DMB* and *HLA-DRB6* (Fig. 3d), and were enriched in MHC class II antigen presentation, interferon/gamma signaling, and costimulation by CD28 pathways, as well as PD-1 signaling (Fig. 3g and Supplementary Table 5).

**Figure 3.**
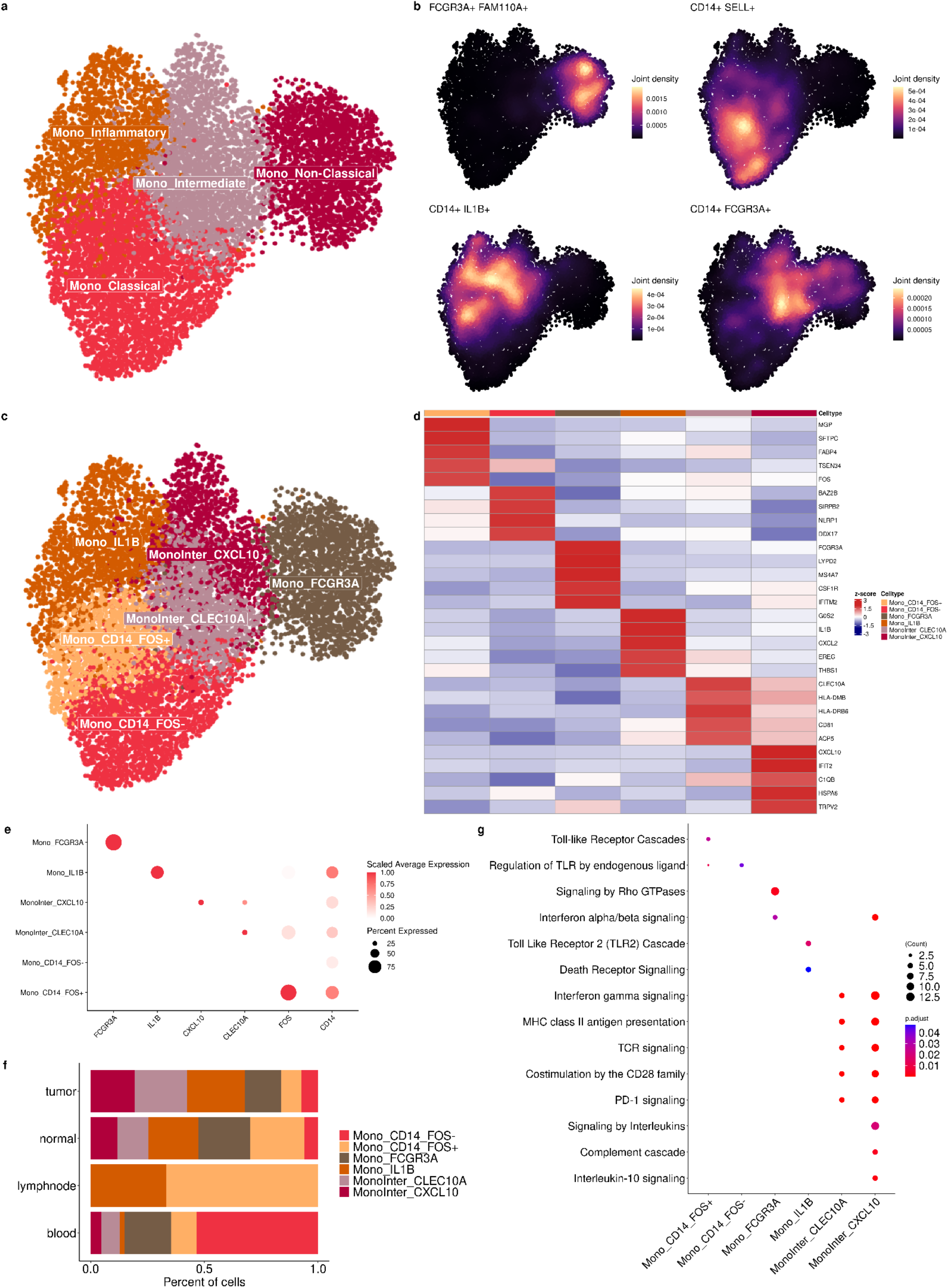
Unraveling the complexity of monocyte subpopulations across the conditions. **a** UMAP of Monocytes subpopulations colored by major classifications: Mono_Inflammatory, Mono_Intermediate, Mono_Non-Classical, and Classical. **b** Density plots highlighting the gene marker expression and gene co- expression of each cell type. **c** UMAP of monocyte subpopulations colored by the six states identified. **D** Heatmap showing the DEGs per cluster. The color scale represents the scaled expression of genes. **e** Dotplot showing the mean expression of gene markers of monocyte subpopulations. Dot size indicates the percent of expressing cells, and the dot color the scaled average expression. **f** Barplot showing the distribution of monocytes across sample types. **g** Enrichment pathways analysis of monocyte subpopulations using Reactome database. The size of each circle represents the number of genes belonging to a given pathway and such circles are colored by p-adjust.

Macrophages are the largest group of mononuclear phagocytes in this study, corresponding to 51.6% of all myeloid cells (Fig. 1f). We identified three main clusters of macrophages based on their ontogeny markers: 1) recruited macrophages distinguished by the expression of monocyte markers such as *VCAN*, *FCN1*, *CD300E*, and *S100A9;* 2) resident-tissue macrophages (RTM), expressing *LYVE1*, *FOLR2*, and *PLTP;* and 3) RTM-like cells expressing markers of both groups (Fig. 4a-c). Although recruited macrophages were the most common type in tumors, RTMs were surprisingly shown to be expanded in tumors (Supplementary Fig. 5a, b), particularly in lung samples (Supplementary Fig. 5c-f). RTM-like cells were found in all conditions except uveal melanoma, melanoma, liver tumors, and normal breast tissue (Supplementary Fig. 5c-f). Also, we observed an RTM-like enrichment in ovarian samples predominantly in normal tissue compared to tumors (Supplementary Fig. 5e).

**Figure 4.**
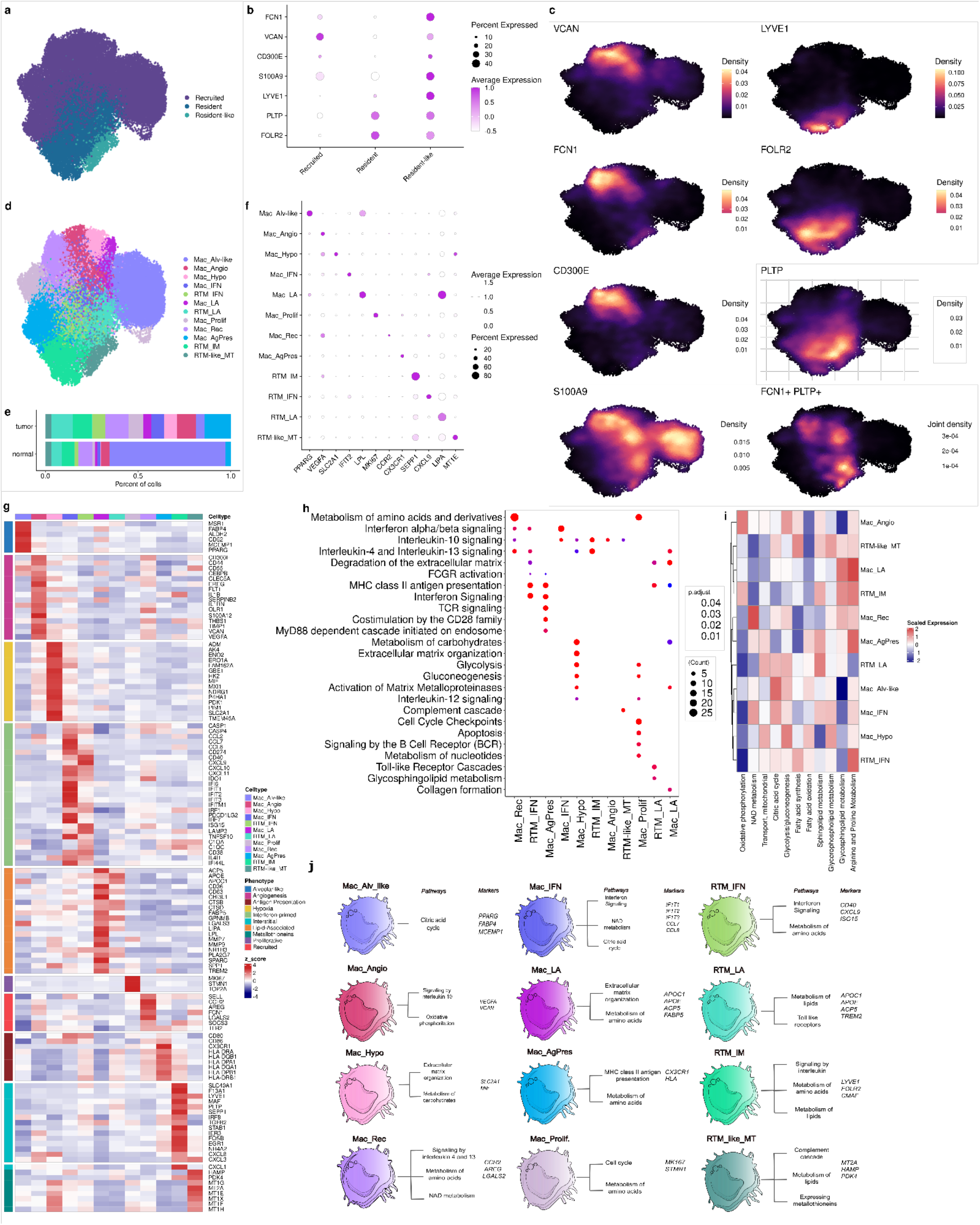
Diverse landscape of macrophage states heterogeneity across distinct cancer types. **a** UMAP of macrophages color-coded by the ontogeny subdivision. **b** Dotplot showing the mean expression of the resident and recruited-macrophages related gene markers. Dot size indicates the percent of expressing cells, and the dot color is related to the scaled average expression. **c** Density plots highlighting the gene mark expression and gene co-expression for each cell type. **d** UMAP of macrophages colored by the twelve states/subpopulations identified. **e** Barplot showing the distribution of macrophage subpopulations across sample types. **f** Dotplot showing the mean expression of gene markers related to each macrophage subpopulation. Dot size indicates the percent of expressing cells, and the dot color the scaled average expression. **g** Heatmap showing gene signatures per subpopulation. The color scale represents the scaled expression of each gene. **h** Dotplot representing the functional enrichment analysis of macrophage subpopulations using the Reactome database. The size of each circle represents the number of genes belonging to a given pathway and such circles are colored by p-adjust. **i** Heatmap showing selected metabolic pathways for each macrophage state. The color scale represents the median effect size (Cohen’s d, the difference between two means) for an increase (red) or decrease (blue) in a given metabolic pathway. **j** Schematic overview of the diverse phenotypes and functional signatures of macrophage subpopulations characterized in this study.

In addition to classifying macrophages based on origin, we also distinguished 12 subpopulations based on gene programs (Fig. 4d-g), with greater diversity in tumors than normal samples (Fig. 4e and Supplementary Fig. 5d). The subpopulations were stratified according to their functional gene signatures, mostly following a consensus model for TAM diversity^22^. The alveolar-like macrophage (Mac_Alv-like, n = 8,816), marked by the expression of *PPARG* and *MCEMP1,* was mostly found in lung samples, as expected (Fig. 4e and Supplementary Fig. 5d-f). The pro-angiogenic macrophage subpopulation (Mac_Angio, n = 3,108) displayed high expression of genes associated with angiogenesis, such as *VEGFA*, *VCAN*, and *EREG,* enriched in lung and ovarian tumors (Supplementary Fig. 5d-f). Hypoxia-associated macrophages (Mac_Hypo, n = 1,802) were mostly found in tumor samples (lung and ovary) (Supplementary Fig. 5d-f) and presented an enrichment of genes associated with hypoxia, such as *SLC2A1* and *ERO1* (Fig. 4f, g).

Two populations expressing high levels of interferon-primed genes, such as *CCL8, IDO1,* and *CXCL9,* were named RTM_IFN (n = 2,061) and Mac_IFN (n = 1,981), abundant in ovarian and lung tumor samples, respectively (Fig. 4d-g and Supplementary Fig. 5d-f). Two subpopulations revealing a lipid-associated (LA) metabolism transcriptional signature (Fig. 4d-g), including genes such as *FABP5*, *LPL*, and *LIPA,* named RTM_LA (n = 3,569) and Mac_LA (n = 1,249), were found enriched in breast, lung, and colorectal tumors (Supplementary Fig. 5d-f). Another subpopulation expressing high levels of *MKI67* and *STMN1* was classified as Mac_Prolif (n = 2,234) and found to be enriched in liver, melanoma, and ovary tumor samples (Fig. 4d-g, Supplementary Fig. 5e,f). We also identified a large population of macrophages in uveal melanoma metastatic samples (Supplementary Fig. 5e) expressing high levels of *CCR2* (Fig. 4d,f), a receptor for monocyte chemoattractant protein-1, and *LGALS2*. Therefore, this population was named recruited macrophage Mac_Rec (n = 4,073).

We identified one recruited subpopulation displaying a pro-inflammatory phenotype, expressing the receptor of the chemokine CX3CL1 (*CX3CR1)* and genes from the antigen-presentation pathway (*HLA-A/C*, *HLA-DQA1/B1*). This population was named Mac_AgPres (n = 3,965) (Fig. 4d-g). Additionally, a population expressing high levels of resident markers such as *LYVE1, FOLR2,* and *PLTP* resembled perivascular macrophages with specific expressions of genes such as *MAF* and *SEPP1*. This subpopulation was named RTM_IM (n = 3,507) and was found to be enriched in normal tissues (Fig. 4e, Supplementary Fig. 5d). A RTM-like macrophage subpopulation was found to express high levels of metallothioneins (Fig. 4d-g), named RTM-like_MT (n = 1,209), and found predominantly in ovary samples (Supplementary Fig. 5e,f).

By examining the enriched pathways in each subpopulation, we were able to identify distinct roles for each group. Enriched pathways in Mac_IFN, Mac_Rec, RTM_IFN, and Mac_AgPres subpopulations were linked to the interferon signaling pathway. Antigen presentation genes were enriched in RTM_IFN, RTM_LA, Mac_LA, and Mac_AgPres, with the latter also expressing genes related to phagocytosis, such as the MyD88-dependent cascade initiated on endosomes and FCGR activation (Fig. 4h). Interestingly, Mac_Rec, Mac_IFN, Mac_Hypo, RTM_IM, Mac_Angio, RTM-like_MT, and RTM_IFN showed enrichment of genes involved in the IL-10, -4, or 13 pathways, which have been associated with pro-tumoral processes (Fig. 4h). Mac_Hypo, RTM_LA, Mac_LA, and RTM_IFN displayed pathways related to extracellular cellular matrix remodeling (Fig. 4h and Supplementary Table 5).

Macrophages have the ability to utilize diverse energy sources for their metabolism, which is influenced by their role in inflammation and the TME (Fig 4i). Certain subpopulations, namely Mac_Hypo and Mac_Angio, appear to rely on glucose metabolism. Among these, Mac_Angio also exhibits enrichment in genes involved in the oxidative phosphorylation pathway. On the other hand, subpopulations like Mac_LA, RTM_LA, and RTM-like_MT expressed more genes associated with fatty acid lipid metabolism. Notably, NAD metabolism is highly enriched in the Mac_Rec and Mac_IFN subpopulations. The Mac_LA, RTM_IFN, and Mac_AgPres subpopulations expressed more genes related to amino acid metabolism. Interestingly, the metabolic profiles of the different RTM subpopulations appeared to vary significantly from one another. For a comprehensive overview of the major metabolic pathways, please refer to Supplementary Fig. 6.

The consistency and reproducibility of our findings are validated by the strong correlations we observed between our identified populations and those described by other research groups^23,24^. Moreover, our approach has uncovered previously unidentified subpopulations. When assessing cluster purity, we found that our technique produced highly homogeneous clusters compared to previous integration approaches, indicating its effectiveness (Supplementary Fig. 7 and Supplementary Table 6). Overall, our approach expanded the catalog of mononuclear phagocytes in the TME, enabling the identification of rare populations such as cDC2_CXCL8. Additionally, our analysis led to the discovery of distinct phenotypes of RTMs, including interstitial, lipid-associated metabolism, and interferon-primed macrophages. Furthermore, we unveiled new subpopulations derived from monocytes, such as Mac_Hypo and RTM-like_MT. This led to valuable insights into the diversity and complexity of mononuclear phagocyte populations in the TME, especially macrophages (Fig. 4j), underscoring the importance of employing deep scRNA-Seq analysis to uncover previously unknown subpopulations.

### A plethora of macrophages phenotypes and functional states co-exist in the TME

Since M1 and M2 polarization gene programs are well-known^25–27^ and reflect important macrophage-related features, we attempted to assign these profiles to the 12 macrophage subpopulations identified (Fig. 5a, b). Although the RTM-IFN and RTM_IM were, respectively, the subpopulations displaying the higher scores for M1 and M2 gene signatures (Supplementary 8a, b and Supplementary Table 7), there was no clear overrepresentation of genes associated with either polarization profile in any of the subpopulations. In effect, populations such as the RTM-IFN express high levels of genes encoding pro-inflammatory cytokines such as *CXCL9*, *CXCL10,* and *CXCL11*, as well as immunosuppressive molecules such as PDL-1 and PD-L2 (Fig. 5a, b). Although M1/M2 ratio scores (Supplementary Fig. 8c) discriminate the subpopulations regarding their polarization states, this dichotomy oversimplifies the highly specialized, transcriptomically dynamic, and extremely heterogeneous nature of macrophages *in vivo*.

**Figure 5.**
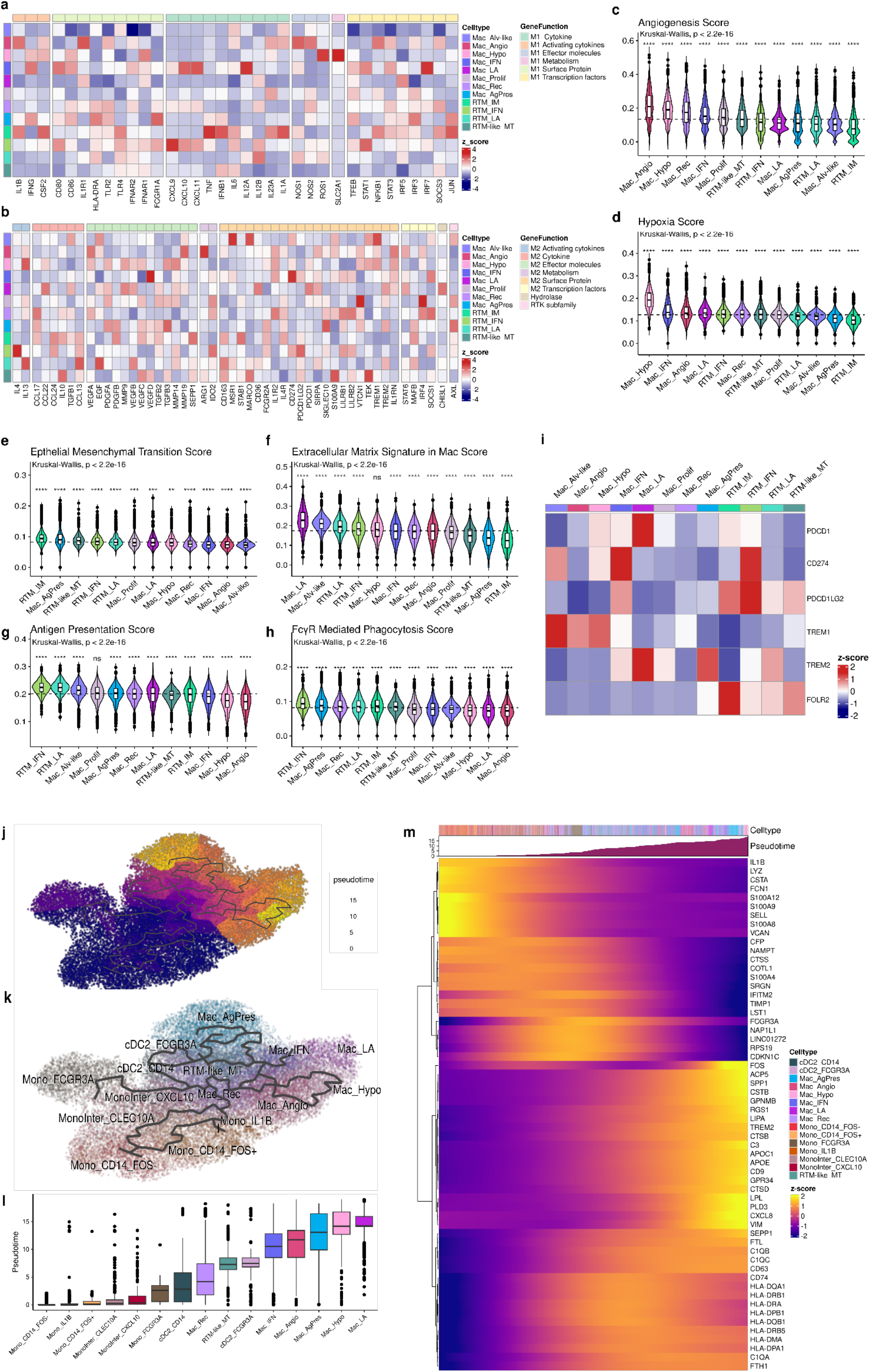
Macrophages exhibited diverse phenotypes and functional states in the TME. **a-b** Heatmap showing the score signature of M1 **(a)** and M2 **(b)** markers across the macrophage subpopulations. The color scale represents the scaled expression of each gene. **c-h** Violin plot showing the gene signature of five hallmarks of cancer: angiogenesis **(c)**, hypoxia **(d)**, EMT **(e)**, ECM **(f)**, antigen presentation **(g)**, and phagocytosis **(h)** (Supplementary Table 4). Boxes span the first to third quartiles; the horizontal line inside the boxes represents the median black and dots represent outlier samples in each group. For statistical significance, the Kruskall–Wallis test followed by Wilcoxon was performed to compare each of the ten groups against “all” (i.e. base-mean). Ns non-significant. *p < 0.05; **p < 0.01; ***p < 0.001; ****p < 0.0001. **i** Heatmap of immunosuppressive gene signature. The color scale represents the scaled expression of each gene. **j-m** trajectory inference of monocytes-derived cells in the TME. UMAP color-coded by the pseudotime showing simulated trajectories **(j)** and by the subpopulations **(k)**; Boxplot comparing estimated pseudotime across the subpopulations arranged in ascending order. Boxes span the first to third quartiles; the horizontal line inside the boxes represents the median black and the dots represent outlier samples in each group **(l)**. Heatmap showing expression variation of the indicated transcripts (only genes with q-value = 0 and morans_I > 0.25, are depicted **(m)**. EMT: Epithelial-Mesenchymal Transition; ECM: extracellular matrix.

To gain further insights into the roles of macrophage subpopulations in the TME, we investigated their associations with cancer hallmark signatures^28^ (Supplementary Table 4). Consistent with their names, Mac_Angio exhibited higher scores for the Angiogenesis signature, while Mac_Hypo showed elevated scores for the Hypoxia signature (Fig. 5c, d). Notably, RTM_IM demonstrated enrichment in signatures associated with the induction of epithelial-mesenchymal transition (EMT) (Fig. 5e). The extracellular matrix (ECM) signature in macrophages^29^ was predominantly observed in Mac_LA, Mac_Alv-like, and RTM_LA subpopulations (Fig. 5f and Supplementary Table 4). In terms of antigen presentation^28^, the RTM_IFN subpopulation exhibited the highest scores, while Mac_AgPres also exhibited FcγR-mediated phagocytosis signatures (Fig. 5g, h).

To explore the potential immunosuppressive role of macrophages, we examined the co-expression of checkpoint genes and immune regulators. We found that *TREM2*, a marker for immunosuppressive tumor-associated macrophages (TAMs)^30^, was highly expressed in both resident and recruited LA populations, as well as in Mac_AgPres (Fig. 5i). However, only Mac_LA exhibited high levels of co-expression with the *PDCD1* gene (programmed death 1 PD-1). Furthermore, Interferon-primed subpopulations demonstrated elevated expression of *CD274* and *PDCD1LG2*, the genes encoding PD-L1 and PD-L2, respectively (Fig. 5i). RTM-IM, which expressed high levels of *FOLR2*, also displayed high expression of the PD-L2 coding-gene. Interestingly, *TREM1*, typically highly expressed in macrophages in inflamed tissues^31,32^, was mainly expressed by the Mac_Alv-like, Mac_Angio and Mac_Hypo subpopulations (Fig. 5i). We also evaluated the myeloid-derived suppressor cell signature proposed by Alshetaiwi et al (2020) in all myeloid-derived suppressor cells (MDSCs). However, none of the subpopulations resembled that particular signature (Supplementary Fig. 9). Nevertheless, all resident subpopulations (RTM_IM, RTM_IFN, and RTM_LA) exhibited higher suppressive profiles based on *CD84* expression, followed by Mono_CD14_FOS^-^ (Supplementary Fig. 9).

To elucidate the intricate differentiation trajectory in the monocyte-macrophage and monocyte-DC axes, we employed an unsupervised pseudo-time analysis strategy. By using Mono_CD14_FOS^-^ as the “root” for trajectory inference, our analysis supported the notion that CD14^hi^ monocytes serve as precursor cells for both Mono_CD16^hi^ and monocyte-derived cells in tissues (Fig. 5j-m). Intermediate monocytes (MonoInter_CXCL10 and MonoInter_CLEC10A) can potentially give rise to both cDC2 and Mac_Rec subpopulations^34^. Notably, the Mac_Rec subpopulation appears to be the earliest monocyte-derived macrophage subpopulation ontologically, acting as a crucial precursor for other macrophage states, including RTM-like. Two distinct phenotypes, originating from two main branches, were identified as the most distant from the root: 1) Mac_AgPres and 2) Mac_LA, both of which are characterized by high levels of *TREM2* expression. Through the analysis of the genes that co-vary over pseudo-time, we were able to identify a set of monocyte-related genes (*VCAN, SELL,* and *FCN1*) at the beginning of the trajectory and both specific and general phenotype markers of the macrophage lineages (*APOE*, *TREM2*, *SPP1*, *HLA-DRA*, *C1QA,* and *CD63*) in late branches (Fig. 5k-m).

### Deciphering the clinical impact of TREM2 macrophages in the TME

The clinical significance of MDC subpopulations as potential targets has garnered increased interest due to their higher abundance in the TME. To further explore their clinical impact, we conducted an investigation using larger cohorts to assess MDC subpopulations in different tumor types. We utilized a set of 47 subpopulation signatures to estimate the proportions of cell types through deconvolution analysis of bulk RNA-Seq samples obtained from nine tumor types available in The Cancer Genome Atlas (TCGA) database. To validate the applicability of our signature matrix in estimating cell populations within the TCGA data, we calculated the correlation between bulk RNA-Seq and scRNA-Seq data. Encouragingly, we observed a significant correlation in all TCGA cohorts (Pearson R^2^ > 0.7, p-value ≤ 0.05) (Supplementary Fig. 10a-i).

As expected, all the tumor samples were enriched with malignant cells (∼50%), except for lung adenocarcinoma (LUAD), which had very similar proportions of malignant cells (16% of total cells) and fibroblasts (19.9% of total cells) and a higher content of epithelial cells (43.4% of total cells) (Fig. 6a). All proportions of subpopulations across tumor types are shown in Supplementary Fig. 11 a-m. Breast cancer (BRCA) presented a higher proportion of non-epithelial cells, and together with LUAD, these tumors were the most enriched in MDCs (Fig. 6a). Stratifying for mononuclear phagocyte subpopulations, macrophages were among the most abundant cell types (∼ 12% of total). The RTM_IFN subpopulation was more commonly found in different tumor types, especially in skin cutaneous melanoma (SKCM) (7.6% for metastatic and 5.6% for primary tumor) and uveal melanoma (UVM) samples (1.9% of total cells), whereas RTM_IM was predominantly found in lung tumors (6.06% in LUAD and 3.9% in lung squamous cell carcinoma). The Mac_LA subpopulation was mainly found in LUAD and BRCA tumors, while Mac_AgPres was mainly found in colorectal tumors (2.6% of total cells) (Fig. 6b). Mast cells and neutrophils constituted a small proportion of cells predicted in the TME and were mainly enriched in BRCA tumors (Supplementary Fig. 12).

**Figure 6.**
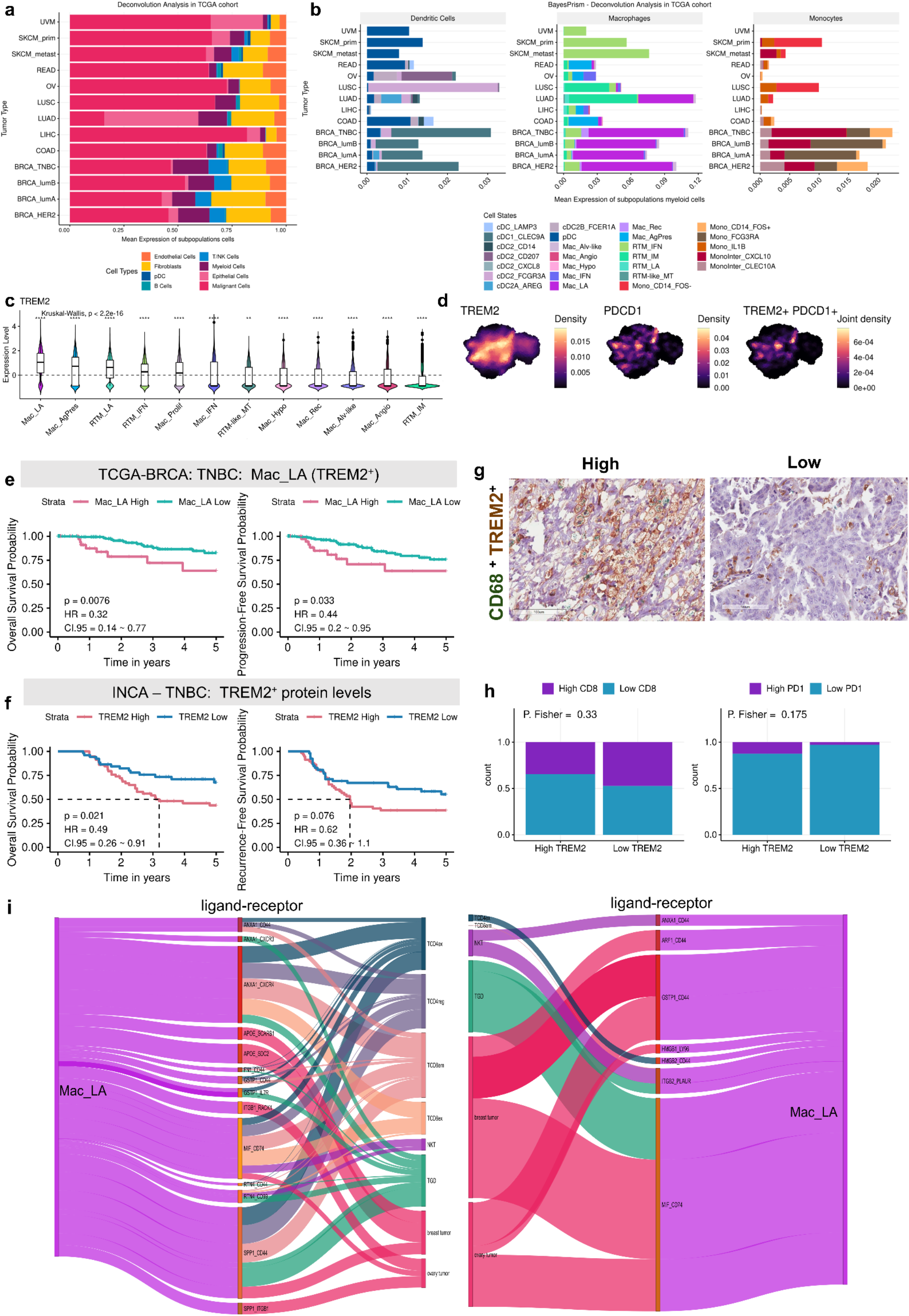
Clinical significance of *TREM2*-expressing macrophage subpopulations in different tumor types. **a** Barplot showing the distribution of predicted cell types across the bulk RNA-Seq profiled samples from TCGA database. **b** Barplot showing the distribution of mononuclear phagocyte states across different tumor types. **c** Violin plot showing the *TREM2* gene expression across macrophage populations. **d** Density plots showing the expression of PD1 encoding gene and TREM2 in macrophages, as well as their co- expression. **e** Kaplan–Meier curves for Overall and Progression-Free survival for the TNBC patients from TCGA-BRCA cohort. The patients were divided into low-Mac_LA (TREM2+) and high-Mac_LA (TREM2+) groups using the surv_cutpoints function. P-values were calculated using the log-rank test. **f** Kaplan–Meier curves for Overall and Recurrence-Free Survival analysis for TNBC patients from INCA cohort. P-values were calculated using the log-rank test. The patients were divided into LOW-TREM2+ and HIGH-TREM2+ groups using the surv_cutpoints function after ImageJ analysis. **g** IHC representative of CD68+ TREM2+ expression in TNBC samples from INCA cohort. Image obtained by Aperio ImageScope v12.4.6.5003. **h** Proportions of CD8+ and PD1+ markers in LOW-TREM2+ and HIGH-TREM2+ groups in the TNBC samples from the INCA cohort. **i** Sankey diagram representing the putative cell-cell interactions between Mac_LA and other cells. In the middle are the pairs of ligands and receptors, and the thickness of the lines represents the mean expression of both ligand and receptor converted into z-score. HR, hazard ratio. CI, confidence interval.

We then conducted an investigation to assess the impact of immunosuppressive *TREM2^+^* macrophage subpopulations (Fig. 6c), particularly Mac_LA, due to its high abundance in tumors and co-expression with the gene encoding PD-L1 (Fig. 6d). Patients with triple-negative breast cancer (TNBC) were categorized into groups based on a high and low abundance of Mac_LA in their TME. Those with a high quantity of Mac_LA demonstrated significantly worse overall survival (OS) and progression-free survival (PFS) compared to those with low levels of Mac_LA (OS, log-rank test, p-value = 0.0076; PFS, log-rank test, p-value = 0.033) (Fig. 6e). However, this trend was divergent in other BRCA subtypes (Supplementary Fig. 13a-c) as well as HGSOC samples (Supplementary Fig. 14a, b). We also evaluated global *TREM2* expression and protein levels as clinical biomarkers in different cohorts. Global *TREM2* gene expression was associated with poorer OS compared to the lower expression group in patients diagnosed with different BRCA subtypes (TNBC, log-rank test, p-value = 0.0031, Supplementary Fig. 13d-g) and HGSOC (OS, log-rank test, p-value = 0.013, Supplementary Fig. 14c, d). When protein levels were assessed using immunohistochemistry, only the TNBC cohort was significantly associated with TREM2^+^ protein levels (Figure 6f, g and Supplementary Fig. 14e, g), consistent with our bioinformatic deconvolution analysis.

Interestingly, in HGSOC the higher intratumoral expression of CD68^+^ and TREM2^+^ was accompanied by increased PD-1 and CD8 levels, suggesting the involvement of the PD-1 receptor in the clinical response associated with TREM2-expressing macrophages (Supplementary Fig. 14h). The same process was observed in TNBC, when we evaluated the pan protein level of TREM2 (Fig. 6h). Notably, PD-1 expression was significantly associated with poorer OS and recurrence-free survival (RFS) in TNBC patients (OS, log-rank test, p-value = 0.0053; RFS, log-rank test, p-value = 0.03) (Supplementary Fig. 15a). No difference in OS and RFS were observed for CD8^+^ and PD-L1 (Supplementary Fig. 15b, c).

To identify communication axes involving Mac_LA, we conducted a ligand-receptor inference analysis between these cells and other key players in tumor progression, including malignant cells and T/NK cells. We first clustered and annotated these cells based on canonical gene markers and DEGs, resulting in 11 cell types (Supplementary Fig. 16a, b). Mac_LA exhibited 714 putative intercellular communications, as indicated by ligand-receptor pairings between Mac_LA and lymphocytes, as well as malignant cells (Supplementary Table 8). Among these interactions, CD8^+^ T cells emerged as the most representative cell type (Supplementary Fig. 16c). Notably, the axes of major expression in these interactions were SPP1_CD44, MIF_CD74, ANXA1_CXCR4 and GSTP1_CD44 (Fig. 6i). These findings shed light on potential communication pathways involving Mac_LA and provide valuable insights into its role in tumor microenvironment interactions.

### MDC subpopulations can display distinct clinical outcomes depending on their niche

To further investigate the clinical impact of other MDC subpopulations inferred in the TME across different tumor types, we conducted a univariate COX regression analysis (Supplementary Fig. 17a). Mono_FCG3RA subpopulation, enriched in BRCA tumors, especially the Luminal B subtype (representing 1.22% of total cells) (Fig. 6b), exhibited a significant association with poor OS in TNBC when assessed through uni and multivariate regression analysis (log-rank p-value = 0.0027, Supplementary Fig. 17a-c), reinforcing its independent value as a prognostic marker. Conversely, in patients with the Luminal A subtype, a higher percentage of this population was correlated with improved OS (log-rank p-value = 0.0011, Supplementary Fig. 17a, c, e), indicating that its clinical impact depends on the specific TME context. Furthermore, Mono_FCG3RA was markedly reduced in TNBC patients with “tumor-free” status (p-value = 0.0095), whereas this was not observed in the Luminal A subtype (Supplementary Fig. 17f).

Another subpopulation with distinct impacts depending on the TME niche is the resident macrophage RTM_IM. In HGSOC and TNBC, RTM_IM is associated with poor OS, while in lung adenocarcinoma (LUAD), it is linked to improved OS (univariate COX regression and log-rank, p-value < 0.05, Fig. 7a and Supplementary Fig. 17a). Of note, no significant clinical impact was observed in other BRCA subtypes (Supplementary Fig. 18 a-c). Patients with TNBC and HGSOC were categorized into groups, and those with a high quantity of RTM_IM demonstrated significantly worse OS compared to those with low levels of RTM_IM (log-rank test, p-value = 0.0025, and log-rank test, p-value = 0.017, respectively), but there was no significant difference for progression-free survival (PFS) (log-rank test, p-value = 0.34, and log-rank test, p-value = 0.65, respectively) (Fig. 7b, c). When analyzing FOLR2 as a biomarker, high protein levels were associated with poor OS in TNBC (log-rank test, p-value = 0.0088, Fig.7d); in HGSOC the high expression was associated with a poor PFS (log-rank test, p-value = 0.0034, Fig. 7e). The global expression of *FOLR2* showed distinct results depending on the analyzed cohort (Supplementary Fig. 19 a-c). Representative images of tissue microarrays, and the relationship between FOLR2 and CD8 or PD1 are provided in Figure 7f-i.

**Figure 7.**
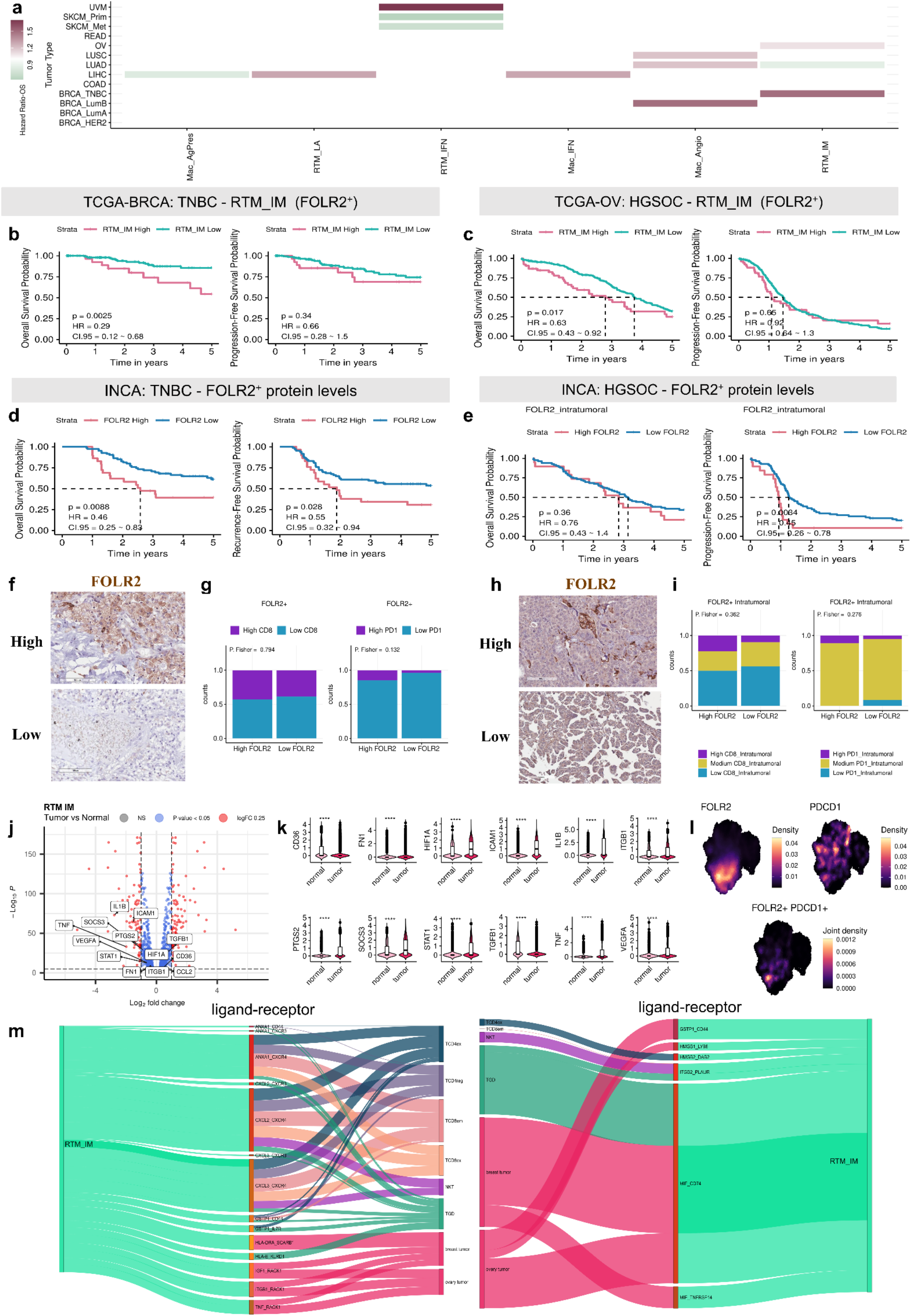
Dual role of the resident macrophage subpopulation RTM_IM FOLR2+ in TME: Implications for prognosis and therapeutic targeting. **a** Univariate cox regression analysis for predicted macrophage subpopulations in bulk RNAseq samples from different cancer types. Color scale represents the Hazard Ratios (HR) that were calculated relative to the patient’s overall survival; **b-e** Kaplan–Meier curves for Overall and Progression-Free survival for the TNBC **(b)** and HGSOC **(c)** patients from the TCGA cohort. The patients were divided into LOW-RTM_IM (*FOLR2*+) and HIGH-RTM_IM (*FOLR2*+) groups using the surv_cutpoints function. P-values were calculated using the log-rank test. Kaplan–Meier curves for Overall and Progression-Free survival for the FOLR2 protein through IHC for TNBC **(d)** and HGSOC **(e)** patients from the INCA cohort. **f-i** Representative IHC of FOLR2 expression (image obtained by Aperio ImageScope v12.4.6.5003) and Barplot showing the proportions of CD8+ and PD1+ in the HIGH-FOLR2 and LOW- FOLR2 groups in TNBC **(f-g)** and HGSOC samples **(h-i)** from INCA cohort. **j** Volcano Plot showing DEGs in RTM-IM when in tumor vs normal niche (p-value =< 0.05, cutoff FC < 1). **k** Violin plot of DEGs in RTM-IM cells comparing tumor and normal niches (**** p < 0.0001). **l** Density plots highlighting the co-expression of FOLR2 and PDCD1 genes in the macrophage populations. **m** Sankey diagram representing the putative cell-cell interactions between RTM_IM and other cells. Pairs of ligands and receptors are represented in the middle, and the thickness of the lines represents the mean expression of both ligand and receptor converted into z-score. HR, hazard ratio. CI, confidence interval.

Due to its dual role in prognosis associated with niche variability, we compared DEGs between RTM_IM cells in normal and tumor samples. Protumorigenic factors associated with hypoxia and angiogenesis, like *HIF1A* and *VEGFA*, as well as genes related to immunosuppressive phenotypes, such as *IL1B* and *TGFB1*, were highly expressed in the tumor context (Fig. 7j, k). Since RTM_IM exhibits high levels of *FOLR2* and the immunosuppressive gene *PDCD1* (Fig. 7l), we also investigated the association with other immune markers. Interestingly, an increase in CD8^+^ T cell frequency was observed in tumor samples displaying high global levels of FOLR2 (Fig. 7g) and intratumoral FOLR2 macrophages (Fig. 7i) accompanied by higher levels of PD-1^+^. No significant difference in OS and PFS was observed for PD-1^+^ and CD8^+^ cells, or CD68^+^ macrophage (Supplementary Fig. S20a-h). These findings support the notion that FOLR2^+^ macrophages are reprogrammed during tumor formation by activating immunosuppressive gene programs, potentially leading to the exhaustion of CD8^+^ T lymphocytes in the context of TNBC and ovarian tumors. This highlights RTM_IM as a promising new therapeutic target. Analyses of OS (uni and multivariate) from other tumor types are described in Supplementary Tables 9 and 10.

In our study, we further explored the clinical impact of RTM_IM by investigating the communication axes between this subpopulation and malignant cells, as well as T/NK cells. We identified 889 intercellular communications involving the “RTM_IM” cell type (Supplementary Fig.16 c, Supplementary Table 8). Notably, the connections with CD8^+^ T and CD4^+^ T cells were particularly enriched, with ANXA1_CXCR4, CXCL2_CXCR4, and CXCL3_CXCR4 axes being prevalent in these communications (Fig. 7m). Additionally, similar to Mac_LA, MIF_CD74 appeared to be an important axis ligand involved in cell-cell communication between RTM_IM and tumor cells (Fig. 7m).

In comparison to previous studies, our method of integrating scRNA-Seq datasets, clustering, and characterizing cell subpopulations significantly expanded the catalog of mononuclear phagocytes in the TME. This approach allowed us to identify rare and poorly described subpopulations, including cDC2_CXCL8, Mono_CD14_FOS^+^, and Mono_CD14_FOS^-^. Furthermore, our analysis led to the identification of distinct RTM phenotypes, such as interstitial, lipid-associated metabolism, and interferon-primed macrophages, as well as unveiling new monocyte-derived subpopulations like Mac_Hypo and RTM-like_MT. Moreover, we unveiled the clinical impact of specific TREM2^+^ (Mac_LA) and FOLR2^+^ (RTM_IM) subpopulations in both TNBC and HGSOC patients, revealing new possibilities for targeted treatments.

## Discussion

Integration of single-cell data has proven to be a game changer for uncovering valuable biological information from large and complex volumes of data. By pooling data from diverse sources, it is possible to unveil previously hidden cellular heterogeneity and discover novel cell states and phenotypes by enhancing the robustness and reliability of the findings. Here we have thoroughly integrated and characterized cell subpopulations in the TME, yielding a large and comprehensive dataset comprising different tumors, technologies, and sample types, generated across multiple conditions and donors. Notably, we expanded the classification of myeloid-derived cell types including monocytes, macrophages, cDCs, neutrophils, and mast cells to 29 distinct subpopulations. We also distinguished resident-tissue macrophages from those originating from monocytes. Through a rigorous methodology and extensive curation, we identified more homogeneous clusters compared to previous approaches^23,24^, unraveling high-quality clusters representing a wide range of cells found in the TME of solid tumors, including rare states never previously characterized such as Mono_CD14_FOS^+/-^, Mac_Hypo, and RTM-like_MT. These findings offer valuable insights for future research and the development of novel immune checkpoint blockades and target therapies.

While no integrative approach has previously described different mast cell phenotypes/states in the TME, our findings show that they are linked to a better prognosis in LUAD, although they can display gene programs involved in immune response inhibition, such as PD-1 signaling. For tumor-associated neutrophils, their single-cell characterization has been limited due to technological constraints. Nonetheless, our work overcomes this limitation by assembling a substantial collection of these cells within the TME comprising data from different technologies, including InDrop Seq^35^. Among the identified neutrophil subpopulations, neutrophils expressing high levels of *CXCL8* were mainly found in tumors. This cytokine has been previously associated with tumor growth and metastasis^36–38^, and CXCL8-expressing neutrophils were associated with an anti-inflammatory profile involving IL-4 and -13 signaling, and with poor prognostics in LUAD. Nevertheless, the absence of neutrophils in normal tissues used in this work might introduce a potential bias, leading to the observation of only pro-tumorigenic activities in these cells.

Mononuclear phagocytes play a pivotal role in development and homeostasis maintenance^39^. Their remarkable plasticity enables them to actively participate in various critical processes, including organogenesis, tissue repair, and immune defense. In addition to producing essential growth and inflammatory factors that orchestrate lymphocyte responses, they are involved in extracellular matrix maintenance, angiogenesis, tissue enervation, and the clearance of apoptotic cells^40,41^. Interestingly, in the context of tumor development, which shares similarities with organ development, these functions are also highly prevalent. Notably, in lung metastasis, mononuclear phagocytes account for more than 60% of the TME cell population, underscoring their significant involvement in the tumor microenvironment and potentially impacting disease progression.

In our study, we identified eight distinct states of cDCs based on their unique gene signatures. Among them, one cDC type 1, expressing *CLEC9A* and *CADM1,* six subpopulations of cDC type 2, distinguished by *CD1C* and *CLEC10A* expression, and one mature subpopulation expressing *LAMP3* and *CCR7*. Interestingly the cCD2_CXCL8, a newly described phenotype, exhibits contrasting prognostic implications in different tumor types. Moreover, we identified six monocytes displaying either pro- or anti-tumoral properties, regulating a variety of processes from angiogenesis to immune modulation in a context-dependent manner. Two of these subpopulations, Mono_CD14_FOS^+^ and Mono_CD14_FOS^-^, are described for the first time. The transcription factor c-Fos has been shown to positively regulate monocyte development^42,43^. Resting monocytes typically have low or undetectable c-Fos mRNA levels, and their activation is associated with a rapid and transient increase in c-Fos mRNA levels^42^. Furthermore, c-Fos overexpression stimulates growth arrest and terminal differentiation into macrophages^42,44^. In addition, the Mono_FCGR3A associates with a worse prognosis in the TNBC subtype, aligning with the findings of Chen et al., who have linked non-classical monocytes with immunosuppressive functions^45^. However, an opposite impact is observed in the LumA subtype, where the Mono_FCG3RA enrichment is associated with better prognosis, reinforcing that MDC association with cancer prognosis is context-dependent.

Macrophages are versatile immune cells present in various tissues and their ontogeny and functional diversity have been the subject of extensive research. Studies have revealed that TAMs can be derived from circulating monocytes recruited to the tumor site or from RTMs that adapt to the tumor milieu^46^. These TAMs undergo polarization into various phenotypes influenced by their environment, such as the M1 and M2 states, classically associated with pro-inflammatory and immunosuppressive functions, respectively^47–49^. However, our study challenges the traditional binary division of macrophages since none of the states identified clearly displayed M1 or M2 signatures. Instead, we observed that some populations simultaneously expressed genes associated with both signatures, highlighting the complex nature of macrophage plasticity in the TME. To address this complexity and to follow recent efforts to standardize nomenclatures, our in-depth analysis identified twelve distinct TAM states, each characterized by a unique transcriptional profile and ontogeny. We based our annotation mainly on a recently proposed consensus model of TAM diversity gene signatures^22^, and report new phenotypes, such as Mac_Hypo, Mac_AgPres and three different profiles of RTMs (RTM_LA, RTM_IFN and RTM_IM), which were frequently lumped together in a single group of cells in multiple TME studies^22,50^. We also report RTM-like_MT, a subpopulation that displays markers associated with both ontogeny types, supporting the notion that under steady-state conditions, RTMs are mostly maintained via self-renewal processes. However, circulating monocytes can give rise to self-renewing RTMs if an adequate habitat is provided ^51,52^.

One of the novel subpopulations we identified Mac_Hypo is predominantly found in lung and ovary tumor samples. This subpopulation shows enrichment of pathways involved in hypoxia, a condition of low oxygen, as well as matrix remodeling. Given its gene program, glycolysis was found as the main metabolic pathway in this subpopulation, potentially associated with a M2-like polarization, a phenotype known to induce angiogenesis, tumor progression, and metastasis^53^. Furthermore, hypoxia can enhance macrophage-mediated T-cell exclusion in a HIF-1α-dependent manner, and immune suppression, as well as upregulate PD-L1, PD-1, and CTLA-4 on CD8^+^ T cells, contributing to T cell exhaustion^54^.

We also investigated the expression of the members of the Triggering Receptor Expressed on Myeloid cells (TREM) family, *TREM1* and *TREM2*, associated with TAMs ^55,56^. Our catalog revealed that TREM2^+^ macrophages^23^ can be further expanded into at least four different cell states including, RTM_LA, Mac_LA, Mac_IFN, and Mac_AgPres. Similarly, *TREM1* can be divided into three subpopulations (Mac_Alv-like, Mac_Angio, and Mac_Hypo). These findings indicate that there is no single phenotype for TREM1^+^ or TREM2^+^ macrophages.

The Mac_LA subpopulation stands out for its significant expression of *TREM2* among the other Mac subpopulations, and its specific implication in clinical outcome is notable in TNBC, but not other BRCA subtypes or HGSOC, despite the association of *TREM2* expression or protein levels with poor overall survival, consistent with findings from other research groups^57–59^. These results suggest that in different tumor types, other TREM2-expressing macrophage subpopulations might play central roles in tumor progression within their respective microenvironments.

Notably, other subpopulations expressing high levels of *TREM2*, such as RTM_LA and Mac_AgPres, are respectively associated with poor and good prognosis in LIHC. The co-expression of *TREM2* and the receptor *PD-1* in Mac_LA raises the possibility that this subpopulation may indeed have immunosuppressive roles in the TME, in line with previous findings^60^. Of note, an important axis of communication between malignant cells and these macrophages is the MIF:CD74 interaction axis. The activation of CD74 receptors in Mac_LA may contribute to the activation of an anti-inflammatory profile^61,62^. In mouse models of sarcoma, colorectal, BRCA, and gynecological cancers TREM2 deletion or blockade with a monoclonal antibody has been shown to reduce tumor growth, enhance antitumor CD8^+^ T cell responses, including the effectiveness of anti–PD-1 treatment, and modified the TAM landscape^60,63^. Our findings strongly support the notion that distinct TAM phenotypes expressing TREM2 can coexist in the tumor microenvironment, exerting different roles, and contributing to the unclear dual function of TREM2 in cancer prognosis^64^. Phase I clinical trial has been leading in advanced refractory tumors using specific TREM2 mAb against tumor associated macrophages expressing TREM2^65^. The complex and context-dependent function of TREM2-expressing macrophages in the tumor microenvironment underscores the need for further research to fully understand its contribution to cancer progression and its potential as a therapeutic target. The identification of distinct TAM phenotypes expressing TREM2 and their differential impact on cancer prognosis opens new avenues for precision medicine approaches and personalized treatment strategies.

In addition to *TREM2-*expressing macrophages, we also investigated the clinical impact of RTM_IM, a subpopulation expressing high levels of folate receptor β (FRβ), which is encoded by the *FOLR2* gene and serves as a marker for resident macrophages in the TME. High levels of this subpopulation are associated with a poor prognosis in TNBC and HGSOC, two very lethal tumor types, although a beneficial role has been seen for other breast cancer subtypes^8^, demonstrating its importance as a target for new immunotherapies in a context-dependent maner^66–68^ likely through the induction of PD-1 and CD8^+^ T cell exhaustion^69^.

In addition, RTM_IM expresses of β1 integrins (ITGB1), Insulin-like Growth Factor 1 (IGF1), and Tumor necrosis factor (TNF) that are involved in interaction with the receptor for activated C kinase (RACK1). These pathways are involved, in general, in therapy resistance and cell progression^69–71^. Our results also show the potential of utilizing FOLR2 as a biomarker for patient stratification and prognosis in these tumor types, in line with other studies^8,63,72,73^. Several trials involving targeting FOLR2 are currently ongoing, reflecting the current level of research interest using this approach^74,75^.

In conclusion, we have characterized in high-resolution the landscape of MDCs in solid tumors by effectively integrating a large dataset through a pipeline that uses cluster-specific gene expression analysis to infer cell type, distribution, function, ontogeny, phenotype, cellular interactions, and disease associations. This comprehensive reference atlas serves as a resource for the scientific community and provides novel insights into the identities and characteristics of mononuclear phagocytes in the TME, presenting new avenues for understanding and manipulating their behavior in cancer. Our findings also demonstrate that the reprogramming of TAMs is highly dependent on the microenvironment, leading to varied TAM expression profiles across different cancer types, with distinct clinical impacts. This highlights the importance of understanding the specificities of subpopulations/states in the TME context. By shedding light onto the intricate landscape of mononuclear phagocytes in the TME, our work pushes the development of more effective and targeted therapeutic interventions against cancer.

## Methods

### Data download and pre-processing

Pre-processed scRNA-Seq data (Table S1) from patients with breast cancer (GSE114727), hepatocellular carcinoma (GSE140228, GSE125449), lung cancer (GSE127465), melanoma (GSE115979), ovarian cancer (GSE154600, GSE72056), uveal melanoma (GSE139829), skin (GSE130973) and metastasis from uveal melanoma and lung (GSE158803) samples were obtained from the public repository Gene Expression Omnibus (GEO) using the GEOquery Bioconductor package^76^. Additionally, datasets from lung, ovarian, and breast cancer were downloaded from Qian et al.^77^, on its platform (blueprint.lambrechtslab.org/). We downloaded datasets from the Human Lung Cell Atlas project^78^ on the Synapse platform (SYN21560407). Finally, the dataset containing PBMCs (10x Genomics standard) was downloaded from the company platform (support.10xgenomics.com/single-cell-gene-expression/datasets/1.1.0/pbmc3k).

Raw gene expression matrices were analyzed with the Seurat package (v4.0)^79^ for each study. We converted all gene symbols to Ensembl format (hg v38). Different filters of quality control were applied by technology: for 10x Genomics data the cutoff points were percentage of mitochondrial genes expressed <10, number of counts (nCounts) per cell >200, and ratio of nCounts by number of features (nFeatures) <5; for Smart-Seq2 were percentage of mitochondrial genes expressed <15, nCounts > 200, and ratio nCounts/nFeature < 1000; and for inDrop were percentage of mitochondrial genes expressed <15, nCount >200, and ratio nCounts/nFeature <1000. The remaining cells were submitted to the doublet removal step through the Scrublet package^80^. Through the doublet score histogram distribution, a cutoff value was determined for each library by manually setting the threshold (Optimal pK), eliminating cells with a higher probability of being doublets.

### Integration and cluster annotation

The datasets were concatenated with Scanpy (v1.7.2)^81^, and the 3000 highly variably expressed genes were selected using flavor = “Seurat”. The integration was performed with scVI (v0.6.8)^82^ from scvi-tools (v0.19.0)^83^. For training the variational autoencoder neural network, we used the following hyperparameters: n_latent=20, n_layers=4, dropout_rate=0.1. After training the scVI model and corrected for batch effects using “study” and “tissue origin” as co-variables in order to remove the technical bias, the clusters were calculated through the Leiden algorithm^84^ Different resolutions were evaluated, ranging from 0.3 to 2.0, and the resolution of 0.6 was selectedfor the annotation of broad cell types based on gene profiles. The latent space generated by scVI was then projected to a two dimenson space using the UMAP dimensional reduction method^85^. To determine the broad cell types, we use canonical markers (Supplementary Table 3). We found and removed three undefined/contaminated subsets with low quality (clusters 17, 19, and 29). After this first level of annotation, we applied DecontX^86^, from the celda package (v1.14.1), using default parameters, to estimate and remove ambient RNA contamination in individual droplets. For the other levels of annotation, we subset the broad cell types and then rerun highly variable genes and scVI models with the same parameters, except for batch correction, which was performed by each sample. We also excluding samples with a contribution of fewer than ten cells. In order to classify clusters in cell types or states, we apply DEG analysis by Wilcoxon rank-sum and MAST^87^, using logfc.threshold = 0.25, and min.pct = 0.1. The FDR method was used to adjust p-values. In addition, we evaluated the purity of the clusters through the ROGUE algorithm (v1.0)^88^ with default parameters.

Copy Number Variation inference

The copy number variation (CNV) analysis was used to distinguish epithelial from malignant cells. The CNV signal for individual cells was estimated using the inferCNVpy (v0.4.0) (github.com/icbi-lab/infercnvpy), inspired by the inferCNV^89^, by comparing all immune and stromal cells in the TME with epithelial cells, in order to classify malignant cells. Epithelial cells were classified according to the CNV signal: cells with a score lower than the median score (0.004 for all samples) were classified as “normal”, while those with a higher score were classified as “malignant”.

### *In silico* validation

In order to validate our findings, we correlated the populations we have defined with those from previous works. Another set of pre-processed scRNA-Seq data was downloaded^23,24^ (Supplementary Table 6). Those two datasets were selected because they are considered robust amounts of data, well-annotated regarding mononuclear phagocytes, and with a great diversity of tumor types, including those we have worked with in this paper. We used the same strategy and quality control filters that were used with our main datasets. The datasets were analyzed separately, using the authors’ annotations for macrophages, monocytes, and cDCs subpopulations. To perform the correlation, expression data was log-normalized using the NormalizedData function of the Seurat package (v4.0)^79^, followed by the AverageExpression function from the same package to determine the average expression of each gene for each subpopulation. Cor function^90^ from the stats R library was used to correlate the expression of all genes between subpopulations, and ggplot2^91^ was used to plot the correlation graphs. The purity of clusters was verified using ROGUE^88^ with default parameters.

### Metabolic characterization

We characterized the metabolic pathways of each cluster using Compass^92^. Cells were subsetted into clusters prior to running. The following parameters were used: --calc-metabolites --microcluster-size 10 --lambda 0. Raw reaction penalties were converted to reaction scores as described in the original publication, and reactions with zero variance were removed. Pairwise comparison was performed for each cluster (e.g. Cluster 1 vs. Cluster 2; Cluster 1 vs. Cluster n; etc) applying Wilcoxon’s test and calculating the Cohen’s d effect size between means. The FDR method was used to adjust p-values. Reactions with confidence scores bellow 4 (most confident) or with an adjusted p-value greater than 0.1 were removed. Cohen’s d median scores were calculated for each pairwise comparison and metabolic pathway. Those median scores were again aggregated by median comparing the median Cohen’s d value across all clusters to obtain a single value per cluster and metabolic pathway, which were z-scored to compare across macrophage clusters.

Reaction scores were calculated for each myeloid cluster as previously described and used to perform a pairwise comparison between all clusters. Reactions were filtered for adjusted p-value below 0.1 and a measure of the effect size was calculated using the median of Cohen’s values for each metabolic pathway. Next, Cohen’s medians for each metabolic pathway were summarized for each cluster to reflect the activity of the respective pathway in the cluster. The metabolic reactions Glycolysis/Gluconeogenesis, Citric Acid Cycle, NAD Metabolism, Oxidative Phosphorylation, Arginine and Proline Metabolism, Transport, Mitochondrial, Fatty Acid Oxidation, Fatty Acid Synthesis, Glycerophospholipid Metabolism, Glycosphingolipid Metabolism, Sphingolipid Metabolism are used to classified the macrophage based in the metabolic pathway.

### Trajectory analysis through pseudo-time

We employed Monocle 3 (1.3.1)^93^ with the UMAP generated by the integration with scVI as a “partition”. We set the parameter use_partition = FALSE of the function “learn graph” in order to create a linear trajectory, with default parameters. To order cells, the root used to infer the pseudotime was the Mono_CD14_FOS^-^ cluster, with default parameters. We also used Monocle 3 to perform gene module analysis based on graph autocorrelation to identify genes that co-vary over pseudotime.

### Scores and Pathway Enrichment

The top hundred genes identified in the DEGs analysis, sorted by the log fold change for each cluster, were selected to perform the functional enrichment analysis from the curated database Reactome (2021)^94^, through the clusterProfiler (v3.0)^95^. We considered significant pathways those with a p-adjusted ≤ 0.05. Some pathways were selected for plotting based on scientific experience of authors. The redundancy was removed from the enriched pathway set by selecting the pathway with the lowest p-value. All enriched pathways by subpopulations are available in Supplementary Table 5. The gene signature scores were computed by the AddModuleScore_UCell^96^ function. The gene set used for the scores is available in Supplementary Table 4.

### Cell-cell communication

To perform cell communication analyses, we only considered cells detected with more than 200 genes and genes with detection in more than 50 cells. The CellComm algorithm^97^ was used to infer communications between pairs of cell types including macrophages and tumors or lymphocytes. We used as a set of possible intercellular communications the 688 ligands and 857 receptors contained in the NicheNet database^98^ resulting in 12650 possible pairs. In communications between macrophages and tumors, we used cells classified as “MacAlv-like”, “Mac_Angio”, “Mac_Hypo”, “Mac_IFN”, “Mac_LA”, “Mac_Rec”, “Mac_AgPres”, “RTM-like_MT”, RTM_IFN”, “RTM_IM” and “RTM_LA” and breast, colorectal, liver, lung, melanoma and ovarian tumor cells. In communications between macrophages and lymphocytes, we used cells classified as macrophages “Mac_IFN”, “Mac_LA”, “Mac_Rec”, “RTM_IFN” and “RTM_IM” and lymphocytes “NKT”, “TCD4ex”, “TCD4reg”, “TCD8em”, “TCD8ex” and “TGD”. For consideration of intercellular communication, the expression of ligands and receptors should be greater than 0.25 normalized counts and show a p-value less than 0.01 in a permutation test of cluster annotation containing 1000 permutations. In order to display the main figures we applied the following filters: expression values of the ligands, receptors and mean of both higher than 1 and the p-value lower than 0.002. All interactions are shown in the Supplementary Table 8.

### Deconvolution analysis

To better understand the heterogeneity of the TME, as well as the cellular composition in each tumor type, we applied deconvolution methods to predict the abundance of cell types by modeling gene expression levels through bulk RNA-seq data. For these analyses, gene expression data (HTSeq counts), and clinical-pathological information from the database portal TCGA (http://cancergenome.nih.gov/) were downloaded using the TCGAbiolinks^99^ package of nine different tumor types, using the identifiers: OV (n=354 samples), LUAD (n =510), LUSC (n=496), UVM (n=77), SKCM (divided into Metastatic (n=385) and Primary (n= 103) tumor samples), COAD (n=454), READ (n=170), LIHC (n = 369) and BRCA (n = 1195) which comprises Luminal A (n= 569), Luminal B (n=210), HER2 (n=81) and TNBC (n= 188) subtypes. The single-cell signatures identified in this work was used as a reference using BayesPrism package^100^ (v2.0) to predict the relative frequency of cell types for each patient of each tumor type. We considered 47 cell signatures, including cell types identified in the first level of annotation, such as fibroblasts, endothelial cells, B lymphocytes, mast cells, pDC, malignant cells, and normal epithelial cells, and cell states/subtypes identified in other levels of clustering, including macrophages (n =12, except the Mac_Prolif subpopulation), monocytes (n = 6), cDC (n =8), T cells (n = 9), NK cells (n = 2), and neutrophil (n = 3) subpopulations. The gene expression counts matrix was filtered to only contain the 12,854 genes that make up the signatures of the cell types. Only cell types representing more than 1% of in each tumor type were considered. Ribosomal and mitochondrial genes with high expression were removed using the *cleanup.genes()* function. In addition, genes expressed in less than 5 cells were excluded. After that, to get more consistent results, we filtered for protein-coding genes and used the function *get.exp.stat()* to select differentially expressed genes between cell states of different cell types using the default parameters, except for the parameter *pseudo.count* (0.1 for 10x data and 10 for Smart-seq data). Finally, we set up the prism object using the *new.prism()* function for each tumor type by setting the following parameters:

*reference=**scRNAfiltered_matrix**,mixture=**bulk_matrix**,input.type=**“count.matrix”**,cell.type.labels = **celltype**, cell.state.labels = **states**, key=**“Malignant Cells”**, outlier.cut=**0.01**, outlier.fraction=**0.1***. And then running BayesPrism, *run.prism(prism = **prism object**, n.cores=**70**)*.

### Study design, ethical considerations, and patient selection

This retrospective study was approved by the Ethics in Human Research Committee of the Brazilian National Cancer Institute (INCA), Rio de Janeiro, Brazil, and was conducted following the Good Clinical Practice Guidelines. For the ovary cohort, 193 all women diagnosed with HGSOC at INCA between 2001 and 2017, regardless of adjuvant or neoadjuvant treatment, were included in our study. The TNBC cohort, previously described by our collaborators^101^consisted of 112 women who underwent neoadjuvant chemotherapy and thereafter curative surgery between 2010 and 2014. Clinical data regarding age at diagnosis, staging, surgery, histological subtype, chemotherapy, and survival were retrospectively obtained from the medical records. The two INCA cohorts are described in more detail in Supplementary Table 11.

### Immunohistochemistry

The blocks were cut with a microtome into fine slivers of 4-mm sections to form the tissue microarray which encompasses the most representative areas of greatest tumor cellularity in formalin-fixed paraffin-embedded tissue specimens. The immunohistochemistry was processed using a NOVOLINK™ Polymer Detection Systems (Leica), followed by 3,3′-Diaminobenzidine (DAB). Antigen retrieval was performed with the Trilogy (Cell Marque) reagent. The samples were immunostained for Ki67 (clone 30-9 at 1:7, Ventana- Roche), CD8 (clone SP57 at 1:7, Ventana- Roche), PD-1 (clone NAT105, Cell Marque, diluted 1:100), CD68 (clone KP-1 at 1:20, Ventana- Roche), FOLR2 (OTI4G6 at 1:200, ThermoFisher) and TREM2 (clone D814C, 1:100, Cell Signaling Technology). For double staining, TREM2 was combined with anti-CD68. Briefly, after completing the first immune reaction (TREM2) with DAB, the second immune reaction (CD86) was visualized using MACH4 MR-AP (Biocare Medical), using green substract (Polydetector HRP Green Substrate-chromogen Bio-SB - ref BSB0129t. The entire analysis was carried out at the Division of Pathology - INCA by experienced pathologists.

For the HGSOC cohort, tissue microarrays were assessed by determining the percent of immune cells and their intratumoral distribution with the markers TREM2 and CD68 (low = 0 cells, high ≥ 10% cell), FOLR2 (low =< 25%, high > 75% cells), CD68 (low = 0 cells, high ≥ 10% cells), PD-1 (low = 0-1% cells, high 3-30% cells), CD8 (low = 0-1% cell, high >=3% cells).For the TNBC cohort, FOLR2 and the TREM2 were acessed by ImageJusing the IHC Profiler ImageJ Plugin^102^. The separation of the color layers was based on hematoxylin and 3,3′-Diaminobenzidine. For each patient, we calculated the average of the percentage values obtained from ImageJ for all the spots. Then, the score for each average value was calculated based on the IHC Profiler tutorial, as follows: percentage of high positive (4 x % area of high positive staining), positive (3 x % area of positive staining), and low positive (2 x % area of low positive staining) staining. Finally, the score per sample was defined as the higher value among them.

### Survival and statistical analysis

To compare means (scores and clinical impact), statistical analyzes were performed using the Kruskal-Wallis method for global comparisons, followed by the Wilcoxon test for paired comparisons. The Kaplan-Meier method was used to visualize the difference between the curves of groups through the Survival (v3.5) and Survminer (v0.4.9) packages^103,104^. The surv_cutpoint function was applied to split samples in high and low groups. The hazard ratios were calculated using the univariate and multivariate Cox proportional-hazards model. All statistical significance testing in this study was two-sided. To control for multiple hypothesis testing, we applied the Benjamini–Hochberg method to correct *P* values and the FDR *Q* values were calculated. Results were considered statistically significant at *P* value or FDR *Q* value of <0.05.

Regarding the clinical parameters used, the clinical annotation of TCGA dataset, and the times calculated for OS and PFS were downloaded from the PanCanAtlas study. For the INCA cohort, OS was calculated considering the day from the diagnosis to the date of death of any cause if the patient was known to be alive on the last day of data collection. The PFS was calculated from the last chemotherapy data or the last registration to recurrence data if the patient was known to be alive or the date of death. The Kaplan-Meier method was used to visualize the difference between the curves of the high and low groups. The statistical analyses were conducted using R project version 3.5.3

## Data Availability

All data used in this work have been listed in Supplementary Tables 1 and 6, including accession numbers. The integrated dataset is publicly available and can be downloaded via CELL×GENE | Collections through the following link: cellxgene.cziscience.com/collections/3f7c572c-cd73-4b51-a313-207c7f20f188.

## Code Availability

The code for reproducing the analyses and panels of this paper is available in the GitHub repository: github.com/bioinformatics-inca/MyeloidDerivedCells_in_TME.

## Acknowledgements

We would like to thank the Bioinformatics Core Facility, the National Tumor Bank, and the Division of Pathology, all three at the Brazilian National Cancer Institute (INCA-RJ), for their support. This work was supported by INCA/MS fellowships, Chan Zuckerberg Initiative - Inflammation Network (MAM, MB, and PMMMV), and Fundação de Amparo à Pesquisa do Estado do Rio de Janeiro (FAPERJ - E-26/201.322/2022 (272260), E-26/211.648/2021 (269502), and E-26/210.302/2022 (270388) for M.B). This publication is part of the Human Cell Atlas – www.humancellatlas.org/publications.

## Author contributions

Conceptualization: MB, MAM, PMMMV

Single-cell integration, and cells annotation: GRM, GRG

Pathways analysis: MAP, NET

Pseudo-time analysis: GRG In silico validation: LOS

Cell-Cell communication analysis: MF, MMD, LOS, ELR

Deconvolution and clinical association: CET, NGT, GRM

Immunohistochemistry analysis: NGT, NCB, FCM, FRR, CET, JGVC, AFS

Cohort obtention: CBPC, ACM, JLS, NGT, NET

Funding acquisition: MB, MAM, PMMMV

Supervision: MB, MAM, PMMMV

Writing – original draft: GRG, GRM, NGT, CET, LOS, NET

Writing – review & editing: MB, MAM, PMMMV, ACM, ELR

