## Supplementary Figures for "Identification of novel myeloid-derived cell states with implication in cancer outcome"

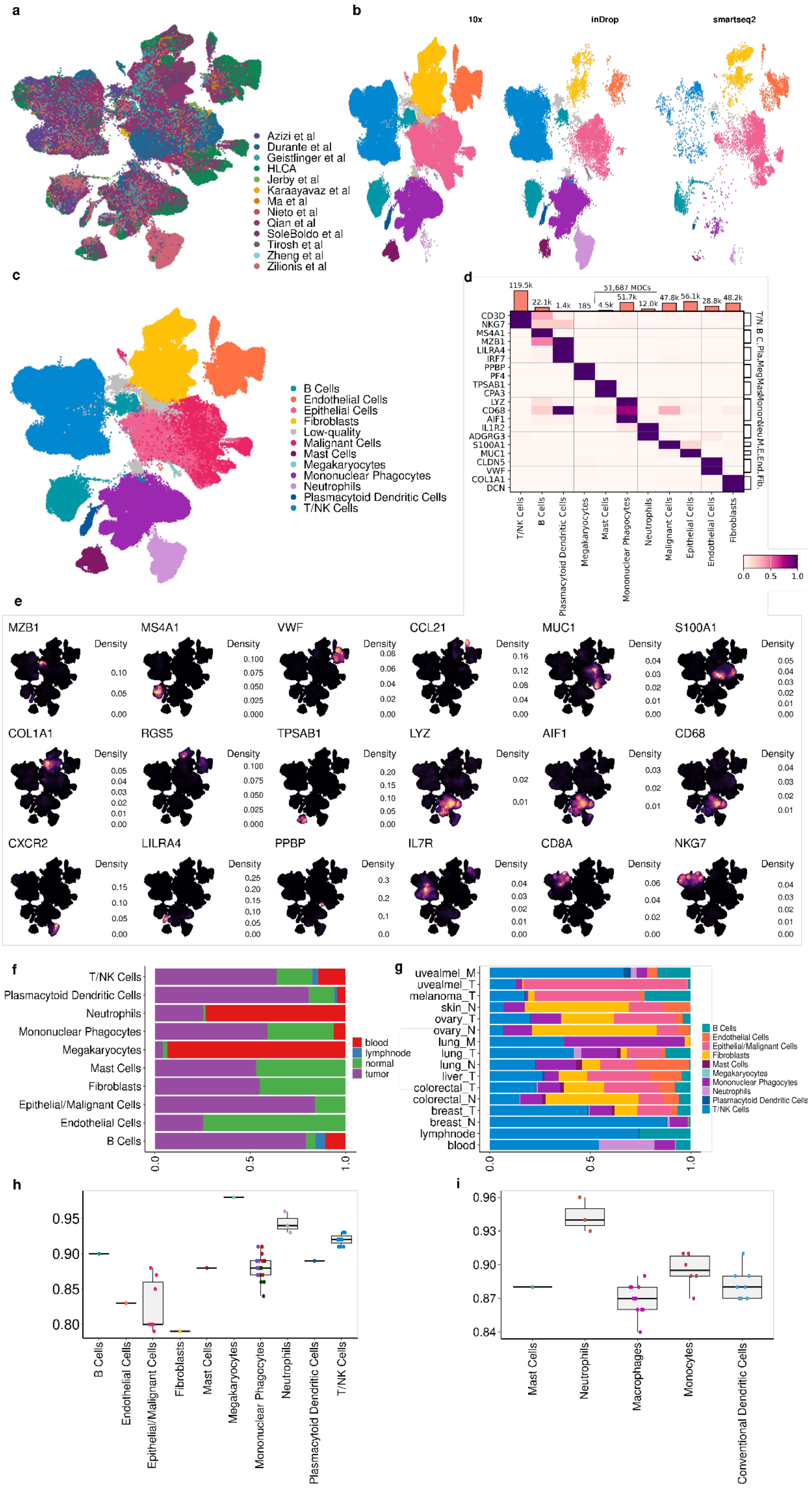

a
Azizi et al
Durante et al
Geistlinger et al
HLCA
Jerby et al
Karaayavaz et al
Ma et al
Nieto et al
Qian et al
SoleBoldo et al
Tirosh et al
Zheng et al
Zilionis et al
b
10x
inDrop
smartseq2
c
B Cells
Endothelial Cells
Epithelial Cells
Fibroblasts
Low-quality
Malignant Cells
Mast Cells
Megakaryocytes
Mononuclear Phagocytes
Neutrophils
Plasmacytoid Dendritic Cells
T/NK Cells
d
119.5k
22.1k
1.4k
185
51,687 MDCs
4.5k
51.7k
12.0k
47.8k
56.1k
28.8k
48.2k
CD3D
NKG7
MS4A1
MZB1
LILRA4
IRF7
PPBP
PF4
TPSAB1
CPA3
LYZ
CD68
AIF1
IL1R2
ADGRG3
S100A1
MUC1
CLDN5
VWF
COL1A1
DCN
T/N B C. Pla. Meg Mas Monon Neu M. E. End. Fib.
T/NK Cells
B Cells
Plasmacytoid Dendritic Cells
Megakaryocytes
Mast Cells
Mononuclear Phagocytes
Neutrophils
Malignant Cells
Epithelial Cells
Endothelial Cells
Fibroblasts
0.0
0.5
1.0
e
MZB1
MS4A1
VWF
CCL21
MUC1
S100A1
COL1A1
RGS5
TPSAB1
LYZ
AIF1
CD68
CXCR2
LILRA4
PPBP
IL7R
CD8A
NKG7
Density
f
T/NK Cells
Plasmacytoid Dendritic Cells
Neutrophils
Mononuclear Phagocytes
Megakaryocytes
Mast Cells
Fibroblasts
Epithelial/Malignant Cells
Endothelial Cells
B Cells
blood
lymphnode
normal
tumor
0.0
0.5
1.0
g
uvealmel_M
uvealmel_T
melanoma_T
skin_N
ovary_T
ovary_N
lung_M
lung_T
lung_N
liver_T
colorectal_T
colorectal_N
breast_T
breast_N
lymphnode
blood
B Cells
Endothelial Cells
Epithelial/Malignant Cells
Fibroblasts
Mast Cells
Megakaryocytes
Mononuclear Phagocytes
Neutrophils
Plasmacytoid Dendritic Cells
T/NK Cells
h
0.95
0.90
0.85
0.80
B Cells
Endothelial Cells
Epithelial/Malignant Cells
Fibroblasts
Mast Cells
Megakaryocytes
Mononuclear Phagocytes
Neutrophils
Plasmacytoid Dendritic Cells
T/NK Cells
i
0.96
0.93
0.90
0.87
0.84
Mast Cells
Neutrophils
Macrophages
Monocytes
Conventional Dendritic Cells

**Supplementary Figure 1. Integrative profiling of scRNA-seq datasets across eight human cancer types. a** UMAP representing all cell populations and their distribution across the 13 different datasets (Supplementary Table 1). **b** UMAP split by technologies, showing the integration by cell similarities. **c** UMAP color-coded by the eleven broad cell types and the low-quality cluster. B cells (n = 22,102) were identified by the expression of *MZB1* and *MS4A1*; Endothelial cells (n = 28,775) were marked by *VWF* and *CCL21*; *MUC* and *S100A1* expression allowed the identification of normal (n = 56,060) and malignant (n = 47,827) epithelial cells, which were further distinguished by the amount of copy number variation events observed in the malignant cells. Fibroblasts (n = 48,159) were identified by *COL1A1* and *DCN*; Mast cells (n = 4,539), identified by *TPSAB1* and *CPA3*; Mononuclear Phagocytes were positive for *LYZ*, *AIF1*, and *CD68*; Neutrophils (n = 12,033), marked by *CXCR2* and *FCGR3B*; Plasmacytoid Dendritic Cells (pDC) (n = 1,365) by *LILRA4* and *IRF7*; megakaryocytes (n = 185) marked by *PPBP* expression; and T and Natural Killer (NK) cells (n = 119,472) identified by the expression of *CD3E*, *CD4/IL7R* or *CD8A,* or *NKG7*; **d** Heatmap showing the marker genes across the broad cell types and the number of cells per type. The color scale reflects the log10-normalized gene expression range. **e** Density plots highlight the gene marker expression of each cell type. **f** Barplot showing the proportion of sample type origin by cell types. **g** Barplot showing the proportion of broad cell types across conditions and sample types (normal (_N), primary tumor (_T), and tumor metastasis (_M)). **h-i** ROGUE values for nine identified broad cell types **(h)** and for MDCs **(i)**. Each point represents an individual cluster. The center line indicates the median ROUGE value.

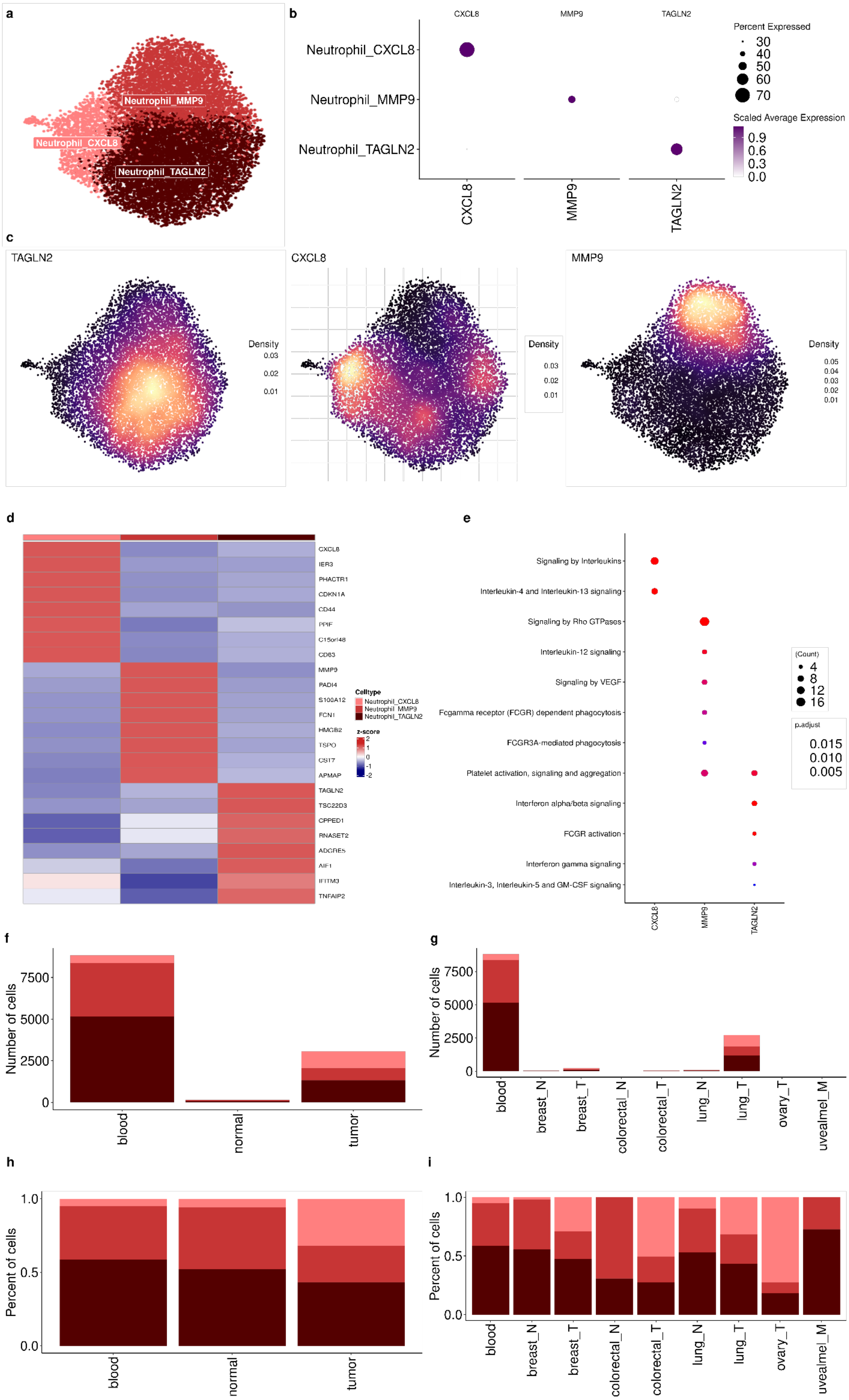
a
Neutrophil_MMP9
Neutrophil_CXCL8
Neutrophil_TAGLN2
b
CXCL8
MMP9
TAGLN2
Neutrophil_CXCL8
Neutrophil_MMP9
Neutrophil_TAGLN2
Percent Expressed
30
40
50
60
70
Scaled Average Expression
0.9
0.6
0.3
0.0
c
TAGLN2
CXCL8
MMP9
Density
d
CXCL8
IER3
PHACTR1
CDKN1A
CD44
PPIF
C15orf48
CD63
MMP9
PADI4
S100A12
FCN1
HMGB2
TSPO
CST7
APMAP
TAGLN2
TSC22D3
CPPED1
RNASET2
ADGRE5
AIF1
IFITM3
TNFAIP2
Celltype
Neutrophil_CXCL8
Neutrophil_MMP9
Neutrophil_TAGLN2
z-score
e
Signaling by Interleukins
Interleukin-4 and Interleukin-13 signaling
Signaling by Rho GTPases
Interleukin-12 signaling
Signaling by VEGF
Fcgamma receptor (FCGR) dependent phagocytosis
FCGR3A-mediated phagocytosis
Platelet activation, signaling and aggregation
Interferon alpha/beta signaling
FCGR activation
Interferon gamma signaling
Interleukin-3, Interleukin-5 and GM-CSF signaling
(Count)
4
8
12
16
p.adjust
0.015
0.010
0.005
f
Number of cells
blood
normal
tumor
g
Number of cells
blood
breast_N
breast_T
colorectal_N
colorectal_T
lung_N
lung_T
ovary_T
uvealmel_M
h
Percent of cells
blood
normal
tumor
i
Percent of cells
blood
breast_N
breast_T
colorectal_N
colorectal_T
lung_N
lung_T
ovary_T
uvealmel_M

**Supplementary Figure 2. Neutrophils across tumor samples. a** UMAP of neutrophils colored by distinct subpopulations. **b** Dotplot showing the mean expression of gene markers from neutrophil subpopulations. Dot size indicates the fraction of expressing cells, colored based on scaled expression levels. **c** Density plots highlight the gene marker expression for each cell state. **d** Heatmap showing the marker genes across neutrophils subpopulation. **e** Enrichment pathways analysis of neutrophil subpopulations using Reactome database. The size of each circle represents the number of genes belonging to a given pathway and such circles are colored by p-adjust. **f-i** Barplot showing the number of neutrophils across sample **(f-h)** and conditions **(g-i)**.

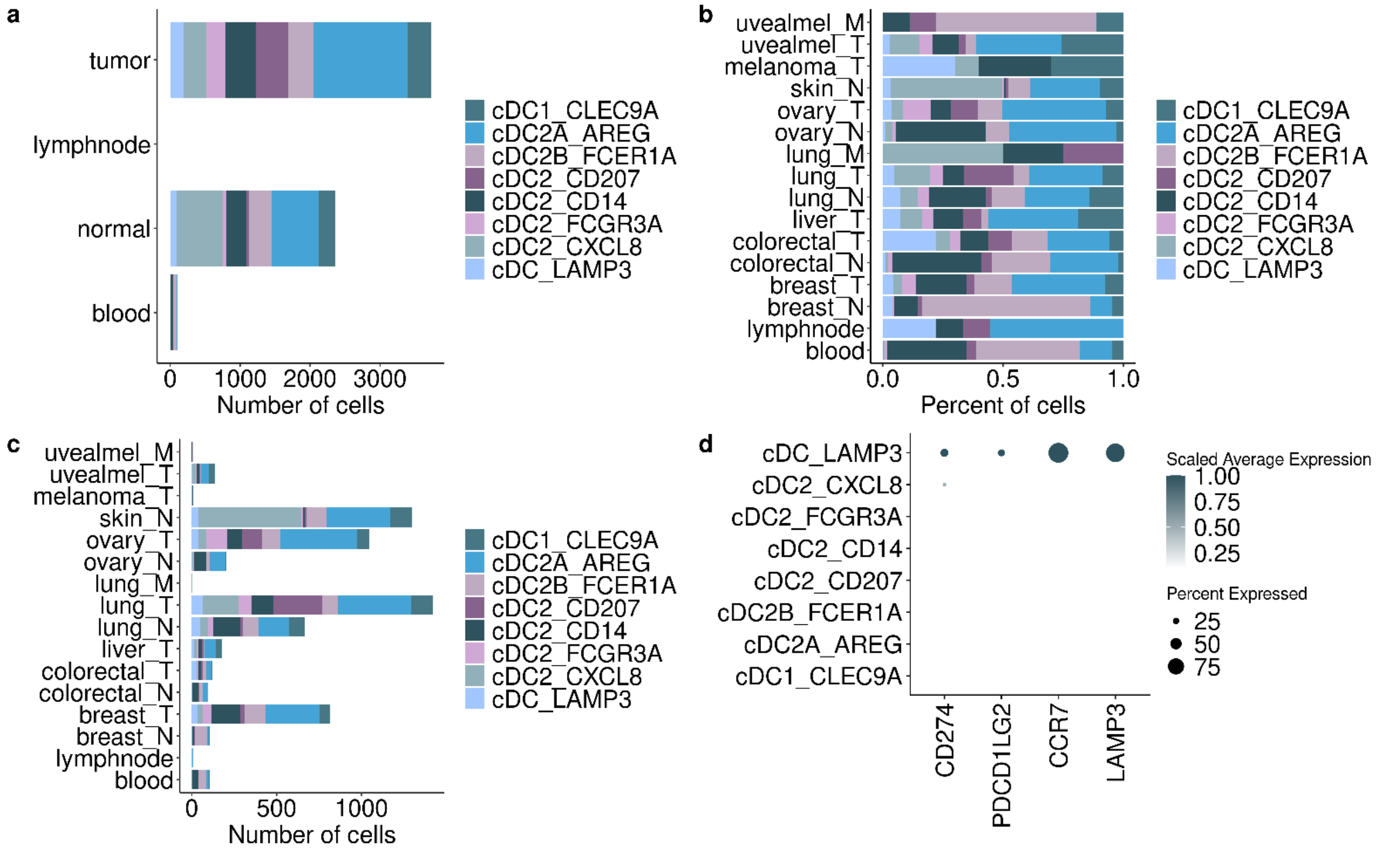

a
tumor
lymphnode
normal
blood
0
1000
2000
3000
Number of cells
cDC1_CLEC9A
cDC2A_AREG
cDC2B_FCER1A
cDC2_CD207
cDC2_CD14
cDC2_FCGR3A
cDC2_CXCL8
cDC_LAMP3
b
uvealmel_M
uvealmel_T
melanoma_T
skin_N
ovary_T
ovary_N
lung_M
lung_T
lung_N
liver_T
colorectal_T
colorectal_N
breast_T
breast_N
lymphnode
blood
0.0
0.5
1.0
Percent of cells
c
0
500
1000
Number of cells
d
Scaled Average Expression
1.00
0.75
0.50
0.25
Percent Expressed
25
50
75
CD274
PDCD1LG2
CCR7
LAMP3

**Supplementary Figure 3. Distribution of cDCs across samples. a-c** Barplot showing the distribution of cDC subpopulations across sample types **(a)** and conditions **(b-c)**. **d** Dotplot showing the expression of immunosuppressive genes in cDC_LAMP3. Dot size indicates the percent of expressing cells, and the dot color is the scaled average expression.

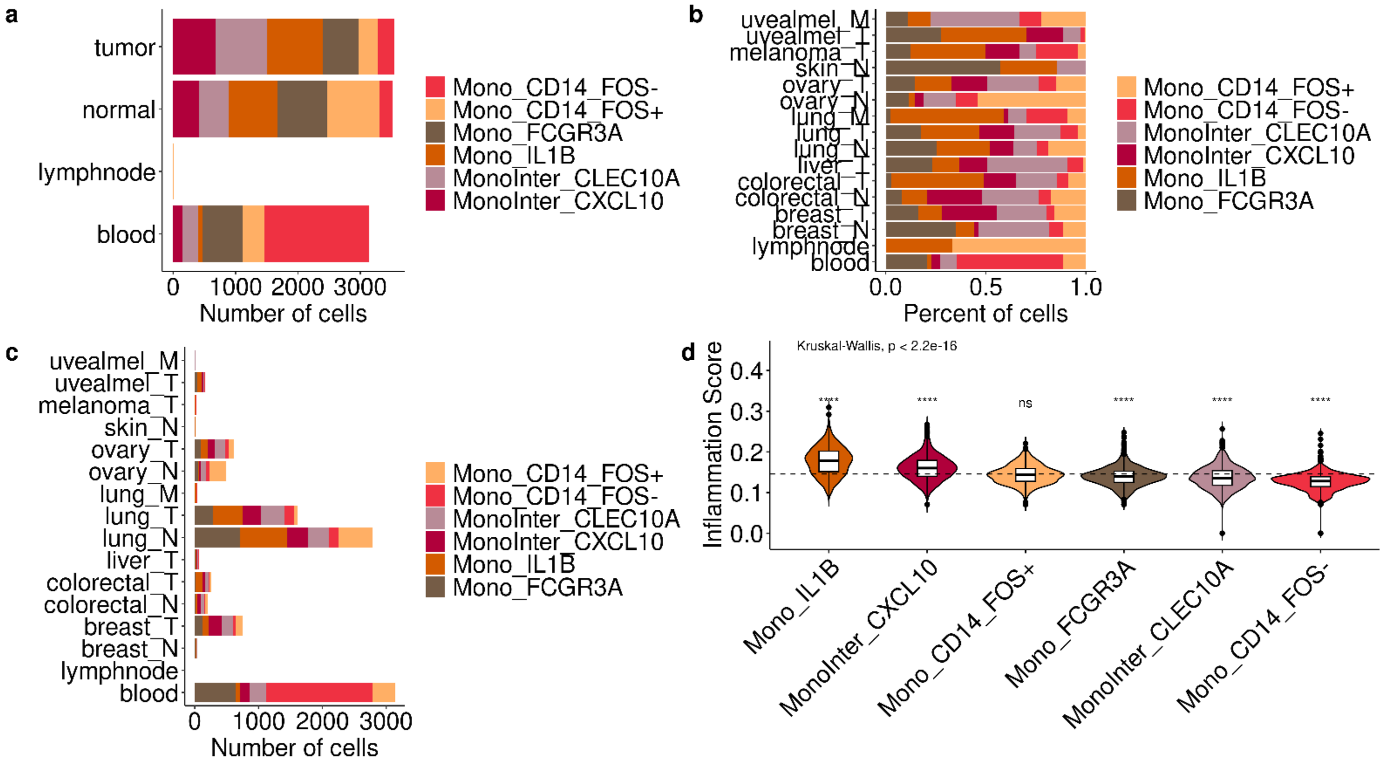
a
tumor
normal
lymphnode
blood
0
1000
2000
3000
Number of cells
Mono_CD14_FOS-
Mono_CD14_FOS+
Mono_FCGR3A
Mono_IL1B
MonoInter_CLEC10A
MonoInter_CXCL10
b
uvealmel_M
uvealmel_T
melanoma_T
skin_N
ovary_T
ovary_N
lung_M
lung_T
lung_N
liver_T
colorectal_T
colorectal_N
breast_T
breast_N
lymphnode
blood
0.0
0.5
1.0
Percent of cells
c
Number of cells
d
Kruskal-Wallis, p < 2.2e-16
Inflammation Score
0.0
0.1
0.2
0.3
0.4
****
ns
Mono_IL1B
MonoInter_CXCL10
Mono_CD14_FOS+
Mono_FCGR3A
MonoInter_CLEC10A
Mono_CD14_FOS-

**Supplementary Figure 4. Monocytes' distribution across samples.** **a-c** Barplot showing the distribution of monocyte subpopulations across sample types **(a)** and conditions **(b-c)**. **d** Violin plot demonstrating the inflammation score among Monocytes subpopulations (Supplementary Table 4). Boxes span the first to third quartiles; the horizontal line inside the boxes represents the median black and dots represent outlier samples in each group. For statistical significance, Kruskall–Wallis test followed by Wilcoxon was performed to compare each of the ten groups against "all" (i.e. base-mean). Ns non-significant. *$p < 0.05$; **$p < 0.01$; ***$p < 0.001$; ****$p < 0.0001$.

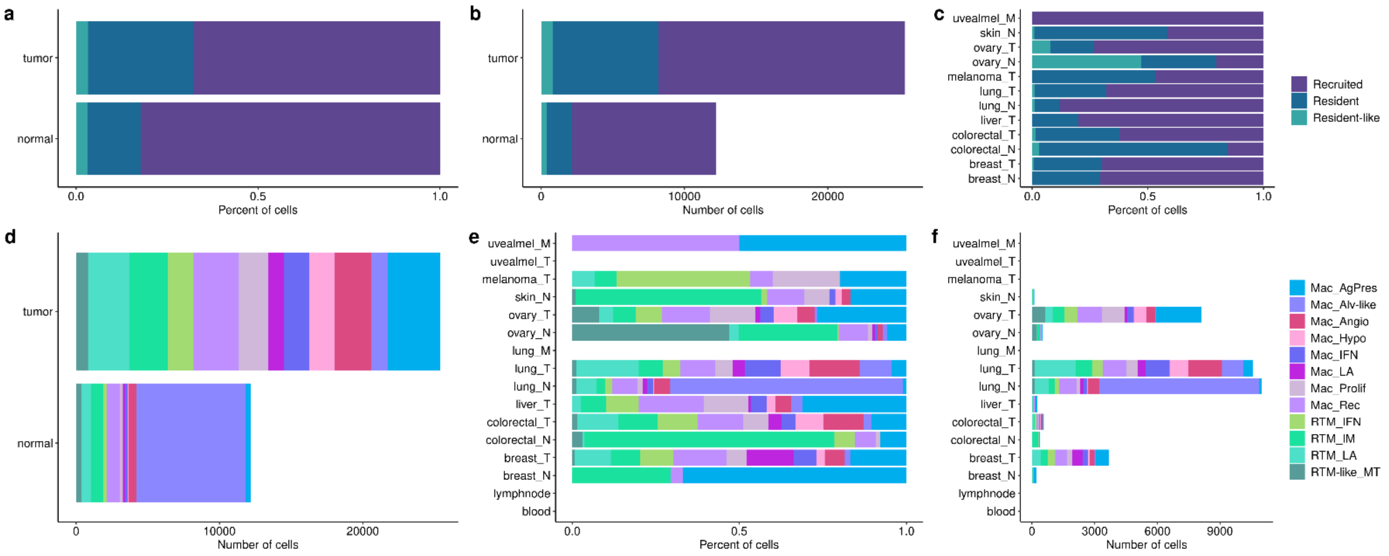

a
tumor
normal
0.0
0.5
1.0
Percent of cells
b
0
10000
20000
Number of cells
c
uvealmel_M
skin_N
ovary_T
ovary_N
melanoma_T
lung_T
lung_N
liver_T
colorectal_T
colorectal_N
breast_T
breast_N
Recruited
Resident
Resident-like
d
e
f
uvealmel_T
lung_M
lymphnode
blood
3000
6000
9000
Mac_AgPres
Mac_Alv-like
Mac_Angio
Mac_Hypo
Mac_IFN
Mac_LA
Mac_Prolif
Mac_Rec
RTM_IFN
RTM_IM
RTM_LA
RTM-like_MT

**Supplementary Figure 5. Distribution of macrophage subpopulations across tumor samples. a-c** Barplot showing the distribution of recruited and resident-tissue macrophages across samples **(a-b)** and conditions **(c)**. **d-f** Barplot showing the distribution of macrophage subpopulations across samples **(d)** and conditions **(e-f)**.

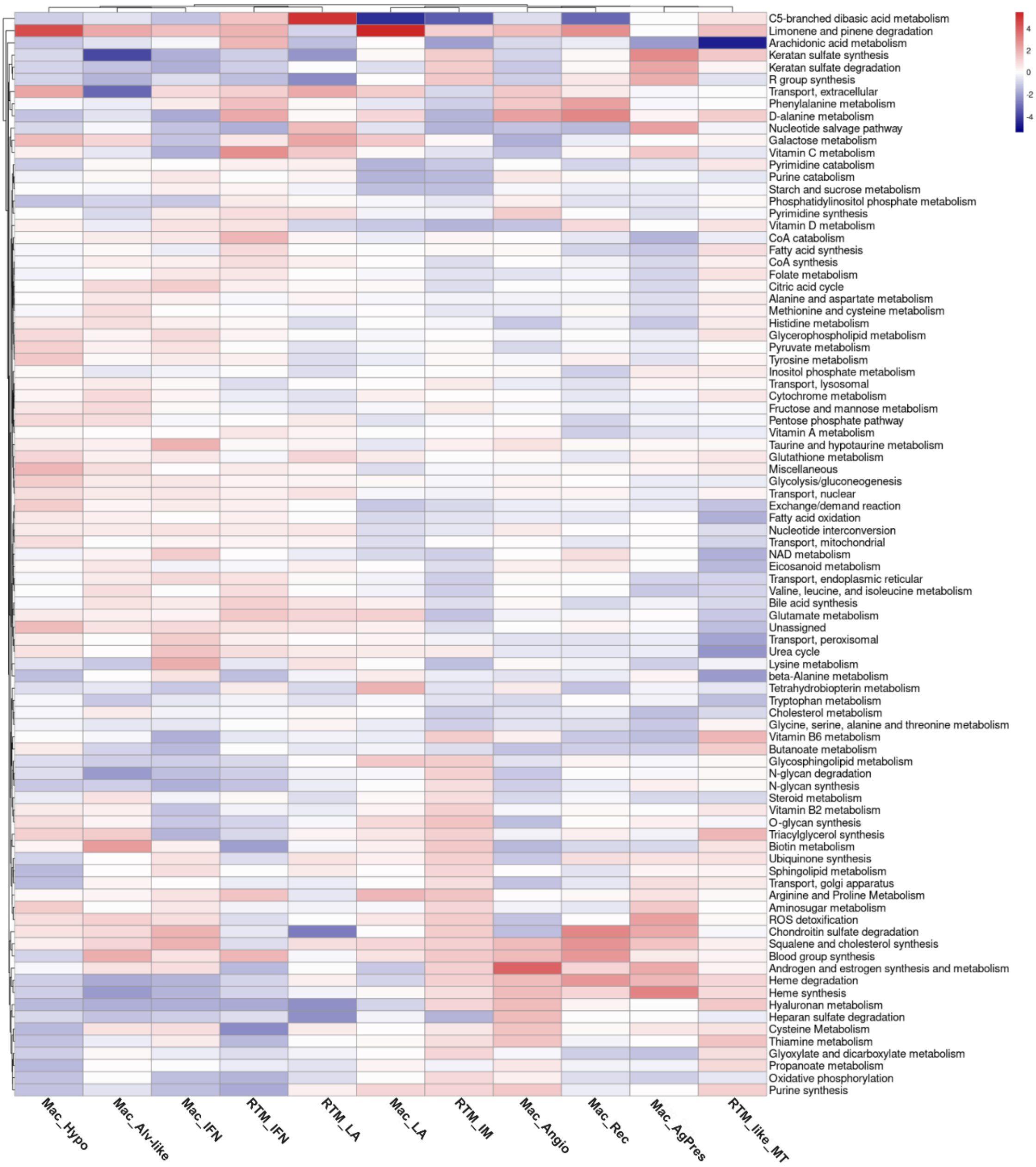

C5-branched dibasic acid metabolism
Limonene and pinene degradation
Arachidonic acid metabolism
Keratan sulfate synthesis
Keratan sulfate degradation
R group synthesis
Transport, extracellular
Phenylalanine metabolism
D-alanine metabolism
Nucleotide salvage pathway
Galactose metabolism
Vitamin C metabolism
Pyrimidine catabolism
Purine catabolism
Starch and sucrose metabolism
Phosphatidylinositol phosphate metabolism
Pyrimidine synthesis
Vitamin D metabolism
CoA catabolism
Fatty acid synthesis
CoA synthesis
Folate metabolism
Citric acid cycle
Alanine and aspartate metabolism
Methionine and cysteine metabolism
Histidine metabolism
Glycerophospholipid metabolism
Pyruvate metabolism
Tyrosine metabolism
Inositol phosphate metabolism
Transport, lysosomal
Cytochrome metabolism
Fructose and mannose metabolism
Pentose phosphate pathway
Vitamin A metabolism
Taurine and hypotaurine metabolism
Glutathione metabolism
Miscellaneous
Glycolysis/gluconeogenesis
Transport, nuclear
Exchange/demand reaction
Fatty acid oxidation
Nucleotide interconversion
Transport, mitochondrial
NAD metabolism
Eicosanoid metabolism
Transport, endoplasmic reticular
Valine, leucine, and isoleucine metabolism
Bile acid synthesis
Glutamate metabolism
Unassigned
Transport, peroxisomal
Urea cycle
Lysine metabolism
beta-Alanine metabolism
Tetrahydrobiopterin metabolism
Tryptophan metabolism
Cholesterol metabolism
Glycine, serine, alanine and threonine metabolism
Vitamin B6 metabolism
Butanoate metabolism
Glycosphingolipid metabolism
N-glycan degradation
N-glycan synthesis
Steroid metabolism
Vitamin B2 metabolism
O-glycan synthesis
Triacylglycerol synthesis
Biotin metabolism
Ubiquinone synthesis
Sphingolipid metabolism
Transport, golgi apparatus
Arginine and Proline Metabolism
Aminosugar metabolism
ROS detoxification
Chondroitin sulfate degradation
Squalene and cholesterol synthesis
Blood group synthesis
Androgen and estrogen synthesis and metabolism
Heme degradation
Heme synthesis
Hyaluronan metabolism
Heparan sulfate degradation
Cysteine Metabolism
Thiamine metabolism
Glyoxylate and dicarboxylate metabolism
Propanoate metabolism
Oxidative phosphorylation
Purine synthesis
Mac_Hypo
Mac_Alv-like
Mac_IFN
RTM_IFN
RTM_LA
Mac_LA
RTM_IM
Mac_Angio
Mac_Rec
Mac_AgPres
RTM_like_MT
4
2
0
-2
-4

**Supplementary Figure 6. Overview of the main metabolic pathways in the macrophage subpopulations.** Heatmap showing the main metabolism signatures summarized for each macrophage subpopulation/state. The color scale represents the median effect size (Cohen's d, the difference between two means) for an increase (red) or decrease (blue) in a given metabolic pathway.

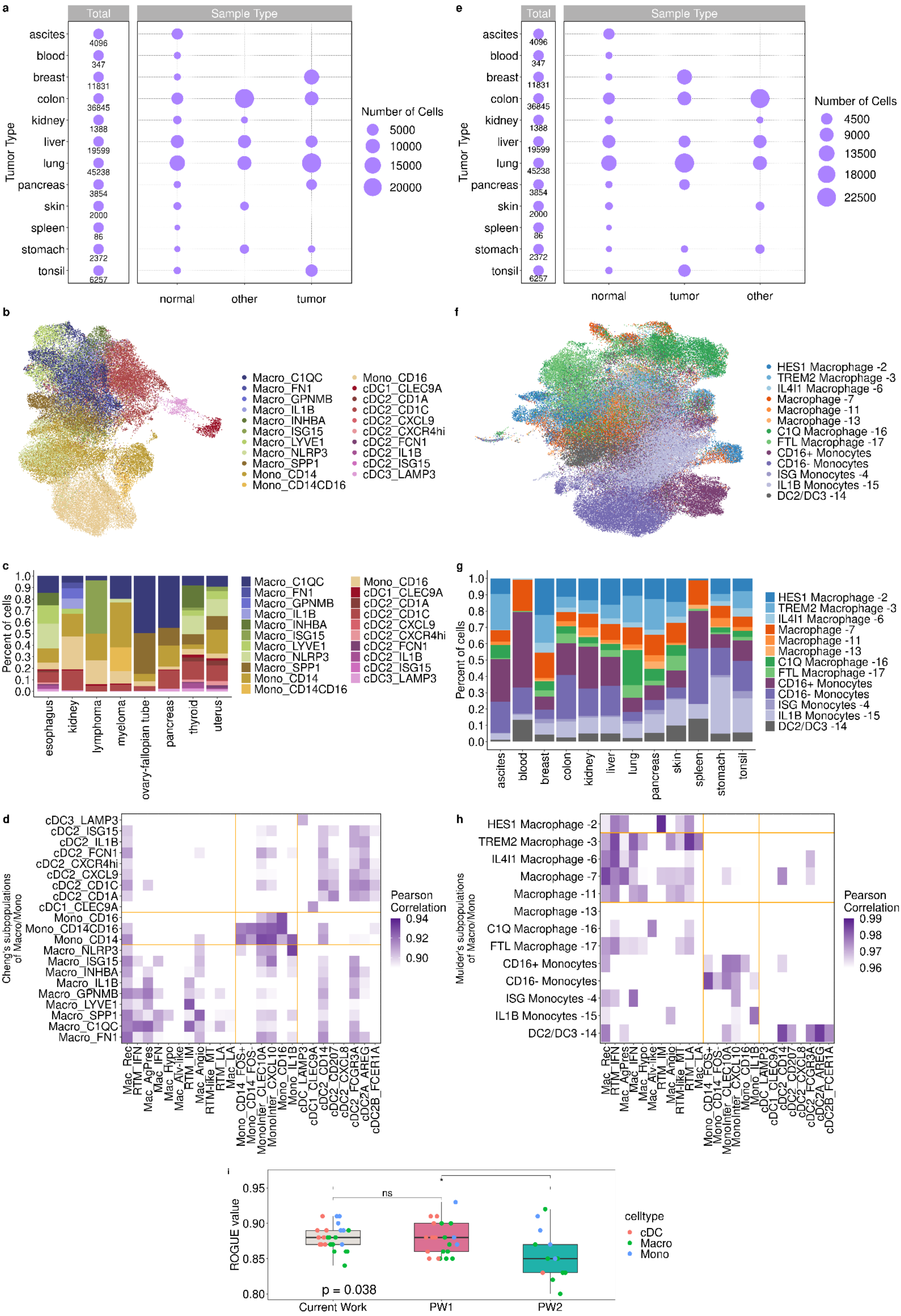
a
Total
Sample Type
ascites
blood
breast
colon
kidney
liver
lung
pancreas
skin
spleen
stomach
tonsil
Tumor Type
4096
347
11831
36845
1388
19599
45238
3854
2000
86
2372
6257
normal
other
tumor
Number of Cells
5000
10000
15000
20000
e
Total
Sample Type
Number of Cells
4500
9000
13500
18000
22500
normal
tumor
other
b
Macro_C1QC
Macro_FN1
Macro_GPNMB
Macro_IL1B
Macro_INHBA
Macro_ISG15
Macro_LYVE1
Macro_NLRP3
Macro_SPP1
Mono_CD14
Mono_CD14CD16
Mono_CD16
cDC1_CLEC9A
cDC2_CD1A
cDC2_CD1C
cDC2_CXCL9
cDC2_CXCR4hi
cDC2_FCN1
cDC2_IL1B
cDC2_ISG15
cDC3_LAMP3
f
HES1 Macrophage -2
TREM2 Macrophage -3
IL4I1 Macrophage -6
Macrophage -7
Macrophage -11
Macrophage -13
C1Q Macrophage -16
FTL Macrophage -17
CD16+ Monocytes
CD16- Monocytes
ISG Monocytes -4
IL1B Monocytes -15
DC2/DC3 -14
c
Percent of cells
esophagus
kidney
lymphoma
myeloma
ovary-fallopian tube
pancreas
thyroid
uterus
g
Percent of cells
ascites
blood
breast
colon
kidney
liver
lung
pancreas
skin
spleen
stomach
tonsil
d
Cheng's subpopulations of Macro/Mono
Pearson Correlation
0.94
0.92
0.90
Mac_Rec
RTM_IFN
Mac_AgPres
Mac_IFN
Mac_Hypo
Mac_Alv-like
RTM_IM
Mac_Angio
RTM-like_MT
RTM_LA
Mac_LA
Mono_CD14_FOS+
Mono_CD14_FOS-
MonoInter_CLEC10A
MonoInter_CXCL10
Mono_CD16
Mono_IL1B
cDC_LAMP3
cDC1_CLEC9A
cDC2_CD14
cDC2_CD207
cDC2_CXCL8
cDC2_FCGR3A
cDC2A_AREG
cDC2B_FCER1A
h
Mulder's subpopulations of Macro/Mono
Pearson Correlation
0.99
0.98
0.97
0.96
i
ROGUE value
0.95
0.90
0.85
0.80
ns
*
p = 0.038
Current Work
PW1
PW2
celltype
cDC
Macro
Mono

**Supplementary Figure 7. Validation of robustness and novelty: comparing our identified populations with literature datasets.** **a** Dotplot showing the number of cells for each tumor and sample type in Cheng and colleagues (2021) dataset; **b** UMAP showing the mononuclear phagocytes subpopulations described in their work; **c-d** Stacked Barplot showing the distribution of mononuclear phagocytes subpopulations by each tumor type **(c)**, and the Pearson correlation between mononuclear phagocytes subpopulations described in this work and the aforementioned **(d)**. **e** Dotplot showing the number of cells for each tumor and sample type in Mulder and colleagues (2021) dataset; **f** UMAP showing the mononuclear phagocytes subpopulations described in their work; **g-h** Stacked Barplot showing the distribution of mononuclear phagocytes subpopulations by each tumor type **g**, and the Pearson correlation between mononuclear phagocytes subpopulations described in this work and the aforementioned (**h)**. **i** Boxplot showing a comparison of ROGUE values between mononuclear phagocyte populations described by this work, Cheng and colleagues (2021) work (PW1), and Mulder and colleagues (2021) work (PW2). Statistical significance for values of $p < 0.05$.

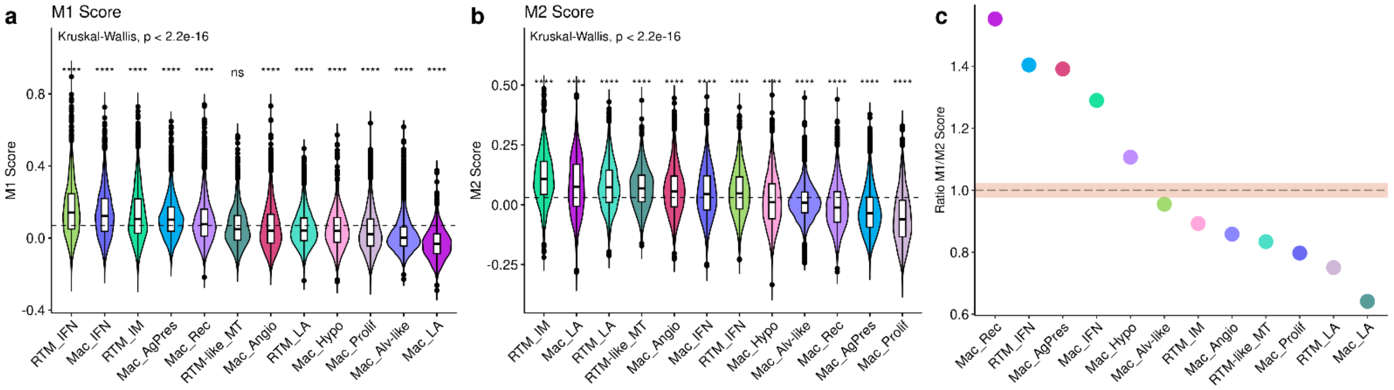
a
M1 Score
Kruskal-Wallis, p < 2.2e-16
M1 Score
0.8
0.4
0.0
-0.4
ns
RTM_IFN
Mac_IFN
RTM_IM
Mac_AgPres
Mac_Rec
RTM-like_MT
Mac_Angio
RTM_LA
Mac_Hypo
Mac_Prolif
Mac_Alv-like
Mac_LA
b
M2 Score
Kruskal-Wallis, p < 2.2e-16
M2 Score
0.50
0.25
0.00
-0.25
RTM_IM
Mac_LA
RTM_LA
RTM-like_MT
Mac_Angio
Mac_IFN
RTM_IFN
Mac_Hypo
Mac_Alv-like
Mac_Rec
Mac_AgPres
Mac_Prolif
c
Ratio M1/M2 Score
1.4
1.2
1.0
0.8
0.6
Mac_Rec
RTM_IFN
Mac_AgPres
Mac_IFN
Mac_Hypo
Mac_Alv-like
RTM_IM
Mac_Angio
RTM-like_MT
Mac_Prolif
RTM_LA
Mac_LA

**Supplementary Figure 8. Macrophages subpopulations express different genes from both classic polarization states. a-b** Violin plot showing the distribution and score signature of M1 **(a)** and M2 **(b)** markers across the macrophage states. Boxes span the first to third quartiles; the horizontal line inside the boxes represents the median black and the dots represent outlier samples in each group. For statistical significance, we performed the Kruskall–Wallis test followed by Wilcoxon to compare each of the thirteen groups against “all” (i.e. base-mean). Ns non-significant. $*p < 0.05$; $**p < 0.01$; $***p < 0.001$; $****p < 0.0001$. **c** Scatter plot of the ratio of M1 and M2 scores.

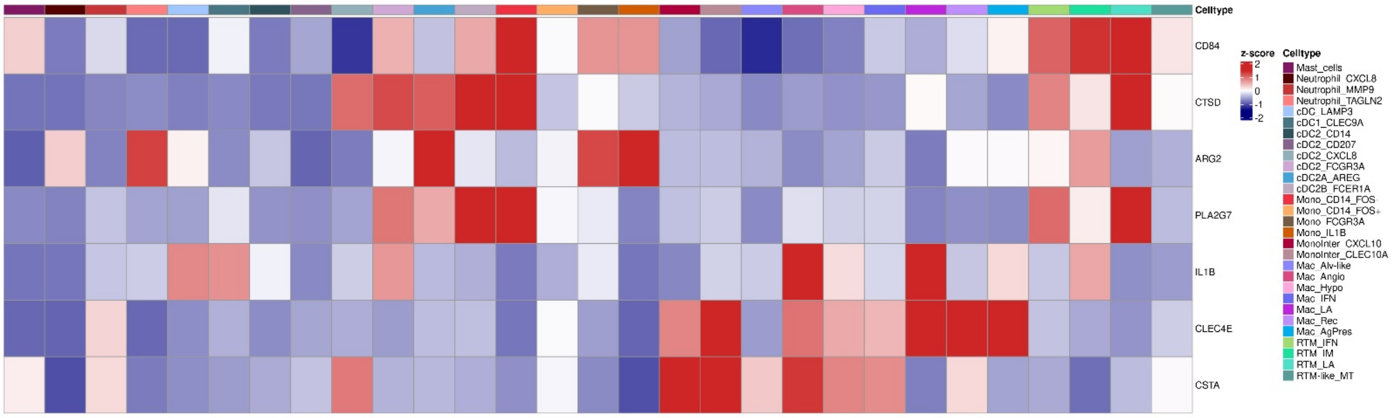
Celltype
CD84
CTSD
ARG2
PLA2G7
IL1B
CLEC4E
CSTA
z-score
2
1
0
-1
-2
Celltype
Mast_cells
Neutrophil_CXCL8
Neutrophil_MMP9
Neutrophil_TAGLN2
cDC_LAMP3
cDC1_CLEC9A
cDC2_CD14
cDC2_CD207
cDC2_CXCL8
cDC2_FCGR3A
cDC2A_AREG
cDC2B_FCER1A
Mono_CD14_FOS-
Mono_CD14_FOS+
Mono_FCGR3A
Mono_IL1B
MonoInter_CXCL10
MonoInter_CLEC10A
Mac_Alv-like
Mac_Angio
Mac_Hypo
Mac_IFN
Mac_LA
Mac_Rec
Mac_AgPres
RTM_IFN
RTM_IM
RTM_LA
RTM-like_MT

**Supplementary Figure 9. Overview of the myeloid-derived suppressor cell signature across MDC subpopulations.** Heatmap showing MDSC signature for each MDC. The color scale represents the scaled expression of each gene.

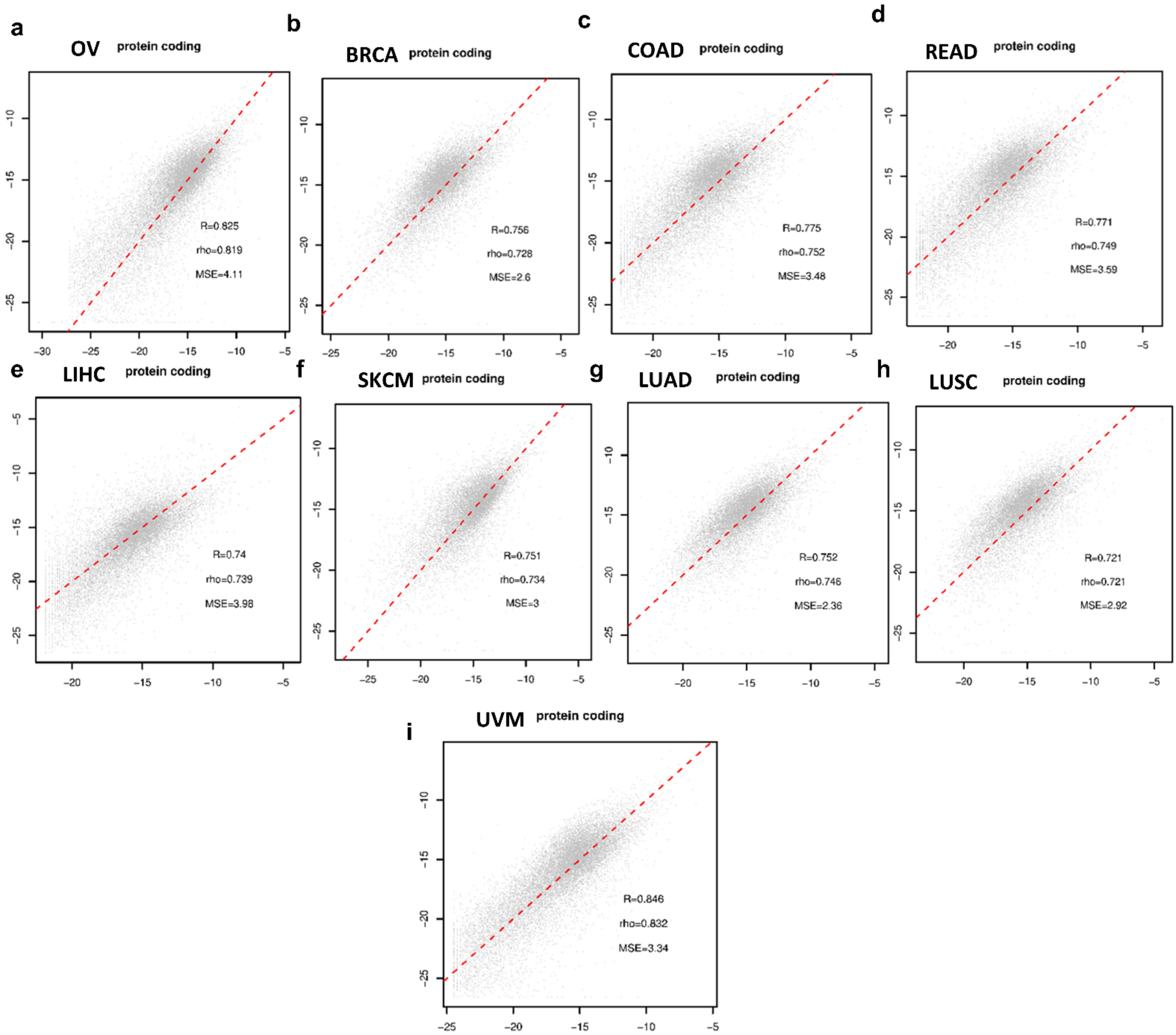
a
OV protein coding
R=0.825
rho=0.819
MSE=4.11
b
BRCA protein coding
R=0.756
rho=0.728
MSE=2.6
c
COAD protein coding
R=0.775
rho=0.752
MSE=3.48
d
READ protein coding
R=0.771
rho=0.749
MSE=3.59
e
LIHC protein coding
R=0.74
rho=0.739
MSE=3.98
f
SKCM protein coding
R=0.751
rho=0.734
MSE=3
g
LUAD protein coding
R=0.752
rho=0.746
MSE=2.36
h
LUSC protein coding
R=0.721
rho=0.721
MSE=2.92
i
UVM protein coding
R=0.846
rho=0.832
MSE=3.34

**Supplementary Figure 10. Correlation between bulk RNA-seq and scRNA-seq cohorts. a-i** Pearson's correlation between bulk RNA-seq and sc-RNA-seq gene expression values. R2 values were found above 0.7 for all tumor types: OV **(a)**; BRCA **(b)**; COAD **(c)**; READ **(d)**; LIHC **(e)**; SKCM **(f)**; LUAD **(g)**; LUSC **(h)**; and UVM **(i).** OV: Ovarian carcinoma; BRCA: Breast carcinoma; COAD: Colon adenocarcinoma; READ: Rectum adenocarcinoma; LIHC: Liver hepatocellular carcinoma; SKCM: Skin cutaneous melanoma; LUAD: Lung adenocarcinoma; LUSC: Lung squamous cell carcinoma (LUSC); UVM: Uveal melanoma.

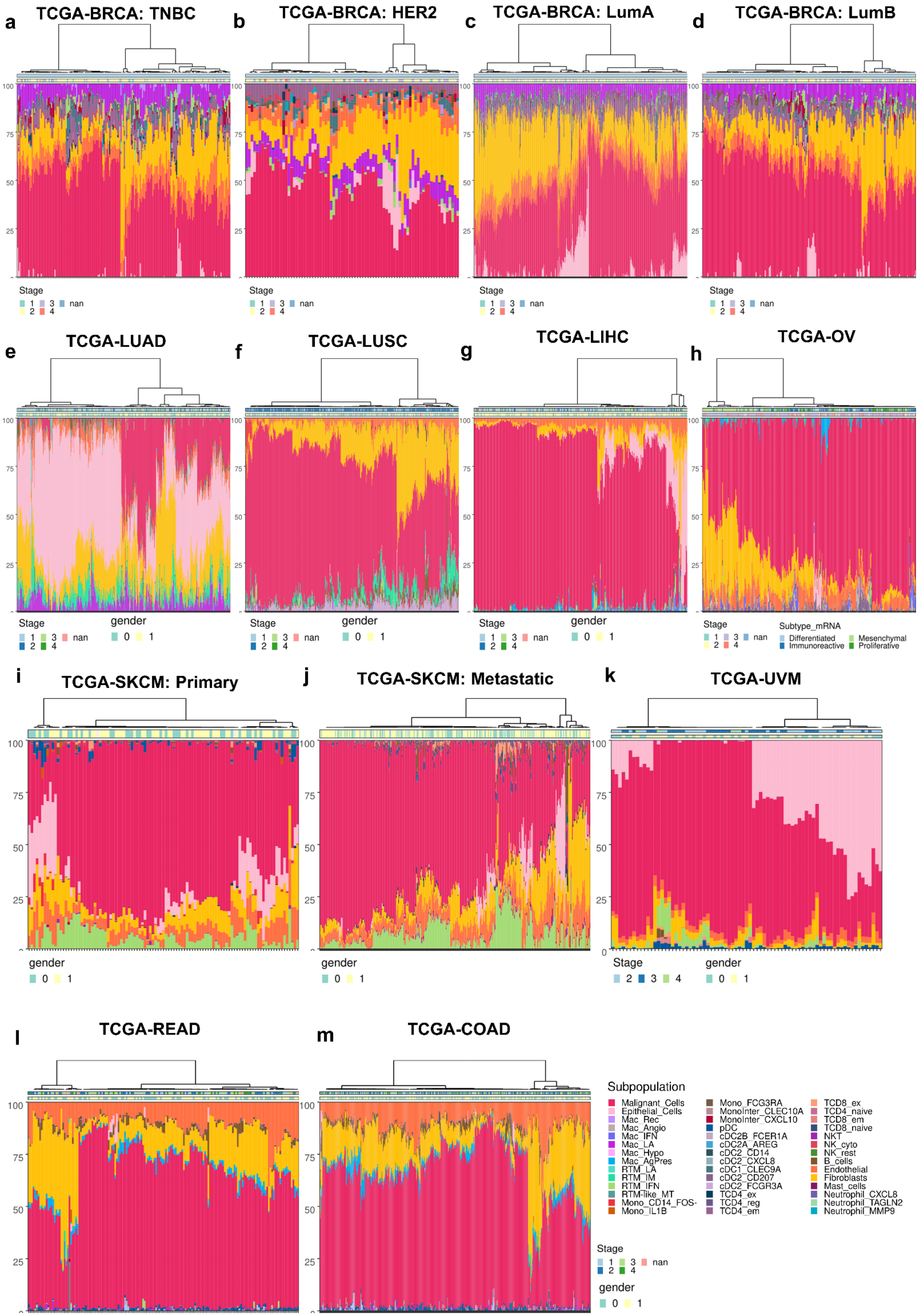

a
TCGA-BRCA: TNBC
b
TCGA-BRCA: HER2
c
TCGA-BRCA: LumA
d
TCGA-BRCA: LumB
100
75
50
25
0
Stage
1 3 nan
2 4
e
TCGA-LUAD
f
TCGA-LUSC
g
TCGA-LIHC
h
TCGA-OV
gender
0 1
Subtype_mRNA
Differentiated
Mesenchymal
Immunoreactive
Proliferative
i
TCGA-SKCM: Primary
j
TCGA-SKCM: Metastatic
k
TCGA-UVM
Stage
2 3 4
l
TCGA-READ
m
TCGA-COAD
Subpopulation
Malignant_Cells
Epithelial_Cells
Mac_Rec
Mac_Angio
Mac_IFN
Mac_LA
Mac_Hypo
Mac_AgPres
RTM_LA
RTM_IM
RTM_IFN
RTM-like_MT
Mono_CD14_FOS-
Mono_IL1B
Mono_FCG3RA
MonoInter_CLEC10A
MonoInter_CXCL10
pDC
cDC2B_FCER1A
cDC2A_AREG
cDC2_CD14
cDC2_CXCL8
cDC1_CLEC9A
cDC2_CD207
cDC2_FCGR3A
TCD4_ex
TCD4_reg
TCD4_em
TCD8_ex
TCD4_naive
TCD8_em
TCD8_naive
NKT
NK_cyto
NK_rest
B_cells
Endothelial
Fibroblasts
Mast_cells
Neutrophil_CXCL8
Neutrophil_TAGLN2
Neutrophil_MMP9

**Supplementary Figure 11. Proportion of subpopulation cells predicted by deconvolution analysis.** Barplot showing the proportion of predicted subpopulations through deconvolution using bulk RNA-Seq from TCGA for **a** TNBC, **b** HER2, **c** Luminal A, **d** Luminal B, **e** Lung Adenocarcinoma (LUAD), **f** Lung squamous cell carcinoma (LUSC), **g** Liver Hepatocellular Carcinoma (LIHC), **h** High-Grade Serous Epithelial Ovarian Cancer, **i** Skin cutaneous melanoma (SKCM - primary tumor), **j** Skin cutaneous melanoma (SKCM - metastatic tumor), **k** Uveal Melanoma (UVM), **l** Rectum adenocarcinoma (READ) and **m** Colon adenocarcinoma (COAD).

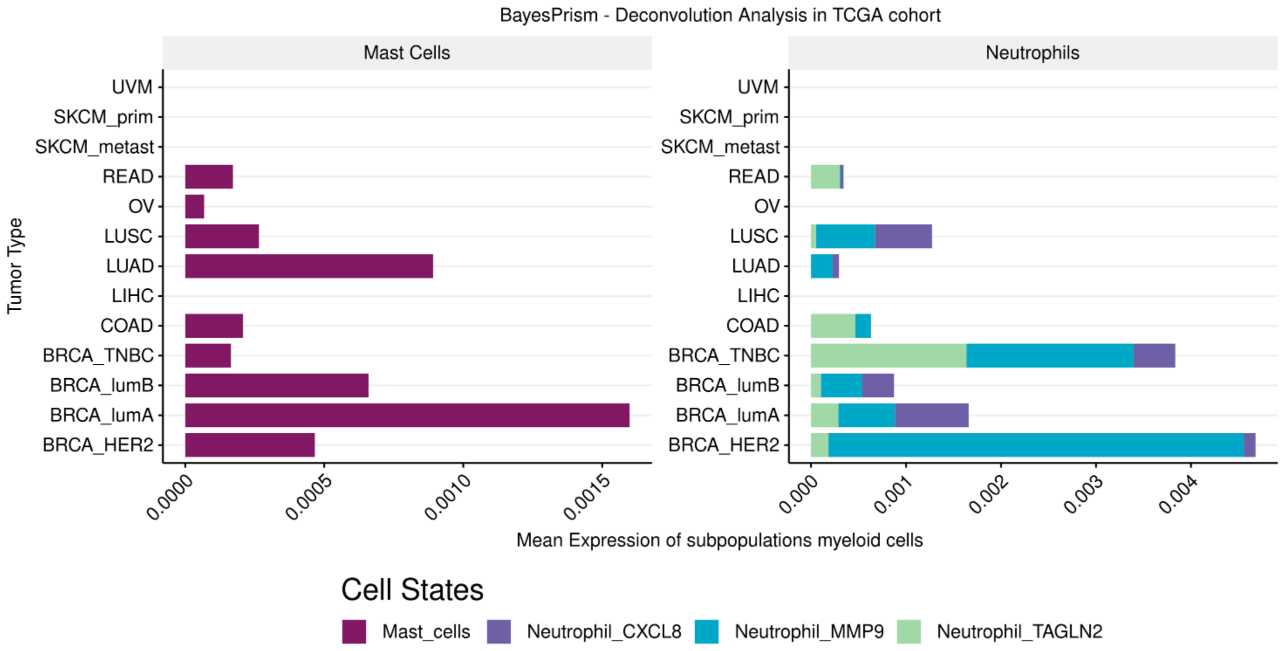

BayesPrism - Deconvolution Analysis in TCGA cohort
Mast Cells
Neutrophils
Tumor Type
UVM
SKCM_prim
SKCM_metast
READ
OV
LUSC
LUAD
LIHC
COAD
BRCA_TNBC
BRCA_lumB
BRCA_lumA
BRCA_HER2
0.0000
0.0005
0.0010
0.0015
0.000
0.001
0.002
0.003
0.004
Mean Expression of subpopulations myeloid cells
Cell States
Mast_cells
Neutrophil_CXCL8
Neutrophil_MMP9
Neutrophil_TAGLN2

**Supplementary Figure 12. Prediction of Mast Cells and Neutrophils enrichment across different tumor types.** Barplot showing the estimation of mast cells and neutrophils calculated through deconvolution analysis in bulk RNA-Seq samples from the TCGA database.

### TCGA-BRCA: TNBC: Mac_LA (TREM2⁺) - Overall Survival and Progression-Free Survival

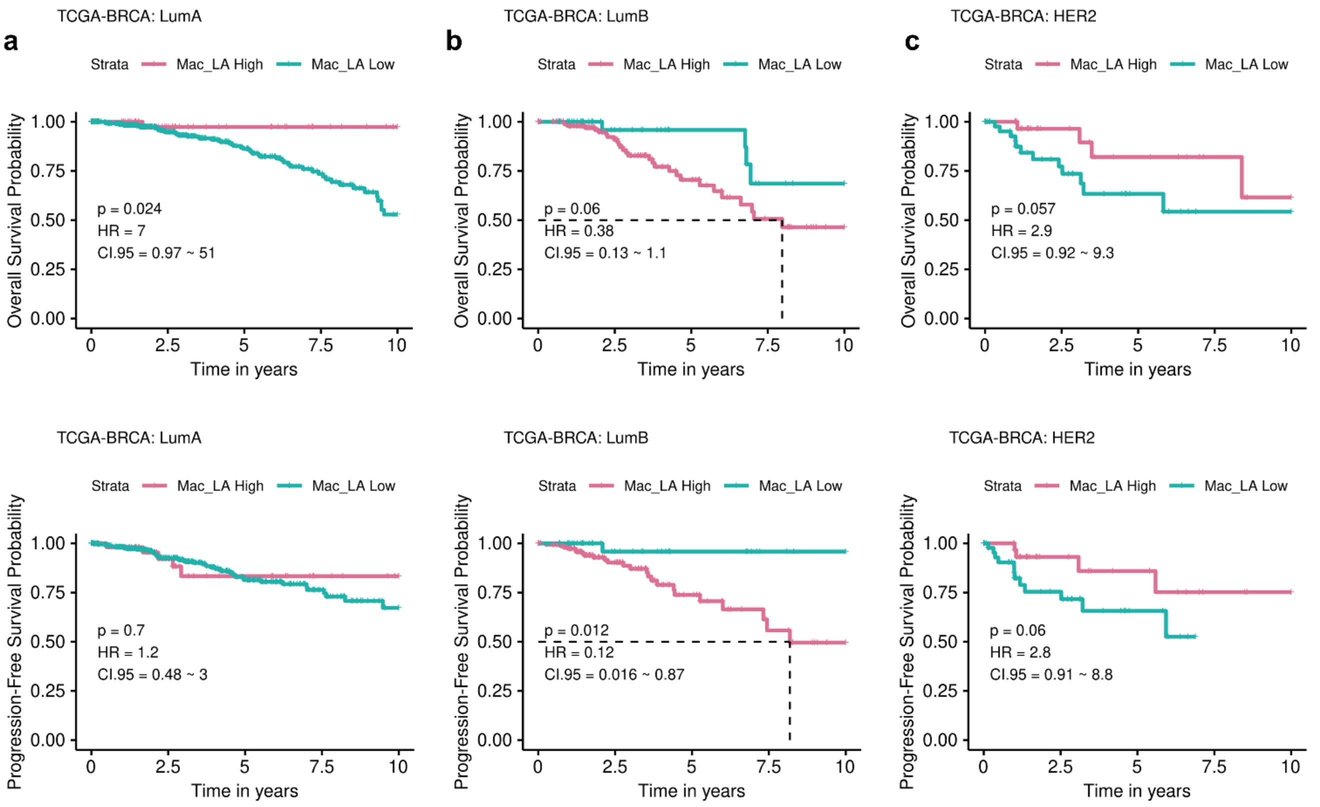


### TCGA-BRCA: TREM2⁺ expression - Overall Survival and Progression-Free Survival

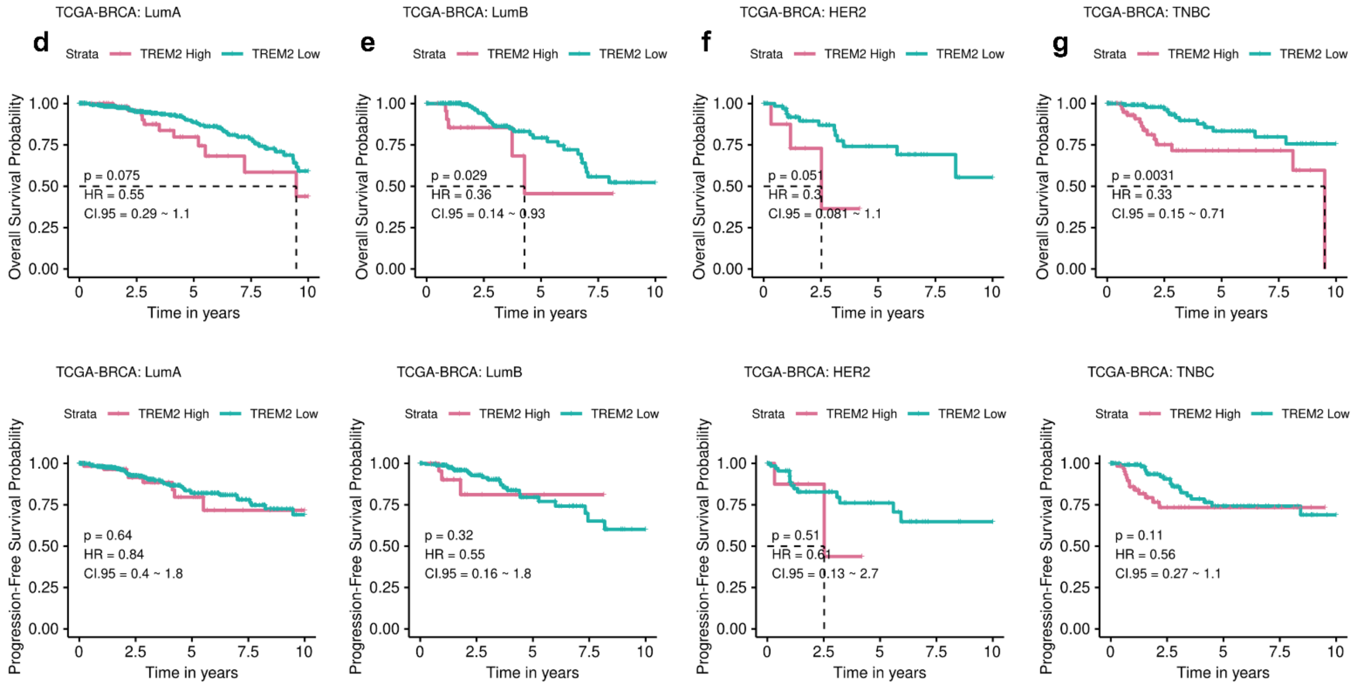

**Supplementary Figure 13. Investigation of Mac_LA and TREM2 prognostic values in breast cancers.** **a-c** Kaplan–Meier curves for Overall and Progression-Free survival for the Luminal A **(a)**, Luminal B **(b)**, and HER2 **(c)** subtypes from the TCGA-BRCA cohort. The patients were divided into low-Mac_LA ($TREM2^{+}$) and high-Mac_LA ($TREM2^{+}$) groups. P-values were calculated using the log-rank test. **d-g** Kaplan–Meier curves for Overall and Progression-Free survival for Luminal A **(d)**, Luminal B **(e)**, HER2 **(f)**, and TNBC **(g)** subtypes grouping samples by pan HIGH- and LOW-*TREM2* expression. The cutoff was obtained using the surv_cutpoint function. HR, hazard ratio. CI, confidence interval.

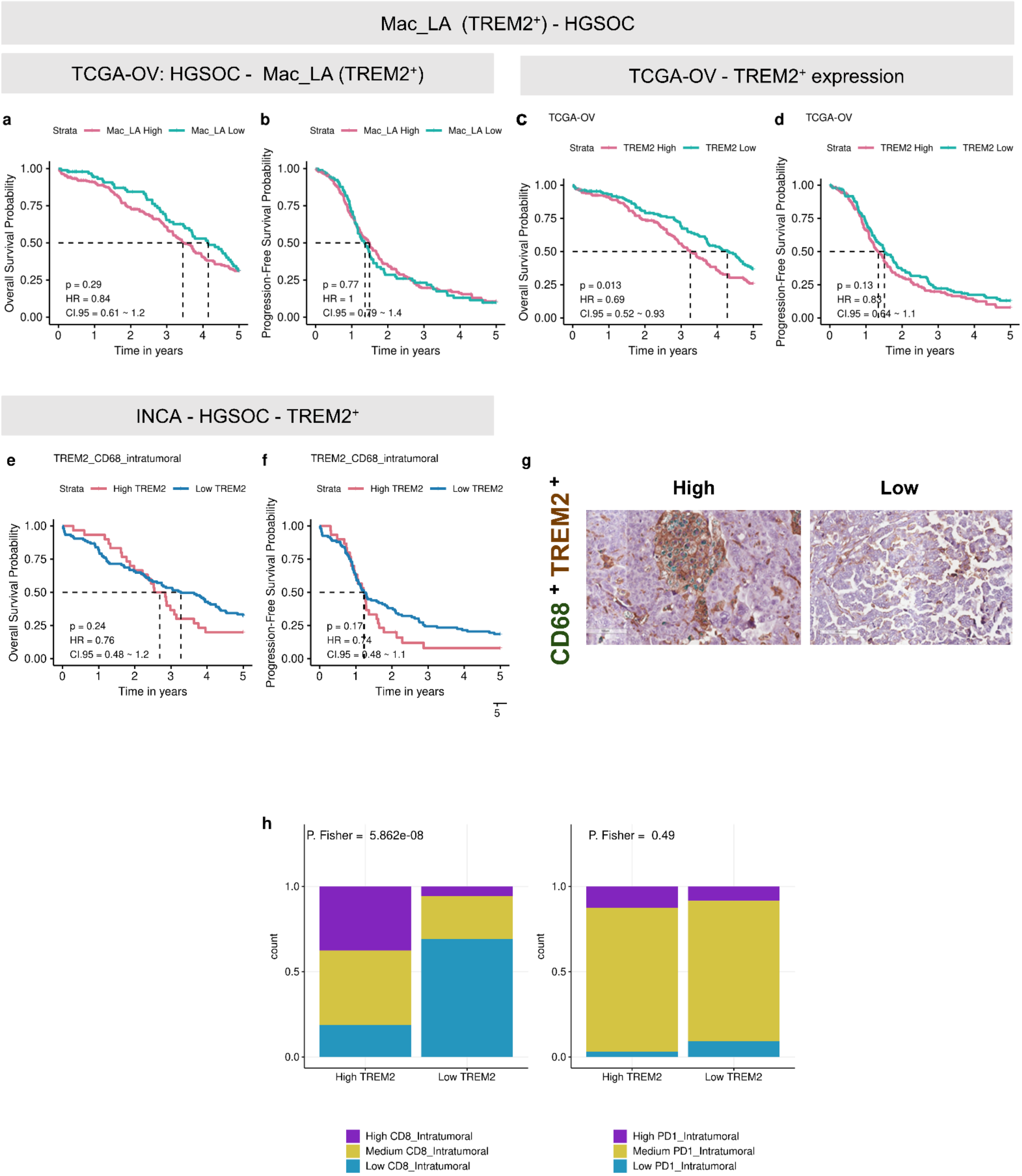
Mac_LA (TREM2+) - HGSOC
TCGA-OV: HGSOC - Mac_LA (TREM2+)
TCGA-OV - TREM2+ expression
a
Strata Mac_LA High Mac_LA Low
Overall Survival Probability
p = 0.29
HR = 0.84
CI.95 = 0.61 ~ 1.2
Time in years
b
Strata Mac_LA High Mac_LA Low
Progression-Free Survival Probability
p = 0.77
HR = 1
Time in years
c
TCGA-OV
Strata TREM2 High TREM2 Low
Overall Survival Probability
p = 0.013
HR = 0.69
CI.95 = 0.52 ~ 0.93
Time in years
d
TCGA-OV
Strata TREM2 High TREM2 Low
Progression-Free Survival Probability
p = 0.13
Time in years
INCA - HGSOC - TREM2+
e
TREM2_CD68_intratumoral
Strata High TREM2 Low TREM2
Overall Survival Probability
p = 0.24
HR = 0.76
CI.95 = 0.48 ~ 1.2
Time in years
f
TREM2_CD68_intratumoral
Strata High TREM2 Low TREM2
Progression-Free Survival Probability
Time in years
g
CD68+ TREM2+
High
Low
h
P. Fisher = 5.862e-08
P. Fisher = 0.49
count
High TREM2
Low TREM2
High CD8_Intratumoral
Medium CD8_Intratumoral
Low CD8_Intratumoral
High PD1_Intratumoral
Medium PD1_Intratumoral
Low PD1_Intratumoral

**Supplementary Figure 14. Clinical impact of Mac_LA and TREM2 in HGSOC tumors. a-b** Kaplan–Meier curves for Overall **(a)** and Progression-Free survival **(b)** curves for HIGH- and LOW- Mac_LA (TREM2$^{+}$) the HGSOC tumors from TCGA cohort; Groups were defined using the surv_cutpoint function. **c-d** Kaplan–Meier curves for Overall **(c)** and Progression-Free Survival **(c)** pan *TREM2* expressing from TCGA cohort. **e-f** Evaluating the presence of TREM2-expressing macrophages through the detection of CD68 and TREM2 protein levels by IHC (INCA cohort). Kaplan–Meier curves for Overall **(e)** and Progression-Free Survival **(f). g** Representative IHC of CD68 and TREM2 expression in the HGSOC cohort. Image obtained by Aperio ImageScope v12.4.6.5003. **h** Proportions of CD8 and PD1 markers in the HIGH and LOW groups of macrophages (CD68$^{+}$ and TREM2$^{+}$) in HGSOC (INCA cohort). The patients were divided into high and low groups of each marker in the IHC analysis. HR, hazard ratio. CI, confidence interval.

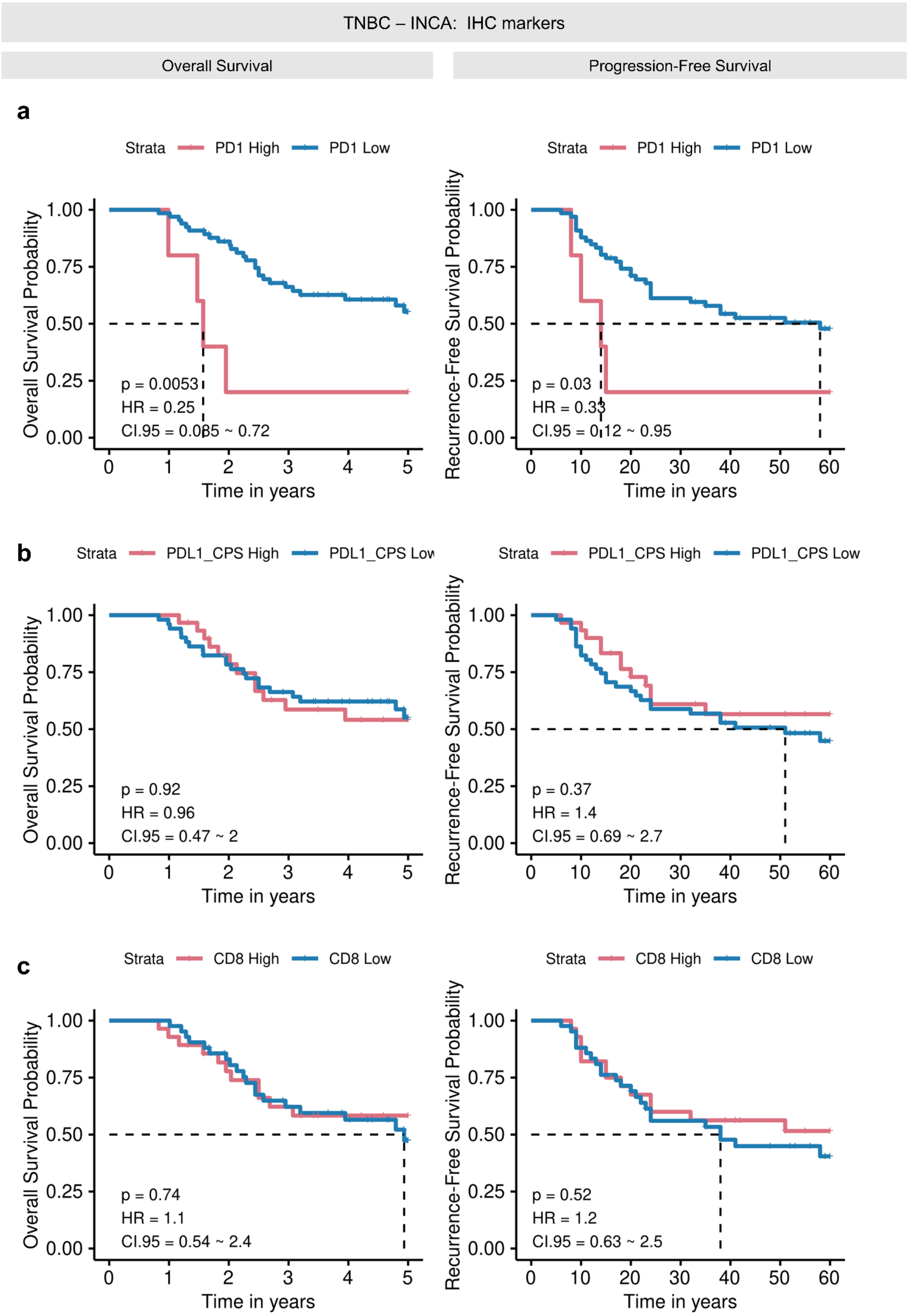

TNBC – INCA: IHC markers
Overall Survival
Progression-Free Survival
a
Strata PD1 High PD1 Low
Overall Survival Probability
1.00
0.75
0.50
0.25
0.00
p = 0.0053
HR = 0.25
CI.95 = 0.085 ~ 0.72
0 1 2 3 4 5
Time in years
Recurrence-Free Survival Probability
p = 0.03
HR = 0.33
CI.95 = 0.12 ~ 0.95
0 10 20 30 40 50 60
Time in years
b
Strata PDL1_CPS High PDL1_CPS Low
p = 0.92
HR = 0.96
CI.95 = 0.47 ~ 2
p = 0.37
HR = 1.4
CI.95 = 0.69 ~ 2.7
c
Strata CD8 High CD8 Low
p = 0.74
HR = 1.1
CI.95 = 0.54 ~ 2.4
p = 0.52
HR = 1.2
CI.95 = 0.63 ~ 2.5

**Supplementary Figure 15. Clinical impact of PD-1, PD-L1, and CD8 protein levels in the Brazilian TNBC Cohort. a-c** Kaplan–Meier curves for Overall and Recurrence-Free survival for the PD-1 **(a)**, PD-L1 **(b)**, and CD8 **(c)** protein levels measured by IHC. The patients were divided into high and low groups of each marker. P-values were calculated using the log-rank test. HR, hazard ratio. CI, confidence interval.

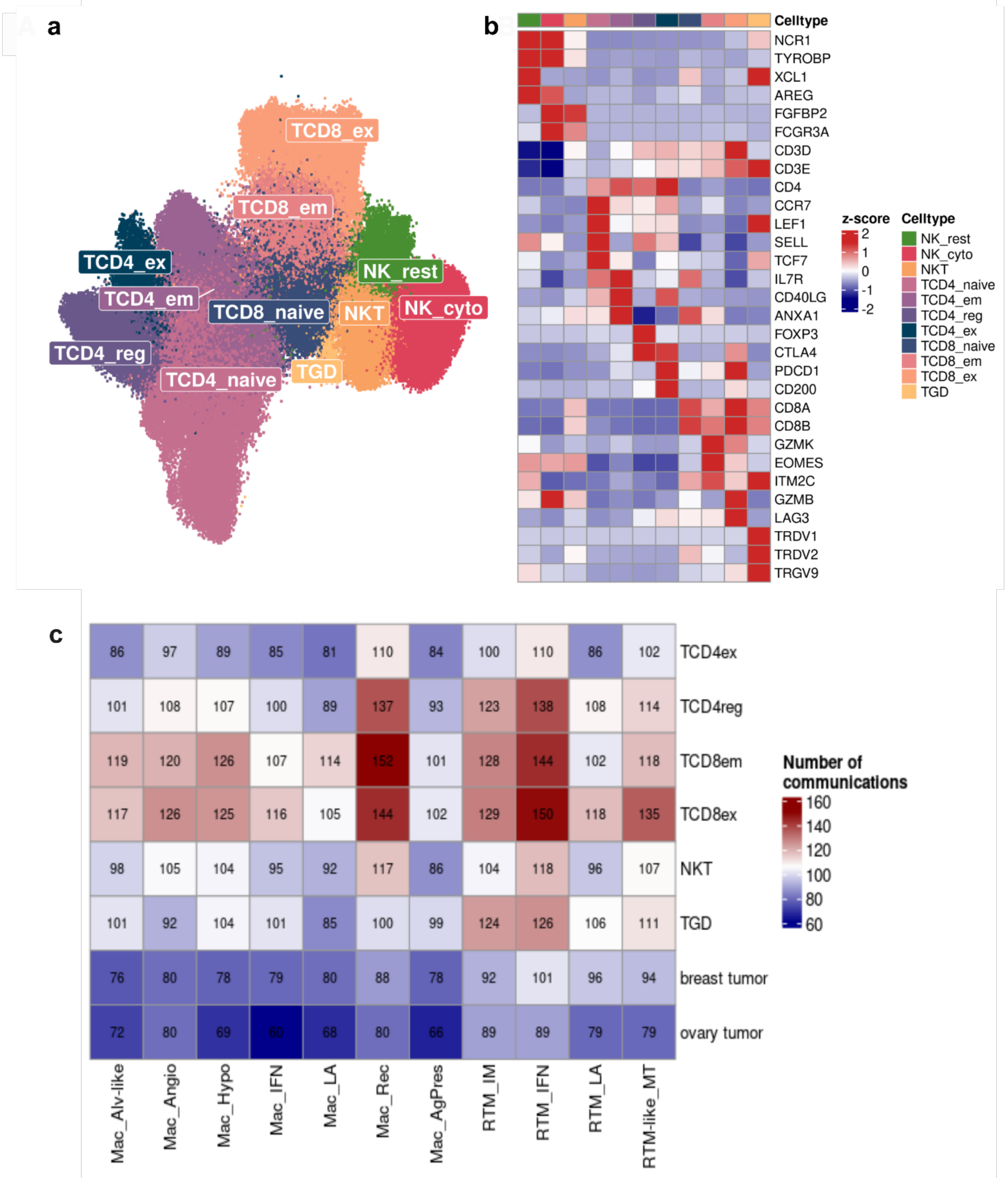
a
TCD8_ex
TCD8_em
TCD4_ex
NK_rest
TCD4_em
TCD8_naive
NKT
NK_cyto
TCD4_reg
TCD4_naive
TGD
b
Celltype
NCR1
TYROBP
XCL1
AREG
FGFBP2
FCGR3A
CD3D
CD3E
CD4
CCR7
LEF1
SELL
TCF7
IL7R
CD40LG
ANXA1
FOXP3
CTLA4
PDCD1
CD200
CD8A
CD8B
GZMK
EOMES
ITM2C
GZMB
LAG3
TRDV1
TRDV2
TRGV9
z-score
2
1
0
-1
-2
Celltype
NK_rest
NK_cyto
NKT
TCD4_naive
TCD4_em
TCD4_reg
TCD4_ex
TCD8_naive
TCD8_em
TCD8_ex
TGD
c
86 97 89 85 81 110 84 100 110 86 102 TCD4ex
101 108 107 100 89 137 93 123 138 108 114 TCD4reg
119 120 126 107 114 152 101 128 144 102 118 TCD8em
117 126 125 116 105 144 102 129 150 118 135 TCD8ex
98 105 104 95 92 117 86 104 118 96 107 NKT
101 92 104 101 85 100 99 124 126 106 111 TGD
76 80 78 79 80 88 78 92 101 96 94 breast tumor
72 80 69 60 68 80 66 89 89 79 79 ovary tumor
Mac_Alv-like
Mac_Angio
Mac_Hypo
Mac_IFN
Mac_LA
Mac_Rec
Mac_AgPres
RTM_IM
RTM_IFN
RTM_LA
RTM-like_MT
Number of communications
160
140
120
100
80
60

**Supplementary Figure 16. T and NK cells characterization. a** UMAP of T and Natural Killer (NK) subpopulations colored by the eleven states identified. **b** Heatmap showing the average expression of the canonical markers for each subpopulation. The clusters were annotated based on canonical gene markers, yielding 11 broad cell types, as mentioned: Resting NK cells - NK_rest (*XCL1* and *AREG*); NK_cyto (*GZMB*, *FGFBP2*, and *FCGR3A*); NKT (*CD8A*, *CD8B*, and *FGFBP2*); TCD4_ex (*CTLA4*, *PDCD1*, and *CD200*); TCD4_em (*IL7R*, *CD40LG* and *ANXA1*); TCD4_naive (*CCR7*, *LEF1*, *SELL* and *TCF7*); TCD4_reg (*FOXP3* and *CTLA4*); TCD8_em (*GZMK*, *EMOS*, and *ITM2C*); TCD8_ex (*LAG3* and *GZMB*); TCD8_naive (*IL7R* and *ANXA1*); and TGD (*TRDV1*, *TRDV2*, and *TRGV9*). **c** Overall number of predicted communication axis retrieved from CellComm algorithm for each cell pair. NK, natural killer, NK_cyto, cytotoxic NK, NKT, Natural killer T cells, TCD4_ex, exhausting T $CD4^{+}$ lymphocytes, TCD4_em, effector/memory T $CD4^{+}$ lymphocytes, TCD4_reg, regulatory T $CD4^{+}$ lymphocytes, TCD8_em, effector/memory T $CD8^{+}$ lymphocytes, TCD8_ex, exhausting T $CD8^{+}$ lymphocytes, TGD, γδ T lymphocytes.

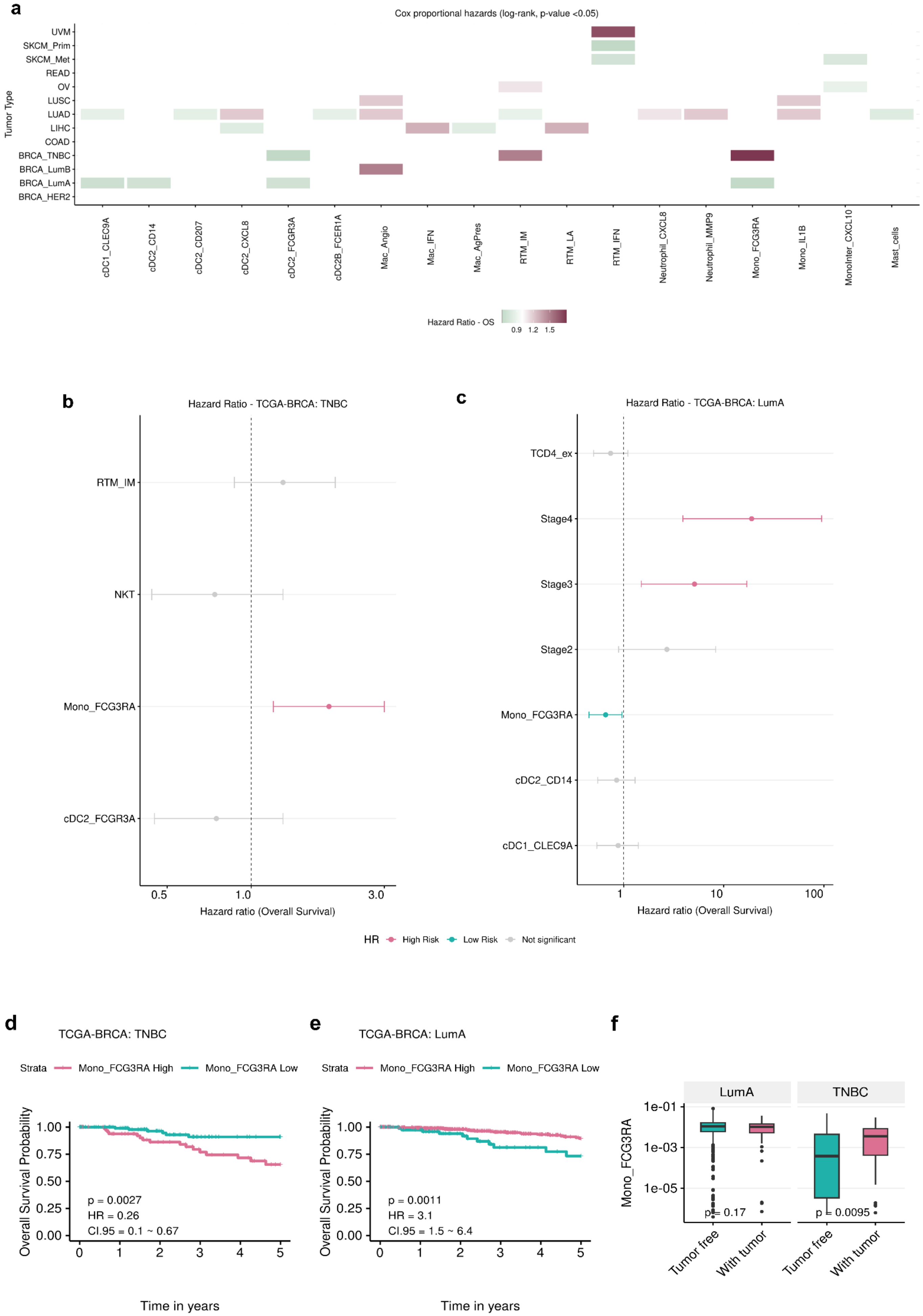

a
Cox proportional hazards (log-rank, p-value <0.05)
Tumor Type
UVM
SKCM_Prim
SKCM_Met
READ
OV
LUSC
LUAD
LIHC
COAD
BRCA_TNBC
BRCA_LumB
BRCA_LumA
BRCA_HER2
cDC1_CLEC9A
cDC2_CD14
cDC2_CD207
cDC2_CXCL8
cDC2_FCGR3A
cDC2B_FCER1A
Mac_Angio
Mac_IFN
Mac_AgPres
RTM_IM
RTM_LA
RTM_IFN
Neutrophil_CXCL8
Neutrophil_MMP9
Mono_FCG3RA
Mono_IL1B
MonoInter_CXCL10
Mast_cells
Hazard Ratio - OS
0.9 1.2 1.5
b
Hazard Ratio - TCGA-BRCA: TNBC
RTM_IM
NKT
Mono_FCG3RA
cDC2_FCGR3A
0.5
1.0
3.0
Hazard ratio (Overall Survival)
c
Hazard Ratio - TCGA-BRCA: LumA
TCD4_ex
Stage4
Stage3
Stage2
Mono_FCG3RA
cDC2_CD14
cDC1_CLEC9A
1
10
100
Hazard ratio (Overall Survival)
HR High Risk Low Risk Not significant
d
TCGA-BRCA: TNBC
Strata Mono_FCG3RA High Mono_FCG3RA Low
Overall Survival Probability
1.00
0.75
0.50
0.25
0.00
p = 0.0027
HR = 0.26
CI.95 = 0.1 ~ 0.67
0 1 2 3 4 5
Time in years
e
TCGA-BRCA: LumA
Strata Mono_FCG3RA High Mono_FCG3RA Low
Overall Survival Probability
1.00
0.75
0.50
0.25
0.00
p = 0.0011
HR = 3.1
CI.95 = 1.5 ~ 6.4
0 1 2 3 4 5
Time in years
f
LumA
TNBC
Mono_FCG3RA
1e-01
1e-03
1e-05
p = 0.17
p = 0.0095
Tumor free
With tumor
Tumor free
With tumor

**Supplementary Figure 17. Mono_FCGR3A is associated with the clinical outcome in BRCA tumors from TCGA**. **a** Univariate cox regression analysis for predicted cell subpopulation enrichment in bulk RNAseq samples from different cancer types. Color scale represents the Hazard Ratios (HR) that were calculated relative to the patient's overall survival; **b-c** Multivariate cox regression analysis for TCGA-BRCA:TNBC cohort **(b)** and TCGA-BRCA: Luminal A **(c)** cohort. Light green = Low Risk ($HR < 1$; $p < 0.05$); Pink = High Risk ($HR > 1$, $p < 0.05$); Gray = Not significant. **d-e** Kaplan–Meier curves for Overall and Progression-Free survival for the TNBC **(d)** and Luminal A **(e)** patients from TCGA-BRCA cohort. The patients were divided into LOW-Mono_FCGR3A and HIGH-Mono_FCGR3A groups using the surv_cutpoints function. P-values were calculated using the log-rank test. **f** Boxplot demonstrating the clinical impact of Mono_FCGR3A in relation to tumor status (tumor-free and with tumor) in both breast cancer subtypes TNBC and Luminal-A. For statistical significance, we performed the Wilcoxon test to compare the groups. HR, hazard ratio. CI, confidence interval.

TCGA-BRCA - RTM_IM (FOLR2 +) – Overall Survival and Progression-Free Survival

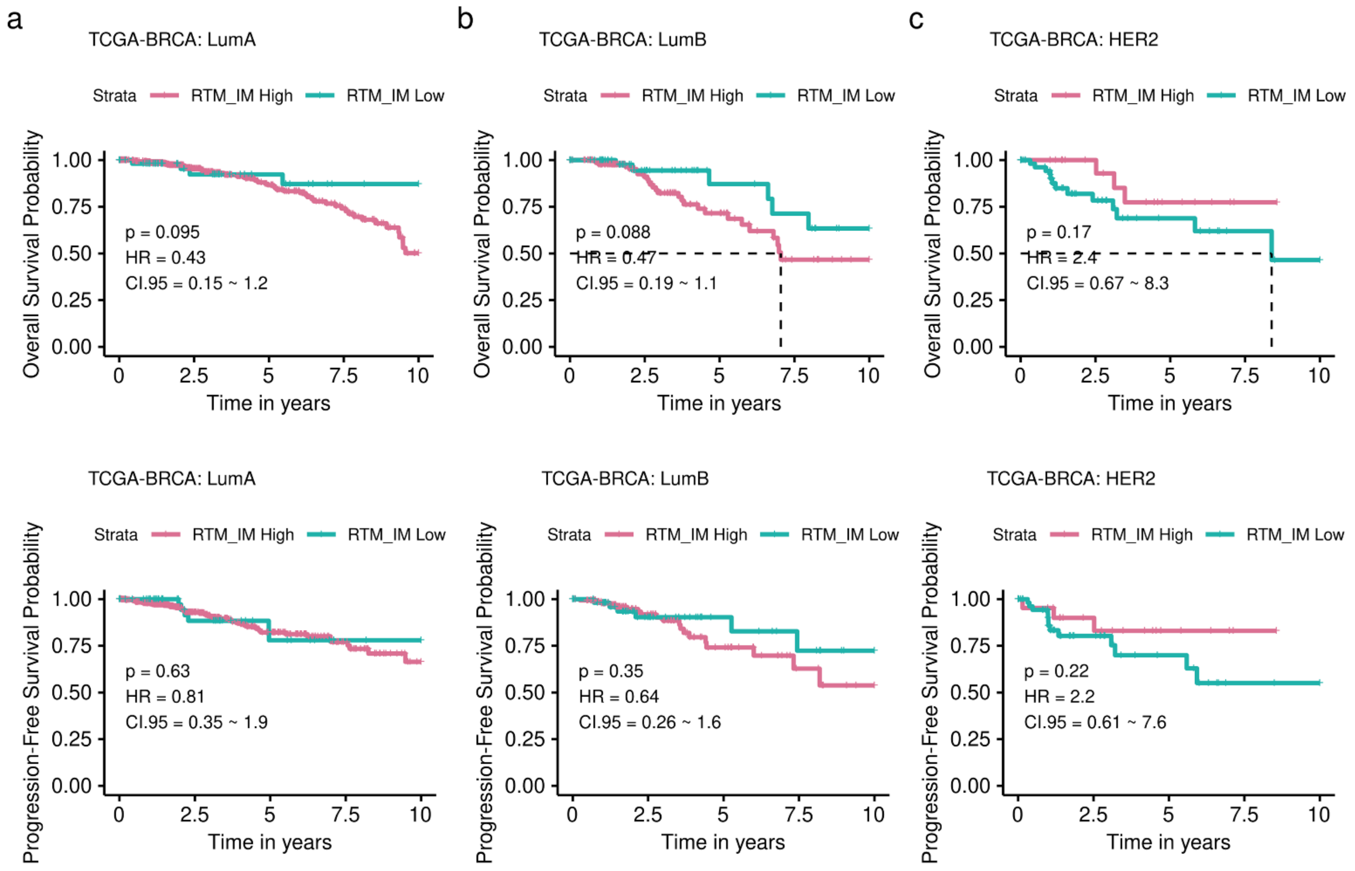

**Supplementary Figure 18. Clinical impact of RTM_IM (FOLR2+) on breast tumors. a-c** Kaplan–Meier curves for Overall and Progression-Free survival for the Luminal A **(a)**, Luminal B **(b)**, and HER2 **(c)** patients from TCGA-BRCA cohort. The patients were divided into LOW-RTM_IM (*FOLR2+*) and HIGH-RTM_IM (*FOLR2+*) groups using the surv_cutpoints function. P-values were calculated using the log-rank test. HR, hazard ratio. CI, confidence interval.

### METABRIC - FOLR2 + (zscore) expression - Overall Survival

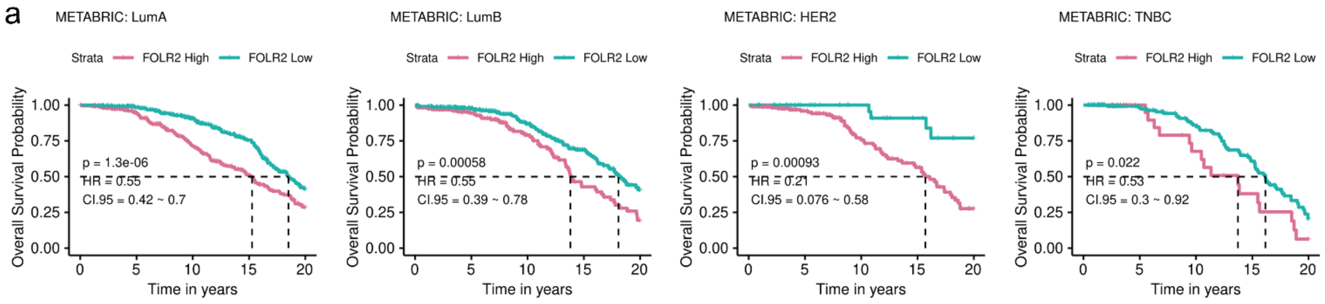


### TCGA-BRCA - FOLR2 + expression. - Overall Survival and Progression-Free Survival

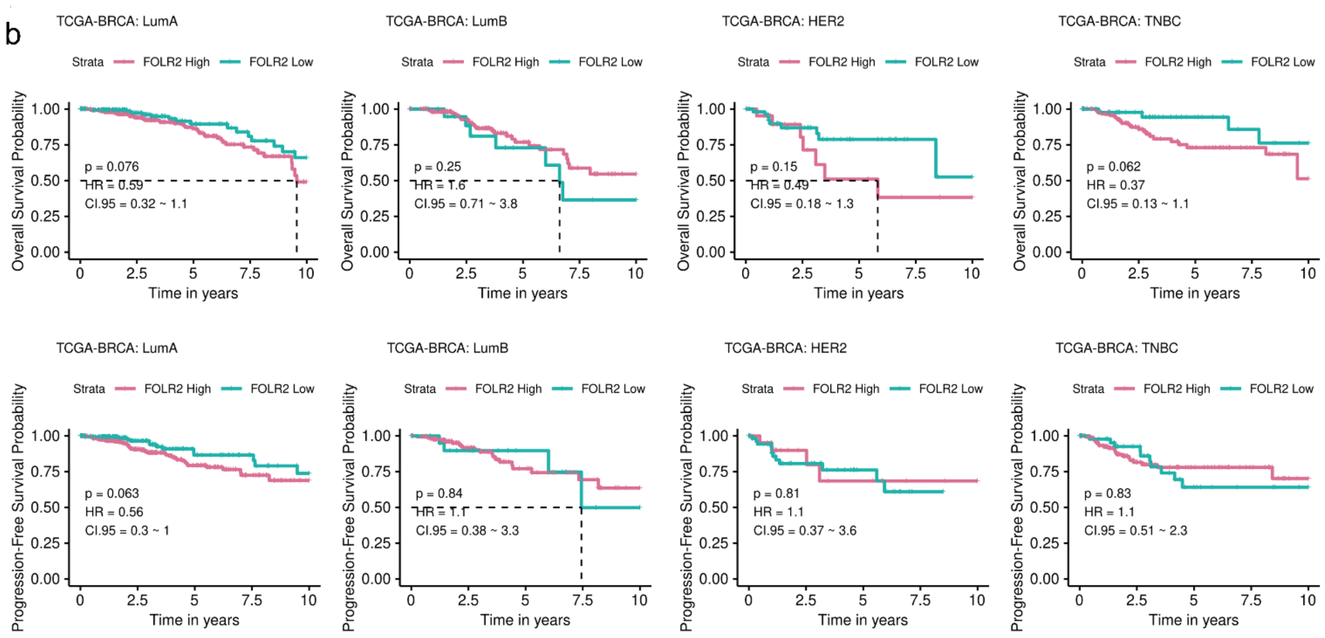


### TCGA-OV: HGSOC - FOLR2 + expression - Overall Survival and Progression-Free Survival

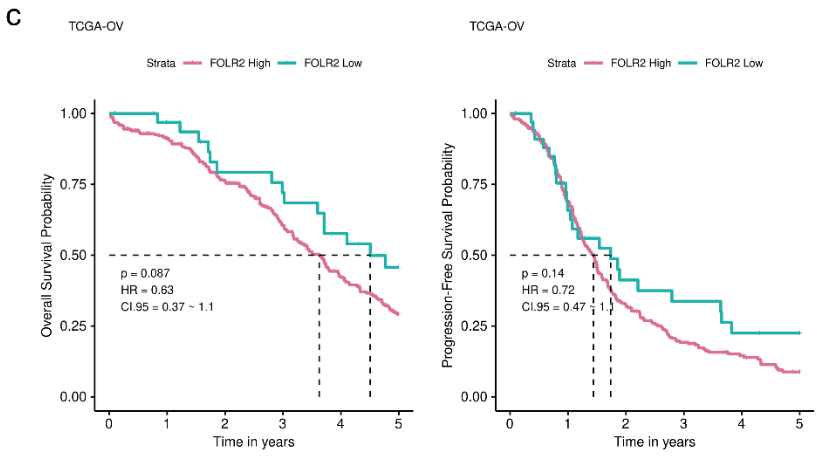

**Supplementary Figure 19. Clinical Impact of *FOLR2* gene expression in different cohorts. a-c** Kaplan–Meier curves for Overall and Progression-Free survival for breast cancer subtypes using Metabric **(a),** BRCA-TCGA cohorts **(b),** and HGSOC patients from the TCGA cohort **(c)**. The patients were divided into LOW-*FOLR2*$^{+}$ and HIGH-*FOLR2*$^{+}$ groups using the surv_cutpoints function. P-values were calculated using the log-rank test. HR, hazard ratio. CI, confidence interval.

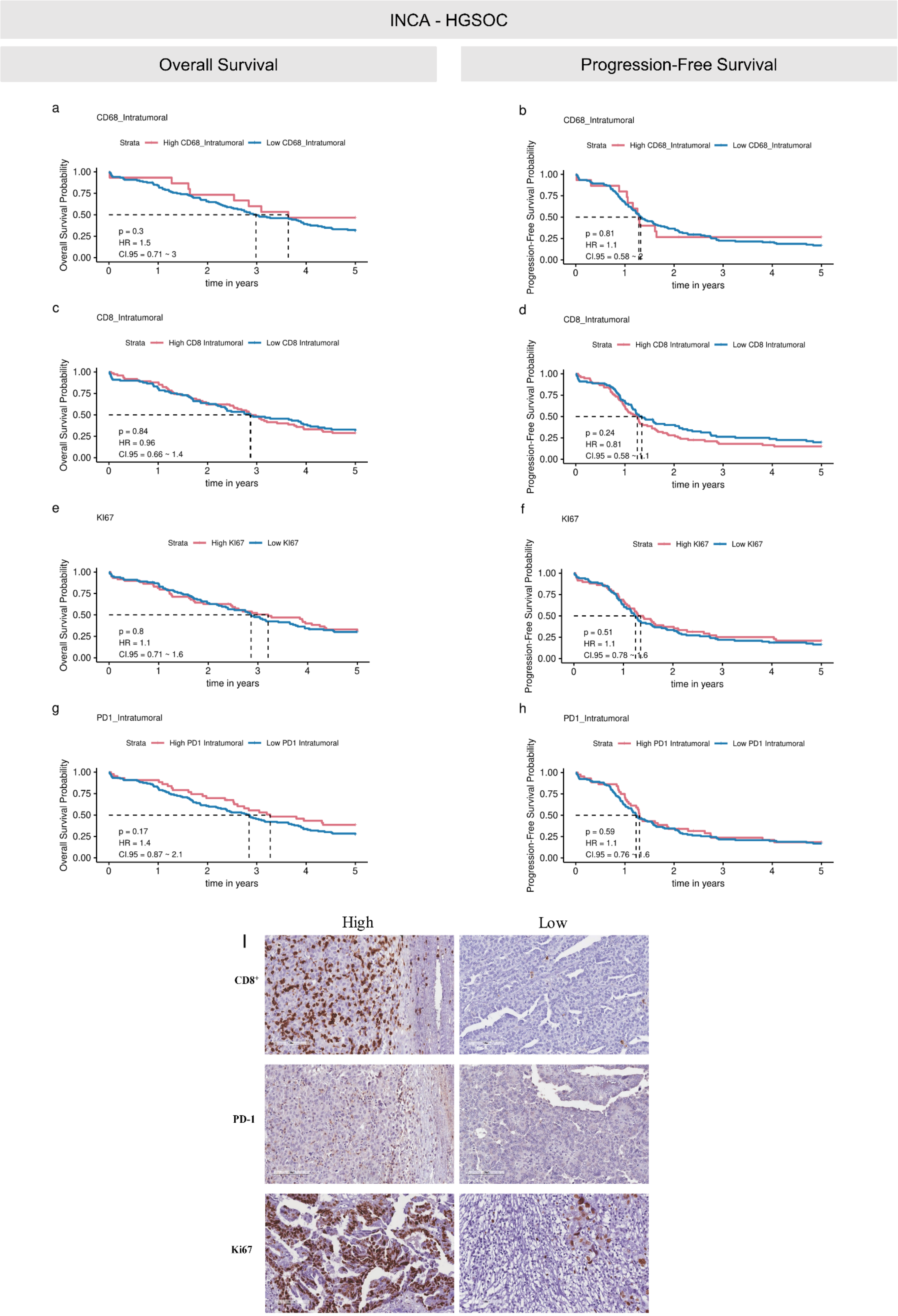
INCA - HGSOC
Overall Survival
Progression-Free Survival
a
CD68_Intratumoral
Strata High CD68_Intratumoral Low CD68_Intratumoral
Overall Survival Probability
p = 0.3
HR = 1.5
CI.95 = 0.71 ~ 3
time in years
b
CD68_Intratumoral
Strata High CD68_Intratumoral Low CD68_Intratumoral
Progression-Free Survival Probability
p = 0.81
HR = 1.1
CI.95 = 0.58 ~ 2
time in years
c
CD8_Intratumoral
Strata High CD8 Intratumoral Low CD8 Intratumoral
Overall Survival Probability
p = 0.84
HR = 0.96
CI.95 = 0.66 ~ 1.4
time in years
d
CD8_Intratumoral
Strata High CD8 Intratumoral Low CD8 Intratumoral
Progression-Free Survival Probability
p = 0.24
HR = 0.81
CI.95 = 0.58 ~ 1.1
time in years
e
KI67
Strata High KI67 Low KI67
Overall Survival Probability
p = 0.8
HR = 1.1
CI.95 = 0.71 ~ 1.6
time in years
f
KI67
Strata High KI67 Low KI67
Progression-Free Survival Probability
p = 0.51
HR = 1.1
CI.95 = 0.78 ~ 1.6
time in years
g
PD1_Intratumoral
Strata High PD1 Intratumoral Low PD1 Intratumoral
Overall Survival Probability
p = 0.17
HR = 1.4
CI.95 = 0.87 ~ 2.1
time in years
h
PD1_Intratumoral
Strata High PD1 Intratumoral Low PD1 Intratumoral
Progression-Free Survival Probability
p = 0.59
HR = 1.1
CI.95 = 0.76 ~ 1.6
time in years
High
Low
I
CD8+
PD-1
Ki67

**Supplementary Figure 20. Clinical impact of CD68, CD8, Ki-67, and PD-1 in the Brazilian HGSOC Cohort. a-h** Kaplan–Meier curves for Overall and Recurrence-Free survival for HGSOC patients from INCA cohort considering intratumoral CD68 **(a-b),** CD8 **(c-d)**, Ki67 **(e-f)**, and PD1 **(g-h)** markers through IHC. The patients were divided into LOW- and HIGH groups for each marker using the surv_cutpoints function. P-values were calculated using the log-rank test. HR, hazard ratio. CI, confidence interval. **i** Representative images of high and low CD8, PD-1 and Ki67$^{+}$ IHC staining on human ovarian tissue. Images were collected at 20X magnification.
