## Supplementary Table 1 for "Identification of novel myeloid-derived cell states with implication in cancer outcome"

**Supplementary Table 1: Data summary for the single-cell analysis**

| Study | Link to Download | Paper | Tissue | Sample Type | Technology | total cells* |
|---|---|---|---|---|---|---|
| Azizi et. al 2018 | GSE114727 | https://doi.org/10.1016/j.cell.2018.05.060 | Breast | Normal/Tumor/Lymph node/Blood | InDrop | 45054 |
| Durante et al | GSE139829 | https://doi.org/10.1038/s41467-019-14256-1 | Uveal Melanoma | Tumor | 10x | 56759 |
| Geistlinger et al., 2020 | GSE154600 | https://doi.org/10.1158/0008-5472.can-20-0521 | Ovary | Tumor | 10x | 42293 |
| Human Lung Cell Atlas | https://www.synapse.org/#!Synapse:syn21560407 | https://hlca.ds.czbiohub.org/ | Lung | Normal | Smart-seq/10x | 47077 / 8563 |
| Jerby-Arnon et al. 2018 | GSE115979 | https://doi.org/10.1016/j.cell.2018.09.006 | Melanoma | Tumor | Smart-seq | 2637 |
| Karaayvaz et al 2018 | GSE118390 | https://doi.org/10.1038/s41467-018-06052-0 | Breast | Tumor | Smart-seq | 1168 |
| Ma et al 2019 | GSE125449 | https://doi.org/10.1016/j.ccell.2019.08.007 | Liver | Tumor | 10x | 7871 |
| Nieto et al. 2020 | GSE158803 | https://doi.org/10.1101/2020.10.26.354829 | NSCLC metastasis/ uveal melanoma metastasis/uveal melanoma | Tumor | 10x | 2274 |
| Qian et. al 2020 | blueprint.lambrechtslab.org/ | https://doi.org/10.1038/s41422-020-0355-0 | Ovary/Lung/Colorectal/Breast | Normal/Tumor** | 10x | 155819 |
| Solé-Boldo et. al 2020 | GSE130973 | https://doi.org/10.1038/s42003-020-0922-4 | Skin | Normal | 10x | 14754 |
| Tirosh et al, 2018 | GSE115978 | https://doi.org/10.1126/science.aad0501 | Melanoma | Tumor | Smart-seq | 4231 |
| Zheng et al | https://support.10xgenomics.com/single-cell-gene-expression/datasets/1.1.0/pbmc3k? | https://doi.org/10.1038/ncomms14049 | PBMC | Normal | 10x | 2661 |

| | | | | | | |
|---|---|---|---|---|---|---|
| Zilionis et al 2019 | GSE127465 | https://doi.org/10.1016/j.immuni.2019.03.009 | Lung | Tumor | InDrop | 52577 |
| **13 studies** | | | **8 sites** | **4 conditions** | **3 technologies** | **452.321 cells** |

*after quality control

**except for breast - only tumor cells
