## Supplementary Table 2 for "Identification of novel myeloid-derived cell states with implication in cancer outcome"

**Supplementary Table 2: Clinical information of 142 patients involved in this study**

| Study | Patient | Gender | Age | Sample Type | Histologycal Subtype | Grade/Stag | Treatment | Single cell technology | Total cells* |
|---|---|---|---|---|---|---|---|---|---|
| Azizi *et al*. 2018 | BC1 | F | 38 | Blood, normal and tumor | Breast Ductal | I | Naïve | InDrop | 47,016 CD45+ |
| | BC2 | | 60 | Lymph node, normal and | | II | | | |
| | BC3 | | 43 | normal and tumor | | III | | | |
| | BC4 | | 52 | Blood and tumor | | I | | | |
| | BC5 | | 78 | Tumor | | III | | | |
| | BC6 | | 58 | Tumor | | II | | | |
| | BC7 | | 65 | Normal and tumor | | II | | | |
| | BC8 | | 72 | Normal and tumor | | III | | | |
| Durante *et al*. 2020 | UMM059 | F | 86 | Tumor | Primary Uveal melanoma | II | N/A | 10X-Chromium | 59,915 |
| | UMM061 | F | 71 | | | II | | | |
| | UMM062 | M | 69 | | | IA | | | |
| | UMM063 | M | 66 | | | II | | | |
| | UMM064 | M | 77 | | | II | | | |
| | UMM065 | F | 44 | | | IA | | | |
| | UMM066 | M | 53 | | | II | | | |
| | UMM069 | F | 80 | | | II | | | |
| | BSSR0022 | F | 68 | | | IB | | | |
| | UMM067L | M | 73 | Tumor-liver interface | Uveal melanoma (Metastasis) | II | | | |
| | UMM041L | F | 63 | | | II | | | |
| Geistlinger *et al.* 2020 | Tumor59 | F | N/A | Omentum | HGSOC | III | After chemotherapy | 10X-Chromium | 42293 |
| | Tumor76 | | | | | | | | |
| | Tumor77 | | | | | | | | |
| | Tumor89 | | | | | | | | |
| | Tumor90 | | | | | | | | |
| Human Lung Cell Atlas (2020) | Patient 1 | M | 75 | Blood, normal and tumor | Lung | N/A | N/A | Smart-seq2/10X-Chromium | 9,404/ 65,662 |
| | Patient 2 | | 46 | | | | | | |
| | Patient 3 | F | 51 | | | | | | |
| | Mel126 | M | 63 | Soft tissue | Melanoma (metastasis) | | Ipilimumab, nivolumab | | |
| | Mel04.3 | | 81 | Skin | | | Ipilimumab | | |
| | Mel110 | | 74 | R adrenal metastasis | | | ipilimumab + angiopoietin 2 inhibitor, Temezlolamide, Pembrolizumab | | |
| | Mel121.1 | | 74 | Skin | | | S/p Pembrolizumab | | |
| | Mel106 | | 67 | Necrotic L axillary lymph nodes | | | Prior treatment: nivolumab + ipilimumab | | |
| | Mel75 | | 81 | Soft tissue | | | Ipilimumab + nivolumab, WDVAX, Pembrolizumab | | |

| | | | | | | | | | |
|---|---|---|---|---|---|---|---|---|---|
| | Mel98 | F | 47 | L thigh soft tissue metastasis | | N/A | S/p IFN, s/p ipilimumab+GMCSF | Smart-seq | 2,987 |
| | Mel102 | F | 72 | Fragmented pieces of (R) adrenal gland metastasis | | | S/p nivolumab + ipilimumab | | |
| | Mel129 | M | 63 | Skin | Melanoma (primary) | | Naïve | | |
| | Mel129 | M | 63 | Skin | Melanoma (primary) | | Naïve | | |
| | Mel116 | M | 85 | Lymph node | Melanoma (metastasis) | | Naïve | | |
| | Mel103 | M | 58 | Lymph node | Melanoma (metastasis) | | Naïve | | |
| | Mel105 | M | 77 | Skin | Melanoma (primary) | | Naïve | | |
| | Mel112 | M | 76 | Bulky (L) axillary metastasis | | | Naïve | | |
| | Mel194 | M | 68 | L anterior shoulder subcutaneous | Melanoma (metastasis) | | Nivolumab + lirilumab (anti-kit), Nivolumab, Ipilimumab, Pan-RAF-inhibitor, Pembrolizumab | | |
| Jerby-Arnon *et al.* 2018 | Mel478 | F | 54 | Transanal rectal mass | | | Naïve | | |
| | Mel128 | M | 37 | Lymph node | | | Naïve | | |
| | Mel53 | F | 77 | Subcutaneous back lesion | | | Naïve | | |
| | Mel58 | M | 83 | Subcutaneous leg lesion | | | Ipilimumab | | |
| | Mel60 | M | 60 | Spleen | | | Trametinib, ipilimumab | | |
| | Mel71 | M | 79 | Transverse colon | | | Naïve | | |
| | Mel72 | F | 57 | External iliac lymph node | | | IL-2, nivolumab, ipilimumab + anti- KIR-Ab | | |
| | Mel74 | M | 63 | Terminal Ileum | Melanoma (metastasis) | | Nivolumab | | |
| | Mel75 | M | 80 | Subcutaneous leg lesion | | | Ipilimumab + nivolumab, WDVAX | | |
| | Mel78 | M | 73 | Small bowel | | N/A | WDVAX, ipilimumab + nivolumab | Smart-seq | 4,199 |
| | Mel79 | M | 74 | Axillary lymph node | | | Naïve | | |
| | Mel80 | F | 86 | Axillary lymph node | | | Naïve | | |
| | Mel81 | F | 43 | Axillary lymph node | | | Naïve | | |
| | Mel82 | F | 73 | Axillary lymph node | | | Naïve | | |
| | Mel84 | M | 67 | Acral primary | Melanoma (primary) | | Naïve | | |
| | Mel88 | F | 54 | Cutanoues met | | | Tremelimumab + MEDI3617 | | |
| | Mel89 | M | 67 | Axillary lymph node | Melanoma (metastasis) | | None | | |
| Tirosh et al. 2016 | Mel94 | F | 54 | Iliac lymph node | | | IFN, ipilimumab + nivolumab | | |
| | 39 | | 64 | | | III | | | |
| | 58 | | 46 | | | III | | | |
| | 81 | F | 45 | Tumor | Non-metastatic TNBC invasive ductal | III | Naïve | Smart-seq | 1168 |
| | 84 | | 52 | | | III | | | |
| | 89 | | 44 | | | II | | | |
| Karaayvaz et al 2018 | 124 | | 42 | | | III | | | |

| | | | | | | | | | |
|---|---|---|---|---|---|---|---|---|---|
| | H18 | M | 61 | | | IV | durvalumab/tremelimumab | | |
| | H21 | M | 77 | | | III | resection | | |
| | H23 | F | 41 | | | III | durvalumab/tremelimumab | | |
| | C25 | F | 47 | | | IV | durvalumab/tremelimumab | | |
| | C26 | M | 63 | | | IV | durvalumab/tremelimumab | | |
| | H28 | F | 63 | | | III | durvalumab/tremelimumab | | |
| | C29 | M | 61 | | | IV | durvalumab/tremelimumab | | |
| | H30 | M | 63 | | | IV | durvalumab/tremelimumab | | |
| | C35 | F | 64 | | | IV | pembrolizumab | | |
| | H37 | M | 65 | Tumor | Liver | IV | durvalumab/tremelimumab | 10X-Chromium | 7871 |
| | H38 | M | 74 | | | I | resection | | |
| | C39 | M | 61 | | | IV | pembrolizumab | | |
| | H34 | M | 63 | | | IV | durvalumab/tremelimumab | | |
| | C42 | M | 67 | | | IV | pembrolizumab | | |
| | C46 | M | 69 | | | IV | pembrolizumab | | |
| | C56 | F | 52 | | | IV | durvalumab/tremelimumab | | |
| | C60 | F | 80 | | | III | resection | | |
| | H65 | F | 62 | | | IV | durvalumab/tremelimumab | | |
| Ma et al 2019 | C66 | F | 71 | | | IV | pembrolizumab | | |
| | P1 | | | | | | | | |
| | P2 | | | | | | | | |
| | P3 | | | | | | | | |
| | P4 | N/A | N/A | Metastasis | Uveal Melanoma | N/A | N/A | 10X-Chromium | 2274 |
| | P5 | | | | | | | | |
| | P6 | | | | | | | | |
| Nieto et al. 2020* | P7 | | | | | | | | |
| | LC_1 | F | 70-75 | | Squamous cell carcinoma | IIA | | | |
| | LC_2 | M | 86-90 | | Squamous cell carcinoma | IB | | | |
| | LC_3 | M | 66-70 | | Adenocarcinoma | IIIB | | | |
| | LC_4 | F | 60-65 | | Adenocarcinoma | IIB | | | |
| | LC_5 | M | 60-65 | | Large cell carcinoma | IA3 | | | |
| | LC_6 | M | 60-65 | | Adenocarcinoma | IIIA | | | |
| | LC_7 | M | 60-65 | | Squamous cell carcinoma | IB | | | |
| | LC_8 | F | 50-55 | | Pleiomorphic carcinoma | IIB | | | |
| | OvC_1 | | 70-75 | | HGSOC | IIIC | | | |
| | OvC_2 | | 50-55 | | HGSOC | IVB | | | |
| | OvC_3 | F | 60-65 | | HGSOC | IVB | | | |
| | OvC_4 | | 80-85 | | HGSOC + clear cell carcinoma (mix) | IVB | | | |
| | OvC_5 | | 60-65 | | HGSOC | IA | | | |

| | | | | | | | | | |
|---|---|---|---|---|---|---|---|---|---|
| | CRC_1 | F | 80-85 | | Right caecum, global moderately differentiated adenocarcinoma with mixed glandulair, mucinous growth pattern, moderate budding | IIB | | | |
| | CRC_2 | F | 86-90 | | Left rectosigmoid, moderately differentiated adenocarcinoma NST | IIIB | | | |
| | CRC_3 | F | 50-55 | | Left sigmoid, moderately differentiated adenocarcinoma | IVA | | | |
| | CRC_4 | M | 80-85 | Tumor | Left sigmoid, moderately differentiated adenocarcinoma | I | Naïve | 10X-Chromium | 155819 |
| | CRC_5 | M | 50-55 | | Left sigmoid, moderately differentiated adenocarcinoma | IIA | | | |
| | CRC_6 | F | 76-80 | | Right caecum,moderately differentiated adenocarcinoma NOS | IIIB | | | |
| | CRC_7 | M | 80-85 | | Right ascending,moderately differentiated adenocarcinoma | IIA | | | |
| | BC_1 | | 40-45 | | Invasive ductal carcinoma | | | | |
| | BC_2 | | 50-55 | | Invasive ductal carcinoma | | | | |
| | BC_3 | | 50-55 | | Invasive ductal carcinoma | | | | |
| | BC_4 | | 86-90 | | Invasive ductal carcinoma | | | | |
| | BC_5 | | 60-65 | | Invasive apocrine carcinoma | | | | |
| | BC_6 | | 70-75 | | Invasive ductal carcinoma | III | | | |
| | BC_7 | | 80-85 | | Metaplastic carcinoma | | | | |
| | BC_8 | F | 60-65 | | | | | | |
| | BC_9 | | 40-45 | | | | | | |
| | BC_10 | | 50-55 | | | | | | |
| | BC_11 | | 46-50 | | | | | | |
| | BC_12 | | 76-80 | | Invasive ductal carcinoma | | | | |
| | BC_13 | | 60-65 | | | | | | |
| | BC_14 | | 46-50 | | | II | | | |
| Qian et al | BC_15 | | 66-70 | | | III | | | |
| 2020 | BC_16 | | 60-65 | | | III | | | |
| | Donor 1 | | 25 | | | | | | |
| | Donor 2 | | 27 | | | | | | |
| | Donor 3 | M | 53 | Normal | Skin | N/A | N/A | 10X-Chromium | 14754 |

| | | | | | | | | | |
|---|---|---|---|---|---|---|---|---|---|
| Solé-Boldo et. al | Donor 4 | | 70 | | | | | | |
| 2020 | Donor 5 | | 69 | | | | | | |
| Zheng et al | Donor A | N/A | N/A | Normal | PBMC | N/A | N/A | 10X-Chromium | 2661 |
| | NSC004 | M | 79 | | Lung (Squamous) | | Naïve | | |
| | NSC009 | F | 74 | | Lung (Squamous) | | Chemo and XRT | | |
| | NSC016 | F | 61 | | Lung (Adeno) | | Naïve | | |
| | NSC018 | M | 83 | Tumor | Lung (Adeno) | | Naïve | inDrop | 52577 |
| | NSC019 | F | 72 | | Lung (Adeno) | | Naïve | | |
| | NSC020 | M | 76 | | Lung (Adeno) | | Naïve | | |
| Zilionis et al 2019 | NSC021 | F | 63 | | Lung (Adeno) | | Naïve | | |

*Download in 2021/03
