## Supplementary Table 3 for "Identification of novel myeloid-derived cell states with implication in cancer outcome"

**Supplementary Table 3: Canonical gene markers**

| Cell Type | ***Gene Markers*** | Reference |
| --- | --- | --- |
| T lymphocytes and Natural Killer | *CD3E, CD4 or CD8A,* and *NKG7* | (Solé-Boldo et al. 2020) (Qian et al. 2020) (Schelker et al. 2017) |
| Mononuclear phagocytes | *LYZ, AIF1,* and *CD68* | (Solé-Boldo et al. 2020) (Tirosh et al. 2016) |
| Neutrophils | *CXCR2* and *FCGR3B* | (Schupp et al. 2020) (Salcher et al. 2022) |
| Mast cells | *TPSAB1* and *CPA3* | (Gentles et al. 2020) (Chen et al. 2020) |
| Megakaryocytes | *PPBP* | (Kammers et al. 2021) |
| B lymphocytes | *MS4A1* and *MZB1* | (Izar et al. 2020) (Schelker et al. 2017) |
| Endothelial cells | *VWF* and *CCL21* | (Solé-Boldo et al. 2020) |
| Fibroblasts | *COL1A1* and *COL1A2* | (Muhl et al. 2020) |
| Plasmacytoid dendritic cells | *LILRA4* and *IRF7* | (Qian et al. 2020) (Schelker et al. 2017) |
| Normal and malignant cells | *EPCAM* and *CDH1* | (Qian et al. 2020) (Izar et al. 2020) (Schelker et al. 2017) |

**Supplementary Table 3: Canonical gene markers**

| Cell Type | Possible phenotype | Gene Markers | Reference |
|---|---|---|---|
| Neutrophil_TAGLN2 | Neutrophils, proinflammatory | *TAGLN2* | |
| Neutrophil_MMP9 | Neutroplhils, anti-tumoral | *MMP9* | (Sebastian et al. 2022; Hong et al. 2022) |
| Neutrophil_CXCL8 | Neutroplhils, anti-inflammatory | *CXCL8* | (Hong et al. 2022) |
| cDC1_CLEC9A | cDC type 1, proinflammatory | *CLEC9A* and *CADM1* | (Villani et al. 2017; Ginhoux, Guilliams e Merad, 2022) |
| cDC2A_AREG | cDC type 2A, anti-inflammatory | *CD1C, CLEC10A, AREG* | (Brown et al. 2019; Shin et al. 2020) |
| cDC2B_FCER1A | cDC type 2B, proinflammatory | *CD1C, CLEC10A, FCER1A* | (Brown et al. 2019; Shin et al. 2020) |
| cDC2_CD14 | cDC type 2, proinflammatory | *CD1C* and *CD14* | (Villani et al. 2017; Cabeza-Cabrerizo et al. 2021) |
| cDC2_FCGR3A | cDC type 2, immunosuppressive process | *CD1C* and *FCGR3A* | (Villani et al. 2017; Cabeza-Cabrerizo et al. 2021) |
| cDC2_CXCL8 | cDC type 2, proinflammatory | *G0S2, RIT1, SELK, INSIG1,* and *CXCL8* | (Bourdely et al. 2019) |
| cDC_CD207 | cDC type 2, anti- and pro-inflammatory | *CD1A* and *CD207* | (Collin et al. 2017) |
| cDC_LAMP3 | cDC type 2, proinflammatory | *LAMP3 and CCR7* | (Lavin et al. 2017; Zhang, et al. 2019) |
| Mono_FCG3RA | Nonclassical monocyte | *FCGR3A, S100A6,* and *CXCR4* | (Villani et al. 2017) |
| Mono_CD14_FOS- | Classical monocyte | *CD14, SELL,* and *FCN1* | (Villani et al. 2017; McEvoy et al. 2021) |
| Mono_CD14_FOS+ | Classical monocyte | *CD14, SELL, FCN1* and *FOS* | (Villani et al. 2017; McEvoy et al. 2021) |
| Mono_IL1B | Monocytes, proinflammatory | *IL1B, CXCL2,* and *NFKBIA* | (Zhang et al. 2018) |
| MonoInter_CXCL10 | Intermediate monocyte | *CD16, CD14, CXCL10* | (Villani et al. 2017; Metcalf et al. 2017; Rizzo et al. 2020) |
| MonoInter_CLEC10A | Intermediate monocyte | *CD16, CD14* and *CLEC10A* | (Villani et al. 2017; Rizzo et al 2020) |
| Mac_Alv-like | Macrophage | *FABP4, CRIP1, MCEMP1* | (Liao et al. 2020ç Qian et al. 2020) |

| | | | |
|---|---|---|---|
| Mac_Angio | Macrophage, pro-angiogenic | *VCAN, TIMP1* and *EREG* | (Ma, Black e Qian, 2022) |
| RTM_IFN | Resident macrophage, interferon pathways | *CXCL9, IFI27, STAT1* | (Ma, Black e Qian, 2022) |
| Mac_Reg | Macrophage, immunosuppressive and immune regulator | *RGS1, C3* and *CX3CR1* | (Ma, Black e Qian, 2022) |
| Mac_Rec | Macrophage, proinflammatory | *AREG, FCN1* and *BTG1* | (Qian et al., 2020 |
| Mac_IFN | Macrophage, interferon pathways | *CCL2, CXCL10, CCL7* and *CCL8* | (Ma, Black e Qian, 2022) |
| Mac_Hypo | Macrophage, hypoxia | *GOS2, SPP1, C15orf48, CSTB* and *MMP9* | (Bassez et al. 2021) |
| RTM_IM | Resident macrophage, interstitials | *SEPP1,F13A1, SLC40A1, PLTP* and *FOLR2* | (Chakarov et al. 2019) |
| RTM_like_MT | Resident-like macrophage, metallothioneins | *LYVE1, MT2A, MT1X* and *S100A8* | (Bassez et al. 2021) |
| Mac_Prolif | Macrophage | *STMN1 and MKI67* | (Ma, Black e Qian, 2022) |
| RTM_LA | Resident macrophage, suppressive profile | *GPNMB, APOE* and *LGMN* | |
| Mac_LA | Macrophage, suppressive profile | *SPP1, CSTB* and *MMP9* | (Ma, Black e Qian, 2022) |
| NK_rest | Natural killer, resting | *XCL1* and *AREG* | (Qian et al. 2020) |
| NK_cyto | Natural killer, cytotoxic | *GZMB, FGFBP2,* and *FCGR3A* | (Qian et al. 2020) |
| NKT | Natural killer T | *CD8A, CD8B,* and *FGFBP2* | (Zhou et al. 2020; Shen et al. 2020) |
| TCD4_naive | CD4, naïve | *CD4, CCR7, LEF1, SELL* and *TCF7* | (Cano-Gamez et al. 2020; Zhao et al. 2020) |
| TCD8_naive | CD8, naïve | *CD8, IL7R* and *ANXA1* | (Zheng et al. 2017) |
| TCD4_em | T CD4, memory effector | *CD4, L7R, CD40LG* and A*NXA1* | (Cano-Gamez et al. 2020; Li et al. 2019; Szabo et al. 2019) |
| TCD8_em | T CD8, memory effector | *CD8, GZMK, EMOS,* and *ITM2C* | (Andreatta et al. 2021; McLane et al. 2013) |
| TCD4_ex | T CD4, exhausted | *CD4, CTLA4, PDCD1,* and *CD200* | (Miggelbrink et al. 2021) |
| TCD8_ex | T CD4, exhausted | *CD8, LAG3* and *GZMB* | (Andreatta et al. 2021) |

| | | | |
|---|---|---|---|
| TCD4_reg | T regulatory | *CD4, FOXP3* and *CTLA4* | (Cano-Gamez et al. 2020; Li et al. 2019; Szabo et al. 2019) |
| TGD | γδ T lymphocytes | *TRDV1, TRDV2,* and *TRGV9* | (Wang et al.2021) |
