## Supplementary Table 4 for "Identification of novel myeloid-derived cell states with implication in cancer outcome"

**Supplementary Table 4: Summary of hallmark gene signatures**

| Gene Signature | Gene | Reference |
|---|---|---|
| Hallmark Inflammatory Response | ABCA1, ABI1, ACVR1B, ACVR2A, ADM, ADORA2B, ADRM1, AHR, APLNR, AQP9, ATP2A2, ATP2B1, ATP2C1, AXL, BDKRB1, BEST1, BST2, BTG2, C3AR1, C5AR1, CALCRL, CCL17, CCL2, CCL20, CCL22, CCL24, CCL5, CCL7, CCR7, CCRL2, CD14, CD40, CD48, CD55, CD69, CD70, CD82, CDKN1A, CHST2, CLEC5A, CMKLR1, CSF1, CSF3, CSF3R, CX3CL1, CXCL10, CXCL11, CXCL6, CXCL9, CXCR6, CYBB, DCBLD2, EBI3, EDN1, EIF2AK2, EMP3, ADGRE1, EREG, F3, FFAR2, FPR1, FZD5, GABBR1, GCH1, GNA15, GNAI3, GP1BA, GPC3, GPR132, GPR183, HAS2, HBEGF, HIF1A, HPN, HRH1, ICAM1, ICAM4, ICOSLG, IFITM1, IFNAR1, IFNGR2, IL10, IL10RA, IL12B, IL15, IL15RA, IL18, IL18R1, IL18RAP, IL1A, IL1B, IL1R1, IL2RB, IL4R, IL6, IL7R, CXCL8, INHBA, IRAK2, IRF1, IRF7, ITGA5, ITGB3, ITGB8, KCNA3, KCNJ2, KCNMB2, KIF1B, KLF6, LAMP3, LCK, LCP2, LDLR, LIF, LPAR1, LTA, LY6E, LYN, MARCO, MEFV, MEP1A, MET, MMP14, MSR1, MXD1, MYC, NAMPT, NDP, NFKB1, NFKBIA, NLRP3, NMI, NMUR1, NOD2, NPFFR2, OLR1, OPRK1, OSM, OSMR, P2RX4, P2RX7, P2RY2, PCDH7, PDE4B, PDPN, PIK3R5, PLAUR, PROK2, PSEN1, PTAFR, PTGER2, PTGER4, PTGIR, PTPRE, PVR, RAF1, RASGRP1, RELA, RGS1, RGS16, RHOG, RIPK2, RNF144B, ROS1, RTP4, SCARF1, SCN1B, SELE, SELL, SELENOS, SEMA4D, SERPINE1, SGMS2, SLAMF1, SLC11A2, SLC1A2, SLC28A2, SLC31A1, SLC31A2, SLC4A4, SLC7A1, SLC7A2, SPHK1, SRI, STAB1, TACR1, TACR3, TAPBP, TIMP1, TLR1, TLR2, TLR3, TNFAIP6, TNFRSF1B, TNFRSF9, TNFSF10, TNFSF15, TNFSF9, TPBG, VIP | MsigDB |
| Angiogenesis | APP, CCND2, CXCL6, ITGAV, JAG1, OLR1, PTK2, SLCO2A1, SPP1, THBD, TIMP1, VAV2, VCAN, VEGFA | MsigDB |
| Hypoxia | ADM, ADORA2B, AK4, AKAP12, ALDOA, ALDOB, ALDOC, AMPD3, ANGPTL4, ANKZF1, ANXA2, ATF3, ATP7A, B3GALT6, B4GALNT2, BCAN, BCL2, BGN, BHLHE40, BNIP3L, BRS3, BTG1, CA12, CASP6, CAV1, CCNG2, CCRN4L, CDKN1A, CDKN1B, CDKN1C, CHST2, CHST3, CITED2, COL5A1, CP, CSRP2, CTGF, CXCR4, CXCR7, CYR61, DCN, DDIT3, DDIT4, DPYSL4, DTNA, DUSP1, EDN2, EFNA1, EFNA3, EGFR, ENO1, ENO2, ENO3, ERO1L, ERRFI1, ETS1, EXT1, F3, FAM162A, FBP1, FOS, FOSL2, FOXO3, GAA, GALK1, GAPDH, GAPDHS, GBE1, GCK, GCNT2, GLRX, GPC1, GPC3, GPC4, GPI, GRHPR, GYS1, HAS1, HDLBP, HEXA, HK1, HK2, HMOX1, HOXB9, HS3ST1, HSPA5, IDS, IER3, IGFBP1, IGFBP3, IL6, ILVBL, INHA, IRS2, ISG20, JMJD6, JUN, KDELR3, KDM3A, KIF5A, KLF6, KLF7, KLHL24, LALBA, LARGE, LDHA, LDHC, LOX, LXN, MAFF, MAP3K1, MIF, MT1E, MT2A, MXI1, MYH9, NAGK, NCAN, NDRG1, NDST1, NDST2, NEDD4L, NFIL3, NR3C1, P4HA1, P4HA2, PAM, PCK1, PDGFB, PDK1, PDK3, PFKFB3, PFKL, PFKP, PGAM2, PGF, PGK1, PGM1, PGM2, PHKG1, PIM1, PKLR, PKP1, PLAC8, PLAUR, PLIN2, PNRC1, PPARGC1A, PPFIA4, PPP1R15A, PPP1R3C, PRDX5, PRKCA, PRKCDBP, PTRF, PYGM, RBPJ, RORA, RRAGD, S100A4, SAP30, SCARB1, SDC2, SDC3, SDC4, | MsigDB |
| Epithelial-Mesenchymal Transition | ABI3BP, ACTA2, ADAM12, BGN, BMP1, CADM1, CALD1, CAP2, CCN1, CCN2, CDH11, CDH2, CDH6, COMP, CTHRC1, CXCL1, CXCL12, CXCL8, DAB2, DCN, DKK1, DPYSL3, DST, ECM2, EFEMP2, ELN, FAP, FBLN1, FBLN2, FBLN, FBN1, FBN2, FERMT2, FGF2, FMOD, FSTL1, FSTL3, FUCA1, FZD8, GADD4A, GADD4B, GAS1, GEM, GJA1, GLIPR1, GPC1, HTRA1, ID2, IGFBP3, IGFBP4, ITGAV, JUN, LAMA2, LAMA3, LAMC2, LOX, LOXL1, LOXL2, LRP1, LRRC1, LUM, MAGEE1, MATN2, MATN3, MCM7, MEST, MFAP, MGP, MMP14, MMP2, MMP3, MSX1, MXRA, MYL9, MYLK, NID2, NNMT, NTM, P3H1, PCOLCE, PDGFRB, PFN2, PMEPA1, POSTN, PRRX1, PRSS2, PTHLH, RGS4, RHOB, SAT1, SERPINE2, SFRP1, SFRP4, SGCB, SGCD, SGCG, SLIT2, SLIT3, SNAI2, SPARC, SPOCK1, TAGLN, TFPI2, TGFB1, TGFBR3, THBS2, THY1, TIMP3, TNC, TNFAIP3, TPM1, TPM2, VCAM1, VEGFA, VEGFC, WIPF1 | MsigDB |
| Extracellular Matrix Signature in Macrophages | SDC4, PLXDC2, CLEC2D, LGALS3, ANXA1, PLXNA1, SDC1, SFTPA2, SEMA7A, PLXNB2, ANXA7, LGALS9C, PLXNA3, SEMA3C, LGALS8, LGALS1, LGALS9, ANXA11, ANXA6, PLXNC1, CLEC5A, GREM1, ELFN1, CLEC11A, ANXA4, ANXA2, ANXA5, GPC4, PLXND1, | Etich et al. 2019 |
| Antigen Presentation | ACTR1A, ACTR1B, AP1B1, AP1M2, AP2B1, AP2M1, CAPZB, CD74, CTSF, CTSH, CTSO, DCTN2, DNM3, DYNC1I1, DYNC1I2, DYNC1LI1, HLA-DMA, HLA-DMB, HLA-DOA, HLA-DOB, HLA-DPA1, HLA-DPB1, HLA-DQA1, HLA-DQA2, HLA-DQB1, HLA-DQB2, HLA-DRA, HLA-DRB1, HLA-DRB3, HLA-DRB4, HLA-DRB5, KIF26A, KIF2A, KIF3B, KIF3C, KIF4B, KIF5B, KIFAP3, KLC1, KLC4, SAR1B, SEC13, SPTBN2, TUBA1A, TUBA1B, TUBA3C, TUBA3D, TUBA4A, TUBA4B, TUBA8, TUBB2A, TUBB2B, TUBB3, TUBB4A, TUBB4B, TUBB8, TUBB8B | MsigDB |
| Phagocytosis | AKT3, PLA2G4B, DNM1L, WASF2, VAV3, ARPC1A, CFL2, WASF3, PLA2G4E, CRK, CRKL, DNM1, DNM2, DOCK2, PIKFYVE, AKT1, AKT2, FCGR1A, FCGR2A, FCGR2B, FCGR3A, PIP5K1C, PIK3R5, DNM3, AMPH, PLA2G4D, HCK, INPP5D, LIMK2, LYN, MARCKS, MYO10, PAK1, ASAP1, PIK3CA, PIK3CB, PIK3CD, PIK3CG, PIK3R1, PIK3R2, PLA2G4A, PLCG1, PLCG2, PLD1, PLD2, PRKCA, PRKCG, MAPK1, MAPK3, SPHK2, PTPRC, RAC1, RAF1, RPS6KB1, NCF1, SYK, VAV1, VAV2, WAS, PIP5K1A, PIP5K1B, PIP4K2B, PLA2G6, PIK3R3, SCIN, PLPP1, ASAP2, WASL, FCGR2C, GAB2 | MsigDB |
