## Supplementary Table 5 for "Identification of novel myeloid-derived cell states with implication in cancer outcome"

**Supplementary Table 5: Enrichment pathway analysis**

**Mast cells**

| Cluster | ID | Description | GeneRatio | BgRatio | pvalue | p.adjust | qvalue | geneID | Count |
|---|---|---|---|---|---|---|---|---|---|
| normal | R-HSA-72706 | GTP hydrolysis and joining of the 60S ribosomal subunit | 97/1399 | 112/10704 | 2.18522929593212e-70 | 2.92602202725311e-67 | 1.98280805588788e-67 | 6136/6230/6154/1973/6222/6144/6159/6170/6193/6128/6166/3646/6147/6142/25873/6209/11224/51065/6231/6203/6223/6146/6171/6132/7311/8664/6157/6235/6207/6201/6155/6210/4736/6161/6152/27335/6217/9045/6165/6129/6133/6141/51386/3921/6139/6208/6224/6138/6156/6158/6188/8667/6227/6160/6181/6 | 97 |
| normal | R-HSA-156827 | L13a-mediated translational silencing of Ceruloplasmin expression | 96/1399 | 111/10704 | 1.53897100492482e-69 | 1.03034108779716e-66 | 6.98206845392206e-67 | 6136/6230/6154/1973/6222/6144/6159/6170/6193/6128/6166/3646/6147/6142/25873/6209/11224/51065/6231/6203/6223/6146/6171/6132/7311/8664/6157/6235/6207/6201/6155/6210/4736/6161/6152/27335/6217/9045/6165/6129/6133/6141/51386/3921/6139/6208/6224/6138/6156/6158/6188/8667/6227/6160/6181/6 | 96 |
| normal | R-HSA-72613 | Eukaryotic Translation Initiation | 98/1399 | 119/10704 | 9.09660948778127e-67 | 3.04509002603478e-64 | 2.06349404696512e-64 | 6136/6230/6154/1973/6222/6144/6159/6170/6193/6128/6166/3646/6147/6142/25873/6209/11224/51065/6231/6203/6223/6146/6171/6132/7311/8664/6157/6235/6207/6201/6155/6210/4736/6161/6152/27335/6217/9045/6165/6129/6133/6141/51386/3921/6139/6208/6224/6138/6156/6158/6188/8667/6227/6160/6181/6 | 98 |
| normal | R-HSA-72737 | Cap-dependent Translation Initiation | 98/1399 | 119/10704 | 9.09660948778127e-67 | 3.04509002603478e-64 | 2.06349404696512e-64 | 6136/6230/6154/1973/6222/6144/6159/6170/6193/6128/6166/3646/6147/6142/25873/6209/11224/51065/6231/6203/6223/6146/6171/6132/7311/8664/6157/6235/6207/6201/6155/6210/4736/6161/6152/27335/6217/9045/6165/6129/6133/6141/51386/3921/6139/6208/6224/6138/6156/6158/6188/8667/6227/6160/6181/6 | 98 |
| normal | R-HSA-72689 | Formation of a pool of free 40S subunits | 88/1399 | 101/10704 | 2.23802787394266e-64 | 5.99343864641843e-62 | 4.06143163650225e-62 | 6136/6230/6154/6222/6144/6159/6170/6193/6128/6166/3646/6147/6142/25873/6209/11224/51065/6231/6203/6223/6146/6171/6132/7311/8664/6157/6235/6207/6201/6155/6210/4736/6161/6152/27335/6217/9045/6165/6129/6133/6141/51386/3921/6139/6208/6224/6138/6156/6158/6188/8667/6227/6160/6181/6175/6 | 88 |
| normal | R-HSA-156902 | Peptide chain elongation | 80/1399 | 89/10704 | 5.2710947327034e-61 | 1.14827840603813e-58 | 7.78126634963144e-59 | 6136/6230/6154/6222/6144/6159/6170/6193/6128/6166/6147/6142/25873/6209/11224/51065/6231/6203/6223/6146/6171/6132/7311/6157/6235/6207/6201/6155/6210/4736/6161/6152/6217/9045/6165/6129/6133/6141/3921/6139/6208/6224/6138/6156/6158/6188/6227/6160/6181/6175/6234/6192/6228/6143/6187/61 | 80 |
| normal | R-HSA-156842 | Eukaryotic Translation Elongation | 82/1399 | 93/10704 | 6.00294909803353e-61 | 1.14827840603813e-58 | 7.78126634963144e-59 | 1937/6136/6230/6154/6222/6144/6159/6170/6193/6128/6166/6147/6142/25873/6209/11224/51065/6231/6203/6223/6146/6171/6132/7311/6157/6235/6207/6201/6155/6210/4736/6161/1936/6152/6217/9045/6165/6129/6133/6141/3921/6139/6208/6224/6138/6156/6158/6188/6227/6160/6181/6175/6234/6192/6228/614 | 82 |
| normal | R-HSA-72766 | Translation | 152/1399 | 291/10704 | 9.9537200545829e-60 | 1.66600389413581e-57 | 1.12896140619085e-57 | 1937/116540/6136/6230/6154/1973/6222/6144/6159/6170/6193/6748/6128/6166/3646/6727/6147/6142/25873/6209/11224/6150/51065/6231/6203/6223/6146/6171/6132/7311/8664/6157/6235/6207/51504/6201/6155/6210/4736/6161/1936/6152/27335/6217/28972/9045/6165/6129/6133/6141/51386/3921/6139/90480/ | 152 |
| normal | R-HSA-192823 | Viral mRNA Translation | 79/1399 | 89/10704 | 2.97569173818811e-59 | 4.4271680415932e-57 | 3.0000541266879e-57 | 6136/6230/6154/6222/6144/6159/6170/6193/6128/6166/6147/6142/25873/6209/11224/51065/6231/6203/6223/6146/6171/6132/7311/6157/6235/6207/6201/6155/6210/4736/6161/6152/6217/9045/6165/6129/6133/6141/3921/6139/6208/6224/6138/6156/6158/6188/6227/6160/6181/6175/6234/6192/6228/6143/6187/61 | 79 |
| normal | R-HSA-1799339 | SRP-dependent cotranslational protein targeting to membrane | 89/1399 | 112/10704 | 4.26005760461267e-58 | 5.70421713257637e-56 | 3.86544174229066e-56 | 6136/6230/6154/6222/6144/6159/6170/6193/6748/6128/6166/6727/6147/6142/25873/6209/11224/51065/6231/6203/6223/6146/6171/6132/7311/6157/6235/6207/6201/6155/6210/4736/6161/6152/6217/28972/9045/6165/6129/6133/6141/3921/6139/6208/6224/6138/6156/6158/6188/9789/6227/6160/6181/6175/6234/61 | 89 |
| normal | R-HSA-2408557 | Selenocysteine synthesis | 79/1399 | 93/10704 | 5.13957963302805e-56 | 5.7349142738538e-54 | 3.88624354707911e-54 | 6136/6230/6154/6222/6144/6159/6170/6193/6128/6166/6147/6142/25873/6209/11224/51065/6231/6203/6223/6146/6171/6132/7311/6157/6235/6207/6201/6155/6210/4736/6161/6152/6217/9045/6165/6129/6133/6141/3921/6139/6208/6224/6138/6156/6158/6188/6227/6160/6181/6175/6234/6192/6228/6143/6187/61 | 79 |
| normal | R-HSA-72764 | Eukaryotic Translation Termination | 79/1399 | 93/10704 | 5.13957963302805e-56 | 5.7349142738538e-54 | 3.88624354707911e-54 | 6136/6230/6154/6222/6144/6159/6170/6193/6128/6166/6147/6142/25873/6209/11224/51065/6231/6203/6223/6146/6171/6132/7311/6157/6235/6207/51504/6201/6155/6210/4736/6161/6152/6217/9045/6165/6129/6133/6141/3921/6139/6208/6224/6138/6156/6158/6188/6227/6160/6181/6175/6234/6192/6228/6143/61 | 79 |
| normal | R-HSA-9010553 | Regulation of expression of SLITs and ROBOs | 110/1399 | 171/10704 | 7.88610635600989e-56 | 8.12268954669019e-54 | 5.50431067115832e-54 | 6136/6230/6154/5687/6222/6144/6159/6170/6193/6128/6166/6147/6142/25873/6209/11224/51065/6231/6203/6223/6146/6171/6132/7311/6157/6235/5689/6207/5699/5700/6201/6155/6210/4736/6161/6152/6217/9045/6165/6129/6133/6141/3921/6139/7316/5685/6208/6224/6138/6156/9978/6158/6188/6227/6160/61 | 110 |
| normal | R-HSA-9633012 | Response of EIF2AK4 (GCN2) to amino acid deficiency | 82/1399 | 101/10704 | 5.63631825129573e-55 | 5.39073581320355e-53 | 3.65301228016309e-53 | 6136/6230/6154/6222/6144/6159/6170/6193/6128/6166/6147/6142/25873/6209/11224/51065/6231/6203/6223/6146/6171/6132/7311/6157/6235/6207/6201/6155/6210/4736/6161/6152/6217/9045/6165/6129/6133/6141/3921/6139/6208/6224/6138/6156/6158/6188/6227/6160/6181/6175/1051/6234/6192/6228/6143/61 | 82 |
| normal | R-HSA-975956 | Nonsense Mediated Decay (NMD) independent of the Exon Junction Complex (EJC) | 79/1399 | 95/10704 | 1.47149041043651e-54 | 1.31355043971632e-52 | 8.9012262020 7908e-53 | 6136/6230/6154/6222/6144/6159/6170/6193/6128/6166/6147/6142/25873/6209/11224/51065/6231/6203/6223/6146/6171/6132/7311/6157/6235/6207/6201/6155/6210/4736/6161/6152/6217/9045/6165/6129/6133/6141/3921/6139/6208/6224/6138/6156/6158/6188/6227/6160/6181/6175/6234/6192/6228/6143/6187/61 | 79 |
| normal | R-HSA-927802 | Nonsense-Mediated Decay (NMD) | 84/1399 | 115/10704 | 1.19028178419703e-49 | 9.37521946494017e-48 | 6.35308295961513e-48 | 6136/6230/6154/6222/6144/6159/6170/6193/6128/6166/6147/6142/25873/6209/11224/51065/6231/6203/6223/6146/6171/6132/7311/6157/6235/6207/6201/6155/6210/4736/6161/6152/6217/9045/6165/6129/6133/6141/3921/6139/6208/6224/6138/6156/6158/6188/6227/6160/6181/6175/6234/6192/4116/6228/6143/61 | 84 |
| normal | R-HSA-975957 | Nonsense Mediated Decay (NMD) enhanced by the Exon Junction Complex (EJC) | 84/1399 | 115/10704 | 1.19028178419703e-49 | 9.37521946494017e-48 | 6.35308295961513e-48 | 6136/6230/6154/6222/6144/6159/6170/6193/6128/6166/6147/6142/25873/6209/11224/51065/6231/6203/6223/6146/6171/6132/7311/6157/6235/6207/6201/6155/6210/4736/6161/6152/6217/9045/6165/6129/6133/6141/3921/6139/6208/6224/6138/6156/6158/6188/6227/6160/6181/6175/6234/6192/4116/6228/6143/61 | 84 |
| normal | R-HSA-376176 | Signaling by ROBO receptors | 118/1399 | 218/10704 | 1.79843550257203e-48 | 1.33783618774664e-46 | 9.06579767963209e-47 | 6136/6230/6154/5687/6222/6144/6159/6170/6193/6128/6166/5216/6147/6142/25873/6209/11224/51065/6231/6203/6223/6146/6171/6132/7311/6157/6235/5689/6207/5699/5700/6201/6155/6210/4736/6161/6152/6217/9045/6165/6129/6133/6141/3921/6139/7316/5685/6208/6224/6138/6156/5879/998/9978/6158/618 | 118 |
| normal | R-HSA-168273 | Influenza Viral RNA Transcription and Replication | 88/1399 | 135/10704 | 1.81760936844804e-45 | 1.28093628650101e-43 | 8.68021759336404e-44 | 6136/6230/6154/6222/6144/6159/6170/6193/6128/6166/6147/6142/25873/6209/11224/51065/6231/6203/6223/6146/6171/6132/7311/6157/6235/6207/6201/6155/6210/4736/6161/6152/6217/9045/6165/6129/6133/6141/3921/6139/6208/6224/6138/6156/6158/6188/6227/6160/6181/6175/6234/6192/6228/6143/6187/61 | 88 |
| normal | R-HSA-168255 | Influenza Infection | 94/1399 | 156/10704 | 3.59206287487816e-44 | 2.40488609473093e-42 | 1.62966220954999e-42 | 6136/6230/6154/6222/6144/6159/6170/6193/6128/6166/6147/6142/25873/6209/11224/51065/6231/6203/6223/6146/6171/6132/7311/6157/6235/6207/6201/6155/6210/4736/6161/6152/6217/9045/6165/6129/6133/6141/293/3921/6139/6208/6224/6138/6156/6158/6188/6227/6160/6181/6175/6234/6192/6228/6143/6187 | 94 |
| normal | R-HSA-2408522 | Selenoamino acid metabolism | 80/1399 | 118/10704 | 2.32112319416959e-43 | 1.47999236047289e-41 | 1.0029113751249e-41 | 6136/6230/6154/6222/6144/6159/6170/6193/6128/6166/6147/6142/25873/6209/11224/51065/6231/6203/6223/6146/6171/6132/7311/6157/6235/6207/6201/6155/6210/4736/6161/6152/6217/9045/6165/6129/6133/6141/3921/6139/6208/6224/6138/6156/6158/6188/6227/6160/6181/6175/6234/6192/6228/6143/6187/61 | 80 |
| normal | R-HSA-8868773 | rRNA processing in the nucleus and cytosol | 101/1399 | 194/10704 | 1.54531251931387e-39 | 9.40533392436941e-38 | 6.37348991219404e-38 | 6136/6230/6154/6222/6144/6159/6170/6193/6128/6166/6147/6142/25873/6209/11224/51065/6231/6203/6223/6146/6171/6132/7311/6157/6235/6207/51504/6201/6155/6210/4736/6161/6152/6217/9045/6165/6129/6133/6141/3921/6139/6208/6224/6138/6156/6158/6188/6227/6160/6181/6175/6234/6192/6228/6143/61 | 101 |
| normal | R-HSA-6791226 | Major pathway of rRNA processing in the nucleolus and cytosol | 98/1399 | 184/10704 | 1.6371519847612e-39 | 9.53107177215326e-38 | 6.45869570189545e-38 | 6136/6230/6154/6222/6144/6159/6170/6193/6128/6166/6147/6142/25873/6209/11224/51065/6231/6203/6223/6146/6171/6132/7311/6157/6235/6207/6201/6155/6210/4736/6161/6152/6217/9045/6165/6129/6133/6141/3921/6139/6208/6224/6138/6156/6158/6188/6227/6160/6181/6175/6234/6192/6228/6143/6187/61 | 98 |
| normal | R-HSA-72312 | rRNA processing | 103/1399 | 204/10704 | 9.66413400310396e-39 | 5.39178142923175e-37 | 3.65372083801562e-37 | 6136/6230/6154/6222/6144/6159/6170/6193/6128/6166/6147/6142/25873/6209/11224/51065/6231/6203/6223/6146/6171/6132/7311/6157/6235/6207/51504/6201/6155/6210/4736/6161/6152/6217/9045/6165/6129/6133/6141/3921/6139/6208/6224/6138/6156/6158/6188/6227/6160/6181/6175/6234/6192/6228/6143/61 | 103 |
| normal | R-HSA-72649 | Translation initiation complex formation | 49/1399 | 58/10704 | 7.66925203681312e-35 | 3.94966479895876e-33 | 2.67647581203761e-33 | 6230/1973/6222/6193/3646/6209/51065/6231/6203/6223/8664/6235/6207/6201/6210/27335/6217/51386/3921/6208/6224/6188/8667/6227/6234/6192/6228/1974/6187/6194/6229/6189/6191/6233/8665/6232/6206/8666/8668/6202/1968/7458/8894/10480/1964/1965/6205/26986/1975 | 49 |
| normal | R-HSA-72702 | Ribosomal scanning and start codon recognition | 49/1399 | 58/10704 | 7.66925203681312e-35 | 3.94966479895876e-33 | 2.67647581203761e-33 | 6230/1973/6222/6193/3646/6209/51065/6231/6203/6223/8664/6235/6207/6201/6210/27335/6217/51386/3921/6208/6224/6188/8667/6227/6234/6192/6228/1974/6187/6194/6229/6189/6191/6233/8665/6232/6206/8666/8668/6202/1968/7458/8894/10480/1983/1964/1965/6205/1975 | 49 |
| normal | R-HSA-72662 | Activation of the mRNA upon binding of the cap-binding complex and eIFs, and subsequent binding to 43S | 49/1399 | 59/10704 | 3.96284978824481e-34 | 1.96527995054067e-32 | 1.33176472415868e-32 | 6230/1973/6222/6193/3646/6209/51065/6231/6203/6223/8664/6235/6207/6201/6210/27335/6217/51386/3921/6208/6224/6188/8667/6227/6234/6192/6228/1974/6187/6194/6229/6189/6191/6233/8665/6232/6206/8666/8668/6202/1968/7458/8894/10480/1964/1965/6205/26986/1975 | 49 |
| normal | R-HSA-72695 | Formation of the ternary complex, and subsequently, the 43S complex | 44/1399 | 51/10704 | 3.29419300055103e-32 | 1.57533015276351e-30 | 1.06751667912593e-30 | 6230/6222/6193/3646/6209/51065/6231/6203/6223/8664/6235/6207/6201/6210/27335/6217/51386/3921/6208/6224/6188/8667/6227/6234/6192/6228/6187/6194/6229/6189/6191/6233/8665/6232/6206/8666/8668/6202/1968/8894/10480/1964/1965/6205 | 44 |
| normal | R-HSA-72163 | mRNA Splicing - Major Pathway | 89/1399 | 183/10704 | 5.77474346591933e-32 | 2.66633844857447e-30 | 1.8068344347087e-30 | 7307/4904/56259/6430/6633/10949/6637/6428/6635/3178/8683/4116/3188/51691/10914/4670/220988/6628/51639/3181/11157/27258/22827/27316/3190/3192/1655/10250/5440/5441/10286/4809/3185/25949/6431/25804/9410/22826/51503/6632/10084/84991/51729/22938/85313/23350/23020/6629/6432/51690/834 | 89 |
| normal | R-HSA-72172 | mRNA Splicing | 91/1399 | 191/10704 | 8.81911698284831e-32 | 3.9362658800113e-30 | 2.66739608393517e-30 | 7307/4904/56259/6430/6633/10949/6637/6428/6635/3178/8683/4116/3188/51691/10914/4670/220988/6628/51639/3181/11157/27258/22827/27316/3190/3192/1655/10250/5440/5441/10286/4809/3185/25949/6431/25804/9410/22826/51503/6632/10084/84991/51729/22938/85313/23350/23020/6629/6432/51690/834 | 91 |
| normal | R-HSA-611105 | Respiratory electron transport | 61/1399 | 101/10704 | 4.48185329563539e-29 | 1.93587147188897e-27 | 1.31183617685491e-27 | 51079/4697/7381/1329/1337/4707/1327/4729/54205/4725/25915/1340/1345/1349/126328/9167/4715/4713/4710/4724/27089/1351/7388/1350/7386/4698/4694/4706/4717/55967/4708/6392/374291/4712/2109/4716/4711/54539/4726/6390/7384/1537/4723/29796/4709/84987/29078/55863/6391/4702/4720/4695/90639 | 61 |
| normal | R-HSA-6798695 | Neutrophil degranulation | 151/1399 | 480/10704 | 4.39309791970062e-27 | 1.83823691077473e-25 | 1.24567447591511e-25 | 527/3958/1535/4282/226/2207/10577/26578/8673/2495/6727/3146/1471/966/2950/9535/5689/7305/5269/9588/6282/2512/1992/84418/967/5879/1476/3107/3608/5478/684/5908/6993/2547/10092/9218/5036/5788/391/960/258010/8655/5702/3848/6386/965/10694/5660/5695/6280/4125/7879/5223/29108/3105/5315/1 | 151 |
| normal | R-HSA-71291 | Metabolism of amino acids and derivatives | 128/1399 | 374/10704 | 7.81799453209062e-27 | 3.17221050862707e-25 | 2.1496367740549e-25 | 6136/6230/6154/5687/6222/6144/6159/6170/6193/6128/6166/6147/6142/25873/6209/11224/51065/6231/6203/6223/6146/6171/6132/7311/6157/6235/5689/6207/5699/5700/6201/6155/4946/6210/4736/6161/6152/6217/9045/6165/6129/6133/6141/3921/6139/5685/6208/6224/6138/6156/6158/6188/6227/6160/6181/617 | 128 |
| normal | R-HSA-72203 | Processing of Capped Intron-Containing Pre-mRNA | 98/1399 | 244/10704 | 8.37982588339049e-27 | 3.3001726052529e-25 | 2.23634981779647e-25 | 7307/4904/56259/6430/6633/10949/6637/6428/6635/3178/8683/4116/3188/51691/10914/4670/220988/6628/51639/3181/11157/27258/22827/27316/3190/3192/1655/9589/10250/5440/5441/10286/4809/3185/25949/6431/25804/9410/22826/80145/51503/6632/10084/84991/51729/22938/85313/23350/23020/6629/643 | 98 |
| normal | R-HSA-1428517 | The citric acid (TCA) cycle and respiratory electron transport | 80/1399 | 176/10704 | 2.52586515596478e-26 | 9.66323841096242e-25 | 6.54825793817036e-25 | 51079/4697/7381/1329/1337/4707/1327/4729/54205/4725/25915/1340/1345/1349/126328/9167/4715/4713/3421/4710/7351/4724/3945/27089/1351/7388/1350/7386/4698/4694/4706/4717/55967/3939/4708/6392/374291/51660/4712/2109/4716/4711/54539/1431/4726/6390/7384/1537/682/4723/29796/4709/3420/880 | 80 |
| normal | R-HSA-163200 | Respiratory electron transport, ATP synthesis by chemiosmotic coupling, and heat production by uncoupling proteins. | 62/1399 | 125/10704 | 3.5658402332212e-23 | 1.32629446452311e-21 | 8.98758561706631e-22 | 51079/4697/7381/1329/1337/4707/1327/4729/54205/4725/25915/1340/1345/1349/126328/9167/4715/4713/4710/7351/4724/27089/1351/7388/1350/7386/4698/4694/4706/4717/55967/4708/6392/374291/4712/2109/4716/4711/54539/4726/6390/7384/1537/4723/29796/4709/84987/29078/55863/6391/4702/4720/4695/ | 62 |
| normal | R-HSA-2454202 | Fc epsilon receptor (FCERI) signaling | 58/1399 | 134/10704 | 3.73469606049136e-18 | 1.35155622297241e-16 | 9.15877099329601e-17 | 2205/5687/2207/2353/805/7311/5689/5699/5700/6500/7316/5685/5879/5688/6233/7322/5694/3937/5702/5695/5684/5705/4214/4792/7314/801/5290/5530/2206/5693/7462/5710/9861/5714/808/5686/5708/7334/5713/10768/3551/5701/27040/5718/5690/5594/6464/4067/115727/10213/7321/5058/5717/5719/2534/372 | 58 |
| normal | R-HSA-1236974 | ER-Phagosome pathway | 42/1399 | 83/10704 | 1.61554108571512e-16 | 5.69265661519091e-15 | 3.85760780023943e-15 | 5687/8673/7311/5689/5699/5700/7316/5685/3107/5688/10952/6233/5694/5702/5695/23480/5684/5705/3105/7314/3106/5693/5710/9861/5714/5686/8773/5708/2923/5713/3551/5701/6810/4615/5718/5690/567/811/10213/5717/5719/5707 | 42 |
| normal | R-HSA-1236975 | Antigen processing-Cross presentation | 46/1399 | 98/10704 | 2.48361463993798e-16 | 8.52707693045373e-15 | 5.77834229318904e-15 | 1535/5687/8673/7311/5689/5699/5700/7316/5685/3107/5688/10952/4012/6233/5694/5702/5695/23480/5684/5705/3105/7314/3106/5693/5710/9861/5714/5686/8773/5708/2923/5713/3551/5701/6810/4615/5718/5690/567/811/1514/1520/10213/5717/5719/5707 | 46 |
| normal | R-HSA-1234174 | Cellular response to hypoxia | 39/1399 | 75/10704 | 5.90802985696287e-16 | 1.97771299461832e-14 | 1.34018993071105e-14 | 5687/7311/5689/5699/5700/7316/5685/9978/25994/7323/5688/6233/3091/7322/5694/5702/5695/5684/7422/5705/7314/5693/5710/9861/5714/5686/5708/5713/5701/2034/5718/5690/54583/10213/7321/5717/5719/2033/5707 | 39 |
| normal | R-HSA-1234176 | Oxygen-dependent proline hydroxylation of Hypoxia-inducible Factor Alpha | 36/1399 | 66/10704 | 1.06373066679391e-15 | 3.47398868984645e-14 | 2.35413564769281e-14 | 5687/7311/5689/5699/5700/7316/5685/9978/7323/5688/6233/3091/7322/5694/5702/5695/5684/5705/7314/5693/5710/9861/5714/5686/5708/5713/5701/2034/5718/5690/54583/10213/7321/5717/5719/5707 | 36 |
| normal | R-HSA-9604323 | Negative regulation of NOTCH4 signaling | 32/1399 | 54/10704 | 1.62756910002433e-15 | 5.18884529745853e-14 | 3.51620191534079e-14 | 5687/7311/5689/5699/5700/6500/7316/5685/9978/5688/6233/5694/5702/5695/5684/5705/7314/5693/5710/9861/207/5714/5686/5708/5713/5701/5718/5690/10213/5717/5719/5707 | 32 |
| normal | R-HSA-162909 | Host Interactions of HIV factors | 53/1399 | 130/10704 | 2.33403294575022e-15 | 7.26807003339429e-14 | 4.92518090388419e-14 | 5687/7311/5689/5699/5700/6500/4869/293/7316/5685/5879/9978/5478/5902/5688/5901/6233/8815/5694/1173/5702/5695/5684/5705/3105/7314/5693/5710/9861/1175/8905/375/5714/5686/5708/1025/5713/8907/292/5701/5718/5690/567/53371/5903/10213/130340/3837/5717/5719/2534/9844/5707 | 53 |
| normal | R-HSA-6799198 | Complex I biogenesis | 32/1399 | 55/10704 | 3.40913721784144e-15 | 1.01440771881993e-13 | 6.87409656556565e-14 | 51079/4707/4729/4725/25915/126328/4715/4713/4710/4724/4698/4694/4706/4717/55967/4708/374291/4712/4716/4711/54539/4726/4723/4709/29078/55863/4702/4720/4695/4728/4722/4704 | 32 |
| normal | R-HSA-8941858 | Regulation of RUNX3 expression and activity | 32/1399 | 55/10704 | 3.40913721784144e-15 | 1.01440771881993e-13 | 6.87409656556565e-14 | 5687/7311/5689/5699/5700/7316/5685/5688/6233/5694/5702/5695/5684/5705/7314/5693/864/5710/9861/5714/5686/5708/5713/7040/5701/5718/5690/10213/5717/5719/2033/5707 | 32 |
| normal | R-HSA-2871837 | FCERI mediated NF-kB activation | 40/1399 | 82/10704 | 4.3072604714708e-15 | 1.25378734158683e-13 | 8.49624376752364e-14 | 2205/5687/2207/7311/5689/5699/5700/6500/7316/5685/5688/6233/7322/5694/5702/5695/5684/5705/4792/7314/2206/5693/5710/9861/5714/5686/5708/7334/5713/3551/5701/5718/5690/4067/115727/10213/7321/5717/5719/5707 | 40 |
| normal | R-HSA-8854050 | FBXL7 down-regulates AURKA during mitotic entry and in early mitosis | 31/1399 | 54/10704 | 1.54972331751356e-14 | 4.41506281308651e-13 | 2.99185106315048e-13 | 5687/7311/5689/5699/5700/6500/7316/5685/9978/5688/6233/5694/5702/5695/5684/5705/7314/5693/5710/9861/5714/5686/5708/5713/5701/5718/5690/10213/5717/5719/5707 | 31 |
| normal | R-HSA-5607764 | CLEC7A (Dectin-1) signaling | 44/1399 | 101/10704 | 3.25876612822839e-14 | 9.036336556364e-13 | 6.12344020406884e-13 | 5687/805/7311/5689/5699/5700/6500/7316/5971/5685/5688/6233/7322/5694/5702/5695/5684/5705/29108/4792/7314/801/5530/5693/5710/9861/5714/808/5686/5708/7334/5713/10768/3551/5701/5718/5690/10213/7321/4791/5717/5719/6850/5707 | 44 |
| normal | R-HSA-180534 | Vpu mediated degradation of CD4 | 30/1399 | 52/10704 | 3.44177120518718e-14 | 9.036336556364e-13 | 6.12344020406884e-13 | 5687/7311/5689/5699/5700/7316/5685/5688/6233/5694/5702/5695/5684/5705/7314/5693/5710/9861/5714/5686/5708/5713/5701/5718/5690/10213/5717/5719/5707 | 30 |
| normal | R-HSA-349425 | Autodegradation of the E3 ubiquitin ligase COP1 | 30/1399 | 52/10704 | 3.44177120518718e-14 | 9.036336556364e-13 | 6.12344020406884e-13 | 5687/7311/5689/5699/5700/7316/5685/5688/6233/5694/5702/5695/5684/5705/7314/5693/5710/9861/5714/5686/5708/5713/472/5701/5718/5690/10213/5717/5719/5707 | 30 |
| normal | R-HSA-75815 | Ubiquitin-dependent degradation of Cyclin D | 30/1399 | 52/10704 | 3.44177120518718e-14 | 9.036336556364e-13 | 6.12344020406884e-13 | 5687/7311/5689/5699/5700/7316/5685/5688/6233/5694/5702/5695/5684/5705/7314/5693/5710/9861/5714/5686/5708/5713/5701/5718/5690/1019/10213/5717/5719/5707 | 30 |
| normal | R-HSA-1168372 | Downstream signaling events of B Cell Receptor (BCR) | 39/1399 | 83/10704 | 4.54785838128148e-14 | 1.17107353317998e-12 | 7.93573668960453e-13 | 5687/805/7311/5689/5699/5700/6500/7316/5685/5478/5688/6233/5694/5702/2280/5695/5684/5705/4792/7314/801/5530/5693/5710/9861/5714/808/5686/5708/5713/3551/5701/5718/5690/5579/10213/5717/5719/5707 | 39 |
| normal | R-HSA-351202 | Metabolism of polyamines | 32/1399 | 59/10704 | 5.20501142674293e-14 | 1.29065005563126e-12 | 8.74604259230488e-13 | 5687/5689/5699/5700/4946/5685/5688/6303/5694/6611/5702/5695/5684/5705/4947/5693/5710/9861/5714/5686/5708/5713/6723/5701/5718/5690/4953/10213/262/5717/5719/5707 | 32 |
| normal | R-HSA-5676590 | NIK-->noncanonical NF-kB signaling | 32/1399 | 59/10704 | 5.20501142674293e-14 | 1.29065005563126e-12 | 8.74604259230488e-13 | 5687/7311/5689/5699/5700/6500/7316/5971/5685/5688/6233/5694/5702/5695/5684/5705/7314/5693/5710/9861/5714/5686/5708/5713/5701/5718/5690/10213/4791/5717/5719/5707 | 32 |
| normal | R-HSA-450408 | AUF1 (hnRNP D0) binds and destabilizes mRNA | 31/1399 | 56/10704 | 6.1048623510918e-14 | 1.48625648874762e-12 | 1.0071562385916e-12 | 5687/7311/5689/5699/5700/7316/5685/5688/6233/5694/5702/5695/5684/5705/7314/5693/5710/9861/3184/5714/5686/5708/5713/26986/5701/5718/5690/10213/5717/5719/5707 | 31 |
| normal | R-HSA-211733 | Regulation of activated PAK-2p34 by proteasome mediated degradation | 29/1399 | 50/10704 | 7.63161755181359e-14 | 1.82477426819257e-12 | 1.23655156572619e-12 | 5687/7311/5689/5699/5700/7316/5685/5688/6233/5694/5702/5695/5684/5705/7314/5693/5710/9861/5714/5686/5708/5713/5701/5718/5690/10213/5717/5719/5707 | 29 |
| normal | R-HSA-5607761 | Dectin-1 mediated noncanonical NF-kB signaling | 32/1399 | 60/10704 | 9.77026831437428e-14 | 2.29515601279775e-12 | 1.55530402345164e-12 | 5687/7311/5689/5699/5700/6500/7316/5971/5685/5688/6233/5694/5702/5695/5684/5705/7314/5693/5710/9861/5714/5686/5708/5713/5701/5718/5690/10213/4791/5717/5719/5707 | 32 |
| normal | R-HSA-69541 | Stabilization of p53 | 31/1399 | 57/10704 | 1.17245424010515e-13 | 2.70675211638067e-12 | 1.83422060793219e-12 | 5687/7311/5689/5699/5700/7316/5685/5688/6233/51230/5694/5702/5695/5684/5705/7314/5693/5710/9861/5714/5686/5708/5713/472/5701/5718/5690/10213/5717/5719/5707 | 31 |
| normal | R-HSA-983705 | Signaling by the B Cell Receptor (BCR) | 46/1399 | 112/10704 | 1.20112716434778e-13 | 2.72594792044351e-12 | 1.84722857389435e-12 | 5687/805/7311/5689/5699/5700/6500/7316/5685/5478/5688/6233/5694/5702/2280/5695/5684/5705/4792/7314/801/5530/5693/5710/9861/5714/808/5686/5708/5713/10768/3551/30011/5701/5718/5690/5579/4067/10213/5777/80228/5717/5719/2534/6850/5707 | 46 |
| normal | R-HSA-180585 | Vif-mediated degradation of APOBEC3G | 30/1399 | 54/10704 | 1.36963171825631e-13 | 3.056561451242e-12 | 2.07126761602972e-12 | 5687/7311/5689/5699/5700/7316/5685/9978/5688/6233/5694/5702/5695/5684/5705/7314/5693/5710/9861/5714/5686/5708/5713/5701/5718/5690/10213/5717/5719/5707 | 30 |
| normal | R-HSA-174084 | Autodegradation of Cdh1 by Cdh1:APC/C | 33/1399 | 64/10704 | 1.43879141557766e-13 | 3.15826509091555e-12 | 2.14018671307669e-12 | 5687/7311/5689/5699/5700/119504/7316/5685/5688/246184/6233/5694/5702/5695/5684/5705/7314/5693/5710/9861/5714/5686/5708/5713/5701/51529/5718/5690/10213/7321/5717/5719/5707 | 33 |
| normal | R-HSA-162906 | HIV Infection | 73/1399 | 234/10704 | 2.29706389548884e-13 | 4.96091702590251e-12 | 3.36174716113986e-12 | 5687/7311/5689/51652/5699/5700/6500/4869/902/293/7316/5685/5879/9978/5478/6879/5902/5688/2547/5901/6233/8815/5694/51160/128866/1173/5702/5440/5441/7251/27243/5695/5684/5705/3105/7314/2958/5693/7520/5710/9861/1175/8905/375/5714/25978/5686/5708/1025/5713/8907/9397/5439/292/5701/5151 | 73 |
| normal | R-HSA-174113 | SCF-beta-TrCP mediated degradation of Emi1 | 30/1399 | 55/10704 | 2.63985749472421e-13 | 5.61074473878685e-12 | 3.80210051871724e-12 | 5687/7311/5689/5699/5700/6500/7316/5685/5688/6233/5694/5702/5695/5684/5705/7314/5693/5710/9861/5714/5686/5708/5713/5701/5718/5690/10213/5717/5719/5707 | 30 |
| normal | R-HSA-69601 | Ubiquitin Mediated Degradation of Phosphorylated Cdc25A | 29/1399 | 52/10704 | 3.07049686509723e-13 | 6.22938682176544e-12 | 4.22132104898535e-12 | 5687/7311/5689/5699/5700/7316/5685/5688/6233/5694/5702/5695/5684/5705/7314/5693/5710/9861/5714/5686/5708/5713/5701/5718/5690/10213/5717/5719/5707 | 29 |
| normal | R-HSA-69610 | p53-Independent DNA Damage Response | 29/1399 | 52/10704 | 3.07049686509723e-13 | 6.22938682176544e-12 | 4.22132104898535e-12 | 5687/7311/5689/5699/5700/7316/5685/5688/6233/5694/5702/5695/5684/5705/7314/5693/5710/9861/5714/5686/5708/5713/5701/5718/5690/10213/5717/5719/5707 | 29 |
| normal | R-HSA-69613 | p53-Independent G1/S DNA damage checkpoint | 29/1399 | 52/10704 | 3.07049686509723e-13 | 6.22938682176544e-12 | 4.22132104898535e-12 | 5687/7311/5689/5699/5700/7316/5685/5688/6233/5694/5702/5695/5684/5705/7314/5693/5710/9861/5714/5686/5708/5713/5701/5718/5690/10213/5717/5719/5707 | 29 |
| normal | R-HSA-5362768 | Hh mutants that don't undergo autocatalytic processing are degraded by ERAD | 30/1399 | 56/10704 | 4.98143971735694e-13 | 9.9554444500611e-12 | 6.74627028493587e-12 | 5687/7311/5689/5699/5700/7316/5685/5688/6233/5694/5702/5695/5684/5705/7314/5693/5710/9861/5714/5686/5708/5713/51009/5701/5718/5690/10213/5717/5719/5707 | 30 |
| normal | R-HSA-8852276 | The role of GTSE1 in G2/M progression after G2 checkpoint | 36/1399 | 77/10704 | 5.23129392699727e-13 | 1.0301033188602e-11 | 6.98045722147314e-12 | 5687/7846/84617/7311/5689/5699/5700/10376/7316/5685/5688/6233/5694/5702/5695/5684/5705/7314/22919/5693/5710/9861/5714/5686/5708/3326/10383/5713/5701/5718/5690/3320/10213/5717/5719/5707 | 36 |
| normal | R-HSA-169911 | Regulation of Apoptosis | 29/1399 | 53/10704 | 5.94110526495237e-13 | 1.15291883330018e-11 | 7.81271203415553e-12 | 5687/7311/5689/5699/5700/7316/5685/5688/6233/5694/5702/5695/5684/5705/7314/5693/5710/9861/5714/5686/5708/5713/5701/5718/5690/10213/5717/5719/5707 | 29 |
| normal | R-HSA-187577 | SCF(Skp2)-mediated degradation of p27/p21 | 31/1399 | 60/10704 | 7.38233356866604e-13 | 1.35410200663614e-11 | 9.17602240258129e-12 | 5687/7311/5689/5699/5700/6500/7316/5685/5688/6233/5694/5702/5695/5684/5705/7314/5693/5710/9861/5714/5686/5708/5713/5701/5718/5690/1019/10213/5717/5719/5707 | 31 |
| normal | R-HSA-5610780 | Degradation of GLI1 by the proteasome | 31/1399 | 60/10704 | 7.38233356866604e-13 | 1.35410200663614e-11 | 9.17602240258129e-12 | 5687/7311/5689/5699/5700/6500/7316/5685/9978/5688/6233/5694/5702/5695/5684/5705/7314/5693/5710/9861/5714/5686/5708/5713/5701/5718/5690/10213/5717/5719/5707 | 31 |
| normal | R-HSA-5610783 | Degradation of GLI2 by the proteasome | 31/1399 | 60/10704 | 7.38233356866604e-13 | 1.35410200663614e-11 | 9.17602240258129e-12 | 5687/7311/5689/5699/5700/6500/7316/5685/9978/5688/6233/5694/5702/5695/5684/5705/7314/5693/5710/9861/5714/5686/5708/5713/5701/5718/5690/10213/5717/5719/5707 | 31 |
| normal | R-HSA-5610785 | GLI3 is processed to GLI3R by the proteasome | 31/1399 | 60/10704 | 7.38233356866604e-13 | 1.35410200663614e-11 | 9.17602240258129e-12 | 5687/7311/5689/5699/5700/6500/7316/5685/9978/5688/6233/5694/5702/5695/5684/5705/7314/5693/5710/9861/5714/5686/5708/5713/5701/5718/5690/10213/5717/5719/5707 | 31 |
| normal | R-HSA-1169091 | Activation of NF-kappaB in B cells | 33/1399 | 67/10704 | 7.74152710796857e-13 | 1.40079794561756e-11 | 9.49245571418052e-12 | 5687/7311/5689/5699/5700/6500/7316/5685/5688/6233/5694/5702/5695/5684/5705/4792/7314/5693/5710/9861/5714/5686/5708/5713/3551/5701/5718/5690/5579/10213/5717/5719/5707 | 33 |
| normal | R-HSA-4641258 | Degradation of DVL | 30/1399 | 57/10704 | 9.21364580319221e-13 | 1.64494289739658e-11 | 1.11468949927743e-11 | 5687/7311/5689/5699/5700/7316/5685/9978/5688/6233/5694/5702/5695/5684/5705/7314/5693/5710/9861/5714/5686/5708/5713/5701/5718/5690/10213/5717/5719/5707 | 30 |
| normal | R-HSA-5678895 | Defective CFTR causes cystic fibrosis | 31/1399 | 61/10704 | 1.31504795793467e-12 | 2.29110984709435e-11 | 1.55256215415693e-11 | 5687/7311/5689/5699/5700/7316/5685/5688/6233/5694/5702/6048/5695/5684/5705/7314/5693/5710/9861/5714/5686/5708/5713/51009/5701/5718/5690/10213/5717/5719/5707 | 31 |
| normal | R-HSA-174154 | APC/C:Cdc20 mediated degradation of Securin | 33/1399 | 68/10704 | 1.31751649160765e-12 | 2.29110984709435e-11 | 1.55256215415693e-11 | 5687/7311/5689/5699/5700/119504/7316/5685/5688/246184/6233/5694/5702/5695/5684/5705/7314/5693/5710/9861/5714/5686/5708/5713/5701/51529/5718/5690/10213/7321/5717/5719/5707 | 33 |
| normal | R-HSA-350562 | Regulation of ornithine decarboxylase (ODC) | 28/1399 | 51/10704 | 1.33642927497148e-12 | 2.29420358870104e-11 | 1.55465861676844e-11 | 5687/5689/5699/5700/4946/5685/5688/5694/5702/5695/5684/5705/4947/5693/5710/9861/5714/5686/5708/5713/5701/5718/5690/4953/10213/5717/5719/5707 | 28 |
| normal | R-HSA-4641257 | Degradation of AXIN | 29/1399 | 55/10704 | 2.08409590194072e-12 | 3.5324106489856e-11 | 2.39372507324837e-11 | 5687/7311/5689/5699/5700/7316/5685/5688/6233/5694/5702/5695/5684/5705/7314/5693/5710/9861/5714/5686/5708/5713/5701/5718/5690/10213/5717/5719/5707 | 29 |
| normal | R-HSA-450531 | Regulation of mRNA stability by proteins that bind AU-rich elements | 38/1399 | 88/10704 | 2.57922725839383e-12 | 4.31698162373667e-11 | 2.9253867062309e-11 | 5687/7311/5689/5699/5700/7316/5685/5688/7538/6233/5694/11340/5702/5695/5684/5705/7314/51010/5693/5710/9861/3184/207/5714/5686/5708/5713/26986/5701/5718/5690/54512/9261/10213/5717/1432/5719/5707 | 38 |
| normal | R-HSA-4086400 | PCP/CE pathway | 39/1399 | 92/10704 | 2.70141881532108e-12 | 4.46567875767274e-11 | 3.02615077167871e-11 | 5687/5216/7311/5689/5699/5700/7316/5685/5879/5688/6233/5694/1173/5702/5695/5684/5705/7314/5693/1212/5710/9861/1175/5714/5686/5708/5713/409/5701/1213/5718/5690/5579/5880/10213/387/5717/5719/5707 | 39 |
| normal | R-HSA-5387390 | Hh mutants abrogate ligand secretion | 30/1399 | 59/10704 | 2.98060538526994e-12 | 4.86711050106884e-11 | 3.29817951489433e-11 | 5687/7311/5689/5699/5700/7316/5685/5688/6233/5694/5702/5695/5684/5705/7314/5693/5710/9861/5714/5686/5708/5713/51009/5701/5718/5690/10213/5717/5719/5707 | 30 |
| normal | R-HSA-195253 | Degradation of beta-catenin by the destruction complex | 36/1399 | 83/10704 | 8.30179129860179e-12 | 1.33928898178648e-10 | 9.0756424849648e-11 | 5687/7311/5689/5699/5700/6500/7316/5685/9978/5688/6233/5694/5702/5695/5684/5705/7314/5693/5710/9861/5714/5686/5708/5713/5701/5527/5525/5718/5690/10213/5515/5717/5719/1487/3065/5707 | 36 |
| normal | R-HSA-5621481 | C-type lectin receptors (CLRs) | 50/1399 | 142/10704 | 1.06826574159525e-11 | 1.70286646190005e-10 | 1.15394118954274e-10 | 5687/2207/805/7311/5689/5699/5700/6500/7316/5971/5685/5688/6233/7322/5694/5702/5695/5684/5705/29108/4792/7314/801/5530/5693/5710/9861/5714/808/5686/5708/7334/5713/10768/3551/5701/5718/5690/4067/10213/7321/5894/5058/4791/5717/5719/2033/2534/6850/5707 | 50 |
| normal | R-HSA-8948751 | Regulation of PTEN stability and activity | 32/1399 | 69/10704 | 1.35232944280646e-11 | 2.13031661637394e-10 | 1.44360121324974e-10 | 5687/7311/5689/5699/5700/10273/7316/5685/5688/6233/5694/5702/5695/5684/5705/7314/5693/5710/9861/207/5714/5686/5708/5713/23608/5701/5718/5690/10213/5717/5719/5707 | 32 |
| normal | R-HSA-174184 | Cdc20:Phospho-APC/C mediated degradation of Cyclin A | 33/1399 | 73/10704 | 1.55137423731585e-11 | 2.36055693609765e-10 | 1.5996227183807e-10 | 5687/7311/5689/5699/5700/119504/7316/5685/5688/246184/6233/5694/5702/5695/5684/5705/7314/5693/5710/9861/5714/5686/5708/5713/5701/51529/5718/5690/10213/7321/5717/5719/5707 | 33 |
| normal | R-HSA-69017 | CDK-mediated phosphorylation and removal of Cdc6 | 33/1399 | 73/10704 | 1.55137423731585e-11 | 2.36055693609765e-10 | 1.5996227183807e-10 | 5687/7311/5689/5699/5700/119504/7316/5685/5688/246184/6233/5694/5702/5695/5684/5705/7314/5693/5710/9861/5714/5686/5708/5713/5701/51529/5718/5690/10213/7321/5717/5719/5707 | 33 |
| normal | R-HSA-8939902 | Regulation of RUNX2 expression and activity | 33/1399 | 73/10704 | 1.55137423731585e-11 | 2.36055693609765e-10 | 1.5996227183807e-10 | 5687/7311/5689/5699/5700/6500/10273/7316/5685/9978/5688/6233/5694/5702/5695/5684/5705/7314/5693/5710/9861/5714/5686/5708/5713/5701/5718/5690/10213/5717/5719/2908/5707 | 33 |
| normal | R-HSA-69563 | p53-Dependent G1 DNA Damage Response | 31/1399 | 66/10704 | 1.86861224734614e-11 | 2.78007977688498e-10 | 1.88391082714898e-10 | 5687/7311/5689/5699/5700/7316/5685/5688/6233/51230/5694/5702/5695/5684/5705/7314/5693/5710/9861/5714/5686/5708/5713/472/5701/5718/5690/10213/5717/5719/5707 | 31 |
| normal | R-HSA-69580 | p53-Dependent G1/S DNA damage checkpoint | 31/1399 | 66/10704 | 1.86861224734614e-11 | 2.78007977688498e-10 | 1.88391082714898e-10 | 5687/7311/5689/5699/5700/7316/5685/5688/6233/51230/5694/5702/5695/5684/5705/7314/5693/5710/9861/5714/5686/5708/5713/472/5701/5718/5690/10213/5717/5719/5707 | 31 |
| normal | R-HSA-72165 | mRNA Splicing - Minor Pathway | 27/1399 | 52/10704 | 1.94350232147995e-11 | 2.85972484446335e-10 | 1.93788201401471e-10 | 4904/6633/6637/6635/6628/51639/5440/5441/4809/6431/9410/6632/23020/6432/83443/6636/6634/24148/10081/5439/10262/6427/85437/10992/10907/2962/5436 | 27 |
| normal | R-HSA-5687128 | MAPK6/MAPK4 signaling | 37/1399 | 89/10704 | 1.97895948022572e-11 | 2.88024646089373e-10 | 1.95178841184733e-10 | 5687/7311/5689/5699/5700/7316/5685/5879/998/192669/5688/6233/5694/5702/5695/5684/5705/7314/5693/5710/9861/3337/5714/5686/5708/5713/896/5701/5718/5690/23112/10213/5058/5717/5719/3725/5707 | 37 |
| normal | R-HSA-68827 | CDT1 association with the CDC6:ORC:origin complex | 29/1399 | 59/10704 | 2.04039997227543e-11 | 2.93773716438365e-10 | 1.9907467754402e-10 | 5687/7311/5689/5699/5700/7316/5685/5688/6233/5694/5702/5695/5684/5705/7314/5693/5710/9861/5714/5686/5708/5713/5701/5718/5690/10213/5717/5719/5707 | 29 |
| normal | R-HSA-174178 | APC/C:Cdh1 mediated degradation of Cdc20 and other APC/C:Cdh1 targeted proteins in late mitosis/early G1 | 33/1399 | 74/10704 | 2.45290863863311e-11 | 3.45731017592604e-10 | 2.34283351412935e-10 | 5687/7311/5689/5699/5700/119504/7316/5685/5688/246184/6233/5694/5702/5695/5684/5705/7314/5693/5710/9861/5714/5686/5708/5713/5701/51529/5718/5690/10213/7321/5717/5719/5707 | 33 |
| normal | R-HSA-179419 | APC:Cdc20 mediated degradation of cell cycle proteins prior to satisfation of the cell cycle checkpoint | 33/1399 | 74/10704 | 2.45290863863311e-11 | 3.45731017592604e-10 | 2.34283351412935e-10 | 5687/7311/5689/5699/5700/119504/7316/5685/5688/246184/6233/5694/5702/5695/5684/5705/7314/5693/5710/9861/5714/5686/5708/5713/5701/51529/5718/5690/10213/7321/5717/5719/5707 | 33 |
| normal | R-HSA-9013694 | Signaling by NOTCH4 | 35/1399 | 82/10704 | 2.82200289420238e-11 | 3.9361061201427e-10 | 2.6672878232483e-10 | 5687/7311/5689/5699/5700/6500/7316/5685/9978/5688/6233/5694/5702/5695/5684/5705/7314/22938/5693/5710/9861/207/5714/5686/5708/5713/5701/5718/5690/51107/10213/5717/5719/2033/5707 | 35 |
| normal | R-HSA-5389840 | Mitochondrial translation elongation | 36/1399 | 87/10704 | 4.3523471633251e-11 | 6.00803386772403e-10 | 4.07132203449402e-10 | 116540/6150/90480/55052/9553/116541/28957/64983/54998/64928/57129/122704/92259/54534/51258/29074/28998/29093/7284/65005/78988/10240/64975/128308/64969/64976/11222/29088/51081/79590/84545/219927/10884/64968/54948/51642 | 36 |
| normal | R-HSA-69615 | G1/S DNA Damage Checkpoints | 31/1399 | 68/10704 | 4.9062773122291e-11 | 6.70357685823955e-10 | 4.54265418167721e-10 | 5687/7311/5689/5699/5700/7316/5685/5688/6233/51230/5694/5702/5695/5684/5705/7314/5693/5710/9861/5714/5686/5708/5713/472/5701/5718/5690/10213/5717/5719/5707 | 31 |
| normal | R-HSA-176409 | APC/C:Cdc20 mediated degradation of mitotic proteins | 33/1399 | 76/10704 | 5.94138406843044e-11 | 8.03587198750339e-10 | 5.44547907175655e-10 | 5687/7311/5689/5699/5700/119504/7316/5685/5688/246184/6233/5694/5702/5695/5684/5705/7314/5693/5710/9861/5714/5686/5708/5713/5701/51529/5718/5690/10213/7321/5717/5719/5707 | 33 |
| normal | R-HSA-5358346 | Hedgehog ligand biogenesis | 30/1399 | 65/10704 | 6.86302509750209e-11 | 9.18959060555529e-10 | 6.22729224636505e-10 | 5687/7311/5689/5699/5700/7316/5685/5688/6233/5694/5702/5695/5684/5705/7314/5693/5710/9861/5714/5686/5708/5713/51009/5701/5718/5690/10213/5717/5719/5707 | 30 |
| normal | R-HSA-6807070 | PTEN Regulation | 48/1399 | 140/10704 | 8.11674187960564e-11 | 1.07607102740514e-09 | 7.29195570632628e-10 | 5687/389541/7311/5689/5699/5700/10273/7316/5685/10542/192669/5688/6233/5694/5702/5695/23466/6009/5684/5705/5931/55004/7314/10014/5693/5710/9861/28956/207/5714/5686/5708/84232/5713/23512/23608/10670/5701/5718/5690/5594/23112/10213/5717/5719/3725/3065/5707 | 48 |
| normal | R-HSA-8950505 | Gene and protein expression by JAK-STAT signaling after Interleukin-12 stimulation | 22/1399 | 38/10704 | 8.49964385837453e-11 | 1.11578658101603e-09 | 7.5610866934147e-10 | 4282/1072/9987/998/5478/9446/6175/6647/5908/3181/27250/3185/3313/302/375/6888/6950/6627/9049/829/3936/1265 | 22 |
| normal | R-HSA-176814 | Activation of APC/C and APC/C:Cdc20 mediated degradation of mitotic proteins | 33/1399 | 77/10704 | 9.10873530488754e-11 | 1.18413558963538e-09 | 8.02425123435162e-10 | 5687/7311/5689/5699/5700/119504/7316/5685/5688/246184/6233/5694/5702/5695/5684/5705/7314/5693/5710/9861/5714/5686/5708/5713/5701/51529/5718/5690/10213/7321/5717/5719/5707 | 33 |
| normal | R-HSA-176408 | Regulation of APC/C activators between G1/S and early anaphase | 34/1399 | 81/10704 | 9.43774774249596e-11 | 1.21511002184635e-09 | 8.23414833404e-10 | 5687/7311/5689/5699/5700/6500/119504/7316/5685/5688/246184/6233/5694/5702/5695/5684/5705/7314/5693/5710/9861/5714/5686/5708/5713/5701/51529/5718/5690/10213/7321/5717/5719/5707 | 34 |
| normal | R-HSA-9020702 | Interleukin-1 signaling | 39/1399 | 103/10704 | 1.6849568251013e-10 | 2.1487211322006e-09 | 1.45607296565145e-09 | 5687/3146/7311/5689/5699/5700/6500/7316/5685/9978/5688/6233/5694/5702/5695/5684/5705/4792/7314/5693/5710/9861/5714/5686/5708/7334/5713/3551/5701/4615/5718/5690/10213/8878/4791/5717/5719/5604/5707 | 39 |
| normal | R-HSA-1236978 | Cross-presentation of soluble exogenous antigens (endosomes) | 25/1399 | 49/10704 | 1.78582489077214e-10 | 2.25586747994708e-09 | 1.52868029378906e-09 | 5687/5689/5699/5700/5685/5688/5694/5702/5695/5684/5705/5693/5710/9861/5714/5686/5708/5713/5701/5718/5690/10213/5717/5719/5707 | 25 |
| normal | R-HSA-5357801 | Programmed Cell Death | 58/1399 | 190/10704 | 1.80733624907247e-10 | 2.26170396028789e-09 | 1.53263536320755e-09 | 5687/4000/3146/7311/5689/5699/5700/54205/10273/140735/7316/5685/5688/708/6233/7533/5694/8655/5702/6386/5695/10134/5684/7431/5705/11140/7314/2934/5693/5710/9861/6709/207/5714/3148/10971/5686/5708/5713/8743/581/5701/5718/330/5690/5594/6304/118/28988/3320/10213/7532/3837/329/5717/571 | 58 |
| normal | R-HSA-68949 | Orc1 removal from chromatin | 31/1399 | 71/10704 | 1.90908958114508e-10 | 2.36691754551228e-09 | 1.60393296193671e-09 | 5687/7311/5689/5699/5700/6500/7316/5685/9978/5688/6233/5694/5702/5695/5684/5705/7314/5693/5710/9861/5714/5686/5708/5713/5701/5718/5690/10213/5717/5719/5707 | 31 |
| normal | R-HSA-5368286 | Mitochondrial translation initiation | 35/1399 | 87/10704 | 2.06921697468099e-10 | 2.51880139008894e-09 | 1.70685649011963e-09 | 116540/6150/90480/55052/9553/116541/28957/64983/54998/64928/57129/122704/92259/54534/51258/29074/28998/29093/65005/78988/10240/64975/128308/64969/64976/11222/29088/51081/79590/84545/219927/10884/64968/54948/51642 | 35 |
| normal | R-HSA-5419276 | Mitochondrial translation termination | 35/1399 | 87/10704 | 2.06921697468099e-10 | 2.51880139008894e-09 | 1.70685649011963e-09 | 116540/6150/90480/55052/9553/116541/28957/64983/54998/64928/57129/122704/92259/54534/51258/29074/28998/29093/65005/78988/10240/64975/128308/64969/64976/11222/29088/51081/79590/84545/219927/10884/64968/54948/51642 | 35 |
| normal | R-HSA-4608870 | Asymmetric localization of PCP proteins | 29/1399 | 64/10704 | 2.47638308177071e-10 | 2.98727652837026e-09 | 2.02431694308805e-09 | 5687/7311/5689/5699/5700/7316/5685/5688/6233/5694/5702/5695/5684/5705/7314/5693/5710/9861/5714/5686/5708/5713/5701/5718/5690/10213/5717/5719/5707 | 29 |
| normal | R-HSA-9609507 | Protein localization | 52/1399 | 163/10704 | 2.72615812587525e-10 | 3.25048025936387e-09 | 2.20267598252557e-09 | 100188893/7311/51142/401505/293/7316/6844/131118/3421/7323/54543/10952/10063/9218/6233/7322/51024/6342/819/23480/10245/3313/1431/7314/56993/373156/1537/9804/1528/10455/83752/11001/54469/9409/4580/5621/847/6811/9512/79145/27230/7416/90639/118424/5828/10440/2010/7321/26515/10651/3 | 52 |

| | | | | | | | | | |
|---|---|---|---|---|---|---|---|---|---|
| normal | R-HSA-5658442 | Regulation of RAS by GAPs | 30/1399 | 68/10704 | 2.74312374389931e-10 | 3.25048025936387e-09 | 2.20267598252557e-09 | 5687/7311/5689/5699/5700/7316/5685/9978/5688/6233/5694/5702/5695/5684/5705/7314/5693/5710/9861/5714/5686/5708/5713/5701/5718/5690/10213/5717/5719/5707 | 30 |
| normal | R-HSA-5368287 | Mitochondrial translation | 36/1399 | 93/10704 | 4.12945262373648e-10 | 4.85029566945899e-09 | 3.28678500615037e-09 | 116540/6150/90480/55052/9553/116541/28957/6183/64949/54498/64928/57129/122704/92259/54534/51258/29074/28998/29093/7284/65005/78988/10240/64975/128308/64969/64976/11222/29088/51081/79590/84545/219927/10884/64968/64948/51642 | 36 |
| normal | R-HSA-5632684 | Hedgehog 'on' state | 34/1399 | 85/10704 | 4.50366563741331e-10 | 5.24383329434471e-09 | 3.55346432901627e-09 | 5687/7311/5689/5699/5700/7316/5685/9978/5688/6233/5694/5702/5695/5684/5705/7314/5693/5710/9861/5714/5686/5708/5713/408/409/156/5701/5718/5690/79577/10213/5717/5719/5707 | 34 |
| normal | R-HSA-68882 | Mitotic Anaphase | 66/1399 | 236/10704 | 6.54296468142798e-10 | 7.55261181761594e-09 | 5.11799959654495e-09 | 5687/4000/7846/84617/7311/5689/51652/5699/5700/119504/10376/140735/7316/5685/5688/246184/7341/11243/5901/10726/6233/6232/8815/5694/128866/8655/5702/27243/5695/5684/5705/9126/7314/1781/22919/5693/5710/9861/1060/5714/25978/5686/5708/10383/5713/7329/5701/51529/5527/5525/5718/5690/5 | 66 |
| normal | R-HSA-2555396 | Mitotic Metaphase and Anaphase | 66/1399 | 237/10704 | 7.94059045896758e-10 | 9.08756463637401e-09 | 6.15815472391368e-09 | 5687/4000/7846/84617/7311/5689/51652/5699/5700/119504/10376/140735/7316/5685/5688/246184/7341/11243/5901/10726/6233/6232/8815/5694/128866/8655/5702/27243/5695/5684/5705/9126/7314/1781/22919/5693/5710/9861/1060/5714/25978/5686/5708/10383/5713/7329/5701/51529/5527/5525/5718/5690/5 | 66 |
| normal | R-HSA-449147 | Signaling by Interleukins | 107/1399 | 461/10704 | 8.01183669977589e-10 | 9.09139774661010e-09 | 6.16075221695524e-09 | 4282/10538/5687/9021/2353/301/3146/7311/5689/5699/5700/6500/1072/9987/8651/7316/6844/5685/998/9978/5478/9446/6175/6813/5292/6647/5778/5688/5908/3181/6233/3091/27250/5694/4170/3976/5702/7076/3185/5695/5684/7431/7422/5705/3313/3568/4792/6347/7314/3726/8660/1511/3687/5290/302/5693/5 | 107 |
| normal | R-HSA-69052 | Switching of origins to a post-replicative state | 35/1399 | 91/10704 | 8.86695331707951e-10 | 9.97718528703317e-09 | 6.76100288308053e-09 | 5687/7311/5689/5699/5700/119504/7316/5685/9978/5688/246184/6233/5694/5702/5695/5684/5705/7314/5693/5710/9861/5714/5686/5708/5713/5701/5718/5690/10213/7321/5717/5719/5707 | 35 |
| normal | R-HSA-69202 | Cyclin E associated events during G1/S transition | 33/1399 | 83/10704 | 9.78321628388481e-10 | 1.09164383676681e-08 | 7.39748459360413e-09 | 5687/7311/5689/5699/5700/6500/902/7316/5685/5688/6233/5694/5702/5695/5684/5705/7314/5693/5710/9861/207/5714/5686/5708/5713/5701/5718/5690/1019/10213/5717/5719/5707 | 33 |
| normal | R-HSA-174143 | APC/C-mediated degradation of cell cycle proteins | 34/1399 | 88/10704 | 1.33971599266514e-09 | 1.47039320834313e-08 | 9.96406545019283e-09 | 5687/7311/5689/5699/5700/6500/119504/7316/5685/5688/246184/6233/5694/5702/5695/5684/5705/7314/5693/5710/9861/5714/5686/5708/5713/5701/51529/5718/5690/10213/7321/5717/5719/5707 | 34 |
| normal | R-HSA-453276 | Regulation of mitotic cell cycle | 34/1399 | 88/10704 | 1.33971599266514e-09 | 1.47039320834313e-08 | 9.96406545019283e-09 | 5687/7311/5689/5699/5700/6500/119504/7316/5685/5688/246184/6233/5694/5702/5695/5684/5705/7314/5693/5710/9861/5714/5686/5708/5713/5701/51529/5718/5690/10213/7321/5717/5719/5707 | 34 |
| normal | R-HSA-3858494 | Beta-catenin independent WNT signaling | 47/1399 | 146/10704 | 1.40964750389053e-09 | 1.53456748594261e-08 | 1.03989400800482e-08 | 5687/805/5216/7311/5689/5699/5700/7316/5685/5879/192669/5688/6233/5694/1173/5702/5695/5684/5705/7314/801/5530/5693/1212/5710/9861/1175/5714/808/5686/5708/5713/409/5701/1213/5718/5690/2783/2782/23112/5579/5880/10213/387/5717/5719/5707 | 47 |
| normal | R-HSA-68867 | Assembly of the pre-replicative complex | 29/1399 | 68/10704 | 1.44634384825139e-09 | 1.56181807484565e-08 | 1.05836026926375e-08 | 5687/7311/5689/5699/5700/7316/5685/5688/6233/5694/5702/5695/5684/5705/7314/5693/5710/9861/5714/5686/5708/5713/5701/5718/5690/10213/5717/5719/5707 | 29 |
| normal | R-HSA-69656 | Cyclin A:Cdk2-associated events at S phase entry | 33/1399 | 85/10704 | 2.02188537992733e-09 | 2.16584361897815e-08 | 1.46767595578725e-08 | 5687/7311/5689/5699/5700/6500/902/7316/5685/5688/6233/5694/5702/5695/5684/5705/7314/5693/5710/9861/207/5714/5686/5708/5713/5701/5718/5690/1019/10213/5717/5719/5707 | 33 |
| normal | R-HSA-202403 | TCR signaling | 41/1399 | 120/10704 | 2.18274478723718e-09 | 2.30603018640797e-08 | 1.56267286717001e-08 | 5687/7311/5689/5699/5700/7316/5685/4738/11047/5688/6233/7322/5788/5694/3937/5702/5695/5684/5705/4792/7314/7408/5290/5693/5710/9861/5714/5686/5708/7334/5713/3551/5701/27040/5718/5690/10213/7321/5058/5717/5719/5707 | 41 |
| normal | R-HSA-5619084 | ABC transporter disorders | 31/1399 | 77/10704 | 2.18719816037194e-09 | 2.30603018640797e-08 | 1.56267286717001e-08 | 5687/7311/5689/5699/5700/7316/5685/5688/6233/5694/5702/6048/5695/5684/5705/7314/5693/5710/9861/5714/5686/5708/5713/51009/5701/5718/5690/10213/5717/5719/5707 | 31 |
| normal | R-HSA-202424 | Downstream TCR signaling | 36/1399 | 98/10704 | 2.23536775817462e-09 | 2.33840424077798e-08 | 1.58461102594286e-08 | 5687/7311/5689/5699/5700/6500/7316/5685/5688/6233/7322/5694/5702/5695/5684/5705/4792/7314/5290/5693/5710/9861/5714/5686/5708/7334/5713/3551/5701/5718/5690/10213/7321/5717/5719/5707 | 36 |
| normal | R-HSA-109581 | Apoptosis | 53/1399 | 178/10704 | 2.95871433600828e-09 | 3.07109969092365e-08 | 2.08111934527878e-08 | 5687/4000/3146/7311/5689/5699/5700/54205/140735/7316/5685/5688/708/6233/7533/5694/8655/5702/5695/10134/5684/7431/5705/7314/2934/5693/5710/9861/6709/207/5714/3148/10971/5686/5708/5713/8743/581/5701/5718/5690/5594/6304/118/28988/10213/7532/3837/329/5717/5719/7159/5707 | 53 |
| normal | R-HSA-5668541 | TNFR2 non-canonical NF-kB pathway | 36/1399 | 102/10704 | 7.76259522743852e-09 | 7.93443893858029e-08 | 5.37674333953556e-08 | 5687/7311/5689/5699/5700/6500/7316/5971/5685/5688/6233/5694/3604/7293/5702/5695/5684/5705/7314/5693/5710/9861/5714/5686/5708/5713/5701/5718/330/5690/10213/329/4791/5717/5719/5707 | 36 |
| normal | R-HSA-5689603 | UCH proteinases | 36/1399 | 102/10704 | 7.76259522743852e-09 | 7.93443893858029e-08 | 5.37674333953556e-08 | 5687/7311/5689/5699/5700/7316/5685/5688/11047/5688/6233/5694/5702/5695/5684/5705/7314/5693/5710/9861/5714/60/5686/5708/125476/7314/7048/7040/5701/5718/5690/10213/5717/5719/5707 | 36 |
| normal | R-HSA-5696398 | Nucleotide Excision Repair | 38/1399 | 111/10704 | 7.91406534892798e-09 | 8.02797992592013e-08 | 5.44013104527585e-08 | 6613/7311/902/7316/9978/3150/7341/6233/56655/5440/5441/7314/6118/51138/5886/5887/10980/60/125476/7334/1069/142/8533/7336/5439/7329/2067/5425/91603/7507/6917/10450/10987/10920/9318/5436/2033/86 | 38 |
| normal | R-HSA-8951664 | Neddylation | 63/1399 | 234/10704 | 8.22826526377249e-09 | 8.28394525427922e-08 | 5.61358500781313e-08 | 5687/9021/170622/7311/5689/5699/5700/6500/7316/5685/9978/4738/7323/5688/6233/3091/7322/5694/5702/5695/5684/5705/5931/7314/51138/8883/5693/79176/5710/9861/5714/10980/26224/9616/5686/5708/5713/5929/8533/5701/8835/2034/54951/9306/5718/5690/25793/80204/84259/23369/10213/7321/79016/1 | 63 |
| normal | R-HSA-2173795 | Downregulation of SMAD2/3:SMAD4 transcriptional activity | 15/1399 | 23/10704 | 9.06308418529001e-09 | 9.05632068962935e-08 | 6.13698237841319e-08 | 7311/7316/7323/6233/7050/7314/51592/9611/142/6497/9612/6498/7321/5494/3065 | 15 |
| normal | R-HSA-446652 | Interleukin-1 family signaling | 44/1399 | 140/10704 | 1.07499497942995e-08 | 1.06623576107904e-07 | 7.22530738611007e-08 | 5687/3146/7311/5689/5699/5700/6500/7316/5685/9978/5778/5688/6233/5694/5702/5695/5684/5705/4792/7314/1511/5693/5710/9861/9173/5714/5686/5708/7334/5713/3551/5701/4615/5718/5690/8809/10213/8878/5777/4791/5717/5719/5604/5707 | 44 |
| normal | R-HSA-76002 | Platelet activation, signaling and aggregation | 68/1399 | 263/10704 | 1.26908552369349e-08 | 1.24948935067467e-07 | 8.46711858687142e-08 | 226/2207/8407/805/5216/308/2769/1072/967/5879/998/5478/6813/5874/6647/928/5908/7078/3674/2771/7114/391/3937/10916/1191/7076/1521/5660/7422/801/14/5290/5552/51368/207/808/5906/7040/6814/408/409/51706/6810/1/10487/27040/2776/5594/55664/2773/2783/2782/5770/5579/6464/4067/5880/334/5777 | 68 |
| normal | R-HSA-5610787 | Hedgehog 'off' state | 38/1399 | 113/10704 | 1.38481196038381e-08 | 1.35346767923644e-07 | 9.17178570765154e-08 | 5687/7846/84617/7311/5689/5699/5700/10376/7316/5685/9978/5688/6233/5694/5702/5695/5684/5705/7314/5693/5573/5710/9861/2778/5714/5686/5708/10383/5713/5701/5718/5690/10213/5717/5719/5707 | 38 |
| normal | R-HSA-9612973 | Autophagy | 46/1399 | 151/10704 | 1.49160675747215e-08 | 1.44729909453276e-07 | 9.80751353883287e-08 | 100188893/11337/7846/389541/84617/7311/51652/10376/401505/11345/140735/7316/123/10542/54543/11315/6233/51160/128866/8655/7251/27243/6009/7431/55004/7314/1781/56993/9804/28956/64422/25978/7334/3326/10383/4792/10672/80/9140/2080/2080/2034/28106/7416/3320/8878/56270/9821 | 46 |
| normal | R-HSA-2467813 | Separation of Sister Chromatids | 54/1399 | 191/10704 | 1.57098874949736e-08 | 1.51334815509134e-07 | 1.02551480656321e-07 | 5687/7846/84617/7311/5689/5699/5700/119504/10376/140735/7316/5685/5688/246184/11243/10726/6233/6232/5694/8655/5702/5695/5684/5705/9126/7314/1781/22919/5693/5710/9861/5714/5686/5708/10383/5713/5701/51529/5527/5525/5718/5690/23063/5903/10213/7321/23122/5515/5717/5719/81565/25 | 54 |
| normal | R-HSA-5628897 | TP53 Regulates Metabolic Genes | 32/1399 | 87/10704 | 1.67234469626401e-08 | 1.59947824878394e-07 | 1.08388054750344e-07 | 4697/7295/389541/1329/1337/1327/54205/1340/1345/1349/9167/5052/10542/192669/1351/1350/25824/7533/6009/7001/55004/28956/207/10971/54541/84987/10670/90639/23112/9377/7532/1153 | 32 |
| normal | R-HSA-5663202 | Diseases of signal transduction by growth factor receptors and second messengers | 90/1399 | 387/10704 | 1.69802775942368e-08 | 1.61252423394915e-07 | 1.09272111132752e-07 | 5687/4000/805/7311/5689/5699/5700/6500/5037/7316/5685/5879/9978/5688/5908/3674/6233/5694/391/3164/5702/5440/9208/5441/2280/5695/5684/5705/11221/11140/7314/22938/801/10014/8660/8036/5290/5693/5710/9861/207/26127/5714/60/808/5686/5708/5500/9611/5713/7048/5906/7040/408/409/5439/51009/9 | 90 |
| normal | R-HSA-9020591 | Interleukin-12 signaling | 22/1399 | 47/10704 | 1.74365252143352e-08 | 1.64419065225474e-07 | 1.11417974312612e-07 | 4282/1072/9987/998/5478/9446/6175/6647/5908/3181/27250/3185/3313/302/375/6888/6950/6627/9049/829/3936/1265 | 22 |
| normal | R-HSA-8878166 | Transcriptional regulation by RUNX2 | 39/1399 | 121/10704 | 3.38560934491462e-08 | 3.17061477510966e-07 | 2.14858258165061e-07 | 3958/5687/7311/5689/5699/5700/6500/10273/7316/5685/9978/5688/6233/5694/5702/5695/5684/5705/7314/5693/5710/9861/207/5714/5686/5708/10432/5713/581/5701/5718/5690/5594/1019/10213/5717/5719/2908/5707 | 39 |
| normal | R-HSA-69002 | DNA Replication Pre-Initiation | 31/1399 | 85/10704 | 3.51898210427344e-08 | 3.27216460495822e-07 | 2.21737030254657e-07 | 5687/7311/5689/5699/5700/7316/5685/5688/6233/56655/5694/5702/5695/5684/5705/7314/6118/5693/5710/9861/5714/5686/5708/5713/5701/5718/5690/10213/5717/5719/5707 | 31 |
| normal | R-HSA-382556 | ABC-family proteins mediated transport | 35/1399 | 103/10704 | 3.80696104843518e-08 | 3.51553161645152e-07 | 2.38228705898448e-07 | 5687/7311/5689/5699/5700/7316/5685/5688/6233/5694/5702/6048/5695/5684/5705/1968/7314/8894/5693/5710/9861/1965/5714/5686/5708/5713/51009/5701/5718/5690/10213/5717/5719/10257/5707 | 35 |
| normal | R-HSA-1632852 | Macroautophagy | 42/1399 | 136/10704 | 4.07175274638305e-08 | 3.73429926534719e-07 | 2.53053415096048e-07 | 100188893/11337/7846/389541/84617/7311/51652/10376/401505/11345/140735/7316/123/10542/54543/11315/6233/128866/8655/27243/6009/7431/55004/7314/1781/56993/9804/28956/64422/25978/7334/10383/472/9140/10670/2034/26100/7416/3320/8878/56270/9821 | 42 |
| normal | R-HSA-69275 | G2/M Transition | 54/1399 | 196/10704 | 4.13302135723836e-08 | 3.76470448798786e-07 | 2.55113813815 93e-07 | 5687/7846/84617/7311/5689/5699/5700/6500/10376/902/7316/5685/9978/80097/5688/6233/5694/8655/5702/5695/5684/5705/984/7314/1781/22919/5693/5710/9861/5714/5686/11258/5708/5500/1069/3326/10383/5713/4659/5701/5718/5690/11064/10142/3320/10213/7532/5515/5717/5719/2033/10121/5108/5707 | 54 |
| normal | R-HSA-170834 | Signaling by TGF-beta Receptor Complex | 28/1399 | 73/10704 | 5.57537710039512e-08 | 4.13045617684571e-07 | 2.80510317250406e-07 | 7311/10273/7316/4738/7323/6233/23645/2280/7050/7314/22938/3726/11171/51592/1025/5500/9611/7048/142/7040/6497/5499/9612/6498/7321/387/5494/3065 | 28 |
| normal | R-HSA-453274 | Mitotic G2-G2/M phases | 54/1399 | 198/10704 | 6.00313169029483e-08 | 5.39476062637905e-07 | 3.65573967999586e-07 | 5687/7846/84617/7311/5689/5699/5700/6500/10376/902/7316/5685/9978/80097/5688/6233/5694/8655/5702/5695/5684/5705/984/7314/1781/22919/5693/5710/9861/5714/5686/11258/5708/5500/1069/3326/10383/5713/4659/5701/5718/5690/11064/10142/3320/10213/7532/5515/5717/5719/2033/10121/5108/5707 | 54 |
| normal | R-HSA-8878159 | Transcriptional regulation by RUNX3 | 33/1399 | 96/10704 | 6.7690768555943e-08 | 6.04252927309385e-07 | 4.09469771896301e-07 | 5687/7311/5689/5699/5700/7316/5685/5688/6233/5694/5702/5695/5684/5705/7314/22938/5693/864/5710/9861/5714/5686/5708/5713/7040/5701/5718/5690/10213/5717/5719/2033/5707 | 33 |
| normal | R-HSA-69239 | Synthesis of DNA | 38/1399 | 120/10704 | 8.57913987587149e-08 | 7.60759489655094e-07 | 5.15525867758886e-07 | 5687/7311/5689/5699/5700/119504/7316/5685/9978/5688/246184/6233/56655/5694/5702/5695/5684/5705/7314/6118/5693/5710/9861/5714/5686/5708/5713/5701/51529/5718/5690/5425/10213/7321/5717/5719/5707 | 38 |
| normal | R-HSA-73856 | RNA Polymerase II Transcription Termination | 26/1399 | 67/10704 | 1.04628696764 86e-07 | 9.21696216895711e-07 | 6.24584048554776e-07 | 7307/6430/6637/6428/6635/8683/4116/10914/6628/10250/6431/80145/6432/6636/6634/23283/55339/10569/6427/9295/10212/199746/10921/51585/55110/51362 | 26 |
| normal | R-HSA-5358351 | Signaling by Hedgehog | 43/1399 | 149/10704 | 2.42194569137818e-07 | 2.11959822271933e-06 | 1.43633793324251e-06 | 5687/7846/84617/7311/5689/5699/5700/6500/10376/7316/5685/9978/5688/6233/55081/5694/5702/5695/5684/5705/7314/5693/5573/5710/9861/2778/5714/5686/5708/10383/5713/408/409/51009/156/5701/5718/5690/79577/10213/5717/5719/5707 | 43 |
| normal | R-HSA-5696394 | DNA Damage Recognition in GG-NER | 18/1399 | 38/10704 | 2.79824664128233e-07 | 2.43302094329678e-06 | 1.64872768611144e-06 | 7311/7316/9978/6233/7314/51138/5886/5887/10980/60/125476/1069/142/8533/10987/10920/9318/86 | 18 |
| normal | R-HSA-6781823 | Formation of TC-NER Pre-Incision Complex | 22/1399 | 54/10704 | 3.67454103217065e-07 | 3.17432931746871e-06 | 2.15107257706696e-06 | 7311/902/7316/9978/3150/6233/5440/5441/7314/5138/10980/8533/5439/91603/7507/6917/10450/10987/10920/9318/5436/2033 | 22 |
| normal | R-HSA-5696399 | Global Genome Nucleotide Excision Repair (GG-NER) | 29/1399 | 84/10704 | 3.77219109964123e-07 | 3.23779736052539e-06 | 2.19408146281426e-06 | 6613/7311/902/7316/9978/7341/6233/56655/7314/6118/51138/5886/5887/10980/60/125476/7334/1069/142/8533/7336/7329/2067/5425/7507/10987/10920/9318/86 | 29 |
| normal | R-HSA-1257604 | PIP3 activates AKT signaling | 64/1399 | 264/10704 | 4.35442918699558e-07 | 3.71374565693445e-06 | 2.51660607387877e-06 | 5687/389541/7311/5689/5699/5700/10273/7316/5685/5879/10542/192669/5688/6233/5694/391/3164/5702/5695/23466/6009/5684/5705/5931/55004/7314/10014/8660/5290/5693/5710/9861/28956/207/9173/5714/5686/5708/84232/117145/5713/23512/23608/10670/5701/5527/5525/4615/5718/5690/253260/5594/231 | 64 |
| normal | R-HSA-8866654 | E3 ubiquitin ligases ubiquitinate target proteins | 23/1399 | 59/10704 | 5.14815671815662e-07 | 4.36289990228590e-06 | 2.95650305866157e-06 | 7311/7316/7323/6233/7322/7332/51255/3105/7314/80349/23168/123169/7334/7320/7336/10670/7319/9646/118424/5828/51322/79577/7321 | 23 |
| normal | R-HSA-71406 | Pyruvate metabolism and Citric Acid (TCA) cycle | 22/1399 | 55/10704 | 5.37468003943815e-07 | 4.52622425962747e-06 | 3.06717920820634e-06 | 3421/3945/3939/6392/51660/1431/6390/6832/3420/8801/6391/7416/2739/4191/3029/9123/5162/8802/6389/3419/5160/25874 | 22 |
| normal | R-HSA-69306 | DNA Replication | 38/1399 | 128/10704 | 5.49230071685936e-07 | 4.59636916242168e-06 | 3.11471264337682e-06 | 5687/7311/5689/5699/5700/119504/7316/5685/9978/5688/246184/6233/56655/5694/5702/5695/5684/5705/7314/6118/5693/5710/9861/5714/5686/5708/5713/5701/51529/5718/5690/5425/10213/7321/5717/5719/5707 | 38 |
| normal | R-HSA-983169 | Class I MHC mediated antigen processing & presentation | 82/1399 | 370/10704 | 6.48264129061514e-07 | 5.39146378144949e-06 | 3.65350558516525e-06 | 1535/5687/9021/8673/7311/5689/5699/5700/6500/10273/8651/7316/5685/9978/3107/7323/84676/5688/246184/10952/4012/6233/7322/5694/7332/5702/5695/55819/10134/23480/5684/5705/3105/7314/3106/5693/7916/5710/9861/5714/57674/26224/9616/55585/5686/8773/5708/2923/7334/5713/7320/3551/9520/73 | 82 |
| normal | R-HSA-447115 | Interleukin-12 family signaling | 22/1399 | 57/10704 | 1.11051659845688e-06 | 9.17889953909734e-06 | 6.22004748453434e-06 | 4282/1072/9987/998/5478/9446/6175/6647/5908/3181/27250/3185/3313/302/375/6888/6950/6627/9049/829/3936/1265 | 22 |
| normal | R-HSA-8852135 | Protein ubiquitination | 27/1399 | 79/10704 | 1.18737198479617e-06 | 9.75393305001882e-06 | 6.60971682850692e-06 | 7311/7316/7323/6233/7322/7332/51255/3105/7314/80349/23168/123169/7334/7320/7336/10670/7317/7319/10477/9646/118424/5828/51322/79577/7321/7327/55284 | 27 |
| normal | R-HSA-9006925 | Intracellular signaling by second messengers | 70/1399 | 306/10704 | 1.29160472865074e-06 | 1.05454800711179e-05 | 7.14610575158497e-06 | 5687/805/389541/7311/5689/5699/5700/10273/7316/5685/5879/10542/192669/5688/6233/5694/391/3164/5702/5695/23466/6009/5684/5705/5931/55004/7314/801/10014/8660/5290/5693/5573/5710/9861/28956/207/9173/5714/5686/5708/84232/117145/5713/10768/23512/23608/156/10670/5701/5527/5525/4615 | 70 |
| normal | R-HSA-5683057 | MAPK family signaling cascades | 73/1399 | 325/10704 | 1.58628846501319e-06 | 1.28731124691295e-05 | 8.72332157474558e-06 | 5687/1843/805/4023/7311/5689/5699/5700/5037/7316/5685/5879/9978/192669/5778/5688/5908/3674/6233/5694/5702/5695/5684/5705/11221/3568/7314/801/8660/8036/5290/5693/5710/9861/6709/28956/5714/60/808/5686/5708/5500/5713/5906/408/409/896/1439/5701/5245/5527/27040/5525/5125/5718/5 | 73 |
| normal | R-HSA-5673001 | RAF/MAP kinase cascade | 65/1399 | 280/10704 | 1.80937912295865e-06 | 1.45949316002508e-05 | 9.89020167400355e-06 | 5687/1843/805/4023/7311/5689/5699/5700/5037/7316/5685/9978/5778/5688/5908/3674/6233/5694/5702/5695/5684/5705/11221/3568/7314/801/8660/8036/5290/5693/5710/9861/6709/28956/5714/60/808/5686/5708/5500/5713/5906/408/409/1439/5701/5245/5527/27040/5525/5125/5718/5690/5594/8826/6464/11572 | 65 |
| normal | R-HSA-76005 | Response to elevated platelet cytosolic Ca2+ | 38/1399 | 134/10704 | 1.92421454865506e-06 | 1.54282831176594e-05 | 1.04549192621536e-05 | 226/8407/805/5216/308/1072/967/5478/6813/5874/6647/928/7078/3674/7114/10916/1191/7076/1521/5660/7422/801/5552/51368/808/7040/6814/51706/6810/1/10487/55664/5579/334/10490/7873/5627/10257 | 38 |
| normal | R-HSA-3371497 | HSP90 chaperone cycle for steroid hormone receptors (SHR) | 21/1399 | 55/10704 | 2.38798215093036e-06 | 1.90327863100938e-05 | 1.28974975820925e-05 | 7846/84617/10376/140735/830/8655/832/1781/10728/10294/3337/11258/3326/10383/55860/829/3320/10671/2289/2908/10121 | 21 |
| normal | R-HSA-5696395 | Formation of Incision Complex in GG-NER | 18/1399 | 43/10704 | 2.65328355461375e-06 | 2.10024614947912e-05 | 1.42322407204984e-05 | 6613/7311/902/7316/9978/7341/6233/7314/6118/5886/5887/7334/1069/142/7336/7329/2067/7507 | 18 |
| normal | R-HSA-389957 | Prefoldin mediated transfer of substrate to CCT/TriC | 14/1399 | 28/10704 | 2.66648129508178e-06 | 2.10024614979012e-05 | 1.42322407204984e-05 | 5204/7846/84617/5202/10694/60/10574/10383/6950/10575/5203/7411/22948/7203 | 14 |
| normal | R-HSA-69242 | S Phase | 43/1399 | 162/10704 | 2.87528112582956e-06 | 2.25146283091e-05 | 1.52569549428444e-05 | 5687/7311/5689/5699/5700/6500/119504/902/7316/5685/9978/5688/246184/6233/56655/5694/5702/5695/5684/5705/9126/7314/6118/5693/5710/9861/207/5714/5686/5708/5713/5701/51529/5718/5690/5425/1019/23063/10213/7321/5717/5719/5707 | 43 |
| normal | R-HSA-69206 | G1/S Transition | 37/1399 | 131/10704 | 2.90673964511777e-06 | 2.26286301442598e-05 | 1.53342079197767e-05 | 5687/7311/5689/5699/5700/6500/902/7316/5685/5688/6233/56655/5694/5702/5695/5684/5705/7314/6118/5693/5710/9861/207/5714/5686/5708/5713/5701/5718/5690/1019/10213/5515/5717/5719/3065/5707 | 37 |
| normal | R-HSA-389958 | Cooperation of Prefoldin and TriC/CCT in actin and tubulin folding | 15/1399 | 32/10704 | 3.28738092067787e-06 | 2.54439482820097e-05 | 1.72419978924551e-05 | 5204/7846/84617/10376/5202/10694/60/10574/10383/6950/10575/5203/7411/22948/7203 | 15 |
| normal | R-HSA-5684996 | MAPK1/MAPK3 signaling | 65/1399 | 286/10704 | 3.89125705474624e-06 | 2.99447884845127e-05 | 2.02919756877874e-05 | 5687/1843/805/4023/7311/5689/5699/5700/5037/7316/5685/9978/5778/5688/5908/3674/6233/5694/5702/5695/5684/5705/11221/3568/7314/801/8660/8036/5290/5693/5710/9861/6709/28956/5714/60/808/5686/5708/5500/5713/5906/408/409/1439/5701/5245/5527/27040/5525/5125/5718/5690/5594/8826/6464/11572 | 65 |
| normal | R-HSA-2173793 | Transcriptional activity of SMAD2/SMAD3:SMAD4 heterotrimer | 18/1399 | 44/10704 | 3.94548899127739e-06 | 3.01886271960961e-05 | 2.04572120931013e-05 | 7311/7316/6233/7050/7314/22938/3726/51592/1025/9611/142/6497/9612/6498/7321/5494/3065 | 18 |
| normal | R-HSA-6781827 | Transcription-Coupled Nucleotide Excision Repair (TC-NER) | 26/1399 | 79/10704 | 4.16278631181027e-06 | 3.16702890426929e-05 | 2.14612547893568e-05 | 7311/902/7316/9978/3150/6233/56655/5440/5441/7314/6118/51138/10980/8533/5439/2067/5425/91603/7507/6917/10450/10987/10920/9318/5436/2033 | 26 |
| normal | R-HSA-5205685 | PINK1-PRKN Mediated Mitophagy | 12/1399 | 22/10704 | 4.32685115992402e-06 | 3.27325067973913e-05 | 2.21810627407345e-05 | 100188893/7311/401505/7316/54543/6233/7314/56993/9804/9140/7416/8878 | 12 |
| normal | R-HSA-9607240 | FLT3 Signaling | 66/1399 | 294/10704 | 5.10506334592161e-06 | 3.84026956190395e-05 | 2.60234453233852e-05 | 5687/1843/805/4023/7311/5689/5699/5700/5037/7316/5685/9978/5778/5688/5908/3674/6233/5694/5702/5695/5684/5705/11221/3568/7314/801/8660/8036/5290/5693/5710/9861/6709/28956/5714/60/808/5686/5708/5500/117145/5713/5906/408/409/1439/5701/5245/5527/27040/5525/5125/5718/5690/5594/8826/646 | 66 |
| normal | R-HSA-72187 | mRNA 3'-end processing | 21/1399 | 58/10704 | 6.41931970256713e-06 | 4.77526060096521e-05 | 3.23593776819466e-05 | 7307/6430/6428/8683/4116/10914/10250/6431/80145/6432/23283/55339/10569/6427/9295/10212/199746/10921/51585/55110/51362 | 21 |
| normal | R-HSA-917937 | Iron uptake and transport | 21/1399 | 58/10704 | 6.41931970256713e-06 | 4.77526060096521e-05 | 3.23593776819466e-05 | 527/8992/2495/9296/9550/7311/2512/6500/533/7316/4738/6233/7314/10312/529/537/4891/9114/523/51382/23545 | 21 |
| normal | R-HSA-9663891 | Selective autophagy | 26/1399 | 81/10704 | 6.99028325415126e-06 | 5.17126479409346e-05 | 3.50428515195044e-05 | 100188893/7846/84617/7311/10376/401505/140735/7316/123/54543/11315/6233/8655/7431/7314/1781/56993/9804/7334/10383/472/9140/2034/7416/3320/8878 | 26 |
| normal | R-HSA-9637690 | Response of Mtb to phagocytosis | 12/1399 | 23/10704 | 7.97226539170546e-06 | 5.86530958381833e-05 | 3.97460541795842e-05 | 7311/7316/6233/7879/2023/7314/1511/57674/5230/5868/5594/3837 | 12 |
| normal | R-HSA-3700989 | Transcriptional Regulation by TP53 | 77/1399 | 365/10704 | 9.90574905940587e-06 | 7.21123499879425e-05 | 4.88666685190098e-05 | 4697/7295/2353/389541/1329/1337/7311/1327/4869/54205/902/1340/1345/1349/9167/7316/64769/5052/10542/6879/192669/1351/1350/124446/6233/51230/25824/7533/28984/3622/5440/5441/6009/7001/5595/55004/27166/5300/7314/6118/3276/28956/207/10971/665/1025/54541/84987/581/5439/900/472/10670/5 | 77 |
| normal | R-HSA-168927 | TICAM1, RIP1-mediated IKK complex recruitment | 11/1399 | 20/10704 | 9.90938939341406e-06 | 7.21123499879425e-05 | 4.88666685190098e-05 | 7311/7316/7323/6233/7322/7314/7334/3551/330/7321/329 | 11 |
| normal | R-HSA-1268020 | Mitochondrial protein import | 22/1399 | 64/10704 | 1.02721185295256e-05 | 7.43479281677553e-05 | 5.03815998432493e-05 | 100188893/51142/401505/293/131118/3421/54543/10063/10245/3313/1431/56993/1537/9804/4580/9512/79145/7416/90639/10440/26515/10651 | 22 |
| normal | R-HSA-9635486 | Infection with Mycobacterium tuberculosis | 13/1399 | 27/10704 | 1.03851013749687e-05 | 7.47615631241028e-05 | 5.06618980488004e-05 | 7311/7316/6233/7879/2023/7314/1511/57674/5230/5868/5594/567/3837 | 13 |
| normal | R-HSA-199977 | ER to Golgi Anterograde Transport | 40/1399 | 154/10704 | 1.11358412918581e-05 | 7.97373876459787e-05 | 5.40337472197112e-05 | 126003/7846/84617/966/10376/381/140735/11316/830/8655/79090/10342/10959/1604/832/58485/1781/10960/11014/6709/5861/375/11258/10383/54732/27095/6811/9117/23423/10175/55860/51272/22872/829/23243/10671/23256/10121/372/1314 | 40 |
| normal | R-HSA-453279 | Mitotic G1 phase and G1/S transition | 39/1399 | 149/10704 | 1.17609395507579e-05 | 8.37654152046003e-05 | 5.67633252673759e-05 | 5687/7311/5689/5699/5700/6500/902/7316/5685/5688/6233/56655/5694/5702/5695/5684/5705/7314/6118/5693/5710/896/5701/5718/5690/1019/4067/10213/5515/5717/5719/3065/5707 | 39 |
| normal | R-HSA-5627123 | RHO GTPases activate PAKs | 12/1399 | 24/10704 | 1.40519944824122e-05 | 9.94012816684067e-05 | 6.73589126199179e-05 | 10627/4637/805/103910/5879/998/4627/801/808/5500/4659/5058 | 12 |
| normal | R-HSA-114608 | Platelet degranulation | 35/1399 | 129/10704 | 1.41047375033587e-05 | 9.94012816684067e-05 | 6.73589126199179e-05 | 226/8407/805/5216/308/1072/967/5478/6813/5874/6647/928/7078/3674/7114/10916/1191/7076/1521/5660/7422/801/5552/51368/808/7040/51706/1/10487/55664/334/10490/7873/5627/10257 | 35 |
| normal | R-HSA-2871796 | FCERI mediated MAPK activation | 14/1399 | 32/10704 | 1.88093878992476e-05 | 0.000131862672236087 | 8.93562544455853e-05 | 2205/2207/2353/5879/3937/4214/2206/27040/5594/6464/4067/5058/3725/6850 | 14 |
| normal | R-HSA-9679506 | SARS-CoV Infections | 38/1399 | 147/10704 | 2.05895764385131e-05 | 0.000143590848831972 | 9.73038097039379e-05 | 1603/10284/7311/51652/51185/7316/3459/7341/6233/1655/128866/1173/2280/27243/7422/5926/5931/7314/3460/6484/10728/1175/25978/3326/23476/6185/7329/25805/55741/1514/483/23193/85437/3320/3615/2908/3688/3065 | 38 |
| normal | R-HSA-6807878 | COPI-mediated anterograde transport | 29/1399 | 101/10704 | 2.38543945682902e-05 | 0.000165497587186221 | 0.000112148830749202 | 7846/84617/966/10376/381/140735/11316/830/8655/10959/1604/832/1781/11014/6709/5861/375/11258/10383/6811/23423/55860/51272/829/10671/10121/372/1314 | 29 |
| normal | R-HSA-199992 | trans-Golgi Network Vesicle Budding | 23/1399 | 72/10704 | 2.54999044827024e-05 | 0.000176001918053291 | 0.000119267051894137 | 8673/2495/2512/6844/4074/2647/6643/1176/27243/2799/1212/8905/375/8773/8402/8907/408/5878/6810/1213/27131/23557/130340 | 23 |
| normal | R-HSA-3299685 | Detoxification of Reactive Oxygen Species | 15/1399 | 37/10704 | 2.86915278653491e-05 | 0.000197015158008731 | 0.000133506596598818 | 1535/7295/2876/2950/9588/54205/5052/6647/25824/7001/9973/10935/847/475/30001 | 15 |
| normal | R-HSA-9006936 | Signaling by TGFB family members | 29/1399 | 102/10704 | 2.92364863671446e-05 | 0.000199732934926566 | 0.000135348288122872 | 7311/10589/10273/7316/4738/7323/6233/23645/2280/7050/7314/22938/3726/11171/51592/1025/5500/9611/7048/142/7040/6497/5499/9612/6498/7321/387/5494/3065 | 29 |
| normal | R-HSA-71403 | Citric acid cycle (TCA cycle) | 11/1399 | 22/10704 | 3.2381833021515 9e-05 | 0.000218986234423282 | 0.000148395215654156 | 3421/6392/1431/6390/3420/8801/6391/4191/8802/6389/3419 | 11 |
| normal | R-HSA-9609523 | Insertion of tail-anchored proteins into the endoplasmic reticulum membrane | 11/1399 | 22/10704 | 3.2381833021515 9e-05 | 0.000218986234423282 | 0.000148395215654156 | 6844/10952/9218/819/23480/1528/5621/6811/27230/118424/2010 | 11 |
| normal | R-HSA-77387 | Insulin receptor recycling | 12/1399 | 26/10704 | 3.89934002803705e-05 | 0.000262372678268423 | 0.000177795879617452 | 527/8992/9296/9550/533/10312/529/537/9114/523/51382/23545 | 12 |
| normal | R-HSA-8866652 | Synthesis of active ubiquitin: roles of E1 and E2 enzymes | 13/1399 | 30/10704 | 4.23825825151619e-05 | 0.000283751383993009 | 0.000192283084884577 | 7311/7316/6233/7322/7332/7314/7320/7317/7319/10477/7321/7327/55284 | 13 |
| normal | R-HSA-2730905 | Role of LAT2/NTAL/LAB on calcium mobilization | 9/1399 | 16/10704 | 5.23032617926936e-05 | 0.000348428196718491 | 0.000236111084919099 | 2205/2207/5290/2206/7462/6464/4067/2534/6850 | 9 |
| normal | R-HSA-390450 | Folding of actin by CCT/TriC | 7/1399 | 10/10704 | 5.37325400011212e-05 | 0.000356117757951283 | 0.000241324236264339 | 10694/60/10574/6950/10575/22948/7203 | 7 |
| normal | R-HSA-2029482 | Regulation of actin dynamics for phagocytic cup formation | 20/1399 | 61/10704 | 5.59384350127568e-05 | 0.000368973224049662 | 0.000250033347062465 | 10094/1072/55845/5879/998/10092/10109/4627/3071/147179/10093/60/7456/3326/5594/10163/3320/5058/9844/6850 | 20 |
| normal | R-HSA-532668 | N-glycan trimming in the ER and Calnexin/Calreticulin cycle | 14/1399 | 35/10704 | 6.31875367962166e-05 | 0.000414745364593203 | 0.000281050860259746 | 7311/5700/7316/6233/51035/6048/7314/5887/11236/2923/51009/55741/811/23193 | 14 |
| normal | R-HSA-391251 | Protein folding | 28/1399 | 101/10704 | 6.54782598806564e-05 | 0.000427684829171702 | 0.000289819050152122 | 5204/7846/84617/402/10376/5202/2771/6902/1155/10694/83752/22803/60/10574/26224/10528/10383/6950/10575/5203/7411/2773/2783/2782/22948/6903/7203/10238 | 28 |
| normal | R-HSA-5627117 | RHO GTPases Activate ROCKs | 10/1399 | 20/10704 | 7.48668225237349e-05 | 0.000486634364044277 | 0.000329765973507713 | 10627/4637/103910/1072/389/4627/5500/4659/5058/387 | 10 |
| normal | R-HSA-68886 | M Phase | 82/1399 | 418/10704 | 8.07192560042025e-05 | 0.000522140501399165 | 0.000353826588379499 | 5687/4000/7846/84617/7311/5689/51652/5699/5700/119504/10376/140735/7316/5685/80097/5688/246184/7341/11243/5901/10726/6233/6232/8815/5694/2029/128866/8655/5702/27243/5695/4926/5862/5684/5705/9126/7314/1781/22919/5693/5710/9861/23175/1060/5861/5714/25978/5686/11258/5708/1069/10383 | 82 |
| normal | R-HSA-2173789 | TGF-beta receptor signaling activates SMADs | 13/1399 | 32/10704 | 9.54456871071033e-05 | 0.000611491745857 | 0.000414375131132325 | 7311/10273/7316/4738/6233/23645/2280/7314/11171/5500/7048/7040/5499 | 13 |
| normal | R-HSA-917729 | Endosomal Sorting Complex Required For Transport (ESCRT) | 13/1399 | 32/10704 | 9.54456871071033e-05 | 0.000611491745857 | 0.000414375131132325 | 7311/51652/7316/6233/51160/128866/7251/27243/7314/25978/11267/84313/51510 | 13 |
| normal | R-HSA-983168 | Antigen processing: Ubiquitination & Proteasome degradation | 64/1399 | 309/10704 | 9.73671228281808e-05 | 0.000620831356040481 | 0.000420704083099638 | 5687/9021/7311/5689/5699/5700/6500/10273/8651/7316/5685/9978/7323/84676/5688/246184/4012/6233/7322/5694/7332/5702/5695/55819/5684/5705/3106/7314/5693/79176/5710/9861/5714/57674/26224/9616/55585/5686/5708/7334/5713/7320/9520/7336/23608/5701/51529/7317/5718/5690/7319/10477/25793/80204/164 | 64 |
| normal | R-HSA-446203 | Asparagine N-linked glycosylation | 63/1399 | 304/10704 | 0.000108237182084984 | 0.000686870079676744 | 0.000465454981087852 | 126003/1603/7846/5973/84617/966/7311/5700/10376/381/140735/7316/6233/11316/8813/830/51035/8655/6048/79090/10342/10959/1604/832/7314/58485/1781/10960/5887/11014/6484/55907/6709/11236/5861/79868/375/11258/2923/10383/6185/54732/27095/10000/6811/9117/23423/10175/55860/55741/811/5127/63 | 63 |
| normal | R-HSA-8878171 | Transcriptional regulation by RUNX1 | 52/1399 | 239/10704 | 0.000118692006339455 | 0.000749661199083636 | 0.000508006065081038 | 3958/5687/9021/7311/5689/5699/5700/1997/902/7316/5685/192669/5688/3674/6233/5694/57492/5702/84661/55904/5695/23466/5684/5705/7314/3276/5693/8289/5710/9861/5714/5686/5708/2623/5713/5929/6595/896/5701/5718/5690/6688/23112/5579/10213/5717/5719/2033/86/3065/7707/5707 | 52 |
| normal | R-HSA-1236977 | Endosomal/Vacuolar pathway | 7/1399 | 11/10704 | 0.000131040583814415 | 0.000819922157605145 | 0.000555617231913553 | 3107/4012/3105/3106/567/1514/1520 | 7 |
| normal | R-HSA-9664873 | Pexophagy | 7/1399 | 11/10704 | 0.000131040583814415 | 0.000819922157605145 | 0.000555617231913553 | 7311/7316/6233/7314/472/2034/8878 | 7 |
| normal | R-HSA-429914 | Deadenylation-dependent mRNA decay | 18/1399 | 55/10704 | 0.000137214874791 1 | 0.000832804948411763 | 0.000564347232347 58 | 1973/1974/11157/27258/11340/25804/10605/51010/51690/10767/23658/80349/255967/26986/27257/1975/54512/9125 | 18 |
| normal | R-HSA-2871809 | FCERI mediated Ca+2 mobilization | 13/1399 | 33/10704 | 0.000138789862368414 | 0.000860368637552341 | 0.000583025640163609 | 2205/2207/805/3937/801/5530/2206/808/10768/27040/6464/4067/6850 | 13 |
| normal | R-HSA-937041 | IKK complex recruitment mediated by RIP1 | 11/1399 | 25/10704 | 0.000140457156991703 | 0.000867397844326681 | 0.000587788956259266 | 7311/7316/7323/6233/7322/7314/7334/3551/330/7321/329 | 11 |
| normal | R-HSA-162588 | Budding and maturation of HIV virion | 12/1399 | 29/10704 | 0.000143641401349604 | 0.000878245828343012 | 0.000595140052695787 | 7311/51652/7316/5478/6233/51160/128866/7251/27243/7314/25978/51510 | 12 |
| normal | R-HSA-5205647 | Mitophagy | 12/1399 | 29/10704 | 0.000143641401349604 | 0.000878245828343012 | 0.000595140052695787 | 100188893/7311/401505/7316/54543/6233/7314/56993/9804/9140/7416/8878 | 12 |
| normal | R-HSA-6790901 | rRNA modification in the nucleus and cytosol | 19/1399 | 61/10704 | 0.000184132595842431 | 0.00112069793560461 | 0.000759436830699403 | 6203/51504/6201/6208/6187/6194/4809/55505/51602/10199/10528/55651/56902/10436/51096/27341/55272/23160/51118 | 19 |
| normal | R-HSA-2029480 | Fcgamma receptor (FCGR) dependent phagocytosis | 24/1399 | 86/10704 | 0.000190371773785539 | 0.00114796501503357 | 0.000777914266702515 | 10094/1072/55845/5879/998/10092/10109/4627/5290/3071/147179/10093/60/7456/3326/10768/5594/4067/10163/3320/5058/2534/9844/6850 | 24 |
| normal | R-HSA-2132295 | MHC class II antigen presentation | 31/1399 | 123/10704 | 0.000190493924433665 | 0.00114796501503357 | 0.000777914266702515 | 10437/7846/84617/10376/140735/830/1173/8655/7879/832/1781/972/1175/1509/8905/375/11258/3799/10383/8907/1213/55860/1514/8722/22872/10120/1520/829/130340/10671/10121 | 31 |
| normal | R-HSA-6782135 | Dual incision in TC-NER | 20/1399 | 66/10704 | 0.000191184614154209 | 0.00114796501503357 | 0.000777914266702515 | 7311/902/7316/9978/3150/6233/56655/5440/5441/7314/6118/5439/2067/5425/91603/7507/6917/10450/5436/2033 | 20 |
| normal | R-HSA-69481 | G2/M Checkpoints | 39/1399 | 168/10704 | 0.000203454191455129 | 0.00121618376052865 | 0.000824142448469553 | 5687/7311/5689/5699/5700/7316/5685/5688/7341/6233/7533/5694/5702/5695/5684/5705/7314/165918/6118/5693/5710/9861/5714/10971/5686/5708/7334/5713/7336/472/5701/5718/5690/10213/4683/7532/5717/5719/5707 | 39 |
| normal | R-HSA-9613829 | Chaperone Mediated Autophagy | 10/1399 | 22/10704 | 0.000204458447830609 | 0.00121675494064527 | 0.000824525950657303 | 7311/7316/123/11315/6233/7431/7314/3326/1915/3320 | 10 |
| normal | R-HSA-2173788 | Downregulation of TGF-beta receptor signaling | 11/1399 | 26/10704 | 0.000215062227780013 | 0.00126858529069355 | 0.000859650546470535 | 7311/10273/7316/6233/23645/7314/11171/5500/7048/7040/5499 | 11 |
| normal | R-HSA-901042 | Calnexin/calreticulin cycle | 11/1399 | 28/10704 | 0.000215062227780013 | 0.00126858529069355 | 0.000859650546470535 | 7311/7316/6233/6048/7314/11236/2923/51009/55741/811/23193 | 11 |

| | | | | | | | | | |
|---|---|---|---|---|---|---|---|---|---|
| normal | R-HSA-9646399 | Aggrephagy | 15/1399 | 43/10704 | 0.000216043319829503 | 0.00126878072478818 | 0.000859784587687126 | 7846/84617/7311/10376/140735/7316/11315/6233/8655/7431/7314/1781/7334/10383/3320 | 15 |
| normal | R-HSA-8939236 | RUNX1 regulates transcription of genes involved in differentiation of HSCs | 32/1399 | 130/10704 | 0.000241237405198972 | 0.00141055408542107 | 0.000955856783642906 | 5687/7311/5689/5699/5700/902/7316/5685/5688/6233/5694/5702/5695/5684/5705/7314/5693/5710/9861/5714/5686/5708/2623/5713/5701/5718/5690/6688/10213/5717/5719/5707 | 32 |
| normal | R-HSA-114604 | GPVI-mediated activation cascade | 13/1399 | 35/10704 | 0.000277750035275106 | 0.00160304869497141 | 0.00108630004721934 | 2207/5879/998/391/3937/5290/27040/4067/5880/5777/387/2534/6850 | 13 |
| normal | R-HSA-432720 | Lysosome Vesicle Biogenesis | 13/1399 | 35/10704 | 0.000277750035275106 | 0.00160304869497141 | 0.00108630004721934 | 8673/6844/4074/2647/27243/2799/1212/8905/375/8907/408/1213/130340 | 13 |
| normal | R-HSA-9615710 | Late endosomal microautophagy | 13/1399 | 35/10704 | 0.000277750035275106 | 0.00160304869497141 | 0.00108630004721934 | 7311/51652/7316/123/11315/6233/51160/128866/7251/27243/7431/7314/25978 | 13 |
| normal | R-HSA-5689877 | Josephin domain DUBs | 7/1399 | 12/10704 | 0.000278977800135033 | 0.0016032243535657 | 0.00108641908161915 | 7311/7316/6233/7314/5886/5887/126119 | 7 |
| normal | R-HSA-8876725 | Protein methylation | 9/1399 | 19/10704 | 0.000292053635813783 | 0.00167119580493443 | 0.00113247968543176 | 805/6187/2109/801/1938/808/1915/151194/22944 | 9 |
| normal | R-HSA-1834949 | Cytosolic sensors of pathogen-associated DNA | 19/1399 | 63/10704 | 0.000293891001754349 | 0.00167455340999606 | 0.00113475495414221 | 7311/7316/81844/2547/6233/5440/9208/5441/84265/4792/7314/3428/7520/6778/51082/3551/4615/4791/2033 | 19 |
| normal | R-HSA-917977 | Transferrin endocytosis and recycling | 12/1399 | 31/10704 | 0.000303885232163295 | 0.00172416239774005 | 0.00116837230207297 | 527/8992/9296/9550/533/10312/529/537/9114/523/51382/23545 | 12 |
| normal | R-HSA-157118 | Signaling by NOTCH | 50/1399 | 236/10704 | 0.0003125629118575 | 0.00176591451045229 | 0.00119666546756014 | 5687/4904/7311/5689/5699/5700/6500/7316/5685/9978/192669/5688/6233/3091/5694/5702/5695/5684/5705/10959/5087/7314/22938/10014/5693/6484/5710/9861/207/5714/5686/5708/9611/5713/408/409/5701/9612/5718/5690/51107/488/23112/10213/5717/5719/2033/3725/3065/5707 | 50 |
| normal | R-HSA-9664407 | Parasite infection | 18/1399 | 59/10704 | 0.000359630814413855 | 0.00200644025208396 | 0.00135965685098571 | 10094/55845/5879/998/10092/10109/4627/3071/147179/10093/60/7456/5594/4067/10163/2534/9844/6850 | 18 |
| normal | R-HSA-9664417 | Leishmania phagocytosis | 18/1399 | 59/10704 | 0.000359630814413855 | 0.00200644025208396 | 0.00135965685098571 | 10094/55845/5879/998/10092/10109/4627/3071/147179/10093/60/7456/5594/4067/10163/2534/9844/6850 | 18 |
| normal | R-HSA-9664422 | FCGR3A-mediated phagocytosis | 18/1399 | 59/10704 | 0.000359630814413855 | 0.00200644025208396 | 0.00135965685098571 | 10094/55845/5879/998/10092/10109/4627/3071/147179/10093/60/7456/5594/4067/10163/2534/9844/6850 | 18 |
| normal | R-HSA-1222556 | ROS and RNS production in phagocytes | 13/1399 | 36/10704 | 0.000383269837684908 | 0.0021147561659088 | 0.00143305673126572 | 527/1535/8992/9296/9550/533/10312/529/9114/523/5880/51382/23545 | 13 |
| normal | R-HSA-5663213 | RHO GTPases Activate WASPs and WAVEs | 13/1399 | 36/10704 | 0.000383269837684908 | 0.0021147561659088 | 0.00143305673126572 | 10094/55845/5879/998/10092/10109/3071/147179/10093/60/7456/5594/10163 | 13 |
| normal | R-HSA-5689880 | Ub-specific processing proteases | 47/1399 | 220/10704 | 0.000383783232497322 | 0.0021147561659088 | 0.00143305673126572 | 5687/23770/7311/5689/5699/5700/8724/7316/5685/11047/5688/6233/3091/5694/7417/5702/5695/5684/5705/4792/7314/9804/5693/5710/9861/5714/5686/5708/23326/5713/7419/154/408/409/5701/10600/5718/330/5690/7416/10213/55031/329/5717/5719/9736/5707 | 47 |
| normal | R-HSA-948021 | Transport to the Golgi and subsequent modification | 41/1399 | 185/10704 | 0.000402271845084243 | 0.00219803070474937 | 0.00148948741597733 | 126003/7846/84617/966/10376/381/140735/11316/830/8655/79090/10342/10959/1604/832/58485/1781/10960/11014/6484/6709/5861/375/11258/10383/54732/27095/6811/9117/23423/10175/55860/51272/22872/829/23243/10671/23256/10121/372/1314 | 41 |
| normal | R-HSA-175474 | Assembly Of The HIV Virion | 8/1399 | 16/10704 | 0.000405461974662506 | 0.00219803070474937 | 0.00148948741597733 | 7311/7316/5478/6233/51160/7251/7314/9397 | 8 |
| normal | R-HSA-193639 | p75NTR signals via NF-kB | 8/1399 | 16/10704 | 0.000405461974662506 | 0.00219803070474937 | 0.00148948741597733 | 7311/7316/6233/4792/7314/3551/4615/8878 | 8 |
| normal | R-HSA-2173791 | TGF-beta receptor signaling in EMT (epithelial to mesenchymal transition) | 8/1399 | 16/10704 | 0.000405461974662506 | 0.00219803070474937 | 0.00148948741597733 | 7311/7316/6233/2280/7314/7048/7040/387 | 8 |
| normal | R-HSA-5626467 | RHO GTPases activate IQGAPs | 12/1399 | 32/10704 | 0.000428948959197008 | 0.00230797507156693 | 0.00156399081143877 | 7846/805/84617/10376/5879/998/801/60/808/10383/8826/10788 | 12 |
| normal | R-HSA-69620 | Cell Cycle Checkpoints | 59/1399 | 294/10704 | 0.000429190285900048 | 0.00230797507156693 | 0.00156399081143877 | 5687/7311/5689/5699/5700/119504/140735/7316/5685/5688/246184/7341/11243/10726/6233/51230/6232/7533/5694/8655/5702/5695/5684/5705/7314/1781/165918/6118/22919/5693/5710/9861/1060/5714/10971/5686/5708/7334/5713/7336/472/5701/51529/5527/5525/5718/5690/5903/10213/4683/7321/23122/7532/ | 59 |
| normal | R-HSA-9639288 | Amino acids regulate mTORC1 | 17/1399 | 55/10704 | 0.000438716044222696 | 0.00234976313285676 | 0.00159230833734722 | 527/8992/389541/9296/9550/533/10542/6009/55004/28956/10312/529/10670/9114/523/96459/51382 | 17 |
| normal | R-HSA-159236 | Transport of Mature mRNA derived from an Intron-Containing Transcript | 21/1399 | 75/10704 | 0.000446822082261882 | 0.00238364449461617 | 0.00161526791742396 | 7307/6430/6428/8683/4116/10250/6431/80145/6432/29107/10569/6427/10482/9295/53371/10212/199746/5903/10921/55110/51362 | 21 |
| normal | R-HSA-373755 | Semaphorin interactions | 19/1399 | 65/10704 | 0.000456754932840044 | 0.00242696371060642 | 0.00164462302467885 | 10627/4637/103910/7305/1072/5879/389/5788/10154/1808/4627/10507/3326/23365/3320/5058/387/3688/2534 | 19 |
| normal | R-HSA-164952 | The role of Nef in HIV-1 replication and disease pathogenesis | 11/1399 | 28/10704 | 0.000465892586202332 | 0.00246573190879416 | 0.00167089415147248 | 5879/1173/3105/1175/8905/375/8907/567/130340/2534/9844 | 11 |
| normal | R-HSA-5625900 | RHO GTPases activate CIT | 9/1399 | 20/10704 | 0.000469751937596549 | 0.00247636946630622 | 0.00167810265316297 | 10627/4637/103910/5879/389/4627/5500/4659/387 | 9 |
| normal | R-HSA-8856828 | Clathrin-mediated endocytosis | 34/1399 | 146/10704 | 0.00047630258492297 | 0.00250105553416415 | 0.00169483107617585 | 8673/7311/382/10094/7316/6844/4074/4738/10092/6233/1173/10109/7314/51138/1212/10093/1175/10980/60/5868/2060/154/8533/30011/408/409/5878/156/1213/50618/5869/10987/10920/9318 | 34 |
| normal | R-HSA-209543 | p75NTR recruits signalling complexes | 7/1399 | 13/10704 | 0.000536332104620956 | 0.00278352204685062 | 0.00188624346871997 | 7311/7316/6233/7314/3551/4615/8878 | 7 |
| normal | R-HSA-209560 | NF-kB is activated and signals survival | 7/1399 | 13/10704 | 0.000536332104620956 | 0.00278352204685062 | 0.00188624346871997 | 7311/7316/6233/4792/7314/3551/8878 | 7 |
| normal | R-HSA-9637687 | Suppression of phagosomal maturation | 7/1399 | 13/10704 | 0.000536332104620956 | 0.00278352204685062 | 0.00188624346871997 | 7311/7316/6233/7879/7314/5868/3837 | 7 |
| normal | R-HSA-432722 | Golgi Associated Vesicle Biogenesis | 17/1399 | 56/10704 | 0.000554408952609009 | 0.00286623006773538 | 0.00194229025461884 | 8673/2495/2512/6844/2647/6643/1176/8905/375/84062/8907/408/5878/1213/27131/23557/130340 | 17 |
| normal | R-HSA-5688426 | Deubiquitination | 59/1399 | 297/10704 | 0.000562079341828211 | 0.00289470861041529 | 0.00196158863423449 | 5687/23770/7311/5689/5699/5700/8724/7316/5685/4738/11047/5688/6233/3091/5694/7417/5702/5695/5684/5705/4792/7314/9804/5886/5693/5887/5710/9861/5714/60/5686/5708/125476/23326/5713/7419/7048/154/7040/408/409/5701/10600/5718/330/5690/7416/10213/55031/7321/387/329/126119/5717/5719/2033 | 59 |
| normal | R-HSA-6802946 | Signaling by moderate kinase activity BRAF mutants | 15/1399 | 47/10704 | 0.000644759274619078 | 0.00327019950270813 | 0.00221603865518997 | 805/5037/5908/3674/801/60/808/5906/408/409/5245/5594/8826/5894/5604 | 15 |
| normal | R-HSA-6802949 | Signaling by RAS mutants | 15/1399 | 47/10704 | 0.000644759274619078 | 0.00327019950270813 | 0.00221603865518997 | 805/5037/5908/3674/801/60/808/5906/408/409/5245/5594/8826/5894/5604 | 15 |
| normal | R-HSA-6802955 | Paradoxical activation of RAF signaling by kinase inactive BRAF | 15/1399 | 47/10704 | 0.000644759274619078 | 0.00327019950270813 | 0.00221603865518997 | 805/5037/5908/3674/801/60/808/5906/408/409/5245/5594/8826/5894/5604 | 15 |
| normal | R-HSA-9649948 | Signaling downstream of RAS mutants | 15/1399 | 47/10704 | 0.000644759274619078 | 0.00327019950270813 | 0.00221603865518997 | 805/5037/5908/3674/801/60/808/5906/408/409/5245/5594/8826/5894/5604 | 15 |
| normal | R-HSA-5213460 | RIPK1-mediated regulated necrosis | 11/1399 | 29/10704 | 0.000662823865834556 | 0.00332404927472835 | 0.00225252975497492 | 7311/10273/7316/6233/6386/11140/7314/8743/330/3320/329 | 11 |
| normal | R-HSA-5218859 | Regulated Necrosis | 11/1399 | 29/10704 | 0.000662823865834556 | 0.00332404927472835 | 0.00225252975497492 | 7311/10273/7316/6233/6386/11140/7314/8743/330/3320/329 | 11 |
| normal | R-HSA-5675482 | Regulation of necroptotic cell death | 11/1399 | 29/10704 | 0.000662823865834556 | 0.00332404927472835 | 0.00225252975497492 | 7311/10273/7316/6233/6386/11140/7314/8743/330/3320/329 | 11 |
| normal | R-HSA-1445148 | Translocation of SLC2A4 (GLUT4) to the plasma membrane | 20/1399 | 72/10704 | 0.000677457822930221 | 0.00338476128695361 | 0.00229367102657443 | 7846/805/84617/10376/6844/5879/4012/7533/4627/801/8766/207/60/808/10971/8773/10383/6814/6810/7532 | 20 |
| normal | R-HSA-389960 | Formation of tubulin folding intermediates by CCT/TriC | 10/1399 | 25/10704 | 0.000713275482534084 | 0.00355046792235367 | 0.00240596151807623 | 7846/84617/10376/10694/10574/10383/6950/10575/22948/7203 | 10 |
| normal | R-HSA-4420097 | VEGFA-VEGFR2 Pathway | 25/1399 | 99/10704 | 0.0007441239639211 | 0.00369030365811242 | 0.00250072068966857 | 1535/805/55845/5879/998/7422/801/5290/3071/207/60/2185/808/117145/10768/253260/9261/5579/10163/3320/5058/387/1432/2534/9844 | 25 |
| normal | R-HSA-195721 | Signaling by WNT | 64/1399 | 332/10704 | 0.000770122355112614 | 0.00380514329703243 | 0.00257854134825043 | 5687/805/5216/7311/5689/5699/5700/6500/8724/7316/5685/5879/9978/51699/192669/5688/6233/5694/1173/5702/5695/5684/5705/7314/801/5530/5693/1212/864/5710/9861/1175/207/9580/5714/808/5686/5708/123169/5713/5929/409/5701/5527/1213/5525/5718/5690/2783/2782/23112/5579/5880/79577/10213/387/ | 64 |
| normal | R-HSA-5675221 | Negative regulation of MAPK pathway | 14/1399 | 43/10704 | 0.000774921159803797 | 0.00381477732712237 | 0.00258506981327737 | 1843/7311/5037/7316/5778/6233/11221/7314/5527/5525/5594/5894/5515/5604 | 14 |
| normal | R-HSA-381119 | Unfolded Protein Response (UPR) | 24/1399 | 94/10704 | 0.000795414788973763 | 0.00390132015544274 | 0.00264371524231881 | 4000/1051/11340/1968/6347/8894/51010/37/1965/10488/5611/64764/27230/811/118/1200/9114/54512/22872/10897/56005/6464/64061/3068 | 24 |
| normal | R-HSA-195258 | RHO GTPase Effectors | 63/1399 | 327/10704 | 0.000856578041333557 | 0.00418597899250231 | 0.00283661264552258 | 1535/7846/10627/4637/805/5216/84617/103910/10094/1072/10376/140735/55845/5879/998/389/10092/11243/10726/6232/3895/7533/391/10109/8655/6280/5300/4627/1781/801/22919/3071/147179/10093/1060/60/808/752/10971/7456/3799/5500/10383/4659/91010/5527/5525/5594/5579/8826/10788/10163/5880/5 | 63 |
| normal | R-HSA-72202 | Transport of Mature Transcript to Cytoplasm | 22/1399 | 84/10704 | 0.000897095801302882 | 0.00436804101070749 | 0.002959986 9118587 | 7307/6430/6428/8683/4116/10250/6431/80145/6432/29107/55339/10569/6427/10482/9295/53371/10212/199746/5903/10921/55110/51362 | 22 |
| normal | R-HSA-390466 | Chaperonin-mediated protein folding | 24/1399 | 95/10704 | 0.000935077574637892 | 0.00450455743940069 | 0.00305249676723666 | 5204/7846/84617/10376/5202/2771/10694/83752/22803/60/10574/26224/10528/10383/6950/10575/5203/7411/2773/2783/2782/22948/7203/10238 | 24 |
| normal | R-HSA-174490 | Membrane binding and targetting of GAG proteins | 7/1399 | 14/10704 | 0.000952046120500669 | 0.00450455743940069 | 0.00305249676723666 | 7311/7316/6233/51160/7251/7314/9397 | 7 |
| normal | R-HSA-174495 | Synthesis And Processing Of GAG, GAGPOL Polyproteins | 7/1399 | 14/10704 | 0.000952046120500669 | 0.00450455743940069 | 0.00305249676723666 | 7311/7316/6233/51160/7251/7314/9397 | 7 |
| normal | R-HSA-877312 | Regulation of IFNG signaling | 7/1399 | 14/10704 | 0.000952046120500669 | 0.00450455743940069 | 0.00305249676723666 | 9021/8651/3459/7341/3460/5770/5777 | 7 |
| normal | R-HSA-9005891 | Loss of function of MECP2 in Rett syndrome | 7/1399 | 14/10704 | 0.000952046120500669 | 0.00450455743940069 | 0.00305249676723666 | 805/801/808/9611/9612/4204/3065 | 7 |
| normal | R-HSA-9005895 | Pervasive developmental disorders | 7/1399 | 14/10704 | 0.000952046120500669 | 0.00450455743940069 | 0.00305249676723666 | 805/801/808/9611/9612/4204/3065 | 7 |
| normal | R-HSA-9675151 | Disorders of Developmental Biology | 7/1399 | 14/10704 | 0.000952046120500669 | 0.00450455743940069 | 0.00305249676723666 | 805/801/808/9611/9612/4204/3065 | 7 |
| normal | R-HSA-9697154 | Disorders of Nervous System Development | 7/1399 | 14/10704 | 0.000952046120500669 | 0.00450455743940069 | 0.00305249676723666 | 805/801/808/9611/9612/4204/3065 | 7 |
| normal | R-HSA-8848021 | Signaling by PTK6 | 16/1399 | 54/10704 | 0.00107039339221228 | 0.00502897106025349 | 0.00340786372700642 | 9021/7311/7316/5879/6233/3091/7314/10657/207/2034/1019/5770/116985/387/2908/9844 | 16 |
| normal | R-HSA-9006927 | Signaling by Non-Receptor Tyrosine Kinases | 16/1399 | 54/10704 | 0.00107039339221228 | 0.00502897106025349 | 0.00340786372700642 | 9021/7311/7316/5879/6233/3091/7314/10657/207/2034/1019/5770/116985/387/2908/9844 | 16 |
| normal | R-HSA-194138 | Signaling by VEGF | 26/1399 | 108/10704 | 0.00126477639946174 | 0.00586862811061793 | 0.00397685423635286 | 1535/805/55845/5879/998/7422/801/14/5290/3071/207/60/2185/808/117145/10768/253260/9261/5579/10163/3320/5058/387/1432/2534/9844 | 26 |
| normal | R-HSA-70268 | Pyruvate metabolism | 11/1399 | 31/10704 | 0.00126556041849675 | 0.00586862811061793 | 0.00397685423635286 | 3945/3939/51660/682/7416/2739/3029/9123/5162/5160/25874 | 11 |
| normal | R-HSA-2025928 | Calcineurin activates NFAT | 6/1399 | 11/10704 | 0.00126664191483837 | 0.00586862811061793 | 0.00397685423635286 | 805/5478/2280/801/5530/808 | 6 |
| normal | R-HSA-5625970 | RHO GTPases activate KTN1 | 6/1399 | 11/10704 | 0.00126664191483837 | 0.00586862811061793 | 0.00397685423635286 | 5879/998/3895/391/3799/387 | 6 |
| normal | R-HSA-3899300 | SUMOylation of transcription cofactors | 14/1399 | 45/10704 | 0.00128209092073333 | 0.00590736062526661 | 0.004003101182327 6 | 6613/4869/7341/11315/1655/3622/10210/10521/6294/7329/9612/10155/2033/1487 | 14 |
| normal | R-HSA-6782210 | Gap-filling DNA repair synthesis and ligation in TC-NER | 18/1399 | 65/10704 | 0.00128382519936713 | 0.00590736062526661 | 0.004003101182327 6 | 7311/902/7316/9978/3150/6233/56655/5440/5441/7314/6118/5439/5425/91603/6917/10450/5436/2033 | 18 |
| normal | R-HSA-2682334 | EPH-Ephrin signaling | 23/1399 | 92/10704 | 0.00137768365436977 | 0.00631752881233259 | 0.00428105014443669 | 10627/4637/103910/10094/1072/5879/998/10092/1173/10109/6386/4627/1212/10093/1175/60/1213/51107/7074/4067/5058/387/2534 | 23 |
| normal | R-HSA-6804757 | Regulation of TP53 Degradation | 12/1399 | 36/10704 | 0.00144249998112519 | 0.00659217568166085 | 0.00446716358444373 | 7311/7316/6233/51230/7314/207/900/472/5527/253260/5515/6446 | 12 |
| normal | R-HSA-8949215 | Mitochondrial calcium ion transport | 9/1399 | 23/10704 | 0.00158427985024242 | 0.00721547863766871 | 0.00488954253816314 | 91689/7417/55486/30968/10730/7419/5245/9512/7416 | 9 |
| normal | R-HSA-168643 | Nucleotide-binding domain, leucine rich repeat containing receptor (NLR) signaling pathways | 16/1399 | 56/10704 | 0.00163959672177069 | 0.00742069062240117 | 0.00502860368421823 | 7295/7311/7316/6233/29108/7314/14/7334/3326/3551/330/10910/329/4791/1432/837 | 16 |
| normal | R-HSA-6811436 | COPI-independent Golgi-to-ER retrograde traffic | 15/1399 | 51/10704 | 0.00164865857283914 | 0.00742069062240117 | 0.00502860368421823 | 7846/84617/10376/140735/830/8655/832/1781/11258/10383/55860/829/10671/10121/22931 | 15 |
| normal | R-HSA-9678108 | SARS-CoV-1 Infection | 15/1399 | 51/10704 | 0.00164865857283914 | 0.00742069062240117 | 0.00502860368421823 | 7311/51652/7316/7341/6233/1655/128866/27243/7314/6484/25978/7329/1514/23193/85437 | 15 |
| normal | R-HSA-1236382 | Constitutive Signaling by Ligand-Responsive EGFR Cancer Variants | 8/1399 | 19/10704 | 0.00165704742053618 | 0.00742069062240117 | 0.00502860368421823 | 7311/7316/6233/11140/7314/5290/6464/3320 | 8 |
| normal | R-HSA-5637815 | Signaling by Ligand-Responsive EGFR Variants in Cancer | 8/1399 | 19/10704 | 0.00165704742053618 | 0.00742069062240117 | 0.00502860368421823 | 7311/7316/6233/11140/7314/5290/6464/3320 | 8 |
| normal | R-HSA-5619115 | Disorders of transmembrane transporters | 37/1399 | 175/10704 | 0.00185903287147942 | 0.00829748338303648 | 0.00562275907093073 | 5687/7311/5689/5699/5700/7316/5685/5688/6233/5694/5702/6048/5695/5684/6520/5705/7314/5693/682/5710/9861/5714/5686/5708/5713/51009/4891/1836/5701/5718/5690/53371/5903/10213/5717/5719/5707 | 37 |
| normal | R-HSA-168164 | Toll Like Receptor 3 (TLR3) Cascade | 23/1399 | 94/10704 | 0.00187232042552636 | 0.00832902674345447 | 0.0056441343130048 | 2353/3146/7311/6500/7316/7323/6233/10010/7322/4792/7314/7334/3551/330/5594/9261/7321/5515/329/4791/1432/3725/5604 | 23 |
| normal | R-HSA-6806003 | Regulation of TP53 Expression and Degradation | 12/1399 | 37/10704 | 0.001884826863522 | 0.00835689791475482 | 0.00566302110964086 | 7311/7316/6233/51230/7314/207/900/472/5527/253260/5515/6446 | 12 |
| normal | R-HSA-194315 | Signaling by Rho GTPases | 81/1399 | 455/10704 | 0.00198738153054911 | 0.008777125528361 | 0.00594778680511551 | 1535/7846/10627/4637/805/5216/84617/103910/10094/1072/10376/140735/55845/5879/998/389/10092/11243/399/10726/6232/3895/397/7533/391/93663/10109/8655/6280/7204/5300/4627/1781/801/22919/3071/147179/10093/396/1060/392/60/808/752/10971/7456/3799/5500/23365/11214/10383/9181/9459/1124/4 | 81 |
| normal | R-HSA-3928662 | EPHB-mediated forward signaling | 13/1399 | 42/10704 | 0.00199271557925448 | 0.008777125528361 | 0.00594778680511551 | 10094/1072/5879/998/10092/10109/10093/60/7074/4067/5058/387/2534 | 13 |
| normal | R-HSA-8856825 | Cargo recognition for clathrin-mediated endocytosis | 25/1399 | 106/10704 | 0.00210696630711413 | 0.00924992749254369 | 0.00626817931572868 | 8673/7311/7316/6844/4074/4738/6233/1173/7314/51138/1212/1175/10980/2060/154/8533/30011/408/409/156/1213/50618/10987/10920/9318 | 25 |
| normal | R-HSA-9694516 | SARS-CoV-2 Infection | 18/1399 | 68/10704 | 0.00223890915977703 | 0.00978879399943352 | 0.0066333402205194 | 1603/7311/51652/7316/7341/6233/1655/128866/27243/7314/6484/25978/6185/7329/55741/1514/23193/85437 | 18 |
| normal | R-HSA-166208 | mTORC1-mediated signalling | 9/1399 | 24/10704 | 0.00224433140987759 | 0.00978879399943352 | 0.0066333402205194 | 389541/10542/6194/2280/6009/55004/28956/10670/1975 | 9 |
| normal | R-HSA-3371511 | HSF1 activation | 6/1399 | 12/10704 | 0.00225430602954171 | 0.00980037588816995 | 0.00664118864478794 | 6118/10728/3326/1915/3281/3320 | 6 |
| normal | R-HSA-9033241 | Peroxisomal protein import | 17/1399 | 63/10704 | 0.00234974043215394 | 0.0101822085393337 | 0.00689993613529788 | 7311/7316/7323/6233/7322/6342/7314/373156/10455/83752/11001/54469/847/5828/7321/3295/5264 | 17 |
| normal | R-HSA-5684264 | MAP3K8 (TPL2)-dependent MAPK1/3 activation | 7/1399 | 16/10704 | 0.00250208812635513 | 0.01080740645545 | 0.00732359920175932 | 7311/6500/7316/6233/7314/3551/5604 | 7 |
| normal | R-HSA-9656223 | Signaling by RAF1 mutants | 13/1399 | 43/10704 | 0.00252042649434976 | 0.0108516111766377 | 0.00735355436835165 | 805/5908/3674/801/60/808/5906/408/409/5594/8826/5894/5604 | 13 |
| normal | R-HSA-400685 | Sema4D in semaphorin signaling | 9/1399 | 25/10704 | 0.00310555036900833 | 0.0133279870003274 | 0.00903166132957214 | 10627/4637/103910/5879/389/4627/10507/23365/387 | 9 |
| normal | R-HSA-9679191 | Potential therapeutics for SARS | 20/1399 | 81/10704 | 0.00321392974340557 | 0.013749047688243 | 0.00931699155478595 | 10284/51185/3459/1173/2280/7422/5926/5931/3460/10728/1175/3326/23476/25855/483/3320/3615/2908/3688/3065 | 20 |
| normal | R-HSA-164938 | Nef-mediates down modulation of cell surface receptors by recruiting them to clathrin adapters | 8/1399 | 21/10704 | 0.00350777356111357 | 0.0148636354377566 | 0.0100722878403728 | 1173/3105/1175/8905/375/8907/567/130340 | 8 |
| normal | R-HSA-416572 | Sema4D induced cell migration and growth-cone collapse | 8/1399 | 21/10704 | 0.00350777356111357 | 0.0148636354377566 | 0.0100722878403728 | 10627/4637/103910/389/4627/10507/23365/387 | 8 |
| normal | R-HSA-901032 | ER Quality Control Compartment (ERQC) | 8/1399 | 21/10704 | 0.00350777356111357 | 0.0148636354377566 | 0.0100722878403728 | 7311/7316/6233/6048/7314/11236/51009/55741 | 8 |
| normal | R-HSA-203615 | eNOS activation | 6/1399 | 13/10704 | 0.00372685510804521 | 0.0157421419232572 | 0.0106676045264319 | 805/23564/801/207/808/3320 | 6 |
| normal | R-HSA-202131 | Metabolism of nitric oxide: NOS3 activation and regulation | 7/1399 | 17/10704 | 0.00377858192260272 | 0.0159104440074373 | 0.0107816538142454 | 805/23564/801/51070/207/808/3320 | 7 |
| normal | R-HSA-166166 | MyD88-independent TLR4 cascade | 23/1399 | 99/10704 | 0.003808115120496 | 0.0159345817073255 | 0.0107980106377222 | 2353/3146/7311/6500/7316/7323/6233/10010/7322/4792/7314/7334/3551/330/5594/9261/7321/5515/329/4791/1432/3725/5604 | 23 |
| normal | R-HSA-937061 | TRIF(TICAM1)-mediated TLR4 signaling | 23/1399 | 99/10704 | 0.003808115120496 | 0.0159345817073255 | 0.0107980106377222 | 2353/3146/7311/6500/7316/7323/6233/10010/7322/4792/7314/7334/3551/330/5594/9261/7321/5515/329/4791/1432/3725/5604 | 23 |
| normal | R-HSA-5674135 | MAP2K and MAPK activation | 12/1399 | 40/10704 | 0.00391539191317903 | 0.0163324291954726 | 0.0110676105235623 | 5037/5908/3674/28956/60/5906/408/409/5594/8826/5894/5604 | 12 |
| normal | R-HSA-6803529 | FGFR2 alternative splicing | 9/1399 | 26/10704 | 0.00420719886431869 | 0.0174951530413749 | 0.0118555260576748 | 3178/4670/5440/5441/3185/5439/5725/2962/5436 | 9 |
| normal | R-HSA-112382 | Formation of RNA Pol II elongation complex | 16/1399 | 61/10704 | 0.00424085196818315 | 0.0175262369919668 | 0.0118765899823713 | 902/5440/5441/80349/1025/23168/123169/5439/6830/11198/9646/6917/79577/2962/5436/27125 | 16 |
| normal | R-HSA-75955 | RNA Polymerase II Transcription Elongation | 16/1399 | 61/10704 | 0.00424085196818315 | 0.0175262369919668 | 0.0118765899823713 | 902/5440/5441/80349/1025/23168/123169/5439/6830/11198/9646/6917/79577/2962/5436/27125 | 16 |
| normal | R-HSA-5673000 | RAF activation | 11/1399 | 36/10704 | 0.00484217086110302 | 0.0198885484141624 | 0.0134774016217979 | 805/801/8036/808/5500/5245/5527/5525/5894/5515/5604 | 11 |
| normal | R-HSA-6802948 | Signaling by high-kinase activity BRAF mutants | 11/1399 | 36/10704 | 0.00484217086110302 | 0.0198885484141624 | 0.0134774016217979 | 5037/5908/3674/60/5906/408/409/5594/8826/5894/5604 | 11 |
| normal | R-HSA-389977 | Post-chaperonin tubulin folding pathway | 8/1399 | 22/10704 | 0.00488969542949254 | 0.0199612871344223 | 0.013526692747826 | 7846/84617/402/10376/6902/1155/10383/6903 | 8 |
| normal | R-HSA-912631 | Regulation of signaling by CBL | 8/1399 | 22/10704 | 0.00488969542949254 | 0.0199612871344223 | 0.013526692747826 | 7311/7316/6233/7314/5290/4067/2534/6850 | 8 |
| normal | R-HSA-6802957 | Oncogenic MAPK signaling | 20/1399 | 84/10704 | 0.00501046331349838 | 0.0203921288047852 | 0.0138186510197908 | 4000/805/5037/5908/3674/11221/801/8036/60/808/5500/5906/408/409/5245/5594/10142/8826/5894/5604 | 20 |
| normal | R-HSA-111367 | SLBP independent Processing of Histone Pre-mRNAs | 5/1399 | 10/10704 | 0.00539906360983881 | 0.0218409249956924 | 0.0148004224254446 | 6637/6635/6628/6636/6634 | 5 |
| normal | R-HSA-164940 | Nef mediated downregulation of MHC class I complex cell surface expression | 5/1399 | 10/10704 | 0.00539906360983881 | 0.0218409249956924 | 0.0148004224254446 | 3105/8905/8907/567/130340 | 5 |
| normal | R-HSA-9648002 | RAS processing | 9/1399 | 27/10704 | 0.00559132662355117 | 0.0225505612919729 | 0.015281304849401 1 | 805/402/7311/7316/6233/7314/801/808/51125 | 9 |
| normal | R-HSA-3371568 | Attenuation phase | 6/1399 | 14/10704 | 0.00580796184870362 | 0.0232840147168088 | 0.0157783268628507 | 10728/3337/3326/3281/3320/2033 | 6 |
| normal | R-HSA-975110 | TRAF6 mediated IRF7 activation in TLR7/8 or 9 signaling | 6/1399 | 14/10704 | 0.00580796184870362 | 0.0232840147168088 | 0.0157783268628507 | 7311/7316/6233/7314/7334/4615 | 6 |
| normal | R-HSA-983695 | Antigen activates B Cell Receptor (BCR) leading to generation of second messengers | 10/1399 | 32/10704 | 0.00596425420631468 | 0.0238392130813593 | 0.0161545549908665 | 805/801/808/10768/30011/4067/5777/80228/2534/6850 | 10 |
| normal | R-HSA-168638 | NOD1/2 Signaling Pathway | 11/1399 | 37/10704 | 0.00609335486007962 | 0.0242827445168054 | 0.01645511243542 8 | 7311/7316/6233/7314/14/7334/3551/330/329/1432/837 | 11 |
| normal | R-HSA-193704 | p75 NTR receptor-mediated signalling | 22/1399 | 97/10704 | 0.00621292813838828 | 0.0246857886566822 | 0.0167282338131835 | 7311/7316/5879/57142/6233/7204/4792/7314/396/23365/11214/9181/9459/3551/27018/4615/51107/7074/55701/8878/387/3065 | 22 |
| normal | R-HSA-3108232 | SUMO E3 ligases SUMOylate target proteins | 36/1399 | 182/10704 | 0.00666878725910743 | 0.0264186572187717 | 0.0179025058154799 | 6613/4869/7341/11315/7376/3190/1655/3622/4929/9126/4792/165918/51602/10210/10521/1069/23512/142/3183/6294/7329/9612/54780/23468/6304/10155/53371/4286/5903/6672/4791/2033/2908/23137/1487/3065 | 36 |
| normal | R-HSA-2122947 | NOTCH1 Intracellular Domain Regulates Transcription | 13/1399 | 48/10704 | 0.00712903243635567 | 0.0281586266438945 | 0.0190815896914721 | 7311/6500/7316/9978/6233/3091/7314/22938/10014/9611/9612/2033/3065 | 13 |
| normal | R-HSA-162587 | HIV Life Cycle | 31/1399 | 152/10704 | 0.0072157678799771 | 0.0284173917390863 | 0.0192569409056974 | 7311/51652/902/7316/5478/6879/5902/2547/5901/6233/8815/51160/128866/5440/5441/7251/27243/7314/2958/7520/25978/1025/9397/5439/51510/11198/53371/6917/5903/2962/5436 | 31 |
| normal | R-HSA-75067 | Processing of Capped Intronless Pre-mRNA | 9/1399 | 28/10704 | 0.00730217902886791 | 0.0286733657467863 | 0.0194304007497581 | 6637/6635/10914/6628/6636/6634/23283/55339/51585 | 9 |
| normal | R-HSA-400253 | Circadian Clock | 17/1399 | 70/10704 | 0.00753044598389358 | 0.0294832373462968 | 0.0199792072579756 | 7311/6500/7316/6233/3091/10135/7314/26224/5500/9611/5499/1390/8553/64784/5187/2033/2908 | 17 |

| | | | | | | | | | |
|---|---|---|---|---|---|---|---|---|---|
| normal | R-HSA-6804760 | Regulation of TP53 Activity through Methylation | 7/1399 | 19/10704 | 0.00773283872092409 | 0.0301873791466978 | 0.0204563663570249 | 7311/7316/6233/7314/472/79813/2033 | 7 |
| normal | R-HSA-2565942 | Regulation of PLK1 Activity at G2/M Transition | 20/1399 | 88/10704 | 0.00861824460752751 | 0.0330851692355956 | 0.0224200431438099 | 7846/7311/6500/7316/6233/8655/7314/1781/22919/11258/5500/1069/10383/4659/11064/10142/3320/7532/10121/5108 | 20 |
| normal | R-HSA-430039 | mRNA decay by 5' to 3' exoribonuclease | 6/1399 | 15/10704 | 0.00862339362451297 | 0.0330851692355956 | 0.0224200431438099 | 11157/27258/25804/51690/23658/27257 | 6 |
| normal | R-HSA-5099900 | WNT5A-dependent internalization of FZD4 | 6/1399 | 15/10704 | 0.00862339362451297 | 0.0330851692355956 | 0.0224200431438099 | 1173/1212/1175/409/1213/5579 | 6 |
| normal | R-HSA-8875360 | InlB-mediated entry of Listeria monocytogenes into host cell | 6/1399 | 15/10704 | 0.00862339362451297 | 0.0330851692355956 | 0.0224200431438099 | 7311/7316/6233/7314/2060/30011 | 6 |
| normal | R-HSA-937039 | IRAK1 recruits IKK complex | 6/1399 | 15/10704 | 0.00862339362451297 | 0.0330851692355956 | 0.0224200431438099 | 7311/7316/6233/7314/7334/3551 | 6 |
| normal | R-HSA-975144 | IRAK1 recruits IKK complex upon TLR7/8 or 9 stimulation | 6/1399 | 15/10704 | 0.00862339362451297 | 0.0330851692355956 | 0.0224200431438099 | 7311/7316/6233/7314/7334/3551 | 6 |
| normal | R-HSA-1234158 | Regulation of gene expression by Hypoxia-inducible Factor | 5/1399 | 11/10704 | 0.00884274835567251 | 0.0333121757295379 | 0.02257387325804 | 25994/3091/7422/2034/2033 | 5 |
| normal | R-HSA-4755510 | SUMOylation of immune response proteins | 5/1399 | 11/10704 | 0.00884274835567251 | 0.0333121757295379 | 0.02257387325804 | 7341/4792/10210/7329/4791 | 5 |
| normal | R-HSA-77286 | mitochondrial fatty acid beta-oxidation of saturated fatty acids | 5/1399 | 11/10704 | 0.00884274835567251 | 0.0333121757295379 | 0.02257387325804 | 37/1892/3030/34/3032 | 5 |
| normal | R-HSA-77588 | SLBP Dependent Processing of Replication-Dependent Histone Pre-mRNAs | 5/1399 | 11/10704 | 0.00884274835567251 | 0.0333121757295379 | 0.02257387325804 | 6637/6635/6628/6636/6634 | 5 |
| normal | R-HSA-174048 | APC/C:Cdc20 mediated degradation of Cyclin B | 8/1399 | 24/10704 | 0.00885670990270014 | 0.0333121757295379 | 0.02257387325804 | 7311/119504/7316/246184/6233/7314/51529/7321 | 8 |
| normal | R-HSA-5668599 | RHO GTPases Activate NADPH Oxidases | 8/1399 | 24/10704 | 0.00885670990270014 | 0.0333121757295379 | 0.02257387325804 | 1535/5879/6280/5300/5594/5579/5880/1432 | 8 |
| normal | R-HSA-982772 | Growth hormone receptor signaling | 8/1399 | 24/10704 | 0.00885670990270014 | 0.0333121757295379 | 0.02257387325804 | 9021/8651/8660/8835/5594/5770/4067/5777 | 8 |
| normal | R-HSA-390471 | Association of TriC/CCT with target proteins during biosynthesis | 11/1399 | 39/10704 | 0.0093455815179092 | 0.035052475217032 | 0.0237531808003471 | 10694/83752/22803/10574/26224/10528/6950/10575/22948/7203/10238 | 11 |
| normal | R-HSA-5218920 | VEGFR2 mediated vascular permeability | 9/1399 | 29/10704 | 0.00938552163003689 | 0.0351039482196073 | 0.0237880612910667 | 805/5879/801/207/808/117145/253260/3320/5058 | 9 |
| normal | R-HSA-70263 | Gluconeogenesis | 10/1399 | 34/10704 | 0.00950266909093127 | 0.0353446497576583 | 0.0239511718022888 | 226/7167/2597/5223/2023/4190/5230/2203/8402/4191 | 10 |
| normal | R-HSA-936440 | Negative regulators of DDX58/IFIH1 signaling | 10/1399 | 34/10704 | 0.00950266909093127 | 0.0353446497576583 | 0.0239511718022888 | 7311/7316/7323/6233/7322/8887/5300/7314/9140/7321 | 10 |
| normal | R-HSA-168898 | Toll-like Receptor Cascades | 31/1399 | 155/10704 | 0.00960134874642742 | 0.0356127589237294 | 0.0241328549917494 | 2353/3146/7311/6500/8651/7316/7323/6233/10010/7322/6280/4214/4792/7314/7334/3551/8411/4615/330/5594/1514/10695/9261/1520/7321/5515/329/4791/1432/3725/5604 | 31 |
| normal | R-HSA-3371556 | Cellular response to heat stress | 20/1399 | 89/10704 | 0.00978877814665231 | 0.0362076628131697 | 0.0245359894225481 | 6767/3313/10049/6118/10728/3337/10808/3326/472/1915/51501/3281/573/5594/53371/9261/3320/5903/91582/2033 | 20 |
| normal | R-HSA-381038 | XBP1(S) activates chaperone genes | 13/1399 | 50/10704 | 0.0102308035837727 | 0.037738418729123 | 0.0255733005341803 | 4000/37/5611/27230/118/1200/9114/22872/10897/56005/6464/64061/3068 | 13 |
| normal | R-HSA-6802952 | Signaling by BRAF and RAF fusions | 16/1399 | 67/10704 | 0.0110419518660237 | 0.0406186086500158 | 0.0275250506319041 | 4000/805/5037/5908/3674/801/60/808/5906/408/409/5594/10142/8826/5894/5604 | 16 |
| normal | R-HSA-2990846 | SUMOylation | 36/1399 | 188/10704 | 0.0111462656895763 | 0.040889999337925 | 0.0277089575325588 | 6613/4869/7341/11315/7376/3190/1655/3622/4929/9126/4792/165918/51602/10210/10521/1069/23512/142/3183/6294/7329/9612/54780/23468/6304/10155/53371/4286/5903/6672/4791/2033/2908/23137/1487/3065 | 36 |
| normal | R-HSA-201681 | TCF dependent signaling in response to WNT | 43/1399 | 233/10704 | 0.011368673659339 | 0.041591950901243 | 0.0281846324255112 | 5687/7311/5689/5699/5700/7316/5685/9978/5688/6233/5694/5702/5695/5684/5705/7314/5693/864/5710/9861/207/9580/5714/5686/5708/123169/5713/5929/5701/5527/5525/5718/5690/79577/10213/5515/5717/5719/2033/9736/1487/3065/5707 | 43 |
| normal | R-HSA-445355 | Smooth Muscle Contraction | 11/1399 | 40/10704 | 0.0114053086093586 | 0.0416122839998124 | 0.0281984110749092 | 10627/4637/805/301/103910/309/7170/801/302/808/5058 | 11 |
| normal | R-HSA-1643713 | Signaling by EGFR in Cancer | 8/1399 | 25/10704 | 0.0115631379185551 | 0.0418888939927831 | 0.0283858548184262 | 7311/7316/6233/11140/7314/5290/6464/3320 | 8 |
| normal | R-HSA-983170 | Antigen Presentation: Folding, assembly and peptide loading of class I MHC | 8/1399 | 25/10704 | 0.0115631379185551 | 0.0418888939927831 | 0.0283858548184262 | 3107/10134/3105/3106/2923/567/811/22872 | 8 |
| normal | R-HSA-5654738 | Signaling by FGFR2 | 17/1399 | 73/10704 | 0.0115749744416204 | 0.0418888939927831 | 0.0283858548184262 | 7311/7316/3178/4670/6233/5440/5441/3185/7314/5290/5439/5594/6464/5725/2962/5436/5515 | 17 |
| normal | R-HSA-5357905 | Regulation of TNFR1 signaling | 10/1399 | 35/10704 | 0.0117823837126376 | 0.0425245600841557 | 0.02881661160531 6 | 7311/10399/7316/6233/8887/7314/3551/330/329/81858 | 10 |
| normal | R-HSA-110314 | Recognition of DNA damage by PCNA-containing replication complex | 9/1399 | 30/10704 | 0.0118879025623611 | 0.0425612340401111 | 0.0288414635765699 | 7311/7316/9978/6233/56655/7314/6118/7320/5425 | 9 |
| normal | R-HSA-2424491 | DAP12 signaling | 9/1399 | 30/10704 | 0.0118879025623611 | 0.0425612340401111 | 0.0288414635765699 | 7305/5879/3937/5290/27040/567/6464/2534/6850 | 9 |
| normal | R-HSA-5357956 | TNFR1-induced NFkappaB signaling pathway | 9/1399 | 30/10704 | 0.0118879025623611 | 0.0425612340401111 | 0.0288414635765699 | 7311/10399/7316/6233/7314/3551/330/329/81858 | 9 |
| normal | R-HSA-162599 | Late Phase of HIV Life Cycle | 28/1399 | 139/10704 | 0.0121619730374763 | 0.0429884618810519 | 0.0291309729503296 | 7311/51652/902/7316/5478/6879/5902/5901/6233/51160/128866/5440/5441/7251/27243/7314/2958/25978/1025/9397/5439/51510/11198/53371/6917/5903/2962/5436 | 28 |
| normal | R-HSA-168142 | Toll Like Receptor 10 (TLR10) Cascade | 19/1399 | 85/10704 | 0.0122956975477995 | 0.0429884618810519 | 0.0291309729503296 | 2353/3146/7311/6500/7316/6233/4214/4792/7314/7334/3551/4615/5594/9261/5515/4791/1432/3725/5604 | 19 |
| normal | R-HSA-168176 | Toll Like Receptor 5 (TLR5) Cascade | 19/1399 | 85/10704 | 0.0122956975477995 | 0.0429884618810519 | 0.0291309729503296 | 2353/3146/7311/6500/7316/6233/4214/4792/7314/7334/3551/4615/5594/9261/5515/4791/1432/3725/5604 | 19 |
| normal | R-HSA-975871 | MyD88 cascade initiated on plasma membrane | 19/1399 | 85/10704 | 0.0122956975477995 | 0.0429884618810519 | 0.0291309729503296 | 2353/3146/7311/6500/7316/6233/4214/4792/7314/7334/3551/4615/5594/9261/5515/4791/1432/3725/5604 | 19 |
| normal | R-HSA-110312 | Translesion synthesis by REV1 | 6/1399 | 16/10704 | 0.0122961769234077 | 0.0429884618810519 | 0.0291309729503296 | 7311/7316/6233/7314/6118/5980 | 6 |
| normal | R-HSA-1295596 | Spry regulation of FGF signaling | 6/1399 | 16/10704 | 0.0122961769234077 | 0.0429884618810519 | 0.0291309729503296 | 7311/7316/6233/7314/5594/5515 | 6 |
| normal | R-HSA-205043 | NRIF signals cell death from the nucleus | 6/1399 | 16/10704 | 0.0122961769234077 | 0.0429884618810519 | 0.0291309729503296 | 7311/7316/6233/7314/51107/8878 | 6 |
| normal | R-HSA-399954 | Sema3A PAK dependent Axon repulsion | 6/1399 | 16/10704 | 0.0122961769234077 | 0.0429884618810519 | 0.0291309729503296 | 1072/5879/3326/3320/5058/2534 | 6 |
| normal | R-HSA-8949664 | Processing of SMDT1 | 6/1399 | 16/10704 | 0.0122961769234077 | 0.0429884618810519 | 0.0291309729503296 | 91689/55486/30968/10730/5245/9512 | 6 |
| normal | R-HSA-6811442 | Intra-Golgi and retrograde Golgi-to-ER traffic | 38/1399 | 202/10704 | 0.0123491182374156 | 0.0430611180205717 | 0.029180208115823 2 | 7846/84617/10376/381/140735/4074/11316/830/8655/10959/832/1781/9648/11014/5861/375/11258/3799/10383/54732/6811/400/23423/55860/51272/7110/27333/829/2803/738/8677/9367/10671/10121/1523/372/22931/1314 | 38 |
| normal | R-HSA-5663220 | RHO GTPases Activate Formins | 28/1399 | 140/10704 | 0.0133651035895775 | 0.046482788847907 | 0.0314988907565708 | 7846/5216/84617/10376/140735/5879/998/389/11243/10726/6232/8655/1781/22919/1060/60/752/10383/91010/5527/5525/5903/23122/387/5515/81565/25936/3688 | 28 |
| normal | R-HSA-1474151 | Tetrahydrobiopterin (BH4) synthesis, recycling, salvage and regulation | 5/1399 | 12/10704 | 0.0135487785886231 | 0.0467572539437276 | 0.0316848810184295 | 805/801/207/808/3320 | 5 |
| normal | R-HSA-264870 | Caspase-mediated cleavage of cytoskeletal proteins | 5/1399 | 12/10704 | 0.0135487785886231 | 0.0467572539437276 | 0.0316848810184295 | 7431/2934/6709/118/28988 | 5 |
| normal | R-HSA-9013973 | TICAM1-dependent activation of IRF3/IRF7 | 5/1399 | 12/10704 | 0.0135487785886231 | 0.0467572539437276 | 0.0316848810184295 | 7311/7316/6233/10010/7314 | 5 |
| normal | R-HSA-165159 | MTOR signalling | 11/1399 | 41/10704 | 0.0137941593845879 | 0.0473599472204185 | 0.0320932938988253 | 389541/10542/6194/2280/6009/55004/28956/207/10670/1975/5494 | 11 |
| normal | R-HSA-5696400 | Dual Incision in GG-NER | 11/1399 | 41/10704 | 0.0137941593845879 | 0.0473599472204185 | 0.0320932938988253 | 7311/7316/9978/6233/56655/7314/6118/142/2067/5425/7507 | 11 |
| normal | R-HSA-6807004 | Negative regulation of MET activity | 7/1399 | 21/10704 | 0.0141134282449444 | 0.0483321749871626 | 0.0327521204661249 | 7311/7316/6233/7314/2060/30011/5770 | 7 |
| normal | R-HSA-381070 | IRE1alpha activates chaperones | 13/1399 | 52/10704 | 0.0142934321320594 | 0.0488237388388457 | 0.033085226900739 | 4000/37/5611/27230/118/1200/9114/22872/10897/56005/6464/64061/3068 | 13 |
| normal | R-HSA-73887 | Death Receptor Signalling | 28/1399 | 141/10704 | 0.0146622566215523 | 0.0499561364281386 | 0.0338525919586931 | 7311/10399/7316/5879/57142/6233/7204/8887/4792/7314/396/23365/11214/8743/9181/9459/3551/27018/4615/330/51107/7074/55701/8878/387/329/81858/3065 | 28 |
| tumor | R-HSA-202430 | Translocation of ZAP-70 to Immunological synapse | 4/15 | 19/10704 | 9.55928114247554e-09 | 7.76358358351943e-07 | 5.57542156022546e-07 | 3123/3113/3127/3117 | 4 |
| tumor | R-HSA-202427 | Phosphorylation of CD3 and TCR zeta chains | 4/15 | 22/10704 | 1.79962939677995e-08 | 7.76358358351943e-07 | 5.57542156022546e-07 | 3123/3113/3127/3117 | 4 |
| tumor | R-HSA-389948 | PD-1 signaling | 4/15 | 23/10704 | 2.17670567762227e-08 | 7.76358358351943e-07 | 5.57542156022546e-07 | 3123/3113/3127/3117 | 4 |
| tumor | R-HSA-202433 | Generation of second messenger molecules | 4/15 | 34/10704 | 1.12971726670634e-07 | 3.02199368843947e-06 | 2.1702463281464e-06 | 3123/3113/3127/3117 | 4 |
| tumor | R-HSA-2132295 | MHC class II antigen presentation | 5/15 | 123/10704 | 5.05758141221637e-07 | 1.0823224222143e-05 | 7.77270406509042e-06 | 10484/3123/3113/3127/3117 | 5 |
| tumor | R-HSA-388841 | Costimulation by the CD28 family | 4/15 | 69/10704 | 2.04602119945592e-06 | 3.6487378056964e-05 | 2.62034293965408e-05 | 3123/3113/3127/3117 | 4 |
| tumor | R-HSA-877300 | Interferon gamma signaling | 4/15 | 91/10704 | 6.21159510632818e-06 | 9.49486680538736e-05 | 6.81874350017981e-05 | 3123/3113/3127/3117 | 4 |
| tumor | R-HSA-202424 | Downstream TCR signaling | 4/15 | 98/10704 | 8.3469575063917e-06 | 0.000111640556647989 | 8.01747234166571e-05 | 3123/3113/3127/3117 | 4 |
| tumor | R-HSA-202403 | TCR signaling | 4/15 | 120/10704 | 1.86395326711247e-05 | 0.000221603332867816 | 0.000159144547952293 | 3123/3113/3127/3117 | 4 |
| tumor | R-HSA-913531 | Interferon Signaling | 4/15 | 199/10704 | 0.000134729545499429 | 0.00144160613684389 | 0.00103529019173246 | 3123/3113/3127/3117 | 4 |
| tumor | R-HSA-3371568 | Attenuation phase | 2/15 | 14/10704 | 0.000165191497182412 | 0.00160686274531982 | 0.00115396931046087 | 3304/3303 | 2 |
| tumor | R-HSA-3371571 | HSF1-dependent transactivation | 2/15 | 24/10704 | 0.000496979866653548 | 0.00443140381099414 | 0.00318241493558851 | 3304/3303 | 2 |
| tumor | R-HSA-168274 | Export of Viral Ribonucleoproteins from Nucleus | 2/15 | 33/10704 | 0.000943842818361632 | 0.00776855242805343 | 0.00557898993849386 | 3304/3303 | 2 |
| tumor | R-HSA-168255 | Influenza Infection | 3/15 | 156/10704 | 0.00121475489325743 | 0.00928419811275322 | 0.00666745166975883 | 3304/3303/6173 | 3 |
| tumor | R-HSA-3371497 | HSP90 chaperone cycle for steroid hormone receptors (SHR) | 2/15 | 55/10704 | 0.00260772625138782 | 0.018070443933574 | 0.0129772986438849 | 3304/3303 | 2 |
| tumor | R-HSA-450408 | AUF1 (hnRNP D0) binds and destabilizes mRNA | 2/15 | 56/10704 | 0.00270212245735687 | 0.018070443933574 | 0.0129772986438849 | 3304/3303 | 2 |
| tumor | R-HSA-3371453 | Regulation of HSF1-mediated heat shock response | 2/15 | 69/10704 | 0.00407330826404978 | 0.0256378814266663 | 0.01841185778796 5 | 3304/3303 | 2 |
| tumor | R-HSA-450531 | Regulation of mRNA stability by proteins that bind AU-rich elements | 2/15 | 88/10704 | 0.00654519911088012 | 0.0376767080266153 | 0.0270575473285088 | 3304/3303 | 2 |
| tumor | R-HSA-3371556 | Cellular response to heat stress | 2/15 | 89/10704 | 0.00669025656547375 | 0.0376767080266153 | 0.0270575473285088 | 3304/3303 | 2 |
| tumor | R-HSA-8878159 | Transcriptional regulation by RUNX3 | 2/15 | 96/10704 | 0.00774656855589726 | 0.0414441417740503 | 0.0297631318200263 | 6696/6046 | 2 |

**Supplementary Table 5: Enrichment pathway analysis**

**Neutrophils**

| Cluster | ID | Description | GeneRatio | BgRatio | pvalue | p.adjust | qvalue | geneID | Count |
|---|---|---|---|---|---|---|---|---|---|
| Neutrophil_TAGLN2 | R-HSA-6798695 | Neutrophil degranulation | 17/46 | 480/10704 | 4.75354102939002e-12 | 1.64472519616895e-09 | 1.23091694024205e-09 | 3579/55313/8635/3107/5788/976/2215/2212/3577/4332/5724/5879/5265/2268/11031/5906/3837 | 17 |
| Neutrophil_TAGLN2 | R-HSA-9664323 | FCGR3A-mediated IL10 synthesis | 6/46 | 40/10704 | 1.54523459209549e-08 | 2.6732558443252e-06 | 2.00067215608153e-06 | 2212/4067/3055/2268/805/5573 | 6 |
| Neutrophil_TAGLN2 | R-HSA-2029481 | FCGR activation | 4/46 | 12/10704 | 1.44088892614154e-07 | 1.66182522814991e-05 | 1.24371465203797e-05 | 2212/4067/3055/2268 | 4 |
| Neutrophil_TAGLN2 | R-HSA-909733 | Interferon alpha/beta signaling | 6/46 | 70/10704 | 4.79349315788426e-07 | 4.14637158156988e-05 | 3.10315609694613e-05 | 3107/10410/3133/3437/3659/4600 | 6 |
| Neutrophil_TAGLN2 | R-HSA-913531 | Interferon Signaling | 8/46 | 199/10704 | 1.76518576737545e-06 | 0.000122150855102381 | 9.14180418472337e-05 | 3107/10410/3133/3437/5724/3659/4600/3837 | 8 |
| Neutrophil_TAGLN2 | R-HSA-9662851 | Anti-inflammatory response favouring Leishmania parasite infection | 7/46 | 169/10704 | 6.88284777378926e-06 | 0.000340209332818726 | 0.000254613616895061 | 2790/2212/4067/3055/2268/805/5573 | 7 |
| Neutrophil_TAGLN2 | R-HSA-9664433 | Leishmania parasite growth and survival | 7/46 | 169/10704 | 6.88284777378926e-06 | 0.000340209332818726 | 0.000254613616895061 | 2790/2212/4067/3055/2268/805/5573 | 7 |
| Neutrophil_TAGLN2 | R-HSA-9658195 | Leishmania infection | 8/46 | 252/10704 | 1.01590958344459e-05 | 0.000439380894839783 | 0.000328833891483379 | 2790/2212/4067/3055/5879/2268/805/5573 | 8 |
| Neutrophil_TAGLN2 | R-HSA-2029480 | Fcgamma receptor (FCGR) dependent phagocytosis | 5/46 | 86/10704 | 3.14968401761867e-05 | 0.00121087852232896 | 0.00090622487524467 | 2212/4067/3055/5879/2268 | 5 |
| Neutrophil_TAGLN2 | R-HSA-9664407 | Parasite infection | 4/46 | 59/10704 | 0.000114306324643376 | 0.00309435591625452 | 0.00231582462853244 | 4067/3055/5879/2268 | 4 |
| Neutrophil_TAGLN2 | R-HSA-9664417 | Leishmania phagocytosis | 4/46 | 59/10704 | 0.000114306324643376 | 0.00309435591625452 | 0.00231582462853244 | 4067/3055/5879/2268 | 4 |
| Neutrophil_TAGLN2 | R-HSA-9664422 | FCGR3A-mediated phagocytosis | 4/46 | 59/10704 | 0.000114306324643376 | 0.00309435591625452 | 0.00231582462853244 | 4067/3055/5879/2268 | 4 |
| Neutrophil_TAGLN2 | R-HSA-76002 | Platelet activation, signaling and aggregation | 7/46 | 263/10704 | 0.000117692931099536 | 0.00309435591625452 | 0.00231582462853244 | 8407/2790/4067/5879/5265/805/5906 | 7 |
| Neutrophil_TAGLN2 | R-HSA-449147 | Signaling by Interleukins | 9/46 | 461/10704 | 0.000125205152680819 | 0.00309435591625452 | 0.00231582462853244 | 1441/6648/4067/1439/3055/1052/5724/4170/4478 | 9 |
| Neutrophil_TAGLN2 | R-HSA-877300 | Interferon gamma signaling | 4/46 | 91/10704 | 0.00060723110799957 | 0.0140067975578567 | 0.0104827264959926 | 3107/3133/5724/3659 | 4 |
| Neutrophil_TAGLN2 | R-HSA-381676 | Glucagon-like Peptide-1 (GLP1) regulates insulin secretion | 3/46 | 42/10704 | 0.000758318967248905 | 0.0163986476667576 | 0.0122727938120546 | 2790/5906/5573 | 3 |
| Neutrophil_TAGLN2 | R-HSA-2172127 | DAP12 interactions | 3/46 | 45/10704 | 0.000928924774765388 | 0.0176558114425893 | 0.0132136587005688 | 3107/3133/5879 | 3 |
| Neutrophil_TAGLN2 | R-HSA-1236977 | Endosomal/Vacuolar pathway | 2/46 | 11/10704 | 0.000969538778639296 | 0.0176558114425893 | 0.0132136587005688 | 3107/3133 | 2 |
| Neutrophil_TAGLN2 | R-HSA-8875555 | MET activates RAP1 and RAC1 | 2/46 | 11/10704 | 0.000969538778639296 | 0.0176558114425893 | 0.0132136587005688 | 5879/5906 | 2 |
| Neutrophil_TAGLN2 | R-HSA-512988 | Interleukin-3, Interleukin-5 and GM-CSF signaling | 3/46 | 48/10704 | 0.00112210511643394 | 0.0194124185143071 | 0.0145283083496184 | 4067/1439/3055 | 3 |
| Neutrophil_TAGLN2 | R-HSA-442720 | CREB1 phosphorylation through the activation of Adenylate Cyclase | 2/46 | 14/10704 | 0.00159102765144384 | 0.0262140746380746 | 0.0196186868298338 | 805/5573 | 2 |
| Neutrophil_MMP9 | R-HSA-6798695 | Neutrophil degranulation | 31/89 | 480/10704 | 4.71525729183317e-20 | 2.19730989799426e-17 | 1.91588348910274e-17 | 4318/6283/6279/978/2219/6286/4311/6280/6402/10970/25797/6515/10487/2896/1509/83716/290/5873/25801/8972/64386/5660/3303/240/3689/4069/1265/6556/3482/51411/1604 | 31 |
| Neutrophil_MMP9 | R-HSA-195258 | RHO GTPase Effectors | 17/89 | 327/10704 | 1.19675519228733e-09 | 2.78843959802947e-07 | 2.43130265380478e-07 | 6279/6280/653361/4627/1072/4637/5216/7529/71/10095/801/5880/10627/4689/5579/998/7846 | 17 |
| Neutrophil_MMP9 | R-HSA-5668599 | RHO GTPases Activate NADPH Oxidases | 6/89 | 24/10704 | 3.32521358613635e-08 | 5.16516510379847e-06 | 4.50362261139871e-06 | 6279/6280/653361/5880/4689/5579 | 6 |
| Neutrophil_MMP9 | R-HSA-194315 | Signaling by Rho GTPases | 17/89 | 455/10704 | 1.60497620520096e-07 | 1.86979727905912e-05 | 1.63031793475676e-05 | 6279/6280/653361/4627/1072/4637/5216/7529/71/10095/801/5880/10627/4689/5579/998/7846 | 17 |
| Neutrophil_MMP9 | R-HSA-2682334 | EPH-Ephrin signaling | 8/89 | 92/10704 | 8.74477947264054e-07 | 8.15013446850098e-05 | 7.10628395039842e-05 | 4318/4627/1072/4637/71/10095/10627/998 | 8 |
| Neutrophil_MMP9 | R-HSA-5627123 | RHO GTPases activate PAKs | 5/89 | 24/10704 | 1.33138401019868e-06 | 0.000103404158125431 | 9.0160390866086e-05 | 4627/4637/801/10627/998 | 5 |
| Neutrophil_MMP9 | R-HSA-8950505 | Gene and protein expression by JAK-STAT signaling after Interleukin-12 stimulation | 5/89 | 38/10704 | 1.43459941377092e-05 | 0.000904757340497769 | 0.000788878096752064 | 1072/3936/1265/998/829 | 5 |
| Neutrophil_MMP9 | R-HSA-373755 | Semaphorin interactions | 6/89 | 65/10704 | 1.55323148583308e-05 | 0.000904757340497769 | 0.000788878096752064 | 4627/1072/4637/10627/7094/10154 | 6 |
| Neutrophil_MMP9 | R-HSA-445355 | Smooth Muscle Contraction | 5/89 | 40/10704 | 1.85621570387425e-05 | 0.000910734546689376 | 0.000794089756092385 | 4637/801/10627/7170/7094 | 5 |
| Neutrophil_MMP9 | R-HSA-5627117 | RHO GTPases Activate ROCKs | 4/89 | 20/10704 | 1.95436598002012e-05 | 0.000910734546689376 | 0.000794089756092385 | 4627/1072/4637/10627 | 4 |
| Neutrophil_MMP9 | R-HSA-9020591 | Interleukin-12 signaling | 5/89 | 47/10704 | 4.13367839536608e-05 | 0.00175117648385508 | 0.0015268898187668 | 1072/3936/1265/998/829 | 5 |
| Neutrophil_MMP9 | R-HSA-449147 | Signaling by Interleukins | 13/89 | 461/10704 | 0.00010436390713821 | 0.00378905487456402 | 0.00330376142214075 | 4318/6283/7431/1072/3936/240/3689/7850/8660/1265/23765/998/829 | 13 |
| Neutrophil_MMP9 | R-HSA-447115 | Interleukin-12 family signaling | 5/89 | 57/10704 | 0.000105703247573674 | 0.00378905487456402 | 0.00330376142214075 | 1072/3936/1265/998/829 | 5 |
| Neutrophil_MMP9 | R-HSA-76005 | Response to elevated platelet cytosolic Ca2+ | 7/89 | 134/10704 | 0.000120875989059696 | 0.00402344363584416 | 0.00350813020880019 | 10487/1072/5216/5660/801/5579/7094 | 7 |
| Neutrophil_MMP9 | R-HSA-5626467 | RHO GTPases activate IQGAPs | 4/89 | 32/10704 | 0.000134414096931382 | 0.00417579794466828 | 0.00364097132740446 | 71/801/998/7846 | 4 |
| Neutrophil_MMP9 | R-HSA-2029482 | Regulation of actin dynamics for phagocytic cup formation | 5/89 | 61/10704 | 0.000146317854766624 | 0.00426150752007794 | 0.00371570341709981 | 4627/1072/71/10095/998 | 5 |
| Neutrophil_MMP9 | R-HSA-4420097 | VEGFA-VEGFR2 Pathway | 6/89 | 99/10704 | 0.000168134846115971 | 0.00460887284059074 | 0.00401857898456748 | 653361/71/801/4689/5579/998 | 6 |
| Neutrophil_MMP9 | R-HSA-1222556 | ROS and RNS production in phagocytes | 4/89 | 36/10704 | 0.000214667394794659 | 0.00555750033190617 | 0.00484570844390283 | 653361/5880/4689/6556 | 4 |
| Neutrophil_MMP9 | R-HSA-194138 | Signaling by VEGF | 6/89 | 108/10704 | 0.000270501975965292 | 0.00663441688420138 | 0.00578469599571207 | 653361/71/801/4689/5579/998 | 6 |
| Neutrophil_MMP9 | R-HSA-1445148 | Translocation of SLC2A4 (GLUT4) to the plasma membrane | 5/89 | 72/10704 | 0.000320313987795318 | 0.00721922331876725 | 0.00629460176427413 | 4627/7529/71/801/7846 | 5 |
| Neutrophil_MMP9 | R-HSA-76002 | Platelet activation, signaling and aggregation | 9/89 | 263/10704 | 0.000325329806210541 | 0.00721922331876725 | 0.00629460176427413 | 10487/1072/5216/5660/801/5880/5579/998/7094 | 9 |
| Neutrophil_MMP9 | R-HSA-3928662 | EPHB-mediated forward signaling | 4/89 | 42/10704 | 0.000392699704884255 | 0.00831809374891194 | 0.00725273139164222 | 1072/71/10095/998 | 4 |
| Neutrophil_MMP9 | R-HSA-2022377 | Metabolism of Angiotensinogen to Angiotensins | 3/89 | 18/10704 | 0.000414260189465247 | 0.00835393551396585 | 0.00728398262568515 | 4311/1509/290 | 3 |
| Neutrophil_MMP9 | R-HSA-9656223 | Signaling by RAF1 mutants | 4/89 | 43/10704 | 0.000430245605869486 | 0.00835393551396585 | 0.00728398262568515 | 7529/71/801/7094 | 4 |
| Neutrophil_MMP9 | R-HSA-5625900 | RHO GTPases activate CIT | 3/89 | 20/10704 | 0.000571826096259192 | 0.00974265849548703 | 0.00849484115486333 | 4627/4637/10627 | 3 |
| Neutrophil_MMP9 | R-HSA-6802946 | Signaling by moderate kinase activity BRAF mutants | 4/89 | 47/10704 | 0.000606302781907991 | 0.00974265849548703 | 0.00849484115486333 | 7529/71/801/7094 | 4 |
| Neutrophil_MMP9 | R-HSA-6802949 | Signaling by RAS mutants | 4/89 | 47/10704 | 0.000606302781907991 | 0.00974265849548703 | 0.00849484115486333 | 7529/71/801/7094 | 4 |
| Neutrophil_MMP9 | R-HSA-6802955 | Paradoxical activation of RAF signaling by kinase inactive BRAF | 4/89 | 47/10704 | 0.000606302781907991 | 0.00974265849548703 | 0.00849484115486333 | 7529/71/801/7094 | 4 |
| Neutrophil_MMP9 | R-HSA-9649948 | Signaling downstream of RAS mutants | 4/89 | 47/10704 | 0.000606302781907991 | 0.00974265849548703 | 0.00849484115486333 | 7529/71/801/7094 | 4 |
| Neutrophil_MMP9 | R-HSA-2142691 | Synthesis of Leukotrienes (LT) and Eoxins (EX) | 3/89 | 21/10704 | 0.000663132243482335 | 0.009968375014928 | 0.00869164842051549 | 241/4051/240 | 3 |
| Neutrophil_MMP9 | R-HSA-416572 | Sema4D induced cell migration and growth-cone collapse | 3/89 | 21/10704 | 0.000663132243482335 | 0.009968375014928 | 0.00869164842051549 | 4627/4637/10627 | 3 |
| Neutrophil_MMP9 | R-HSA-114608 | Platelet degranulation | 6/89 | 129/10704 | 0.000699735870754437 | 0.0101899036178615 | 0.00888480414839515 | 10487/1072/5216/5660/801/7094 | 6 |
| Neutrophil_MMP9 | R-HSA-2029480 | Fcgamma receptor (FCGR) dependent phagocytosis | 5/89 | 86/10704 | 0.000727880041686762 | 0.0102785484674555 | 0.00896209556909378 | 4627/1072/71/10095/998 | 5 |
| Neutrophil_MMP9 | R-HSA-3371497 | HSP90 chaperone cycle for steroid hormone receptors (SHR) | 4/89 | 55/10704 | 0.0011021986911979 | 0.0149059876218613 | 0.0129968629366128 | 3303/832/829/7846 | 4 |
| Neutrophil_MMP9 | R-HSA-400685 | Sema4D in semaphorin signaling | 3/89 | 25/10704 | 0.00111954842653465 | 0.0149059876218613 | 0.0129968629366128 | 4627/4637/10627 | 3 |
| Neutrophil_MMP9 | R-HSA-9664407 | Parasite infection | 4/89 | 59/10704 | 0.00143419764345963 | 0.0175877921540049 | 0.015335188099042 | 4627/71/10095/998 | 4 |
| Neutrophil_MMP9 | R-HSA-9664417 | Leishmania phagocytosis | 4/89 | 59/10704 | 0.00143419764345963 | 0.0175877921540049 | 0.015335188099042 | 4627/71/10095/998 | 4 |
| Neutrophil_MMP9 | R-HSA-9664422 | FCGR3A-mediated phagocytosis | 4/89 | 59/10704 | 0.00143419764345963 | 0.0175877921540049 | 0.015335188099042 | 4627/71/10095/998 | 4 |
| Neutrophil_MMP9 | R-HSA-3928663 | EPHA-mediated growth cone collapse | 3/89 | 30/10704 | 0.00191787219359997 | 0.0229161139030151 | 0.0199810706269795 | 4627/4637/10627 | 3 |
| Neutrophil_MMP9 | R-HSA-6802952 | Signaling by BRAF and RAF fusions | 4/89 | 67/10704 | 0.00229662783982854 | 0.0267557143340025 | 0.0233289038466794 | 7529/71/801/7094 | 4 |
| Neutrophil_MMP9 | R-HSA-5663213 | RHO GTPases Activate WASPs and WAVEs | 3/89 | 36/10704 | 0.00325384125050662 | 0.0361021433984782 | 0.0314782637267056 | 71/10095/998 | 3 |
| Neutrophil_MMP9 | R-HSA-6802948 | Signaling by high-kinase activity BRAF mutants | 3/89 | 36/10704 | 0.00325384125050662 | 0.0361021433984782 | 0.0314782637267056 | 7529/71/7094 | 3 |
| Neutrophil_MMP9 | R-HSA-3299685 | Detoxification of Reactive Oxygen Species | 3/89 | 37/10704 | 0.00351984186042146 | 0.0379252558329979 | 0.0330678761046695 | 653361/7295/4689 | 3 |
| Neutrophil_MMP9 | R-HSA-2025928 | Calcineurin activates NFAT | 2/89 | 11/10704 | 0.00358092544345903 | 0.0379252558329979 | 0.0330678761046695 | 2280/801 | 2 |
| Neutrophil_MMP9 | R-HSA-5674135 | MAP2K and MAPK activation | 3/89 | 40/10704 | 0.0043961915373745 | 0.0455250056981449 | 0.0396942674485745 | 7529/71/7094 | 3 |
| Neutrophil_MMP9 | R-HSA-9658195 | Leishmania infection | 7/89 | 252/10704 | 0.00492984278595847 | 0.0498107126328871 | 0.0434310708748463 | 7295/4627/2778/71/10095/801/998 | 7 |
| Neutrophil_MMP9 | R-HSA-879415 | Advanced glycosylation endproduct receptor signaling | 2/89 | 13/10704 | 0.00502382723979763 | 0.0498107126328871 | 0.0434310708748463 | 6283/829 | 2 |
| Neutrophil_CXCL8 | R-HSA-6798695 | Neutrophil degranulation | 17/53 | 480/10704 | 6.55526888810955e-11 | 2.31400991750267e-08 | 2.09768604419506e-08 | 2495/960/25798/1509/1476/5329/3105/6515/963/427/3320/2207/967/1915/54918/7879/3687 | 17 |
| Neutrophil_CXCL8 | R-HSA-6785807 | Interleukin-4 and Interleukin-13 signaling | 8/53 | 108/10704 | 5.02858112987348e-08 | 8.87544569422669e-06 | 8.04572980779756e-06 | 3576/1026/3553/5008/3726/3320/2353/3687 | 8 |
| Neutrophil_CXCL8 | R-HSA-449147 | Signaling by Interleukins | 12/53 | 461/10704 | 1.88144070926019e-06 | 0.000221382856789615 | 0.000200687008987753 | 3576/3557/1026/4792/8878/6648/3553/5008/3726/3320/2353/3687 | 12 |
| Neutrophil_CXCL8 | R-HSA-2871796 | FCERI mediated MAPK activation | 3/53 | 32/10704 | 0.000513690906260674 | 0.0453332224775044 | 0.0410952725008539 | 3937/2207/2353 | 3 |

**Supplementary Table 5: Enrichment pathway analysis**

**Macrophages**

| Cluster | ID | Description | GeneRatio | BgRatio | pvalue | p.adjust | qvalue | geneID | Count |
|---|---|---|---|---|---|---|---|---|---|
| Mac_Alv-like | R-HSA-6798695 | Neutrophil degranulation | 21/81 | 480/10704 | 3.5077340032768e-11 | 1.90820729778258e-08 | 1.49909474245303e-08 | 199675/5265/3958/56729/527/6282/2495/4048/1535/4257/6279/6280/2171/4973/1675/3240/9535/2896/7130/2512/3107 | 21 |
| Mac_Alv-like | R-HSA-3000480 | Scavenging by Class A Receptors | 4/81 | 19/10704 | 1.08186036082461e-05 | 0.00271304863163917 | 0.00213138108445338 | 8685/2495/4481/2512 | 4 |
| Mac_Alv-like | R-HSA-2173782 | Binding and Uptake of Ligands by Scavenger Receptors | 5/81 | 42/10704 | 1.49616652480101e-05 | 0.00271304863163917 | 0.00213138108445338 | 8685/2495/4481/3240/2512 | 5 |
| Mac_Alv-like | R-HSA-114608 | Platelet degranulation | 7/81 | 129/10704 | 5.20179732986713e-05 | 0.0070744443686193 | 0.00555770977875278 | 5265/6813/7114/8407/710/928/1675 | 7 |
| Mac_Alv-like | R-HSA-76005 | Response to elevated platelet cytosolic Ca2+ | 7/81 | 134/10704 | 6.6273466903188e-05 | 0.00721055319906686 | 0.00566463738161986 | 5265/6813/7114/8407/710/928/1675 | 7 |
| Mac_Alv-like | R-HSA-9018678 | Biosynthesis of specialized proresolving mediators (SPMs) | 3/81 | 19/10704 | 0.000370682755669359 | 0.0336085698473553 | 0.0264030173336421 | 241/4048/2879 | 3 |
| Mac_Rec | R-HSA-156902 | Peptide chain elongation | 27/80 | 89/10704 | 9.85361904366498e-38 | 1.34501899946027e-35 | 1.04240917251403e-35 | 6203/6202/6165/6229/6187/6137/6141/6210/6135/6201/6204/6164/6217/23521/6130/6134/6132/6167/6173/6189/6188/6194/6232/6222/6142/6191/6122 | 27 |
| Mac_Rec | R-HSA-192823 | Viral mRNA Translation | 27/80 | 89/10704 | 9.85361904366498e-38 | 1.34501899946027e-35 | 1.04240917251403e-35 | 6203/6202/6165/6229/6187/6137/6141/6210/6135/6201/6204/6164/6217/23521/6130/6134/6132/6167/6173/6189/6188/6194/6232/6222/6142/6191/6122 | 27 |
| Mac_Rec | R-HSA-156842 | Eukaryotic Translation Elongation | 27/80 | 93/10704 | 3.91554740927724e-37 | 2.13788888546538e-35 | 1.65689479845205e-35 | 6203/6202/6165/6229/6187/6137/6141/6210/6135/6201/6204/6164/6217/23521/6130/6134/6132/6167/6173/6189/6188/6194/6232/6222/6142/6191/6122 | 27 |
| Mac_Rec | R-HSA-2408557 | Selenocysteine synthesis | 27/80 | 93/10704 | 3.91554740927724e-37 | 2.13788888546538e-35 | 1.65689479845205e-35 | 6203/6202/6165/6229/6187/6137/6141/6210/6135/6201/6204/6164/6217/23521/6130/6134/6132/6167/6173/6189/6188/6194/6232/6222/6142/6191/6122 | 27 |
| Mac_Rec | R-HSA-72764 | Eukaryotic Translation Termination | 27/80 | 93/10704 | 3.91554740927724e-37 | 2.13788888546538e-35 | 1.65689479845205e-35 | 6203/6202/6165/6229/6187/6137/6141/6210/6135/6201/6204/6164/6217/23521/6130/6134/6132/6167/6173/6189/6188/6194/6232/6222/6142/6191/6122 | 27 |
| Mac_Rec | R-HSA-975956 | Nonsense Mediated Decay (NMD) independent of the Exon Junction Complex (EJC) | 27/80 | 95/10704 | 7.60097131130803e-37 | 3.45844194664515e-35 | 2.6803425150402e-35 | 6203/6202/6165/6229/6187/6137/6141/6210/6135/6201/6204/6164/6217/23521/6130/6134/6132/6167/6173/6189/6188/6194/6232/6222/6142/6191/6122 | 27 |
| Mac_Rec | R-HSA-72689 | Formation of a pool of free 40S subunits | 27/80 | 101/10704 | 5.05154789112517e-36 | 1.72384071784646e-34 | 1.33600148173179e-34 | 6203/6202/6165/6229/6187/6137/6141/6210/6135/6201/6204/6164/6217/23521/6130/6134/6132/6167/6173/6189/6188/6194/6232/6222/6142/6191/6122 | 27 |
| Mac_Rec | R-HSA-9633012 | Response of EIF2AK4 (GCN2) to amino acid deficiency | 27/80 | 101/10704 | 5.05154789112517e-36 | 1.72384071784646e-34 | 1.33600148173179e-34 | 6203/6202/6165/6229/6187/6137/6141/6210/6135/6201/6204/6164/6217/23521/6130/6134/6132/6167/6173/6189/6188/6194/6232/6222/6142/6191/6122 | 27 |
| Mac_Rec | R-HSA-156827 | L13a-mediated translational silencing of Ceruloplasmin expression | 27/80 | 111/10704 | 8.98523715295751e-35 | 2.72552193639711e-33 | 2.11231890964264e-33 | 6203/6202/6165/6229/6187/6137/6141/6210/6135/6201/6204/6164/6217/23521/6130/6134/6132/6167/6173/6189/6188/6194/6232/6222/6142/6191/6122 | 27 |
| Mac_Rec | R-HSA-1799339 | SRP-dependent cotranslational protein targeting to membrane | 27/80 | 112/10704 | 1.17822968700021e-34 | 2.92415185955506e-33 | 2.2662599721248e-33 | 6203/6202/6165/6229/6187/6137/6141/6210/6135/6201/6204/6164/6217/23521/6130/6134/6132/6167/6173/6189/6188/6194/6232/6222/6142/6191/6122 | 27 |
| Mac_Rec | R-HSA-72706 | GTP hydrolysis and joining of the 60S ribosomal subunit | 27/80 | 112/10704 | 1.17822968700021e-34 | 2.92415185955506e-33 | 2.2662599721248e-33 | 6203/6202/6165/6229/6187/6137/6141/6210/6135/6201/6204/6164/6217/23521/6130/6134/6132/6167/6173/6189/6188/6194/6232/6222/6142/6191/6122 | 27 |
| Mac_Rec | R-HSA-927802 | Nonsense-Mediated Decay (NMD) | 27/80 | 115/10704 | 2.61284478876152e-34 | 5.48697405639919e-33 | 4.25248423110174e-33 | 6203/6202/6165/6229/6187/6137/6141/6210/6135/6201/6204/6164/6217/23521/6130/6134/6132/6167/6173/6189/6188/6194/6232/6222/6142/6191/6122 | 27 |
| Mac_Rec | R-HSA-975957 | Nonsense Mediated Decay (NMD) enhanced by the Exon Junction Complex (EJC) | 27/80 | 115/10704 | 2.61284478876152e-34 | 5.48697405639919e-33 | 4.25248423110174e-33 | 6203/6202/6165/6229/6187/6137/6141/6210/6135/6201/6204/6164/6217/23521/6130/6134/6132/6167/6173/6189/6188/6194/6232/6222/6142/6191/6122 | 27 |
| Mac_Rec | R-HSA-2408522 | Selenoamino acid metabolism | 27/80 | 118/10704 | 5.65805090948131e-34 | 1.10331992734886e-32 | 8.55088896846423e-33 | 6203/6202/6165/6229/6187/6137/6141/6210/6135/6201/6204/6164/6217/23521/6130/6134/6132/6167/6173/6189/6188/6194/6232/6222/6142/6191/6122 | 27 |
| Mac_Rec | R-HSA-72613 | Eukaryotic Translation Initiation | 27/80 | 119/10704 | 7.28326409814278e-34 | 1.24270693674561e-32 | 9.63115844557039e-33 | 6203/6202/6165/6229/6187/6137/6141/6210/6135/6201/6204/6164/6217/23521/6130/6134/6132/6167/6173/6189/6188/6194/6232/6222/6142/6191/6122 | 27 |
| Mac_Rec | R-HSA-72737 | Cap-dependent Translation Initiation | 27/80 | 119/10704 | 7.28326409814278e-34 | 1.24270693674561e-32 | 9.63115844557039e-33 | 6203/6202/6165/6229/6187/6137/6141/6210/6135/6201/6204/6164/6217/23521/6130/6134/6132/6167/6173/6189/6188/6194/6232/6222/6142/6191/6122 | 27 |
| Mac_Rec | R-HSA-168273 | Influenza Viral RNA Transcription and Replication | 27/80 | 135/10704 | 3.05481925710238e-32 | 4.90568033640559e-31 | 3.80197319305002e-31 | 6203/6202/6165/6229/6187/6137/6141/6210/6135/6201/6204/6164/6217/23521/6130/6134/6132/6167/6173/6189/6188/6194/6232/6222/6142/6191/6122 | 27 |
| Mac_Rec | R-HSA-168255 | Influenza Infection | 28/80 | 156/10704 | 4.73971650492053e-32 | 7.1885700324628e-31 | 5.57124571631009e-31 | 6203/6202/6165/6229/6187/6137/293/6141/6210/6135/6201/6204/6164/6217/23521/6130/6134/6132/6167/6173/6189/6188/6194/6232/6222/6142/6191/6122 | 28 |
| Mac_Rec | R-HSA-9010553 | Regulation of expression of SLITs and ROBOs | 27/80 | 171/10704 | 2.82421865403426e-29 | 4.05795627658607e-28 | 3.14497478925699e-28 | 6203/6202/6165/6229/6187/6137/6141/6210/6135/6201/6204/6164/6217/23521/6130/6134/6132/6167/6173/6189/6188/6194/6232/6222/6142/6191/6122 | 27 |
| Mac_Rec | R-HSA-6791226 | Major pathway of rRNA processing in the nucleolus and cytosol | 27/80 | 184/10704 | 2.25272510806872e-28 | 3.0749697725138e-27 | 2.38314603537796e-27 | 6203/6202/6165/6229/6187/6137/6141/6210/6135/6201/6204/6164/6217/23521/6130/6134/6132/6167/6173/6189/6188/6194/6232/6222/6142/6191/6122 | 27 |
| Mac_Rec | R-HSA-376176 | Signaling by ROBO receptors | 28/80 | 218/10704 | 8.90808991646366e-28 | 1.15805168914028e-26 | 8.97506803613632e-27 | 7852/6203/6202/6165/6229/6187/6137/6141/6210/6135/6201/6204/6164/6217/23521/6130/6134/6132/6167/6173/6189/6188/6194/6232/6222/6142/6191/6122 | 28 |
| Mac_Rec | R-HSA-8868773 | rRNA processing in the nucleus and cytosol | 27/80 | 194/10704 | 9.98437449133099e-28 | 1.23897010733335e-26 | 9.60219747730875e-27 | 6203/6202/6165/6229/6187/6137/6141/6210/6135/6201/6204/6164/6217/23521/6130/6134/6132/6167/6173/6189/6188/6194/6232/6222/6142/6191/6122 | 27 |
| Mac_Rec | R-HSA-72312 | rRNA processing | 27/80 | 204/10704 | 4.07255919555057e-27 | 4.83395069732741e-26 | 3.74638168560945e-26 | 6203/6202/6165/6229/6187/6137/6141/6210/6135/6201/6204/6164/6217/23521/6130/6134/6132/6167/6173/6189/6188/6194/6232/6222/6142/6191/6122 | 27 |
| Mac_Rec | R-HSA-72766 | Translation | 28/80 | 291/10704 | 3.2637201989123e-24 | 3.71248172626274e-23 | 2.87722701746216e-23 | 5464/6203/6202/6165/6229/6187/6137/6141/6210/6135/6201/6204/6164/6217/23521/6130/6134/6132/6167/6173/6189/6188/6194/6232/6222/6142/6191/6122 | 28 |
| Mac_Rec | R-HSA-71291 | Metabolism of amino acids and derivatives | 27/80 | 374/10704 | 5.24179778078538e-20 | 5.72404317661764e-19 | 4.43621622710679e-19 | 6203/6202/6165/6229/6187/6137/6141/6210/6135/6201/6204/6164/6217/23521/6130/6134/6132/6167/6173/6189/6188/6194/6232/6222/6142/6191/6122 | 27 |
| Mac_Rec | R-HSA-72695 | Formation of the ternary complex, and subsequently, the 43S complex | 14/80 | 51/10704 | 5.33869610806326e-19 | 5.60563091346642e-18 | 4.34444501101504e-18 | 6203/6202/6229/6187/6210/6201/6204/6217/6189/6188/6194/6232/6222/6191 | 14 |
| Mac_Rec | R-HSA-72649 | Translation initiation complex formation | 14/80 | 58/10704 | 4.02241806076563e-18 | 3.92185760924648e-17 | 3.03949635418756e-17 | 6203/6202/6229/6187/6210/6201/6204/6217/6189/6188/6194/6232/6222/6191 | 14 |
| Mac_Rec | R-HSA-72702 | Ribosomal scanning and start codon recognition | 14/80 | 58/10704 | 4.02241806076563e-18 | 3.92185760924648e-17 | 3.03949635418756e-17 | 6203/6202/6229/6187/6210/6201/6204/6217/6189/6188/6194/6232/6222/6191 | 14 |
| Mac_Rec | R-HSA-72662 | Activation of the mRNA upon binding of the cap-binding complex and eIFs, and subsequent binding to 4 | 14/80 | 59/10704 | 5.24336486015774e-18 | 4.9359951959416e-17 | 3.82546764751981e-17 | 6203/6202/6229/6187/6210/6201/6204/6217/6189/6188/6194/6232/6222/6191 | 14 |
| Mac_Rec | R-HSA-179812 | GRB2 events in EGFR signaling | 3/80 | 13/10704 | 0.000108940844598015 | 0.000991361685841937 | 0.000768319640849159 | 374/2069/1839 | 3 |
| Mac_Rec | R-HSA-180336 | SHC1 events in EGFR signaling | 3/80 | 14/10704 | 0.00013790588910444 | 0.00121446153953265 | 0.00094122525331044 | 374/2069/1839 | 3 |
| Mac_Rec | R-HSA-6785807 | Interleukin-4 and Interleukin-13 signaling | 6/80 | 108/10704 | 0.000150513212500818 | 0.0012840658441476 | 0.000995169595811328 | 7076/9021/1052/7422/3726/7133 | 6 |
| Mac_Rec | R-HSA-180292 | GAB1 signalosome | 3/80 | 17/10704 | 0.000253492233366487 | 0.00209707211239548 | 0.00162526120914398 | 374/2069/1839 | 3 |
| Mac_Rec | R-HSA-9634638 | Estrogen-dependent nuclear events downstream of ESR-membrane signaling | 3/80 | 24/10704 | 0.000726603243985329 | 0.00583419663552926 | 0.00452158675049694 | 374/2069/1839 | 3 |
| Mac_Rec | R-HSA-1643713 | Signaling by EGFR in Cancer | 3/80 | 25/10704 | 0.000821256924697794 | 0.00640580401264279 | 0.00496459073275959 | 374/2069/1839 | 3 |
| Mac_Rec | R-HSA-380108 | Chemokine receptors bind chemokines | 4/80 | 59/10704 | 0.000963816859449372 | 0.00730894451749107 | 0.00566453768272877 | 7852/4283/6373/3627 | 4 |
| Mac_Rec | R-HSA-6790901 | rRNA modification in the nucleus and cytosol | 4/80 | 61/10704 | 0.00109272316596946 | 0.00806252498134223 | 0.00624857343840288 | 6203/6187/6201/6194 | 4 |
| Mac_Rec | R-HSA-182971 | EGFR downregulation | 3/80 | 31/10704 | 0.00155411104591476 | 0.0111650609351244 | 0.00865308366284948 | 374/2069/1839 | 3 |
| Mac_Rec | R-HSA-909733 | Interferon alpha/beta signaling | 4/80 | 70/10704 | 0.0018249072815138 | 0.0127743509705966 | 0.00990030670942709 | 9021/10581/3669/10410 | 4 |
| Mac_Rec | R-HSA-1250342 | PI3K events in ERBB4 signaling | 2/80 | 10/10704 | 0.00238774219953213 | 0.0162963405118068 | 0.0126298995291042 | 2069/1839 | 2 |
| Mac_Rec | R-HSA-8847993 | ERBB2 Activates PTK6 Signaling | 2/80 | 13/10704 | 0.00407900867560045 | 0.0271602284985103 | 0.0210495698022 | 2069/1839 | 2 |
| Mac_Rec | R-HSA-1250347 | SHC1 events in ERBB4 signaling | 2/80 | 14/10704 | 0.00473585914001644 | 0.0307830844101069 | 0.0238573355173761 | 2069/1839 | 2 |
| Mac_Rec | R-HSA-6783783 | Interleukin-10 signaling | 3/80 | 47/10704 | 0.00514594542190751 | 0.0326707697716454 | 0.0253203189670357 | 7076/7133/3627 | 3 |
| Mac_Rec | R-HSA-6785631 | ERBB2 Regulates Cell Motility | 2/80 | 15/10704 | 0.00543807515640384 | 0.033740784493142 | 0.0261495958477792 | 2069/1839 | 2 |
| Mac_Rec | R-HSA-177929 | Signaling by EGFR | 3/80 | 50/10704 | 0.00612137897222305 | 0.0359254014138982 | 0.0278427055492329 | 374/2069/1839 | 3 |
| Mac_Rec | R-HSA-1963640 | GRB2 events in ERBB2 signaling | 2/80 | 16/10704 | 0.00618495921777735 | 0.0359254014138982 | 0.0278427055492329 | 2069/1839 | 2 |
| Mac_Rec | R-HSA-1963642 | PI3K events in ERBB2 signaling | 2/80 | 16/10704 | 0.00618495921777735 | 0.0359254014138982 | 0.0278427055492329 | 2069/1839 | 2 |
| Mac_Rec | R-HSA-9022699 | MECP2 regulates neuronal receptors and channels | 2/80 | 17/10704 | 0.00697582148236546 | 0.0396749846809535 | 0.0307486867972688 | 6515/2289 | 2 |
| Mac_Rec | R-HSA-8848021 | Signaling by PTK6 | 3/80 | 54/10704 | 0.0075831890934854 | 0.0414042124504303 | 0.0320888633219067 | 2069/9021/1839 | 3 |
| Mac_Rec | R-HSA-9006927 | Signaling by Non-Receptor Tyrosine Kinases | 3/80 | 54/10704 | 0.0075831890934854 | 0.0414042124504303 | 0.0320888633219067 | 2069/9021/1839 | 3 |
| Mac_Rec | R-HSA-5686938 | Regulation of TLR by endogenous ligand | 2/80 | 19/10704 | 0.00868675914109907 | 0.0464997106964715 | 0.0360379481395441 | 6279/6280 | 2 |
| Mac_Rec | R-HSA-6798695 | Neutrophil degranulation | 9/80 | 480/10704 | 0.00933642267696047 | 0.0490162190540425 | 0.0379882785034222 | 2219/10875/23601/6515/4069/23406/7133/6279/6280 | 9 |
| Mac_Reg | R-HSA-202430 | Translocation of ZAP-70 to Immunological synapse | 7/74 | 19/10704 | 2.66267672238378e-11 | 1.44050810680962e-08 | 1.26126792112916e-08 | 3117/3119/3115/3113/3123/3122/3127 | 7 |
| Mac_Reg | R-HSA-202427 | Phosphorylation of CD3 and TCR zeta chains | 7/74 | 22/10704 | 8.86478359592746e-11 | 2.28541717928968e-08 | 2.00104627041611e-08 | 3117/3119/3115/3113/3123/3122/3127 | 7 |
| Mac_Reg | R-HSA-389948 | PD-1 signaling | 7/74 | 23/10704 | 1.26732930459687e-10 | 2.28541717928968e-08 | 2.00104627041611e-08 | 3117/3119/3115/3113/3123/3122/3127 | 7 |
| Mac_Reg | R-HSA-202433 | Generation of second messenger molecules | 7/74 | 34/10704 | 2.61785636779759e-09 | 3.54065073744624e-07 | 3.10009306712872e-07 | 3117/3119/3115/3113/3123/3122/3127 | 7 |
| Mac_Reg | R-HSA-2132295 | MHC class II antigen presentation | 10/74 | 123/10704 | 1.07234914014155e-08 | 1.16028176963316e-06 | 1.01590971171305e-06 | 3117/3119/3109/3115/3113/3111/3123/972/3122/3127 | 10 |
| Mac_Reg | R-HSA-202424 | Downstream TCR signaling | 9/74 | 98/10704 | 2.10580143676505e-08 | 1.89873096214982e-06 | 1.66247481849872e-06 | 3117/3119/3115/3113/3635/3123/3122/7316/3127 | 9 |
| Mac_Reg | R-HSA-202403 | TCR signaling | 9/74 | 120/10704 | 1.23980165733622e-07 | 9.58189566598425e-06 | 8.38963527520753e-06 | 3117/3119/3115/3113/3635/3123/3122/7316/3127 | 9 |
| Mac_Reg | R-HSA-877300 | Interferon gamma signaling | 8/74 | 91/10704 | 1.90297899412495e-07 | 1.286889544777e-05 | 1.1267638781003e-05 | 3117/3119/3115/3113/2209/3123/3122/3127 | 8 |
| Mac_Reg | R-HSA-388841 | Costimulation by the CD28 family | 7/74 | 69/10704 | 4.33177274437671e-07 | 2.60387672745311e-05 | 2.27988039177721e-05 | 3117/3119/3115/3113/3123/3122/3127 | 7 |
| Mac_Reg | R-HSA-913531 | Interferon Signaling | 10/74 | 199/10704 | 1.00482600073e-06 | 5.43610866394932e-05 | 4.75970210872107e-05 | 3117/3119/1958/3115/3113/2209/3123/3122/7316/3127 | 10 |
| Mac_Reg | R-HSA-198725 | Nuclear Events (kinase and transcription factor activation) | 6/74 | 61/10704 | 3.65212602172949e-06 | 0.000179618197977787 | 0.000157268584667777 | 4208/6446/1958/2353/1959/3726 | 6 |
| Mac_Reg | R-HSA-9031628 | NGF-stimulated transcription | 5/74 | 39/10704 | 6.60218149118686e-06 | 0.000297648348894341 | 0.000260612427283692 | 6446/1958/2353/1959/3726 | 5 |
| Mac_Reg | R-HSA-198933 | Immunoregulatory interactions between a Lymphoid and a non-Lymphoid cell | 7/74 | 132/10704 | 3.34806282589984e-05 | 0.00139330922216293 | 0.00121994192036836 | 718/1277/1278/54209/11314/2209/2213 | 7 |
| Mac_Reg | R-HSA-187037 | Signaling by NTRK1 (TRKA) | 6/74 | 115/10704 | 0.000137918729914488 | 0.00532957377740985 | 0.00466642319259545 | 4208/6446/1958/2353/1959/3726 | 6 |
| Mac_Reg | R-HSA-166520 | Signaling by NTRKs | 6/74 | 134/10704 | 0.00031731191952755 | 0.0114443832309603 | 0.0100203764061332 | 4208/6446/1958/2353/1959/3726 | 6 |
| Mac_Reg | R-HSA-202733 | Cell surface interactions at the vascular wall | 6/74 | 137/10704 | 0.000357434734415044 | 0.0120857619574087 | 0.0105819493741296 | 1277/1278/3635/972/962/6404 | 6 |
| Mac_Reg | R-HSA-975138 | TRAF6 mediated induction of NFkB and MAP kinases upon TLR7/8 or 9 activation | 5/74 | 91/10704 | 0.000403492933232867 | 0.0123717459043809 | 0.0108323487828999 | 4208/6285/2353/929/7316 | 5 |
| Mac_Reg | R-HSA-975155 | MyD88 dependent cascade initiated on endosome | 5/74 | 92/10704 | 0.000424405614891681 | 0.0123717459043809 | 0.0108323487828999 | 4208/6285/2353/929/7316 | 5 |
| Mac_Reg | R-HSA-168181 | Toll Like Receptor 7/8 (TLR7/8) Cascade | 5/74 | 93/10704 | 0.000446126716486636 | 0.0123717459043809 | 0.0108323487828999 | 4208/6285/2353/929/7316 | 5 |
| Mac_Reg | R-HSA-166058 | MyD88:MAL(TIRAP) cascade initiated on plasma membrane | 5/74 | 95/10704 | 0.000492075236212121 | 0.0123717459043809 | 0.0108323487828999 | 4208/6285/2353/929/7316 | 5 |
| Mac_Reg | R-HSA-168188 | Toll Like Receptor TLR6:TLR2 Cascade | 5/74 | 95/10704 | 0.000492075236212121 | 0.0123717459043809 | 0.0108323487828999 | 4208/6285/2353/929/7316 | 5 |
| Mac_Reg | R-HSA-168138 | Toll Like Receptor 9 (TLR9) Cascade | 5/74 | 96/10704 | 0.000516343663574603 | 0.0123717459043809 | 0.0108323487828999 | 4208/6285/2353/929/7316 | 5 |
| Mac_Reg | R-HSA-168179 | Toll Like Receptor TLR1:TLR2 Cascade | 5/74 | 98/10704 | 0.000567572781039322 | 0.0123717459043809 | 0.0108323487828999 | 4208/6285/2353/929/7316 | 5 |
| Mac_Reg | R-HSA-181438 | Toll Like Receptor 2 (TLR2) Cascade | 5/74 | 98/10704 | 0.000567572781039322 | 0.0123717459043809 | 0.0108323487828999 | 4208/6285/2353/929/7316 | 5 |
| Mac_Reg | R-HSA-166166 | MyD88-independent TLR4 cascade | 5/74 | 99/10704 | 0.000594575588750285 | 0.0123717459043809 | 0.0108323487828999 | 4208/6285/2353/929/7316 | 5 |
| Mac_Reg | R-HSA-937061 | TRIF(TICAM1)-mediated TLR4 signaling | 5/74 | 99/10704 | 0.000594575588750285 | 0.0123717459043809 | 0.0108323487828999 | 4208/6285/2353/929/7316 | 5 |
| Mac_Reg | R-HSA-2029485 | Role of phospholipids in phagocytosis | 3/74 | 25/10704 | 0.000653982880521354 | 0.0131038791985945 | 0.0114733838687957 | 122618/2209/2212 | 3 |
| Mac_Reg | R-HSA-380108 | Chemokine receptors bind chemokines | 4/74 | 59/10704 | 0.000718613651879879 | 0.0138846423452505 | 0.0121569978701483 | 1524/7852/2920/6387 | 4 |
| Mac_Reg | R-HSA-449147 | Signaling by Interleukins | 10/74 | 461/10704 | 0.00120408744848767 | 0.0224624589528217 | 0.0196674900841906 | 1278/4208/1436/2920/3635/6285/2353/3726/7124/7316 | 10 |
| Mac_Reg | R-HSA-418594 | G alpha (i) signalling events | 9/74 | 405/10704 | 0.00182001275257482 | 0.0328208966380993 | 0.0287370434617077 | 5996/718/1524/1880/7852/2920/6001/6387/9249 | 9 |
| Mac_Reg | R-HSA-166016 | Toll Like Receptor 4 (TLR4) Cascade | 5/74 | 129/10704 | 0.00194946637254462 | 0.0340213325015044 | 0.0297881109556902 | 4208/6285/2353/929/7316 | 5 |
| Mac_Reg | R-HSA-9664873 | Pexophagy | 2/74 | 11/10704 | 0.00249075041482623 | 0.042109249200656 | 0.0368696607457831 | 4077/7316 | 2 |
| Mac_Reg | R-HSA-168142 | Toll Like Receptor 10 (TLR10) Cascade | 4/74 | 85/10704 | 0.00279320461440204 | 0.0427226245732406 | 0.0374067147737295 | 4208/6285/2353/7316 | 4 |
| Mac_Reg | R-HSA-168176 | Toll Like Receptor 5 (TLR5) Cascade | 4/74 | 85/10704 | 0.00279320461440204 | 0.0427226245732406 | 0.0374067147737295 | 4208/6285/2353/7316 | 4 |
| Mac_Reg | R-HSA-975871 | MyD88 cascade initiated on plasma membrane | 4/74 | 85/10704 | 0.00279320461440204 | 0.0427226245732406 | 0.0374067147737295 | 4208/6285/2353/7316 | 4 |
| Mac_Reg | R-HSA-2029481 | FCGR activation | 2/74 | 12/10704 | 0.0029755546909195 | 0.0427226245732406 | 0.0374067147737295 | 2209/2212 | 2 |
| Mac_Reg | R-HSA-430116 | GP1b-IX-V activation signalling | 2/74 | 12/10704 | 0.0029755546909195 | 0.0427226245732406 | 0.0374067147737295 | 1277/1278 | 2 |
| Mac_Reg | R-HSA-2173782 | Binding and Uptake of Ligands by Scavenger Receptors | 3/74 | 42/10704 | 0.00300084978518141 | 0.0427226245732406 | 0.0374067147737295 | 1277/1278/23166 | 3 |
| RTM_IM | R-HSA-6785807 | Interleukin-4 and Interleukin-13 signaling | 11/80 | 108/10704 | 3.85479597816328e-10 | 1.13716481355817e-07 | 1.01441999425349e-07 | 2162/5743/3576/3383/1052/7124/6347/3553/3726/5008/3162 | 11 |
| RTM_IM | R-HSA-6783783 | Interleukin-10 signaling | 8/80 | 47/10704 | 1.69198217210217e-09 | 2.49567370385071e-07 | 2.22629233171339e-07 | 5743/3576/3383/7124/2920/2919/6347/3553 | 8 |
| RTM_IM | R-HSA-449147 | Signaling by Interleukins | 18/80 | 461/10704 | 5.59535208297784e-09 | 5.50209621492821e-07 | 4.90820358155951e-07 | 2162/3559/5743/3576/10148/3383/1052/7124/2920/2919/6347/3553/3726/23291/5008/200081/4792/3162 | 18 |
| RTM_IM | R-HSA-380108 | Chemokine receptors bind chemokines | 7/80 | 59/10704 | 2.48987340654507e-07 | 1.83628163732699e-05 | 1.63807460956913e-05 | 6387/6357/3576/2920/2919/6347/2921 | 7 |
| RTM_IM | R-HSA-9031628 | NGF-stimulated transcription | 4/80 | 39/10704 | 0.000195041665169792 | 0.0115074582450177 | 0.0102653507984101 | 1958/2354/1959/3726 | 4 |
| RTM_IM | R-HSA-375276 | Peptide ligand-binding receptors | 7/80 | 201/10704 | 0.000739487550288425 | 0.0363581378891809 | 0.0324336644863344 | 6387/6357/3576/2920/2919/6347/2921 | 7 |
| RTM_IM | R-HSA-380994 | ATF4 activates genes in response to endoplasmic reticulum stress | 3/80 | 27/10704 | 0.00103325606108879 | 0.0402941667451238 | 0.0359448409858375 | 3576/6347/467 | 3 |
| RTM_IM | R-HSA-198725 | Nuclear Events (kinase and transcription factor activation) | 4/80 | 61/10704 | 0.00109272316596946 | 0.0402941667451238 | 0.0359448409858375 | 1958/2354/1959/3726 | 4 |
| RTM_IM | R-HSA-5357956 | TNFR1-induced NFkappaB signaling pathway | 3/80 | 30/10704 | 0.00141126884610367 | 0.0462582566222869 | 0.0412651709387038 | 7124/330/7128 | 3 |
| RTM_LA | R-HSA-6798695 | Neutrophil degranulation | 32/85 | 480/10704 | 7.18322058582317e-22 | 3.8142901310721e-19 | 3.44038459636794e-19 | 1522/2517/2760/8804/3417/966/1118/5912/2799/3916/427/1675/7077/1509/1075/3074/1520/4069/3920/2512/2896/6386/1508/3903/3689/5660/4860/3304/3303/84061/3312/5611 | 32 |
| RTM_LA | R-HSA-1679131 | Trafficking and processing of endosomal TLR | 5/85 | 13/10704 | 3.43257297771476e-08 | 9.11348125583269e-06 | 8.2201089729485e-06 | 5641/1514/1520/1508/7184 | 5 |
| RTM_LA | R-HSA-9029569 | NR1H3 & NR1H2 regulate gene expression linked to cholesterol transport and efflux | 5/85 | 37/10704 | 1.00093256741493e-05 | 0.00173998382678631 | 0.00156941746691996 | 348/5360/341/19/10123 | 5 |
| RTM_LA | R-HSA-8963898 | Plasma lipoprotein assembly | 4/85 | 19/10704 | 1.31072228006502e-05 | 0.00173998382678631 | 0.00156941746691996 | 348/341/19/2 | 4 |
| RTM_LA | R-HSA-174824 | Plasma lipoprotein assembly, remodeling, and clearance | 6/85 | 70/10704 | 1.83219224436914e-05 | 0.00194578816352003 | 0.00175504730776412 | 348/3988/5360/341/19/2 | 6 |
| RTM_LA | R-HSA-9613829 | Chaperone Mediated Autophagy | 4/85 | 22/10704 | 2.42912389723067e-05 | 0.00214977464904914 | 0.0019390374969122 | 6035/3920/3312/7316 | 4 |
| RTM_LA | R-HSA-9024446 | NR1H2 and NR1H3-mediated signaling | 5/85 | 47/10704 | 3.30968526970397e-05 | 0.00251063268316116 | 0.00226452150032377 | 348/5360/341/19/10123 | 5 |
| RTM_LA | R-HSA-2132295 | MHC class II antigen presentation | 7/85 | 123/10704 | 5.23888152104157e-05 | 0.00347730760959134 | 0.00313643564746568 | 5641/1514/1509/1075/3109/1520/1508 | 7 |
| RTM_LA | R-HSA-1236977 | Endosomal/Vacuolar pathway | 3/85 | 11/10704 | 7.61623690248168e-05 | 0.00449357977246419 | 0.00405308513523879 | 1514/1520/3133 | 3 |
| RTM_LA | R-HSA-917937 | Iron uptake and transport | 5/85 | 58/10704 | 9.23279390204789e-05 | 0.00490261356198743 | 0.00442202234255978 | 3162/7037/2512/526/7316 | 5 |
| RTM_LA | R-HSA-3371568 | Attenuation phase | 3/85 | 14/10704 | 0.000165146616844664 | 0.00797207759495607 | 0.00719059432194471 | 3304/3303/3312 | 3 |
| RTM_LA | R-HSA-168898 | Toll-like Receptor Cascades | 7/85 | 155/10704 | 0.000223574007270961 | 0.00989314982174001 | 0.00892334853581466 | 5641/1514/1520/1508/3689/7184/7316 | 7 |
| RTM_LA | R-HSA-199992 | trans-Golgi Network Vesicle Budding | 5/85 | 72/10704 | 0.000258447762777985 | 0.0105565970796239 | 0.0095217596812942 | 1522/2799/7037/2512/3312 | 5 |
| RTM_LA | R-HSA-2173782 | Binding and Uptake of Ligands by Scavenger Receptors | 4/85 | 42/10704 | 0.00032937025845151 | 0.0124925433741251 | 0.0112679298943938 | 348/9332/2512/7184 | 4 |
| RTM_LA | R-HSA-3000480 | Scavenging by Class A Receptors | 3/85 | 19/10704 | 0.000427196488781892 | 0.015122755702879 | 0.0136403089400534 | 348/2512/7184 | 3 |
| RTM_LA | R-HSA-1660662 | Glycosphingolipid metabolism | 4/85 | 46/10704 | 0.000468751543951872 | 0.0155566918649027 | 0.0140317074011909 | 2760/427/3074/5660 | 4 |
| RTM_LA | R-HSA-114608 | Platelet degranulation | 6/85 | 129/10704 | 0.000547663308680042 | 0.0171064245240648 | 0.0154295235572396 | 2/1675/3920/334/5660/3309 | 6 |
| RTM_LA | R-HSA-76005 | Response to elevated platelet cytosolic Ca2+ | 6/85 | 134/10704 | 0.000669677363295991 | 0.0197554822172317 | 0.0178189006023202 | 2/1675/3920/334/5660/3309 | 6 |
| RTM_LA | R-HSA-1474228 | Degradation of the extracellular matrix | 6/85 | 140/10704 | 0.000842831502476682 | 0.0223705078332624 | 0.0201775816515699 | 1514/2/7077/1509/1520/1508 | 6 |
| RTM_LA | R-HSA-3371571 | HSF1-dependent transactivation | 3/85 | 24/10704 | 0.000867101821240161 | 0.0223705078332624 | 0.0201775816515699 | 3304/3303/3312 | 3 |
| RTM_LA | R-HSA-381119 | Unfolded Protein Response (UPR) | 5/85 | 94/10704 | 0.000884709349338063 | 0.0223705078332624 | 0.0201775816515699 | 1200/9709/7184/5611/3309 | 5 |
| RTM_LA | R-HSA-450408 | AUF1 (hnRNP D0) binds and destabilizes mRNA | 4/85 | 56/10704 | 0.000993388850124828 | 0.0239767860983108 | 0.0216264003860272 | 3304/3303/3312/7316 | 4 |
| RTM_LA | R-HSA-8856828 | Clathrin-mediated endocytosis | 6/85 | 146/10704 | 0.00104870943383951 | 0.0242115091029905 | 0.0218381140685116 | 1601/920/7037/950/3312/7316 | 6 |
| RTM_LA | R-HSA-8856825 | Cargo recognition for clathrin-mediated endocytosis | 5/85 | 106/10704 | 0.00151655523759985 | 0.0335537846318967 | 0.0302645891714005 | 1601/920/7037/950/7316 | 5 |
| RTM_LA | R-HSA-3371453 | Regulation of HSF1-mediated heat shock response | 4/85 | 69/10704 | 0.00216249901600437 | 0.0436632665972933 | 0.0393830633398125 | 3304/3303/3312/3309 | 4 |
| RTM_LA | R-HSA-5694530 | Cargo concentration in the ER | 3/85 | 33/10704 | 0.00222016609816745 | 0.0436632665972933 | 0.0393830633398125 | 1522/966/1075 | 3 |
| RTM_LA | R-HSA-8964043 | Plasma lipoprotein clearance | 3/85 | 33/10704 | 0.00222016609816745 | 0.0436632665972933 | 0.0393830633398125 | 348/3988/341 | 3 |
| RTM_LA | R-HSA-9006931 | Signaling by Nuclear Receptors | 8/85 | 299/10704 | 0.00251831103322957 | 0.0446508855808396 | 0.0402738684493647 | 348/5360/341/5166/19/10123/216/1509 | 8 |
| RTM_LA | R-HSA-432720 | Lysosome Vesicle Biogenesis | 3/85 | 35/10704 | 0.0026330764622155 | 0.0446508855808396 | 0.0402738684493647 | 1522/2799/3312 | 3 |
| RTM_LA | R-HSA-9615710 | Late endosomal microautophagy | 3/85 | 35/10704 | 0.0026330764622155 | 0.0446508855808396 | 0.0402738684493647 | 6035/3312/7316 | 3 |
| RTM_LA | R-HSA-381183 | ATF6 (ATF6-alpha) activates chaperone genes | 2/85 | 10/10704 | 0.00269082549639711 | 0.0446508855808396 | 0.0402738684493647 | 7184/3309 | 2 |
| RTM_LA | R-HSA-8964058 | HDL remodeling | 2/85 | 10/10704 | 0.00269082549639711 | 0.0446508855808396 | 0.0402738684493647 | 348/5360 | 2 |
| Mac_Angio | R-HSA-6783783 | Interleukin-10 signaling | 4/32 | 47/10704 | 1.07219207951727e-05 | 0.00188705805995039 | 0.00160264500306792 | 7076/3557/6347/2920 | 4 |
| Mac_Prolif | R-HSA-168255 | Influenza Infection | 17/78 | 156/10704 | 7.29710351363707e-16 | 4.20313162385495e-13 | 2.54246448738302e-13 | 3838/142/3921/6187/5901/6130/6188/6194/6175/6129/6191/6125/3320/6124/6122/6173/6139 | 17 |
| Mac_Prolif | R-HSA-156842 | Eukaryotic Translation Elongation | 14/78 | 93/10704 | 3.30465184525824e-15 | 9.51739731434374e-13 | 5.75705137252883e-13 | 3921/6187/6130/6188/6194/1933/6175/6129/6191/6125/6124/6122/6173/6139 | 14 |
| Mac_Prolif | R-HSA-168273 | Influenza Viral RNA Transcription and Replication | 15/78 | 135/10704 | 3.2800474250565e-14 | 6.01053050706681e-12 | 3.63575584400423e-12 | 142/3921/6187/6130/6188/6194/6175/6129/6191/6125/3320/6124/6122/6173/6139 | 15 |
| Mac_Prolif | R-HSA-156902 | Peptide chain elongation | 13/78 | 89/10704 | 5.21747439849549e-14 | 6.01053050706681e-12 | 3.63575584400423e-12 | 3921/6187/6130/6188/6194/6175/6129/6191/6125/6124/6122/6173/6139 | 13 |
| Mac_Prolif | R-HSA-192823 | Viral mRNA Translation | 13/78 | 89/10704 | 5.21747439849549e-14 | 6.01053050706681e-12 | 3.63575584400423e-12 | 3921/6187/6130/6188/6194/6175/6129/6191/6125/6124/6122/6173/6139 | 13 |
| Mac_Prolif | R-HSA-2408557 | Selenocysteine synthesis | 13/78 | 93/10704 | 9.41528555610786e-14 | 7.74743497188304e-12 | 4.68640529183715e-12 | 3921/6187/6130/6188/6194/6175/6129/6191/6125/6124/6122/6173/6139 | 13 |
| Mac_Prolif | R-HSA-72764 | Eukaryotic Translation Termination | 13/78 | 93/10704 | 9.41528555610786e-14 | 7.74743497188304e-12 | 4.68640529183715e-12 | 3921/6187/6130/6188/6194/6175/6129/6191/6125/6124/6122/6173/6139 | 13 |
| Mac_Prolif | R-HSA-975956 | Nonsense Mediated Decay (NMD) independent of the Exon Junction Complex (EJC) | 13/78 | 95/10704 | 1.25154276859892e-13 | 9.01110793391221e-12 | 5.45079811060845e-12 | 3921/6187/6130/6188/6194/6175/6129/6191/6125/6124/6122/6173/6139 | 13 |
| Mac_Prolif | R-HSA-72689 | Formation of a pool of free 40S subunits | 13/78 | 101/10704 | 2.82859073571642e-13 | 1.62926826377266e-11 | 9.85540561602247e-12 | 3921/6187/6130/6188/6194/6175/6129/6191/6125/6124/6122/6173/6139 | 13 |
| Mac_Prolif | R-HSA-9633012 | Response of EIF2AK4 (GCN2) to amino acid deficiency | 13/78 | 101/10704 | 2.82859073571642e-13 | 1.62926826377266e-11 | 9.85540561602247e-12 | 3921/6187/6130/6188/6194/6175/6129/6191/6125/6124/6122/6173/6139 | 13 |
| Mac_Prolif | R-HSA-156827 | L13a-mediated translational silencing of Ceruloplasmin expression | 13/78 | 111/10704 | 9.83335858996656e-13 | 4.55745664630419e-11 | 2.7567948646321e-11 | 3921/6187/6130/6188/6194/6175/6129/6191/6125/6124/6122/6173/6139 | 13 |
| Mac_Prolif | R-HSA-1799339 | SRP-dependent cotranslational protein targeting to membrane | 13/78 | 112/10704 | 1.10614504127549e-12 | 4.55745664630419e-11 | 2.7567948646321e-11 | 3921/6187/6130/6188/6194/6175/6129/6191/6125/6124/6122/6173/6139 | 13 |
| Mac_Prolif | R-HSA-72706 | GTP hydrolysis and joining of the 60S ribosomal subunit | 13/78 | 112/10704 | 1.10614504127549e-12 | 4.55745664630419e-11 | 2.7567948646321e-11 | 3921/6187/6130/6188/6194/6175/6129/6191/6125/6124/6122/6173/6139 | 13 |
| Mac_Prolif | R-HSA-9010553 | Regulation of expression of SLITs and ROBOs | 15/78 | 171/10704 | 1.10771515708782e-12 | 4.55745664630419e-11 | 2.7567948646321e-11 | 3921/6187/6130/6188/6194/5698/5685/6175/6129/6191/6125/6124/6122/6173/6139 | 15 |

| | | | | | | | | | |
|---|---|---|---|---|---|---|---|---|---|
| Mac_Prolif | R-HSA-927802 | Nonsense-Mediated Decay (NMD) | 13/78 | 115/10704 | 1.5636982536372e-12 | 5.62931371309391e-11 | 3.40515869706521e-11 | 3921/6187/6130/6188/6194/6175/6129/6191/6125/6124/6122/6173/6139 | 13 |
| Mac_Prolif | R-HSA-975957 | Nonsense Mediated Decay (NMD) enhanced by the Exon Junction Complex (EJC) | 13/78 | 115/10704 | 1.5636982536372e-12 | 5.62931371309391e-11 | 3.40515869706521e-11 | 3921/6187/6130/6188/6194/6175/6129/6191/6125/6124/6122/6173/6139 | 13 |
| Mac_Prolif | R-HSA-2408522 | Selenoamino acid metabolism | 13/78 | 118/10704 | 2.18876424393645e-12 | 7.40688099698466e-11 | 4.4804049890386e-11 | 3921/6187/6130/6188/6194/6175/6129/6191/6125/6124/6122/6173/6139 | 13 |
| Mac_Prolif | R-HSA-72613 | Eukaryotic Translation Initiation | 13/78 | 119/10704 | 2.44324199553314e-12 | 7.40688099698466e-11 | 4.4804049890386e-11 | 3921/6187/6130/6188/6194/6175/6129/6191/6125/6124/6122/6173/6139 | 13 |
| Mac_Prolif | R-HSA-72737 | Cap-dependent Translation Initiation | 13/78 | 119/10704 | 2.44324199553314e-12 | 7.40688099698466e-11 | 4.4804049890386e-11 | 3921/6187/6130/6188/6194/6175/6129/6191/6125/6124/6122/6173/6139 | 13 |
| Mac_Prolif | R-HSA-68886 | M Phase | 20/78 | 418/10704 | 1.06312581210859e-11 | 3.06180233887273e-10 | 1.85207707267338e-10 | 11065/9232/983/1063/891/10051/332/11130/991/7112/203068/5901/10376/84790/80097/6418/7531/5698/5685/3320 | 20 |
| Mac_Prolif | R-HSA-376176 | Signaling by ROBO receptors | 15/78 | 218/10704 | 3.73659569374878e-11 | 1.02489481885681e-09 | 6.19956478511703e-10 | 3921/6187/6130/6188/6194/5698/5685/6175/6129/6191/6125/6124/6122/6173/6139 | 15 |
| Mac_Prolif | R-HSA-6791226 | Major pathway of rRNA processing in the nucleolus and cytosol | 13/78 | 184/10704 | 6.22557367199517e-10 | 1.62996837957692e-08 | 9.85964060014547e-09 | 3921/6187/6130/6188/6194/6175/6129/6191/6125/6124/6122/6173/6139 | 13 |
| Mac_Prolif | R-HSA-8868773 | rRNA processing in the nucleus and cytosol | 13/78 | 194/10704 | 1.19692975649046e-09 | 2.99752843364568e-08 | 1.81319793774985e-08 | 3921/6187/6130/6188/6194/6175/6129/6191/6125/6124/6122/6173/6139 | 13 |
| Mac_Prolif | R-HSA-68882 | Mitotic Anaphase | 14/78 | 236/10704 | 1.27738287332934e-09 | 3.06571889599042e-08 | 1.85444618891233e-08 | 11065/9232/983/1063/891/332/11130/991/7112/5901/10376/84790/5698/5685 | 14 |
| Mac_Prolif | R-HSA-2555396 | Mitotic Metaphase and Anaphase | 14/78 | 237/10704 | 1.34988881504461e-09 | 3.11014382986277e-08 | 1.88131872749375e-08 | 11065/9232/983/1063/891/332/11130/991/7112/5901/10376/84790/5698/5685 | 14 |
| Mac_Prolif | R-HSA-68877 | Mitotic Prometaphase | 13/78 | 204/10704 | 2.21783330870374e-09 | 4.73137772523464e-08 | 2.8619993184442e-08 | 983/1063/891/10051/332/11130/991/203068/10376/84790/80097/7531/3320 | 13 |
| Mac_Prolif | R-HSA-72312 | rRNA processing | 13/78 | 204/10704 | 2.21783330870374e-09 | 4.73137772523464e-08 | 2.8619993184442e-08 | 3921/6187/6130/6188/6194/6175/6129/6191/6125/6124/6122/6173/6139 | 13 |
| Mac_Prolif | R-HSA-453279 | Mitotic G1 phase and G1/S transition | 11/78 | 149/10704 | 9.17685768520814e-09 | 1.88781072381425e-07 | 1.14193229090372e-07 | 7153/1163/983/7298/7083/891/6241/5111/4176/5698/5685 | 11 |
| Mac_Prolif | R-HSA-72766 | Translation | 14/78 | 291/10704 | 1.89220756253884e-08 | 3.75831571042197e-07 | 2.27339638185247e-07 | 3921/6187/6130/6188/6194/1933/6175/6129/6191/6125/6124/6122/6173/6139 | 14 |
| Mac_Prolif | R-HSA-75035 | Chk1/Chk2(Cds1) mediated inactivation of Cyclin B:Cdk1 complex | 5/78 | 13/10704 | 2.21891357042744e-08 | 4.26031405522068e-07 | 2.57705400635608e-07 | 983/891/7533/7531/7534 | 5 |
| Mac_Prolif | R-HSA-69206 | G1/S Transition | 10/78 | 131/10704 | 3.31621001246774e-08 | 6.16173215219812e-07 | 3.7272173654561e-07 | 1163/983/7298/7083/891/6241/5111/4176/5698/5685 | 10 |
| Mac_Prolif | R-HSA-71291 | Metabolism of amino acids and derivatives | 15/78 | 374/10704 | 6.26490819367882e-08 | 1.12768347486219e-06 | 6.82133096088056e-07 | 3921/6187/6130/6188/6194/5698/5685/6175/6129/6191/6125/6124/6122/6173/6139 | 15 |
| Mac_Prolif | R-HSA-69275 | G2/M Transition | 11/78 | 196/10704 | 1.56380409410606e-07 | 2.72954896425786e-06 | 1.65109778356972e-06 | 983/1063/891/203068/10376/84790/80097/7531/5698/5685/3320 | 11 |
| Mac_Prolif | R-HSA-69620 | Cell Cycle Checkpoints | 13/78 | 294/10704 | 1.72868664598665e-07 | 2.85279804053207e-06 | 1.72565086150606e-06 | 11065/983/1063/891/332/11130/4176/991/7533/7531/5698/5685/7534 | 13 |
| Mac_Prolif | R-HSA-453274 | Mitotic G2-G2/M phases | 11/78 | 198/10704 | 1.73347103157331e-07 | 2.85279804053207e-06 | 1.72565086150606e-06 | 983/1063/891/203068/10376/84790/80097/7531/5698/5685/3320 | 11 |
| Mac_Prolif | R-HSA-1445148 | Translocation of SLC2A4 (GLUT4) to the plasma membrane | 7/78 | 72/10704 | 8.36088544245214e-07 | 1.33774167079234e-05 | 8.09196807442005e-06 | 10376/84790/7533/808/7531/805/7534 | 7 |
| Mac_Prolif | R-HSA-2467813 | Separation of Sister Chromatids | 10/78 | 191/10704 | 1.13580335895574e-06 | 1.76816955340137e-05 | 1.06956162678336e-05 | 11065/9232/1063/332/11130/991/10376/84790/5698/5685 | 10 |
| Mac_Prolif | R-HSA-176409 | APC/C:Cdc20 mediated degradation of mitotic proteins | 7/78 | 76/10704 | 1.21226017378633e-06 | 1.83753121079191e-05 | 1.1115183310894e-05 | 11065/9232/983/891/991/5698/5685 | 7 |
| Mac_Prolif | R-HSA-176814 | Activation of APC/C and APC/C:Cdc20 mediated degradation of mitotic proteins | 7/78 | 77/10704 | 1.32574914542335e-06 | 1.90907876940963e-05 | 1.15479728193455e-05 | 11065/9232/983/891/991/5698/5685 | 7 |
| Mac_Prolif | R-HSA-8852276 | The role of GTSE1 in G2/M progression after G2 checkpoint | 7/78 | 77/10704 | 1.32574914542335e-06 | 1.90907876940963e-05 | 1.15479728193455e-05 | 983/891/10376/84790/5698/5685/3320 | 7 |
| Mac_Prolif | R-HSA-69205 | G1/S-Specific Transcription | 5/78 | 28/10704 | 1.55570999246156e-06 | 2.18558281867771e-05 | 1.32205393454372e-05 | 983/7298/7083/6241/5111 | 5 |
| Mac_Prolif | R-HSA-174143 | APC/C-mediated degradation of cell cycle proteins | 7/78 | 88/10704 | 3.28280596100355e-06 | 4.39743310125127e-05 | 2.65999699655368e-05 | 11065/9232/983/891/991/5698/5685 | 7 |
| Mac_Prolif | R-HSA-453276 | Regulation of mitotic cell cycle | 7/78 | 88/10704 | 3.28280596100355e-06 | 4.39743310125127e-05 | 2.65999699655368e-05 | 11065/9232/983/891/991/5698/5685 | 7 |
| Mac_Prolif | R-HSA-2500257 | Resolution of Sister Chromatid Cohesion | 8/78 | 126/10704 | 3.46907260740407e-06 | 4.54133141332897e-05 | 2.74704074892523e-05 | 983/1063/891/332/11130/991/10376/84790 | 8 |
| Mac_Prolif | R-HSA-380320 | Recruitment of NuMA to mitotic centrosomes | 7/78 | 94/10704 | 5.1115329989857e-06 | 6.5427622387017e-05 | 3.95770157348367e-05 | 983/203068/10376/84790/80097/7531/3320 | 7 |
| Mac_Prolif | R-HSA-2995410 | Nuclear Envelope (NE) Reassembly | 6/78 | 75/10704 | 1.66419486494963e-05 | 0.000208386139611084 | 0.000126052288397787 | 983/891/7112/5901/10376/84790 | 6 |
| Mac_Prolif | R-HSA-195258 | RHO GTPase Effectors | 11/78 | 327/10704 | 2.28347596473788e-05 | 0.000279847267167877 | 0.000169278957296358 | 1063/332/11130/991/10376/84790/7533/808/7531/805/7534 | 11 |
| Mac_Prolif | R-HSA-174048 | APC/C:Cdc20 mediated degradation of Cyclin B | 4/78 | 24/10704 | 2.48182304349097e-05 | 0.000297818765218917 | 0.00018014987442884 | 11065/983/891/991 | 4 |
| Mac_Prolif | R-HSA-176408 | Regulation of APC/C activators between G1/S and early anaphase | 6/78 | 81/10704 | 2.59103861241002e-05 | 0.00030457923280575 | 0.000184239265458156 | 11065/983/891/991/5698/5685 | 6 |
| Mac_Prolif | R-HSA-69481 | G2/M Checkpoints | 8/78 | 168/10704 | 2.87234913242905e-05 | 0.000330894620055827 | 0.000200157381649266 | 983/891/4176/7533/7531/5698/5685/7534 | 8 |
| Mac_Prolif | R-HSA-72695 | Formation of the ternary complex, and subsequently, the 43S complex | 5/78 | 51/10704 | 3.26265041749152e-05 | 0.000368487576563748 | 0.000222897273104168 | 3921/6187/6188/6194/6191 | 5 |
| Mac_Prolif | R-HSA-162909 | Host Interactions of HIV factors | 7/78 | 130/10704 | 4.27892957738409e-05 | 0.000467952041298422 | 0.000283063095156757 | 5902/5901/292/8905/5698/5685/5478 | 7 |
| Mac_Prolif | R-HSA-442729 | CREB1 phosphorylation through the activation of CaMKII/CaMKK/CaMKIV cascasde | 3/78 | 10/10704 | 4.30580871333617e-05 | 0.000467952041298422 | 0.000283063095156757 | 3838/808/805 | 3 |
| Mac_Prolif | R-HSA-9619483 | Activation of AMPK downstream of NMDARs | 4/78 | 29/10704 | 5.39606732892817e-05 | 0.000575580515085671 | 0.000348167307188153 | 10376/84790/808/805 | 4 |
| Mac_Prolif | R-HSA-2025928 | Calcineurin activates NFAT | 3/78 | 11/10704 | 5.88942666842175e-05 | 0.000604499305267875 | 0.000365660215723075 | 808/5478/805 | 3 |
| Mac_Prolif | R-HSA-2514853 | Condensation of Prometaphase Chromosomes | 3/78 | 11/10704 | 5.88942666842175e-05 | 0.000604499305267875 | 0.000365660215723075 | 983/891/10051 | 3 |
| Mac_Prolif | R-HSA-72649 | Translation initiation complex formation | 5/78 | 58/10704 | 6.11622999084295e-05 | 0.000604499305267875 | 0.000365660215723075 | 3921/6187/6188/6194/6191 | 5 |
| Mac_Prolif | R-HSA-72702 | Ribosomal scanning and start codon recognition | 5/78 | 58/10704 | 6.11622999084295e-05 | 0.000604499305267875 | 0.000365660215723075 | 3921/6187/6188/6194/6191 | 5 |
| Mac_Prolif | R-HSA-499943 | Interconversion of nucleotide di- and triphosphates | 4/78 | 30/10704 | 6.19191996715358e-05 | 0.000604499305267875 | 0.000365660215723075 | 7298/6241/1841/1854 | 4 |
| Mac_Prolif | R-HSA-72662 | Activation of the mRNA upon binding of the cap-binding complex and eIFs, and subsequent binding to | 5/78 | 59/10704 | 6.64473221108269e-05 | 0.000637894292263938 | 0.000385860765240065 | 3921/6187/6188/6194/6191 | 5 |
| Mac_Prolif | R-HSA-5357801 | Programmed Cell Death | 8/78 | 190/10704 | 6.89675455000945e-05 | 0.000651234528000892 | 0.000393930242632119 | 3148/3146/7533/7531/5698/5685/3320/7534 | 8 |
| Mac_Prolif | R-HSA-111932 | CaMK IV-mediated phosphorylation of CREB | 3/78 | 12/10704 | 7.81138402284969e-05 | 0.000714183682089115 | 0.000432008038690601 | 3838/808/805 | 3 |
| Mac_Prolif | R-HSA-1474151 | Tetrahydrobiopterin (BH4) synthesis, recycling, salvage and regulation | 3/78 | 12/10704 | 7.81138402284969e-05 | 0.000714183682089115 | 0.000432008038690601 | 808/3320/805 | 3 |
| Mac_Prolif | R-HSA-5626467 | RHO GTPases activate IQGAPs | 4/78 | 32/10704 | 8.03556704727884e-05 | 0.000723201034255096 | 0.000437462613922582 | 10376/84790/808/805 | 4 |
| Mac_Prolif | R-HSA-203615 | eNOS activation | 3/78 | 13/10704 | 0.000101015547559543 | 0.000881131475797597 | 0.000532994368583707 | 808/3320/805 | 3 |
| Mac_Prolif | R-HSA-194315 | Signaling by Rho GTPases | 12/78 | 455/10704 | 0.000101722599516707 | 0.000881131475797597 | 0.000532994368583707 | 1063/332/11130/991/10376/84790/7533/808/7531/397/805/7534 | 12 |
| Mac_Prolif | R-HSA-70263 | Gluconeogenesis | 4/78 | 34/10704 | 0.00010249272374729 | 0.000881131475797597 | 0.000532994368583707 | 5223/7167/2023/2597 | 4 |
| Mac_Prolif | R-HSA-174154 | APC/C:Cdc20 mediated degradation of Securin | 5/78 | 68/10704 | 0.00013146876387155 | 0.00111361776455901 | 0.000673624780827911 | 11065/9232/991/5698/5685 | 5 |
| Mac_Prolif | R-HSA-111447 | Activation of BAD and translocation to mitochondria | 3/78 | 15/10704 | 0.000159026424125391 | 0.00130856028994607 | 0.000791545058428638 | 7533/7531/7534 | 3 |
| Mac_Prolif | R-HSA-4419969 | Depolymerisation of the Nuclear Lamina | 3/78 | 15/10704 | 0.000159026424125391 | 0.00130856028994607 | 0.000791545058428638 | 983/891/7112 | 3 |
| Mac_Prolif | R-HSA-70171 | Glycolysis | 5/78 | 71/10704 | 0.000161437749161864 | 0.00130969216221456 | 0.000792229725316188 | 5223/7167/5315/2023/2597 | 5 |
| Mac_Prolif | R-HSA-174184 | Cdc20:Phospho-APC/C mediated degradation of Cyclin A | 5/78 | 73/10704 | 0.000184142558966931 | 0.00147314047173545 | 0.000891099225410151 | 11065/983/991/5698/5685 | 5 |
| Mac_Prolif | R-HSA-174178 | APC/C:Cdh1 mediated degradation of Cdc20 and other APC/C:Cdh1 targeted proteins in late mitosis/ear | 5/78 | 74/10704 | 0.000196370341864095 | 0.00152850428261782 | 0.000924588665106904 | 11065/9232/991/5698/5685 | 5 |
| Mac_Prolif | R-HSA-179419 | APC:Cdc20 mediated degradation of cell cycle proteins prior to satisfation of the cell cycle checkpoint | 5/78 | 74/10704 | 0.000196370341864095 | 0.00152850428261782 | 0.000924588665106904 | 11065/983/991/5698/5685 | 5 |
| Mac_Prolif | R-HSA-9648025 | EML4 and NUDC in mitotic spindle formation | 6/78 | 117/10704 | 0.000202936335228278 | 0.00155855105455317 | 0.000942763887165754 | 1063/332/11130/991/10376/84790 | 6 |
| Mac_Prolif | R-HSA-202131 | Metabolism of nitric oxide: NOS3 activation and regulation | 3/78 | 17/10704 | 0.000235182563516411 | 0.00178243627086122 | 0.00107819153080238 | 808/3320/805 | 3 |
| Mac_Prolif | R-HSA-6804114 | TP53 Regulates Transcription of Genes Involved in G2 Cell Cycle Arrest | 3/78 | 18/10704 | 0.000280741425159423 | 0.00208807646750106 | 0.00126307257080199 | 983/891/5111 | 3 |
| Mac_Prolif | R-HSA-438064 | Post NMDA receptor activation events | 5/78 | 80/10704 | 0.000283261962118904 | 0.00208807646750106 | 0.00126307257080199 | 3838/10376/84790/808/805 | 5 |
| Mac_Prolif | R-HSA-1852241 | Organelle biogenesis and maintenance | 9/78 | 296/10704 | 0.000286385487773018 | 0.00208807646750106 | 0.00126307257080199 | 983/3418/203068/10376/84790/808/7531/3320/805 | 9 |
| Mac_Prolif | R-HSA-109581 | Apoptosis | 7/78 | 178/10704 | 0.000305545752928485 | 0.00219992942108509 | 0.00133073216078064 | 3148/3146/7533/7531/5698/5685/7534 | 7 |
| Mac_Prolif | R-HSA-380270 | Recruitment of mitotic centrosome proteins and complexes | 5/78 | 82/10704 | 0.000317881153292327 | 0.00223292127190708 | 0.00135068885416894 | 983/203068/80097/7531/3320 | 5 |
| Mac_Prolif | R-HSA-380287 | Centrosome maturation | 5/78 | 82/10704 | 0.000317881153292327 | 0.00223292127190708 | 0.00135068885416894 | 983/203068/80097/7531/3320 | 5 |
| Mac_Prolif | R-HSA-2995383 | Initiation of Nuclear Envelope (NE) Reformation | 3/78 | 19/10704 | 0.000331635418796496 | 0.00227407144317597 | 0.00137558049651178 | 983/891/7112 | 3 |
| Mac_Prolif | R-HSA-8876725 | Protein methylation | 3/78 | 19/10704 | 0.000331635418796496 | 0.00227407144317597 | 0.00137558049651178 | 6187/808/805 | 3 |
| Mac_Prolif | R-HSA-1168372 | Downstream signaling events of B Cell Receptor (BCR) | 5/78 | 83/10704 | 0.000336348608784961 | 0.00227925645482515 | 0.00137871689792968 | 808/5698/5685/5478/805 | 5 |
| Mac_Prolif | R-HSA-176412 | Phosphorylation of the APC/C | 3/78 | 20/10704 | 0.000388117679124593 | 0.00256960670316972 | 0.00155434908397145 | 11065/983/891 | 3 |
| Mac_Prolif | R-HSA-8978934 | Metabolism of cofactors | 3/78 | 20/10704 | 0.000388117679124593 | 0.00256960670316972 | 0.00155434908397145 | 808/3320/805 | 3 |
| Mac_Prolif | R-HSA-2565942 | Regulation of PLK1 Activity at G2/M Transition | 5/78 | 88/10704 | 0.000441168873112974 | 0.00288765080583037 | 0.00174673321770807 | 983/891/203068/7531/3320 | 5 |
| Mac_Prolif | R-HSA-72172 | mRNA Splicing | 7/78 | 191/10704 | 0.000467930103467159 | 0.00302840156850656 | 0.00183187302480934 | 79622/25804/6628/220988/3178/6428/3181 | 7 |
| Mac_Prolif | R-HSA-70326 | Glucose metabolism | 5/78 | 91/10704 | 0.00051488663927673 | 0.00329527449137107 | 0.00199330383158594 | 5223/7167/5315/2023/2597 | 5 |
| Mac_Prolif | R-HSA-5663220 | RHO GTPases Activate Formins | 6/78 | 140/10704 | 0.000533330215063729 | 0.00337580443820558 | 0.00204201620805199 | 1063/332/11130/991/10376/84790 | 6 |
| Mac_Prolif | R-HSA-442755 | Activation of NMDA receptors and postsynaptic events | 5/78 | 93/10704 | 0.000568942165592374 | 0.00356207268892617 | 0.00215468943719766 | 3838/10376/84790/808/805 | 5 |
| Mac_Prolif | R-HSA-69473 | G2/M DNA damage checkpoint | 5/78 | 95/10704 | 0.000627156261403 | 0.00388432265127019 | 0.00234961768561848 | 983/891/7533/7531/7534 | 5 |
| Mac_Prolif | R-HSA-15869 | Metabolism of nucleotides | 5/78 | 101/10704 | 0.000828779432738216 | 0.00507847822614056 | 0.00307195959951119 | 7298/7083/6241/1841/1854 | 5 |
| Mac_Prolif | R-HSA-1538133 | G0 and Early G1 | 3/78 | 27/10704 | 0.000959959730152566 | 0.00582038741650398 | 0.00352073873330193 | 7153/983/5111 | 3 |
| Mac_Prolif | R-HSA-69242 | S Phase | 6/78 | 162/10704 | 0.00114534659490553 | 0.00687207956943316 | 0.00415690485651019 | 11065/1163/5111/4176/5698/5685 | 6 |
| Mac_Prolif | R-HSA-5218920 | VEGFR2 mediated vascular permeability | 3/78 | 29/10704 | 0.00118672426169676 | 0.00704693994574571 | 0.00426267748911153 | 808/3320/805 | 3 |
| Mac_Prolif | R-HSA-114452 | Activation of BH3-only proteins | 3/78 | 30/10704 | 0.00131170260192179 | 0.00767503699485463 | 0.00464261192488465 | 7533/7531/7534 | 3 |
| Mac_Prolif | R-HSA-983705 | Signaling by the B Cell Receptor (BCR) | 5/78 | 112/10704 | 0.00131914698349064 | 0.00767503699485463 | 0.00464261192488465 | 808/5698/5685/5478/805 | 5 |
| Mac_Prolif | R-HSA-69618 | Mitotic Spindle Checkpoint | 5/78 | 113/10704 | 0.00137245215370545 | 0.00790532440534336 | 0.00478191224080529 | 11065/1063/332/11130/991 | 5 |
| Mac_Prolif | R-HSA-5339562 | Uptake and actions of bacterial toxins | 3/78 | 31/10704 | 0.001444667091206 | 0.00823889351024412 | 0.00498368741208115 | 808/3320/805 | 3 |
| Mac_Prolif | R-HSA-162906 | HIV Infection | 7/78 | 234/10704 | 0.0015423004135826 | 0.00870946115905466 | 0.0052683326821036 | 5902/5901/292/8905/5698/5685/5478 | 7 |
| Mac_Prolif | R-HSA-380259 | Loss of Nlp from mitotic centrosomes | 4/78 | 70/10704 | 0.00166207441240418 | 0.0092053352071616 | 0.0055682857338642 | 983/203068/7531/3320 | 4 |
| Mac_Prolif | R-HSA-380284 | Loss of proteins required for interphase microtubule organization from the centrosome | 4/78 | 70/10704 | 0.00166207441240418 | 0.0092053352071616 | 0.0055682857338642 | 983/203068/7531/3320 | 4 |
| Mac_Prolif | R-HSA-69239 | Synthesis of DNA | 5/78 | 120/10704 | 0.00179128950143576 | 0.00982650240787615 | 0.00594402832055374 | 11065/5111/4176/5698/5685 | 5 |
| Mac_Prolif | R-HSA-8854518 | AURKA Activation by TPX2 | 4/78 | 73/10704 | 0.00194081228842288 | 0.010541570480839 | 0.00637657132521514 | 983/203068/7531/3320 | 4 |
| Mac_Prolif | R-HSA-72203 | Processing of Capped Intron-Containing Pre-mRNA | 7/78 | 244/10704 | 0.00195824312751698 | 0.010541570480839 | 0.00637657132521514 | 79622/25804/6628/220988/3178/6428/3181 | 7 |
| Mac_Prolif | R-HSA-2299718 | Condensation of Prophase Chromosomes | 4/78 | 74/10704 | 0.0020404946853691 | 0.0108826383219685 | 0.00658288246449484 | 983/891/10051/6418 | 4 |
| Mac_Prolif | R-HSA-72163 | mRNA Splicing - Major Pathway | 6/78 | 183/10704 | 0.00213295395054449 | 0.0112713896836112 | 0.00681803725379261 | 25804/6628/220988/3178/6428/3181 | 6 |
| Mac_Prolif | R-HSA-2465910 | MASTL Facilitates Mitotic Progression | 2/78 | 10/10704 | 0.0022713752465656 | 0.0118937467456526 | 0.00719449958481545 | 983/891 | 2 |
| Mac_Prolif | R-HSA-69306 | DNA Replication | 5/78 | 128/10704 | 0.00237710360394028 | 0.0123352403231495 | 0.00746155801710983 | 11065/5111/4176/5698/5685 | 5 |
| Mac_Prolif | R-HSA-111933 | Calmodulin induced events | 3/78 | 37/10704 | 0.00242016875389613 | 0.01233643541809 | 0.00746228092724377 | 3838/808/805 | 3 |
| Mac_Prolif | R-HSA-111997 | CaM pathway | 3/78 | 37/10704 | 0.00242016875389613 | 0.01233643541809 | 0.00746228092724377 | 3838/808/805 | 3 |
| Mac_Prolif | R-HSA-8950505 | Gene and protein expression by JAK-STAT signaling after Interleukin-12 stimulation | 3/78 | 38/10704 | 0.0026139342446391 | 0.0132072467097554 | 0.00798903264058672 | 3181/6175/5478 | 3 |
| Mac_Prolif | R-HSA-111996 | Ca-dependent events | 3/78 | 39/10704 | 0.0028170262917291 | 0.0141096273394431 | 0.00853488057265291 | 3838/808/805 | 3 |
| Mac_Prolif | R-HSA-3371511 | HSF1 activation | 2/78 | 12/10704 | 0.00330002245580139 | 0.0161085841910305 | 0.00974404489625567 | 7531/3320 | 2 |
| Mac_Prolif | R-HSA-451306 | Ionotropic activity of kainate receptors | 2/78 | 12/10704 | 0.00330002245580139 | 0.0161085841910305 | 0.00974404489625567 | 808/805 | 2 |
| Mac_Prolif | R-HSA-451308 | Activation of Ca-permeable Kainate Receptor | 2/78 | 12/10704 | 0.00330002245580139 | 0.0161085841910305 | 0.00974404489625567 | 808/805 | 2 |
| Mac_Prolif | R-HSA-5617833 | Cilium Assembly | 6/78 | 201/10704 | 0.00340367637715986 | 0.0164749377583536 | 0.009965651312162 | 983/203068/10376/84790/7531/3320 | 6 |
| Mac_Prolif | R-HSA-68875 | Mitotic Prophase | 5/78 | 142/10704 | 0.0037208464752046 | 0.0175893012517646 | 0.0106397271826281 | 983/891/10051/7112/6418 | 5 |
| Mac_Prolif | R-HSA-1489509 | DAG and IP3 signaling | 3/78 | 43/10704 | 0.00372551172346403 | 0.0175893012517646 | 0.0106397271826281 | 3838/808/805 | 3 |
| Mac_Prolif | R-HSA-9646399 | Aggrephagy | 3/78 | 43/10704 | 0.00372551172346403 | 0.0175893012517646 | 0.0106397271826281 | 10376/84790/3320 | 3 |
| Mac_Prolif | R-HSA-450531 | Regulation of mRNA stability by proteins that bind AU-rich elements | 4/78 | 88/10704 | 0.00383010692873046 | 0.0176050094762006 | 0.010649229050845 | 6418/5698/5685/7534 | 4 |
| Mac_Prolif | R-HSA-140342 | Apoptosis induced DNA fragmentation | 2/78 | 13/10704 | 0.00388166007548173 | 0.0176050094762006 | 0.010649229050845 | 3148/3146 | 2 |
| Mac_Prolif | R-HSA-425561 | Sodium/Calcium exchangers | 2/78 | 13/10704 | 0.00388166007548173 | 0.0176050094762006 | 0.010649229050845 | 808/805 | 2 |
| Mac_Prolif | R-HSA-5210891 | Uptake and function of anthrax toxins | 2/78 | 13/10704 | 0.00388166007548173 | 0.0176050094762006 | 0.010649229050845 | 808/805 | 2 |
| Mac_Prolif | R-HSA-5607763 | CLEC7A (Dectin-1) induces NFAT activation | 2/78 | 13/10704 | 0.00388166007548173 | 0.0176050094762006 | 0.010649229050845 | 808/805 | 2 |
| Mac_Prolif | R-HSA-4615885 | SUMOylation of DNA replication proteins | 3/78 | 45/10704 | 0.00423929622408347 | 0.0190768330083756 | 0.0115395316625956 | 7153/332/5111 | 3 |
| Mac_Prolif | R-HSA-69052 | Switching of origins to a post-replicative state | 4/78 | 91/10704 | 0.00431841232949942 | 0.0190896911992437 | 0.0115473095521741 | 11065/4176/5698/5685 | 4 |
| Mac_Prolif | R-HSA-162658 | Golgi Cisternae Pericentriolar Stack Reorganization | 2/78 | 14/10704 | 0.00450728819982142 | 0.0190896911992437 | 0.0115473095521741 | 983/891 | 2 |
| Mac_Prolif | R-HSA-418359 | Reduction of cytosolic Ca++ levels | 2/78 | 14/10704 | 0.00450728819982142 | 0.0190896911992437 | 0.0115473095521741 | 808/805 | 2 |
| Mac_Prolif | R-HSA-442720 | CREB1 phosphorylation through the activation of Adenylate Cyclase | 2/78 | 14/10704 | 0.00450728819982142 | 0.0190896911992437 | 0.0115473095521741 | 808/805 | 2 |
| Mac_Prolif | R-HSA-9005891 | Loss of function of MECP2 in Rett syndrome | 2/78 | 14/10704 | 0.00450728819982142 | 0.0190896911992437 | 0.0115473095521741 | 808/805 | 2 |
| Mac_Prolif | R-HSA-9005895 | Pervasive developmental disorders | 2/78 | 14/10704 | 0.00450728819982142 | 0.0190896911992437 | 0.0115473095521741 | 808/805 | 2 |
| Mac_Prolif | R-HSA-9675151 | Disorders of Developmental Biology | 2/78 | 14/10704 | 0.00450728819982142 | 0.0190896911992437 | 0.0115473095521741 | 808/805 | 2 |
| Mac_Prolif | R-HSA-9697154 | Disorders of Nervous System Development | 2/78 | 14/10704 | 0.00450728819982142 | 0.0190896911992437 | 0.0115473095521741 | 808/805 | 2 |
| Mac_Prolif | R-HSA-9020591 | Interleukin-12 signaling | 3/78 | 47/10704 | 0.004794097476048 | 0.020156205446742 | 0.0121924415257156 | 3181/6175/5478 | 3 |
| Mac_Prolif | R-HSA-6791312 | TP53 Regulates Transcription of Cell Cycle Genes | 3/78 | 48/10704 | 0.00508716574640143 | 0.0212333874632407 | 0.0128440264077717 | 983/891/5111 | 3 |
| Mac_Prolif | R-HSA-141424 | Amplification of signal from the kinetochores | 4/78 | 96/10704 | 0.00522286410061538 | 0.0214883551568176 | 0.0129982557692007 | 1063/332/11130/991 | 4 |
| Mac_Prolif | R-HSA-141444 | Amplification of signal from unattached kinetochores via a MAD2 inhibitory signal | 4/78 | 96/10704 | 0.00522286410061538 | 0.0214883551568176 | 0.0129982557692007 | 1063/332/11130/991 | 4 |
| Mac_Prolif | R-HSA-71387 | Metabolism of carbohydrates | 7/78 | 295/10704 | 0.00558976040300768 | 0.022787006233612 | 0.0137838067677733 | 5223/808/7167/5315/2023/2597/805 | 7 |
| Mac_Prolif | R-HSA-5620912 | Anchoring of the basal body to the plasma membrane | 4/78 | 98/10704 | 0.00561763000898073 | 0.022787006233612 | 0.0137838067677733 | 983/203068/7531/3320 | 4 |
| Mac_Prolif | R-HSA-156711 | Polo-like kinase mediated events | 2/78 | 16/10704 | 0.00588790184810084 | 0.023716303947595 | 0.0143459367811658 | 1063/891 | 2 |
| Mac_Prolif | R-HSA-449147 | Signaling by Interleukins | 9/78 | 461/10704 | 0.00614934080792478 | 0.0245973632316991 | 0.014878887481163 | 332/3146/5698/5685/3181/6175/5478/3320/7534 | 9 |
| Mac_Prolif | R-HSA-5607764 | CLEC7A (Dectin-1) signaling | 4/78 | 101/10704 | 0.00624644113624419 | 0.0248134489274252 | 0.0150095972130441 | 808/5698/5685/805 | 4 |
| Mac_Prolif | R-HSA-2980766 | Nuclear Envelope Breakdown | 3/78 | 52/10704 | 0.00636635174057328 | 0.0251165657710288 | 0.0151929518826947 | 983/891/7112 | 3 |
| Mac_Prolif | R-HSA-70221 | Glycogen breakdown (glycogenolysis) | 2/78 | 17/10704 | 0.00664159775271209 | 0.026024219765729 | 0.0157419896609216 | 808/805 | 2 |
| Mac_Prolif | R-HSA-109606 | Intrinsic Pathway for Apoptosis | 3/78 | 53/10704 | 0.0067133651477245 | 0.0261276913857386 | 0.0158045794018265 | 7533/7531/7534 | 3 |
| Mac_Prolif | R-HSA-9604323 | Negative regulation of NOTCH4 signaling | 3/78 | 54/10704 | 0.00707144757224811 | 0.0273366026954021 | 0.0165358470251793 | 5698/5685/7534 | 3 |
| Mac_Prolif | R-HSA-190840 | Microtubule-dependent trafficking of connexons from Golgi to the plasma membrane | 2/78 | 18/10704 | 0.00743670479365617 | 0.0280119857007472 | 0.0169443846252692 | 10376/84790 | 2 |
| Mac_Prolif | R-HSA-112043 | PLC beta mediated events | 3/78 | 55/10704 | 0.00744068370176098 | 0.0280119857007472 | 0.0169443846252692 | 3838/808/805 | 3 |
| Mac_Prolif | R-HSA-174113 | SCF-beta-TrCP mediated degradation of Emi1 | 3/78 | 55/10704 | 0.00744068370176098 | 0.0280119857007472 | 0.0169443846252692 | 991/5698/5685 | 3 |
| Mac_Prolif | R-HSA-3371497 | HSP90 chaperone cycle for steroid hormone receptors (SHR) | 3/78 | 55/10704 | 0.00744068370176098 | 0.0280119857007472 | 0.0169443846252692 | 10376/84790/3320 | 3 |
| Mac_Prolif | R-HSA-112040 | G-protein mediated events | 3/78 | 56/10704 | 0.00782115463948883 | 0.0290644198215843 | 0.0175809995631294 | 3838/808/805 | 3 |
| Mac_Prolif | R-HSA-2151201 | Transcriptional activation of mitochondrial biogenesis | 3/78 | 56/10704 | 0.00782115463948883 | 0.0290644198215843 | 0.0175809995631294 | 3418/808/805 | 3 |
| Mac_Prolif | R-HSA-447115 | Interleukin-12 family signaling | 3/78 | 57/10704 | 0.00821293796269578 | 0.0301583221367534 | 0.0182426985147394 | 3181/6175/5478 | 3 |
| Mac_Prolif | R-HSA-1362277 | Transcription of E2F targets under negative control by DREAM complex | 2/78 | 19/10704 | 0.00827259530834555 | 0.0301583221367534 | 0.0182426985147394 | 7153/5111 | 2 |
| Mac_Prolif | R-HSA-190872 | Transport of connexons to the plasma membrane | 2/78 | 19/10704 | 0.00827259530834555 | 0.0301583221367534 | 0.0182426985147394 | 10376/84790 | 2 |
| Mac_Prolif | R-HSA-163615 | PKA activation | 2/78 | 20/10704 | 0.00914864837825899 | 0.0329351341617324 | 0.0199223856131824 | 808/805 | 2 |
| Mac_Prolif | R-HSA-176407 | Conversion from APC/C:Cdc20 to APC/C:Cdh1 in late anaphase | 2/78 | 20/10704 | 0.00914864837825899 | 0.0329351341617324 | 0.0199223856131824 | 11065/991 | 2 |
| Mac_Prolif | R-HSA-5610787 | Hedgehog 'off' state | 4/78 | 113/10704 | 0.00922702780162088 | 0.0330109814517617 | 0.0199682654615006 | 10376/84790/5698/5685 | 4 |
| Mac_Prolif | R-HSA-187577 | SCF(Skp2)-mediated degradation of p27/p21 | 3/78 | 60/10704 | 0.00945688633538299 | 0.0336244847480284 | 0.0203393721703169 | 1163/5698/5685 | 3 |
| Mac_Prolif | R-HSA-141405 | Inhibition of the proteolytic activity of APC/C required for the onset of anaphase by mitotic spindle chec | 2/78 | 21/10704 | 0.0100642497660343 | 0.0349217341279262 | 0.0211240752857156 | 11065/991 | 2 |
| Mac_Prolif | R-HSA-141430 | Inactivation of APC/C via direct inhibition of the APC/C complex | 2/78 | 21/10704 | 0.0100642497660343 | 0.0349217341279262 | 0.0211240752857156 | 11065/991 | 2 |
| Mac_Prolif | R-HSA-442982 | Ras activation upon Ca2+ influx through NMDA receptor | 2/78 | 21/10704 | 0.0100642497660343 | 0.0349217341279262 | 0.0211240752857156 | 808/805 | 2 |
| Mac_Prolif | R-HSA-5218921 | VEGFR2 mediated cell proliferation | 2/78 | 21/10704 | 0.0100642497660343 | 0.0349217341279262 | 0.0211240752857156 | 808/805 | 2 |
| Mac_Prolif | R-HSA-111931 | PKA-mediated phosphorylation of CREB | 2/78 | 22/10704 | 0.0110187918531015 | 0.0371159304525525 | 0.0224513395098591 | 808/805 | 2 |
| Mac_Prolif | R-HSA-113510 | E2F mediated regulation of DNA replication | 2/78 | 22/10704 | 0.0110187918531015 | 0.0371159304525525 | 0.0224513395098591 | 983/891 | 2 |
| Mac_Prolif | R-HSA-389977 | Post-chaperonin tubulin folding pathway | 2/78 | 22/10704 | 0.0110187918531015 | 0.0371159304525525 | 0.0224513395098591 | 10376/84790 | 2 |

| | | | | | | | | | |
|---|---|---|---|---|---|---|---|---|---|
| Mac_Prolif | R-HSA-438066 | Unblocking of NMDA receptors, glutamate binding and activation | 2/78 | 22/10704 | 0.0110187918531015 | 0.0371159304525525 | 0.0224513395098591 | 808/805 | 2 |
| Mac_Prolif | R-HSA-9617324 | Negative regulation of NMDA receptor-mediated neuronal transmission | 2/78 | 22/10704 | 0.0110187918531015 | 0.0371159304525525 | 0.0224513395098591 | 808/805 | 2 |
| Mac_Prolif | R-HSA-174084 | Autodegradation of Cdh1 by Cdh1:APC/C | 3/78 | 64/10704 | 0.0112779703938015 | 0.0377680869001723 | 0.0228458274195121 | 11065/5698/5685 | 3 |
| Mac_Prolif | R-HSA-8878166 | Transcriptional regulation by RUNX2 | 4/78 | 121/10704 | 0.0116559937934522 | 0.0388083955203957 | 0.0234751076704148 | 983/891/5698/5685 | 4 |
| Mac_Prolif | R-HSA-9637690 | Response of Mtb to phagocytosis | 2/78 | 23/10704 | 0.0120116735778524 | 0.0397627814990977 | 0.0240524135164498 | 11151/2023 | 2 |
| Mac_Prolif | R-HSA-5627123 | RHO GTPases activate PAKs | 2/78 | 24/10704 | 0.0130423003743416 | 0.0426838921342088 | 0.0258193865066212 | 808/805 | 2 |
| Mac_Prolif | R-HSA-9620244 | Long-term potentiation | 2/78 | 24/10704 | 0.0130423003743416 | 0.0426838921342088 | 0.0258193865066212 | 808/805 | 2 |
| Mac_Prolif | R-HSA-68867 | Assembly of the pre-replicative complex | 3/78 | 68/10704 | 0.0132879993699392 | 0.0432423030343784 | 0.0261571679539094 | 4176/5698/5685 | 3 |
| Mac_Prolif | R-HSA-389960 | Formation of tubulin folding intermediates by CCT/TriC | 2/78 | 25/10704 | 0.0141100841115151 | 0.0454045164705738 | 0.0274650858036548 | 10376/84790 | 2 |
| Mac_Prolif | R-HSA-69273 | Cyclin A/B1/B2 associated events during G2/M transition | 2/78 | 25/10704 | 0.0141100841115151 | 0.0454045164705738 | 0.0274650858036548 | 983/891 | 2 |
| Mac_Prolif | R-HSA-68949 | Orc1 removal from chromatin | 3/78 | 71/10704 | 0.0149212379641982 | 0.0477479614854343 | 0.0288826302114012 | 4176/5698/5685 | 3 |
| Mac_Prolif | R-HSA-110373 | Resolution of AP sites via the multiple-nucleotide patch replacement pathway | 2/78 | 26/10704 | 0.0152144430329625 | 0.0478880829889966 | 0.0289673893811365 | 5111/142 | 2 |
| Mac_Prolif | R-HSA-179409 | APC-Cdc20 mediated degradation of Nek2A | 2/78 | 26/10704 | 0.0152144430329625 | 0.0478880829889966 | 0.0289673893811365 | 11065/991 | 2 |
| Mac_Prolif | R-HSA-180024 | DARPP-32 events | 2/78 | 26/10704 | 0.0152144430329625 | 0.0478880829889966 | 0.0289673893811365 | 808/805 | 2 |
| RTM_IFN | R-HSA-913531 | Interferon Signaling | 18/75 | 199/10704 | 1.2279757181133e-15 | 5.25573607352494e-13 | 4.54997318711456e-13 | 3429/6772/115362/2633/10410/9636/3383/2634/3117/8519/3105/10581/3133/3119/3115/3113/3106/567 | 18 |
| RTM_IFN | R-HSA-877300 | Interferon gamma signaling | 13/75 | 91/10704 | 4.12230422561768e-14 | 8.82173104282183e-12 | 7.63711098640749e-12 | 6772/115362/2633/3383/2634/3117/3105/3133/3119/3115/3113/3106/567 | 13 |
| RTM_IFN | R-HSA-909733 | Interferon alpha/beta signaling | 10/75 | 70/10704 | 4.35076392859657e-11 | 6.20708987146445e-09 | 5.37357509777542e-09 | 3429/6772/10410/9636/2634/8519/3105/10581/3133/3106 | 10 |
| RTM_IFN | R-HSA-2132295 | MHC class II antigen presentation | 11/75 | 123/10704 | 7.72601898010404e-10 | 8.26684030871132e-08 | 7.15673337104374e-08 | 5641/1514/3117/1075/3109/3119/3115/3113/1508/7846/972 | 11 |
| RTM_IFN | R-HSA-1236977 | Endosomal/Vacuolar pathway | 5/75 | 11/10704 | 6.5959993179483e-09 | 5.64617541616374e-07 | 4.88798265245853e-07 | 1514/3105/3133/3106/567 | 5 |
| RTM_IFN | R-HSA-1236975 | Antigen processing-Cross presentation | 9/75 | 98/10704 | 2.37502814922927e-08 | 1.69418674645021e-06 | 1.46668405005036e-06 | 5721/1514/5698/653361/3105/929/3133/3106/567 | 9 |
| RTM_IFN | R-HSA-202430 | Translocation of ZAP-70 to Immunological synapse | 5/75 | 19/10704 | 1.58914271857628e-07 | 9.7164726221521e-06 | 8.41170281111053e-06 | 920/3117/3119/3115/3113 | 5 |
| RTM_IFN | R-HSA-202427 | Phosphorylation of CD3 and TCR zeta chains | 5/75 | 22/10704 | 3.54046283572328e-07 | 1.89414761711196e-05 | 1.63979331338763e-05 | 920/3117/3119/3115/3113 | 5 |
| RTM_IFN | R-HSA-389948 | PD-1 signaling | 5/75 | 23/10704 | 4.49929216497282e-07 | 2.13966338512041e-05 | 1.85234016616425e-05 | 920/3117/3119/3115/3113 | 5 |
| RTM_IFN | R-HSA-1236974 | ER-Phagosome pathway | 7/75 | 83/10704 | 1.69194919641202e-06 | 7.24154256064346e-05 | 6.26911702249508e-05 | 5721/5698/3105/929/3133/3106/567 | 7 |
| RTM_IFN | R-HSA-202433 | Generation of second messenger molecules | 5/75 | 34/10704 | 3.50385724057381e-06 | 0.0001300040490722 | 0.000112546545187935 | 920/3117/3119/3115/3113 | 5 |
| RTM_IFN | R-HSA-198933 | Immunoregulatory interactions between a Lymphoid and a non-Lymphoid cell | 8/75 | 132/10704 | 3.64497333847289e-06 | 0.0001300040490722 | 0.000112546545187935 | 11006/3383/8519/3105/2214/3133/3106/567 | 8 |
| RTM_IFN | R-HSA-202424 | Downstream TCR signaling | 7/75 | 98/10704 | 5.18554803597798e-06 | 0.000170724196876814 | 0.000147798616086174 | 5721/5698/920/3117/3119/3115/3113 | 7 |
| RTM_IFN | R-HSA-202403 | TCR signaling | 7/75 | 120/10704 | 1.9725590191148e-05 | 0.000603039471557953 | 0.000522060732878503 | 5721/5698/920/3117/3119/3115/3113 | 7 |
| RTM_IFN | R-HSA-6798695 | Neutrophil degranulation | 13/75 | 480/10704 | 2.49398134172369e-05 | 0.000672899901547228 | 0.000582540003306996 | 1522/10875/8804/4318/1075/3105/8635/929/6518/2212/1508/3106/567 | 13 |
| RTM_IFN | R-HSA-983170 | Antigen Presentation: Folding, assembly and peptide loading of class I MHC | 4/75 | 25/10704 | 2.51551365064384e-05 | 0.000672899901547228 | 0.000582540003306996 | 3105/3133/3106/567 | 4 |
| RTM_IFN | R-HSA-380108 | Chemokine receptors bind chemokines | 5/75 | 59/10704 | 5.50130746873037e-05 | 0.00138503505683329 | 0.00119904658141987 | 4283/3627/6373/6357/6347 | 5 |
| RTM_IFN | R-HSA-1679131 | Trafficking and processing of endosomal TLR | 3/75 | 13/10704 | 8.98504903279417e-05 | 0.00213644499224217 | 0.00184955395294944 | 5641/1514/1508 | 3 |
| RTM_IFN | R-HSA-166786 | Creation of C4 and C2 activators | 3/75 | 14/10704 | 0.000113779637241982 | 0.00250286055851017 | 0.00216676565812981 | 713/714/712 | 3 |
| RTM_IFN | R-HSA-388841 | Costimulation by the CD28 family | 5/75 | 69/10704 | 0.000116956100864961 | 0.00250286055851017 | 0.00216676565812981 | 920/3117/3119/3115/3113 | 5 |
| RTM_IFN | R-HSA-3299685 | Detoxification of Reactive Oxygen Species | 4/75 | 37/10704 | 0.000123238724616461 | 0.00251172257789739 | 0.00217443764736813 | 475/653361/5052/2876 | 4 |
| RTM_IFN | R-HSA-983169 | Class I MHC mediated antigen processing & presentation | 10/75 | 370/10704 | 0.000241050817901687 | 0.00468953409372374 | 0.00405980324887053 | 5721/1514/5698/57674/653361/3105/929/3133/3106/567 | 10 |
| RTM_IFN | R-HSA-8963898 | Plasma lipoprotein assembly | 3/75 | 19/10704 | 0.000295351746711943 | 0.00549611076490051 | 0.00475806932918096 | 2/348/19 | 3 |
| RTM_IFN | R-HSA-977606 | Regulation of Complement cascade | 4/75 | 47/10704 | 0.000315669303086142 | 0.0056294359050362 | 0.00487349099501412 | 713/714/710/712 | 4 |
| RTM_IFN | R-HSA-449147 | Signaling by Interleukins | 11/75 | 461/10704 | 0.000339164081927904 | 0.00580648908260572 | 0.00502676870899462 | 3627/9235/6772/5721/5698/920/3383/6347/4318/1436/3162 | 11 |
| RTM_IFN | R-HSA-164938 | Nef-mediates down modulation of cell surface receptors by recruiting them to clathrin adapters | 3/75 | 21/10704 | 0.000401321678058166 | 0.00660637223880365 | 0.00571924010835929 | 920/3105/567 | 3 |
| RTM_IFN | R-HSA-166663 | Initial triggering of complement | 3/75 | 23/10704 | 0.000529038340792146 | 0.00838623740218661 | 0.00726009730833666 | 713/714/712 | 3 |
| RTM_IFN | R-HSA-449836 | Other interleukin signaling | 3/75 | 24/10704 | 0.000601580568520111 | 0.00919558869023598 | 0.00796076541801049 | 9235/920/1436 | 3 |
| RTM_IFN | R-HSA-166658 | Complement cascade | 4/75 | 58/10704 | 0.000708475089707446 | 0.0104561151170616 | 0.00905202292475576 | 713/714/710/712 | 4 |
| RTM_IFN | R-HSA-6785807 | Interleukin-4 and Interleukin-13 signaling | 5/75 | 108/10704 | 0.000938716796023299 | 0.013175712644079 | 0.0114064211773631 | 6772/3383/6347/4318/3162 | 5 |
| RTM_IFN | R-HSA-164952 | The role of Nef in HIV-1 replication and disease pathogenesis | 3/75 | 28/10704 | 0.00095431563543563 | 0.013175712644079 | 0.0114064211773631 | 920/3105/567 | 3 |
| RTM_IFN | R-HSA-174824 | Plasma lipoprotein assembly, remodeling, and clearance | 4/75 | 70/10704 | 0.00143720552213146 | 0.0192226238585082 | 0.0166413270983642 | 5360/2/348/19 | 4 |
| RTM_IFN | R-HSA-164940 | Nef mediated downregulation of MHC class I complex cell surface expression | 2/75 | 10/10704 | 0.00210206595901321 | 0.025721243220495 | 0.0222672838505023 | 3105/567 | 2 |
| RTM_IFN | R-HSA-8964058 | HDL remodeling | 2/75 | 10/10704 | 0.00210206595901321 | 0.025721243220495 | 0.0222672838505023 | 5360/348 | 2 |
| RTM_IFN | R-HSA-162909 | Host Interactions of HIV factors | 5/75 | 130/10704 | 0.00214000486495849 | 0.025721243220495 | 0.0222672838505023 | 5721/5698/920/3105/567 | 5 |
| RTM_IFN | R-HSA-9029569 | NR1H3 & NR1H2 regulate gene expression linked to cholesterol transport and efflux | 3/75 | 37/10704 | 0.00216346905592948 | 0.025721243220495 | 0.0222672838505023 | 5360/348/19 | 3 |
| RTM_IFN | R-HSA-1474228 | Degradation of the extracellular matrix | 5/75 | 140/10704 | 0.00295337543798301 | 0.0341633699312629 | 0.0295757654102423 | 2/1514/4318/6696/1508 | 5 |
| RTM_IFN | R-HSA-2029481 | FCGR activation | 2/75 | 12/10704 | 0.00305517357382434 | 0.034410902357811 | 0.0297900581159604 | 2214/2212 | 2 |
| RTM_IFN | R-HSA-2172127 | DAP12 interactions | 3/75 | 45/10704 | 0.00379590492789184 | 0.0416576233112233 | 0.0360636581543301 | 3133/3106/567 | 3 |
| RTM_IFN | R-HSA-6783783 | Interleukin-10 signaling | 3/75 | 47/10704 | 0.00429444389241582 | 0.0448298045354628 | 0.0388098652151571 | 3627/3383/6347 | 3 |
| RTM_IFN | R-HSA-9024446 | NR1H2 and NR1H3-mediated signaling | 3/75 | 47/10704 | 0.00429444389241582 | 0.0448298045354628 | 0.0388098652151571 | 5360/348/19 | 3 |
| Mac_IFN | R-HSA-909733 | Interferon alpha/beta signaling | 15/33 | 70/10704 | 3.26450056192827e-25 | 2.77482547763903e-23 | 2.06178982858628e-23 | 9636/91543/3434/3669/3433/4599/3437/2537/4600/10410/8519/4940/4938/3665/10581 | 15 |
| Mac_IFN | R-HSA-913531 | Interferon Signaling | 17/33 | 199/10704 | 1.70871183803833e-21 | 7.2620253116629e-20 | 5.39593212012104e-20 | 9636/91543/3434/3669/3433/4599/3437/115362/2537/4600/2633/10410/8519/4940/4938/3665/10581 | 17 |
| Mac_IFN | R-HSA-1169410 | Antiviral mechanism by IFN-stimulated genes | 6/33 | 80/10704 | 1.3589064861321e-07 | 3.85023504404094e-06 | 2.8608557602781e-06 | 9636/3434/4599/4600/4940/4938 | 6 |
| Mac_IFN | R-HSA-877300 | Interferon gamma signaling | 5/33 | 91/10704 | 7.81832751136551e-06 | 0.000166139459616517 | 0.000123447276495245 | 115362/2633/4940/4938/3665 | 5 |
| Mac_IFN | R-HSA-6783783 | Interleukin-10 signaling | 4/33 | 47/10704 | 1.21616455279589e-05 | 0.000206747973975302 | 0.000153620785616323 | 3627/6347/3557/7076 | 4 |
| Mac_IFN | R-HSA-380108 | Chemokine receptors bind chemokines | 4/33 | 59/10704 | 3.023501258226e-05 | 0.00042832934491535 | 0.000318263290339579 | 3627/6354/6347/6373 | 4 |
| Mac_IFN | R-HSA-1169408 | ISG15 antiviral mechanism | 4/33 | 72/10704 | 6.64448825232072e-05 | 0.00080683071635323 | 0.000599502699457508 | 9636/3434/4599/4600 | 4 |
| Mac_IFN | R-HSA-375276 | Peptide ligand-binding receptors | 4/33 | 201/10704 | 0.0032236314376315 | 0.0342510840248347 | 0.0254497218760382 | 3627/6354/6347/6373 | 4 |
| Mac_Hypo | R-HSA-70171 | Glycolysis | 8/63 | 71/10704 | 7.24076945323556e-09 | 1.46987619900682e-06 | 1.23474173834122e-06 | 2026/3099/7167/2597/2023/5223/5230/5315 | 8 |
| Mac_Hypo | R-HSA-70263 | Gluconeogenesis | 6/63 | 34/10704 | 3.85359018665144e-08 | 3.57060636703278e-06 | 2.99942043795339e-06 | 2026/7167/2597/2023/5223/5230 | 6 |
| Mac_Hypo | R-HSA-70326 | Glucose metabolism | 8/63 | 91/10704 | 5.27675817788096e-08 | 3.57060636703278e-06 | 2.99942043795339e-06 | 2026/3099/7167/2597/2023/5223/5230/5315 | 8 |
| Mac_Hypo | R-HSA-71387 | Metabolism of carbohydrates | 12/63 | 295/10704 | 1.16656079457255e-07 | 5.9202960324557e-06 | 4.97323286107245e-06 | 6513/2026/3099/6383/2632/1462/7167/2597/2023/5223/5230/5315 | 12 |
| Mac_Hypo | R-HSA-1474244 | Extracellular matrix organization | 11/63 | 301/10704 | 1.2122406408325e-06 | 4.92169700177996e-05 | 4.13437860662874e-05 | 6696/4318/101/6383/5033/871/1462/4316/4323/7076/1514 | 11 |
| Mac_Hypo | R-HSA-1474228 | Degradation of the extracellular matrix | 7/63 | 140/10704 | 1.69190940084443e-05 | 0.0005724293472857 | 0.000480858461292629 | 6696/4318/101/4316/4323/7076/1514 | 7 |
| Mac_Hypo | R-HSA-1592389 | Activation of Matrix Metalloproteinases | 4/63 | 33/10704 | 3.92295216588331e-05 | 0.00113765612810616 | 0.000955666542666311 | 4318/4316/4323/7076 | 4 |
| Mac_Hypo | R-HSA-1474290 | Collagen formation | 5/63 | 90/10704 | 0.000179857905984777 | 0.00456389436436373 | 0.0038338132591492 | 4318/5033/871/4316/1514 | 5 |
| Mac_Hypo | R-HSA-8957275 | Post-translational protein phosphorylation | 5/63 | 108/10704 | 0.000420656446708091 | 0.00913712299273672 | 0.00767546759047627 | 6696/6383/1462/3956/7076 | 5 |
| Mac_Hypo | R-HSA-6798695 | Neutrophil degranulation | 10/63 | 480/10704 | 0.000450104580922991 | 0.00913712299273672 | 0.00767546759047627 | 1476/4318/101/6515/4282/23601/3310/5223/3304/5315 | 10 |
| Mac_Hypo | R-HSA-1442490 | Collagen degradation | 4/63 | 64/10704 | 0.000531474642531888 | 0.00980812299854302 | 0.00823912842968094 | 4318/4316/4323/1514 | 4 |
| Mac_Hypo | R-HSA-3371453 | Regulation of HSF1-mediated heat shock response | 4/63 | 69/10704 | 0.000707433030590723 | 0.0119674087674931 | 0.0100529956978682 | 9531/3310/10808/3304 | 4 |
| Mac_Hypo | R-HSA-381426 | Regulation of Insulin-like Growth Factor (IGF) transport and uptake by Insulin-like Growth Factor Bind | 5/63 | 125/10704 | 0.000819904761178708 | 0.0128031281937906 | 0.010755026017081 | 6696/6383/1462/3956/7076 | 5 |
| Mac_Hypo | R-HSA-8950505 | Gene and protein expression by JAK-STAT signaling after Interleukin-12 stimulation | 3/63 | 38/10704 | 0.0014153270186682 | 0.0205222417706889 | 0.0172393215807706 | 5898/4282/6648 | 3 |
| Mac_Hypo | R-HSA-3371556 | Cellular response to heat stress | 4/63 | 89/10704 | 0.00183033613841262 | 0.0247705490731841 | 0.0208080318893224 | 9531/3310/10808/3304 | 4 |
| Mac_Hypo | R-HSA-6783783 | Interleukin-10 signaling | 3/63 | 47/10704 | 0.00262001272454842 | 0.0312860342990194 | 0.0262812421905167 | 3576/3557/7076 | 3 |
| Mac_Hypo | R-HSA-9020591 | Interleukin-12 signaling | 3/63 | 47/10704 | 0.00262001272454842 | 0.0312860342990194 | 0.0262812421905167 | 5898/4282/6648 | 3 |
| Mac_Hypo | R-HSA-6785807 | Interleukin-4 and Interleukin-13 signaling | 4/63 | 108/10704 | 0.00369716919262074 | 0.0416958525612228 | 0.0350258134037754 | 4318/3576/7422/7076 | 4 |
| Mac_Hypo | R-HSA-447115 | Interleukin-12 family signaling | 3/63 | 57/10704 | 0.0045347244174329 | 0.0484499503546779 | 0.0406994656855474 | 5898/4282/6648 | 3 |
| Mac_LA | R-HSA-6798695 | Neutrophil degranulation | 26/87 | 480/10704 | 3.64874652675651e-15 | 1.39746991974774e-12 | 1.06773845730348e-12 | 1116/1476/948/4318/1118/2171/2760/25798/10493/3958/2934/3916/2268/7077/64114/2799/10159/3689/5315/3074/1509/302/967/5476/968/1508 | 26 |
| Mac_LA | R-HSA-114608 | Platelet degranulation | 10/87 | 129/10704 | 8.29993510960227e-08 | 1.16798099378119e-05 | 8.92397186398714e-06 | 7078/948/6678/2316/928/813/808/2335/967/334 | 10 |
| Mac_LA | R-HSA-76005 | Response to elevated platelet cytosolic Ca2+ | 10/87 | 134/10704 | 1.19085662206867e-07 | 1.16798099378119e-05 | 8.92397186398714e-06 | 7078/948/6678/2316/928/813/808/2335/967/334 | 10 |
| Mac_LA | R-HSA-1474244 | Extracellular matrix organization | 14/87 | 301/10704 | 1.21982349219968e-07 | 1.16798099378119e-05 | 8.92397186398714e-06 | 4318/6383/6678/6696/1291/3693/1513/4316/7077/5034/3689/1509/2335/1508 | 14 |
| Mac_LA | R-HSA-76002 | Platelet activation, signaling and aggregation | 12/87 | 263/10704 | 1.29944335951786e-06 | 9.95373613390682e-05 | 7.60516324096769e-05 | 7078/948/6678/2316/11343/928/813/5880/808/2335/967/334 | 12 |
| Mac_LA | R-HSA-1474228 | Degradation of the extracellular matrix | 9/87 | 140/10704 | 1.86948316608588e-06 | 0.000119335342101815 | 9.11783017845393e-05 | 4318/6696/1291/1513/4316/7077/1509/2335/1508 | 9 |
| Mac_LA | R-HSA-8957275 | Post-translational protein phosphorylation | 8/87 | 108/10704 | 2.49741491350262e-06 | 0.000136644273124501 | 0.000104403209917854 | 6383/56975/6696/813/348/5034/2335/334 | 8 |
| Mac_LA | R-HSA-381426 | Regulation of Insulin-like Growth Factor (IGF) transport and uptake by Insulin-like Growth Factor Bind | 8/87 | 125/10704 | 7.46074587936148e-06 | 0.000357183208974431 | 0.000272906230850328 | 6383/56975/6696/813/348/5034/2335/334 | 8 |
| Mac_LA | R-HSA-1442490 | Collagen degradation | 6/87 | 64/10704 | 1.24485890702733e-05 | 0.000529756623768297 | 0.000404761141700115 | 4318/1291/1513/4316/1509/1508 | 6 |
| Mac_LA | R-HSA-2173782 | Binding and Uptake of Ligands by Scavenger Receptors | 5/87 | 42/10704 | 2.12065125858863e-05 | 0.000812209432039444 | 0.000620569526197514 | 948/6678/348/811/4035 | 5 |
| Mac_LA | R-HSA-975634 | Retinoid metabolism and transport | 5/87 | 44/10704 | 2.6730044118267e-05 | 0.000930691536117843 | 0.000711095910514664 | 4023/2239/6383/348/4035 | 5 |
| Mac_LA | R-HSA-6806667 | Metabolism of fat-soluble vitamins | 5/87 | 48/10704 | 4.10834001069416e-05 | 0.00131124518674655 | 0.00100185835348507 | 4023/2239/6383/348/4035 | 5 |
| Mac_LA | R-HSA-1592389 | Activation of Matrix Metalloproteinases | 4/87 | 33/10704 | 0.000139160362317913 | 0.00382604706624457 | 0.00292329554601069 | 4318/1513/4316/7077 | 4 |
| Mac_LA | R-HSA-1679131 | Trafficking and processing of endosomal TLR | 3/87 | 13/10704 | 0.000139855506337922 | 0.00382604706624457 | 0.00292329554601069 | 1513/5641/1508 | 3 |
| Mac_LA | R-HSA-2187338 | Visual phototransduction | 6/87 | 104/10704 | 0.000194423153501146 | 0.00496427118606259 | 0.00379295695953113 | 4023/2239/6383/348/808/4035 | 6 |
| Mac_LA | R-HSA-6785807 | Interleukin-4 and Interleukin-13 signaling | 6/87 | 108/10704 | 0.000238922409747495 | 0.00571920518333067 | 0.00436976512564498 | 948/4318/6367/7431/3689/2335 | 6 |
| Mac_LA | R-HSA-216083 | Integrin cell surface interactions | 5/87 | 85/10704 | 0.000621918346636715 | 0.0140114545154036 | 0.0107054675148611 | 6696/1291/3693/3689/2335 | 5 |
| Mac_LA | R-HSA-983712 | Ion channel transport | 7/87 | 186/10704 | 0.000773929383282201 | 0.0159989801267078 | 0.0122240386841412 | 493/80727/523/476/808/537/6446 | 7 |
| Mac_LA | R-HSA-1474290 | Collagen formation | 5/87 | 90/10704 | 0.000807255019956185 | 0.0159989801267078 | 0.0122240386841412 | 4318/1291/4316/5034/1508 | 5 |
| Mac_LA | R-HSA-202733 | Cell surface interactions at the vascular wall | 6/87 | 137/10704 | 0.000850837296335254 | 0.0159989801267078 | 0.0122240386841412 | 6383/8832/4267/3689/9123/2335 | 6 |
| Mac_LA | R-HSA-196854 | Metabolism of vitamins and cofactors | 7/87 | 190/10704 | 0.000877228675354735 | 0.0159989801267078 | 0.0122240386841412 | 54/4023/2239/6383/348/808/4035 | 7 |
| Mac_LA | R-HSA-3000170 | Syndecan interactions | 3/87 | 27/10704 | 0.00131744216594146 | 0.0229354704343445 | 0.0175238718723314 | 6383/3693/2335 | 3 |
| Mac_LA | R-HSA-2022090 | Assembly of collagen fibrils and other multimeric structures | 4/87 | 61/10704 | 0.00149297550001518 | 0.0248612876741658 | 0.0189952946912686 | 4318/1291/4316/1508 | 4 |
| Mac_LA | R-HSA-174824 | Plasma lipoprotein assembly, remodeling, and clearance | 4/87 | 70/10704 | 0.00248189288911804 | 0.0396068740221754 | 0.0302616764550357 | 4023/3988/348/5034 | 4 |
| Mac_LA | R-HSA-71387 | Metabolism of carbohydrates | 8/87 | 295/10704 | 0.00268315964244264 | 0.039958282466949 | 0.030530170470831 | 2239/6383/219855/2203/808/2799/5315/3074 | 8 |
| Mac_LA | R-HSA-8963888 | Chylomicron assembly | 2/87 | 10/10704 | 0.00281690241934106 | 0.039958282466949 | 0.030530170470831 | 348/5034 | 2 |
| Mac_LA | R-HSA-8963901 | Chylomicron remodeling | 2/87 | 10/10704 | 0.00281690241934106 | 0.039958282466949 | 0.030530170470831 | 4023/348 | 2 |
| Mac_LA | R-HSA-2132295 | MHC class II antigen presentation | 5/87 | 123/10704 | 0.0032206247630713 | 0.0440535458662967 | 0.0336591610576624 | 1513/5641/1509/5476/1508 | 5 |
| Mac_LA | R-HSA-3000178 | ECM proteoglycans | 4/87 | 76/10704 | 0.00334719628998168 | 0.0442060751401029 | 0.0337757012201418 | 6678/1291/3693/2335 | 4 |
| Mac_LA | R-HSA-111465 | Apoptotic cleavage of cellular proteins | 3/87 | 38/10704 | 0.00356290667882929 | 0.0454864419330539 | 0.0347539669022646 | 2934/10134/7431 | 3 |
| Mac_LA | R-HSA-5668914 | Diseases of metabolism | 7/87 | 245/10704 | 0.00372802671870744 | 0.0460591688149984 | 0.0351915595178495 | 2239/6383/10417/1593/2799/3074/5476 | 7 |
| Mac_LA | R-HSA-449147 | Signaling by Interleukins | 10/87 | 461/10704 | 0.00408193051291881 | 0.0464597068010049 | 0.0354975910146472 | 948/4318/6367/3557/7431/5034/3689/3936/302/2335 | 10 |
| Mac_LA | R-HSA-264870 | Caspase-mediated cleavage of cytoskeletal proteins | 2/87 | 12/10704 | 0.00408804247512988 | 0.0464597068010049 | 0.0354975910146472 | 2934/7431 | 2 |
| Mac_LA | R-HSA-445355 | Smooth Muscle Contraction | 3/87 | 40/10704 | 0.00412436039486728 | 0.0464597068010049 | 0.0354975910146472 | 3693/808/302 | 3 |
| Mac_LA | R-HSA-198933 | Immunoregulatory interactions between a Lymphoid and a non-Lymphoid cell | 5/87 | 132/10704 | 0.00435594563811432 | 0.0470674164895478 | 0.0359619122827932 | 54209/4267/10870/3689/975 | 5 |
| Mac_LA | R-HSA-3560782 | Diseases associated with glycosaminoglycan metabolism | 3/87 | 41/10704 | 0.00442409136716377 | 0.0470674164895478 | 0.0359619122827932 | 2239/6383/3074 | 3 |
| Mac_LA | R-HSA-2022857 | Keratan sulfate degradation | 2/87 | 13/10704 | 0.00480589465127388 | 0.0478631478483788 | 0.0365698917187008 | 2799/3074 | 2 |
| Mac_LA | R-HSA-9656223 | Signaling by RAF1 mutants | 3/87 | 43/10704 | 0.00506237526172789 | 0.0478631478483788 | 0.0365698917187008 | 808/2335/71 | 3 |
| Mac_LA | R-HSA-397014 | Muscle contraction | 6/87 | 197/10704 | 0.00529017677157272 | 0.0478631478483788 | 0.0365698917187008 | 493/3693/476/808/7431/302 | 6 |
| Mac_LA | R-HSA-193775 | Synthesis of bile acids and bile salts via 24-hydroxycholesterol | 2/87 | 14/10704 | 0.00557738486377646 | 0.0478631478483788 | 0.0365698917187008 | 80270/1593 | 2 |
| Mac_LA | R-HSA-3656237 | Defective EXT2 causes exostoses 2 | 2/87 | 14/10704 | 0.00557738486377646 | 0.0478631478483788 | 0.0365698917187008 | 2239/6383 | 2 |
| Mac_LA | R-HSA-3656253 | Defective EXT1 causes exostoses 1, TRPS2 and CHDS | 2/87 | 14/10704 | 0.00557738486377646 | 0.0478631478483788 | 0.0365698917187008 | 2239/6383 | 2 |
| Mac_LA | R-HSA-418359 | Reduction of cytosolic Ca++ levels | 2/87 | 14/10704 | 0.00557738486377646 | 0.0478631478483788 | 0.0365698917187008 | 493/808 | 2 |
| Mac_LA | R-HSA-418346 | Platelet homeostasis | 4/87 | 88/10704 | 0.00565240661610104 | 0.0478631478483788 | 0.0365698917187008 | 493/2778/2268/808 | 4 |
| Mac_LA | R-HSA-3781865 | Diseases of glycosylation | 5/87 | 143/10704 | 0.0061006496322197 | 0.0478631478483788 | 0.0365698917187008 | 2239/6383/10417/3074/5476 | 5 |
| Mac_LA | R-HSA-1660662 | Glycosphingolipid metabolism | 3/87 | 46/10704 | 0.00611910749258343 | 0.0478631478483788 | 0.0365698917187008 | 2760/3074/5476 | 3 |
| Mac_LA | R-HSA-193807 | Synthesis of bile acids and bile salts via 27-hydroxycholesterol | 2/87 | 15/10704 | 0.00640161420716179 | 0.0478631478483788 | 0.0365698917187008 | 80270/1593 | 2 |
| Mac_LA | R-HSA-6802946 | Signaling by moderate kinase activity BRAF mutants | 3/87 | 47/10704 | 0.00649839083058929 | 0.0478631478483788 | 0.0365698917187008 | 808/2335/71 | 3 |
| Mac_LA | R-HSA-6802949 | Signaling by RAS mutants | 3/87 | 47/10704 | 0.00649839083058929 | 0.0478631478483788 | 0.0365698917187008 | 808/2335/71 | 3 |
| Mac_LA | R-HSA-6802955 | Paradoxical activation of RAF signaling by kinase inactive BRAF | 3/87 | 47/10704 | 0.00649839083058929 | 0.0478631478483788 | 0.0365698917187008 | 808/2335/71 | 3 |
| Mac_LA | R-HSA-9020591 | Interleukin-12 signaling | 3/87 | 47/10704 | 0.00649839083058929 | 0.0478631478483788 | 0.0365698917187008 | 5034/3936/302 | 3 |
| Mac_LA | R-HSA-9649948 | Signaling downstream of RAS mutants | 3/87 | 47/10704 | 0.00649839083058929 | 0.0478631478483788 | 0.0365698917187008 | 808/2335/71 | 3 |
| RTM-like_MT | R-HSA-166663 | Initial triggering of complement | 4/38 | 23/10704 | 1.13921640736194e-06 | 0.000267579710745769 | 0.000235621810069655 | 2219/1675/5199/717 | 4 |
| RTM-like_MT | R-HSA-166658 | Complement cascade | 5/38 | 58/10704 | 1.71525455606262e-06 | 0.000267579710745769 | 0.000235621810069655 | 2219/1675/5199/717/728 | 5 |
| RTM-like_MT | R-HSA-5686938 | Regulation of TLR by endogenous ligand | 3/38 | 19/10704 | 3.84627315835682e-05 | 0.0040001240846911 | 0.0035223764713373 | 6279/6280/23643 | 3 |
| RTM-like_MT | R-HSA-6783783 | Interleukin-10 signaling | 3/38 | 47/10704 | 0.000600962918963999 | 0.0375000861433535 | 0.033021330915706 | 2920/7076/3586 | 3 |
| RTM-like_MT | R-HSA-977606 | Regulation of Complement cascade | 3/38 | 47/10704 | 0.000600962918963999 | 0.0375000861433535 | 0.033021330915706 | 5199/717/728 | 3 |

**Supplementary Table 5: Enrichment pathway analysis**
**Dendritic cells**

| Cluster | ID | Description | GeneRatio | BgRatio | pvalue | p.adjust | qvalue | geneID | Count |
|---|---|---|---|---|---|---|---|---|---|
| cDC2B_FCER1A | R-HSA-198933 | Immunoregulatory interactions between a Lymphoid and a non-Lymphoid cell | 4/11 | 132/10704 | 6.81940193217667e-06 | 0.000231859665694007 | 0.000114853085173502 | 911/120425/912/10870 | 4 |
| cDC2A_AREG | R-HSA-6785807 | Interleukin-4 and Interleukin-13 signaling | 5/35 | 108/10704 | 2.43049079324897e-05 | 0.00422905398025321 | 0.00355619179222744 | 1052/7422/3726/3553/3576 | 5 |
| cDC2A_AREG | R-HSA-380994 | ATF4 activates genes in response to endoplasmic reticulum stress | 3/35 | 27/10704 | 8.8774404965677e-05 | 0.0077233732320139 | 0.00649454857380479 | 9709/1051/3576 | 3 |
| cDC2A_AREG | R-HSA-381042 | PERK regulates gene expression | 3/35 | 32/10704 | 0.000148859890909803 | 0.00863387367276858 | 0.00726018415314478 | 9709/1051/3576 | 3 |
| cDC2A_AREG | R-HSA-6783783 | Interleukin-10 signaling | 3/35 | 47/10704 | 0.000470572874258382 | 0.0183925557385234 | 0.0154662144443724 | 7850/3553/3576 | 3 |
| cDC2A_AREG | R-HSA-449147 | Signaling by Interleukins | 7/35 | 461/10704 | 0.000618201011363809 | 0.0183925557385234 | 0.0154662144443724 | 7850/1326/1052/7422/3726/3553/3576 | 7 |
| cDC2A_AREG | R-HSA-383280 | Nuclear Receptor transcription pathway | 3/35 | 52/10704 | 0.000634226059949083 | 0.0183925557385234 | 0.0154662144443724 | 8013/4929/3164 | 3 |
| cDC2A_AREG | R-HSA-179812 | GRB2 events in EGFR signaling | 2/35 | 13/10704 | 0.000792079103963135 | 0.0196888234413694 | 0.0165562399174249 | 374/1839 | 2 |
| cDC2A_AREG | R-HSA-180336 | SHC1 events in EGFR signaling | 2/35 | 14/10704 | 0.000922196576733824 | 0.0200577755439607 | 0.0168664900218423 | 374/1839 | 2 |
| cDC2A_AREG | R-HSA-180292 | GAB1 signalosome | 2/35 | 17/10704 | 0.00136976643267048 | 0.0264821510316293 | 0.0222687174434149 | 374/1839 | 2 |
| cDC2A_AREG | R-HSA-9634638 | Estrogen-dependent nuclear events downstream of ESR-membrane signaling | 2/35 | 24/10704 | 0.00274019925824087 | 0.0470176494481974 | 0.0395369224035053 | 374/1839 | 2 |
| cDC2A_AREG | R-HSA-1643713 | Signaling by EGFR in Cancer | 2/35 | 25/10704 | 0.00297238013752972 | 0.0470176494481974 | 0.0395369224035053 | 374/1839 | 2 |
| cDC2_CD14 | R-HSA-6798695 | Neutrophil degranulation | 20/79 | 480/10704 | 1.65383332997614e-10 | 1.02041516459528e-07 | 8.82624735050422e-08 | 2219/6280/929/6279/1520/4048/1536/968/1075/4069/10577/2207/7305/1512/963/55313/5269/2512/5689/5795 | 20 |
| cDC2_CD14 | R-HSA-166786 | Creation of C4 and C2 activators | 4/79 | 14/10704 | 2.6008486798848e-06 | 0.00080236181774446 | 0.000694015937211364 | 2219/712/714/713 | 4 |
| cDC2_CD14 | R-HSA-6807004 | Negative regulation of MET activity | 4/79 | 21/10704 | 1.49515303792732e-05 | 0.00307503141467052 | 0.00265979856220755 | 30011/7316/5795/2885 | 4 |
| cDC2_CD14 | R-HSA-166663 | Initial triggering of complement | 4/79 | 23/10704 | 2.18745683604681e-05 | 0.00322107781020726 | 0.00278612377339432 | 2219/712/714/713 | 4 |
| cDC2_CD14 | R-HSA-5668599 | RHO GTPases Activate NADPH Oxidases | 4/79 | 24/10704 | 2.61027375219389e-05 | 0.00322107781020726 | 0.00278612377339432 | 6280/6279/1536/4689 | 4 |
| cDC2_CD14 | R-HSA-5621480 | Dectin-2 family | 4/79 | 27/10704 | 4.23927643025752e-05 | 0.00435938926244815 | 0.00377072482480801 | 26253/10462/2207/6850 | 4 |
| cDC2_CD14 | R-HSA-1236975 | Antigen processing-Cross presentation | 6/79 | 98/10704 | 8.1853545550622e-05 | 0.00651430672857627 | 0.00563465582425688 | 929/1520/1536/4689/5689/7316 | 6 |
| cDC2_CD14 | R-HSA-2871796 | FCERI mediated MAPK activation | 4/79 | 32/10704 | 8.44642687659808e-05 | 0.00651430672857627 | 0.00563465582425688 | 2353/2207/6850/2885 | 4 |
| cDC2_CD14 | R-HSA-3299685 | Detoxification of Reactive Oxygen Species | 4/79 | 37/10704 | 0.000150848936422256 | 0.00926423082058387 | 0.00801324750667239 | 1536/4689/2876/2878 | 4 |
| cDC2_CD14 | R-HSA-977225 | Amyloid fiber formation | 6/79 | 110/10704 | 0.000155286308929757 | 0.00926423082058387 | 0.00801324750667239 | 6653/4069/7045/9445/7316/9246 | 6 |
| cDC2_CD14 | R-HSA-8875360 | InlB-mediated entry of Listeria monocytogenes into host cell | 3/79 | 15/10704 | 0.00016516456892451 | 0.00926423082058387 | 0.00801324750667239 | 30011/7316/2885 | 3 |
| cDC2_CD14 | R-HSA-2730905 | Role of LAT2/NTAL/LAB on calcium mobilization | 3/79 | 16/10704 | 0.000202200388131107 | 0.0103964699564078 | 0.00899259620898872 | 2207/6850/2885 | 3 |
| cDC2_CD14 | R-HSA-5686938 | Regulation of TLR by endogenous ligand | 3/79 | 19/10704 | 0.000344339758917174 | 0.0163428947116843 | 0.0141360532608103 | 6280/929/6279 | 3 |
| cDC2_CD14 | R-HSA-8876384 | Listeria monocytogenes entry into host cells | 3/79 | 20/10704 | 0.000402957652076261 | 0.0177589193807895 | 0.0153608668874184 | 30011/7316/2885 | 3 |
| cDC2_CD14 | R-HSA-2454202 | Fc epsilon receptor (FCERI) signaling | 6/79 | 134/10704 | 0.000452447088957932 | 0.0186106569258029 | 0.0160975911650296 | 2353/2207/6850/5689/7316/2885 | 6 |
| cDC2_CD14 | R-HSA-912631 | Regulation of signaling by CBL | 3/79 | 22/10704 | 0.000538591671491702 | 0.0207694413318988 | 0.0179648669372561 | 6850/7316/2885 | 3 |
| cDC2_CD14 | R-HSA-5621481 | C-type lectin receptors (CLRs) | 6/79 | 142/10704 | 0.000615523353783604 | 0.0223398770167343 | 0.0193232408896772 | 26253/10462/2207/6850/5689/7316 | 6 |
| cDC2_CD14 | R-HSA-76002 | Platelet activation, signaling and aggregation | 8/79 | 263/10704 | 0.000689731625706556 | 0.0236424673922747 | 0.0204499376744575 | 388/2162/2207/6850/2/2771/7076/2885 | 8 |
| cDC2_CD14 | R-HSA-166658 | Complement cascade | 4/79 | 58/10704 | 0.000861836110638125 | 0.0279869936980907 | 0.0242078065425778 | 2219/712/714/713 | 4 |
| cDC2_CD14 | R-HSA-168898 | Toll-like Receptor Cascades | 6/79 | 155/10704 | 0.000974060901009308 | 0.0300497787961371 | 0.0259920461479852 | 6280/929/6279/1520/2353/7316 | 6 |
| cDC2_CD14 | R-HSA-2424491 | DAP12 signaling | 3/79 | 30/10704 | 0.00136091080380167 | 0.0391709403106321 | 0.0338815435255318 | 7305/6850/2885 | 3 |
| cDC2_CD14 | R-HSA-983705 | Signaling by the B Cell Receptor (BCR) | 5/79 | 112/10704 | 0.00139669479227537 | 0.0391709403106321 | 0.0338815435255318 | 6850/30011/5689/7316/2885 | 5 |
| cDC2_CD14 | R-HSA-182971 | EGFR downregulation | 3/79 | 31/10704 | 0.00149875969348796 | 0.0402058578644379 | 0.0347767123385993 | 30011/7316/2885 | 3 |
| cDC2_CD14 | R-HSA-983695 | Antigen activates B Cell Receptor (BCR) leading to generation of second messengers | 3/79 | 32/10704 | 0.00164506611673506 | 0.0422919080843973 | 0.036581075490556 | 6850/30011/2885 | 3 |
| cDC2_CD14 | R-HSA-2871809 | FCERI mediated Ca+2 mobilization | 3/79 | 33/10704 | 0.00180001492840186 | 0.0444243684329578 | 0.0384255818399891 | 2207/6850/2885 | 3 |
| cDC2_CD14 | R-HSA-449147 | Signaling by Interleukins | 10/79 | 461/10704 | 0.00199434771600647 | 0.0473274054144613 | 0.0409366110127645 | 2162/2353/920/6850/3587/5689/7316/942/7076/2885 | 10 |
| cDC2_CD14 | R-HSA-114604 | GPVI-mediated activation cascade | 3/79 | 35/10704 | 0.00213655269119239 | 0.0488241855728038 | 0.0422312754165513 | 388/2207/6850 | 3 |
| cDC2_CXCL8 | R-HSA-449147 | Signaling by Interleukins | 11/62 | 461/10704 | 5.82655878516046e-05 | 0.0266273736481833 | 0.024594211293151 | 3576/2920/84868/6648/3553/3320/6364/5717/6776/1846/4215 | 11 |
| cDC2_CXCL8 | R-HSA-6783783 | Interleukin-10 signaling | 4/62 | 47/10704 | 0.00015104910503285 | 0.0300205171992638 | 0.0277282676423006 | 3576/2920/3553/6364 | 4 |
| cDC2_CXCL8 | R-HSA-9660826 | Purinergic signaling in leishmaniasis infection | 3/62 | 24/10704 | 0.000343441510874101 | 0.0300205171992638 | 0.0277282676423006 | 3326/3553/114548 | 3 |
| cDC2_CXCL8 | R-HSA-9664424 | Cell recruitment (pro-inflammatory response) | 3/62 | 24/10704 | 0.000343441510874101 | 0.0300205171992638 | 0.0277282676423006 | 3326/3553/114548 | 3 |
| cDC2_CXCL8 | R-HSA-380108 | Chemokine receptors bind chemokines | 4/62 | 59/10704 | 0.000365941550490541 | 0.0300205171992638 | 0.0277282676423006 | 3576/2920/2921/6364 | 4 |
| cDC2_CXCL8 | R-HSA-6798695 | Neutrophil degranulation | 10/62 | 480/10704 | 0.000394142457758387 | 0.0300205171992638 | 0.0277282676423006 | 2896/9588/2720/1794/3326/3320/8826/976/5717/79169 | 10 |
| cDC2_CD207 | R-HSA-6798695 | Neutrophil degranulation | 21/78 | 480/10704 | 1.59509276367136e-11 | 6.12515621249802e-09 | 5.42331539648262e-09 | 240/1509/5315/2934/5777/10487/3689/101/8560/5265/5788/6556/23601/203068/226/5660/2992/3608/2212/10159/1508 | 21 |
| cDC2_CD207 | R-HSA-449147 | Signaling by Interleukins | 13/78 | 461/10704 | 2.51975436299604e-05 | 0.00483792837695239 | 0.00428358241709327 | 6285/116379/240/3055/5777/3689/920/1436/6888/375/3091/829/821 | 13 |
| cDC2_CD207 | R-HSA-2029481 | FCGR activation | 3/78 | 12/10704 | 7.81138402284969e-05 | 0.00999857154924761 | 0.00885290189256299 | 3055/2214/2212 | 3 |
| cDC2_CD207 | R-HSA-2132295 | MHC class II antigen presentation | 6/78 | 123/10704 | 0.000266370550669177 | 0.020273664970251 | 0.0179506408590764 | 3120/1509/375/829/821/1508 | 6 |
| cDC2_CD207 | R-HSA-416700 | Other semaphorin interactions | 3/78 | 19/10704 | 0.000331635418796496 | 0.020273664970251 | 0.0179506408590764 | 5788/54209/10154 | 3 |
| cDC2_CD207 | R-HSA-114608 | Platelet degranulation | 6/78 | 129/10704 | 0.000344465470722767 | 0.020273664970251 | 0.0179506408590764 | 10487/5265/226/5660/334/308 | 6 |
| cDC2_CD207 | R-HSA-198933 | Immunoregulatory interactions between a Lymphoid and a non-Lymphoid cell | 6/78 | 132/10704 | 0.000389709783655481 | 0.020273664970251 | 0.0179506408590764 | 909/3689/2213/2214/54209/29992 | 6 |
| cDC2_CD207 | R-HSA-76005 | Response to elevated platelet cytosolic Ca2+ | 6/78 | 134/10704 | 0.000422368020213562 | 0.020273664970251 | 0.0179506408590764 | 10487/5265/226/5660/334/308 | 6 |
| cDC2_CD207 | R-HSA-202427 | Phosphorylation of CD3 and TCR zeta chains | 3/78 | 22/10704 | 0.000518828895361945 | 0.022136699535443 | 0.0196002027136735 | 3120/920/5788 | 3 |
| cDC2_CD207 | R-HSA-389948 | PD-1 signaling | 3/78 | 23/10704 | 0.00059353374321705 | 0.0227916957395347 | 0.0201801472693797 | 3120/5777/920 | 3 |
| cDC2_CD207 | R-HSA-447115 | Interleukin-12 family signaling | 4/78 | 57/10704 | 0.000769054821763758 | 0.0268470046870257 | 0.0237707853999707 | 6888/375/829/821 | 4 |
| cDC2_CD207 | R-HSA-2142753 | Arachidonic acid metabolism | 4/78 | 59/10704 | 0.000876403594518938 | 0.028044915024606 | 0.0248314351780366 | 241/240/5742/6916 | 4 |
| cDC2_CD207 | R-HSA-164952 | The role of Nef in HIV-1 replication and disease pathogenesis | 3/78 | 28/10704 | 0.00106954182151668 | 0.0315926199586465 | 0.0279726322550515 | 3055/920/375 | 3 |
| cDC1_CLEC9A | R-HSA-1236974 | ER-Phagosome pathway | 9/69 | 83/10704 | 2.52695961116622e-09 | 1.16998229996996e-06 | 1.01078384446649e-06 | 6890/5698/5720/6892/3107/3106/811/3105/4615 | 9 |
| cDC1_CLEC9A | R-HSA-983170 | Antigen Presentation: Folding, assembly and peptide loading of class I MHC | 6/69 | 25/10704 | 9.24428268906261e-09 | 1.73029014083566e-06 | 1.49485109359452e-06 | 6890/6892/3107/3106/811/3105 | 6 |
| cDC1_CLEC9A | R-HSA-1236975 | Antigen processing-Cross presentation | 9/69 | 98/10704 | 1.12113832019589e-08 | 1.73029014083566e-06 | 1.49485109359452e-06 | 6890/5698/5720/6892/3107/3106/811/3105/4615 | 9 |
| cDC1_CLEC9A | R-HSA-6798695 | Neutrophil degranulation | 15/69 | 480/10704 | 2.90523812770659e-07 | 3.36281313282038e-05 | 2.9052381277066e-05 | 8895/2517/1727/966/290/1265/54509/2950/10875/3107/3106/3312/2760/3105/1471 | 15 |
| cDC1_CLEC9A | R-HSA-449147 | Signaling by Interleukins | 14/69 | 461/10704 | 1.08770845588883e-06 | 0.000100721803015305 | 8.70166764711061e-05 | 595/1846/6285/5698/867/5608/5720/604/1265/3603/3936/7534/3312/4615 | 14 |
| cDC1_CLEC9A | R-HSA-1236977 | Endosomal/Vacuolar pathway | 3/69 | 11/10704 | 4.07666266897377e-05 | 0.00314582469289143 | 0.00271777511264918 | 3107/3106/3105 | 3 |
| cDC1_CLEC9A | R-HSA-983169 | Class I MHC mediated antigen processing & presentation | 10/69 | 370/10704 | 0.000118846733941036 | 0.00786086254495707 | 0.00679124193948775 | 6890/51676/5698/5720/6892/3107/3106/811/3105/4615 | 10 |
| cDC1_CLEC9A | R-HSA-877300 | Interferon gamma signaling | 5/69 | 91/10704 | 0.000291043204702804 | 0.0168441254721748 | 0.0145521602351402 | 3394/817/3107/3106/3105 | 5 |
| cDC1_CLEC9A | R-HSA-9020702 | Interleukin-1 signaling | 5/69 | 103/10704 | 0.000516183220851292 | 0.0265547590282387 | 0.0229414764822797 | 6285/5698/5608/5720/4615 | 5 |
| cDC1_CLEC9A | R-HSA-909733 | Interferon alpha/beta signaling | 4/69 | 70/10704 | 0.00105272876791962 | 0.0487413419546786 | 0.0421091507167849 | 3394/3107/3106/3105 | 4 |
| cDC2_FCGR3A | R-HSA-913531 | Interferon Signaling | 17/72 | 199/10704 | 1.05421910072718e-14 | 3.76356218959603e-12 | 2.1861175036132e-12 | 2633/2537/115361/2209/2634/6772/115362/3437/9636/3429/9246/684/4599/10410/5696/3105/8651 | 17 |
| cDC2_FCGR3A | R-HSA-909733 | Interferon alpha/beta signaling | 12/72 | 70/10704 | 2.57523670732334e-14 | 4.59679752257215e-12 | 2.67011384917209e-12 | 2537/2634/6772/3437/9636/3429/684/4599/10410/5696/3105/8651 | 12 |
| cDC2_FCGR3A | R-HSA-877300 | Interferon gamma signaling | 8/72 | 91/10704 | 1.53230812422823e-07 | 1.8234466678316e-05 | 1.05917438762443e-05 | 2633/115361/2209/2634/6772/115362/3105/8651 | 8 |
| cDC2_FCGR3A | R-HSA-449147 | Signaling by Interleukins | 15/72 | 461/10704 | 3.11901965425728e-07 | 2.78372504142462e-05 | 1.61696545233864e-05 | 3627/6772/5721/3553/6648/5685/5698/5696/9235/7124/9446/8651/6347/3557/4792 | 15 |
| cDC2_FCGR3A | R-HSA-1236975 | Antigen processing-Cross presentation | 7/72 | 98/10704 | 3.93562361817293e-06 | 0.000281003526337547 | 0.000163224811111593 | 2209/1536/5721/5685/5698/5696/3105 | 7 |
| cDC2_FCGR3A | R-HSA-9020702 | Interleukin-1 signaling | 7/72 | 103/10704 | 5.48839010357638e-06 | 0.000326559211162795 | 0.000189686464983254 | 5721/3553/5685/5698/5696/3557/4792 | 7 |
| cDC2_FCGR3A | R-HSA-6783783 | Interleukin-10 signaling | 5/72 | 47/10704 | 1.47310985475825e-05 | 0.00075128602592671 | 0.000436394949454701 | 3627/3553/7124/6347/3557 | 5 |
| cDC2_FCGR3A | R-HSA-1236978 | Cross-presentation of soluble exogenous antigens (endosomes) | 5/72 | 49/10704 | 1.81226225346e-05 | 0.000808722030606523 | 0.000469757452541604 | 2209/5721/5685/5698/5696 | 5 |
| cDC2_FCGR3A | R-HSA-446652 | Interleukin-1 family signaling | 7/72 | 140/10704 | 4.08494511360808e-05 | 0.00162036156173121 | 0.000941209575883968 | 5721/3553/5685/5698/5696/3557/4792 | 7 |
| cDC2_FCGR3A | R-HSA-5607764 | CLEC7A (Dectin-1) signaling | 6/72 | 101/10704 | 5.74038478092887e-05 | 0.00196902273804484 | 0.00114373427508428 | 5721/3553/5685/5698/5696/4792 | 6 |
| cDC2_FCGR3A | R-HSA-5668541 | TNFR2 non-canonical NF-kB pathway | 6/72 | 102/10704 | 6.06701683991407e-05 | 0.00196902273804484 | 0.00114373427508428 | 10673/5721/5685/5698/5696/7124 | 6 |
| cDC2_FCGR3A | R-HSA-1169091 | Activation of NF-kappaB in B cells | 5/72 | 67/10704 | 8.35657944490245e-05 | 0.00248608238485848 | 0.00144407557074191 | 5721/5685/5698/5696/4792 | 5 |
| cDC2_FCGR3A | R-HSA-166786 | Creation of C4 and C2 activators | 3/72 | 14/10704 | 0.000100726568706694 | 0.00276610654063766 | 0.00160673150082742 | 713/712/714 | 3 |
| cDC2_FCGR3A | R-HSA-8939902 | Regulation of RUNX2 expression and activity | 5/72 | 73/10704 | 0.000125970939042995 | 0.00321225894559638 | 0.00186588533770452 | 6772/5721/5685/5698/5696 | 5 |
| cDC2_FCGR3A | R-HSA-202403 | TCR signaling | 6/72 | 120/10704 | 0.000149652779209189 | 0.00356173614517869 | 0.00206888403538317 | 5721/5685/5698/5696/3937/4792 | 6 |
| cDC2_FCGR3A | R-HSA-2871837 | FCERI mediated NF-kB activation | 5/72 | 82/10704 | 0.000218365232870376 | 0.00456545212888156 | 0.00265190644077744 | 5721/5685/5698/5696/4792 | 5 |
| cDC2_FCGR3A | R-HSA-1168372 | Downstream signaling events of B Cell Receptor (BCR) | 5/72 | 83/10704 | 0.000231157686306207 | 0.00456545212888156 | 0.00265190644077744 | 5721/5685/5698/5696/4792 | 5 |
| cDC2_FCGR3A | R-HSA-1236974 | ER-Phagosome pathway | 5/72 | 83/10704 | 0.000231157686306207 | 0.00456545212888156 | 0.00265190644077744 | 5721/5685/5698/5696/3105 | 5 |
| cDC2_FCGR3A | R-HSA-977606 | Regulation of Complement cascade | 4/72 | 47/10704 | 0.000269773753332008 | 0.00456545212888156 | 0.00265190644077744 | 713/710/712/714 | 4 |
| cDC2_FCGR3A | R-HSA-2454202 | Fc epsilon receptor (FCERI) signaling | 6/72 | 134/10704 | 0.000273135973095435 | 0.00456545212888156 | 0.00265190644077744 | 5721/5685/5698/5696/3937/4792 | 6 |
| cDC2_FCGR3A | R-HSA-211733 | Regulation of activated PAK-2p34 by proteasome mediated degradation | 4/72 | 50/10704 | 0.000343071494874433 | 0.00456545212888156 | 0.00265190644077744 | 5721/5685/5698/5696 | 4 |
| cDC2_FCGR3A | R-HSA-350562 | Regulation of ornithine decarboxylase (ODC) | 4/72 | 51/10704 | 0.000370388784874095 | 0.00456545212888156 | 0.00265190644077744 | 5721/5685/5698/5696 | 4 |
| cDC2_FCGR3A | R-HSA-5621481 | C-type lectin receptors (CLRs) | 6/72 | 142/10704 | 0.000373222316563327 | 0.00456545212888156 | 0.00265190644077744 | 5721/3553/5685/5698/5696/4792 | 6 |
| cDC2_FCGR3A | R-HSA-180534 | Vpu mediated degradation of CD4 | 4/72 | 52/10704 | 0.000399228136165921 | 0.00456545212888156 | 0.00265190644077744 | 5721/5685/5698/5696 | 4 |
| cDC2_FCGR3A | R-HSA-349425 | Autodegradation of the E3 ubiquitin ligase COP1 | 4/72 | 52/10704 | 0.000399228136165921 | 0.00456545212888156 | 0.00265190644077744 | 5721/5685/5698/5696 | 4 |
| cDC2_FCGR3A | R-HSA-69601 | Ubiquitin Mediated Degradation of Phosphorylated Cdc25A | 4/72 | 52/10704 | 0.000399228136165921 | 0.00456545212888156 | 0.00265190644077744 | 5721/5685/5698/5696 | 4 |
| cDC2_FCGR3A | R-HSA-69610 | p53-Independent DNA Damage Response | 4/72 | 52/10704 | 0.000399228136165921 | 0.00456545212888156 | 0.00265190644077744 | 5721/5685/5698/5696 | 4 |
| cDC2_FCGR3A | R-HSA-69613 | p53-Independent G1/S DNA damage checkpoint | 4/72 | 52/10704 | 0.000399228136165921 | 0.00456545212888156 | 0.00265190644077744 | 5721/5685/5698/5696 | 4 |
| cDC2_FCGR3A | R-HSA-75815 | Ubiquitin-dependent degradation of Cyclin D | 4/72 | 52/10704 | 0.000399228136165921 | 0.00456545212888156 | 0.00265190644077744 | 5721/5685/5698/5696 | 4 |
| cDC2_FCGR3A | R-HSA-169911 | Regulation of Apoptosis | 4/72 | 53/10704 | 0.00042963778634521 | 0.00456545212888156 | 0.00265190644077744 | 5721/5685/5698/5696 | 4 |
| cDC2_FCGR3A | R-HSA-180585 | Vif-mediated degradation of APOBEC3G | 4/72 | 54/10704 | 0.000461666220062068 | 0.00456545212888156 | 0.00265190644077744 | 5721/5685/5698/5696 | 4 |
| cDC2_FCGR3A | R-HSA-8854050 | FBXL7 down-regulates AURKA during mitotic entry and in early mitosis | 4/72 | 54/10704 | 0.000461666220062068 | 0.00456545212888156 | 0.00265190644077744 | 5721/5685/5698/5696 | 4 |
| cDC2_FCGR3A | R-HSA-9604323 | Negative regulation of NOTCH4 signaling | 4/72 | 54/10704 | 0.000461666220062068 | 0.00456545212888156 | 0.00265190644077744 | 5721/5685/5698/5696 | 4 |
| cDC2_FCGR3A | R-HSA-166663 | Initial triggering of complement | 3/72 | 23/10704 | 0.000469229301370622 | 0.00456545212888156 | 0.00265190644077744 | 713/712/714 | 3 |
| cDC2_FCGR3A | R-HSA-174113 | SCF-beta-TrCP mediated degradation of Emi1 | 4/72 | 55/10704 | 0.000495362149035569 | 0.00456545212888156 | 0.00265190644077744 | 5721/5685/5698/5696 | 4 |
| cDC2_FCGR3A | R-HSA-4641257 | Degradation of AXIN | 4/72 | 55/10704 | 0.000495362149035569 | 0.00456545212888156 | 0.00265190644077744 | 5721/5685/5698/5696 | 4 |
| cDC2_FCGR3A | R-HSA-8941858 | Regulation of RUNX3 expression and activity | 4/72 | 55/10704 | 0.000495362149035569 | 0.00456545212888156 | 0.00265190644077744 | 5721/5685/5698/5696 | 4 |
| cDC2_FCGR3A | R-HSA-202424 | Downstream TCR signaling | 5/72 | 98/10704 | 0.000500125633193441 | 0.00456545212888156 | 0.00265190644077744 | 5721/5685/5698/5696/4792 | 5 |
| cDC2_FCGR3A | R-HSA-450408 | AUF1 (hnRNP D0) binds and destabilizes mRNA | 4/72 | 56/10704 | 0.000530774492458524 | 0.00456545212888156 | 0.00265190644077744 | 5721/5685/5698/5696 | 4 |
| cDC2_FCGR3A | R-HSA-5362768 | Hh mutants that don't undergo autocatalytic processing are degraded by ERAD | 4/72 | 56/10704 | 0.000530774492458524 | 0.00456545212888156 | 0.00265190644077744 | 5721/5685/5698/5696 | 4 |
| cDC2_FCGR3A | R-HSA-447115 | Interleukin-12 family signaling | 4/72 | 57/10704 | 0.000567952357787529 | 0.00456545212888156 | 0.00265190644077744 | 6772/5721/6648/9446 | 4 |
| cDC2_FCGR3A | R-HSA-4641258 | Degradation of DVL | 4/72 | 57/10704 | 0.000567952357787529 | 0.00456545212888156 | 0.00265190644077744 | 5721/5685/5698/5696 | 4 |
| cDC2_FCGR3A | R-HSA-69541 | Stabilization of p53 | 4/72 | 57/10704 | 0.000567952357787529 | 0.00456545212888156 | 0.00265190644077744 | 5721/5685/5698/5696 | 4 |
| cDC2_FCGR3A | R-HSA-166658 | Complement cascade | 4/72 | 58/10704 | 0.000606945021913026 | 0.00456545212888156 | 0.00265190644077744 | 713/710/712/714 | 4 |
| cDC2_FCGR3A | R-HSA-351202 | Metabolism of polyamines | 4/72 | 59/10704 | 0.000647801912704159 | 0.00456545212888156 | 0.00265190644077744 | 5721/5685/5698/5696 | 4 |
| cDC2_FCGR3A | R-HSA-380108 | Chemokine receptors bind chemokines | 4/72 | 59/10704 | 0.000647801912704159 | 0.00456545212888156 | 0.00265190644077744 | 3627/4283/6373/6347 | 4 |
| cDC2_FCGR3A | R-HSA-5387390 | Hh mutants abrogate ligand secretion | 4/72 | 59/10704 | 0.000647801912704159 | 0.00456545212888156 | 0.00265190644077744 | 5721/5685/5698/5696 | 4 |
| cDC2_FCGR3A | R-HSA-5676590 | NIK-->noncanonical NF-kB signaling | 4/72 | 59/10704 | 0.000647801912704159 | 0.00456545212888156 | 0.00265190644077744 | 5721/5685/5698/5696 | 4 |
| cDC2_FCGR3A | R-HSA-68827 | CDT1 association with the CDC6:ORC:origin complex | 4/72 | 59/10704 | 0.000647801912704159 | 0.00456545212888156 | 0.00265190644077744 | 5721/5685/5698/5696 | 4 |
| cDC2_FCGR3A | R-HSA-187577 | SCF(Skp2)-mediated degradation of p27/p21 | 4/72 | 60/10704 | 0.000690572590923261 | 0.00456545212888156 | 0.00265190644077744 | 5721/5685/5698/5696 | 4 |
| cDC2_FCGR3A | R-HSA-5607761 | Dectin-1 mediated noncanonical NF-kB signaling | 4/72 | 60/10704 | 0.000690572590923261 | 0.00456545212888156 | 0.00265190644077744 | 5721/5685/5698/5696 | 4 |
| cDC2_FCGR3A | R-HSA-5610780 | Degradation of GLI1 by the proteasome | 4/72 | 60/10704 | 0.000690572590923261 | 0.00456545212888156 | 0.00265190644077744 | 5721/5685/5698/5696 | 4 |
| cDC2_FCGR3A | R-HSA-5610783 | Degradation of GLI2 by the proteasome | 4/72 | 60/10704 | 0.000690572590923261 | 0.00456545212888156 | 0.00265190644077744 | 5721/5685/5698/5696 | 4 |
| cDC2_FCGR3A | R-HSA-5610785 | GLI3 is processed to GLI3R by the proteasome | 4/72 | 60/10704 | 0.000690572590923261 | 0.00456545212888156 | 0.00265190644077744 | 5721/5685/5698/5696 | 4 |
| cDC2_FCGR3A | R-HSA-5678895 | Defective CFTR causes cystic fibrosis | 4/72 | 61/10704 | 0.000735306732504901 | 0.00477280915462272 | 0.00277235265652566 | 5721/5685/5698/5696 | 4 |
| cDC2_FCGR3A | R-HSA-6785807 | Interleukin-4 and Interleukin-13 signaling | 5/72 | 108/10704 | 0.000779380861626469 | 0.00496855299286874 | 0.00288605319060929 | 6772/3553/7124/8651/6347 | 5 |
| cDC2_FCGR3A | R-HSA-983169 | Class I MHC mediated antigen processing & presentation | 9/72 | 370/10704 | 0.000793982251138867 | 0.00497283620450133 | 0.00288854115372774 | 2209/1536/9246/5721/5685/5698/5696/3105/8651 | 9 |
| cDC2_FCGR3A | R-HSA-174084 | Autodegradation of Cdh1 by Cdh1:APC/C | 4/72 | 64/10704 | 0.000881788062240629 | 0.00533556505457465 | 0.00309923725711693 | 5721/5685/5698/5696 | 4 |
| cDC2_FCGR3A | R-HSA-4608870 | Asymmetric localization of PCP proteins | 4/72 | 64/10704 | 0.000881788062240629 | 0.00533556505457465 | 0.00309923725711693 | 5721/5685/5698/5696 | 4 |
| cDC2_FCGR3A | R-HSA-983705 | Signaling by the B Cell Receptor (BCR) | 5/72 | 112/10704 | 0.000918725015106218 | 0.005466413839882 | 0.00317524259606886 | 5721/5685/5698/5696/4792 | 5 |
| cDC2_FCGR3A | R-HSA-5358346 | Hedgehog ligand biogenesis | 4/72 | 65/10704 | 0.000934874519822841 | 0.00547131481273367 | 0.00317808939439344 | 5721/5685/5698/5696 | 4 |
| cDC2_FCGR3A | R-HSA-1234176 | Oxygen-dependent proline hydroxylation of Hypoxia-inducible Factor Alpha | 4/72 | 66/10704 | 0.000990173952678829 | 0.00552331407978659 | 0.00320829389272581 | 5721/5685/5698/5696 | 4 |
| cDC2_FCGR3A | R-HSA-69563 | p53-Dependent G1 DNA Damage Response | 4/72 | 66/10704 | 0.000990173952678829 | 0.00552331407978659 | 0.00320829389272581 | 5721/5685/5698/5696 | 4 |
| cDC2_FCGR3A | R-HSA-69580 | p53-Dependent G1/S DNA damage checkpoint | 4/72 | 66/10704 | 0.000990173952678829 | 0.00552331407978659 | 0.00320829389272581 | 5721/5685/5698/5696 | 4 |
| cDC2_FCGR3A | R-HSA-174154 | APC/C:Cdc20 mediated degradation of Securin | 4/72 | 68/10704 | 0.00110761180358895 | 0.00581496196884196 | 0.00337770163013966 | 5721/5685/5698/5696 | 4 |
| cDC2_FCGR3A | R-HSA-5658442 | Regulation of RAS by GAPs | 4/72 | 68/10704 | 0.00110761180358895 | 0.00581496196884196 | 0.00337770163013966 | 5721/5685/5698/5696 | 4 |
| cDC2_FCGR3A | R-HSA-68867 | Assembly of the pre-replicative complex | 4/72 | 68/10704 | 0.00110761180358895 | 0.00581496196884196 | 0.00337770163013966 | 5721/5685/5698/5696 | 4 |
| cDC2_FCGR3A | R-HSA-69615 | G1/S DNA Damage Checkpoints | 4/72 | 68/10704 | 0.00110761180358895 | 0.00581496196884196 | 0.00337770163013966 | 5721/5685/5698/5696 | 4 |
| cDC2_FCGR3A | R-HSA-8948751 | Regulation of PTEN stability and activity | 4/72 | 69/10704 | 0.00116985023864248 | 0.00605270340862846 | 0.00351579705587441 | 5721/5685/5698/5696 | 4 |
| cDC2_FCGR3A | R-HSA-8878166 | Transcriptional regulation by RUNX2 | 5/72 | 121/10704 | 0.00129932658128899 | 0.00654474512539597 | 0.00380160633850216 | 6772/5721/5685/5698/5696 | 5 |
| cDC2_FCGR3A | R-HSA-68949 | Orc1 removal from chromatin | 4/72 | 71/10704 | 0.00130161597731965 | 0.00654474512539597 | 0.00380160633850216 | 5721/5685/5698/5696 | 4 |

| | | | | | | | | | |
|---|---|---|---|---|---|---|---|---|---|
| cDC2_FCGR3A | R-HSA-1169408 | ISG15 antiviral mechanism | 4/72 | 72/10704 | 0.00137124308319619 | 0.00679908028751445 | 0.00394934045891301 | 6772/9636/9246/4599 | 4 |
| cDC2_FCGR3A | R-HSA-174184 | Cdc20:Phospho-APC/C mediated degradation of Cyclin A | 4/72 | 73/10704 | 0.00144343277271232 | 0.00696358783592295 | 0.00404489695909427 | 5721/5685/5698/5696 | 4 |
| cDC2_FCGR3A | R-HSA-69017 | CDK-mediated phosphorylation and removal of Cdc6 | 4/72 | 73/10704 | 0.00144343277271232 | 0.00696358783592295 | 0.00404489695909427 | 5721/5685/5698/5696 | 4 |
| cDC2_FCGR3A | R-HSA-936440 | Negative regulators of DDX58/IFIH1 signaling | 3/72 | 34/10704 | 0.00150365205417048 | 0.00703908844840458 | 0.0040887525411638 | 9636/9246/7128 | 3 |
| cDC2_FCGR3A | R-HSA-174178 | APC/C:Cdh1 mediated degradation of Cdc20 and other APC/C:Cdh1 targeted proteins in late mitosis/early G1 | 4/72 | 74/10704 | 0.00151823476338138 | 0.00703908844840458 | 0.0040887525411638 | 5721/5685/5698/5696 | 4 |
| cDC2_FCGR3A | R-HSA-179419 | APC:Cdc20 mediated degradation of cell cycle proteins prior to satisfation of the cell cycle checkpoint | 4/72 | 74/10704 | 0.00151823476338138 | 0.00703908844840458 | 0.0040887525411638 | 5721/5685/5698/5696 | 4 |
| cDC2_FCGR3A | R-HSA-1234174 | Cellular response to hypoxia | 4/72 | 75/10704 | 0.00159569867535394 | 0.00730339009104305 | 0.00424227583056312 | 5721/5685/5698/5696 | 4 |
| cDC2_FCGR3A | R-HSA-176409 | APC/C:Cdc20 mediated degradation of mitotic proteins | 4/72 | 76/10704 | 0.00167587401859161 | 0.00757325347642031 | 0.00439902973567684 | 5721/5685/5698/5696 | 4 |
| cDC2_FCGR3A | R-HSA-176814 | Activation of APC/C and APC/C:Cdc20 mediated degradation of mitotic proteins | 4/72 | 77/10704 | 0.00175881018033137 | 0.00758920014796435 | 0.0044082925818929 | 5721/5685/5698/5696 | 4 |
| cDC2_FCGR3A | R-HSA-5619084 | ABC transporter disorders | 4/72 | 77/10704 | 0.00175881018033137 | 0.00758920014796435 | 0.0044082925818929 | 5721/5685/5698/5696 | 4 |
| cDC2_FCGR3A | R-HSA-8852276 | The role of GTSE1 in G2/M progression after G2 checkpoint | 4/72 | 77/10704 | 0.00175881018033137 | 0.00758920014796435 | 0.0044082925818929 | 5721/5685/5698/5696 | 4 |
| cDC2_FCGR3A | R-HSA-1222556 | ROS and RNS production in phagocytes | 3/72 | 36/10704 | 0.00177695888912922 | 0.00758920014796435 | 0.0044082925818929 | 1536/526/9296 | 3 |
| cDC2_FCGR3A | R-HSA-162909 | Host Interactions of HIV factors | 5/72 | 130/10704 | 0.0017856941524622 | 0.00758920014796435 | 0.0044082925818929 | 5721/5685/5698/5696/3105 | 5 |
| cDC2_FCGR3A | R-HSA-168928 | DDX58/IFIH1-mediated induction of interferon-alpha/beta | 4/72 | 78/10704 | 0.00184455641283626 | 0.00774713693391227 | 0.00450003236320424 | 9636/9246/7128/4792 | 4 |
| cDC2_FCGR3A | R-HSA-3299685 | Detoxification of Reactive Oxygen Species | 3/72 | 37/10704 | 0.00192447699377533 | 0.00798881728811389 | 0.00464041576222449 | 1536/5052/6648 | 3 |
| cDC2_FCGR3A | R-HSA-1169410 | Antiviral mechanism by IFN-stimulated genes | 4/72 | 80/10704 | 0.00202467535279899 | 0.00830815058562345 | 0.0048259049546449 | 6772/9636/9246/4599 | 4 |
| cDC2_FCGR3A | R-HSA-8950505 | Gene and protein expression by JAK-STAT signaling after Interleukin-12 stimulation | 3/72 | 38/10704 | 0.00207941594917171 | 0.00843581243016252 | 0.00490005911467497 | 5721/6648/9446 | 3 |
| cDC2_FCGR3A | R-HSA-176408 | Regulation of APC/C activators between G1/S and early anaphase | 4/72 | 81/10704 | 0.00211914578359523 | 0.00850039376116291 | 0.0049375720800504 | 5721/5685/5698/5696 | 4 |
| cDC2_FCGR3A | R-HSA-9013694 | Signaling by NOTCH4 | 4/72 | 82/10704 | 0.00221662170927343 | 0.0087925994467846 | 0.00510730382136685 | 5721/5685/5698/5696 | 4 |
| cDC2_FCGR3A | R-HSA-195253 | Degradation of beta-catenin by the destruction complex | 4/72 | 83/10704 | 0.00231715153322114 | 0.00899155540608639 | 0.00522287016069297 | 5721/5685/5698/5696 | 4 |
| cDC2_FCGR3A | R-HSA-69202 | Cyclin E associated events during G1/S transition | 4/72 | 83/10704 | 0.00231715153322114 | 0.00899155540608639 | 0.00522287016069297 | 5721/5685/5698/5696 | 4 |
| cDC2_FCGR3A | R-HSA-5632684 | Hedgehog 'on' state | 4/72 | 85/10704 | 0.00252756546568908 | 0.00949832496053687 | 0.00551723431291689 | 5721/5685/5698/5696 | 4 |
| cDC2_FCGR3A | R-HSA-69002 | DNA Replication Pre-Initiation | 4/72 | 85/10704 | 0.00252756546568908 | 0.00949832496053687 | 0.00551723431291689 | 5721/5685/5698/5696 | 4 |
| cDC2_FCGR3A | R-HSA-69656 | Cyclin A:Cdk2-associated events at S phase entry | 4/72 | 85/10704 | 0.00252756546568908 | 0.00949832496053687 | 0.00551723431291689 | 5721/5685/5698/5696 | 4 |
| cDC2_FCGR3A | R-HSA-174143 | APC/C-mediated degradation of cell cycle proteins | 4/72 | 88/10704 | 0.00286728848460856 | 0.0104451223367883 | 0.00606719475260888 | 5721/5685/5698/5696 | 4 |
| cDC2_FCGR3A | R-HSA-450531 | Regulation of mRNA stability by proteins that bind AU-rich elements | 4/72 | 88/10704 | 0.00286728848460856 | 0.0104451223367883 | 0.00606719475260888 | 5721/5685/5698/5696 | 4 |
| cDC2_FCGR3A | R-HSA-453276 | Regulation of mitotic cell cycle | 4/72 | 88/10704 | 0.00286728848460856 | 0.0104451223367883 | 0.00606719475260888 | 5721/5685/5698/5696 | 4 |
| cDC2_FCGR3A | R-HSA-5687128 | MAPK6/MAPK4 signaling | 4/72 | 89/10704 | 0.00298714610798801 | 0.0107718299045628 | 0.00625696739259583 | 5721/5685/5698/5696 | 4 |
| cDC2_FCGR3A | R-HSA-69052 | Switching of origins to a post-replicative state | 4/72 | 91/10704 | 0.00323706731359999 | 0.011556330309552 | 0.00671265537662314 | 5721/5685/5698/5696 | 4 |
| cDC2_FCGR3A | R-HSA-5357786 | TNFR1-induced proapoptotic signaling | 2/72 | 13/10704 | 0.00331747006116666 | 0.0117261070478861 | 0.00681127255914363 | 7124/7128 | 2 |
| cDC2_FCGR3A | R-HSA-4086400 | PCP/CE pathway | 4/72 | 92/10704 | 0.00336722362892722 | 0.0117852827012453 | 0.00684564556138971 | 5721/5685/5698/5696 | 4 |
| cDC2_FCGR3A | R-HSA-9020591 | Interleukin-12 signaling | 3/72 | 47/10704 | 0.00382793801092395 | 0.0132282024003641 | 0.00768378556058299 | 5721/6648/9446 | 3 |
| cDC2_FCGR3A | R-HSA-877312 | Regulation of IFNG signaling | 2/72 | 14/10704 | 0.00385359397657665 | 0.0132282024003641 | 0.00768378556058299 | 6772/8651 | 2 |
| cDC2_FCGR3A | R-HSA-8878159 | Transcriptional regulation by RUNX3 | 4/72 | 96/10704 | 0.00392355539144641 | 0.0133400883309178 | 0.00774877606130269 | 5721/5685/5698/5696 | 4 |
| cDC2_FCGR3A | R-HSA-9648895 | Response of EIF2AK1 (HRI) to heme deficiency | 2/72 | 15/10704 | 0.00442717751074343 | 0.0149103997295793 | 0.00866091330304325 | 22809/467 | 2 |
| cDC2_FCGR3A | R-HSA-6798695 | Neutrophil degranulation | 9/72 | 480/10704 | 0.00469050587121397 | 0.0156496317385363 | 0.00909030650889475 | 5265/1536/684/10577/1075/51316/967/3105/6282 | 9 |
| cDC2_FCGR3A | R-HSA-5689603 | UCH proteinases | 4/72 | 102/10704 | 0.00487009830803968 | 0.0160983805182423 | 0.00935096848619705 | 5721/5685/5698/5696 | 4 |
| cDC2_FCGR3A | R-HSA-157118 | Signaling by NOTCH | 6/72 | 236/10704 | 0.00500780202252915 | 0.0163619244666206 | 0.00950405165833482 | 4192/6772/5721/5685/5698/5696 | 6 |
| cDC2_FCGR3A | R-HSA-382556 | ABC-family proteins mediated transport | 4/72 | 103/10704 | 0.00504148933145172 | 0.0163619244666206 | 0.00950405165833482 | 5721/5685/5698/5696 | 4 |
| cDC2_FCGR3A | R-HSA-109581 | Apoptosis | 5/72 | 178/10704 | 0.00685035197182726 | 0.0220322130985795 | 0.0127977177662396 | 8743/5721/5685/5698/5696 | 5 |
| cDC2_FCGR3A | R-HSA-5610787 | Hedgehog 'off' state | 4/72 | 113/10704 | 0.00698270388538481 | 0.0222573686346641 | 0.0129285024945565 | 5721/5685/5698/5696 | 4 |
| cDC2_FCGR3A | R-HSA-69239 | Synthesis of DNA | 4/72 | 120/10704 | 0.00860157486847058 | 0.0271748869738407 | 0.0157849114959358 | 5721/5685/5698/5696 | 4 |
| cDC2_FCGR3A | R-HSA-5357801 | Programmed Cell Death | 5/72 | 190/10704 | 0.00895606758319897 | 0.0280466326947547 | 0.0162912771365669 | 8743/5721/5685/5698/5696 | 5 |
| cDC2_FCGR3A | R-HSA-8862803 | Deregulated CDK5 triggers multiple neurodegenerative pathways in Alzheimer's disease models | 2/72 | 22/10704 | 0.00944859102165811 | 0.0290788534028616 | 0.0168908569080458 | 5052/6648 | 2 |
| cDC2_FCGR3A | R-HSA-8863678 | Neurodegenerative Diseases | 2/72 | 22/10704 | 0.00944859102165811 | 0.0290788534028616 | 0.0168908569080458 | 5052/6648 | 2 |
| cDC2_FCGR3A | R-HSA-69306 | DNA Replication | 4/72 | 128/10704 | 0.0107323263825338 | 0.0327473548595261 | 0.0190217570612609 | 5721/5685/5698/5696 | 4 |
| cDC2_FCGR3A | R-HSA-114608 | Platelet degranulation | 4/72 | 129/10704 | 0.0110205311489017 | 0.0333417764420161 | 0.0193670351144838 | 710/5265/5341/967 | 4 |
| cDC2_FCGR3A | R-HSA-982772 | Growth hormone receptor signaling | 2/72 | 24/10704 | 0.0111919580076918 | 0.0335758740230755 | 0.0195030139541379 | 6772/8651 | 2 |
| cDC2_FCGR3A | R-HSA-8939236 | RUNX1 regulates transcription of genes involved in differentiation of HSCs | 4/72 | 130/10704 | 0.0113137005773601 | 0.0336582592176462 | 0.0195508685415784 | 5721/5685/5698/5696 | 4 |
| cDC2_FCGR3A | R-HSA-69206 | G1/S Transition | 4/72 | 131/10704 | 0.0116118653985155 | 0.0342598012171076 | 0.0199002825881474 | 5721/5685/5698/5696 | 4 |
| cDC2_FCGR3A | R-HSA-76005 | Response to elevated platelet cytosolic Ca2+ | 4/72 | 134/10704 | 0.0125366324640259 | 0.0366850638496496 | 0.0213090301588706 | 710/5265/5341/967 | 4 |
| cDC2_FCGR3A | R-HSA-77387 | Insulin receptor recycling | 2/72 | 26/10704 | 0.0130654991387194 | 0.0376159934880874 | 0.021849773602103 | 526/9296 | 2 |
| cDC2_FCGR3A | R-HSA-912694 | Regulation of IFNA signaling | 2/72 | 26/10704 | 0.0130654991387194 | 0.0376159934880874 | 0.021849773602103 | 6772/8651 | 2 |
| cDC2_FCGR3A | R-HSA-380994 | ATF4 activates genes in response to endoplasmic reticulum stress | 2/72 | 27/10704 | 0.014049924566459 | 0.0401265845618069 | 0.0233080853860415 | 467/6347 | 2 |
| cDC2_FCGR3A | R-HSA-6807070 | PTEN Regulation | 4/72 | 140/10704 | 0.0145249781142498 | 0.0411541046570412 | 0.0239049347410795 | 5721/5685/5698/5696 | 4 |
| cDC2_FCGR3A | R-HSA-5688426 | Deubiquitination | 6/72 | 297/10704 | 0.0146500682001991 | 0.0411816877753629 | 0.0239209567794383 | 5721/5685/5698/5696/7128/4792 | 6 |
| cDC2_FCGR3A | R-HSA-73887 | Death Receptor Signalling | 4/72 | 141/10704 | 0.0148746828916637 | 0.0414864202525308 | 0.0240979648820539 | 8743/7124/7128/4792 | 4 |
| cDC2_FCGR3A | R-HSA-5689880 | Ub-specific processing proteases | 5/72 | 220/10704 | 0.0160958949839945 | 0.0442454942066268 | 0.0257006114070632 | 5721/5685/5698/5696/4792 | 5 |
| cDC2_FCGR3A | R-HSA-9645723 | Diseases of programmed cell death | 2/72 | 29/10704 | 0.0161118046130574 | 0.0442454942066268 | 0.0257006114070632 | 5052/6648 | 2 |
| cDC2_FCGR3A | R-HSA-3858494 | Beta-catenin independent WNT signaling | 4/72 | 146/10704 | 0.0167032309390428 | 0.0455194919483839 | 0.0264406307351662 | 5721/5685/5698/5696 | 4 |
| cDC2_FCGR3A | R-HSA-5357956 | TNFR1-induced NFkappaB signaling pathway | 2/72 | 30/10704 | 0.0171883606629671 | 0.0464867027021156 | 0.027002448569414 | 7124/7128 | 2 |
| cDC2_FCGR3A | R-HSA-983168 | Antigen processing: Ubiquitination & Proteasome degradation | 6/72 | 309/10704 | 0.0174872449006469 | 0.0469394468385784 | 0.027265431305322 | 9246/5721/5685/5698/5696/8651 | 6 |
| cDC2_FCGR3A | R-HSA-453279 | Mitotic G1 phase and G1/S transition | 4/72 | 149/10704 | 0.0178652356675803 | 0.0472436232098235 | 0.0274421163860688 | 5721/5685/5698/5696 | 4 |
| cDC2_FCGR3A | R-HSA-5358351 | Signaling by Hedgehog | 4/72 | 149/10704 | 0.0178652356675803 | 0.0472436232098235 | 0.0274421163860688 | 5721/5685/5698/5696 | 4 |
| cDC2_FCGR3A | R-HSA-917977 | Transferrin endocytosis and recycling | 2/72 | 31/10704 | 0.0182947358401533 | 0.0480236815804024 | 0.0278952241525557 | 526/9296 | 2 |
| cDC_LAMP3 | R-HSA-380108 | Chemokine receptors bind chemokines | 5/66 | 59/10704 | 2.95838487157303e-05 | 0.0100585085633483 | 0.00952911337580364 | 1236/6363/6367/6361/4283 | 5 |
| cDC_LAMP3 | R-HSA-449147 | Signaling by Interleukins | 10/66 | 461/10704 | 0.000480590808555777 | 0.0496100919710178 | 0.046999034498859 | 6624/6363/9235/3575/10148/6367/1846/1026/57162/1438 | 10 |
| cDC_LAMP3 | R-HSA-5213460 | RIPK1-mediated regulated necrosis | 3/66 | 29/10704 | 0.00072956017604438 | 0.0496100919710178 | 0.046999034498859 | 330/8837/57162 | 3 |
| cDC_LAMP3 | R-HSA-5218859 | Regulated Necrosis | 3/66 | 29/10704 | 0.00072956017604438 | 0.0496100919710178 | 0.046999034498859 | 330/8837/57162 | 3 |
| cDC_LAMP3 | R-HSA-5675482 | Regulation of necroptotic cell death | 3/66 | 29/10704 | 0.00072956017604438 | 0.0496100919710178 | 0.046999034498859 | 330/8837/57162 | 3 |

**Supplementary Table 5: Enrichment pathway analysis**

**Monocytes**

| Cluster | ID | Description | GeneRatio | BgRatio | pvalue | p.adjust | qvalue | geneID | Count |
|---|---|---|---|---|---|---|---|---|---|
| Mono_CD14_FOS- | R-HSA-6798695 | Neutrophil degranulation | 19/69 | 480/10704 | 9.75755412854525e-11 | 3.46393171563356e-08 | 3.2867550748784e-08 | 6283/6279/6402/6280/948/3684/4332/83716/4069/683/5873/1509/6036/150372/2219/11322/945/23218/64333 | 19 |
| Mono_CD14_FOS- | R-HSA-5686938 | Regulation of TLR by endogenous ligand | 3/69 | 19/10704 | 0.000230717197364171 | 0.0409523025321404 | 0.0388576332402815 | 6279/6280/948 | 3 |
| Mono_CD14_FOS- | R-HSA-6803157 | Antimicrobial peptides | 5/69 | 95/10704 | 0.000355482742992541 | 0.042065457920784 | 0.0399138518447765 | 6279/6280/729230/4069/6039 | 5 |
| Mono_CD14_FOS+ | R-HSA-6798695 | Neutrophil degranulation | 8/17 | 480/10704 | 2.62353281108147e-07 | 1.9676496083111e-05 | 1.46365514723493e-05 | 6283/6279/4257/4332/6280/6402/25797/4069 | 8 |
| Mono_CD14_FOS+ | R-HSA-5686938 | Regulation of TLR by endogenous ligand | 2/17 | 19/10704 | 0.000399589483249287 | 0.010520293052906 | 0.00782562149900375 | 6279/6280 | 2 |
| Mono_CD14_FOS+ | R-HSA-6803157 | Antimicrobial peptides | 3/17 | 95/10704 | 0.00042081172211624 | 0.010520293052906 | 0.00782562149900375 | 6279/6280/4069 | 3 |
| Mono_CD14_FOS+ | R-HSA-5668599 | RHO GTPases Activate NADPH Oxidases | 2/17 | 24/10704 | 0.000641946542725846 | 0.0120364976761096 | 0.00895346493801838 | 6279/6280 | 2 |
| Mono_CD14_FOS+ | R-HSA-168898 | Toll-like Receptor Cascades | 3/17 | 155/10704 | 0.00174458850479225 | 0.0261688275718838 | 0.0194659348955767 | 6283/6279/6280 | 3 |
| Mono_CD16 | R-HSA-114604 | GPVI-mediated activation cascade | 5/72 | 35/10704 | 3.3190586134693e-06 | 0.00140728085211098 | 0.00118088611721329 | 5777/4067/5880/391/2207 | 5 |
| Mono_CD16 | R-HSA-6798695 | Neutrophil degranulation | 13/72 | 480/10704 | 1.57958229537202e-05 | 0.00265516240269154 | 0.00222801611745218 | 5777/5175/1075/51411/11314/7133/81619/391/719/23406/2207/5660/51382 | 13 |
| Mono_CD16 | R-HSA-202733 | Cell surface interactions at the vascular wall | 7/72 | 137/10704 | 3.55466660874656e-05 | 0.00265516240269154 | 0.00222801611745218 | 6693/5777/5175/6404/4067/2207/483 | 7 |
| Mono_CD16 | R-HSA-9664407 | Parasite infection | 5/72 | 59/10704 | 4.51627735349458e-05 | 0.00265516240269154 | 0.00222801611745218 | 2214/4067/10109/71/60 | 5 |
| Mono_CD16 | R-HSA-9664417 | Leishmania phagocytosis | 5/72 | 59/10704 | 4.51627735349458e-05 | 0.00265516240269154 | 0.00222801611745218 | 2214/4067/10109/71/60 | 5 |
| Mono_CD16 | R-HSA-9664422 | FCGR3A-mediated phagocytosis | 5/72 | 59/10704 | 4.51627735349458e-05 | 0.00265516240269154 | 0.00222801611745218 | 2214/4067/10109/71/60 | 5 |
| Mono_CD16 | R-HSA-194315 | Signaling by Rho GTPases | 12/72 | 455/10704 | 4.53377956551813e-05 | 0.00265516240269154 | 0.00222801611745218 | 389/51466/7846/5880/114793/397/391/103910/10109/392/71/60 | 12 |
| Mono_CD16 | R-HSA-210990 | PECAM1 interactions | 3/72 | 12/10704 | 6.14703865309476e-05 | 0.00265516240269154 | 0.00222801611745218 | 5777/5175/4067 | 3 |
| Mono_CD16 | R-HSA-195258 | RHO GTPase Effectors | 10/72 | 327/10704 | 6.15242239528858e-05 | 0.00265516240269154 | 0.00222801611745218 | 389/51466/7846/5880/114793/391/103910/10109/71/60 | 10 |
| Mono_CD16 | R-HSA-76002 | Platelet activation, signaling and aggregation | 9/72 | 263/10704 | 6.26217547804608e-05 | 0.00265516240269154 | 0.00222801611745218 | 5777/5175/6647/4067/5880/391/2207/5660/2771 | 9 |
| Mono_CD16 | R-HSA-3928662 | EPHB-mediated forward signaling | 4/72 | 42/10704 | 0.000173636915813873 | 0.00669291384591657 | 0.0056161988081425 | 4067/10109/71/60 | 4 |
| Mono_CD16 | R-HSA-2029480 | Fcgamma receptor (FCGR) dependent phagocytosis | 5/72 | 86/10704 | 0.000272978321568639 | 0.00964523402875858 | 0.00809356777984211 | 2214/4067/10109/71/60 | 5 |
| Mono_CD16 | R-HSA-5663220 | RHO GTPases Activate Formins | 6/72 | 140/10704 | 0.000345870692077237 | 0.0112807056492883 | 0.0094659347305349 | 389/51466/7846/114793/71/60 | 6 |
| Mono_CD16 | R-HSA-2682334 | EPH-Ephrin signaling | 5/72 | 92/10704 | 0.000373628376400878 | 0.0113156022567123 | 0.00949521738522533 | 4067/103910/10109/71/60 | 5 |
| Mono_CD16 | R-HSA-400685 | Sema4D in semaphorin signaling | 3/72 | 25/10704 | 0.000603539470844386 | 0.0170600490425347 | 0.0143155327119581 | 389/6237/103910 | 3 |
| Mono_CD16 | R-HSA-2029482 | Regulation of actin dynamics for phagocytic cup formation | 4/72 | 61/10704 | 0.000735306732504901 | 0.0194856284113799 | 0.0163508997096485 | 2214/10109/71/60 | 4 |
| Mono_CD16 | R-HSA-909733 | Interferon alpha/beta signaling | 4/72 | 70/10704 | 0.00123450165332746 | 0.029642144181882 | 0.0248734973522247 | 4938/10410/5777/10581 | 4 |
| Mono_CD16 | R-HSA-5626467 | RHO GTPases activate IQGAPs | 3/72 | 32/10704 | 0.00125839291338178 | 0.029642144181882 | 0.0248734973522247 | 7846/71/60 | 3 |
| Mono_CD16 | R-HSA-9658195 | Leishmania infection | 7/72 | 252/10704 | 0.00147596005855963 | 0.0317509106819577 | 0.026643018397472 | 2214/4067/719/10109/71/2771/60 | 7 |
| Mono_CD16 | R-HSA-418594 | G alpha (i) signalling events | 9/72 | 405/10704 | 0.00149768446613008 | 0.0317509106819577 | 0.026643018397472 | 10287/8698/1524/58191/5499/719/7852/5660/2771 | 9 |
| Mono_CD16 | R-HSA-5663213 | RHO GTPases Activate WASPs and WAVEs | 3/72 | 36/10704 | 0.00177695888912922 | 0.0358776461424186 | 0.0301058699010364 | 10109/71/60 | 3 |
| Mono_CD16 | R-HSA-194840 | Rho GTPase cycle | 5/72 | 138/10704 | 0.00231998195680359 | 0.0435002321716309 | 0.0365021809185979 | 389/5880/397/391/392 | 5 |
| Mono_CD16 | R-HSA-196025 | Formation of annular gap junctions | 2/72 | 11/10704 | 0.00235968240553658 | 0.0435002321716309 | 0.0365021809185979 | 71/60 | 2 |
| Mono_CD16 | R-HSA-190873 | Gap junction degradation | 2/72 | 12/10704 | 0.0028193248093377 | 0.0478157487663674 | 0.0401234436023639 | 71/60 | 2 |
| Mono_CD16 | R-HSA-2029481 | FCGR activation | 2/72 | 12/10704 | 0.0028193248093377 | 0.0478157487663674 | 0.0401234436023639 | 2214/4067 | 2 |
| Mono_IL1B | R-HSA-6783783 | Interleukin-10 signaling | 10/84 | 47/10704 | 2.08883229931862e-12 | 7.54068460054024e-10 | 5.60686564553947e-10 | 3553/3576/2920/6364/3557/5743/3383/7076/3552/7850 | 10 |
| Mono_IL1B | R-HSA-449147 | Signaling by Interleukins | 18/84 | 461/10704 | 1.27669866324425e-08 | 2.30444108715588e-06 | 1.71346399540676e-06 | 3553/3576/2920/6364/3557/6648/1026/3091/5743/3383/4790/5055/7076/3552/4170/4792/7850/8767 | 18 |
| Mono_IL1B | R-HSA-6785807 | Interleukin-4 and Interleukin-13 signaling | 9/84 | 108/10704 | 1.51770810648381e-07 | 1.82630875480218e-05 | 1.35794935843288e-05 | 3553/3576/1026/3091/5743/3383/7076/3552/4170 | 9 |
| Mono_IL1B | R-HSA-6798695 | Neutrophil degranulation | 16/84 | 480/10704 | 7.92041898603663e-07 | 7.14817813489806e-05 | 5.31501800378774e-05 | 6515/4973/5329/4860/7130/290/4790/23601/1604/2204/960/1654/7097/2683/1992/2495 | 16 |
| Mono_IL1B | R-HSA-9020702 | Interleukin-1 signaling | 7/84 | 103/10704 | 1.53570490614845e-05 | 0.00110877894223918 | 0.000824431054879693 | 3553/3557/4790/3552/4792/7850/8767 | 7 |
| Mono_IL1B | R-HSA-5260271 | Diseases of Immune System | 4/84 | 24/10704 | 3.32708002798951e-05 | 0.00133452876678246 | 0.000992287025891608 | 4790/4792/7097/7099 | 4 |
| Mono_IL1B | R-HSA-5602358 | Diseases associated with the TLR signaling cascade | 4/84 | 24/10704 | 3.32708002798951e-05 | 0.00133452876678246 | 0.000992287025891608 | 4790/4792/7097/7099 | 4 |
| Mono_IL1B | R-HSA-9660826 | Purinergic signaling in leishmaniasis infection | 4/84 | 24/10704 | 3.32708002798951e-05 | 0.00133452876678246 | 0.000992287025891608 | 3553/114548/4790/3552 | 4 |
| Mono_IL1B | R-HSA-9664424 | Cell recruitment (pro-inflammatory response) | 4/84 | 24/10704 | 3.32708002798951e-05 | 0.00133452876678246 | 0.000992287025891608 | 3553/114548/4790/3552 | 4 |
| Mono_IL1B | R-HSA-446652 | Interleukin-1 family signaling | 7/84 | 140/10704 | 0.000110257770737095 | 0.00379014351325458 | 0.00281815598740318 | 3553/3557/4790/3552/4792/7850/8767 | 7 |
| Mono_IL1B | R-HSA-179812 | GRB2 events in EGFR signaling | 3/84 | 13/10704 | 0.000125988150025083 | 0.00379014351325458 | 0.00281815598740318 | 2069/374/1839 | 3 |
| Mono_IL1B | R-HSA-75205 | Dissolution of Fibrin Clot | 3/84 | 13/10704 | 0.000125988150025083 | 0.00379014351325458 | 0.00281815598740318 | 5329/5055/5271 | 3 |
| Mono_IL1B | R-HSA-180336 | SHC1 events in EGFR signaling | 3/84 | 14/10704 | 0.000159441060211576 | 0.00442755559510606 | 0.00329210286266816 | 2069/374/1839 | 3 |
| Mono_IL1B | R-HSA-193639 | p75NTR signals via NF-kB | 3/84 | 16/10704 | 0.000242526605718912 | 0.00625372176175195 | 0.00464994619987388 | 4790/4792/8767 | 3 |
| Mono_IL1B | R-HSA-180292 | GAB1 signalosome | 3/84 | 17/10704 | 0.000292831454731205 | 0.007047477010531 | 0.00524014182150577 | 2069/374/1839 | 3 |
| Mono_IL1B | R-HSA-9634638 | Estrogen-dependent nuclear events downstream of ESR-membrane signaling | 3/84 | 24/10704 | 0.000837728634706182 | 0.0166099377986271 | 0.0123502963657965 | 2069/374/1839 | 3 |
| Mono_IL1B | R-HSA-166058 | MyD88:MAL(TIRAP) cascade initiated on plasma membrane | 5/84 | 95/10704 | 0.000879604679542438 | 0.0166099377986271 | 0.0123502963657965 | 4790/4792/7097/8767/7099 | 5 |
| Mono_IL1B | R-HSA-168188 | Toll Like Receptor TLR6:TLR2 Cascade | 5/84 | 95/10704 | 0.000879604679542438 | 0.0166099377986271 | 0.0123502963657965 | 4790/4792/7097/8767/7099 | 5 |
| Mono_IL1B | R-HSA-1643713 | Signaling by EGFR in Cancer | 3/84 | 25/10704 | 0.000946595445062209 | 0.0166099377986271 | 0.0123502963657965 | 2069/374/1839 | 3 |
| Mono_IL1B | R-HSA-168643 | Nucleotide-binding domain, leucine rich repeat containing receptor (NLR) signaling pathways | 4/84 | 56/10704 | 0.000950308136250262 | 0.0166099377986271 | 0.0123502963657965 | 114548/7128/4790/8767 | 4 |
| Mono_IL1B | R-HSA-168179 | Toll Like Receptor TLR1:TLR2 Cascade | 5/84 | 98/10704 | 0.00101223997664763 | 0.0166099377986271 | 0.0123502963657965 | 4790/4792/7097/8767/7099 | 5 |
| Mono_IL1B | R-HSA-181438 | Toll Like Receptor 2 (TLR2) Cascade | 5/84 | 98/10704 | 0.00101223997664763 | 0.0166099377986271 | 0.0123502963657965 | 4790/4792/7097/8767/7099 | 5 |
| Mono_IL1B | R-HSA-380108 | Chemokine receptors bind chemokines | 4/84 | 59/10704 | 0.00115677004776218 | 0.0174363433125928 | 0.0129647690471239 | 3576/2920/6364/2921 | 4 |
| Mono_IL1B | R-HSA-2219528 | PI3K/AKT Signaling in Cancer | 5/84 | 101/10704 | 0.00115920287950755 | 0.0174363433125928 | 0.0129647690471239 | 2069/374/1026/1839/3164 | 5 |
| Mono_IL1B | R-HSA-182971 | EGFR downregulation | 3/84 | 31/10704 | 0.00178832012358107 | 0.0258233425845107 | 0.0192009108005546 | 2069/374/1839 | 3 |
| Mono_IL1B | R-HSA-445989 | TAK1 activates NFkB by phosphorylation and activation of IKKs complex | 3/84 | 33/10704 | 0.00214629449123735 | 0.0298004735129494 | 0.0221581010228957 | 4790/4792/8767 | 3 |
| Mono_IL1B | R-HSA-1250342 | PI3K events in ERBB4 signaling | 2/84 | 10/10704 | 0.00262882278559396 | 0.0338930366285507 | 0.0252011206889647 | 2069/1839 | 2 |
| Mono_IL1B | R-HSA-5602498 | MyD88 deficiency (TLR2/4) | 2/84 | 10/10704 | 0.00262882278559396 | 0.0338930366285507 | 0.0252011206889647 | 7097/7099 | 2 |
| Mono_IL1B | R-HSA-5603041 | IRAK4 deficiency (TLR2/4) | 2/84 | 11/10704 | 0.00319666882038548 | 0.0392092305645299 | 0.0291539693656659 | 7097/7099 | 2 |
| Mono_IL1B | R-HSA-114608 | Platelet degranulation | 5/84 | 129/10704 | 0.00339597626820698 | 0.0392092305645299 | 0.0291539693656659 | 7057/7076/5341/3309/5552 | 5 |
| Mono_IL1B | R-HSA-166016 | Toll Like Receptor 4 (TLR4) Cascade | 5/84 | 129/10704 | 0.00339597626820698 | 0.0392092305645299 | 0.0291539693656659 | 4790/4792/7097/8767/7099 | 5 |
| Mono_IL1B | R-HSA-9031628 | NGF-stimulated transcription | 3/84 | 39/10704 | 0.0034756104655539 | 0.0392092305645299 | 0.0291539693656659 | 2354/1958/10221 | 3 |
| Mono_IL1B | R-HSA-76005 | Response to elevated platelet cytosolic Ca2+ | 5/84 | 134/10704 | 0.00399593564081761 | 0.0437131141313684 | 0.0325028257865548 | 7057/7076/5341/3309/5552 | 5 |
| Mono_IL1B | R-HSA-2559583 | Cellular Senescence | 6/84 | 197/10704 | 0.00445522174507886 | 0.0449997862090821 | 0.0334595290372239 | 3576/1026/23135/2114/4790/3552 | 6 |
| Mono_IL1B | R-HSA-209560 | NF-kB is activated and signals survival | 2/84 | 13/10704 | 0.00448751330616885 | 0.0449997862090821 | 0.0334595290372239 | 4790/4792 | 2 |
| Mono_IL1B | R-HSA-8847993 | ERBB2 Activates PTK6 Signaling | 2/84 | 13/10704 | 0.00448751330616885 | 0.0449997862090821 | 0.0334595290372239 | 2069/1839 | 2 |
| Mono_IL1B | R-HSA-1257604 | PIP3 activates AKT signaling | 7/84 | 264/10704 | 0.00461353599657641 | 0.0450131485071374 | 0.033469464555533 | 8870/2069/374/1026/1839/1958/3164 | 7 |
| Mono_IL1B | R-HSA-73887 | Death Receptor Signalling | 5/84 | 141/10704 | 0.00495861466588268 | 0.0471068393258855 | 0.0350262254792267 | 7128/4790/4792/7185/8767 | 5 |
| Mono_IL1B | R-HSA-1250347 | SHC1 events in ERBB4 signaling | 2/84 | 14/10704 | 0.00520885868049507 | 0.0482153329143262 | 0.0358504443596827 | 2069/1839 | 2 |
| Mono_IL1B | R-HSA-975138 | TRAF6 mediated induction of NFkB and MAP kinases upon TLR7/8 or 9 activation | 4/84 | 91/10704 | 0.00562074713156668 | 0.0486814417123121 | 0.0361970189142429 | 4790/4792/8767/7099 | 4 |
| Mono_IL1B | R-HSA-8856828 | Clathrin-mediated endocytosis | 5/84 | 146/10704 | 0.00574012599399567 | 0.0486814417123121 | 0.0361970189142429 | 2069/374/1839/7037/51429 | 5 |
| Mono_IL1B | R-HSA-975155 | MyD88 dependent cascade initiated on endosome | 4/84 | 92/10704 | 0.00584167278809162 | 0.0486814417123121 | 0.0361970189142429 | 4790/4792/8767/7099 | 4 |
| Mono_IL1B | R-HSA-6785631 | ERBB2 Regulates Cell Motility | 2/84 | 15/10704 | 0.0059797332881941 | 0.0486814417123121 | 0.0361970189142429 | 2069/1839 | 2 |
| Mono_IL1B | R-HSA-844456 | The NLRP3 inflammasome | 2/84 | 15/10704 | 0.0059797332881941 | 0.0486814417123121 | 0.0361970189142429 | 114548/4790 | 2 |
| Mono_IL1B | R-HSA-168181 | Toll Like Receptor 7/8 (TLR7/8) Cascade | 4/84 | 93/10704 | 0.00606832375915248 | 0.0486814417123121 | 0.0361970189142429 | 4790/4792/8767/7099 | 4 |
| MonoInter_CLEC10A | R-HSA-202430 | Translocation of ZAP-70 to Immunological synapse | 5/12 | 19/10704 | 7.81212929367748e-12 | 4.66185371344341e-10 | 3.00764755706026e-10 | 3115/3117/3113/3127/3122 | 5 |
| MonoInter_CLEC10A | R-HSA-202427 | Phosphorylation of CD3 and TCR zeta chains | 5/12 | 22/10704 | 1.76632244412953e-11 | 4.66185371344341e-10 | 3.00764755706026e-10 | 3115/3117/3113/3127/3122 | 5 |
| MonoInter_CLEC10A | R-HSA-389948 | PD-1 signaling | 5/12 | 23/10704 | 2.2557356677952e-11 | 4.66185371344341e-10 | 3.00764755706026e-10 | 3115/3117/3113/3127/3122 | 5 |
| MonoInter_CLEC10A | R-HSA-202433 | Generation of second messenger molecules | 5/12 | 34/10704 | 1.85417765756962e-10 | 2.87397536923291e-09 | 1.85417765756962e-09 | 3115/3117/3113/3127/3122 | 5 |
| MonoInter_CLEC10A | R-HSA-2132295 | MHC class II antigen presentation | 6/12 | 123/10704 | 1.77880945322873e-09 | 2.20572372200362e-08 | 1.42304756258298e-08 | 3115/3117/3113/3109/3127/3122 | 6 |
| MonoInter_CLEC10A | R-HSA-388841 | Costimulation by the CD28 family | 5/12 | 69/10704 | 7.34685038231316e-09 | 7.59174539505694e-08 | 4.89790025487544e-08 | 3115/3117/3113/3127/3122 | 5 |
| MonoInter_CLEC10A | R-HSA-877300 | Interferon gamma signaling | 5/12 | 91/10704 | 3.00366206486452e-08 | 2.66038640030857e-07 | 1.71637832277973e-07 | 3115/3117/3113/3127/3122 | 5 |
| MonoInter_CLEC10A | R-HSA-202424 | Downstream TCR signaling | 5/12 | 98/10704 | 4.36946456471948e-08 | 3.3863350376576e-07 | 2.18473228235974e-07 | 3115/3117/3113/3127/3122 | 5 |
| MonoInter_CLEC10A | R-HSA-202403 | TCR signaling | 5/12 | 120/10704 | 1.21147960436785e-07 | 8.34574838564522e-07 | 5.38435379719046e-07 | 3115/3117/3113/3127/3122 | 5 |
| MonoInter_CLEC10A | R-HSA-913531 | Interferon Signaling | 5/12 | 199/10704 | 1.50466172103383e-06 | 9.32890267040977e-06 | 6.01864688413534e-06 | 3115/3117/3113/3127/3122 | 5 |
| MonoInter_CXCL10 | R-HSA-913531 | Interferon Signaling | 19/83 | 199/10704 | 4.97689291239804e-16 | 2.42872374125024e-13 | 1.99075716495921e-13 | 2633/9636/2537/3117/3115/3119/3669/3113/2209/3434/3665/4599/3118/3123/3122/3429/3127/3383/10437 | 19 |
| MonoInter_CXCL10 | R-HSA-877300 | Interferon gamma signaling | 13/83 | 91/10704 | 1.63089810248188e-13 | 3.9793913700558e-11 | 3.26179620496377e-11 | 2633/3117/3115/3119/3113/2209/3665/3118/3123/3122/3127/3383/10437 | 13 |
| MonoInter_CXCL10 | R-HSA-202430 | Translocation of ZAP-70 to Immunological synapse | 8/83 | 19/10704 | 6.52843373282268e-13 | 1.06195855387249e-10 | 8.70457831043024e-11 | 3117/3115/3119/3113/3118/3123/3122/3127 | 8 |
| MonoInter_CXCL10 | R-HSA-202427 | Phosphorylation of CD3 and TCR zeta chains | 8/83 | 22/10704 | 2.71070172043356e-12 | 3.30705609892895e-10 | 2.71070172043356e-10 | 3117/3115/3119/3113/3118/3123/3122/3127 | 8 |
| MonoInter_CXCL10 | R-HSA-389948 | PD-1 signaling | 8/83 | 23/10704 | 4.13050116391758e-12 | 4.03136913598355e-10 | 3.30440093113406e-10 | 3117/3115/3119/3113/3118/3123/3122/3127 | 8 |
| MonoInter_CXCL10 | R-HSA-202433 | Generation of second messenger molecules | 8/83 | 34/10704 | 1.42784691651855e-10 | 1.16131549210176e-08 | 9.51897944345702e-09 | 3117/3115/3119/3113/3118/3123/3122/3127 | 8 |
| MonoInter_CXCL10 | R-HSA-2132295 | MHC class II antigen presentation | 11/83 | 123/10704 | 2.35676595394916e-09 | 1.54144536281139e-07 | 1.26347980558311e-07 | 3117/3115/3119/3113/3118/3123/1508/3122/3127/972/10437 | 11 |
| MonoInter_CXCL10 | R-HSA-388841 | Costimulation by the CD28 family | 9/83 | 69/10704 | 2.52695961116622e-09 | 1.54144536281139e-07 | 1.26347980558311e-07 | 3117/3115/3119/3113/3118/3123/3122/3127/6714 | 9 |
| MonoInter_CXCL10 | R-HSA-202424 | Downstream TCR signaling | 10/83 | 98/10704 | 3.63256823622445e-09 | 1.96965922141948e-07 | 1.61447477165531e-07 | 5721/3117/3115/3119/3113/5698/3118/3123/3122/3127 | 10 |
| MonoInter_CXCL10 | R-HSA-202403 | TCR signaling | 10/83 | 120/10704 | 2.62401916018234e-08 | 1.28052135016898e-06 | 1.04960766407294e-06 | 5721/3117/3115/3119/3113/5698/3118/3123/3122/3127 | 10 |
| MonoInter_CXCL10 | R-HSA-909733 | Interferon alpha/beta signaling | 7/83 | 70/10704 | 1.05420424017704e-06 | 4.67683335642176e-05 | 3.83346996428013e-05 | 9636/2537/3669/3434/3665/4599/3429 | 7 |
| MonoInter_CXCL10 | R-HSA-6783783 | Interleukin-10 signaling | 5/83 | 47/10704 | 2.94881695992066e-05 | 0.00110694052033945 | 0.000907328295360204 | 3627/6347/7076/1230/3383 | 5 |
| MonoInter_CXCL10 | R-HSA-977606 | Regulation of Complement cascade | 5/83 | 47/10704 | 2.94881695992066e-05 | 0.00110694052033945 | 0.000907328295360204 | 710/712/713/714/975 | 5 |
| MonoInter_CXCL10 | R-HSA-166658 | Complement cascade | 5/83 | 58/10704 | 8.24010396523491e-05 | 0.00287226481073903 | 0.00235431541863855 | 710/712/713/714/975 | 5 |
| MonoInter_CXCL10 | R-HSA-166786 | Creation of C4 and C2 activators | 3/83 | 14/10704 | 0.000153865575180494 | 0.00500576004587206 | 0.00410308200481316 | 712/713/714 | 3 |
| MonoInter_CXCL10 | R-HSA-180292 | GAB1 signalosome | 3/83 | 17/10704 | 0.000282650711478717 | 0.00862084670010088 | 0.00706626778696793 | 374/1839/6714 | 3 |
| MonoInter_CXCL10 | R-HSA-9009391 | Extra-nuclear estrogen signaling | 5/83 | 77/10704 | 0.000316697342042799 | 0.00909107664216977 | 0.00745170216571292 | 3315/374/1839/8877/6714 | 5 |
| MonoInter_CXCL10 | R-HSA-198933 | Immunoregulatory interactions between a Lymphoid and a non-Lymphoid cell | 6/83 | 132/10704 | 0.000544661504329713 | 0.0147663785618278 | 0.0121035889851047 | 2209/10870/10859/975/3383/342510 | 6 |
| MonoInter_CXCL10 | R-HSA-72695 | Formation of the ternary complex, and subsequently, the 43S complex | 4/83 | 51/10704 | 0.000636593296725389 | 0.0163503962527363 | 0.0134019641415871 | 6222/6231/3921/8668 | 4 |
| MonoInter_CXCL10 | R-HSA-166663 | Initial triggering of complement | 3/83 | 23/10704 | 0.000711839995822981 | 0.0173688958980807 | 0.0142367999164596 | 712/713/714 | 3 |
| MonoInter_CXCL10 | R-HSA-449147 | Signaling by Interleukins | 11/83 | 461/10704 | 0.000820441093390119 | 0.0190654882654466 | 0.015627449397907 | 3627/5721/5698/6347/7076/9446/2335/1230/3725/3383/6810 | 11 |
| MonoInter_CXCL10 | R-HSA-5683057 | MAPK family signaling cascades | 9/83 | 325/10704 | 0.000894841054002276 | 0.019849201561505 | 0.0162698373454959 | 5721/5698/3315/374/8682/1839/2335/3725/6714 | 9 |
| MonoInter_CXCL10 | R-HSA-1236975 | Antigen processing-Cross presentation | 5/83 | 98/10704 | 0.000958855397344346 | 0.0199935000008564 | 0.0163881147548003 | 5721/2209/5698/6892/6810 | 5 |
| MonoInter_CXCL10 | R-HSA-72649 | Translation initiation complex formation | 4/83 | 58/10704 | 0.00103726818710725 | 0.0199935000008564 | 0.0163881147548003 | 6222/6231/3921/8668 | 4 |
| MonoInter_CXCL10 | R-HSA-72702 | Ribosomal scanning and start codon recognition | 4/83 | 58/10704 | 0.00103726818710725 | 0.0199935000008564 | 0.0163881147548003 | 6222/6231/3921/8668 | 4 |
| MonoInter_CXCL10 | R-HSA-380108 | Chemokine receptors bind chemokines | 4/83 | 59/10704 | 0.00110619774594902 | 0.0199935000008564 | 0.0163881147548003 | 3627/6373/6347/1230 | 4 |
| MonoInter_CXCL10 | R-HSA-72662 | Activation of the mRNA upon binding of the cap-binding complex and eIFs, and subsequent binding to 43S | 4/83 | 59/10704 | 0.00110619774594902 | 0.0199935000008564 | 0.0163881147548003 | 6222/6231/3921/8668 | 4 |
| MonoInter_CXCL10 | R-HSA-8939211 | ESR-mediated signaling | 7/83 | 223/10704 | 0.001681966921247 | 0.0293142806274476 | 0.0240280988749571 | 3315/374/3326/1839/3725/8877/6714 | 7 |

**Supplementary Table 5: Enrichment pathway analysis - Association of phenotypic and metabolic characteristics of the identified macrophage clusters**

| Cell Type | General metabolic pathways (Reactome) | Enriched Pathways (Reactome) | p-value |
|---|---|---|---|
| | | Selenoamino acid metabolism | 5.65805090948131e-34 |
| Mac_Rec | Metabolism of amino acids and derivatives | Selenocysteine synthesis | 3.91554740927724e-37 |
| | | Glucose metabolism | 5.27675817788096e-08 |
| | | Glycolysis | 7.24076945323556e-09 |
| Mac_Hypo | Metabolism of carbohydrates | Gluconeogenesis | 3.85359018665144e-08 |
| | | Glucose metabolism | 0.00051488663927673 |
| | Metabolism of carbohydrates | Glycolysis | 0.000161437749161864 |
| | | Selenoamino acid metabolism | 2.18876424393645e-12 |
| Mac_Prolif | Metabolism of amino acids and derivatives | Selenocysteine synthesis | 9.41528555610786e-14 |
| RTM_LA | Metabolism of lipids | Glycosphingolipid metabolism | 0.000468751543951872 |
| | Metabolism of lipids | Glycosphingolipid metabolism | 0.00611910749258343 |
| | Metabolism of vitamins and cofactors | Metabolism of fat-soluble vitamins | 4.10834001069416e-05 |
| Mac_LA | Metabolism of carbohydrates | - | 0.00268315964244264 |
