## Supplementary Table 6 for "Identification of novel myeloid-derived cell states with implication in cancer outcome"

**Supplementary Table 6: Summary of datasets used for validation**

**Cheng and colleagues (2021) datasets**

| Study | Link to Download | Paper | Tissue | Sample Type | Technology | Total of Myeloid Cells |
|---|---|---|---|---|---|---|
| Cheng et al., 2021 | GSE154763 | https://doi.org/10.1016/j.cell.2021.01.010 | Esophagus | Normal/Tumor | 10x | 7,673 |
| Zhang et al., 2019 | | https://doi.org/10.1016/j.cell.2019.10.003 | Kidney | Normal/Tumor/Blood/Lymph node | 10x and Smart-seq2 | 8,669 |
| Cheng et al., 2021 | | https://doi.org/10.1016/j.cell.2021.01.010 | Lymphoma | Tumor/Blood | 10x | 615 |
| Cheng et al., 2021 | | https://doi.org/10.1016/j.cell.2021.01.010 | Myeloma | Tumor/Blood | 10x | 7,619 |
| Cheng et al., 2021 | | https://doi.org/10.1016/j.cell.2021.01.010 | Ovary-Fallopian Tube | Normal/Tumor | 10x | 3,888 |
| Cheng et al., 2021 and Peng et al., 2019 | | https://doi.org/10.1016/j.cell.2021.01.010 https://doi.org/10.1038/s41422-019-0195-y | Pancreas | Normal/Tumor | 10x | 7,904 |
| Cheng et al., 2021 | | https://doi.org/10.1016/j.cell.2021.01.010 | Thyroid | Normal/Tumor | 10x | 5,312 |
| Cheng et al., 2021 | | https://doi.org/10.1016/j.cell.2021.01.010 | Uterus | Normal/Tumor | 10x | 8,808 |
| **3 studies** | | | **8 sites** | **4 conditions** | **2 technologies** | **50,488 cells** |

**Mulder and colleagues (2021) datasets**

| Study | Link to Download | Paper | Tissue | Sample Type | Technology | Total of Myeloid Cells |
|---|---|---|---|---|---|---|
| MacParland et al., 2018 | GSE1154+B15:B3769 | https://doi.org/10.1038/s41467-018-06318-7 | Liver | Normal | 10x | 1,200 |

| | | | | | | |
|---|---|---|---|---|---|---|
| Aizarani et al., 2019 | GSE124395 | https://doi.org/10.1038/s41586-019-1373-2 | Liver | Normal | 10x | 1,685 |
| Ramachandran et al., 2019 | GSE136103 | https://doi.org/10.1038/s41586-019-1631-3 | Liver | Normal | 10x | 5,938 |
| Sharma et al., 2020 | GSE156337 | https://doi.org/10.1016/j.cell.2020.08.040 | Liver | Normal/Tumor | 10x | 4,991 |
| Zheng et al., 2017 | GSE98638 | https://doi.org/10.1016/j.cell.2017.05.035 | Liver | Normal/Tumor | 10x and Smart-seq2 | 11,310 |
| Mulder et al., 2021 | GSE178209 | https://doi.org/10.1016/j.immuni.2021.07.007 | Liver | Normal/Tumor | Smart-seq2 | 514 |
| Brown et al., 2019 | GSE137710 | https://doi.org/10.1016/j.cell.2019.09.035 | Spleen | Normal | 10x | 3,914 |
| Mulder et al., 2021 | GSE178209 | https://doi.org/10.1016/j.immuni.2021.07.007 | Spleen | Normal | Smart-seq2 | 560 |
| Tang-Huau et al., 2018 | GSE115007 | https://doi.org/10.1038/s41467-018-04985-0 | Tonsil | Normal | Dropseq | 1,782 |
| Mulder et al., 2021 | GSE178209 | https://doi.org/10.1016/j.immuni.2021.07.007 | Tonsil | Normal | Smart-seq2 | 37 |
| Cillo et al., 2020 | GSE139324 | https://doi.org/10.1016/j.immuni.2019.11.014 | Head Neck/Tonsil | Normal/Tumor | 10x | 8429 |
| Kim et al., 2020 | GSE131907 | https://doi.org/10.1038/s41467-020-16164-1 | Skin | Normal | 10x | 881 |
| Unpublished | - | - | Skin | Normal | Smart-seq2 | 1,370 |
| Cheng et al., 2018 | EGAS00001002927 | https://doi.org/10.1016/j.celrep.2018.09.006 | Skin | Normal | 10x | 1,303 |

| | | | | | | |
|---|---|---|---|---|---|---|
| He et al., 2020 | GSE147424 | https://doi.org/10.1016/j.jaci.2020.01.042 | Skin | Normal | 10x | 1,227 |
| Xue et al., 2019 | - | https://doi.org/10.4049/jimmunol.202.Supp.177.8 | Skin | Normal | 10x | 698 |
| Smillie et al., 2019 | GSE14580 | https://doi.org/10.1016/j.cell.2019.06.029 | Colon | Normal | 10x | 19,975 |
| Zhang et al., 2020 | GSE146771 | https://doi.org/10.1016/j.cell.2020.03.048 | Colon | Normal/Tumor | 10x and Smart-seq2 | 13,531 |
| James et al., 2020 | E-MTAB-8007, E-MTAB-8474, E-MTAB-8476, E-MTAB-8484 and E-MTAB-8486 | https://doi.org/10.1038/s41590-020-0602-z | Colon | Normal | 10x | 638 |
| Lee et al., 2020 | GSE132465, GSE132257 and GSE144735 | https://doi.org/10.1038/s41588-020-0636-z | Colon | Normal/Tumor | 10x | 6,245 |
| Reyfman et al., 2019 | - | https://doi.org/10.1164/rccm.201712-2410OC | Lung | Normal | 10x | 11,520 |
| Zilionis et al., 2019 | GSE127465 | https://doi.org/10.1016/j.immuni.2019.03.009 | Lung | Normal/Tumor | 10x | 9,185 |
| Braga et al., 2019 | GSE130148 | https://doi.org/10.1038/s41591-019-0468-5 | Lung | Normal | 10x | 3,087 |
| Lambrechts et al., 2018 | E-MTAB-6149 and E-MTAB-6653 | https://doi.org/10.1038/s41591-018-0096-5 | Lung | Normal/Tumor | 10x | 9,690 |
| Kim et al., 2020 | GSE131907 | https://doi.org/10.1038/s41467-020-16164-1 | Lung | Normal/Tumor | 10x | 18,729 |
| Mulder et al., 2021 | GSE178209 | https://doi.org/10.1016/j.immuni.2021.07.007 | Lung | Normal | Smart-seq2 | 206 |

| | | | | | | |
|---|---|---|---|---|---|---|
| Maier et al., 2020 | GSE131957 | https://doi.org/10.1038/s41586-020-2134-y | Lung | Normal/Tumor | 10x, CITE-seq | 6,684 |
| Arazi et al., 2019 | phs001457.v1.p1 | https://doi.org/10.1038/s41590-019-0398-x | Kidney | Normal | CEL-seq2 | 393 |
| Stewart et al., 2019 | ERP120466 | https://doi.org/10.1126/science.aat5031 | Kidney | Normal | 10x | 602 |
| Liao et al., 2020 | GSE145926 | https://doi.org/10.1038/s41591-020-0901-9 | Kidney | Normal | 10x | 694 |
| Peng et al., 2019 | CRA001160 | https://doi.org/10.1038/s41422-019-0195-y | Pancreas | Normal/Tumor | 10x | 5,426 |
| Tang-Huau et al., 2018 | GSE115007 | https://doi.org/10.1038/s41467-018-04985-0 | Ascite | Tumor | Dropseq | 5,440 |
| Unpublished | - | - | Stomach | Normal/Tumor | 10x | 3,398 |
| Azizi et al., 2018 | GSE114727, GSE114725 | https://doi.org/10.1016/j.cell.2018.05.060 | Breast | Normal/Tumor | 10x | 4,922 |
| Unpublished | - | - | Breast | Tumor | 10x, CITE-seq | 8,788 |
| Dutertre et al., 2019 | GSE132566 | https://doi.org/10.1016/j.immuni.2019.08.008 | Blood | Normal | Smart-seq2 | 320 |
| Villani et al., 2017 | GSE94820 | https://doi.org/10.1126/science.aah4573 | Blood | Normal | Smart-seq2 | 374 |
| **32 studies** | | | **12 sites** | **2 conditions** | **5 technologies** | **151,701 cells** |
