## Supplementary Table 7 for "Identification of novel myeloid-derived cell states with implication in cancer outcome"

**Supplementary Table 7: M1 and M2 gene signatures**

**M1 Signature**

| Gene name | Gene symbol | Function |
|---|---|---|
| Interleukin 1 Beta | IL1B | M1 Activating cytokines |
| Interferon Gamma | INFG | M1 Activating cytokines |
| Colony Stimulating Factor 2 | CSF2 | M1 Activating cytokines |
| CD80 Molecule | CD80 | M1 Surface Protein |
| CD86 Molecule | CD86 | M1 Surface Protein |
| Interleukin 1 Receptor Type 1 | IL1R1 | M1 Surface Protein |
| Major Histocompatibility Complex, Class II, DR | HLA-DR | M1 Surface Protein |
| Toll Like Receptor 2 | TLR2 | M1 Surface Protein |
| Toll Like Receptor 4 | TLR4 | M1 Surface Protein |
| Interferon Alpha/Beta Receptor 2 | IFNR2 | M1 Surface Protein |
| Interferon Alpha/Beta Receptor 1 | IFNR1 | M1 Surface Protein |
| Fc Gamma Receptor Ia | FCGR1A | M1 Surface Protein |
| C-X-C Motif Chemokine Ligand 9 | CXCL9 | M1 Cytokines |
| C-X-C Motif Chemokine Ligand 10 | CXCL10 | M1 Cytokines |
| C-X-C Motif Chemokine Ligand 11 | CXCL11 | M1 Cytokines |
| Tumor Necrosis Factor | TNF | M1 Cytokines |
| Interleukin 6 | IL-6 | M1 Cytokines |
| Interleukin 12A | IL-12A | M1 Cytokines |
| Interleukin 12B | IL-12B | M1 Cytokines |
| Interleukin 23A | IL-23A | M1 Cytokines |
| Interleukin 1 alpha | IL1A | M1 Cytokines |
| Nitric Oxide Synthase 1 | NOS1 | M1 Effector molecules |
| Nitric Oxide Synthase 2 | NOS2 | M1 Effector molecules |
| ROS Proto-Oncogene 1, Receptor Tyrosine Kinase | ROS1 | M1 Effector molecules |
| Solute Carrier Family 2 Member 1 | SLC2A1 | M1 Metabolism |
| Transcription Factor EB | TFEB | M1 Transcription factors |
| Signal Transducer And Activator Of Transcription 1 | STAT1 | M1 Transcription factors |
| Nuclear Factor Kappa B Subunit 1 | NFKB1 | M1 Transcription factors |
| Signal Transducer And Activator Of Transcription 3 | STAT3 | M1 Transcription factors |
| interferon regulatory transcription 3 | IFR3 | M1 Transcription factors |
| interferon regulatory transcription 5 | IFR5 | M1 Transcription factors |
| interferon regulatory transcription 7 | IFR7 | M1 Transcription factors |
| Suppressor Of Cytokine Signaling 3 | SOCS3 | M1 Transcription factors |
| Jun Proto-Oncogene, AP-1 Transcription Factor Subunit | JUN | M1 Transcription factors |

**Supplementary Table 7: M1 and M2 gene signatures**

**M2 Signature**

| Gene name | Gene symbol | Function |
|---|---|---|
| Interleukin 4 | IL4 | M2 Activating cytokines |
| Interleukin 13 | IL13 | M2 Activating cytokines |
| C-C Motif Chemokine Ligand 17 | CCL17 | M2 Cytokines |
| C-C Motif Chemokine Ligand 22 | CCL22 | M2 Cytokines |
| C-C Motif Chemokine Ligand 24 | CCL24 | M2 Cytokines |
| Interleukin 10 | IL10 | M2 Cytokines |
| Transforming Growth Factor Beta 1 | TGFB1 | M2 Cytokines |
| C-C Motif Chemokine Ligand 23 | CCL13 | M2 Cytokines |
| Vascular endothelial growth factor A | VEGFA | M2 Effector molecules |
| Epidermal growth factor | EGF | M2 Effector molecules |
| Platelet derived growth factor subunit A | PDGFA | M2 Effector molecules |
| Platelet derived growth factor subunit B | PDGFB | M2 Effector molecules |
| Matrix metallopeptidase 9 | MMP9 | M2 Effector molecules |
| Vascular endothelial growth factor B | VEGFB | M2 Effector molecules |
| Vascular endothelial growth factor C | VEGFC | M2 Effector molecules |
| Vascular endothelial growth factor D | VEGFD | M2 Effector molecules |
| Transforming growth factor Beta-2 | TGFB2 | M2 Effector molecules |
| Transforming growth factor Beta-3 | TGFB3 | M2 Effector molecules |
| Matrix metallopeptidase 14 | MMP14 | M2 Effector molecules |
| Matrix metallopeptidase 19 | MMP19 | M2 Effector molecules |
| Selenoprotein P | SEPP1 | M2 Effector molecules |
| Arginase 1 | ARG1 | M2 Metabolism |
| Indoleamine 2,3-dioxygenase 2 | IDO2 | M2 Metabolism |
| CD163 molecule | CD163 | M2 Surface Protein |
| Macrophage Scavenger Receptor 1 | MSR1 | M2 Surface Protein |
| Stabilin 1 | STAB1 | M2 Surface Protein |
| Macrophage Receptor With Collagenous Structure | MARCO | M2 Surface Protein |
| CD36 Molecule | CD36 | M2 Surface Protein |

| | | |
|---|---|---|
| Fc Gamma Receptor IIa | FCGR2A | M2 Surface Protein |
| Interleukin 1 Receptor Type 2 | IL1R2 | M2 Surface Protein |
| Interleukin 4 Receptor | IL4R | M2 Surface Protein |
| Programmed Cell Death 1 Ligand 1 | CD274 | M2 Surface Protein |
| Programmed Cell Death 1 Ligand 2 | PDCD1LG2 | M2 Surface Protein |
| Programmed Cell Death 1 | PDCD1 | M2 Surface Protein |
| Signal Regulatory Protein Alpha | SIRPA | M2 Surface Protein |
| Sialic Acid Binding Ig Like Lectin 10 | SIGLEC10 | M2 Surface Protein |
| S100 Calcium Binding Protein A9 | S100A9 | M2 Surface Protein |
| Leukocyte Immunoglobulin Like Receptor B1 | LILRB1 | M2 Surface Protein |
| Leukocyte Immunoglobulin Like Receptor B2 | LILRB2 | M2 Surface Protein |
| V-Set Domain Containing T Cell Activation Inhibitor 1) | VTCN1 | M2 Surface Protein |
| TEK Receptor Tyrosine Kinase | TEK | M2 Surface Protein |
| Triggering Receptor Expressed On Myeloid Cells 1 | TREM1 | M2 Surface Protein |
| Triggering Receptor Expressed On Myeloid Cells 2 | TREM2 | M2 Surface Protein |
| Interleukin 1 Receptor Antagonist | IL1RN | M2 Surface Protein |
| Signal transducer and activator of transcription 6 | STAT6 | M2 Transcription factors |
| MAF bZIP transcription factor B | MAFB | M2 Transcription factors |
| Interferon regulatory factor 4 | IFR4 | M2 Transcription factors |
| Suppressor Of Cytokine Signaling 1 | SOCS1 | M2 Transcription factors |
| Chitinase 3 Like 1 | CHI3L1 | Hydrolase |
| AXL Receptor Tyrosine Kinase | AXL | RTK subfamily |
