## Supplementary Table 8 for "Identification of novel myeloid-derived cell states with implication in cancer outcome"

**Supplementary Table 8: Prediction of communication axes between macrophages and lymphocytes and malignant cells**

**MacAlv-like vs. lymphocytes**

| celltype_01 | celltype_02 | ligand | receptor | ligand_mean_expression | receptor_mean_expression | ligand_receptor_mean_expression | pvalue |
|---|---|---|---|---|---|---|---|
| MacAlv-like | NKT | ADAM17 | ITGB1 | 0.594200 | 0.488000 | 0.541100 | 0.000999 |
| MacAlv-like | NKT | ADAM17 | TNFRSF1B | 0.594200 | 0.494900 | 0.544600 | 0.000999 |
| MacAlv-like | NKT | AIMP1 | RACK1 | 0.302300 | 2.432700 | 1.367500 | 0.000999 |
| MacAlv-like | NKT | ANXA1 | CD44 | 1.783100 | 0.672400 | 1.227800 | 0.000999 |
| MacAlv-like | NKT | ANXA1 | CXCR4 | 1.783100 | 2.037300 | 1.910200 | 0.000999 |
| MacAlv-like | NKT | ANXA1 | F2R | 1.783100 | 0.297800 | 1.040500 | 0.000999 |
| MacAlv-like | NKT | ANXA1 | ITGA4 | 1.783100 | 0.294600 | 1.038800 | 0.000999 |
| MacAlv-like | NKT | APOE | SORL1 | 2.499500 | 0.413700 | 1.456600 | 0.000999 |
| MacAlv-like | NKT | CCL20 | CXCR4 | 0.294400 | 2.037300 | 1.165800 | 0.000999 |
| MacAlv-like | NKT | CCL20 | S1PR4 | 0.294400 | 0.443500 | 0.369000 | 0.000999 |
| MacAlv-like | NKT | CCL20 | S1PR5 | 0.294400 | 0.508000 | 0.401200 | 0.000999 |
| MacAlv-like | NKT | CD58 | CD2 | 0.295000 | 1.146500 | 0.720700 | 0.000999 |
| MacAlv-like | NKT | CXCL16 | CXCR4 | 1.132400 | 2.037300 | 1.584800 | 0.000999 |
| MacAlv-like | NKT | CXCL16 | S1PR4 | 1.132400 | 0.443500 | 0.787900 | 0.000999 |
| MacAlv-like | NKT | CXCL16 | S1PR5 | 1.132400 | 0.508000 | 0.820200 | 0.000999 |
| MacAlv-like | NKT | CXCL2 | CXCR4 | 0.292800 | 2.037300 | 1.165000 | 0.000999 |
| MacAlv-like | NKT | CXCL2 | S1PR4 | 0.292800 | 0.443500 | 0.368200 | 0.000999 |
| MacAlv-like | NKT | CXCL2 | S1PR5 | 0.292800 | 0.508000 | 0.400400 | 0.000999 |
| MacAlv-like | NKT | CXCL3 | CXCR4 | 0.649800 | 2.037300 | 1.343500 | 0.000999 |
| MacAlv-like | NKT | CXCL3 | S1PR4 | 0.649800 | 0.443500 | 0.546700 | 0.000999 |
| MacAlv-like | NKT | CXCL3 | S1PR5 | 0.649800 | 0.508000 | 0.578900 | 0.000999 |
| MacAlv-like | NKT | FN1 | CD44 | 1.493500 | 0.672400 | 1.083000 | 0.000999 |
| MacAlv-like | NKT | FN1 | ITGA4 | 1.493500 | 0.294600 | 0.894000 | 0.000999 |
| MacAlv-like | NKT | FN1 | ITGB1 | 1.493500 | 0.488000 | 0.990800 | 0.000999 |
| MacAlv-like | NKT | FN1 | ITGB7 | 1.493500 | 0.472100 | 0.982800 | 0.000999 |
| MacAlv-like | NKT | FN1 | RPSA | 1.493500 | 2.363900 | 1.928700 | 0.000999 |
| MacAlv-like | NKT | GRN | TNFRSF1B | 2.472000 | 0.494900 | 1.483500 | 0.000999 |

| | | | | | | | |
|---|---|---|---|---|---|---|---|
| MacAlv-like | NKT | GSTP1 | CD44 | 1.773100 | 0.672400 | 1.222800 | 0.000999 |
| MacAlv-like | NKT | GSTP1 | IL7R | 1.773100 | 0.593300 | 1.183200 | 0.000999 |
| MacAlv-like | NKT | GSTP1 | ITGA4 | 1.773100 | 0.294600 | 1.033800 | 0.000999 |
| MacAlv-like | NKT | HLA-DRA | PTPRC | 4.039300 | 1.879700 | 2.959500 | 0.000999 |
| MacAlv-like | NKT | HSP90B1 | ITGB1 | 0.760900 | 0.488000 | 0.624400 | 0.000999 |
| MacAlv-like | NKT | HSP90B1 | TNFRSF1B | 0.760900 | 0.494900 | 0.627900 | 0.000999 |
| MacAlv-like | NKT | INHBA | TGFBR3 | 0.624000 | 0.412800 | 0.518400 | 0.000999 |
| MacAlv-like | NKT | LGALS3 | CD6 | 2.969500 | 0.488100 | 1.728800 | 0.000999 |
| MacAlv-like | NKT | LGALS3 | ITGB1 | 2.969500 | 0.488000 | 1.728700 | 0.000999 |
| MacAlv-like | NKT | LGALS3 | SPN | 2.969500 | 0.639800 | 1.804600 | 0.000999 |
| MacAlv-like | NKT | LRPAP1 | SORL1 | 0.831100 | 0.413700 | 0.622400 | 0.000999 |
| MacAlv-like | NKT | NAMPT | ANXA6 | 0.773600 | 0.844000 | 0.808800 | 0.000999 |
| MacAlv-like | NKT | NMB | F2R | 0.304200 | 0.297800 | 0.301000 | 0.000999 |
| MacAlv-like | NKT | NMB | GPR65 | 0.304200 | 0.489800 | 0.397000 | 0.000999 |
| MacAlv-like | NKT | PECAM1 | CXCR4 | 0.645200 | 2.037300 | 1.341300 | 0.000999 |
| MacAlv-like | NKT | RPS19 | ITGA4 | 3.667200 | 0.294600 | 1.980900 | 0.000999 |
| MacAlv-like | NKT | RPS19 | RPSA | 3.667200 | 2.363900 | 3.015500 | 0.000999 |
| MacAlv-like | NKT | RTN4 | CD44 | 1.382900 | 0.672400 | 1.027600 | 0.000999 |
| MacAlv-like | NKT | RTN4 | CD99 | 1.382900 | 1.454700 | 1.418800 | 0.000999 |
| MacAlv-like | NKT | THBS1 | CD47 | 0.473400 | 0.557600 | 0.515500 | 0.000999 |
| MacAlv-like | NKT | THBS1 | ITGA4 | 0.473400 | 0.294600 | 0.384000 | 0.000999 |
| MacAlv-like | NKT | THBS1 | ITGB1 | 0.473400 | 0.488000 | 0.480700 | 0.000999 |
| MacAlv-like | NKT | TNFSF13B | ITGB1 | 0.444100 | 0.488000 | 0.466000 | 0.000999 |
| MacAlv-like | NKT | ITGB1 | ITGA4 | 0.480400 | 0.294600 | 0.387500 | 0.006990 |
| MacAlv-like | NKT | ITGB1 | RACK1 | 0.480400 | 2.432700 | 1.456500 | 0.008990 |
| MacAlv-like | TCD4ex | ADAM17 | ITGB1 | 0.594200 | 0.461500 | 0.527900 | 0.000999 |
| MacAlv-like | TCD4ex | ADAM17 | TNFRSF1B | 0.594200 | 0.383000 | 0.488600 | 0.000999 |
| MacAlv-like | TCD4ex | AIMP1 | RACK1 | 0.302300 | 2.443400 | 1.372900 | 0.000999 |
| MacAlv-like | TCD4ex | ANXA1 | CCR7 | 1.783100 | 0.484400 | 1.133800 | 0.000999 |
| MacAlv-like | TCD4ex | ANXA1 | CD4 | 1.783100 | 0.586400 | 1.184800 | 0.000999 |

| | | | | | | | |
|---|---|---|---|---|---|---|---|
| MacAlv-like | TCD4ex | ANXA1 | CD44 | 1.783100 | 1.024400 | 1.403800 | 0.000999 |
| MacAlv-like | TCD4ex | ANXA1 | CXCR4 | 1.783100 | 2.420100 | 2.101600 | 0.000999 |
| MacAlv-like | TCD4ex | ARF1 | CD4 | 0.641600 | 0.586400 | 0.614000 | 0.000999 |
| MacAlv-like | TCD4ex | CALM1 | RPSA | 1.526800 | 2.489500 | 2.008100 | 0.000999 |
| MacAlv-like | TCD4ex | CALM1 | TNFRSF1B | 1.526800 | 0.383000 | 0.954900 | 0.000999 |
| MacAlv-like | TCD4ex | CALR | IL7R | 0.820700 | 1.235300 | 1.028000 | 0.000999 |
| MacAlv-like | TCD4ex | CCL20 | CCR7 | 0.294400 | 0.484400 | 0.389400 | 0.000999 |
| MacAlv-like | TCD4ex | CCL20 | CXCR4 | 0.294400 | 2.420100 | 1.357200 | 0.000999 |
| MacAlv-like | TCD4ex | CD58 | CD2 | 0.295000 | 1.573100 | 0.934000 | 0.000999 |
| MacAlv-like | TCD4ex | CD86 | CD28 | 0.330400 | 0.325600 | 0.328000 | 0.000999 |
| MacAlv-like | TCD4ex | CD86 | CTLA4 | 0.330400 | 0.629400 | 0.479900 | 0.000999 |
| MacAlv-like | TCD4ex | CXCL16 | CCR7 | 1.132400 | 0.484400 | 0.808400 | 0.000999 |
| MacAlv-like | TCD4ex | CXCL16 | CXCR4 | 1.132400 | 2.420100 | 1.776200 | 0.000999 |
| MacAlv-like | TCD4ex | CXCL2 | CCR7 | 0.292800 | 0.484400 | 0.388600 | 0.000999 |
| MacAlv-like | TCD4ex | CXCL2 | CXCR4 | 0.292800 | 2.420100 | 1.356400 | 0.000999 |
| MacAlv-like | TCD4ex | CXCL3 | CCR7 | 0.649800 | 0.484400 | 0.567100 | 0.000999 |
| MacAlv-like | TCD4ex | CXCL3 | CXCR4 | 0.649800 | 2.420100 | 1.534900 | 0.000999 |
| MacAlv-like | TCD4ex | FN1 | CD44 | 1.493500 | 1.024400 | 1.259000 | 0.000999 |
| MacAlv-like | TCD4ex | FN1 | ITGB1 | 1.493500 | 0.461500 | 0.977500 | 0.000999 |
| MacAlv-like | TCD4ex | FN1 | RPSA | 1.493500 | 2.489500 | 1.991500 | 0.000999 |
| MacAlv-like | TCD4ex | GRN | TNFRSF1B | 2.472000 | 0.383000 | 1.427500 | 0.000999 |
| MacAlv-like | TCD4ex | GSTP1 | CD44 | 1.773100 | 1.024400 | 1.398700 | 0.000999 |
| MacAlv-like | TCD4ex | GSTP1 | IL7R | 1.773100 | 1.235300 | 1.504200 | 0.000999 |
| MacAlv-like | TCD4ex | HLA-DMA | CD4 | 1.709300 | 0.586400 | 1.147800 | 0.000999 |
| MacAlv-like | TCD4ex | HLA-DRA | CD4 | 4.039300 | 0.586400 | 2.312900 | 0.000999 |
| MacAlv-like | TCD4ex | HLA-DRA | PDCD1 | 4.039300 | 0.457000 | 2.248200 | 0.000999 |
| MacAlv-like | TCD4ex | HLA-DRA | PTPRC | 4.039300 | 1.749300 | 2.894300 | 0.000999 |
| MacAlv-like | TCD4ex | HSP90B1 | ITGB1 | 0.760900 | 0.461500 | 0.611200 | 0.000999 |
| MacAlv-like | TCD4ex | HSP90B1 | TNFRSF1B | 0.760900 | 0.383000 | 0.572000 | 0.000999 |
| MacAlv-like | TCD4ex | ITGB1 | RACK1 | 0.480400 | 2.443400 | 1.461900 | 0.000999 |

| | | | | | | | |
|---|---|---|---|---|---|---|---|
| MacAlv-like | TCD4ex | LGALS3 | CD6 | 2.969500 | 0.533800 | 1.751600 | 0.000999 |
| MacAlv-like | TCD4ex | LGALS3 | ITGB1 | 2.969500 | 0.461500 | 1.715500 | 0.000999 |
| MacAlv-like | TCD4ex | NAMPT | ANXA6 | 0.773600 | 0.538600 | 0.656100 | 0.000999 |
| MacAlv-like | TCD4ex | PECAM1 | CXCR4 | 0.645200 | 2.420100 | 1.532600 | 0.000999 |
| MacAlv-like | TCD4ex | RPS19 | CD4 | 3.667200 | 0.586400 | 2.126800 | 0.000999 |
| MacAlv-like | TCD4ex | RPS19 | RPSA | 3.667200 | 2.489500 | 3.078300 | 0.000999 |
| MacAlv-like | TCD4ex | RTN4 | CD44 | 1.382900 | 1.024400 | 1.203600 | 0.000999 |
| MacAlv-like | TCD4ex | RTN4 | CD99 | 1.382900 | 0.916500 | 1.149700 | 0.000999 |
| MacAlv-like | TCD4ex | TGFB1 | ITGB1 | 0.365800 | 0.461500 | 0.413600 | 0.000999 |
| MacAlv-like | TCD4ex | THBS1 | CD47 | 0.473400 | 0.345600 | 0.409500 | 0.000999 |
| MacAlv-like | TCD4ex | THBS1 | ITGB1 | 0.473400 | 0.461500 | 0.467500 | 0.000999 |
| MacAlv-like | TCD4ex | TNFSF12 | TNFRSF25 | 0.297400 | 0.325900 | 0.311600 | 0.000999 |
| MacAlv-like | TCD4ex | TNFSF13B | ITGB1 | 0.444100 | 0.461500 | 0.452800 | 0.000999 |
| MacAlv-like | TCD4reg | ADAM17 | ITGB1 | 0.594200 | 0.400100 | 0.497100 | 0.000999 |
| MacAlv-like | TCD4reg | ADAM17 | TNFRSF1B | 0.594200 | 0.798200 | 0.696200 | 0.000999 |
| MacAlv-like | TCD4reg | AIMP1 | FAS | 0.302300 | 0.272700 | 0.287500 | 0.000999 |
| MacAlv-like | TCD4reg | AIMP1 | RACK1 | 0.302300 | 2.529800 | 1.416000 | 0.000999 |
| MacAlv-like | TCD4reg | ANXA1 | CCR7 | 1.783100 | 0.309800 | 1.046500 | 0.000999 |
| MacAlv-like | TCD4reg | ANXA1 | CD4 | 1.783100 | 0.429000 | 1.106000 | 0.000999 |
| MacAlv-like | TCD4reg | ANXA1 | CD44 | 1.783100 | 1.089600 | 1.436400 | 0.000999 |
| MacAlv-like | TCD4reg | ANXA1 | CXCR3 | 1.783100 | 0.335700 | 1.059400 | 0.000999 |
| MacAlv-like | TCD4reg | ANXA1 | CXCR4 | 1.783100 | 2.296200 | 2.039600 | 0.000999 |
| MacAlv-like | TCD4reg | ANXA1 | CXCR6 | 1.783100 | 0.394500 | 1.088800 | 0.000999 |
| MacAlv-like | TCD4reg | ANXA1 | FAS | 1.783100 | 0.272700 | 1.027900 | 0.000999 |
| MacAlv-like | TCD4reg | CALM1 | RPSA | 1.526800 | 2.499900 | 2.013400 | 0.000999 |
| MacAlv-like | TCD4reg | CALM1 | TNFRSF1B | 1.526800 | 0.798200 | 1.162500 | 0.000999 |
| MacAlv-like | TCD4reg | CALR | FAS | 0.820700 | 0.272700 | 0.546700 | 0.000999 |
| MacAlv-like | TCD4reg | CALR | IL7R | 0.820700 | 0.968200 | 0.894500 | 0.000999 |
| MacAlv-like | TCD4reg | CCL20 | CCR7 | 0.294400 | 0.309800 | 0.302100 | 0.000999 |
| MacAlv-like | TCD4reg | CCL20 | CXCR3 | 0.294400 | 0.335700 | 0.315000 | 0.000999 |

| | | | | | | | |
|---|---|---|---|---|---|---|---|
| MacAlv-like | TCD4reg | CCL20 | CXCR4 | 0.294400 | 2.296200 | 1.295300 | 0.000999 |
| MacAlv-like | TCD4reg | CCL20 | CXCR6 | 0.294400 | 0.394500 | 0.344500 | 0.000999 |
| MacAlv-like | TCD4reg | CCL20 | S1PR4 | 0.294400 | 0.274400 | 0.284400 | 0.000999 |
| MacAlv-like | TCD4reg | CD58 | CD2 | 0.295000 | 1.656600 | 0.975800 | 0.000999 |
| MacAlv-like | TCD4reg | CD86 | CD28 | 0.330400 | 0.313000 | 0.321700 | 0.000999 |
| MacAlv-like | TCD4reg | CD86 | CTLA4 | 0.330400 | 0.790900 | 0.560700 | 0.000999 |
| MacAlv-like | TCD4reg | CXCL16 | CCR7 | 1.132400 | 0.309800 | 0.721100 | 0.000999 |
| MacAlv-like | TCD4reg | CXCL16 | CXCR3 | 1.132400 | 0.335700 | 0.734000 | 0.000999 |
| MacAlv-like | TCD4reg | CXCL16 | CXCR4 | 1.132400 | 2.296200 | 1.714300 | 0.000999 |
| MacAlv-like | TCD4reg | CXCL16 | CXCR6 | 1.132400 | 0.394500 | 0.763400 | 0.000999 |
| MacAlv-like | TCD4reg | CXCL16 | S1PR4 | 1.132400 | 0.274400 | 0.703400 | 0.000999 |
| MacAlv-like | TCD4reg | CXCL2 | CCR7 | 0.292800 | 0.309800 | 0.301300 | 0.000999 |
| MacAlv-like | TCD4reg | CXCL2 | CXCR3 | 0.292800 | 0.335700 | 0.314200 | 0.000999 |
| MacAlv-like | TCD4reg | CXCL2 | CXCR4 | 0.292800 | 2.296200 | 1.294500 | 0.000999 |
| MacAlv-like | TCD4reg | CXCL2 | CXCR6 | 0.292800 | 0.394500 | 0.343700 | 0.000999 |
| MacAlv-like | TCD4reg | CXCL2 | S1PR4 | 0.292800 | 0.274400 | 0.283600 | 0.000999 |
| MacAlv-like | TCD4reg | CXCL3 | CCR7 | 0.649800 | 0.309800 | 0.479800 | 0.000999 |
| MacAlv-like | TCD4reg | CXCL3 | CXCR3 | 0.649800 | 0.335700 | 0.492700 | 0.000999 |
| MacAlv-like | TCD4reg | CXCL3 | CXCR4 | 0.649800 | 2.296200 | 1.473000 | 0.000999 |
| MacAlv-like | TCD4reg | CXCL3 | CXCR6 | 0.649800 | 0.394500 | 0.522200 | 0.000999 |
| MacAlv-like | TCD4reg | CXCL3 | S1PR4 | 0.649800 | 0.274400 | 0.462100 | 0.000999 |
| MacAlv-like | TCD4reg | FN1 | CD44 | 1.493500 | 1.089600 | 1.291600 | 0.000999 |
| MacAlv-like | TCD4reg | FN1 | ITGB1 | 1.493500 | 0.400100 | 0.946800 | 0.000999 |
| MacAlv-like | TCD4reg | FN1 | RPSA | 1.493500 | 2.499900 | 1.996700 | 0.000999 |
| MacAlv-like | TCD4reg | GRN | TNFRSF1B | 2.472000 | 0.798200 | 1.635100 | 0.000999 |
| MacAlv-like | TCD4reg | GSTP1 | CD44 | 1.773100 | 1.089600 | 1.431400 | 0.000999 |
| MacAlv-like | TCD4reg | GSTP1 | FAS | 1.773100 | 0.272700 | 1.022900 | 0.000999 |
| MacAlv-like | TCD4reg | GSTP1 | IL7R | 1.773100 | 0.968200 | 1.370600 | 0.000999 |
| MacAlv-like | TCD4reg | HLA-DMA | CD4 | 1.709300 | 0.429000 | 1.069100 | 0.000999 |
| MacAlv-like | TCD4reg | HLA-DRA | CD4 | 4.039300 | 0.429000 | 2.234200 | 0.000999 |

| | | | | | | | |
|---|---|---|---|---|---|---|---|
| MacAlv-like | TCD4reg | HLA-DRA | PTPRC | 4.039300 | 1.660300 | 2.849800 | 0.000999 |
| MacAlv-like | TCD4reg | HSP90B1 | ITGB1 | 0.760900 | 0.400100 | 0.580500 | 0.000999 |
| MacAlv-like | TCD4reg | HSP90B1 | TNFRSF1B | 0.760900 | 0.798200 | 0.779500 | 0.000999 |
| MacAlv-like | TCD4reg | ITGB1 | RACK1 | 0.480400 | 2.529800 | 1.505100 | 0.000999 |
| MacAlv-like | TCD4reg | LGALS3 | CD6 | 2.969500 | 0.265100 | 1.617300 | 0.000999 |
| MacAlv-like | TCD4reg | LGALS3 | FAS | 2.969500 | 0.272700 | 1.621100 | 0.000999 |
| MacAlv-like | TCD4reg | LGALS3 | ITGB1 | 2.969500 | 0.400100 | 1.684800 | 0.000999 |
| MacAlv-like | TCD4reg | NAMPT | ANXA6 | 0.773600 | 0.455700 | 0.614600 | 0.000999 |
| MacAlv-like | TCD4reg | NMB | P2RY10 | 0.304200 | 0.282400 | 0.293300 | 0.000999 |
| MacAlv-like | TCD4reg | PECAM1 | CXCR4 | 0.645200 | 2.296200 | 1.470700 | 0.000999 |
| MacAlv-like | TCD4reg | RPS19 | RPSA | 3.667200 | 2.499900 | 3.083600 | 0.000999 |
| MacAlv-like | TCD4reg | RTN4 | CD44 | 1.382900 | 1.089600 | 1.236200 | 0.000999 |
| MacAlv-like | TCD4reg | RTN4 | CD99 | 1.382900 | 0.660300 | 1.021600 | 0.000999 |
| MacAlv-like | TCD4reg | TGFB1 | ITGB1 | 0.365800 | 0.400100 | 0.382900 | 0.000999 |
| MacAlv-like | TCD4reg | THBS1 | CD47 | 0.473400 | 0.416000 | 0.444700 | 0.000999 |
| MacAlv-like | TCD4reg | THBS1 | ITGB1 | 0.473400 | 0.400100 | 0.436800 | 0.000999 |
| MacAlv-like | TCD4reg | TNFSF13B | ITGB1 | 0.444100 | 0.400100 | 0.422100 | 0.000999 |
| MacAlv-like | TCD8em | ADAM17 | ITGB1 | 0.594200 | 0.255700 | 0.425000 | 0.000999 |
| MacAlv-like | TCD8em | ADAM17 | TNFRSF1B | 0.594200 | 0.303800 | 0.449000 | 0.000999 |
| MacAlv-like | TCD8em | AIMP1 | RACK1 | 0.302300 | 2.437400 | 1.369800 | 0.000999 |
| MacAlv-like | TCD8em | ANXA1 | CD44 | 1.783100 | 1.119000 | 1.451100 | 0.000999 |
| MacAlv-like | TCD8em | ANXA1 | CXCR3 | 1.783100 | 0.418900 | 1.101000 | 0.000999 |
| MacAlv-like | TCD8em | ANXA1 | CXCR4 | 1.783100 | 2.838900 | 2.311000 | 0.000999 |
| MacAlv-like | TCD8em | ANXA1 | CXCR6 | 1.783100 | 0.339900 | 1.061500 | 0.000999 |
| MacAlv-like | TCD8em | ANXA1 | F2R | 1.783100 | 0.269500 | 1.026300 | 0.000999 |
| MacAlv-like | TCD8em | ANXA1 | ITGA4 | 1.783100 | 0.282300 | 1.032700 | 0.000999 |
| MacAlv-like | TCD8em | CALR | IL7R | 0.820700 | 0.718300 | 0.769500 | 0.000999 |
| MacAlv-like | TCD8em | CCL20 | CXCR3 | 0.294400 | 0.418900 | 0.356600 | 0.000999 |
| MacAlv-like | TCD8em | CCL20 | CXCR4 | 0.294400 | 2.838900 | 1.566700 | 0.000999 |
| MacAlv-like | TCD8em | CCL20 | CXCR6 | 0.294400 | 0.339900 | 0.317100 | 0.000999 |

| | | | | | | | |
|---|---|---|---|---|---|---|---|
| MacAlv-like | TCD8em | CCL20 | S1PR4 | 0.294400 | 0.267700 | 0.281100 | 0.000999 |
| MacAlv-like | TCD8em | CD58 | CD2 | 0.295000 | 1.596100 | 0.945500 | 0.000999 |
| MacAlv-like | TCD8em | CXCL16 | CXCR3 | 1.132400 | 0.418900 | 0.775600 | 0.000999 |
| MacAlv-like | TCD8em | CXCL16 | CXCR4 | 1.132400 | 2.838900 | 1.985600 | 0.000999 |
| MacAlv-like | TCD8em | CXCL16 | CXCR6 | 1.132400 | 0.339900 | 0.736100 | 0.000999 |
| MacAlv-like | TCD8em | CXCL16 | S1PR4 | 1.132400 | 0.267700 | 0.700000 | 0.000999 |
| MacAlv-like | TCD8em | CXCL2 | CXCR3 | 0.292800 | 0.418900 | 0.355800 | 0.000999 |
| MacAlv-like | TCD8em | CXCL2 | CXCR4 | 0.292800 | 2.838900 | 1.565900 | 0.000999 |
| MacAlv-like | TCD8em | CXCL2 | CXCR6 | 0.292800 | 0.339900 | 0.316300 | 0.000999 |
| MacAlv-like | TCD8em | CXCL2 | S1PR4 | 0.292800 | 0.267700 | 0.280300 | 0.000999 |
| MacAlv-like | TCD8em | CXCL3 | CXCR3 | 0.649800 | 0.418900 | 0.534300 | 0.000999 |
| MacAlv-like | TCD8em | CXCL3 | CXCR4 | 0.649800 | 2.838900 | 1.744400 | 0.000999 |
| MacAlv-like | TCD8em | CXCL3 | CXCR6 | 0.649800 | 0.339900 | 0.494800 | 0.000999 |
| MacAlv-like | TCD8em | CXCL3 | S1PR4 | 0.649800 | 0.267700 | 0.458800 | 0.000999 |
| MacAlv-like | TCD8em | FN1 | CD44 | 1.493500 | 1.119000 | 1.306300 | 0.000999 |
| MacAlv-like | TCD8em | FN1 | ITGA4 | 1.493500 | 0.282300 | 0.887900 | 0.000999 |
| MacAlv-like | TCD8em | FN1 | ITGB1 | 1.493500 | 0.255700 | 0.874600 | 0.000999 |
| MacAlv-like | TCD8em | FN1 | RPSA | 1.493500 | 2.368600 | 1.931100 | 0.000999 |
| MacAlv-like | TCD8em | GRN | TNFRSF1B | 2.472000 | 0.303800 | 1.387900 | 0.000999 |
| MacAlv-like | TCD8em | GSTP1 | CD44 | 1.773100 | 1.119000 | 1.446100 | 0.000999 |
| MacAlv-like | TCD8em | GSTP1 | IL7R | 1.773100 | 0.718300 | 1.245700 | 0.000999 |
| MacAlv-like | TCD8em | GSTP1 | ITGA4 | 1.773100 | 0.282300 | 1.027700 | 0.000999 |
| MacAlv-like | TCD8em | HLA-DRA | PTPRC | 4.039300 | 1.791200 | 2.915300 | 0.000999 |
| MacAlv-like | TCD8em | HSP90B1 | ITGB1 | 0.760900 | 0.255700 | 0.508300 | 0.000999 |
| MacAlv-like | TCD8em | HSP90B1 | TNFRSF1B | 0.760900 | 0.303800 | 0.532300 | 0.000999 |
| MacAlv-like | TCD8em | ITGB1 | CD47 | 0.480400 | 0.427600 | 0.454000 | 0.000999 |
| MacAlv-like | TCD8em | ITGB1 | ITGA4 | 0.480400 | 0.282300 | 0.381300 | 0.000999 |
| MacAlv-like | TCD8em | ITGB1 | RACK1 | 0.480400 | 2.437400 | 1.458900 | 0.000999 |
| MacAlv-like | TCD8em | LGALS3 | CD6 | 2.969500 | 0.420400 | 1.695000 | 0.000999 |
| MacAlv-like | TCD8em | LGALS3 | ITGB1 | 2.969500 | 0.255700 | 1.612600 | 0.000999 |

| | | | | | | | |
|---|---|---|---|---|---|---|---|
| MacAlv-like | TCD8em | LGALS3 | SPN | 2.969500 | 0.260300 | 1.614900 | 0.000999 |
| MacAlv-like | TCD8em | NAMPT | ANXA6 | 0.773600 | 0.552000 | 0.662800 | 0.000999 |
| MacAlv-like | TCD8em | NMB | F2R | 0.304200 | 0.269500 | 0.286900 | 0.000999 |
| MacAlv-like | TCD8em | NMB | GPR65 | 0.304200 | 0.262500 | 0.283300 | 0.000999 |
| MacAlv-like | TCD8em | PECAM1 | CXCR4 | 0.645200 | 2.838900 | 1.742100 | 0.000999 |
| MacAlv-like | TCD8em | RPS19 | ITGA4 | 3.667200 | 0.282300 | 1.974800 | 0.000999 |
| MacAlv-like | TCD8em | RPS19 | RPSA | 3.667200 | 2.368600 | 3.017900 | 0.000999 |
| MacAlv-like | TCD8em | RTN4 | CD44 | 1.382900 | 1.119000 | 1.251000 | 0.000999 |
| MacAlv-like | TCD8em | RTN4 | CD99 | 1.382900 | 1.152500 | 1.267700 | 0.000999 |
| MacAlv-like | TCD8em | TGFB1 | ITGB1 | 0.365800 | 0.255700 | 0.310800 | 0.000999 |
| MacAlv-like | TCD8em | THBS1 | CD47 | 0.473400 | 0.427600 | 0.450500 | 0.000999 |
| MacAlv-like | TCD8em | THBS1 | ITGA4 | 0.473400 | 0.282300 | 0.377900 | 0.000999 |
| MacAlv-like | TCD8em | THBS1 | ITGB1 | 0.473400 | 0.255700 | 0.364600 | 0.000999 |
| MacAlv-like | TCD8em | TNFSF13B | ITGB1 | 0.444100 | 0.255700 | 0.349900 | 0.000999 |
| MacAlv-like | TCD8em | CALM1 | RPSA | 1.526800 | 2.368600 | 1.947700 | 0.005000 |
| MacAlv-like | TCD8ex | ADAM17 | ITGB1 | 0.594200 | 0.299500 | 0.446800 | 0.000999 |
| MacAlv-like | TCD8ex | ADAM17 | TNFRSF1B | 0.594200 | 0.643600 | 0.618900 | 0.000999 |
| MacAlv-like | TCD8ex | AIMP1 | RACK1 | 0.302300 | 2.335300 | 1.318800 | 0.000999 |
| MacAlv-like | TCD8ex | ANXA1 | CD44 | 1.783100 | 0.821900 | 1.302500 | 0.000999 |
| MacAlv-like | TCD8ex | ANXA1 | CXCR3 | 1.783100 | 0.364700 | 1.073900 | 0.000999 |
| MacAlv-like | TCD8ex | ANXA1 | CXCR4 | 1.783100 | 2.347200 | 2.065200 | 0.000999 |
| MacAlv-like | TCD8ex | ANXA1 | CXCR6 | 1.783100 | 0.662600 | 1.222800 | 0.000999 |
| MacAlv-like | TCD8ex | ANXA1 | ITGA4 | 1.783100 | 0.443200 | 1.113200 | 0.000999 |
| MacAlv-like | TCD8ex | ANXA1 | VCAM1 | 1.783100 | 0.446200 | 1.114700 | 0.000999 |
| MacAlv-like | TCD8ex | APOE | VCAM1 | 2.499500 | 0.446200 | 1.472800 | 0.000999 |
| MacAlv-like | TCD8ex | CALM1 | RPSA | 1.526800 | 2.269700 | 1.898200 | 0.000999 |
| MacAlv-like | TCD8ex | CALM1 | TNFRSF1B | 1.526800 | 0.643600 | 1.085200 | 0.000999 |
| MacAlv-like | TCD8ex | CCL20 | CXCR3 | 0.294400 | 0.364700 | 0.329500 | 0.000999 |
| MacAlv-like | TCD8ex | CCL20 | CXCR4 | 0.294400 | 2.347200 | 1.320800 | 0.000999 |
| MacAlv-like | TCD8ex | CCL20 | CXCR6 | 0.294400 | 0.662600 | 0.478500 | 0.000999 |

| | | | | | | | |
|---|---|---|---|---|---|---|---|
| MacAlv-like | TCD8ex | CD58 | CD2 | 0.295000 | 2.050200 | 1.172600 | 0.000999 |
| MacAlv-like | TCD8ex | CD86 | CTLA4 | 0.330400 | 0.488300 | 0.409400 | 0.000999 |
| MacAlv-like | TCD8ex | CXCL16 | CXCR3 | 1.132400 | 0.364700 | 0.748500 | 0.000999 |
| MacAlv-like | TCD8ex | CXCL16 | CXCR4 | 1.132400 | 2.347200 | 1.739800 | 0.000999 |
| MacAlv-like | TCD8ex | CXCL16 | CXCR6 | 1.132400 | 0.662600 | 0.897500 | 0.000999 |
| MacAlv-like | TCD8ex | CXCL2 | CXCR3 | 0.292800 | 0.364700 | 0.328700 | 0.000999 |
| MacAlv-like | TCD8ex | CXCL2 | CXCR4 | 0.292800 | 2.347200 | 1.320000 | 0.000999 |
| MacAlv-like | TCD8ex | CXCL2 | CXCR6 | 0.292800 | 0.662600 | 0.477700 | 0.000999 |
| MacAlv-like | TCD8ex | CXCL3 | CXCR3 | 0.649800 | 0.364700 | 0.507200 | 0.000999 |
| MacAlv-like | TCD8ex | CXCL3 | CXCR4 | 0.649800 | 2.347200 | 1.498500 | 0.000999 |
| MacAlv-like | TCD8ex | CXCL3 | CXCR6 | 0.649800 | 0.662600 | 0.656200 | 0.000999 |
| MacAlv-like | TCD8ex | FN1 | CD44 | 1.493500 | 0.821900 | 1.157700 | 0.000999 |
| MacAlv-like | TCD8ex | FN1 | ITGA4 | 1.493500 | 0.443200 | 0.968400 | 0.000999 |
| MacAlv-like | TCD8ex | FN1 | ITGB1 | 1.493500 | 0.299500 | 0.896500 | 0.000999 |
| MacAlv-like | TCD8ex | FN1 | ITGB7 | 1.493500 | 0.300700 | 0.897100 | 0.000999 |
| MacAlv-like | TCD8ex | FN1 | RPSA | 1.493500 | 2.269700 | 1.881600 | 0.000999 |
| MacAlv-like | TCD8ex | GRN | TNFRSF1B | 2.472000 | 0.643600 | 1.557800 | 0.000999 |
| MacAlv-like | TCD8ex | GSTP1 | CD44 | 1.773100 | 0.821900 | 1.297500 | 0.000999 |
| MacAlv-like | TCD8ex | GSTP1 | ITGA4 | 1.773100 | 0.443200 | 1.108200 | 0.000999 |
| MacAlv-like | TCD8ex | GSTP1 | VCAM1 | 1.773100 | 0.446200 | 1.109600 | 0.000999 |
| MacAlv-like | TCD8ex | HLA-DRA | PDCD1 | 4.039300 | 0.610900 | 2.325100 | 0.000999 |
| MacAlv-like | TCD8ex | HLA-DRA | PTPRC | 4.039300 | 1.676200 | 2.857800 | 0.000999 |
| MacAlv-like | TCD8ex | HSP90B1 | ITGB1 | 0.760900 | 0.299500 | 0.530200 | 0.000999 |
| MacAlv-like | TCD8ex | HSP90B1 | TNFRSF1B | 0.760900 | 0.643600 | 0.702300 | 0.000999 |
| MacAlv-like | TCD8ex | ITGB1 | CD47 | 0.480400 | 0.395200 | 0.437800 | 0.000999 |
| MacAlv-like | TCD8ex | ITGB1 | ITGA4 | 0.480400 | 0.443200 | 0.461800 | 0.000999 |
| MacAlv-like | TCD8ex | ITGB1 | RACK1 | 0.480400 | 2.335300 | 1.407800 | 0.000999 |
| MacAlv-like | TCD8ex | ITGB1 | VCAM1 | 0.480400 | 0.446200 | 0.463300 | 0.000999 |
| MacAlv-like | TCD8ex | LGALS3 | CD6 | 2.969500 | 0.318000 | 1.643800 | 0.000999 |
| MacAlv-like | TCD8ex | LGALS3 | ITGB1 | 2.969500 | 0.299500 | 1.634500 | 0.000999 |

| | | | | | | | |
|---|---|---|---|---|---|---|---|
| MacAlv-like | TCD8ex | LGALS3 | SPN | 2.969500 | 0.348600 | 1.659100 | 0.000999 |
| MacAlv-like | TCD8ex | NAMPT | ANXA6 | 0.773600 | 0.567800 | 0.670700 | 0.000999 |
| MacAlv-like | TCD8ex | NMB | GPR65 | 0.304200 | 0.390400 | 0.347300 | 0.000999 |
| MacAlv-like | TCD8ex | PECAM1 | CXCR4 | 0.645200 | 2.347200 | 1.496200 | 0.000999 |
| MacAlv-like | TCD8ex | RTN4 | CD44 | 1.382900 | 0.821900 | 1.102400 | 0.000999 |
| MacAlv-like | TCD8ex | RTN4 | CD99 | 1.382900 | 1.204900 | 1.293900 | 0.000999 |
| MacAlv-like | TCD8ex | TGFB1 | ITGB1 | 0.365800 | 0.299500 | 0.332600 | 0.000999 |
| MacAlv-like | TCD8ex | THBS1 | CD47 | 0.473400 | 0.395200 | 0.434300 | 0.000999 |
| MacAlv-like | TCD8ex | THBS1 | ITGA4 | 0.473400 | 0.443200 | 0.458300 | 0.000999 |
| MacAlv-like | TCD8ex | THBS1 | ITGB1 | 0.473400 | 0.299500 | 0.386500 | 0.000999 |
| MacAlv-like | TCD8ex | TNFSF13B | ITGB1 | 0.444100 | 0.299500 | 0.371800 | 0.000999 |
| MacAlv-like | TGD | AIMP1 | RACK1 | 0.302300 | 2.728700 | 1.515500 | 0.000999 |
| MacAlv-like | TGD | ANXA1 | CXCR3 | 1.783100 | 1.174800 | 1.478900 | 0.000999 |
| MacAlv-like | TGD | ANXA1 | CXCR4 | 1.783100 | 1.489400 | 1.636300 | 0.000999 |
| MacAlv-like | TGD | ANXA1 | ITGA4 | 1.783100 | 0.396800 | 1.090000 | 0.000999 |
| MacAlv-like | TGD | APOE | SORL1 | 2.499500 | 0.331800 | 1.415600 | 0.000999 |
| MacAlv-like | TGD | ARF1 | ITGA4 | 0.641600 | 0.396800 | 0.519200 | 0.000999 |
| MacAlv-like | TGD | CALM1 | RPSA | 1.526800 | 2.881400 | 2.204100 | 0.000999 |
| MacAlv-like | TGD | CALR | IL7R | 0.820700 | 1.386200 | 1.103500 | 0.000999 |
| MacAlv-like | TGD | CCL20 | CXCR3 | 0.294400 | 1.174800 | 0.734600 | 0.000999 |
| MacAlv-like | TGD | CCL20 | CXCR4 | 0.294400 | 1.489400 | 0.891900 | 0.000999 |
| MacAlv-like | TGD | CD58 | CD2 | 0.295000 | 1.452200 | 0.873600 | 0.000999 |
| MacAlv-like | TGD | CD99 | CD99 | 0.507100 | 1.715400 | 1.111200 | 0.000999 |
| MacAlv-like | TGD | CXCL16 | CXCR3 | 1.132400 | 1.174800 | 1.153600 | 0.000999 |
| MacAlv-like | TGD | CXCL16 | CXCR4 | 1.132400 | 1.489400 | 1.310900 | 0.000999 |
| MacAlv-like | TGD | CXCL2 | CXCR3 | 0.292800 | 1.174800 | 0.733800 | 0.000999 |
| MacAlv-like | TGD | CXCL2 | CXCR4 | 0.292800 | 1.489400 | 0.891100 | 0.000999 |
| MacAlv-like | TGD | CXCL3 | CXCR3 | 0.649800 | 1.174800 | 0.912300 | 0.000999 |
| MacAlv-like | TGD | CXCL3 | CXCR4 | 0.649800 | 1.489400 | 1.069600 | 0.000999 |
| MacAlv-like | TGD | FN1 | ITGA4 | 1.493500 | 0.396800 | 0.945200 | 0.000999 |

| | | | | | | | |
|---|---|---|---|---|---|---|---|
| MacAlv-like | TGD | FN1 | ITGB7 | 1.493500 | 0.459000 | 0.976300 | 0.000999 |
| MacAlv-like | TGD | FN1 | RPSA | 1.493500 | 2.881400 | 2.187400 | 0.000999 |
| MacAlv-like | TGD | GRN | TNFRSF1A | 2.472000 | 0.309900 | 1.391000 | 0.000999 |
| MacAlv-like | TGD | GSTP1 | IL7R | 1.773100 | 1.386200 | 1.579600 | 0.000999 |
| MacAlv-like | TGD | GSTP1 | ITGA4 | 1.773100 | 0.396800 | 1.085000 | 0.000999 |
| MacAlv-like | TGD | HLA-A | CD8A | 2.470500 | 1.746000 | 2.108300 | 0.000999 |
| MacAlv-like | TGD | HLA-DRA | PTPRC | 4.039300 | 1.865300 | 2.952300 | 0.000999 |
| MacAlv-like | TGD | HLA-E | CD8A | 1.262800 | 1.746000 | 1.504400 | 0.000999 |
| MacAlv-like | TGD | HLA-E | KLRC2 | 1.262800 | 0.808200 | 1.035500 | 0.000999 |
| MacAlv-like | TGD | HLA-E | KLRD1 | 1.262800 | 1.345100 | 1.303900 | 0.000999 |
| MacAlv-like | TGD | HLA-F | CD8A | 0.320200 | 1.746000 | 1.033100 | 0.000999 |
| MacAlv-like | TGD | HMGB1 | ITGA4 | 1.171800 | 0.396800 | 0.784300 | 0.000999 |
| MacAlv-like | TGD | ITGB1 | ITGA4 | 0.480400 | 0.396800 | 0.438600 | 0.000999 |
| MacAlv-like | TGD | ITGB1 | PTK2B | 0.480400 | 0.366800 | 0.423600 | 0.000999 |
| MacAlv-like | TGD | ITGB1 | RACK1 | 0.480400 | 2.728700 | 1.604500 | 0.000999 |
| MacAlv-like | TGD | ITGB2 | ICAM1 | 0.731600 | 0.291500 | 0.511600 | 0.000999 |
| MacAlv-like | TGD | ITGB2 | ITGAL | 0.731600 | 0.373100 | 0.552400 | 0.000999 |
| MacAlv-like | TGD | LGALS3 | CD6 | 2.969500 | 0.468400 | 1.719000 | 0.000999 |
| MacAlv-like | TGD | LGALS3 | ITGB1 | 2.969500 | 0.365700 | 1.667600 | 0.000999 |
| MacAlv-like | TGD | LGALS3 | SPN | 2.969500 | 0.564400 | 1.766900 | 0.000999 |
| MacAlv-like | TGD | LGALS3 | TGFBR2 | 2.969500 | 0.367700 | 1.668600 | 0.000999 |
| MacAlv-like | TGD | LRPAP1 | SORL1 | 0.831100 | 0.331800 | 0.581400 | 0.000999 |
| MacAlv-like | TGD | MIF | ITGA4 | 0.585100 | 0.396800 | 0.491000 | 0.000999 |
| MacAlv-like | TGD | MIF | TNFRSF14 | 0.585100 | 0.464200 | 0.524600 | 0.000999 |
| MacAlv-like | TGD | NAMPT | ANXA6 | 0.773600 | 0.855300 | 0.814400 | 0.000999 |
| MacAlv-like | TGD | NMB | GPR65 | 0.304200 | 0.536800 | 0.420500 | 0.000999 |
| MacAlv-like | TGD | PECAM1 | CXCR4 | 0.645200 | 1.489400 | 1.067300 | 0.000999 |
| MacAlv-like | TGD | PTPRC | CD2 | 0.690800 | 1.452200 | 1.071500 | 0.000999 |
| MacAlv-like | TGD | PTPRC | CD8A | 0.690800 | 1.746000 | 1.218400 | 0.000999 |
| MacAlv-like | TGD | PTPRC | CXCR4 | 0.690800 | 1.489400 | 1.090100 | 0.000999 |

| | | | | | | | |
|---|---|---|---|---|---|---|---|
| MacAlv-like | TGD | RPS19 | ICAM1 | 3.667200 | 0.291500 | 1.979300 | 0.000999 |
| MacAlv-like | TGD | RPS19 | ITGA4 | 3.667200 | 0.396800 | 2.032000 | 0.000999 |
| MacAlv-like | TGD | RPS19 | RPSA | 3.667200 | 2.881400 | 3.274300 | 0.000999 |
| MacAlv-like | TGD | RTN4 | CD99 | 1.382900 | 1.715400 | 1.549100 | 0.000999 |
| MacAlv-like | TGD | THBS1 | ITGA4 | 0.473400 | 0.396800 | 0.435100 | 0.000999 |
| MacAlv-like | TGD | ADAM17 | TNFRSF1A | 0.594200 | 0.309900 | 0.452000 | 0.003000 |
| NKT | MacAlv-like | ALOX5AP | ALOX5 | 0.502100 | 0.873900 | 0.688000 | 0.000999 |
| NKT | MacAlv-like | ANXA1 | C3AR1 | 1.225900 | 0.301300 | 0.763600 | 0.000999 |
| NKT | MacAlv-like | ANXA1 | C5AR1 | 1.225900 | 0.535500 | 0.880700 | 0.000999 |
| NKT | MacAlv-like | ANXA1 | CD44 | 1.225900 | 1.485400 | 1.355700 | 0.000999 |
| NKT | MacAlv-like | ANXA1 | FPR1 | 1.225900 | 0.600900 | 0.913400 | 0.000999 |
| NKT | MacAlv-like | ANXA1 | FPR2 | 1.225900 | 0.361100 | 0.793500 | 0.000999 |
| NKT | MacAlv-like | ANXA1 | FPR3 | 1.225900 | 0.340000 | 0.783000 | 0.000999 |
| NKT | MacAlv-like | ARF1 | CD4 | 0.799100 | 0.262400 | 0.530700 | 0.000999 |
| NKT | MacAlv-like | ARF1 | CD44 | 0.799100 | 1.485400 | 1.142300 | 0.000999 |
| NKT | MacAlv-like | CALM1 | TNFRSF1A | 1.933500 | 0.270600 | 1.102100 | 0.000999 |
| NKT | MacAlv-like | CALR | CD44 | 0.947900 | 1.485400 | 1.216600 | 0.000999 |
| NKT | MacAlv-like | CALR | LRP1 | 0.947900 | 0.514900 | 0.731400 | 0.000999 |
| NKT | MacAlv-like | CD2 | CD4 | 1.146500 | 0.262400 | 0.704400 | 0.000999 |
| NKT | MacAlv-like | CD320 | TGFBR2 | 0.254500 | 0.259700 | 0.257100 | 0.000999 |
| NKT | MacAlv-like | GSTP1 | CD44 | 0.790900 | 1.485400 | 1.138100 | 0.000999 |
| NKT | MacAlv-like | GZMB | IGF2R | 1.836100 | 0.265800 | 1.050900 | 0.000999 |
| NKT | MacAlv-like | HMGB1 | LY96 | 1.195000 | 0.944100 | 1.069500 | 0.000999 |
| NKT | MacAlv-like | HMGB2 | CD44 | 0.646700 | 1.485400 | 1.066000 | 0.000999 |
| NKT | MacAlv-like | HMGB2 | DAB2 | 0.646700 | 0.738700 | 0.692700 | 0.000999 |
| NKT | MacAlv-like | HSP90B1 | LRP1 | 0.613800 | 0.514900 | 0.564400 | 0.000999 |
| NKT | MacAlv-like | IL16 | CD4 | 0.355600 | 0.262400 | 0.309000 | 0.000999 |
| NKT | MacAlv-like | IL16 | FCGR1A | 0.355600 | 0.490300 | 0.422900 | 0.000999 |
| NKT | MacAlv-like | ITGA4 | CD44 | 0.294600 | 1.485400 | 0.890000 | 0.000999 |
| NKT | MacAlv-like | ITGA4 | TFRC | 0.294600 | 0.617300 | 0.455900 | 0.000999 |

| | | | | | | | |
|---|---|---|---|---|---|---|---|
| NKT | MacAlv-like | ITGB1 | CD151 | 0.488000 | 0.513700 | 0.500800 | 0.000999 |
| NKT | MacAlv-like | ITGB1 | CD46 | 0.488000 | 0.364700 | 0.426400 | 0.000999 |
| NKT | MacAlv-like | ITGB1 | DAB2 | 0.488000 | 0.738700 | 0.613300 | 0.000999 |
| NKT | MacAlv-like | ITGB1 | ENG | 0.488000 | 0.293500 | 0.390700 | 0.000999 |
| NKT | MacAlv-like | ITGB1 | PLAUR | 0.488000 | 0.658200 | 0.573100 | 0.000999 |
| NKT | MacAlv-like | ITGB2 | PLAUR | 1.648100 | 0.658200 | 1.153200 | 0.000999 |
| NKT | MacAlv-like | LRPAP1 | LRP1 | 0.280700 | 0.514900 | 0.397800 | 0.000999 |
| NKT | MacAlv-like | LRPAP1 | SORT1 | 0.280700 | 0.283500 | 0.282100 | 0.000999 |
| NKT | MacAlv-like | LTB | TNFRSF1A | 0.260500 | 0.270600 | 0.265500 | 0.000999 |
| NKT | MacAlv-like | MIF | CD74 | 0.665700 | 4.191700 | 2.428700 | 0.000999 |
| NKT | MacAlv-like | PTPRC | CD4 | 1.879700 | 0.262400 | 1.071000 | 0.000999 |
| NKT | MacAlv-like | PTPRC | CD44 | 1.879700 | 1.485400 | 1.682500 | 0.000999 |
| NKT | MacAlv-like | RPS19 | C5AR1 | 3.533600 | 0.535500 | 2.034600 | 0.000999 |
| NKT | MacAlv-like | RPS19 | CD4 | 3.533600 | 0.262400 | 1.898000 | 0.000999 |
| NKT | MacAlv-like | RTN4 | CD44 | 0.437400 | 1.485400 | 0.961400 | 0.000999 |
| NKT | MacAlv-like | TGFB1 | ENG | 0.544100 | 0.293500 | 0.418800 | 0.000999 |
| NKT | MacAlv-like | TGFB1 | TGFBR2 | 0.544100 | 0.259700 | 0.401900 | 0.000999 |
| NKT | MacAlv-like | TIMP1 | LRP1 | 0.300500 | 0.514900 | 0.407700 | 0.000999 |
| NKT | MacAlv-like | XCL2 | FPR2 | 0.277400 | 0.361100 | 0.319300 | 0.000999 |
| NKT | MacAlv-like | XCL2 | PTAFR | 0.277400 | 0.307700 | 0.292600 | 0.000999 |
| NKT | MacAlv-like | HMGB1 | AGTRAP | 1.195000 | 0.266000 | 0.730500 | 0.003000 |
| NKT | MacAlv-like | ITGB1 | CD47 | 0.488000 | 0.618300 | 0.553200 | 0.008990 |
| TCD4ex | MacAlv-like | ALOX5AP | ALOX5 | 1.119000 | 0.873900 | 0.996400 | 0.000999 |
| TCD4ex | MacAlv-like | ARF1 | CD44 | 0.682700 | 1.485400 | 1.084000 | 0.000999 |
| TCD4ex | MacAlv-like | CALM1 | TNFRSF1A | 1.295700 | 0.270600 | 0.783200 | 0.000999 |
| TCD4ex | MacAlv-like | CALR | CD44 | 0.748000 | 1.485400 | 1.116700 | 0.000999 |
| TCD4ex | MacAlv-like | CALR | LRP1 | 0.748000 | 0.514900 | 0.631500 | 0.000999 |
| TCD4ex | MacAlv-like | CD2 | CD4 | 1.573100 | 0.262400 | 0.917700 | 0.000999 |
| TCD4ex | MacAlv-like | CXCL13 | C3AR1 | 2.336700 | 0.301300 | 1.319000 | 0.000999 |
| TCD4ex | MacAlv-like | CXCL13 | C5AR1 | 2.336700 | 0.535500 | 1.436100 | 0.000999 |

| | | | | | | | |
|---|---|---|---|---|---|---|---|
| TCD4ex | MacAlv-like | CXCL13 | FPR1 | 2.336700 | 0.600900 | 1.468800 | 0.000999 |
| TCD4ex | MacAlv-like | CXCL13 | FPR2 | 2.336700 | 0.361100 | 1.348900 | 0.000999 |
| TCD4ex | MacAlv-like | HMGB1 | AGTRAP | 1.387600 | 0.266000 | 0.826800 | 0.000999 |
| TCD4ex | MacAlv-like | HMGB1 | LY96 | 1.387600 | 0.944100 | 1.165800 | 0.000999 |
| TCD4ex | MacAlv-like | HMGB2 | CD44 | 1.024300 | 1.485400 | 1.254800 | 0.000999 |
| TCD4ex | MacAlv-like | HMGB2 | DAB2 | 1.024300 | 0.738700 | 0.881500 | 0.000999 |
| TCD4ex | MacAlv-like | HSP90B1 | LRP1 | 0.394400 | 0.514900 | 0.454700 | 0.000999 |
| TCD4ex | MacAlv-like | ICAM3 | ITGB2 | 0.573000 | 0.731600 | 0.652300 | 0.000999 |
| TCD4ex | MacAlv-like | ITGB1 | CD151 | 0.461500 | 0.513700 | 0.487600 | 0.000999 |
| TCD4ex | MacAlv-like | ITGB1 | CD46 | 0.461500 | 0.364700 | 0.413100 | 0.000999 |
| TCD4ex | MacAlv-like | ITGB1 | CD47 | 0.461500 | 0.618300 | 0.539900 | 0.000999 |
| TCD4ex | MacAlv-like | ITGB1 | DAB2 | 0.461500 | 0.738700 | 0.600100 | 0.000999 |
| TCD4ex | MacAlv-like | ITGB1 | ENG | 0.461500 | 0.293500 | 0.377500 | 0.000999 |
| TCD4ex | MacAlv-like | ITGB1 | IGF2R | 0.461500 | 0.265800 | 0.363600 | 0.000999 |
| TCD4ex | MacAlv-like | ITGB1 | PLAUR | 0.461500 | 0.658200 | 0.559900 | 0.000999 |
| TCD4ex | MacAlv-like | ITGB2 | PLAUR | 0.787100 | 0.658200 | 0.722600 | 0.000999 |
| TCD4ex | MacAlv-like | LRPAP1 | LRP1 | 0.280900 | 0.514900 | 0.397900 | 0.000999 |
| TCD4ex | MacAlv-like | LTB | ITGB1 | 0.965500 | 0.480400 | 0.722900 | 0.000999 |
| TCD4ex | MacAlv-like | LTB | TNFRSF1A | 0.965500 | 0.270600 | 0.618100 | 0.000999 |
| TCD4ex | MacAlv-like | MIF | CD74 | 0.699900 | 4.191700 | 2.445800 | 0.000999 |
| TCD4ex | MacAlv-like | NMB | FPR2 | 0.639100 | 0.361100 | 0.500100 | 0.000999 |
| TCD4ex | MacAlv-like | NMB | GPR65 | 0.639100 | 0.308500 | 0.473800 | 0.000999 |
| TCD4ex | MacAlv-like | NMB | PTAFR | 0.639100 | 0.307700 | 0.473400 | 0.000999 |
| TCD4ex | MacAlv-like | PTPRC | CD4 | 1.749300 | 0.262400 | 1.005800 | 0.000999 |
| TCD4ex | MacAlv-like | PTPRC | CD44 | 1.749300 | 1.485400 | 1.617400 | 0.000999 |
| TCD4ex | MacAlv-like | RPS19 | C5AR1 | 3.632900 | 0.535500 | 2.084200 | 0.000999 |
| TCD4ex | MacAlv-like | SELL | SELPLG | 0.561800 | 0.349200 | 0.455500 | 0.000999 |
| TCD4ex | MacAlv-like | SPP1 | CD44 | 0.331400 | 1.485400 | 0.908400 | 0.000999 |
| TCD4ex | MacAlv-like | SPP1 | ITGB1 | 0.331400 | 0.480400 | 0.405900 | 0.000999 |
| TCD4ex | MacAlv-like | TIMP1 | LRP1 | 0.337400 | 0.514900 | 0.426200 | 0.000999 |

| | | | | | | | |
|---|---|---|---|---|---|---|---|
| TCD4reg | MacAlv-like | ALOX5AP | ALOX5 | 0.458900 | 0.873900 | 0.666400 | 0.000999 |
| TCD4reg | MacAlv-like | APOE | LRP1 | 0.293400 | 0.514900 | 0.404200 | 0.000999 |
| TCD4reg | MacAlv-like | ARF1 | CD44 | 0.718500 | 1.485400 | 1.101900 | 0.000999 |
| TCD4reg | MacAlv-like | CALM1 | TNFRSF1A | 1.534000 | 0.270600 | 0.902300 | 0.000999 |
| TCD4reg | MacAlv-like | CALR | CD44 | 0.683500 | 1.485400 | 1.084400 | 0.000999 |
| TCD4reg | MacAlv-like | CALR | LRP1 | 0.683500 | 0.514900 | 0.599200 | 0.000999 |
| TCD4reg | MacAlv-like | CD2 | CD4 | 1.656600 | 0.262400 | 0.959500 | 0.000999 |
| TCD4reg | MacAlv-like | GSTP1 | CD44 | 0.491700 | 1.485400 | 0.988500 | 0.000999 |
| TCD4reg | MacAlv-like | HMGB1 | AGTRAP | 1.254700 | 0.266000 | 0.760400 | 0.000999 |
| TCD4reg | MacAlv-like | HMGB1 | LY96 | 1.254700 | 0.944100 | 1.099400 | 0.000999 |
| TCD4reg | MacAlv-like | HMGB2 | CD44 | 0.616000 | 1.485400 | 1.050700 | 0.000999 |
| TCD4reg | MacAlv-like | HMGB2 | DAB2 | 0.616000 | 0.738700 | 0.677400 | 0.000999 |
| TCD4reg | MacAlv-like | HSP90B1 | LRP1 | 0.422100 | 0.514900 | 0.468500 | 0.000999 |
| TCD4reg | MacAlv-like | ICAM3 | ITGB2 | 0.666300 | 0.731600 | 0.698900 | 0.000999 |
| TCD4reg | MacAlv-like | IL16 | FCGR1A | 0.252800 | 0.490300 | 0.371600 | 0.000999 |
| TCD4reg | MacAlv-like | ITGB1 | CD151 | 0.400100 | 0.513700 | 0.456900 | 0.000999 |
| TCD4reg | MacAlv-like | ITGB1 | CD46 | 0.400100 | 0.364700 | 0.382400 | 0.000999 |
| TCD4reg | MacAlv-like | ITGB1 | CD47 | 0.400100 | 0.618300 | 0.509200 | 0.000999 |
| TCD4reg | MacAlv-like | ITGB1 | DAB2 | 0.400100 | 0.738700 | 0.569400 | 0.000999 |
| TCD4reg | MacAlv-like | ITGB1 | ENG | 0.400100 | 0.293500 | 0.346800 | 0.000999 |
| TCD4reg | MacAlv-like | ITGB1 | IGF2R | 0.400100 | 0.265800 | 0.332900 | 0.000999 |
| TCD4reg | MacAlv-like | ITGB1 | PLAUR | 0.400100 | 0.658200 | 0.529200 | 0.000999 |
| TCD4reg | MacAlv-like | ITGB2 | PLAUR | 0.623800 | 0.658200 | 0.641000 | 0.000999 |
| TCD4reg | MacAlv-like | LGALS3 | FCGR2A | 0.449200 | 0.785400 | 0.617300 | 0.000999 |
| TCD4reg | MacAlv-like | LRPAP1 | LRP1 | 0.310100 | 0.514900 | 0.412500 | 0.000999 |
| TCD4reg | MacAlv-like | LRPAP1 | SORT1 | 0.310100 | 0.283500 | 0.296800 | 0.000999 |
| TCD4reg | MacAlv-like | LTB | ITGB1 | 2.242000 | 0.480400 | 1.361200 | 0.000999 |
| TCD4reg | MacAlv-like | LTB | TNFRSF1A | 2.242000 | 0.270600 | 1.256300 | 0.000999 |
| TCD4reg | MacAlv-like | MIF | CD74 | 0.629500 | 4.191700 | 2.410600 | 0.000999 |
| TCD4reg | MacAlv-like | PSAP | LRP1 | 0.593800 | 0.514900 | 0.554400 | 0.000999 |

| | | | | | | | |
|---|---|---|---|---|---|---|---|
| TCD4reg | MacAlv-like | PTPRC | CD44 | 1.660300 | 1.485400 | 1.572900 | 0.000999 |
| TCD4reg | MacAlv-like | RPS19 | C5AR1 | 3.689700 | 0.535500 | 2.112600 | 0.000999 |
| TCD4reg | MacAlv-like | RTN4 | CD44 | 0.352800 | 1.485400 | 0.919100 | 0.000999 |
| TCD4reg | MacAlv-like | SELL | SELPLG | 0.706200 | 0.349200 | 0.527700 | 0.000999 |
| TCD4reg | MacAlv-like | SPP1 | CD44 | 0.312500 | 1.485400 | 0.899000 | 0.000999 |
| TCD4reg | MacAlv-like | SPP1 | ITGB1 | 0.312500 | 0.480400 | 0.396400 | 0.000999 |
| TCD4reg | MacAlv-like | TIMP1 | LRP1 | 0.436200 | 0.514900 | 0.475600 | 0.000999 |
| TCD8em | MacAlv-like | ALOX5AP | ALOX5 | 0.797300 | 0.873900 | 0.835600 | 0.000999 |
| TCD8em | MacAlv-like | ANXA1 | C3AR1 | 1.069300 | 0.301300 | 0.685300 | 0.000999 |
| TCD8em | MacAlv-like | ANXA1 | C5AR1 | 1.069300 | 0.535500 | 0.802400 | 0.000999 |
| TCD8em | MacAlv-like | ANXA1 | CD4 | 1.069300 | 0.262400 | 0.665800 | 0.000999 |
| TCD8em | MacAlv-like | ANXA1 | CD44 | 1.069300 | 1.485400 | 1.277300 | 0.000999 |
| TCD8em | MacAlv-like | ANXA1 | FPR1 | 1.069300 | 0.600900 | 0.835100 | 0.000999 |
| TCD8em | MacAlv-like | ANXA1 | FPR2 | 1.069300 | 0.361100 | 0.715200 | 0.000999 |
| TCD8em | MacAlv-like | ANXA1 | FPR3 | 1.069300 | 0.340000 | 0.704600 | 0.000999 |
| TCD8em | MacAlv-like | APOE | LRP1 | 0.279100 | 0.514900 | 0.397000 | 0.000999 |
| TCD8em | MacAlv-like | ARF1 | CD4 | 0.668400 | 0.262400 | 0.465400 | 0.000999 |
| TCD8em | MacAlv-like | ARF1 | CD44 | 0.668400 | 1.485400 | 1.076900 | 0.000999 |
| TCD8em | MacAlv-like | CALM1 | TNFRSF1A | 1.526200 | 0.270600 | 0.898400 | 0.000999 |
| TCD8em | MacAlv-like | CALR | CD44 | 0.714100 | 1.485400 | 1.099800 | 0.000999 |
| TCD8em | MacAlv-like | CALR | LRP1 | 0.714100 | 0.514900 | 0.614500 | 0.000999 |
| TCD8em | MacAlv-like | CD2 | CD4 | 1.596100 | 0.262400 | 0.929200 | 0.000999 |
| TCD8em | MacAlv-like | GSTP1 | CD44 | 0.767000 | 1.485400 | 1.126200 | 0.000999 |
| TCD8em | MacAlv-like | GZMB | IGF2R | 0.478100 | 0.265800 | 0.371900 | 0.000999 |
| TCD8em | MacAlv-like | HMGB1 | AGTRAP | 1.363600 | 0.266000 | 0.814800 | 0.000999 |
| TCD8em | MacAlv-like | HMGB1 | LY96 | 1.363600 | 0.944100 | 1.153800 | 0.000999 |
| TCD8em | MacAlv-like | HMGB2 | CD44 | 0.721200 | 1.485400 | 1.103300 | 0.000999 |
| TCD8em | MacAlv-like | HMGB2 | DAB2 | 0.721200 | 0.738700 | 0.729900 | 0.000999 |
| TCD8em | MacAlv-like | HSP90B1 | ITGB1 | 0.471000 | 0.480400 | 0.475700 | 0.000999 |
| TCD8em | MacAlv-like | HSP90B1 | LRP1 | 0.471000 | 0.514900 | 0.493000 | 0.000999 |

| | | | | | | | |
|---|---|---|---|---|---|---|---|
| TCD8em | MacAlv-like | IFNG | IFNGR1 | 0.396700 | 0.437700 | 0.417200 | 0.000999 |
| TCD8em | MacAlv-like | IFNG | IFNGR2 | 0.396700 | 0.422900 | 0.409800 | 0.000999 |
| TCD8em | MacAlv-like | IFNG | PTPN6 | 0.396700 | 0.563800 | 0.480200 | 0.000999 |
| TCD8em | MacAlv-like | IL16 | CD4 | 0.408500 | 0.262400 | 0.335400 | 0.000999 |
| TCD8em | MacAlv-like | IL16 | FCGR1A | 0.408500 | 0.490300 | 0.449400 | 0.000999 |
| TCD8em | MacAlv-like | ITGA4 | CD44 | 0.282300 | 1.485400 | 0.883900 | 0.000999 |
| TCD8em | MacAlv-like | ITGA4 | TFRC | 0.282300 | 0.617300 | 0.449800 | 0.000999 |
| TCD8em | MacAlv-like | ITGB1 | CD151 | 0.255700 | 0.513700 | 0.384700 | 0.000999 |
| TCD8em | MacAlv-like | ITGB1 | CD46 | 0.255700 | 0.364700 | 0.310200 | 0.000999 |
| TCD8em | MacAlv-like | ITGB1 | CD47 | 0.255700 | 0.618300 | 0.437000 | 0.000999 |
| TCD8em | MacAlv-like | ITGB1 | DAB2 | 0.255700 | 0.738700 | 0.497200 | 0.000999 |
| TCD8em | MacAlv-like | ITGB1 | ENG | 0.255700 | 0.293500 | 0.274600 | 0.000999 |
| TCD8em | MacAlv-like | ITGB1 | IGF2R | 0.255700 | 0.265800 | 0.260800 | 0.000999 |
| TCD8em | MacAlv-like | ITGB1 | PLAUR | 0.255700 | 0.658200 | 0.457000 | 0.000999 |
| TCD8em | MacAlv-like | ITGB2 | PLAUR | 1.001800 | 0.658200 | 0.830000 | 0.000999 |
| TCD8em | MacAlv-like | LGALS3 | CLEC7A | 0.268900 | 0.581400 | 0.425200 | 0.000999 |
| TCD8em | MacAlv-like | LGALS3 | FCGR2A | 0.268900 | 0.785400 | 0.527200 | 0.000999 |
| TCD8em | MacAlv-like | LGALS3 | TFRC | 0.268900 | 0.617300 | 0.443100 | 0.000999 |
| TCD8em | MacAlv-like | LRPAP1 | LRP1 | 0.271900 | 0.514900 | 0.393400 | 0.000999 |
| TCD8em | MacAlv-like | LRPAP1 | SORT1 | 0.271900 | 0.283500 | 0.277700 | 0.000999 |
| TCD8em | MacAlv-like | LTB | ITGB1 | 0.635200 | 0.480400 | 0.557800 | 0.000999 |
| TCD8em | MacAlv-like | LTB | TNFRSF1A | 0.635200 | 0.270600 | 0.452900 | 0.000999 |
| TCD8em | MacAlv-like | MIF | CD74 | 0.606300 | 4.191700 | 2.399000 | 0.000999 |
| TCD8em | MacAlv-like | PSAP | LRP1 | 0.501200 | 0.514900 | 0.508100 | 0.000999 |
| TCD8em | MacAlv-like | PTPRC | CD4 | 1.791200 | 0.262400 | 1.026800 | 0.000999 |
| TCD8em | MacAlv-like | PTPRC | CD44 | 1.791200 | 1.485400 | 1.638300 | 0.000999 |
| TCD8em | MacAlv-like | RPS19 | C5AR1 | 3.536000 | 0.535500 | 2.035800 | 0.000999 |
| TCD8em | MacAlv-like | RPS19 | CD4 | 3.536000 | 0.262400 | 1.899200 | 0.000999 |
| TCD8em | MacAlv-like | RTN4 | CD44 | 0.363200 | 1.485400 | 0.924300 | 0.000999 |
| TCD8em | MacAlv-like | SELL | CD4 | 0.258000 | 0.262400 | 0.260200 | 0.000999 |

| | | | | | | | |
|---|---|---|---|---|---|---|---|
| TCD8em | MacAlv-like | SELL | SELPLG | 0.258000 | 0.349200 | 0.303600 | 0.000999 |
| TCD8em | MacAlv-like | SPP1 | CD44 | 0.360800 | 1.485400 | 0.923100 | 0.000999 |
| TCD8em | MacAlv-like | SPP1 | ITGB1 | 0.360800 | 0.480400 | 0.420600 | 0.000999 |
| TCD8em | MacAlv-like | TIMP1 | LRP1 | 0.266700 | 0.514900 | 0.390800 | 0.000999 |
| TCD8em | MacAlv-like | TNF | TNFRSF1A | 0.266700 | 0.270600 | 0.268700 | 0.000999 |
| TCD8em | MacAlv-like | XCL2 | FPR2 | 0.295000 | 0.361100 | 0.328000 | 0.000999 |
| TCD8em | MacAlv-like | XCL2 | GPR65 | 0.295000 | 0.308500 | 0.301700 | 0.000999 |
| TCD8em | MacAlv-like | XCL2 | PTAFR | 0.295000 | 0.307700 | 0.301300 | 0.000999 |
| TCD8ex | MacAlv-like | ALOX5AP | ALOX5 | 1.502400 | 0.873900 | 1.188100 | 0.000999 |
| TCD8ex | MacAlv-like | ANXA1 | C3AR1 | 0.564500 | 0.301300 | 0.432900 | 0.000999 |
| TCD8ex | MacAlv-like | ANXA1 | C5AR1 | 0.564500 | 0.535500 | 0.550000 | 0.000999 |
| TCD8ex | MacAlv-like | ANXA1 | CD44 | 0.564500 | 1.485400 | 1.024900 | 0.000999 |
| TCD8ex | MacAlv-like | ANXA1 | FPR1 | 0.564500 | 0.600900 | 0.582700 | 0.000999 |
| TCD8ex | MacAlv-like | ANXA1 | FPR2 | 0.564500 | 0.361100 | 0.462800 | 0.000999 |
| TCD8ex | MacAlv-like | ANXA1 | FPR3 | 0.564500 | 0.340000 | 0.452200 | 0.000999 |
| TCD8ex | MacAlv-like | APOE | LRP1 | 0.281400 | 0.514900 | 0.398100 | 0.000999 |
| TCD8ex | MacAlv-like | ARF1 | CD4 | 0.783500 | 0.262400 | 0.522900 | 0.000999 |
| TCD8ex | MacAlv-like | ARF1 | CD44 | 0.783500 | 1.485400 | 1.134500 | 0.000999 |
| TCD8ex | MacAlv-like | CALM1 | TNFRSF1A | 1.462700 | 0.270600 | 0.866700 | 0.000999 |
| TCD8ex | MacAlv-like | CALR | CD44 | 0.963400 | 1.485400 | 1.224400 | 0.000999 |
| TCD8ex | MacAlv-like | CALR | IL7R | 0.963400 | 0.251000 | 0.607200 | 0.000999 |
| TCD8ex | MacAlv-like | CALR | LRP1 | 0.963400 | 0.514900 | 0.739200 | 0.000999 |
| TCD8ex | MacAlv-like | CD2 | CD4 | 2.050200 | 0.262400 | 1.156300 | 0.000999 |
| TCD8ex | MacAlv-like | CXCL13 | C3AR1 | 1.774900 | 0.301300 | 1.038100 | 0.000999 |
| TCD8ex | MacAlv-like | CXCL13 | C5AR1 | 1.774900 | 0.535500 | 1.155200 | 0.000999 |
| TCD8ex | MacAlv-like | CXCL13 | FPR1 | 1.774900 | 0.600900 | 1.187900 | 0.000999 |
| TCD8ex | MacAlv-like | CXCL13 | FPR2 | 1.774900 | 0.361100 | 1.068000 | 0.000999 |
| TCD8ex | MacAlv-like | FASLG | TNFRSF1A | 0.261300 | 0.270600 | 0.265900 | 0.000999 |
| TCD8ex | MacAlv-like | GSTP1 | CD44 | 0.828600 | 1.485400 | 1.157000 | 0.000999 |
| TCD8ex | MacAlv-like | GZMB | IGF2R | 1.979400 | 0.265800 | 1.122600 | 0.000999 |

| | | | | | | | |
|---|---|---|---|---|---|---|---|
| TCD8ex | MacAlv-like | HMGB1 | AGTRAP | 1.541400 | 0.266000 | 0.903700 | 0.000999 |
| TCD8ex | MacAlv-like | HMGB1 | LY96 | 1.541400 | 0.944100 | 1.242800 | 0.000999 |
| TCD8ex | MacAlv-like | HMGB2 | CD44 | 0.850000 | 1.485400 | 1.167700 | 0.000999 |
| TCD8ex | MacAlv-like | HMGB2 | DAB2 | 0.850000 | 0.738700 | 0.794300 | 0.000999 |
| TCD8ex | MacAlv-like | HSP90B1 | ITGB1 | 0.566600 | 0.480400 | 0.523500 | 0.000999 |
| TCD8ex | MacAlv-like | HSP90B1 | LRP1 | 0.566600 | 0.514900 | 0.540800 | 0.000999 |
| TCD8ex | MacAlv-like | IFNG | IFNGR1 | 0.857500 | 0.437700 | 0.647600 | 0.000999 |
| TCD8ex | MacAlv-like | IFNG | IFNGR2 | 0.857500 | 0.422900 | 0.640200 | 0.000999 |
| TCD8ex | MacAlv-like | IFNG | PTPN6 | 0.857500 | 0.563800 | 0.710700 | 0.000999 |
| TCD8ex | MacAlv-like | ITGA4 | CD44 | 0.443200 | 1.485400 | 0.964300 | 0.000999 |
| TCD8ex | MacAlv-like | ITGA4 | TFRC | 0.443200 | 0.617300 | 0.530300 | 0.000999 |
| TCD8ex | MacAlv-like | ITGB1 | CD151 | 0.299500 | 0.513700 | 0.406600 | 0.000999 |
| TCD8ex | MacAlv-like | ITGB1 | CD46 | 0.299500 | 0.364700 | 0.332100 | 0.000999 |
| TCD8ex | MacAlv-like | ITGB1 | CD47 | 0.299500 | 0.618300 | 0.458900 | 0.000999 |
| TCD8ex | MacAlv-like | ITGB1 | DAB2 | 0.299500 | 0.738700 | 0.519100 | 0.000999 |
| TCD8ex | MacAlv-like | ITGB1 | ENG | 0.299500 | 0.293500 | 0.296500 | 0.000999 |
| TCD8ex | MacAlv-like | ITGB1 | IGF2R | 0.299500 | 0.265800 | 0.282600 | 0.000999 |
| TCD8ex | MacAlv-like | ITGB1 | PLAUR | 0.299500 | 0.658200 | 0.478900 | 0.000999 |
| TCD8ex | MacAlv-like | ITGB2 | PLAUR | 0.763500 | 0.658200 | 0.710800 | 0.000999 |
| TCD8ex | MacAlv-like | LGALS3 | CLEC7A | 0.382400 | 0.581400 | 0.481900 | 0.000999 |
| TCD8ex | MacAlv-like | LGALS3 | FCGR2A | 0.382400 | 0.785400 | 0.583900 | 0.000999 |
| TCD8ex | MacAlv-like | LRPAP1 | LRP1 | 0.297100 | 0.514900 | 0.406000 | 0.000999 |
| TCD8ex | MacAlv-like | LRPAP1 | SORT1 | 0.297100 | 0.283500 | 0.290300 | 0.000999 |
| TCD8ex | MacAlv-like | LTB | ITGB1 | 0.457600 | 0.480400 | 0.469000 | 0.000999 |
| TCD8ex | MacAlv-like | LTB | TNFRSF1A | 0.457600 | 0.270600 | 0.364100 | 0.000999 |
| TCD8ex | MacAlv-like | MIF | CD74 | 0.749000 | 4.191700 | 2.470400 | 0.000999 |
| TCD8ex | MacAlv-like | PSAP | LRP1 | 0.501600 | 0.514900 | 0.508300 | 0.000999 |
| TCD8ex | MacAlv-like | PTPRC | CD4 | 1.676200 | 0.262400 | 0.969300 | 0.000999 |
| TCD8ex | MacAlv-like | PTPRC | CD44 | 1.676200 | 1.485400 | 1.580800 | 0.000999 |
| TCD8ex | MacAlv-like | RPS19 | C5AR1 | 3.835300 | 0.535500 | 2.185400 | 0.000999 |

| | | | | | | | |
|---|---|---|---|---|---|---|---|
| TCD8ex | MacAlv-like | RPS19 | CD4 | 3.835300 | 0.262400 | 2.048800 | 0.000999 |
| TCD8ex | MacAlv-like | RTN4 | CD44 | 0.303300 | 1.485400 | 0.894300 | 0.000999 |
| TCD8ex | MacAlv-like | SPP1 | CD44 | 0.297800 | 1.485400 | 0.891600 | 0.000999 |
| TCD8ex | MacAlv-like | SPP1 | ITGB1 | 0.297800 | 0.480400 | 0.389100 | 0.000999 |
| TCD8ex | MacAlv-like | VCAM1 | IGF2R | 0.446200 | 0.265800 | 0.356000 | 0.000999 |
| TCD8ex | MacAlv-like | VCAM1 | ITGB1 | 0.446200 | 0.480400 | 0.463300 | 0.000999 |
| TCD8ex | MacAlv-like | VCAM1 | TFRC | 0.446200 | 0.617300 | 0.531800 | 0.000999 |
| TCD8ex | MacAlv-like | XCL2 | FPR2 | 0.315700 | 0.361100 | 0.338400 | 0.000999 |
| TCD8ex | MacAlv-like | XCL2 | PTAFR | 0.315700 | 0.307700 | 0.311700 | 0.000999 |
| TGD | MacAlv-like | ARF1 | CD4 | 0.945300 | 0.262400 | 0.603800 | 0.000999 |
| TGD | MacAlv-like | ARF1 | CD44 | 0.945300 | 1.485400 | 1.215300 | 0.000999 |
| TGD | MacAlv-like | CALM1 | TNFRSF1A | 1.758700 | 0.270600 | 1.014700 | 0.000999 |
| TGD | MacAlv-like | CALR | LRP1 | 0.962700 | 0.514900 | 0.738800 | 0.000999 |
| TGD | MacAlv-like | CD2 | CD4 | 1.452200 | 0.262400 | 0.857300 | 0.000999 |
| TGD | MacAlv-like | CD2 | PTPRC | 1.452200 | 0.690800 | 1.071500 | 0.000999 |
| TGD | MacAlv-like | CD99 | CD99 | 1.715400 | 0.507100 | 1.111200 | 0.000999 |
| TGD | MacAlv-like | COL1A1 | CD44 | 0.451800 | 1.485400 | 0.968600 | 0.000999 |
| TGD | MacAlv-like | COL1A1 | ITGB1 | 0.451800 | 0.480400 | 0.466100 | 0.000999 |
| TGD | MacAlv-like | HMGB1 | AGTRAP | 1.752500 | 0.266000 | 1.009300 | 0.000999 |
| TGD | MacAlv-like | HMGB1 | LY96 | 1.752500 | 0.944100 | 1.348300 | 0.000999 |
| TGD | MacAlv-like | HMGB2 | DAB2 | 0.327500 | 0.738700 | 0.533100 | 0.000999 |
| TGD | MacAlv-like | HSP90B1 | LRP1 | 0.933800 | 0.514900 | 0.724400 | 0.000999 |
| TGD | MacAlv-like | ICAM1 | ITGB2 | 0.291500 | 0.731600 | 0.511600 | 0.000999 |
| TGD | MacAlv-like | ICAM1 | SPN | 0.291500 | 0.541200 | 0.416400 | 0.000999 |
| TGD | MacAlv-like | ICAM3 | ITGB2 | 0.881600 | 0.731600 | 0.806600 | 0.000999 |
| TGD | MacAlv-like | IL16 | CD4 | 0.935700 | 0.262400 | 0.599000 | 0.000999 |
| TGD | MacAlv-like | IL16 | FCGR1A | 0.935700 | 0.490300 | 0.713000 | 0.000999 |
| TGD | MacAlv-like | ITGA4 | CD44 | 0.396800 | 1.485400 | 0.941100 | 0.000999 |
| TGD | MacAlv-like | ITGA4 | CXCR4 | 0.396800 | 0.738800 | 0.567800 | 0.000999 |
| TGD | MacAlv-like | ITGA4 | RPSA | 0.396800 | 1.773500 | 1.085200 | 0.000999 |

| | | | | | | | |
|---|---|---|---|---|---|---|---|
| TGD | MacAlv-like | ITGA4 | TFRC | 0.396800 | 0.617300 | 0.507100 | 0.000999 |
| TGD | MacAlv-like | ITGB2 | PLAUR | 1.444400 | 0.658200 | 1.051300 | 0.000999 |
| TGD | MacAlv-like | LTB | ITGB1 | 1.682900 | 0.480400 | 1.081600 | 0.000999 |
| TGD | MacAlv-like | LTB | TNFRSF1A | 1.682900 | 0.270600 | 0.976800 | 0.000999 |
| TGD | MacAlv-like | MIF | CD74 | 1.633500 | 4.191700 | 2.912600 | 0.000999 |
| TGD | MacAlv-like | PTPRC | CD4 | 1.865300 | 0.262400 | 1.063800 | 0.000999 |
| TGD | MacAlv-like | PTPRC | CD44 | 1.865300 | 1.485400 | 1.675300 | 0.000999 |
| TGD | MacAlv-like | PTPRC | CXCR4 | 1.865300 | 0.738800 | 1.302100 | 0.000999 |
| TGD | MacAlv-like | RPS19 | C5AR1 | 3.890200 | 0.535500 | 2.212900 | 0.000999 |
| TGD | MacAlv-like | RPS19 | CD4 | 3.890200 | 0.262400 | 2.076300 | 0.000999 |
| TGD | MacAlv-like | SELL | CD4 | 0.355500 | 0.262400 | 0.308900 | 0.000999 |
| TGD | MacAlv-like | SELL | SELPLG | 0.355500 | 0.349200 | 0.352400 | 0.000999 |
| TGD | MacAlv-like | SEMA4D | PTPRC | 0.565800 | 0.690800 | 0.628300 | 0.000999 |
| TGD | MacAlv-like | TGFB1 | ENG | 0.770300 | 0.293500 | 0.531900 | 0.000999 |
| TGD | MacAlv-like | TGFB1 | ITGB1 | 0.770300 | 0.480400 | 0.625300 | 0.000999 |
| TGD | MacAlv-like | TGFB1 | TGFBR2 | 0.770300 | 0.259700 | 0.515000 | 0.000999 |
| TGD | MacAlv-like | XCL1 | FPR2 | 1.582200 | 0.361100 | 0.971600 | 0.000999 |
| TGD | MacAlv-like | XCL1 | GPR65 | 1.582200 | 0.308500 | 0.945300 | 0.000999 |
| TGD | MacAlv-like | XCL1 | PTAFR | 1.582200 | 0.307700 | 0.944900 | 0.000999 |
| TGD | MacAlv-like | XCL2 | FPR2 | 0.486400 | 0.361100 | 0.423800 | 0.000999 |
| TGD | MacAlv-like | XCL2 | GPR65 | 0.486400 | 0.308500 | 0.397500 | 0.000999 |
| TGD | MacAlv-like | XCL2 | PTAFR | 0.486400 | 0.307700 | 0.397100 | 0.000999 |
| TGD | MacAlv-like | CALR | CD44 | 0.962700 | 1.485400 | 1.224000 | 0.002000 |
| TGD | MacAlv-like | HMGB2 | CD44 | 0.327500 | 1.485400 | 0.906400 | 0.003000 |
| TGD | MacAlv-like | HSP90B1 | ITGB1 | 0.933800 | 0.480400 | 0.707100 | 0.003000 |

| Supplementary Table 8: Communication results between macrophages and lymphocytes subpopulations, and malignant cells | | | | | | | |
|---|---|---|---|---|---|---|---|
| **MacAlv-like vs. malignant cells** | | | | | | | |
| celltype_01 | celltype_02 | ligand | receptor | ligand_mean_expression | receptor_mean_expression | ligand_receptor_mean_expression | pvalue |
| MacAlv-like | breasttumor | ADAM17 | ITGB1 | 0.5943 | 0.2939 | 0.4441 | 0.000999 |
| MacAlv-like | breasttumor | ADAM17 | TNFRSF1A | 0.5943 | 0.2536 | 0.4239 | 0.000999 |
| MacAlv-like | breasttumor | AIMP1 | RACK1 | 0.3018 | 2.6334 | 1.4676 | 0.000999 |
| MacAlv-like | breasttumor | AIMP1 | RRBP1 | 0.3018 | 0.2835 | 0.2926 | 0.000999 |
| MacAlv-like | breasttumor | ANXA1 | CD44 | 1.7842 | 0.3247 | 1.0545 | 0.000999 |
| MacAlv-like | breasttumor | APOE | SCARB1 | 2.5005 | 0.2628 | 1.3817 | 0.000999 |
| MacAlv-like | breasttumor | APOE | SDC2 | 2.5005 | 1.071 | 1.7858 | 0.000999 |
| MacAlv-like | breasttumor | CALM1 | MYLK | 1.5254 | 0.4064 | 0.9659 | 0.000999 |
| MacAlv-like | breasttumor | CALM1 | RPSA | 1.5254 | 2.358 | 1.9417 | 0.000999 |
| MacAlv-like | breasttumor | CALM1 | TNFRSF1A | 1.5254 | 0.2536 | 0.8895 | 0.000999 |
| MacAlv-like | breasttumor | CALM2 | MYLK | 2.0764 | 0.4064 | 1.2414 | 0.000999 |
| MacAlv-like | breasttumor | CALM3 | MYLK | 0.4437 | 0.4064 | 0.425 | 0.000999 |
| MacAlv-like | breasttumor | FN1 | ITGB1 | 1.4938 | 0.2939 | 0.8938 | 0.000999 |
| MacAlv-like | breasttumor | FN1 | RPSA | 1.4938 | 2.358 | 1.9259 | 0.000999 |
| MacAlv-like | breasttumor | FN1 | SDC2 | 1.4938 | 1.071 | 1.2824 | 0.000999 |
| MacAlv-like | breasttumor | GRN | TNFRSF1A | 2.4715 | 0.2536 | 1.3625 | 0.000999 |
| MacAlv-like | breasttumor | HLA-DRA | SCARB1 | 4.0391 | 0.2628 | 2.1509 | 0.000999 |
| MacAlv-like | breasttumor | ITGB1 | RACK1 | 0.479 | 2.6334 | 1.5562 | 0.000999 |
| MacAlv-like | breasttumor | LGALS3 | ITGB1 | 2.969 | 0.2939 | 1.6314 | 0.000999 |
| MacAlv-like | breasttumor | RPS19 | RPSA | 3.6665 | 2.358 | 3.0123 | 0.000999 |
| MacAlv-like | breasttumor | RTN4 | CD99 | 1.3825 | 0.3871 | 0.8848 | 0.000999 |
| MacAlv-like | breasttumor | SERPINA1 | SDC2 | 2.4202 | 1.071 | 1.7456 | 0.000999 |
| MacAlv-like | breasttumor | TGFB1 | ITGB1 | 0.3653 | 0.2939 | 0.3296 | 0.000999 |
| MacAlv-like | breasttumor | TGFB1 | SDC2 | 0.3653 | 1.071 | 0.7182 | 0.000999 |
| MacAlv-like | breasttumor | THBS1 | ITGB1 | 0.474 | 0.2939 | 0.3839 | 0.000999 |
| MacAlv-like | breasttumor | TNFSF13B | ITGB1 | 0.4439 | 0.2939 | 0.3689 | 0.000999 |
| MacAlv-like | breasttumor | FN1 | CD44 | 1.4938 | 0.3247 | 0.9093 | 0.003 |

| | | | | | | | |
|---|---|---|---|---|---|---|---|
| MacAlv-like | colorectaltumor | AIMP1 | RACK1 | 0.3018 | 2.7146 | 1.5082 | 0.000999 |
| MacAlv-like | colorectaltumor | APOE | SDC1 | 2.5005 | 0.2575 | 1.379 | 0.000999 |
| MacAlv-like | colorectaltumor | CALM1 | RPSA | 1.5254 | 2.251 | 1.8882 | 0.000999 |
| MacAlv-like | colorectaltumor | FN1 | ITGB1 | 1.4938 | 0.4214 | 0.9576 | 0.000999 |
| MacAlv-like | colorectaltumor | FN1 | RPSA | 1.4938 | 2.251 | 1.8724 | 0.000999 |
| MacAlv-like | colorectaltumor | FN1 | SDC1 | 1.4938 | 0.2575 | 0.8756 | 0.000999 |
| MacAlv-like | colorectaltumor | ITGB1 | CDH17 | 0.479 | 0.254 | 0.3665 | 0.000999 |
| MacAlv-like | colorectaltumor | ITGB1 | RACK1 | 0.479 | 2.7146 | 1.5968 | 0.000999 |
| MacAlv-like | colorectaltumor | ITGB1 | SDC1 | 0.479 | 0.2575 | 0.3682 | 0.000999 |
| MacAlv-like | colorectaltumor | LGALS3 | ITGB1 | 2.969 | 0.4214 | 1.6952 | 0.000999 |
| MacAlv-like | colorectaltumor | LGALS3 | ITGB4 | 2.969 | 0.2913 | 1.6301 | 0.000999 |
| MacAlv-like | colorectaltumor | PECAM1 | SDC1 | 0.6441 | 0.2575 | 0.4508 | 0.000999 |
| MacAlv-like | colorectaltumor | RPS19 | RPSA | 3.6665 | 2.251 | 2.9588 | 0.000999 |
| MacAlv-like | colorectaltumor | THBS1 | SDC1 | 0.474 | 0.2575 | 0.3657 | 0.000999 |
| MacAlv-like | colorectaltumor | ITGB1 | CD151 | 0.479 | 0.6439 | 0.5615 | 0.003 |
| MacAlv-like | livertumor | ADAM17 | ITGB1 | 0.5943 | 1.6044 | 1.0994 | 0.000999 |
| MacAlv-like | livertumor | AIMP1 | RACK1 | 0.3018 | 2.7655 | 1.5337 | 0.000999 |
| MacAlv-like | livertumor | ANXA1 | MET | 1.7842 | 0.2983 | 1.0412 | 0.000999 |
| MacAlv-like | livertumor | ANXA1 | PTPRF | 1.7842 | 0.384 | 1.0841 | 0.000999 |
| MacAlv-like | livertumor | APOE | SDC1 | 2.5005 | 0.5354 | 1.518 | 0.000999 |
| MacAlv-like | livertumor | APOE | SDC4 | 2.5005 | 0.6068 | 1.5537 | 0.000999 |
| MacAlv-like | livertumor | ARF1 | INSR | 0.6415 | 0.3039 | 0.4727 | 0.000999 |
| MacAlv-like | livertumor | CALM1 | INSR | 1.5254 | 0.3039 | 0.9147 | 0.000999 |
| MacAlv-like | livertumor | CALM1 | RPSA | 1.5254 | 2.6317 | 2.0785 | 0.000999 |
| MacAlv-like | livertumor | CALR | ITGA3 | 0.8198 | 0.3913 | 0.6056 | 0.000999 |
| MacAlv-like | livertumor | CD99 | EPHB4 | 0.5077 | 0.4535 | 0.4806 | 0.000999 |
| MacAlv-like | livertumor | CD99 | INSR | 0.5077 | 0.3039 | 0.4058 | 0.000999 |
| MacAlv-like | livertumor | FN1 | ITGA2 | 1.4938 | 0.3983 | 0.946 | 0.000999 |
| MacAlv-like | livertumor | FN1 | ITGA3 | 1.4938 | 0.3913 | 0.9425 | 0.000999 |
| MacAlv-like | livertumor | FN1 | ITGB1 | 1.4938 | 1.6044 | 1.5491 | 0.000999 |

| | | | | | | | |
|---|---|---|---|---|---|---|---|
| MacAlv-like | livertumor | FN1 | RPSA | 1.4938 | 2.6317 | 2.0628 | 0.000999 |
| MacAlv-like | livertumor | FN1 | SDC1 | 1.4938 | 0.5354 | 1.0146 | 0.000999 |
| MacAlv-like | livertumor | HLA-A | ERBB3 | 2.47 | 0.2644 | 1.3672 | 0.000999 |
| MacAlv-like | livertumor | HLA-DRA | EPHB4 | 4.0391 | 0.4535 | 2.2463 | 0.000999 |
| MacAlv-like | livertumor | HLA-E | EPHB4 | 1.2621 | 0.4535 | 0.8578 | 0.000999 |
| MacAlv-like | livertumor | HLA-E | ITGA6 | 1.2621 | 0.5211 | 0.8916 | 0.000999 |
| MacAlv-like | livertumor | HLA-E | PLXNB2 | 1.2621 | 0.265 | 0.7635 | 0.000999 |
| MacAlv-like | livertumor | HMGB2 | CDH1 | 0.2514 | 0.4896 | 0.3705 | 0.000999 |
| MacAlv-like | livertumor | HMGB2 | DSG2 | 0.2514 | 0.7479 | 0.4997 | 0.000999 |
| MacAlv-like | livertumor | HSP90B1 | ITGB1 | 0.7612 | 1.6044 | 1.1828 | 0.000999 |
| MacAlv-like | livertumor | ITGB1 | CD151 | 0.479 | 1.1085 | 0.7938 | 0.000999 |
| MacAlv-like | livertumor | ITGB1 | ITGA2 | 0.479 | 0.3983 | 0.4387 | 0.000999 |
| MacAlv-like | livertumor | ITGB1 | ITGA3 | 0.479 | 0.3913 | 0.4352 | 0.000999 |
| MacAlv-like | livertumor | ITGB1 | ITGA6 | 0.479 | 0.5211 | 0.5001 | 0.000999 |
| MacAlv-like | livertumor | ITGB1 | RACK1 | 0.479 | 2.7655 | 1.6223 | 0.000999 |
| MacAlv-like | livertumor | ITGB1 | SDC1 | 0.479 | 0.5354 | 0.5072 | 0.000999 |
| MacAlv-like | livertumor | ITGB1 | VTN | 0.479 | 0.7399 | 0.6095 | 0.000999 |
| MacAlv-like | livertumor | LGALS3 | ITGA2 | 2.969 | 0.3983 | 1.6836 | 0.000999 |
| MacAlv-like | livertumor | LGALS3 | ITGB1 | 2.969 | 1.6044 | 2.2867 | 0.000999 |
| MacAlv-like | livertumor | LGALS3 | ITGB4 | 2.969 | 0.5518 | 1.7604 | 0.000999 |
| MacAlv-like | livertumor | NAMPT | INSR | 0.7729 | 0.3039 | 0.5384 | 0.000999 |
| MacAlv-like | livertumor | PECAM1 | SDC1 | 0.6441 | 0.5354 | 0.5898 | 0.000999 |
| MacAlv-like | livertumor | PECAM1 | SDC4 | 0.6441 | 0.6068 | 0.6255 | 0.000999 |
| MacAlv-like | livertumor | PTPRC | INSR | 0.6901 | 0.3039 | 0.497 | 0.000999 |
| MacAlv-like | livertumor | PTPRC | MET | 0.6901 | 0.2983 | 0.4942 | 0.000999 |
| MacAlv-like | livertumor | RPS19 | RPSA | 3.6665 | 2.6317 | 3.1491 | 0.000999 |
| MacAlv-like | livertumor | RTN4 | CD99 | 1.3825 | 0.5906 | 0.9866 | 0.000999 |
| MacAlv-like | livertumor | TGFB1 | ITGB1 | 0.3653 | 1.6044 | 0.9849 | 0.000999 |
| MacAlv-like | livertumor | TGFB1 | VTN | 0.3653 | 0.7399 | 0.5526 | 0.000999 |
| MacAlv-like | livertumor | THBS1 | ITGA3 | 0.474 | 0.3913 | 0.4327 | 0.000999 |

| | | | | | | | |
|---|---|---|---|---|---|---|---|
| MacAlv-like | livertumor | THBS1 | ITGB1 | 0.474 | 1.6044 | 1.0392 | 0.000999 |
| MacAlv-like | livertumor | THBS1 | SDC1 | 0.474 | 0.5354 | 0.5047 | 0.000999 |
| MacAlv-like | livertumor | TNFSF12 | TNFRSF12A | 0.2978 | 1.0484 | 0.6731 | 0.000999 |
| MacAlv-like | livertumor | TNFSF13B | ITGB1 | 0.4439 | 1.6044 | 1.0242 | 0.000999 |
| MacAlv-like | livertumor | LGALS3 | TFRC | 2.969 | 0.2899 | 1.6294 | 0.002 |
| MacAlv-like | livertumor | AIMP1 | RRBP1 | 0.3018 | 0.355 | 0.3284 | 0.004 |
| MacAlv-like | lungtumor | ANXA1 | CXCR4 | 1.7842 | 0.2684 | 1.0263 | 0.000999 |
| MacAlv-like | lungtumor | APOE | SDC1 | 2.5005 | 0.2715 | 1.386 | 0.000999 |
| MacAlv-like | lungtumor | APOE | SDC4 | 2.5005 | 0.3516 | 1.4261 | 0.000999 |
| MacAlv-like | lungtumor | CALM1 | RPSA | 1.5254 | 1.6057 | 1.5656 | 0.000999 |
| MacAlv-like | lungtumor | CXCL16 | CXCR4 | 1.1328 | 0.2684 | 0.7006 | 0.000999 |
| MacAlv-like | lungtumor | CXCL3 | CXCR4 | 0.6506 | 0.2684 | 0.4595 | 0.000999 |
| MacAlv-like | lungtumor | FN1 | CD44 | 1.4938 | 0.3802 | 0.937 | 0.000999 |
| MacAlv-like | lungtumor | FN1 | RPSA | 1.4938 | 1.6057 | 1.5498 | 0.000999 |
| MacAlv-like | lungtumor | FN1 | SDC1 | 1.4938 | 0.2715 | 0.8827 | 0.000999 |
| MacAlv-like | lungtumor | ITGB1 | CD151 | 0.479 | 0.6776 | 0.5783 | 0.000999 |
| MacAlv-like | lungtumor | ITGB1 | CD46 | 0.479 | 0.434 | 0.4565 | 0.000999 |
| MacAlv-like | lungtumor | ITGB1 | CD47 | 0.479 | 0.3255 | 0.4023 | 0.000999 |
| MacAlv-like | lungtumor | ITGB1 | RACK1 | 0.479 | 2.2108 | 1.3449 | 0.000999 |
| MacAlv-like | lungtumor | ITGB1 | SDC1 | 0.479 | 0.2715 | 0.3753 | 0.000999 |
| MacAlv-like | lungtumor | PECAM1 | CXCR4 | 0.6441 | 0.2684 | 0.4562 | 0.000999 |
| MacAlv-like | lungtumor | PECAM1 | SDC1 | 0.6441 | 0.2715 | 0.4578 | 0.000999 |
| MacAlv-like | lungtumor | PECAM1 | SDC4 | 0.6441 | 0.3516 | 0.4978 | 0.000999 |
| MacAlv-like | lungtumor | PTPRC | CD44 | 0.6901 | 0.3802 | 0.5352 | 0.000999 |
| MacAlv-like | lungtumor | PTPRC | CXCR4 | 0.6901 | 0.2684 | 0.4793 | 0.000999 |
| MacAlv-like | lungtumor | RTN4 | CD44 | 1.3825 | 0.3802 | 0.8813 | 0.000999 |
| MacAlv-like | lungtumor | RTN4 | CD99 | 1.3825 | 0.5938 | 0.9882 | 0.000999 |
| MacAlv-like | lungtumor | THBS1 | CD47 | 0.474 | 0.3255 | 0.3998 | 0.000999 |
| MacAlv-like | lungtumor | THBS1 | SDC1 | 0.474 | 0.2715 | 0.3728 | 0.000999 |
| MacAlv-like | lungtumor | TNFSF12 | TNFRSF12A | 0.2978 | 0.441 | 0.3694 | 0.000999 |

| | | | | | | | |
|---|---|---|---|---|---|---|---|
| MacAlv-like | lungtumor | RPS19 | RPSA | 3.6665 | 1.6057 | 2.6361 | 0.00599 |
| MacAlv-like | melanomatumo | AIMP1 | RACK1 | 0.3018 | 4.145 | 2.2234 | 0.000999 |
| MacAlv-like | melanomatumo | ANXA1 | EDNRB | 1.7842 | 0.5666 | 1.1754 | 0.000999 |
| MacAlv-like | melanomatumo | CALM1 | RPSA | 1.5254 | 3.7787 | 2.652 | 0.000999 |
| MacAlv-like | melanomatumo | FN1 | RPSA | 1.4938 | 3.7787 | 2.6362 | 0.000999 |
| MacAlv-like | melanomatumo | ITGB1 | RACK1 | 0.479 | 4.145 | 2.312 | 0.000999 |
| MacAlv-like | melanomatumo | MIF | TNFRSF14 | 0.585 | 0.5069 | 0.5459 | 0.000999 |
| MacAlv-like | melanomatumo | NAMPT | ANXA6 | 0.7729 | 0.3625 | 0.5677 | 0.000999 |
| MacAlv-like | melanomatumo | NMB | EDNRB | 0.3046 | 0.5666 | 0.4356 | 0.000999 |
| MacAlv-like | melanomatumo | RPS19 | RPSA | 3.6665 | 3.7787 | 3.7226 | 0.000999 |
| MacAlv-like | ovarytumor | ADAM17 | ITGB1 | 0.5943 | 0.2783 | 0.4363 | 0.000999 |
| MacAlv-like | ovarytumor | AIMP1 | RACK1 | 0.3018 | 2.9543 | 1.628 | 0.000999 |
| MacAlv-like | ovarytumor | ANXA1 | CXCR4 | 1.7842 | 0.3688 | 1.0765 | 0.000999 |
| MacAlv-like | ovarytumor | CALM1 | RPSA | 1.5254 | 2.9542 | 2.2398 | 0.000999 |
| MacAlv-like | ovarytumor | CCL20 | CXCR4 | 0.2948 | 0.3688 | 0.3318 | 0.000999 |
| MacAlv-like | ovarytumor | CXCL16 | CXCR4 | 1.1328 | 0.3688 | 0.7508 | 0.000999 |
| MacAlv-like | ovarytumor | CXCL2 | CXCR4 | 0.2923 | 0.3688 | 0.3305 | 0.000999 |
| MacAlv-like | ovarytumor | CXCL3 | CXCR4 | 0.6506 | 0.3688 | 0.5097 | 0.000999 |
| MacAlv-like | ovarytumor | FN1 | ITGB1 | 1.4938 | 0.2783 | 0.886 | 0.000999 |
| MacAlv-like | ovarytumor | FN1 | RPSA | 1.4938 | 2.9542 | 2.224 | 0.000999 |
| MacAlv-like | ovarytumor | ITGB1 | CD151 | 0.479 | 0.4981 | 0.4886 | 0.000999 |
| MacAlv-like | ovarytumor | ITGB1 | CD47 | 0.479 | 0.4931 | 0.486 | 0.000999 |
| MacAlv-like | ovarytumor | ITGB1 | RACK1 | 0.479 | 2.9543 | 1.7166 | 0.000999 |
| MacAlv-like | ovarytumor | LGALS3 | ITGB1 | 2.969 | 0.2783 | 1.6236 | 0.000999 |
| MacAlv-like | ovarytumor | PECAM1 | CXCR4 | 0.6441 | 0.3688 | 0.5064 | 0.000999 |
| MacAlv-like | ovarytumor | PTPRC | CXCR4 | 0.6901 | 0.3688 | 0.5294 | 0.000999 |
| MacAlv-like | ovarytumor | TGFB1 | ITGB1 | 0.3653 | 0.2783 | 0.3218 | 0.000999 |
| MacAlv-like | ovarytumor | THBS1 | CD47 | 0.474 | 0.4931 | 0.4835 | 0.000999 |
| MacAlv-like | ovarytumor | THBS1 | ITGB1 | 0.474 | 0.2783 | 0.3761 | 0.000999 |
| MacAlv-like | ovarytumor | TNFSF12 | TNFRSF12A | 0.2978 | 0.3901 | 0.3439 | 0.000999 |

| | | | | | | | |
|---|---|---|---|---|---|---|---|
| MacAlv-like | ovarytumor | TNFSF13B | ITGB1 | 0.4439 | 0.2783 | 0.3611 | 0.000999 |
| MacAlv-like | ovarytumor | RPS19 | RPSA | 3.6665 | 2.9542 | 3.3104 | 0.003 |
| MacAlv-like | ovarytumor | HSP90B1 | ITGB1 | 0.7612 | 0.2783 | 0.5197 | 0.00799 |
| breasttumor | MacAlv-like | A2M | LRP1 | 0.3129 | 0.5137 | 0.4133 | 0.000999 |
| breasttumor | MacAlv-like | APP | ALOX5 | 0.5409 | 0.8725 | 0.7067 | 0.000999 |
| breasttumor | MacAlv-like | APP | C3AR1 | 0.5409 | 0.3017 | 0.4213 | 0.000999 |
| breasttumor | MacAlv-like | APP | C5AR1 | 0.5409 | 0.5353 | 0.5381 | 0.000999 |
| breasttumor | MacAlv-like | APP | CXCR4 | 0.5409 | 0.7383 | 0.6396 | 0.000999 |
| breasttumor | MacAlv-like | APP | FPR1 | 0.5409 | 0.601 | 0.571 | 0.000999 |
| breasttumor | MacAlv-like | APP | FPR2 | 0.5409 | 0.3607 | 0.4508 | 0.000999 |
| breasttumor | MacAlv-like | APP | LRP1 | 0.5409 | 0.5137 | 0.5273 | 0.000999 |
| breasttumor | MacAlv-like | APP | TGFBR2 | 0.5409 | 0.2593 | 0.4001 | 0.000999 |
| breasttumor | MacAlv-like | ARF1 | CD4 | 1.6219 | 0.2618 | 0.9418 | 0.000999 |
| breasttumor | MacAlv-like | ARF1 | CD44 | 1.6219 | 1.4851 | 1.5535 | 0.000999 |
| breasttumor | MacAlv-like | CALR | CD44 | 1.1835 | 1.4851 | 1.3343 | 0.000999 |
| breasttumor | MacAlv-like | CALR | IL7R | 1.1835 | 0.2513 | 0.7174 | 0.000999 |
| breasttumor | MacAlv-like | CALR | LRP1 | 1.1835 | 0.5137 | 0.8486 | 0.000999 |
| breasttumor | MacAlv-like | COL2A1 | CD44 | 0.3935 | 1.4851 | 0.9393 | 0.000999 |
| breasttumor | MacAlv-like | COL2A1 | ITGB1 | 0.3935 | 0.479 | 0.4363 | 0.000999 |
| breasttumor | MacAlv-like | GSTP1 | CD44 | 2.5435 | 1.4851 | 2.0143 | 0.000999 |
| breasttumor | MacAlv-like | GSTP1 | IL7R | 2.5435 | 0.2513 | 1.3974 | 0.000999 |
| breasttumor | MacAlv-like | HEBP1 | C3AR1 | 0.4062 | 0.3017 | 0.3539 | 0.000999 |
| breasttumor | MacAlv-like | HEBP1 | C5AR1 | 0.4062 | 0.5353 | 0.4707 | 0.000999 |
| breasttumor | MacAlv-like | HEBP1 | CXCR4 | 0.4062 | 0.7383 | 0.5723 | 0.000999 |
| breasttumor | MacAlv-like | HEBP1 | FPR1 | 0.4062 | 0.601 | 0.5036 | 0.000999 |
| breasttumor | MacAlv-like | HEBP1 | FPR2 | 0.4062 | 0.3607 | 0.3834 | 0.000999 |
| breasttumor | MacAlv-like | HEBP1 | FPR3 | 0.4062 | 0.3404 | 0.3733 | 0.000999 |
| breasttumor | MacAlv-like | HMGB1 | LY96 | 1.1815 | 0.9431 | 1.0623 | 0.000999 |
| breasttumor | MacAlv-like | HMGB2 | CD44 | 0.3139 | 1.4851 | 0.8995 | 0.000999 |
| breasttumor | MacAlv-like | HMGB2 | DAB2 | 0.3139 | 0.7389 | 0.5264 | 0.000999 |

| | | | | | | | |
|---|---|---|---|---|---|---|---|
| breasttumor | MacAlv-like | HSP90B1 | ITGB1 | 0.7786 | 0.479 | 0.6288 | 0.000999 |
| breasttumor | MacAlv-like | HSP90B1 | LRP1 | 0.7786 | 0.5137 | 0.6461 | 0.000999 |
| breasttumor | MacAlv-like | IGF2 | IGF2R | 1.9535 | 0.2661 | 1.1098 | 0.000999 |
| breasttumor | MacAlv-like | ITGB1 | CD47 | 0.2939 | 0.6172 | 0.4555 | 0.000999 |
| breasttumor | MacAlv-like | ITGB1 | DAB2 | 0.2939 | 0.7389 | 0.5164 | 0.000999 |
| breasttumor | MacAlv-like | ITGB1 | PLAUR | 0.2939 | 0.6578 | 0.4758 | 0.000999 |
| breasttumor | MacAlv-like | MDK | LRP1 | 0.592 | 0.5137 | 0.5528 | 0.000999 |
| breasttumor | MacAlv-like | MIF | CD74 | 3.2716 | 4.1914 | 3.7315 | 0.000999 |
| breasttumor | MacAlv-like | NPW | C3AR1 | 0.5478 | 0.3017 | 0.4248 | 0.000999 |
| breasttumor | MacAlv-like | NPW | C5AR1 | 0.5478 | 0.5353 | 0.5416 | 0.000999 |
| breasttumor | MacAlv-like | NPW | CXCR4 | 0.5478 | 0.7383 | 0.6431 | 0.000999 |
| breasttumor | MacAlv-like | NPW | FPR1 | 0.5478 | 0.601 | 0.5744 | 0.000999 |
| breasttumor | MacAlv-like | NPW | FPR2 | 0.5478 | 0.3607 | 0.4543 | 0.000999 |
| breasttumor | MacAlv-like | RPS19 | C5AR1 | 3.691 | 0.5353 | 2.1132 | 0.000999 |
| breasttumor | MacAlv-like | RPS19 | CD4 | 3.691 | 0.2618 | 1.9764 | 0.000999 |
| breasttumor | MacAlv-like | SERPINE2 | LRP1 | 0.6001 | 0.5137 | 0.5569 | 0.000999 |
| breasttumor | MacAlv-like | TF | CD4 | 0.2688 | 0.2618 | 0.2653 | 0.000999 |
| breasttumor | MacAlv-like | TF | IGF2R | 0.2688 | 0.2661 | 0.2675 | 0.000999 |
| breasttumor | MacAlv-like | TF | IL7R | 0.2688 | 0.2513 | 0.26 | 0.000999 |
| breasttumor | MacAlv-like | TF | TFRC | 0.2688 | 0.6181 | 0.4435 | 0.000999 |
| breasttumor | MacAlv-like | TIMP1 | LRP1 | 0.7045 | 0.5137 | 0.6091 | 0.000999 |
| breasttumor | MacAlv-like | ITGB1 | ENG | 0.2939 | 0.2939 | 0.2939 | 0.005 |
| colorectaltumor | MacAlv-like | APP | ALOX5 | 0.4966 | 0.8725 | 0.6845 | 0.000999 |
| colorectaltumor | MacAlv-like | APP | C3AR1 | 0.4966 | 0.3017 | 0.3992 | 0.000999 |
| colorectaltumor | MacAlv-like | APP | C5AR1 | 0.4966 | 0.5353 | 0.516 | 0.000999 |
| colorectaltumor | MacAlv-like | APP | CXCR4 | 0.4966 | 0.7383 | 0.6175 | 0.000999 |
| colorectaltumor | MacAlv-like | APP | FPR1 | 0.4966 | 0.601 | 0.5488 | 0.000999 |
| colorectaltumor | MacAlv-like | APP | FPR2 | 0.4966 | 0.3607 | 0.4287 | 0.000999 |
| colorectaltumor | MacAlv-like | APP | LRP1 | 0.4966 | 0.5137 | 0.5051 | 0.000999 |
| colorectaltumor | MacAlv-like | APP | TGFBR2 | 0.4966 | 0.2593 | 0.378 | 0.000999 |

| | | | | | | | |
|---|---|---|---|---|---|---|---|
| colorectaltumor | MacAlv-like | ARF1 | CD44 | 0.6489 | 1.4851 | 1.067 | 0.000999 |
| colorectaltumor | MacAlv-like | COL1A1 | CD44 | 0.4875 | 1.4851 | 0.9863 | 0.000999 |
| colorectaltumor | MacAlv-like | COL1A1 | ITGB1 | 0.4875 | 0.479 | 0.4833 | 0.000999 |
| colorectaltumor | MacAlv-like | GSTP1 | CD44 | 2.8039 | 1.4851 | 2.1445 | 0.000999 |
| colorectaltumor | MacAlv-like | GSTP1 | IL7R | 2.8039 | 0.2513 | 1.5276 | 0.000999 |
| colorectaltumor | MacAlv-like | HMGB1 | AGTRAP | 2.155 | 0.2657 | 1.2103 | 0.000999 |
| colorectaltumor | MacAlv-like | HMGB1 | LY96 | 2.155 | 0.9431 | 1.549 | 0.000999 |
| colorectaltumor | MacAlv-like | HMGB2 | CD44 | 0.5347 | 1.4851 | 1.0099 | 0.000999 |
| colorectaltumor | MacAlv-like | HMGB2 | DAB2 | 0.5347 | 0.7389 | 0.6368 | 0.000999 |
| colorectaltumor | MacAlv-like | MDK | LRP1 | 0.7548 | 0.5137 | 0.6343 | 0.000999 |
| colorectaltumor | MacAlv-like | MIF | CD74 | 1.6741 | 4.1914 | 2.9327 | 0.000999 |
| colorectaltumor | MacAlv-like | RPS19 | C5AR1 | 4.0956 | 0.5353 | 2.3154 | 0.000999 |
| colorectaltumor | MacAlv-like | RPS19 | CD4 | 4.0956 | 0.2618 | 2.1787 | 0.000999 |
| colorectaltumor | MacAlv-like | RPS19 | RPSA | 4.0956 | 1.7739 | 2.9347 | 0.000999 |
| colorectaltumor | MacAlv-like | TIMP1 | LRP1 | 0.8931 | 0.5137 | 0.7034 | 0.000999 |
| colorectaltumor | MacAlv-like | AIMP1 | RACK1 | 0.5265 | 2.1551 | 1.3408 | 0.00899 |
| livertumor | MacAlv-like | AGRN | LRP1 | 0.3492 | 0.5137 | 0.4315 | 0.000999 |
| livertumor | MacAlv-like | AGT | ANPEP | 0.4701 | 0.274 | 0.372 | 0.000999 |
| livertumor | MacAlv-like | AGT | ATP6AP2 | 0.4701 | 0.6172 | 0.5436 | 0.000999 |
| livertumor | MacAlv-like | AGT | C3AR1 | 0.4701 | 0.3017 | 0.3859 | 0.000999 |
| livertumor | MacAlv-like | AGT | C5AR1 | 0.4701 | 0.5353 | 0.5027 | 0.000999 |
| livertumor | MacAlv-like | AGT | CXCR4 | 0.4701 | 0.7383 | 0.6042 | 0.000999 |
| livertumor | MacAlv-like | AGT | FPR1 | 0.4701 | 0.601 | 0.5355 | 0.000999 |
| livertumor | MacAlv-like | AGT | FPR2 | 0.4701 | 0.3607 | 0.4154 | 0.000999 |
| livertumor | MacAlv-like | AGT | GPR65 | 0.4701 | 0.3089 | 0.3895 | 0.000999 |
| livertumor | MacAlv-like | AGT | PTAFR | 0.4701 | 0.3081 | 0.3891 | 0.000999 |
| livertumor | MacAlv-like | APOA1 | DERL1 | 0.2888 | 0.3024 | 0.2956 | 0.000999 |
| livertumor | MacAlv-like | APOA1 | IL7R | 0.2888 | 0.2513 | 0.27 | 0.000999 |
| livertumor | MacAlv-like | APOA1 | LRP1 | 0.2888 | 0.5137 | 0.4012 | 0.000999 |
| livertumor | MacAlv-like | APOA1 | MSR1 | 0.2888 | 1.4629 | 0.8758 | 0.000999 |

| | | | | | | | |
|---|---|---|---|---|---|---|---|
| livertumor | MacAlv-like | APP | ALOX5 | 0.5626 | 0.8725 | 0.7175 | 0.000999 |
| livertumor | MacAlv-like | APP | C3AR1 | 0.5626 | 0.3017 | 0.4321 | 0.000999 |
| livertumor | MacAlv-like | APP | C5AR1 | 0.5626 | 0.5353 | 0.5489 | 0.000999 |
| livertumor | MacAlv-like | APP | CXCR4 | 0.5626 | 0.7383 | 0.6504 | 0.000999 |
| livertumor | MacAlv-like | APP | FPR1 | 0.5626 | 0.601 | 0.5818 | 0.000999 |
| livertumor | MacAlv-like | APP | FPR2 | 0.5626 | 0.3607 | 0.4616 | 0.000999 |
| livertumor | MacAlv-like | APP | LRP1 | 0.5626 | 0.5137 | 0.5381 | 0.000999 |
| livertumor | MacAlv-like | APP | TGFBR2 | 0.5626 | 0.2593 | 0.4109 | 0.000999 |
| livertumor | MacAlv-like | ARF1 | CD44 | 0.6936 | 1.4851 | 1.0893 | 0.000999 |
| livertumor | MacAlv-like | C3 | C3AR1 | 0.6584 | 0.3017 | 0.4801 | 0.000999 |
| livertumor | MacAlv-like | C3 | C5AR1 | 0.6584 | 0.5353 | 0.5969 | 0.000999 |
| livertumor | MacAlv-like | C3 | CXCR4 | 0.6584 | 0.7383 | 0.6984 | 0.000999 |
| livertumor | MacAlv-like | C3 | FPR1 | 0.6584 | 0.601 | 0.6297 | 0.000999 |
| livertumor | MacAlv-like | C3 | FPR2 | 0.6584 | 0.3607 | 0.5096 | 0.000999 |
| livertumor | MacAlv-like | C3 | ITGB2 | 0.6584 | 0.7326 | 0.6955 | 0.000999 |
| livertumor | MacAlv-like | C3 | LRP1 | 0.6584 | 0.5137 | 0.5861 | 0.000999 |
| livertumor | MacAlv-like | CALR | CD44 | 1.0416 | 1.4851 | 1.2634 | 0.000999 |
| livertumor | MacAlv-like | CALR | IL7R | 1.0416 | 0.2513 | 0.6465 | 0.000999 |
| livertumor | MacAlv-like | CALR | LRP1 | 1.0416 | 0.5137 | 0.7776 | 0.000999 |
| livertumor | MacAlv-like | CCL20 | C3AR1 | 1.0877 | 0.3017 | 0.6947 | 0.000999 |
| livertumor | MacAlv-like | CCL20 | C5AR1 | 1.0877 | 0.5353 | 0.8115 | 0.000999 |
| livertumor | MacAlv-like | CCL20 | CXCR4 | 1.0877 | 0.7383 | 0.913 | 0.000999 |
| livertumor | MacAlv-like | CCL20 | FPR1 | 1.0877 | 0.601 | 0.8443 | 0.000999 |
| livertumor | MacAlv-like | CCL20 | FPR2 | 1.0877 | 0.3607 | 0.7242 | 0.000999 |
| livertumor | MacAlv-like | CDH1 | RACK1 | 0.4896 | 2.1551 | 1.3224 | 0.000999 |
| livertumor | MacAlv-like | CXCL2 | C5AR1 | 0.3364 | 0.5353 | 0.4358 | 0.000999 |
| livertumor | MacAlv-like | CXCL2 | CXCR4 | 0.3364 | 0.7383 | 0.5373 | 0.000999 |
| livertumor | MacAlv-like | CXCL2 | FPR1 | 0.3364 | 0.601 | 0.4687 | 0.000999 |
| livertumor | MacAlv-like | DSG2 | CD44 | 0.7479 | 1.4851 | 1.1165 | 0.000999 |
| livertumor | MacAlv-like | DSG2 | CXCR4 | 0.7479 | 0.7383 | 0.7431 | 0.000999 |

| | | | | | | | |
|---|---|---|---|---|---|---|---|
| livertumor | MacAlv-like | DSG2 | DAB2 | 0.7479 | 0.7389 | 0.7434 | 0.000999 |
| livertumor | MacAlv-like | FGA | ITGB2 | 0.2688 | 0.7326 | 0.5007 | 0.000999 |
| livertumor | MacAlv-like | HMGB1 | AGTRAP | 1.4361 | 0.2657 | 0.8509 | 0.000999 |
| livertumor | MacAlv-like | HMGB1 | LY96 | 1.4361 | 0.9431 | 1.1896 | 0.000999 |
| livertumor | MacAlv-like | HSP90B1 | LRP1 | 0.8951 | 0.5137 | 0.7044 | 0.000999 |
| livertumor | MacAlv-like | HSPG2 | LRP1 | 0.2746 | 0.5137 | 0.3942 | 0.000999 |
| livertumor | MacAlv-like | ITGB1 | CD151 | 1.6044 | 0.5136 | 1.059 | 0.000999 |
| livertumor | MacAlv-like | ITGB1 | CD46 | 1.6044 | 0.3652 | 0.9848 | 0.000999 |
| livertumor | MacAlv-like | ITGB1 | CD47 | 1.6044 | 0.6172 | 1.1108 | 0.000999 |
| livertumor | MacAlv-like | ITGB1 | DAB2 | 1.6044 | 0.7389 | 1.1717 | 0.000999 |
| livertumor | MacAlv-like | ITGB1 | ENG | 1.6044 | 0.2939 | 0.9492 | 0.000999 |
| livertumor | MacAlv-like | ITGB1 | IGF2R | 1.6044 | 0.2661 | 0.9353 | 0.000999 |
| livertumor | MacAlv-like | ITGB1 | PLAUR | 1.6044 | 0.6578 | 1.1311 | 0.000999 |
| livertumor | MacAlv-like | ITGB1 | RACK1 | 1.6044 | 2.1551 | 1.8798 | 0.000999 |
| livertumor | MacAlv-like | LAMC2 | CD151 | 0.4847 | 0.5136 | 0.4992 | 0.000999 |
| livertumor | MacAlv-like | LAMC2 | ITGB1 | 0.4847 | 0.479 | 0.4818 | 0.000999 |
| livertumor | MacAlv-like | MDK | LRP1 | 1.6101 | 0.5137 | 1.0619 | 0.000999 |
| livertumor | MacAlv-like | MIF | CD74 | 1.7638 | 4.1914 | 2.9776 | 0.000999 |
| livertumor | MacAlv-like | NTS | FPR2 | 0.3425 | 0.3607 | 0.3516 | 0.000999 |
| livertumor | MacAlv-like | NTS | GPR65 | 0.3425 | 0.3089 | 0.3257 | 0.000999 |
| livertumor | MacAlv-like | NTS | PTAFR | 0.3425 | 0.3081 | 0.3253 | 0.000999 |
| livertumor | MacAlv-like | NTS | SORT1 | 0.3425 | 0.2831 | 0.3128 | 0.000999 |
| livertumor | MacAlv-like | OCLN | TGFBR2 | 0.3091 | 0.2593 | 0.2842 | 0.000999 |
| livertumor | MacAlv-like | PTPRF | RACK1 | 0.384 | 2.1551 | 1.2696 | 0.000999 |
| livertumor | MacAlv-like | PTPRF | TGFBR2 | 0.384 | 0.2593 | 0.3217 | 0.000999 |
| livertumor | MacAlv-like | RPS19 | C5AR1 | 4.0772 | 0.5353 | 2.3062 | 0.000999 |
| livertumor | MacAlv-like | RPS19 | CD4 | 4.0772 | 0.2618 | 2.1695 | 0.000999 |
| livertumor | MacAlv-like | RPS19 | RPSA | 4.0772 | 1.7739 | 2.9255 | 0.000999 |
| livertumor | MacAlv-like | SPP1 | CD44 | 1.0048 | 1.4851 | 1.2449 | 0.000999 |
| livertumor | MacAlv-like | SPP1 | ITGB1 | 1.0048 | 0.479 | 0.7419 | 0.000999 |

| | | | | | | | |
|---|---|---|---|---|---|---|---|
| livertumor | MacAlv-like | TF | CD4 | 0.523 | 0.2618 | 0.3924 | 0.000999 |
| livertumor | MacAlv-like | TF | IGF2R | 0.523 | 0.2661 | 0.3946 | 0.000999 |
| livertumor | MacAlv-like | TF | IL7R | 0.523 | 0.2513 | 0.3871 | 0.000999 |
| livertumor | MacAlv-like | TF | TFRC | 0.523 | 0.6181 | 0.5706 | 0.000999 |
| livertumor | MacAlv-like | TFPI | LRP1 | 0.5043 | 0.5137 | 0.509 | 0.000999 |
| livertumor | MacAlv-like | TIMP1 | LRP1 | 1.0257 | 0.5137 | 0.7697 | 0.000999 |
| livertumor | MacAlv-like | TTR | IL7R | 0.4579 | 0.2513 | 0.3546 | 0.000999 |
| livertumor | MacAlv-like | VEGFA | AXL | 0.5485 | 0.4377 | 0.4931 | 0.000999 |
| livertumor | MacAlv-like | VTN | ITGB1 | 0.7399 | 0.479 | 0.6095 | 0.000999 |
| livertumor | MacAlv-like | VTN | PLAUR | 0.7399 | 0.6578 | 0.6988 | 0.000999 |
| livertumor | MacAlv-like | CXCL2 | C3AR1 | 0.3364 | 0.3017 | 0.319 | 0.005 |
| livertumor | MacAlv-like | CXCL2 | FPR2 | 0.3364 | 0.3607 | 0.3485 | 0.005 |
| livertumor | MacAlv-like | ARF1 | CD4 | 0.6936 | 0.2618 | 0.4777 | 0.00699 |
| lungtumor | MacAlv-like | ANXA1 | C3AR1 | 1.4026 | 0.3017 | 0.8522 | 0.000999 |
| lungtumor | MacAlv-like | ANXA1 | C5AR1 | 1.4026 | 0.5353 | 0.969 | 0.000999 |
| lungtumor | MacAlv-like | ANXA1 | CD4 | 1.4026 | 0.2618 | 0.8322 | 0.000999 |
| lungtumor | MacAlv-like | ANXA1 | CD44 | 1.4026 | 1.4851 | 1.4439 | 0.000999 |
| lungtumor | MacAlv-like | ANXA1 | CXCR4 | 1.4026 | 0.7383 | 1.0705 | 0.000999 |
| lungtumor | MacAlv-like | ANXA1 | FPR1 | 1.4026 | 0.601 | 1.0018 | 0.000999 |
| lungtumor | MacAlv-like | ANXA1 | FPR2 | 1.4026 | 0.3607 | 0.8817 | 0.000999 |
| lungtumor | MacAlv-like | ANXA1 | FPR3 | 1.4026 | 0.3404 | 0.8715 | 0.000999 |
| lungtumor | MacAlv-like | ARF1 | CD4 | 0.8791 | 0.2618 | 0.5705 | 0.000999 |
| lungtumor | MacAlv-like | ARF1 | CD44 | 0.8791 | 1.4851 | 1.1821 | 0.000999 |
| lungtumor | MacAlv-like | CALR | CD44 | 0.7795 | 1.4851 | 1.1323 | 0.000999 |
| lungtumor | MacAlv-like | CALR | IL7R | 0.7795 | 0.2513 | 0.5154 | 0.000999 |
| lungtumor | MacAlv-like | CALR | LRP1 | 0.7795 | 0.5137 | 0.6466 | 0.000999 |
| lungtumor | MacAlv-like | CCL20 | C3AR1 | 0.2885 | 0.3017 | 0.2951 | 0.000999 |
| lungtumor | MacAlv-like | CCL20 | C5AR1 | 0.2885 | 0.5353 | 0.4119 | 0.000999 |
| lungtumor | MacAlv-like | CCL20 | CXCR4 | 0.2885 | 0.7383 | 0.5134 | 0.000999 |
| lungtumor | MacAlv-like | CCL20 | FPR1 | 0.2885 | 0.601 | 0.4448 | 0.000999 |

| | | | | | | | |
|---|---|---|---|---|---|---|---|
| lungtumor | MacAlv-like | CCL20 | FPR2 | 0.2885 | 0.3607 | 0.3246 | 0.000999 |
| lungtumor | MacAlv-like | CXCL10 | C3AR1 | 0.3417 | 0.3017 | 0.3217 | 0.000999 |
| lungtumor | MacAlv-like | CXCL10 | C5AR1 | 0.3417 | 0.5353 | 0.4385 | 0.000999 |
| lungtumor | MacAlv-like | CXCL10 | CXCR4 | 0.3417 | 0.7383 | 0.54 | 0.000999 |
| lungtumor | MacAlv-like | CXCL10 | FPR1 | 0.3417 | 0.601 | 0.4713 | 0.000999 |
| lungtumor | MacAlv-like | CXCL10 | FPR2 | 0.3417 | 0.3607 | 0.3512 | 0.000999 |
| lungtumor | MacAlv-like | FGA | ITGB1 | 0.4445 | 0.479 | 0.4618 | 0.000999 |
| lungtumor | MacAlv-like | FGA | ITGB2 | 0.4445 | 0.7326 | 0.5885 | 0.000999 |
| lungtumor | MacAlv-like | GSTP1 | CD44 | 2.4093 | 1.4851 | 1.9472 | 0.000999 |
| lungtumor | MacAlv-like | GSTP1 | IL7R | 2.4093 | 0.2513 | 1.3303 | 0.000999 |
| lungtumor | MacAlv-like | HMGB1 | LY96 | 1.1943 | 0.9431 | 1.0687 | 0.000999 |
| lungtumor | MacAlv-like | HMGB2 | CD44 | 0.5593 | 1.4851 | 1.0222 | 0.000999 |
| lungtumor | MacAlv-like | HMGB2 | DAB2 | 0.5593 | 0.7389 | 0.6491 | 0.000999 |
| lungtumor | MacAlv-like | HSP90B1 | ITGB1 | 0.6141 | 0.479 | 0.5466 | 0.000999 |
| lungtumor | MacAlv-like | HSP90B1 | LRP1 | 0.6141 | 0.5137 | 0.5639 | 0.000999 |
| lungtumor | MacAlv-like | MDK | LRP1 | 1.2896 | 0.5137 | 0.9016 | 0.000999 |
| lungtumor | MacAlv-like | MIF | CD74 | 1.4585 | 4.1914 | 2.825 | 0.000999 |
| lungtumor | MacAlv-like | RPS19 | C5AR1 | 3.5104 | 0.5353 | 2.0229 | 0.000999 |
| lungtumor | MacAlv-like | RPS19 | CD4 | 3.5104 | 0.2618 | 1.8861 | 0.000999 |
| lungtumor | MacAlv-like | RTN4 | CD44 | 0.4013 | 1.4851 | 0.9432 | 0.000999 |
| lungtumor | MacAlv-like | SAA1 | C3AR1 | 0.2539 | 0.3017 | 0.2778 | 0.000999 |
| lungtumor | MacAlv-like | SAA1 | C5AR1 | 0.2539 | 0.5353 | 0.3946 | 0.000999 |
| lungtumor | MacAlv-like | SAA1 | CXCR4 | 0.2539 | 0.7383 | 0.4961 | 0.000999 |
| lungtumor | MacAlv-like | SAA1 | FPR1 | 0.2539 | 0.601 | 0.4274 | 0.000999 |
| lungtumor | MacAlv-like | SAA1 | FPR2 | 0.2539 | 0.3607 | 0.3073 | 0.000999 |
| lungtumor | MacAlv-like | SAA1 | GPR65 | 0.2539 | 0.3089 | 0.2814 | 0.000999 |
| lungtumor | MacAlv-like | SAA1 | PTAFR | 0.2539 | 0.3081 | 0.281 | 0.000999 |
| lungtumor | MacAlv-like | SAA1 | SELENOS | 0.2539 | 0.4612 | 0.3576 | 0.000999 |
| lungtumor | MacAlv-like | SPP1 | CD44 | 1.4874 | 1.4851 | 1.4862 | 0.000999 |
| lungtumor | MacAlv-like | SPP1 | ITGB1 | 1.4874 | 0.479 | 0.9832 | 0.000999 |

| | | | | | | | |
|---|---|---|---|---|---|---|---|
| lungtumor | MacAlv-like | TFPI | LRP1 | 0.3615 | 0.5137 | 0.4376 | 0.000999 |
| lungtumor | MacAlv-like | TIMP1 | LRP1 | 0.6195 | 0.5137 | 0.5666 | 0.000999 |
| lungtumor | MacAlv-like | RPS19 | RPSA | 3.5104 | 1.7739 | 2.6421 | 0.00899 |
| melanomatumo | MacAlv-like | ARF1 | CD4 | 1.8545 | 0.2618 | 1.0581 | 0.000999 |
| melanomatumo | MacAlv-like | ARF1 | CD44 | 1.8545 | 1.4851 | 1.6698 | 0.000999 |
| melanomatumo | MacAlv-like | CD320 | TGFBR2 | 0.2588 | 0.2593 | 0.2591 | 0.000999 |
| melanomatumo | MacAlv-like | GSTP1 | CD44 | 2.3341 | 1.4851 | 1.9096 | 0.000999 |
| melanomatumo | MacAlv-like | GSTP1 | IL7R | 2.3341 | 0.2513 | 1.2927 | 0.000999 |
| melanomatumo | MacAlv-like | HAS2 | CD44 | 0.3078 | 1.4851 | 0.8964 | 0.000999 |
| melanomatumo | MacAlv-like | HMGB1 | AGTRAP | 1.6757 | 0.2657 | 0.9707 | 0.000999 |
| melanomatumo | MacAlv-like | HMGB1 | LY96 | 1.6757 | 0.9431 | 1.3094 | 0.000999 |
| melanomatumo | MacAlv-like | MDK | LRP1 | 0.3759 | 0.5137 | 0.4448 | 0.000999 |
| melanomatumo | MacAlv-like | MIF | CD74 | 2.6939 | 4.1914 | 3.4427 | 0.000999 |
| melanomatumo | MacAlv-like | SERPINE2 | LRP1 | 0.9323 | 0.5137 | 0.723 | 0.000999 |
| melanomatumo | MacAlv-like | HEBP1 | C5AR1 | 0.3689 | 0.5353 | 0.4521 | 0.004 |
| melanomatumo | MacAlv-like | HEBP1 | CXCR4 | 0.3689 | 0.7383 | 0.5536 | 0.004 |
| melanomatumo | MacAlv-like | HEBP1 | FPR1 | 0.3689 | 0.601 | 0.4849 | 0.004 |
| melanomatumo | MacAlv-like | HEBP1 | C3AR1 | 0.3689 | 0.3017 | 0.3353 | 0.00599 |
| melanomatumo | MacAlv-like | HEBP1 | FPR2 | 0.3689 | 0.3607 | 0.3648 | 0.00599 |
| melanomatumo | MacAlv-like | HEBP1 | FPR3 | 0.3689 | 0.3404 | 0.3546 | 0.00599 |
| ovarytumor | MacAlv-like | ANXA1 | C3AR1 | 1.0314 | 0.3017 | 0.6665 | 0.000999 |
| ovarytumor | MacAlv-like | ANXA1 | C5AR1 | 1.0314 | 0.5353 | 0.7833 | 0.000999 |
| ovarytumor | MacAlv-like | ANXA1 | CD44 | 1.0314 | 1.4851 | 1.2582 | 0.000999 |
| ovarytumor | MacAlv-like | ANXA1 | CXCR4 | 1.0314 | 0.7383 | 0.8849 | 0.000999 |
| ovarytumor | MacAlv-like | ANXA1 | FPR1 | 1.0314 | 0.601 | 0.8162 | 0.000999 |
| ovarytumor | MacAlv-like | ANXA1 | FPR2 | 1.0314 | 0.3607 | 0.696 | 0.000999 |
| ovarytumor | MacAlv-like | ANXA1 | FPR3 | 1.0314 | 0.3404 | 0.6859 | 0.000999 |
| ovarytumor | MacAlv-like | APP | ALOX5 | 0.2907 | 0.8725 | 0.5816 | 0.000999 |
| ovarytumor | MacAlv-like | APP | C3AR1 | 0.2907 | 0.3017 | 0.2962 | 0.000999 |
| ovarytumor | MacAlv-like | APP | C5AR1 | 0.2907 | 0.5353 | 0.413 | 0.000999 |

| | | | | | | | |
|---|---|---|---|---|---|---|---|
| ovarytumor | MacAlv-like | APP | CXCR4 | 0.2907 | 0.7383 | 0.5145 | 0.000999 |
| ovarytumor | MacAlv-like | APP | FPR1 | 0.2907 | 0.601 | 0.4458 | 0.000999 |
| ovarytumor | MacAlv-like | APP | FPR2 | 0.2907 | 0.3607 | 0.3257 | 0.000999 |
| ovarytumor | MacAlv-like | APP | LRP1 | 0.2907 | 0.5137 | 0.4022 | 0.000999 |
| ovarytumor | MacAlv-like | APP | TGFBR2 | 0.2907 | 0.2593 | 0.275 | 0.000999 |
| ovarytumor | MacAlv-like | ARF1 | CD4 | 0.7364 | 0.2618 | 0.4991 | 0.000999 |
| ovarytumor | MacAlv-like | ARF1 | CD44 | 0.7364 | 1.4851 | 1.1107 | 0.000999 |
| ovarytumor | MacAlv-like | CALR | CD44 | 0.7423 | 1.4851 | 1.1137 | 0.000999 |
| ovarytumor | MacAlv-like | CALR | IL7R | 0.7423 | 0.2513 | 0.4968 | 0.000999 |
| ovarytumor | MacAlv-like | CALR | LRP1 | 0.7423 | 0.5137 | 0.628 | 0.000999 |
| ovarytumor | MacAlv-like | COL1A1 | CD44 | 0.3504 | 1.4851 | 0.9177 | 0.000999 |
| ovarytumor | MacAlv-like | COL1A1 | ITGB1 | 0.3504 | 0.479 | 0.4147 | 0.000999 |
| ovarytumor | MacAlv-like | GSTP1 | CD44 | 2.6185 | 1.4851 | 2.0518 | 0.000999 |
| ovarytumor | MacAlv-like | GSTP1 | IL7R | 2.6185 | 0.2513 | 1.4349 | 0.000999 |
| ovarytumor | MacAlv-like | HMGB1 | AGTRAP | 1.7272 | 0.2657 | 0.9964 | 0.000999 |
| ovarytumor | MacAlv-like | HMGB1 | LY96 | 1.7272 | 0.9431 | 1.3351 | 0.000999 |
| ovarytumor | MacAlv-like | HMGB2 | CD44 | 0.4974 | 1.4851 | 0.9912 | 0.000999 |
| ovarytumor | MacAlv-like | HMGB2 | DAB2 | 0.4974 | 0.7389 | 0.6182 | 0.000999 |
| ovarytumor | MacAlv-like | HSP90B1 | ITGB1 | 0.6365 | 0.479 | 0.5578 | 0.000999 |
| ovarytumor | MacAlv-like | HSP90B1 | LRP1 | 0.6365 | 0.5137 | 0.5751 | 0.000999 |
| ovarytumor | MacAlv-like | ITGB1 | CD46 | 0.2783 | 0.3652 | 0.3217 | 0.000999 |
| ovarytumor | MacAlv-like | ITGB1 | DAB2 | 0.2783 | 0.7389 | 0.5086 | 0.000999 |
| ovarytumor | MacAlv-like | ITGB1 | ENG | 0.2783 | 0.2939 | 0.2861 | 0.000999 |
| ovarytumor | MacAlv-like | ITGB1 | IGF2R | 0.2783 | 0.2661 | 0.2722 | 0.000999 |
| ovarytumor | MacAlv-like | ITGB1 | PLAUR | 0.2783 | 0.6578 | 0.468 | 0.000999 |
| ovarytumor | MacAlv-like | LGALS3 | FCGR2A | 0.5097 | 0.7842 | 0.647 | 0.000999 |
| ovarytumor | MacAlv-like | LRPAP1 | LRP1 | 0.3826 | 0.5137 | 0.4481 | 0.000999 |
| ovarytumor | MacAlv-like | LRPAP1 | SORT1 | 0.3826 | 0.2831 | 0.3329 | 0.000999 |
| ovarytumor | MacAlv-like | MDK | LRP1 | 1.5321 | 0.5137 | 1.0229 | 0.000999 |
| ovarytumor | MacAlv-like | MIF | CD74 | 0.8565 | 4.1914 | 2.5239 | 0.000999 |

| | | | | | | | |
|---|---|---|---|---|---|---|---|
| ovarytumor | MacAlv-like | RPS19 | C5AR1 | 3.9325 | 0.5353 | 2.2339 | 0.000999 |
| ovarytumor | MacAlv-like | RPS19 | CD4 | 3.9325 | 0.2618 | 2.0971 | 0.000999 |
| ovarytumor | MacAlv-like | RTN4 | CD44 | 0.4992 | 1.4851 | 0.9921 | 0.000999 |
| ovarytumor | MacAlv-like | SERPING1 | LRP1 | 0.3678 | 0.5137 | 0.4407 | 0.000999 |
| ovarytumor | MacAlv-like | SPP1 | CD44 | 0.3946 | 1.4851 | 0.9399 | 0.000999 |
| ovarytumor | MacAlv-like | SPP1 | ITGB1 | 0.3946 | 0.479 | 0.4368 | 0.000999 |
| ovarytumor | MacAlv-like | TIMP1 | LRP1 | 0.7779 | 0.5137 | 0.6458 | 0.000999 |
| ovarytumor | MacAlv-like | ANXA1 | CD4 | 1.0314 | 0.2618 | 0.6466 | 0.004 |
| ovarytumor | MacAlv-like | ITGB1 | CD47 | 0.2783 | 0.6172 | 0.4477 | 0.00699 |

| Supplementary Table 8: Communication results between macrophages and lymphocytes subpopulations, and malignant cells | | | | | | | |
|---|---|---|---|---|---|---|---|
| **MacAngio vs. lymphocytes** | | | | | | | |
| celltype_01 | celltype_02 | ligand | receptor | ligand_mean_expression | receptor_mean_expression | ligand_receptor_mean_expression | pvalue |
| MacAngio | NKT | ADAM17 | ITGB1 | 0.273 | 0.488 | 0.3805 | 0.000999 |
| MacAngio | NKT | ADAM17 | TNFRSF1B | 0.273 | 0.4949 | 0.3839 | 0.000999 |
| MacAngio | NKT | ADM | PTGER2 | 0.4998 | 0.3627 | 0.4312 | 0.000999 |
| MacAngio | NKT | ALCAM | CD6 | 0.3699 | 0.4881 | 0.429 | 0.000999 |
| MacAngio | NKT | ANXA1 | CXCR4 | 1.3087 | 2.0373 | 1.673 | 0.000999 |
| MacAngio | NKT | ANXA1 | F2R | 1.3087 | 0.2978 | 0.8033 | 0.000999 |
| MacAngio | NKT | APOE | SORL1 | 1.2725 | 0.4137 | 0.8431 | 0.000999 |
| MacAngio | NKT | CALR | IL7R | 1.0589 | 0.5933 | 0.8261 | 0.000999 |
| MacAngio | NKT | CCL20 | CXCR4 | 0.7173 | 2.0373 | 1.3773 | 0.000999 |
| MacAngio | NKT | CCL20 | S1PR4 | 0.7173 | 0.4435 | 0.5804 | 0.000999 |
| MacAngio | NKT | CCL20 | S1PR5 | 0.7173 | 0.508 | 0.6127 | 0.000999 |
| MacAngio | NKT | COL1A1 | ITGB1 | 0.2951 | 0.488 | 0.3915 | 0.000999 |
| MacAngio | NKT | CXCL16 | CXCR4 | 0.6569 | 2.0373 | 1.3471 | 0.000999 |
| MacAngio | NKT | CXCL16 | S1PR4 | 0.6569 | 0.4435 | 0.5502 | 0.000999 |
| MacAngio | NKT | CXCL16 | S1PR5 | 0.6569 | 0.508 | 0.5825 | 0.000999 |
| MacAngio | NKT | CXCL2 | CXCR4 | 0.8421 | 2.0373 | 1.4397 | 0.000999 |
| MacAngio | NKT | CXCL2 | S1PR4 | 0.8421 | 0.4435 | 0.6428 | 0.000999 |
| MacAngio | NKT | CXCL2 | S1PR5 | 0.8421 | 0.508 | 0.6751 | 0.000999 |
| MacAngio | NKT | CXCL3 | CXCR4 | 0.8499 | 2.0373 | 1.4436 | 0.000999 |
| MacAngio | NKT | CXCL3 | S1PR4 | 0.8499 | 0.4435 | 0.6467 | 0.000999 |
| MacAngio | NKT | CXCL3 | S1PR5 | 0.8499 | 0.508 | 0.679 | 0.000999 |
| MacAngio | NKT | FN1 | CD44 | 1.5266 | 0.6724 | 1.0995 | 0.000999 |
| MacAngio | NKT | FN1 | ITGA4 | 1.5266 | 0.2946 | 0.9106 | 0.000999 |
| MacAngio | NKT | FN1 | ITGB1 | 1.5266 | 0.488 | 1.0073 | 0.000999 |
| MacAngio | NKT | FN1 | ITGB7 | 1.5266 | 0.4721 | 0.9993 | 0.000999 |
| MacAngio | NKT | FN1 | RPSA | 1.5266 | 2.3639 | 1.9452 | 0.000999 |
| MacAngio | NKT | GRN | TNFRSF1B | 1.4052 | 0.4949 | 0.95 | 0.000999 |

| | | | | | | | |
|---|---|---|---|---|---|---|---|
| MacAngio | NKT | GSTP1 | CD44 | 1.3386 | 0.6724 | 1.0055 | 0.000999 |
| MacAngio | NKT | GSTP1 | IL7R | 1.3386 | 0.5933 | 0.966 | 0.000999 |
| MacAngio | NKT | GSTP1 | ITGA4 | 1.3386 | 0.2946 | 0.8166 | 0.000999 |
| MacAngio | NKT | HBEGF | CD44 | 0.5321 | 0.6724 | 0.6022 | 0.000999 |
| MacAngio | NKT | HLA-DRA | PTPRC | 3.3459 | 1.8797 | 2.6128 | 0.000999 |
| MacAngio | NKT | HSP90B1 | ITGB1 | 0.8423 | 0.488 | 0.6651 | 0.000999 |
| MacAngio | NKT | HSP90B1 | TNFRSF1B | 0.8423 | 0.4949 | 0.6686 | 0.000999 |
| MacAngio | NKT | ICAM1 | IL2RG | 0.2944 | 1.1524 | 0.7234 | 0.000999 |
| MacAngio | NKT | ICAM1 | ITGAL | 0.2944 | 0.6778 | 0.4861 | 0.000999 |
| MacAngio | NKT | ICAM1 | ITGB2 | 0.2944 | 1.6481 | 0.9713 | 0.000999 |
| MacAngio | NKT | ICAM1 | SPN | 0.2944 | 0.6398 | 0.4671 | 0.000999 |
| MacAngio | NKT | ITGB1 | ITGA4 | 0.5437 | 0.2946 | 0.4191 | 0.000999 |
| MacAngio | NKT | ITGB1 | RACK1 | 0.5437 | 2.4327 | 1.4882 | 0.000999 |
| MacAngio | NKT | LGALS3 | CD6 | 1.49 | 0.4881 | 0.9891 | 0.000999 |
| MacAngio | NKT | LGALS3 | ITGB1 | 1.49 | 0.488 | 0.989 | 0.000999 |
| MacAngio | NKT | LGALS3 | SPN | 1.49 | 0.6398 | 1.0649 | 0.000999 |
| MacAngio | NKT | LRPAP1 | SORL1 | 0.4668 | 0.4137 | 0.4402 | 0.000999 |
| MacAngio | NKT | MIF | CD74 | 1.1844 | 1.6792 | 1.4318 | 0.000999 |
| MacAngio | NKT | MIF | ITGA4 | 1.1844 | 0.2946 | 0.7395 | 0.000999 |
| MacAngio | NKT | MIF | TNFRSF14 | 1.1844 | 0.2848 | 0.7346 | 0.000999 |
| MacAngio | NKT | NAMPT | ANXA6 | 1.2297 | 0.844 | 1.0368 | 0.000999 |
| MacAngio | NKT | RTN4 | CD44 | 0.9484 | 0.6724 | 0.8104 | 0.000999 |
| MacAngio | NKT | RTN4 | CD99 | 0.9484 | 1.4547 | 1.2016 | 0.000999 |
| MacAngio | NKT | SPP1 | CD44 | 2.926 | 0.6724 | 1.7992 | 0.000999 |
| MacAngio | NKT | SPP1 | ITGA4 | 2.926 | 0.2946 | 1.6103 | 0.000999 |
| MacAngio | NKT | SPP1 | ITGB1 | 2.926 | 0.488 | 1.707 | 0.000999 |
| MacAngio | NKT | THBS1 | CD47 | 0.4513 | 0.5576 | 0.5044 | 0.000999 |
| MacAngio | NKT | THBS1 | ITGA4 | 0.4513 | 0.2946 | 0.3729 | 0.000999 |
| MacAngio | NKT | THBS1 | ITGB1 | 0.4513 | 0.488 | 0.4696 | 0.000999 |
| MacAngio | NKT | TNFSF13B | ITGB1 | 0.2934 | 0.488 | 0.3907 | 0.000999 |

| | | | | | | | |
|---|---|---|---|---|---|---|---|
| MacAngio | NKT | ANXA1 | ITGA4 | 1.3087 | 0.2946 | 0.8016 | 0.002 |
| MacAngio | NKT | ITGB1 | CD47 | 0.5437 | 0.5576 | 0.5507 | 0.00599 |
| MacAngio | TCD4ex | ADAM17 | ITGB1 | 0.273 | 0.4615 | 0.3672 | 0.000999 |
| MacAngio | TCD4ex | ADAM17 | TNFRSF1B | 0.273 | 0.383 | 0.328 | 0.000999 |
| MacAngio | TCD4ex | ADM | PTGER4 | 0.4998 | 0.4696 | 0.4847 | 0.000999 |
| MacAngio | TCD4ex | ALCAM | CD6 | 0.3699 | 0.5338 | 0.4519 | 0.000999 |
| MacAngio | TCD4ex | ANXA1 | CCR7 | 1.3087 | 0.4844 | 0.8966 | 0.000999 |
| MacAngio | TCD4ex | ANXA1 | CD4 | 1.3087 | 0.5864 | 0.9476 | 0.000999 |
| MacAngio | TCD4ex | ANXA1 | CD44 | 1.3087 | 1.0244 | 1.1666 | 0.000999 |
| MacAngio | TCD4ex | ANXA1 | CXCR4 | 1.3087 | 2.4201 | 1.8644 | 0.000999 |
| MacAngio | TCD4ex | CALM1 | RPSA | 1.3243 | 2.4895 | 1.9069 | 0.000999 |
| MacAngio | TCD4ex | CALR | IL7R | 1.0589 | 1.2353 | 1.1471 | 0.000999 |
| MacAngio | TCD4ex | CCL20 | CCR7 | 0.7173 | 0.4844 | 0.6008 | 0.000999 |
| MacAngio | TCD4ex | CCL20 | CXCR4 | 0.7173 | 2.4201 | 1.5687 | 0.000999 |
| MacAngio | TCD4ex | CD86 | CD28 | 0.3542 | 0.3256 | 0.3399 | 0.000999 |
| MacAngio | TCD4ex | CD86 | CTLA4 | 0.3542 | 0.6294 | 0.4918 | 0.000999 |
| MacAngio | TCD4ex | COL1A1 | ITGB1 | 0.2951 | 0.4615 | 0.3783 | 0.000999 |
| MacAngio | TCD4ex | CXCL16 | CCR7 | 0.6569 | 0.4844 | 0.5706 | 0.000999 |
| MacAngio | TCD4ex | CXCL16 | CXCR4 | 0.6569 | 2.4201 | 1.5385 | 0.000999 |
| MacAngio | TCD4ex | CXCL2 | CCR7 | 0.8421 | 0.4844 | 0.6632 | 0.000999 |
| MacAngio | TCD4ex | CXCL2 | CXCR4 | 0.8421 | 2.4201 | 1.6311 | 0.000999 |
| MacAngio | TCD4ex | CXCL3 | CCR7 | 0.8499 | 0.4844 | 0.6672 | 0.000999 |
| MacAngio | TCD4ex | CXCL3 | CXCR4 | 0.8499 | 2.4201 | 1.635 | 0.000999 |
| MacAngio | TCD4ex | FN1 | CD44 | 1.5266 | 1.0244 | 1.2755 | 0.000999 |
| MacAngio | TCD4ex | FN1 | ITGB1 | 1.5266 | 0.4615 | 0.994 | 0.000999 |
| MacAngio | TCD4ex | FN1 | RPSA | 1.5266 | 2.4895 | 2.008 | 0.000999 |
| MacAngio | TCD4ex | GRN | TNFRSF1B | 1.4052 | 0.383 | 0.8941 | 0.000999 |
| MacAngio | TCD4ex | GSTP1 | CD44 | 1.3386 | 1.0244 | 1.1815 | 0.000999 |
| MacAngio | TCD4ex | GSTP1 | IL7R | 1.3386 | 1.2353 | 1.2869 | 0.000999 |
| MacAngio | TCD4ex | HBEGF | CD44 | 0.5321 | 1.0244 | 0.7782 | 0.000999 |

| | | | | | | | |
|---|---|---|---|---|---|---|---|
| MacAngio | TCD4ex | HLA-DMA | CD4 | 0.8597 | 0.5864 | 0.7231 | 0.000999 |
| MacAngio | TCD4ex | HLA-DRA | CD4 | 3.3459 | 0.5864 | 1.9662 | 0.000999 |
| MacAngio | TCD4ex | HLA-DRA | PDCD1 | 3.3459 | 0.457 | 1.9015 | 0.000999 |
| MacAngio | TCD4ex | HLA-DRA | PTPRC | 3.3459 | 1.7493 | 2.5476 | 0.000999 |
| MacAngio | TCD4ex | HSP90B1 | ITGB1 | 0.8423 | 0.4615 | 0.6519 | 0.000999 |
| MacAngio | TCD4ex | HSP90B1 | TNFRSF1B | 0.8423 | 0.383 | 0.6127 | 0.000999 |
| MacAngio | TCD4ex | ICAM1 | IL2RG | 0.2944 | 0.9695 | 0.6319 | 0.000999 |
| MacAngio | TCD4ex | ICAM1 | ITGAL | 0.2944 | 0.2603 | 0.2774 | 0.000999 |
| MacAngio | TCD4ex | ICAM1 | ITGB2 | 0.2944 | 0.7871 | 0.5407 | 0.000999 |
| MacAngio | TCD4ex | ITGB1 | RACK1 | 0.5437 | 2.4434 | 1.4936 | 0.000999 |
| MacAngio | TCD4ex | ITGB2 | ITGAL | 0.9554 | 0.2603 | 0.6079 | 0.000999 |
| MacAngio | TCD4ex | LGALS3 | CD6 | 1.49 | 0.5338 | 1.0119 | 0.000999 |
| MacAngio | TCD4ex | LGALS3 | ITGB1 | 1.49 | 0.4615 | 0.9757 | 0.000999 |
| MacAngio | TCD4ex | MIF | TNFRSF14 | 1.1844 | 0.2806 | 0.7325 | 0.000999 |
| MacAngio | TCD4ex | NAMPT | ANXA6 | 1.2297 | 0.5386 | 0.8841 | 0.000999 |
| MacAngio | TCD4ex | RTN4 | CD44 | 0.9484 | 1.0244 | 0.9864 | 0.000999 |
| MacAngio | TCD4ex | RTN4 | CD99 | 0.9484 | 0.9165 | 0.9325 | 0.000999 |
| MacAngio | TCD4ex | SPP1 | CD44 | 2.926 | 1.0244 | 1.9752 | 0.000999 |
| MacAngio | TCD4ex | SPP1 | ITGB1 | 2.926 | 0.4615 | 1.6938 | 0.000999 |
| MacAngio | TCD4ex | TGFB1 | ITGB1 | 0.2752 | 0.4615 | 0.3683 | 0.000999 |
| MacAngio | TCD4ex | THBS1 | CD47 | 0.4513 | 0.3456 | 0.3985 | 0.000999 |
| MacAngio | TCD4ex | THBS1 | ITGB1 | 0.4513 | 0.4615 | 0.4564 | 0.000999 |
| MacAngio | TCD4ex | TNFSF13B | ITGB1 | 0.2934 | 0.4615 | 0.3775 | 0.000999 |
| MacAngio | TCD4reg | ADAM17 | ITGB1 | 0.273 | 0.4001 | 0.3365 | 0.000999 |
| MacAngio | TCD4reg | ADAM17 | TNFRSF1B | 0.273 | 0.7982 | 0.5356 | 0.000999 |
| MacAngio | TCD4reg | ALCAM | CD6 | 0.3699 | 0.2651 | 0.3175 | 0.000999 |
| MacAngio | TCD4reg | ANXA1 | CCR7 | 1.3087 | 0.3098 | 0.8093 | 0.000999 |
| MacAngio | TCD4reg | ANXA1 | CD4 | 1.3087 | 0.429 | 0.8688 | 0.000999 |
| MacAngio | TCD4reg | ANXA1 | CD44 | 1.3087 | 1.0896 | 1.1992 | 0.000999 |
| MacAngio | TCD4reg | ANXA1 | CXCR3 | 1.3087 | 0.3357 | 0.8222 | 0.000999 |

| | | | | | | | |
|---|---|---|---|---|---|---|---|
| MacAngio | TCD4reg | ANXA1 | CXCR4 | 1.3087 | 2.2962 | 1.8024 | 0.000999 |
| MacAngio | TCD4reg | ANXA1 | CXCR6 | 1.3087 | 0.3945 | 0.8516 | 0.000999 |
| MacAngio | TCD4reg | ANXA1 | FAS | 1.3087 | 0.2727 | 0.7907 | 0.000999 |
| MacAngio | TCD4reg | CALR | CD44 | 1.0589 | 1.0896 | 1.0743 | 0.000999 |
| MacAngio | TCD4reg | CALR | FAS | 1.0589 | 0.2727 | 0.6658 | 0.000999 |
| MacAngio | TCD4reg | CALR | IL7R | 1.0589 | 0.9682 | 1.0135 | 0.000999 |
| MacAngio | TCD4reg | CCL20 | CCR7 | 0.7173 | 0.3098 | 0.5136 | 0.000999 |
| MacAngio | TCD4reg | CCL20 | CXCR3 | 0.7173 | 0.3357 | 0.5265 | 0.000999 |
| MacAngio | TCD4reg | CCL20 | CXCR4 | 0.7173 | 2.2962 | 1.5067 | 0.000999 |
| MacAngio | TCD4reg | CCL20 | CXCR6 | 0.7173 | 0.3945 | 0.5559 | 0.000999 |
| MacAngio | TCD4reg | CCL20 | S1PR4 | 0.7173 | 0.2744 | 0.4959 | 0.000999 |
| MacAngio | TCD4reg | CD86 | CD28 | 0.3542 | 0.313 | 0.3336 | 0.000999 |
| MacAngio | TCD4reg | CD86 | CTLA4 | 0.3542 | 0.7909 | 0.5726 | 0.000999 |
| MacAngio | TCD4reg | COL1A1 | ITGB1 | 0.2951 | 0.4001 | 0.3476 | 0.000999 |
| MacAngio | TCD4reg | CXCL16 | CCR7 | 0.6569 | 0.3098 | 0.4834 | 0.000999 |
| MacAngio | TCD4reg | CXCL16 | CXCR3 | 0.6569 | 0.3357 | 0.4963 | 0.000999 |
| MacAngio | TCD4reg | CXCL16 | CXCR4 | 0.6569 | 2.2962 | 1.4765 | 0.000999 |
| MacAngio | TCD4reg | CXCL16 | CXCR6 | 0.6569 | 0.3945 | 0.5257 | 0.000999 |
| MacAngio | TCD4reg | CXCL16 | S1PR4 | 0.6569 | 0.2744 | 0.4656 | 0.000999 |
| MacAngio | TCD4reg | CXCL2 | CCR7 | 0.8421 | 0.3098 | 0.576 | 0.000999 |
| MacAngio | TCD4reg | CXCL2 | CXCR3 | 0.8421 | 0.3357 | 0.5889 | 0.000999 |
| MacAngio | TCD4reg | CXCL2 | CXCR4 | 0.8421 | 2.2962 | 1.5691 | 0.000999 |
| MacAngio | TCD4reg | CXCL2 | CXCR6 | 0.8421 | 0.3945 | 0.6183 | 0.000999 |
| MacAngio | TCD4reg | CXCL2 | S1PR4 | 0.8421 | 0.2744 | 0.5582 | 0.000999 |
| MacAngio | TCD4reg | CXCL3 | CCR7 | 0.8499 | 0.3098 | 0.5799 | 0.000999 |
| MacAngio | TCD4reg | CXCL3 | CXCR3 | 0.8499 | 0.3357 | 0.5928 | 0.000999 |
| MacAngio | TCD4reg | CXCL3 | CXCR4 | 0.8499 | 2.2962 | 1.573 | 0.000999 |
| MacAngio | TCD4reg | CXCL3 | CXCR6 | 0.8499 | 0.3945 | 0.6222 | 0.000999 |
| MacAngio | TCD4reg | CXCL3 | S1PR4 | 0.8499 | 0.2744 | 0.5622 | 0.000999 |
| MacAngio | TCD4reg | FN1 | CD44 | 1.5266 | 1.0896 | 1.3081 | 0.000999 |

| | | | | | | | |
|---|---|---|---|---|---|---|---|
| MacAngio | TCD4reg | FN1 | ITGB1 | 1.5266 | 0.4001 | 0.9633 | 0.000999 |
| MacAngio | TCD4reg | FN1 | RPSA | 1.5266 | 2.4999 | 2.0132 | 0.000999 |
| MacAngio | TCD4reg | GRN | TNFRSF1B | 1.4052 | 0.7982 | 1.1017 | 0.000999 |
| MacAngio | TCD4reg | GSTP1 | CD44 | 1.3386 | 1.0896 | 1.2141 | 0.000999 |
| MacAngio | TCD4reg | GSTP1 | FAS | 1.3386 | 0.2727 | 0.8057 | 0.000999 |
| MacAngio | TCD4reg | GSTP1 | IL7R | 1.3386 | 0.9682 | 1.1534 | 0.000999 |
| MacAngio | TCD4reg | HBEGF | CD44 | 0.5321 | 1.0896 | 0.8109 | 0.000999 |
| MacAngio | TCD4reg | HLA-DMA | CD4 | 0.8597 | 0.429 | 0.6443 | 0.000999 |
| MacAngio | TCD4reg | HLA-DRA | CD4 | 3.3459 | 0.429 | 1.8874 | 0.000999 |
| MacAngio | TCD4reg | HLA-DRA | PTPRC | 3.3459 | 1.6603 | 2.5031 | 0.000999 |
| MacAngio | TCD4reg | HSP90B1 | ITGB1 | 0.8423 | 0.4001 | 0.6212 | 0.000999 |
| MacAngio | TCD4reg | HSP90B1 | TNFRSF1B | 0.8423 | 0.7982 | 0.8202 | 0.000999 |
| MacAngio | TCD4reg | ICAM1 | IL2RA | 0.2944 | 0.7115 | 0.5029 | 0.000999 |
| MacAngio | TCD4reg | ICAM1 | IL2RG | 0.2944 | 1.1601 | 0.7273 | 0.000999 |
| MacAngio | TCD4reg | ICAM1 | ITGB2 | 0.2944 | 0.6238 | 0.4591 | 0.000999 |
| MacAngio | TCD4reg | ITGB1 | CD47 | 0.5437 | 0.416 | 0.4799 | 0.000999 |
| MacAngio | TCD4reg | ITGB1 | RACK1 | 0.5437 | 2.5298 | 1.5368 | 0.000999 |
| MacAngio | TCD4reg | LGALS3 | CD6 | 1.49 | 0.2651 | 0.8775 | 0.000999 |
| MacAngio | TCD4reg | LGALS3 | FAS | 1.49 | 0.2727 | 0.8814 | 0.000999 |
| MacAngio | TCD4reg | LGALS3 | ITGB1 | 1.49 | 0.4001 | 0.945 | 0.000999 |
| MacAngio | TCD4reg | MIF | CD74 | 1.1844 | 2.0716 | 1.628 | 0.000999 |
| MacAngio | TCD4reg | MIF | TNFRSF14 | 1.1844 | 0.3843 | 0.7844 | 0.000999 |
| MacAngio | TCD4reg | NAMPT | ANXA6 | 1.2297 | 0.4557 | 0.8427 | 0.000999 |
| MacAngio | TCD4reg | RTN4 | CD44 | 0.9484 | 1.0896 | 1.019 | 0.000999 |
| MacAngio | TCD4reg | RTN4 | CD99 | 0.9484 | 0.6603 | 0.8044 | 0.000999 |
| MacAngio | TCD4reg | SPP1 | CD44 | 2.926 | 1.0896 | 2.0078 | 0.000999 |
| MacAngio | TCD4reg | SPP1 | ITGB1 | 2.926 | 0.4001 | 1.6631 | 0.000999 |
| MacAngio | TCD4reg | TGFB1 | ITGB1 | 0.2752 | 0.4001 | 0.3376 | 0.000999 |
| MacAngio | TCD4reg | THBS1 | CD47 | 0.4513 | 0.416 | 0.4336 | 0.000999 |
| MacAngio | TCD4reg | THBS1 | ITGB1 | 0.4513 | 0.4001 | 0.4257 | 0.000999 |

| | | | | | | | |
|---|---|---|---|---|---|---|---|
| MacAngio | TCD4reg | TNFSF13B | ITGB1 | 0.2934 | 0.4001 | 0.3468 | 0.000999 |
| MacAngio | TCD8em | ADAM17 | ITGB1 | 0.273 | 0.2557 | 0.2644 | 0.000999 |
| MacAngio | TCD8em | ADAM17 | TNFRSF1B | 0.273 | 0.3038 | 0.2884 | 0.000999 |
| MacAngio | TCD8em | ADM | PTGER4 | 0.4998 | 0.2847 | 0.3923 | 0.000999 |
| MacAngio | TCD8em | ALCAM | CD6 | 0.3699 | 0.4204 | 0.3952 | 0.000999 |
| MacAngio | TCD8em | ANXA1 | CD44 | 1.3087 | 1.119 | 1.2139 | 0.000999 |
| MacAngio | TCD8em | ANXA1 | CXCR3 | 1.3087 | 0.4189 | 0.8638 | 0.000999 |
| MacAngio | TCD8em | ANXA1 | CXCR4 | 1.3087 | 2.8389 | 2.0738 | 0.000999 |
| MacAngio | TCD8em | ANXA1 | CXCR6 | 1.3087 | 0.3399 | 0.8243 | 0.000999 |
| MacAngio | TCD8em | ANXA1 | F2R | 1.3087 | 0.2695 | 0.7891 | 0.000999 |
| MacAngio | TCD8em | ANXA1 | ITGA4 | 1.3087 | 0.2823 | 0.7955 | 0.000999 |
| MacAngio | TCD8em | CALR | CD44 | 1.0589 | 1.119 | 1.089 | 0.000999 |
| MacAngio | TCD8em | CALR | IL7R | 1.0589 | 0.7183 | 0.8886 | 0.000999 |
| MacAngio | TCD8em | CCL20 | CXCR3 | 0.7173 | 0.4189 | 0.5681 | 0.000999 |
| MacAngio | TCD8em | CCL20 | CXCR4 | 0.7173 | 2.8389 | 1.7781 | 0.000999 |
| MacAngio | TCD8em | CCL20 | CXCR6 | 0.7173 | 0.3399 | 0.5286 | 0.000999 |
| MacAngio | TCD8em | CCL20 | S1PR4 | 0.7173 | 0.2677 | 0.4925 | 0.000999 |
| MacAngio | TCD8em | CXCL16 | CXCR3 | 0.6569 | 0.4189 | 0.5379 | 0.000999 |
| MacAngio | TCD8em | CXCL16 | CXCR4 | 0.6569 | 2.8389 | 1.7479 | 0.000999 |
| MacAngio | TCD8em | CXCL16 | CXCR6 | 0.6569 | 0.3399 | 0.4984 | 0.000999 |
| MacAngio | TCD8em | CXCL16 | S1PR4 | 0.6569 | 0.2677 | 0.4623 | 0.000999 |
| MacAngio | TCD8em | CXCL2 | CXCR3 | 0.8421 | 0.4189 | 0.6305 | 0.000999 |
| MacAngio | TCD8em | CXCL2 | CXCR4 | 0.8421 | 2.8389 | 1.8405 | 0.000999 |
| MacAngio | TCD8em | CXCL2 | CXCR6 | 0.8421 | 0.3399 | 0.591 | 0.000999 |
| MacAngio | TCD8em | CXCL2 | S1PR4 | 0.8421 | 0.2677 | 0.5549 | 0.000999 |
| MacAngio | TCD8em | CXCL3 | CXCR3 | 0.8499 | 0.4189 | 0.6344 | 0.000999 |
| MacAngio | TCD8em | CXCL3 | CXCR4 | 0.8499 | 2.8389 | 1.8444 | 0.000999 |
| MacAngio | TCD8em | CXCL3 | CXCR6 | 0.8499 | 0.3399 | 0.5949 | 0.000999 |
| MacAngio | TCD8em | CXCL3 | S1PR4 | 0.8499 | 0.2677 | 0.5588 | 0.000999 |
| MacAngio | TCD8em | FN1 | CD44 | 1.5266 | 1.119 | 1.3228 | 0.000999 |

| | | | | | | | |
|---|---|---|---|---|---|---|---|
| MacAngio | TCD8em | FN1 | ITGA4 | 1.5266 | 0.2823 | 0.9044 | 0.000999 |
| MacAngio | TCD8em | FN1 | ITGB1 | 1.5266 | 0.2557 | 0.8911 | 0.000999 |
| MacAngio | TCD8em | FN1 | RPSA | 1.5266 | 2.3686 | 1.9476 | 0.000999 |
| MacAngio | TCD8em | GRN | TNFRSF1B | 1.4052 | 0.3038 | 0.8545 | 0.000999 |
| MacAngio | TCD8em | GSTP1 | CD44 | 1.3386 | 1.119 | 1.2288 | 0.000999 |
| MacAngio | TCD8em | GSTP1 | IL7R | 1.3386 | 0.7183 | 1.0285 | 0.000999 |
| MacAngio | TCD8em | GSTP1 | ITGA4 | 1.3386 | 0.2823 | 0.8105 | 0.000999 |
| MacAngio | TCD8em | HBEGF | CD44 | 0.5321 | 1.119 | 0.8256 | 0.000999 |
| MacAngio | TCD8em | HLA-DRA | PTPRC | 3.3459 | 1.7912 | 2.5686 | 0.000999 |
| MacAngio | TCD8em | HSP90B1 | ITGB1 | 0.8423 | 0.2557 | 0.549 | 0.000999 |
| MacAngio | TCD8em | HSP90B1 | TNFRSF1B | 0.8423 | 0.3038 | 0.573 | 0.000999 |
| MacAngio | TCD8em | ICAM1 | IL2RG | 0.2944 | 0.9501 | 0.6223 | 0.000999 |
| MacAngio | TCD8em | ICAM1 | ITGAL | 0.2944 | 0.3162 | 0.3053 | 0.000999 |
| MacAngio | TCD8em | ICAM1 | ITGB2 | 0.2944 | 1.0018 | 0.6481 | 0.000999 |
| MacAngio | TCD8em | ICAM1 | SPN | 0.2944 | 0.2603 | 0.2774 | 0.000999 |
| MacAngio | TCD8em | ITGB1 | CD47 | 0.5437 | 0.4276 | 0.4856 | 0.000999 |
| MacAngio | TCD8em | ITGB1 | ITGA4 | 0.5437 | 0.2823 | 0.413 | 0.000999 |
| MacAngio | TCD8em | ITGB1 | RACK1 | 0.5437 | 2.4374 | 1.4905 | 0.000999 |
| MacAngio | TCD8em | LGALS3 | CD6 | 1.49 | 0.4204 | 0.9552 | 0.000999 |
| MacAngio | TCD8em | LGALS3 | ITGB1 | 1.49 | 0.2557 | 0.8729 | 0.000999 |
| MacAngio | TCD8em | LGALS3 | SPN | 1.49 | 0.2603 | 0.8752 | 0.000999 |
| MacAngio | TCD8em | MIF | CD74 | 1.1844 | 2.4538 | 1.8191 | 0.000999 |
| MacAngio | TCD8em | MIF | ITGA4 | 1.1844 | 0.2823 | 0.7334 | 0.000999 |
| MacAngio | TCD8em | MIF | TNFRSF14 | 1.1844 | 0.2915 | 0.7379 | 0.000999 |
| MacAngio | TCD8em | NAMPT | ANXA6 | 1.2297 | 0.552 | 0.8908 | 0.000999 |
| MacAngio | TCD8em | RTN4 | CD44 | 0.9484 | 1.119 | 1.0337 | 0.000999 |
| MacAngio | TCD8em | RTN4 | CD99 | 0.9484 | 1.1525 | 1.0505 | 0.000999 |
| MacAngio | TCD8em | SPP1 | CD44 | 2.926 | 1.119 | 2.0225 | 0.000999 |
| MacAngio | TCD8em | SPP1 | ITGA4 | 2.926 | 0.2823 | 1.6042 | 0.000999 |
| MacAngio | TCD8em | SPP1 | ITGB1 | 2.926 | 0.2557 | 1.5909 | 0.000999 |

| | | | | | | | |
|---|---|---|---|---|---|---|---|
| MacAngio | TCD8em | THBS1 | CD47 | 0.4513 | 0.4276 | 0.4394 | 0.000999 |
| MacAngio | TCD8em | THBS1 | ITGA4 | 0.4513 | 0.2823 | 0.3668 | 0.000999 |
| MacAngio | TCD8em | THBS1 | ITGB1 | 0.4513 | 0.2557 | 0.3535 | 0.000999 |
| MacAngio | TCD8em | TNFSF13B | ITGB1 | 0.2934 | 0.2557 | 0.2746 | 0.000999 |
| MacAngio | TCD8ex | ADAM17 | ITGB1 | 0.273 | 0.2995 | 0.2862 | 0.000999 |
| MacAngio | TCD8ex | ADAM17 | TNFRSF1B | 0.273 | 0.6436 | 0.4583 | 0.000999 |
| MacAngio | TCD8ex | ADM | PTGER4 | 0.4998 | 0.3089 | 0.4043 | 0.000999 |
| MacAngio | TCD8ex | ALCAM | CD6 | 0.3699 | 0.318 | 0.344 | 0.000999 |
| MacAngio | TCD8ex | ANXA1 | CD44 | 1.3087 | 0.8219 | 1.0653 | 0.000999 |
| MacAngio | TCD8ex | ANXA1 | CXCR3 | 1.3087 | 0.3647 | 0.8367 | 0.000999 |
| MacAngio | TCD8ex | ANXA1 | CXCR4 | 1.3087 | 2.3472 | 1.828 | 0.000999 |
| MacAngio | TCD8ex | ANXA1 | CXCR6 | 1.3087 | 0.6626 | 0.9856 | 0.000999 |
| MacAngio | TCD8ex | ANXA1 | ITGA4 | 1.3087 | 0.4432 | 0.876 | 0.000999 |
| MacAngio | TCD8ex | ANXA1 | VCAM1 | 1.3087 | 0.4462 | 0.8775 | 0.000999 |
| MacAngio | TCD8ex | APOE | VCAM1 | 1.2725 | 0.4462 | 0.8593 | 0.000999 |
| MacAngio | TCD8ex | CCL20 | CXCR3 | 0.7173 | 0.3647 | 0.541 | 0.000999 |
| MacAngio | TCD8ex | CCL20 | CXCR4 | 0.7173 | 2.3472 | 1.5323 | 0.000999 |
| MacAngio | TCD8ex | CCL20 | CXCR6 | 0.7173 | 0.6626 | 0.6899 | 0.000999 |
| MacAngio | TCD8ex | CD86 | CTLA4 | 0.3542 | 0.4883 | 0.4212 | 0.000999 |
| MacAngio | TCD8ex | COL1A1 | CD44 | 0.2951 | 0.8219 | 0.5585 | 0.000999 |
| MacAngio | TCD8ex | COL1A1 | ITGB1 | 0.2951 | 0.2995 | 0.2973 | 0.000999 |
| MacAngio | TCD8ex | CXCL16 | CXCR3 | 0.6569 | 0.3647 | 0.5108 | 0.000999 |
| MacAngio | TCD8ex | CXCL16 | CXCR4 | 0.6569 | 2.3472 | 1.5021 | 0.000999 |
| MacAngio | TCD8ex | CXCL16 | CXCR6 | 0.6569 | 0.6626 | 0.6597 | 0.000999 |
| MacAngio | TCD8ex | CXCL2 | CXCR3 | 0.8421 | 0.3647 | 0.6034 | 0.000999 |
| MacAngio | TCD8ex | CXCL2 | CXCR4 | 0.8421 | 2.3472 | 1.5947 | 0.000999 |
| MacAngio | TCD8ex | CXCL2 | CXCR6 | 0.8421 | 0.6626 | 0.7523 | 0.000999 |
| MacAngio | TCD8ex | CXCL3 | CXCR3 | 0.8499 | 0.3647 | 0.6073 | 0.000999 |
| MacAngio | TCD8ex | CXCL3 | CXCR4 | 0.8499 | 2.3472 | 1.5986 | 0.000999 |
| MacAngio | TCD8ex | CXCL3 | CXCR6 | 0.8499 | 0.6626 | 0.7562 | 0.000999 |

| | | | | | | | |
|---|---|---|---|---|---|---|---|
| MacAngio | TCD8ex | FN1 | CD44 | 1.5266 | 0.8219 | 1.1743 | 0.000999 |
| MacAngio | TCD8ex | FN1 | ITGA4 | 1.5266 | 0.4432 | 0.9849 | 0.000999 |
| MacAngio | TCD8ex | FN1 | ITGB1 | 1.5266 | 0.2995 | 0.913 | 0.000999 |
| MacAngio | TCD8ex | FN1 | ITGB7 | 1.5266 | 0.3007 | 0.9136 | 0.000999 |
| MacAngio | TCD8ex | FN1 | RPSA | 1.5266 | 2.2697 | 1.8981 | 0.000999 |
| MacAngio | TCD8ex | GRN | TNFRSF1B | 1.4052 | 0.6436 | 1.0244 | 0.000999 |
| MacAngio | TCD8ex | GSTP1 | CD44 | 1.3386 | 0.8219 | 1.0803 | 0.000999 |
| MacAngio | TCD8ex | GSTP1 | ITGA4 | 1.3386 | 0.4432 | 0.8909 | 0.000999 |
| MacAngio | TCD8ex | GSTP1 | VCAM1 | 1.3386 | 0.4462 | 0.8924 | 0.000999 |
| MacAngio | TCD8ex | HBEGF | CD44 | 0.5321 | 0.8219 | 0.677 | 0.000999 |
| MacAngio | TCD8ex | HLA-DRA | PDCD1 | 3.3459 | 0.6109 | 1.9784 | 0.000999 |
| MacAngio | TCD8ex | HLA-DRA | PTPRC | 3.3459 | 1.6762 | 2.511 | 0.000999 |
| MacAngio | TCD8ex | HSP90B1 | ITGB1 | 0.8423 | 0.2995 | 0.5709 | 0.000999 |
| MacAngio | TCD8ex | HSP90B1 | TNFRSF1B | 0.8423 | 0.6436 | 0.7429 | 0.000999 |
| MacAngio | TCD8ex | ICAM1 | IL2RG | 0.2944 | 1.2126 | 0.7535 | 0.000999 |
| MacAngio | TCD8ex | ICAM1 | ITGB2 | 0.2944 | 0.7635 | 0.5289 | 0.000999 |
| MacAngio | TCD8ex | ICAM1 | SPN | 0.2944 | 0.3486 | 0.3215 | 0.000999 |
| MacAngio | TCD8ex | ITGB1 | CD47 | 0.5437 | 0.3952 | 0.4694 | 0.000999 |
| MacAngio | TCD8ex | ITGB1 | ITGA4 | 0.5437 | 0.4432 | 0.4935 | 0.000999 |
| MacAngio | TCD8ex | ITGB1 | RACK1 | 0.5437 | 2.3353 | 1.4395 | 0.000999 |
| MacAngio | TCD8ex | ITGB1 | VCAM1 | 0.5437 | 0.4462 | 0.495 | 0.000999 |
| MacAngio | TCD8ex | LGALS3 | CD6 | 1.49 | 0.318 | 0.904 | 0.000999 |
| MacAngio | TCD8ex | LGALS3 | ITGB1 | 1.49 | 0.2995 | 0.8947 | 0.000999 |
| MacAngio | TCD8ex | LGALS3 | SPN | 1.49 | 0.3486 | 0.9193 | 0.000999 |
| MacAngio | TCD8ex | MIF | CD74 | 1.1844 | 2.5058 | 1.8451 | 0.000999 |
| MacAngio | TCD8ex | MIF | ITGA4 | 1.1844 | 0.4432 | 0.8138 | 0.000999 |
| MacAngio | TCD8ex | MIF | TNFRSF14 | 1.1844 | 0.3094 | 0.7469 | 0.000999 |
| MacAngio | TCD8ex | NAMPT | ANXA6 | 1.2297 | 0.5678 | 0.8987 | 0.000999 |
| MacAngio | TCD8ex | RTN4 | CD44 | 0.9484 | 0.8219 | 0.8852 | 0.000999 |
| MacAngio | TCD8ex | RTN4 | CD99 | 0.9484 | 1.2049 | 1.0766 | 0.000999 |

| | | | | | | | |
|---|---|---|---|---|---|---|---|
| MacAngio | TCD8ex | SPP1 | CD44 | 2.926 | 0.8219 | 1.874 | 0.000999 |
| MacAngio | TCD8ex | SPP1 | ITGA4 | 2.926 | 0.4432 | 1.6846 | 0.000999 |
| MacAngio | TCD8ex | SPP1 | ITGB1 | 2.926 | 0.2995 | 1.6128 | 0.000999 |
| MacAngio | TCD8ex | TGFB1 | ITGB1 | 0.2752 | 0.2995 | 0.2873 | 0.000999 |
| MacAngio | TCD8ex | THBS1 | CD47 | 0.4513 | 0.3952 | 0.4232 | 0.000999 |
| MacAngio | TCD8ex | THBS1 | ITGA4 | 0.4513 | 0.4432 | 0.4472 | 0.000999 |
| MacAngio | TCD8ex | THBS1 | ITGB1 | 0.4513 | 0.2995 | 0.3754 | 0.000999 |
| MacAngio | TCD8ex | TNFSF13B | ITGB1 | 0.2934 | 0.2995 | 0.2965 | 0.000999 |
| MacAngio | TGD | ADM | PTGDR | 0.4998 | 0.7153 | 0.6075 | 0.000999 |
| MacAngio | TGD | ALCAM | CD6 | 0.3699 | 0.4684 | 0.4192 | 0.000999 |
| MacAngio | TGD | ANXA1 | CXCR3 | 1.3087 | 1.1748 | 1.2417 | 0.000999 |
| MacAngio | TGD | ANXA1 | CXCR4 | 1.3087 | 1.4894 | 1.3991 | 0.000999 |
| MacAngio | TGD | ANXA1 | ITGA4 | 1.3087 | 0.3968 | 0.8528 | 0.000999 |
| MacAngio | TGD | APOE | SORL1 | 1.2725 | 0.3318 | 0.8021 | 0.000999 |
| MacAngio | TGD | ARF1 | ITGA4 | 0.648 | 0.3968 | 0.5224 | 0.000999 |
| MacAngio | TGD | CALM1 | RPSA | 1.3243 | 2.8814 | 2.1028 | 0.000999 |
| MacAngio | TGD | CALR | IL7R | 1.0589 | 1.3862 | 1.2226 | 0.000999 |
| MacAngio | TGD | CCL20 | CXCR3 | 0.7173 | 1.1748 | 0.946 | 0.000999 |
| MacAngio | TGD | CCL20 | CXCR4 | 0.7173 | 1.4894 | 1.1033 | 0.000999 |
| MacAngio | TGD | CD99 | CD99 | 0.5878 | 1.7154 | 1.1516 | 0.000999 |
| MacAngio | TGD | CXCL16 | CXCR3 | 0.6569 | 1.1748 | 0.9158 | 0.000999 |
| MacAngio | TGD | CXCL16 | CXCR4 | 0.6569 | 1.4894 | 1.0731 | 0.000999 |
| MacAngio | TGD | CXCL2 | CXCR3 | 0.8421 | 1.1748 | 1.0084 | 0.000999 |
| MacAngio | TGD | CXCL2 | CXCR4 | 0.8421 | 1.4894 | 1.1657 | 0.000999 |
| MacAngio | TGD | CXCL3 | CXCR3 | 0.8499 | 1.1748 | 1.0123 | 0.000999 |
| MacAngio | TGD | CXCL3 | CXCR4 | 0.8499 | 1.4894 | 1.1696 | 0.000999 |
| MacAngio | TGD | FN1 | ITGA4 | 1.5266 | 0.3968 | 0.9617 | 0.000999 |
| MacAngio | TGD | FN1 | ITGB7 | 1.5266 | 0.459 | 0.9928 | 0.000999 |
| MacAngio | TGD | FN1 | RPSA | 1.5266 | 2.8814 | 2.204 | 0.000999 |
| MacAngio | TGD | GRN | TNFRSF1A | 1.4052 | 0.3099 | 0.8575 | 0.000999 |

| | | | | | | | |
|---|---|---|---|---|---|---|---|
| MacAngio | TGD | GSTP1 | IL7R | 1.3386 | 1.3862 | 1.3624 | 0.000999 |
| MacAngio | TGD | GSTP1 | ITGA4 | 1.3386 | 0.3968 | 0.8677 | 0.000999 |
| MacAngio | TGD | HLA-A | CD8A | 2.3272 | 1.746 | 2.0366 | 0.000999 |
| MacAngio | TGD | HLA-DRA | PTPRC | 3.3459 | 1.8653 | 2.6056 | 0.000999 |
| MacAngio | TGD | HLA-E | CD8A | 1.1716 | 1.746 | 1.4588 | 0.000999 |
| MacAngio | TGD | HLA-E | KLRC2 | 1.1716 | 0.8082 | 0.9899 | 0.000999 |
| MacAngio | TGD | HLA-E | KLRD1 | 1.1716 | 1.3451 | 1.2583 | 0.000999 |
| MacAngio | TGD | HMGB1 | ITGA4 | 1.0175 | 0.3968 | 0.7072 | 0.000999 |
| MacAngio | TGD | ICAM1 | IL2RG | 0.2944 | 1.561 | 0.9277 | 0.000999 |
| MacAngio | TGD | ICAM1 | ITGAL | 0.2944 | 0.3731 | 0.3338 | 0.000999 |
| MacAngio | TGD | ICAM1 | ITGB2 | 0.2944 | 1.4444 | 0.8694 | 0.000999 |
| MacAngio | TGD | ICAM1 | SPN | 0.2944 | 0.5644 | 0.4294 | 0.000999 |
| MacAngio | TGD | ITGB1 | CD46 | 0.5437 | 0.3085 | 0.4261 | 0.000999 |
| MacAngio | TGD | ITGB1 | ITGA4 | 0.5437 | 0.3968 | 0.4703 | 0.000999 |
| MacAngio | TGD | ITGB1 | PTK2B | 0.5437 | 0.3668 | 0.4553 | 0.000999 |
| MacAngio | TGD | ITGB1 | RACK1 | 0.5437 | 2.7287 | 1.6362 | 0.000999 |
| MacAngio | TGD | ITGB2 | ITGAL | 0.9554 | 0.3731 | 0.6643 | 0.000999 |
| MacAngio | TGD | LGALS3 | CD6 | 1.49 | 0.4684 | 0.9792 | 0.000999 |
| MacAngio | TGD | LGALS3 | SPN | 1.49 | 0.5644 | 1.0272 | 0.000999 |
| MacAngio | TGD | LGALS3 | TGFBR2 | 1.49 | 0.3677 | 0.9288 | 0.000999 |
| MacAngio | TGD | MIF | ITGA4 | 1.1844 | 0.3968 | 0.7906 | 0.000999 |
| MacAngio | TGD | NAMPT | ANXA6 | 1.2297 | 0.8553 | 1.0425 | 0.000999 |
| MacAngio | TGD | PTPRC | CD2 | 0.4851 | 1.4522 | 0.9687 | 0.000999 |
| MacAngio | TGD | PTPRC | CD8A | 0.4851 | 1.746 | 1.1156 | 0.000999 |
| MacAngio | TGD | RPS19 | ITGA4 | 3.3281 | 0.3968 | 1.8625 | 0.000999 |
| MacAngio | TGD | RPS19 | RPSA | 3.3281 | 2.8814 | 3.1047 | 0.000999 |
| MacAngio | TGD | RTN4 | CD99 | 0.9484 | 1.7154 | 1.3319 | 0.000999 |
| MacAngio | TGD | SPP1 | ITGA4 | 2.926 | 0.3968 | 1.6614 | 0.000999 |
| MacAngio | TGD | SPP1 | ITGB1 | 2.926 | 0.3657 | 1.6458 | 0.000999 |
| MacAngio | TGD | TGFB1 | TGFBR2 | 0.2752 | 0.3677 | 0.3214 | 0.000999 |

| | | | | | | | |
|---|---|---|---|---|---|---|---|
| MacAngio | TGD | THBS1 | ITGA4 | 0.4513 | 0.3968 | 0.424 | 0.000999 |
| MacAngio | TGD | VEGFA | PTK2B | 0.6881 | 0.3668 | 0.5274 | 0.000999 |
| MacAngio | TGD | MIF | TNFRSF14 | 1.1844 | 0.4642 | 0.8243 | 0.002 |
| MacAngio | TGD | LRPAP1 | SORL1 | 0.4668 | 0.3318 | 0.3993 | 0.00699 |
| NKT | MacAngio | ALOX5AP | ALOX5 | 0.5021 | 0.3129 | 0.4075 | 0.000999 |
| NKT | MacAngio | ANXA1 | C3AR1 | 1.2259 | 0.3698 | 0.7979 | 0.000999 |
| NKT | MacAngio | ANXA1 | C5AR1 | 1.2259 | 0.6392 | 0.9326 | 0.000999 |
| NKT | MacAngio | ANXA1 | CCR1 | 1.2259 | 0.2766 | 0.7513 | 0.000999 |
| NKT | MacAngio | ANXA1 | CD44 | 1.2259 | 1.6468 | 1.4364 | 0.000999 |
| NKT | MacAngio | ANXA1 | FPR1 | 1.2259 | 0.5081 | 0.867 | 0.000999 |
| NKT | MacAngio | ANXA1 | FPR3 | 1.2259 | 0.5507 | 0.8883 | 0.000999 |
| NKT | MacAngio | ARF1 | CD44 | 0.7991 | 1.6468 | 1.223 | 0.000999 |
| NKT | MacAngio | CALM1 | TNFRSF1B | 1.9335 | 0.4897 | 1.2116 | 0.000999 |
| NKT | MacAngio | CALR | CD44 | 0.9479 | 1.6468 | 1.2974 | 0.000999 |
| NKT | MacAngio | CALR | ITGA5 | 0.9479 | 0.2751 | 0.6115 | 0.000999 |
| NKT | MacAngio | CALR | LRP1 | 0.9479 | 0.3542 | 0.6511 | 0.000999 |
| NKT | MacAngio | CD6 | ALCAM | 0.4881 | 0.3699 | 0.429 | 0.000999 |
| NKT | MacAngio | GSTP1 | CD44 | 0.7909 | 1.6468 | 1.2189 | 0.000999 |
| NKT | MacAngio | HMGB1 | AGTRAP | 1.195 | 0.302 | 0.7485 | 0.000999 |
| NKT | MacAngio | HMGB1 | LY96 | 1.195 | 0.5458 | 0.8704 | 0.000999 |
| NKT | MacAngio | HMGB1 | TLR2 | 1.195 | 0.3171 | 0.756 | 0.000999 |
| NKT | MacAngio | HMGB2 | CD44 | 0.6467 | 1.6468 | 1.1468 | 0.000999 |
| NKT | MacAngio | HMGB2 | DAB2 | 0.6467 | 0.5892 | 0.6179 | 0.000999 |
| NKT | MacAngio | HSP90B1 | LRP1 | 0.6138 | 0.3542 | 0.484 | 0.000999 |
| NKT | MacAngio | ITGA4 | CD44 | 0.2946 | 1.6468 | 0.9707 | 0.000999 |
| NKT | MacAngio | ITGA4 | TFRC | 0.2946 | 0.4794 | 0.387 | 0.000999 |
| NKT | MacAngio | ITGAL | ICAM1 | 0.6778 | 0.2944 | 0.4861 | 0.000999 |
| NKT | MacAngio | ITGB1 | CD151 | 0.488 | 0.3151 | 0.4016 | 0.000999 |
| NKT | MacAngio | ITGB1 | CD36 | 0.488 | 0.2591 | 0.3735 | 0.000999 |
| NKT | MacAngio | ITGB1 | DAB2 | 0.488 | 0.5892 | 0.5386 | 0.000999 |

| | | | | | | | |
|---|---|---|---|---|---|---|---|
| NKT | MacAngio | ITGB1 | ENG | 0.488 | 0.4057 | 0.4468 | 0.000999 |
| NKT | MacAngio | ITGB1 | ITGA5 | 0.488 | 0.2751 | 0.3815 | 0.000999 |
| NKT | MacAngio | ITGB1 | ITGAX | 0.488 | 0.438 | 0.463 | 0.000999 |
| NKT | MacAngio | ITGB1 | PLAUR | 0.488 | 1.4699 | 0.9789 | 0.000999 |
| NKT | MacAngio | ITGB2 | ICAM1 | 1.6481 | 0.2944 | 0.9713 | 0.000999 |
| NKT | MacAngio | ITGB2 | ITGAM | 1.6481 | 0.3193 | 0.9837 | 0.000999 |
| NKT | MacAngio | ITGB2 | ITGAX | 1.6481 | 0.438 | 1.0431 | 0.000999 |
| NKT | MacAngio | ITGB2 | PLAUR | 1.6481 | 1.4699 | 1.559 | 0.000999 |
| NKT | MacAngio | LRPAP1 | LRP1 | 0.2807 | 0.3542 | 0.3175 | 0.000999 |
| NKT | MacAngio | LTB | ITGB1 | 0.2605 | 0.5437 | 0.4021 | 0.000999 |
| NKT | MacAngio | MIF | CD74 | 0.6657 | 3.2987 | 1.9822 | 0.000999 |
| NKT | MacAngio | PTPRC | CD44 | 1.8797 | 1.6468 | 1.7633 | 0.000999 |
| NKT | MacAngio | RPS19 | C5AR1 | 3.5336 | 0.6392 | 2.0864 | 0.000999 |
| NKT | MacAngio | RPS19 | ICAM1 | 3.5336 | 0.2944 | 1.914 | 0.000999 |
| NKT | MacAngio | RTN4 | CD44 | 0.4374 | 1.6468 | 1.0421 | 0.000999 |
| NKT | MacAngio | TGFB1 | ENG | 0.5441 | 0.4057 | 0.4749 | 0.000999 |
| NKT | MacAngio | TGFB1 | FCN1 | 0.5441 | 0.6982 | 0.6211 | 0.000999 |
| NKT | MacAngio | TGFB1 | ITGB1 | 0.5441 | 0.5437 | 0.5439 | 0.000999 |
| NKT | MacAngio | TGFB1 | SDC2 | 0.5441 | 0.3815 | 0.4628 | 0.000999 |
| NKT | MacAngio | TGFB1 | TLR2 | 0.5441 | 0.3171 | 0.4306 | 0.000999 |
| TCD4ex | MacAngio | ALOX5AP | ALOX5 | 1.119 | 0.3129 | 0.7159 | 0.000999 |
| TCD4ex | MacAngio | ANXA1 | C5AR1 | 0.5174 | 0.6392 | 0.5783 | 0.000999 |
| TCD4ex | MacAngio | ANXA1 | CD44 | 0.5174 | 1.6468 | 1.0821 | 0.000999 |
| TCD4ex | MacAngio | ANXA1 | FPR1 | 0.5174 | 0.5081 | 0.5128 | 0.000999 |
| TCD4ex | MacAngio | ANXA1 | FPR3 | 0.5174 | 0.5507 | 0.5341 | 0.000999 |
| TCD4ex | MacAngio | ARF1 | CD44 | 0.6827 | 1.6468 | 1.1648 | 0.000999 |
| TCD4ex | MacAngio | CALR | CD44 | 0.748 | 1.6468 | 1.1974 | 0.000999 |
| TCD4ex | MacAngio | CALR | ITGA5 | 0.748 | 0.2751 | 0.5115 | 0.000999 |
| TCD4ex | MacAngio | CALR | LRP1 | 0.748 | 0.3542 | 0.5511 | 0.000999 |
| TCD4ex | MacAngio | CD40LG | ITGA5 | 0.3948 | 0.2751 | 0.335 | 0.000999 |

| | | | | | | | |
|---|---|---|---|---|---|---|---|
| TCD4ex | MacAngio | CD6 | ALCAM | 0.5338 | 0.3699 | 0.4519 | 0.000999 |
| TCD4ex | MacAngio | CXCL13 | C3AR1 | 2.3367 | 0.3698 | 1.3532 | 0.000999 |
| TCD4ex | MacAngio | CXCL13 | C5AR1 | 2.3367 | 0.6392 | 1.4879 | 0.000999 |
| TCD4ex | MacAngio | CXCL13 | CCR1 | 2.3367 | 0.2766 | 1.3067 | 0.000999 |
| TCD4ex | MacAngio | CXCL13 | FPR1 | 2.3367 | 0.5081 | 1.4224 | 0.000999 |
| TCD4ex | MacAngio | GSTP1 | CD44 | 0.4481 | 1.6468 | 1.0475 | 0.000999 |
| TCD4ex | MacAngio | HMGB1 | AGTRAP | 1.3876 | 0.302 | 0.8448 | 0.000999 |
| TCD4ex | MacAngio | HMGB1 | LY96 | 1.3876 | 0.5458 | 0.9667 | 0.000999 |
| TCD4ex | MacAngio | HMGB1 | TLR2 | 1.3876 | 0.3171 | 0.8523 | 0.000999 |
| TCD4ex | MacAngio | HMGB2 | CD44 | 1.0243 | 1.6468 | 1.3356 | 0.000999 |
| TCD4ex | MacAngio | HMGB2 | DAB2 | 1.0243 | 0.5892 | 0.8067 | 0.000999 |
| TCD4ex | MacAngio | HSP90B1 | LRP1 | 0.3944 | 0.3542 | 0.3743 | 0.000999 |
| TCD4ex | MacAngio | ICAM3 | ITGB2 | 0.573 | 0.9554 | 0.7642 | 0.000999 |
| TCD4ex | MacAngio | ITGAL | ICAM1 | 0.2603 | 0.2944 | 0.2774 | 0.000999 |
| TCD4ex | MacAngio | ITGB1 | CD151 | 0.4615 | 0.3151 | 0.3883 | 0.000999 |
| TCD4ex | MacAngio | ITGB1 | CD36 | 0.4615 | 0.2591 | 0.3603 | 0.000999 |
| TCD4ex | MacAngio | ITGB1 | CD47 | 0.4615 | 0.4747 | 0.4681 | 0.000999 |
| TCD4ex | MacAngio | ITGB1 | DAB2 | 0.4615 | 0.5892 | 0.5253 | 0.000999 |
| TCD4ex | MacAngio | ITGB1 | ENG | 0.4615 | 0.4057 | 0.4336 | 0.000999 |
| TCD4ex | MacAngio | ITGB1 | ITGA5 | 0.4615 | 0.2751 | 0.3683 | 0.000999 |
| TCD4ex | MacAngio | ITGB1 | ITGAX | 0.4615 | 0.438 | 0.4498 | 0.000999 |
| TCD4ex | MacAngio | ITGB1 | PLAUR | 0.4615 | 1.4699 | 0.9657 | 0.000999 |
| TCD4ex | MacAngio | ITGB2 | ICAM1 | 0.7871 | 0.2944 | 0.5407 | 0.000999 |
| TCD4ex | MacAngio | ITGB2 | ITGAM | 0.7871 | 0.3193 | 0.5532 | 0.000999 |
| TCD4ex | MacAngio | ITGB2 | ITGAX | 0.7871 | 0.438 | 0.6125 | 0.000999 |
| TCD4ex | MacAngio | ITGB2 | PLAUR | 0.7871 | 1.4699 | 1.1285 | 0.000999 |
| TCD4ex | MacAngio | LRPAP1 | LRP1 | 0.2809 | 0.3542 | 0.3176 | 0.000999 |
| TCD4ex | MacAngio | LTB | ITGB1 | 0.9655 | 0.5437 | 0.7546 | 0.000999 |
| TCD4ex | MacAngio | MIF | CD74 | 0.6999 | 3.2987 | 1.9993 | 0.000999 |
| TCD4ex | MacAngio | PTPRC | CD44 | 1.7493 | 1.6468 | 1.6981 | 0.000999 |

| | | | | | | | |
|---|---|---|---|---|---|---|---|
| TCD4ex | MacAngio | RPS19 | C5AR1 | 3.6329 | 0.6392 | 2.136 | 0.000999 |
| TCD4ex | MacAngio | RPS19 | ICAM1 | 3.6329 | 0.2944 | 1.9637 | 0.000999 |
| TCD4ex | MacAngio | RTN4 | CD44 | 0.355 | 1.6468 | 1.0009 | 0.000999 |
| TCD4ex | MacAngio | ANXA1 | C3AR1 | 0.5174 | 0.3698 | 0.4436 | 0.002 |
| TCD4ex | MacAngio | AREG | ICAM1 | 0.3427 | 0.2944 | 0.3186 | 0.002 |
| TCD4ex | MacAngio | CALM1 | TNFRSF1B | 1.2957 | 0.4897 | 0.8927 | 0.004 |
| TCD4reg | MacAngio | ALOX5AP | ALOX5 | 0.4589 | 0.3129 | 0.3859 | 0.000999 |
| TCD4reg | MacAngio | APOE | LRP1 | 0.2934 | 0.3542 | 0.3238 | 0.000999 |
| TCD4reg | MacAngio | APOE | SDC2 | 0.2934 | 0.3815 | 0.3375 | 0.000999 |
| TCD4reg | MacAngio | ARF1 | CD44 | 0.7185 | 1.6468 | 1.1827 | 0.000999 |
| TCD4reg | MacAngio | CALR | CD44 | 0.6835 | 1.6468 | 1.1652 | 0.000999 |
| TCD4reg | MacAngio | CALR | ITGA5 | 0.6835 | 0.2751 | 0.4793 | 0.000999 |
| TCD4reg | MacAngio | CALR | LRP1 | 0.6835 | 0.3542 | 0.5189 | 0.000999 |
| TCD4reg | MacAngio | CD6 | ALCAM | 0.2651 | 0.3699 | 0.3175 | 0.000999 |
| TCD4reg | MacAngio | GSTP1 | CD44 | 0.4917 | 1.6468 | 1.0692 | 0.000999 |
| TCD4reg | MacAngio | HMGB1 | AGTRAP | 1.2547 | 0.302 | 0.7784 | 0.000999 |
| TCD4reg | MacAngio | HMGB1 | LY96 | 1.2547 | 0.5458 | 0.9003 | 0.000999 |
| TCD4reg | MacAngio | HMGB1 | TLR2 | 1.2547 | 0.3171 | 0.7859 | 0.000999 |
| TCD4reg | MacAngio | HMGB2 | CD44 | 0.616 | 1.6468 | 1.1314 | 0.000999 |
| TCD4reg | MacAngio | HMGB2 | DAB2 | 0.616 | 0.5892 | 0.6026 | 0.000999 |
| TCD4reg | MacAngio | HSP90B1 | LRP1 | 0.4221 | 0.3542 | 0.3882 | 0.000999 |
| TCD4reg | MacAngio | ICAM3 | ITGB2 | 0.6663 | 0.9554 | 0.8108 | 0.000999 |
| TCD4reg | MacAngio | ITGB1 | CD151 | 0.4001 | 0.3151 | 0.3576 | 0.000999 |
| TCD4reg | MacAngio | ITGB1 | CD36 | 0.4001 | 0.2591 | 0.3296 | 0.000999 |
| TCD4reg | MacAngio | ITGB1 | DAB2 | 0.4001 | 0.5892 | 0.4946 | 0.000999 |
| TCD4reg | MacAngio | ITGB1 | ENG | 0.4001 | 0.4057 | 0.4029 | 0.000999 |
| TCD4reg | MacAngio | ITGB1 | ITGA5 | 0.4001 | 0.2751 | 0.3376 | 0.000999 |
| TCD4reg | MacAngio | ITGB1 | ITGAX | 0.4001 | 0.438 | 0.4191 | 0.000999 |
| TCD4reg | MacAngio | ITGB1 | PLAUR | 0.4001 | 1.4699 | 0.935 | 0.000999 |
| TCD4reg | MacAngio | ITGB2 | ICAM1 | 0.6238 | 0.2944 | 0.4591 | 0.000999 |

| | | | | | | | |
|---|---|---|---|---|---|---|---|
| TCD4reg | MacAngio | ITGB2 | ITGAM | 0.6238 | 0.3193 | 0.4716 | 0.000999 |
| TCD4reg | MacAngio | ITGB2 | ITGAX | 0.6238 | 0.438 | 0.5309 | 0.000999 |
| TCD4reg | MacAngio | ITGB2 | PLAUR | 0.6238 | 1.4699 | 1.0469 | 0.000999 |
| TCD4reg | MacAngio | LGALS3 | CLEC7A | 0.4492 | 0.5046 | 0.4769 | 0.000999 |
| TCD4reg | MacAngio | LGALS3 | FCGR2A | 0.4492 | 0.7158 | 0.5825 | 0.000999 |
| TCD4reg | MacAngio | LGALS3 | FCGR2B | 0.4492 | 0.4543 | 0.4517 | 0.000999 |
| TCD4reg | MacAngio | LGALS3 | TFRC | 0.4492 | 0.4794 | 0.4643 | 0.000999 |
| TCD4reg | MacAngio | LRPAP1 | LRP1 | 0.3101 | 0.3542 | 0.3322 | 0.000999 |
| TCD4reg | MacAngio | LTB | ITGB1 | 2.242 | 0.5437 | 1.3929 | 0.000999 |
| TCD4reg | MacAngio | MIF | CD74 | 0.6295 | 3.2987 | 1.9641 | 0.000999 |
| TCD4reg | MacAngio | PTPRC | CD44 | 1.6603 | 1.6468 | 1.6536 | 0.000999 |
| TCD4reg | MacAngio | RPS19 | C5AR1 | 3.6897 | 0.6392 | 2.1644 | 0.000999 |
| TCD4reg | MacAngio | RPS19 | ICAM1 | 3.6897 | 0.2944 | 1.992 | 0.000999 |
| TCD4reg | MacAngio | RTN4 | CD44 | 0.3528 | 1.6468 | 0.9998 | 0.000999 |
| TCD4reg | MacAngio | SPP1 | CD44 | 0.3125 | 1.6468 | 0.9797 | 0.000999 |
| TCD4reg | MacAngio | HSP90B1 | ITGB1 | 0.4221 | 0.5437 | 0.4829 | 0.002 |
| TCD8em | MacAngio | ALOX5AP | ALOX5 | 0.7973 | 0.3129 | 0.5551 | 0.000999 |
| TCD8em | MacAngio | ANXA1 | C3AR1 | 1.0693 | 0.3698 | 0.7195 | 0.000999 |
| TCD8em | MacAngio | ANXA1 | C5AR1 | 1.0693 | 0.6392 | 0.8542 | 0.000999 |
| TCD8em | MacAngio | ANXA1 | CCR1 | 1.0693 | 0.2766 | 0.6729 | 0.000999 |
| TCD8em | MacAngio | ANXA1 | CD44 | 1.0693 | 1.6468 | 1.358 | 0.000999 |
| TCD8em | MacAngio | ANXA1 | FPR1 | 1.0693 | 0.5081 | 0.7887 | 0.000999 |
| TCD8em | MacAngio | ANXA1 | FPR3 | 1.0693 | 0.5507 | 0.81 | 0.000999 |
| TCD8em | MacAngio | APOE | LRP1 | 0.2791 | 0.3542 | 0.3167 | 0.000999 |
| TCD8em | MacAngio | APOE | SDC2 | 0.2791 | 0.3815 | 0.3303 | 0.000999 |
| TCD8em | MacAngio | APOE | SDC4 | 0.2791 | 0.2598 | 0.2695 | 0.000999 |
| TCD8em | MacAngio | ARF1 | CD44 | 0.6684 | 1.6468 | 1.1576 | 0.000999 |
| TCD8em | MacAngio | CALM1 | TNFRSF1B | 1.5262 | 0.4897 | 1.0079 | 0.000999 |
| TCD8em | MacAngio | CALR | CD44 | 0.7141 | 1.6468 | 1.1805 | 0.000999 |
| TCD8em | MacAngio | CALR | ITGA5 | 0.7141 | 0.2751 | 0.4946 | 0.000999 |

| | | | | | | | |
|---|---|---|---|---|---|---|---|
| TCD8em | MacAngio | CALR | LRP1 | 0.7141 | 0.3542 | 0.5342 | 0.000999 |
| TCD8em | MacAngio | CD6 | ALCAM | 0.4204 | 0.3699 | 0.3952 | 0.000999 |
| TCD8em | MacAngio | GSTP1 | CD44 | 0.767 | 1.6468 | 1.2069 | 0.000999 |
| TCD8em | MacAngio | HMGB1 | AGTRAP | 1.3636 | 0.302 | 0.8328 | 0.000999 |
| TCD8em | MacAngio | HMGB1 | LY96 | 1.3636 | 0.5458 | 0.9547 | 0.000999 |
| TCD8em | MacAngio | HMGB1 | TLR2 | 1.3636 | 0.3171 | 0.8403 | 0.000999 |
| TCD8em | MacAngio | HMGB2 | CD44 | 0.7212 | 1.6468 | 1.184 | 0.000999 |
| TCD8em | MacAngio | HMGB2 | DAB2 | 0.7212 | 0.5892 | 0.6552 | 0.000999 |
| TCD8em | MacAngio | HSP90B1 | ITGB1 | 0.471 | 0.5437 | 0.5074 | 0.000999 |
| TCD8em | MacAngio | HSP90B1 | LRP1 | 0.471 | 0.3542 | 0.4126 | 0.000999 |
| TCD8em | MacAngio | HSP90B1 | TNFRSF1B | 0.471 | 0.4897 | 0.4804 | 0.000999 |
| TCD8em | MacAngio | IFNG | IFNGR1 | 0.3967 | 0.4333 | 0.415 | 0.000999 |
| TCD8em | MacAngio | IFNG | IFNGR2 | 0.3967 | 0.6578 | 0.5272 | 0.000999 |
| TCD8em | MacAngio | IFNG | PTPN6 | 0.3967 | 0.363 | 0.3799 | 0.000999 |
| TCD8em | MacAngio | ITGA4 | CD44 | 0.2823 | 1.6468 | 0.9646 | 0.000999 |
| TCD8em | MacAngio | ITGA4 | TFRC | 0.2823 | 0.4794 | 0.3809 | 0.000999 |
| TCD8em | MacAngio | ITGAL | ICAM1 | 0.3162 | 0.2944 | 0.3053 | 0.000999 |
| TCD8em | MacAngio | ITGB1 | CD151 | 0.2557 | 0.3151 | 0.2854 | 0.000999 |
| TCD8em | MacAngio | ITGB1 | CD36 | 0.2557 | 0.2591 | 0.2574 | 0.000999 |
| TCD8em | MacAngio | ITGB1 | DAB2 | 0.2557 | 0.5892 | 0.4225 | 0.000999 |
| TCD8em | MacAngio | ITGB1 | ENG | 0.2557 | 0.4057 | 0.3307 | 0.000999 |
| TCD8em | MacAngio | ITGB1 | ITGA5 | 0.2557 | 0.2751 | 0.2654 | 0.000999 |
| TCD8em | MacAngio | ITGB1 | ITGAX | 0.2557 | 0.438 | 0.3469 | 0.000999 |
| TCD8em | MacAngio | ITGB1 | PLAUR | 0.2557 | 1.4699 | 0.8628 | 0.000999 |
| TCD8em | MacAngio | ITGB2 | ICAM1 | 1.0018 | 0.2944 | 0.6481 | 0.000999 |
| TCD8em | MacAngio | ITGB2 | ITGAM | 1.0018 | 0.3193 | 0.6606 | 0.000999 |
| TCD8em | MacAngio | ITGB2 | ITGAX | 1.0018 | 0.438 | 0.7199 | 0.000999 |
| TCD8em | MacAngio | ITGB2 | PLAUR | 1.0018 | 1.4699 | 1.2359 | 0.000999 |
| TCD8em | MacAngio | LGALS3 | CLEC7A | 0.2689 | 0.5046 | 0.3868 | 0.000999 |
| TCD8em | MacAngio | LGALS3 | FCGR2A | 0.2689 | 0.7158 | 0.4924 | 0.000999 |

| | | | | | | | |
|---|---|---|---|---|---|---|---|
| TCD8em | MacAngio | LGALS3 | FCGR2B | 0.2689 | 0.4543 | 0.3616 | 0.000999 |
| TCD8em | MacAngio | LGALS3 | ITGB1 | 0.2689 | 0.5437 | 0.4063 | 0.000999 |
| TCD8em | MacAngio | LGALS3 | TFRC | 0.2689 | 0.4794 | 0.3742 | 0.000999 |
| TCD8em | MacAngio | LRPAP1 | LRP1 | 0.2719 | 0.3542 | 0.313 | 0.000999 |
| TCD8em | MacAngio | LTB | ITGB1 | 0.6352 | 0.5437 | 0.5894 | 0.000999 |
| TCD8em | MacAngio | MIF | CD74 | 0.6063 | 3.2987 | 1.9525 | 0.000999 |
| TCD8em | MacAngio | PSAP | LRP1 | 0.5012 | 0.3542 | 0.4277 | 0.000999 |
| TCD8em | MacAngio | PTPRC | CD44 | 1.7912 | 1.6468 | 1.719 | 0.000999 |
| TCD8em | MacAngio | RPS19 | C5AR1 | 3.536 | 0.6392 | 2.0876 | 0.000999 |
| TCD8em | MacAngio | RPS19 | ICAM1 | 3.536 | 0.2944 | 1.9152 | 0.000999 |
| TCD8em | MacAngio | RTN4 | CD44 | 0.3632 | 1.6468 | 1.005 | 0.000999 |
| TCD8em | MacAngio | SPP1 | CD44 | 0.3608 | 1.6468 | 1.0038 | 0.000999 |
| TCD8em | MacAngio | TNF | TNFRSF1B | 0.2667 | 0.4897 | 0.3782 | 0.000999 |
| TCD8ex | MacAngio | ALOX5AP | ALOX5 | 1.5024 | 0.3129 | 0.9076 | 0.000999 |
| TCD8ex | MacAngio | ANXA1 | C3AR1 | 0.5645 | 0.3698 | 0.4671 | 0.000999 |
| TCD8ex | MacAngio | ANXA1 | C5AR1 | 0.5645 | 0.6392 | 0.6018 | 0.000999 |
| TCD8ex | MacAngio | ANXA1 | CCR1 | 0.5645 | 0.2766 | 0.4206 | 0.000999 |
| TCD8ex | MacAngio | ANXA1 | CD44 | 0.5645 | 1.6468 | 1.1057 | 0.000999 |
| TCD8ex | MacAngio | ANXA1 | FPR1 | 0.5645 | 0.5081 | 0.5363 | 0.000999 |
| TCD8ex | MacAngio | ANXA1 | FPR3 | 0.5645 | 0.5507 | 0.5576 | 0.000999 |
| TCD8ex | MacAngio | APOE | LRP1 | 0.2814 | 0.3542 | 0.3178 | 0.000999 |
| TCD8ex | MacAngio | APOE | SDC2 | 0.2814 | 0.3815 | 0.3314 | 0.000999 |
| TCD8ex | MacAngio | APOE | SDC4 | 0.2814 | 0.2598 | 0.2706 | 0.000999 |
| TCD8ex | MacAngio | AREG | ICAM1 | 0.2758 | 0.2944 | 0.2851 | 0.000999 |
| TCD8ex | MacAngio | ARF1 | CD44 | 0.7835 | 1.6468 | 1.2152 | 0.000999 |
| TCD8ex | MacAngio | CALR | CD44 | 0.9634 | 1.6468 | 1.3051 | 0.000999 |
| TCD8ex | MacAngio | CALR | ITGA5 | 0.9634 | 0.2751 | 0.6192 | 0.000999 |
| TCD8ex | MacAngio | CALR | LRP1 | 0.9634 | 0.3542 | 0.6588 | 0.000999 |
| TCD8ex | MacAngio | CD6 | ALCAM | 0.318 | 0.3699 | 0.344 | 0.000999 |
| TCD8ex | MacAngio | CXCL13 | C3AR1 | 1.7749 | 0.3698 | 1.0724 | 0.000999 |

| | | | | | | | |
|---|---|---|---|---|---|---|---|
| TCD8ex | MacAngio | CXCL13 | C5AR1 | 1.7749 | 0.6392 | 1.2071 | 0.000999 |
| TCD8ex | MacAngio | CXCL13 | CCR1 | 1.7749 | 0.2766 | 1.0258 | 0.000999 |
| TCD8ex | MacAngio | CXCL13 | FPR1 | 1.7749 | 0.5081 | 1.1415 | 0.000999 |
| TCD8ex | MacAngio | GSTP1 | CD44 | 0.8286 | 1.6468 | 1.2377 | 0.000999 |
| TCD8ex | MacAngio | HMGB1 | AGTRAP | 1.5414 | 0.302 | 0.9217 | 0.000999 |
| TCD8ex | MacAngio | HMGB1 | LY96 | 1.5414 | 0.5458 | 1.0436 | 0.000999 |
| TCD8ex | MacAngio | HMGB1 | TLR2 | 1.5414 | 0.3171 | 0.9293 | 0.000999 |
| TCD8ex | MacAngio | HMGB2 | CD44 | 0.85 | 1.6468 | 1.2484 | 0.000999 |
| TCD8ex | MacAngio | HMGB2 | DAB2 | 0.85 | 0.5892 | 0.7196 | 0.000999 |
| TCD8ex | MacAngio | HSP90B1 | ITGB1 | 0.5666 | 0.5437 | 0.5552 | 0.000999 |
| TCD8ex | MacAngio | HSP90B1 | LRP1 | 0.5666 | 0.3542 | 0.4604 | 0.000999 |
| TCD8ex | MacAngio | ICAM3 | ITGB2 | 0.5405 | 0.9554 | 0.748 | 0.000999 |
| TCD8ex | MacAngio | IFNG | IFNGR1 | 0.8575 | 0.4333 | 0.6454 | 0.000999 |
| TCD8ex | MacAngio | IFNG | IFNGR2 | 0.8575 | 0.6578 | 0.7577 | 0.000999 |
| TCD8ex | MacAngio | ITGA4 | CD44 | 0.4432 | 1.6468 | 1.045 | 0.000999 |
| TCD8ex | MacAngio | ITGA4 | TFRC | 0.4432 | 0.4794 | 0.4613 | 0.000999 |
| TCD8ex | MacAngio | ITGB1 | CD151 | 0.2995 | 0.3151 | 0.3073 | 0.000999 |
| TCD8ex | MacAngio | ITGB1 | CD36 | 0.2995 | 0.2591 | 0.2793 | 0.000999 |
| TCD8ex | MacAngio | ITGB1 | DAB2 | 0.2995 | 0.5892 | 0.4443 | 0.000999 |
| TCD8ex | MacAngio | ITGB1 | ENG | 0.2995 | 0.4057 | 0.3526 | 0.000999 |
| TCD8ex | MacAngio | ITGB1 | ITGA5 | 0.2995 | 0.2751 | 0.2873 | 0.000999 |
| TCD8ex | MacAngio | ITGB1 | ITGAX | 0.2995 | 0.438 | 0.3688 | 0.000999 |
| TCD8ex | MacAngio | ITGB1 | PLAUR | 0.2995 | 1.4699 | 0.8847 | 0.000999 |
| TCD8ex | MacAngio | ITGB2 | ICAM1 | 0.7635 | 0.2944 | 0.5289 | 0.000999 |
| TCD8ex | MacAngio | ITGB2 | ITGAM | 0.7635 | 0.3193 | 0.5414 | 0.000999 |
| TCD8ex | MacAngio | ITGB2 | ITGAX | 0.7635 | 0.438 | 0.6007 | 0.000999 |
| TCD8ex | MacAngio | ITGB2 | PLAUR | 0.7635 | 1.4699 | 1.1167 | 0.000999 |
| TCD8ex | MacAngio | LGALS3 | CLEC7A | 0.3824 | 0.5046 | 0.4435 | 0.000999 |
| TCD8ex | MacAngio | LGALS3 | FCGR2A | 0.3824 | 0.7158 | 0.5491 | 0.000999 |
| TCD8ex | MacAngio | LGALS3 | FCGR2B | 0.3824 | 0.4543 | 0.4183 | 0.000999 |

| | | | | | | | |
|---|---|---|---|---|---|---|---|
| TCD8ex | MacAngio | LGALS3 | TFRC | 0.3824 | 0.4794 | 0.4309 | 0.000999 |
| TCD8ex | MacAngio | LRPAP1 | LRP1 | 0.2971 | 0.3542 | 0.3257 | 0.000999 |
| TCD8ex | MacAngio | LTB | ITGB1 | 0.4576 | 0.5437 | 0.5007 | 0.000999 |
| TCD8ex | MacAngio | MIF | CD74 | 0.749 | 3.2987 | 2.0239 | 0.000999 |
| TCD8ex | MacAngio | PTPRC | CD44 | 1.6762 | 1.6468 | 1.6615 | 0.000999 |
| TCD8ex | MacAngio | RPS19 | C5AR1 | 3.8353 | 0.6392 | 2.2372 | 0.000999 |
| TCD8ex | MacAngio | RPS19 | ICAM1 | 3.8353 | 0.2944 | 2.0648 | 0.000999 |
| TCD8ex | MacAngio | RTN4 | CD44 | 0.3033 | 1.6468 | 0.9751 | 0.000999 |
| TCD8ex | MacAngio | SPP1 | CD44 | 0.2978 | 1.6468 | 0.9723 | 0.000999 |
| TCD8ex | MacAngio | VCAM1 | ITGA5 | 0.4462 | 0.2751 | 0.3606 | 0.000999 |
| TCD8ex | MacAngio | VCAM1 | ITGB1 | 0.4462 | 0.5437 | 0.495 | 0.000999 |
| TCD8ex | MacAngio | VCAM1 | ITGB2 | 0.4462 | 0.9554 | 0.7008 | 0.000999 |
| TCD8ex | MacAngio | VCAM1 | TFRC | 0.4462 | 0.4794 | 0.4628 | 0.000999 |
| TCD8ex | MacAngio | IFNG | PTPN6 | 0.8575 | 0.363 | 0.6103 | 0.003 |
| TCD8ex | MacAngio | LGALS3 | ITGB1 | 0.3824 | 0.5437 | 0.4631 | 0.003 |
| TGD | MacAngio | ARF1 | CD44 | 0.9453 | 1.6468 | 1.2961 | 0.000999 |
| TGD | MacAngio | CALM1 | TNFRSF1B | 1.7587 | 0.4897 | 1.1242 | 0.000999 |
| TGD | MacAngio | CD2 | PTPRC | 1.4522 | 0.4851 | 0.9687 | 0.000999 |
| TGD | MacAngio | CD6 | ALCAM | 0.4684 | 0.3699 | 0.4192 | 0.000999 |
| TGD | MacAngio | CD99 | CD99 | 1.7154 | 0.5878 | 1.1516 | 0.000999 |
| TGD | MacAngio | COL1A1 | CD36 | 0.4518 | 0.2591 | 0.3554 | 0.000999 |
| TGD | MacAngio | COL1A1 | CD44 | 0.4518 | 1.6468 | 1.0493 | 0.000999 |
| TGD | MacAngio | COL1A1 | ITGA5 | 0.4518 | 0.2751 | 0.3635 | 0.000999 |
| TGD | MacAngio | COL1A1 | ITGB1 | 0.4518 | 0.5437 | 0.4978 | 0.000999 |
| TGD | MacAngio | HMGB1 | AGTRAP | 1.7525 | 0.302 | 1.0272 | 0.000999 |
| TGD | MacAngio | HMGB1 | LY96 | 1.7525 | 0.5458 | 1.1491 | 0.000999 |
| TGD | MacAngio | HMGB1 | TLR2 | 1.7525 | 0.3171 | 1.0348 | 0.000999 |
| TGD | MacAngio | ICAM3 | ITGB2 | 0.8816 | 0.9554 | 0.9185 | 0.000999 |
| TGD | MacAngio | ITGA4 | CD44 | 0.3968 | 1.6468 | 1.0218 | 0.000999 |
| TGD | MacAngio | ITGA4 | CXCR4 | 0.3968 | 1.0335 | 0.7152 | 0.000999 |

| | | | | | | | |
|---|---|---|---|---|---|---|---|
| TGD | MacAngio | ITGA4 | RPSA | 0.3968 | 1.8199 | 1.1084 | 0.000999 |
| TGD | MacAngio | ITGA4 | TFRC | 0.3968 | 0.4794 | 0.4381 | 0.000999 |
| TGD | MacAngio | ITGAL | ICAM1 | 0.3731 | 0.2944 | 0.3338 | 0.000999 |
| TGD | MacAngio | ITGB2 | ICAM1 | 1.4444 | 0.2944 | 0.8694 | 0.000999 |
| TGD | MacAngio | ITGB2 | ITGAM | 1.4444 | 0.3193 | 0.8819 | 0.000999 |
| TGD | MacAngio | ITGB2 | ITGAX | 1.4444 | 0.438 | 0.9412 | 0.000999 |
| TGD | MacAngio | ITGB2 | PLAUR | 1.4444 | 1.4699 | 1.4572 | 0.000999 |
| TGD | MacAngio | LTB | ITGB1 | 1.6829 | 0.5437 | 1.1133 | 0.000999 |
| TGD | MacAngio | MIF | CD74 | 1.6335 | 3.2987 | 2.4661 | 0.000999 |
| TGD | MacAngio | PTPRC | CD44 | 1.8653 | 1.6468 | 1.7561 | 0.000999 |
| TGD | MacAngio | PTPRC | CXCR4 | 1.8653 | 1.0335 | 1.4494 | 0.000999 |
| TGD | MacAngio | RPS19 | C5AR1 | 3.8902 | 0.6392 | 2.2647 | 0.000999 |
| TGD | MacAngio | RPS19 | ICAM1 | 3.8902 | 0.2944 | 2.0923 | 0.000999 |
| TGD | MacAngio | RPS19 | RPSA | 3.8902 | 1.8199 | 2.8551 | 0.000999 |
| TGD | MacAngio | SEMA4D | PTPRC | 0.5658 | 0.4851 | 0.5254 | 0.000999 |
| TGD | MacAngio | TGFB1 | ENG | 0.7703 | 0.4057 | 0.588 | 0.000999 |
| TGD | MacAngio | TGFB1 | FCN1 | 0.7703 | 0.6982 | 0.7343 | 0.000999 |
| TGD | MacAngio | TGFB1 | ITGB1 | 0.7703 | 0.5437 | 0.657 | 0.000999 |
| TGD | MacAngio | TGFB1 | SDC2 | 0.7703 | 0.3815 | 0.5759 | 0.000999 |
| TGD | MacAngio | TGFB1 | TLR2 | 0.7703 | 0.3171 | 0.5437 | 0.000999 |
| TGD | MacAngio | CALM1 | RPSA | 1.7587 | 1.8199 | 1.7893 | 0.003 |

| Supplementary Table 8: Communication results between macrophages and lymphocytes subpopulations, and malignant cells | | | | | | | |
|---|---|---|---|---|---|---|---|
| **MacAngio vs. malignant cells** | | | | | | | |
| celltype_01 | celltype_02 | ligand | receptor | ligand_mean_expression | receptor_mean_expression | ligand_receptor_mean_expression | pvalue |
| MacAngio | breasttumor | ADAM17 | TNFRSF1A | 0.2751 | 0.2536 | 0.2643 | 0.000999 |
| MacAngio | breasttumor | APOE | SCARB1 | 1.2713 | 0.2628 | 0.767 | 0.000999 |
| MacAngio | breasttumor | APOE | SDC2 | 1.2713 | 1.071 | 1.1712 | 0.000999 |
| MacAngio | breasttumor | CALM1 | MYLK | 1.3231 | 0.4064 | 0.8647 | 0.000999 |
| MacAngio | breasttumor | CALM1 | RPSA | 1.3231 | 2.358 | 1.8405 | 0.000999 |
| MacAngio | breasttumor | CALM1 | TNFRSF1A | 1.3231 | 0.2536 | 0.7883 | 0.000999 |
| MacAngio | breasttumor | CALM2 | MYLK | 1.4916 | 0.4064 | 0.949 | 0.000999 |
| MacAngio | breasttumor | CALM3 | MYLK | 0.6655 | 0.4064 | 0.536 | 0.000999 |
| MacAngio | breasttumor | CD99 | CD99 | 0.5897 | 0.3871 | 0.4884 | 0.000999 |
| MacAngio | breasttumor | FN1 | CD44 | 1.5252 | 0.3247 | 0.925 | 0.000999 |
| MacAngio | breasttumor | FN1 | ITGB1 | 1.5252 | 0.2939 | 0.9095 | 0.000999 |
| MacAngio | breasttumor | FN1 | RPSA | 1.5252 | 2.358 | 1.9416 | 0.000999 |
| MacAngio | breasttumor | FN1 | SDC2 | 1.5252 | 1.071 | 1.2981 | 0.000999 |
| MacAngio | breasttumor | GRN | TNFRSF1A | 1.4063 | 0.2536 | 0.8299 | 0.000999 |
| MacAngio | breasttumor | HLA-DRA | SCARB1 | 3.3453 | 0.2628 | 1.8041 | 0.000999 |
| MacAngio | breasttumor | ITGB1 | CD151 | 0.5432 | 0.3569 | 0.4501 | 0.000999 |
| MacAngio | breasttumor | ITGB1 | RACK1 | 0.5432 | 2.6334 | 1.5883 | 0.000999 |
| MacAngio | breasttumor | LGALS3 | ITGB1 | 1.4887 | 0.2939 | 0.8913 | 0.000999 |
| MacAngio | breasttumor | RTN4 | CD99 | 0.9475 | 0.3871 | 0.6673 | 0.000999 |
| MacAngio | breasttumor | SERPINA1 | SDC2 | 1.2518 | 1.071 | 1.1614 | 0.000999 |
| MacAngio | breasttumor | SPP1 | CD44 | 2.927 | 0.3247 | 1.6258 | 0.000999 |
| MacAngio | breasttumor | SPP1 | ITGB1 | 2.927 | 0.2939 | 1.6104 | 0.000999 |
| MacAngio | breasttumor | TGFB1 | SDC2 | 0.2749 | 1.071 | 0.673 | 0.000999 |
| MacAngio | breasttumor | THBS1 | ITGB1 | 0.4532 | 0.2939 | 0.3736 | 0.000999 |
| MacAngio | breasttumor | CALR | MYLK | 1.0603 | 0.4064 | 0.7333 | 0.002 |
| MacAngio | breasttumor | TGFB1 | ITGB1 | 0.2749 | 0.2939 | 0.2844 | 0.005 |
| MacAngio | colorectaltumor | APOE | SDC1 | 1.2713 | 0.2575 | 0.7644 | 0.000999 |

| | | | | | | | |
|---|---|---|---|---|---|---|---|
| MacAngio | colorectaltumor | CALM1 | RPSA | 1.3231 | 2.251 | 1.7871 | 0.000999 |
| MacAngio | colorectaltumor | COL1A1 | SDC1 | 0.2972 | 0.2575 | 0.2773 | 0.000999 |
| MacAngio | colorectaltumor | FN1 | RPSA | 1.5252 | 2.251 | 1.8881 | 0.000999 |
| MacAngio | colorectaltumor | FN1 | SDC1 | 1.5252 | 0.2575 | 0.8913 | 0.000999 |
| MacAngio | colorectaltumor | ITGB1 | CD151 | 0.5432 | 0.6439 | 0.5936 | 0.000999 |
| MacAngio | colorectaltumor | ITGB1 | CD46 | 0.5432 | 0.3583 | 0.4508 | 0.000999 |
| MacAngio | colorectaltumor | ITGB1 | CDH17 | 0.5432 | 0.254 | 0.3986 | 0.000999 |
| MacAngio | colorectaltumor | ITGB1 | RACK1 | 0.5432 | 2.7146 | 1.6289 | 0.000999 |
| MacAngio | colorectaltumor | ITGB1 | SDC1 | 0.5432 | 0.2575 | 0.4003 | 0.000999 |
| MacAngio | colorectaltumor | LGALS3 | ITGB4 | 1.4887 | 0.2913 | 0.89 | 0.000999 |
| MacAngio | colorectaltumor | SPP1 | ITGB1 | 2.927 | 0.4214 | 1.6742 | 0.000999 |
| MacAngio | colorectaltumor | SPP1 | SDC1 | 2.927 | 0.2575 | 1.5922 | 0.000999 |
| MacAngio | colorectaltumor | THBS1 | SDC1 | 0.4532 | 0.2575 | 0.3553 | 0.000999 |
| MacAngio | colorectaltumor | FN1 | ITGB1 | 1.5252 | 0.4214 | 0.9733 | 0.002 |
| MacAngio | colorectaltumor | RPS19 | RPSA | 3.3275 | 2.251 | 2.7893 | 0.002 |
| MacAngio | livertumor | ADAM17 | ITGB1 | 0.2751 | 1.6044 | 0.9398 | 0.000999 |
| MacAngio | livertumor | ADM | RAMP1 | 0.4993 | 0.4499 | 0.4746 | 0.000999 |
| MacAngio | livertumor | ANXA1 | MET | 1.3105 | 0.2983 | 0.8044 | 0.000999 |
| MacAngio | livertumor | ANXA1 | PTPRF | 1.3105 | 0.384 | 0.8473 | 0.000999 |
| MacAngio | livertumor | APOE | SDC1 | 1.2713 | 0.5354 | 0.9034 | 0.000999 |
| MacAngio | livertumor | APOE | SDC4 | 1.2713 | 0.6068 | 0.9391 | 0.000999 |
| MacAngio | livertumor | AREG | ERBB3 | 0.8518 | 0.2644 | 0.5581 | 0.000999 |
| MacAngio | livertumor | ARF1 | INSR | 0.6474 | 0.3039 | 0.4757 | 0.000999 |
| MacAngio | livertumor | CALM1 | INSR | 1.3231 | 0.3039 | 0.8135 | 0.000999 |
| MacAngio | livertumor | CALM1 | RPSA | 1.3231 | 2.6317 | 1.9774 | 0.000999 |
| MacAngio | livertumor | CALR | ITGA3 | 1.0603 | 0.3913 | 0.7258 | 0.000999 |
| MacAngio | livertumor | CD99 | EPHB4 | 0.5897 | 0.4535 | 0.5216 | 0.000999 |
| MacAngio | livertumor | CD99 | INSR | 0.5897 | 0.3039 | 0.4468 | 0.000999 |
| MacAngio | livertumor | COL1A1 | ITGA2 | 0.2972 | 0.3983 | 0.3478 | 0.000999 |
| MacAngio | livertumor | COL1A1 | ITGA3 | 0.2972 | 0.3913 | 0.3443 | 0.000999 |

| | | | | | | | |
|---|---|---|---|---|---|---|---|
| MacAngio | livertumor | COL1A1 | ITGB1 | 0.2972 | 1.6044 | 0.9508 | 0.000999 |
| MacAngio | livertumor | COL1A1 | SDC1 | 0.2972 | 0.5354 | 0.4163 | 0.000999 |
| MacAngio | livertumor | EREG | ERBB3 | 0.6003 | 0.2644 | 0.4323 | 0.000999 |
| MacAngio | livertumor | FN1 | ITGA2 | 1.5252 | 0.3983 | 0.9617 | 0.000999 |
| MacAngio | livertumor | FN1 | ITGA3 | 1.5252 | 0.3913 | 0.9582 | 0.000999 |
| MacAngio | livertumor | FN1 | ITGB1 | 1.5252 | 1.6044 | 1.5648 | 0.000999 |
| MacAngio | livertumor | FN1 | RPSA | 1.5252 | 2.6317 | 2.0784 | 0.000999 |
| MacAngio | livertumor | FN1 | SDC1 | 1.5252 | 0.5354 | 1.0303 | 0.000999 |
| MacAngio | livertumor | FN1 | TFRC | 1.5252 | 0.2899 | 0.9075 | 0.000999 |
| MacAngio | livertumor | HBEGF | ERBB3 | 0.5316 | 0.2644 | 0.398 | 0.000999 |
| MacAngio | livertumor | HLA-A | ERBB3 | 2.328 | 0.2644 | 1.2962 | 0.000999 |
| MacAngio | livertumor | HLA-DRA | EPHB4 | 3.3453 | 0.4535 | 1.8994 | 0.000999 |
| MacAngio | livertumor | HLA-E | EPHB4 | 1.1729 | 0.4535 | 0.8132 | 0.000999 |
| MacAngio | livertumor | HLA-E | ITGA6 | 1.1729 | 0.5211 | 0.847 | 0.000999 |
| MacAngio | livertumor | HLA-E | PLXNB2 | 1.1729 | 0.265 | 0.7189 | 0.000999 |
| MacAngio | livertumor | HMGB2 | CDH1 | 0.386 | 0.4896 | 0.4378 | 0.000999 |
| MacAngio | livertumor | HMGB2 | DSG2 | 0.386 | 0.7479 | 0.5669 | 0.000999 |
| MacAngio | livertumor | HSP90B1 | ITGB1 | 0.8415 | 1.6044 | 1.223 | 0.000999 |
| MacAngio | livertumor | ITGB1 | CD151 | 0.5432 | 1.1085 | 0.8259 | 0.000999 |
| MacAngio | livertumor | ITGB1 | ITGA6 | 0.5432 | 0.5211 | 0.5322 | 0.000999 |
| MacAngio | livertumor | ITGB1 | RACK1 | 0.5432 | 2.7655 | 1.6544 | 0.000999 |
| MacAngio | livertumor | ITGB1 | SDC1 | 0.5432 | 0.5354 | 0.5393 | 0.000999 |
| MacAngio | livertumor | ITGB1 | VTN | 0.5432 | 0.7399 | 0.6416 | 0.000999 |
| MacAngio | livertumor | LGALS3 | ITGA2 | 1.4887 | 0.3983 | 0.9435 | 0.000999 |
| MacAngio | livertumor | LGALS3 | ITGB1 | 1.4887 | 1.6044 | 1.5466 | 0.000999 |
| MacAngio | livertumor | LGALS3 | ITGB4 | 1.4887 | 0.5518 | 1.0202 | 0.000999 |
| MacAngio | livertumor | NAMPT | INSR | 1.2286 | 0.3039 | 0.7662 | 0.000999 |
| MacAngio | livertumor | PTPRC | INSR | 0.4847 | 0.3039 | 0.3943 | 0.000999 |
| MacAngio | livertumor | PTPRC | MET | 0.4847 | 0.2983 | 0.3915 | 0.000999 |
| MacAngio | livertumor | RPS19 | RPSA | 3.3275 | 2.6317 | 2.9796 | 0.000999 |

| | | | | | | | |
|---|---|---|---|---|---|---|---|
| MacAngio | livertumor | SPP1 | ITGB1 | 2.927 | 1.6044 | 2.2657 | 0.000999 |
| MacAngio | livertumor | SPP1 | SDC1 | 2.927 | 0.5354 | 1.7312 | 0.000999 |
| MacAngio | livertumor | TGFB1 | ITGB1 | 0.2749 | 1.6044 | 0.9397 | 0.000999 |
| MacAngio | livertumor | TGFB1 | VTN | 0.2749 | 0.7399 | 0.5074 | 0.000999 |
| MacAngio | livertumor | THBS1 | ITGA3 | 0.4532 | 0.3913 | 0.4223 | 0.000999 |
| MacAngio | livertumor | THBS1 | ITGB1 | 0.4532 | 1.6044 | 1.0288 | 0.000999 |
| MacAngio | livertumor | THBS1 | SDC1 | 0.4532 | 0.5354 | 0.4943 | 0.000999 |
| MacAngio | livertumor | TNFSF13B | ITGB1 | 0.2932 | 1.6044 | 0.9488 | 0.000999 |
| MacAngio | livertumor | VEGFA | AXL | 0.6875 | 0.3099 | 0.4987 | 0.000999 |
| MacAngio | livertumor | VEGFA | VTN | 0.6875 | 0.7399 | 0.7137 | 0.000999 |
| MacAngio | livertumor | THBS1 | CD47 | 0.4532 | 0.5289 | 0.4911 | 0.002 |
| MacAngio | lungtumor | APOE | SDC1 | 1.2713 | 0.2715 | 0.7714 | 0.000999 |
| MacAngio | lungtumor | APOE | SDC4 | 1.2713 | 0.3516 | 0.8115 | 0.000999 |
| MacAngio | lungtumor | CALM1 | RPSA | 1.3231 | 1.6057 | 1.4644 | 0.000999 |
| MacAngio | lungtumor | CCL20 | CXCR4 | 0.7166 | 0.2684 | 0.4925 | 0.000999 |
| MacAngio | lungtumor | COL1A1 | SDC1 | 0.2972 | 0.2715 | 0.2844 | 0.000999 |
| MacAngio | lungtumor | CXCL16 | CXCR4 | 0.6563 | 0.2684 | 0.4623 | 0.000999 |
| MacAngio | lungtumor | CXCL2 | CXCR4 | 0.8413 | 0.2684 | 0.5549 | 0.000999 |
| MacAngio | lungtumor | CXCL3 | CXCR4 | 0.8491 | 0.2684 | 0.5588 | 0.000999 |
| MacAngio | lungtumor | FN1 | CD44 | 1.5252 | 0.3802 | 0.9527 | 0.000999 |
| MacAngio | lungtumor | FN1 | RPSA | 1.5252 | 1.6057 | 1.5655 | 0.000999 |
| MacAngio | lungtumor | FN1 | SDC1 | 1.5252 | 0.2715 | 0.8983 | 0.000999 |
| MacAngio | lungtumor | HBEGF | CD44 | 0.5316 | 0.3802 | 0.4559 | 0.000999 |
| MacAngio | lungtumor | ITGB1 | CD151 | 0.5432 | 0.6776 | 0.6104 | 0.000999 |
| MacAngio | lungtumor | ITGB1 | CD46 | 0.5432 | 0.434 | 0.4886 | 0.000999 |
| MacAngio | lungtumor | ITGB1 | CD47 | 0.5432 | 0.3255 | 0.4344 | 0.000999 |
| MacAngio | lungtumor | ITGB1 | RACK1 | 0.5432 | 2.2108 | 1.377 | 0.000999 |
| MacAngio | lungtumor | ITGB1 | SDC1 | 0.5432 | 0.2715 | 0.4074 | 0.000999 |
| MacAngio | lungtumor | PTPRC | CXCR4 | 0.4847 | 0.2684 | 0.3765 | 0.000999 |
| MacAngio | lungtumor | RTN4 | CD44 | 0.9475 | 0.3802 | 0.6639 | 0.000999 |

| | | | | | | | |
|---|---|---|---|---|---|---|---|
| MacAngio | lungtumor | RTN4 | CD99 | 0.9475 | 0.5938 | 0.7707 | 0.000999 |
| MacAngio | lungtumor | SPP1 | CD44 | 2.927 | 0.3802 | 1.6536 | 0.000999 |
| MacAngio | lungtumor | SPP1 | SDC1 | 2.927 | 0.2715 | 1.5992 | 0.000999 |
| MacAngio | lungtumor | THBS1 | CD47 | 0.4532 | 0.3255 | 0.3894 | 0.000999 |
| MacAngio | lungtumor | THBS1 | SDC1 | 0.4532 | 0.2715 | 0.3624 | 0.000999 |
| MacAngio | lungtumor | PTPRC | CD44 | 0.4847 | 0.3802 | 0.4324 | 0.002 |
| MacAngio | melanomatumo | ANXA1 | EDNRB | 1.3105 | 0.5666 | 0.9386 | 0.000999 |
| MacAngio | melanomatumo | CALM1 | RPSA | 1.3231 | 3.7787 | 2.5509 | 0.000999 |
| MacAngio | melanomatumo | FN1 | RPSA | 1.5252 | 3.7787 | 2.6519 | 0.000999 |
| MacAngio | melanomatumo | ITGB1 | RACK1 | 0.5432 | 4.145 | 2.3441 | 0.000999 |
| MacAngio | melanomatumo | NAMPT | ANXA6 | 1.2286 | 0.3625 | 0.7955 | 0.000999 |
| MacAngio | melanomatumo | RPS19 | RPSA | 3.3275 | 3.7787 | 3.5531 | 0.000999 |
| MacAngio | melanomatumo | SPP1 | NCSTN | 2.927 | 0.2614 | 1.5942 | 0.000999 |
| MacAngio | melanomatumo | MIF | TNFRSF14 | 1.1833 | 0.5069 | 0.8451 | 0.002 |
| MacAngio | ovarytumor | ADAM17 | ITGB1 | 0.2751 | 0.2783 | 0.2767 | 0.000999 |
| MacAngio | ovarytumor | ANXA1 | CXCR4 | 1.3105 | 0.3688 | 0.8396 | 0.000999 |
| MacAngio | ovarytumor | AREG | FOLR1 | 0.8518 | 0.5619 | 0.7069 | 0.000999 |
| MacAngio | ovarytumor | CALM1 | RPSA | 1.3231 | 2.9542 | 2.1387 | 0.000999 |
| MacAngio | ovarytumor | CCL20 | CXCR4 | 0.7166 | 0.3688 | 0.5427 | 0.000999 |
| MacAngio | ovarytumor | CXCL16 | CXCR4 | 0.6563 | 0.3688 | 0.5125 | 0.000999 |
| MacAngio | ovarytumor | CXCL2 | CXCR4 | 0.8413 | 0.3688 | 0.6051 | 0.000999 |
| MacAngio | ovarytumor | CXCL3 | CXCR4 | 0.8491 | 0.3688 | 0.609 | 0.000999 |
| MacAngio | ovarytumor | FN1 | ITGB1 | 1.5252 | 0.2783 | 0.9017 | 0.000999 |
| MacAngio | ovarytumor | FN1 | RPSA | 1.5252 | 2.9542 | 2.2397 | 0.000999 |
| MacAngio | ovarytumor | HSP90B1 | ITGB1 | 0.8415 | 0.2783 | 0.5599 | 0.000999 |
| MacAngio | ovarytumor | ITGB1 | CD151 | 0.5432 | 0.4981 | 0.5207 | 0.000999 |
| MacAngio | ovarytumor | ITGB1 | CD47 | 0.5432 | 0.4931 | 0.5181 | 0.000999 |
| MacAngio | ovarytumor | ITGB1 | RACK1 | 0.5432 | 2.9543 | 1.7487 | 0.000999 |
| MacAngio | ovarytumor | LGALS3 | ITGB1 | 1.4887 | 0.2783 | 0.8835 | 0.000999 |
| MacAngio | ovarytumor | PTPRC | CXCR4 | 0.4847 | 0.3688 | 0.4267 | 0.000999 |

| | | | | | | | |
|---|---|---|---|---|---|---|---|
| MacAngio | ovarytumor | SPP1 | ITGB1 | 2.927 | 0.2783 | 1.6026 | 0.000999 |
| MacAngio | ovarytumor | TGFB1 | ITGB1 | 0.2749 | 0.2783 | 0.2766 | 0.000999 |
| MacAngio | ovarytumor | THBS1 | CD47 | 0.4532 | 0.4931 | 0.4732 | 0.000999 |
| MacAngio | ovarytumor | THBS1 | ITGB1 | 0.4532 | 0.2783 | 0.3658 | 0.000999 |
| MacAngio | ovarytumor | TNFSF13B | ITGB1 | 0.2932 | 0.2783 | 0.2857 | 0.000999 |
| breasttumor | MacAngio | A2M | LRP1 | 0.3129 | 0.3563 | 0.3346 | 0.000999 |
| breasttumor | MacAngio | ANXA1 | CD44 | 0.4092 | 1.6454 | 1.0273 | 0.000999 |
| breasttumor | MacAngio | ANXA1 | CXCR4 | 0.4092 | 1.035 | 0.7221 | 0.000999 |
| breasttumor | MacAngio | APP | ALOX5 | 0.5409 | 0.3126 | 0.4268 | 0.000999 |
| breasttumor | MacAngio | APP | C3AR1 | 0.5409 | 0.3695 | 0.4552 | 0.000999 |
| breasttumor | MacAngio | APP | C5AR1 | 0.5409 | 0.6386 | 0.5898 | 0.000999 |
| breasttumor | MacAngio | APP | CCR1 | 0.5409 | 0.2788 | 0.4099 | 0.000999 |
| breasttumor | MacAngio | APP | CD36 | 0.5409 | 0.2588 | 0.3999 | 0.000999 |
| breasttumor | MacAngio | APP | CXCR4 | 0.5409 | 1.035 | 0.788 | 0.000999 |
| breasttumor | MacAngio | APP | FPR1 | 0.5409 | 0.5101 | 0.5255 | 0.000999 |
| breasttumor | MacAngio | APP | LRP1 | 0.5409 | 0.3563 | 0.4486 | 0.000999 |
| breasttumor | MacAngio | ARF1 | CD44 | 1.6219 | 1.6454 | 1.6336 | 0.000999 |
| breasttumor | MacAngio | CALM1 | TNFRSF1B | 0.9015 | 0.4892 | 0.6954 | 0.000999 |
| breasttumor | MacAngio | CALR | CD44 | 1.1835 | 1.6454 | 1.4144 | 0.000999 |
| breasttumor | MacAngio | CALR | ITGA5 | 1.1835 | 0.2748 | 0.7292 | 0.000999 |
| breasttumor | MacAngio | CALR | LRP1 | 1.1835 | 0.3563 | 0.7699 | 0.000999 |
| breasttumor | MacAngio | CD99 | CD99 | 0.3871 | 0.5897 | 0.4884 | 0.000999 |
| breasttumor | MacAngio | COL2A1 | CD44 | 0.3935 | 1.6454 | 1.0194 | 0.000999 |
| breasttumor | MacAngio | COL2A1 | ITGB1 | 0.3935 | 0.5432 | 0.4684 | 0.000999 |
| breasttumor | MacAngio | GSTP1 | CD44 | 2.5435 | 1.6454 | 2.0945 | 0.000999 |
| breasttumor | MacAngio | HEBP1 | C3AR1 | 0.4062 | 0.3695 | 0.3878 | 0.000999 |
| breasttumor | MacAngio | HEBP1 | C5AR1 | 0.4062 | 0.6386 | 0.5224 | 0.000999 |
| breasttumor | MacAngio | HEBP1 | CCR1 | 0.4062 | 0.2788 | 0.3425 | 0.000999 |
| breasttumor | MacAngio | HEBP1 | CXCR4 | 0.4062 | 1.035 | 0.7206 | 0.000999 |
| breasttumor | MacAngio | HEBP1 | FPR1 | 0.4062 | 0.5101 | 0.4581 | 0.000999 |

| | | | | | | | |
|---|---|---|---|---|---|---|---|
| breasttumor | MacAngio | HEBP1 | FPR3 | 0.4062 | 0.5502 | 0.4782 | 0.000999 |
| breasttumor | MacAngio | HMGB1 | AGTRAP | 1.1815 | 0.3017 | 0.7416 | 0.000999 |
| breasttumor | MacAngio | HMGB1 | LY96 | 1.1815 | 0.5477 | 0.8646 | 0.000999 |
| breasttumor | MacAngio | HMGB1 | TLR2 | 1.1815 | 0.3168 | 0.7491 | 0.000999 |
| breasttumor | MacAngio | HMGB2 | CD44 | 0.3139 | 1.6454 | 0.9796 | 0.000999 |
| breasttumor | MacAngio | HMGB2 | DAB2 | 0.3139 | 0.5886 | 0.4513 | 0.000999 |
| breasttumor | MacAngio | HSP90B1 | ITGB1 | 0.7786 | 0.5432 | 0.6609 | 0.000999 |
| breasttumor | MacAngio | HSP90B1 | LRP1 | 0.7786 | 0.3563 | 0.5674 | 0.000999 |
| breasttumor | MacAngio | HSP90B1 | TNFRSF1B | 0.7786 | 0.4892 | 0.6339 | 0.000999 |
| breasttumor | MacAngio | ITGB1 | CD47 | 0.2939 | 0.4767 | 0.3853 | 0.000999 |
| breasttumor | MacAngio | ITGB1 | DAB2 | 0.2939 | 0.5886 | 0.4413 | 0.000999 |
| breasttumor | MacAngio | ITGB1 | ENG | 0.2939 | 0.4053 | 0.3496 | 0.000999 |
| breasttumor | MacAngio | ITGB1 | ITGAX | 0.2939 | 0.4376 | 0.3657 | 0.000999 |
| breasttumor | MacAngio | ITGB1 | PLAUR | 0.2939 | 1.4686 | 0.8812 | 0.000999 |
| breasttumor | MacAngio | LGALS3 | FCGR2A | 0.8391 | 0.7152 | 0.7771 | 0.000999 |
| breasttumor | MacAngio | LRPAP1 | LRP1 | 0.2756 | 0.3563 | 0.3159 | 0.000999 |
| breasttumor | MacAngio | MDK | LRP1 | 0.592 | 0.3563 | 0.4742 | 0.000999 |
| breasttumor | MacAngio | MDK | SDC4 | 0.592 | 0.2596 | 0.4258 | 0.000999 |
| breasttumor | MacAngio | MIF | CD74 | 3.2716 | 3.2958 | 3.2837 | 0.000999 |
| breasttumor | MacAngio | NPW | C3AR1 | 0.5478 | 0.3695 | 0.4586 | 0.000999 |
| breasttumor | MacAngio | NPW | C5AR1 | 0.5478 | 0.6386 | 0.5932 | 0.000999 |
| breasttumor | MacAngio | NPW | CCR1 | 0.5478 | 0.2788 | 0.4133 | 0.000999 |
| breasttumor | MacAngio | NPW | CXCR4 | 0.5478 | 1.035 | 0.7914 | 0.000999 |
| breasttumor | MacAngio | NPW | FPR1 | 0.5478 | 0.5101 | 0.5289 | 0.000999 |
| breasttumor | MacAngio | RPS19 | C5AR1 | 3.691 | 0.6386 | 2.1648 | 0.000999 |
| breasttumor | MacAngio | RPS19 | ICAM1 | 3.691 | 0.2941 | 1.9926 | 0.000999 |
| breasttumor | MacAngio | RTN4 | CD44 | 0.3318 | 1.6454 | 0.9886 | 0.000999 |
| breasttumor | MacAngio | SERPINE2 | LRP1 | 0.6001 | 0.3563 | 0.4782 | 0.000999 |
| breasttumor | MacAngio | TF | TFRC | 0.2688 | 0.479 | 0.3739 | 0.000999 |
| colorectaltumor | MacAngio | APP | ALOX5 | 0.4966 | 0.3126 | 0.4046 | 0.000999 |

| | | | | | | | |
|---|---|---|---|---|---|---|---|
| colorectaltumor | MacAngio | APP | C3AR1 | 0.4966 | 0.3695 | 0.433 | 0.000999 |
| colorectaltumor | MacAngio | APP | C5AR1 | 0.4966 | 0.6386 | 0.5676 | 0.000999 |
| colorectaltumor | MacAngio | APP | CCR1 | 0.4966 | 0.2788 | 0.3877 | 0.000999 |
| colorectaltumor | MacAngio | APP | CD36 | 0.4966 | 0.2588 | 0.3777 | 0.000999 |
| colorectaltumor | MacAngio | APP | CXCR4 | 0.4966 | 1.035 | 0.7658 | 0.000999 |
| colorectaltumor | MacAngio | APP | FPR1 | 0.4966 | 0.5101 | 0.5033 | 0.000999 |
| colorectaltumor | MacAngio | APP | LRP1 | 0.4966 | 0.3563 | 0.4265 | 0.000999 |
| colorectaltumor | MacAngio | ARF1 | CD44 | 0.6489 | 1.6454 | 1.1471 | 0.000999 |
| colorectaltumor | MacAngio | COL1A1 | CD36 | 0.4875 | 0.2588 | 0.3732 | 0.000999 |
| colorectaltumor | MacAngio | COL1A1 | CD44 | 0.4875 | 1.6454 | 1.0664 | 0.000999 |
| colorectaltumor | MacAngio | COL1A1 | ITGA5 | 0.4875 | 0.2748 | 0.3812 | 0.000999 |
| colorectaltumor | MacAngio | COL1A1 | ITGB1 | 0.4875 | 0.5432 | 0.5154 | 0.000999 |
| colorectaltumor | MacAngio | GSTP1 | CD44 | 2.8039 | 1.6454 | 2.2246 | 0.000999 |
| colorectaltumor | MacAngio | HMGB1 | AGTRAP | 2.155 | 0.3017 | 1.2284 | 0.000999 |
| colorectaltumor | MacAngio | HMGB1 | LY96 | 2.155 | 0.5477 | 1.3513 | 0.000999 |
| colorectaltumor | MacAngio | HMGB1 | TLR2 | 2.155 | 0.3168 | 1.2359 | 0.000999 |
| colorectaltumor | MacAngio | HMGB2 | CD44 | 0.5347 | 1.6454 | 1.09 | 0.000999 |
| colorectaltumor | MacAngio | HMGB2 | DAB2 | 0.5347 | 0.5886 | 0.5617 | 0.000999 |
| colorectaltumor | MacAngio | ITGB1 | PLAUR | 0.4214 | 1.4686 | 0.945 | 0.000999 |
| colorectaltumor | MacAngio | MDK | LRP1 | 0.7548 | 0.3563 | 0.5556 | 0.000999 |
| colorectaltumor | MacAngio | MDK | SDC4 | 0.7548 | 0.2596 | 0.5072 | 0.000999 |
| colorectaltumor | MacAngio | MIF | CD74 | 1.6741 | 3.2958 | 2.4849 | 0.000999 |
| colorectaltumor | MacAngio | RPS19 | C5AR1 | 4.0956 | 0.6386 | 2.3671 | 0.000999 |
| colorectaltumor | MacAngio | RPS19 | ICAM1 | 4.0956 | 0.2941 | 2.1949 | 0.000999 |
| colorectaltumor | MacAngio | RPS19 | RPSA | 4.0956 | 1.8207 | 2.9581 | 0.000999 |
| colorectaltumor | MacAngio | LGALS3 | FCGR2A | 1.5563 | 0.7152 | 1.1357 | 0.003 |
| colorectaltumor | MacAngio | AIMP1 | RACK1 | 0.5265 | 2.0487 | 1.2876 | 0.00799 |
| livertumor | MacAngio | AGRN | LRP1 | 0.3492 | 0.3563 | 0.3528 | 0.000999 |
| livertumor | MacAngio | AGT | ANPEP | 0.4701 | 0.5585 | 0.5143 | 0.000999 |
| livertumor | MacAngio | AGT | ATP6AP2 | 0.4701 | 0.5 | 0.485 | 0.000999 |

| | | | | | | | |
|---|---|---|---|---|---|---|---|
| livertumor | MacAngio | AGT | C3AR1 | 0.4701 | 0.3695 | 0.4198 | 0.000999 |
| livertumor | MacAngio | AGT | C5AR1 | 0.4701 | 0.6386 | 0.5543 | 0.000999 |
| livertumor | MacAngio | AGT | CCR1 | 0.4701 | 0.2788 | 0.3744 | 0.000999 |
| livertumor | MacAngio | AGT | CXCR4 | 0.4701 | 1.035 | 0.7525 | 0.000999 |
| livertumor | MacAngio | AGT | FPR1 | 0.4701 | 0.5101 | 0.4901 | 0.000999 |
| livertumor | MacAngio | APOA1 | ABCA1 | 0.2888 | 0.2716 | 0.2802 | 0.000999 |
| livertumor | MacAngio | APOA1 | CD36 | 0.2888 | 0.2588 | 0.2738 | 0.000999 |
| livertumor | MacAngio | APOA1 | LRP1 | 0.2888 | 0.3563 | 0.3225 | 0.000999 |
| livertumor | MacAngio | APOA1 | MSR1 | 0.2888 | 0.4416 | 0.3652 | 0.000999 |
| livertumor | MacAngio | APOA1 | SDC4 | 0.2888 | 0.2596 | 0.2742 | 0.000999 |
| livertumor | MacAngio | APP | ALOX5 | 0.5626 | 0.3126 | 0.4376 | 0.000999 |
| livertumor | MacAngio | APP | C3AR1 | 0.5626 | 0.3695 | 0.466 | 0.000999 |
| livertumor | MacAngio | APP | C5AR1 | 0.5626 | 0.6386 | 0.6006 | 0.000999 |
| livertumor | MacAngio | APP | CCR1 | 0.5626 | 0.2788 | 0.4207 | 0.000999 |
| livertumor | MacAngio | APP | CD36 | 0.5626 | 0.2588 | 0.4107 | 0.000999 |
| livertumor | MacAngio | APP | CXCR4 | 0.5626 | 1.035 | 0.7988 | 0.000999 |
| livertumor | MacAngio | APP | FPR1 | 0.5626 | 0.5101 | 0.5363 | 0.000999 |
| livertumor | MacAngio | APP | LRP1 | 0.5626 | 0.3563 | 0.4594 | 0.000999 |
| livertumor | MacAngio | ARF1 | CD44 | 0.6936 | 1.6454 | 1.1695 | 0.000999 |
| livertumor | MacAngio | C3 | C3AR1 | 0.6584 | 0.3695 | 0.5139 | 0.000999 |
| livertumor | MacAngio | C3 | C5AR1 | 0.6584 | 0.6386 | 0.6485 | 0.000999 |
| livertumor | MacAngio | C3 | CCR1 | 0.6584 | 0.2788 | 0.4686 | 0.000999 |
| livertumor | MacAngio | C3 | CXCR4 | 0.6584 | 1.035 | 0.8467 | 0.000999 |
| livertumor | MacAngio | C3 | FPR1 | 0.6584 | 0.5101 | 0.5843 | 0.000999 |
| livertumor | MacAngio | C3 | ITGAM | 0.6584 | 0.3191 | 0.4888 | 0.000999 |
| livertumor | MacAngio | C3 | ITGAX | 0.6584 | 0.4376 | 0.548 | 0.000999 |
| livertumor | MacAngio | C3 | ITGB2 | 0.6584 | 0.9546 | 0.8065 | 0.000999 |
| livertumor | MacAngio | C3 | LRP1 | 0.6584 | 0.3563 | 0.5074 | 0.000999 |
| livertumor | MacAngio | CALR | CD44 | 1.0416 | 1.6454 | 1.3435 | 0.000999 |
| livertumor | MacAngio | CCL20 | C3AR1 | 1.0877 | 0.3695 | 0.7286 | 0.000999 |

| | | | | | | | |
|---|---|---|---|---|---|---|---|
| livertumor | MacAngio | CCL20 | C5AR1 | 1.0877 | 0.6386 | 0.8631 | 0.000999 |
| livertumor | MacAngio | CCL20 | CCR1 | 1.0877 | 0.2788 | 0.6832 | 0.000999 |
| livertumor | MacAngio | CCL20 | CXCR4 | 1.0877 | 1.035 | 1.0613 | 0.000999 |
| livertumor | MacAngio | CCL20 | FPR1 | 1.0877 | 0.5101 | 0.7989 | 0.000999 |
| livertumor | MacAngio | CDH1 | RACK1 | 0.4896 | 2.0487 | 1.2692 | 0.000999 |
| livertumor | MacAngio | CFH | ITGAM | 0.3373 | 0.3191 | 0.3282 | 0.000999 |
| livertumor | MacAngio | DSG2 | CD44 | 0.7479 | 1.6454 | 1.1966 | 0.000999 |
| livertumor | MacAngio | DSG2 | CXCR4 | 0.7479 | 1.035 | 0.8914 | 0.000999 |
| livertumor | MacAngio | DSG2 | DAB2 | 0.7479 | 0.5886 | 0.6683 | 0.000999 |
| livertumor | MacAngio | FGA | ITGAX | 0.2688 | 0.4376 | 0.3532 | 0.000999 |
| livertumor | MacAngio | FGA | ITGB2 | 0.2688 | 0.9546 | 0.6117 | 0.000999 |
| livertumor | MacAngio | GSTP1 | CD44 | 1.7219 | 1.6454 | 1.6836 | 0.000999 |
| livertumor | MacAngio | HMGB1 | AGTRAP | 1.4361 | 0.3017 | 0.8689 | 0.000999 |
| livertumor | MacAngio | HMGB1 | LY96 | 1.4361 | 0.5477 | 0.9919 | 0.000999 |
| livertumor | MacAngio | HMGB1 | TLR2 | 1.4361 | 0.3168 | 0.8764 | 0.000999 |
| livertumor | MacAngio | HSPG2 | LRP1 | 0.2746 | 0.3563 | 0.3155 | 0.000999 |
| livertumor | MacAngio | ITGB1 | CD151 | 1.6044 | 0.3149 | 0.9597 | 0.000999 |
| livertumor | MacAngio | ITGB1 | CD36 | 1.6044 | 0.2588 | 0.9316 | 0.000999 |
| livertumor | MacAngio | ITGB1 | CD47 | 1.6044 | 0.4767 | 1.0406 | 0.000999 |
| livertumor | MacAngio | ITGB1 | DAB2 | 1.6044 | 0.5886 | 1.0965 | 0.000999 |
| livertumor | MacAngio | ITGB1 | ENG | 1.6044 | 0.4053 | 1.0049 | 0.000999 |
| livertumor | MacAngio | ITGB1 | ITGA5 | 1.6044 | 0.2748 | 0.9396 | 0.000999 |
| livertumor | MacAngio | ITGB1 | ITGAX | 1.6044 | 0.4376 | 1.021 | 0.000999 |
| livertumor | MacAngio | ITGB1 | PLAUR | 1.6044 | 1.4686 | 1.5365 | 0.000999 |
| livertumor | MacAngio | ITGB1 | RACK1 | 1.6044 | 2.0487 | 1.8266 | 0.000999 |
| livertumor | MacAngio | LAMC2 | CD151 | 0.4847 | 0.3149 | 0.3998 | 0.000999 |
| livertumor | MacAngio | LAMC2 | ITGB1 | 0.4847 | 0.5432 | 0.514 | 0.000999 |
| livertumor | MacAngio | MDK | LRP1 | 1.6101 | 0.3563 | 0.9832 | 0.000999 |
| livertumor | MacAngio | MDK | SDC4 | 1.6101 | 0.2596 | 0.9349 | 0.000999 |
| livertumor | MacAngio | MIF | CD74 | 1.7638 | 3.2958 | 2.5298 | 0.000999 |

| | | | | | | | |
|---|---|---|---|---|---|---|---|
| livertumor | MacAngio | PTPRF | RACK1 | 0.384 | 2.0487 | 1.2164 | 0.000999 |
| livertumor | MacAngio | RPS19 | C5AR1 | 4.0772 | 0.6386 | 2.3579 | 0.000999 |
| livertumor | MacAngio | RPS19 | ICAM1 | 4.0772 | 0.2941 | 2.1857 | 0.000999 |
| livertumor | MacAngio | RPS19 | RPSA | 4.0772 | 1.8207 | 2.9489 | 0.000999 |
| livertumor | MacAngio | RTN4 | CD44 | 1.0046 | 1.6454 | 1.325 | 0.000999 |
| livertumor | MacAngio | SERPINA1 | LRP1 | 1.604 | 0.3563 | 0.9802 | 0.000999 |
| livertumor | MacAngio | SERPINA1 | SDC2 | 1.604 | 0.3836 | 0.9938 | 0.000999 |
| livertumor | MacAngio | SERPING1 | LRP1 | 0.3498 | 0.3563 | 0.353 | 0.000999 |
| livertumor | MacAngio | TF | SDC2 | 0.523 | 0.3836 | 0.4533 | 0.000999 |
| livertumor | MacAngio | TF | TFRC | 0.523 | 0.479 | 0.501 | 0.000999 |
| livertumor | MacAngio | TFPI | LRP1 | 0.5043 | 0.3563 | 0.4303 | 0.000999 |
| livertumor | MacAngio | VTN | ITGA5 | 0.7399 | 0.2748 | 0.5074 | 0.000999 |
| livertumor | MacAngio | VTN | ITGB1 | 0.7399 | 0.5432 | 0.6416 | 0.000999 |
| livertumor | MacAngio | VTN | PLAUR | 0.7399 | 1.4686 | 1.1043 | 0.000999 |
| livertumor | MacAngio | VTN | SDC2 | 0.7399 | 0.3836 | 0.5617 | 0.000999 |
| livertumor | MacAngio | VTN | SDC4 | 0.7399 | 0.2596 | 0.4998 | 0.000999 |
| livertumor | MacAngio | HSP90B1 | TNFRSF1B | 0.8951 | 0.4892 | 0.6922 | 0.002 |
| livertumor | MacAngio | HSP90B1 | LRP1 | 0.8951 | 0.3563 | 0.6257 | 0.00999 |
| lungtumor | MacAngio | ANXA1 | C3AR1 | 1.4026 | 0.3695 | 0.886 | 0.000999 |
| lungtumor | MacAngio | ANXA1 | C5AR1 | 1.4026 | 0.6386 | 1.0206 | 0.000999 |
| lungtumor | MacAngio | ANXA1 | CCR1 | 1.4026 | 0.2788 | 0.8407 | 0.000999 |
| lungtumor | MacAngio | ANXA1 | CD44 | 1.4026 | 1.6454 | 1.524 | 0.000999 |
| lungtumor | MacAngio | ANXA1 | CXCR4 | 1.4026 | 1.035 | 1.2188 | 0.000999 |
| lungtumor | MacAngio | ANXA1 | FPR1 | 1.4026 | 0.5101 | 0.9563 | 0.000999 |
| lungtumor | MacAngio | ANXA1 | FPR3 | 1.4026 | 0.5502 | 0.9764 | 0.000999 |
| lungtumor | MacAngio | ARF1 | CD44 | 0.8791 | 1.6454 | 1.2622 | 0.000999 |
| lungtumor | MacAngio | CALM1 | TNFRSF1B | 0.8426 | 0.4892 | 0.6659 | 0.000999 |
| lungtumor | MacAngio | CALR | CD44 | 0.7795 | 1.6454 | 1.2125 | 0.000999 |
| lungtumor | MacAngio | CALR | ITGA5 | 0.7795 | 0.2748 | 0.5272 | 0.000999 |
| lungtumor | MacAngio | CALR | LRP1 | 0.7795 | 0.3563 | 0.5679 | 0.000999 |

| | | | | | | | |
|---|---|---|---|---|---|---|---|
| lungtumor | MacAngio | CCL20 | C3AR1 | 0.2885 | 0.3695 | 0.329 | 0.000999 |
| lungtumor | MacAngio | CCL20 | C5AR1 | 0.2885 | 0.6386 | 0.4636 | 0.000999 |
| lungtumor | MacAngio | CCL20 | CCR1 | 0.2885 | 0.2788 | 0.2837 | 0.000999 |
| lungtumor | MacAngio | CCL20 | CXCR4 | 0.2885 | 1.035 | 0.6618 | 0.000999 |
| lungtumor | MacAngio | CCL20 | FPR1 | 0.2885 | 0.5101 | 0.3993 | 0.000999 |
| lungtumor | MacAngio | CXCL10 | C3AR1 | 0.3417 | 0.3695 | 0.3556 | 0.000999 |
| lungtumor | MacAngio | CXCL10 | C5AR1 | 0.3417 | 0.6386 | 0.4901 | 0.000999 |
| lungtumor | MacAngio | CXCL10 | CCR1 | 0.3417 | 0.2788 | 0.3102 | 0.000999 |
| lungtumor | MacAngio | CXCL10 | CXCR4 | 0.3417 | 1.035 | 0.6883 | 0.000999 |
| lungtumor | MacAngio | CXCL10 | FPR1 | 0.3417 | 0.5101 | 0.4259 | 0.000999 |
| lungtumor | MacAngio | FGA | ITGAX | 0.4445 | 0.4376 | 0.4411 | 0.000999 |
| lungtumor | MacAngio | FGA | ITGB1 | 0.4445 | 0.5432 | 0.4939 | 0.000999 |
| lungtumor | MacAngio | FGA | ITGB2 | 0.4445 | 0.9546 | 0.6995 | 0.000999 |
| lungtumor | MacAngio | GRN | TNFRSF1B | 0.4026 | 0.4892 | 0.4459 | 0.000999 |
| lungtumor | MacAngio | GSTP1 | CD44 | 2.4093 | 1.6454 | 2.0273 | 0.000999 |
| lungtumor | MacAngio | HMGB1 | AGTRAP | 1.1943 | 0.3017 | 0.748 | 0.000999 |
| lungtumor | MacAngio | HMGB1 | LY96 | 1.1943 | 0.5477 | 0.871 | 0.000999 |
| lungtumor | MacAngio | HMGB1 | TLR2 | 1.1943 | 0.3168 | 0.7555 | 0.000999 |
| lungtumor | MacAngio | HMGB2 | CD44 | 0.5593 | 1.6454 | 1.1023 | 0.000999 |
| lungtumor | MacAngio | HMGB2 | DAB2 | 0.5593 | 0.5886 | 0.574 | 0.000999 |
| lungtumor | MacAngio | HSP90B1 | ITGB1 | 0.6141 | 0.5432 | 0.5787 | 0.000999 |
| lungtumor | MacAngio | HSP90B1 | LRP1 | 0.6141 | 0.3563 | 0.4852 | 0.000999 |
| lungtumor | MacAngio | HSP90B1 | TNFRSF1B | 0.6141 | 0.4892 | 0.5517 | 0.000999 |
| lungtumor | MacAngio | LGALS3 | CLEC7A | 0.9915 | 0.5041 | 0.7478 | 0.000999 |
| lungtumor | MacAngio | LGALS3 | FCGR2A | 0.9915 | 0.7152 | 0.8534 | 0.000999 |
| lungtumor | MacAngio | LGALS3 | FCGR2B | 0.9915 | 0.4538 | 0.7227 | 0.000999 |
| lungtumor | MacAngio | LGALS3 | ITGB1 | 0.9915 | 0.5432 | 0.7674 | 0.000999 |
| lungtumor | MacAngio | LGALS3 | TFRC | 0.9915 | 0.479 | 0.7352 | 0.000999 |
| lungtumor | MacAngio | MDK | LRP1 | 1.2896 | 0.3563 | 0.823 | 0.000999 |
| lungtumor | MacAngio | MDK | SDC4 | 1.2896 | 0.2596 | 0.7746 | 0.000999 |

| | | | | | | | |
|---|---|---|---|---|---|---|---|
| lungtumor | MacAngio | MIF | CD74 | 1.4585 | 3.2958 | 2.3772 | 0.000999 |
| lungtumor | MacAngio | RPS19 | C5AR1 | 3.5104 | 0.6386 | 2.0745 | 0.000999 |
| lungtumor | MacAngio | RPS19 | ICAM1 | 3.5104 | 0.2941 | 1.9023 | 0.000999 |
| lungtumor | MacAngio | RPS19 | RPSA | 3.5104 | 1.8207 | 2.6656 | 0.000999 |
| lungtumor | MacAngio | RTN4 | CD44 | 0.4013 | 1.6454 | 1.0233 | 0.000999 |
| lungtumor | MacAngio | SAA1 | C3AR1 | 0.2539 | 0.3695 | 0.3117 | 0.000999 |
| lungtumor | MacAngio | SAA1 | C5AR1 | 0.2539 | 0.6386 | 0.4462 | 0.000999 |
| lungtumor | MacAngio | SAA1 | CCR1 | 0.2539 | 0.2788 | 0.2663 | 0.000999 |
| lungtumor | MacAngio | SAA1 | CXCR4 | 0.2539 | 1.035 | 0.6444 | 0.000999 |
| lungtumor | MacAngio | SAA1 | FPR1 | 0.2539 | 0.5101 | 0.382 | 0.000999 |
| lungtumor | MacAngio | SERPINA1 | LRP1 | 0.3624 | 0.3563 | 0.3593 | 0.000999 |
| lungtumor | MacAngio | SPP1 | CD44 | 1.4874 | 1.6454 | 1.5664 | 0.000999 |
| lungtumor | MacAngio | TFPI | LRP1 | 0.3615 | 0.3563 | 0.3589 | 0.000999 |
| lungtumor | MacAngio | SERPINA1 | SDC2 | 0.3624 | 0.3836 | 0.373 | 0.002 |
| melanomatumo | MacAngio | ARF1 | CD44 | 1.8545 | 1.6454 | 1.7499 | 0.000999 |
| melanomatumo | MacAngio | GSTP1 | CD44 | 2.3341 | 1.6454 | 1.9897 | 0.000999 |
| melanomatumo | MacAngio | HAS2 | CD44 | 0.3078 | 1.6454 | 0.9766 | 0.000999 |
| melanomatumo | MacAngio | HEBP1 | C3AR1 | 0.3689 | 0.3695 | 0.3692 | 0.000999 |
| melanomatumo | MacAngio | HEBP1 | C5AR1 | 0.3689 | 0.6386 | 0.5037 | 0.000999 |
| melanomatumo | MacAngio | HEBP1 | CCR1 | 0.3689 | 0.2788 | 0.3238 | 0.000999 |
| melanomatumo | MacAngio | HEBP1 | CXCR4 | 0.3689 | 1.035 | 0.7019 | 0.000999 |
| melanomatumo | MacAngio | HEBP1 | FPR1 | 0.3689 | 0.5101 | 0.4395 | 0.000999 |
| melanomatumo | MacAngio | HEBP1 | FPR3 | 0.3689 | 0.5502 | 0.4595 | 0.000999 |
| melanomatumo | MacAngio | HMGB1 | AGTRAP | 1.6757 | 0.3017 | 0.9887 | 0.000999 |
| melanomatumo | MacAngio | HMGB1 | LY96 | 1.6757 | 0.5477 | 1.1117 | 0.000999 |
| melanomatumo | MacAngio | HMGB1 | TLR2 | 1.6757 | 0.3168 | 0.9962 | 0.000999 |
| melanomatumo | MacAngio | MIF | CD74 | 2.6939 | 3.2958 | 2.9949 | 0.000999 |
| melanomatumo | MacAngio | SERPINE2 | LRP1 | 0.9323 | 0.3563 | 0.6443 | 0.000999 |
| ovarytumor | MacAngio | ANXA1 | C3AR1 | 1.0314 | 0.3695 | 0.7004 | 0.000999 |
| ovarytumor | MacAngio | ANXA1 | C5AR1 | 1.0314 | 0.6386 | 0.835 | 0.000999 |

| | | | | | | | |
|---|---|---|---|---|---|---|---|
| ovarytumor | MacAngio | ANXA1 | CCR1 | 1.0314 | 0.2788 | 0.6551 | 0.000999 |
| ovarytumor | MacAngio | ANXA1 | CD44 | 1.0314 | 1.6454 | 1.3384 | 0.000999 |
| ovarytumor | MacAngio | ANXA1 | CXCR4 | 1.0314 | 1.035 | 1.0332 | 0.000999 |
| ovarytumor | MacAngio | ANXA1 | FPR1 | 1.0314 | 0.5101 | 0.7707 | 0.000999 |
| ovarytumor | MacAngio | ANXA1 | FPR3 | 1.0314 | 0.5502 | 0.7908 | 0.000999 |
| ovarytumor | MacAngio | APOE | LRP1 | 0.837 | 0.3563 | 0.5967 | 0.000999 |
| ovarytumor | MacAngio | APOE | SDC2 | 0.837 | 0.3836 | 0.6103 | 0.000999 |
| ovarytumor | MacAngio | APOE | SDC4 | 0.837 | 0.2596 | 0.5483 | 0.000999 |
| ovarytumor | MacAngio | APP | ALOX5 | 0.2907 | 0.3126 | 0.3017 | 0.000999 |
| ovarytumor | MacAngio | APP | C3AR1 | 0.2907 | 0.3695 | 0.3301 | 0.000999 |
| ovarytumor | MacAngio | APP | C5AR1 | 0.2907 | 0.6386 | 0.4647 | 0.000999 |
| ovarytumor | MacAngio | APP | CCR1 | 0.2907 | 0.2788 | 0.2847 | 0.000999 |
| ovarytumor | MacAngio | APP | CD36 | 0.2907 | 0.2588 | 0.2748 | 0.000999 |
| ovarytumor | MacAngio | APP | CXCR4 | 0.2907 | 1.035 | 0.6628 | 0.000999 |
| ovarytumor | MacAngio | APP | FPR1 | 0.2907 | 0.5101 | 0.4004 | 0.000999 |
| ovarytumor | MacAngio | APP | LRP1 | 0.2907 | 0.3563 | 0.3235 | 0.000999 |
| ovarytumor | MacAngio | ARF1 | CD44 | 0.7364 | 1.6454 | 1.1909 | 0.000999 |
| ovarytumor | MacAngio | CALM1 | TNFRSF1B | 1.0087 | 0.4892 | 0.749 | 0.000999 |
| ovarytumor | MacAngio | CALR | CD44 | 0.7423 | 1.6454 | 1.1938 | 0.000999 |
| ovarytumor | MacAngio | CALR | ITGA5 | 0.7423 | 0.2748 | 0.5085 | 0.000999 |
| ovarytumor | MacAngio | CALR | LRP1 | 0.7423 | 0.3563 | 0.5493 | 0.000999 |
| ovarytumor | MacAngio | CCL2 | CCR1 | 0.3526 | 0.2788 | 0.3157 | 0.000999 |
| ovarytumor | MacAngio | COL1A1 | CD36 | 0.3504 | 0.2588 | 0.3046 | 0.000999 |
| ovarytumor | MacAngio | COL1A1 | CD44 | 0.3504 | 1.6454 | 0.9979 | 0.000999 |
| ovarytumor | MacAngio | COL1A1 | ITGA5 | 0.3504 | 0.2748 | 0.3126 | 0.000999 |
| ovarytumor | MacAngio | COL1A1 | ITGB1 | 0.3504 | 0.5432 | 0.4468 | 0.000999 |
| ovarytumor | MacAngio | GRN | TNFRSF1B | 0.3896 | 0.4892 | 0.4394 | 0.000999 |
| ovarytumor | MacAngio | GSTP1 | CD44 | 2.6185 | 1.6454 | 2.1319 | 0.000999 |
| ovarytumor | MacAngio | HMGB1 | AGTRAP | 1.7272 | 0.3017 | 1.0145 | 0.000999 |
| ovarytumor | MacAngio | HMGB1 | LY96 | 1.7272 | 0.5477 | 1.1374 | 0.000999 |

| | | | | | | | |
|---|---|---|---|---|---|---|---|
| ovarytumor | MacAngio | HMGB1 | TLR2 | 1.7272 | 0.3168 | 1.022 | 0.000999 |
| ovarytumor | MacAngio | HMGB2 | CD44 | 0.4974 | 1.6454 | 1.0714 | 0.000999 |
| ovarytumor | MacAngio | HMGB2 | DAB2 | 0.4974 | 0.5886 | 0.543 | 0.000999 |
| ovarytumor | MacAngio | HSP90B1 | ITGB1 | 0.6365 | 0.5432 | 0.5899 | 0.000999 |
| ovarytumor | MacAngio | HSP90B1 | LRP1 | 0.6365 | 0.3563 | 0.4964 | 0.000999 |
| ovarytumor | MacAngio | HSP90B1 | TNFRSF1B | 0.6365 | 0.4892 | 0.5629 | 0.000999 |
| ovarytumor | MacAngio | ITGB1 | CD36 | 0.2783 | 0.2588 | 0.2685 | 0.000999 |
| ovarytumor | MacAngio | ITGB1 | DAB2 | 0.2783 | 0.5886 | 0.4335 | 0.000999 |
| ovarytumor | MacAngio | ITGB1 | ENG | 0.2783 | 0.4053 | 0.3418 | 0.000999 |
| ovarytumor | MacAngio | ITGB1 | ITGA5 | 0.2783 | 0.2748 | 0.2765 | 0.000999 |
| ovarytumor | MacAngio | ITGB1 | ITGAX | 0.2783 | 0.4376 | 0.358 | 0.000999 |
| ovarytumor | MacAngio | ITGB1 | PLAUR | 0.2783 | 1.4686 | 0.8734 | 0.000999 |
| ovarytumor | MacAngio | LGALS3 | CLEC7A | 0.5097 | 0.5041 | 0.5069 | 0.000999 |
| ovarytumor | MacAngio | LGALS3 | FCGR2A | 0.5097 | 0.7152 | 0.6125 | 0.000999 |
| ovarytumor | MacAngio | LGALS3 | FCGR2B | 0.5097 | 0.4538 | 0.4818 | 0.000999 |
| ovarytumor | MacAngio | LGALS3 | TFRC | 0.5097 | 0.479 | 0.4943 | 0.000999 |
| ovarytumor | MacAngio | LRPAP1 | LRP1 | 0.3826 | 0.3563 | 0.3695 | 0.000999 |
| ovarytumor | MacAngio | MDK | LRP1 | 1.5321 | 0.3563 | 0.9442 | 0.000999 |
| ovarytumor | MacAngio | MDK | SDC4 | 1.5321 | 0.2596 | 0.8958 | 0.000999 |
| ovarytumor | MacAngio | MIF | CD74 | 0.8565 | 3.2958 | 2.0761 | 0.000999 |
| ovarytumor | MacAngio | RPS19 | C5AR1 | 3.9325 | 0.6386 | 2.2856 | 0.000999 |
| ovarytumor | MacAngio | RPS19 | ICAM1 | 3.9325 | 0.2941 | 2.1133 | 0.000999 |
| ovarytumor | MacAngio | RTN4 | CD44 | 0.4992 | 1.6454 | 1.0723 | 0.000999 |
| ovarytumor | MacAngio | RTN4 | CD99 | 0.4992 | 0.5897 | 0.5444 | 0.000999 |
| ovarytumor | MacAngio | SERPING1 | LRP1 | 0.3678 | 0.3563 | 0.3621 | 0.000999 |
| ovarytumor | MacAngio | SPP1 | CD44 | 0.3946 | 1.6454 | 1.02 | 0.000999 |
| ovarytumor | MacAngio | LGALS3 | ITGB1 | 0.5097 | 0.5432 | 0.5265 | 0.00599 |

| Supplementary Table 8: Communication results between macrophages and lymphocytes subpopulations, and malignant cells | | | | | | | |
|---|---|---|---|---|---|---|---|
| **MacHypo vs. lymphocytes** | | | | | | | |
| celltype_01 | celltype_02 | ligand | receptor | ligand_mean_expression | receptor_mean_expression | ligand_receptor_mean_expression | pvalue |
| MacHypo | NKT | A2M | TNFRSF14 | 0.3313 | 0.2848 | 0.3081 | 0.000999 |
| MacHypo | NKT | ADM | PTGER2 | 0.9374 | 0.3627 | 0.65 | 0.000999 |
| MacHypo | NKT | AIMP1 | RACK1 | 0.3406 | 2.4327 | 1.3867 | 0.000999 |
| MacHypo | NKT | ALCAM | CD6 | 0.2569 | 0.4881 | 0.3725 | 0.000999 |
| MacHypo | NKT | ANXA1 | CD44 | 1.6214 | 0.6724 | 1.1469 | 0.000999 |
| MacHypo | NKT | ANXA1 | CXCR4 | 1.6214 | 2.0373 | 1.8294 | 0.000999 |
| MacHypo | NKT | ANXA1 | F2R | 1.6214 | 0.2978 | 0.9596 | 0.000999 |
| MacHypo | NKT | ANXA1 | ITGA4 | 1.6214 | 0.2946 | 0.958 | 0.000999 |
| MacHypo | NKT | APOE | SORL1 | 2.2472 | 0.4137 | 1.3304 | 0.000999 |
| MacHypo | NKT | APP | CXCR4 | 0.2783 | 2.0373 | 1.1578 | 0.000999 |
| MacHypo | NKT | APP | F2R | 0.2783 | 0.2978 | 0.2881 | 0.000999 |
| MacHypo | NKT | APP | SORL1 | 0.2783 | 0.4137 | 0.346 | 0.000999 |
| MacHypo | NKT | APP | TNFRSF14 | 0.2783 | 0.2848 | 0.2816 | 0.000999 |
| MacHypo | NKT | CCL20 | CXCR4 | 0.6366 | 2.0373 | 1.337 | 0.000999 |
| MacHypo | NKT | CCL20 | S1PR4 | 0.6366 | 0.4435 | 0.5401 | 0.000999 |
| MacHypo | NKT | CCL20 | S1PR5 | 0.6366 | 0.508 | 0.5723 | 0.000999 |
| MacHypo | NKT | COL1A1 | CD44 | 0.3651 | 0.6724 | 0.5188 | 0.000999 |
| MacHypo | NKT | COL1A1 | ITGB1 | 0.3651 | 0.488 | 0.4265 | 0.000999 |
| MacHypo | NKT | CXCL16 | CXCR4 | 0.6914 | 2.0373 | 1.3643 | 0.000999 |
| MacHypo | NKT | CXCL16 | S1PR4 | 0.6914 | 0.4435 | 0.5674 | 0.000999 |
| MacHypo | NKT | CXCL16 | S1PR5 | 0.6914 | 0.508 | 0.5997 | 0.000999 |
| MacHypo | NKT | CXCL2 | CXCR4 | 0.6881 | 2.0373 | 1.3627 | 0.000999 |
| MacHypo | NKT | CXCL2 | S1PR4 | 0.6881 | 0.4435 | 0.5658 | 0.000999 |
| MacHypo | NKT | CXCL2 | S1PR5 | 0.6881 | 0.508 | 0.5981 | 0.000999 |
| MacHypo | NKT | CXCL3 | CXCR4 | 0.7216 | 2.0373 | 1.3794 | 0.000999 |
| MacHypo | NKT | CXCL3 | S1PR4 | 0.7216 | 0.4435 | 0.5826 | 0.000999 |
| MacHypo | NKT | CXCL3 | S1PR5 | 0.7216 | 0.508 | 0.6148 | 0.000999 |

| | | | | | | | |
|---|---|---|---|---|---|---|---|
| MacHypo | NKT | FN1 | CD44 | 1.342 | 0.6724 | 1.0072 | 0.000999 |
| MacHypo | NKT | FN1 | ITGA4 | 1.342 | 0.2946 | 0.8183 | 0.000999 |
| MacHypo | NKT | FN1 | ITGB1 | 1.342 | 0.488 | 0.915 | 0.000999 |
| MacHypo | NKT | FN1 | ITGB7 | 1.342 | 0.4721 | 0.9071 | 0.000999 |
| MacHypo | NKT | FN1 | RPSA | 1.342 | 2.3639 | 1.853 | 0.000999 |
| MacHypo | NKT | GRN | TNFRSF1B | 1.0708 | 0.4949 | 0.7829 | 0.000999 |
| MacHypo | NKT | GSTP1 | CD44 | 1.3428 | 0.6724 | 1.0076 | 0.000999 |
| MacHypo | NKT | GSTP1 | IL7R | 1.3428 | 0.5933 | 0.9681 | 0.000999 |
| MacHypo | NKT | GSTP1 | ITGA4 | 1.3428 | 0.2946 | 0.8187 | 0.000999 |
| MacHypo | NKT | HBEGF | CD44 | 0.2764 | 0.6724 | 0.4744 | 0.000999 |
| MacHypo | NKT | HLA-DRA | PTPRC | 3.2012 | 1.8797 | 2.5404 | 0.000999 |
| MacHypo | NKT | HSP90B1 | ITGB1 | 0.8573 | 0.488 | 0.6726 | 0.000999 |
| MacHypo | NKT | HSP90B1 | TNFRSF1B | 0.8573 | 0.4949 | 0.6761 | 0.000999 |
| MacHypo | NKT | ICAM1 | IL2RG | 0.3685 | 1.1524 | 0.7604 | 0.000999 |
| MacHypo | NKT | ICAM1 | ITGAL | 0.3685 | 0.6778 | 0.5232 | 0.000999 |
| MacHypo | NKT | ICAM1 | ITGB2 | 0.3685 | 1.6481 | 1.0083 | 0.000999 |
| MacHypo | NKT | ICAM1 | SPN | 0.3685 | 0.6398 | 0.5041 | 0.000999 |
| MacHypo | NKT | INHBA | TGFBR3 | 0.3596 | 0.4128 | 0.3862 | 0.000999 |
| MacHypo | NKT | LGALS3 | CD6 | 1.669 | 0.4881 | 1.0786 | 0.000999 |
| MacHypo | NKT | LGALS3 | ITGB1 | 1.669 | 0.488 | 1.0785 | 0.000999 |
| MacHypo | NKT | LGALS3 | SPN | 1.669 | 0.6398 | 1.1544 | 0.000999 |
| MacHypo | NKT | LRPAP1 | SORL1 | 0.4248 | 0.4137 | 0.4192 | 0.000999 |
| MacHypo | NKT | MIF | CD74 | 1.9478 | 1.6792 | 1.8135 | 0.000999 |
| MacHypo | NKT | MIF | ITGA4 | 1.9478 | 0.2946 | 1.1212 | 0.000999 |
| MacHypo | NKT | MIF | TNFRSF14 | 1.9478 | 0.2848 | 1.1163 | 0.000999 |
| MacHypo | NKT | MMP9 | CD44 | 0.8087 | 0.6724 | 0.7405 | 0.000999 |
| MacHypo | NKT | MMP9 | ITGB1 | 0.8087 | 0.488 | 0.6483 | 0.000999 |
| MacHypo | NKT | NAMPT | ANXA6 | 0.8899 | 0.844 | 0.8669 | 0.000999 |
| MacHypo | NKT | NMB | F2R | 0.356 | 0.2978 | 0.3269 | 0.000999 |
| MacHypo | NKT | NMB | GPR65 | 0.356 | 0.4898 | 0.4229 | 0.000999 |

| | | | | | | | |
|---|---|---|---|---|---|---|---|
| MacHypo | NKT | RTN4 | CD44 | 0.9167 | 0.6724 | 0.7946 | 0.000999 |
| MacHypo | NKT | RTN4 | CD99 | 0.9167 | 1.4547 | 1.1857 | 0.000999 |
| MacHypo | NKT | SPP1 | CD44 | 4.3573 | 0.6724 | 2.5149 | 0.000999 |
| MacHypo | NKT | SPP1 | ITGA4 | 4.3573 | 0.2946 | 2.3259 | 0.000999 |
| MacHypo | NKT | SPP1 | ITGB1 | 4.3573 | 0.488 | 2.4226 | 0.000999 |
| MacHypo | NKT | TNFSF13B | ITGB1 | 0.3932 | 0.488 | 0.4406 | 0.000999 |
| MacHypo | TCD4ex | A2M | TNFRSF14 | 0.3313 | 0.2806 | 0.306 | 0.000999 |
| MacHypo | TCD4ex | ADM | PTGER4 | 0.9374 | 0.4696 | 0.7035 | 0.000999 |
| MacHypo | TCD4ex | AIMP1 | RACK1 | 0.3406 | 2.4434 | 1.392 | 0.000999 |
| MacHypo | TCD4ex | ALCAM | CD6 | 0.2569 | 0.5338 | 0.3953 | 0.000999 |
| MacHypo | TCD4ex | ANXA1 | CCR7 | 1.6214 | 0.4844 | 1.0529 | 0.000999 |
| MacHypo | TCD4ex | ANXA1 | CD4 | 1.6214 | 0.5864 | 1.1039 | 0.000999 |
| MacHypo | TCD4ex | ANXA1 | CD44 | 1.6214 | 1.0244 | 1.3229 | 0.000999 |
| MacHypo | TCD4ex | ANXA1 | CXCR4 | 1.6214 | 2.4201 | 2.0207 | 0.000999 |
| MacHypo | TCD4ex | APP | CCR7 | 0.2783 | 0.4844 | 0.3814 | 0.000999 |
| MacHypo | TCD4ex | APP | CXCR4 | 0.2783 | 2.4201 | 1.3492 | 0.000999 |
| MacHypo | TCD4ex | APP | TNFRSF14 | 0.2783 | 0.2806 | 0.2795 | 0.000999 |
| MacHypo | TCD4ex | CALR | IL7R | 0.9194 | 1.2353 | 1.0773 | 0.000999 |
| MacHypo | TCD4ex | CCL20 | CCR7 | 0.6366 | 0.4844 | 0.5605 | 0.000999 |
| MacHypo | TCD4ex | CCL20 | CXCR4 | 0.6366 | 2.4201 | 1.5283 | 0.000999 |
| MacHypo | TCD4ex | CD86 | CD28 | 0.566 | 0.3256 | 0.4458 | 0.000999 |
| MacHypo | TCD4ex | CD86 | CTLA4 | 0.566 | 0.6294 | 0.5977 | 0.000999 |
| MacHypo | TCD4ex | COL1A1 | CD44 | 0.3651 | 1.0244 | 0.6948 | 0.000999 |
| MacHypo | TCD4ex | COL1A1 | ITGB1 | 0.3651 | 0.4615 | 0.4133 | 0.000999 |
| MacHypo | TCD4ex | CXCL16 | CCR7 | 0.6914 | 0.4844 | 0.5879 | 0.000999 |
| MacHypo | TCD4ex | CXCL16 | CXCR4 | 0.6914 | 2.4201 | 1.5557 | 0.000999 |
| MacHypo | TCD4ex | CXCL2 | CCR7 | 0.6881 | 0.4844 | 0.5863 | 0.000999 |
| MacHypo | TCD4ex | CXCL2 | CXCR4 | 0.6881 | 2.4201 | 1.5541 | 0.000999 |
| MacHypo | TCD4ex | CXCL3 | CCR7 | 0.7216 | 0.4844 | 0.603 | 0.000999 |
| MacHypo | TCD4ex | CXCL3 | CXCR4 | 0.7216 | 2.4201 | 1.5708 | 0.000999 |

| | | | | | | | |
|---|---|---|---|---|---|---|---|
| MacHypo | TCD4ex | FN1 | CD44 | 1.342 | 1.0244 | 1.1832 | 0.000999 |
| MacHypo | TCD4ex | FN1 | ITGB1 | 1.342 | 0.4615 | 0.9018 | 0.000999 |
| MacHypo | TCD4ex | FN1 | RPSA | 1.342 | 2.4895 | 1.9158 | 0.000999 |
| MacHypo | TCD4ex | GRN | TNFRSF1B | 1.0708 | 0.383 | 0.7269 | 0.000999 |
| MacHypo | TCD4ex | GSTP1 | CD44 | 1.3428 | 1.0244 | 1.1836 | 0.000999 |
| MacHypo | TCD4ex | GSTP1 | IL7R | 1.3428 | 1.2353 | 1.289 | 0.000999 |
| MacHypo | TCD4ex | HBEGF | CD44 | 0.2764 | 1.0244 | 0.6504 | 0.000999 |
| MacHypo | TCD4ex | HLA-DMA | CD4 | 0.8533 | 0.5864 | 0.7199 | 0.000999 |
| MacHypo | TCD4ex | HLA-DRA | CD4 | 3.2012 | 0.5864 | 1.8938 | 0.000999 |
| MacHypo | TCD4ex | HLA-DRA | PDCD1 | 3.2012 | 0.457 | 1.8291 | 0.000999 |
| MacHypo | TCD4ex | HLA-DRA | PTPRC | 3.2012 | 1.7493 | 2.4753 | 0.000999 |
| MacHypo | TCD4ex | HSP90B1 | ITGB1 | 0.8573 | 0.4615 | 0.6594 | 0.000999 |
| MacHypo | TCD4ex | HSP90B1 | TNFRSF1B | 0.8573 | 0.383 | 0.6202 | 0.000999 |
| MacHypo | TCD4ex | ICAM1 | IL2RG | 0.3685 | 0.9695 | 0.669 | 0.000999 |
| MacHypo | TCD4ex | ICAM1 | ITGAL | 0.3685 | 0.2603 | 0.3144 | 0.000999 |
| MacHypo | TCD4ex | ICAM1 | ITGB2 | 0.3685 | 0.7871 | 0.5778 | 0.000999 |
| MacHypo | TCD4ex | ITGB2 | ITGAL | 1.0357 | 0.2603 | 0.648 | 0.000999 |
| MacHypo | TCD4ex | LGALS3 | CD6 | 1.669 | 0.5338 | 1.1014 | 0.000999 |
| MacHypo | TCD4ex | LGALS3 | ITGB1 | 1.669 | 0.4615 | 1.0652 | 0.000999 |
| MacHypo | TCD4ex | MIF | CD74 | 1.9478 | 1.6094 | 1.7786 | 0.000999 |
| MacHypo | TCD4ex | MIF | TNFRSF14 | 1.9478 | 0.2806 | 1.1142 | 0.000999 |
| MacHypo | TCD4ex | MMP9 | CD44 | 0.8087 | 1.0244 | 0.9165 | 0.000999 |
| MacHypo | TCD4ex | MMP9 | ITGB1 | 0.8087 | 0.4615 | 0.6351 | 0.000999 |
| MacHypo | TCD4ex | NAMPT | ANXA6 | 0.8899 | 0.5386 | 0.7143 | 0.000999 |
| MacHypo | TCD4ex | RTN4 | CD44 | 0.9167 | 1.0244 | 0.9706 | 0.000999 |
| MacHypo | TCD4ex | RTN4 | CD99 | 0.9167 | 0.9165 | 0.9166 | 0.000999 |
| MacHypo | TCD4ex | SPP1 | CD44 | 4.3573 | 1.0244 | 2.6909 | 0.000999 |
| MacHypo | TCD4ex | SPP1 | ITGB1 | 4.3573 | 0.4615 | 2.4094 | 0.000999 |
| MacHypo | TCD4ex | TNFSF13B | ITGB1 | 0.3932 | 0.4615 | 0.4273 | 0.000999 |
| MacHypo | TCD4reg | A2M | TNFRSF14 | 0.3313 | 0.3843 | 0.3578 | 0.000999 |

| | | | | | | | |
|---|---|---|---|---|---|---|---|
| MacHypo | TCD4reg | AIMP1 | FAS | 0.3406 | 0.2727 | 0.3067 | 0.000999 |
| MacHypo | TCD4reg | AIMP1 | RACK1 | 0.3406 | 2.5298 | 1.4352 | 0.000999 |
| MacHypo | TCD4reg | ALCAM | CD6 | 0.2569 | 0.2651 | 0.261 | 0.000999 |
| MacHypo | TCD4reg | ANXA1 | CCR7 | 1.6214 | 0.3098 | 0.9656 | 0.000999 |
| MacHypo | TCD4reg | ANXA1 | CD4 | 1.6214 | 0.429 | 1.0252 | 0.000999 |
| MacHypo | TCD4reg | ANXA1 | CD44 | 1.6214 | 1.0896 | 1.3555 | 0.000999 |
| MacHypo | TCD4reg | ANXA1 | CXCR3 | 1.6214 | 0.3357 | 0.9785 | 0.000999 |
| MacHypo | TCD4reg | ANXA1 | CXCR4 | 1.6214 | 2.2962 | 1.9588 | 0.000999 |
| MacHypo | TCD4reg | ANXA1 | CXCR6 | 1.6214 | 0.3945 | 1.008 | 0.000999 |
| MacHypo | TCD4reg | ANXA1 | FAS | 1.6214 | 0.2727 | 0.9471 | 0.000999 |
| MacHypo | TCD4reg | APP | CCR7 | 0.2783 | 0.3098 | 0.2941 | 0.000999 |
| MacHypo | TCD4reg | APP | CXCR3 | 0.2783 | 0.3357 | 0.307 | 0.000999 |
| MacHypo | TCD4reg | APP | CXCR4 | 0.2783 | 2.2962 | 1.2873 | 0.000999 |
| MacHypo | TCD4reg | APP | CXCR6 | 0.2783 | 0.3945 | 0.3364 | 0.000999 |
| MacHypo | TCD4reg | APP | TNFRSF14 | 0.2783 | 0.3843 | 0.3313 | 0.000999 |
| MacHypo | TCD4reg | CALR | CD44 | 0.9194 | 1.0896 | 1.0045 | 0.000999 |
| MacHypo | TCD4reg | CALR | FAS | 0.9194 | 0.2727 | 0.5961 | 0.000999 |
| MacHypo | TCD4reg | CALR | IL7R | 0.9194 | 0.9682 | 0.9438 | 0.000999 |
| MacHypo | TCD4reg | CCL20 | CCR7 | 0.6366 | 0.3098 | 0.4732 | 0.000999 |
| MacHypo | TCD4reg | CCL20 | CXCR3 | 0.6366 | 0.3357 | 0.4862 | 0.000999 |
| MacHypo | TCD4reg | CCL20 | CXCR4 | 0.6366 | 2.2962 | 1.4664 | 0.000999 |
| MacHypo | TCD4reg | CCL20 | CXCR6 | 0.6366 | 0.3945 | 0.5156 | 0.000999 |
| MacHypo | TCD4reg | CCL20 | S1PR4 | 0.6366 | 0.2744 | 0.4555 | 0.000999 |
| MacHypo | TCD4reg | CD86 | CD28 | 0.566 | 0.313 | 0.4395 | 0.000999 |
| MacHypo | TCD4reg | CD86 | CTLA4 | 0.566 | 0.7909 | 0.6785 | 0.000999 |
| MacHypo | TCD4reg | COL1A1 | CD44 | 0.3651 | 1.0896 | 0.7274 | 0.000999 |
| MacHypo | TCD4reg | COL1A1 | ITGB1 | 0.3651 | 0.4001 | 0.3826 | 0.000999 |
| MacHypo | TCD4reg | CXCL16 | CCR7 | 0.6914 | 0.3098 | 0.5006 | 0.000999 |
| MacHypo | TCD4reg | CXCL16 | CXCR3 | 0.6914 | 0.3357 | 0.5135 | 0.000999 |
| MacHypo | TCD4reg | CXCL16 | CXCR4 | 0.6914 | 2.2962 | 1.4938 | 0.000999 |

| | | | | | | | |
|---|---|---|---|---|---|---|---|
| MacHypo | TCD4reg | CXCL16 | CXCR6 | 0.6914 | 0.3945 | 0.5429 | 0.000999 |
| MacHypo | TCD4reg | CXCL16 | S1PR4 | 0.6914 | 0.2744 | 0.4829 | 0.000999 |
| MacHypo | TCD4reg | CXCL2 | CCR7 | 0.6881 | 0.3098 | 0.499 | 0.000999 |
| MacHypo | TCD4reg | CXCL2 | CXCR3 | 0.6881 | 0.3357 | 0.5119 | 0.000999 |
| MacHypo | TCD4reg | CXCL2 | CXCR4 | 0.6881 | 2.2962 | 1.4921 | 0.000999 |
| MacHypo | TCD4reg | CXCL2 | CXCR6 | 0.6881 | 0.3945 | 0.5413 | 0.000999 |
| MacHypo | TCD4reg | CXCL2 | S1PR4 | 0.6881 | 0.2744 | 0.4813 | 0.000999 |
| MacHypo | TCD4reg | CXCL3 | CCR7 | 0.7216 | 0.3098 | 0.5157 | 0.000999 |
| MacHypo | TCD4reg | CXCL3 | CXCR3 | 0.7216 | 0.3357 | 0.5286 | 0.000999 |
| MacHypo | TCD4reg | CXCL3 | CXCR4 | 0.7216 | 2.2962 | 1.5089 | 0.000999 |
| MacHypo | TCD4reg | CXCL3 | CXCR6 | 0.7216 | 0.3945 | 0.5581 | 0.000999 |
| MacHypo | TCD4reg | CXCL3 | S1PR4 | 0.7216 | 0.2744 | 0.498 | 0.000999 |
| MacHypo | TCD4reg | FN1 | CD44 | 1.342 | 1.0896 | 1.2158 | 0.000999 |
| MacHypo | TCD4reg | FN1 | ITGB1 | 1.342 | 0.4001 | 0.8711 | 0.000999 |
| MacHypo | TCD4reg | FN1 | RPSA | 1.342 | 2.4999 | 1.921 | 0.000999 |
| MacHypo | TCD4reg | GRN | TNFRSF1B | 1.0708 | 0.7982 | 0.9345 | 0.000999 |
| MacHypo | TCD4reg | GSTP1 | CD44 | 1.3428 | 1.0896 | 1.2162 | 0.000999 |
| MacHypo | TCD4reg | GSTP1 | FAS | 1.3428 | 0.2727 | 0.8077 | 0.000999 |
| MacHypo | TCD4reg | GSTP1 | IL7R | 1.3428 | 0.9682 | 1.1555 | 0.000999 |
| MacHypo | TCD4reg | HBEGF | CD44 | 0.2764 | 1.0896 | 0.683 | 0.000999 |
| MacHypo | TCD4reg | HLA-DMA | CD4 | 0.8533 | 0.429 | 0.6411 | 0.000999 |
| MacHypo | TCD4reg | HLA-DRA | CD4 | 3.2012 | 0.429 | 1.8151 | 0.000999 |
| MacHypo | TCD4reg | HLA-DRA | PTPRC | 3.2012 | 1.6603 | 2.4308 | 0.000999 |
| MacHypo | TCD4reg | HSP90B1 | ITGB1 | 0.8573 | 0.4001 | 0.6287 | 0.000999 |
| MacHypo | TCD4reg | HSP90B1 | TNFRSF1B | 0.8573 | 0.7982 | 0.8277 | 0.000999 |
| MacHypo | TCD4reg | ICAM1 | IL2RA | 0.3685 | 0.7115 | 0.54 | 0.000999 |
| MacHypo | TCD4reg | ICAM1 | IL2RG | 0.3685 | 1.1601 | 0.7643 | 0.000999 |
| MacHypo | TCD4reg | ICAM1 | ITGB2 | 0.3685 | 0.6238 | 0.4962 | 0.000999 |
| MacHypo | TCD4reg | LGALS3 | CD6 | 1.669 | 0.2651 | 0.967 | 0.000999 |
| MacHypo | TCD4reg | LGALS3 | FAS | 1.669 | 0.2727 | 0.9708 | 0.000999 |

| | | | | | | | |
|---|---|---|---|---|---|---|---|
| MacHypo | TCD4reg | LGALS3 | ITGB1 | 1.669 | 0.4001 | 1.0345 | 0.000999 |
| MacHypo | TCD4reg | MIF | CD74 | 1.9478 | 2.0716 | 2.0097 | 0.000999 |
| MacHypo | TCD4reg | MIF | TNFRSF14 | 1.9478 | 0.3843 | 1.166 | 0.000999 |
| MacHypo | TCD4reg | MMP9 | CD44 | 0.8087 | 1.0896 | 0.9491 | 0.000999 |
| MacHypo | TCD4reg | MMP9 | ITGB1 | 0.8087 | 0.4001 | 0.6044 | 0.000999 |
| MacHypo | TCD4reg | NAMPT | ANXA6 | 0.8899 | 0.4557 | 0.6728 | 0.000999 |
| MacHypo | TCD4reg | NMB | P2RY10 | 0.356 | 0.2824 | 0.3192 | 0.000999 |
| MacHypo | TCD4reg | RTN4 | CD44 | 0.9167 | 1.0896 | 1.0032 | 0.000999 |
| MacHypo | TCD4reg | RTN4 | CD99 | 0.9167 | 0.6603 | 0.7885 | 0.000999 |
| MacHypo | TCD4reg | SPP1 | CD44 | 4.3573 | 1.0896 | 2.7235 | 0.000999 |
| MacHypo | TCD4reg | SPP1 | ITGB1 | 4.3573 | 0.4001 | 2.3787 | 0.000999 |
| MacHypo | TCD4reg | TNFSF13B | ITGB1 | 0.3932 | 0.4001 | 0.3966 | 0.000999 |
| MacHypo | TCD4reg | CD99 | CD99 | 0.765 | 0.6603 | 0.7127 | 0.004 |
| MacHypo | TCD4reg | ITGB1 | RACK1 | 0.4597 | 2.5298 | 1.4947 | 0.005 |
| MacHypo | TCD8em | A2M | TNFRSF14 | 0.3313 | 0.2915 | 0.3114 | 0.000999 |
| MacHypo | TCD8em | ADM | PTGER4 | 0.9374 | 0.2847 | 0.6111 | 0.000999 |
| MacHypo | TCD8em | AIMP1 | RACK1 | 0.3406 | 2.4374 | 1.389 | 0.000999 |
| MacHypo | TCD8em | ALCAM | CD6 | 0.2569 | 0.4204 | 0.3387 | 0.000999 |
| MacHypo | TCD8em | ANXA1 | CD44 | 1.6214 | 1.119 | 1.3702 | 0.000999 |
| MacHypo | TCD8em | ANXA1 | CXCR3 | 1.6214 | 0.4189 | 1.0202 | 0.000999 |
| MacHypo | TCD8em | ANXA1 | CXCR4 | 1.6214 | 2.8389 | 2.2302 | 0.000999 |
| MacHypo | TCD8em | ANXA1 | CXCR6 | 1.6214 | 0.3399 | 0.9806 | 0.000999 |
| MacHypo | TCD8em | ANXA1 | F2R | 1.6214 | 0.2695 | 0.9455 | 0.000999 |
| MacHypo | TCD8em | ANXA1 | ITGA4 | 1.6214 | 0.2823 | 0.9519 | 0.000999 |
| MacHypo | TCD8em | APP | CXCR3 | 0.2783 | 0.4189 | 0.3486 | 0.000999 |
| MacHypo | TCD8em | APP | CXCR4 | 0.2783 | 2.8389 | 1.5586 | 0.000999 |
| MacHypo | TCD8em | APP | CXCR6 | 0.2783 | 0.3399 | 0.3091 | 0.000999 |
| MacHypo | TCD8em | APP | F2R | 0.2783 | 0.2695 | 0.2739 | 0.000999 |
| MacHypo | TCD8em | APP | TNFRSF14 | 0.2783 | 0.2915 | 0.2849 | 0.000999 |
| MacHypo | TCD8em | CALR | CD44 | 0.9194 | 1.119 | 1.0192 | 0.000999 |

| | | | | | | | |
|---|---|---|---|---|---|---|---|
| MacHypo | TCD8em | CALR | IL7R | 0.9194 | 0.7183 | 0.8189 | 0.000999 |
| MacHypo | TCD8em | CCL20 | CXCR3 | 0.6366 | 0.4189 | 0.5278 | 0.000999 |
| MacHypo | TCD8em | CCL20 | CXCR4 | 0.6366 | 2.8389 | 1.7378 | 0.000999 |
| MacHypo | TCD8em | CCL20 | CXCR6 | 0.6366 | 0.3399 | 0.4882 | 0.000999 |
| MacHypo | TCD8em | CCL20 | S1PR4 | 0.6366 | 0.2677 | 0.4522 | 0.000999 |
| MacHypo | TCD8em | COL1A1 | ITGB1 | 0.3651 | 0.2557 | 0.3104 | 0.000999 |
| MacHypo | TCD8em | CXCL16 | CXCR3 | 0.6914 | 0.4189 | 0.5551 | 0.000999 |
| MacHypo | TCD8em | CXCL16 | CXCR4 | 0.6914 | 2.8389 | 1.7651 | 0.000999 |
| MacHypo | TCD8em | CXCL16 | CXCR6 | 0.6914 | 0.3399 | 0.5156 | 0.000999 |
| MacHypo | TCD8em | CXCL16 | S1PR4 | 0.6914 | 0.2677 | 0.4795 | 0.000999 |
| MacHypo | TCD8em | CXCL2 | CXCR3 | 0.6881 | 0.4189 | 0.5535 | 0.000999 |
| MacHypo | TCD8em | CXCL2 | CXCR4 | 0.6881 | 2.8389 | 1.7635 | 0.000999 |
| MacHypo | TCD8em | CXCL2 | CXCR6 | 0.6881 | 0.3399 | 0.514 | 0.000999 |
| MacHypo | TCD8em | CXCL2 | S1PR4 | 0.6881 | 0.2677 | 0.4779 | 0.000999 |
| MacHypo | TCD8em | CXCL3 | CXCR3 | 0.7216 | 0.4189 | 0.5702 | 0.000999 |
| MacHypo | TCD8em | CXCL3 | CXCR4 | 0.7216 | 2.8389 | 1.7803 | 0.000999 |
| MacHypo | TCD8em | CXCL3 | CXCR6 | 0.7216 | 0.3399 | 0.5307 | 0.000999 |
| MacHypo | TCD8em | CXCL3 | S1PR4 | 0.7216 | 0.2677 | 0.4947 | 0.000999 |
| MacHypo | TCD8em | FN1 | CD44 | 1.342 | 1.119 | 1.2305 | 0.000999 |
| MacHypo | TCD8em | FN1 | ITGA4 | 1.342 | 0.2823 | 0.8122 | 0.000999 |
| MacHypo | TCD8em | FN1 | ITGB1 | 1.342 | 0.2557 | 0.7989 | 0.000999 |
| MacHypo | TCD8em | FN1 | RPSA | 1.342 | 2.3686 | 1.8553 | 0.000999 |
| MacHypo | TCD8em | GRN | TNFRSF1B | 1.0708 | 0.3038 | 0.6873 | 0.000999 |
| MacHypo | TCD8em | GSTP1 | CD44 | 1.3428 | 1.119 | 1.2309 | 0.000999 |
| MacHypo | TCD8em | GSTP1 | IL7R | 1.3428 | 0.7183 | 1.0305 | 0.000999 |
| MacHypo | TCD8em | GSTP1 | ITGA4 | 1.3428 | 0.2823 | 0.8126 | 0.000999 |
| MacHypo | TCD8em | HBEGF | CD44 | 0.2764 | 1.119 | 0.6977 | 0.000999 |
| MacHypo | TCD8em | HLA-DRA | PTPRC | 3.2012 | 1.7912 | 2.4962 | 0.000999 |
| MacHypo | TCD8em | HSP90B1 | ITGB1 | 0.8573 | 0.2557 | 0.5565 | 0.000999 |
| MacHypo | TCD8em | HSP90B1 | TNFRSF1B | 0.8573 | 0.3038 | 0.5805 | 0.000999 |

| | | | | | | | |
|---|---|---|---|---|---|---|---|
| MacHypo | TCD8em | ICAM1 | IL2RG | 0.3685 | 0.9501 | 0.6593 | 0.000999 |
| MacHypo | TCD8em | ICAM1 | ITGAL | 0.3685 | 0.3162 | 0.3424 | 0.000999 |
| MacHypo | TCD8em | ICAM1 | ITGB2 | 0.3685 | 1.0018 | 0.6852 | 0.000999 |
| MacHypo | TCD8em | ICAM1 | SPN | 0.3685 | 0.2603 | 0.3144 | 0.000999 |
| MacHypo | TCD8em | ITGB1 | CD47 | 0.4597 | 0.4276 | 0.4436 | 0.000999 |
| MacHypo | TCD8em | ITGB1 | ITGA4 | 0.4597 | 0.2823 | 0.371 | 0.000999 |
| MacHypo | TCD8em | ITGB1 | RACK1 | 0.4597 | 2.4374 | 1.4485 | 0.000999 |
| MacHypo | TCD8em | LGALS3 | CD6 | 1.669 | 0.4204 | 1.0447 | 0.000999 |
| MacHypo | TCD8em | LGALS3 | ITGB1 | 1.669 | 0.2557 | 0.9624 | 0.000999 |
| MacHypo | TCD8em | LGALS3 | SPN | 1.669 | 0.2603 | 0.9647 | 0.000999 |
| MacHypo | TCD8em | MIF | CD74 | 1.9478 | 2.4538 | 2.2008 | 0.000999 |
| MacHypo | TCD8em | MIF | ITGA4 | 1.9478 | 0.2823 | 1.1151 | 0.000999 |
| MacHypo | TCD8em | MIF | TNFRSF14 | 1.9478 | 0.2915 | 1.1196 | 0.000999 |
| MacHypo | TCD8em | MMP9 | CD44 | 0.8087 | 1.119 | 0.9638 | 0.000999 |
| MacHypo | TCD8em | MMP9 | ITGB1 | 0.8087 | 0.2557 | 0.5322 | 0.000999 |
| MacHypo | TCD8em | NAMPT | ANXA6 | 0.8899 | 0.552 | 0.721 | 0.000999 |
| MacHypo | TCD8em | NMB | F2R | 0.356 | 0.2695 | 0.3128 | 0.000999 |
| MacHypo | TCD8em | NMB | GPR65 | 0.356 | 0.2625 | 0.3092 | 0.000999 |
| MacHypo | TCD8em | RTN4 | CD44 | 0.9167 | 1.119 | 1.0179 | 0.000999 |
| MacHypo | TCD8em | RTN4 | CD99 | 0.9167 | 1.1525 | 1.0346 | 0.000999 |
| MacHypo | TCD8em | SPP1 | CD44 | 4.3573 | 1.119 | 2.7382 | 0.000999 |
| MacHypo | TCD8em | SPP1 | ITGA4 | 4.3573 | 0.2823 | 2.3198 | 0.000999 |
| MacHypo | TCD8em | SPP1 | ITGB1 | 4.3573 | 0.2557 | 2.3065 | 0.000999 |
| MacHypo | TCD8em | TNFSF13B | ITGB1 | 0.3932 | 0.2557 | 0.3245 | 0.000999 |
| MacHypo | TCD8em | COL1A1 | CD44 | 0.3651 | 1.119 | 0.7421 | 0.003 |
| MacHypo | TCD8ex | A2M | TNFRSF14 | 0.3313 | 0.3094 | 0.3204 | 0.000999 |
| MacHypo | TCD8ex | ADM | PTGER4 | 0.9374 | 0.3089 | 0.6232 | 0.000999 |
| MacHypo | TCD8ex | AIMP1 | RACK1 | 0.3406 | 2.3353 | 1.338 | 0.000999 |
| MacHypo | TCD8ex | ALCAM | CD6 | 0.2569 | 0.318 | 0.2875 | 0.000999 |
| MacHypo | TCD8ex | ANXA1 | CD44 | 1.6214 | 0.8219 | 1.2217 | 0.000999 |

| | | | | | | | |
|---|---|---|---|---|---|---|---|
| MacHypo | TCD8ex | ANXA1 | CXCR3 | 1.6214 | 0.3647 | 0.993 | 0.000999 |
| MacHypo | TCD8ex | ANXA1 | CXCR4 | 1.6214 | 2.3472 | 1.9843 | 0.000999 |
| MacHypo | TCD8ex | ANXA1 | CXCR6 | 1.6214 | 0.6626 | 1.142 | 0.000999 |
| MacHypo | TCD8ex | ANXA1 | ITGA4 | 1.6214 | 0.4432 | 1.0323 | 0.000999 |
| MacHypo | TCD8ex | ANXA1 | VCAM1 | 1.6214 | 0.4462 | 1.0338 | 0.000999 |
| MacHypo | TCD8ex | APOE | VCAM1 | 2.2472 | 0.4462 | 1.3467 | 0.000999 |
| MacHypo | TCD8ex | APP | CXCR3 | 0.2783 | 0.3647 | 0.3215 | 0.000999 |
| MacHypo | TCD8ex | APP | CXCR4 | 0.2783 | 2.3472 | 1.3128 | 0.000999 |
| MacHypo | TCD8ex | APP | CXCR6 | 0.2783 | 0.6626 | 0.4704 | 0.000999 |
| MacHypo | TCD8ex | APP | TNFRSF14 | 0.2783 | 0.3094 | 0.2939 | 0.000999 |
| MacHypo | TCD8ex | CCL20 | CXCR3 | 0.6366 | 0.3647 | 0.5006 | 0.000999 |
| MacHypo | TCD8ex | CCL20 | CXCR4 | 0.6366 | 2.3472 | 1.4919 | 0.000999 |
| MacHypo | TCD8ex | CCL20 | CXCR6 | 0.6366 | 0.6626 | 0.6496 | 0.000999 |
| MacHypo | TCD8ex | CD86 | CTLA4 | 0.566 | 0.4883 | 0.5271 | 0.000999 |
| MacHypo | TCD8ex | COL1A1 | CD44 | 0.3651 | 0.8219 | 0.5935 | 0.000999 |
| MacHypo | TCD8ex | COL1A1 | ITGB1 | 0.3651 | 0.2995 | 0.3323 | 0.000999 |
| MacHypo | TCD8ex | CXCL16 | CXCR3 | 0.6914 | 0.3647 | 0.528 | 0.000999 |
| MacHypo | TCD8ex | CXCL16 | CXCR4 | 0.6914 | 2.3472 | 1.5193 | 0.000999 |
| MacHypo | TCD8ex | CXCL16 | CXCR6 | 0.6914 | 0.6626 | 0.677 | 0.000999 |
| MacHypo | TCD8ex | CXCL2 | CXCR3 | 0.6881 | 0.3647 | 0.5264 | 0.000999 |
| MacHypo | TCD8ex | CXCL2 | CXCR4 | 0.6881 | 2.3472 | 1.5177 | 0.000999 |
| MacHypo | TCD8ex | CXCL2 | CXCR6 | 0.6881 | 0.6626 | 0.6753 | 0.000999 |
| MacHypo | TCD8ex | CXCL3 | CXCR3 | 0.7216 | 0.3647 | 0.5431 | 0.000999 |
| MacHypo | TCD8ex | CXCL3 | CXCR4 | 0.7216 | 2.3472 | 1.5344 | 0.000999 |
| MacHypo | TCD8ex | CXCL3 | CXCR6 | 0.7216 | 0.6626 | 0.6921 | 0.000999 |
| MacHypo | TCD8ex | FN1 | CD44 | 1.342 | 0.8219 | 1.082 | 0.000999 |
| MacHypo | TCD8ex | FN1 | ITGA4 | 1.342 | 0.4432 | 0.8926 | 0.000999 |
| MacHypo | TCD8ex | FN1 | ITGB1 | 1.342 | 0.2995 | 0.8208 | 0.000999 |
| MacHypo | TCD8ex | FN1 | ITGB7 | 1.342 | 0.3007 | 0.8214 | 0.000999 |
| MacHypo | TCD8ex | FN1 | RPSA | 1.342 | 2.2697 | 1.8059 | 0.000999 |

| | | | | | | | |
|---|---|---|---|---|---|---|---|
| MacHypo | TCD8ex | GRN | TNFRSF1B | 1.0708 | 0.6436 | 0.8572 | 0.000999 |
| MacHypo | TCD8ex | GSTP1 | CD44 | 1.3428 | 0.8219 | 1.0824 | 0.000999 |
| MacHypo | TCD8ex | GSTP1 | ITGA4 | 1.3428 | 0.4432 | 0.893 | 0.000999 |
| MacHypo | TCD8ex | GSTP1 | VCAM1 | 1.3428 | 0.4462 | 0.8945 | 0.000999 |
| MacHypo | TCD8ex | HBEGF | CD44 | 0.2764 | 0.8219 | 0.5492 | 0.000999 |
| MacHypo | TCD8ex | HLA-DRA | PDCD1 | 3.2012 | 0.6109 | 1.906 | 0.000999 |
| MacHypo | TCD8ex | HLA-DRA | PTPRC | 3.2012 | 1.6762 | 2.4387 | 0.000999 |
| MacHypo | TCD8ex | HSP90B1 | ITGB1 | 0.8573 | 0.2995 | 0.5784 | 0.000999 |
| MacHypo | TCD8ex | HSP90B1 | TNFRSF1B | 0.8573 | 0.6436 | 0.7505 | 0.000999 |
| MacHypo | TCD8ex | ICAM1 | IL2RG | 0.3685 | 1.2126 | 0.7905 | 0.000999 |
| MacHypo | TCD8ex | ICAM1 | ITGB2 | 0.3685 | 0.7635 | 0.566 | 0.000999 |
| MacHypo | TCD8ex | ICAM1 | SPN | 0.3685 | 0.3486 | 0.3586 | 0.000999 |
| MacHypo | TCD8ex | ITGB1 | CD47 | 0.4597 | 0.3952 | 0.4274 | 0.000999 |
| MacHypo | TCD8ex | ITGB1 | ITGA4 | 0.4597 | 0.4432 | 0.4515 | 0.000999 |
| MacHypo | TCD8ex | ITGB1 | RACK1 | 0.4597 | 2.3353 | 1.3975 | 0.000999 |
| MacHypo | TCD8ex | ITGB1 | VCAM1 | 0.4597 | 0.4462 | 0.4529 | 0.000999 |
| MacHypo | TCD8ex | LGALS3 | CD6 | 1.669 | 0.318 | 0.9935 | 0.000999 |
| MacHypo | TCD8ex | LGALS3 | ITGB1 | 1.669 | 0.2995 | 0.9842 | 0.000999 |
| MacHypo | TCD8ex | LGALS3 | SPN | 1.669 | 0.3486 | 1.0088 | 0.000999 |
| MacHypo | TCD8ex | MIF | CD74 | 1.9478 | 2.5058 | 2.2268 | 0.000999 |
| MacHypo | TCD8ex | MIF | ITGA4 | 1.9478 | 0.4432 | 1.1955 | 0.000999 |
| MacHypo | TCD8ex | MIF | TNFRSF14 | 1.9478 | 0.3094 | 1.1286 | 0.000999 |
| MacHypo | TCD8ex | MMP9 | CD44 | 0.8087 | 0.8219 | 0.8153 | 0.000999 |
| MacHypo | TCD8ex | MMP9 | ITGB1 | 0.8087 | 0.2995 | 0.5541 | 0.000999 |
| MacHypo | TCD8ex | NAMPT | ANXA6 | 0.8899 | 0.5678 | 0.7288 | 0.000999 |
| MacHypo | TCD8ex | NMB | GPR65 | 0.356 | 0.3904 | 0.3732 | 0.000999 |
| MacHypo | TCD8ex | RTN4 | CD44 | 0.9167 | 0.8219 | 0.8693 | 0.000999 |
| MacHypo | TCD8ex | RTN4 | CD99 | 0.9167 | 1.2049 | 1.0608 | 0.000999 |
| MacHypo | TCD8ex | SPP1 | CD44 | 4.3573 | 0.8219 | 2.5896 | 0.000999 |
| MacHypo | TCD8ex | SPP1 | ITGA4 | 4.3573 | 0.4432 | 2.4003 | 0.000999 |

| | | | | | | | |
|---|---|---|---|---|---|---|---|
| MacHypo | TCD8ex | SPP1 | ITGB1 | 4.3573 | 0.2995 | 2.3284 | 0.000999 |
| MacHypo | TCD8ex | TNFSF13B | ITGB1 | 0.3932 | 0.2995 | 0.3463 | 0.000999 |
| MacHypo | TGD | A2M | TNFRSF14 | 0.3313 | 0.4642 | 0.3977 | 0.000999 |
| MacHypo | TGD | ADM | PTGDR | 0.9374 | 0.7153 | 0.8263 | 0.000999 |
| MacHypo | TGD | AIMP1 | RACK1 | 0.3406 | 2.7287 | 1.5347 | 0.000999 |
| MacHypo | TGD | ALCAM | CD6 | 0.2569 | 0.4684 | 0.3627 | 0.000999 |
| MacHypo | TGD | ANXA1 | CXCR3 | 1.6214 | 1.1748 | 1.3981 | 0.000999 |
| MacHypo | TGD | ANXA1 | CXCR4 | 1.6214 | 1.4894 | 1.5554 | 0.000999 |
| MacHypo | TGD | ANXA1 | ITGA4 | 1.6214 | 0.3968 | 1.0091 | 0.000999 |
| MacHypo | TGD | APOE | SORL1 | 2.2472 | 0.3318 | 1.2895 | 0.000999 |
| MacHypo | TGD | APP | CXCR3 | 0.2783 | 1.1748 | 0.7266 | 0.000999 |
| MacHypo | TGD | APP | CXCR4 | 0.2783 | 1.4894 | 0.8839 | 0.000999 |
| MacHypo | TGD | APP | SORL1 | 0.2783 | 0.3318 | 0.3051 | 0.000999 |
| MacHypo | TGD | APP | TGFBR2 | 0.2783 | 0.3677 | 0.323 | 0.000999 |
| MacHypo | TGD | ARF1 | ITGA4 | 0.6074 | 0.3968 | 0.5021 | 0.000999 |
| MacHypo | TGD | CALM1 | RPSA | 1.2536 | 2.8814 | 2.0675 | 0.000999 |
| MacHypo | TGD | CALR | IL7R | 0.9194 | 1.3862 | 1.1528 | 0.000999 |
| MacHypo | TGD | CCL20 | CXCR3 | 0.6366 | 1.1748 | 0.9057 | 0.000999 |
| MacHypo | TGD | CCL20 | CXCR4 | 0.6366 | 1.4894 | 1.063 | 0.000999 |
| MacHypo | TGD | CD99 | CD99 | 0.765 | 1.7154 | 1.2402 | 0.000999 |
| MacHypo | TGD | CXCL16 | CXCR3 | 0.6914 | 1.1748 | 0.9331 | 0.000999 |
| MacHypo | TGD | CXCL16 | CXCR4 | 0.6914 | 1.4894 | 1.0904 | 0.000999 |
| MacHypo | TGD | CXCL2 | CXCR3 | 0.6881 | 1.1748 | 0.9314 | 0.000999 |
| MacHypo | TGD | CXCL2 | CXCR4 | 0.6881 | 1.4894 | 1.0887 | 0.000999 |
| MacHypo | TGD | CXCL3 | CXCR3 | 0.7216 | 1.1748 | 0.9482 | 0.000999 |
| MacHypo | TGD | CXCL3 | CXCR4 | 0.7216 | 1.4894 | 1.1055 | 0.000999 |
| MacHypo | TGD | FN1 | ITGA4 | 1.342 | 0.3968 | 0.8694 | 0.000999 |
| MacHypo | TGD | FN1 | ITGB1 | 1.342 | 0.3657 | 0.8539 | 0.000999 |
| MacHypo | TGD | FN1 | ITGB7 | 1.342 | 0.459 | 0.9005 | 0.000999 |
| MacHypo | TGD | FN1 | RPSA | 1.342 | 2.8814 | 2.1117 | 0.000999 |

| | | | | | | | |
|---|---|---|---|---|---|---|---|
| MacHypo | TGD | GRN | TNFRSF1A | 1.0708 | 0.3099 | 0.6904 | 0.000999 |
| MacHypo | TGD | GSTP1 | IL7R | 1.3428 | 1.3862 | 1.3645 | 0.000999 |
| MacHypo | TGD | GSTP1 | ITGA4 | 1.3428 | 0.3968 | 0.8698 | 0.000999 |
| MacHypo | TGD | HLA-A | CD8A | 2.5037 | 1.746 | 2.1248 | 0.000999 |
| MacHypo | TGD | HLA-DRA | PTPRC | 3.2012 | 1.8653 | 2.5333 | 0.000999 |
| MacHypo | TGD | HLA-E | CD8A | 1.1724 | 1.746 | 1.4592 | 0.000999 |
| MacHypo | TGD | HLA-E | KLRC2 | 1.1724 | 0.8082 | 0.9903 | 0.000999 |
| MacHypo | TGD | HLA-E | KLRD1 | 1.1724 | 1.3451 | 1.2587 | 0.000999 |
| MacHypo | TGD | ICAM1 | IL2RG | 0.3685 | 1.561 | 0.9648 | 0.000999 |
| MacHypo | TGD | ICAM1 | ITGAL | 0.3685 | 0.3731 | 0.3708 | 0.000999 |
| MacHypo | TGD | ICAM1 | ITGB2 | 0.3685 | 1.4444 | 0.9065 | 0.000999 |
| MacHypo | TGD | ICAM1 | SPN | 0.3685 | 0.5644 | 0.4664 | 0.000999 |
| MacHypo | TGD | ITGB1 | ITGA4 | 0.4597 | 0.3968 | 0.4283 | 0.000999 |
| MacHypo | TGD | ITGB1 | PTK2B | 0.4597 | 0.3668 | 0.4132 | 0.000999 |
| MacHypo | TGD | ITGB1 | RACK1 | 0.4597 | 2.7287 | 1.5942 | 0.000999 |
| MacHypo | TGD | ITGB2 | ITGAL | 1.0357 | 0.3731 | 0.7044 | 0.000999 |
| MacHypo | TGD | LGALS3 | CD6 | 1.669 | 0.4684 | 1.0687 | 0.000999 |
| MacHypo | TGD | LGALS3 | ITGB1 | 1.669 | 0.3657 | 1.0173 | 0.000999 |
| MacHypo | TGD | LGALS3 | SPN | 1.669 | 0.5644 | 1.1167 | 0.000999 |
| MacHypo | TGD | LGALS3 | TGFBR2 | 1.669 | 0.3677 | 1.0183 | 0.000999 |
| MacHypo | TGD | LRPAP1 | SORL1 | 0.4248 | 0.3318 | 0.3783 | 0.000999 |
| MacHypo | TGD | MIF | ITGA4 | 1.9478 | 0.3968 | 1.1723 | 0.000999 |
| MacHypo | TGD | MIF | TNFRSF14 | 1.9478 | 0.4642 | 1.206 | 0.000999 |
| MacHypo | TGD | NAMPT | ANXA6 | 0.8899 | 0.8553 | 0.8726 | 0.000999 |
| MacHypo | TGD | NMB | GPR65 | 0.356 | 0.5368 | 0.4464 | 0.000999 |
| MacHypo | TGD | PTPRC | CD2 | 0.4793 | 1.4522 | 0.9658 | 0.000999 |
| MacHypo | TGD | PTPRC | CD8A | 0.4793 | 1.746 | 1.1127 | 0.000999 |
| MacHypo | TGD | RPS19 | ITGA4 | 3.5438 | 0.3968 | 1.9703 | 0.000999 |
| MacHypo | TGD | RPS19 | RPSA | 3.5438 | 2.8814 | 3.2126 | 0.000999 |
| MacHypo | TGD | RTN4 | CD99 | 0.9167 | 1.7154 | 1.3161 | 0.000999 |

| | | | | | | | |
|---|---|---|---|---|---|---|---|
| MacHypo | TGD | SPP1 | CD44 | 4.3573 | 1.1342 | 2.7458 | 0.000999 |
| MacHypo | TGD | SPP1 | ITGA4 | 4.3573 | 0.3968 | 2.3771 | 0.000999 |
| MacHypo | TGD | SPP1 | ITGB1 | 4.3573 | 0.3657 | 2.3615 | 0.000999 |
| MacHypo | TGD | VEGFA | PTK2B | 0.5814 | 0.3668 | 0.4741 | 0.000999 |
| MacHypo | TGD | APP | TNFRSF14 | 0.2783 | 0.4642 | 0.3713 | 0.002 |
| MacHypo | TGD | ITGB1 | CD47 | 0.4597 | 0.4765 | 0.4681 | 0.004 |
| MacHypo | TGD | HMGB1 | ITGA4 | 0.9251 | 0.3968 | 0.6609 | 0.005 |
| MacHypo | TGD | ITGB1 | CD46 | 0.4597 | 0.3085 | 0.3841 | 0.005 |
| MacHypo | TGD | MMP9 | ITGB1 | 0.8087 | 0.3657 | 0.5872 | 0.00599 |
| NKT | MacHypo | ALOX5AP | ALOX5 | 0.5021 | 0.2606 | 0.3814 | 0.000999 |
| NKT | MacHypo | ANXA1 | C3AR1 | 1.2259 | 0.2925 | 0.7592 | 0.000999 |
| NKT | MacHypo | ANXA1 | C5AR1 | 1.2259 | 0.4508 | 0.8384 | 0.000999 |
| NKT | MacHypo | ANXA1 | CCR1 | 1.2259 | 0.2564 | 0.7412 | 0.000999 |
| NKT | MacHypo | ANXA1 | CD4 | 1.2259 | 0.2639 | 0.7449 | 0.000999 |
| NKT | MacHypo | ANXA1 | CD44 | 1.2259 | 1.5822 | 1.4041 | 0.000999 |
| NKT | MacHypo | ANXA1 | FPR3 | 1.2259 | 0.3793 | 0.8026 | 0.000999 |
| NKT | MacHypo | ARF1 | CD4 | 0.7991 | 0.2639 | 0.5315 | 0.000999 |
| NKT | MacHypo | ARF1 | CD44 | 0.7991 | 1.5822 | 1.1907 | 0.000999 |
| NKT | MacHypo | CALR | CD44 | 0.9479 | 1.5822 | 1.2651 | 0.000999 |
| NKT | MacHypo | CD2 | CD4 | 1.1465 | 0.2639 | 0.7052 | 0.000999 |
| NKT | MacHypo | CD6 | ALCAM | 0.4881 | 0.2569 | 0.3725 | 0.000999 |
| NKT | MacHypo | GSTP1 | CD44 | 0.7909 | 1.5822 | 1.1865 | 0.000999 |
| NKT | MacHypo | HMGB1 | AGTRAP | 1.195 | 0.263 | 0.729 | 0.000999 |
| NKT | MacHypo | HMGB1 | LY96 | 1.195 | 0.7779 | 0.9864 | 0.000999 |
| NKT | MacHypo | HMGB1 | TLR2 | 1.195 | 0.2562 | 0.7256 | 0.000999 |
| NKT | MacHypo | HMGB2 | CD44 | 0.6467 | 1.5822 | 1.1145 | 0.000999 |
| NKT | MacHypo | HMGB2 | DAB2 | 0.6467 | 0.6139 | 0.6303 | 0.000999 |
| NKT | MacHypo | IL16 | CD4 | 0.3556 | 0.2639 | 0.3097 | 0.000999 |
| NKT | MacHypo | ITGA4 | CD44 | 0.2946 | 1.5822 | 0.9384 | 0.000999 |
| NKT | MacHypo | ITGA4 | TFRC | 0.2946 | 0.4916 | 0.3931 | 0.000999 |

| | | | | | | | |
|---|---|---|---|---|---|---|---|
| NKT | MacHypo | ITGAL | ICAM1 | 0.6778 | 0.3685 | 0.5232 | 0.000999 |
| NKT | MacHypo | ITGB1 | CD151 | 0.488 | 0.3292 | 0.4086 | 0.000999 |
| NKT | MacHypo | ITGB1 | DAB2 | 0.488 | 0.6139 | 0.5509 | 0.000999 |
| NKT | MacHypo | ITGB1 | ITGAX | 0.488 | 0.3626 | 0.4253 | 0.000999 |
| NKT | MacHypo | ITGB1 | PLAUR | 0.488 | 1.5015 | 0.9947 | 0.000999 |
| NKT | MacHypo | ITGB2 | ICAM1 | 1.6481 | 0.3685 | 1.0083 | 0.000999 |
| NKT | MacHypo | ITGB2 | ITGAM | 1.6481 | 0.328 | 0.9881 | 0.000999 |
| NKT | MacHypo | ITGB2 | ITGAX | 1.6481 | 0.3626 | 1.0053 | 0.000999 |
| NKT | MacHypo | ITGB2 | PLAUR | 1.6481 | 1.5015 | 1.5748 | 0.000999 |
| NKT | MacHypo | MIF | CD74 | 0.6657 | 3.1474 | 1.9065 | 0.000999 |
| NKT | MacHypo | PTPRC | CD4 | 1.8797 | 0.2639 | 1.0718 | 0.000999 |
| NKT | MacHypo | PTPRC | CD44 | 1.8797 | 1.5822 | 1.7309 | 0.000999 |
| NKT | MacHypo | PTPRC | IFNAR1 | 1.8797 | 0.3706 | 1.1251 | 0.000999 |
| NKT | MacHypo | RPS19 | C5AR1 | 3.5336 | 0.4508 | 1.9922 | 0.000999 |
| NKT | MacHypo | RPS19 | CD4 | 3.5336 | 0.2639 | 1.8988 | 0.000999 |
| NKT | MacHypo | RPS19 | ICAM1 | 3.5336 | 0.3685 | 1.9511 | 0.000999 |
| NKT | MacHypo | RTN4 | CD44 | 0.4374 | 1.5822 | 1.0098 | 0.000999 |
| NKT | MacHypo | TGFB1 | FCN1 | 0.5441 | 0.3143 | 0.4292 | 0.000999 |
| NKT | MacHypo | TGFB1 | SDC2 | 0.5441 | 0.6066 | 0.5753 | 0.000999 |
| NKT | MacHypo | TGFB1 | TLR2 | 0.5441 | 0.2562 | 0.4001 | 0.000999 |
| TCD4ex | MacHypo | ALOX5AP | ALOX5 | 1.119 | 0.2606 | 0.6898 | 0.000999 |
| TCD4ex | MacHypo | ANXA1 | C5AR1 | 0.5174 | 0.4508 | 0.4841 | 0.000999 |
| TCD4ex | MacHypo | ANXA1 | CD44 | 0.5174 | 1.5822 | 1.0498 | 0.000999 |
| TCD4ex | MacHypo | ANXA1 | FPR3 | 0.5174 | 0.3793 | 0.4484 | 0.000999 |
| TCD4ex | MacHypo | APOE | SDC2 | 0.2557 | 0.6066 | 0.4311 | 0.000999 |
| TCD4ex | MacHypo | AREG | ICAM1 | 0.3427 | 0.3685 | 0.3556 | 0.000999 |
| TCD4ex | MacHypo | ARF1 | CD44 | 0.6827 | 1.5822 | 1.1324 | 0.000999 |
| TCD4ex | MacHypo | BTLA | TNFRSF14 | 0.3493 | 0.3382 | 0.3438 | 0.000999 |
| TCD4ex | MacHypo | CALR | CD44 | 0.748 | 1.5822 | 1.1651 | 0.000999 |
| TCD4ex | MacHypo | CD6 | ALCAM | 0.5338 | 0.2569 | 0.3953 | 0.000999 |

| | | | | | | | |
|---|---|---|---|---|---|---|---|
| TCD4ex | MacHypo | CXCL13 | C3AR1 | 2.3367 | 0.2925 | 1.3146 | 0.000999 |
| TCD4ex | MacHypo | CXCL13 | C5AR1 | 2.3367 | 0.4508 | 1.3937 | 0.000999 |
| TCD4ex | MacHypo | CXCL13 | CCR1 | 2.3367 | 0.2564 | 1.2965 | 0.000999 |
| TCD4ex | MacHypo | GSTP1 | CD44 | 0.4481 | 1.5822 | 1.0151 | 0.000999 |
| TCD4ex | MacHypo | HMGB1 | AGTRAP | 1.3876 | 0.263 | 0.8253 | 0.000999 |
| TCD4ex | MacHypo | HMGB1 | LY96 | 1.3876 | 0.7779 | 1.0827 | 0.000999 |
| TCD4ex | MacHypo | HMGB1 | TLR2 | 1.3876 | 0.2562 | 0.8219 | 0.000999 |
| TCD4ex | MacHypo | HMGB2 | CD44 | 1.0243 | 1.5822 | 1.3033 | 0.000999 |
| TCD4ex | MacHypo | HMGB2 | DAB2 | 1.0243 | 0.6139 | 0.8191 | 0.000999 |
| TCD4ex | MacHypo | ICAM3 | ITGB2 | 0.573 | 1.0357 | 0.8044 | 0.000999 |
| TCD4ex | MacHypo | ITGAL | ICAM1 | 0.2603 | 0.3685 | 0.3144 | 0.000999 |
| TCD4ex | MacHypo | ITGB1 | CD151 | 0.4615 | 0.3292 | 0.3954 | 0.000999 |
| TCD4ex | MacHypo | ITGB1 | DAB2 | 0.4615 | 0.6139 | 0.5377 | 0.000999 |
| TCD4ex | MacHypo | ITGB1 | ITGAX | 0.4615 | 0.3626 | 0.412 | 0.000999 |
| TCD4ex | MacHypo | ITGB1 | PLAUR | 0.4615 | 1.5015 | 0.9815 | 0.000999 |
| TCD4ex | MacHypo | ITGB2 | ICAM1 | 0.7871 | 0.3685 | 0.5778 | 0.000999 |
| TCD4ex | MacHypo | ITGB2 | ITGAM | 0.7871 | 0.328 | 0.5575 | 0.000999 |
| TCD4ex | MacHypo | ITGB2 | ITGAX | 0.7871 | 0.3626 | 0.5748 | 0.000999 |
| TCD4ex | MacHypo | ITGB2 | PLAUR | 0.7871 | 1.5015 | 1.1443 | 0.000999 |
| TCD4ex | MacHypo | LTB | ITGB1 | 0.9655 | 0.4597 | 0.7126 | 0.000999 |
| TCD4ex | MacHypo | MIF | CD74 | 0.6999 | 3.1474 | 1.9236 | 0.000999 |
| TCD4ex | MacHypo | PTPRC | CD44 | 1.7493 | 1.5822 | 1.6658 | 0.000999 |
| TCD4ex | MacHypo | PTPRC | IFNAR1 | 1.7493 | 0.3706 | 1.06 | 0.000999 |
| TCD4ex | MacHypo | RPS19 | C5AR1 | 3.6329 | 0.4508 | 2.0419 | 0.000999 |
| TCD4ex | MacHypo | RPS19 | ICAM1 | 3.6329 | 0.3685 | 2.0007 | 0.000999 |
| TCD4ex | MacHypo | RTN4 | CD44 | 0.355 | 1.5822 | 0.9686 | 0.000999 |
| TCD4reg | MacHypo | ALOX5AP | ALOX5 | 0.4589 | 0.2606 | 0.3597 | 0.000999 |
| TCD4reg | MacHypo | APOE | SDC2 | 0.2934 | 0.6066 | 0.45 | 0.000999 |
| TCD4reg | MacHypo | ARF1 | CD44 | 0.7185 | 1.5822 | 1.1504 | 0.000999 |
| TCD4reg | MacHypo | CALR | CD44 | 0.6835 | 1.5822 | 1.1329 | 0.000999 |

| | | | | | | | |
|---|---|---|---|---|---|---|---|
| TCD4reg | MacHypo | CD6 | ALCAM | 0.2651 | 0.2569 | 0.261 | 0.000999 |
| TCD4reg | MacHypo | GSTP1 | CD44 | 0.4917 | 1.5822 | 1.0369 | 0.000999 |
| TCD4reg | MacHypo | HMGB1 | AGTRAP | 1.2547 | 0.263 | 0.7589 | 0.000999 |
| TCD4reg | MacHypo | HMGB1 | LY96 | 1.2547 | 0.7779 | 1.0163 | 0.000999 |
| TCD4reg | MacHypo | HMGB1 | TLR2 | 1.2547 | 0.2562 | 0.7555 | 0.000999 |
| TCD4reg | MacHypo | HMGB2 | CD44 | 0.616 | 1.5822 | 1.0991 | 0.000999 |
| TCD4reg | MacHypo | HMGB2 | DAB2 | 0.616 | 0.6139 | 0.615 | 0.000999 |
| TCD4reg | MacHypo | ICAM3 | ITGB2 | 0.6663 | 1.0357 | 0.851 | 0.000999 |
| TCD4reg | MacHypo | ITGB1 | CD151 | 0.4001 | 0.3292 | 0.3647 | 0.000999 |
| TCD4reg | MacHypo | ITGB1 | DAB2 | 0.4001 | 0.6139 | 0.507 | 0.000999 |
| TCD4reg | MacHypo | ITGB1 | ITGAX | 0.4001 | 0.3626 | 0.3813 | 0.000999 |
| TCD4reg | MacHypo | ITGB1 | PLAUR | 0.4001 | 1.5015 | 0.9508 | 0.000999 |
| TCD4reg | MacHypo | ITGB2 | ICAM1 | 0.6238 | 0.3685 | 0.4962 | 0.000999 |
| TCD4reg | MacHypo | ITGB2 | ITGAM | 0.6238 | 0.328 | 0.4759 | 0.000999 |
| TCD4reg | MacHypo | ITGB2 | ITGAX | 0.6238 | 0.3626 | 0.4932 | 0.000999 |
| TCD4reg | MacHypo | ITGB2 | PLAUR | 0.6238 | 1.5015 | 1.0627 | 0.000999 |
| TCD4reg | MacHypo | LGALS3 | CLEC7A | 0.4492 | 0.4087 | 0.4289 | 0.000999 |
| TCD4reg | MacHypo | LGALS3 | FCGR2A | 0.4492 | 0.7828 | 0.616 | 0.000999 |
| TCD4reg | MacHypo | LGALS3 | FCGR2B | 0.4492 | 0.7745 | 0.6118 | 0.000999 |
| TCD4reg | MacHypo | LGALS3 | TFRC | 0.4492 | 0.4916 | 0.4704 | 0.000999 |
| TCD4reg | MacHypo | LTB | ITGB1 | 2.242 | 0.4597 | 1.3509 | 0.000999 |
| TCD4reg | MacHypo | MIF | CD74 | 0.6295 | 3.1474 | 1.8885 | 0.000999 |
| TCD4reg | MacHypo | PTPRC | CD44 | 1.6603 | 1.5822 | 1.6213 | 0.000999 |
| TCD4reg | MacHypo | PTPRC | IFNAR1 | 1.6603 | 0.3706 | 1.0155 | 0.000999 |
| TCD4reg | MacHypo | RPS19 | C5AR1 | 3.6897 | 0.4508 | 2.0702 | 0.000999 |
| TCD4reg | MacHypo | RPS19 | ICAM1 | 3.6897 | 0.3685 | 2.0291 | 0.000999 |
| TCD4reg | MacHypo | RTN4 | CD44 | 0.3528 | 1.5822 | 0.9675 | 0.000999 |
| TCD4reg | MacHypo | CD99 | CD99 | 0.6603 | 0.765 | 0.7127 | 0.004 |
| TCD8em | MacHypo | ALOX5AP | ALOX5 | 0.7973 | 0.2606 | 0.529 | 0.000999 |
| TCD8em | MacHypo | ANXA1 | C3AR1 | 1.0693 | 0.2925 | 0.6809 | 0.000999 |

| | | | | | | | |
|---|---|---|---|---|---|---|---|
| TCD8em | MacHypo | ANXA1 | C5AR1 | 1.0693 | 0.4508 | 0.76 | 0.000999 |
| TCD8em | MacHypo | ANXA1 | CCR1 | 1.0693 | 0.2564 | 0.6628 | 0.000999 |
| TCD8em | MacHypo | ANXA1 | CD4 | 1.0693 | 0.2639 | 0.6666 | 0.000999 |
| TCD8em | MacHypo | ANXA1 | CD44 | 1.0693 | 1.5822 | 1.3257 | 0.000999 |
| TCD8em | MacHypo | ANXA1 | FPR3 | 1.0693 | 0.3793 | 0.7243 | 0.000999 |
| TCD8em | MacHypo | APOE | SDC2 | 0.2791 | 0.6066 | 0.4429 | 0.000999 |
| TCD8em | MacHypo | ARF1 | CD4 | 0.6684 | 0.2639 | 0.4661 | 0.000999 |
| TCD8em | MacHypo | ARF1 | CD44 | 0.6684 | 1.5822 | 1.1253 | 0.000999 |
| TCD8em | MacHypo | CALM1 | TNFRSF1B | 1.5262 | 0.4397 | 0.9829 | 0.000999 |
| TCD8em | MacHypo | CALR | CD44 | 0.7141 | 1.5822 | 1.1482 | 0.000999 |
| TCD8em | MacHypo | CD2 | CD4 | 1.5961 | 0.2639 | 0.93 | 0.000999 |
| TCD8em | MacHypo | CD6 | ALCAM | 0.4204 | 0.2569 | 0.3387 | 0.000999 |
| TCD8em | MacHypo | GSTP1 | CD44 | 0.767 | 1.5822 | 1.1746 | 0.000999 |
| TCD8em | MacHypo | HLA-DMA | CD4 | 0.2969 | 0.2639 | 0.2804 | 0.000999 |
| TCD8em | MacHypo | HMGB1 | AGTRAP | 1.3636 | 0.263 | 0.8133 | 0.000999 |
| TCD8em | MacHypo | HMGB1 | LY96 | 1.3636 | 0.7779 | 1.0707 | 0.000999 |
| TCD8em | MacHypo | HMGB1 | TLR2 | 1.3636 | 0.2562 | 0.8099 | 0.000999 |
| TCD8em | MacHypo | HMGB2 | CD44 | 0.7212 | 1.5822 | 1.1517 | 0.000999 |
| TCD8em | MacHypo | HMGB2 | DAB2 | 0.7212 | 0.6139 | 0.6675 | 0.000999 |
| TCD8em | MacHypo | HSP90B1 | ITGB1 | 0.471 | 0.4597 | 0.4654 | 0.000999 |
| TCD8em | MacHypo | HSP90B1 | TNFRSF1B | 0.471 | 0.4397 | 0.4554 | 0.000999 |
| TCD8em | MacHypo | IFNG | IFNGR1 | 0.3967 | 0.467 | 0.4319 | 0.000999 |
| TCD8em | MacHypo | IFNG | IFNGR2 | 0.3967 | 0.5438 | 0.4702 | 0.000999 |
| TCD8em | MacHypo | IL16 | CD4 | 0.4085 | 0.2639 | 0.3362 | 0.000999 |
| TCD8em | MacHypo | ITGA4 | CD44 | 0.2823 | 1.5822 | 0.9323 | 0.000999 |
| TCD8em | MacHypo | ITGA4 | TFRC | 0.2823 | 0.4916 | 0.387 | 0.000999 |
| TCD8em | MacHypo | ITGAL | ICAM1 | 0.3162 | 0.3685 | 0.3424 | 0.000999 |
| TCD8em | MacHypo | ITGB1 | CD151 | 0.2557 | 0.3292 | 0.2925 | 0.000999 |
| TCD8em | MacHypo | ITGB1 | DAB2 | 0.2557 | 0.6139 | 0.4348 | 0.000999 |
| TCD8em | MacHypo | ITGB1 | ITGAX | 0.2557 | 0.3626 | 0.3092 | 0.000999 |

| | | | | | | | |
|---|---|---|---|---|---|---|---|
| TCD8em | MacHypo | ITGB1 | PLAUR | 0.2557 | 1.5015 | 0.8786 | 0.000999 |
| TCD8em | MacHypo | ITGB2 | ICAM1 | 1.0018 | 0.3685 | 0.6852 | 0.000999 |
| TCD8em | MacHypo | ITGB2 | ITGAM | 1.0018 | 0.328 | 0.6649 | 0.000999 |
| TCD8em | MacHypo | ITGB2 | ITGAX | 1.0018 | 0.3626 | 0.6822 | 0.000999 |
| TCD8em | MacHypo | ITGB2 | PLAUR | 1.0018 | 1.5015 | 1.2517 | 0.000999 |
| TCD8em | MacHypo | LGALS3 | CLEC7A | 0.2689 | 0.4087 | 0.3388 | 0.000999 |
| TCD8em | MacHypo | LGALS3 | FCGR2A | 0.2689 | 0.7828 | 0.5258 | 0.000999 |
| TCD8em | MacHypo | LGALS3 | FCGR2B | 0.2689 | 0.7745 | 0.5217 | 0.000999 |
| TCD8em | MacHypo | LGALS3 | ITGB1 | 0.2689 | 0.4597 | 0.3643 | 0.000999 |
| TCD8em | MacHypo | LGALS3 | TFRC | 0.2689 | 0.4916 | 0.3802 | 0.000999 |
| TCD8em | MacHypo | LTB | ITGB1 | 0.6352 | 0.4597 | 0.5474 | 0.000999 |
| TCD8em | MacHypo | MIF | CD74 | 0.6063 | 3.1474 | 1.8769 | 0.000999 |
| TCD8em | MacHypo | PTPRC | CD4 | 1.7912 | 0.2639 | 1.0275 | 0.000999 |
| TCD8em | MacHypo | PTPRC | CD44 | 1.7912 | 1.5822 | 1.6867 | 0.000999 |
| TCD8em | MacHypo | PTPRC | IFNAR1 | 1.7912 | 0.3706 | 1.0809 | 0.000999 |
| TCD8em | MacHypo | RPS19 | C5AR1 | 3.536 | 0.4508 | 1.9934 | 0.000999 |
| TCD8em | MacHypo | RPS19 | CD4 | 3.536 | 0.2639 | 1.9 | 0.000999 |
| TCD8em | MacHypo | RPS19 | ICAM1 | 3.536 | 0.3685 | 1.9523 | 0.000999 |
| TCD8em | MacHypo | RTN4 | CD44 | 0.3632 | 1.5822 | 0.9727 | 0.000999 |
| TCD8em | MacHypo | SELL | CD4 | 0.258 | 0.2639 | 0.2609 | 0.000999 |
| TCD8em | MacHypo | TNF | NRP1 | 0.2667 | 0.2592 | 0.263 | 0.000999 |
| TCD8em | MacHypo | TNF | TNFRSF1B | 0.2667 | 0.4397 | 0.3532 | 0.000999 |
| TCD8em | MacHypo | SPP1 | CD44 | 0.3608 | 1.5822 | 0.9715 | 0.004 |
| TCD8ex | MacHypo | ALOX5AP | ALOX5 | 1.5024 | 0.2606 | 0.8815 | 0.000999 |
| TCD8ex | MacHypo | ANXA1 | C3AR1 | 0.5645 | 0.2925 | 0.4285 | 0.000999 |
| TCD8ex | MacHypo | ANXA1 | C5AR1 | 0.5645 | 0.4508 | 0.5076 | 0.000999 |
| TCD8ex | MacHypo | ANXA1 | CCR1 | 0.5645 | 0.2564 | 0.4104 | 0.000999 |
| TCD8ex | MacHypo | ANXA1 | CD4 | 0.5645 | 0.2639 | 0.4142 | 0.000999 |
| TCD8ex | MacHypo | ANXA1 | CD44 | 0.5645 | 1.5822 | 1.0733 | 0.000999 |
| TCD8ex | MacHypo | ANXA1 | FPR3 | 0.5645 | 0.3793 | 0.4719 | 0.000999 |

| | | | | | | | |
|---|---|---|---|---|---|---|---|
| TCD8ex | MacHypo | APOE | SDC2 | 0.2814 | 0.6066 | 0.444 | 0.000999 |
| TCD8ex | MacHypo | AREG | ICAM1 | 0.2758 | 0.3685 | 0.3222 | 0.000999 |
| TCD8ex | MacHypo | ARF1 | CD4 | 0.7835 | 0.2639 | 0.5237 | 0.000999 |
| TCD8ex | MacHypo | ARF1 | CD44 | 0.7835 | 1.5822 | 1.1829 | 0.000999 |
| TCD8ex | MacHypo | CALR | CD44 | 0.9634 | 1.5822 | 1.2728 | 0.000999 |
| TCD8ex | MacHypo | CALR | IL7R | 0.9634 | 0.3899 | 0.6766 | 0.000999 |
| TCD8ex | MacHypo | CD2 | CD4 | 2.0502 | 0.2639 | 1.157 | 0.000999 |
| TCD8ex | MacHypo | CD6 | ALCAM | 0.318 | 0.2569 | 0.2875 | 0.000999 |
| TCD8ex | MacHypo | CXCL13 | C3AR1 | 1.7749 | 0.2925 | 1.0337 | 0.000999 |
| TCD8ex | MacHypo | CXCL13 | C5AR1 | 1.7749 | 0.4508 | 1.1129 | 0.000999 |
| TCD8ex | MacHypo | CXCL13 | CCR1 | 1.7749 | 0.2564 | 1.0156 | 0.000999 |
| TCD8ex | MacHypo | GSTP1 | CD44 | 0.8286 | 1.5822 | 1.2054 | 0.000999 |
| TCD8ex | MacHypo | GSTP1 | IL7R | 0.8286 | 0.3899 | 0.6092 | 0.000999 |
| TCD8ex | MacHypo | HLA-DMA | CD4 | 0.3991 | 0.2639 | 0.3315 | 0.000999 |
| TCD8ex | MacHypo | HLA-DRA | CD4 | 1.247 | 0.2639 | 0.7554 | 0.000999 |
| TCD8ex | MacHypo | HMGB1 | AGTRAP | 1.5414 | 0.263 | 0.9022 | 0.000999 |
| TCD8ex | MacHypo | HMGB1 | LY96 | 1.5414 | 0.7779 | 1.1597 | 0.000999 |
| TCD8ex | MacHypo | HMGB1 | TLR2 | 1.5414 | 0.2562 | 0.8988 | 0.000999 |
| TCD8ex | MacHypo | HMGB2 | CD44 | 0.85 | 1.5822 | 1.2161 | 0.000999 |
| TCD8ex | MacHypo | HMGB2 | DAB2 | 0.85 | 0.6139 | 0.7319 | 0.000999 |
| TCD8ex | MacHypo | HSP90B1 | ITGB1 | 0.5666 | 0.4597 | 0.5132 | 0.000999 |
| TCD8ex | MacHypo | ICAM3 | ITGB2 | 0.5405 | 1.0357 | 0.7881 | 0.000999 |
| TCD8ex | MacHypo | IFNG | IFNGR1 | 0.8575 | 0.467 | 0.6623 | 0.000999 |
| TCD8ex | MacHypo | IFNG | IFNGR2 | 0.8575 | 0.5438 | 0.7007 | 0.000999 |
| TCD8ex | MacHypo | ITGA4 | CD44 | 0.4432 | 1.5822 | 1.0127 | 0.000999 |
| TCD8ex | MacHypo | ITGA4 | TFRC | 0.4432 | 0.4916 | 0.4674 | 0.000999 |
| TCD8ex | MacHypo | ITGB1 | CD151 | 0.2995 | 0.3292 | 0.3144 | 0.000999 |
| TCD8ex | MacHypo | ITGB1 | DAB2 | 0.2995 | 0.6139 | 0.4567 | 0.000999 |
| TCD8ex | MacHypo | ITGB1 | ITGAX | 0.2995 | 0.3626 | 0.331 | 0.000999 |
| TCD8ex | MacHypo | ITGB1 | PLAUR | 0.2995 | 1.5015 | 0.9005 | 0.000999 |

| | | | | | | | |
|---|---|---|---|---|---|---|---|
| TCD8ex | MacHypo | ITGB2 | ICAM1 | 0.7635 | 0.3685 | 0.566 | 0.000999 |
| TCD8ex | MacHypo | ITGB2 | ITGAM | 0.7635 | 0.328 | 0.5457 | 0.000999 |
| TCD8ex | MacHypo | ITGB2 | ITGAX | 0.7635 | 0.3626 | 0.563 | 0.000999 |
| TCD8ex | MacHypo | ITGB2 | PLAUR | 0.7635 | 1.5015 | 1.1325 | 0.000999 |
| TCD8ex | MacHypo | LGALS3 | CLEC7A | 0.3824 | 0.4087 | 0.3955 | 0.000999 |
| TCD8ex | MacHypo | LGALS3 | FCGR2A | 0.3824 | 0.7828 | 0.5826 | 0.000999 |
| TCD8ex | MacHypo | LGALS3 | FCGR2B | 0.3824 | 0.7745 | 0.5784 | 0.000999 |
| TCD8ex | MacHypo | LGALS3 | TFRC | 0.3824 | 0.4916 | 0.437 | 0.000999 |
| TCD8ex | MacHypo | LTB | ITGB1 | 0.4576 | 0.4597 | 0.4586 | 0.000999 |
| TCD8ex | MacHypo | MIF | CD74 | 0.749 | 3.1474 | 1.9482 | 0.000999 |
| TCD8ex | MacHypo | PTPRC | CD4 | 1.6762 | 0.2639 | 0.97 | 0.000999 |
| TCD8ex | MacHypo | PTPRC | CD44 | 1.6762 | 1.5822 | 1.6292 | 0.000999 |
| TCD8ex | MacHypo | PTPRC | IFNAR1 | 1.6762 | 0.3706 | 1.0234 | 0.000999 |
| TCD8ex | MacHypo | RPS19 | C5AR1 | 3.8353 | 0.4508 | 2.143 | 0.000999 |
| TCD8ex | MacHypo | RPS19 | CD4 | 3.8353 | 0.2639 | 2.0496 | 0.000999 |
| TCD8ex | MacHypo | RPS19 | ICAM1 | 3.8353 | 0.3685 | 2.1019 | 0.000999 |
| TCD8ex | MacHypo | RTN4 | CD44 | 0.3033 | 1.5822 | 0.9427 | 0.000999 |
| TCD8ex | MacHypo | SPP1 | CD44 | 0.2978 | 1.5822 | 0.94 | 0.000999 |
| TCD8ex | MacHypo | VCAM1 | ITGB1 | 0.4462 | 0.4597 | 0.4529 | 0.000999 |
| TCD8ex | MacHypo | VCAM1 | ITGB2 | 0.4462 | 1.0357 | 0.741 | 0.000999 |
| TCD8ex | MacHypo | VCAM1 | TFRC | 0.4462 | 0.4916 | 0.4689 | 0.000999 |
| TGD | MacHypo | ARF1 | CD4 | 0.9453 | 0.2639 | 0.6046 | 0.000999 |
| TGD | MacHypo | ARF1 | CD44 | 0.9453 | 1.5822 | 1.2637 | 0.000999 |
| TGD | MacHypo | CALM1 | TNFRSF1B | 1.7587 | 0.4397 | 1.0992 | 0.000999 |
| TGD | MacHypo | CD2 | CD4 | 1.4522 | 0.2639 | 0.8581 | 0.000999 |
| TGD | MacHypo | CD2 | PTPRC | 1.4522 | 0.4793 | 0.9658 | 0.000999 |
| TGD | MacHypo | CD6 | ALCAM | 0.4684 | 0.2569 | 0.3627 | 0.000999 |
| TGD | MacHypo | CD99 | CD99 | 1.7154 | 0.765 | 1.2402 | 0.000999 |
| TGD | MacHypo | HMGB1 | AGTRAP | 1.7525 | 0.263 | 1.0077 | 0.000999 |
| TGD | MacHypo | HMGB1 | LY96 | 1.7525 | 0.7779 | 1.2652 | 0.000999 |

| | | | | | | | |
|---|---|---|---|---|---|---|---|
| TGD | MacHypo | HMGB1 | TLR2 | 1.7525 | 0.2562 | 1.0043 | 0.000999 |
| TGD | MacHypo | ICAM3 | ITGB2 | 0.8816 | 1.0357 | 0.9586 | 0.000999 |
| TGD | MacHypo | IL16 | CD4 | 0.9357 | 0.2639 | 0.5998 | 0.000999 |
| TGD | MacHypo | ITGA4 | CD44 | 0.3968 | 1.5822 | 0.9895 | 0.000999 |
| TGD | MacHypo | ITGA4 | CXCR4 | 0.3968 | 1.1308 | 0.7638 | 0.000999 |
| TGD | MacHypo | ITGA4 | TFRC | 0.3968 | 0.4916 | 0.4442 | 0.000999 |
| TGD | MacHypo | ITGAL | ICAM1 | 0.3731 | 0.3685 | 0.3708 | 0.000999 |
| TGD | MacHypo | ITGB1 | PLAUR | 0.3657 | 1.5015 | 0.9336 | 0.000999 |
| TGD | MacHypo | ITGB2 | ICAM1 | 1.4444 | 0.3685 | 0.9065 | 0.000999 |
| TGD | MacHypo | ITGB2 | ITGAM | 1.4444 | 0.328 | 0.8862 | 0.000999 |
| TGD | MacHypo | ITGB2 | ITGAX | 1.4444 | 0.3626 | 0.9035 | 0.000999 |
| TGD | MacHypo | ITGB2 | PLAUR | 1.4444 | 1.5015 | 1.473 | 0.000999 |
| TGD | MacHypo | LTB | ITGB1 | 1.6829 | 0.4597 | 1.0713 | 0.000999 |
| TGD | MacHypo | PTPRC | CD4 | 1.8653 | 0.2639 | 1.0646 | 0.000999 |
| TGD | MacHypo | PTPRC | CD44 | 1.8653 | 1.5822 | 1.7238 | 0.000999 |
| TGD | MacHypo | PTPRC | CXCR4 | 1.8653 | 1.1308 | 1.4981 | 0.000999 |
| TGD | MacHypo | PTPRC | IFNAR1 | 1.8653 | 0.3706 | 1.118 | 0.000999 |
| TGD | MacHypo | RPS19 | C5AR1 | 3.8902 | 0.4508 | 2.1705 | 0.000999 |
| TGD | MacHypo | RPS19 | CD4 | 3.8902 | 0.2639 | 2.077 | 0.000999 |
| TGD | MacHypo | RPS19 | ICAM1 | 3.8902 | 0.3685 | 2.1294 | 0.000999 |
| TGD | MacHypo | SELL | CD4 | 0.3555 | 0.2639 | 0.3097 | 0.000999 |
| TGD | MacHypo | TGFB1 | FCN1 | 0.7703 | 0.3143 | 0.5423 | 0.000999 |
| TGD | MacHypo | TGFB1 | ITGB1 | 0.7703 | 0.4597 | 0.615 | 0.000999 |
| TGD | MacHypo | TGFB1 | SDC2 | 0.7703 | 0.6066 | 0.6885 | 0.000999 |
| TGD | MacHypo | TGFB1 | TLR2 | 0.7703 | 0.2562 | 0.5133 | 0.000999 |
| TGD | MacHypo | COL1A1 | CD44 | 0.4518 | 1.5822 | 1.017 | 0.002 |
| TGD | MacHypo | CALM1 | RPSA | 1.7587 | 1.9137 | 1.8362 | 0.004 |
| TGD | MacHypo | CALR | CD44 | 0.9627 | 1.5822 | 1.2724 | 0.00799 |

| Supplementary Table 8: Communication results between macrophages and lymphocytes subpopulations, and malignant cells | | | | | | | |
|---|---|---|---|---|---|---|---|
| **MacHypo vs. malignant cells** | | | | | | | |
| celltype_01 | celltype_02 | ligand | receptor | ligand_mean_expression | receptor_mean_expression | ligand_receptor_mean_expression | pvalue |
| MacHypo | breasttumor | AIMP1 | RACK1 | 0.3406 | 2.6334 | 1.487 | 0.000999 |
| MacHypo | breasttumor | AIMP1 | RRBP1 | 0.3406 | 0.2835 | 0.312 | 0.000999 |
| MacHypo | breasttumor | ANXA1 | CD44 | 1.6213 | 0.3247 | 0.973 | 0.000999 |
| MacHypo | breasttumor | APOE | SCARB1 | 2.2471 | 0.2628 | 1.2549 | 0.000999 |
| MacHypo | breasttumor | APOE | SDC2 | 2.2471 | 1.071 | 1.659 | 0.000999 |
| MacHypo | breasttumor | CALM1 | MYLK | 1.2535 | 0.4064 | 0.8299 | 0.000999 |
| MacHypo | breasttumor | CALM1 | RPSA | 1.2535 | 2.358 | 1.8057 | 0.000999 |
| MacHypo | breasttumor | CALM1 | TNFRSF1A | 1.2535 | 0.2536 | 0.7535 | 0.000999 |
| MacHypo | breasttumor | CALM2 | MYLK | 1.3534 | 0.4064 | 0.8799 | 0.000999 |
| MacHypo | breasttumor | CALM3 | MYLK | 0.7395 | 0.4064 | 0.5729 | 0.000999 |
| MacHypo | breasttumor | CD99 | CD99 | 0.765 | 0.3871 | 0.576 | 0.000999 |
| MacHypo | breasttumor | COL1A1 | ITGB1 | 0.3651 | 0.2939 | 0.3295 | 0.000999 |
| MacHypo | breasttumor | FN1 | CD44 | 1.342 | 0.3247 | 0.8334 | 0.000999 |
| MacHypo | breasttumor | FN1 | ITGB1 | 1.342 | 0.2939 | 0.8179 | 0.000999 |
| MacHypo | breasttumor | FN1 | RPSA | 1.342 | 2.358 | 1.85 | 0.000999 |
| MacHypo | breasttumor | FN1 | SDC2 | 1.342 | 1.071 | 1.2065 | 0.000999 |
| MacHypo | breasttumor | GRN | TNFRSF1A | 1.0708 | 0.2536 | 0.6622 | 0.000999 |
| MacHypo | breasttumor | HLA-DRA | SCARB1 | 3.2011 | 0.2628 | 1.7319 | 0.000999 |
| MacHypo | breasttumor | ITGB1 | CD151 | 0.4596 | 0.3569 | 0.4083 | 0.000999 |
| MacHypo | breasttumor | ITGB1 | RACK1 | 0.4596 | 2.6334 | 1.5465 | 0.000999 |
| MacHypo | breasttumor | LGALS3 | ITGB1 | 1.6689 | 0.2939 | 0.9814 | 0.000999 |
| MacHypo | breasttumor | MMP9 | CD44 | 0.8086 | 0.3247 | 0.5667 | 0.000999 |
| MacHypo | breasttumor | MMP9 | ITGB1 | 0.8086 | 0.2939 | 0.5512 | 0.000999 |
| MacHypo | breasttumor | RTN4 | CD99 | 0.9167 | 0.3871 | 0.6519 | 0.000999 |
| MacHypo | breasttumor | SERPINA1 | SDC2 | 0.9051 | 1.071 | 0.9881 | 0.000999 |
| MacHypo | breasttumor | SPP1 | CD44 | 4.3572 | 0.3247 | 2.341 | 0.000999 |
| MacHypo | breasttumor | SPP1 | ITGB1 | 4.3572 | 0.2939 | 2.3255 | 0.000999 |

| | | | | | | | |
|---|---|---|---|---|---|---|---|
| MacHypo | breasttumor | TNFSF13B | ITGB1 | 0.3932 | 0.2939 | 0.3435 | 0.000999 |
| MacHypo | colorectaltumor | AIMP1 | RACK1 | 0.3406 | 2.7146 | 1.5276 | 0.000999 |
| MacHypo | colorectaltumor | APOE | SDC1 | 2.2471 | 0.2575 | 1.2523 | 0.000999 |
| MacHypo | colorectaltumor | APP | SDC1 | 0.2783 | 0.2575 | 0.2679 | 0.000999 |
| MacHypo | colorectaltumor | CALM1 | RPSA | 1.2535 | 2.251 | 1.7523 | 0.000999 |
| MacHypo | colorectaltumor | COL1A1 | SDC1 | 0.3651 | 0.2575 | 0.3113 | 0.000999 |
| MacHypo | colorectaltumor | FN1 | ITGB1 | 1.342 | 0.4214 | 0.8817 | 0.000999 |
| MacHypo | colorectaltumor | FN1 | RPSA | 1.342 | 2.251 | 1.7965 | 0.000999 |
| MacHypo | colorectaltumor | FN1 | SDC1 | 1.342 | 0.2575 | 0.7997 | 0.000999 |
| MacHypo | colorectaltumor | ITGB1 | CD151 | 0.4596 | 0.6439 | 0.5518 | 0.000999 |
| MacHypo | colorectaltumor | ITGB1 | CDH17 | 0.4596 | 0.254 | 0.3568 | 0.000999 |
| MacHypo | colorectaltumor | ITGB1 | RACK1 | 0.4596 | 2.7146 | 1.5871 | 0.000999 |
| MacHypo | colorectaltumor | ITGB1 | SDC1 | 0.4596 | 0.2575 | 0.3586 | 0.000999 |
| MacHypo | colorectaltumor | LGALS3 | ITGB4 | 1.6689 | 0.2913 | 0.9801 | 0.000999 |
| MacHypo | colorectaltumor | MMP9 | ITGB1 | 0.8086 | 0.4214 | 0.615 | 0.000999 |
| MacHypo | colorectaltumor | SPP1 | ITGB1 | 4.3572 | 0.4214 | 2.3893 | 0.000999 |
| MacHypo | colorectaltumor | SPP1 | SDC1 | 4.3572 | 0.2575 | 2.3073 | 0.000999 |
| MacHypo | colorectaltumor | ITGB1 | CD46 | 0.4596 | 0.3583 | 0.409 | 0.002 |
| MacHypo | colorectaltumor | SPP1 | CD44 | 4.3572 | 0.2512 | 2.3042 | 0.002 |
| MacHypo | livertumor | ADM | RAMP1 | 0.9373 | 0.4499 | 0.6936 | 0.000999 |
| MacHypo | livertumor | AIMP1 | RACK1 | 0.3406 | 2.7655 | 1.5531 | 0.000999 |
| MacHypo | livertumor | AIMP1 | RRBP1 | 0.3406 | 0.355 | 0.3478 | 0.000999 |
| MacHypo | livertumor | ANXA1 | MET | 1.6213 | 0.2983 | 0.9598 | 0.000999 |
| MacHypo | livertumor | ANXA1 | PTPRF | 1.6213 | 0.384 | 1.0027 | 0.000999 |
| MacHypo | livertumor | APOE | SDC1 | 2.2471 | 0.5354 | 1.3913 | 0.000999 |
| MacHypo | livertumor | APOE | SDC4 | 2.2471 | 0.6068 | 1.427 | 0.000999 |
| MacHypo | livertumor | APP | EPHB4 | 0.2783 | 0.4535 | 0.3659 | 0.000999 |
| MacHypo | livertumor | APP | ITGA6 | 0.2783 | 0.5211 | 0.3997 | 0.000999 |
| MacHypo | livertumor | APP | SDC1 | 0.2783 | 0.5354 | 0.4069 | 0.000999 |
| MacHypo | livertumor | AREG | ERBB3 | 0.3573 | 0.2644 | 0.3108 | 0.000999 |

| | | | | | | | |
|---|---|---|---|---|---|---|---|
| MacHypo | livertumor | ARF1 | INSR | 0.6073 | 0.3039 | 0.4556 | 0.000999 |
| MacHypo | livertumor | CALM1 | INSR | 1.2535 | 0.3039 | 0.7787 | 0.000999 |
| MacHypo | livertumor | CALM1 | RPSA | 1.2535 | 2.6317 | 1.9426 | 0.000999 |
| MacHypo | livertumor | CALR | ITGA3 | 0.9193 | 0.3913 | 0.6553 | 0.000999 |
| MacHypo | livertumor | CD99 | EPHB4 | 0.765 | 0.4535 | 0.6092 | 0.000999 |
| MacHypo | livertumor | CD99 | INSR | 0.765 | 0.3039 | 0.5344 | 0.000999 |
| MacHypo | livertumor | COL1A1 | ITGA2 | 0.3651 | 0.3983 | 0.3817 | 0.000999 |
| MacHypo | livertumor | COL1A1 | ITGA3 | 0.3651 | 0.3913 | 0.3782 | 0.000999 |
| MacHypo | livertumor | COL1A1 | ITGB1 | 0.3651 | 1.6044 | 0.9848 | 0.000999 |
| MacHypo | livertumor | COL1A1 | SDC1 | 0.3651 | 0.5354 | 0.4503 | 0.000999 |
| MacHypo | livertumor | FN1 | ITGA2 | 1.342 | 0.3983 | 0.8701 | 0.000999 |
| MacHypo | livertumor | FN1 | ITGA3 | 1.342 | 0.3913 | 0.8666 | 0.000999 |
| MacHypo | livertumor | FN1 | ITGB1 | 1.342 | 1.6044 | 1.4732 | 0.000999 |
| MacHypo | livertumor | FN1 | RPSA | 1.342 | 2.6317 | 1.9868 | 0.000999 |
| MacHypo | livertumor | FN1 | SDC1 | 1.342 | 0.5354 | 0.9387 | 0.000999 |
| MacHypo | livertumor | FN1 | TFRC | 1.342 | 0.2899 | 0.8159 | 0.000999 |
| MacHypo | livertumor | HAMP | SLC40A1 | 0.2525 | 0.4972 | 0.3749 | 0.000999 |
| MacHypo | livertumor | HBEGF | ERBB3 | 0.2764 | 0.2644 | 0.2704 | 0.000999 |
| MacHypo | livertumor | HLA-A | ERBB3 | 2.5035 | 0.2644 | 1.3839 | 0.000999 |
| MacHypo | livertumor | HLA-DRA | EPHB4 | 3.2011 | 0.4535 | 1.8273 | 0.000999 |
| MacHypo | livertumor | HLA-E | EPHB4 | 1.1723 | 0.4535 | 0.8129 | 0.000999 |
| MacHypo | livertumor | HLA-E | ITGA6 | 1.1723 | 0.5211 | 0.8467 | 0.000999 |
| MacHypo | livertumor | HLA-E | PLXNB2 | 1.1723 | 0.265 | 0.7186 | 0.000999 |
| MacHypo | livertumor | HMGB2 | CDH1 | 0.4281 | 0.4896 | 0.4589 | 0.000999 |
| MacHypo | livertumor | HMGB2 | DSG2 | 0.4281 | 0.7479 | 0.588 | 0.000999 |
| MacHypo | livertumor | HSP90B1 | ITGB1 | 0.8572 | 1.6044 | 1.2308 | 0.000999 |
| MacHypo | livertumor | ITGB1 | CD151 | 0.4596 | 1.1085 | 0.7841 | 0.000999 |
| MacHypo | livertumor | ITGB1 | VTN | 0.4596 | 0.7399 | 0.5998 | 0.000999 |
| MacHypo | livertumor | LGALS3 | ITGA2 | 1.6689 | 0.3983 | 1.0336 | 0.000999 |
| MacHypo | livertumor | LGALS3 | ITGB1 | 1.6689 | 1.6044 | 1.6367 | 0.000999 |

| | | | | | | | |
|---|---|---|---|---|---|---|---|
| MacHypo | livertumor | LGALS3 | ITGB4 | 1.6689 | 0.5518 | 1.1103 | 0.000999 |
| MacHypo | livertumor | LGALS3 | TFRC | 1.6689 | 0.2899 | 0.9794 | 0.000999 |
| MacHypo | livertumor | MMP9 | CDH1 | 0.8086 | 0.4896 | 0.6491 | 0.000999 |
| MacHypo | livertumor | MMP9 | ITGB1 | 0.8086 | 1.6044 | 1.2065 | 0.000999 |
| MacHypo | livertumor | NAMPT | INSR | 0.8899 | 0.3039 | 0.5969 | 0.000999 |
| MacHypo | livertumor | PTPRC | INSR | 0.4793 | 0.3039 | 0.3916 | 0.000999 |
| MacHypo | livertumor | PTPRC | MET | 0.4793 | 0.2983 | 0.3888 | 0.000999 |
| MacHypo | livertumor | RPS19 | RPSA | 3.5437 | 2.6317 | 3.0877 | 0.000999 |
| MacHypo | livertumor | SPP1 | CD44 | 4.3572 | 0.8016 | 2.5794 | 0.000999 |
| MacHypo | livertumor | SPP1 | ITGB1 | 4.3572 | 1.6044 | 2.9808 | 0.000999 |
| MacHypo | livertumor | SPP1 | SDC1 | 4.3572 | 0.5354 | 2.4463 | 0.000999 |
| MacHypo | livertumor | TNFSF13B | ITGB1 | 0.3932 | 1.6044 | 0.9988 | 0.000999 |
| MacHypo | livertumor | VEGFA | AXL | 0.5813 | 0.3099 | 0.4456 | 0.000999 |
| MacHypo | livertumor | VEGFA | VTN | 0.5813 | 0.7399 | 0.6606 | 0.000999 |
| MacHypo | lungtumor | APOE | SDC1 | 2.2471 | 0.2715 | 1.2593 | 0.000999 |
| MacHypo | lungtumor | APOE | SDC4 | 2.2471 | 0.3516 | 1.2993 | 0.000999 |
| MacHypo | lungtumor | APP | SDC1 | 0.2783 | 0.2715 | 0.2749 | 0.000999 |
| MacHypo | lungtumor | CALM1 | RPSA | 1.2535 | 1.6057 | 1.4296 | 0.000999 |
| MacHypo | lungtumor | CCL20 | CXCR4 | 0.6366 | 0.2684 | 0.4525 | 0.000999 |
| MacHypo | lungtumor | CD99 | CD99 | 0.765 | 0.5938 | 0.6794 | 0.000999 |
| MacHypo | lungtumor | COL1A1 | SDC1 | 0.3651 | 0.2715 | 0.3183 | 0.000999 |
| MacHypo | lungtumor | CXCL16 | CXCR4 | 0.6913 | 0.2684 | 0.4799 | 0.000999 |
| MacHypo | lungtumor | CXCL2 | CXCR4 | 0.6881 | 0.2684 | 0.4782 | 0.000999 |
| MacHypo | lungtumor | CXCL3 | CXCR4 | 0.7215 | 0.2684 | 0.495 | 0.000999 |
| MacHypo | lungtumor | FN1 | CD44 | 1.342 | 0.3802 | 0.8611 | 0.000999 |
| MacHypo | lungtumor | FN1 | RPSA | 1.342 | 1.6057 | 1.4738 | 0.000999 |
| MacHypo | lungtumor | FN1 | SDC1 | 1.342 | 0.2715 | 0.8067 | 0.000999 |
| MacHypo | lungtumor | ITGB1 | CD151 | 0.4596 | 0.6776 | 0.5686 | 0.000999 |
| MacHypo | lungtumor | ITGB1 | CD46 | 0.4596 | 0.434 | 0.4468 | 0.000999 |
| MacHypo | lungtumor | ITGB1 | CD47 | 0.4596 | 0.3255 | 0.3926 | 0.000999 |

| | | | | | | | |
|---|---|---|---|---|---|---|---|
| MacHypo | lungtumor | ITGB1 | RACK1 | 0.4596 | 2.2108 | 1.3352 | 0.000999 |
| MacHypo | lungtumor | ITGB1 | SDC1 | 0.4596 | 0.2715 | 0.3656 | 0.000999 |
| MacHypo | lungtumor | MMP9 | CD44 | 0.8086 | 0.3802 | 0.5944 | 0.000999 |
| MacHypo | lungtumor | PTPRC | CD44 | 0.4793 | 0.3802 | 0.4297 | 0.000999 |
| MacHypo | lungtumor | PTPRC | CXCR4 | 0.4793 | 0.2684 | 0.3738 | 0.000999 |
| MacHypo | lungtumor | RTN4 | CD44 | 0.9167 | 0.3802 | 0.6484 | 0.000999 |
| MacHypo | lungtumor | RTN4 | CD99 | 0.9167 | 0.5938 | 0.7552 | 0.000999 |
| MacHypo | lungtumor | SPP1 | CD44 | 4.3572 | 0.3802 | 2.3687 | 0.000999 |
| MacHypo | lungtumor | SPP1 | SDC1 | 4.3572 | 0.2715 | 2.3143 | 0.000999 |
| MacHypo | melanomatumo | AIMP1 | RACK1 | 0.3406 | 4.145 | 2.2428 | 0.000999 |
| MacHypo | melanomatumo | ANXA1 | EDNRB | 1.6213 | 0.5666 | 1.094 | 0.000999 |
| MacHypo | melanomatumo | APP | EDNRB | 0.2783 | 0.5666 | 0.4225 | 0.000999 |
| MacHypo | melanomatumo | CALM1 | RPSA | 1.2535 | 3.7787 | 2.5161 | 0.000999 |
| MacHypo | melanomatumo | FN1 | RPSA | 1.342 | 3.7787 | 2.5603 | 0.000999 |
| MacHypo | melanomatumo | ITGB1 | RACK1 | 0.4596 | 4.145 | 2.3023 | 0.000999 |
| MacHypo | melanomatumo | NAMPT | ANXA6 | 0.8899 | 0.3625 | 0.6262 | 0.000999 |
| MacHypo | melanomatumo | NMB | EDNRB | 0.356 | 0.5666 | 0.4613 | 0.000999 |
| MacHypo | melanomatumo | RPS19 | RPSA | 3.5437 | 3.7787 | 3.6612 | 0.000999 |
| MacHypo | melanomatumo | SPP1 | NCSTN | 4.3572 | 0.2614 | 2.3093 | 0.000999 |
| MacHypo | melanomatumo | CALR | NCSTN | 0.9193 | 0.2614 | 0.5904 | 0.00699 |
| MacHypo | ovarytumor | AIMP1 | RACK1 | 0.3406 | 2.9543 | 1.6474 | 0.000999 |
| MacHypo | ovarytumor | ANXA1 | CXCR4 | 1.6213 | 0.3688 | 0.9951 | 0.000999 |
| MacHypo | ovarytumor | AREG | FOLR1 | 0.3573 | 0.5619 | 0.4596 | 0.000999 |
| MacHypo | ovarytumor | CALM1 | RPSA | 1.2535 | 2.9542 | 2.1039 | 0.000999 |
| MacHypo | ovarytumor | CCL20 | CXCR4 | 0.6366 | 0.3688 | 0.5027 | 0.000999 |
| MacHypo | ovarytumor | CXCL16 | CXCR4 | 0.6913 | 0.3688 | 0.53 | 0.000999 |
| MacHypo | ovarytumor | CXCL2 | CXCR4 | 0.6881 | 0.3688 | 0.5284 | 0.000999 |
| MacHypo | ovarytumor | CXCL3 | CXCR4 | 0.7215 | 0.3688 | 0.5452 | 0.000999 |
| MacHypo | ovarytumor | FN1 | ITGB1 | 1.342 | 0.2783 | 0.8101 | 0.000999 |
| MacHypo | ovarytumor | FN1 | RPSA | 1.342 | 2.9542 | 2.1481 | 0.000999 |

| | | | | | | | |
|---|---|---|---|---|---|---|---|
| MacHypo | ovarytumor | HSP90B1 | ITGB1 | 0.8572 | 0.2783 | 0.5678 | 0.000999 |
| MacHypo | ovarytumor | ITGB1 | CD151 | 0.4596 | 0.4981 | 0.4789 | 0.000999 |
| MacHypo | ovarytumor | ITGB1 | CD47 | 0.4596 | 0.4931 | 0.4764 | 0.000999 |
| MacHypo | ovarytumor | ITGB1 | RACK1 | 0.4596 | 2.9543 | 1.707 | 0.000999 |
| MacHypo | ovarytumor | LGALS3 | ITGB1 | 1.6689 | 0.2783 | 0.9736 | 0.000999 |
| MacHypo | ovarytumor | MIF | CD74 | 1.9477 | 1.4431 | 1.6954 | 0.000999 |
| MacHypo | ovarytumor | MMP9 | ITGB1 | 0.8086 | 0.2783 | 0.5434 | 0.000999 |
| MacHypo | ovarytumor | PTPRC | CXCR4 | 0.4793 | 0.3688 | 0.424 | 0.000999 |
| MacHypo | ovarytumor | SPP1 | ITGB1 | 4.3572 | 0.2783 | 2.3177 | 0.000999 |
| MacHypo | ovarytumor | TNFSF13B | ITGB1 | 0.3932 | 0.2783 | 0.3357 | 0.000999 |
| breasttumor | MacHypo | A2M | TNFRSF14 | 0.3129 | 0.3382 | 0.3255 | 0.000999 |
| breasttumor | MacHypo | ANXA1 | CD44 | 0.4092 | 1.5821 | 0.9956 | 0.000999 |
| breasttumor | MacHypo | ANXA1 | CXCR4 | 0.4092 | 1.1308 | 0.77 | 0.000999 |
| breasttumor | MacHypo | APP | ALOX5 | 0.5409 | 0.2606 | 0.4008 | 0.000999 |
| breasttumor | MacHypo | APP | C3AR1 | 0.5409 | 0.2925 | 0.4167 | 0.000999 |
| breasttumor | MacHypo | APP | C5AR1 | 0.5409 | 0.4508 | 0.4959 | 0.000999 |
| breasttumor | MacHypo | APP | CCR1 | 0.5409 | 0.2563 | 0.3986 | 0.000999 |
| breasttumor | MacHypo | APP | CXCR4 | 0.5409 | 1.1308 | 0.8359 | 0.000999 |
| breasttumor | MacHypo | APP | TNFRSF14 | 0.5409 | 0.3382 | 0.4396 | 0.000999 |
| breasttumor | MacHypo | ARF1 | CD4 | 1.6219 | 0.2639 | 0.9429 | 0.000999 |
| breasttumor | MacHypo | ARF1 | CD44 | 1.6219 | 1.5821 | 1.602 | 0.000999 |
| breasttumor | MacHypo | CALM1 | TNFRSF1B | 0.9015 | 0.4397 | 0.6706 | 0.000999 |
| breasttumor | MacHypo | CALR | CD44 | 1.1835 | 1.5821 | 1.3828 | 0.000999 |
| breasttumor | MacHypo | CALR | IL7R | 1.1835 | 0.3898 | 0.7867 | 0.000999 |
| breasttumor | MacHypo | CD99 | CD99 | 0.3871 | 0.765 | 0.576 | 0.000999 |
| breasttumor | MacHypo | COL2A1 | CD44 | 0.3935 | 1.5821 | 0.9878 | 0.000999 |
| breasttumor | MacHypo | COL2A1 | ITGB1 | 0.3935 | 0.4596 | 0.4266 | 0.000999 |
| breasttumor | MacHypo | GSTP1 | CD44 | 2.5435 | 1.5821 | 2.0628 | 0.000999 |
| breasttumor | MacHypo | GSTP1 | IL7R | 2.5435 | 0.3898 | 1.4667 | 0.000999 |
| breasttumor | MacHypo | HEBP1 | C3AR1 | 0.4062 | 0.2925 | 0.3493 | 0.000999 |

| | | | | | | | |
|---|---|---|---|---|---|---|---|
| breasttumor | MacHypo | HEBP1 | C5AR1 | 0.4062 | 0.4508 | 0.4285 | 0.000999 |
| breasttumor | MacHypo | HEBP1 | CCR1 | 0.4062 | 0.2563 | 0.3313 | 0.000999 |
| breasttumor | MacHypo | HEBP1 | CXCR4 | 0.4062 | 1.1308 | 0.7685 | 0.000999 |
| breasttumor | MacHypo | HEBP1 | FPR3 | 0.4062 | 0.3793 | 0.3927 | 0.000999 |
| breasttumor | MacHypo | HMGB1 | AGTRAP | 1.1815 | 0.263 | 0.7222 | 0.000999 |
| breasttumor | MacHypo | HMGB1 | LY96 | 1.1815 | 0.7778 | 0.9796 | 0.000999 |
| breasttumor | MacHypo | HMGB1 | TLR2 | 1.1815 | 0.2562 | 0.7188 | 0.000999 |
| breasttumor | MacHypo | HMGB2 | CD44 | 0.3139 | 1.5821 | 0.948 | 0.000999 |
| breasttumor | MacHypo | HMGB2 | DAB2 | 0.3139 | 0.6138 | 0.4639 | 0.000999 |
| breasttumor | MacHypo | HSP90B1 | TNFRSF1B | 0.7786 | 0.4397 | 0.6091 | 0.000999 |
| breasttumor | MacHypo | ITGB1 | CD47 | 0.2939 | 0.3564 | 0.3251 | 0.000999 |
| breasttumor | MacHypo | ITGB1 | DAB2 | 0.2939 | 0.6138 | 0.4538 | 0.000999 |
| breasttumor | MacHypo | ITGB1 | ITGAX | 0.2939 | 0.3625 | 0.3282 | 0.000999 |
| breasttumor | MacHypo | ITGB1 | PLAUR | 0.2939 | 1.5014 | 0.8976 | 0.000999 |
| breasttumor | MacHypo | LGALS3 | FCGR2A | 0.8391 | 0.7827 | 0.8109 | 0.000999 |
| breasttumor | MacHypo | LGALS3 | FCGR2B | 0.8391 | 0.7744 | 0.8068 | 0.000999 |
| breasttumor | MacHypo | MIF | CD74 | 3.2716 | 3.1473 | 3.2094 | 0.000999 |
| breasttumor | MacHypo | MIF | TNFRSF14 | 3.2716 | 0.3382 | 1.8049 | 0.000999 |
| breasttumor | MacHypo | NPW | C3AR1 | 0.5478 | 0.2925 | 0.4201 | 0.000999 |
| breasttumor | MacHypo | NPW | C5AR1 | 0.5478 | 0.4508 | 0.4993 | 0.000999 |
| breasttumor | MacHypo | NPW | CCR1 | 0.5478 | 0.2563 | 0.4021 | 0.000999 |
| breasttumor | MacHypo | NPW | CXCR4 | 0.5478 | 1.1308 | 0.8393 | 0.000999 |
| breasttumor | MacHypo | RPS19 | C5AR1 | 3.691 | 0.4508 | 2.0709 | 0.000999 |
| breasttumor | MacHypo | RPS19 | CD4 | 3.691 | 0.2639 | 1.9774 | 0.000999 |
| breasttumor | MacHypo | RPS19 | ICAM1 | 3.691 | 0.3685 | 2.0297 | 0.000999 |
| breasttumor | MacHypo | RTN4 | CD44 | 0.3318 | 1.5821 | 0.957 | 0.000999 |
| breasttumor | MacHypo | TF | CD4 | 0.2688 | 0.2639 | 0.2663 | 0.000999 |
| breasttumor | MacHypo | TF | IL7R | 0.2688 | 0.3898 | 0.3293 | 0.000999 |
| breasttumor | MacHypo | TF | TFRC | 0.2688 | 0.4916 | 0.3802 | 0.000999 |
| breasttumor | MacHypo | HSP90B1 | ITGB1 | 0.7786 | 0.4596 | 0.6191 | 0.002 |

| | | | | | | | |
|---|---|---|---|---|---|---|---|
| colorectaltumor | MacHypo | APP | ALOX5 | 0.4966 | 0.2606 | 0.3786 | 0.000999 |
| colorectaltumor | MacHypo | APP | C3AR1 | 0.4966 | 0.2925 | 0.3945 | 0.000999 |
| colorectaltumor | MacHypo | APP | C5AR1 | 0.4966 | 0.4508 | 0.4737 | 0.000999 |
| colorectaltumor | MacHypo | APP | CCR1 | 0.4966 | 0.2563 | 0.3765 | 0.000999 |
| colorectaltumor | MacHypo | APP | CXCR4 | 0.4966 | 1.1308 | 0.8137 | 0.000999 |
| colorectaltumor | MacHypo | APP | TNFRSF14 | 0.4966 | 0.3382 | 0.4174 | 0.000999 |
| colorectaltumor | MacHypo | ARF1 | CD44 | 0.6489 | 1.5821 | 1.1155 | 0.000999 |
| colorectaltumor | MacHypo | COL1A1 | CD44 | 0.4875 | 1.5821 | 1.0348 | 0.000999 |
| colorectaltumor | MacHypo | GSTP1 | CD44 | 2.8039 | 1.5821 | 2.193 | 0.000999 |
| colorectaltumor | MacHypo | GSTP1 | IL7R | 2.8039 | 0.3898 | 1.5969 | 0.000999 |
| colorectaltumor | MacHypo | HMGB1 | AGTRAP | 2.155 | 0.263 | 1.209 | 0.000999 |
| colorectaltumor | MacHypo | HMGB1 | LY96 | 2.155 | 0.7778 | 1.4664 | 0.000999 |
| colorectaltumor | MacHypo | HMGB1 | TLR2 | 2.155 | 0.2562 | 1.2056 | 0.000999 |
| colorectaltumor | MacHypo | HMGB2 | CD44 | 0.5347 | 1.5821 | 1.0584 | 0.000999 |
| colorectaltumor | MacHypo | HMGB2 | DAB2 | 0.5347 | 0.6138 | 0.5743 | 0.000999 |
| colorectaltumor | MacHypo | ITGB1 | PLAUR | 0.4214 | 1.5014 | 0.9614 | 0.000999 |
| colorectaltumor | MacHypo | MIF | CD74 | 1.6741 | 3.1473 | 2.4107 | 0.000999 |
| colorectaltumor | MacHypo | RPS19 | C5AR1 | 4.0956 | 0.4508 | 2.2732 | 0.000999 |
| colorectaltumor | MacHypo | RPS19 | CD4 | 4.0956 | 0.2639 | 2.1797 | 0.000999 |
| colorectaltumor | MacHypo | RPS19 | ICAM1 | 4.0956 | 0.3685 | 2.232 | 0.000999 |
| colorectaltumor | MacHypo | RPS19 | RPSA | 4.0956 | 1.9136 | 3.0046 | 0.000999 |
| colorectaltumor | MacHypo | ITGB1 | DAB2 | 0.4214 | 0.6138 | 0.5176 | 0.004 |
| livertumor | MacHypo | AGT | ANPEP | 0.4701 | 0.3075 | 0.3888 | 0.000999 |
| livertumor | MacHypo | AGT | ATP6AP2 | 0.4701 | 0.6634 | 0.5667 | 0.000999 |
| livertumor | MacHypo | AGT | C3AR1 | 0.4701 | 0.2925 | 0.3813 | 0.000999 |
| livertumor | MacHypo | AGT | C5AR1 | 0.4701 | 0.4508 | 0.4604 | 0.000999 |
| livertumor | MacHypo | AGT | CCR1 | 0.4701 | 0.2563 | 0.3632 | 0.000999 |
| livertumor | MacHypo | AGT | CXCR4 | 0.4701 | 1.1308 | 0.8004 | 0.000999 |
| livertumor | MacHypo | APOA1 | ABCA1 | 0.2888 | 0.4872 | 0.388 | 0.000999 |
| livertumor | MacHypo | APOA1 | IL7R | 0.2888 | 0.3898 | 0.3393 | 0.000999 |

| | | | | | | | |
|---|---|---|---|---|---|---|---|
| livertumor | MacHypo | APOA1 | MSR1 | 0.2888 | 0.4769 | 0.3828 | 0.000999 |
| livertumor | MacHypo | APP | ALOX5 | 0.5626 | 0.2606 | 0.4116 | 0.000999 |
| livertumor | MacHypo | APP | C3AR1 | 0.5626 | 0.2925 | 0.4275 | 0.000999 |
| livertumor | MacHypo | APP | C5AR1 | 0.5626 | 0.4508 | 0.5067 | 0.000999 |
| livertumor | MacHypo | APP | CCR1 | 0.5626 | 0.2563 | 0.4094 | 0.000999 |
| livertumor | MacHypo | APP | CXCR4 | 0.5626 | 1.1308 | 0.8467 | 0.000999 |
| livertumor | MacHypo | APP | TNFRSF14 | 0.5626 | 0.3382 | 0.4504 | 0.000999 |
| livertumor | MacHypo | AREG | ICAM1 | 0.448 | 0.3685 | 0.4082 | 0.000999 |
| livertumor | MacHypo | ARF1 | CD4 | 0.6936 | 0.2639 | 0.4787 | 0.000999 |
| livertumor | MacHypo | ARF1 | CD44 | 0.6936 | 1.5821 | 1.1378 | 0.000999 |
| livertumor | MacHypo | C3 | C3AR1 | 0.6584 | 0.2925 | 0.4755 | 0.000999 |
| livertumor | MacHypo | C3 | C5AR1 | 0.6584 | 0.4508 | 0.5546 | 0.000999 |
| livertumor | MacHypo | C3 | CCR1 | 0.6584 | 0.2563 | 0.4574 | 0.000999 |
| livertumor | MacHypo | C3 | CXCR4 | 0.6584 | 1.1308 | 0.8946 | 0.000999 |
| livertumor | MacHypo | C3 | ITGAM | 0.6584 | 0.328 | 0.4932 | 0.000999 |
| livertumor | MacHypo | C3 | ITGAX | 0.6584 | 0.3625 | 0.5105 | 0.000999 |
| livertumor | MacHypo | C3 | ITGB2 | 0.6584 | 1.0356 | 0.847 | 0.000999 |
| livertumor | MacHypo | CALR | CD44 | 1.0416 | 1.5821 | 1.3119 | 0.000999 |
| livertumor | MacHypo | CALR | IL7R | 1.0416 | 0.3898 | 0.7157 | 0.000999 |
| livertumor | MacHypo | CCL20 | C3AR1 | 1.0877 | 0.2925 | 0.6901 | 0.000999 |
| livertumor | MacHypo | CCL20 | C5AR1 | 1.0877 | 0.4508 | 0.7692 | 0.000999 |
| livertumor | MacHypo | CCL20 | CCR1 | 1.0877 | 0.2563 | 0.672 | 0.000999 |
| livertumor | MacHypo | CCL20 | CXCR4 | 1.0877 | 1.1308 | 1.1092 | 0.000999 |
| livertumor | MacHypo | CDH1 | RACK1 | 0.4896 | 2.3023 | 1.396 | 0.000999 |
| livertumor | MacHypo | CFH | ITGAM | 0.3373 | 0.328 | 0.3327 | 0.000999 |
| livertumor | MacHypo | DSG2 | CD44 | 0.7479 | 1.5821 | 1.165 | 0.000999 |
| livertumor | MacHypo | DSG2 | CXCR4 | 0.7479 | 1.1308 | 0.9393 | 0.000999 |
| livertumor | MacHypo | DSG2 | DAB2 | 0.7479 | 0.6138 | 0.6809 | 0.000999 |
| livertumor | MacHypo | FGA | ITGAX | 0.2688 | 0.3625 | 0.3157 | 0.000999 |
| livertumor | MacHypo | FGA | ITGB2 | 0.2688 | 1.0356 | 0.6522 | 0.000999 |

| | | | | | | | |
|---|---|---|---|---|---|---|---|
| livertumor | MacHypo | GSTP1 | CD44 | 1.7219 | 1.5821 | 1.652 | 0.000999 |
| livertumor | MacHypo | GSTP1 | IL7R | 1.7219 | 0.3898 | 1.0559 | 0.000999 |
| livertumor | MacHypo | HMGB1 | AGTRAP | 1.4361 | 0.263 | 0.8495 | 0.000999 |
| livertumor | MacHypo | HMGB1 | LY96 | 1.4361 | 0.7778 | 1.1069 | 0.000999 |
| livertumor | MacHypo | HMGB1 | TLR2 | 1.4361 | 0.2562 | 0.8461 | 0.000999 |
| livertumor | MacHypo | HSP90B1 | TNFRSF1B | 0.8951 | 0.4397 | 0.6674 | 0.000999 |
| livertumor | MacHypo | ITGB1 | CD151 | 1.6044 | 0.3292 | 0.9668 | 0.000999 |
| livertumor | MacHypo | ITGB1 | CD47 | 1.6044 | 0.3564 | 0.9804 | 0.000999 |
| livertumor | MacHypo | ITGB1 | DAB2 | 1.6044 | 0.6138 | 1.1091 | 0.000999 |
| livertumor | MacHypo | ITGB1 | ITGAX | 1.6044 | 0.3625 | 0.9835 | 0.000999 |
| livertumor | MacHypo | ITGB1 | PLAUR | 1.6044 | 1.5014 | 1.5529 | 0.000999 |
| livertumor | MacHypo | ITGB1 | RACK1 | 1.6044 | 2.3023 | 1.9534 | 0.000999 |
| livertumor | MacHypo | MIF | CD74 | 1.7638 | 3.1473 | 2.4555 | 0.000999 |
| livertumor | MacHypo | PTPRF | RACK1 | 0.384 | 2.3023 | 1.3432 | 0.000999 |
| livertumor | MacHypo | RPS19 | C5AR1 | 4.0772 | 0.4508 | 2.264 | 0.000999 |
| livertumor | MacHypo | RPS19 | CD4 | 4.0772 | 0.2639 | 2.1705 | 0.000999 |
| livertumor | MacHypo | RPS19 | ICAM1 | 4.0772 | 0.3685 | 2.2228 | 0.000999 |
| livertumor | MacHypo | RPS19 | RPSA | 4.0772 | 1.9136 | 2.9954 | 0.000999 |
| livertumor | MacHypo | RTN4 | CD44 | 1.0046 | 1.5821 | 1.2933 | 0.000999 |
| livertumor | MacHypo | SEMA3C | NRP2 | 0.2576 | 0.3119 | 0.2847 | 0.000999 |
| livertumor | MacHypo | SERPINA1 | SDC2 | 1.604 | 0.6065 | 1.1053 | 0.000999 |
| livertumor | MacHypo | TF | CD4 | 0.523 | 0.2639 | 0.3934 | 0.000999 |
| livertumor | MacHypo | TF | IL7R | 0.523 | 0.3898 | 0.4564 | 0.000999 |
| livertumor | MacHypo | TF | SDC2 | 0.523 | 0.6065 | 0.5648 | 0.000999 |
| livertumor | MacHypo | TF | TFRC | 0.523 | 0.4916 | 0.5073 | 0.000999 |
| livertumor | MacHypo | TTR | IL7R | 0.4579 | 0.3898 | 0.4239 | 0.000999 |
| livertumor | MacHypo | VTN | ITGB1 | 0.7399 | 0.4596 | 0.5998 | 0.000999 |
| livertumor | MacHypo | VTN | PLAUR | 0.7399 | 1.5014 | 1.1207 | 0.000999 |
| livertumor | MacHypo | VTN | SDC2 | 0.7399 | 0.6065 | 0.6732 | 0.000999 |
| livertumor | MacHypo | SEMA3C | NRP1 | 0.2576 | 0.2591 | 0.2584 | 0.002 |

| | | | | | | | |
|---|---|---|---|---|---|---|---|
| livertumor | MacHypo | CXCL2 | CXCR4 | 0.3364 | 1.1308 | 0.7336 | 0.004 |
| livertumor | MacHypo | LAMC2 | CD151 | 0.4847 | 0.3292 | 0.4069 | 0.004 |
| livertumor | MacHypo | RTN4 | CD99 | 1.0046 | 0.765 | 0.8848 | 0.005 |
| lungtumor | MacHypo | ANXA1 | C3AR1 | 1.4026 | 0.2925 | 0.8475 | 0.000999 |
| lungtumor | MacHypo | ANXA1 | C5AR1 | 1.4026 | 0.4508 | 0.9267 | 0.000999 |
| lungtumor | MacHypo | ANXA1 | CCR1 | 1.4026 | 0.2563 | 0.8295 | 0.000999 |
| lungtumor | MacHypo | ANXA1 | CD4 | 1.4026 | 0.2639 | 0.8333 | 0.000999 |
| lungtumor | MacHypo | ANXA1 | CD44 | 1.4026 | 1.5821 | 1.4924 | 0.000999 |
| lungtumor | MacHypo | ANXA1 | CXCR4 | 1.4026 | 1.1308 | 1.2667 | 0.000999 |
| lungtumor | MacHypo | ANXA1 | FPR3 | 1.4026 | 0.3793 | 0.891 | 0.000999 |
| lungtumor | MacHypo | AREG | ICAM1 | 0.3537 | 0.3685 | 0.3611 | 0.000999 |
| lungtumor | MacHypo | ARF1 | CD4 | 0.8791 | 0.2639 | 0.5715 | 0.000999 |
| lungtumor | MacHypo | ARF1 | CD44 | 0.8791 | 1.5821 | 1.2306 | 0.000999 |
| lungtumor | MacHypo | CALM1 | RPSA | 0.8426 | 1.9136 | 1.3781 | 0.000999 |
| lungtumor | MacHypo | CALM1 | TNFRSF1B | 0.8426 | 0.4397 | 0.6411 | 0.000999 |
| lungtumor | MacHypo | CALR | CD44 | 0.7795 | 1.5821 | 1.1808 | 0.000999 |
| lungtumor | MacHypo | CALR | IL7R | 0.7795 | 0.3898 | 0.5847 | 0.000999 |
| lungtumor | MacHypo | CCL20 | C3AR1 | 0.2885 | 0.2925 | 0.2905 | 0.000999 |
| lungtumor | MacHypo | CCL20 | C5AR1 | 0.2885 | 0.4508 | 0.3697 | 0.000999 |
| lungtumor | MacHypo | CCL20 | CCR1 | 0.2885 | 0.2563 | 0.2724 | 0.000999 |
| lungtumor | MacHypo | CCL20 | CXCR4 | 0.2885 | 1.1308 | 0.7097 | 0.000999 |
| lungtumor | MacHypo | CD99 | CD99 | 0.5938 | 0.765 | 0.6794 | 0.000999 |
| lungtumor | MacHypo | CXCL10 | C3AR1 | 0.3417 | 0.2925 | 0.3171 | 0.000999 |
| lungtumor | MacHypo | CXCL10 | C5AR1 | 0.3417 | 0.4508 | 0.3962 | 0.000999 |
| lungtumor | MacHypo | CXCL10 | CCR1 | 0.3417 | 0.2563 | 0.299 | 0.000999 |
| lungtumor | MacHypo | CXCL10 | CXCR4 | 0.3417 | 1.1308 | 0.7362 | 0.000999 |
| lungtumor | MacHypo | FGA | ITGAX | 0.4445 | 0.3625 | 0.4035 | 0.000999 |
| lungtumor | MacHypo | FGA | ITGB1 | 0.4445 | 0.4596 | 0.4521 | 0.000999 |
| lungtumor | MacHypo | FGA | ITGB2 | 0.4445 | 1.0356 | 0.7401 | 0.000999 |
| lungtumor | MacHypo | GRN | TNFRSF1B | 0.4026 | 0.4397 | 0.4211 | 0.000999 |

| | | | | | | | |
|---|---|---|---|---|---|---|---|
| lungtumor | MacHypo | GSTP1 | CD44 | 2.4093 | 1.5821 | 1.9957 | 0.000999 |
| lungtumor | MacHypo | GSTP1 | IL7R | 2.4093 | 0.3898 | 1.3996 | 0.000999 |
| lungtumor | MacHypo | HMGB1 | LY96 | 1.1943 | 0.7778 | 0.9861 | 0.000999 |
| lungtumor | MacHypo | HMGB1 | TLR2 | 1.1943 | 0.2562 | 0.7252 | 0.000999 |
| lungtumor | MacHypo | HMGB2 | CD44 | 0.5593 | 1.5821 | 1.0707 | 0.000999 |
| lungtumor | MacHypo | HMGB2 | DAB2 | 0.5593 | 0.6138 | 0.5866 | 0.000999 |
| lungtumor | MacHypo | HSP90B1 | ITGB1 | 0.6141 | 0.4596 | 0.5369 | 0.000999 |
| lungtumor | MacHypo | HSP90B1 | TNFRSF1B | 0.6141 | 0.4397 | 0.5269 | 0.000999 |
| lungtumor | MacHypo | LGALS3 | CLEC7A | 0.9915 | 0.4087 | 0.7001 | 0.000999 |
| lungtumor | MacHypo | LGALS3 | FCGR2A | 0.9915 | 0.7827 | 0.8871 | 0.000999 |
| lungtumor | MacHypo | LGALS3 | FCGR2B | 0.9915 | 0.7744 | 0.883 | 0.000999 |
| lungtumor | MacHypo | LGALS3 | TFRC | 0.9915 | 0.4916 | 0.7415 | 0.000999 |
| lungtumor | MacHypo | MIF | CD74 | 1.4585 | 3.1473 | 2.3029 | 0.000999 |
| lungtumor | MacHypo | MIF | TNFRSF14 | 1.4585 | 0.3382 | 0.8983 | 0.000999 |
| lungtumor | MacHypo | RPS19 | C5AR1 | 3.5104 | 0.4508 | 1.9806 | 0.000999 |
| lungtumor | MacHypo | RPS19 | CD4 | 3.5104 | 0.2639 | 1.8871 | 0.000999 |
| lungtumor | MacHypo | RPS19 | ICAM1 | 3.5104 | 0.3685 | 1.9395 | 0.000999 |
| lungtumor | MacHypo | RPS19 | RPSA | 3.5104 | 1.9136 | 2.712 | 0.000999 |
| lungtumor | MacHypo | RTN4 | CD44 | 0.4013 | 1.5821 | 0.9917 | 0.000999 |
| lungtumor | MacHypo | SAA1 | C3AR1 | 0.2539 | 0.2925 | 0.2732 | 0.000999 |
| lungtumor | MacHypo | SAA1 | C5AR1 | 0.2539 | 0.4508 | 0.3523 | 0.000999 |
| lungtumor | MacHypo | SAA1 | CCR1 | 0.2539 | 0.2563 | 0.2551 | 0.000999 |
| lungtumor | MacHypo | SAA1 | CXCR4 | 0.2539 | 1.1308 | 0.6923 | 0.000999 |
| lungtumor | MacHypo | SERPINA1 | SDC2 | 0.3624 | 0.6065 | 0.4845 | 0.000999 |
| lungtumor | MacHypo | SPP1 | CD44 | 1.4874 | 1.5821 | 1.5348 | 0.000999 |
| lungtumor | MacHypo | HMGB1 | AGTRAP | 1.1943 | 0.263 | 0.7286 | 0.003 |
| melanomatumo | MacHypo | ARF1 | CD4 | 1.8545 | 0.2639 | 1.0592 | 0.000999 |
| melanomatumo | MacHypo | ARF1 | CD44 | 1.8545 | 1.5821 | 1.7183 | 0.000999 |
| melanomatumo | MacHypo | GSTP1 | CD44 | 2.3341 | 1.5821 | 1.9581 | 0.000999 |
| melanomatumo | MacHypo | GSTP1 | IL7R | 2.3341 | 0.3898 | 1.362 | 0.000999 |

| | | | | | | | |
|---|---|---|---|---|---|---|---|
| melanomatumo | MacHypo | HAS2 | CD44 | 0.3078 | 1.5821 | 0.9449 | 0.000999 |
| melanomatumo | MacHypo | HMGB1 | AGTRAP | 1.6757 | 0.263 | 0.9693 | 0.000999 |
| melanomatumo | MacHypo | HMGB1 | LY96 | 1.6757 | 0.7778 | 1.2268 | 0.000999 |
| melanomatumo | MacHypo | HMGB1 | TLR2 | 1.6757 | 0.2562 | 0.9659 | 0.000999 |
| melanomatumo | MacHypo | MIF | CD74 | 2.6939 | 3.1473 | 2.9206 | 0.000999 |
| melanomatumo | MacHypo | MIF | TNFRSF14 | 2.6939 | 0.3382 | 1.5161 | 0.000999 |
| melanomatumo | MacHypo | HEBP1 | CXCR4 | 0.3689 | 1.1308 | 0.7498 | 0.002 |
| melanomatumo | MacHypo | HEBP1 | C5AR1 | 0.3689 | 0.4508 | 0.4098 | 0.003 |
| melanomatumo | MacHypo | HEBP1 | C3AR1 | 0.3689 | 0.2925 | 0.3307 | 0.005 |
| melanomatumo | MacHypo | HEBP1 | FPR3 | 0.3689 | 0.3793 | 0.3741 | 0.005 |
| melanomatumo | MacHypo | HEBP1 | CCR1 | 0.3689 | 0.2563 | 0.3126 | 0.00599 |
| ovarytumor | MacHypo | ANXA1 | C3AR1 | 1.0314 | 0.2925 | 0.6619 | 0.000999 |
| ovarytumor | MacHypo | ANXA1 | C5AR1 | 1.0314 | 0.4508 | 0.7411 | 0.000999 |
| ovarytumor | MacHypo | ANXA1 | CCR1 | 1.0314 | 0.2563 | 0.6439 | 0.000999 |
| ovarytumor | MacHypo | ANXA1 | CD4 | 1.0314 | 0.2639 | 0.6476 | 0.000999 |
| ovarytumor | MacHypo | ANXA1 | CD44 | 1.0314 | 1.5821 | 1.3067 | 0.000999 |
| ovarytumor | MacHypo | ANXA1 | CXCR4 | 1.0314 | 1.1308 | 1.0811 | 0.000999 |
| ovarytumor | MacHypo | ANXA1 | FPR3 | 1.0314 | 0.3793 | 0.7053 | 0.000999 |
| ovarytumor | MacHypo | APOE | SDC2 | 0.837 | 0.6065 | 0.7218 | 0.000999 |
| ovarytumor | MacHypo | APP | ALOX5 | 0.2907 | 0.2606 | 0.2757 | 0.000999 |
| ovarytumor | MacHypo | APP | C3AR1 | 0.2907 | 0.2925 | 0.2916 | 0.000999 |
| ovarytumor | MacHypo | APP | C5AR1 | 0.2907 | 0.4508 | 0.3707 | 0.000999 |
| ovarytumor | MacHypo | APP | CCR1 | 0.2907 | 0.2563 | 0.2735 | 0.000999 |
| ovarytumor | MacHypo | APP | CXCR4 | 0.2907 | 1.1308 | 0.7107 | 0.000999 |
| ovarytumor | MacHypo | APP | TNFRSF14 | 0.2907 | 0.3382 | 0.3144 | 0.000999 |
| ovarytumor | MacHypo | ARF1 | CD4 | 0.7364 | 0.2639 | 0.5001 | 0.000999 |
| ovarytumor | MacHypo | ARF1 | CD44 | 0.7364 | 1.5821 | 1.1593 | 0.000999 |
| ovarytumor | MacHypo | CALM1 | TNFRSF1B | 1.0087 | 0.4397 | 0.7242 | 0.000999 |
| ovarytumor | MacHypo | CALR | CD44 | 0.7423 | 1.5821 | 1.1622 | 0.000999 |
| ovarytumor | MacHypo | CALR | IL7R | 0.7423 | 0.3898 | 0.566 | 0.000999 |

| | | | | | | | |
|---|---|---|---|---|---|---|---|
| ovarytumor | MacHypo | CCL2 | CCR1 | 0.3526 | 0.2563 | 0.3045 | 0.000999 |
| ovarytumor | MacHypo | COL1A1 | CD44 | 0.3504 | 1.5821 | 0.9662 | 0.000999 |
| ovarytumor | MacHypo | COL1A1 | ITGB1 | 0.3504 | 0.4596 | 0.405 | 0.000999 |
| ovarytumor | MacHypo | GRN | TNFRSF1B | 0.3896 | 0.4397 | 0.4146 | 0.000999 |
| ovarytumor | MacHypo | GSTP1 | CD44 | 2.6185 | 1.5821 | 2.1003 | 0.000999 |
| ovarytumor | MacHypo | GSTP1 | IL7R | 2.6185 | 0.3898 | 1.5042 | 0.000999 |
| ovarytumor | MacHypo | HLA-DMA | CD4 | 0.2537 | 0.2639 | 0.2588 | 0.000999 |
| ovarytumor | MacHypo | HLA-DRA | PTPRC | 1.2369 | 0.4793 | 0.8581 | 0.000999 |
| ovarytumor | MacHypo | HMGB1 | AGTRAP | 1.7272 | 0.263 | 0.9951 | 0.000999 |
| ovarytumor | MacHypo | HMGB1 | LY96 | 1.7272 | 0.7778 | 1.2525 | 0.000999 |
| ovarytumor | MacHypo | HMGB1 | TLR2 | 1.7272 | 0.2562 | 0.9917 | 0.000999 |
| ovarytumor | MacHypo | HMGB2 | CD44 | 0.4974 | 1.5821 | 1.0398 | 0.000999 |
| ovarytumor | MacHypo | HMGB2 | DAB2 | 0.4974 | 0.6138 | 0.5556 | 0.000999 |
| ovarytumor | MacHypo | HSP90B1 | ITGB1 | 0.6365 | 0.4596 | 0.5481 | 0.000999 |
| ovarytumor | MacHypo | HSP90B1 | TNFRSF1B | 0.6365 | 0.4397 | 0.5381 | 0.000999 |
| ovarytumor | MacHypo | ITGB1 | DAB2 | 0.2783 | 0.6138 | 0.446 | 0.000999 |
| ovarytumor | MacHypo | ITGB1 | ITGAX | 0.2783 | 0.3625 | 0.3204 | 0.000999 |
| ovarytumor | MacHypo | ITGB1 | PLAUR | 0.2783 | 1.5014 | 0.8899 | 0.000999 |
| ovarytumor | MacHypo | LGALS3 | CLEC7A | 0.5097 | 0.4087 | 0.4592 | 0.000999 |
| ovarytumor | MacHypo | LGALS3 | FCGR2A | 0.5097 | 0.7827 | 0.6462 | 0.000999 |
| ovarytumor | MacHypo | LGALS3 | FCGR2B | 0.5097 | 0.7744 | 0.6421 | 0.000999 |
| ovarytumor | MacHypo | LGALS3 | TFRC | 0.5097 | 0.4916 | 0.5006 | 0.000999 |
| ovarytumor | MacHypo | MIF | CD74 | 0.8565 | 3.1473 | 2.0019 | 0.000999 |
| ovarytumor | MacHypo | MIF | TNFRSF14 | 0.8565 | 0.3382 | 0.5973 | 0.000999 |
| ovarytumor | MacHypo | RPS19 | C5AR1 | 3.9325 | 0.4508 | 2.1916 | 0.000999 |
| ovarytumor | MacHypo | RPS19 | CD4 | 3.9325 | 0.2639 | 2.0982 | 0.000999 |
| ovarytumor | MacHypo | RPS19 | ICAM1 | 3.9325 | 0.3685 | 2.1505 | 0.000999 |
| ovarytumor | MacHypo | RTN4 | CD44 | 0.4992 | 1.5821 | 1.0406 | 0.000999 |
| ovarytumor | MacHypo | RTN4 | CD99 | 0.4992 | 0.765 | 0.6321 | 0.000999 |
| ovarytumor | MacHypo | SPP1 | CD44 | 0.3946 | 1.5821 | 0.9884 | 0.000999 |

| Supplementary Table 8: Communication results between macrophages and lymphocytes subpopulations, and malignant cells | | | | | | | |
|---|---|---|---|---|---|---|---|
| **MacIFN vs. lymphocytes** | | | | | | | |
| celltype_01 | celltype_02 | ligand | receptor | ligand_mean_expression | receptor_mean_expression | ligand_receptor_mean_expression | pvalue |
| MacIFN | NKT | A2M | TNFRSF14 | 0.3025 | 0.2848 | 0.2936 | 0.000999 |
| MacIFN | NKT | ADM | PTGER2 | 0.627 | 0.3627 | 0.4948 | 0.000999 |
| MacIFN | NKT | AIMP1 | RACK1 | 0.2858 | 2.4327 | 1.3593 | 0.000999 |
| MacIFN | NKT | APOE | SORL1 | 2.0054 | 0.4137 | 1.2095 | 0.000999 |
| MacIFN | NKT | CCL20 | CXCR4 | 0.33 | 2.0373 | 1.1836 | 0.000999 |
| MacIFN | NKT | CCL20 | S1PR4 | 0.33 | 0.4435 | 0.3868 | 0.000999 |
| MacIFN | NKT | CCL20 | S1PR5 | 0.33 | 0.508 | 0.419 | 0.000999 |
| MacIFN | NKT | COL1A1 | CD44 | 0.3983 | 0.6724 | 0.5354 | 0.000999 |
| MacIFN | NKT | COL1A1 | ITGB1 | 0.3983 | 0.488 | 0.4431 | 0.000999 |
| MacIFN | NKT | CXCL10 | CXCR4 | 1.153 | 2.0373 | 1.5951 | 0.000999 |
| MacIFN | NKT | CXCL10 | S1PR4 | 1.153 | 0.4435 | 0.7982 | 0.000999 |
| MacIFN | NKT | CXCL10 | S1PR5 | 1.153 | 0.508 | 0.8305 | 0.000999 |
| MacIFN | NKT | CXCL11 | CXCR4 | 0.347 | 2.0373 | 1.1921 | 0.000999 |
| MacIFN | NKT | CXCL11 | S1PR4 | 0.347 | 0.4435 | 0.3953 | 0.000999 |
| MacIFN | NKT | CXCL11 | S1PR5 | 0.347 | 0.508 | 0.4275 | 0.000999 |
| MacIFN | NKT | CXCL16 | CXCR4 | 0.7145 | 2.0373 | 1.3759 | 0.000999 |
| MacIFN | NKT | CXCL16 | S1PR4 | 0.7145 | 0.4435 | 0.579 | 0.000999 |
| MacIFN | NKT | CXCL16 | S1PR5 | 0.7145 | 0.508 | 0.6113 | 0.000999 |
| MacIFN | NKT | CXCL2 | CXCR4 | 0.4907 | 2.0373 | 1.264 | 0.000999 |
| MacIFN | NKT | CXCL2 | S1PR4 | 0.4907 | 0.4435 | 0.4671 | 0.000999 |
| MacIFN | NKT | CXCL2 | S1PR5 | 0.4907 | 0.508 | 0.4994 | 0.000999 |
| MacIFN | NKT | CXCL3 | CXCR4 | 0.3451 | 2.0373 | 1.1912 | 0.000999 |
| MacIFN | NKT | CXCL3 | S1PR4 | 0.3451 | 0.4435 | 0.3943 | 0.000999 |
| MacIFN | NKT | CXCL3 | S1PR5 | 0.3451 | 0.508 | 0.4266 | 0.000999 |
| MacIFN | NKT | CXCL9 | CXCR4 | 0.3037 | 2.0373 | 1.1705 | 0.000999 |
| MacIFN | NKT | CXCL9 | S1PR4 | 0.3037 | 0.4435 | 0.3736 | 0.000999 |
| MacIFN | NKT | CXCL9 | S1PR5 | 0.3037 | 0.508 | 0.4059 | 0.000999 |

| | | | | | | | |
|---|---|---|---|---|---|---|---|
| MacIFN | NKT | FN1 | CD44 | 1.1317 | 0.6724 | 0.9021 | 0.000999 |
| MacIFN | NKT | FN1 | ITGA4 | 1.1317 | 0.2946 | 0.7131 | 0.000999 |
| MacIFN | NKT | FN1 | ITGB1 | 1.1317 | 0.488 | 0.8098 | 0.000999 |
| MacIFN | NKT | FN1 | ITGB7 | 1.1317 | 0.4721 | 0.8019 | 0.000999 |
| MacIFN | NKT | FN1 | RPSA | 1.1317 | 2.3639 | 1.7478 | 0.000999 |
| MacIFN | NKT | GRN | TNFRSF1B | 1.3662 | 0.4949 | 0.9306 | 0.000999 |
| MacIFN | NKT | GSTP1 | CD44 | 1.2812 | 0.6724 | 0.9768 | 0.000999 |
| MacIFN | NKT | GSTP1 | IL7R | 1.2812 | 0.5933 | 0.9373 | 0.000999 |
| MacIFN | NKT | GSTP1 | ITGA4 | 1.2812 | 0.2946 | 0.7879 | 0.000999 |
| MacIFN | NKT | HBEGF | CD44 | 0.2793 | 0.6724 | 0.4759 | 0.000999 |
| MacIFN | NKT | HLA-DRA | PTPRC | 3.8224 | 1.8797 | 2.8511 | 0.000999 |
| MacIFN | NKT | ICAM1 | IL2RG | 0.3372 | 1.1524 | 0.7448 | 0.000999 |
| MacIFN | NKT | ICAM1 | ITGAL | 0.3372 | 0.6778 | 0.5075 | 0.000999 |
| MacIFN | NKT | ICAM1 | ITGB2 | 0.3372 | 1.6481 | 0.9927 | 0.000999 |
| MacIFN | NKT | ICAM1 | SPN | 0.3372 | 0.6398 | 0.4885 | 0.000999 |
| MacIFN | NKT | LGALS3 | CD6 | 1.5031 | 0.4881 | 0.9956 | 0.000999 |
| MacIFN | NKT | LGALS3 | ITGB1 | 1.5031 | 0.488 | 0.9955 | 0.000999 |
| MacIFN | NKT | LGALS3 | SPN | 1.5031 | 0.6398 | 1.0714 | 0.000999 |
| MacIFN | NKT | LRPAP1 | SORL1 | 0.4177 | 0.4137 | 0.4157 | 0.000999 |
| MacIFN | NKT | MIF | CD74 | 1.1447 | 1.6792 | 1.412 | 0.000999 |
| MacIFN | NKT | MIF | ITGA4 | 1.1447 | 0.2946 | 0.7196 | 0.000999 |
| MacIFN | NKT | MIF | TNFRSF14 | 1.1447 | 0.2848 | 0.7148 | 0.000999 |
| MacIFN | NKT | NAMPT | ANXA6 | 0.9717 | 0.844 | 0.9078 | 0.000999 |
| MacIFN | NKT | RTN4 | CD44 | 0.9008 | 0.6724 | 0.7866 | 0.000999 |
| MacIFN | NKT | RTN4 | CD99 | 0.9008 | 1.4547 | 1.1777 | 0.000999 |
| MacIFN | NKT | SPP1 | CD44 | 3.5345 | 0.6724 | 2.1034 | 0.000999 |
| MacIFN | NKT | SPP1 | ITGA4 | 3.5345 | 0.2946 | 1.9145 | 0.000999 |
| MacIFN | NKT | SPP1 | ITGB1 | 3.5345 | 0.488 | 2.0112 | 0.000999 |
| MacIFN | NKT | TNFSF13B | ITGB1 | 0.9552 | 0.488 | 0.7216 | 0.000999 |
| MacIFN | TCD4ex | A2M | TNFRSF14 | 0.3025 | 0.2806 | 0.2915 | 0.000999 |

| | | | | | | | |
|---|---|---|---|---|---|---|---|
| MacIFN | TCD4ex | ADM | PTGER4 | 0.627 | 0.4696 | 0.5483 | 0.000999 |
| MacIFN | TCD4ex | AIMP1 | RACK1 | 0.2858 | 2.4434 | 1.3646 | 0.000999 |
| MacIFN | TCD4ex | ANXA1 | CCR7 | 1.0061 | 0.4844 | 0.7453 | 0.000999 |
| MacIFN | TCD4ex | ANXA1 | CD4 | 1.0061 | 0.5864 | 0.7963 | 0.000999 |
| MacIFN | TCD4ex | ANXA1 | CD44 | 1.0061 | 1.0244 | 1.0152 | 0.000999 |
| MacIFN | TCD4ex | ANXA1 | CXCR4 | 1.0061 | 2.4201 | 1.7131 | 0.000999 |
| MacIFN | TCD4ex | CALM1 | RPSA | 1.377 | 2.4895 | 1.9333 | 0.000999 |
| MacIFN | TCD4ex | CALR | IL7R | 0.904 | 1.2353 | 1.0696 | 0.000999 |
| MacIFN | TCD4ex | CCL20 | CCR7 | 0.33 | 0.4844 | 0.4072 | 0.000999 |
| MacIFN | TCD4ex | CCL20 | CXCR4 | 0.33 | 2.4201 | 1.375 | 0.000999 |
| MacIFN | TCD4ex | CD86 | CD28 | 0.5457 | 0.3256 | 0.4356 | 0.000999 |
| MacIFN | TCD4ex | CD86 | CTLA4 | 0.5457 | 0.6294 | 0.5875 | 0.000999 |
| MacIFN | TCD4ex | COL1A1 | CD44 | 0.3983 | 1.0244 | 0.7113 | 0.000999 |
| MacIFN | TCD4ex | COL1A1 | ITGB1 | 0.3983 | 0.4615 | 0.4299 | 0.000999 |
| MacIFN | TCD4ex | CXCL10 | CCR7 | 1.153 | 0.4844 | 0.8187 | 0.000999 |
| MacIFN | TCD4ex | CXCL10 | CXCR4 | 1.153 | 2.4201 | 1.7865 | 0.000999 |
| MacIFN | TCD4ex | CXCL11 | CCR7 | 0.347 | 0.4844 | 0.4157 | 0.000999 |
| MacIFN | TCD4ex | CXCL11 | CXCR4 | 0.347 | 2.4201 | 1.3835 | 0.000999 |
| MacIFN | TCD4ex | CXCL16 | CCR7 | 0.7145 | 0.4844 | 0.5995 | 0.000999 |
| MacIFN | TCD4ex | CXCL16 | CXCR4 | 0.7145 | 2.4201 | 1.5673 | 0.000999 |
| MacIFN | TCD4ex | CXCL2 | CCR7 | 0.4907 | 0.4844 | 0.4876 | 0.000999 |
| MacIFN | TCD4ex | CXCL2 | CXCR4 | 0.4907 | 2.4201 | 1.4554 | 0.000999 |
| MacIFN | TCD4ex | CXCL3 | CCR7 | 0.3451 | 0.4844 | 0.4147 | 0.000999 |
| MacIFN | TCD4ex | CXCL3 | CXCR4 | 0.3451 | 2.4201 | 1.3826 | 0.000999 |
| MacIFN | TCD4ex | CXCL9 | CCR7 | 0.3037 | 0.4844 | 0.3941 | 0.000999 |
| MacIFN | TCD4ex | CXCL9 | CXCR4 | 0.3037 | 2.4201 | 1.3619 | 0.000999 |
| MacIFN | TCD4ex | FN1 | CD44 | 1.1317 | 1.0244 | 1.0781 | 0.000999 |
| MacIFN | TCD4ex | FN1 | ITGB1 | 1.1317 | 0.4615 | 0.7966 | 0.000999 |
| MacIFN | TCD4ex | FN1 | RPSA | 1.1317 | 2.4895 | 1.8106 | 0.000999 |
| MacIFN | TCD4ex | GRN | TNFRSF1B | 1.3662 | 0.383 | 0.8746 | 0.000999 |

| | | | | | | | |
|---|---|---|---|---|---|---|---|
| MacIFN | TCD4ex | GSTP1 | CD44 | 1.2812 | 1.0244 | 1.1528 | 0.000999 |
| MacIFN | TCD4ex | GSTP1 | IL7R | 1.2812 | 1.2353 | 1.2582 | 0.000999 |
| MacIFN | TCD4ex | HBEGF | CD44 | 0.2793 | 1.0244 | 0.6519 | 0.000999 |
| MacIFN | TCD4ex | HLA-DMA | CD4 | 1.0847 | 0.5864 | 0.8356 | 0.000999 |
| MacIFN | TCD4ex | HLA-DRA | CD4 | 3.8224 | 0.5864 | 2.2044 | 0.000999 |
| MacIFN | TCD4ex | HLA-DRA | PDCD1 | 3.8224 | 0.457 | 2.1397 | 0.000999 |
| MacIFN | TCD4ex | HLA-DRA | PTPRC | 3.8224 | 1.7493 | 2.7859 | 0.000999 |
| MacIFN | TCD4ex | HSP90B1 | ITGB1 | 0.6822 | 0.4615 | 0.5719 | 0.000999 |
| MacIFN | TCD4ex | HSP90B1 | TNFRSF1B | 0.6822 | 0.383 | 0.5326 | 0.000999 |
| MacIFN | TCD4ex | ICAM1 | IL2RG | 0.3372 | 0.9695 | 0.6533 | 0.000999 |
| MacIFN | TCD4ex | ICAM1 | ITGAL | 0.3372 | 0.2603 | 0.2988 | 0.000999 |
| MacIFN | TCD4ex | ICAM1 | ITGB2 | 0.3372 | 0.7871 | 0.5621 | 0.000999 |
| MacIFN | TCD4ex | LGALS3 | CD6 | 1.5031 | 0.5338 | 1.0184 | 0.000999 |
| MacIFN | TCD4ex | LGALS3 | ITGB1 | 1.5031 | 0.4615 | 0.9823 | 0.000999 |
| MacIFN | TCD4ex | MIF | TNFRSF14 | 1.1447 | 0.2806 | 0.7127 | 0.000999 |
| MacIFN | TCD4ex | NAMPT | ANXA6 | 0.9717 | 0.5386 | 0.7551 | 0.000999 |
| MacIFN | TCD4ex | RTN4 | CD44 | 0.9008 | 1.0244 | 0.9626 | 0.000999 |
| MacIFN | TCD4ex | RTN4 | CD99 | 0.9008 | 0.9165 | 0.9086 | 0.000999 |
| MacIFN | TCD4ex | SPP1 | CD44 | 3.5345 | 1.0244 | 2.2794 | 0.000999 |
| MacIFN | TCD4ex | SPP1 | ITGB1 | 3.5345 | 0.4615 | 1.998 | 0.000999 |
| MacIFN | TCD4ex | TGFB1 | ITGB1 | 0.2566 | 0.4615 | 0.359 | 0.000999 |
| MacIFN | TCD4ex | TNFSF13B | ITGB1 | 0.9552 | 0.4615 | 0.7084 | 0.000999 |
| MacIFN | TCD4ex | ITGB2 | ITGAL | 0.926 | 0.2603 | 0.5932 | 0.002 |
| MacIFN | TCD4ex | MIF | CD74 | 1.1447 | 1.6094 | 1.3771 | 0.003 |
| MacIFN | TCD4ex | CALR | CD44 | 0.904 | 1.0244 | 0.9642 | 0.00599 |
| MacIFN | TCD4reg | A2M | TNFRSF14 | 0.3025 | 0.3843 | 0.3434 | 0.000999 |
| MacIFN | TCD4reg | ANXA1 | CCR7 | 1.0061 | 0.3098 | 0.658 | 0.000999 |
| MacIFN | TCD4reg | ANXA1 | CD4 | 1.0061 | 0.429 | 0.7175 | 0.000999 |
| MacIFN | TCD4reg | ANXA1 | CD44 | 1.0061 | 1.0896 | 1.0479 | 0.000999 |
| MacIFN | TCD4reg | ANXA1 | CXCR3 | 1.0061 | 0.3357 | 0.6709 | 0.000999 |

| | | | | | | | |
|---|---|---|---|---|---|---|---|
| MacIFN | TCD4reg | ANXA1 | CXCR4 | 1.0061 | 2.2962 | 1.6511 | 0.000999 |
| MacIFN | TCD4reg | ANXA1 | CXCR6 | 1.0061 | 0.3945 | 0.7003 | 0.000999 |
| MacIFN | TCD4reg | ANXA1 | FAS | 1.0061 | 0.2727 | 0.6394 | 0.000999 |
| MacIFN | TCD4reg | CALR | CD44 | 0.904 | 1.0896 | 0.9968 | 0.000999 |
| MacIFN | TCD4reg | CALR | FAS | 0.904 | 0.2727 | 0.5884 | 0.000999 |
| MacIFN | TCD4reg | CALR | IL7R | 0.904 | 0.9682 | 0.9361 | 0.000999 |
| MacIFN | TCD4reg | CCL20 | CCR7 | 0.33 | 0.3098 | 0.3199 | 0.000999 |
| MacIFN | TCD4reg | CCL20 | CXCR3 | 0.33 | 0.3357 | 0.3328 | 0.000999 |
| MacIFN | TCD4reg | CCL20 | CXCR4 | 0.33 | 2.2962 | 1.3131 | 0.000999 |
| MacIFN | TCD4reg | CCL20 | CXCR6 | 0.33 | 0.3945 | 0.3623 | 0.000999 |
| MacIFN | TCD4reg | CCL20 | S1PR4 | 0.33 | 0.2744 | 0.3022 | 0.000999 |
| MacIFN | TCD4reg | CCL7 | CXCR3 | 0.3065 | 0.3357 | 0.3211 | 0.000999 |
| MacIFN | TCD4reg | CD86 | CD28 | 0.5457 | 0.313 | 0.4293 | 0.000999 |
| MacIFN | TCD4reg | CD86 | CTLA4 | 0.5457 | 0.7909 | 0.6683 | 0.000999 |
| MacIFN | TCD4reg | COL1A1 | CD44 | 0.3983 | 1.0896 | 0.744 | 0.000999 |
| MacIFN | TCD4reg | COL1A1 | ITGB1 | 0.3983 | 0.4001 | 0.3992 | 0.000999 |
| MacIFN | TCD4reg | CXCL10 | CCR7 | 1.153 | 0.3098 | 0.7314 | 0.000999 |
| MacIFN | TCD4reg | CXCL10 | CXCR3 | 1.153 | 0.3357 | 0.7443 | 0.000999 |
| MacIFN | TCD4reg | CXCL10 | CXCR4 | 1.153 | 2.2962 | 1.7246 | 0.000999 |
| MacIFN | TCD4reg | CXCL10 | CXCR6 | 1.153 | 0.3945 | 0.7737 | 0.000999 |
| MacIFN | TCD4reg | CXCL10 | S1PR4 | 1.153 | 0.2744 | 0.7137 | 0.000999 |
| MacIFN | TCD4reg | CXCL11 | CCR7 | 0.347 | 0.3098 | 0.3284 | 0.000999 |
| MacIFN | TCD4reg | CXCL11 | CXCR3 | 0.347 | 0.3357 | 0.3413 | 0.000999 |
| MacIFN | TCD4reg | CXCL11 | CXCR4 | 0.347 | 2.2962 | 1.3216 | 0.000999 |
| MacIFN | TCD4reg | CXCL11 | CXCR6 | 0.347 | 0.3945 | 0.3708 | 0.000999 |
| MacIFN | TCD4reg | CXCL11 | S1PR4 | 0.347 | 0.2744 | 0.3107 | 0.000999 |
| MacIFN | TCD4reg | CXCL16 | CCR7 | 0.7145 | 0.3098 | 0.5122 | 0.000999 |
| MacIFN | TCD4reg | CXCL16 | CXCR3 | 0.7145 | 0.3357 | 0.5251 | 0.000999 |
| MacIFN | TCD4reg | CXCL16 | CXCR4 | 0.7145 | 2.2962 | 1.5053 | 0.000999 |
| MacIFN | TCD4reg | CXCL16 | CXCR6 | 0.7145 | 0.3945 | 0.5545 | 0.000999 |

| | | | | | | | |
|---|---|---|---|---|---|---|---|
| MacIFN | TCD4reg | CXCL16 | S1PR4 | 0.7145 | 0.2744 | 0.4945 | 0.000999 |
| MacIFN | TCD4reg | CXCL2 | CCR7 | 0.4907 | 0.3098 | 0.4003 | 0.000999 |
| MacIFN | TCD4reg | CXCL2 | CXCR3 | 0.4907 | 0.3357 | 0.4132 | 0.000999 |
| MacIFN | TCD4reg | CXCL2 | CXCR4 | 0.4907 | 2.2962 | 1.3934 | 0.000999 |
| MacIFN | TCD4reg | CXCL2 | CXCR6 | 0.4907 | 0.3945 | 0.4426 | 0.000999 |
| MacIFN | TCD4reg | CXCL2 | S1PR4 | 0.4907 | 0.2744 | 0.3826 | 0.000999 |
| MacIFN | TCD4reg | CXCL3 | CCR7 | 0.3451 | 0.3098 | 0.3275 | 0.000999 |
| MacIFN | TCD4reg | CXCL3 | CXCR3 | 0.3451 | 0.3357 | 0.3404 | 0.000999 |
| MacIFN | TCD4reg | CXCL3 | CXCR4 | 0.3451 | 2.2962 | 1.3206 | 0.000999 |
| MacIFN | TCD4reg | CXCL3 | CXCR6 | 0.3451 | 0.3945 | 0.3698 | 0.000999 |
| MacIFN | TCD4reg | CXCL3 | S1PR4 | 0.3451 | 0.2744 | 0.3097 | 0.000999 |
| MacIFN | TCD4reg | CXCL9 | CCR7 | 0.3037 | 0.3098 | 0.3068 | 0.000999 |
| MacIFN | TCD4reg | CXCL9 | CXCR3 | 0.3037 | 0.3357 | 0.3197 | 0.000999 |
| MacIFN | TCD4reg | CXCL9 | CXCR4 | 0.3037 | 2.2962 | 1.3 | 0.000999 |
| MacIFN | TCD4reg | CXCL9 | CXCR6 | 0.3037 | 0.3945 | 0.3491 | 0.000999 |
| MacIFN | TCD4reg | CXCL9 | S1PR4 | 0.3037 | 0.2744 | 0.2891 | 0.000999 |
| MacIFN | TCD4reg | FN1 | CD44 | 1.1317 | 1.0896 | 1.1107 | 0.000999 |
| MacIFN | TCD4reg | FN1 | ITGB1 | 1.1317 | 0.4001 | 0.7659 | 0.000999 |
| MacIFN | TCD4reg | FN1 | RPSA | 1.1317 | 2.4999 | 1.8158 | 0.000999 |
| MacIFN | TCD4reg | GRN | TNFRSF1B | 1.3662 | 0.7982 | 1.0822 | 0.000999 |
| MacIFN | TCD4reg | GSTP1 | CD44 | 1.2812 | 1.0896 | 1.1854 | 0.000999 |
| MacIFN | TCD4reg | GSTP1 | FAS | 1.2812 | 0.2727 | 0.777 | 0.000999 |
| MacIFN | TCD4reg | GSTP1 | IL7R | 1.2812 | 0.9682 | 1.1247 | 0.000999 |
| MacIFN | TCD4reg | HBEGF | CD44 | 0.2793 | 1.0896 | 0.6845 | 0.000999 |
| MacIFN | TCD4reg | HLA-DMA | CD4 | 1.0847 | 0.429 | 0.7568 | 0.000999 |
| MacIFN | TCD4reg | HLA-DRA | CD4 | 3.8224 | 0.429 | 2.1257 | 0.000999 |
| MacIFN | TCD4reg | HLA-DRA | PTPRC | 3.8224 | 1.6603 | 2.7414 | 0.000999 |
| MacIFN | TCD4reg | HSP90B1 | ITGB1 | 0.6822 | 0.4001 | 0.5412 | 0.000999 |
| MacIFN | TCD4reg | HSP90B1 | TNFRSF1B | 0.6822 | 0.7982 | 0.7402 | 0.000999 |
| MacIFN | TCD4reg | ICAM1 | IL2RA | 0.3372 | 0.7115 | 0.5243 | 0.000999 |

| | | | | | | | |
|---|---|---|---|---|---|---|---|
| MacIFN | TCD4reg | ICAM1 | IL2RG | 0.3372 | 1.1601 | 0.7487 | 0.000999 |
| MacIFN | TCD4reg | ICAM1 | ITGB2 | 0.3372 | 0.6238 | 0.4805 | 0.000999 |
| MacIFN | TCD4reg | LGALS3 | CD6 | 1.5031 | 0.2651 | 0.8841 | 0.000999 |
| MacIFN | TCD4reg | LGALS3 | FAS | 1.5031 | 0.2727 | 0.8879 | 0.000999 |
| MacIFN | TCD4reg | LGALS3 | ITGB1 | 1.5031 | 0.4001 | 0.9516 | 0.000999 |
| MacIFN | TCD4reg | MIF | CD74 | 1.1447 | 2.0716 | 1.6082 | 0.000999 |
| MacIFN | TCD4reg | MIF | TNFRSF14 | 1.1447 | 0.3843 | 0.7645 | 0.000999 |
| MacIFN | TCD4reg | NAMPT | ANXA6 | 0.9717 | 0.4557 | 0.7137 | 0.000999 |
| MacIFN | TCD4reg | RTN4 | CD44 | 0.9008 | 1.0896 | 0.9952 | 0.000999 |
| MacIFN | TCD4reg | RTN4 | CD99 | 0.9008 | 0.6603 | 0.7805 | 0.000999 |
| MacIFN | TCD4reg | SPP1 | CD44 | 3.5345 | 1.0896 | 2.312 | 0.000999 |
| MacIFN | TCD4reg | SPP1 | ITGB1 | 3.5345 | 0.4001 | 1.9673 | 0.000999 |
| MacIFN | TCD4reg | TNFSF13B | ITGB1 | 0.9552 | 0.4001 | 0.6777 | 0.000999 |
| MacIFN | TCD4reg | TGFB1 | ITGB1 | 0.2566 | 0.4001 | 0.3283 | 0.002 |
| MacIFN | TCD8em | A2M | TNFRSF14 | 0.3025 | 0.2915 | 0.297 | 0.000999 |
| MacIFN | TCD8em | ADM | PTGER4 | 0.627 | 0.2847 | 0.4559 | 0.000999 |
| MacIFN | TCD8em | CALR | CD44 | 0.904 | 1.119 | 1.0115 | 0.000999 |
| MacIFN | TCD8em | CALR | IL7R | 0.904 | 0.7183 | 0.8112 | 0.000999 |
| MacIFN | TCD8em | CCL20 | CXCR3 | 0.33 | 0.4189 | 0.3744 | 0.000999 |
| MacIFN | TCD8em | CCL20 | CXCR4 | 0.33 | 2.8389 | 1.5845 | 0.000999 |
| MacIFN | TCD8em | CCL20 | CXCR6 | 0.33 | 0.3399 | 0.3349 | 0.000999 |
| MacIFN | TCD8em | CCL20 | S1PR4 | 0.33 | 0.2677 | 0.2989 | 0.000999 |
| MacIFN | TCD8em | CCL7 | CXCR3 | 0.3065 | 0.4189 | 0.3627 | 0.000999 |
| MacIFN | TCD8em | COL1A1 | CD44 | 0.3983 | 1.119 | 0.7587 | 0.000999 |
| MacIFN | TCD8em | COL1A1 | ITGB1 | 0.3983 | 0.2557 | 0.327 | 0.000999 |
| MacIFN | TCD8em | CXCL10 | CXCR3 | 1.153 | 0.4189 | 0.7859 | 0.000999 |
| MacIFN | TCD8em | CXCL10 | CXCR4 | 1.153 | 2.8389 | 1.9959 | 0.000999 |
| MacIFN | TCD8em | CXCL10 | CXCR6 | 1.153 | 0.3399 | 0.7464 | 0.000999 |
| MacIFN | TCD8em | CXCL10 | S1PR4 | 1.153 | 0.2677 | 0.7103 | 0.000999 |
| MacIFN | TCD8em | CXCL11 | CXCR3 | 0.347 | 0.4189 | 0.3829 | 0.000999 |

| | | | | | | | |
|---|---|---|---|---|---|---|---|
| MacIFN | TCD8em | CXCL11 | CXCR4 | 0.347 | 2.8389 | 1.5929 | 0.000999 |
| MacIFN | TCD8em | CXCL11 | CXCR6 | 0.347 | 0.3399 | 0.3434 | 0.000999 |
| MacIFN | TCD8em | CXCL11 | S1PR4 | 0.347 | 0.2677 | 0.3073 | 0.000999 |
| MacIFN | TCD8em | CXCL16 | CXCR3 | 0.7145 | 0.4189 | 0.5667 | 0.000999 |
| MacIFN | TCD8em | CXCL16 | CXCR4 | 0.7145 | 2.8389 | 1.7767 | 0.000999 |
| MacIFN | TCD8em | CXCL16 | CXCR6 | 0.7145 | 0.3399 | 0.5272 | 0.000999 |
| MacIFN | TCD8em | CXCL16 | S1PR4 | 0.7145 | 0.2677 | 0.4911 | 0.000999 |
| MacIFN | TCD8em | CXCL2 | CXCR3 | 0.4907 | 0.4189 | 0.4548 | 0.000999 |
| MacIFN | TCD8em | CXCL2 | CXCR4 | 0.4907 | 2.8389 | 1.6648 | 0.000999 |
| MacIFN | TCD8em | CXCL2 | CXCR6 | 0.4907 | 0.3399 | 0.4153 | 0.000999 |
| MacIFN | TCD8em | CXCL2 | S1PR4 | 0.4907 | 0.2677 | 0.3792 | 0.000999 |
| MacIFN | TCD8em | CXCL3 | CXCR3 | 0.3451 | 0.4189 | 0.382 | 0.000999 |
| MacIFN | TCD8em | CXCL3 | CXCR4 | 0.3451 | 2.8389 | 1.592 | 0.000999 |
| MacIFN | TCD8em | CXCL3 | CXCR6 | 0.3451 | 0.3399 | 0.3425 | 0.000999 |
| MacIFN | TCD8em | CXCL3 | S1PR4 | 0.3451 | 0.2677 | 0.3064 | 0.000999 |
| MacIFN | TCD8em | CXCL9 | CXCR3 | 0.3037 | 0.4189 | 0.3613 | 0.000999 |
| MacIFN | TCD8em | CXCL9 | CXCR4 | 0.3037 | 2.8389 | 1.5713 | 0.000999 |
| MacIFN | TCD8em | CXCL9 | CXCR6 | 0.3037 | 0.3399 | 0.3218 | 0.000999 |
| MacIFN | TCD8em | CXCL9 | S1PR4 | 0.3037 | 0.2677 | 0.2857 | 0.000999 |
| MacIFN | TCD8em | FN1 | CD44 | 1.1317 | 1.119 | 1.1254 | 0.000999 |
| MacIFN | TCD8em | FN1 | ITGA4 | 1.1317 | 0.2823 | 0.707 | 0.000999 |
| MacIFN | TCD8em | FN1 | ITGB1 | 1.1317 | 0.2557 | 0.6937 | 0.000999 |
| MacIFN | TCD8em | FN1 | RPSA | 1.1317 | 2.3686 | 1.7502 | 0.000999 |
| MacIFN | TCD8em | GRN | TNFRSF1B | 1.3662 | 0.3038 | 0.835 | 0.000999 |
| MacIFN | TCD8em | GSTP1 | CD44 | 1.2812 | 1.119 | 1.2001 | 0.000999 |
| MacIFN | TCD8em | GSTP1 | IL7R | 1.2812 | 0.7183 | 0.9998 | 0.000999 |
| MacIFN | TCD8em | GSTP1 | ITGA4 | 1.2812 | 0.2823 | 0.7818 | 0.000999 |
| MacIFN | TCD8em | HBEGF | CD44 | 0.2793 | 1.119 | 0.6992 | 0.000999 |
| MacIFN | TCD8em | HLA-DRA | PTPRC | 3.8224 | 1.7912 | 2.8068 | 0.000999 |
| MacIFN | TCD8em | HSP90B1 | ITGB1 | 0.6822 | 0.2557 | 0.469 | 0.000999 |

| | | | | | | | |
|---|---|---|---|---|---|---|---|
| MacIFN | TCD8em | HSP90B1 | TNFRSF1B | 0.6822 | 0.3038 | 0.493 | 0.000999 |
| MacIFN | TCD8em | ICAM1 | IL2RG | 0.3372 | 0.9501 | 0.6437 | 0.000999 |
| MacIFN | TCD8em | ICAM1 | ITGAL | 0.3372 | 0.3162 | 0.3267 | 0.000999 |
| MacIFN | TCD8em | ICAM1 | ITGB2 | 0.3372 | 1.0018 | 0.6695 | 0.000999 |
| MacIFN | TCD8em | ICAM1 | SPN | 0.3372 | 0.2603 | 0.2988 | 0.000999 |
| MacIFN | TCD8em | LGALS3 | CD6 | 1.5031 | 0.4204 | 0.9617 | 0.000999 |
| MacIFN | TCD8em | LGALS3 | ITGB1 | 1.5031 | 0.2557 | 0.8794 | 0.000999 |
| MacIFN | TCD8em | LGALS3 | SPN | 1.5031 | 0.2603 | 0.8817 | 0.000999 |
| MacIFN | TCD8em | MIF | CD74 | 1.1447 | 2.4538 | 1.7993 | 0.000999 |
| MacIFN | TCD8em | MIF | ITGA4 | 1.1447 | 0.2823 | 0.7135 | 0.000999 |
| MacIFN | TCD8em | MIF | TNFRSF14 | 1.1447 | 0.2915 | 0.7181 | 0.000999 |
| MacIFN | TCD8em | NAMPT | ANXA6 | 0.9717 | 0.552 | 0.7618 | 0.000999 |
| MacIFN | TCD8em | RTN4 | CD44 | 0.9008 | 1.119 | 1.0099 | 0.000999 |
| MacIFN | TCD8em | RTN4 | CD99 | 0.9008 | 1.1525 | 1.0266 | 0.000999 |
| MacIFN | TCD8em | SPP1 | CD44 | 3.5345 | 1.119 | 2.3268 | 0.000999 |
| MacIFN | TCD8em | SPP1 | ITGA4 | 3.5345 | 0.2823 | 1.9084 | 0.000999 |
| MacIFN | TCD8em | SPP1 | ITGB1 | 3.5345 | 0.2557 | 1.8951 | 0.000999 |
| MacIFN | TCD8em | TNFSF13B | ITGB1 | 0.9552 | 0.2557 | 0.6055 | 0.000999 |
| MacIFN | TCD8em | AIMP1 | RACK1 | 0.2858 | 2.4374 | 1.3616 | 0.003 |
| MacIFN | TCD8ex | A2M | TNFRSF14 | 0.3025 | 0.3094 | 0.306 | 0.000999 |
| MacIFN | TCD8ex | ADM | PTGER4 | 0.627 | 0.3089 | 0.4679 | 0.000999 |
| MacIFN | TCD8ex | ANXA1 | CD44 | 1.0061 | 0.8219 | 0.914 | 0.000999 |
| MacIFN | TCD8ex | ANXA1 | CXCR3 | 1.0061 | 0.3647 | 0.6854 | 0.000999 |
| MacIFN | TCD8ex | ANXA1 | CXCR4 | 1.0061 | 2.3472 | 1.6767 | 0.000999 |
| MacIFN | TCD8ex | ANXA1 | CXCR6 | 1.0061 | 0.6626 | 0.8343 | 0.000999 |
| MacIFN | TCD8ex | ANXA1 | ITGA4 | 1.0061 | 0.4432 | 0.7247 | 0.000999 |
| MacIFN | TCD8ex | ANXA1 | VCAM1 | 1.0061 | 0.4462 | 0.7262 | 0.000999 |
| MacIFN | TCD8ex | APOE | VCAM1 | 2.0054 | 0.4462 | 1.2258 | 0.000999 |
| MacIFN | TCD8ex | CCL20 | CXCR3 | 0.33 | 0.3647 | 0.3473 | 0.000999 |
| MacIFN | TCD8ex | CCL20 | CXCR4 | 0.33 | 2.3472 | 1.3386 | 0.000999 |

| | | | | | | | |
|---|---|---|---|---|---|---|---|
| MacIFN | TCD8ex | CCL20 | CXCR6 | 0.33 | 0.6626 | 0.4963 | 0.000999 |
| MacIFN | TCD8ex | CCL7 | CXCR3 | 0.3065 | 0.3647 | 0.3356 | 0.000999 |
| MacIFN | TCD8ex | CD86 | CTLA4 | 0.5457 | 0.4883 | 0.517 | 0.000999 |
| MacIFN | TCD8ex | COL1A1 | CD44 | 0.3983 | 0.8219 | 0.6101 | 0.000999 |
| MacIFN | TCD8ex | COL1A1 | ITGB1 | 0.3983 | 0.2995 | 0.3489 | 0.000999 |
| MacIFN | TCD8ex | CXCL10 | CXCR3 | 1.153 | 0.3647 | 0.7588 | 0.000999 |
| MacIFN | TCD8ex | CXCL10 | CXCR4 | 1.153 | 2.3472 | 1.7501 | 0.000999 |
| MacIFN | TCD8ex | CXCL10 | CXCR6 | 1.153 | 0.6626 | 0.9078 | 0.000999 |
| MacIFN | TCD8ex | CXCL11 | CXCR3 | 0.347 | 0.3647 | 0.3558 | 0.000999 |
| MacIFN | TCD8ex | CXCL11 | CXCR4 | 0.347 | 2.3472 | 1.3471 | 0.000999 |
| MacIFN | TCD8ex | CXCL11 | CXCR6 | 0.347 | 0.6626 | 0.5048 | 0.000999 |
| MacIFN | TCD8ex | CXCL16 | CXCR3 | 0.7145 | 0.3647 | 0.5396 | 0.000999 |
| MacIFN | TCD8ex | CXCL16 | CXCR4 | 0.7145 | 2.3472 | 1.5309 | 0.000999 |
| MacIFN | TCD8ex | CXCL16 | CXCR6 | 0.7145 | 0.6626 | 0.6885 | 0.000999 |
| MacIFN | TCD8ex | CXCL2 | CXCR3 | 0.4907 | 0.3647 | 0.4277 | 0.000999 |
| MacIFN | TCD8ex | CXCL2 | CXCR4 | 0.4907 | 2.3472 | 1.419 | 0.000999 |
| MacIFN | TCD8ex | CXCL2 | CXCR6 | 0.4907 | 0.6626 | 0.5766 | 0.000999 |
| MacIFN | TCD8ex | CXCL3 | CXCR3 | 0.3451 | 0.3647 | 0.3549 | 0.000999 |
| MacIFN | TCD8ex | CXCL3 | CXCR4 | 0.3451 | 2.3472 | 1.3462 | 0.000999 |
| MacIFN | TCD8ex | CXCL3 | CXCR6 | 0.3451 | 0.6626 | 0.5038 | 0.000999 |
| MacIFN | TCD8ex | CXCL9 | CXCR3 | 0.3037 | 0.3647 | 0.3342 | 0.000999 |
| MacIFN | TCD8ex | CXCL9 | CXCR4 | 0.3037 | 2.3472 | 1.3255 | 0.000999 |
| MacIFN | TCD8ex | CXCL9 | CXCR6 | 0.3037 | 0.6626 | 0.4832 | 0.000999 |
| MacIFN | TCD8ex | FN1 | CD44 | 1.1317 | 0.8219 | 0.9768 | 0.000999 |
| MacIFN | TCD8ex | FN1 | ITGA4 | 1.1317 | 0.4432 | 0.7875 | 0.000999 |
| MacIFN | TCD8ex | FN1 | ITGB1 | 1.1317 | 0.2995 | 0.7156 | 0.000999 |
| MacIFN | TCD8ex | FN1 | ITGB7 | 1.1317 | 0.3007 | 0.7162 | 0.000999 |
| MacIFN | TCD8ex | FN1 | RPSA | 1.1317 | 2.2697 | 1.7007 | 0.000999 |
| MacIFN | TCD8ex | GRN | TNFRSF1B | 1.3662 | 0.6436 | 1.0049 | 0.000999 |
| MacIFN | TCD8ex | GSTP1 | CD44 | 1.2812 | 0.8219 | 1.0516 | 0.000999 |

| | | | | | | | |
|---|---|---|---|---|---|---|---|
| MacIFN | TCD8ex | GSTP1 | ITGA4 | 1.2812 | 0.4432 | 0.8622 | 0.000999 |
| MacIFN | TCD8ex | GSTP1 | VCAM1 | 1.2812 | 0.4462 | 0.8637 | 0.000999 |
| MacIFN | TCD8ex | HBEGF | CD44 | 0.2793 | 0.8219 | 0.5506 | 0.000999 |
| MacIFN | TCD8ex | HLA-DRA | PDCD1 | 3.8224 | 0.6109 | 2.2167 | 0.000999 |
| MacIFN | TCD8ex | HLA-DRA | PTPRC | 3.8224 | 1.6762 | 2.7493 | 0.000999 |
| MacIFN | TCD8ex | ICAM1 | IL2RG | 0.3372 | 1.2126 | 0.7749 | 0.000999 |
| MacIFN | TCD8ex | ICAM1 | ITGB2 | 0.3372 | 0.7635 | 0.5503 | 0.000999 |
| MacIFN | TCD8ex | ICAM1 | SPN | 0.3372 | 0.3486 | 0.3429 | 0.000999 |
| MacIFN | TCD8ex | LGALS3 | CD6 | 1.5031 | 0.318 | 0.9106 | 0.000999 |
| MacIFN | TCD8ex | LGALS3 | ITGB1 | 1.5031 | 0.2995 | 0.9013 | 0.000999 |
| MacIFN | TCD8ex | LGALS3 | SPN | 1.5031 | 0.3486 | 0.9259 | 0.000999 |
| MacIFN | TCD8ex | MIF | CD74 | 1.1447 | 2.5058 | 1.8253 | 0.000999 |
| MacIFN | TCD8ex | MIF | ITGA4 | 1.1447 | 0.4432 | 0.794 | 0.000999 |
| MacIFN | TCD8ex | MIF | TNFRSF14 | 1.1447 | 0.3094 | 0.7271 | 0.000999 |
| MacIFN | TCD8ex | NAMPT | ANXA6 | 0.9717 | 0.5678 | 0.7697 | 0.000999 |
| MacIFN | TCD8ex | RTN4 | CD44 | 0.9008 | 0.8219 | 0.8613 | 0.000999 |
| MacIFN | TCD8ex | RTN4 | CD99 | 0.9008 | 1.2049 | 1.0528 | 0.000999 |
| MacIFN | TCD8ex | SPP1 | CD44 | 3.5345 | 0.8219 | 2.1782 | 0.000999 |
| MacIFN | TCD8ex | SPP1 | ITGA4 | 3.5345 | 0.4432 | 1.9889 | 0.000999 |
| MacIFN | TCD8ex | SPP1 | ITGB1 | 3.5345 | 0.2995 | 1.917 | 0.000999 |
| MacIFN | TCD8ex | TGFB1 | ITGB1 | 0.2566 | 0.2995 | 0.278 | 0.000999 |
| MacIFN | TCD8ex | TNFSF13B | ITGB1 | 0.9552 | 0.2995 | 0.6274 | 0.000999 |
| MacIFN | TCD8ex | HSP90B1 | TNFRSF1B | 0.6822 | 0.6436 | 0.6629 | 0.00999 |
| MacIFN | TGD | A2M | TNFRSF14 | 0.3025 | 0.4642 | 0.3833 | 0.000999 |
| MacIFN | TGD | ADM | PTGDR | 0.627 | 0.7153 | 0.6711 | 0.000999 |
| MacIFN | TGD | AIMP1 | RACK1 | 0.2858 | 2.7287 | 1.5073 | 0.000999 |
| MacIFN | TGD | ANXA1 | CXCR3 | 1.0061 | 1.1748 | 1.0904 | 0.000999 |
| MacIFN | TGD | ANXA1 | CXCR4 | 1.0061 | 1.4894 | 1.2477 | 0.000999 |
| MacIFN | TGD | ANXA1 | ITGA4 | 1.0061 | 0.3968 | 0.7015 | 0.000999 |
| MacIFN | TGD | APOE | SORL1 | 2.0054 | 0.3318 | 1.1686 | 0.000999 |

| | | | | | | | |
|---|---|---|---|---|---|---|---|
| MacIFN | TGD | CALM1 | RPSA | 1.377 | 2.8814 | 2.1292 | 0.000999 |
| MacIFN | TGD | CALR | IL7R | 0.904 | 1.3862 | 1.1451 | 0.000999 |
| MacIFN | TGD | CCL20 | CXCR3 | 0.33 | 1.1748 | 0.7524 | 0.000999 |
| MacIFN | TGD | CCL20 | CXCR4 | 0.33 | 1.4894 | 0.9097 | 0.000999 |
| MacIFN | TGD | CCL7 | CXCR3 | 0.3065 | 1.1748 | 0.7406 | 0.000999 |
| MacIFN | TGD | CD99 | CD99 | 0.4553 | 1.7154 | 1.0854 | 0.000999 |
| MacIFN | TGD | CXCL10 | CXCR3 | 1.153 | 1.1748 | 1.1639 | 0.000999 |
| MacIFN | TGD | CXCL10 | CXCR4 | 1.153 | 1.4894 | 1.3212 | 0.000999 |
| MacIFN | TGD | CXCL11 | CXCR3 | 0.347 | 1.1748 | 0.7609 | 0.000999 |
| MacIFN | TGD | CXCL11 | CXCR4 | 0.347 | 1.4894 | 0.9182 | 0.000999 |
| MacIFN | TGD | CXCL16 | CXCR3 | 0.7145 | 1.1748 | 0.9446 | 0.000999 |
| MacIFN | TGD | CXCL16 | CXCR4 | 0.7145 | 1.4894 | 1.102 | 0.000999 |
| MacIFN | TGD | CXCL2 | CXCR3 | 0.4907 | 1.1748 | 0.8328 | 0.000999 |
| MacIFN | TGD | CXCL2 | CXCR4 | 0.4907 | 1.4894 | 0.9901 | 0.000999 |
| MacIFN | TGD | CXCL3 | CXCR3 | 0.3451 | 1.1748 | 0.7599 | 0.000999 |
| MacIFN | TGD | CXCL3 | CXCR4 | 0.3451 | 1.4894 | 0.9172 | 0.000999 |
| MacIFN | TGD | CXCL9 | CXCR3 | 0.3037 | 1.1748 | 0.7393 | 0.000999 |
| MacIFN | TGD | CXCL9 | CXCR4 | 0.3037 | 1.4894 | 0.8966 | 0.000999 |
| MacIFN | TGD | FN1 | CD44 | 1.1317 | 1.1342 | 1.133 | 0.000999 |
| MacIFN | TGD | FN1 | ITGA4 | 1.1317 | 0.3968 | 0.7643 | 0.000999 |
| MacIFN | TGD | FN1 | ITGB1 | 1.1317 | 0.3657 | 0.7487 | 0.000999 |
| MacIFN | TGD | FN1 | ITGB7 | 1.1317 | 0.459 | 0.7954 | 0.000999 |
| MacIFN | TGD | FN1 | RPSA | 1.1317 | 2.8814 | 2.0065 | 0.000999 |
| MacIFN | TGD | GRN | TNFRSF1A | 1.3662 | 0.3099 | 0.838 | 0.000999 |
| MacIFN | TGD | GSTP1 | IL7R | 1.2812 | 1.3862 | 1.3337 | 0.000999 |
| MacIFN | TGD | GSTP1 | ITGA4 | 1.2812 | 0.3968 | 0.839 | 0.000999 |
| MacIFN | TGD | HLA-A | CD8A | 2.9543 | 1.746 | 2.3502 | 0.000999 |
| MacIFN | TGD | HLA-DRA | PTPRC | 3.8224 | 1.8653 | 2.8439 | 0.000999 |
| MacIFN | TGD | HLA-E | CD8A | 1.6113 | 1.746 | 1.6787 | 0.000999 |
| MacIFN | TGD | HLA-E | KLRD1 | 1.6113 | 1.3451 | 1.4782 | 0.000999 |

| | | | | | | | |
|---|---|---|---|---|---|---|---|
| MacIFN | TGD | HLA-F | CD8A | 0.3189 | 1.746 | 1.0325 | 0.000999 |
| MacIFN | TGD | ICAM1 | IL2RG | 0.3372 | 1.561 | 0.9491 | 0.000999 |
| MacIFN | TGD | ICAM1 | ITGAL | 0.3372 | 0.3731 | 0.3552 | 0.000999 |
| MacIFN | TGD | ICAM1 | ITGB2 | 0.3372 | 1.4444 | 0.8908 | 0.000999 |
| MacIFN | TGD | ICAM1 | SPN | 0.3372 | 0.5644 | 0.4508 | 0.000999 |
| MacIFN | TGD | ITGB1 | ITGA4 | 0.2698 | 0.3968 | 0.3333 | 0.000999 |
| MacIFN | TGD | ITGB1 | PTK2B | 0.2698 | 0.3668 | 0.3183 | 0.000999 |
| MacIFN | TGD | ITGB1 | RACK1 | 0.2698 | 2.7287 | 1.4993 | 0.000999 |
| MacIFN | TGD | LGALS3 | CD6 | 1.5031 | 0.4684 | 0.9857 | 0.000999 |
| MacIFN | TGD | LGALS3 | ITGB1 | 1.5031 | 0.3657 | 0.9344 | 0.000999 |
| MacIFN | TGD | LGALS3 | SPN | 1.5031 | 0.5644 | 1.0337 | 0.000999 |
| MacIFN | TGD | LGALS3 | TGFBR2 | 1.5031 | 0.3677 | 0.9354 | 0.000999 |
| MacIFN | TGD | LRPAP1 | SORL1 | 0.4177 | 0.3318 | 0.3747 | 0.000999 |
| MacIFN | TGD | NAMPT | ANXA6 | 0.9717 | 0.8553 | 0.9135 | 0.000999 |
| MacIFN | TGD | PTPRC | CD2 | 0.4998 | 1.4522 | 0.976 | 0.000999 |
| MacIFN | TGD | PTPRC | CD8A | 0.4998 | 1.746 | 1.1229 | 0.000999 |
| MacIFN | TGD | RPS19 | RPSA | 3.2866 | 2.8814 | 3.084 | 0.000999 |
| MacIFN | TGD | RTN4 | CD99 | 0.9008 | 1.7154 | 1.3081 | 0.000999 |
| MacIFN | TGD | SPP1 | CD44 | 3.5345 | 1.1342 | 2.3343 | 0.000999 |
| MacIFN | TGD | SPP1 | ITGA4 | 3.5345 | 0.3968 | 1.9657 | 0.000999 |
| MacIFN | TGD | SPP1 | ITGB1 | 3.5345 | 0.3657 | 1.9501 | 0.000999 |
| MacIFN | TGD | TNFSF13B | ITGB1 | 0.9552 | 0.3657 | 0.6604 | 0.000999 |
| MacIFN | TGD | VEGFA | PTK2B | 0.5137 | 0.3668 | 0.4402 | 0.000999 |
| MacIFN | TGD | HLA-E | KLRC2 | 1.6113 | 0.8082 | 1.2098 | 0.002 |
| MacIFN | TGD | GSTP1 | CD44 | 1.2812 | 1.1342 | 1.2077 | 0.004 |
| MacIFN | TGD | ARF1 | ITGA4 | 0.5726 | 0.3968 | 0.4847 | 0.005 |
| MacIFN | TGD | RTN4 | CD44 | 0.9008 | 1.1342 | 1.0175 | 0.005 |
| NKT | MacIFN | ANXA1 | C3AR1 | 1.2259 | 0.2967 | 0.7613 | 0.000999 |
| NKT | MacIFN | ANXA1 | C5AR1 | 1.2259 | 0.4502 | 0.8381 | 0.000999 |
| NKT | MacIFN | ANXA1 | CCR1 | 1.2259 | 0.3151 | 0.7705 | 0.000999 |

| | | | | | | | |
|---|---|---|---|---|---|---|---|
| NKT | MacIFN | ANXA1 | CD4 | 1.2259 | 0.33 | 0.778 | 0.000999 |
| NKT | MacIFN | ANXA1 | CD44 | 1.2259 | 1.0945 | 1.1602 | 0.000999 |
| NKT | MacIFN | ANXA1 | FPR1 | 1.2259 | 0.3577 | 0.7918 | 0.000999 |
| NKT | MacIFN | ANXA1 | FPR3 | 1.2259 | 0.4448 | 0.8354 | 0.000999 |
| NKT | MacIFN | ARF1 | CD4 | 0.7991 | 0.33 | 0.5646 | 0.000999 |
| NKT | MacIFN | ARF1 | CD44 | 0.7991 | 1.0945 | 0.9468 | 0.000999 |
| NKT | MacIFN | CALM1 | TNFRSF1A | 1.9335 | 0.2739 | 1.1037 | 0.000999 |
| NKT | MacIFN | CALR | CD44 | 0.9479 | 1.0945 | 1.0212 | 0.000999 |
| NKT | MacIFN | CD2 | CD4 | 1.1465 | 0.33 | 0.7383 | 0.000999 |
| NKT | MacIFN | GSTP1 | CD44 | 0.7909 | 1.0945 | 0.9427 | 0.000999 |
| NKT | MacIFN | HMGB1 | LY96 | 1.195 | 0.8119 | 1.0034 | 0.000999 |
| NKT | MacIFN | HMGB1 | TLR2 | 1.195 | 0.29 | 0.7425 | 0.000999 |
| NKT | MacIFN | HMGB2 | CD44 | 0.6467 | 1.0945 | 0.8706 | 0.000999 |
| NKT | MacIFN | HMGB2 | DAB2 | 0.6467 | 0.4741 | 0.5604 | 0.000999 |
| NKT | MacIFN | IL16 | CD4 | 0.3556 | 0.33 | 0.3428 | 0.000999 |
| NKT | MacIFN | IL16 | FCGR1A | 0.3556 | 0.4375 | 0.3965 | 0.000999 |
| NKT | MacIFN | ITGA4 | CD44 | 0.2946 | 1.0945 | 0.6945 | 0.000999 |
| NKT | MacIFN | ITGA4 | TFRC | 0.2946 | 0.5198 | 0.4072 | 0.000999 |
| NKT | MacIFN | ITGAL | ICAM1 | 0.6778 | 0.3372 | 0.5075 | 0.000999 |
| NKT | MacIFN | ITGB1 | DAB2 | 0.488 | 0.4741 | 0.481 | 0.000999 |
| NKT | MacIFN | ITGB1 | ITGAX | 0.488 | 0.2606 | 0.3743 | 0.000999 |
| NKT | MacIFN | ITGB1 | PLAUR | 0.488 | 1.2551 | 0.8715 | 0.000999 |
| NKT | MacIFN | ITGB2 | ICAM1 | 1.6481 | 0.3372 | 0.9927 | 0.000999 |
| NKT | MacIFN | ITGB2 | ITGAX | 1.6481 | 0.2606 | 0.9544 | 0.000999 |
| NKT | MacIFN | ITGB2 | PLAUR | 1.6481 | 1.2551 | 1.4516 | 0.000999 |
| NKT | MacIFN | LTB | TNFRSF1A | 0.2605 | 0.2739 | 0.2672 | 0.000999 |
| NKT | MacIFN | MIF | CD74 | 0.6657 | 3.8444 | 2.255 | 0.000999 |
| NKT | MacIFN | PTPRC | CD4 | 1.8797 | 0.33 | 1.1049 | 0.000999 |
| NKT | MacIFN | PTPRC | CD44 | 1.8797 | 1.0945 | 1.4871 | 0.000999 |
| NKT | MacIFN | PTPRC | IFNAR1 | 1.8797 | 0.2575 | 1.0686 | 0.000999 |

| | | | | | | | |
|---|---|---|---|---|---|---|---|
| NKT | MacIFN | RPS19 | C5AR1 | 3.5336 | 0.4502 | 1.9919 | 0.000999 |
| NKT | MacIFN | RPS19 | CD4 | 3.5336 | 0.33 | 1.9318 | 0.000999 |
| NKT | MacIFN | RPS19 | ICAM1 | 3.5336 | 0.3372 | 1.9354 | 0.000999 |
| NKT | MacIFN | RTN4 | CD44 | 0.4374 | 1.0945 | 0.766 | 0.000999 |
| NKT | MacIFN | TGFB1 | FCN1 | 0.5441 | 0.5801 | 0.5621 | 0.000999 |
| NKT | MacIFN | TGFB1 | TLR2 | 0.5441 | 0.29 | 0.417 | 0.000999 |
| TCD4ex | MacIFN | ANXA1 | C3AR1 | 0.5174 | 0.2967 | 0.4071 | 0.000999 |
| TCD4ex | MacIFN | ANXA1 | C5AR1 | 0.5174 | 0.4502 | 0.4838 | 0.000999 |
| TCD4ex | MacIFN | ANXA1 | CCR1 | 0.5174 | 0.3151 | 0.4163 | 0.000999 |
| TCD4ex | MacIFN | ANXA1 | FPR1 | 0.5174 | 0.3577 | 0.4376 | 0.000999 |
| TCD4ex | MacIFN | ANXA1 | FPR3 | 0.5174 | 0.4448 | 0.4811 | 0.000999 |
| TCD4ex | MacIFN | AREG | ICAM1 | 0.3427 | 0.3372 | 0.34 | 0.000999 |
| TCD4ex | MacIFN | CALM1 | TNFRSF1A | 1.2957 | 0.2739 | 0.7848 | 0.000999 |
| TCD4ex | MacIFN | CXCL13 | C3AR1 | 2.3367 | 0.2967 | 1.3167 | 0.000999 |
| TCD4ex | MacIFN | CXCL13 | C5AR1 | 2.3367 | 0.4502 | 1.3934 | 0.000999 |
| TCD4ex | MacIFN | CXCL13 | CCR1 | 2.3367 | 0.3151 | 1.3259 | 0.000999 |
| TCD4ex | MacIFN | CXCL13 | FPR1 | 2.3367 | 0.3577 | 1.3472 | 0.000999 |
| TCD4ex | MacIFN | HMGB1 | LY96 | 1.3876 | 0.8119 | 1.0997 | 0.000999 |
| TCD4ex | MacIFN | HMGB1 | TLR2 | 1.3876 | 0.29 | 0.8388 | 0.000999 |
| TCD4ex | MacIFN | HMGB2 | DAB2 | 1.0243 | 0.4741 | 0.7492 | 0.000999 |
| TCD4ex | MacIFN | ICAM3 | ITGB2 | 0.573 | 0.926 | 0.7495 | 0.000999 |
| TCD4ex | MacIFN | ITGAL | ICAM1 | 0.2603 | 0.3372 | 0.2988 | 0.000999 |
| TCD4ex | MacIFN | ITGB1 | DAB2 | 0.4615 | 0.4741 | 0.4678 | 0.000999 |
| TCD4ex | MacIFN | ITGB1 | ITGAX | 0.4615 | 0.2606 | 0.361 | 0.000999 |
| TCD4ex | MacIFN | ITGB1 | PLAUR | 0.4615 | 1.2551 | 0.8583 | 0.000999 |
| TCD4ex | MacIFN | ITGB2 | ICAM1 | 0.7871 | 0.3372 | 0.5621 | 0.000999 |
| TCD4ex | MacIFN | ITGB2 | ITGAX | 0.7871 | 0.2606 | 0.5238 | 0.000999 |
| TCD4ex | MacIFN | ITGB2 | PLAUR | 0.7871 | 1.2551 | 1.0211 | 0.000999 |
| TCD4ex | MacIFN | LTB | TNFRSF1A | 0.9655 | 0.2739 | 0.6197 | 0.000999 |
| TCD4ex | MacIFN | MIF | CD74 | 0.6999 | 3.8444 | 2.2721 | 0.000999 |

| | | | | | | | |
|---|---|---|---|---|---|---|---|
| TCD4ex | MacIFN | PTPRC | CD44 | 1.7493 | 1.0945 | 1.4219 | 0.000999 |
| TCD4ex | MacIFN | PTPRC | IFNAR1 | 1.7493 | 0.2575 | 1.0034 | 0.000999 |
| TCD4ex | MacIFN | RPS19 | C5AR1 | 3.6329 | 0.4502 | 2.0415 | 0.000999 |
| TCD4ex | MacIFN | RPS19 | ICAM1 | 3.6329 | 0.3372 | 1.9851 | 0.000999 |
| TCD4ex | MacIFN | ITGB1 | CD47 | 0.4615 | 0.457 | 0.4593 | 0.002 |
| TCD4reg | MacIFN | CALM1 | TNFRSF1A | 1.534 | 0.2739 | 0.9039 | 0.000999 |
| TCD4reg | MacIFN | HMGB1 | LY96 | 1.2547 | 0.8119 | 1.0333 | 0.000999 |
| TCD4reg | MacIFN | HMGB1 | TLR2 | 1.2547 | 0.29 | 0.7724 | 0.000999 |
| TCD4reg | MacIFN | HMGB2 | DAB2 | 0.616 | 0.4741 | 0.5451 | 0.000999 |
| TCD4reg | MacIFN | ICAM3 | ITGB2 | 0.6663 | 0.926 | 0.7961 | 0.000999 |
| TCD4reg | MacIFN | IL16 | FCGR1A | 0.2528 | 0.4375 | 0.3452 | 0.000999 |
| TCD4reg | MacIFN | ITGB1 | DAB2 | 0.4001 | 0.4741 | 0.4371 | 0.000999 |
| TCD4reg | MacIFN | ITGB1 | ITGAX | 0.4001 | 0.2606 | 0.3303 | 0.000999 |
| TCD4reg | MacIFN | ITGB1 | PLAUR | 0.4001 | 1.2551 | 0.8276 | 0.000999 |
| TCD4reg | MacIFN | ITGB2 | ICAM1 | 0.6238 | 0.3372 | 0.4805 | 0.000999 |
| TCD4reg | MacIFN | ITGB2 | ITGAX | 0.6238 | 0.2606 | 0.4422 | 0.000999 |
| TCD4reg | MacIFN | ITGB2 | PLAUR | 0.6238 | 1.2551 | 0.9394 | 0.000999 |
| TCD4reg | MacIFN | LGALS3 | CLEC7A | 0.4492 | 0.5065 | 0.4778 | 0.000999 |
| TCD4reg | MacIFN | LGALS3 | FCGR2A | 0.4492 | 0.7338 | 0.5915 | 0.000999 |
| TCD4reg | MacIFN | LGALS3 | FCGR2B | 0.4492 | 0.5644 | 0.5068 | 0.000999 |
| TCD4reg | MacIFN | LGALS3 | TFRC | 0.4492 | 0.5198 | 0.4845 | 0.000999 |
| TCD4reg | MacIFN | LTB | TNFRSF1A | 2.242 | 0.2739 | 1.258 | 0.000999 |
| TCD4reg | MacIFN | MIF | CD74 | 0.6295 | 3.8444 | 2.237 | 0.000999 |
| TCD4reg | MacIFN | PTPRC | IFNAR1 | 1.6603 | 0.2575 | 0.9589 | 0.000999 |
| TCD4reg | MacIFN | RPS19 | C5AR1 | 3.6897 | 0.4502 | 2.0699 | 0.000999 |
| TCD4reg | MacIFN | RPS19 | ICAM1 | 3.6897 | 0.3372 | 2.0134 | 0.000999 |
| TCD8em | MacIFN | ANXA1 | C3AR1 | 1.0693 | 0.2967 | 0.683 | 0.000999 |
| TCD8em | MacIFN | ANXA1 | C5AR1 | 1.0693 | 0.4502 | 0.7597 | 0.000999 |
| TCD8em | MacIFN | ANXA1 | CCR1 | 1.0693 | 0.3151 | 0.6922 | 0.000999 |
| TCD8em | MacIFN | ANXA1 | CD4 | 1.0693 | 0.33 | 0.6997 | 0.000999 |

| | | | | | | | |
|---|---|---|---|---|---|---|---|
| TCD8em | MacIFN | ANXA1 | FPR1 | 1.0693 | 0.3577 | 0.7135 | 0.000999 |
| TCD8em | MacIFN | ANXA1 | FPR3 | 1.0693 | 0.4448 | 0.757 | 0.000999 |
| TCD8em | MacIFN | ARF1 | CD4 | 0.6684 | 0.33 | 0.4992 | 0.000999 |
| TCD8em | MacIFN | CALM1 | TNFRSF1A | 1.5262 | 0.2739 | 0.9 | 0.000999 |
| TCD8em | MacIFN | CALM1 | TNFRSF1B | 1.5262 | 0.439 | 0.9826 | 0.000999 |
| TCD8em | MacIFN | CD2 | CD4 | 1.5961 | 0.33 | 0.9631 | 0.000999 |
| TCD8em | MacIFN | HLA-DMA | CD4 | 0.2969 | 0.33 | 0.3135 | 0.000999 |
| TCD8em | MacIFN | HLA-DRA | CD4 | 0.952 | 0.33 | 0.641 | 0.000999 |
| TCD8em | MacIFN | HMGB1 | LY96 | 1.3636 | 0.8119 | 1.0877 | 0.000999 |
| TCD8em | MacIFN | HMGB1 | TLR2 | 1.3636 | 0.29 | 0.8268 | 0.000999 |
| TCD8em | MacIFN | HMGB2 | DAB2 | 0.7212 | 0.4741 | 0.5976 | 0.000999 |
| TCD8em | MacIFN | IFNG | IFNGR1 | 0.3967 | 0.3707 | 0.3837 | 0.000999 |
| TCD8em | MacIFN | IFNG | IFNGR2 | 0.3967 | 0.7235 | 0.5601 | 0.000999 |
| TCD8em | MacIFN | IL16 | CD4 | 0.4085 | 0.33 | 0.3693 | 0.000999 |
| TCD8em | MacIFN | IL16 | FCGR1A | 0.4085 | 0.4375 | 0.423 | 0.000999 |
| TCD8em | MacIFN | ITGA4 | TFRC | 0.2823 | 0.5198 | 0.4011 | 0.000999 |
| TCD8em | MacIFN | ITGAL | ICAM1 | 0.3162 | 0.3372 | 0.3267 | 0.000999 |
| TCD8em | MacIFN | ITGB1 | DAB2 | 0.2557 | 0.4741 | 0.3649 | 0.000999 |
| TCD8em | MacIFN | ITGB1 | ITGAX | 0.2557 | 0.2606 | 0.2582 | 0.000999 |
| TCD8em | MacIFN | ITGB1 | PLAUR | 0.2557 | 1.2551 | 0.7554 | 0.000999 |
| TCD8em | MacIFN | ITGB2 | ICAM1 | 1.0018 | 0.3372 | 0.6695 | 0.000999 |
| TCD8em | MacIFN | ITGB2 | ITGAX | 1.0018 | 0.2606 | 0.6312 | 0.000999 |
| TCD8em | MacIFN | ITGB2 | PLAUR | 1.0018 | 1.2551 | 1.1284 | 0.000999 |
| TCD8em | MacIFN | LGALS3 | CLEC7A | 0.2689 | 0.5065 | 0.3877 | 0.000999 |
| TCD8em | MacIFN | LGALS3 | FCGR2A | 0.2689 | 0.7338 | 0.5014 | 0.000999 |
| TCD8em | MacIFN | LGALS3 | FCGR2B | 0.2689 | 0.5644 | 0.4167 | 0.000999 |
| TCD8em | MacIFN | LGALS3 | TFRC | 0.2689 | 0.5198 | 0.3944 | 0.000999 |
| TCD8em | MacIFN | LTB | TNFRSF1A | 0.6352 | 0.2739 | 0.4545 | 0.000999 |
| TCD8em | MacIFN | MIF | CD74 | 0.6063 | 3.8444 | 2.2254 | 0.000999 |
| TCD8em | MacIFN | PTPRC | CD4 | 1.7912 | 0.33 | 1.0606 | 0.000999 |

| | | | | | | | |
|---|---|---|---|---|---|---|---|
| TCD8em | MacIFN | PTPRC | IFNAR1 | 1.7912 | 0.2575 | 1.0243 | 0.000999 |
| TCD8em | MacIFN | RPS19 | C5AR1 | 3.536 | 0.4502 | 1.9931 | 0.000999 |
| TCD8em | MacIFN | RPS19 | CD4 | 3.536 | 0.33 | 1.933 | 0.000999 |
| TCD8em | MacIFN | RPS19 | ICAM1 | 3.536 | 0.3372 | 1.9366 | 0.000999 |
| TCD8em | MacIFN | SELL | CD4 | 0.258 | 0.33 | 0.294 | 0.000999 |
| TCD8em | MacIFN | TNF | TNFRSF1A | 0.2667 | 0.2739 | 0.2703 | 0.000999 |
| TCD8em | MacIFN | TNF | TNFRSF1B | 0.2667 | 0.439 | 0.3529 | 0.000999 |
| TCD8em | MacIFN | HSP90B1 | TNFRSF1B | 0.471 | 0.439 | 0.455 | 0.002 |
| TCD8ex | MacIFN | AIMP1 | RRBP1 | 0.2508 | 0.2512 | 0.251 | 0.000999 |
| TCD8ex | MacIFN | ANXA1 | C3AR1 | 0.5645 | 0.2967 | 0.4306 | 0.000999 |
| TCD8ex | MacIFN | ANXA1 | C5AR1 | 0.5645 | 0.4502 | 0.5073 | 0.000999 |
| TCD8ex | MacIFN | ANXA1 | CCR1 | 0.5645 | 0.3151 | 0.4398 | 0.000999 |
| TCD8ex | MacIFN | ANXA1 | CD4 | 0.5645 | 0.33 | 0.4473 | 0.000999 |
| TCD8ex | MacIFN | ANXA1 | CD44 | 0.5645 | 1.0945 | 0.8295 | 0.000999 |
| TCD8ex | MacIFN | ANXA1 | FPR1 | 0.5645 | 0.3577 | 0.4611 | 0.000999 |
| TCD8ex | MacIFN | ANXA1 | FPR3 | 0.5645 | 0.4448 | 0.5046 | 0.000999 |
| TCD8ex | MacIFN | AREG | ICAM1 | 0.2758 | 0.3372 | 0.3065 | 0.000999 |
| TCD8ex | MacIFN | ARF1 | CD4 | 0.7835 | 0.33 | 0.5568 | 0.000999 |
| TCD8ex | MacIFN | ARF1 | CD44 | 0.7835 | 1.0945 | 0.939 | 0.000999 |
| TCD8ex | MacIFN | CALM1 | TNFRSF1A | 1.4627 | 0.2739 | 0.8683 | 0.000999 |
| TCD8ex | MacIFN | CALR | CD44 | 0.9634 | 1.0945 | 1.0289 | 0.000999 |
| TCD8ex | MacIFN | CD2 | CD4 | 2.0502 | 0.33 | 1.1901 | 0.000999 |
| TCD8ex | MacIFN | CXCL13 | C3AR1 | 1.7749 | 0.2967 | 1.0358 | 0.000999 |
| TCD8ex | MacIFN | CXCL13 | C5AR1 | 1.7749 | 0.4502 | 1.1126 | 0.000999 |
| TCD8ex | MacIFN | CXCL13 | CCR1 | 1.7749 | 0.3151 | 1.045 | 0.000999 |
| TCD8ex | MacIFN | CXCL13 | FPR1 | 1.7749 | 0.3577 | 1.0663 | 0.000999 |
| TCD8ex | MacIFN | FASLG | TNFRSF1A | 0.2613 | 0.2739 | 0.2676 | 0.000999 |
| TCD8ex | MacIFN | GSTP1 | CD44 | 0.8286 | 1.0945 | 0.9616 | 0.000999 |
| TCD8ex | MacIFN | HLA-DMA | CD4 | 0.3991 | 0.33 | 0.3646 | 0.000999 |
| TCD8ex | MacIFN | HLA-DRA | CD4 | 1.247 | 0.33 | 0.7885 | 0.000999 |

| | | | | | | | |
|---|---|---|---|---|---|---|---|
| TCD8ex | MacIFN | HMGB1 | LY96 | 1.5414 | 0.8119 | 1.1767 | 0.000999 |
| TCD8ex | MacIFN | HMGB1 | TLR2 | 1.5414 | 0.29 | 0.9157 | 0.000999 |
| TCD8ex | MacIFN | HMGB2 | CD44 | 0.85 | 1.0945 | 0.9723 | 0.000999 |
| TCD8ex | MacIFN | HMGB2 | DAB2 | 0.85 | 0.4741 | 0.6621 | 0.000999 |
| TCD8ex | MacIFN | ICAM3 | ITGB2 | 0.5405 | 0.926 | 0.7332 | 0.000999 |
| TCD8ex | MacIFN | IFNG | IFNGR1 | 0.8575 | 0.3707 | 0.6141 | 0.000999 |
| TCD8ex | MacIFN | IFNG | IFNGR2 | 0.8575 | 0.7235 | 0.7905 | 0.000999 |
| TCD8ex | MacIFN | ITGA4 | CD44 | 0.4432 | 1.0945 | 0.7689 | 0.000999 |
| TCD8ex | MacIFN | ITGA4 | TFRC | 0.4432 | 0.5198 | 0.4815 | 0.000999 |
| TCD8ex | MacIFN | ITGB1 | DAB2 | 0.2995 | 0.4741 | 0.3868 | 0.000999 |
| TCD8ex | MacIFN | ITGB1 | ITGAX | 0.2995 | 0.2606 | 0.28 | 0.000999 |
| TCD8ex | MacIFN | ITGB1 | PLAUR | 0.2995 | 1.2551 | 0.7773 | 0.000999 |
| TCD8ex | MacIFN | ITGB2 | ICAM1 | 0.7635 | 0.3372 | 0.5503 | 0.000999 |
| TCD8ex | MacIFN | ITGB2 | ITGAX | 0.7635 | 0.2606 | 0.512 | 0.000999 |
| TCD8ex | MacIFN | ITGB2 | PLAUR | 0.7635 | 1.2551 | 1.0093 | 0.000999 |
| TCD8ex | MacIFN | LGALS3 | CLEC7A | 0.3824 | 0.5065 | 0.4444 | 0.000999 |
| TCD8ex | MacIFN | LGALS3 | FCGR2A | 0.3824 | 0.7338 | 0.5581 | 0.000999 |
| TCD8ex | MacIFN | LGALS3 | FCGR2B | 0.3824 | 0.5644 | 0.4734 | 0.000999 |
| TCD8ex | MacIFN | LGALS3 | TFRC | 0.3824 | 0.5198 | 0.4511 | 0.000999 |
| TCD8ex | MacIFN | LTB | TNFRSF1A | 0.4576 | 0.2739 | 0.3657 | 0.000999 |
| TCD8ex | MacIFN | MIF | CD74 | 0.749 | 3.8444 | 2.2967 | 0.000999 |
| TCD8ex | MacIFN | PTPRC | CD4 | 1.6762 | 0.33 | 1.0031 | 0.000999 |
| TCD8ex | MacIFN | PTPRC | CD44 | 1.6762 | 1.0945 | 1.3853 | 0.000999 |
| TCD8ex | MacIFN | PTPRC | IFNAR1 | 1.6762 | 0.2575 | 0.9668 | 0.000999 |
| TCD8ex | MacIFN | RPS19 | C5AR1 | 3.8353 | 0.4502 | 2.1427 | 0.000999 |
| TCD8ex | MacIFN | RPS19 | CD4 | 3.8353 | 0.33 | 2.0827 | 0.000999 |
| TCD8ex | MacIFN | RPS19 | ICAM1 | 3.8353 | 0.3372 | 2.0862 | 0.000999 |
| TCD8ex | MacIFN | RTN4 | CD44 | 0.3033 | 1.0945 | 0.6989 | 0.000999 |
| TCD8ex | MacIFN | VCAM1 | TFRC | 0.4462 | 0.5198 | 0.483 | 0.000999 |
| TCD8ex | MacIFN | VCAM1 | ITGB2 | 0.4462 | 0.926 | 0.6861 | 0.002 |

| | | | | | | | |
|---|---|---|---|---|---|---|---|
| TGD | MacIFN | ARF1 | CD4 | 0.9453 | 0.33 | 0.6377 | 0.000999 |
| TGD | MacIFN | ARF1 | CD44 | 0.9453 | 1.0945 | 1.0199 | 0.000999 |
| TGD | MacIFN | CALM1 | TNFRSF1A | 1.7587 | 0.2739 | 1.0163 | 0.000999 |
| TGD | MacIFN | CALM1 | TNFRSF1B | 1.7587 | 0.439 | 1.0989 | 0.000999 |
| TGD | MacIFN | CD2 | CD4 | 1.4522 | 0.33 | 0.8911 | 0.000999 |
| TGD | MacIFN | CD2 | PTPRC | 1.4522 | 0.4998 | 0.976 | 0.000999 |
| TGD | MacIFN | CD99 | CD99 | 1.7154 | 0.4553 | 1.0854 | 0.000999 |
| TGD | MacIFN | HMGB1 | LY96 | 1.7525 | 0.8119 | 1.2822 | 0.000999 |
| TGD | MacIFN | HMGB1 | TLR2 | 1.7525 | 0.29 | 1.0212 | 0.000999 |
| TGD | MacIFN | HSP90B1 | TNFRSF1B | 0.9338 | 0.439 | 0.6864 | 0.000999 |
| TGD | MacIFN | ICAM3 | ITGB2 | 0.8816 | 0.926 | 0.9038 | 0.000999 |
| TGD | MacIFN | IL16 | CD4 | 0.9357 | 0.33 | 0.6329 | 0.000999 |
| TGD | MacIFN | IL16 | FCGR1A | 0.9357 | 0.4375 | 0.6866 | 0.000999 |
| TGD | MacIFN | ITGA4 | CD44 | 0.3968 | 1.0945 | 0.7457 | 0.000999 |
| TGD | MacIFN | ITGA4 | TFRC | 0.3968 | 0.5198 | 0.4583 | 0.000999 |
| TGD | MacIFN | ITGAL | ICAM1 | 0.3731 | 0.3372 | 0.3552 | 0.000999 |
| TGD | MacIFN | ITGB1 | DAB2 | 0.3657 | 0.4741 | 0.4199 | 0.000999 |
| TGD | MacIFN | ITGB1 | ITGAX | 0.3657 | 0.2606 | 0.3131 | 0.000999 |
| TGD | MacIFN | ITGB1 | PLAUR | 0.3657 | 1.2551 | 0.8104 | 0.000999 |
| TGD | MacIFN | ITGB2 | ICAM1 | 1.4444 | 0.3372 | 0.8908 | 0.000999 |
| TGD | MacIFN | ITGB2 | ITGAX | 1.4444 | 0.2606 | 0.8525 | 0.000999 |
| TGD | MacIFN | ITGB2 | PLAUR | 1.4444 | 1.2551 | 1.3498 | 0.000999 |
| TGD | MacIFN | LTB | ITGB1 | 1.6829 | 0.2698 | 0.9764 | 0.000999 |
| TGD | MacIFN | LTB | TNFRSF1A | 1.6829 | 0.2739 | 0.9784 | 0.000999 |
| TGD | MacIFN | MIF | CD74 | 1.6335 | 3.8444 | 2.7389 | 0.000999 |
| TGD | MacIFN | PTPRC | CD4 | 1.8653 | 0.33 | 1.0977 | 0.000999 |
| TGD | MacIFN | PTPRC | CD44 | 1.8653 | 1.0945 | 1.4799 | 0.000999 |
| TGD | MacIFN | PTPRC | CXCR4 | 1.8653 | 1.1937 | 1.5295 | 0.000999 |
| TGD | MacIFN | PTPRC | IFNAR1 | 1.8653 | 0.2575 | 1.0614 | 0.000999 |
| TGD | MacIFN | RPS19 | C5AR1 | 3.8902 | 0.4502 | 2.1702 | 0.000999 |

| | | | | | | | |
|---|---|---|---|---|---|---|---|
| TGD | MacIFN | RPS19 | CD4 | 3.8902 | 0.33 | 2.1101 | 0.000999 |
| TGD | MacIFN | RPS19 | ICAM1 | 3.8902 | 0.3372 | 2.1137 | 0.000999 |
| TGD | MacIFN | SELL | CD4 | 0.3555 | 0.33 | 0.3428 | 0.000999 |
| TGD | MacIFN | TGFB1 | FCN1 | 0.7703 | 0.5801 | 0.6752 | 0.000999 |
| TGD | MacIFN | TGFB1 | ITGB1 | 0.7703 | 0.2698 | 0.5201 | 0.000999 |
| TGD | MacIFN | TGFB1 | TLR2 | 0.7703 | 0.29 | 0.5302 | 0.000999 |
| TGD | MacIFN | ITGA4 | CXCR4 | 0.3968 | 1.1937 | 0.7953 | 0.002 |

| Supplementary Table 8: Communication results between macrophages and lymphocytes subpopulations, and malignant cells | | | | | | | |
|---|---|---|---|---|---|---|---|
| **MacIFN vs. malignant cells** | | | | | | | |
| celltype_01 | celltype_02 | ligand | receptor | ligand_mean_expression | receptor_mean_expression | ligand_receptor_mean_expression | pvalue |
| MacIFN | breasttumor | AIMP1 | RACK1 | 0.282 | 2.6334 | 1.4577 | 0.000999 |
| MacIFN | breasttumor | ANXA1 | CD44 | 1.0054 | 0.3247 | 0.6651 | 0.000999 |
| MacIFN | breasttumor | APOE | SCARB1 | 2.0034 | 0.2628 | 1.1331 | 0.000999 |
| MacIFN | breasttumor | APOE | SDC2 | 2.0034 | 1.071 | 1.5372 | 0.000999 |
| MacIFN | breasttumor | CALM1 | MYLK | 1.3759 | 0.4064 | 0.8911 | 0.000999 |
| MacIFN | breasttumor | CALM1 | RPSA | 1.3759 | 2.358 | 1.8669 | 0.000999 |
| MacIFN | breasttumor | CALM1 | TNFRSF1A | 1.3759 | 0.2536 | 0.8147 | 0.000999 |
| MacIFN | breasttumor | CALM2 | MYLK | 1.6125 | 0.4064 | 1.0094 | 0.000999 |
| MacIFN | breasttumor | CALM3 | MYLK | 0.7317 | 0.4064 | 0.569 | 0.000999 |
| MacIFN | breasttumor | COL1A1 | ITGB1 | 0.3927 | 0.2939 | 0.3433 | 0.000999 |
| MacIFN | breasttumor | FN1 | CD44 | 1.1314 | 0.3247 | 0.728 | 0.000999 |
| MacIFN | breasttumor | FN1 | ITGB1 | 1.1314 | 0.2939 | 0.7126 | 0.000999 |
| MacIFN | breasttumor | FN1 | RPSA | 1.1314 | 2.358 | 1.7447 | 0.000999 |
| MacIFN | breasttumor | FN1 | SDC2 | 1.1314 | 1.071 | 1.1012 | 0.000999 |
| MacIFN | breasttumor | GRN | TNFRSF1A | 1.3664 | 0.2536 | 0.81 | 0.000999 |
| MacIFN | breasttumor | HLA-DRA | SCARB1 | 3.8213 | 0.2628 | 2.042 | 0.000999 |
| MacIFN | breasttumor | ITGB1 | RACK1 | 0.2705 | 2.6334 | 1.4519 | 0.000999 |
| MacIFN | breasttumor | LGALS3 | ITGB1 | 1.5068 | 0.2939 | 0.9003 | 0.000999 |
| MacIFN | breasttumor | PTPRC | CD44 | 0.5011 | 0.3247 | 0.4129 | 0.000999 |
| MacIFN | breasttumor | RTN4 | CD44 | 0.8998 | 0.3247 | 0.6123 | 0.000999 |
| MacIFN | breasttumor | RTN4 | CD99 | 0.8998 | 0.3871 | 0.6434 | 0.000999 |
| MacIFN | breasttumor | SERPINA1 | SDC2 | 1.0423 | 1.071 | 1.0567 | 0.000999 |
| MacIFN | breasttumor | SPP1 | CD44 | 3.5301 | 0.3247 | 1.9274 | 0.000999 |
| MacIFN | breasttumor | SPP1 | ITGB1 | 3.5301 | 0.2939 | 1.912 | 0.000999 |
| MacIFN | breasttumor | TGFB1 | ITGB1 | 0.2572 | 0.2939 | 0.2755 | 0.000999 |
| MacIFN | breasttumor | TGFB1 | SDC2 | 0.2572 | 1.071 | 0.6641 | 0.000999 |
| MacIFN | breasttumor | TNFSF13B | ITGB1 | 0.9576 | 0.2939 | 0.6257 | 0.000999 |

| | | | | | | | |
|---|---|---|---|---|---|---|---|
| MacIFN | breasttumor | HBEGF | CD44 | 0.28 | 0.3247 | 0.3024 | 0.002 |
| MacIFN | breasttumor | AIMP1 | RRBP1 | 0.282 | 0.2835 | 0.2827 | 0.005 |
| MacIFN | colorectaltumor | AIMP1 | RACK1 | 0.282 | 2.7146 | 1.4983 | 0.000999 |
| MacIFN | colorectaltumor | APOE | SDC1 | 2.0034 | 0.2575 | 1.1304 | 0.000999 |
| MacIFN | colorectaltumor | CALM1 | RPSA | 1.3759 | 2.251 | 1.8135 | 0.000999 |
| MacIFN | colorectaltumor | FN1 | ITGB1 | 1.1314 | 0.4214 | 0.7764 | 0.000999 |
| MacIFN | colorectaltumor | FN1 | RPSA | 1.1314 | 2.251 | 1.6912 | 0.000999 |
| MacIFN | colorectaltumor | FN1 | SDC1 | 1.1314 | 0.2575 | 0.6944 | 0.000999 |
| MacIFN | colorectaltumor | ITGB1 | CD151 | 0.2705 | 0.6439 | 0.4572 | 0.000999 |
| MacIFN | colorectaltumor | ITGB1 | RACK1 | 0.2705 | 2.7146 | 1.4925 | 0.000999 |
| MacIFN | colorectaltumor | RTN4 | CD99 | 0.8998 | 0.3443 | 0.622 | 0.000999 |
| MacIFN | colorectaltumor | SPP1 | CD44 | 3.5301 | 0.2512 | 1.8907 | 0.000999 |
| MacIFN | colorectaltumor | SPP1 | ITGB1 | 3.5301 | 0.4214 | 1.9758 | 0.000999 |
| MacIFN | colorectaltumor | SPP1 | SDC1 | 3.5301 | 0.2575 | 1.8938 | 0.000999 |
| MacIFN | colorectaltumor | TGFB1 | ITGB1 | 0.2572 | 0.4214 | 0.3393 | 0.000999 |
| MacIFN | colorectaltumor | TNFSF13B | ITGB1 | 0.9576 | 0.4214 | 0.6895 | 0.000999 |
| MacIFN | colorectaltumor | HSP90B1 | ITGB1 | 0.6773 | 0.4214 | 0.5493 | 0.002 |
| MacIFN | colorectaltumor | ITGB1 | CDH17 | 0.2705 | 0.254 | 0.2622 | 0.002 |
| MacIFN | colorectaltumor | LGALS3 | ITGB4 | 1.5068 | 0.2913 | 0.8991 | 0.004 |
| MacIFN | colorectaltumor | COL1A1 | SDC1 | 0.3927 | 0.2575 | 0.3251 | 0.00899 |
| MacIFN | livertumor | A2M | TNFRSF14 | 0.3032 | 0.2788 | 0.291 | 0.000999 |
| MacIFN | livertumor | ADM | RAMP1 | 0.6215 | 0.4499 | 0.5357 | 0.000999 |
| MacIFN | livertumor | AIMP1 | RACK1 | 0.282 | 2.7655 | 1.5237 | 0.000999 |
| MacIFN | livertumor | ANXA1 | CD44 | 1.0054 | 0.8016 | 0.9035 | 0.000999 |
| MacIFN | livertumor | ANXA1 | MET | 1.0054 | 0.2983 | 0.6518 | 0.000999 |
| MacIFN | livertumor | ANXA1 | PTPRF | 1.0054 | 0.384 | 0.6947 | 0.000999 |
| MacIFN | livertumor | APOE | SDC1 | 2.0034 | 0.5354 | 1.2694 | 0.000999 |
| MacIFN | livertumor | APOE | SDC4 | 2.0034 | 0.6068 | 1.3051 | 0.000999 |
| MacIFN | livertumor | CALM1 | INSR | 1.3759 | 0.3039 | 0.8399 | 0.000999 |
| MacIFN | livertumor | CALM1 | RPSA | 1.3759 | 2.6317 | 2.0038 | 0.000999 |

| | | | | | | | |
|---|---|---|---|---|---|---|---|
| MacIFN | livertumor | CD99 | EPHB4 | 0.4533 | 0.4535 | 0.4534 | 0.000999 |
| MacIFN | livertumor | COL1A1 | ITGA2 | 0.3927 | 0.3983 | 0.3955 | 0.000999 |
| MacIFN | livertumor | COL1A1 | ITGA3 | 0.3927 | 0.3913 | 0.392 | 0.000999 |
| MacIFN | livertumor | COL1A1 | ITGB1 | 0.3927 | 1.6044 | 0.9986 | 0.000999 |
| MacIFN | livertumor | COL1A1 | SDC1 | 0.3927 | 0.5354 | 0.4641 | 0.000999 |
| MacIFN | livertumor | FN1 | CD44 | 1.1314 | 0.8016 | 0.9665 | 0.000999 |
| MacIFN | livertumor | FN1 | ITGA2 | 1.1314 | 0.3983 | 0.7648 | 0.000999 |
| MacIFN | livertumor | FN1 | ITGA3 | 1.1314 | 0.3913 | 0.7613 | 0.000999 |
| MacIFN | livertumor | FN1 | ITGB1 | 1.1314 | 1.6044 | 1.3679 | 0.000999 |
| MacIFN | livertumor | FN1 | RPSA | 1.1314 | 2.6317 | 1.8815 | 0.000999 |
| MacIFN | livertumor | FN1 | SDC1 | 1.1314 | 0.5354 | 0.8334 | 0.000999 |
| MacIFN | livertumor | FN1 | TFRC | 1.1314 | 0.2899 | 0.7106 | 0.000999 |
| MacIFN | livertumor | HAMP | SLC40A1 | 0.4127 | 0.4972 | 0.455 | 0.000999 |
| MacIFN | livertumor | HBEGF | ERBB3 | 0.28 | 0.2644 | 0.2722 | 0.000999 |
| MacIFN | livertumor | HLA-A | ERBB3 | 2.9518 | 0.2644 | 1.6081 | 0.000999 |
| MacIFN | livertumor | HLA-DRA | EPHB4 | 3.8213 | 0.4535 | 2.1374 | 0.000999 |
| MacIFN | livertumor | HLA-E | EPHB4 | 1.6121 | 0.4535 | 1.0328 | 0.000999 |
| MacIFN | livertumor | HLA-E | ITGA6 | 1.6121 | 0.5211 | 1.0666 | 0.000999 |
| MacIFN | livertumor | HLA-E | PLXNB2 | 1.6121 | 0.265 | 0.9386 | 0.000999 |
| MacIFN | livertumor | HMGB2 | CDH1 | 0.5207 | 0.4896 | 0.5052 | 0.000999 |
| MacIFN | livertumor | HMGB2 | DSG2 | 0.5207 | 0.7479 | 0.6343 | 0.000999 |
| MacIFN | livertumor | HSP90B1 | ITGB1 | 0.6773 | 1.6044 | 1.1409 | 0.000999 |
| MacIFN | livertumor | LGALS3 | ITGA2 | 1.5068 | 0.3983 | 0.9526 | 0.000999 |
| MacIFN | livertumor | LGALS3 | ITGB1 | 1.5068 | 1.6044 | 1.5556 | 0.000999 |
| MacIFN | livertumor | LGALS3 | ITGB4 | 1.5068 | 0.5518 | 1.0293 | 0.000999 |
| MacIFN | livertumor | NAMPT | INSR | 0.9695 | 0.3039 | 0.6367 | 0.000999 |
| MacIFN | livertumor | PTPRC | INSR | 0.5011 | 0.3039 | 0.4025 | 0.000999 |
| MacIFN | livertumor | PTPRC | MET | 0.5011 | 0.2983 | 0.3997 | 0.000999 |
| MacIFN | livertumor | RPS19 | RPSA | 3.2857 | 2.6317 | 2.9587 | 0.000999 |
| MacIFN | livertumor | SPP1 | CD44 | 3.5301 | 0.8016 | 2.1659 | 0.000999 |

| | | | | | | | |
|---|---|---|---|---|---|---|---|
| MacIFN | livertumor | SPP1 | ITGB1 | 3.5301 | 1.6044 | 2.5673 | 0.000999 |
| MacIFN | livertumor | SPP1 | SDC1 | 3.5301 | 0.5354 | 2.0328 | 0.000999 |
| MacIFN | livertumor | TGFB1 | ITGB1 | 0.2572 | 1.6044 | 0.9308 | 0.000999 |
| MacIFN | livertumor | TGFB1 | VTN | 0.2572 | 0.7399 | 0.4986 | 0.000999 |
| MacIFN | livertumor | TNFSF13B | ITGB1 | 0.9576 | 1.6044 | 1.281 | 0.000999 |
| MacIFN | livertumor | VEGFA | VTN | 0.5118 | 0.7399 | 0.6258 | 0.000999 |
| MacIFN | livertumor | LGALS3 | TFRC | 1.5068 | 0.2899 | 0.8983 | 0.002 |
| MacIFN | livertumor | CALR | ITGA3 | 0.8992 | 0.3913 | 0.6452 | 0.004 |
| MacIFN | livertumor | CD99 | INSR | 0.4533 | 0.3039 | 0.3786 | 0.005 |
| MacIFN | livertumor | CD99 | CD99 | 0.4533 | 0.5906 | 0.5219 | 0.00799 |
| MacIFN | lungtumor | APOE | SDC1 | 2.0034 | 0.2715 | 1.1374 | 0.000999 |
| MacIFN | lungtumor | APOE | SDC4 | 2.0034 | 0.3516 | 1.1775 | 0.000999 |
| MacIFN | lungtumor | CALM1 | RPSA | 1.3759 | 1.6057 | 1.4908 | 0.000999 |
| MacIFN | lungtumor | COL1A1 | CD44 | 0.3927 | 0.3802 | 0.3864 | 0.000999 |
| MacIFN | lungtumor | COL1A1 | SDC1 | 0.3927 | 0.2715 | 0.3321 | 0.000999 |
| MacIFN | lungtumor | CXCL10 | CXCR4 | 1.1558 | 0.2684 | 0.7121 | 0.000999 |
| MacIFN | lungtumor | CXCL11 | CXCR4 | 0.3478 | 0.2684 | 0.3081 | 0.000999 |
| MacIFN | lungtumor | CXCL16 | CXCR4 | 0.7131 | 0.2684 | 0.4907 | 0.000999 |
| MacIFN | lungtumor | CXCL2 | CXCR4 | 0.492 | 0.2684 | 0.3802 | 0.000999 |
| MacIFN | lungtumor | CXCL3 | CXCR4 | 0.3459 | 0.2684 | 0.3072 | 0.000999 |
| MacIFN | lungtumor | CXCL9 | CXCR4 | 0.3045 | 0.2684 | 0.2865 | 0.000999 |
| MacIFN | lungtumor | FN1 | CD44 | 1.1314 | 0.3802 | 0.7558 | 0.000999 |
| MacIFN | lungtumor | FN1 | RPSA | 1.1314 | 1.6057 | 1.3685 | 0.000999 |
| MacIFN | lungtumor | FN1 | SDC1 | 1.1314 | 0.2715 | 0.7014 | 0.000999 |
| MacIFN | lungtumor | HBEGF | CD44 | 0.28 | 0.3802 | 0.3301 | 0.000999 |
| MacIFN | lungtumor | PTPRC | CD44 | 0.5011 | 0.3802 | 0.4406 | 0.000999 |
| MacIFN | lungtumor | PTPRC | CXCR4 | 0.5011 | 0.2684 | 0.3847 | 0.000999 |
| MacIFN | lungtumor | RTN4 | CD44 | 0.8998 | 0.3802 | 0.64 | 0.000999 |
| MacIFN | lungtumor | RTN4 | CD99 | 0.8998 | 0.5938 | 0.7468 | 0.000999 |
| MacIFN | lungtumor | SPP1 | CD44 | 3.5301 | 0.3802 | 1.9552 | 0.000999 |

| | | | | | | | |
|---|---|---|---|---|---|---|---|
| MacIFN | lungtumor | SPP1 | SDC1 | 3.5301 | 0.2715 | 1.9008 | 0.000999 |
| MacIFN | melanomatumo | AIMP1 | RACK1 | 0.282 | 4.145 | 2.2135 | 0.000999 |
| MacIFN | melanomatumo | ANXA1 | EDNRB | 1.0054 | 0.5666 | 0.786 | 0.000999 |
| MacIFN | melanomatumo | CALM1 | RPSA | 1.3759 | 3.7787 | 2.5773 | 0.000999 |
| MacIFN | melanomatumo | FN1 | RPSA | 1.1314 | 3.7787 | 2.455 | 0.000999 |
| MacIFN | melanomatumo | ITGB1 | RACK1 | 0.2705 | 4.145 | 2.2078 | 0.000999 |
| MacIFN | melanomatumo | NAMPT | ANXA6 | 0.9695 | 0.3625 | 0.666 | 0.000999 |
| MacIFN | melanomatumo | RPS19 | RPSA | 3.2857 | 3.7787 | 3.5322 | 0.000999 |
| MacIFN | melanomatumo | SPP1 | ITGB1 | 3.5301 | 0.3547 | 1.9424 | 0.000999 |
| MacIFN | melanomatumo | SPP1 | NCSTN | 3.5301 | 0.2614 | 1.8958 | 0.000999 |
| MacIFN | melanomatumo | SPP1 | CD44 | 3.5301 | 1.2653 | 2.3977 | 0.003 |
| MacIFN | melanomatumo | CALR | NCSTN | 0.8992 | 0.2614 | 0.5803 | 0.004 |
| MacIFN | melanomatumo | RTN4 | CD99 | 0.8998 | 0.6845 | 0.7921 | 0.005 |
| MacIFN | melanomatumo | A2M | TNFRSF14 | 0.3032 | 0.5069 | 0.4051 | 0.00999 |
| MacIFN | ovarytumor | AIMP1 | RACK1 | 0.282 | 2.9543 | 1.6181 | 0.000999 |
| MacIFN | ovarytumor | AREG | FOLR1 | 0.3335 | 0.5619 | 0.4477 | 0.000999 |
| MacIFN | ovarytumor | CALM1 | RPSA | 1.3759 | 2.9542 | 2.165 | 0.000999 |
| MacIFN | ovarytumor | CCL20 | CXCR4 | 0.3308 | 0.3688 | 0.3498 | 0.000999 |
| MacIFN | ovarytumor | CXCL10 | CXCR4 | 1.1558 | 0.3688 | 0.7623 | 0.000999 |
| MacIFN | ovarytumor | CXCL11 | CXCR4 | 0.3478 | 0.3688 | 0.3583 | 0.000999 |
| MacIFN | ovarytumor | CXCL16 | CXCR4 | 0.7131 | 0.3688 | 0.5409 | 0.000999 |
| MacIFN | ovarytumor | CXCL2 | CXCR4 | 0.492 | 0.3688 | 0.4304 | 0.000999 |
| MacIFN | ovarytumor | CXCL3 | CXCR4 | 0.3459 | 0.3688 | 0.3574 | 0.000999 |
| MacIFN | ovarytumor | CXCL9 | CXCR4 | 0.3045 | 0.3688 | 0.3366 | 0.000999 |
| MacIFN | ovarytumor | FN1 | ITGB1 | 1.1314 | 0.2783 | 0.7048 | 0.000999 |
| MacIFN | ovarytumor | FN1 | RPSA | 1.1314 | 2.9542 | 2.0428 | 0.000999 |
| MacIFN | ovarytumor | LGALS3 | ITGB1 | 1.5068 | 0.2783 | 0.8926 | 0.000999 |
| MacIFN | ovarytumor | PTPRC | CXCR4 | 0.5011 | 0.3688 | 0.4349 | 0.000999 |
| MacIFN | ovarytumor | SPP1 | ITGB1 | 3.5301 | 0.2783 | 1.9042 | 0.000999 |
| MacIFN | ovarytumor | TGFB1 | ITGB1 | 0.2572 | 0.2783 | 0.2677 | 0.000999 |

| | | | | | | | |
|---|---|---|---|---|---|---|---|
| MacIFN | ovarytumor | TNFSF13B | ITGB1 | 0.9576 | 0.2783 | 0.6179 | 0.000999 |
| breasttumor | MacIFN | ANXA1 | C3AR1 | 0.4092 | 0.292 | 0.3506 | 0.000999 |
| breasttumor | MacIFN | ANXA1 | C5AR1 | 0.4092 | 0.4513 | 0.4302 | 0.000999 |
| breasttumor | MacIFN | ANXA1 | CCR1 | 0.4092 | 0.3127 | 0.3609 | 0.000999 |
| breasttumor | MacIFN | ANXA1 | CD4 | 0.4092 | 0.3309 | 0.37 | 0.000999 |
| breasttumor | MacIFN | ANXA1 | CD44 | 0.4092 | 1.0927 | 0.7509 | 0.000999 |
| breasttumor | MacIFN | ANXA1 | CXCR4 | 0.4092 | 1.1921 | 0.8006 | 0.000999 |
| breasttumor | MacIFN | ANXA1 | FPR1 | 0.4092 | 0.3587 | 0.3839 | 0.000999 |
| breasttumor | MacIFN | ANXA1 | FPR3 | 0.4092 | 0.4459 | 0.4275 | 0.000999 |
| breasttumor | MacIFN | APP | C3AR1 | 0.5409 | 0.292 | 0.4165 | 0.000999 |
| breasttumor | MacIFN | APP | C5AR1 | 0.5409 | 0.4513 | 0.4961 | 0.000999 |
| breasttumor | MacIFN | APP | CCR1 | 0.5409 | 0.3127 | 0.4268 | 0.000999 |
| breasttumor | MacIFN | APP | CXCR4 | 0.5409 | 1.1921 | 0.8665 | 0.000999 |
| breasttumor | MacIFN | APP | FPR1 | 0.5409 | 0.3587 | 0.4498 | 0.000999 |
| breasttumor | MacIFN | ARF1 | CD4 | 1.6219 | 0.3309 | 0.9764 | 0.000999 |
| breasttumor | MacIFN | ARF1 | CD44 | 1.6219 | 1.0927 | 1.3573 | 0.000999 |
| breasttumor | MacIFN | CALM1 | TNFRSF1B | 0.9015 | 0.4401 | 0.6708 | 0.000999 |
| breasttumor | MacIFN | CALR | CD44 | 1.1835 | 1.0927 | 1.1381 | 0.000999 |
| breasttumor | MacIFN | COL2A1 | CD44 | 0.3935 | 1.0927 | 0.7431 | 0.000999 |
| breasttumor | MacIFN | GSTP1 | CD44 | 2.5435 | 1.0927 | 1.8181 | 0.000999 |
| breasttumor | MacIFN | HEBP1 | C3AR1 | 0.4062 | 0.292 | 0.3491 | 0.000999 |
| breasttumor | MacIFN | HEBP1 | C5AR1 | 0.4062 | 0.4513 | 0.4287 | 0.000999 |
| breasttumor | MacIFN | HEBP1 | CCR1 | 0.4062 | 0.3127 | 0.3594 | 0.000999 |
| breasttumor | MacIFN | HEBP1 | CXCR4 | 0.4062 | 1.1921 | 0.7991 | 0.000999 |
| breasttumor | MacIFN | HEBP1 | FPR1 | 0.4062 | 0.3587 | 0.3824 | 0.000999 |
| breasttumor | MacIFN | HEBP1 | FPR3 | 0.4062 | 0.4459 | 0.426 | 0.000999 |
| breasttumor | MacIFN | HMGB1 | LY96 | 1.1815 | 0.8107 | 0.9961 | 0.000999 |
| breasttumor | MacIFN | HMGB1 | TLR2 | 1.1815 | 0.2907 | 0.7361 | 0.000999 |
| breasttumor | MacIFN | HMGB2 | CD44 | 0.3139 | 1.0927 | 0.7033 | 0.000999 |
| breasttumor | MacIFN | HMGB2 | DAB2 | 0.3139 | 0.4721 | 0.393 | 0.000999 |

| | | | | | | | |
|---|---|---|---|---|---|---|---|
| breasttumor | MacIFN | HSP90B1 | TNFRSF1B | 0.7786 | 0.4401 | 0.6093 | 0.000999 |
| breasttumor | MacIFN | ITGB1 | CD47 | 0.2939 | 0.4549 | 0.3744 | 0.000999 |
| breasttumor | MacIFN | ITGB1 | DAB2 | 0.2939 | 0.4721 | 0.383 | 0.000999 |
| breasttumor | MacIFN | ITGB1 | ITGAX | 0.2939 | 0.2612 | 0.2776 | 0.000999 |
| breasttumor | MacIFN | ITGB1 | PLAUR | 0.2939 | 1.2582 | 0.776 | 0.000999 |
| breasttumor | MacIFN | LGALS3 | CLEC7A | 0.8391 | 0.5077 | 0.6734 | 0.000999 |
| breasttumor | MacIFN | LGALS3 | FCGR2A | 0.8391 | 0.7311 | 0.7851 | 0.000999 |
| breasttumor | MacIFN | LGALS3 | FCGR2B | 0.8391 | 0.5626 | 0.7009 | 0.000999 |
| breasttumor | MacIFN | LGALS3 | TFRC | 0.8391 | 0.5211 | 0.6801 | 0.000999 |
| breasttumor | MacIFN | MIF | CD74 | 3.2716 | 3.8419 | 3.5568 | 0.000999 |
| breasttumor | MacIFN | NPW | C3AR1 | 0.5478 | 0.292 | 0.4199 | 0.000999 |
| breasttumor | MacIFN | NPW | C5AR1 | 0.5478 | 0.4513 | 0.4996 | 0.000999 |
| breasttumor | MacIFN | NPW | CCR1 | 0.5478 | 0.3127 | 0.4303 | 0.000999 |
| breasttumor | MacIFN | NPW | CXCR4 | 0.5478 | 1.1921 | 0.87 | 0.000999 |
| breasttumor | MacIFN | NPW | FPR1 | 0.5478 | 0.3587 | 0.4532 | 0.000999 |
| breasttumor | MacIFN | RPS19 | C5AR1 | 3.691 | 0.4513 | 2.0712 | 0.000999 |
| breasttumor | MacIFN | RPS19 | CD4 | 3.691 | 0.3309 | 2.0109 | 0.000999 |
| breasttumor | MacIFN | RPS19 | ICAM1 | 3.691 | 0.3381 | 2.0145 | 0.000999 |
| breasttumor | MacIFN | RTN4 | CD44 | 0.3318 | 1.0927 | 0.7122 | 0.000999 |
| breasttumor | MacIFN | TF | CD4 | 0.2688 | 0.3309 | 0.2998 | 0.000999 |
| breasttumor | MacIFN | TF | TFRC | 0.2688 | 0.5211 | 0.3949 | 0.000999 |
| colorectaltumor | MacIFN | ANXA1 | CXCR4 | 0.5916 | 1.1921 | 0.8918 | 0.000999 |
| colorectaltumor | MacIFN | APP | C3AR1 | 0.4966 | 0.292 | 0.3943 | 0.000999 |
| colorectaltumor | MacIFN | APP | C5AR1 | 0.4966 | 0.4513 | 0.474 | 0.000999 |
| colorectaltumor | MacIFN | APP | CCR1 | 0.4966 | 0.3127 | 0.4047 | 0.000999 |
| colorectaltumor | MacIFN | APP | CXCR4 | 0.4966 | 1.1921 | 0.8444 | 0.000999 |
| colorectaltumor | MacIFN | APP | FPR1 | 0.4966 | 0.3587 | 0.4276 | 0.000999 |
| colorectaltumor | MacIFN | ARF1 | CD4 | 0.6489 | 0.3309 | 0.4899 | 0.000999 |
| colorectaltumor | MacIFN | ARF1 | CD44 | 0.6489 | 1.0927 | 0.8708 | 0.000999 |
| colorectaltumor | MacIFN | COL1A1 | CD44 | 0.4875 | 1.0927 | 0.7901 | 0.000999 |

| | | | | | | | |
|---|---|---|---|---|---|---|---|
| colorectaltumor | MacIFN | GSTP1 | CD44 | 2.8039 | 1.0927 | 1.9483 | 0.000999 |
| colorectaltumor | MacIFN | HMGB1 | LY96 | 2.155 | 0.8107 | 1.4828 | 0.000999 |
| colorectaltumor | MacIFN | HMGB1 | TLR2 | 2.155 | 0.2907 | 1.2228 | 0.000999 |
| colorectaltumor | MacIFN | HMGB2 | CD44 | 0.5347 | 1.0927 | 0.8137 | 0.000999 |
| colorectaltumor | MacIFN | ITGB1 | CD47 | 0.4214 | 0.4549 | 0.4382 | 0.000999 |
| colorectaltumor | MacIFN | ITGB1 | DAB2 | 0.4214 | 0.4721 | 0.4467 | 0.000999 |
| colorectaltumor | MacIFN | ITGB1 | ITGAX | 0.4214 | 0.2612 | 0.3413 | 0.000999 |
| colorectaltumor | MacIFN | ITGB1 | PLAUR | 0.4214 | 1.2582 | 0.8398 | 0.000999 |
| colorectaltumor | MacIFN | LGALS3 | CLEC7A | 1.5563 | 0.5077 | 1.032 | 0.000999 |
| colorectaltumor | MacIFN | LGALS3 | FCGR2A | 1.5563 | 0.7311 | 1.1437 | 0.000999 |
| colorectaltumor | MacIFN | LGALS3 | FCGR2B | 1.5563 | 0.5626 | 1.0594 | 0.000999 |
| colorectaltumor | MacIFN | MIF | CD74 | 1.6741 | 3.8419 | 2.758 | 0.000999 |
| colorectaltumor | MacIFN | RPS19 | C5AR1 | 4.0956 | 0.4513 | 2.2734 | 0.000999 |
| colorectaltumor | MacIFN | RPS19 | CD4 | 4.0956 | 0.3309 | 2.2132 | 0.000999 |
| colorectaltumor | MacIFN | RPS19 | ICAM1 | 4.0956 | 0.3381 | 2.2168 | 0.000999 |
| colorectaltumor | MacIFN | HMGB2 | DAB2 | 0.5347 | 0.4721 | 0.5034 | 0.002 |
| colorectaltumor | MacIFN | LGALS3 | TFRC | 1.5563 | 0.5211 | 1.0387 | 0.004 |
| colorectaltumor | MacIFN | HSP90B1 | TNFRSF1B | 0.5887 | 0.4401 | 0.5144 | 0.005 |
| colorectaltumor | MacIFN | AIMP1 | RRBP1 | 0.5265 | 0.2518 | 0.3891 | 0.00699 |
| livertumor | MacIFN | AGT | ATP6AP2 | 0.4701 | 0.4586 | 0.4643 | 0.000999 |
| livertumor | MacIFN | AGT | C3AR1 | 0.4701 | 0.292 | 0.381 | 0.000999 |
| livertumor | MacIFN | AGT | C5AR1 | 0.4701 | 0.4513 | 0.4607 | 0.000999 |
| livertumor | MacIFN | AGT | CCR1 | 0.4701 | 0.3127 | 0.3914 | 0.000999 |
| livertumor | MacIFN | AGT | CXCR4 | 0.4701 | 1.1921 | 0.8311 | 0.000999 |
| livertumor | MacIFN | AGT | FPR1 | 0.4701 | 0.3587 | 0.4144 | 0.000999 |
| livertumor | MacIFN | ANXA1 | CXCR4 | 0.3041 | 1.1921 | 0.7481 | 0.000999 |
| livertumor | MacIFN | APOA1 | ABCA1 | 0.2888 | 0.2736 | 0.2812 | 0.000999 |
| livertumor | MacIFN | APOA1 | MSR1 | 0.2888 | 0.6639 | 0.4763 | 0.000999 |
| livertumor | MacIFN | APP | C3AR1 | 0.5626 | 0.292 | 0.4273 | 0.000999 |
| livertumor | MacIFN | APP | C5AR1 | 0.5626 | 0.4513 | 0.5069 | 0.000999 |

| | | | | | | | |
|---|---|---|---|---|---|---|---|
| livertumor | MacIFN | APP | CCR1 | 0.5626 | 0.3127 | 0.4376 | 0.000999 |
| livertumor | MacIFN | APP | CXCR4 | 0.5626 | 1.1921 | 0.8773 | 0.000999 |
| livertumor | MacIFN | APP | FPR1 | 0.5626 | 0.3587 | 0.4606 | 0.000999 |
| livertumor | MacIFN | AREG | ICAM1 | 0.448 | 0.3381 | 0.393 | 0.000999 |
| livertumor | MacIFN | ARF1 | CD4 | 0.6936 | 0.3309 | 0.5122 | 0.000999 |
| livertumor | MacIFN | ARF1 | CD44 | 0.6936 | 1.0927 | 0.8931 | 0.000999 |
| livertumor | MacIFN | C3 | C3AR1 | 0.6584 | 0.292 | 0.4752 | 0.000999 |
| livertumor | MacIFN | C3 | C5AR1 | 0.6584 | 0.4513 | 0.5549 | 0.000999 |
| livertumor | MacIFN | C3 | CCR1 | 0.6584 | 0.3127 | 0.4856 | 0.000999 |
| livertumor | MacIFN | C3 | CXCR4 | 0.6584 | 1.1921 | 0.9253 | 0.000999 |
| livertumor | MacIFN | C3 | FPR1 | 0.6584 | 0.3587 | 0.5085 | 0.000999 |
| livertumor | MacIFN | C3 | ITGAX | 0.6584 | 0.2612 | 0.4598 | 0.000999 |
| livertumor | MacIFN | C3 | ITGB2 | 0.6584 | 0.9251 | 0.7918 | 0.000999 |
| livertumor | MacIFN | CALR | CD44 | 1.0416 | 1.0927 | 1.0671 | 0.000999 |
| livertumor | MacIFN | CCL20 | C3AR1 | 1.0877 | 0.292 | 0.6898 | 0.000999 |
| livertumor | MacIFN | CCL20 | C5AR1 | 1.0877 | 0.4513 | 0.7695 | 0.000999 |
| livertumor | MacIFN | CCL20 | CCR1 | 1.0877 | 0.3127 | 0.7002 | 0.000999 |
| livertumor | MacIFN | CCL20 | CXCR4 | 1.0877 | 1.1921 | 1.1399 | 0.000999 |
| livertumor | MacIFN | CCL20 | FPR1 | 1.0877 | 0.3587 | 0.7232 | 0.000999 |
| livertumor | MacIFN | CXCL2 | C5AR1 | 0.3364 | 0.4513 | 0.3938 | 0.000999 |
| livertumor | MacIFN | CXCL2 | CXCR4 | 0.3364 | 1.1921 | 0.7642 | 0.000999 |
| livertumor | MacIFN | CXCL3 | C5AR1 | 0.2543 | 0.4513 | 0.3528 | 0.000999 |
| livertumor | MacIFN | CXCL3 | CXCR4 | 0.2543 | 1.1921 | 0.7232 | 0.000999 |
| livertumor | MacIFN | CXCL3 | FPR1 | 0.2543 | 0.3587 | 0.3065 | 0.000999 |
| livertumor | MacIFN | DSG2 | CD44 | 0.7479 | 1.0927 | 0.9203 | 0.000999 |
| livertumor | MacIFN | DSG2 | CXCR4 | 0.7479 | 1.1921 | 0.97 | 0.000999 |
| livertumor | MacIFN | DSG2 | DAB2 | 0.7479 | 0.4721 | 0.61 | 0.000999 |
| livertumor | MacIFN | FGA | ITGAX | 0.2688 | 0.2612 | 0.265 | 0.000999 |
| livertumor | MacIFN | FGA | ITGB2 | 0.2688 | 0.9251 | 0.5969 | 0.000999 |
| livertumor | MacIFN | GSTP1 | CD44 | 1.7219 | 1.0927 | 1.4073 | 0.000999 |

| | | | | | | | |
|---|---|---|---|---|---|---|---|
| livertumor | MacIFN | HMGB1 | LY96 | 1.4361 | 0.8107 | 1.1234 | 0.000999 |
| livertumor | MacIFN | HMGB1 | TLR2 | 1.4361 | 0.2907 | 0.8634 | 0.000999 |
| livertumor | MacIFN | HSP90B1 | TNFRSF1B | 0.8951 | 0.4401 | 0.6676 | 0.000999 |
| livertumor | MacIFN | ITGB1 | CD47 | 1.6044 | 0.4549 | 1.0297 | 0.000999 |
| livertumor | MacIFN | ITGB1 | DAB2 | 1.6044 | 0.4721 | 1.0383 | 0.000999 |
| livertumor | MacIFN | ITGB1 | ITGAX | 1.6044 | 0.2612 | 0.9328 | 0.000999 |
| livertumor | MacIFN | ITGB1 | PLAUR | 1.6044 | 1.2582 | 1.4313 | 0.000999 |
| livertumor | MacIFN | ITGB1 | RACK1 | 1.6044 | 1.885 | 1.7447 | 0.000999 |
| livertumor | MacIFN | MIF | CD74 | 1.7638 | 3.8419 | 2.8028 | 0.000999 |
| livertumor | MacIFN | RPS19 | C5AR1 | 4.0772 | 0.4513 | 2.2643 | 0.000999 |
| livertumor | MacIFN | RPS19 | CD4 | 4.0772 | 0.3309 | 2.204 | 0.000999 |
| livertumor | MacIFN | RPS19 | ICAM1 | 4.0772 | 0.3381 | 2.2076 | 0.000999 |
| livertumor | MacIFN | RTN4 | CD44 | 1.0046 | 1.0927 | 1.0486 | 0.000999 |
| livertumor | MacIFN | TF | CD4 | 0.523 | 0.3309 | 0.4269 | 0.000999 |
| livertumor | MacIFN | TF | TFRC | 0.523 | 0.5211 | 0.5221 | 0.000999 |
| livertumor | MacIFN | VTN | PLAUR | 0.7399 | 1.2582 | 0.9991 | 0.000999 |
| livertumor | MacIFN | CXCL3 | CCR1 | 0.2543 | 0.3127 | 0.2835 | 0.002 |
| livertumor | MacIFN | CXCL3 | C3AR1 | 0.2543 | 0.292 | 0.2731 | 0.004 |
| livertumor | MacIFN | CD99 | CD99 | 0.5906 | 0.4533 | 0.5219 | 0.00799 |
| lungtumor | MacIFN | ANXA1 | C3AR1 | 1.4026 | 0.292 | 0.8473 | 0.000999 |
| lungtumor | MacIFN | ANXA1 | C5AR1 | 1.4026 | 0.4513 | 0.927 | 0.000999 |
| lungtumor | MacIFN | ANXA1 | CCR1 | 1.4026 | 0.3127 | 0.8577 | 0.000999 |
| lungtumor | MacIFN | ANXA1 | CD4 | 1.4026 | 0.3309 | 0.8667 | 0.000999 |
| lungtumor | MacIFN | ANXA1 | CD44 | 1.4026 | 1.0927 | 1.2476 | 0.000999 |
| lungtumor | MacIFN | ANXA1 | CXCR4 | 1.4026 | 1.1921 | 1.2974 | 0.000999 |
| lungtumor | MacIFN | ANXA1 | FPR1 | 1.4026 | 0.3587 | 0.8806 | 0.000999 |
| lungtumor | MacIFN | ANXA1 | FPR3 | 1.4026 | 0.4459 | 0.9243 | 0.000999 |
| lungtumor | MacIFN | AREG | ICAM1 | 0.3537 | 0.3381 | 0.3459 | 0.000999 |
| lungtumor | MacIFN | ARF1 | CD4 | 0.8791 | 0.3309 | 0.605 | 0.000999 |
| lungtumor | MacIFN | ARF1 | CD44 | 0.8791 | 1.0927 | 0.9859 | 0.000999 |

| | | | | | | | |
|---|---|---|---|---|---|---|---|
| lungtumor | MacIFN | CALM1 | TNFRSF1B | 0.8426 | 0.4401 | 0.6414 | 0.000999 |
| lungtumor | MacIFN | CALR | CD44 | 0.7795 | 1.0927 | 0.9361 | 0.000999 |
| lungtumor | MacIFN | CCL20 | C3AR1 | 0.2885 | 0.292 | 0.2903 | 0.000999 |
| lungtumor | MacIFN | CCL20 | C5AR1 | 0.2885 | 0.4513 | 0.3699 | 0.000999 |
| lungtumor | MacIFN | CCL20 | CCR1 | 0.2885 | 0.3127 | 0.3006 | 0.000999 |
| lungtumor | MacIFN | CCL20 | CXCR4 | 0.2885 | 1.1921 | 0.7403 | 0.000999 |
| lungtumor | MacIFN | CCL20 | FPR1 | 0.2885 | 0.3587 | 0.3236 | 0.000999 |
| lungtumor | MacIFN | CXCL10 | C3AR1 | 0.3417 | 0.292 | 0.3168 | 0.000999 |
| lungtumor | MacIFN | CXCL10 | C5AR1 | 0.3417 | 0.4513 | 0.3965 | 0.000999 |
| lungtumor | MacIFN | CXCL10 | CCR1 | 0.3417 | 0.3127 | 0.3272 | 0.000999 |
| lungtumor | MacIFN | CXCL10 | CXCR4 | 0.3417 | 1.1921 | 0.7669 | 0.000999 |
| lungtumor | MacIFN | CXCL10 | FPR1 | 0.3417 | 0.3587 | 0.3502 | 0.000999 |
| lungtumor | MacIFN | FGA | ITGAX | 0.4445 | 0.2612 | 0.3529 | 0.000999 |
| lungtumor | MacIFN | FGA | ITGB2 | 0.4445 | 0.9251 | 0.6848 | 0.000999 |
| lungtumor | MacIFN | GRN | TNFRSF1B | 0.4026 | 0.4401 | 0.4214 | 0.000999 |
| lungtumor | MacIFN | GSTP1 | CD44 | 2.4093 | 1.0927 | 1.751 | 0.000999 |
| lungtumor | MacIFN | HLA-DRA | PTPRC | 0.9609 | 0.5011 | 0.731 | 0.000999 |
| lungtumor | MacIFN | HMGB1 | LY96 | 1.1943 | 0.8107 | 1.0025 | 0.000999 |
| lungtumor | MacIFN | HMGB1 | TLR2 | 1.1943 | 0.2907 | 0.7425 | 0.000999 |
| lungtumor | MacIFN | HMGB2 | CD44 | 0.5593 | 1.0927 | 0.826 | 0.000999 |
| lungtumor | MacIFN | HMGB2 | DAB2 | 0.5593 | 0.4721 | 0.5157 | 0.000999 |
| lungtumor | MacIFN | HSP90B1 | TNFRSF1B | 0.6141 | 0.4401 | 0.5271 | 0.000999 |
| lungtumor | MacIFN | LGALS3 | CLEC7A | 0.9915 | 0.5077 | 0.7496 | 0.000999 |
| lungtumor | MacIFN | LGALS3 | FCGR2A | 0.9915 | 0.7311 | 0.8613 | 0.000999 |
| lungtumor | MacIFN | LGALS3 | FCGR2B | 0.9915 | 0.5626 | 0.7771 | 0.000999 |
| lungtumor | MacIFN | LGALS3 | TFRC | 0.9915 | 0.5211 | 0.7563 | 0.000999 |
| lungtumor | MacIFN | MIF | CD74 | 1.4585 | 3.8419 | 2.6502 | 0.000999 |
| lungtumor | MacIFN | RPS19 | C5AR1 | 3.5104 | 0.4513 | 1.9809 | 0.000999 |
| lungtumor | MacIFN | RPS19 | CD4 | 3.5104 | 0.3309 | 1.9206 | 0.000999 |
| lungtumor | MacIFN | RPS19 | ICAM1 | 3.5104 | 0.3381 | 1.9242 | 0.000999 |

| | | | | | | | |
|---|---|---|---|---|---|---|---|
| lungtumor | MacIFN | RTN4 | CD44 | 0.4013 | 1.0927 | 0.747 | 0.000999 |
| lungtumor | MacIFN | SAA1 | C3AR1 | 0.2539 | 0.292 | 0.2729 | 0.000999 |
| lungtumor | MacIFN | SAA1 | C5AR1 | 0.2539 | 0.4513 | 0.3526 | 0.000999 |
| lungtumor | MacIFN | SAA1 | CCR1 | 0.2539 | 0.3127 | 0.2833 | 0.000999 |
| lungtumor | MacIFN | SAA1 | CXCR4 | 0.2539 | 1.1921 | 0.723 | 0.000999 |
| lungtumor | MacIFN | SAA1 | FPR1 | 0.2539 | 0.3587 | 0.3063 | 0.000999 |
| lungtumor | MacIFN | SPP1 | CD44 | 1.4874 | 1.0927 | 1.29 | 0.000999 |
| lungtumor | MacIFN | AIMP1 | RRBP1 | 0.3345 | 0.2518 | 0.2932 | 0.003 |
| melanomatumo | MacIFN | ARF1 | CD4 | 1.8545 | 0.3309 | 1.0927 | 0.000999 |
| melanomatumo | MacIFN | ARF1 | CD44 | 1.8545 | 1.0927 | 1.4736 | 0.000999 |
| melanomatumo | MacIFN | GSTP1 | CD44 | 2.3341 | 1.0927 | 1.7134 | 0.000999 |
| melanomatumo | MacIFN | HAS2 | CD44 | 0.3078 | 1.0927 | 0.7002 | 0.000999 |
| melanomatumo | MacIFN | HEBP1 | C3AR1 | 0.3689 | 0.292 | 0.3304 | 0.000999 |
| melanomatumo | MacIFN | HEBP1 | C5AR1 | 0.3689 | 0.4513 | 0.4101 | 0.000999 |
| melanomatumo | MacIFN | HEBP1 | CCR1 | 0.3689 | 0.3127 | 0.3408 | 0.000999 |
| melanomatumo | MacIFN | HEBP1 | CXCR4 | 0.3689 | 1.1921 | 0.7805 | 0.000999 |
| melanomatumo | MacIFN | HEBP1 | FPR1 | 0.3689 | 0.3587 | 0.3638 | 0.000999 |
| melanomatumo | MacIFN | HEBP1 | FPR3 | 0.3689 | 0.4459 | 0.4074 | 0.000999 |
| melanomatumo | MacIFN | HMGB1 | LY96 | 1.6757 | 0.8107 | 1.2432 | 0.000999 |
| melanomatumo | MacIFN | HMGB1 | TLR2 | 1.6757 | 0.2907 | 0.9832 | 0.000999 |
| melanomatumo | MacIFN | MIF | CD74 | 2.6939 | 3.8419 | 3.2679 | 0.000999 |
| ovarytumor | MacIFN | ANXA1 | C3AR1 | 1.0314 | 0.292 | 0.6617 | 0.000999 |
| ovarytumor | MacIFN | ANXA1 | C5AR1 | 1.0314 | 0.4513 | 0.7413 | 0.000999 |
| ovarytumor | MacIFN | ANXA1 | CCR1 | 1.0314 | 0.3127 | 0.672 | 0.000999 |
| ovarytumor | MacIFN | ANXA1 | CD4 | 1.0314 | 0.3309 | 0.6811 | 0.000999 |
| ovarytumor | MacIFN | ANXA1 | CD44 | 1.0314 | 1.0927 | 1.062 | 0.000999 |
| ovarytumor | MacIFN | ANXA1 | CXCR4 | 1.0314 | 1.1921 | 1.1117 | 0.000999 |
| ovarytumor | MacIFN | ANXA1 | FPR1 | 1.0314 | 0.3587 | 0.695 | 0.000999 |
| ovarytumor | MacIFN | ANXA1 | FPR3 | 1.0314 | 0.4459 | 0.7386 | 0.000999 |
| ovarytumor | MacIFN | APP | C3AR1 | 0.2907 | 0.292 | 0.2914 | 0.000999 |

| | | | | | | | |
|---|---|---|---|---|---|---|---|
| ovarytumor | MacIFN | APP | C5AR1 | 0.2907 | 0.4513 | 0.371 | 0.000999 |
| ovarytumor | MacIFN | APP | CCR1 | 0.2907 | 0.3127 | 0.3017 | 0.000999 |
| ovarytumor | MacIFN | APP | CXCR4 | 0.2907 | 1.1921 | 0.7414 | 0.000999 |
| ovarytumor | MacIFN | APP | FPR1 | 0.2907 | 0.3587 | 0.3247 | 0.000999 |
| ovarytumor | MacIFN | ARF1 | CD4 | 0.7364 | 0.3309 | 0.5336 | 0.000999 |
| ovarytumor | MacIFN | ARF1 | CD44 | 0.7364 | 1.0927 | 0.9145 | 0.000999 |
| ovarytumor | MacIFN | CALM1 | TNFRSF1B | 1.0087 | 0.4401 | 0.7244 | 0.000999 |
| ovarytumor | MacIFN | CALR | CD44 | 0.7423 | 1.0927 | 0.9175 | 0.000999 |
| ovarytumor | MacIFN | CCL2 | CCR1 | 0.3526 | 0.3127 | 0.3327 | 0.000999 |
| ovarytumor | MacIFN | COL1A1 | CD44 | 0.3504 | 1.0927 | 0.7215 | 0.000999 |
| ovarytumor | MacIFN | GRN | TNFRSF1B | 0.3896 | 0.4401 | 0.4148 | 0.000999 |
| ovarytumor | MacIFN | GSTP1 | CD44 | 2.6185 | 1.0927 | 1.8556 | 0.000999 |
| ovarytumor | MacIFN | HLA-DMA | CD4 | 0.2537 | 0.3309 | 0.2923 | 0.000999 |
| ovarytumor | MacIFN | HLA-DRA | CD4 | 1.2369 | 0.3309 | 0.7839 | 0.000999 |
| ovarytumor | MacIFN | HLA-DRA | PTPRC | 1.2369 | 0.5011 | 0.869 | 0.000999 |
| ovarytumor | MacIFN | HMGB1 | LY96 | 1.7272 | 0.8107 | 1.2689 | 0.000999 |
| ovarytumor | MacIFN | HMGB1 | TLR2 | 1.7272 | 0.2907 | 1.009 | 0.000999 |
| ovarytumor | MacIFN | HMGB2 | CD44 | 0.4974 | 1.0927 | 0.795 | 0.000999 |
| ovarytumor | MacIFN | HMGB2 | DAB2 | 0.4974 | 0.4721 | 0.4847 | 0.000999 |
| ovarytumor | MacIFN | HSP90B1 | TNFRSF1B | 0.6365 | 0.4401 | 0.5383 | 0.000999 |
| ovarytumor | MacIFN | ITGB1 | DAB2 | 0.2783 | 0.4721 | 0.3752 | 0.000999 |
| ovarytumor | MacIFN | ITGB1 | ITGAX | 0.2783 | 0.2612 | 0.2698 | 0.000999 |
| ovarytumor | MacIFN | ITGB1 | PLAUR | 0.2783 | 1.2582 | 0.7682 | 0.000999 |
| ovarytumor | MacIFN | LGALS3 | CLEC7A | 0.5097 | 0.5077 | 0.5087 | 0.000999 |
| ovarytumor | MacIFN | LGALS3 | FCGR2A | 0.5097 | 0.7311 | 0.6204 | 0.000999 |
| ovarytumor | MacIFN | LGALS3 | FCGR2B | 0.5097 | 0.5626 | 0.5362 | 0.000999 |
| ovarytumor | MacIFN | LGALS3 | TFRC | 0.5097 | 0.5211 | 0.5154 | 0.000999 |
| ovarytumor | MacIFN | MIF | CD74 | 0.8565 | 3.8419 | 2.3492 | 0.000999 |
| ovarytumor | MacIFN | RPS19 | C5AR1 | 3.9325 | 0.4513 | 2.1919 | 0.000999 |
| ovarytumor | MacIFN | RPS19 | CD4 | 3.9325 | 0.3309 | 2.1317 | 0.000999 |

| | | | | | | | |
|---|---|---|---|---|---|---|---|
| ovarytumor | MacIFN | RPS19 | ICAM1 | 3.9325 | 0.3381 | 2.1353 | 0.000999 |
| ovarytumor | MacIFN | RTN4 | CD44 | 0.4992 | 1.0927 | 0.7959 | 0.000999 |
| ovarytumor | MacIFN | RTN4 | CD99 | 0.4992 | 0.4533 | 0.4762 | 0.000999 |
| ovarytumor | MacIFN | SPP1 | CD44 | 0.3946 | 1.0927 | 0.7436 | 0.000999 |

| Supplementary Table 8: Communication results between macrophages and lymphocytes subpopulations, and malignant cells | | | | | | | |
|---|---|---|---|---|---|---|---|
| **MacLA vs. lymphocytes** | | | | | | | |
| celltype_01 | celltype_02 | ligand | receptor | ligand_mean_expression | receptor_mean_expression | ligand_receptor_mean_expression | pvalue |
| MacLA | NKT | A2M | TNFRSF14 | 0.5014 | 0.2848 | 0.3931 | 0.000999 |
| MacLA | NKT | ADAM9 | ITGB1 | 0.3224 | 0.488 | 0.4052 | 0.000999 |
| MacLA | NKT | ADM | PTGER2 | 0.2957 | 0.3627 | 0.3292 | 0.000999 |
| MacLA | NKT | ALCAM | CD6 | 0.3816 | 0.4881 | 0.4349 | 0.000999 |
| MacLA | NKT | APOE | SORL1 | 3.1824 | 0.4137 | 1.798 | 0.000999 |
| MacLA | NKT | APP | CXCR4 | 0.2852 | 2.0373 | 1.1612 | 0.000999 |
| MacLA | NKT | APP | F2R | 0.2852 | 0.2978 | 0.2915 | 0.000999 |
| MacLA | NKT | APP | SORL1 | 0.2852 | 0.4137 | 0.3494 | 0.000999 |
| MacLA | NKT | APP | TNFRSF14 | 0.2852 | 0.2848 | 0.285 | 0.000999 |
| MacLA | NKT | COPA | CD8A | 0.3451 | 1.2358 | 0.7905 | 0.000999 |
| MacLA | NKT | COPA | ITGA4 | 0.3451 | 0.2946 | 0.3198 | 0.000999 |
| MacLA | NKT | CXCL16 | CXCR4 | 0.6306 | 2.0373 | 1.3339 | 0.000999 |
| MacLA | NKT | CXCL16 | S1PR4 | 0.6306 | 0.4435 | 0.5371 | 0.000999 |
| MacLA | NKT | CXCL16 | S1PR5 | 0.6306 | 0.508 | 0.5693 | 0.000999 |
| MacLA | NKT | CXCL3 | CXCR4 | 0.2783 | 2.0373 | 1.1578 | 0.000999 |
| MacLA | NKT | CXCL3 | S1PR4 | 0.2783 | 0.4435 | 0.3609 | 0.000999 |
| MacLA | NKT | CXCL3 | S1PR5 | 0.2783 | 0.508 | 0.3932 | 0.000999 |
| MacLA | NKT | FN1 | CD44 | 1.0963 | 0.6724 | 0.8844 | 0.000999 |
| MacLA | NKT | FN1 | ITGA4 | 1.0963 | 0.2946 | 0.6954 | 0.000999 |
| MacLA | NKT | FN1 | ITGB1 | 1.0963 | 0.488 | 0.7921 | 0.000999 |
| MacLA | NKT | FN1 | ITGB7 | 1.0963 | 0.4721 | 0.7842 | 0.000999 |
| MacLA | NKT | FN1 | RPSA | 1.0963 | 2.3639 | 1.7301 | 0.000999 |
| MacLA | NKT | GRN | TNFRSF1B | 1.7735 | 0.4949 | 1.1342 | 0.000999 |
| MacLA | NKT | GSTP1 | CD44 | 1.2712 | 0.6724 | 0.9718 | 0.000999 |
| MacLA | NKT | GSTP1 | IL7R | 1.2712 | 0.5933 | 0.9323 | 0.000999 |
| MacLA | NKT | GSTP1 | ITGA4 | 1.2712 | 0.2946 | 0.7829 | 0.000999 |
| MacLA | NKT | HLA-DRA | PTPRC | 3.0446 | 1.8797 | 2.4621 | 0.000999 |

| | | | | | | | |
|---|---|---|---|---|---|---|---|
| MacLA | NKT | LGALS3 | CD6 | 2.3411 | 0.4881 | 1.4146 | 0.000999 |
| MacLA | NKT | LGALS3 | ITGB1 | 2.3411 | 0.488 | 1.4145 | 0.000999 |
| MacLA | NKT | LGALS3 | SPN | 2.3411 | 0.6398 | 1.4904 | 0.000999 |
| MacLA | NKT | MIF | CD74 | 1.2502 | 1.6792 | 1.4647 | 0.000999 |
| MacLA | NKT | MIF | ITGA4 | 1.2502 | 0.2946 | 0.7724 | 0.000999 |
| MacLA | NKT | MIF | TNFRSF14 | 1.2502 | 0.2848 | 0.7675 | 0.000999 |
| MacLA | NKT | MMP9 | CD44 | 0.9836 | 0.6724 | 0.828 | 0.000999 |
| MacLA | NKT | MMP9 | ITGB1 | 0.9836 | 0.488 | 0.7358 | 0.000999 |
| MacLA | NKT | NAMPT | ANXA6 | 0.2975 | 0.844 | 0.5707 | 0.000999 |
| MacLA | NKT | NMB | F2R | 0.3014 | 0.2978 | 0.2996 | 0.000999 |
| MacLA | NKT | NMB | GPR65 | 0.3014 | 0.4898 | 0.3956 | 0.000999 |
| MacLA | NKT | RTN4 | CD44 | 1.0466 | 0.6724 | 0.8595 | 0.000999 |
| MacLA | NKT | RTN4 | CD99 | 1.0466 | 1.4547 | 1.2506 | 0.000999 |
| MacLA | NKT | SPP1 | CD44 | 3.0735 | 0.6724 | 1.873 | 0.000999 |
| MacLA | NKT | SPP1 | ITGA4 | 3.0735 | 0.2946 | 1.684 | 0.000999 |
| MacLA | NKT | SPP1 | ITGB1 | 3.0735 | 0.488 | 1.7807 | 0.000999 |
| MacLA | NKT | AIMP1 | RACK1 | 0.2907 | 2.4327 | 1.3617 | 0.002 |
| MacLA | NKT | LRPAP1 | SORL1 | 0.5574 | 0.4137 | 0.4855 | 0.002 |
| MacLA | NKT | ANXA1 | CXCR4 | 1.5201 | 2.0373 | 1.7787 | 0.003 |
| MacLA | NKT | COL1A1 | ITGB1 | 0.2943 | 0.488 | 0.3911 | 0.003 |
| MacLA | NKT | COL1A1 | CD44 | 0.2943 | 0.6724 | 0.4833 | 0.004 |
| MacLA | NKT | ANXA1 | F2R | 1.5201 | 0.2978 | 0.909 | 0.005 |
| MacLA | NKT | ANXA1 | ITGA4 | 1.5201 | 0.2946 | 0.9073 | 0.005 |
| MacLA | NKT | ANXA1 | CD44 | 1.5201 | 0.6724 | 1.0963 | 0.00699 |
| MacLA | TCD4ex | A2M | TNFRSF14 | 0.5014 | 0.2806 | 0.391 | 0.000999 |
| MacLA | TCD4ex | ADAM9 | ITGB1 | 0.3224 | 0.4615 | 0.392 | 0.000999 |
| MacLA | TCD4ex | ADM | PTGER4 | 0.2957 | 0.4696 | 0.3826 | 0.000999 |
| MacLA | TCD4ex | ALCAM | CD6 | 0.3816 | 0.5338 | 0.4577 | 0.000999 |
| MacLA | TCD4ex | ANXA1 | CCR7 | 1.5201 | 0.4844 | 1.0023 | 0.000999 |
| MacLA | TCD4ex | ANXA1 | CD4 | 1.5201 | 0.5864 | 1.0533 | 0.000999 |

| | | | | | | | |
|---|---|---|---|---|---|---|---|
| MacLA | TCD4ex | ANXA1 | CD44 | 1.5201 | 1.0244 | 1.2723 | 0.000999 |
| MacLA | TCD4ex | ANXA1 | CXCR4 | 1.5201 | 2.4201 | 1.9701 | 0.000999 |
| MacLA | TCD4ex | APP | CCR7 | 0.2852 | 0.4844 | 0.3848 | 0.000999 |
| MacLA | TCD4ex | APP | CXCR4 | 0.2852 | 2.4201 | 1.3526 | 0.000999 |
| MacLA | TCD4ex | APP | TNFRSF14 | 0.2852 | 0.2806 | 0.2829 | 0.000999 |
| MacLA | TCD4ex | CD86 | CD28 | 0.2798 | 0.3256 | 0.3027 | 0.000999 |
| MacLA | TCD4ex | CD86 | CTLA4 | 0.2798 | 0.6294 | 0.4546 | 0.000999 |
| MacLA | TCD4ex | COL1A1 | CD44 | 0.2943 | 1.0244 | 0.6593 | 0.000999 |
| MacLA | TCD4ex | COL1A1 | ITGB1 | 0.2943 | 0.4615 | 0.3779 | 0.000999 |
| MacLA | TCD4ex | CXCL16 | CCR7 | 0.6306 | 0.4844 | 0.5575 | 0.000999 |
| MacLA | TCD4ex | CXCL16 | CXCR4 | 0.6306 | 2.4201 | 1.5253 | 0.000999 |
| MacLA | TCD4ex | CXCL3 | CCR7 | 0.2783 | 0.4844 | 0.3814 | 0.000999 |
| MacLA | TCD4ex | CXCL3 | CXCR4 | 0.2783 | 2.4201 | 1.3492 | 0.000999 |
| MacLA | TCD4ex | FN1 | CD44 | 1.0963 | 1.0244 | 1.0604 | 0.000999 |
| MacLA | TCD4ex | FN1 | ITGB1 | 1.0963 | 0.4615 | 0.7789 | 0.000999 |
| MacLA | TCD4ex | FN1 | RPSA | 1.0963 | 2.4895 | 1.7929 | 0.000999 |
| MacLA | TCD4ex | GRN | TNFRSF1B | 1.7735 | 0.383 | 1.0783 | 0.000999 |
| MacLA | TCD4ex | GSTP1 | CD44 | 1.2712 | 1.0244 | 1.1478 | 0.000999 |
| MacLA | TCD4ex | GSTP1 | IL7R | 1.2712 | 1.2353 | 1.2532 | 0.000999 |
| MacLA | TCD4ex | HLA-DMA | CD4 | 0.896 | 0.5864 | 0.7412 | 0.000999 |
| MacLA | TCD4ex | HLA-DRA | CD4 | 3.0446 | 0.5864 | 1.8155 | 0.000999 |
| MacLA | TCD4ex | HLA-DRA | PDCD1 | 3.0446 | 0.457 | 1.7508 | 0.000999 |
| MacLA | TCD4ex | HLA-DRA | PTPRC | 3.0446 | 1.7493 | 2.397 | 0.000999 |
| MacLA | TCD4ex | HSP90B1 | ITGB1 | 0.7134 | 0.4615 | 0.5874 | 0.000999 |
| MacLA | TCD4ex | HSP90B1 | TNFRSF1B | 0.7134 | 0.383 | 0.5482 | 0.000999 |
| MacLA | TCD4ex | ITGB2 | ITGAL | 1.2219 | 0.2603 | 0.7411 | 0.000999 |
| MacLA | TCD4ex | LGALS3 | CD6 | 2.3411 | 0.5338 | 1.4374 | 0.000999 |
| MacLA | TCD4ex | LGALS3 | ITGB1 | 2.3411 | 0.4615 | 1.4013 | 0.000999 |
| MacLA | TCD4ex | MIF | CD74 | 1.2502 | 1.6094 | 1.4298 | 0.000999 |
| MacLA | TCD4ex | MIF | TNFRSF14 | 1.2502 | 0.2806 | 0.7654 | 0.000999 |

| | | | | | | | |
|---|---|---|---|---|---|---|---|
| MacLA | TCD4ex | MMP9 | CD44 | 0.9836 | 1.0244 | 1.004 | 0.000999 |
| MacLA | TCD4ex | MMP9 | ITGB1 | 0.9836 | 0.4615 | 0.7225 | 0.000999 |
| MacLA | TCD4ex | RTN4 | CD44 | 1.0466 | 1.0244 | 1.0355 | 0.000999 |
| MacLA | TCD4ex | RTN4 | CD99 | 1.0466 | 0.9165 | 0.9815 | 0.000999 |
| MacLA | TCD4ex | SPP1 | CD44 | 3.0735 | 1.0244 | 2.0489 | 0.000999 |
| MacLA | TCD4ex | SPP1 | ITGB1 | 3.0735 | 0.4615 | 1.7675 | 0.000999 |
| MacLA | TCD4ex | TGFB1 | ITGB1 | 0.3223 | 0.4615 | 0.3919 | 0.000999 |
| MacLA | TCD4ex | CALR | IL7R | 0.9648 | 1.2353 | 1.1 | 0.004 |
| MacLA | TCD4reg | A2M | TNFRSF14 | 0.5014 | 0.3843 | 0.4429 | 0.000999 |
| MacLA | TCD4reg | ADAM9 | ITGB1 | 0.3224 | 0.4001 | 0.3612 | 0.000999 |
| MacLA | TCD4reg | ALCAM | CD6 | 0.3816 | 0.2651 | 0.3233 | 0.000999 |
| MacLA | TCD4reg | ANXA1 | CCR7 | 1.5201 | 0.3098 | 0.915 | 0.000999 |
| MacLA | TCD4reg | ANXA1 | CD4 | 1.5201 | 0.429 | 0.9745 | 0.000999 |
| MacLA | TCD4reg | ANXA1 | CD44 | 1.5201 | 1.0896 | 1.3049 | 0.000999 |
| MacLA | TCD4reg | ANXA1 | CXCR3 | 1.5201 | 0.3357 | 0.9279 | 0.000999 |
| MacLA | TCD4reg | ANXA1 | CXCR4 | 1.5201 | 2.2962 | 1.9081 | 0.000999 |
| MacLA | TCD4reg | ANXA1 | CXCR6 | 1.5201 | 0.3945 | 0.9573 | 0.000999 |
| MacLA | TCD4reg | ANXA1 | FAS | 1.5201 | 0.2727 | 0.8964 | 0.000999 |
| MacLA | TCD4reg | APP | CCR7 | 0.2852 | 0.3098 | 0.2975 | 0.000999 |
| MacLA | TCD4reg | APP | CXCR3 | 0.2852 | 0.3357 | 0.3104 | 0.000999 |
| MacLA | TCD4reg | APP | CXCR4 | 0.2852 | 2.2962 | 1.2907 | 0.000999 |
| MacLA | TCD4reg | APP | CXCR6 | 0.2852 | 0.3945 | 0.3398 | 0.000999 |
| MacLA | TCD4reg | APP | TNFRSF14 | 0.2852 | 0.3843 | 0.3347 | 0.000999 |
| MacLA | TCD4reg | CD86 | CD28 | 0.2798 | 0.313 | 0.2964 | 0.000999 |
| MacLA | TCD4reg | CD86 | CTLA4 | 0.2798 | 0.7909 | 0.5354 | 0.000999 |
| MacLA | TCD4reg | CXCL16 | CCR7 | 0.6306 | 0.3098 | 0.4702 | 0.000999 |
| MacLA | TCD4reg | CXCL16 | CXCR3 | 0.6306 | 0.3357 | 0.4831 | 0.000999 |
| MacLA | TCD4reg | CXCL16 | CXCR4 | 0.6306 | 2.2962 | 1.4634 | 0.000999 |
| MacLA | TCD4reg | CXCL16 | CXCR6 | 0.6306 | 0.3945 | 0.5125 | 0.000999 |
| MacLA | TCD4reg | CXCL16 | S1PR4 | 0.6306 | 0.2744 | 0.4525 | 0.000999 |

| | | | | | | | |
|---|---|---|---|---|---|---|---|
| MacLA | TCD4reg | CXCL3 | CCR7 | 0.2783 | 0.3098 | 0.2941 | 0.000999 |
| MacLA | TCD4reg | CXCL3 | CXCR3 | 0.2783 | 0.3357 | 0.307 | 0.000999 |
| MacLA | TCD4reg | CXCL3 | CXCR4 | 0.2783 | 2.2962 | 1.2872 | 0.000999 |
| MacLA | TCD4reg | CXCL3 | CXCR6 | 0.2783 | 0.3945 | 0.3364 | 0.000999 |
| MacLA | TCD4reg | CXCL3 | S1PR4 | 0.2783 | 0.2744 | 0.2764 | 0.000999 |
| MacLA | TCD4reg | FN1 | CD44 | 1.0963 | 1.0896 | 1.093 | 0.000999 |
| MacLA | TCD4reg | FN1 | ITGB1 | 1.0963 | 0.4001 | 0.7482 | 0.000999 |
| MacLA | TCD4reg | FN1 | RPSA | 1.0963 | 2.4999 | 1.7981 | 0.000999 |
| MacLA | TCD4reg | GRN | TNFRSF1B | 1.7735 | 0.7982 | 1.2859 | 0.000999 |
| MacLA | TCD4reg | GSTP1 | CD44 | 1.2712 | 1.0896 | 1.1804 | 0.000999 |
| MacLA | TCD4reg | GSTP1 | FAS | 1.2712 | 0.2727 | 0.772 | 0.000999 |
| MacLA | TCD4reg | GSTP1 | IL7R | 1.2712 | 0.9682 | 1.1197 | 0.000999 |
| MacLA | TCD4reg | HLA-DMA | CD4 | 0.896 | 0.429 | 0.6625 | 0.000999 |
| MacLA | TCD4reg | HLA-DRA | CD4 | 3.0446 | 0.429 | 1.7368 | 0.000999 |
| MacLA | TCD4reg | HLA-DRA | PTPRC | 3.0446 | 1.6603 | 2.3525 | 0.000999 |
| MacLA | TCD4reg | HSP90B1 | ITGB1 | 0.7134 | 0.4001 | 0.5567 | 0.000999 |
| MacLA | TCD4reg | HSP90B1 | TNFRSF1B | 0.7134 | 0.7982 | 0.7558 | 0.000999 |
| MacLA | TCD4reg | LGALS3 | CD6 | 2.3411 | 0.2651 | 1.3031 | 0.000999 |
| MacLA | TCD4reg | LGALS3 | FAS | 2.3411 | 0.2727 | 1.3069 | 0.000999 |
| MacLA | TCD4reg | LGALS3 | ITGB1 | 2.3411 | 0.4001 | 1.3706 | 0.000999 |
| MacLA | TCD4reg | MIF | CD74 | 1.2502 | 2.0716 | 1.6609 | 0.000999 |
| MacLA | TCD4reg | MIF | TNFRSF14 | 1.2502 | 0.3843 | 0.8173 | 0.000999 |
| MacLA | TCD4reg | MMP9 | CD44 | 0.9836 | 1.0896 | 1.0366 | 0.000999 |
| MacLA | TCD4reg | MMP9 | ITGB1 | 0.9836 | 0.4001 | 0.6918 | 0.000999 |
| MacLA | TCD4reg | NMB | P2RY10 | 0.3014 | 0.2824 | 0.2919 | 0.000999 |
| MacLA | TCD4reg | RTN4 | CD44 | 1.0466 | 1.0896 | 1.0681 | 0.000999 |
| MacLA | TCD4reg | RTN4 | CD99 | 1.0466 | 0.6603 | 0.8534 | 0.000999 |
| MacLA | TCD4reg | SPP1 | CD44 | 3.0735 | 1.0896 | 2.0816 | 0.000999 |
| MacLA | TCD4reg | SPP1 | ITGB1 | 3.0735 | 0.4001 | 1.7368 | 0.000999 |
| MacLA | TCD4reg | CALR | IL7R | 0.9648 | 0.9682 | 0.9665 | 0.002 |

| | | | | | | | |
|---|---|---|---|---|---|---|---|
| MacLA | TCD4reg | TGFB1 | ITGB1 | 0.3223 | 0.4001 | 0.3612 | 0.002 |
| MacLA | TCD4reg | CALR | FAS | 0.9648 | 0.2727 | 0.6188 | 0.003 |
| MacLA | TCD4reg | CALR | CD44 | 0.9648 | 1.0896 | 1.0272 | 0.004 |
| MacLA | TCD8em | A2M | TNFRSF14 | 0.5014 | 0.2915 | 0.3964 | 0.000999 |
| MacLA | TCD8em | ADAM9 | ITGB1 | 0.3224 | 0.2557 | 0.2891 | 0.000999 |
| MacLA | TCD8em | ADM | PTGER4 | 0.2957 | 0.2847 | 0.2902 | 0.000999 |
| MacLA | TCD8em | ALCAM | CD6 | 0.3816 | 0.4204 | 0.401 | 0.000999 |
| MacLA | TCD8em | ANXA1 | CD44 | 1.5201 | 1.119 | 1.3196 | 0.000999 |
| MacLA | TCD8em | ANXA1 | CXCR3 | 1.5201 | 0.4189 | 0.9695 | 0.000999 |
| MacLA | TCD8em | ANXA1 | CXCR4 | 1.5201 | 2.8389 | 2.1795 | 0.000999 |
| MacLA | TCD8em | ANXA1 | CXCR6 | 1.5201 | 0.3399 | 0.93 | 0.000999 |
| MacLA | TCD8em | ANXA1 | F2R | 1.5201 | 0.2695 | 0.8948 | 0.000999 |
| MacLA | TCD8em | ANXA1 | ITGA4 | 1.5201 | 0.2823 | 0.9012 | 0.000999 |
| MacLA | TCD8em | APP | CXCR3 | 0.2852 | 0.4189 | 0.352 | 0.000999 |
| MacLA | TCD8em | APP | CXCR4 | 0.2852 | 2.8389 | 1.562 | 0.000999 |
| MacLA | TCD8em | APP | CXCR6 | 0.2852 | 0.3399 | 0.3125 | 0.000999 |
| MacLA | TCD8em | APP | F2R | 0.2852 | 0.2695 | 0.2773 | 0.000999 |
| MacLA | TCD8em | APP | TNFRSF14 | 0.2852 | 0.2915 | 0.2883 | 0.000999 |
| MacLA | TCD8em | COPA | CD8A | 0.3451 | 1.2321 | 0.7886 | 0.000999 |
| MacLA | TCD8em | COPA | ITGA4 | 0.3451 | 0.2823 | 0.3137 | 0.000999 |
| MacLA | TCD8em | CXCL16 | CXCR3 | 0.6306 | 0.4189 | 0.5247 | 0.000999 |
| MacLA | TCD8em | CXCL16 | CXCR4 | 0.6306 | 2.8389 | 1.7347 | 0.000999 |
| MacLA | TCD8em | CXCL16 | CXCR6 | 0.6306 | 0.3399 | 0.4852 | 0.000999 |
| MacLA | TCD8em | CXCL16 | S1PR4 | 0.6306 | 0.2677 | 0.4491 | 0.000999 |
| MacLA | TCD8em | CXCL3 | CXCR3 | 0.2783 | 0.4189 | 0.3486 | 0.000999 |
| MacLA | TCD8em | CXCL3 | CXCR4 | 0.2783 | 2.8389 | 1.5586 | 0.000999 |
| MacLA | TCD8em | CXCL3 | CXCR6 | 0.2783 | 0.3399 | 0.3091 | 0.000999 |
| MacLA | TCD8em | CXCL3 | S1PR4 | 0.2783 | 0.2677 | 0.273 | 0.000999 |
| MacLA | TCD8em | FN1 | CD44 | 1.0963 | 1.119 | 1.1077 | 0.000999 |
| MacLA | TCD8em | FN1 | ITGA4 | 1.0963 | 0.2823 | 0.6893 | 0.000999 |

| | | | | | | | |
|---|---|---|---|---|---|---|---|
| MacLA | TCD8em | FN1 | ITGB1 | 1.0963 | 0.2557 | 0.676 | 0.000999 |
| MacLA | TCD8em | FN1 | RPSA | 1.0963 | 2.3686 | 1.7325 | 0.000999 |
| MacLA | TCD8em | GRN | TNFRSF1B | 1.7735 | 0.3038 | 1.0386 | 0.000999 |
| MacLA | TCD8em | GSTP1 | CD44 | 1.2712 | 1.119 | 1.1951 | 0.000999 |
| MacLA | TCD8em | GSTP1 | IL7R | 1.2712 | 0.7183 | 0.9948 | 0.000999 |
| MacLA | TCD8em | GSTP1 | ITGA4 | 1.2712 | 0.2823 | 0.7768 | 0.000999 |
| MacLA | TCD8em | HLA-DRA | PTPRC | 3.0446 | 1.7912 | 2.4179 | 0.000999 |
| MacLA | TCD8em | ITGB1 | CD47 | 0.4657 | 0.4276 | 0.4467 | 0.000999 |
| MacLA | TCD8em | ITGB1 | ITGA4 | 0.4657 | 0.2823 | 0.374 | 0.000999 |
| MacLA | TCD8em | ITGB1 | RACK1 | 0.4657 | 2.4374 | 1.4515 | 0.000999 |
| MacLA | TCD8em | LGALS3 | CD6 | 2.3411 | 0.4204 | 1.3807 | 0.000999 |
| MacLA | TCD8em | LGALS3 | ITGB1 | 2.3411 | 0.2557 | 1.2984 | 0.000999 |
| MacLA | TCD8em | LGALS3 | SPN | 2.3411 | 0.2603 | 1.3007 | 0.000999 |
| MacLA | TCD8em | MIF | CD74 | 1.2502 | 2.4538 | 1.852 | 0.000999 |
| MacLA | TCD8em | MIF | ITGA4 | 1.2502 | 0.2823 | 0.7663 | 0.000999 |
| MacLA | TCD8em | MIF | TNFRSF14 | 1.2502 | 0.2915 | 0.7708 | 0.000999 |
| MacLA | TCD8em | MMP9 | CD44 | 0.9836 | 1.119 | 1.0513 | 0.000999 |
| MacLA | TCD8em | MMP9 | ITGB1 | 0.9836 | 0.2557 | 0.6197 | 0.000999 |
| MacLA | TCD8em | NAMPT | ANXA6 | 0.2975 | 0.552 | 0.4248 | 0.000999 |
| MacLA | TCD8em | NMB | F2R | 0.3014 | 0.2695 | 0.2854 | 0.000999 |
| MacLA | TCD8em | NMB | GPR65 | 0.3014 | 0.2625 | 0.2819 | 0.000999 |
| MacLA | TCD8em | RTN4 | CD44 | 1.0466 | 1.119 | 1.0828 | 0.000999 |
| MacLA | TCD8em | RTN4 | CD99 | 1.0466 | 1.1525 | 1.0995 | 0.000999 |
| MacLA | TCD8em | SPP1 | CD44 | 3.0735 | 1.119 | 2.0963 | 0.000999 |
| MacLA | TCD8em | SPP1 | ITGA4 | 3.0735 | 0.2823 | 1.6779 | 0.000999 |
| MacLA | TCD8em | SPP1 | ITGB1 | 3.0735 | 0.2557 | 1.6646 | 0.000999 |
| MacLA | TCD8em | CALR | CD44 | 0.9648 | 1.119 | 1.0419 | 0.003 |
| MacLA | TCD8em | CALR | IL7R | 0.9648 | 0.7183 | 0.8416 | 0.003 |
| MacLA | TCD8em | HSP90B1 | ITGB1 | 0.7134 | 0.2557 | 0.4846 | 0.005 |
| MacLA | TCD8em | HSP90B1 | TNFRSF1B | 0.7134 | 0.3038 | 0.5086 | 0.005 |

| | | | | | | | |
|---|---|---|---|---|---|---|---|
| MacLA | TCD8ex | A2M | TNFRSF14 | 0.5014 | 0.3094 | 0.4054 | 0.000999 |
| MacLA | TCD8ex | ADAM9 | ITGB1 | 0.3224 | 0.2995 | 0.3109 | 0.000999 |
| MacLA | TCD8ex | ADM | PTGER4 | 0.2957 | 0.3089 | 0.3023 | 0.000999 |
| MacLA | TCD8ex | ALCAM | CD6 | 0.3816 | 0.318 | 0.3498 | 0.000999 |
| MacLA | TCD8ex | ANXA1 | CD44 | 1.5201 | 0.8219 | 1.171 | 0.000999 |
| MacLA | TCD8ex | ANXA1 | CXCR3 | 1.5201 | 0.3647 | 0.9424 | 0.000999 |
| MacLA | TCD8ex | ANXA1 | CXCR4 | 1.5201 | 2.3472 | 1.9337 | 0.000999 |
| MacLA | TCD8ex | ANXA1 | CXCR6 | 1.5201 | 0.6626 | 1.0913 | 0.000999 |
| MacLA | TCD8ex | ANXA1 | ITGA4 | 1.5201 | 0.4432 | 0.9817 | 0.000999 |
| MacLA | TCD8ex | ANXA1 | VCAM1 | 1.5201 | 0.4462 | 0.9832 | 0.000999 |
| MacLA | TCD8ex | APOE | VCAM1 | 3.1824 | 0.4462 | 1.8143 | 0.000999 |
| MacLA | TCD8ex | APP | CXCR3 | 0.2852 | 0.3647 | 0.3249 | 0.000999 |
| MacLA | TCD8ex | APP | CXCR4 | 0.2852 | 2.3472 | 1.3162 | 0.000999 |
| MacLA | TCD8ex | APP | CXCR6 | 0.2852 | 0.6626 | 0.4739 | 0.000999 |
| MacLA | TCD8ex | APP | TNFRSF14 | 0.2852 | 0.3094 | 0.2973 | 0.000999 |
| MacLA | TCD8ex | CD86 | CTLA4 | 0.2798 | 0.4883 | 0.384 | 0.000999 |
| MacLA | TCD8ex | COL1A1 | CD44 | 0.2943 | 0.8219 | 0.5581 | 0.000999 |
| MacLA | TCD8ex | COL1A1 | ITGB1 | 0.2943 | 0.2995 | 0.2969 | 0.000999 |
| MacLA | TCD8ex | COPA | CD8A | 0.3451 | 1.787 | 1.066 | 0.000999 |
| MacLA | TCD8ex | COPA | ITGA4 | 0.3451 | 0.4432 | 0.3942 | 0.000999 |
| MacLA | TCD8ex | COPA | VCAM1 | 0.3451 | 0.4462 | 0.3957 | 0.000999 |
| MacLA | TCD8ex | CXCL16 | CXCR3 | 0.6306 | 0.3647 | 0.4976 | 0.000999 |
| MacLA | TCD8ex | CXCL16 | CXCR4 | 0.6306 | 2.3472 | 1.4889 | 0.000999 |
| MacLA | TCD8ex | CXCL16 | CXCR6 | 0.6306 | 0.6626 | 0.6466 | 0.000999 |
| MacLA | TCD8ex | CXCL3 | CXCR3 | 0.2783 | 0.3647 | 0.3215 | 0.000999 |
| MacLA | TCD8ex | CXCL3 | CXCR4 | 0.2783 | 2.3472 | 1.3128 | 0.000999 |
| MacLA | TCD8ex | CXCL3 | CXCR6 | 0.2783 | 0.6626 | 0.4704 | 0.000999 |
| MacLA | TCD8ex | FN1 | CD44 | 1.0963 | 0.8219 | 0.9591 | 0.000999 |
| MacLA | TCD8ex | FN1 | ITGA4 | 1.0963 | 0.4432 | 0.7698 | 0.000999 |
| MacLA | TCD8ex | FN1 | ITGB1 | 1.0963 | 0.2995 | 0.6979 | 0.000999 |

| | | | | | | | |
|---|---|---|---|---|---|---|---|
| MacLA | TCD8ex | FN1 | ITGB7 | 1.0963 | 0.3007 | 0.6985 | 0.000999 |
| MacLA | TCD8ex | FN1 | RPSA | 1.0963 | 2.2697 | 1.683 | 0.000999 |
| MacLA | TCD8ex | GRN | TNFRSF1B | 1.7735 | 0.6436 | 1.2086 | 0.000999 |
| MacLA | TCD8ex | GSTP1 | CD44 | 1.2712 | 0.8219 | 1.0466 | 0.000999 |
| MacLA | TCD8ex | GSTP1 | ITGA4 | 1.2712 | 0.4432 | 0.8572 | 0.000999 |
| MacLA | TCD8ex | GSTP1 | VCAM1 | 1.2712 | 0.4462 | 0.8587 | 0.000999 |
| MacLA | TCD8ex | HLA-DRA | PDCD1 | 3.0446 | 0.6109 | 1.8277 | 0.000999 |
| MacLA | TCD8ex | HLA-DRA | PTPRC | 3.0446 | 1.6762 | 2.3604 | 0.000999 |
| MacLA | TCD8ex | LGALS3 | CD6 | 2.3411 | 0.318 | 1.3295 | 0.000999 |
| MacLA | TCD8ex | LGALS3 | ITGB1 | 2.3411 | 0.2995 | 1.3203 | 0.000999 |
| MacLA | TCD8ex | LGALS3 | SPN | 2.3411 | 0.3486 | 1.3448 | 0.000999 |
| MacLA | TCD8ex | MIF | CD74 | 1.2502 | 2.5058 | 1.878 | 0.000999 |
| MacLA | TCD8ex | MIF | ITGA4 | 1.2502 | 0.4432 | 0.8467 | 0.000999 |
| MacLA | TCD8ex | MIF | TNFRSF14 | 1.2502 | 0.3094 | 0.7798 | 0.000999 |
| MacLA | TCD8ex | MMP9 | CD44 | 0.9836 | 0.8219 | 0.9028 | 0.000999 |
| MacLA | TCD8ex | MMP9 | ITGB1 | 0.9836 | 0.2995 | 0.6415 | 0.000999 |
| MacLA | TCD8ex | NMB | GPR65 | 0.3014 | 0.3904 | 0.3459 | 0.000999 |
| MacLA | TCD8ex | RTN4 | CD44 | 1.0466 | 0.8219 | 0.9343 | 0.000999 |
| MacLA | TCD8ex | RTN4 | CD99 | 1.0466 | 1.2049 | 1.1257 | 0.000999 |
| MacLA | TCD8ex | SPP1 | CD44 | 3.0735 | 0.8219 | 1.9477 | 0.000999 |
| MacLA | TCD8ex | SPP1 | ITGA4 | 3.0735 | 0.4432 | 1.7584 | 0.000999 |
| MacLA | TCD8ex | SPP1 | ITGB1 | 3.0735 | 0.2995 | 1.6865 | 0.000999 |
| MacLA | TCD8ex | TGFB1 | ITGB1 | 0.3223 | 0.2995 | 0.3109 | 0.002 |
| MacLA | TGD | A2M | TNFRSF14 | 0.5014 | 0.4642 | 0.4828 | 0.000999 |
| MacLA | TGD | ADAM9 | ITGB1 | 0.3224 | 0.3657 | 0.344 | 0.000999 |
| MacLA | TGD | ADM | PTGDR | 0.2957 | 0.7153 | 0.5055 | 0.000999 |
| MacLA | TGD | AIMP1 | RACK1 | 0.2907 | 2.7287 | 1.5097 | 0.000999 |
| MacLA | TGD | ALCAM | CD6 | 0.3816 | 0.4684 | 0.425 | 0.000999 |
| MacLA | TGD | ANXA1 | CD44 | 1.5201 | 1.1342 | 1.3272 | 0.000999 |
| MacLA | TGD | ANXA1 | CXCR3 | 1.5201 | 1.1748 | 1.3474 | 0.000999 |

| | | | | | | | |
|---|---|---|---|---|---|---|---|
| MacLA | TGD | ANXA1 | CXCR4 | 1.5201 | 1.4894 | 1.5048 | 0.000999 |
| MacLA | TGD | ANXA1 | ITGA4 | 1.5201 | 0.3968 | 0.9585 | 0.000999 |
| MacLA | TGD | APOE | SORL1 | 3.1824 | 0.3318 | 1.7571 | 0.000999 |
| MacLA | TGD | APP | CXCR3 | 0.2852 | 1.1748 | 0.73 | 0.000999 |
| MacLA | TGD | APP | CXCR4 | 0.2852 | 1.4894 | 0.8873 | 0.000999 |
| MacLA | TGD | APP | SORL1 | 0.2852 | 0.3318 | 0.3085 | 0.000999 |
| MacLA | TGD | APP | TGFBR2 | 0.2852 | 0.3677 | 0.3264 | 0.000999 |
| MacLA | TGD | APP | TNFRSF14 | 0.2852 | 0.4642 | 0.3747 | 0.000999 |
| MacLA | TGD | CALR | IL7R | 0.9648 | 1.3862 | 1.1755 | 0.000999 |
| MacLA | TGD | COPA | CD8A | 0.3451 | 1.746 | 1.0456 | 0.000999 |
| MacLA | TGD | COPA | ITGA4 | 0.3451 | 0.3968 | 0.371 | 0.000999 |
| MacLA | TGD | CXCL16 | CXCR3 | 0.6306 | 1.1748 | 0.9027 | 0.000999 |
| MacLA | TGD | CXCL16 | CXCR4 | 0.6306 | 1.4894 | 1.06 | 0.000999 |
| MacLA | TGD | CXCL3 | CXCR3 | 0.2783 | 1.1748 | 0.7265 | 0.000999 |
| MacLA | TGD | CXCL3 | CXCR4 | 0.2783 | 1.4894 | 0.8839 | 0.000999 |
| MacLA | TGD | FN1 | CD44 | 1.0963 | 1.1342 | 1.1153 | 0.000999 |
| MacLA | TGD | FN1 | ITGA4 | 1.0963 | 0.3968 | 0.7466 | 0.000999 |
| MacLA | TGD | FN1 | ITGB1 | 1.0963 | 0.3657 | 0.731 | 0.000999 |
| MacLA | TGD | FN1 | ITGB7 | 1.0963 | 0.459 | 0.7777 | 0.000999 |
| MacLA | TGD | FN1 | RPSA | 1.0963 | 2.8814 | 1.9888 | 0.000999 |
| MacLA | TGD | GRN | TNFRSF1A | 1.7735 | 0.3099 | 1.0417 | 0.000999 |
| MacLA | TGD | GSTP1 | IL7R | 1.2712 | 1.3862 | 1.3287 | 0.000999 |
| MacLA | TGD | GSTP1 | ITGA4 | 1.2712 | 0.3968 | 0.834 | 0.000999 |
| MacLA | TGD | HLA-DRA | PTPRC | 3.0446 | 1.8653 | 2.455 | 0.000999 |
| MacLA | TGD | ITGAM | ICAM1 | 0.3326 | 0.2915 | 0.312 | 0.000999 |
| MacLA | TGD | ITGB1 | RACK1 | 0.4657 | 2.7287 | 1.5972 | 0.000999 |
| MacLA | TGD | LGALS3 | CD6 | 2.3411 | 0.4684 | 1.4047 | 0.000999 |
| MacLA | TGD | LGALS3 | ITGB1 | 2.3411 | 0.3657 | 1.3534 | 0.000999 |
| MacLA | TGD | LGALS3 | SPN | 2.3411 | 0.5644 | 1.4527 | 0.000999 |
| MacLA | TGD | LGALS3 | TGFBR2 | 2.3411 | 0.3677 | 1.3544 | 0.000999 |

| | | | | | | | |
|---|---|---|---|---|---|---|---|
| MacLA | TGD | LRPAP1 | SORL1 | 0.5574 | 0.3318 | 0.4446 | 0.000999 |
| MacLA | TGD | MMP9 | CD44 | 0.9836 | 1.1342 | 1.0589 | 0.000999 |
| MacLA | TGD | MMP9 | ITGB1 | 0.9836 | 0.3657 | 0.6746 | 0.000999 |
| MacLA | TGD | NAMPT | ANXA6 | 0.2975 | 0.8553 | 0.5764 | 0.000999 |
| MacLA | TGD | NMB | GPR65 | 0.3014 | 0.5368 | 0.4191 | 0.000999 |
| MacLA | TGD | RTN4 | CD44 | 1.0466 | 1.1342 | 1.0904 | 0.000999 |
| MacLA | TGD | RTN4 | CD99 | 1.0466 | 1.7154 | 1.381 | 0.000999 |
| MacLA | TGD | SPP1 | CD44 | 3.0735 | 1.1342 | 2.1038 | 0.000999 |
| MacLA | TGD | SPP1 | ITGA4 | 3.0735 | 0.3968 | 1.7352 | 0.000999 |
| MacLA | TGD | SPP1 | ITGB1 | 3.0735 | 0.3657 | 1.7196 | 0.000999 |
| MacLA | TGD | ITGB1 | ITGA4 | 0.4657 | 0.3968 | 0.4313 | 0.002 |
| NKT | MacLA | ANXA1 | C5AR1 | 1.2259 | 0.4636 | 0.8448 | 0.000999 |
| NKT | MacLA | ANXA1 | CCR1 | 1.2259 | 0.3657 | 0.7958 | 0.000999 |
| NKT | MacLA | ANXA1 | CD4 | 1.2259 | 0.3763 | 0.8011 | 0.000999 |
| NKT | MacLA | ANXA1 | CD44 | 1.2259 | 1.4055 | 1.3157 | 0.000999 |
| NKT | MacLA | ANXA1 | FPR3 | 1.2259 | 0.4512 | 0.8386 | 0.000999 |
| NKT | MacLA | ARF1 | CD4 | 0.7991 | 0.3763 | 0.5877 | 0.000999 |
| NKT | MacLA | ARF1 | CD44 | 0.7991 | 1.4055 | 1.1023 | 0.000999 |
| NKT | MacLA | CALR | CD44 | 0.9479 | 1.4055 | 1.1767 | 0.000999 |
| NKT | MacLA | CALR | LRP1 | 0.9479 | 0.3427 | 0.6453 | 0.000999 |
| NKT | MacLA | CD2 | CD4 | 1.1465 | 0.3763 | 0.7614 | 0.000999 |
| NKT | MacLA | CD6 | ALCAM | 0.4881 | 0.3816 | 0.4349 | 0.000999 |
| NKT | MacLA | GSTP1 | CD44 | 0.7909 | 1.4055 | 1.0982 | 0.000999 |
| NKT | MacLA | HLA-DRA | CD4 | 0.6626 | 0.3763 | 0.5195 | 0.000999 |
| NKT | MacLA | HMGB1 | LY96 | 1.195 | 0.8103 | 1.0026 | 0.000999 |
| NKT | MacLA | HMGB2 | CD44 | 0.6467 | 1.4055 | 1.0261 | 0.000999 |
| NKT | MacLA | HMGB2 | DAB2 | 0.6467 | 0.9195 | 0.7831 | 0.000999 |
| NKT | MacLA | HSP90B1 | LRP1 | 0.6138 | 0.3427 | 0.4782 | 0.000999 |
| NKT | MacLA | IL16 | CD4 | 0.3556 | 0.3763 | 0.3659 | 0.000999 |
| NKT | MacLA | ITGA4 | CD44 | 0.2946 | 1.4055 | 0.85 | 0.000999 |

| | | | | | | | |
|---|---|---|---|---|---|---|---|
| NKT | MacLA | ITGA4 | TFRC | 0.2946 | 0.6885 | 0.4915 | 0.000999 |
| NKT | MacLA | ITGB1 | CD151 | 0.488 | 0.2853 | 0.3866 | 0.000999 |
| NKT | MacLA | ITGB1 | CD36 | 0.488 | 0.4467 | 0.4673 | 0.000999 |
| NKT | MacLA | ITGB1 | DAB2 | 0.488 | 0.9195 | 0.7037 | 0.000999 |
| NKT | MacLA | ITGB1 | ENG | 0.488 | 0.3236 | 0.4058 | 0.000999 |
| NKT | MacLA | ITGB1 | ITGAX | 0.488 | 0.4923 | 0.4901 | 0.000999 |
| NKT | MacLA | ITGB1 | PLAUR | 0.488 | 1.1605 | 0.8242 | 0.000999 |
| NKT | MacLA | ITGB2 | ITGAX | 1.6481 | 0.4923 | 1.0702 | 0.000999 |
| NKT | MacLA | ITGB2 | PLAUR | 1.6481 | 1.1605 | 1.4043 | 0.000999 |
| NKT | MacLA | LRPAP1 | LRP1 | 0.2807 | 0.3427 | 0.3117 | 0.000999 |
| NKT | MacLA | MIF | CD74 | 0.6657 | 3.1562 | 1.911 | 0.000999 |
| NKT | MacLA | PSAP | LRP1 | 0.6777 | 0.3427 | 0.5102 | 0.000999 |
| NKT | MacLA | PTPRC | CD4 | 1.8797 | 0.3763 | 1.128 | 0.000999 |
| NKT | MacLA | PTPRC | CD44 | 1.8797 | 1.4055 | 1.6426 | 0.000999 |
| NKT | MacLA | PTPRC | IFNAR1 | 1.8797 | 0.3022 | 1.0909 | 0.000999 |
| NKT | MacLA | RPS19 | C5AR1 | 3.5336 | 0.4636 | 1.9986 | 0.000999 |
| NKT | MacLA | RPS19 | CD4 | 3.5336 | 0.3763 | 1.955 | 0.000999 |
| NKT | MacLA | RTN4 | CD44 | 0.4374 | 1.4055 | 0.9214 | 0.000999 |
| NKT | MacLA | TGFB1 | ENG | 0.5441 | 0.3236 | 0.4338 | 0.000999 |
| NKT | MacLA | TGFB1 | SDC2 | 0.5441 | 0.8668 | 0.7054 | 0.000999 |
| NKT | MacLA | TIMP1 | LRP1 | 0.3005 | 0.3427 | 0.3216 | 0.000999 |
| NKT | MacLA | XCL2 | PTAFR | 0.2774 | 0.2667 | 0.2721 | 0.000999 |
| TCD4ex | MacLA | ANXA1 | C5AR1 | 0.5174 | 0.4636 | 0.4905 | 0.000999 |
| TCD4ex | MacLA | ANXA1 | CCR1 | 0.5174 | 0.3657 | 0.4416 | 0.000999 |
| TCD4ex | MacLA | ANXA1 | CD44 | 0.5174 | 1.4055 | 0.9615 | 0.000999 |
| TCD4ex | MacLA | ANXA1 | FPR3 | 0.5174 | 0.4512 | 0.4843 | 0.000999 |
| TCD4ex | MacLA | APOE | LRP1 | 0.2557 | 0.3427 | 0.2992 | 0.000999 |
| TCD4ex | MacLA | APOE | SDC2 | 0.2557 | 0.8668 | 0.5612 | 0.000999 |
| TCD4ex | MacLA | ARF1 | CD44 | 0.6827 | 1.4055 | 1.0441 | 0.000999 |
| TCD4ex | MacLA | CALR | CD44 | 0.748 | 1.4055 | 1.0767 | 0.000999 |

| | | | | | | | |
|---|---|---|---|---|---|---|---|
| TCD4ex | MacLA | CALR | LRP1 | 0.748 | 0.3427 | 0.5453 | 0.000999 |
| TCD4ex | MacLA | CD6 | ALCAM | 0.5338 | 0.3816 | 0.4577 | 0.000999 |
| TCD4ex | MacLA | CXCL13 | C5AR1 | 2.3367 | 0.4636 | 1.4001 | 0.000999 |
| TCD4ex | MacLA | CXCL13 | CCR1 | 2.3367 | 0.3657 | 1.3512 | 0.000999 |
| TCD4ex | MacLA | GSTP1 | CD44 | 0.4481 | 1.4055 | 0.9268 | 0.000999 |
| TCD4ex | MacLA | HMGB1 | LY96 | 1.3876 | 0.8103 | 1.0989 | 0.000999 |
| TCD4ex | MacLA | HMGB2 | CD44 | 1.0243 | 1.4055 | 1.2149 | 0.000999 |
| TCD4ex | MacLA | HMGB2 | DAB2 | 1.0243 | 0.9195 | 0.9719 | 0.000999 |
| TCD4ex | MacLA | HSP90B1 | LRP1 | 0.3944 | 0.3427 | 0.3685 | 0.000999 |
| TCD4ex | MacLA | ICAM3 | ITGB2 | 0.573 | 1.2219 | 0.8974 | 0.000999 |
| TCD4ex | MacLA | ITGB1 | CD36 | 0.4615 | 0.4467 | 0.4541 | 0.000999 |
| TCD4ex | MacLA | ITGB1 | DAB2 | 0.4615 | 0.9195 | 0.6905 | 0.000999 |
| TCD4ex | MacLA | ITGB1 | ENG | 0.4615 | 0.3236 | 0.3925 | 0.000999 |
| TCD4ex | MacLA | ITGB1 | ITGAX | 0.4615 | 0.4923 | 0.4769 | 0.000999 |
| TCD4ex | MacLA | ITGB1 | PLAUR | 0.4615 | 1.1605 | 0.811 | 0.000999 |
| TCD4ex | MacLA | ITGB2 | ITGAM | 0.7871 | 0.3326 | 0.5598 | 0.000999 |
| TCD4ex | MacLA | ITGB2 | ITGAX | 0.7871 | 0.4923 | 0.6397 | 0.000999 |
| TCD4ex | MacLA | ITGB2 | PLAUR | 0.7871 | 1.1605 | 0.9738 | 0.000999 |
| TCD4ex | MacLA | LRPAP1 | LRP1 | 0.2809 | 0.3427 | 0.3118 | 0.000999 |
| TCD4ex | MacLA | MIF | CD74 | 0.6999 | 3.1562 | 1.9281 | 0.000999 |
| TCD4ex | MacLA | NMB | PTAFR | 0.6391 | 0.2667 | 0.4529 | 0.000999 |
| TCD4ex | MacLA | PSAP | LRP1 | 0.5291 | 0.3427 | 0.4359 | 0.000999 |
| TCD4ex | MacLA | PTPRC | CD44 | 1.7493 | 1.4055 | 1.5774 | 0.000999 |
| TCD4ex | MacLA | PTPRC | IFNAR1 | 1.7493 | 0.3022 | 1.0258 | 0.000999 |
| TCD4ex | MacLA | RPS19 | C5AR1 | 3.6329 | 0.4636 | 2.0483 | 0.000999 |
| TCD4ex | MacLA | RTN4 | CD44 | 0.355 | 1.4055 | 0.8803 | 0.000999 |
| TCD4ex | MacLA | TIMP1 | LRP1 | 0.3374 | 0.3427 | 0.3401 | 0.000999 |
| TCD4ex | MacLA | ITGB1 | CD151 | 0.4615 | 0.2853 | 0.3734 | 0.002 |
| TCD4ex | MacLA | SPP1 | CD44 | 0.3314 | 1.4055 | 0.8684 | 0.005 |
| TCD4reg | MacLA | APOE | LRP1 | 0.2934 | 0.3427 | 0.318 | 0.000999 |

| | | | | | | | |
|---|---|---|---|---|---|---|---|
| TCD4reg | MacLA | APOE | SDC2 | 0.2934 | 0.8668 | 0.5801 | 0.000999 |
| TCD4reg | MacLA | CALR | LRP1 | 0.6835 | 0.3427 | 0.5131 | 0.000999 |
| TCD4reg | MacLA | CD6 | ALCAM | 0.2651 | 0.3816 | 0.3233 | 0.000999 |
| TCD4reg | MacLA | HMGB1 | LY96 | 1.2547 | 0.8103 | 1.0325 | 0.000999 |
| TCD4reg | MacLA | HMGB2 | DAB2 | 0.616 | 0.9195 | 0.7677 | 0.000999 |
| TCD4reg | MacLA | HSP90B1 | LRP1 | 0.4221 | 0.3427 | 0.3824 | 0.000999 |
| TCD4reg | MacLA | ICAM3 | ITGB2 | 0.6663 | 1.2219 | 0.9441 | 0.000999 |
| TCD4reg | MacLA | ITGB1 | CD151 | 0.4001 | 0.2853 | 0.3427 | 0.000999 |
| TCD4reg | MacLA | ITGB1 | CD36 | 0.4001 | 0.4467 | 0.4234 | 0.000999 |
| TCD4reg | MacLA | ITGB1 | DAB2 | 0.4001 | 0.9195 | 0.6598 | 0.000999 |
| TCD4reg | MacLA | ITGB1 | ENG | 0.4001 | 0.3236 | 0.3618 | 0.000999 |
| TCD4reg | MacLA | ITGB1 | ITGAX | 0.4001 | 0.4923 | 0.4462 | 0.000999 |
| TCD4reg | MacLA | ITGB1 | PLAUR | 0.4001 | 1.1605 | 0.7803 | 0.000999 |
| TCD4reg | MacLA | ITGB2 | ITGAM | 0.6238 | 0.3326 | 0.4782 | 0.000999 |
| TCD4reg | MacLA | ITGB2 | ITGAX | 0.6238 | 0.4923 | 0.558 | 0.000999 |
| TCD4reg | MacLA | ITGB2 | PLAUR | 0.6238 | 1.1605 | 0.8922 | 0.000999 |
| TCD4reg | MacLA | LGALS3 | CLEC7A | 0.4492 | 0.3231 | 0.3861 | 0.000999 |
| TCD4reg | MacLA | LGALS3 | FCGR2A | 0.4492 | 0.3966 | 0.4229 | 0.000999 |
| TCD4reg | MacLA | LGALS3 | FCGR2B | 0.4492 | 0.3549 | 0.402 | 0.000999 |
| TCD4reg | MacLA | LGALS3 | TFRC | 0.4492 | 0.6885 | 0.5688 | 0.000999 |
| TCD4reg | MacLA | LRPAP1 | LRP1 | 0.3101 | 0.3427 | 0.3264 | 0.000999 |
| TCD4reg | MacLA | MIF | CD74 | 0.6295 | 3.1562 | 1.8929 | 0.000999 |
| TCD4reg | MacLA | PSAP | LRP1 | 0.5938 | 0.3427 | 0.4683 | 0.000999 |
| TCD4reg | MacLA | PTPRC | CD44 | 1.6603 | 1.4055 | 1.5329 | 0.000999 |
| TCD4reg | MacLA | PTPRC | IFNAR1 | 1.6603 | 0.3022 | 0.9813 | 0.000999 |
| TCD4reg | MacLA | RPS19 | C5AR1 | 3.6897 | 0.4636 | 2.0766 | 0.000999 |
| TCD4reg | MacLA | TIMP1 | LRP1 | 0.4362 | 0.3427 | 0.3894 | 0.000999 |
| TCD4reg | MacLA | ARF1 | CD44 | 0.7185 | 1.4055 | 1.062 | 0.002 |
| TCD4reg | MacLA | CALR | CD44 | 0.6835 | 1.4055 | 1.0445 | 0.002 |
| TCD4reg | MacLA | HMGB2 | CD44 | 0.616 | 1.4055 | 1.0108 | 0.002 |

| | | | | | | | |
|---|---|---|---|---|---|---|---|
| TCD4reg | MacLA | RTN4 | CD44 | 0.3528 | 1.4055 | 0.8791 | 0.003 |
| TCD4reg | MacLA | GSTP1 | CD44 | 0.4917 | 1.4055 | 0.9486 | 0.004 |
| TCD4reg | MacLA | SPP1 | CD44 | 0.3125 | 1.4055 | 0.859 | 0.00599 |
| TCD8em | MacLA | ANXA1 | C5AR1 | 1.0693 | 0.4636 | 0.7664 | 0.000999 |
| TCD8em | MacLA | ANXA1 | CCR1 | 1.0693 | 0.3657 | 0.7175 | 0.000999 |
| TCD8em | MacLA | ANXA1 | CD4 | 1.0693 | 0.3763 | 0.7228 | 0.000999 |
| TCD8em | MacLA | ANXA1 | FPR3 | 1.0693 | 0.4512 | 0.7602 | 0.000999 |
| TCD8em | MacLA | APOE | LRP1 | 0.2791 | 0.3427 | 0.3109 | 0.000999 |
| TCD8em | MacLA | APOE | SDC2 | 0.2791 | 0.8668 | 0.573 | 0.000999 |
| TCD8em | MacLA | ARF1 | CD4 | 0.6684 | 0.3763 | 0.5223 | 0.000999 |
| TCD8em | MacLA | CALR | LRP1 | 0.7141 | 0.3427 | 0.5284 | 0.000999 |
| TCD8em | MacLA | CD2 | CD4 | 1.5961 | 0.3763 | 0.9862 | 0.000999 |
| TCD8em | MacLA | CD6 | ALCAM | 0.4204 | 0.3816 | 0.401 | 0.000999 |
| TCD8em | MacLA | HLA-DMA | CD4 | 0.2969 | 0.3763 | 0.3366 | 0.000999 |
| TCD8em | MacLA | HLA-DRA | CD4 | 0.952 | 0.3763 | 0.6642 | 0.000999 |
| TCD8em | MacLA | HMGB1 | LY96 | 1.3636 | 0.8103 | 1.0869 | 0.000999 |
| TCD8em | MacLA | HMGB2 | DAB2 | 0.7212 | 0.9195 | 0.8203 | 0.000999 |
| TCD8em | MacLA | HSP90B1 | ITGB1 | 0.471 | 0.4657 | 0.4684 | 0.000999 |
| TCD8em | MacLA | HSP90B1 | LRP1 | 0.471 | 0.3427 | 0.4068 | 0.000999 |
| TCD8em | MacLA | IFNG | IFNGR1 | 0.3967 | 0.4968 | 0.4468 | 0.000999 |
| TCD8em | MacLA | IFNG | IFNGR2 | 0.3967 | 0.412 | 0.4044 | 0.000999 |
| TCD8em | MacLA | IFNG | PTPN6 | 0.3967 | 0.5493 | 0.473 | 0.000999 |
| TCD8em | MacLA | IL16 | CD4 | 0.4085 | 0.3763 | 0.3924 | 0.000999 |
| TCD8em | MacLA | ITGA4 | TFRC | 0.2823 | 0.6885 | 0.4854 | 0.000999 |
| TCD8em | MacLA | ITGB1 | CD151 | 0.2557 | 0.2853 | 0.2705 | 0.000999 |
| TCD8em | MacLA | ITGB1 | CD36 | 0.2557 | 0.4467 | 0.3512 | 0.000999 |
| TCD8em | MacLA | ITGB1 | DAB2 | 0.2557 | 0.9195 | 0.5876 | 0.000999 |
| TCD8em | MacLA | ITGB1 | ENG | 0.2557 | 0.3236 | 0.2897 | 0.000999 |
| TCD8em | MacLA | ITGB1 | ITGAX | 0.2557 | 0.4923 | 0.374 | 0.000999 |
| TCD8em | MacLA | ITGB1 | PLAUR | 0.2557 | 1.1605 | 0.7081 | 0.000999 |

| | | | | | | | |
|---|---|---|---|---|---|---|---|
| TCD8em | MacLA | ITGB2 | ITGAM | 1.0018 | 0.3326 | 0.6672 | 0.000999 |
| TCD8em | MacLA | ITGB2 | ITGAX | 1.0018 | 0.4923 | 0.747 | 0.000999 |
| TCD8em | MacLA | ITGB2 | PLAUR | 1.0018 | 1.1605 | 1.0812 | 0.000999 |
| TCD8em | MacLA | LGALS3 | CLEC7A | 0.2689 | 0.3231 | 0.296 | 0.000999 |
| TCD8em | MacLA | LGALS3 | FCGR2A | 0.2689 | 0.3966 | 0.3327 | 0.000999 |
| TCD8em | MacLA | LGALS3 | FCGR2B | 0.2689 | 0.3549 | 0.3119 | 0.000999 |
| TCD8em | MacLA | LGALS3 | TFRC | 0.2689 | 0.6885 | 0.4787 | 0.000999 |
| TCD8em | MacLA | LRPAP1 | LRP1 | 0.2719 | 0.3427 | 0.3073 | 0.000999 |
| TCD8em | MacLA | LTB | ITGB1 | 0.6352 | 0.4657 | 0.5505 | 0.000999 |
| TCD8em | MacLA | MIF | CD74 | 0.6063 | 3.1562 | 1.8813 | 0.000999 |
| TCD8em | MacLA | PSAP | LRP1 | 0.5012 | 0.3427 | 0.422 | 0.000999 |
| TCD8em | MacLA | PTPRC | CD4 | 1.7912 | 0.3763 | 1.0837 | 0.000999 |
| TCD8em | MacLA | PTPRC | CD44 | 1.7912 | 1.4055 | 1.5983 | 0.000999 |
| TCD8em | MacLA | PTPRC | IFNAR1 | 1.7912 | 0.3022 | 1.0467 | 0.000999 |
| TCD8em | MacLA | RPS19 | C5AR1 | 3.536 | 0.4636 | 1.9998 | 0.000999 |
| TCD8em | MacLA | RPS19 | CD4 | 3.536 | 0.3763 | 1.9562 | 0.000999 |
| TCD8em | MacLA | SELL | CD4 | 0.258 | 0.3763 | 0.3171 | 0.000999 |
| TCD8em | MacLA | TIMP1 | LRP1 | 0.2667 | 0.3427 | 0.3047 | 0.000999 |
| TCD8em | MacLA | TNF | NRP1 | 0.2667 | 0.2611 | 0.2639 | 0.000999 |
| TCD8em | MacLA | XCL2 | PTAFR | 0.295 | 0.2667 | 0.2808 | 0.000999 |
| TCD8em | MacLA | ARF1 | CD44 | 0.6684 | 1.4055 | 1.0369 | 0.002 |
| TCD8em | MacLA | HMGB2 | CD44 | 0.7212 | 1.4055 | 1.0633 | 0.002 |
| TCD8em | MacLA | ITGA4 | CD44 | 0.2823 | 1.4055 | 0.8439 | 0.002 |
| TCD8em | MacLA | LGALS3 | ITGB1 | 0.2689 | 0.4657 | 0.3673 | 0.002 |
| TCD8em | MacLA | ANXA1 | CD44 | 1.0693 | 1.4055 | 1.2374 | 0.004 |
| TCD8em | MacLA | CALR | CD44 | 0.7141 | 1.4055 | 1.0598 | 0.004 |
| TCD8em | MacLA | GSTP1 | CD44 | 0.767 | 1.4055 | 1.0862 | 0.004 |
| TCD8em | MacLA | RTN4 | CD44 | 0.3632 | 1.4055 | 0.8843 | 0.004 |
| TCD8em | MacLA | SPP1 | ITGB1 | 0.3608 | 0.4657 | 0.4133 | 0.004 |
| TCD8em | MacLA | SPP1 | CD44 | 0.3608 | 1.4055 | 0.8831 | 0.00699 |

| | | | | | | | |
|---|---|---|---|---|---|---|---|
| TCD8ex | MacLA | ANXA1 | C5AR1 | 0.5645 | 0.4636 | 0.514 | 0.000999 |
| TCD8ex | MacLA | ANXA1 | CCR1 | 0.5645 | 0.3657 | 0.4651 | 0.000999 |
| TCD8ex | MacLA | ANXA1 | CD4 | 0.5645 | 0.3763 | 0.4704 | 0.000999 |
| TCD8ex | MacLA | ANXA1 | CD44 | 0.5645 | 1.4055 | 0.985 | 0.000999 |
| TCD8ex | MacLA | ANXA1 | FPR3 | 0.5645 | 0.4512 | 0.5079 | 0.000999 |
| TCD8ex | MacLA | APOE | LRP1 | 0.2814 | 0.3427 | 0.312 | 0.000999 |
| TCD8ex | MacLA | APOE | SDC2 | 0.2814 | 0.8668 | 0.5741 | 0.000999 |
| TCD8ex | MacLA | ARF1 | CD4 | 0.7835 | 0.3763 | 0.5799 | 0.000999 |
| TCD8ex | MacLA | ARF1 | CD44 | 0.7835 | 1.4055 | 1.0945 | 0.000999 |
| TCD8ex | MacLA | CALR | CD44 | 0.9634 | 1.4055 | 1.1844 | 0.000999 |
| TCD8ex | MacLA | CALR | LRP1 | 0.9634 | 0.3427 | 0.653 | 0.000999 |
| TCD8ex | MacLA | CD2 | CD4 | 2.0502 | 0.3763 | 1.2132 | 0.000999 |
| TCD8ex | MacLA | CD6 | ALCAM | 0.318 | 0.3816 | 0.3498 | 0.000999 |
| TCD8ex | MacLA | CXCL13 | C5AR1 | 1.7749 | 0.4636 | 1.1193 | 0.000999 |
| TCD8ex | MacLA | CXCL13 | CCR1 | 1.7749 | 0.3657 | 1.0703 | 0.000999 |
| TCD8ex | MacLA | GSTP1 | CD44 | 0.8286 | 1.4055 | 1.1171 | 0.000999 |
| TCD8ex | MacLA | HLA-DMA | CD4 | 0.3991 | 0.3763 | 0.3877 | 0.000999 |
| TCD8ex | MacLA | HLA-DRA | CD4 | 1.247 | 0.3763 | 0.8116 | 0.000999 |
| TCD8ex | MacLA | HMGB1 | LY96 | 1.5414 | 0.8103 | 1.1759 | 0.000999 |
| TCD8ex | MacLA | HMGB2 | CD44 | 0.85 | 1.4055 | 1.1278 | 0.000999 |
| TCD8ex | MacLA | HMGB2 | DAB2 | 0.85 | 0.9195 | 0.8847 | 0.000999 |
| TCD8ex | MacLA | HSP90B1 | LRP1 | 0.5666 | 0.3427 | 0.4547 | 0.000999 |
| TCD8ex | MacLA | ICAM3 | ITGB2 | 0.5405 | 1.2219 | 0.8812 | 0.000999 |
| TCD8ex | MacLA | IFNG | IFNGR1 | 0.8575 | 0.4968 | 0.6772 | 0.000999 |
| TCD8ex | MacLA | IFNG | IFNGR2 | 0.8575 | 0.412 | 0.6348 | 0.000999 |
| TCD8ex | MacLA | ITGA4 | CD44 | 0.4432 | 1.4055 | 0.9244 | 0.000999 |
| TCD8ex | MacLA | ITGA4 | TFRC | 0.4432 | 0.6885 | 0.5659 | 0.000999 |
| TCD8ex | MacLA | ITGB1 | CD36 | 0.2995 | 0.4467 | 0.3731 | 0.000999 |
| TCD8ex | MacLA | ITGB1 | DAB2 | 0.2995 | 0.9195 | 0.6095 | 0.000999 |
| TCD8ex | MacLA | ITGB1 | ENG | 0.2995 | 0.3236 | 0.3115 | 0.000999 |

| | | | | | | | |
|---|---|---|---|---|---|---|---|
| TCD8ex | MacLA | ITGB1 | ITGAX | 0.2995 | 0.4923 | 0.3959 | 0.000999 |
| TCD8ex | MacLA | ITGB1 | PLAUR | 0.2995 | 1.1605 | 0.73 | 0.000999 |
| TCD8ex | MacLA | ITGB2 | ITGAM | 0.7635 | 0.3326 | 0.548 | 0.000999 |
| TCD8ex | MacLA | ITGB2 | ITGAX | 0.7635 | 0.4923 | 0.6279 | 0.000999 |
| TCD8ex | MacLA | ITGB2 | PLAUR | 0.7635 | 1.1605 | 0.962 | 0.000999 |
| TCD8ex | MacLA | LGALS3 | CLEC7A | 0.3824 | 0.3231 | 0.3528 | 0.000999 |
| TCD8ex | MacLA | LGALS3 | FCGR2A | 0.3824 | 0.3966 | 0.3895 | 0.000999 |
| TCD8ex | MacLA | LGALS3 | FCGR2B | 0.3824 | 0.3549 | 0.3686 | 0.000999 |
| TCD8ex | MacLA | LGALS3 | TFRC | 0.3824 | 0.6885 | 0.5354 | 0.000999 |
| TCD8ex | MacLA | LRPAP1 | LRP1 | 0.2971 | 0.3427 | 0.3199 | 0.000999 |
| TCD8ex | MacLA | MIF | CD74 | 0.749 | 3.1562 | 1.9526 | 0.000999 |
| TCD8ex | MacLA | PSAP | LRP1 | 0.5016 | 0.3427 | 0.4221 | 0.000999 |
| TCD8ex | MacLA | PTPRC | CD4 | 1.6762 | 0.3763 | 1.0262 | 0.000999 |
| TCD8ex | MacLA | PTPRC | CD44 | 1.6762 | 1.4055 | 1.5408 | 0.000999 |
| TCD8ex | MacLA | PTPRC | IFNAR1 | 1.6762 | 0.3022 | 0.9892 | 0.000999 |
| TCD8ex | MacLA | RPS19 | C5AR1 | 3.8353 | 0.4636 | 2.1494 | 0.000999 |
| TCD8ex | MacLA | RPS19 | CD4 | 3.8353 | 0.3763 | 2.1058 | 0.000999 |
| TCD8ex | MacLA | RTN4 | CD44 | 0.3033 | 1.4055 | 0.8544 | 0.000999 |
| TCD8ex | MacLA | SPP1 | CD44 | 0.2978 | 1.4055 | 0.8517 | 0.000999 |
| TCD8ex | MacLA | VCAM1 | ITGB2 | 0.4462 | 1.2219 | 0.834 | 0.000999 |
| TCD8ex | MacLA | VCAM1 | TFRC | 0.4462 | 0.6885 | 0.5673 | 0.000999 |
| TCD8ex | MacLA | XCL2 | PTAFR | 0.3157 | 0.2667 | 0.2912 | 0.000999 |
| TGD | MacLA | ARF1 | CD4 | 0.9453 | 0.3763 | 0.6608 | 0.000999 |
| TGD | MacLA | ARF1 | CD44 | 0.9453 | 1.4055 | 1.1754 | 0.000999 |
| TGD | MacLA | CALR | LRP1 | 0.9627 | 0.3427 | 0.6527 | 0.000999 |
| TGD | MacLA | CD2 | CD4 | 1.4522 | 0.3763 | 0.9143 | 0.000999 |
| TGD | MacLA | CD6 | ALCAM | 0.4684 | 0.3816 | 0.425 | 0.000999 |
| TGD | MacLA | COL1A1 | CD36 | 0.4518 | 0.4467 | 0.4493 | 0.000999 |
| TGD | MacLA | COL1A1 | CD44 | 0.4518 | 1.4055 | 0.9287 | 0.000999 |
| TGD | MacLA | HMGB1 | LY96 | 1.7525 | 0.8103 | 1.2814 | 0.000999 |

| | | | | | | | |
|---|---|---|---|---|---|---|---|
| TGD | MacLA | HMGB2 | DAB2 | 0.3275 | 0.9195 | 0.6235 | 0.000999 |
| TGD | MacLA | HSP90B1 | LRP1 | 0.9338 | 0.3427 | 0.6382 | 0.000999 |
| TGD | MacLA | ICAM1 | ITGAM | 0.2915 | 0.3326 | 0.312 | 0.000999 |
| TGD | MacLA | ICAM1 | ITGAX | 0.2915 | 0.4923 | 0.3919 | 0.000999 |
| TGD | MacLA | IL16 | CD4 | 0.9357 | 0.3763 | 0.656 | 0.000999 |
| TGD | MacLA | ITGA4 | CD44 | 0.3968 | 1.4055 | 0.9012 | 0.000999 |
| TGD | MacLA | ITGA4 | TFRC | 0.3968 | 0.6885 | 0.5427 | 0.000999 |
| TGD | MacLA | ITGB1 | CD36 | 0.3657 | 0.4467 | 0.4062 | 0.000999 |
| TGD | MacLA | ITGB1 | DAB2 | 0.3657 | 0.9195 | 0.6426 | 0.000999 |
| TGD | MacLA | ITGB1 | ITGAX | 0.3657 | 0.4923 | 0.429 | 0.000999 |
| TGD | MacLA | ITGB1 | PLAUR | 0.3657 | 1.1605 | 0.7631 | 0.000999 |
| TGD | MacLA | ITGB2 | ITGAM | 1.4444 | 0.3326 | 0.8885 | 0.000999 |
| TGD | MacLA | ITGB2 | ITGAX | 1.4444 | 0.4923 | 0.9683 | 0.000999 |
| TGD | MacLA | ITGB2 | PLAUR | 1.4444 | 1.1605 | 1.3025 | 0.000999 |
| TGD | MacLA | LTB | ITGB1 | 1.6829 | 0.4657 | 1.0743 | 0.000999 |
| TGD | MacLA | MIF | CD74 | 1.6335 | 3.1562 | 2.3948 | 0.000999 |
| TGD | MacLA | PTPRC | CD4 | 1.8653 | 0.3763 | 1.1208 | 0.000999 |
| TGD | MacLA | PTPRC | CD44 | 1.8653 | 1.4055 | 1.6354 | 0.000999 |
| TGD | MacLA | PTPRC | IFNAR1 | 1.8653 | 0.3022 | 1.0838 | 0.000999 |
| TGD | MacLA | RPS19 | C5AR1 | 3.8902 | 0.4636 | 2.1769 | 0.000999 |
| TGD | MacLA | RPS19 | CD4 | 3.8902 | 0.3763 | 2.1332 | 0.000999 |
| TGD | MacLA | SELL | CD4 | 0.3555 | 0.3763 | 0.3659 | 0.000999 |
| TGD | MacLA | TGFB1 | ENG | 0.7703 | 0.3236 | 0.547 | 0.000999 |
| TGD | MacLA | TGFB1 | ITGB1 | 0.7703 | 0.4657 | 0.618 | 0.000999 |
| TGD | MacLA | TGFB1 | SDC2 | 0.7703 | 0.8668 | 0.8186 | 0.000999 |
| TGD | MacLA | XCL1 | PTAFR | 1.5822 | 0.2667 | 0.9244 | 0.000999 |
| TGD | MacLA | XCL2 | PTAFR | 0.4864 | 0.2667 | 0.3766 | 0.000999 |
| TGD | MacLA | HMGB2 | CD44 | 0.3275 | 1.4055 | 0.8665 | 0.002 |
| TGD | MacLA | LRPAP1 | LRP1 | 0.3161 | 0.3427 | 0.3294 | 0.002 |

| Supplementary Table 8: Communication results between macrophages and lymphocytes subpopulations, and malignant cells | | | | | | | |
|---|---|---|---|---|---|---|---|
| **MacLA vs. malignant cells** | | | | | | | |
| celltype_01 | celltype_02 | ligand | receptor | ligand_mean_expression | receptor_mean_expression | ligand_receptor_mean_expression | pvalue |
| MacLA | breasttumor | ADAM9 | ITGB1 | 0.3224 | 0.2939 | 0.3081 | 0.000999 |
| MacLA | breasttumor | ANXA1 | CD44 | 1.5201 | 0.3247 | 0.9224 | 0.000999 |
| MacLA | breasttumor | APOE | SCARB1 | 3.1823 | 0.2628 | 1.7225 | 0.000999 |
| MacLA | breasttumor | APOE | SDC2 | 3.1823 | 1.071 | 2.1267 | 0.000999 |
| MacLA | breasttumor | CALM1 | MYLK | 1.3467 | 0.4064 | 0.8765 | 0.000999 |
| MacLA | breasttumor | CALM1 | RPSA | 1.3467 | 2.358 | 1.8523 | 0.000999 |
| MacLA | breasttumor | CALM1 | TNFRSF1A | 1.3467 | 0.2536 | 0.8001 | 0.000999 |
| MacLA | breasttumor | CALM3 | MYLK | 0.8842 | 0.4064 | 0.6453 | 0.000999 |
| MacLA | breasttumor | CD99 | CD99 | 0.8115 | 0.3871 | 0.5993 | 0.000999 |
| MacLA | breasttumor | FN1 | CD44 | 1.0962 | 0.3247 | 0.7105 | 0.000999 |
| MacLA | breasttumor | FN1 | ITGB1 | 1.0962 | 0.2939 | 0.695 | 0.000999 |
| MacLA | breasttumor | FN1 | RPSA | 1.0962 | 2.358 | 1.7271 | 0.000999 |
| MacLA | breasttumor | FN1 | SDC2 | 1.0962 | 1.071 | 1.0836 | 0.000999 |
| MacLA | breasttumor | GRN | TNFRSF1A | 1.7735 | 0.2536 | 1.0135 | 0.000999 |
| MacLA | breasttumor | HLA-DRA | SCARB1 | 3.0445 | 0.2628 | 1.6536 | 0.000999 |
| MacLA | breasttumor | LGALS3 | ITGB1 | 2.341 | 0.2939 | 1.3174 | 0.000999 |
| MacLA | breasttumor | MMP9 | CD44 | 0.9835 | 0.3247 | 0.6541 | 0.000999 |
| MacLA | breasttumor | MMP9 | ITGB1 | 0.9835 | 0.2939 | 0.6387 | 0.000999 |
| MacLA | breasttumor | PTPRC | CD44 | 0.3665 | 0.3247 | 0.3456 | 0.000999 |
| MacLA | breasttumor | RTN4 | CD44 | 1.0465 | 0.3247 | 0.6856 | 0.000999 |
| MacLA | breasttumor | RTN4 | CD99 | 1.0465 | 0.3871 | 0.7168 | 0.000999 |
| MacLA | breasttumor | SERPINA1 | SDC2 | 0.6855 | 1.071 | 0.8783 | 0.000999 |
| MacLA | breasttumor | SPP1 | CD44 | 3.0733 | 0.3247 | 1.699 | 0.000999 |
| MacLA | breasttumor | SPP1 | ITGB1 | 3.0733 | 0.2939 | 1.6836 | 0.000999 |
| MacLA | breasttumor | TGFB1 | ITGB1 | 0.3223 | 0.2939 | 0.3081 | 0.000999 |
| MacLA | breasttumor | TGFB1 | SDC2 | 0.3223 | 1.071 | 0.6967 | 0.000999 |
| MacLA | breasttumor | ITGB1 | RACK1 | 0.4657 | 2.6334 | 1.5495 | 0.00799 |

| | | | | | | | |
|---|---|---|---|---|---|---|---|
| MacLA | colorectaltumor | APOE | SDC1 | 3.1823 | 0.2575 | 1.7199 | 0.000999 |
| MacLA | colorectaltumor | CD99 | CD99 | 0.8115 | 0.3443 | 0.5779 | 0.000999 |
| MacLA | colorectaltumor | FN1 | CD44 | 1.0962 | 0.2512 | 0.6737 | 0.000999 |
| MacLA | colorectaltumor | FN1 | ITGB1 | 1.0962 | 0.4214 | 0.7588 | 0.000999 |
| MacLA | colorectaltumor | FN1 | RPSA | 1.0962 | 2.251 | 1.6736 | 0.000999 |
| MacLA | colorectaltumor | FN1 | SDC1 | 1.0962 | 0.2575 | 0.6768 | 0.000999 |
| MacLA | colorectaltumor | LGALS3 | ITGB1 | 2.341 | 0.4214 | 1.3812 | 0.000999 |
| MacLA | colorectaltumor | LGALS3 | ITGB4 | 2.341 | 0.2913 | 1.3161 | 0.000999 |
| MacLA | colorectaltumor | MMP9 | CD44 | 0.9835 | 0.2512 | 0.6173 | 0.000999 |
| MacLA | colorectaltumor | MMP9 | ITGB1 | 0.9835 | 0.4214 | 0.7024 | 0.000999 |
| MacLA | colorectaltumor | RTN4 | CD99 | 1.0465 | 0.3443 | 0.6954 | 0.000999 |
| MacLA | colorectaltumor | SPP1 | CD44 | 3.0733 | 0.2512 | 1.6622 | 0.000999 |
| MacLA | colorectaltumor | SPP1 | ITGB1 | 3.0733 | 0.4214 | 1.7474 | 0.000999 |
| MacLA | colorectaltumor | SPP1 | SDC1 | 3.0733 | 0.2575 | 1.6654 | 0.000999 |
| MacLA | colorectaltumor | TGFB1 | ITGB1 | 0.3223 | 0.4214 | 0.3718 | 0.000999 |
| MacLA | colorectaltumor | ANXA1 | CD44 | 1.5201 | 0.2512 | 0.8856 | 0.002 |
| MacLA | colorectaltumor | CALM1 | RPSA | 1.3467 | 2.251 | 1.7989 | 0.002 |
| MacLA | colorectaltumor | ITGB1 | RACK1 | 0.4657 | 2.7146 | 1.5901 | 0.002 |
| MacLA | colorectaltumor | ITGB1 | CD151 | 0.4657 | 0.6439 | 0.5548 | 0.00899 |
| MacLA | livertumor | A2M | TNFRSF14 | 0.5014 | 0.2788 | 0.3901 | 0.000999 |
| MacLA | livertumor | ADAM9 | ITGB1 | 0.3224 | 1.6044 | 0.9634 | 0.000999 |
| MacLA | livertumor | ANXA1 | CD44 | 1.5201 | 0.8016 | 1.1608 | 0.000999 |
| MacLA | livertumor | ANXA1 | MET | 1.5201 | 0.2983 | 0.9092 | 0.000999 |
| MacLA | livertumor | ANXA1 | PTPRF | 1.5201 | 0.384 | 0.9521 | 0.000999 |
| MacLA | livertumor | APOE | SDC1 | 3.1823 | 0.5354 | 1.8589 | 0.000999 |
| MacLA | livertumor | APOE | SDC4 | 3.1823 | 0.6068 | 1.8946 | 0.000999 |
| MacLA | livertumor | CALM1 | INSR | 1.3467 | 0.3039 | 0.8253 | 0.000999 |
| MacLA | livertumor | CALM1 | RPSA | 1.3467 | 2.6317 | 1.9892 | 0.000999 |
| MacLA | livertumor | CD99 | EPHB4 | 0.8115 | 0.4535 | 0.6325 | 0.000999 |
| MacLA | livertumor | CD99 | INSR | 0.8115 | 0.3039 | 0.5577 | 0.000999 |

| | | | | | | | |
|---|---|---|---|---|---|---|---|
| MacLA | livertumor | COL1A1 | ITGA2 | 0.2943 | 0.3983 | 0.3463 | 0.000999 |
| MacLA | livertumor | COL1A1 | ITGA3 | 0.2943 | 0.3913 | 0.3428 | 0.000999 |
| MacLA | livertumor | COL1A1 | ITGB1 | 0.2943 | 1.6044 | 0.9494 | 0.000999 |
| MacLA | livertumor | COL1A1 | SDC1 | 0.2943 | 0.5354 | 0.4149 | 0.000999 |
| MacLA | livertumor | FN1 | CD44 | 1.0962 | 0.8016 | 0.9489 | 0.000999 |
| MacLA | livertumor | FN1 | ITGA2 | 1.0962 | 0.3983 | 0.7473 | 0.000999 |
| MacLA | livertumor | FN1 | ITGA3 | 1.0962 | 0.3913 | 0.7438 | 0.000999 |
| MacLA | livertumor | FN1 | ITGB1 | 1.0962 | 1.6044 | 1.3503 | 0.000999 |
| MacLA | livertumor | FN1 | RPSA | 1.0962 | 2.6317 | 1.864 | 0.000999 |
| MacLA | livertumor | FN1 | SDC1 | 1.0962 | 0.5354 | 0.8158 | 0.000999 |
| MacLA | livertumor | FN1 | TFRC | 1.0962 | 0.2899 | 0.693 | 0.000999 |
| MacLA | livertumor | HLA-A | ERBB3 | 2.4335 | 0.2644 | 1.349 | 0.000999 |
| MacLA | livertumor | HLA-DRA | EPHB4 | 3.0445 | 0.4535 | 1.749 | 0.000999 |
| MacLA | livertumor | HLA-E | EPHB4 | 1.2527 | 0.4535 | 0.8531 | 0.000999 |
| MacLA | livertumor | HLA-E | ITGA6 | 1.2527 | 0.5211 | 0.8869 | 0.000999 |
| MacLA | livertumor | HLA-E | PLXNB2 | 1.2527 | 0.265 | 0.7588 | 0.000999 |
| MacLA | livertumor | LGALS3 | ITGA2 | 2.341 | 0.3983 | 1.3696 | 0.000999 |
| MacLA | livertumor | LGALS3 | ITGB1 | 2.341 | 1.6044 | 1.9727 | 0.000999 |
| MacLA | livertumor | LGALS3 | ITGB4 | 2.341 | 0.5518 | 1.4464 | 0.000999 |
| MacLA | livertumor | LGALS3 | TFRC | 2.341 | 0.2899 | 1.3154 | 0.000999 |
| MacLA | livertumor | MMP9 | CD44 | 0.9835 | 0.8016 | 0.8925 | 0.000999 |
| MacLA | livertumor | MMP9 | CDH1 | 0.9835 | 0.4896 | 0.7366 | 0.000999 |
| MacLA | livertumor | MMP9 | ITGB1 | 0.9835 | 1.6044 | 1.294 | 0.000999 |
| MacLA | livertumor | PTPRC | INSR | 0.3665 | 0.3039 | 0.3352 | 0.000999 |
| MacLA | livertumor | PTPRC | MET | 0.3665 | 0.2983 | 0.3324 | 0.000999 |
| MacLA | livertumor | SPP1 | CD44 | 3.0733 | 0.8016 | 1.9375 | 0.000999 |
| MacLA | livertumor | SPP1 | ITGB1 | 3.0733 | 1.6044 | 2.3389 | 0.000999 |
| MacLA | livertumor | SPP1 | SDC1 | 3.0733 | 0.5354 | 1.8044 | 0.000999 |
| MacLA | livertumor | TGFB1 | ITGB1 | 0.3223 | 1.6044 | 0.9634 | 0.000999 |
| MacLA | livertumor | TGFB1 | VTN | 0.3223 | 0.7399 | 0.5311 | 0.000999 |

| | | | | | | | |
|---|---|---|---|---|---|---|---|
| MacLA | livertumor | ADM | RAMP1 | 0.2957 | 0.4499 | 0.3728 | 0.003 |
| MacLA | livertumor | PTPRC | CD44 | 0.3665 | 0.8016 | 0.584 | 0.003 |
| MacLA | livertumor | CD99 | CD99 | 0.8115 | 0.5906 | 0.7011 | 0.005 |
| MacLA | livertumor | AIMP1 | RACK1 | 0.2907 | 2.7655 | 1.5281 | 0.00999 |
| MacLA | lungtumor | APOE | SDC1 | 3.1823 | 0.2715 | 1.7269 | 0.000999 |
| MacLA | lungtumor | APOE | SDC4 | 3.1823 | 0.3516 | 1.767 | 0.000999 |
| MacLA | lungtumor | CALM1 | RPSA | 1.3467 | 1.6057 | 1.4762 | 0.000999 |
| MacLA | lungtumor | COL1A1 | SDC1 | 0.2943 | 0.2715 | 0.2829 | 0.000999 |
| MacLA | lungtumor | CXCL16 | CXCR4 | 0.6305 | 0.2684 | 0.4495 | 0.000999 |
| MacLA | lungtumor | CXCL3 | CXCR4 | 0.2783 | 0.2684 | 0.2733 | 0.000999 |
| MacLA | lungtumor | FN1 | CD44 | 1.0962 | 0.3802 | 0.7382 | 0.000999 |
| MacLA | lungtumor | FN1 | RPSA | 1.0962 | 1.6057 | 1.351 | 0.000999 |
| MacLA | lungtumor | FN1 | SDC1 | 1.0962 | 0.2715 | 0.6839 | 0.000999 |
| MacLA | lungtumor | ITGB1 | CD151 | 0.4657 | 0.6776 | 0.5716 | 0.000999 |
| MacLA | lungtumor | ITGB1 | CD46 | 0.4657 | 0.434 | 0.4499 | 0.000999 |
| MacLA | lungtumor | ITGB1 | CD47 | 0.4657 | 0.3255 | 0.3956 | 0.000999 |
| MacLA | lungtumor | ITGB1 | RACK1 | 0.4657 | 2.2108 | 1.3383 | 0.000999 |
| MacLA | lungtumor | ITGB1 | SDC1 | 0.4657 | 0.2715 | 0.3686 | 0.000999 |
| MacLA | lungtumor | MMP9 | CD44 | 0.9835 | 0.3802 | 0.6818 | 0.000999 |
| MacLA | lungtumor | PTPRC | CD44 | 0.3665 | 0.3802 | 0.3733 | 0.000999 |
| MacLA | lungtumor | PTPRC | CXCR4 | 0.3665 | 0.2684 | 0.3174 | 0.000999 |
| MacLA | lungtumor | RTN4 | CD44 | 1.0465 | 0.3802 | 0.7133 | 0.000999 |
| MacLA | lungtumor | RTN4 | CD99 | 1.0465 | 0.5938 | 0.8202 | 0.000999 |
| MacLA | lungtumor | SPP1 | CD44 | 3.0733 | 0.3802 | 1.7268 | 0.000999 |
| MacLA | lungtumor | SPP1 | SDC1 | 3.0733 | 0.2715 | 1.6724 | 0.000999 |
| MacLA | lungtumor | COL1A1 | CD44 | 0.2943 | 0.3802 | 0.3372 | 0.003 |
| MacLA | melanomatumo | AIMP1 | RACK1 | 0.2907 | 4.145 | 2.2179 | 0.000999 |
| MacLA | melanomatumo | ANXA1 | EDNRB | 1.5201 | 0.5666 | 1.0434 | 0.000999 |
| MacLA | melanomatumo | APP | EDNRB | 0.2852 | 0.5666 | 0.4259 | 0.000999 |
| MacLA | melanomatumo | CALM1 | RPSA | 1.3467 | 3.7787 | 2.5627 | 0.000999 |

| | | | | | | | |
|---|---|---|---|---|---|---|---|
| MacLA | melanomatumo | FN1 | RPSA | 1.0962 | 3.7787 | 2.4374 | 0.000999 |
| MacLA | melanomatumo | ITGB1 | RACK1 | 0.4657 | 4.145 | 2.3053 | 0.000999 |
| MacLA | melanomatumo | NAMPT | ANXA6 | 0.2975 | 0.3625 | 0.33 | 0.000999 |
| MacLA | melanomatumo | NMB | EDNRB | 0.3013 | 0.5666 | 0.434 | 0.000999 |
| MacLA | melanomatumo | RPS19 | RPSA | 3.509 | 3.7787 | 3.6438 | 0.000999 |
| MacLA | melanomatumo | SPP1 | CD44 | 3.0733 | 1.2653 | 2.1693 | 0.000999 |
| MacLA | melanomatumo | SPP1 | ITGB1 | 3.0733 | 0.3547 | 1.714 | 0.000999 |
| MacLA | melanomatumo | SPP1 | NCSTN | 3.0733 | 0.2614 | 1.6674 | 0.000999 |
| MacLA | melanomatumo | CALR | NCSTN | 0.9648 | 0.2614 | 0.6131 | 0.002 |
| MacLA | melanomatumo | LGALS3 | ITGB1 | 2.341 | 0.3547 | 1.3479 | 0.002 |
| MacLA | melanomatumo | A2M | TNFRSF14 | 0.5014 | 0.5069 | 0.5041 | 0.003 |
| MacLA | melanomatumo | APP | TNFRSF14 | 0.2852 | 0.5069 | 0.396 | 0.004 |
| MacLA | ovarytumor | ADAM9 | ITGB1 | 0.3224 | 0.2783 | 0.3003 | 0.000999 |
| MacLA | ovarytumor | ANXA1 | CXCR4 | 1.5201 | 0.3688 | 0.9444 | 0.000999 |
| MacLA | ovarytumor | AREG | FOLR1 | 0.2617 | 0.5619 | 0.4118 | 0.000999 |
| MacLA | ovarytumor | CALM1 | RPSA | 1.3467 | 2.9542 | 2.1504 | 0.000999 |
| MacLA | ovarytumor | COPA | FOLR1 | 0.3451 | 0.5619 | 0.4535 | 0.000999 |
| MacLA | ovarytumor | CXCL16 | CXCR4 | 0.6305 | 0.3688 | 0.4997 | 0.000999 |
| MacLA | ovarytumor | CXCL3 | CXCR4 | 0.2783 | 0.3688 | 0.3235 | 0.000999 |
| MacLA | ovarytumor | FN1 | ITGB1 | 1.0962 | 0.2783 | 0.6873 | 0.000999 |
| MacLA | ovarytumor | FN1 | RPSA | 1.0962 | 2.9542 | 2.0252 | 0.000999 |
| MacLA | ovarytumor | LGALS3 | ITGB1 | 2.341 | 0.2783 | 1.3096 | 0.000999 |
| MacLA | ovarytumor | MIF | CD74 | 1.2501 | 1.4431 | 1.3466 | 0.000999 |
| MacLA | ovarytumor | MMP9 | ITGB1 | 0.9835 | 0.2783 | 0.6309 | 0.000999 |
| MacLA | ovarytumor | PTPRC | CXCR4 | 0.3665 | 0.3688 | 0.3676 | 0.000999 |
| MacLA | ovarytumor | SPP1 | ITGB1 | 3.0733 | 0.2783 | 1.6758 | 0.000999 |
| MacLA | ovarytumor | TGFB1 | ITGB1 | 0.3223 | 0.2783 | 0.3003 | 0.000999 |
| MacLA | ovarytumor | ITGB1 | CD151 | 0.4657 | 0.4981 | 0.4819 | 0.004 |
| MacLA | ovarytumor | ITGB1 | CD47 | 0.4657 | 0.4931 | 0.4794 | 0.004 |
| MacLA | ovarytumor | ITGB1 | RACK1 | 0.4657 | 2.9543 | 1.71 | 0.004 |

| | | | | | | | |
|---|---|---|---|---|---|---|---|
| breasttumor | MacLA | A2M | LRP1 | 0.3129 | 0.3427 | 0.3278 | 0.000999 |
| breasttumor | MacLA | ANXA1 | C5AR1 | 0.4092 | 0.4636 | 0.4364 | 0.000999 |
| breasttumor | MacLA | ANXA1 | CCR1 | 0.4092 | 0.3657 | 0.3874 | 0.000999 |
| breasttumor | MacLA | ANXA1 | CD4 | 0.4092 | 0.3763 | 0.3927 | 0.000999 |
| breasttumor | MacLA | ANXA1 | CD44 | 0.4092 | 1.4054 | 0.9073 | 0.000999 |
| breasttumor | MacLA | ANXA1 | CXCR4 | 0.4092 | 0.64 | 0.5246 | 0.000999 |
| breasttumor | MacLA | ANXA1 | FPR3 | 0.4092 | 0.4512 | 0.4302 | 0.000999 |
| breasttumor | MacLA | APP | C5AR1 | 0.5409 | 0.4636 | 0.5023 | 0.000999 |
| breasttumor | MacLA | APP | CCR1 | 0.5409 | 0.3657 | 0.4533 | 0.000999 |
| breasttumor | MacLA | APP | CD36 | 0.5409 | 0.4466 | 0.4938 | 0.000999 |
| breasttumor | MacLA | APP | CXCR4 | 0.5409 | 0.64 | 0.5905 | 0.000999 |
| breasttumor | MacLA | APP | LRP1 | 0.5409 | 0.3427 | 0.4418 | 0.000999 |
| breasttumor | MacLA | ARF1 | CD4 | 1.6219 | 0.3763 | 0.9991 | 0.000999 |
| breasttumor | MacLA | ARF1 | CD44 | 1.6219 | 1.4054 | 1.5136 | 0.000999 |
| breasttumor | MacLA | CALR | CD44 | 1.1835 | 1.4054 | 1.2945 | 0.000999 |
| breasttumor | MacLA | CALR | LRP1 | 1.1835 | 0.3427 | 0.7631 | 0.000999 |
| breasttumor | MacLA | CD99 | CD99 | 0.3871 | 0.8115 | 0.5993 | 0.000999 |
| breasttumor | MacLA | COL2A1 | CD44 | 0.3935 | 1.4054 | 0.8994 | 0.000999 |
| breasttumor | MacLA | GSTP1 | CD44 | 2.5435 | 1.4054 | 1.9745 | 0.000999 |
| breasttumor | MacLA | HEBP1 | C5AR1 | 0.4062 | 0.4636 | 0.4349 | 0.000999 |
| breasttumor | MacLA | HEBP1 | CCR1 | 0.4062 | 0.3657 | 0.3859 | 0.000999 |
| breasttumor | MacLA | HEBP1 | CXCR4 | 0.4062 | 0.64 | 0.5231 | 0.000999 |
| breasttumor | MacLA | HEBP1 | FPR3 | 0.4062 | 0.4512 | 0.4287 | 0.000999 |
| breasttumor | MacLA | HMGB1 | LY96 | 1.1815 | 0.8103 | 0.9959 | 0.000999 |
| breasttumor | MacLA | HMGB2 | CD44 | 0.3139 | 1.4054 | 0.8596 | 0.000999 |
| breasttumor | MacLA | HMGB2 | DAB2 | 0.3139 | 0.9194 | 0.6166 | 0.000999 |
| breasttumor | MacLA | HSP90B1 | LRP1 | 0.7786 | 0.3427 | 0.5606 | 0.000999 |
| breasttumor | MacLA | ITGB1 | CD36 | 0.2939 | 0.4466 | 0.3703 | 0.000999 |
| breasttumor | MacLA | ITGB1 | CD47 | 0.2939 | 0.339 | 0.3165 | 0.000999 |
| breasttumor | MacLA | ITGB1 | DAB2 | 0.2939 | 0.9194 | 0.6066 | 0.000999 |

| | | | | | | | |
|---|---|---|---|---|---|---|---|
| breasttumor | MacLA | ITGB1 | ENG | 0.2939 | 0.3236 | 0.3087 | 0.000999 |
| breasttumor | MacLA | ITGB1 | ITGAX | 0.2939 | 0.4922 | 0.393 | 0.000999 |
| breasttumor | MacLA | ITGB1 | PLAUR | 0.2939 | 1.1605 | 0.7272 | 0.000999 |
| breasttumor | MacLA | LGALS3 | CLEC7A | 0.8391 | 0.3231 | 0.5811 | 0.000999 |
| breasttumor | MacLA | LGALS3 | FCGR2A | 0.8391 | 0.3966 | 0.6178 | 0.000999 |
| breasttumor | MacLA | LGALS3 | FCGR2B | 0.8391 | 0.3549 | 0.597 | 0.000999 |
| breasttumor | MacLA | LGALS3 | TFRC | 0.8391 | 0.6885 | 0.7638 | 0.000999 |
| breasttumor | MacLA | LRPAP1 | LRP1 | 0.2756 | 0.3427 | 0.3091 | 0.000999 |
| breasttumor | MacLA | MDK | LRP1 | 0.592 | 0.3427 | 0.4673 | 0.000999 |
| breasttumor | MacLA | MIF | CD74 | 3.2716 | 3.1561 | 3.2138 | 0.000999 |
| breasttumor | MacLA | NPW | C5AR1 | 0.5478 | 0.4636 | 0.5057 | 0.000999 |
| breasttumor | MacLA | NPW | CCR1 | 0.5478 | 0.3657 | 0.4568 | 0.000999 |
| breasttumor | MacLA | NPW | CXCR4 | 0.5478 | 0.64 | 0.5939 | 0.000999 |
| breasttumor | MacLA | RPS19 | C5AR1 | 3.691 | 0.4636 | 2.0773 | 0.000999 |
| breasttumor | MacLA | RPS19 | CD4 | 3.691 | 0.3763 | 2.0336 | 0.000999 |
| breasttumor | MacLA | RTN4 | CD44 | 0.3318 | 1.4054 | 0.8686 | 0.000999 |
| breasttumor | MacLA | RTN4 | CD99 | 0.3318 | 0.8115 | 0.5717 | 0.000999 |
| breasttumor | MacLA | SERPINE2 | LRP1 | 0.6001 | 0.3427 | 0.4714 | 0.000999 |
| breasttumor | MacLA | TF | CD4 | 0.2688 | 0.3763 | 0.3225 | 0.000999 |
| breasttumor | MacLA | TF | TFRC | 0.2688 | 0.6885 | 0.4786 | 0.000999 |
| breasttumor | MacLA | TIMP1 | LRP1 | 0.7045 | 0.3427 | 0.5236 | 0.000999 |
| breasttumor | MacLA | COL2A1 | ITGB1 | 0.3935 | 0.4657 | 0.4296 | 0.00699 |
| breasttumor | MacLA | HSP90B1 | ITGB1 | 0.7786 | 0.4657 | 0.6221 | 0.00999 |
| colorectaltumor | MacLA | ANXA1 | CD44 | 0.5916 | 1.4054 | 0.9985 | 0.000999 |
| colorectaltumor | MacLA | APP | C5AR1 | 0.4966 | 0.4636 | 0.4801 | 0.000999 |
| colorectaltumor | MacLA | APP | CCR1 | 0.4966 | 0.3657 | 0.4312 | 0.000999 |
| colorectaltumor | MacLA | APP | CD36 | 0.4966 | 0.4466 | 0.4716 | 0.000999 |
| colorectaltumor | MacLA | APP | CXCR4 | 0.4966 | 0.64 | 0.5683 | 0.000999 |
| colorectaltumor | MacLA | APP | LRP1 | 0.4966 | 0.3427 | 0.4196 | 0.000999 |
| colorectaltumor | MacLA | ARF1 | CD4 | 0.6489 | 0.3763 | 0.5126 | 0.000999 |

| | | | | | | | |
|---|---|---|---|---|---|---|---|
| colorectaltumor | MacLA | ARF1 | CD44 | 0.6489 | 1.4054 | 1.0271 | 0.000999 |
| colorectaltumor | MacLA | CALR | CD44 | 0.4934 | 1.4054 | 0.9494 | 0.000999 |
| colorectaltumor | MacLA | CD99 | CD99 | 0.3443 | 0.8115 | 0.5779 | 0.000999 |
| colorectaltumor | MacLA | COL1A1 | CD36 | 0.4875 | 0.4466 | 0.4671 | 0.000999 |
| colorectaltumor | MacLA | COL1A1 | CD44 | 0.4875 | 1.4054 | 0.9465 | 0.000999 |
| colorectaltumor | MacLA | GSTP1 | CD44 | 2.8039 | 1.4054 | 2.1046 | 0.000999 |
| colorectaltumor | MacLA | HMGB1 | LY96 | 2.155 | 0.8103 | 1.4826 | 0.000999 |
| colorectaltumor | MacLA | HMGB2 | CD44 | 0.5347 | 1.4054 | 0.9701 | 0.000999 |
| colorectaltumor | MacLA | HMGB2 | DAB2 | 0.5347 | 0.9194 | 0.7271 | 0.000999 |
| colorectaltumor | MacLA | ITGB1 | CD36 | 0.4214 | 0.4466 | 0.434 | 0.000999 |
| colorectaltumor | MacLA | ITGB1 | DAB2 | 0.4214 | 0.9194 | 0.6704 | 0.000999 |
| colorectaltumor | MacLA | ITGB1 | ENG | 0.4214 | 0.3236 | 0.3725 | 0.000999 |
| colorectaltumor | MacLA | ITGB1 | ITGAX | 0.4214 | 0.4922 | 0.4568 | 0.000999 |
| colorectaltumor | MacLA | ITGB1 | PLAUR | 0.4214 | 1.1605 | 0.7909 | 0.000999 |
| colorectaltumor | MacLA | MDK | LRP1 | 0.7548 | 0.3427 | 0.5488 | 0.000999 |
| colorectaltumor | MacLA | MIF | CD74 | 1.6741 | 3.1561 | 2.4151 | 0.000999 |
| colorectaltumor | MacLA | RPS19 | C5AR1 | 4.0956 | 0.4636 | 2.2796 | 0.000999 |
| colorectaltumor | MacLA | RPS19 | CD4 | 4.0956 | 0.3763 | 2.2359 | 0.000999 |
| colorectaltumor | MacLA | RTN4 | CD44 | 0.4148 | 1.4054 | 0.9101 | 0.000999 |
| colorectaltumor | MacLA | HSP90B1 | LRP1 | 0.5887 | 0.3427 | 0.4657 | 0.002 |
| colorectaltumor | MacLA | LGALS3 | TFRC | 1.5563 | 0.6885 | 1.1224 | 0.004 |
| livertumor | MacLA | AGRN | LRP1 | 0.3492 | 0.3427 | 0.346 | 0.000999 |
| livertumor | MacLA | AGT | ATP6AP2 | 0.4701 | 0.9619 | 0.716 | 0.000999 |
| livertumor | MacLA | AGT | C5AR1 | 0.4701 | 0.4636 | 0.4668 | 0.000999 |
| livertumor | MacLA | AGT | CCR1 | 0.4701 | 0.3657 | 0.4179 | 0.000999 |
| livertumor | MacLA | AGT | CXCR4 | 0.4701 | 0.64 | 0.555 | 0.000999 |
| livertumor | MacLA | AGT | PTAFR | 0.4701 | 0.2667 | 0.3684 | 0.000999 |
| livertumor | MacLA | APOA1 | ABCA1 | 0.2888 | 0.3336 | 0.3112 | 0.000999 |
| livertumor | MacLA | APOA1 | CD36 | 0.2888 | 0.4466 | 0.3677 | 0.000999 |
| livertumor | MacLA | APOA1 | LRP1 | 0.2888 | 0.3427 | 0.3157 | 0.000999 |

| | | | | | | | |
|---|---|---|---|---|---|---|---|
| livertumor | MacLA | APOA1 | MSR1 | 0.2888 | 0.6064 | 0.4476 | 0.000999 |
| livertumor | MacLA | APP | C5AR1 | 0.5626 | 0.4636 | 0.5131 | 0.000999 |
| livertumor | MacLA | APP | CCR1 | 0.5626 | 0.3657 | 0.4641 | 0.000999 |
| livertumor | MacLA | APP | CD36 | 0.5626 | 0.4466 | 0.5046 | 0.000999 |
| livertumor | MacLA | APP | CXCR4 | 0.5626 | 0.64 | 0.6013 | 0.000999 |
| livertumor | MacLA | APP | LRP1 | 0.5626 | 0.3427 | 0.4526 | 0.000999 |
| livertumor | MacLA | ARF1 | CD4 | 0.6936 | 0.3763 | 0.5349 | 0.000999 |
| livertumor | MacLA | ARF1 | CD44 | 0.6936 | 1.4054 | 1.0495 | 0.000999 |
| livertumor | MacLA | C3 | C5AR1 | 0.6584 | 0.4636 | 0.561 | 0.000999 |
| livertumor | MacLA | C3 | CCR1 | 0.6584 | 0.3657 | 0.5121 | 0.000999 |
| livertumor | MacLA | C3 | CXCR4 | 0.6584 | 0.64 | 0.6492 | 0.000999 |
| livertumor | MacLA | C3 | ITGAM | 0.6584 | 0.3326 | 0.4955 | 0.000999 |
| livertumor | MacLA | C3 | ITGAX | 0.6584 | 0.4922 | 0.5753 | 0.000999 |
| livertumor | MacLA | C3 | ITGB2 | 0.6584 | 1.2219 | 0.9402 | 0.000999 |
| livertumor | MacLA | C3 | LRP1 | 0.6584 | 0.3427 | 0.5006 | 0.000999 |
| livertumor | MacLA | CALR | CD44 | 1.0416 | 1.4054 | 1.2235 | 0.000999 |
| livertumor | MacLA | CALR | LRP1 | 1.0416 | 0.3427 | 0.6922 | 0.000999 |
| livertumor | MacLA | CCL20 | C5AR1 | 1.0877 | 0.4636 | 0.7756 | 0.000999 |
| livertumor | MacLA | CCL20 | CCR1 | 1.0877 | 0.3657 | 0.7267 | 0.000999 |
| livertumor | MacLA | CCL20 | CXCR4 | 1.0877 | 0.64 | 0.8638 | 0.000999 |
| livertumor | MacLA | CFH | ITGAM | 0.3373 | 0.3326 | 0.335 | 0.000999 |
| livertumor | MacLA | CXCL2 | C5AR1 | 0.3364 | 0.4636 | 0.4 | 0.000999 |
| livertumor | MacLA | CXCL2 | CCR1 | 0.3364 | 0.3657 | 0.351 | 0.000999 |
| livertumor | MacLA | CXCL2 | CXCR4 | 0.3364 | 0.64 | 0.4882 | 0.000999 |
| livertumor | MacLA | CXCL3 | C5AR1 | 0.2543 | 0.4636 | 0.3589 | 0.000999 |
| livertumor | MacLA | CXCL3 | CCR1 | 0.2543 | 0.3657 | 0.31 | 0.000999 |
| livertumor | MacLA | CXCL3 | CXCR4 | 0.2543 | 0.64 | 0.4471 | 0.000999 |
| livertumor | MacLA | DSG2 | CD44 | 0.7479 | 1.4054 | 1.0766 | 0.000999 |
| livertumor | MacLA | DSG2 | CXCR4 | 0.7479 | 0.64 | 0.6939 | 0.000999 |
| livertumor | MacLA | DSG2 | DAB2 | 0.7479 | 0.9194 | 0.8336 | 0.000999 |

| | | | | | | | |
|---|---|---|---|---|---|---|---|
| livertumor | MacLA | FGA | ITGAX | 0.2688 | 0.4922 | 0.3805 | 0.000999 |
| livertumor | MacLA | FGA | ITGB2 | 0.2688 | 1.2219 | 0.7453 | 0.000999 |
| livertumor | MacLA | FN1 | SDC2 | 0.3125 | 0.8667 | 0.5896 | 0.000999 |
| livertumor | MacLA | GSTP1 | CD44 | 1.7219 | 1.4054 | 1.5636 | 0.000999 |
| livertumor | MacLA | HMGB1 | LY96 | 1.4361 | 0.8103 | 1.1232 | 0.000999 |
| livertumor | MacLA | HSP90B1 | LRP1 | 0.8951 | 0.3427 | 0.6189 | 0.000999 |
| livertumor | MacLA | HSPG2 | LRP1 | 0.2746 | 0.3427 | 0.3087 | 0.000999 |
| livertumor | MacLA | ITGB1 | CD36 | 1.6044 | 0.4466 | 1.0255 | 0.000999 |
| livertumor | MacLA | ITGB1 | DAB2 | 1.6044 | 0.9194 | 1.2619 | 0.000999 |
| livertumor | MacLA | ITGB1 | ENG | 1.6044 | 0.3236 | 0.964 | 0.000999 |
| livertumor | MacLA | ITGB1 | ITGAX | 1.6044 | 0.4922 | 1.0483 | 0.000999 |
| livertumor | MacLA | ITGB1 | PLAUR | 1.6044 | 1.1605 | 1.3825 | 0.000999 |
| livertumor | MacLA | LRPAP1 | LRP1 | 0.273 | 0.3427 | 0.3078 | 0.000999 |
| livertumor | MacLA | MDK | LRP1 | 1.6101 | 0.3427 | 0.9764 | 0.000999 |
| livertumor | MacLA | MIF | CD74 | 1.7638 | 3.1561 | 2.4599 | 0.000999 |
| livertumor | MacLA | NTS | PTAFR | 0.3425 | 0.2667 | 0.3046 | 0.000999 |
| livertumor | MacLA | RPS19 | C5AR1 | 4.0772 | 0.4636 | 2.2704 | 0.000999 |
| livertumor | MacLA | RPS19 | CD4 | 4.0772 | 0.3763 | 2.2267 | 0.000999 |
| livertumor | MacLA | RTN4 | CD44 | 1.0046 | 1.4054 | 1.205 | 0.000999 |
| livertumor | MacLA | SEMA3C | NRP1 | 0.2576 | 0.2611 | 0.2593 | 0.000999 |
| livertumor | MacLA | SEMA3C | NRP2 | 0.2576 | 0.2826 | 0.2701 | 0.000999 |
| livertumor | MacLA | SERPINA1 | LRP1 | 1.604 | 0.3427 | 0.9733 | 0.000999 |
| livertumor | MacLA | SERPINA1 | SDC2 | 1.604 | 0.8667 | 1.2354 | 0.000999 |
| livertumor | MacLA | SERPING1 | LRP1 | 0.3498 | 0.3427 | 0.3462 | 0.000999 |
| livertumor | MacLA | TF | CD4 | 0.523 | 0.3763 | 0.4496 | 0.000999 |
| livertumor | MacLA | TF | SDC2 | 0.523 | 0.8667 | 0.6949 | 0.000999 |
| livertumor | MacLA | TF | TFRC | 0.523 | 0.6885 | 0.6057 | 0.000999 |
| livertumor | MacLA | TFPI | LRP1 | 0.5043 | 0.3427 | 0.4235 | 0.000999 |
| livertumor | MacLA | VEGFA | NRP1 | 0.5485 | 0.2611 | 0.4048 | 0.000999 |
| livertumor | MacLA | VEGFA | NRP2 | 0.5485 | 0.2826 | 0.4155 | 0.000999 |

| | | | | | | | |
|---|---|---|---|---|---|---|---|
| livertumor | MacLA | VTN | PLAUR | 0.7399 | 1.1605 | 0.9502 | 0.000999 |
| livertumor | MacLA | VTN | SDC2 | 0.7399 | 0.8667 | 0.8033 | 0.000999 |
| livertumor | MacLA | ANXA1 | CXCR4 | 0.3041 | 0.64 | 0.4721 | 0.002 |
| livertumor | MacLA | FN1 | CD44 | 0.3125 | 1.4054 | 0.8589 | 0.002 |
| livertumor | MacLA | TIMP1 | LRP1 | 1.0257 | 0.3427 | 0.6842 | 0.002 |
| livertumor | MacLA | CD99 | CD99 | 0.5906 | 0.8115 | 0.7011 | 0.005 |
| livertumor | MacLA | ITGB1 | CD47 | 1.6044 | 0.339 | 0.9717 | 0.00699 |
| lungtumor | MacLA | ANXA1 | C5AR1 | 1.4026 | 0.4636 | 0.9331 | 0.000999 |
| lungtumor | MacLA | ANXA1 | CCR1 | 1.4026 | 0.3657 | 0.8842 | 0.000999 |
| lungtumor | MacLA | ANXA1 | CD4 | 1.4026 | 0.3763 | 0.8895 | 0.000999 |
| lungtumor | MacLA | ANXA1 | CD44 | 1.4026 | 1.4054 | 1.404 | 0.000999 |
| lungtumor | MacLA | ANXA1 | CXCR4 | 1.4026 | 0.64 | 1.0213 | 0.000999 |
| lungtumor | MacLA | ANXA1 | FPR3 | 1.4026 | 0.4512 | 0.9269 | 0.000999 |
| lungtumor | MacLA | ARF1 | CD4 | 0.8791 | 0.3763 | 0.6277 | 0.000999 |
| lungtumor | MacLA | ARF1 | CD44 | 0.8791 | 1.4054 | 1.1423 | 0.000999 |
| lungtumor | MacLA | CALR | CD44 | 0.7795 | 1.4054 | 1.0925 | 0.000999 |
| lungtumor | MacLA | CALR | LRP1 | 0.7795 | 0.3427 | 0.5611 | 0.000999 |
| lungtumor | MacLA | CCL20 | C5AR1 | 0.2885 | 0.4636 | 0.3761 | 0.000999 |
| lungtumor | MacLA | CCL20 | CCR1 | 0.2885 | 0.3657 | 0.3271 | 0.000999 |
| lungtumor | MacLA | CCL20 | CXCR4 | 0.2885 | 0.64 | 0.4642 | 0.000999 |
| lungtumor | MacLA | CXCL10 | C5AR1 | 0.3417 | 0.4636 | 0.4026 | 0.000999 |
| lungtumor | MacLA | CXCL10 | CCR1 | 0.3417 | 0.3657 | 0.3537 | 0.000999 |
| lungtumor | MacLA | CXCL10 | CXCR4 | 0.3417 | 0.64 | 0.4908 | 0.000999 |
| lungtumor | MacLA | FGA | ITGAX | 0.4445 | 0.4922 | 0.4684 | 0.000999 |
| lungtumor | MacLA | FGA | ITGB1 | 0.4445 | 0.4657 | 0.4551 | 0.000999 |
| lungtumor | MacLA | FGA | ITGB2 | 0.4445 | 1.2219 | 0.8332 | 0.000999 |
| lungtumor | MacLA | GSTP1 | CD44 | 2.4093 | 1.4054 | 1.9073 | 0.000999 |
| lungtumor | MacLA | HLA-DRA | CD4 | 0.9609 | 0.3763 | 0.6686 | 0.000999 |
| lungtumor | MacLA | HLA-DRA | PTPRC | 0.9609 | 0.3665 | 0.6637 | 0.000999 |
| lungtumor | MacLA | HMGB1 | LY96 | 1.1943 | 0.8103 | 1.0023 | 0.000999 |

| | | | | | | | |
|---|---|---|---|---|---|---|---|
| lungtumor | MacLA | HMGB2 | CD44 | 0.5593 | 1.4054 | 0.9824 | 0.000999 |
| lungtumor | MacLA | HMGB2 | DAB2 | 0.5593 | 0.9194 | 0.7394 | 0.000999 |
| lungtumor | MacLA | HSP90B1 | LRP1 | 0.6141 | 0.3427 | 0.4784 | 0.000999 |
| lungtumor | MacLA | LGALS3 | CLEC7A | 0.9915 | 0.3231 | 0.6573 | 0.000999 |
| lungtumor | MacLA | LGALS3 | FCGR2A | 0.9915 | 0.3966 | 0.694 | 0.000999 |
| lungtumor | MacLA | LGALS3 | FCGR2B | 0.9915 | 0.3549 | 0.6732 | 0.000999 |
| lungtumor | MacLA | LGALS3 | TFRC | 0.9915 | 0.6885 | 0.84 | 0.000999 |
| lungtumor | MacLA | MDK | LRP1 | 1.2896 | 0.3427 | 0.8161 | 0.000999 |
| lungtumor | MacLA | MIF | CD74 | 1.4585 | 3.1561 | 2.3073 | 0.000999 |
| lungtumor | MacLA | PSAP | LRP1 | 0.5457 | 0.3427 | 0.4442 | 0.000999 |
| lungtumor | MacLA | RPS19 | C5AR1 | 3.5104 | 0.4636 | 1.987 | 0.000999 |
| lungtumor | MacLA | RPS19 | CD4 | 3.5104 | 0.3763 | 1.9433 | 0.000999 |
| lungtumor | MacLA | RTN4 | CD44 | 0.4013 | 1.4054 | 0.9033 | 0.000999 |
| lungtumor | MacLA | SAA1 | C5AR1 | 0.2539 | 0.4636 | 0.3587 | 0.000999 |
| lungtumor | MacLA | SAA1 | CCR1 | 0.2539 | 0.3657 | 0.3098 | 0.000999 |
| lungtumor | MacLA | SAA1 | CXCR4 | 0.2539 | 0.64 | 0.4469 | 0.000999 |
| lungtumor | MacLA | SAA1 | PTAFR | 0.2539 | 0.2667 | 0.2603 | 0.000999 |
| lungtumor | MacLA | SERPINA1 | LRP1 | 0.3624 | 0.3427 | 0.3525 | 0.000999 |
| lungtumor | MacLA | SERPINA1 | SDC2 | 0.3624 | 0.8667 | 0.6145 | 0.000999 |
| lungtumor | MacLA | SPP1 | CD44 | 1.4874 | 1.4054 | 1.4464 | 0.000999 |
| lungtumor | MacLA | TFPI | LRP1 | 0.3615 | 0.3427 | 0.3521 | 0.000999 |
| lungtumor | MacLA | TIMP1 | LRP1 | 0.6195 | 0.3427 | 0.4811 | 0.000999 |
| lungtumor | MacLA | HSP90B1 | ITGB1 | 0.6141 | 0.4657 | 0.5399 | 0.002 |
| lungtumor | MacLA | LGALS3 | ITGB1 | 0.9915 | 0.4657 | 0.7286 | 0.00799 |
| melanomatumo | MacLA | ARF1 | CD4 | 1.8545 | 0.3763 | 1.1154 | 0.000999 |
| melanomatumo | MacLA | ARF1 | CD44 | 1.8545 | 1.4054 | 1.6299 | 0.000999 |
| melanomatumo | MacLA | GSTP1 | CD44 | 2.3341 | 1.4054 | 1.8697 | 0.000999 |
| melanomatumo | MacLA | HAS2 | CD44 | 0.3078 | 1.4054 | 0.8566 | 0.000999 |
| melanomatumo | MacLA | HEBP1 | CXCR4 | 0.3689 | 0.64 | 0.5044 | 0.000999 |
| melanomatumo | MacLA | HMGB1 | LY96 | 1.6757 | 0.8103 | 1.243 | 0.000999 |

| | | | | | | | |
|---|---|---|---|---|---|---|---|
| melanomatumo | MacLA | MIF | CD74 | 2.6939 | 3.1561 | 2.925 | 0.000999 |
| melanomatumo | MacLA | SERPINE2 | LRP1 | 0.9323 | 0.3427 | 0.6375 | 0.000999 |
| melanomatumo | MacLA | HEBP1 | C5AR1 | 0.3689 | 0.4636 | 0.4162 | 0.002 |
| melanomatumo | MacLA | HEBP1 | CCR1 | 0.3689 | 0.3657 | 0.3673 | 0.002 |
| melanomatumo | MacLA | HEBP1 | FPR3 | 0.3689 | 0.4512 | 0.41 | 0.002 |
| melanomatumo | MacLA | MDK | LRP1 | 0.3759 | 0.3427 | 0.3593 | 0.002 |
| melanomatumo | MacLA | HMGB2 | DAB2 | 0.2735 | 0.9194 | 0.5965 | 0.005 |
| ovarytumor | MacLA | ANXA1 | C5AR1 | 1.0314 | 0.4636 | 0.7475 | 0.000999 |
| ovarytumor | MacLA | ANXA1 | CCR1 | 1.0314 | 0.3657 | 0.6985 | 0.000999 |
| ovarytumor | MacLA | ANXA1 | CD4 | 1.0314 | 0.3763 | 0.7038 | 0.000999 |
| ovarytumor | MacLA | ANXA1 | CD44 | 1.0314 | 1.4054 | 1.2184 | 0.000999 |
| ovarytumor | MacLA | ANXA1 | CXCR4 | 1.0314 | 0.64 | 0.8357 | 0.000999 |
| ovarytumor | MacLA | ANXA1 | FPR3 | 1.0314 | 0.4512 | 0.7413 | 0.000999 |
| ovarytumor | MacLA | APOE | LRP1 | 0.837 | 0.3427 | 0.5899 | 0.000999 |
| ovarytumor | MacLA | APOE | SDC2 | 0.837 | 0.8667 | 0.8519 | 0.000999 |
| ovarytumor | MacLA | APP | C5AR1 | 0.2907 | 0.4636 | 0.3771 | 0.000999 |
| ovarytumor | MacLA | APP | CCR1 | 0.2907 | 0.3657 | 0.3282 | 0.000999 |
| ovarytumor | MacLA | APP | CD36 | 0.2907 | 0.4466 | 0.3687 | 0.000999 |
| ovarytumor | MacLA | APP | CXCR4 | 0.2907 | 0.64 | 0.4653 | 0.000999 |
| ovarytumor | MacLA | APP | LRP1 | 0.2907 | 0.3427 | 0.3167 | 0.000999 |
| ovarytumor | MacLA | ARF1 | CD4 | 0.7364 | 0.3763 | 0.5563 | 0.000999 |
| ovarytumor | MacLA | ARF1 | CD44 | 0.7364 | 1.4054 | 1.0709 | 0.000999 |
| ovarytumor | MacLA | CALR | CD44 | 0.7423 | 1.4054 | 1.0738 | 0.000999 |
| ovarytumor | MacLA | CALR | LRP1 | 0.7423 | 0.3427 | 0.5425 | 0.000999 |
| ovarytumor | MacLA | CCL2 | CCR1 | 0.3526 | 0.3657 | 0.3592 | 0.000999 |
| ovarytumor | MacLA | COL1A1 | CD36 | 0.3504 | 0.4466 | 0.3985 | 0.000999 |
| ovarytumor | MacLA | COL1A1 | CD44 | 0.3504 | 1.4054 | 0.8779 | 0.000999 |
| ovarytumor | MacLA | GSTP1 | CD44 | 2.6185 | 1.4054 | 2.0119 | 0.000999 |
| ovarytumor | MacLA | HLA-DMA | CD4 | 0.2537 | 0.3763 | 0.315 | 0.000999 |
| ovarytumor | MacLA | HLA-DRA | CD4 | 1.2369 | 0.3763 | 0.8066 | 0.000999 |

| | | | | | | | |
|---|---|---|---|---|---|---|---|
| ovarytumor | MacLA | HLA-DRA | PTPRC | 1.2369 | 0.3665 | 0.8017 | 0.000999 |
| ovarytumor | MacLA | HMGB1 | LY96 | 1.7272 | 0.8103 | 1.2687 | 0.000999 |
| ovarytumor | MacLA | HMGB2 | CD44 | 0.4974 | 1.4054 | 0.9514 | 0.000999 |
| ovarytumor | MacLA | HMGB2 | DAB2 | 0.4974 | 0.9194 | 0.7084 | 0.000999 |
| ovarytumor | MacLA | HSP90B1 | LRP1 | 0.6365 | 0.3427 | 0.4896 | 0.000999 |
| ovarytumor | MacLA | ITGB1 | CD36 | 0.2783 | 0.4466 | 0.3625 | 0.000999 |
| ovarytumor | MacLA | ITGB1 | DAB2 | 0.2783 | 0.9194 | 0.5988 | 0.000999 |
| ovarytumor | MacLA | ITGB1 | ENG | 0.2783 | 0.3236 | 0.3009 | 0.000999 |
| ovarytumor | MacLA | ITGB1 | ITGAX | 0.2783 | 0.4922 | 0.3852 | 0.000999 |
| ovarytumor | MacLA | ITGB1 | PLAUR | 0.2783 | 1.1605 | 0.7194 | 0.000999 |
| ovarytumor | MacLA | LGALS3 | CLEC7A | 0.5097 | 0.3231 | 0.4164 | 0.000999 |
| ovarytumor | MacLA | LGALS3 | FCGR2A | 0.5097 | 0.3966 | 0.4531 | 0.000999 |
| ovarytumor | MacLA | LGALS3 | FCGR2B | 0.5097 | 0.3549 | 0.4323 | 0.000999 |
| ovarytumor | MacLA | LGALS3 | TFRC | 0.5097 | 0.6885 | 0.5991 | 0.000999 |
| ovarytumor | MacLA | LRPAP1 | LRP1 | 0.3826 | 0.3427 | 0.3626 | 0.000999 |
| ovarytumor | MacLA | MDK | LRP1 | 1.5321 | 0.3427 | 0.9374 | 0.000999 |
| ovarytumor | MacLA | MIF | CD74 | 0.8565 | 3.1561 | 2.0063 | 0.000999 |
| ovarytumor | MacLA | PSAP | LRP1 | 0.6574 | 0.3427 | 0.5001 | 0.000999 |
| ovarytumor | MacLA | RPS19 | C5AR1 | 3.9325 | 0.4636 | 2.198 | 0.000999 |
| ovarytumor | MacLA | RPS19 | CD4 | 3.9325 | 0.3763 | 2.1544 | 0.000999 |
| ovarytumor | MacLA | RTN4 | CD44 | 0.4992 | 1.4054 | 0.9523 | 0.000999 |
| ovarytumor | MacLA | RTN4 | CD99 | 0.4992 | 0.8115 | 0.6554 | 0.000999 |
| ovarytumor | MacLA | SERPING1 | LRP1 | 0.3678 | 0.3427 | 0.3553 | 0.000999 |
| ovarytumor | MacLA | SPP1 | CD44 | 0.3946 | 1.4054 | 0.9 | 0.000999 |
| ovarytumor | MacLA | TIMP1 | LRP1 | 0.7779 | 0.3427 | 0.5603 | 0.000999 |
| ovarytumor | MacLA | COL1A1 | ITGB1 | 0.3504 | 0.4657 | 0.408 | 0.004 |
| ovarytumor | MacLA | HSP90B1 | ITGB1 | 0.6365 | 0.4657 | 0.5511 | 0.005 |

| Supplementary Table 8: Communication results between macrophages and lymphocytes subpopulations, and malignant cells | | | | | | | |
|---|---|---|---|---|---|---|---|
| **MacRec vs. lymphocytes** | | | | | | | |
| celltype_01 | celltype_02 | ligand | receptor | ligand_mean_expression | receptor_mean_expression | ligand_receptor_mean_expression | pvalue |
| MacRec | NKT | A2M | TNFRSF14 | 0.2825 | 0.2848 | 0.2836 | 0.000999 |
| MacRec | NKT | ADM | PTGER2 | 0.3154 | 0.3627 | 0.3391 | 0.000999 |
| MacRec | NKT | AIMP1 | RACK1 | 0.2829 | 2.4327 | 1.3578 | 0.000999 |
| MacRec | NKT | ALCAM | CD6 | 0.2586 | 0.4881 | 0.3733 | 0.000999 |
| MacRec | NKT | ANXA1 | CXCR4 | 1.2584 | 2.0373 | 1.6479 | 0.000999 |
| MacRec | NKT | APOE | SORL1 | 1.1144 | 0.4137 | 0.764 | 0.000999 |
| MacRec | NKT | APP | CXCR4 | 0.2502 | 2.0373 | 1.1438 | 0.000999 |
| MacRec | NKT | APP | F2R | 0.2502 | 0.2978 | 0.274 | 0.000999 |
| MacRec | NKT | APP | SORL1 | 0.2502 | 0.4137 | 0.3319 | 0.000999 |
| MacRec | NKT | APP | TNFRSF14 | 0.2502 | 0.2848 | 0.2675 | 0.000999 |
| MacRec | NKT | C3 | CXCR4 | 0.4838 | 2.0373 | 1.2605 | 0.000999 |
| MacRec | NKT | C3 | ITGB2 | 0.4838 | 1.6481 | 1.0659 | 0.000999 |
| MacRec | NKT | CALR | IL7R | 1.0088 | 0.5933 | 0.8011 | 0.000999 |
| MacRec | NKT | CCL20 | CXCR4 | 0.3793 | 2.0373 | 1.2083 | 0.000999 |
| MacRec | NKT | CCL20 | S1PR4 | 0.3793 | 0.4435 | 0.4114 | 0.000999 |
| MacRec | NKT | CCL20 | S1PR5 | 0.3793 | 0.508 | 0.4437 | 0.000999 |
| MacRec | NKT | CD58 | CD2 | 0.2951 | 1.1465 | 0.7208 | 0.000999 |
| MacRec | NKT | COL1A1 | CD44 | 0.5365 | 0.6724 | 0.6045 | 0.000999 |
| MacRec | NKT | COL1A1 | ITGB1 | 0.5365 | 0.488 | 0.5122 | 0.000999 |
| MacRec | NKT | CXCL10 | CXCR4 | 0.3391 | 2.0373 | 1.1882 | 0.000999 |
| MacRec | NKT | CXCL10 | S1PR4 | 0.3391 | 0.4435 | 0.3913 | 0.000999 |
| MacRec | NKT | CXCL10 | S1PR5 | 0.3391 | 0.508 | 0.4236 | 0.000999 |
| MacRec | NKT | CXCL16 | CXCR4 | 0.8324 | 2.0373 | 1.4349 | 0.000999 |
| MacRec | NKT | CXCL16 | S1PR4 | 0.8324 | 0.4435 | 0.638 | 0.000999 |
| MacRec | NKT | CXCL16 | S1PR5 | 0.8324 | 0.508 | 0.6702 | 0.000999 |
| MacRec | NKT | CXCL2 | CXCR4 | 0.8239 | 2.0373 | 1.4306 | 0.000999 |
| MacRec | NKT | CXCL2 | S1PR4 | 0.8239 | 0.4435 | 0.6337 | 0.000999 |

| | | | | | | | |
|---|---|---|---|---|---|---|---|
| MacRec | NKT | CXCL2 | S1PR5 | 0.8239 | 0.508 | 0.666 | 0.000999 |
| MacRec | NKT | CXCL3 | CXCR4 | 0.5699 | 2.0373 | 1.3036 | 0.000999 |
| MacRec | NKT | CXCL3 | S1PR4 | 0.5699 | 0.4435 | 0.5067 | 0.000999 |
| MacRec | NKT | CXCL3 | S1PR5 | 0.5699 | 0.508 | 0.539 | 0.000999 |
| MacRec | NKT | FN1 | CD44 | 1.5028 | 0.6724 | 1.0876 | 0.000999 |
| MacRec | NKT | FN1 | ITGA4 | 1.5028 | 0.2946 | 0.8987 | 0.000999 |
| MacRec | NKT | FN1 | ITGB1 | 1.5028 | 0.488 | 0.9954 | 0.000999 |
| MacRec | NKT | FN1 | ITGB7 | 1.5028 | 0.4721 | 0.9874 | 0.000999 |
| MacRec | NKT | FN1 | RPSA | 1.5028 | 2.3639 | 1.9333 | 0.000999 |
| MacRec | NKT | GRN | TNFRSF1B | 1.7446 | 0.4949 | 1.1197 | 0.000999 |
| MacRec | NKT | GSTP1 | CD44 | 1.7225 | 0.6724 | 1.1974 | 0.000999 |
| MacRec | NKT | GSTP1 | IL7R | 1.7225 | 0.5933 | 1.1579 | 0.000999 |
| MacRec | NKT | GSTP1 | ITGA4 | 1.7225 | 0.2946 | 1.0085 | 0.000999 |
| MacRec | NKT | HBEGF | CD44 | 0.7419 | 0.6724 | 0.7071 | 0.000999 |
| MacRec | NKT | HLA-DRA | PTPRC | 4.3438 | 1.8797 | 3.1118 | 0.000999 |
| MacRec | NKT | HMGB1 | ITGA4 | 1.2904 | 0.2946 | 0.7925 | 0.000999 |
| MacRec | NKT | HSP90B1 | ITGB1 | 0.7859 | 0.488 | 0.6369 | 0.000999 |
| MacRec | NKT | HSP90B1 | TNFRSF1B | 0.7859 | 0.4949 | 0.6404 | 0.000999 |
| MacRec | NKT | LGALS3 | CD6 | 0.9368 | 0.4881 | 0.7125 | 0.000999 |
| MacRec | NKT | LGALS3 | ITGB1 | 0.9368 | 0.488 | 0.7124 | 0.000999 |
| MacRec | NKT | LGALS3 | SPN | 0.9368 | 0.6398 | 0.7883 | 0.000999 |
| MacRec | NKT | LRPAP1 | SORL1 | 0.5059 | 0.4137 | 0.4598 | 0.000999 |
| MacRec | NKT | MIF | ITGA4 | 0.8867 | 0.2946 | 0.5906 | 0.000999 |
| MacRec | NKT | MIF | TNFRSF14 | 0.8867 | 0.2848 | 0.5857 | 0.000999 |
| MacRec | NKT | NAMPT | ANXA6 | 1.5176 | 0.844 | 1.1808 | 0.000999 |
| MacRec | NKT | NECTIN2 | TIGIT | 0.2672 | 0.35 | 0.3086 | 0.000999 |
| MacRec | NKT | PECAM1 | CXCR4 | 0.3003 | 2.0373 | 1.1688 | 0.000999 |
| MacRec | NKT | RPS19 | ITGA4 | 3.7755 | 0.2946 | 2.035 | 0.000999 |
| MacRec | NKT | RPS19 | RPSA | 3.7755 | 2.3639 | 3.0697 | 0.000999 |
| MacRec | NKT | RTN4 | CD44 | 1.101 | 0.6724 | 0.8867 | 0.000999 |

| | | | | | | | |
|---|---|---|---|---|---|---|---|
| MacRec | NKT | RTN4 | CD99 | 1.101 | 1.4547 | 1.2778 | 0.000999 |
| MacRec | NKT | SPP1 | CD44 | 1.5024 | 0.6724 | 1.0874 | 0.000999 |
| MacRec | NKT | SPP1 | ITGA4 | 1.5024 | 0.2946 | 0.8985 | 0.000999 |
| MacRec | NKT | SPP1 | ITGB1 | 1.5024 | 0.488 | 0.9952 | 0.000999 |
| MacRec | NKT | THBS1 | CD47 | 0.7038 | 0.5576 | 0.6307 | 0.000999 |
| MacRec | NKT | THBS1 | ITGA4 | 0.7038 | 0.2946 | 0.4992 | 0.000999 |
| MacRec | NKT | THBS1 | ITGB1 | 0.7038 | 0.488 | 0.5959 | 0.000999 |
| MacRec | NKT | TNFSF13B | ITGB1 | 0.8304 | 0.488 | 0.6592 | 0.000999 |
| MacRec | NKT | ANXA1 | F2R | 1.2584 | 0.2978 | 0.7781 | 0.004 |
| MacRec | NKT | ANXA1 | ITGA4 | 1.2584 | 0.2946 | 0.7765 | 0.00799 |
| MacRec | TCD4ex | A2M | TNFRSF14 | 0.2825 | 0.2806 | 0.2816 | 0.000999 |
| MacRec | TCD4ex | ADM | PTGER4 | 0.3154 | 0.4696 | 0.3925 | 0.000999 |
| MacRec | TCD4ex | ALCAM | CD6 | 0.2586 | 0.5338 | 0.3962 | 0.000999 |
| MacRec | TCD4ex | ANXA1 | CCR7 | 1.2584 | 0.4844 | 0.8714 | 0.000999 |
| MacRec | TCD4ex | ANXA1 | CD4 | 1.2584 | 0.5864 | 0.9224 | 0.000999 |
| MacRec | TCD4ex | ANXA1 | CD44 | 1.2584 | 1.0244 | 1.1414 | 0.000999 |
| MacRec | TCD4ex | ANXA1 | CXCR4 | 1.2584 | 2.4201 | 1.8393 | 0.000999 |
| MacRec | TCD4ex | APP | CCR7 | 0.2502 | 0.4844 | 0.3673 | 0.000999 |
| MacRec | TCD4ex | APP | CXCR4 | 0.2502 | 2.4201 | 1.3351 | 0.000999 |
| MacRec | TCD4ex | APP | TNFRSF14 | 0.2502 | 0.2806 | 0.2654 | 0.000999 |
| MacRec | TCD4ex | C3 | CCR7 | 0.4838 | 0.4844 | 0.4841 | 0.000999 |
| MacRec | TCD4ex | C3 | CXCR4 | 0.4838 | 2.4201 | 1.4519 | 0.000999 |
| MacRec | TCD4ex | C3 | ITGB2 | 0.4838 | 0.7871 | 0.6354 | 0.000999 |
| MacRec | TCD4ex | CALR | IL7R | 1.0088 | 1.2353 | 1.122 | 0.000999 |
| MacRec | TCD4ex | CCL20 | CCR7 | 0.3793 | 0.4844 | 0.4318 | 0.000999 |
| MacRec | TCD4ex | CCL20 | CXCR4 | 0.3793 | 2.4201 | 1.3997 | 0.000999 |
| MacRec | TCD4ex | CD58 | CD2 | 0.2951 | 1.5731 | 0.9341 | 0.000999 |
| MacRec | TCD4ex | CD86 | CD28 | 0.5786 | 0.3256 | 0.4521 | 0.000999 |
| MacRec | TCD4ex | CD86 | CTLA4 | 0.5786 | 0.6294 | 0.604 | 0.000999 |
| MacRec | TCD4ex | COL1A1 | CD44 | 0.5365 | 1.0244 | 0.7805 | 0.000999 |

| | | | | | | | |
|---|---|---|---|---|---|---|---|
| MacRec | TCD4ex | COL1A1 | ITGB1 | 0.5365 | 0.4615 | 0.499 | 0.000999 |
| MacRec | TCD4ex | CXCL10 | CCR7 | 0.3391 | 0.4844 | 0.4117 | 0.000999 |
| MacRec | TCD4ex | CXCL10 | CXCR4 | 0.3391 | 2.4201 | 1.3796 | 0.000999 |
| MacRec | TCD4ex | CXCL16 | CCR7 | 0.8324 | 0.4844 | 0.6584 | 0.000999 |
| MacRec | TCD4ex | CXCL16 | CXCR4 | 0.8324 | 2.4201 | 1.6262 | 0.000999 |
| MacRec | TCD4ex | CXCL2 | CCR7 | 0.8239 | 0.4844 | 0.6542 | 0.000999 |
| MacRec | TCD4ex | CXCL2 | CXCR4 | 0.8239 | 2.4201 | 1.622 | 0.000999 |
| MacRec | TCD4ex | CXCL3 | CCR7 | 0.5699 | 0.4844 | 0.5272 | 0.000999 |
| MacRec | TCD4ex | CXCL3 | CXCR4 | 0.5699 | 2.4201 | 1.495 | 0.000999 |
| MacRec | TCD4ex | FN1 | CD44 | 1.5028 | 1.0244 | 1.2636 | 0.000999 |
| MacRec | TCD4ex | FN1 | ITGB1 | 1.5028 | 0.4615 | 0.9821 | 0.000999 |
| MacRec | TCD4ex | FN1 | RPSA | 1.5028 | 2.4895 | 1.9961 | 0.000999 |
| MacRec | TCD4ex | GRN | TNFRSF1B | 1.7446 | 0.383 | 1.0638 | 0.000999 |
| MacRec | TCD4ex | GSTP1 | CD44 | 1.7225 | 1.0244 | 1.3734 | 0.000999 |
| MacRec | TCD4ex | GSTP1 | IL7R | 1.7225 | 1.2353 | 1.4789 | 0.000999 |
| MacRec | TCD4ex | HBEGF | CD44 | 0.7419 | 1.0244 | 0.8831 | 0.000999 |
| MacRec | TCD4ex | HLA-DMA | CD4 | 1.7832 | 0.5864 | 1.1848 | 0.000999 |
| MacRec | TCD4ex | HLA-DRA | CD4 | 4.3438 | 0.5864 | 2.4651 | 0.000999 |
| MacRec | TCD4ex | HLA-DRA | PDCD1 | 4.3438 | 0.457 | 2.4004 | 0.000999 |
| MacRec | TCD4ex | HLA-DRA | PTPRC | 4.3438 | 1.7493 | 3.0466 | 0.000999 |
| MacRec | TCD4ex | HSP90B1 | ITGB1 | 0.7859 | 0.4615 | 0.6237 | 0.000999 |
| MacRec | TCD4ex | HSP90B1 | TNFRSF1B | 0.7859 | 0.383 | 0.5845 | 0.000999 |
| MacRec | TCD4ex | ITGB2 | ITGAL | 1.3879 | 0.2603 | 0.8241 | 0.000999 |
| MacRec | TCD4ex | LGALS3 | CD6 | 0.9368 | 0.5338 | 0.7353 | 0.000999 |
| MacRec | TCD4ex | LGALS3 | ITGB1 | 0.9368 | 0.4615 | 0.6992 | 0.000999 |
| MacRec | TCD4ex | MIF | TNFRSF14 | 0.8867 | 0.2806 | 0.5837 | 0.000999 |
| MacRec | TCD4ex | NAMPT | ANXA6 | 1.5176 | 0.5386 | 1.0281 | 0.000999 |
| MacRec | TCD4ex | NECTIN2 | TIGIT | 0.2672 | 0.853 | 0.5601 | 0.000999 |
| MacRec | TCD4ex | OSM | IL6ST | 0.2991 | 0.7106 | 0.5048 | 0.000999 |
| MacRec | TCD4ex | PECAM1 | CXCR4 | 0.3003 | 2.4201 | 1.3602 | 0.000999 |

| | | | | | | | |
|---|---|---|---|---|---|---|---|
| MacRec | TCD4ex | RPS19 | CD4 | 3.7755 | 0.5864 | 2.181 | 0.000999 |
| MacRec | TCD4ex | RPS19 | RPSA | 3.7755 | 2.4895 | 3.1325 | 0.000999 |
| MacRec | TCD4ex | RTN4 | CD44 | 1.101 | 1.0244 | 1.0627 | 0.000999 |
| MacRec | TCD4ex | RTN4 | CD99 | 1.101 | 0.9165 | 1.0087 | 0.000999 |
| MacRec | TCD4ex | SPP1 | CD44 | 1.5024 | 1.0244 | 1.2634 | 0.000999 |
| MacRec | TCD4ex | SPP1 | ITGB1 | 1.5024 | 0.4615 | 0.9819 | 0.000999 |
| MacRec | TCD4ex | TGFB1 | ITGB1 | 0.317 | 0.4615 | 0.3893 | 0.000999 |
| MacRec | TCD4ex | THBS1 | CD47 | 0.7038 | 0.3456 | 0.5247 | 0.000999 |
| MacRec | TCD4ex | THBS1 | ITGB1 | 0.7038 | 0.4615 | 0.5826 | 0.000999 |
| MacRec | TCD4ex | TNFSF13B | ITGB1 | 0.8304 | 0.4615 | 0.646 | 0.000999 |
| MacRec | TCD4ex | ARF1 | CD4 | 0.7488 | 0.5864 | 0.6676 | 0.003 |
| MacRec | TCD4reg | A2M | TNFRSF14 | 0.2825 | 0.3843 | 0.3334 | 0.000999 |
| MacRec | TCD4reg | AIMP1 | FAS | 0.2829 | 0.2727 | 0.2778 | 0.000999 |
| MacRec | TCD4reg | ALCAM | CD6 | 0.2586 | 0.2651 | 0.2618 | 0.000999 |
| MacRec | TCD4reg | ANXA1 | CCR7 | 1.2584 | 0.3098 | 0.7841 | 0.000999 |
| MacRec | TCD4reg | ANXA1 | CD4 | 1.2584 | 0.429 | 0.8437 | 0.000999 |
| MacRec | TCD4reg | ANXA1 | CD44 | 1.2584 | 1.0896 | 1.174 | 0.000999 |
| MacRec | TCD4reg | ANXA1 | CXCR3 | 1.2584 | 0.3357 | 0.7971 | 0.000999 |
| MacRec | TCD4reg | ANXA1 | CXCR4 | 1.2584 | 2.2962 | 1.7773 | 0.000999 |
| MacRec | TCD4reg | ANXA1 | CXCR6 | 1.2584 | 0.3945 | 0.8265 | 0.000999 |
| MacRec | TCD4reg | ANXA1 | FAS | 1.2584 | 0.2727 | 0.7656 | 0.000999 |
| MacRec | TCD4reg | APP | CCR7 | 0.2502 | 0.3098 | 0.28 | 0.000999 |
| MacRec | TCD4reg | APP | CXCR3 | 0.2502 | 0.3357 | 0.293 | 0.000999 |
| MacRec | TCD4reg | APP | CXCR4 | 0.2502 | 2.2962 | 1.2732 | 0.000999 |
| MacRec | TCD4reg | APP | CXCR6 | 0.2502 | 0.3945 | 0.3224 | 0.000999 |
| MacRec | TCD4reg | APP | TNFRSF14 | 0.2502 | 0.3843 | 0.3173 | 0.000999 |
| MacRec | TCD4reg | C3 | CCR7 | 0.4838 | 0.3098 | 0.3968 | 0.000999 |
| MacRec | TCD4reg | C3 | CXCR3 | 0.4838 | 0.3357 | 0.4097 | 0.000999 |
| MacRec | TCD4reg | C3 | CXCR4 | 0.4838 | 2.2962 | 1.39 | 0.000999 |
| MacRec | TCD4reg | C3 | CXCR6 | 0.4838 | 0.3945 | 0.4391 | 0.000999 |

| | | | | | | | |
|---|---|---|---|---|---|---|---|
| MacRec | TCD4reg | C3 | ITGB2 | 0.4838 | 0.6238 | 0.5538 | 0.000999 |
| MacRec | TCD4reg | CALR | CD44 | 1.0088 | 1.0896 | 1.0492 | 0.000999 |
| MacRec | TCD4reg | CALR | FAS | 1.0088 | 0.2727 | 0.6408 | 0.000999 |
| MacRec | TCD4reg | CALR | IL7R | 1.0088 | 0.9682 | 0.9885 | 0.000999 |
| MacRec | TCD4reg | CCL20 | CCR7 | 0.3793 | 0.3098 | 0.3445 | 0.000999 |
| MacRec | TCD4reg | CCL20 | CXCR3 | 0.3793 | 0.3357 | 0.3575 | 0.000999 |
| MacRec | TCD4reg | CCL20 | CXCR4 | 0.3793 | 2.2962 | 1.3377 | 0.000999 |
| MacRec | TCD4reg | CCL20 | CXCR6 | 0.3793 | 0.3945 | 0.3869 | 0.000999 |
| MacRec | TCD4reg | CCL20 | S1PR4 | 0.3793 | 0.2744 | 0.3268 | 0.000999 |
| MacRec | TCD4reg | CD58 | CD2 | 0.2951 | 1.6566 | 0.9759 | 0.000999 |
| MacRec | TCD4reg | CD86 | CD28 | 0.5786 | 0.313 | 0.4458 | 0.000999 |
| MacRec | TCD4reg | CD86 | CTLA4 | 0.5786 | 0.7909 | 0.6847 | 0.000999 |
| MacRec | TCD4reg | COL1A1 | CD44 | 0.5365 | 1.0896 | 0.8131 | 0.000999 |
| MacRec | TCD4reg | COL1A1 | ITGB1 | 0.5365 | 0.4001 | 0.4683 | 0.000999 |
| MacRec | TCD4reg | CXCL10 | CCR7 | 0.3391 | 0.3098 | 0.3245 | 0.000999 |
| MacRec | TCD4reg | CXCL10 | CXCR3 | 0.3391 | 0.3357 | 0.3374 | 0.000999 |
| MacRec | TCD4reg | CXCL10 | CXCR4 | 0.3391 | 2.2962 | 1.3176 | 0.000999 |
| MacRec | TCD4reg | CXCL10 | CXCR6 | 0.3391 | 0.3945 | 0.3668 | 0.000999 |
| MacRec | TCD4reg | CXCL10 | S1PR4 | 0.3391 | 0.2744 | 0.3067 | 0.000999 |
| MacRec | TCD4reg | CXCL16 | CCR7 | 0.8324 | 0.3098 | 0.5711 | 0.000999 |
| MacRec | TCD4reg | CXCL16 | CXCR3 | 0.8324 | 0.3357 | 0.5841 | 0.000999 |
| MacRec | TCD4reg | CXCL16 | CXCR4 | 0.8324 | 2.2962 | 1.5643 | 0.000999 |
| MacRec | TCD4reg | CXCL16 | CXCR6 | 0.8324 | 0.3945 | 0.6135 | 0.000999 |
| MacRec | TCD4reg | CXCL16 | S1PR4 | 0.8324 | 0.2744 | 0.5534 | 0.000999 |
| MacRec | TCD4reg | CXCL2 | CCR7 | 0.8239 | 0.3098 | 0.5669 | 0.000999 |
| MacRec | TCD4reg | CXCL2 | CXCR3 | 0.8239 | 0.3357 | 0.5798 | 0.000999 |
| MacRec | TCD4reg | CXCL2 | CXCR4 | 0.8239 | 2.2962 | 1.56 | 0.000999 |
| MacRec | TCD4reg | CXCL2 | CXCR6 | 0.8239 | 0.3945 | 0.6092 | 0.000999 |
| MacRec | TCD4reg | CXCL2 | S1PR4 | 0.8239 | 0.2744 | 0.5492 | 0.000999 |
| MacRec | TCD4reg | CXCL3 | CCR7 | 0.5699 | 0.3098 | 0.4399 | 0.000999 |

| | | | | | | | |
|---|---|---|---|---|---|---|---|
| MacRec | TCD4reg | CXCL3 | CXCR3 | 0.5699 | 0.3357 | 0.4528 | 0.000999 |
| MacRec | TCD4reg | CXCL3 | CXCR4 | 0.5699 | 2.2962 | 1.433 | 0.000999 |
| MacRec | TCD4reg | CXCL3 | CXCR6 | 0.5699 | 0.3945 | 0.4822 | 0.000999 |
| MacRec | TCD4reg | CXCL3 | S1PR4 | 0.5699 | 0.2744 | 0.4222 | 0.000999 |
| MacRec | TCD4reg | FN1 | CD44 | 1.5028 | 1.0896 | 1.2962 | 0.000999 |
| MacRec | TCD4reg | FN1 | ITGB1 | 1.5028 | 0.4001 | 0.9514 | 0.000999 |
| MacRec | TCD4reg | FN1 | RPSA | 1.5028 | 2.4999 | 2.0014 | 0.000999 |
| MacRec | TCD4reg | GRN | TNFRSF1B | 1.7446 | 0.7982 | 1.2714 | 0.000999 |
| MacRec | TCD4reg | GSTP1 | CD44 | 1.7225 | 1.0896 | 1.406 | 0.000999 |
| MacRec | TCD4reg | GSTP1 | FAS | 1.7225 | 0.2727 | 0.9976 | 0.000999 |
| MacRec | TCD4reg | GSTP1 | IL7R | 1.7225 | 0.9682 | 1.3453 | 0.000999 |
| MacRec | TCD4reg | HBEGF | CD44 | 0.7419 | 1.0896 | 0.9157 | 0.000999 |
| MacRec | TCD4reg | HLA-DMA | CD4 | 1.7832 | 0.429 | 1.1061 | 0.000999 |
| MacRec | TCD4reg | HLA-DRA | CD4 | 4.3438 | 0.429 | 2.3864 | 0.000999 |
| MacRec | TCD4reg | HLA-DRA | PTPRC | 4.3438 | 1.6603 | 3.0021 | 0.000999 |
| MacRec | TCD4reg | HMGB2 | FAS | 0.7555 | 0.2727 | 0.5141 | 0.000999 |
| MacRec | TCD4reg | HSP90B1 | ITGB1 | 0.7859 | 0.4001 | 0.593 | 0.000999 |
| MacRec | TCD4reg | HSP90B1 | TNFRSF1B | 0.7859 | 0.7982 | 0.792 | 0.000999 |
| MacRec | TCD4reg | LGALS3 | CD6 | 0.9368 | 0.2651 | 0.601 | 0.000999 |
| MacRec | TCD4reg | LGALS3 | FAS | 0.9368 | 0.2727 | 0.6048 | 0.000999 |
| MacRec | TCD4reg | LGALS3 | ITGB1 | 0.9368 | 0.4001 | 0.6685 | 0.000999 |
| MacRec | TCD4reg | MIF | TNFRSF14 | 0.8867 | 0.3843 | 0.6355 | 0.000999 |
| MacRec | TCD4reg | NAMPT | ANXA6 | 1.5176 | 0.4557 | 0.9867 | 0.000999 |
| MacRec | TCD4reg | NECTIN2 | TIGIT | 0.2672 | 1.1883 | 0.7278 | 0.000999 |
| MacRec | TCD4reg | OSM | IL6ST | 0.2991 | 0.2555 | 0.2773 | 0.000999 |
| MacRec | TCD4reg | PECAM1 | CXCR4 | 0.3003 | 2.2962 | 1.2982 | 0.000999 |
| MacRec | TCD4reg | RPS19 | RPSA | 3.7755 | 2.4999 | 3.1377 | 0.000999 |
| MacRec | TCD4reg | RTN4 | CD44 | 1.101 | 1.0896 | 1.0953 | 0.000999 |
| MacRec | TCD4reg | RTN4 | CD99 | 1.101 | 0.6603 | 0.8806 | 0.000999 |
| MacRec | TCD4reg | SPP1 | CD44 | 1.5024 | 1.0896 | 1.296 | 0.000999 |

| | | | | | | | |
|---|---|---|---|---|---|---|---|
| MacRec | TCD4reg | SPP1 | ITGB1 | 1.5024 | 0.4001 | 0.9512 | 0.000999 |
| MacRec | TCD4reg | TGFB1 | ITGB1 | 0.317 | 0.4001 | 0.3585 | 0.000999 |
| MacRec | TCD4reg | THBS1 | CD47 | 0.7038 | 0.416 | 0.5599 | 0.000999 |
| MacRec | TCD4reg | THBS1 | ITGB1 | 0.7038 | 0.4001 | 0.5519 | 0.000999 |
| MacRec | TCD4reg | TNFSF13B | ITGB1 | 0.8304 | 0.4001 | 0.6152 | 0.000999 |
| MacRec | TCD4reg | ARF1 | FAS | 0.7488 | 0.2727 | 0.5108 | 0.004 |
| MacRec | TCD4reg | RPS19 | CD4 | 3.7755 | 0.429 | 2.1022 | 0.00899 |
| MacRec | TCD4reg | HMGB1 | FAS | 1.2904 | 0.2727 | 0.7816 | 0.00999 |
| MacRec | TCD8em | A2M | TNFRSF14 | 0.2825 | 0.2915 | 0.287 | 0.000999 |
| MacRec | TCD8em | ADM | PTGER4 | 0.3154 | 0.2847 | 0.3001 | 0.000999 |
| MacRec | TCD8em | ALCAM | CD6 | 0.2586 | 0.4204 | 0.3395 | 0.000999 |
| MacRec | TCD8em | ANXA1 | CD44 | 1.2584 | 1.119 | 1.1887 | 0.000999 |
| MacRec | TCD8em | ANXA1 | CXCR3 | 1.2584 | 0.4189 | 0.8387 | 0.000999 |
| MacRec | TCD8em | ANXA1 | CXCR4 | 1.2584 | 2.8389 | 2.0487 | 0.000999 |
| MacRec | TCD8em | ANXA1 | CXCR6 | 1.2584 | 0.3399 | 0.7992 | 0.000999 |
| MacRec | TCD8em | ANXA1 | F2R | 1.2584 | 0.2695 | 0.764 | 0.000999 |
| MacRec | TCD8em | ANXA1 | ITGA4 | 1.2584 | 0.2823 | 0.7704 | 0.000999 |
| MacRec | TCD8em | APP | CXCR3 | 0.2502 | 0.4189 | 0.3346 | 0.000999 |
| MacRec | TCD8em | APP | CXCR4 | 0.2502 | 2.8389 | 1.5446 | 0.000999 |
| MacRec | TCD8em | APP | CXCR6 | 0.2502 | 0.3399 | 0.2951 | 0.000999 |
| MacRec | TCD8em | APP | F2R | 0.2502 | 0.2695 | 0.2599 | 0.000999 |
| MacRec | TCD8em | APP | TNFRSF14 | 0.2502 | 0.2915 | 0.2709 | 0.000999 |
| MacRec | TCD8em | ARF1 | ITGA4 | 0.7488 | 0.2823 | 0.5156 | 0.000999 |
| MacRec | TCD8em | C3 | CXCR3 | 0.4838 | 0.4189 | 0.4513 | 0.000999 |
| MacRec | TCD8em | C3 | CXCR4 | 0.4838 | 2.8389 | 1.6613 | 0.000999 |
| MacRec | TCD8em | C3 | CXCR6 | 0.4838 | 0.3399 | 0.4118 | 0.000999 |
| MacRec | TCD8em | C3 | ITGB2 | 0.4838 | 1.0018 | 0.7428 | 0.000999 |
| MacRec | TCD8em | CALR | CD44 | 1.0088 | 1.119 | 1.0639 | 0.000999 |
| MacRec | TCD8em | CALR | IL7R | 1.0088 | 0.7183 | 0.8636 | 0.000999 |
| MacRec | TCD8em | CCL20 | CXCR3 | 0.3793 | 0.4189 | 0.3991 | 0.000999 |

| | | | | | | | |
|---|---|---|---|---|---|---|---|
| MacRec | TCD8em | CCL20 | CXCR4 | 0.3793 | 2.8389 | 1.6091 | 0.000999 |
| MacRec | TCD8em | CCL20 | CXCR6 | 0.3793 | 0.3399 | 0.3596 | 0.000999 |
| MacRec | TCD8em | CCL20 | S1PR4 | 0.3793 | 0.2677 | 0.3235 | 0.000999 |
| MacRec | TCD8em | CD58 | CD2 | 0.2951 | 1.5961 | 0.9456 | 0.000999 |
| MacRec | TCD8em | COL1A1 | CD44 | 0.5365 | 1.119 | 0.8278 | 0.000999 |
| MacRec | TCD8em | COL1A1 | ITGB1 | 0.5365 | 0.2557 | 0.3961 | 0.000999 |
| MacRec | TCD8em | CXCL10 | CXCR3 | 0.3391 | 0.4189 | 0.379 | 0.000999 |
| MacRec | TCD8em | CXCL10 | CXCR4 | 0.3391 | 2.8389 | 1.589 | 0.000999 |
| MacRec | TCD8em | CXCL10 | CXCR6 | 0.3391 | 0.3399 | 0.3395 | 0.000999 |
| MacRec | TCD8em | CXCL10 | S1PR4 | 0.3391 | 0.2677 | 0.3034 | 0.000999 |
| MacRec | TCD8em | CXCL16 | CXCR3 | 0.8324 | 0.4189 | 0.6257 | 0.000999 |
| MacRec | TCD8em | CXCL16 | CXCR4 | 0.8324 | 2.8389 | 1.8357 | 0.000999 |
| MacRec | TCD8em | CXCL16 | CXCR6 | 0.8324 | 0.3399 | 0.5861 | 0.000999 |
| MacRec | TCD8em | CXCL16 | S1PR4 | 0.8324 | 0.2677 | 0.5501 | 0.000999 |
| MacRec | TCD8em | CXCL2 | CXCR3 | 0.8239 | 0.4189 | 0.6214 | 0.000999 |
| MacRec | TCD8em | CXCL2 | CXCR4 | 0.8239 | 2.8389 | 1.8314 | 0.000999 |
| MacRec | TCD8em | CXCL2 | CXCR6 | 0.8239 | 0.3399 | 0.5819 | 0.000999 |
| MacRec | TCD8em | CXCL2 | S1PR4 | 0.8239 | 0.2677 | 0.5458 | 0.000999 |
| MacRec | TCD8em | CXCL3 | CXCR3 | 0.5699 | 0.4189 | 0.4944 | 0.000999 |
| MacRec | TCD8em | CXCL3 | CXCR4 | 0.5699 | 2.8389 | 1.7044 | 0.000999 |
| MacRec | TCD8em | CXCL3 | CXCR6 | 0.5699 | 0.3399 | 0.4549 | 0.000999 |
| MacRec | TCD8em | CXCL3 | S1PR4 | 0.5699 | 0.2677 | 0.4188 | 0.000999 |
| MacRec | TCD8em | FN1 | CD44 | 1.5028 | 1.119 | 1.3109 | 0.000999 |
| MacRec | TCD8em | FN1 | ITGA4 | 1.5028 | 0.2823 | 0.8926 | 0.000999 |
| MacRec | TCD8em | FN1 | ITGB1 | 1.5028 | 0.2557 | 0.8793 | 0.000999 |
| MacRec | TCD8em | FN1 | RPSA | 1.5028 | 2.3686 | 1.9357 | 0.000999 |
| MacRec | TCD8em | GRN | TNFRSF1B | 1.7446 | 0.3038 | 1.0242 | 0.000999 |
| MacRec | TCD8em | GSTP1 | CD44 | 1.7225 | 1.119 | 1.4207 | 0.000999 |
| MacRec | TCD8em | GSTP1 | IL7R | 1.7225 | 0.7183 | 1.2204 | 0.000999 |
| MacRec | TCD8em | GSTP1 | ITGA4 | 1.7225 | 0.2823 | 1.0024 | 0.000999 |

| | | | | | | | |
|---|---|---|---|---|---|---|---|
| MacRec | TCD8em | HBEGF | CD44 | 0.7419 | 1.119 | 0.9304 | 0.000999 |
| MacRec | TCD8em | HLA-DRA | PTPRC | 4.3438 | 1.7912 | 3.0675 | 0.000999 |
| MacRec | TCD8em | HSP90B1 | ITGB1 | 0.7859 | 0.2557 | 0.5208 | 0.000999 |
| MacRec | TCD8em | HSP90B1 | TNFRSF1B | 0.7859 | 0.3038 | 0.5448 | 0.000999 |
| MacRec | TCD8em | ITGB1 | CD47 | 0.4459 | 0.4276 | 0.4367 | 0.000999 |
| MacRec | TCD8em | ITGB1 | ITGA4 | 0.4459 | 0.2823 | 0.3641 | 0.000999 |
| MacRec | TCD8em | ITGB1 | RACK1 | 0.4459 | 2.4374 | 1.4416 | 0.000999 |
| MacRec | TCD8em | ITGB2 | ITGAL | 1.3879 | 0.3162 | 0.8521 | 0.000999 |
| MacRec | TCD8em | LGALS3 | CD6 | 0.9368 | 0.4204 | 0.6786 | 0.000999 |
| MacRec | TCD8em | LGALS3 | ITGB1 | 0.9368 | 0.2557 | 0.5963 | 0.000999 |
| MacRec | TCD8em | LGALS3 | SPN | 0.9368 | 0.2603 | 0.5986 | 0.000999 |
| MacRec | TCD8em | MIF | ITGA4 | 0.8867 | 0.2823 | 0.5845 | 0.000999 |
| MacRec | TCD8em | MIF | TNFRSF14 | 0.8867 | 0.2915 | 0.5891 | 0.000999 |
| MacRec | TCD8em | NAMPT | ANXA6 | 1.5176 | 0.552 | 1.0348 | 0.000999 |
| MacRec | TCD8em | NECTIN2 | TIGIT | 0.2672 | 0.4665 | 0.3669 | 0.000999 |
| MacRec | TCD8em | PECAM1 | CXCR4 | 0.3003 | 2.8389 | 1.5696 | 0.000999 |
| MacRec | TCD8em | RPS19 | ITGA4 | 3.7755 | 0.2823 | 2.0289 | 0.000999 |
| MacRec | TCD8em | RPS19 | RPSA | 3.7755 | 2.3686 | 3.0721 | 0.000999 |
| MacRec | TCD8em | RTN4 | CD44 | 1.101 | 1.119 | 1.11 | 0.000999 |
| MacRec | TCD8em | RTN4 | CD99 | 1.101 | 1.1525 | 1.1267 | 0.000999 |
| MacRec | TCD8em | SPP1 | CD44 | 1.5024 | 1.119 | 1.3107 | 0.000999 |
| MacRec | TCD8em | SPP1 | ITGA4 | 1.5024 | 0.2823 | 0.8924 | 0.000999 |
| MacRec | TCD8em | SPP1 | ITGB1 | 1.5024 | 0.2557 | 0.8791 | 0.000999 |
| MacRec | TCD8em | TGFB1 | ITGB1 | 0.317 | 0.2557 | 0.2864 | 0.000999 |
| MacRec | TCD8em | THBS1 | CD47 | 0.7038 | 0.4276 | 0.5657 | 0.000999 |
| MacRec | TCD8em | THBS1 | ITGA4 | 0.7038 | 0.2823 | 0.4931 | 0.000999 |
| MacRec | TCD8em | THBS1 | ITGB1 | 0.7038 | 0.2557 | 0.4798 | 0.000999 |
| MacRec | TCD8em | TNFSF13B | ITGB1 | 0.8304 | 0.2557 | 0.5431 | 0.000999 |
| MacRec | TCD8em | AIMP1 | RACK1 | 0.2829 | 2.4374 | 1.3601 | 0.002 |
| MacRec | TCD8ex | A2M | TNFRSF14 | 0.2825 | 0.3094 | 0.296 | 0.000999 |

| | | | | | | | |
|---|---|---|---|---|---|---|---|
| MacRec | TCD8ex | ADM | PTGER4 | 0.3154 | 0.3089 | 0.3122 | 0.000999 |
| MacRec | TCD8ex | ALCAM | CD6 | 0.2586 | 0.318 | 0.2883 | 0.000999 |
| MacRec | TCD8ex | ANXA1 | CD44 | 1.2584 | 0.8219 | 1.0402 | 0.000999 |
| MacRec | TCD8ex | ANXA1 | CXCR3 | 1.2584 | 0.3647 | 0.8116 | 0.000999 |
| MacRec | TCD8ex | ANXA1 | CXCR4 | 1.2584 | 2.3472 | 1.8028 | 0.000999 |
| MacRec | TCD8ex | ANXA1 | CXCR6 | 1.2584 | 0.6626 | 0.9605 | 0.000999 |
| MacRec | TCD8ex | ANXA1 | ITGA4 | 1.2584 | 0.4432 | 0.8508 | 0.000999 |
| MacRec | TCD8ex | ANXA1 | VCAM1 | 1.2584 | 0.4462 | 0.8523 | 0.000999 |
| MacRec | TCD8ex | APOE | VCAM1 | 1.1144 | 0.4462 | 0.7803 | 0.000999 |
| MacRec | TCD8ex | APP | CXCR3 | 0.2502 | 0.3647 | 0.3074 | 0.000999 |
| MacRec | TCD8ex | APP | CXCR4 | 0.2502 | 2.3472 | 1.2987 | 0.000999 |
| MacRec | TCD8ex | APP | CXCR6 | 0.2502 | 0.6626 | 0.4564 | 0.000999 |
| MacRec | TCD8ex | APP | TNFRSF14 | 0.2502 | 0.3094 | 0.2798 | 0.000999 |
| MacRec | TCD8ex | C3 | CXCR3 | 0.4838 | 0.3647 | 0.4242 | 0.000999 |
| MacRec | TCD8ex | C3 | CXCR4 | 0.4838 | 2.3472 | 1.4155 | 0.000999 |
| MacRec | TCD8ex | C3 | CXCR6 | 0.4838 | 0.6626 | 0.5732 | 0.000999 |
| MacRec | TCD8ex | C3 | ITGB2 | 0.4838 | 0.7635 | 0.6236 | 0.000999 |
| MacRec | TCD8ex | CCL20 | CXCR3 | 0.3793 | 0.3647 | 0.372 | 0.000999 |
| MacRec | TCD8ex | CCL20 | CXCR4 | 0.3793 | 2.3472 | 1.3632 | 0.000999 |
| MacRec | TCD8ex | CCL20 | CXCR6 | 0.3793 | 0.6626 | 0.5209 | 0.000999 |
| MacRec | TCD8ex | CD58 | CD2 | 0.2951 | 2.0502 | 1.1727 | 0.000999 |
| MacRec | TCD8ex | CD86 | CTLA4 | 0.5786 | 0.4883 | 0.5334 | 0.000999 |
| MacRec | TCD8ex | COL1A1 | CD44 | 0.5365 | 0.8219 | 0.6792 | 0.000999 |
| MacRec | TCD8ex | COL1A1 | ITGB1 | 0.5365 | 0.2995 | 0.418 | 0.000999 |
| MacRec | TCD8ex | CXCL10 | CXCR3 | 0.3391 | 0.3647 | 0.3519 | 0.000999 |
| MacRec | TCD8ex | CXCL10 | CXCR4 | 0.3391 | 2.3472 | 1.3431 | 0.000999 |
| MacRec | TCD8ex | CXCL10 | CXCR6 | 0.3391 | 0.6626 | 0.5008 | 0.000999 |
| MacRec | TCD8ex | CXCL16 | CXCR3 | 0.8324 | 0.3647 | 0.5985 | 0.000999 |
| MacRec | TCD8ex | CXCL16 | CXCR4 | 0.8324 | 2.3472 | 1.5898 | 0.000999 |
| MacRec | TCD8ex | CXCL16 | CXCR6 | 0.8324 | 0.6626 | 0.7475 | 0.000999 |

| | | | | | | | |
|---|---|---|---|---|---|---|---|
| MacRec | TCD8ex | CXCL2 | CXCR3 | 0.8239 | 0.3647 | 0.5943 | 0.000999 |
| MacRec | TCD8ex | CXCL2 | CXCR4 | 0.8239 | 2.3472 | 1.5856 | 0.000999 |
| MacRec | TCD8ex | CXCL2 | CXCR6 | 0.8239 | 0.6626 | 0.7432 | 0.000999 |
| MacRec | TCD8ex | CXCL3 | CXCR3 | 0.5699 | 0.3647 | 0.4673 | 0.000999 |
| MacRec | TCD8ex | CXCL3 | CXCR4 | 0.5699 | 2.3472 | 1.4586 | 0.000999 |
| MacRec | TCD8ex | CXCL3 | CXCR6 | 0.5699 | 0.6626 | 0.6162 | 0.000999 |
| MacRec | TCD8ex | FN1 | CD44 | 1.5028 | 0.8219 | 1.1624 | 0.000999 |
| MacRec | TCD8ex | FN1 | ITGA4 | 1.5028 | 0.4432 | 0.973 | 0.000999 |
| MacRec | TCD8ex | FN1 | ITGB1 | 1.5028 | 0.2995 | 0.9011 | 0.000999 |
| MacRec | TCD8ex | FN1 | ITGB7 | 1.5028 | 0.3007 | 0.9018 | 0.000999 |
| MacRec | TCD8ex | FN1 | RPSA | 1.5028 | 2.2697 | 1.8862 | 0.000999 |
| MacRec | TCD8ex | GRN | TNFRSF1B | 1.7446 | 0.6436 | 1.1941 | 0.000999 |
| MacRec | TCD8ex | GSTP1 | CD44 | 1.7225 | 0.8219 | 1.2722 | 0.000999 |
| MacRec | TCD8ex | GSTP1 | ITGA4 | 1.7225 | 0.4432 | 1.0828 | 0.000999 |
| MacRec | TCD8ex | GSTP1 | VCAM1 | 1.7225 | 0.4462 | 1.0843 | 0.000999 |
| MacRec | TCD8ex | HBEGF | CD44 | 0.7419 | 0.8219 | 0.7819 | 0.000999 |
| MacRec | TCD8ex | HLA-DRA | PDCD1 | 4.3438 | 0.6109 | 2.4774 | 0.000999 |
| MacRec | TCD8ex | HLA-DRA | PTPRC | 4.3438 | 1.6762 | 3.01 | 0.000999 |
| MacRec | TCD8ex | HSP90B1 | ITGB1 | 0.7859 | 0.2995 | 0.5427 | 0.000999 |
| MacRec | TCD8ex | HSP90B1 | TNFRSF1B | 0.7859 | 0.6436 | 0.7148 | 0.000999 |
| MacRec | TCD8ex | ITGB1 | CD47 | 0.4459 | 0.3952 | 0.4205 | 0.000999 |
| MacRec | TCD8ex | ITGB1 | ITGA4 | 0.4459 | 0.4432 | 0.4446 | 0.000999 |
| MacRec | TCD8ex | ITGB1 | RACK1 | 0.4459 | 2.3353 | 1.3906 | 0.000999 |
| MacRec | TCD8ex | ITGB1 | VCAM1 | 0.4459 | 0.4462 | 0.446 | 0.000999 |
| MacRec | TCD8ex | LGALS3 | CD6 | 0.9368 | 0.318 | 0.6274 | 0.000999 |
| MacRec | TCD8ex | LGALS3 | ITGB1 | 0.9368 | 0.2995 | 0.6182 | 0.000999 |
| MacRec | TCD8ex | LGALS3 | SPN | 0.9368 | 0.3486 | 0.6427 | 0.000999 |
| MacRec | TCD8ex | MIF | ITGA4 | 0.8867 | 0.4432 | 0.665 | 0.000999 |
| MacRec | TCD8ex | MIF | TNFRSF14 | 0.8867 | 0.3094 | 0.5981 | 0.000999 |
| MacRec | TCD8ex | NAMPT | ANXA6 | 1.5176 | 0.5678 | 1.0427 | 0.000999 |

| | | | | | | | |
|---|---|---|---|---|---|---|---|
| MacRec | TCD8ex | NECTIN2 | TIGIT | 0.2672 | 0.9854 | 0.6263 | 0.000999 |
| MacRec | TCD8ex | PECAM1 | CXCR4 | 0.3003 | 2.3472 | 1.3238 | 0.000999 |
| MacRec | TCD8ex | RTN4 | CD44 | 1.101 | 0.8219 | 0.9614 | 0.000999 |
| MacRec | TCD8ex | RTN4 | CD99 | 1.101 | 1.2049 | 1.1529 | 0.000999 |
| MacRec | TCD8ex | SPP1 | CD44 | 1.5024 | 0.8219 | 1.1622 | 0.000999 |
| MacRec | TCD8ex | SPP1 | ITGA4 | 1.5024 | 0.4432 | 0.9728 | 0.000999 |
| MacRec | TCD8ex | SPP1 | ITGB1 | 1.5024 | 0.2995 | 0.9009 | 0.000999 |
| MacRec | TCD8ex | TGFB1 | ITGB1 | 0.317 | 0.2995 | 0.3082 | 0.000999 |
| MacRec | TCD8ex | THBS1 | CD47 | 0.7038 | 0.3952 | 0.5495 | 0.000999 |
| MacRec | TCD8ex | THBS1 | ITGA4 | 0.7038 | 0.4432 | 0.5735 | 0.000999 |
| MacRec | TCD8ex | THBS1 | ITGB1 | 0.7038 | 0.2995 | 0.5016 | 0.000999 |
| MacRec | TCD8ex | TNFSF13B | ITGB1 | 0.8304 | 0.2995 | 0.5649 | 0.000999 |
| MacRec | TGD | A2M | TNFRSF14 | 0.2825 | 0.4642 | 0.3733 | 0.000999 |
| MacRec | TGD | ADM | PTGDR | 0.3154 | 0.7153 | 0.5154 | 0.000999 |
| MacRec | TGD | ALCAM | CD6 | 0.2586 | 0.4684 | 0.3635 | 0.000999 |
| MacRec | TGD | ANXA1 | CXCR3 | 1.2584 | 1.1748 | 1.2166 | 0.000999 |
| MacRec | TGD | ANXA1 | ITGA4 | 1.2584 | 0.3968 | 0.8276 | 0.000999 |
| MacRec | TGD | APP | CXCR3 | 0.2502 | 1.1748 | 0.7125 | 0.000999 |
| MacRec | TGD | APP | TGFBR2 | 0.2502 | 0.3677 | 0.309 | 0.000999 |
| MacRec | TGD | APP | TNFRSF14 | 0.2502 | 0.4642 | 0.3572 | 0.000999 |
| MacRec | TGD | AREG | ICAM1 | 1.5622 | 0.2915 | 0.9269 | 0.000999 |
| MacRec | TGD | ARF1 | ITGA4 | 0.7488 | 0.3968 | 0.5728 | 0.000999 |
| MacRec | TGD | C3 | CXCR3 | 0.4838 | 1.1748 | 0.8293 | 0.000999 |
| MacRec | TGD | CALM1 | RPSA | 1.1805 | 2.8814 | 2.0309 | 0.000999 |
| MacRec | TGD | CALR | IL7R | 1.0088 | 1.3862 | 1.1975 | 0.000999 |
| MacRec | TGD | CCL20 | CXCR3 | 0.3793 | 1.1748 | 0.777 | 0.000999 |
| MacRec | TGD | CD58 | CD2 | 0.2951 | 1.4522 | 0.8737 | 0.000999 |
| MacRec | TGD | CD99 | CD99 | 0.6603 | 1.7154 | 1.1879 | 0.000999 |
| MacRec | TGD | CXCL10 | CXCR3 | 0.3391 | 1.1748 | 0.7569 | 0.000999 |
| MacRec | TGD | CXCL16 | CXCR3 | 0.8324 | 1.1748 | 1.0036 | 0.000999 |

| | | | | | | | |
|---|---|---|---|---|---|---|---|
| MacRec | TGD | CXCL2 | CXCR3 | 0.8239 | 1.1748 | 0.9993 | 0.000999 |
| MacRec | TGD | CXCL3 | CXCR3 | 0.5699 | 1.1748 | 0.8723 | 0.000999 |
| MacRec | TGD | FN1 | ITGA4 | 1.5028 | 0.3968 | 0.9498 | 0.000999 |
| MacRec | TGD | FN1 | ITGB7 | 1.5028 | 0.459 | 0.9809 | 0.000999 |
| MacRec | TGD | FN1 | RPSA | 1.5028 | 2.8814 | 2.1921 | 0.000999 |
| MacRec | TGD | GRN | TNFRSF1A | 1.7446 | 0.3099 | 1.0272 | 0.000999 |
| MacRec | TGD | GSTP1 | IL7R | 1.7225 | 1.3862 | 1.5543 | 0.000999 |
| MacRec | TGD | GSTP1 | ITGA4 | 1.7225 | 0.3968 | 1.0596 | 0.000999 |
| MacRec | TGD | HLA-A | CD8A | 2.8028 | 1.746 | 2.2744 | 0.000999 |
| MacRec | TGD | HLA-DRA | PTPRC | 4.3438 | 1.8653 | 3.1046 | 0.000999 |
| MacRec | TGD | HLA-E | CD8A | 1.5633 | 1.746 | 1.6547 | 0.000999 |
| MacRec | TGD | HLA-E | KLRC2 | 1.5633 | 0.8082 | 1.1858 | 0.000999 |
| MacRec | TGD | HLA-E | KLRD1 | 1.5633 | 1.3451 | 1.4542 | 0.000999 |
| MacRec | TGD | HLA-F | CD8A | 0.3045 | 1.746 | 1.0252 | 0.000999 |
| MacRec | TGD | HMGB1 | ITGA4 | 1.2904 | 0.3968 | 0.8436 | 0.000999 |
| MacRec | TGD | ITGB1 | ITGA4 | 0.4459 | 0.3968 | 0.4214 | 0.000999 |
| MacRec | TGD | ITGB2 | ITGAL | 1.3879 | 0.3731 | 0.8805 | 0.000999 |
| MacRec | TGD | LGALS3 | CD6 | 0.9368 | 0.4684 | 0.7026 | 0.000999 |
| MacRec | TGD | LGALS3 | SPN | 0.9368 | 0.5644 | 0.7506 | 0.000999 |
| MacRec | TGD | LGALS3 | TGFBR2 | 0.9368 | 0.3677 | 0.6523 | 0.000999 |
| MacRec | TGD | MIF | ITGA4 | 0.8867 | 0.3968 | 0.6418 | 0.000999 |
| MacRec | TGD | NAMPT | ANXA6 | 1.5176 | 0.8553 | 1.1864 | 0.000999 |
| MacRec | TGD | PTPRC | CD2 | 0.8059 | 1.4522 | 1.129 | 0.000999 |
| MacRec | TGD | PTPRC | CD8A | 0.8059 | 1.746 | 1.276 | 0.000999 |
| MacRec | TGD | RPS19 | ITGA4 | 3.7755 | 0.3968 | 2.0862 | 0.000999 |
| MacRec | TGD | RPS19 | RPSA | 3.7755 | 2.8814 | 3.3284 | 0.000999 |
| MacRec | TGD | RTN4 | CD99 | 1.101 | 1.7154 | 1.4082 | 0.000999 |
| MacRec | TGD | SPP1 | ITGA4 | 1.5024 | 0.3968 | 0.9496 | 0.000999 |
| MacRec | TGD | SPP1 | ITGB1 | 1.5024 | 0.3657 | 0.934 | 0.000999 |
| MacRec | TGD | THBS1 | ITGA4 | 0.7038 | 0.3968 | 0.5503 | 0.000999 |

| | | | | | | | |
|---|---|---|---|---|---|---|---|
| MacRec | TGD | VEGFA | PTK2B | 0.6067 | 0.3668 | 0.4868 | 0.000999 |
| MacRec | TGD | ITGB1 | CD46 | 0.4459 | 0.3085 | 0.3772 | 0.002 |
| MacRec | TGD | PTPRC | ITGAL | 0.8059 | 0.3731 | 0.5895 | 0.002 |
| MacRec | TGD | FN1 | ITGB1 | 1.5028 | 0.3657 | 0.9342 | 0.003 |
| MacRec | TGD | ITGB1 | PTK2B | 0.4459 | 0.3668 | 0.4063 | 0.003 |
| MacRec | TGD | AIMP1 | RACK1 | 0.2829 | 2.7287 | 1.5058 | 0.004 |
| MacRec | TGD | ITGB1 | RACK1 | 0.4459 | 2.7287 | 1.5873 | 0.004 |
| NKT | MacRec | ALOX5AP | ALOX5 | 0.5021 | 0.4156 | 0.4588 | 0.000999 |
| NKT | MacRec | ANXA1 | C3AR1 | 1.2259 | 0.425 | 0.8255 | 0.000999 |
| NKT | MacRec | ANXA1 | C5AR1 | 1.2259 | 0.9166 | 1.0713 | 0.000999 |
| NKT | MacRec | ANXA1 | CD4 | 1.2259 | 0.5246 | 0.8753 | 0.000999 |
| NKT | MacRec | ANXA1 | CD44 | 1.2259 | 1.443 | 1.3345 | 0.000999 |
| NKT | MacRec | ANXA1 | FPR1 | 1.2259 | 0.729 | 0.9775 | 0.000999 |
| NKT | MacRec | ANXA1 | FPR3 | 1.2259 | 0.5944 | 0.9102 | 0.000999 |
| NKT | MacRec | ARF1 | CD4 | 0.7991 | 0.5246 | 0.6619 | 0.000999 |
| NKT | MacRec | ARF1 | CD44 | 0.7991 | 1.443 | 1.1211 | 0.000999 |
| NKT | MacRec | CALM1 | RPSA | 1.9335 | 2.3088 | 2.1212 | 0.000999 |
| NKT | MacRec | CALM1 | TNFRSF1A | 1.9335 | 0.4068 | 1.1702 | 0.000999 |
| NKT | MacRec | CALM1 | TNFRSF1B | 1.9335 | 0.5989 | 1.2662 | 0.000999 |
| NKT | MacRec | CALR | CD44 | 0.9479 | 1.443 | 1.1955 | 0.000999 |
| NKT | MacRec | CALR | LRP1 | 0.9479 | 0.3912 | 0.6696 | 0.000999 |
| NKT | MacRec | CD2 | CD4 | 1.1465 | 0.5246 | 0.8356 | 0.000999 |
| NKT | MacRec | CD6 | ALCAM | 0.4881 | 0.2586 | 0.3733 | 0.000999 |
| NKT | MacRec | GSTP1 | CD44 | 0.7909 | 1.443 | 1.1169 | 0.000999 |
| NKT | MacRec | HMGB1 | AGTRAP | 1.195 | 0.3591 | 0.777 | 0.000999 |
| NKT | MacRec | HMGB1 | LY96 | 1.195 | 0.8566 | 1.0258 | 0.000999 |
| NKT | MacRec | HMGB1 | TLR2 | 1.195 | 0.3974 | 0.7962 | 0.000999 |
| NKT | MacRec | HMGB2 | CD44 | 0.6467 | 1.443 | 1.0449 | 0.000999 |
| NKT | MacRec | HMGB2 | DAB2 | 0.6467 | 0.3042 | 0.4755 | 0.000999 |
| NKT | MacRec | HSP90B1 | LRP1 | 0.6138 | 0.3912 | 0.5025 | 0.000999 |

| | | | | | | | |
|---|---|---|---|---|---|---|---|
| NKT | MacRec | IL16 | CD4 | 0.3556 | 0.5246 | 0.4401 | 0.000999 |
| NKT | MacRec | IL16 | FCGR1A | 0.3556 | 0.603 | 0.4793 | 0.000999 |
| NKT | MacRec | ITGA4 | CD44 | 0.2946 | 1.443 | 0.8688 | 0.000999 |
| NKT | MacRec | ITGA4 | TFRC | 0.2946 | 0.5511 | 0.4228 | 0.000999 |
| NKT | MacRec | ITGB1 | CD151 | 0.488 | 0.2764 | 0.3822 | 0.000999 |
| NKT | MacRec | ITGB1 | CD36 | 0.488 | 0.3023 | 0.3951 | 0.000999 |
| NKT | MacRec | ITGB1 | DAB2 | 0.488 | 0.3042 | 0.3961 | 0.000999 |
| NKT | MacRec | ITGB1 | ENG | 0.488 | 0.3417 | 0.4148 | 0.000999 |
| NKT | MacRec | ITGB1 | ITGAX | 0.488 | 0.3629 | 0.4254 | 0.000999 |
| NKT | MacRec | ITGB1 | PLAUR | 0.488 | 1.5641 | 1.026 | 0.000999 |
| NKT | MacRec | ITGB1 | RACK1 | 0.488 | 2.6054 | 1.5467 | 0.000999 |
| NKT | MacRec | ITGB2 | ITGAM | 1.6481 | 0.3077 | 0.9779 | 0.000999 |
| NKT | MacRec | ITGB2 | ITGAX | 1.6481 | 0.3629 | 1.0055 | 0.000999 |
| NKT | MacRec | ITGB2 | PLAUR | 1.6481 | 1.5641 | 1.6061 | 0.000999 |
| NKT | MacRec | LRPAP1 | LRP1 | 0.2807 | 0.3912 | 0.336 | 0.000999 |
| NKT | MacRec | LTB | TNFRSF1A | 0.2605 | 0.4068 | 0.3336 | 0.000999 |
| NKT | MacRec | MIF | CD74 | 0.6657 | 4.4668 | 2.5663 | 0.000999 |
| NKT | MacRec | PTPRC | CD4 | 1.8797 | 0.5246 | 1.2021 | 0.000999 |
| NKT | MacRec | PTPRC | CD44 | 1.8797 | 1.443 | 1.6613 | 0.000999 |
| NKT | MacRec | RPS19 | C5AR1 | 3.5336 | 0.9166 | 2.2251 | 0.000999 |
| NKT | MacRec | RPS19 | CD4 | 3.5336 | 0.5246 | 2.0291 | 0.000999 |
| NKT | MacRec | RTN4 | CD44 | 0.4374 | 1.443 | 0.9402 | 0.000999 |
| NKT | MacRec | TGFB1 | ENG | 0.5441 | 0.3417 | 0.4429 | 0.000999 |
| NKT | MacRec | TGFB1 | FCN1 | 0.5441 | 1.1325 | 0.8383 | 0.000999 |
| NKT | MacRec | TGFB1 | TLR2 | 0.5441 | 0.3974 | 0.4707 | 0.000999 |
| NKT | MacRec | TIGIT | NECTIN2 | 0.35 | 0.2672 | 0.3086 | 0.000999 |
| NKT | MacRec | XCL2 | LPAR6 | 0.2774 | 0.2983 | 0.2879 | 0.000999 |
| TCD4ex | MacRec | ALOX5AP | ALOX5 | 1.119 | 0.4156 | 0.7673 | 0.000999 |
| TCD4ex | MacRec | ANXA1 | C3AR1 | 0.5174 | 0.425 | 0.4712 | 0.000999 |
| TCD4ex | MacRec | ANXA1 | C5AR1 | 0.5174 | 0.9166 | 0.717 | 0.000999 |

| | | | | | | | |
|---|---|---|---|---|---|---|---|
| TCD4ex | MacRec | ANXA1 | FPR1 | 0.5174 | 0.729 | 0.6232 | 0.000999 |
| TCD4ex | MacRec | ANXA1 | FPR3 | 0.5174 | 0.5944 | 0.5559 | 0.000999 |
| TCD4ex | MacRec | ARF1 | CD44 | 0.6827 | 1.443 | 1.0628 | 0.000999 |
| TCD4ex | MacRec | BTLA | TNFRSF14 | 0.3493 | 0.27 | 0.3097 | 0.000999 |
| TCD4ex | MacRec | CALM1 | TNFRSF1A | 1.2957 | 0.4068 | 0.8513 | 0.000999 |
| TCD4ex | MacRec | CALM1 | TNFRSF1B | 1.2957 | 0.5989 | 0.9473 | 0.000999 |
| TCD4ex | MacRec | CALR | CD44 | 0.748 | 1.443 | 1.0955 | 0.000999 |
| TCD4ex | MacRec | CALR | LRP1 | 0.748 | 0.3912 | 0.5696 | 0.000999 |
| TCD4ex | MacRec | CD2 | CD4 | 1.5731 | 0.5246 | 1.0489 | 0.000999 |
| TCD4ex | MacRec | CD6 | ALCAM | 0.5338 | 0.2586 | 0.3962 | 0.000999 |
| TCD4ex | MacRec | CXCL13 | C3AR1 | 2.3367 | 0.425 | 1.3808 | 0.000999 |
| TCD4ex | MacRec | CXCL13 | C5AR1 | 2.3367 | 0.9166 | 1.6267 | 0.000999 |
| TCD4ex | MacRec | CXCL13 | CXCR4 | 2.3367 | 1.6677 | 2.0022 | 0.000999 |
| TCD4ex | MacRec | CXCL13 | FPR1 | 2.3367 | 0.729 | 1.5328 | 0.000999 |
| TCD4ex | MacRec | HMGB1 | AGTRAP | 1.3876 | 0.3591 | 0.8733 | 0.000999 |
| TCD4ex | MacRec | HMGB1 | LY96 | 1.3876 | 0.8566 | 1.1221 | 0.000999 |
| TCD4ex | MacRec | HMGB1 | TLR2 | 1.3876 | 0.3974 | 0.8925 | 0.000999 |
| TCD4ex | MacRec | HMGB2 | CD44 | 1.0243 | 1.443 | 1.2337 | 0.000999 |
| TCD4ex | MacRec | HMGB2 | DAB2 | 1.0243 | 0.3042 | 0.6643 | 0.000999 |
| TCD4ex | MacRec | HSP90B1 | LRP1 | 0.3944 | 0.3912 | 0.3928 | 0.000999 |
| TCD4ex | MacRec | ICAM3 | ITGB2 | 0.573 | 1.3879 | 0.9805 | 0.000999 |
| TCD4ex | MacRec | ITGB1 | CD151 | 0.4615 | 0.2764 | 0.3689 | 0.000999 |
| TCD4ex | MacRec | ITGB1 | CD36 | 0.4615 | 0.3023 | 0.3819 | 0.000999 |
| TCD4ex | MacRec | ITGB1 | CD47 | 0.4615 | 0.5042 | 0.4828 | 0.000999 |
| TCD4ex | MacRec | ITGB1 | DAB2 | 0.4615 | 0.3042 | 0.3829 | 0.000999 |
| TCD4ex | MacRec | ITGB1 | ENG | 0.4615 | 0.3417 | 0.4016 | 0.000999 |
| TCD4ex | MacRec | ITGB1 | ITGAX | 0.4615 | 0.3629 | 0.4122 | 0.000999 |
| TCD4ex | MacRec | ITGB1 | PLAUR | 0.4615 | 1.5641 | 1.0128 | 0.000999 |
| TCD4ex | MacRec | ITGB1 | RACK1 | 0.4615 | 2.6054 | 1.5334 | 0.000999 |
| TCD4ex | MacRec | ITGB2 | ITGAX | 0.7871 | 0.3629 | 0.575 | 0.000999 |

| | | | | | | | |
|---|---|---|---|---|---|---|---|
| TCD4ex | MacRec | ITGB2 | PLAUR | 0.7871 | 1.5641 | 1.1756 | 0.000999 |
| TCD4ex | MacRec | LRPAP1 | LRP1 | 0.2809 | 0.3912 | 0.3361 | 0.000999 |
| TCD4ex | MacRec | LRPAP1 | SORL1 | 0.2809 | 0.5066 | 0.3938 | 0.000999 |
| TCD4ex | MacRec | LTB | ITGB1 | 0.9655 | 0.4459 | 0.7057 | 0.000999 |
| TCD4ex | MacRec | LTB | TNFRSF1A | 0.9655 | 0.4068 | 0.6862 | 0.000999 |
| TCD4ex | MacRec | MIF | CD74 | 0.6999 | 4.4668 | 2.5833 | 0.000999 |
| TCD4ex | MacRec | NMB | GPR65 | 0.6391 | 0.3011 | 0.4701 | 0.000999 |
| TCD4ex | MacRec | NMB | LPAR6 | 0.6391 | 0.2983 | 0.4687 | 0.000999 |
| TCD4ex | MacRec | PTPRC | CD4 | 1.7493 | 0.5246 | 1.137 | 0.000999 |
| TCD4ex | MacRec | PTPRC | CD44 | 1.7493 | 1.443 | 1.5962 | 0.000999 |
| TCD4ex | MacRec | RPS19 | C5AR1 | 3.6329 | 0.9166 | 2.2748 | 0.000999 |
| TCD4ex | MacRec | SELL | CD4 | 0.5618 | 0.5246 | 0.5432 | 0.000999 |
| TCD4ex | MacRec | TIGIT | NECTIN2 | 0.853 | 0.2672 | 0.5601 | 0.000999 |
| TCD4ex | MacRec | ANXA1 | CD44 | 0.5174 | 1.443 | 0.9802 | 0.004 |
| TCD4ex | MacRec | ITGB2 | ITGAM | 0.7871 | 0.3077 | 0.5474 | 0.004 |
| TCD4ex | MacRec | RTN4 | CD44 | 0.355 | 1.443 | 0.899 | 0.00599 |
| TCD4reg | MacRec | ALOX5AP | ALOX5 | 0.4589 | 0.4156 | 0.4372 | 0.000999 |
| TCD4reg | MacRec | APOE | LRP1 | 0.2934 | 0.3912 | 0.3423 | 0.000999 |
| TCD4reg | MacRec | APOE | SORL1 | 0.2934 | 0.5066 | 0.4 | 0.000999 |
| TCD4reg | MacRec | ARF1 | CD4 | 0.7185 | 0.5246 | 0.6216 | 0.000999 |
| TCD4reg | MacRec | ARF1 | CD44 | 0.7185 | 1.443 | 1.0807 | 0.000999 |
| TCD4reg | MacRec | CALM1 | TNFRSF1A | 1.534 | 0.4068 | 0.9704 | 0.000999 |
| TCD4reg | MacRec | CALR | CD44 | 0.6835 | 1.443 | 1.0632 | 0.000999 |
| TCD4reg | MacRec | CALR | LRP1 | 0.6835 | 0.3912 | 0.5374 | 0.000999 |
| TCD4reg | MacRec | CD2 | CD4 | 1.6566 | 0.5246 | 1.0906 | 0.000999 |
| TCD4reg | MacRec | CD28 | CD4 | 0.313 | 0.5246 | 0.4188 | 0.000999 |
| TCD4reg | MacRec | CD6 | ALCAM | 0.2651 | 0.2586 | 0.2618 | 0.000999 |
| TCD4reg | MacRec | GSTP1 | CD44 | 0.4917 | 1.443 | 0.9673 | 0.000999 |
| TCD4reg | MacRec | HMGB1 | AGTRAP | 1.2547 | 0.3591 | 0.8069 | 0.000999 |
| TCD4reg | MacRec | HMGB1 | LY96 | 1.2547 | 0.8566 | 1.0557 | 0.000999 |

| | | | | | | | |
|---|---|---|---|---|---|---|---|
| TCD4reg | MacRec | HMGB1 | TLR2 | 1.2547 | 0.3974 | 0.8261 | 0.000999 |
| TCD4reg | MacRec | HMGB2 | CD44 | 0.616 | 1.443 | 1.0295 | 0.000999 |
| TCD4reg | MacRec | HMGB2 | DAB2 | 0.616 | 0.3042 | 0.4601 | 0.000999 |
| TCD4reg | MacRec | HSP90B1 | LRP1 | 0.4221 | 0.3912 | 0.4067 | 0.000999 |
| TCD4reg | MacRec | ICAM3 | ITGB2 | 0.6663 | 1.3879 | 1.0271 | 0.000999 |
| TCD4reg | MacRec | IL16 | CD4 | 0.2528 | 0.5246 | 0.3887 | 0.000999 |
| TCD4reg | MacRec | IL16 | FCGR1A | 0.2528 | 0.603 | 0.4279 | 0.000999 |
| TCD4reg | MacRec | ITGB1 | CD151 | 0.4001 | 0.2764 | 0.3382 | 0.000999 |
| TCD4reg | MacRec | ITGB1 | CD36 | 0.4001 | 0.3023 | 0.3512 | 0.000999 |
| TCD4reg | MacRec | ITGB1 | CD47 | 0.4001 | 0.5042 | 0.4521 | 0.000999 |
| TCD4reg | MacRec | ITGB1 | DAB2 | 0.4001 | 0.3042 | 0.3522 | 0.000999 |
| TCD4reg | MacRec | ITGB1 | ENG | 0.4001 | 0.3417 | 0.3709 | 0.000999 |
| TCD4reg | MacRec | ITGB1 | ITGAX | 0.4001 | 0.3629 | 0.3815 | 0.000999 |
| TCD4reg | MacRec | ITGB1 | PLAUR | 0.4001 | 1.5641 | 0.9821 | 0.000999 |
| TCD4reg | MacRec | ITGB2 | ITGAM | 0.6238 | 0.3077 | 0.4657 | 0.000999 |
| TCD4reg | MacRec | ITGB2 | ITGAX | 0.6238 | 0.3629 | 0.4933 | 0.000999 |
| TCD4reg | MacRec | ITGB2 | PLAUR | 0.6238 | 1.5641 | 1.0939 | 0.000999 |
| TCD4reg | MacRec | LGALS3 | CLEC7A | 0.4492 | 0.8554 | 0.6523 | 0.000999 |
| TCD4reg | MacRec | LGALS3 | FCGR2A | 0.4492 | 0.8803 | 0.6647 | 0.000999 |
| TCD4reg | MacRec | LGALS3 | FCGR2B | 0.4492 | 0.7132 | 0.5812 | 0.000999 |
| TCD4reg | MacRec | LGALS3 | TFRC | 0.4492 | 0.5511 | 0.5001 | 0.000999 |
| TCD4reg | MacRec | LRPAP1 | LRP1 | 0.3101 | 0.3912 | 0.3507 | 0.000999 |
| TCD4reg | MacRec | LRPAP1 | SORL1 | 0.3101 | 0.5066 | 0.4084 | 0.000999 |
| TCD4reg | MacRec | LTB | ITGB1 | 2.242 | 0.4459 | 1.344 | 0.000999 |
| TCD4reg | MacRec | LTB | TNFRSF1A | 2.242 | 0.4068 | 1.3244 | 0.000999 |
| TCD4reg | MacRec | MIF | CD74 | 0.6295 | 4.4668 | 2.5482 | 0.000999 |
| TCD4reg | MacRec | PSAP | LRP1 | 0.5938 | 0.3912 | 0.4925 | 0.000999 |
| TCD4reg | MacRec | PTPRC | CD4 | 1.6603 | 0.5246 | 1.0925 | 0.000999 |
| TCD4reg | MacRec | PTPRC | CD44 | 1.6603 | 1.443 | 1.5517 | 0.000999 |
| TCD4reg | MacRec | RPS19 | C5AR1 | 3.6897 | 0.9166 | 2.3032 | 0.000999 |

| | | | | | | | |
|---|---|---|---|---|---|---|---|
| TCD4reg | MacRec | RTN4 | CD44 | 0.3528 | 1.443 | 0.8979 | 0.000999 |
| TCD4reg | MacRec | SELL | CD4 | 0.7062 | 0.5246 | 0.6154 | 0.000999 |
| TCD4reg | MacRec | SPP1 | CD44 | 0.3125 | 1.443 | 0.8778 | 0.000999 |
| TCD4reg | MacRec | TIGIT | NECTIN2 | 1.1883 | 0.2672 | 0.7278 | 0.000999 |
| TCD4reg | MacRec | TIMP1 | LRP1 | 0.4362 | 0.3912 | 0.4137 | 0.000999 |
| TCD4reg | MacRec | RPS19 | CD4 | 3.6897 | 0.5246 | 2.1071 | 0.002 |
| TCD8em | MacRec | ALOX5AP | ALOX5 | 0.7973 | 0.4156 | 0.6065 | 0.000999 |
| TCD8em | MacRec | ANXA1 | C3AR1 | 1.0693 | 0.425 | 0.7471 | 0.000999 |
| TCD8em | MacRec | ANXA1 | C5AR1 | 1.0693 | 0.9166 | 0.9929 | 0.000999 |
| TCD8em | MacRec | ANXA1 | CD4 | 1.0693 | 0.5246 | 0.7969 | 0.000999 |
| TCD8em | MacRec | ANXA1 | CD44 | 1.0693 | 1.443 | 1.2561 | 0.000999 |
| TCD8em | MacRec | ANXA1 | FPR1 | 1.0693 | 0.729 | 0.8991 | 0.000999 |
| TCD8em | MacRec | ANXA1 | FPR3 | 1.0693 | 0.5944 | 0.8318 | 0.000999 |
| TCD8em | MacRec | APOE | LRP1 | 0.2791 | 0.3912 | 0.3352 | 0.000999 |
| TCD8em | MacRec | APOE | SORL1 | 0.2791 | 0.5066 | 0.3929 | 0.000999 |
| TCD8em | MacRec | ARF1 | CD4 | 0.6684 | 0.5246 | 0.5965 | 0.000999 |
| TCD8em | MacRec | ARF1 | CD44 | 0.6684 | 1.443 | 1.0557 | 0.000999 |
| TCD8em | MacRec | CALM1 | TNFRSF1A | 1.5262 | 0.4068 | 0.9665 | 0.000999 |
| TCD8em | MacRec | CALM1 | TNFRSF1B | 1.5262 | 0.5989 | 1.0625 | 0.000999 |
| TCD8em | MacRec | CALR | CD44 | 0.7141 | 1.443 | 1.0786 | 0.000999 |
| TCD8em | MacRec | CALR | LRP1 | 0.7141 | 0.3912 | 0.5527 | 0.000999 |
| TCD8em | MacRec | CD2 | CD4 | 1.5961 | 0.5246 | 1.0604 | 0.000999 |
| TCD8em | MacRec | CD6 | ALCAM | 0.4204 | 0.2586 | 0.3395 | 0.000999 |
| TCD8em | MacRec | CRTAM | NECTIN2 | 0.2993 | 0.2672 | 0.2832 | 0.000999 |
| TCD8em | MacRec | GSTP1 | CD44 | 0.767 | 1.443 | 1.105 | 0.000999 |
| TCD8em | MacRec | HLA-DMA | CD4 | 0.2969 | 0.5246 | 0.4108 | 0.000999 |
| TCD8em | MacRec | HMGB1 | AGTRAP | 1.3636 | 0.3591 | 0.8614 | 0.000999 |
| TCD8em | MacRec | HMGB1 | LY96 | 1.3636 | 0.8566 | 1.1101 | 0.000999 |
| TCD8em | MacRec | HMGB1 | TLR2 | 1.3636 | 0.3974 | 0.8805 | 0.000999 |
| TCD8em | MacRec | HMGB2 | CD44 | 0.7212 | 1.443 | 1.0821 | 0.000999 |

| | | | | | | | |
|---|---|---|---|---|---|---|---|
| TCD8em | MacRec | HMGB2 | DAB2 | 0.7212 | 0.3042 | 0.5127 | 0.000999 |
| TCD8em | MacRec | HSP90B1 | ITGB1 | 0.471 | 0.4459 | 0.4585 | 0.000999 |
| TCD8em | MacRec | HSP90B1 | LRP1 | 0.471 | 0.3912 | 0.4311 | 0.000999 |
| TCD8em | MacRec | HSP90B1 | TNFRSF1B | 0.471 | 0.5989 | 0.5349 | 0.000999 |
| TCD8em | MacRec | ICAM3 | ITGB2 | 0.5282 | 1.3879 | 0.9581 | 0.000999 |
| TCD8em | MacRec | IFNG | IFNGR1 | 0.3967 | 0.7181 | 0.5574 | 0.000999 |
| TCD8em | MacRec | IFNG | IFNGR2 | 0.3967 | 0.7916 | 0.5942 | 0.000999 |
| TCD8em | MacRec | IFNG | PTPN6 | 0.3967 | 0.323 | 0.3598 | 0.000999 |
| TCD8em | MacRec | IL16 | CD4 | 0.4085 | 0.5246 | 0.4665 | 0.000999 |
| TCD8em | MacRec | IL16 | FCGR1A | 0.4085 | 0.603 | 0.5057 | 0.000999 |
| TCD8em | MacRec | ITGA4 | CD44 | 0.2823 | 1.443 | 0.8627 | 0.000999 |
| TCD8em | MacRec | ITGA4 | TFRC | 0.2823 | 0.5511 | 0.4167 | 0.000999 |
| TCD8em | MacRec | ITGB1 | CD151 | 0.2557 | 0.2764 | 0.266 | 0.000999 |
| TCD8em | MacRec | ITGB1 | CD36 | 0.2557 | 0.3023 | 0.279 | 0.000999 |
| TCD8em | MacRec | ITGB1 | DAB2 | 0.2557 | 0.3042 | 0.28 | 0.000999 |
| TCD8em | MacRec | ITGB1 | ENG | 0.2557 | 0.3417 | 0.2987 | 0.000999 |
| TCD8em | MacRec | ITGB1 | ITGAX | 0.2557 | 0.3629 | 0.3093 | 0.000999 |
| TCD8em | MacRec | ITGB1 | PLAUR | 0.2557 | 1.5641 | 0.9099 | 0.000999 |
| TCD8em | MacRec | ITGB1 | RACK1 | 0.2557 | 2.6054 | 1.4306 | 0.000999 |
| TCD8em | MacRec | ITGB2 | ITGAM | 1.0018 | 0.3077 | 0.6547 | 0.000999 |
| TCD8em | MacRec | ITGB2 | ITGAX | 1.0018 | 0.3629 | 0.6823 | 0.000999 |
| TCD8em | MacRec | ITGB2 | PLAUR | 1.0018 | 1.5641 | 1.2829 | 0.000999 |
| TCD8em | MacRec | LGALS3 | CLEC7A | 0.2689 | 0.8554 | 0.5622 | 0.000999 |
| TCD8em | MacRec | LGALS3 | FCGR2A | 0.2689 | 0.8803 | 0.5746 | 0.000999 |
| TCD8em | MacRec | LGALS3 | FCGR2B | 0.2689 | 0.7132 | 0.4911 | 0.000999 |
| TCD8em | MacRec | LGALS3 | ITGB1 | 0.2689 | 0.4459 | 0.3574 | 0.000999 |
| TCD8em | MacRec | LGALS3 | TFRC | 0.2689 | 0.5511 | 0.41 | 0.000999 |
| TCD8em | MacRec | LRPAP1 | LRP1 | 0.2719 | 0.3912 | 0.3315 | 0.000999 |
| TCD8em | MacRec | LRPAP1 | SORL1 | 0.2719 | 0.5066 | 0.3893 | 0.000999 |
| TCD8em | MacRec | LTB | ITGB1 | 0.6352 | 0.4459 | 0.5405 | 0.000999 |

| | | | | | | | |
|---|---|---|---|---|---|---|---|
| TCD8em | MacRec | LTB | TNFRSF1A | 0.6352 | 0.4068 | 0.521 | 0.000999 |
| TCD8em | MacRec | MIF | CD74 | 0.6063 | 4.4668 | 2.5366 | 0.000999 |
| TCD8em | MacRec | PSAP | LRP1 | 0.5012 | 0.3912 | 0.4462 | 0.000999 |
| TCD8em | MacRec | PTPRC | CD4 | 1.7912 | 0.5246 | 1.1579 | 0.000999 |
| TCD8em | MacRec | PTPRC | CD44 | 1.7912 | 1.443 | 1.6171 | 0.000999 |
| TCD8em | MacRec | RPS19 | C5AR1 | 3.536 | 0.9166 | 2.2263 | 0.000999 |
| TCD8em | MacRec | RPS19 | CD4 | 3.536 | 0.5246 | 2.0303 | 0.000999 |
| TCD8em | MacRec | RTN4 | CD44 | 0.3632 | 1.443 | 0.9031 | 0.000999 |
| TCD8em | MacRec | SELL | CD4 | 0.258 | 0.5246 | 0.3913 | 0.000999 |
| TCD8em | MacRec | SPP1 | CD44 | 0.3608 | 1.443 | 0.9019 | 0.000999 |
| TCD8em | MacRec | TIGIT | NECTIN2 | 0.4665 | 0.2672 | 0.3669 | 0.000999 |
| TCD8em | MacRec | TIMP1 | LRP1 | 0.2667 | 0.3912 | 0.329 | 0.000999 |
| TCD8em | MacRec | TNF | RACK1 | 0.2667 | 2.6054 | 1.4361 | 0.000999 |
| TCD8em | MacRec | TNF | TNFRSF1A | 0.2667 | 0.4068 | 0.3368 | 0.000999 |
| TCD8em | MacRec | TNF | TNFRSF1B | 0.2667 | 0.5989 | 0.4328 | 0.000999 |
| TCD8em | MacRec | XCL2 | GPR65 | 0.295 | 0.3011 | 0.298 | 0.000999 |
| TCD8em | MacRec | XCL2 | LPAR6 | 0.295 | 0.2983 | 0.2966 | 0.000999 |
| TCD8ex | MacRec | AIMP1 | RACK1 | 0.2508 | 2.6054 | 1.4281 | 0.000999 |
| TCD8ex | MacRec | ALOX5AP | ALOX5 | 1.5024 | 0.4156 | 0.959 | 0.000999 |
| TCD8ex | MacRec | ANXA1 | C3AR1 | 0.5645 | 0.425 | 0.4947 | 0.000999 |
| TCD8ex | MacRec | ANXA1 | C5AR1 | 0.5645 | 0.9166 | 0.7406 | 0.000999 |
| TCD8ex | MacRec | ANXA1 | CD4 | 0.5645 | 0.5246 | 0.5446 | 0.000999 |
| TCD8ex | MacRec | ANXA1 | CD44 | 0.5645 | 1.443 | 1.0037 | 0.000999 |
| TCD8ex | MacRec | ANXA1 | FPR1 | 0.5645 | 0.729 | 0.6467 | 0.000999 |
| TCD8ex | MacRec | ANXA1 | FPR3 | 0.5645 | 0.5944 | 0.5794 | 0.000999 |
| TCD8ex | MacRec | APOE | LRP1 | 0.2814 | 0.3912 | 0.3363 | 0.000999 |
| TCD8ex | MacRec | APOE | SORL1 | 0.2814 | 0.5066 | 0.394 | 0.000999 |
| TCD8ex | MacRec | ARF1 | CD4 | 0.7835 | 0.5246 | 0.6541 | 0.000999 |
| TCD8ex | MacRec | ARF1 | CD44 | 0.7835 | 1.443 | 1.1133 | 0.000999 |
| TCD8ex | MacRec | CALM1 | TNFRSF1A | 1.4627 | 0.4068 | 0.9348 | 0.000999 |

| | | | | | | | |
|---|---|---|---|---|---|---|---|
| TCD8ex | MacRec | CALR | CD44 | 0.9634 | 1.443 | 1.2032 | 0.000999 |
| TCD8ex | MacRec | CALR | LRP1 | 0.9634 | 0.3912 | 0.6773 | 0.000999 |
| TCD8ex | MacRec | CD2 | CD4 | 2.0502 | 0.5246 | 1.2874 | 0.000999 |
| TCD8ex | MacRec | CD6 | ALCAM | 0.318 | 0.2586 | 0.2883 | 0.000999 |
| TCD8ex | MacRec | CRTAM | NECTIN2 | 0.3877 | 0.2672 | 0.3275 | 0.000999 |
| TCD8ex | MacRec | CXCL13 | C3AR1 | 1.7749 | 0.425 | 1.0999 | 0.000999 |
| TCD8ex | MacRec | CXCL13 | C5AR1 | 1.7749 | 0.9166 | 1.3458 | 0.000999 |
| TCD8ex | MacRec | CXCL13 | FPR1 | 1.7749 | 0.729 | 1.252 | 0.000999 |
| TCD8ex | MacRec | FASLG | TNFRSF1A | 0.2613 | 0.4068 | 0.334 | 0.000999 |
| TCD8ex | MacRec | GSTP1 | CD44 | 0.8286 | 1.443 | 1.1358 | 0.000999 |
| TCD8ex | MacRec | HLA-DMA | CD4 | 0.3991 | 0.5246 | 0.4619 | 0.000999 |
| TCD8ex | MacRec | HMGB1 | AGTRAP | 1.5414 | 0.3591 | 0.9503 | 0.000999 |
| TCD8ex | MacRec | HMGB1 | LY96 | 1.5414 | 0.8566 | 1.199 | 0.000999 |
| TCD8ex | MacRec | HMGB1 | TLR2 | 1.5414 | 0.3974 | 0.9694 | 0.000999 |
| TCD8ex | MacRec | HMGB2 | CD44 | 0.85 | 1.443 | 1.1465 | 0.000999 |
| TCD8ex | MacRec | HMGB2 | DAB2 | 0.85 | 0.3042 | 0.5771 | 0.000999 |
| TCD8ex | MacRec | HSP90B1 | ITGB1 | 0.5666 | 0.4459 | 0.5063 | 0.000999 |
| TCD8ex | MacRec | HSP90B1 | LRP1 | 0.5666 | 0.3912 | 0.4789 | 0.000999 |
| TCD8ex | MacRec | ICAM3 | ITGB2 | 0.5405 | 1.3879 | 0.9642 | 0.000999 |
| TCD8ex | MacRec | IFNG | IFNGR1 | 0.8575 | 0.7181 | 0.7878 | 0.000999 |
| TCD8ex | MacRec | IFNG | IFNGR2 | 0.8575 | 0.7916 | 0.8246 | 0.000999 |
| TCD8ex | MacRec | ITGA4 | CD44 | 0.4432 | 1.443 | 0.9431 | 0.000999 |
| TCD8ex | MacRec | ITGA4 | TFRC | 0.4432 | 0.5511 | 0.4972 | 0.000999 |
| TCD8ex | MacRec | ITGB1 | CD151 | 0.2995 | 0.2764 | 0.2879 | 0.000999 |
| TCD8ex | MacRec | ITGB1 | CD36 | 0.2995 | 0.3023 | 0.3009 | 0.000999 |
| TCD8ex | MacRec | ITGB1 | CD47 | 0.2995 | 0.5042 | 0.4018 | 0.000999 |
| TCD8ex | MacRec | ITGB1 | DAB2 | 0.2995 | 0.3042 | 0.3019 | 0.000999 |
| TCD8ex | MacRec | ITGB1 | ENG | 0.2995 | 0.3417 | 0.3206 | 0.000999 |
| TCD8ex | MacRec | ITGB1 | ITGAX | 0.2995 | 0.3629 | 0.3312 | 0.000999 |
| TCD8ex | MacRec | ITGB1 | PLAUR | 0.2995 | 1.5641 | 0.9318 | 0.000999 |

| | | | | | | | |
|---|---|---|---|---|---|---|---|
| TCD8ex | MacRec | ITGB1 | RACK1 | 0.2995 | 2.6054 | 1.4524 | 0.000999 |
| TCD8ex | MacRec | ITGB2 | ITGAM | 0.7635 | 0.3077 | 0.5356 | 0.000999 |
| TCD8ex | MacRec | ITGB2 | ITGAX | 0.7635 | 0.3629 | 0.5632 | 0.000999 |
| TCD8ex | MacRec | ITGB2 | PLAUR | 0.7635 | 1.5641 | 1.1638 | 0.000999 |
| TCD8ex | MacRec | LGALS3 | CLEC7A | 0.3824 | 0.8554 | 0.6189 | 0.000999 |
| TCD8ex | MacRec | LGALS3 | FCGR2A | 0.3824 | 0.8803 | 0.6313 | 0.000999 |
| TCD8ex | MacRec | LGALS3 | FCGR2B | 0.3824 | 0.7132 | 0.5478 | 0.000999 |
| TCD8ex | MacRec | LGALS3 | TFRC | 0.3824 | 0.5511 | 0.4668 | 0.000999 |
| TCD8ex | MacRec | LRPAP1 | LRP1 | 0.2971 | 0.3912 | 0.3442 | 0.000999 |
| TCD8ex | MacRec | LRPAP1 | SORL1 | 0.2971 | 0.5066 | 0.4019 | 0.000999 |
| TCD8ex | MacRec | LTB | ITGB1 | 0.4576 | 0.4459 | 0.4517 | 0.000999 |
| TCD8ex | MacRec | LTB | TNFRSF1A | 0.4576 | 0.4068 | 0.4322 | 0.000999 |
| TCD8ex | MacRec | MIF | CD74 | 0.749 | 4.4668 | 2.6079 | 0.000999 |
| TCD8ex | MacRec | PSAP | LRP1 | 0.5016 | 0.3912 | 0.4464 | 0.000999 |
| TCD8ex | MacRec | PTPRC | CD4 | 1.6762 | 0.5246 | 1.1004 | 0.000999 |
| TCD8ex | MacRec | PTPRC | CD44 | 1.6762 | 1.443 | 1.5596 | 0.000999 |
| TCD8ex | MacRec | RPS19 | C5AR1 | 3.8353 | 0.9166 | 2.376 | 0.000999 |
| TCD8ex | MacRec | RPS19 | CD4 | 3.8353 | 0.5246 | 2.18 | 0.000999 |
| TCD8ex | MacRec | RTN4 | CD44 | 0.3033 | 1.443 | 0.8731 | 0.000999 |
| TCD8ex | MacRec | SPP1 | CD44 | 0.2978 | 1.443 | 0.8704 | 0.000999 |
| TCD8ex | MacRec | TIGIT | NECTIN2 | 0.9854 | 0.2672 | 0.6263 | 0.000999 |
| TCD8ex | MacRec | VCAM1 | ITGB1 | 0.4462 | 0.4459 | 0.446 | 0.000999 |
| TCD8ex | MacRec | VCAM1 | ITGB2 | 0.4462 | 1.3879 | 0.9171 | 0.000999 |
| TCD8ex | MacRec | VCAM1 | RPSA | 0.4462 | 2.3088 | 1.3775 | 0.000999 |
| TCD8ex | MacRec | VCAM1 | TFRC | 0.4462 | 0.5511 | 0.4987 | 0.000999 |
| TCD8ex | MacRec | XCL2 | LPAR6 | 0.3157 | 0.2983 | 0.307 | 0.000999 |
| TCD8ex | MacRec | ITGA4 | RPSA | 0.4432 | 2.3088 | 1.376 | 0.002 |
| TCD8ex | MacRec | CALM1 | RPSA | 1.4627 | 2.3088 | 1.8858 | 0.00899 |
| TGD | MacRec | ARF1 | CD4 | 0.9453 | 0.5246 | 0.7349 | 0.000999 |
| TGD | MacRec | ARF1 | CD44 | 0.9453 | 1.443 | 1.1941 | 0.000999 |

| | | | | | | | |
|---|---|---|---|---|---|---|---|
| TGD | MacRec | CALM1 | RPSA | 1.7587 | 2.3088 | 2.0338 | 0.000999 |
| TGD | MacRec | CALM1 | TNFRSF1A | 1.7587 | 0.4068 | 1.0828 | 0.000999 |
| TGD | MacRec | CALM1 | TNFRSF1B | 1.7587 | 0.5989 | 1.1788 | 0.000999 |
| TGD | MacRec | CD2 | CD4 | 1.4522 | 0.5246 | 0.9884 | 0.000999 |
| TGD | MacRec | CD2 | PTPRC | 1.4522 | 0.8059 | 1.129 | 0.000999 |
| TGD | MacRec | CD6 | ALCAM | 0.4684 | 0.2586 | 0.3635 | 0.000999 |
| TGD | MacRec | CD99 | CD99 | 1.7154 | 0.6603 | 1.1879 | 0.000999 |
| TGD | MacRec | CRTAM | NECTIN2 | 0.2794 | 0.2672 | 0.2733 | 0.000999 |
| TGD | MacRec | HMGB1 | AGTRAP | 1.7525 | 0.3591 | 1.0558 | 0.000999 |
| TGD | MacRec | HMGB1 | LY96 | 1.7525 | 0.8566 | 1.3045 | 0.000999 |
| TGD | MacRec | HMGB1 | TLR2 | 1.7525 | 0.3974 | 1.075 | 0.000999 |
| TGD | MacRec | HSP90B1 | LRP1 | 0.9338 | 0.3912 | 0.6625 | 0.000999 |
| TGD | MacRec | HSP90B1 | TNFRSF1B | 0.9338 | 0.5989 | 0.7663 | 0.000999 |
| TGD | MacRec | ICAM3 | ITGB2 | 0.8816 | 1.3879 | 1.1348 | 0.000999 |
| TGD | MacRec | IL16 | CD4 | 0.9357 | 0.5246 | 0.7302 | 0.000999 |
| TGD | MacRec | IL16 | FCGR1A | 0.9357 | 0.603 | 0.7693 | 0.000999 |
| TGD | MacRec | ITGA4 | CD44 | 0.3968 | 1.443 | 0.9199 | 0.000999 |
| TGD | MacRec | ITGA4 | CXCR4 | 0.3968 | 1.6677 | 1.0323 | 0.000999 |
| TGD | MacRec | ITGA4 | RPSA | 0.3968 | 2.3088 | 1.3528 | 0.000999 |
| TGD | MacRec | ITGA4 | TFRC | 0.3968 | 0.5511 | 0.474 | 0.000999 |
| TGD | MacRec | ITGB1 | PLAUR | 0.3657 | 1.5641 | 0.9649 | 0.000999 |
| TGD | MacRec | ITGB2 | PLAUR | 1.4444 | 1.5641 | 1.5043 | 0.000999 |
| TGD | MacRec | LTB | ITGB1 | 1.6829 | 0.4459 | 1.0644 | 0.000999 |
| TGD | MacRec | LTB | TNFRSF1A | 1.6829 | 0.4068 | 1.0449 | 0.000999 |
| TGD | MacRec | MIF | CD74 | 1.6335 | 4.4668 | 3.0501 | 0.000999 |
| TGD | MacRec | MIF | TNFRSF14 | 1.6335 | 0.27 | 0.9517 | 0.000999 |
| TGD | MacRec | PTPRC | CD4 | 1.8653 | 0.5246 | 1.195 | 0.000999 |
| TGD | MacRec | PTPRC | CD44 | 1.8653 | 1.443 | 1.6542 | 0.000999 |
| TGD | MacRec | PTPRC | CXCR4 | 1.8653 | 1.6677 | 1.7665 | 0.000999 |
| TGD | MacRec | RPS19 | C5AR1 | 3.8902 | 0.9166 | 2.4034 | 0.000999 |

| | | | | | | | |
|---|---|---|---|---|---|---|---|
| TGD | MacRec | RPS19 | CD4 | 3.8902 | 0.5246 | 2.2074 | 0.000999 |
| TGD | MacRec | SELL | CD4 | 0.3555 | 0.5246 | 0.4401 | 0.000999 |
| TGD | MacRec | SEMA4D | PTPRC | 0.5658 | 0.8059 | 0.6858 | 0.000999 |
| TGD | MacRec | TGFB1 | ENG | 0.7703 | 0.3417 | 0.556 | 0.000999 |
| TGD | MacRec | TGFB1 | FCN1 | 0.7703 | 1.1325 | 0.9514 | 0.000999 |
| TGD | MacRec | TGFB1 | ITGB1 | 0.7703 | 0.4459 | 0.6081 | 0.000999 |
| TGD | MacRec | TGFB1 | TLR2 | 0.7703 | 0.3974 | 0.5839 | 0.000999 |
| TGD | MacRec | XCL1 | GPR65 | 1.5822 | 0.3011 | 0.9416 | 0.000999 |
| TGD | MacRec | XCL1 | LPAR6 | 1.5822 | 0.2983 | 0.9402 | 0.000999 |
| TGD | MacRec | XCL2 | GPR65 | 0.4864 | 0.3011 | 0.3938 | 0.000999 |
| TGD | MacRec | XCL2 | LPAR6 | 0.4864 | 0.2983 | 0.3924 | 0.000999 |
| TGD | MacRec | ICAM1 | ITGAX | 0.2915 | 0.3629 | 0.3272 | 0.00599 |
| TGD | MacRec | HSP90B1 | ITGB1 | 0.9338 | 0.4459 | 0.6899 | 0.00699 |

| Supplementary Table 8: Communication results between macrophages and lymphocytes subpopulations, and malignant cells | | | | | | | |
|---|---|---|---|---|---|---|---|
| **MacRec vs. malignant cells** | | | | | | | |
| celltype_01 | celltype_02 | ligand | receptor | ligand_mean_expression | receptor_mean_expression | ligand_receptor_mean_expression | pvalue |
| MacRec | breasttumor | AIMP1 | RRBP1 | 0.2827 | 0.2835 | 0.2831 | 0.000999 |
| MacRec | breasttumor | APOE | SCARB1 | 1.1155 | 0.2628 | 0.6891 | 0.000999 |
| MacRec | breasttumor | APOE | SDC2 | 1.1155 | 1.071 | 1.0933 | 0.000999 |
| MacRec | breasttumor | CALM1 | MYLK | 1.1815 | 0.4064 | 0.7939 | 0.000999 |
| MacRec | breasttumor | CALM1 | RPSA | 1.1815 | 2.358 | 1.7697 | 0.000999 |
| MacRec | breasttumor | CALM1 | TNFRSF1A | 1.1815 | 0.2536 | 0.7175 | 0.000999 |
| MacRec | breasttumor | CALM2 | MYLK | 1.8166 | 0.4064 | 1.1115 | 0.000999 |
| MacRec | breasttumor | CALM3 | MYLK | 0.6494 | 0.4064 | 0.5279 | 0.000999 |
| MacRec | breasttumor | CALR | MYLK | 1.0086 | 0.4064 | 0.7075 | 0.000999 |
| MacRec | breasttumor | CD99 | CD99 | 0.6596 | 0.3871 | 0.5234 | 0.000999 |
| MacRec | breasttumor | COL1A1 | ITGB1 | 0.5353 | 0.2939 | 0.4146 | 0.000999 |
| MacRec | breasttumor | FN1 | CD44 | 1.5033 | 0.3247 | 0.914 | 0.000999 |
| MacRec | breasttumor | FN1 | ITGB1 | 1.5033 | 0.2939 | 0.8986 | 0.000999 |
| MacRec | breasttumor | FN1 | RPSA | 1.5033 | 2.358 | 1.9306 | 0.000999 |
| MacRec | breasttumor | FN1 | SDC2 | 1.5033 | 1.071 | 1.2872 | 0.000999 |
| MacRec | breasttumor | GRN | TNFRSF1A | 1.7452 | 0.2536 | 0.9994 | 0.000999 |
| MacRec | breasttumor | HLA-DRA | SCARB1 | 4.3425 | 0.2628 | 2.3027 | 0.000999 |
| MacRec | breasttumor | ITGB1 | CD151 | 0.446 | 0.3569 | 0.4014 | 0.000999 |
| MacRec | breasttumor | ITGB1 | RACK1 | 0.446 | 2.6334 | 1.5397 | 0.000999 |
| MacRec | breasttumor | RTN4 | CD99 | 1.1012 | 0.3871 | 0.7441 | 0.000999 |
| MacRec | breasttumor | SERPINA1 | SDC2 | 1.5911 | 1.071 | 1.3311 | 0.000999 |
| MacRec | breasttumor | SPP1 | CD44 | 1.503 | 0.3247 | 0.9139 | 0.000999 |
| MacRec | breasttumor | SPP1 | ITGB1 | 1.503 | 0.2939 | 0.8984 | 0.000999 |
| MacRec | breasttumor | TGFB1 | ITGB1 | 0.3169 | 0.2939 | 0.3054 | 0.000999 |
| MacRec | breasttumor | TGFB1 | SDC2 | 0.3169 | 1.071 | 0.694 | 0.000999 |
| MacRec | breasttumor | THBS1 | ITGB1 | 0.7032 | 0.2939 | 0.4985 | 0.000999 |
| MacRec | breasttumor | TNFSF13B | ITGB1 | 0.8294 | 0.2939 | 0.5616 | 0.000999 |

| | | | | | | | |
|---|---|---|---|---|---|---|---|
| MacRec | breasttumor | RPS19 | RPSA | 3.7751 | 2.358 | 3.0666 | 0.002 |
| MacRec | breasttumor | AIMP1 | RACK1 | 0.2827 | 2.6334 | 1.458 | 0.00699 |
| MacRec | colorectaltumor | APOE | SDC1 | 1.1155 | 0.2575 | 0.6865 | 0.000999 |
| MacRec | colorectaltumor | COL1A1 | SDC1 | 0.5353 | 0.2575 | 0.3964 | 0.000999 |
| MacRec | colorectaltumor | FN1 | ITGB1 | 1.5033 | 0.4214 | 0.9623 | 0.000999 |
| MacRec | colorectaltumor | FN1 | SDC1 | 1.5033 | 0.2575 | 0.8804 | 0.000999 |
| MacRec | colorectaltumor | ITGB1 | CD151 | 0.446 | 0.6439 | 0.5449 | 0.000999 |
| MacRec | colorectaltumor | ITGB1 | CD46 | 0.446 | 0.3583 | 0.4021 | 0.000999 |
| MacRec | colorectaltumor | ITGB1 | CDH17 | 0.446 | 0.254 | 0.35 | 0.000999 |
| MacRec | colorectaltumor | ITGB1 | SDC1 | 0.446 | 0.2575 | 0.3517 | 0.000999 |
| MacRec | colorectaltumor | LGALS3 | ITGB4 | 0.9365 | 0.2913 | 0.6139 | 0.000999 |
| MacRec | colorectaltumor | MDK | SDC1 | 0.2959 | 0.2575 | 0.2767 | 0.000999 |
| MacRec | colorectaltumor | PECAM1 | SDC1 | 0.3002 | 0.2575 | 0.2788 | 0.000999 |
| MacRec | colorectaltumor | SPP1 | ITGB1 | 1.503 | 0.4214 | 0.9622 | 0.000999 |
| MacRec | colorectaltumor | SPP1 | SDC1 | 1.503 | 0.2575 | 0.8802 | 0.000999 |
| MacRec | colorectaltumor | THBS1 | SDC1 | 0.7032 | 0.2575 | 0.4803 | 0.000999 |
| MacRec | colorectaltumor | FN1 | RPSA | 1.5033 | 2.251 | 1.8772 | 0.002 |
| MacRec | colorectaltumor | TNFSF13B | ITGB1 | 0.8294 | 0.4214 | 0.6254 | 0.002 |
| MacRec | livertumor | ADM | RAMP1 | 0.3159 | 0.4499 | 0.3829 | 0.000999 |
| MacRec | livertumor | AIMP1 | RACK1 | 0.2827 | 2.7655 | 1.5241 | 0.000999 |
| MacRec | livertumor | AIMP1 | RRBP1 | 0.2827 | 0.355 | 0.3188 | 0.000999 |
| MacRec | livertumor | ANXA1 | MET | 1.2581 | 0.2983 | 0.7782 | 0.000999 |
| MacRec | livertumor | ANXA1 | PTPRF | 1.2581 | 0.384 | 0.8211 | 0.000999 |
| MacRec | livertumor | APOE | SDC1 | 1.1155 | 0.5354 | 0.8255 | 0.000999 |
| MacRec | livertumor | APOE | SDC4 | 1.1155 | 0.6068 | 0.8612 | 0.000999 |
| MacRec | livertumor | AREG | ERBB3 | 1.5637 | 0.2644 | 0.914 | 0.000999 |
| MacRec | livertumor | ARF1 | INSR | 0.7492 | 0.3039 | 0.5265 | 0.000999 |
| MacRec | livertumor | CALM1 | INSR | 1.1815 | 0.3039 | 0.7427 | 0.000999 |
| MacRec | livertumor | CALM1 | RPSA | 1.1815 | 2.6317 | 1.9066 | 0.000999 |
| MacRec | livertumor | CALR | ITGA3 | 1.0086 | 0.3913 | 0.7 | 0.000999 |

| | | | | | | | |
|---|---|---|---|---|---|---|---|
| MacRec | livertumor | CD99 | EPHB4 | 0.6596 | 0.4535 | 0.5565 | 0.000999 |
| MacRec | livertumor | CD99 | INSR | 0.6596 | 0.3039 | 0.4818 | 0.000999 |
| MacRec | livertumor | COL1A1 | ITGA2 | 0.5353 | 0.3983 | 0.4668 | 0.000999 |
| MacRec | livertumor | COL1A1 | ITGA3 | 0.5353 | 0.3913 | 0.4633 | 0.000999 |
| MacRec | livertumor | COL1A1 | ITGB1 | 0.5353 | 1.6044 | 1.0699 | 0.000999 |
| MacRec | livertumor | COL1A1 | SDC1 | 0.5353 | 0.5354 | 0.5354 | 0.000999 |
| MacRec | livertumor | EREG | ERBB3 | 0.6565 | 0.2644 | 0.4604 | 0.000999 |
| MacRec | livertumor | FN1 | ITGA2 | 1.5033 | 0.3983 | 0.9508 | 0.000999 |
| MacRec | livertumor | FN1 | ITGA3 | 1.5033 | 0.3913 | 0.9473 | 0.000999 |
| MacRec | livertumor | FN1 | ITGB1 | 1.5033 | 1.6044 | 1.5539 | 0.000999 |
| MacRec | livertumor | FN1 | RPSA | 1.5033 | 2.6317 | 2.0675 | 0.000999 |
| MacRec | livertumor | FN1 | SDC1 | 1.5033 | 0.5354 | 1.0194 | 0.000999 |
| MacRec | livertumor | HBEGF | ERBB3 | 0.7403 | 0.2644 | 0.5023 | 0.000999 |
| MacRec | livertumor | HLA-A | ERBB3 | 2.802 | 0.2644 | 1.5332 | 0.000999 |
| MacRec | livertumor | HLA-DRA | EPHB4 | 4.3425 | 0.4535 | 2.398 | 0.000999 |
| MacRec | livertumor | HLA-E | EPHB4 | 1.5621 | 0.4535 | 1.0078 | 0.000999 |
| MacRec | livertumor | HLA-E | ITGA6 | 1.5621 | 0.5211 | 1.0416 | 0.000999 |
| MacRec | livertumor | HLA-E | PLXNB2 | 1.5621 | 0.265 | 0.9135 | 0.000999 |
| MacRec | livertumor | HMGB2 | CDH1 | 0.756 | 0.4896 | 0.6228 | 0.000999 |
| MacRec | livertumor | HMGB2 | DSG2 | 0.756 | 0.7479 | 0.752 | 0.000999 |
| MacRec | livertumor | HSP90B1 | ITGB1 | 0.7853 | 1.6044 | 1.1949 | 0.000999 |
| MacRec | livertumor | ITGB1 | CD151 | 0.446 | 1.1085 | 0.7772 | 0.000999 |
| MacRec | livertumor | ITGB1 | ITGA2 | 0.446 | 0.3983 | 0.4221 | 0.000999 |
| MacRec | livertumor | ITGB1 | ITGA6 | 0.446 | 0.5211 | 0.4835 | 0.000999 |
| MacRec | livertumor | ITGB1 | SDC1 | 0.446 | 0.5354 | 0.4907 | 0.000999 |
| MacRec | livertumor | ITGB1 | VTN | 0.446 | 0.7399 | 0.5929 | 0.000999 |
| MacRec | livertumor | LGALS3 | ITGA2 | 0.9365 | 0.3983 | 0.6674 | 0.000999 |
| MacRec | livertumor | LGALS3 | ITGB1 | 0.9365 | 1.6044 | 1.2705 | 0.000999 |
| MacRec | livertumor | LGALS3 | ITGB4 | 0.9365 | 0.5518 | 0.7441 | 0.000999 |
| MacRec | livertumor | MDK | SDC1 | 0.2959 | 0.5354 | 0.4157 | 0.000999 |

| | | | | | | | |
|---|---|---|---|---|---|---|---|
| MacRec | livertumor | NAMPT | INSR | 1.5182 | 0.3039 | 0.9111 | 0.000999 |
| MacRec | livertumor | NECTIN2 | TNFRSF12A | 0.2668 | 1.0484 | 0.6576 | 0.000999 |
| MacRec | livertumor | OSM | IL6ST | 0.2995 | 0.2743 | 0.2869 | 0.000999 |
| MacRec | livertumor | PECAM1 | SDC1 | 0.3002 | 0.5354 | 0.4178 | 0.000999 |
| MacRec | livertumor | PECAM1 | SDC4 | 0.3002 | 0.6068 | 0.4535 | 0.000999 |
| MacRec | livertumor | PTPRC | INSR | 0.8055 | 0.3039 | 0.5547 | 0.000999 |
| MacRec | livertumor | PTPRC | MET | 0.8055 | 0.2983 | 0.5519 | 0.000999 |
| MacRec | livertumor | RPS19 | RPSA | 3.7751 | 2.6317 | 3.2034 | 0.000999 |
| MacRec | livertumor | SPP1 | ITGB1 | 1.503 | 1.6044 | 1.5537 | 0.000999 |
| MacRec | livertumor | SPP1 | SDC1 | 1.503 | 0.5354 | 1.0192 | 0.000999 |
| MacRec | livertumor | TGFB1 | ITGB1 | 0.3169 | 1.6044 | 0.9607 | 0.000999 |
| MacRec | livertumor | TGFB1 | VTN | 0.3169 | 0.7399 | 0.5284 | 0.000999 |
| MacRec | livertumor | THBS1 | CD47 | 0.7032 | 0.5289 | 0.616 | 0.000999 |
| MacRec | livertumor | THBS1 | ITGA3 | 0.7032 | 0.3913 | 0.5473 | 0.000999 |
| MacRec | livertumor | THBS1 | ITGB1 | 0.7032 | 1.6044 | 1.1538 | 0.000999 |
| MacRec | livertumor | THBS1 | SDC1 | 0.7032 | 0.5354 | 0.6193 | 0.000999 |
| MacRec | livertumor | TNFSF13B | ITGB1 | 0.8294 | 1.6044 | 1.2169 | 0.000999 |
| MacRec | livertumor | VEGFA | AXL | 0.6076 | 0.3099 | 0.4588 | 0.000999 |
| MacRec | livertumor | VEGFA | VTN | 0.6076 | 0.7399 | 0.6737 | 0.000999 |
| MacRec | livertumor | A2M | TNFRSF14 | 0.2824 | 0.2788 | 0.2806 | 0.00699 |
| MacRec | livertumor | MDK | SDC4 | 0.2959 | 0.6068 | 0.4514 | 0.00799 |
| MacRec | livertumor | ITGB1 | ITGA3 | 0.446 | 0.3913 | 0.4186 | 0.00899 |
| MacRec | lungtumor | APOE | SDC1 | 1.1155 | 0.2715 | 0.6935 | 0.000999 |
| MacRec | lungtumor | APOE | SDC4 | 1.1155 | 0.3516 | 0.7336 | 0.000999 |
| MacRec | lungtumor | COL1A1 | SDC1 | 0.5353 | 0.2715 | 0.4034 | 0.000999 |
| MacRec | lungtumor | CXCL16 | CXCR4 | 0.8331 | 0.2684 | 0.5507 | 0.000999 |
| MacRec | lungtumor | CXCL2 | CXCR4 | 0.8241 | 0.2684 | 0.5462 | 0.000999 |
| MacRec | lungtumor | FN1 | CD44 | 1.5033 | 0.3802 | 0.9417 | 0.000999 |
| MacRec | lungtumor | FN1 | RPSA | 1.5033 | 1.6057 | 1.5545 | 0.000999 |
| MacRec | lungtumor | FN1 | SDC1 | 1.5033 | 0.2715 | 0.8874 | 0.000999 |

| | | | | | | | |
|---|---|---|---|---|---|---|---|
| MacRec | lungtumor | HBEGF | CD44 | 0.7403 | 0.3802 | 0.5602 | 0.000999 |
| MacRec | lungtumor | ITGB1 | CD151 | 0.446 | 0.6776 | 0.5618 | 0.000999 |
| MacRec | lungtumor | ITGB1 | CD46 | 0.446 | 0.434 | 0.44 | 0.000999 |
| MacRec | lungtumor | ITGB1 | CD47 | 0.446 | 0.3255 | 0.3857 | 0.000999 |
| MacRec | lungtumor | ITGB1 | SDC1 | 0.446 | 0.2715 | 0.3587 | 0.000999 |
| MacRec | lungtumor | NECTIN2 | TNFRSF12A | 0.2668 | 0.441 | 0.3539 | 0.000999 |
| MacRec | lungtumor | PECAM1 | SDC1 | 0.3002 | 0.2715 | 0.2858 | 0.000999 |
| MacRec | lungtumor | PECAM1 | SDC4 | 0.3002 | 0.3516 | 0.3259 | 0.000999 |
| MacRec | lungtumor | PTPRC | CD44 | 0.8055 | 0.3802 | 0.5928 | 0.000999 |
| MacRec | lungtumor | PTPRC | CXCR4 | 0.8055 | 0.2684 | 0.5369 | 0.000999 |
| MacRec | lungtumor | RTN4 | CD44 | 1.1012 | 0.3802 | 0.7407 | 0.000999 |
| MacRec | lungtumor | RTN4 | CD99 | 1.1012 | 0.5938 | 0.8475 | 0.000999 |
| MacRec | lungtumor | THBS1 | CD47 | 0.7032 | 0.3255 | 0.5144 | 0.000999 |
| MacRec | lungtumor | THBS1 | SDC1 | 0.7032 | 0.2715 | 0.4874 | 0.000999 |
| MacRec | melanomatumo | A2M | TNFRSF14 | 0.2824 | 0.5069 | 0.3946 | 0.000999 |
| MacRec | melanomatumo | AIMP1 | RACK1 | 0.2827 | 4.145 | 2.2139 | 0.000999 |
| MacRec | melanomatumo | ANXA1 | EDNRB | 1.2581 | 0.5666 | 0.9124 | 0.000999 |
| MacRec | melanomatumo | CALM1 | RPSA | 1.1815 | 3.7787 | 2.4801 | 0.000999 |
| MacRec | melanomatumo | FN1 | RPSA | 1.5033 | 3.7787 | 2.641 | 0.000999 |
| MacRec | melanomatumo | ITGB1 | RACK1 | 0.446 | 4.145 | 2.2955 | 0.000999 |
| MacRec | melanomatumo | RPS19 | RPSA | 3.7751 | 3.7787 | 3.7769 | 0.000999 |
| MacRec | melanomatumo | NAMPT | ANXA6 | 1.5182 | 0.3625 | 0.9404 | 0.002 |
| MacRec | melanomatumo | SPP1 | NCSTN | 1.503 | 0.2614 | 0.8822 | 0.005 |
| MacRec | melanomatumo | CALR | NCSTN | 1.0086 | 0.2614 | 0.635 | 0.00899 |
| MacRec | ovarytumor | AIMP1 | RACK1 | 0.2827 | 2.9543 | 1.6185 | 0.000999 |
| MacRec | ovarytumor | AREG | FOLR1 | 1.5637 | 0.5619 | 1.0628 | 0.000999 |
| MacRec | ovarytumor | CALM1 | RPSA | 1.1815 | 2.9542 | 2.0678 | 0.000999 |
| MacRec | ovarytumor | COL1A1 | ITGB1 | 0.5353 | 0.2783 | 0.4068 | 0.000999 |
| MacRec | ovarytumor | CXCL16 | CXCR4 | 0.8331 | 0.3688 | 0.6009 | 0.000999 |
| MacRec | ovarytumor | CXCL2 | CXCR4 | 0.8241 | 0.3688 | 0.5964 | 0.000999 |

| | | | | | | | |
|---|---|---|---|---|---|---|---|
| MacRec | ovarytumor | CXCL3 | CXCR4 | 0.5694 | 0.3688 | 0.4691 | 0.000999 |
| MacRec | ovarytumor | FN1 | ITGB1 | 1.5033 | 0.2783 | 0.8908 | 0.000999 |
| MacRec | ovarytumor | FN1 | RPSA | 1.5033 | 2.9542 | 2.2287 | 0.000999 |
| MacRec | ovarytumor | HSP90B1 | ITGB1 | 0.7853 | 0.2783 | 0.5318 | 0.000999 |
| MacRec | ovarytumor | ITGB1 | CD151 | 0.446 | 0.4981 | 0.472 | 0.000999 |
| MacRec | ovarytumor | ITGB1 | CD47 | 0.446 | 0.4931 | 0.4695 | 0.000999 |
| MacRec | ovarytumor | ITGB1 | RACK1 | 0.446 | 2.9543 | 1.7001 | 0.000999 |
| MacRec | ovarytumor | LGALS3 | ITGB1 | 0.9365 | 0.2783 | 0.6074 | 0.000999 |
| MacRec | ovarytumor | NECTIN2 | TNFRSF12A | 0.2668 | 0.3901 | 0.3285 | 0.000999 |
| MacRec | ovarytumor | PTPRC | CXCR4 | 0.8055 | 0.3688 | 0.5871 | 0.000999 |
| MacRec | ovarytumor | SPP1 | ITGB1 | 1.503 | 0.2783 | 0.8906 | 0.000999 |
| MacRec | ovarytumor | TGFB1 | ITGB1 | 0.3169 | 0.2783 | 0.2976 | 0.000999 |
| MacRec | ovarytumor | THBS1 | CD47 | 0.7032 | 0.4931 | 0.5981 | 0.000999 |
| MacRec | ovarytumor | THBS1 | ITGB1 | 0.7032 | 0.2783 | 0.4907 | 0.000999 |
| MacRec | ovarytumor | TNFSF13B | ITGB1 | 0.8294 | 0.2783 | 0.5538 | 0.000999 |
| breasttumor | MacRec | A2M | LRP1 | 0.3129 | 0.3918 | 0.3524 | 0.000999 |
| breasttumor | MacRec | A2M | TNFRSF14 | 0.3129 | 0.2703 | 0.2916 | 0.000999 |
| breasttumor | MacRec | ANXA1 | CD44 | 0.4092 | 1.4429 | 0.926 | 0.000999 |
| breasttumor | MacRec | ANXA1 | CXCR4 | 0.4092 | 1.6682 | 1.0387 | 0.000999 |
| breasttumor | MacRec | APP | ALOX5 | 0.5409 | 0.4148 | 0.4779 | 0.000999 |
| breasttumor | MacRec | APP | C3AR1 | 0.5409 | 0.4251 | 0.483 | 0.000999 |
| breasttumor | MacRec | APP | C5AR1 | 0.5409 | 0.9163 | 0.7286 | 0.000999 |
| breasttumor | MacRec | APP | CD36 | 0.5409 | 0.3022 | 0.4216 | 0.000999 |
| breasttumor | MacRec | APP | CXCR4 | 0.5409 | 1.6682 | 1.1046 | 0.000999 |
| breasttumor | MacRec | APP | FPR1 | 0.5409 | 0.7287 | 0.6348 | 0.000999 |
| breasttumor | MacRec | APP | LRP1 | 0.5409 | 0.3918 | 0.4664 | 0.000999 |
| breasttumor | MacRec | APP | SORL1 | 0.5409 | 0.5063 | 0.5236 | 0.000999 |
| breasttumor | MacRec | APP | TNFRSF14 | 0.5409 | 0.2703 | 0.4056 | 0.000999 |
| breasttumor | MacRec | ARF1 | CD4 | 1.6219 | 0.5254 | 1.0736 | 0.000999 |
| breasttumor | MacRec | ARF1 | CD44 | 1.6219 | 1.4429 | 1.5324 | 0.000999 |

| | | | | | | | |
|---|---|---|---|---|---|---|---|
| breasttumor | MacRec | CALM1 | TNFRSF1B | 0.9015 | 0.5988 | 0.7502 | 0.000999 |
| breasttumor | MacRec | CALR | CD44 | 1.1835 | 1.4429 | 1.3132 | 0.000999 |
| breasttumor | MacRec | CALR | LRP1 | 1.1835 | 0.3918 | 0.7877 | 0.000999 |
| breasttumor | MacRec | CD99 | CD99 | 0.3871 | 0.6596 | 0.5234 | 0.000999 |
| breasttumor | MacRec | COL2A1 | CD44 | 0.3935 | 1.4429 | 0.9182 | 0.000999 |
| breasttumor | MacRec | COL2A1 | ITGB1 | 0.3935 | 0.446 | 0.4197 | 0.000999 |
| breasttumor | MacRec | GSTP1 | CD44 | 2.5435 | 1.4429 | 1.9932 | 0.000999 |
| breasttumor | MacRec | HEBP1 | C3AR1 | 0.4062 | 0.4251 | 0.4156 | 0.000999 |
| breasttumor | MacRec | HEBP1 | C5AR1 | 0.4062 | 0.9163 | 0.6613 | 0.000999 |
| breasttumor | MacRec | HEBP1 | CXCR4 | 0.4062 | 1.6682 | 1.0372 | 0.000999 |
| breasttumor | MacRec | HEBP1 | FPR1 | 0.4062 | 0.7287 | 0.5674 | 0.000999 |
| breasttumor | MacRec | HEBP1 | FPR3 | 0.4062 | 0.5947 | 0.5004 | 0.000999 |
| breasttumor | MacRec | HMGB1 | LY96 | 1.1815 | 0.8558 | 1.0186 | 0.000999 |
| breasttumor | MacRec | HMGB1 | TLR2 | 1.1815 | 0.398 | 0.7897 | 0.000999 |
| breasttumor | MacRec | HMGB2 | CD44 | 0.3139 | 1.4429 | 0.8784 | 0.000999 |
| breasttumor | MacRec | HSP90B1 | ITGB1 | 0.7786 | 0.446 | 0.6123 | 0.000999 |
| breasttumor | MacRec | HSP90B1 | LRP1 | 0.7786 | 0.3918 | 0.5852 | 0.000999 |
| breasttumor | MacRec | HSP90B1 | TNFRSF1B | 0.7786 | 0.5988 | 0.6887 | 0.000999 |
| breasttumor | MacRec | ITGB1 | CD36 | 0.2939 | 0.3022 | 0.298 | 0.000999 |
| breasttumor | MacRec | ITGB1 | CD47 | 0.2939 | 0.503 | 0.3985 | 0.000999 |
| breasttumor | MacRec | ITGB1 | DAB2 | 0.2939 | 0.3033 | 0.2986 | 0.000999 |
| breasttumor | MacRec | ITGB1 | ENG | 0.2939 | 0.3416 | 0.3177 | 0.000999 |
| breasttumor | MacRec | ITGB1 | ITGAX | 0.2939 | 0.3634 | 0.3286 | 0.000999 |
| breasttumor | MacRec | ITGB1 | PLAUR | 0.2939 | 1.5635 | 0.9287 | 0.000999 |
| breasttumor | MacRec | LGALS3 | CLEC7A | 0.8391 | 0.8551 | 0.8471 | 0.000999 |
| breasttumor | MacRec | LGALS3 | FCGR2A | 0.8391 | 0.8792 | 0.8592 | 0.000999 |
| breasttumor | MacRec | LGALS3 | FCGR2B | 0.8391 | 0.7138 | 0.7764 | 0.000999 |
| breasttumor | MacRec | LGALS3 | TFRC | 0.8391 | 0.5506 | 0.6948 | 0.000999 |
| breasttumor | MacRec | LRPAP1 | LRP1 | 0.2756 | 0.3918 | 0.3337 | 0.000999 |
| breasttumor | MacRec | LRPAP1 | SORL1 | 0.2756 | 0.5063 | 0.391 | 0.000999 |

| | | | | | | | |
|---|---|---|---|---|---|---|---|
| breasttumor | MacRec | MDK | LRP1 | 0.592 | 0.3918 | 0.4919 | 0.000999 |
| breasttumor | MacRec | MIF | CD74 | 3.2716 | 4.4657 | 3.8687 | 0.000999 |
| breasttumor | MacRec | MIF | TNFRSF14 | 3.2716 | 0.2703 | 1.771 | 0.000999 |
| breasttumor | MacRec | NPW | C3AR1 | 0.5478 | 0.4251 | 0.4865 | 0.000999 |
| breasttumor | MacRec | NPW | C5AR1 | 0.5478 | 0.9163 | 0.7321 | 0.000999 |
| breasttumor | MacRec | NPW | CXCR4 | 0.5478 | 1.6682 | 1.108 | 0.000999 |
| breasttumor | MacRec | NPW | FPR1 | 0.5478 | 0.7287 | 0.6383 | 0.000999 |
| breasttumor | MacRec | RPS19 | C5AR1 | 3.691 | 0.9163 | 2.3037 | 0.000999 |
| breasttumor | MacRec | RPS19 | CD4 | 3.691 | 0.5254 | 2.1082 | 0.000999 |
| breasttumor | MacRec | RTN4 | CD44 | 0.3318 | 1.4429 | 0.8873 | 0.000999 |
| breasttumor | MacRec | SERPINE2 | LRP1 | 0.6001 | 0.3918 | 0.496 | 0.000999 |
| breasttumor | MacRec | TF | CD4 | 0.2688 | 0.5254 | 0.3971 | 0.000999 |
| breasttumor | MacRec | TF | TFRC | 0.2688 | 0.5506 | 0.4097 | 0.000999 |
| breasttumor | MacRec | ANXA1 | C5AR1 | 0.4092 | 0.9163 | 0.6628 | 0.002 |
| colorectaltumor | MacRec | AIMP1 | RACK1 | 0.5265 | 2.605 | 1.5658 | 0.000999 |
| colorectaltumor | MacRec | APP | ALOX5 | 0.4966 | 0.4148 | 0.4557 | 0.000999 |
| colorectaltumor | MacRec | APP | C3AR1 | 0.4966 | 0.4251 | 0.4609 | 0.000999 |
| colorectaltumor | MacRec | APP | C5AR1 | 0.4966 | 0.9163 | 0.7065 | 0.000999 |
| colorectaltumor | MacRec | APP | CD36 | 0.4966 | 0.3022 | 0.3994 | 0.000999 |
| colorectaltumor | MacRec | APP | CXCR4 | 0.4966 | 1.6682 | 1.0824 | 0.000999 |
| colorectaltumor | MacRec | APP | FPR1 | 0.4966 | 0.7287 | 0.6127 | 0.000999 |
| colorectaltumor | MacRec | APP | LRP1 | 0.4966 | 0.3918 | 0.4442 | 0.000999 |
| colorectaltumor | MacRec | APP | SORL1 | 0.4966 | 0.5063 | 0.5015 | 0.000999 |
| colorectaltumor | MacRec | APP | TNFRSF14 | 0.4966 | 0.2703 | 0.3835 | 0.000999 |
| colorectaltumor | MacRec | GSTP1 | CD44 | 2.8039 | 1.4429 | 2.1234 | 0.000999 |
| colorectaltumor | MacRec | HMGB1 | AGTRAP | 2.155 | 0.3586 | 1.2568 | 0.000999 |
| colorectaltumor | MacRec | HMGB1 | LY96 | 2.155 | 0.8558 | 1.5054 | 0.000999 |
| colorectaltumor | MacRec | HMGB1 | TLR2 | 2.155 | 0.398 | 1.2765 | 0.000999 |
| colorectaltumor | MacRec | ITGB1 | PLAUR | 0.4214 | 1.5635 | 0.9924 | 0.000999 |
| colorectaltumor | MacRec | LGALS3 | CLEC7A | 1.5563 | 0.8551 | 1.2057 | 0.000999 |

| | | | | | | | |
|---|---|---|---|---|---|---|---|
| colorectaltumor | MacRec | LGALS3 | FCGR2A | 1.5563 | 0.8792 | 1.2178 | 0.000999 |
| colorectaltumor | MacRec | LGALS3 | FCGR2B | 1.5563 | 0.7138 | 1.135 | 0.000999 |
| colorectaltumor | MacRec | LGALS3 | ITGB1 | 1.5563 | 0.446 | 1.0011 | 0.000999 |
| colorectaltumor | MacRec | LGALS3 | TFRC | 1.5563 | 0.5506 | 1.0534 | 0.000999 |
| colorectaltumor | MacRec | MDK | LRP1 | 0.7548 | 0.3918 | 0.5733 | 0.000999 |
| colorectaltumor | MacRec | MIF | CD74 | 1.6741 | 4.4657 | 3.0699 | 0.000999 |
| colorectaltumor | MacRec | MIF | TNFRSF14 | 1.6741 | 0.2703 | 0.9722 | 0.000999 |
| colorectaltumor | MacRec | RPS19 | C5AR1 | 4.0956 | 0.9163 | 2.506 | 0.000999 |
| colorectaltumor | MacRec | RPS19 | CD4 | 4.0956 | 0.5254 | 2.3105 | 0.000999 |
| colorectaltumor | MacRec | RPS19 | RPSA | 4.0956 | 2.3085 | 3.2021 | 0.000999 |
| colorectaltumor | MacRec | COL1A1 | CD44 | 0.4875 | 1.4429 | 0.9652 | 0.003 |
| colorectaltumor | MacRec | ARF1 | CD44 | 0.6489 | 1.4429 | 1.0459 | 0.004 |
| livertumor | MacRec | AGRN | LRP1 | 0.3492 | 0.3918 | 0.3705 | 0.000999 |
| livertumor | MacRec | AGT | ANPEP | 0.4701 | 0.3749 | 0.4225 | 0.000999 |
| livertumor | MacRec | AGT | ATP6AP2 | 0.4701 | 0.7304 | 0.6002 | 0.000999 |
| livertumor | MacRec | AGT | C3AR1 | 0.4701 | 0.4251 | 0.4476 | 0.000999 |
| livertumor | MacRec | AGT | C5AR1 | 0.4701 | 0.9163 | 0.6932 | 0.000999 |
| livertumor | MacRec | AGT | CXCR4 | 0.4701 | 1.6682 | 1.0691 | 0.000999 |
| livertumor | MacRec | AGT | FPR1 | 0.4701 | 0.7287 | 0.5994 | 0.000999 |
| livertumor | MacRec | AGT | GPR65 | 0.4701 | 0.3015 | 0.3858 | 0.000999 |
| livertumor | MacRec | AGT | LPAR6 | 0.4701 | 0.2979 | 0.384 | 0.000999 |
| livertumor | MacRec | APOA1 | CD36 | 0.2888 | 0.3022 | 0.2955 | 0.000999 |
| livertumor | MacRec | APOA1 | LRP1 | 0.2888 | 0.3918 | 0.3403 | 0.000999 |
| livertumor | MacRec | APOA1 | MSR1 | 0.2888 | 0.4062 | 0.3475 | 0.000999 |
| livertumor | MacRec | APP | ALOX5 | 0.5626 | 0.4148 | 0.4887 | 0.000999 |
| livertumor | MacRec | APP | C3AR1 | 0.5626 | 0.4251 | 0.4938 | 0.000999 |
| livertumor | MacRec | APP | C5AR1 | 0.5626 | 0.9163 | 0.7395 | 0.000999 |
| livertumor | MacRec | APP | CD36 | 0.5626 | 0.3022 | 0.4324 | 0.000999 |
| livertumor | MacRec | APP | CXCR4 | 0.5626 | 1.6682 | 1.1154 | 0.000999 |
| livertumor | MacRec | APP | FPR1 | 0.5626 | 0.7287 | 0.6456 | 0.000999 |

| | | | | | | | |
|---|---|---|---|---|---|---|---|
| livertumor | MacRec | APP | LRP1 | 0.5626 | 0.3918 | 0.4772 | 0.000999 |
| livertumor | MacRec | APP | SORL1 | 0.5626 | 0.5063 | 0.5344 | 0.000999 |
| livertumor | MacRec | APP | TNFRSF14 | 0.5626 | 0.2703 | 0.4164 | 0.000999 |
| livertumor | MacRec | C3 | C3AR1 | 0.6584 | 0.4251 | 0.5418 | 0.000999 |
| livertumor | MacRec | C3 | C5AR1 | 0.6584 | 0.9163 | 0.7874 | 0.000999 |
| livertumor | MacRec | C3 | CXCR4 | 0.6584 | 1.6682 | 1.1633 | 0.000999 |
| livertumor | MacRec | C3 | FPR1 | 0.6584 | 0.7287 | 0.6936 | 0.000999 |
| livertumor | MacRec | C3 | ITGAM | 0.6584 | 0.3076 | 0.483 | 0.000999 |
| livertumor | MacRec | C3 | ITGAX | 0.6584 | 0.3634 | 0.5109 | 0.000999 |
| livertumor | MacRec | C3 | ITGB2 | 0.6584 | 1.3869 | 1.0227 | 0.000999 |
| livertumor | MacRec | C3 | LRP1 | 0.6584 | 0.3918 | 0.5251 | 0.000999 |
| livertumor | MacRec | CALR | CD44 | 1.0416 | 1.4429 | 1.2422 | 0.000999 |
| livertumor | MacRec | CCL20 | C3AR1 | 1.0877 | 0.4251 | 0.7564 | 0.000999 |
| livertumor | MacRec | CCL20 | C5AR1 | 1.0877 | 0.9163 | 1.002 | 0.000999 |
| livertumor | MacRec | CCL20 | CXCR4 | 1.0877 | 1.6682 | 1.3779 | 0.000999 |
| livertumor | MacRec | CCL20 | FPR1 | 1.0877 | 0.7287 | 0.9082 | 0.000999 |
| livertumor | MacRec | CDH1 | RACK1 | 0.4896 | 2.605 | 1.5473 | 0.000999 |
| livertumor | MacRec | CFH | ITGAM | 0.3373 | 0.3076 | 0.3224 | 0.000999 |
| livertumor | MacRec | DSG2 | CD44 | 0.7479 | 1.4429 | 1.0954 | 0.000999 |
| livertumor | MacRec | DSG2 | CXCR4 | 0.7479 | 1.6682 | 1.2081 | 0.000999 |
| livertumor | MacRec | DSG2 | DAB2 | 0.7479 | 0.3033 | 0.5256 | 0.000999 |
| livertumor | MacRec | FGA | ITGAX | 0.2688 | 0.3634 | 0.3161 | 0.000999 |
| livertumor | MacRec | FGA | ITGB2 | 0.2688 | 1.3869 | 0.8278 | 0.000999 |
| livertumor | MacRec | HMGB1 | AGTRAP | 1.4361 | 0.3586 | 0.8973 | 0.000999 |
| livertumor | MacRec | HMGB1 | LY96 | 1.4361 | 0.8558 | 1.1459 | 0.000999 |
| livertumor | MacRec | HMGB1 | TLR2 | 1.4361 | 0.398 | 0.917 | 0.000999 |
| livertumor | MacRec | HSP90B1 | LRP1 | 0.8951 | 0.3918 | 0.6435 | 0.000999 |
| livertumor | MacRec | HSP90B1 | TNFRSF1B | 0.8951 | 0.5988 | 0.747 | 0.000999 |
| livertumor | MacRec | HSPG2 | LRP1 | 0.2746 | 0.3918 | 0.3332 | 0.000999 |
| livertumor | MacRec | ITGB1 | CD151 | 1.6044 | 0.2751 | 0.9398 | 0.000999 |

| | | | | | | | |
|---|---|---|---|---|---|---|---|
| livertumor | MacRec | ITGB1 | CD36 | 1.6044 | 0.3022 | 0.9533 | 0.000999 |
| livertumor | MacRec | ITGB1 | CD47 | 1.6044 | 0.503 | 1.0537 | 0.000999 |
| livertumor | MacRec | ITGB1 | DAB2 | 1.6044 | 0.3033 | 0.9539 | 0.000999 |
| livertumor | MacRec | ITGB1 | ENG | 1.6044 | 0.3416 | 0.973 | 0.000999 |
| livertumor | MacRec | ITGB1 | ITGAX | 1.6044 | 0.3634 | 0.9839 | 0.000999 |
| livertumor | MacRec | ITGB1 | PLAUR | 1.6044 | 1.5635 | 1.584 | 0.000999 |
| livertumor | MacRec | ITGB1 | RACK1 | 1.6044 | 2.605 | 2.1047 | 0.000999 |
| livertumor | MacRec | LAMC2 | CD151 | 0.4847 | 0.2751 | 0.3799 | 0.000999 |
| livertumor | MacRec | LAMC2 | ITGB1 | 0.4847 | 0.446 | 0.4653 | 0.000999 |
| livertumor | MacRec | MDK | LRP1 | 1.6101 | 0.3918 | 1.001 | 0.000999 |
| livertumor | MacRec | MIF | CD74 | 1.7638 | 4.4657 | 3.1147 | 0.000999 |
| livertumor | MacRec | MIF | TNFRSF14 | 1.7638 | 0.2703 | 1.0171 | 0.000999 |
| livertumor | MacRec | NTS | GPR65 | 0.3425 | 0.3015 | 0.322 | 0.000999 |
| livertumor | MacRec | NTS | LPAR6 | 0.3425 | 0.2979 | 0.3202 | 0.000999 |
| livertumor | MacRec | PTPRF | RACK1 | 0.384 | 2.605 | 1.4945 | 0.000999 |
| livertumor | MacRec | RPS19 | C5AR1 | 4.0772 | 0.9163 | 2.4968 | 0.000999 |
| livertumor | MacRec | RPS19 | CD4 | 4.0772 | 0.5254 | 2.3013 | 0.000999 |
| livertumor | MacRec | RPS19 | RPSA | 4.0772 | 2.3085 | 3.1929 | 0.000999 |
| livertumor | MacRec | TF | CD4 | 0.523 | 0.5254 | 0.5242 | 0.000999 |
| livertumor | MacRec | TF | TFRC | 0.523 | 0.5506 | 0.5368 | 0.000999 |
| livertumor | MacRec | TFPI | LRP1 | 0.5043 | 0.3918 | 0.448 | 0.000999 |
| livertumor | MacRec | VTN | ITGB1 | 0.7399 | 0.446 | 0.5929 | 0.000999 |
| livertumor | MacRec | VTN | PLAUR | 0.7399 | 1.5635 | 1.1517 | 0.000999 |
| livertumor | MacRec | CXCL3 | CXCR4 | 0.2543 | 1.6682 | 0.9612 | 0.002 |
| livertumor | MacRec | GSTP1 | CD44 | 1.7219 | 1.4429 | 1.5824 | 0.003 |
| livertumor | MacRec | ARF1 | CD44 | 0.6936 | 1.4429 | 1.0682 | 0.00799 |
| lungtumor | MacRec | AIMP1 | RACK1 | 0.3345 | 2.605 | 1.4698 | 0.000999 |
| lungtumor | MacRec | ANXA1 | C3AR1 | 1.4026 | 0.4251 | 0.9139 | 0.000999 |
| lungtumor | MacRec | ANXA1 | C5AR1 | 1.4026 | 0.9163 | 1.1595 | 0.000999 |
| lungtumor | MacRec | ANXA1 | CD4 | 1.4026 | 0.5254 | 0.964 | 0.000999 |

| | | | | | | | |
|---|---|---|---|---|---|---|---|
| lungtumor | MacRec | ANXA1 | CD44 | 1.4026 | 1.4429 | 1.4227 | 0.000999 |
| lungtumor | MacRec | ANXA1 | CXCR4 | 1.4026 | 1.6682 | 1.5354 | 0.000999 |
| lungtumor | MacRec | ANXA1 | FPR1 | 1.4026 | 0.7287 | 1.0657 | 0.000999 |
| lungtumor | MacRec | ANXA1 | FPR3 | 1.4026 | 0.5947 | 0.9986 | 0.000999 |
| lungtumor | MacRec | ARF1 | CD4 | 0.8791 | 0.5254 | 0.7022 | 0.000999 |
| lungtumor | MacRec | ARF1 | CD44 | 0.8791 | 1.4429 | 1.161 | 0.000999 |
| lungtumor | MacRec | CALM1 | RPSA | 0.8426 | 2.3085 | 1.5756 | 0.000999 |
| lungtumor | MacRec | CALM1 | TNFRSF1B | 0.8426 | 0.5988 | 0.7207 | 0.000999 |
| lungtumor | MacRec | CALR | CD44 | 0.7795 | 1.4429 | 1.1112 | 0.000999 |
| lungtumor | MacRec | CALR | LRP1 | 0.7795 | 0.3918 | 0.5857 | 0.000999 |
| lungtumor | MacRec | CCL20 | C3AR1 | 0.2885 | 0.4251 | 0.3568 | 0.000999 |
| lungtumor | MacRec | CCL20 | C5AR1 | 0.2885 | 0.9163 | 0.6024 | 0.000999 |
| lungtumor | MacRec | CCL20 | CXCR4 | 0.2885 | 1.6682 | 0.9784 | 0.000999 |
| lungtumor | MacRec | CCL20 | FPR1 | 0.2885 | 0.7287 | 0.5086 | 0.000999 |
| lungtumor | MacRec | CXCL10 | C3AR1 | 0.3417 | 0.4251 | 0.3834 | 0.000999 |
| lungtumor | MacRec | CXCL10 | C5AR1 | 0.3417 | 0.9163 | 0.629 | 0.000999 |
| lungtumor | MacRec | CXCL10 | CXCR4 | 0.3417 | 1.6682 | 1.0049 | 0.000999 |
| lungtumor | MacRec | CXCL10 | FPR1 | 0.3417 | 0.7287 | 0.5352 | 0.000999 |
| lungtumor | MacRec | FGA | ITGAX | 0.4445 | 0.3634 | 0.4039 | 0.000999 |
| lungtumor | MacRec | FGA | ITGB1 | 0.4445 | 0.446 | 0.4452 | 0.000999 |
| lungtumor | MacRec | FGA | ITGB2 | 0.4445 | 1.3869 | 0.9157 | 0.000999 |
| lungtumor | MacRec | GRN | TNFRSF1B | 0.4026 | 0.5988 | 0.5007 | 0.000999 |
| lungtumor | MacRec | GSTP1 | CD44 | 2.4093 | 1.4429 | 1.9261 | 0.000999 |
| lungtumor | MacRec | HMGB1 | AGTRAP | 1.1943 | 0.3586 | 0.7764 | 0.000999 |
| lungtumor | MacRec | HMGB1 | LY96 | 1.1943 | 0.8558 | 1.025 | 0.000999 |
| lungtumor | MacRec | HMGB1 | TLR2 | 1.1943 | 0.398 | 0.7961 | 0.000999 |
| lungtumor | MacRec | HMGB2 | CD44 | 0.5593 | 1.4429 | 1.0011 | 0.000999 |
| lungtumor | MacRec | HMGB2 | DAB2 | 0.5593 | 0.3033 | 0.4313 | 0.000999 |
| lungtumor | MacRec | HSP90B1 | ITGB1 | 0.6141 | 0.446 | 0.53 | 0.000999 |
| lungtumor | MacRec | HSP90B1 | LRP1 | 0.6141 | 0.3918 | 0.5029 | 0.000999 |

| | | | | | | | |
|---|---|---|---|---|---|---|---|
| lungtumor | MacRec | HSP90B1 | TNFRSF1B | 0.6141 | 0.5988 | 0.6065 | 0.000999 |
| lungtumor | MacRec | LGALS3 | CLEC7A | 0.9915 | 0.8551 | 0.9233 | 0.000999 |
| lungtumor | MacRec | LGALS3 | FCGR2A | 0.9915 | 0.8792 | 0.9354 | 0.000999 |
| lungtumor | MacRec | LGALS3 | FCGR2B | 0.9915 | 0.7138 | 0.8526 | 0.000999 |
| lungtumor | MacRec | LGALS3 | ITGB1 | 0.9915 | 0.446 | 0.7187 | 0.000999 |
| lungtumor | MacRec | LGALS3 | TFRC | 0.9915 | 0.5506 | 0.771 | 0.000999 |
| lungtumor | MacRec | MDK | LRP1 | 1.2896 | 0.3918 | 0.8407 | 0.000999 |
| lungtumor | MacRec | MIF | CD74 | 1.4585 | 4.4657 | 2.9621 | 0.000999 |
| lungtumor | MacRec | MIF | TNFRSF14 | 1.4585 | 0.2703 | 0.8644 | 0.000999 |
| lungtumor | MacRec | RPS19 | C5AR1 | 3.5104 | 0.9163 | 2.2134 | 0.000999 |
| lungtumor | MacRec | RPS19 | CD4 | 3.5104 | 0.5254 | 2.0179 | 0.000999 |
| lungtumor | MacRec | RPS19 | RPSA | 3.5104 | 2.3085 | 2.9095 | 0.000999 |
| lungtumor | MacRec | RTN4 | CD44 | 0.4013 | 1.4429 | 0.9221 | 0.000999 |
| lungtumor | MacRec | SAA1 | C3AR1 | 0.2539 | 0.4251 | 0.3395 | 0.000999 |
| lungtumor | MacRec | SAA1 | C5AR1 | 0.2539 | 0.9163 | 0.5851 | 0.000999 |
| lungtumor | MacRec | SAA1 | CXCR4 | 0.2539 | 1.6682 | 0.961 | 0.000999 |
| lungtumor | MacRec | SAA1 | FPR1 | 0.2539 | 0.7287 | 0.4913 | 0.000999 |
| lungtumor | MacRec | SAA1 | GPR65 | 0.2539 | 0.3015 | 0.2777 | 0.000999 |
| lungtumor | MacRec | SAA1 | LPAR6 | 0.2539 | 0.2979 | 0.2759 | 0.000999 |
| lungtumor | MacRec | SAA1 | SELENOS | 0.2539 | 0.3438 | 0.2989 | 0.000999 |
| lungtumor | MacRec | SPP1 | CD44 | 1.4874 | 1.4429 | 1.4651 | 0.000999 |
| lungtumor | MacRec | SPP1 | ITGB1 | 1.4874 | 0.446 | 0.9667 | 0.000999 |
| lungtumor | MacRec | TFPI | LRP1 | 0.3615 | 0.3918 | 0.3766 | 0.000999 |
| lungtumor | MacRec | CALM1 | TNFRSF1A | 0.8426 | 0.4074 | 0.625 | 0.005 |
| melanomatumo | MacRec | ARF1 | CD4 | 1.8545 | 0.5254 | 1.1899 | 0.000999 |
| melanomatumo | MacRec | ARF1 | CD44 | 1.8545 | 1.4429 | 1.6487 | 0.000999 |
| melanomatumo | MacRec | GSTP1 | CD44 | 2.3341 | 1.4429 | 1.8885 | 0.000999 |
| melanomatumo | MacRec | HAS2 | CD44 | 0.3078 | 1.4429 | 0.8753 | 0.000999 |
| melanomatumo | MacRec | HEBP1 | C3AR1 | 0.3689 | 0.4251 | 0.397 | 0.000999 |
| melanomatumo | MacRec | HEBP1 | C5AR1 | 0.3689 | 0.9163 | 0.6426 | 0.000999 |

| | | | | | | | |
|---|---|---|---|---|---|---|---|
| melanomatumo | MacRec | HEBP1 | CXCR4 | 0.3689 | 1.6682 | 1.0185 | 0.000999 |
| melanomatumo | MacRec | HEBP1 | FPR1 | 0.3689 | 0.7287 | 0.5488 | 0.000999 |
| melanomatumo | MacRec | HEBP1 | FPR3 | 0.3689 | 0.5947 | 0.4818 | 0.000999 |
| melanomatumo | MacRec | HMGB1 | LY96 | 1.6757 | 0.8558 | 1.2657 | 0.000999 |
| melanomatumo | MacRec | HMGB1 | TLR2 | 1.6757 | 0.398 | 1.0368 | 0.000999 |
| melanomatumo | MacRec | MIF | CD74 | 2.6939 | 4.4657 | 3.5798 | 0.000999 |
| melanomatumo | MacRec | MIF | TNFRSF14 | 2.6939 | 0.2703 | 1.4821 | 0.000999 |
| melanomatumo | MacRec | SERPINE2 | LRP1 | 0.9323 | 0.3918 | 0.662 | 0.000999 |
| melanomatumo | MacRec | HMGB1 | AGTRAP | 1.6757 | 0.3586 | 1.0171 | 0.002 |
| ovarytumor | MacRec | ANXA1 | C3AR1 | 1.0314 | 0.4251 | 0.7282 | 0.000999 |
| ovarytumor | MacRec | ANXA1 | C5AR1 | 1.0314 | 0.9163 | 0.9739 | 0.000999 |
| ovarytumor | MacRec | ANXA1 | CD4 | 1.0314 | 0.5254 | 0.7784 | 0.000999 |
| ovarytumor | MacRec | ANXA1 | CD44 | 1.0314 | 1.4429 | 1.2371 | 0.000999 |
| ovarytumor | MacRec | ANXA1 | CXCR4 | 1.0314 | 1.6682 | 1.3498 | 0.000999 |
| ovarytumor | MacRec | ANXA1 | FPR1 | 1.0314 | 0.7287 | 0.88 | 0.000999 |
| ovarytumor | MacRec | ANXA1 | FPR3 | 1.0314 | 0.5947 | 0.813 | 0.000999 |
| ovarytumor | MacRec | APOE | LRP1 | 0.837 | 0.3918 | 0.6144 | 0.000999 |
| ovarytumor | MacRec | APOE | SORL1 | 0.837 | 0.5063 | 0.6717 | 0.000999 |
| ovarytumor | MacRec | APP | ALOX5 | 0.2907 | 0.4148 | 0.3528 | 0.000999 |
| ovarytumor | MacRec | APP | C3AR1 | 0.2907 | 0.4251 | 0.3579 | 0.000999 |
| ovarytumor | MacRec | APP | C5AR1 | 0.2907 | 0.9163 | 0.6035 | 0.000999 |
| ovarytumor | MacRec | APP | CD36 | 0.2907 | 0.3022 | 0.2965 | 0.000999 |
| ovarytumor | MacRec | APP | CXCR4 | 0.2907 | 1.6682 | 0.9795 | 0.000999 |
| ovarytumor | MacRec | APP | FPR1 | 0.2907 | 0.7287 | 0.5097 | 0.000999 |
| ovarytumor | MacRec | APP | LRP1 | 0.2907 | 0.3918 | 0.3413 | 0.000999 |
| ovarytumor | MacRec | APP | SORL1 | 0.2907 | 0.5063 | 0.3985 | 0.000999 |
| ovarytumor | MacRec | APP | TNFRSF14 | 0.2907 | 0.2703 | 0.2805 | 0.000999 |
| ovarytumor | MacRec | ARF1 | CD4 | 0.7364 | 0.5254 | 0.6309 | 0.000999 |
| ovarytumor | MacRec | ARF1 | CD44 | 0.7364 | 1.4429 | 1.0896 | 0.000999 |
| ovarytumor | MacRec | CALM1 | TNFRSF1A | 1.0087 | 0.4074 | 0.708 | 0.000999 |

| | | | | | | | |
|---|---|---|---|---|---|---|---|
| ovarytumor | MacRec | CALM1 | TNFRSF1B | 1.0087 | 0.5988 | 0.8038 | 0.000999 |
| ovarytumor | MacRec | CALR | CD44 | 0.7423 | 1.4429 | 1.0926 | 0.000999 |
| ovarytumor | MacRec | CALR | LRP1 | 0.7423 | 0.3918 | 0.567 | 0.000999 |
| ovarytumor | MacRec | COL1A1 | CD36 | 0.3504 | 0.3022 | 0.3263 | 0.000999 |
| ovarytumor | MacRec | COL1A1 | CD44 | 0.3504 | 1.4429 | 0.8966 | 0.000999 |
| ovarytumor | MacRec | COL1A1 | ITGB1 | 0.3504 | 0.446 | 0.3982 | 0.000999 |
| ovarytumor | MacRec | GRN | TNFRSF1B | 0.3896 | 0.5988 | 0.4942 | 0.000999 |
| ovarytumor | MacRec | GSTP1 | CD44 | 2.6185 | 1.4429 | 2.0307 | 0.000999 |
| ovarytumor | MacRec | HLA-DMA | CD4 | 0.2537 | 0.5254 | 0.3895 | 0.000999 |
| ovarytumor | MacRec | HMGB1 | AGTRAP | 1.7272 | 0.3586 | 1.0429 | 0.000999 |
| ovarytumor | MacRec | HMGB1 | LY96 | 1.7272 | 0.8558 | 1.2915 | 0.000999 |
| ovarytumor | MacRec | HMGB1 | TLR2 | 1.7272 | 0.398 | 1.0626 | 0.000999 |
| ovarytumor | MacRec | HMGB2 | CD44 | 0.4974 | 1.4429 | 0.9701 | 0.000999 |
| ovarytumor | MacRec | HMGB2 | DAB2 | 0.4974 | 0.3033 | 0.4004 | 0.000999 |
| ovarytumor | MacRec | HSP90B1 | ITGB1 | 0.6365 | 0.446 | 0.5412 | 0.000999 |
| ovarytumor | MacRec | HSP90B1 | LRP1 | 0.6365 | 0.3918 | 0.5141 | 0.000999 |
| ovarytumor | MacRec | HSP90B1 | TNFRSF1B | 0.6365 | 0.5988 | 0.6177 | 0.000999 |
| ovarytumor | MacRec | ITGB1 | CD36 | 0.2783 | 0.3022 | 0.2902 | 0.000999 |
| ovarytumor | MacRec | ITGB1 | DAB2 | 0.2783 | 0.3033 | 0.2908 | 0.000999 |
| ovarytumor | MacRec | ITGB1 | ENG | 0.2783 | 0.3416 | 0.31 | 0.000999 |
| ovarytumor | MacRec | ITGB1 | ITGAX | 0.2783 | 0.3634 | 0.3208 | 0.000999 |
| ovarytumor | MacRec | ITGB1 | PLAUR | 0.2783 | 1.5635 | 0.9209 | 0.000999 |
| ovarytumor | MacRec | LGALS3 | CLEC7A | 0.5097 | 0.8551 | 0.6824 | 0.000999 |
| ovarytumor | MacRec | LGALS3 | FCGR2A | 0.5097 | 0.8792 | 0.6945 | 0.000999 |
| ovarytumor | MacRec | LGALS3 | FCGR2B | 0.5097 | 0.7138 | 0.6117 | 0.000999 |
| ovarytumor | MacRec | LGALS3 | TFRC | 0.5097 | 0.5506 | 0.5301 | 0.000999 |
| ovarytumor | MacRec | LRPAP1 | LRP1 | 0.3826 | 0.3918 | 0.3872 | 0.000999 |
| ovarytumor | MacRec | LRPAP1 | SORL1 | 0.3826 | 0.5063 | 0.4445 | 0.000999 |
| ovarytumor | MacRec | MDK | LRP1 | 1.5321 | 0.3918 | 0.9619 | 0.000999 |
| ovarytumor | MacRec | MIF | CD74 | 0.8565 | 4.4657 | 2.6611 | 0.000999 |

| | | | | | | | |
|---|---|---|---|---|---|---|---|
| ovarytumor | MacRec | MIF | TNFRSF14 | 0.8565 | 0.2703 | 0.5634 | 0.000999 |
| ovarytumor | MacRec | RPS19 | C5AR1 | 3.9325 | 0.9163 | 2.4244 | 0.000999 |
| ovarytumor | MacRec | RPS19 | CD4 | 3.9325 | 0.5254 | 2.2289 | 0.000999 |
| ovarytumor | MacRec | RTN4 | CD44 | 0.4992 | 1.4429 | 0.971 | 0.000999 |
| ovarytumor | MacRec | RTN4 | CD99 | 0.4992 | 0.6596 | 0.5794 | 0.000999 |
| ovarytumor | MacRec | SERPING1 | LRP1 | 0.3678 | 0.3918 | 0.3798 | 0.000999 |
| ovarytumor | MacRec | SPP1 | CD44 | 0.3946 | 1.4429 | 0.9187 | 0.000999 |
| ovarytumor | MacRec | LGALS3 | ITGB1 | 0.5097 | 0.446 | 0.4778 | 0.003 |

| Supplementary Table 8: Communication results between macrophages and lymphocytes subpopulations, and malignant cells | | | | | | | |
|---|---|---|---|---|---|---|---|
| **MacAgPres vs. lymphocytes** | | | | | | | |
| celltype_01 | celltype_02 | ligand | receptor | ligand_mean_expression | receptor_mean_expression | ligand_receptor_mean_expression | pvalue |
| MacAgPres | NKT | A2M | TNFRSF14 | 0.9993 | 0.2848 | 0.642 | 0.000999 |
| MacAgPres | NKT | ALCAM | CD6 | 0.2541 | 0.4881 | 0.3711 | 0.000999 |
| MacAgPres | NKT | APOE | SORL1 | 2.1012 | 0.4137 | 1.2574 | 0.000999 |
| MacAgPres | NKT | C3 | CXCR4 | 1.5464 | 2.0373 | 1.7919 | 0.000999 |
| MacAgPres | NKT | C3 | ITGB2 | 1.5464 | 1.6481 | 1.5973 | 0.000999 |
| MacAgPres | NKT | COL1A1 | CD44 | 1.0301 | 0.6724 | 0.8512 | 0.000999 |
| MacAgPres | NKT | COL1A1 | ITGB1 | 1.0301 | 0.488 | 0.759 | 0.000999 |
| MacAgPres | NKT | CXCL16 | CXCR4 | 1.0017 | 2.0373 | 1.5195 | 0.000999 |
| MacAgPres | NKT | CXCL16 | S1PR4 | 1.0017 | 0.4435 | 0.7226 | 0.000999 |
| MacAgPres | NKT | CXCL16 | S1PR5 | 1.0017 | 0.508 | 0.7549 | 0.000999 |
| MacAgPres | NKT | CXCL2 | CXCR4 | 0.5101 | 2.0373 | 1.2737 | 0.000999 |
| MacAgPres | NKT | CXCL2 | S1PR4 | 0.5101 | 0.4435 | 0.4768 | 0.000999 |
| MacAgPres | NKT | CXCL2 | S1PR5 | 0.5101 | 0.508 | 0.5091 | 0.000999 |
| MacAgPres | NKT | CXCL3 | CXCR4 | 0.427 | 2.0373 | 1.2321 | 0.000999 |
| MacAgPres | NKT | CXCL3 | S1PR4 | 0.427 | 0.4435 | 0.4353 | 0.000999 |
| MacAgPres | NKT | CXCL3 | S1PR5 | 0.427 | 0.508 | 0.4675 | 0.000999 |
| MacAgPres | NKT | FN1 | CD44 | 0.9125 | 0.6724 | 0.7925 | 0.000999 |
| MacAgPres | NKT | FN1 | ITGA4 | 0.9125 | 0.2946 | 0.6035 | 0.000999 |
| MacAgPres | NKT | FN1 | ITGB1 | 0.9125 | 0.488 | 0.7002 | 0.000999 |
| MacAgPres | NKT | FN1 | ITGB7 | 0.9125 | 0.4721 | 0.6923 | 0.000999 |
| MacAgPres | NKT | FN1 | RPSA | 0.9125 | 2.3639 | 1.6382 | 0.000999 |
| MacAgPres | NKT | GRN | TNFRSF1B | 1.7572 | 0.4949 | 1.1261 | 0.000999 |
| MacAgPres | NKT | GSTP1 | CD44 | 1.4191 | 0.6724 | 1.0458 | 0.000999 |
| MacAgPres | NKT | GSTP1 | IL7R | 1.4191 | 0.5933 | 1.0062 | 0.000999 |
| MacAgPres | NKT | GSTP1 | ITGA4 | 1.4191 | 0.2946 | 0.8568 | 0.000999 |
| MacAgPres | NKT | HBEGF | CD44 | 0.4872 | 0.6724 | 0.5798 | 0.000999 |
| MacAgPres | NKT | HLA-DRA | PTPRC | 4.7477 | 1.8797 | 3.3137 | 0.000999 |

| | | | | | | | |
|---|---|---|---|---|---|---|---|
| MacAgPres | NKT | HMGB1 | ITGA4 | 1.2957 | 0.2946 | 0.7951 | 0.000999 |
| MacAgPres | NKT | HSP90B1 | ITGB1 | 0.7624 | 0.488 | 0.6252 | 0.000999 |
| MacAgPres | NKT | HSP90B1 | TNFRSF1B | 0.7624 | 0.4949 | 0.6287 | 0.000999 |
| MacAgPres | NKT | LGALS3 | CD6 | 0.6864 | 0.4881 | 0.5873 | 0.000999 |
| MacAgPres | NKT | LGALS3 | ITGB1 | 0.6864 | 0.488 | 0.5872 | 0.000999 |
| MacAgPres | NKT | LGALS3 | SPN | 0.6864 | 0.6398 | 0.6631 | 0.000999 |
| MacAgPres | NKT | LRPAP1 | SORL1 | 0.4575 | 0.4137 | 0.4356 | 0.000999 |
| MacAgPres | NKT | NAMPT | ANXA6 | 0.924 | 0.844 | 0.884 | 0.000999 |
| MacAgPres | NKT | PECAM1 | CXCR4 | 0.2665 | 2.0373 | 1.1519 | 0.000999 |
| MacAgPres | NKT | RPS19 | RPSA | 3.5707 | 2.3639 | 2.9673 | 0.000999 |
| MacAgPres | NKT | RTN4 | CD44 | 0.9772 | 0.6724 | 0.8248 | 0.000999 |
| MacAgPres | NKT | RTN4 | CD99 | 0.9772 | 1.4547 | 1.216 | 0.000999 |
| MacAgPres | NKT | SPP1 | CD44 | 1.3063 | 0.6724 | 0.9894 | 0.000999 |
| MacAgPres | NKT | SPP1 | ITGA4 | 1.3063 | 0.2946 | 0.8004 | 0.000999 |
| MacAgPres | NKT | SPP1 | ITGB1 | 1.3063 | 0.488 | 0.8972 | 0.000999 |
| MacAgPres | NKT | TNFSF13B | ITGB1 | 0.8442 | 0.488 | 0.6661 | 0.000999 |
| MacAgPres | NKT | RPS19 | ITGA4 | 3.5707 | 0.2946 | 1.9326 | 0.005 |
| MacAgPres | TCD4ex | A2M | TNFRSF14 | 0.9993 | 0.2806 | 0.6399 | 0.000999 |
| MacAgPres | TCD4ex | ALCAM | CD6 | 0.2541 | 0.5338 | 0.3939 | 0.000999 |
| MacAgPres | TCD4ex | ANXA1 | CCR7 | 0.957 | 0.4844 | 0.7207 | 0.000999 |
| MacAgPres | TCD4ex | ANXA1 | CD4 | 0.957 | 0.5864 | 0.7717 | 0.000999 |
| MacAgPres | TCD4ex | ANXA1 | CD44 | 0.957 | 1.0244 | 0.9907 | 0.000999 |
| MacAgPres | TCD4ex | ANXA1 | CXCR4 | 0.957 | 2.4201 | 1.6886 | 0.000999 |
| MacAgPres | TCD4ex | C3 | CCR7 | 1.5464 | 0.4844 | 1.0154 | 0.000999 |
| MacAgPres | TCD4ex | C3 | CXCR4 | 1.5464 | 2.4201 | 1.9832 | 0.000999 |
| MacAgPres | TCD4ex | C3 | ITGB2 | 1.5464 | 0.7871 | 1.1667 | 0.000999 |
| MacAgPres | TCD4ex | CALR | CD44 | 0.9 | 1.0244 | 0.9622 | 0.000999 |
| MacAgPres | TCD4ex | CALR | IL7R | 0.9 | 1.2353 | 1.0676 | 0.000999 |
| MacAgPres | TCD4ex | CD86 | CD28 | 0.5864 | 0.3256 | 0.456 | 0.000999 |
| MacAgPres | TCD4ex | CD86 | CTLA4 | 0.5864 | 0.6294 | 0.6079 | 0.000999 |

| | | | | | | | |
|---|---|---|---|---|---|---|---|
| MacAgPres | TCD4ex | COL1A1 | CD44 | 1.0301 | 1.0244 | 1.0272 | 0.000999 |
| MacAgPres | TCD4ex | COL1A1 | ITGB1 | 1.0301 | 0.4615 | 0.7458 | 0.000999 |
| MacAgPres | TCD4ex | CXCL16 | CCR7 | 1.0017 | 0.4844 | 0.7431 | 0.000999 |
| MacAgPres | TCD4ex | CXCL16 | CXCR4 | 1.0017 | 2.4201 | 1.7109 | 0.000999 |
| MacAgPres | TCD4ex | CXCL2 | CCR7 | 0.5101 | 0.4844 | 0.4973 | 0.000999 |
| MacAgPres | TCD4ex | CXCL2 | CXCR4 | 0.5101 | 2.4201 | 1.4651 | 0.000999 |
| MacAgPres | TCD4ex | CXCL3 | CCR7 | 0.427 | 0.4844 | 0.4557 | 0.000999 |
| MacAgPres | TCD4ex | CXCL3 | CXCR4 | 0.427 | 2.4201 | 1.4235 | 0.000999 |
| MacAgPres | TCD4ex | FN1 | CD44 | 0.9125 | 1.0244 | 0.9684 | 0.000999 |
| MacAgPres | TCD4ex | FN1 | ITGB1 | 0.9125 | 0.4615 | 0.687 | 0.000999 |
| MacAgPres | TCD4ex | FN1 | RPSA | 0.9125 | 2.4895 | 1.701 | 0.000999 |
| MacAgPres | TCD4ex | GRN | TNFRSF1B | 1.7572 | 0.383 | 1.0701 | 0.000999 |
| MacAgPres | TCD4ex | GSTP1 | CD44 | 1.4191 | 1.0244 | 1.2217 | 0.000999 |
| MacAgPres | TCD4ex | GSTP1 | IL7R | 1.4191 | 1.2353 | 1.3272 | 0.000999 |
| MacAgPres | TCD4ex | HBEGF | CD44 | 0.4872 | 1.0244 | 0.7558 | 0.000999 |
| MacAgPres | TCD4ex | HLA-DMA | CD4 | 2.1626 | 0.5864 | 1.3745 | 0.000999 |
| MacAgPres | TCD4ex | HLA-DRA | CD4 | 4.7477 | 0.5864 | 2.6671 | 0.000999 |
| MacAgPres | TCD4ex | HLA-DRA | PDCD1 | 4.7477 | 0.457 | 2.6023 | 0.000999 |
| MacAgPres | TCD4ex | HLA-DRA | PTPRC | 4.7477 | 1.7493 | 3.2485 | 0.000999 |
| MacAgPres | TCD4ex | HSP90B1 | ITGB1 | 0.7624 | 0.4615 | 0.612 | 0.000999 |
| MacAgPres | TCD4ex | HSP90B1 | TNFRSF1B | 0.7624 | 0.383 | 0.5727 | 0.000999 |
| MacAgPres | TCD4ex | ITGB2 | ITGAL | 1.28 | 0.2603 | 0.7702 | 0.000999 |
| MacAgPres | TCD4ex | LGALS3 | CD6 | 0.6864 | 0.5338 | 0.6101 | 0.000999 |
| MacAgPres | TCD4ex | LGALS3 | ITGB1 | 0.6864 | 0.4615 | 0.5739 | 0.000999 |
| MacAgPres | TCD4ex | NAMPT | ANXA6 | 0.924 | 0.5386 | 0.7313 | 0.000999 |
| MacAgPres | TCD4ex | PECAM1 | CXCR4 | 0.2665 | 2.4201 | 1.3433 | 0.000999 |
| MacAgPres | TCD4ex | RPS19 | RPSA | 3.5707 | 2.4895 | 3.0301 | 0.000999 |
| MacAgPres | TCD4ex | RTN4 | CD44 | 0.9772 | 1.0244 | 1.0008 | 0.000999 |
| MacAgPres | TCD4ex | RTN4 | CD99 | 0.9772 | 0.9165 | 0.9469 | 0.000999 |
| MacAgPres | TCD4ex | SPP1 | CD44 | 1.3063 | 1.0244 | 1.1654 | 0.000999 |

| | | | | | | | |
|---|---|---|---|---|---|---|---|
| MacAgPres | TCD4ex | SPP1 | ITGB1 | 1.3063 | 0.4615 | 0.8839 | 0.000999 |
| MacAgPres | TCD4ex | TNFSF12 | TNFRSF25 | 0.3095 | 0.3259 | 0.3177 | 0.000999 |
| MacAgPres | TCD4ex | TNFSF13B | ITGB1 | 0.8442 | 0.4615 | 0.6529 | 0.000999 |
| MacAgPres | TCD4reg | A2M | TNFRSF14 | 0.9993 | 0.3843 | 0.6918 | 0.000999 |
| MacAgPres | TCD4reg | ALCAM | CD6 | 0.2541 | 0.2651 | 0.2596 | 0.000999 |
| MacAgPres | TCD4reg | ANXA1 | CCR7 | 0.957 | 0.3098 | 0.6334 | 0.000999 |
| MacAgPres | TCD4reg | ANXA1 | CD4 | 0.957 | 0.429 | 0.693 | 0.000999 |
| MacAgPres | TCD4reg | ANXA1 | CD44 | 0.957 | 1.0896 | 1.0233 | 0.000999 |
| MacAgPres | TCD4reg | ANXA1 | CXCR3 | 0.957 | 0.3357 | 0.6464 | 0.000999 |
| MacAgPres | TCD4reg | ANXA1 | CXCR4 | 0.957 | 2.2962 | 1.6266 | 0.000999 |
| MacAgPres | TCD4reg | ANXA1 | CXCR6 | 0.957 | 0.3945 | 0.6758 | 0.000999 |
| MacAgPres | TCD4reg | ANXA1 | FAS | 0.957 | 0.2727 | 0.6149 | 0.000999 |
| MacAgPres | TCD4reg | C3 | CCR7 | 1.5464 | 0.3098 | 0.9281 | 0.000999 |
| MacAgPres | TCD4reg | C3 | CXCR3 | 1.5464 | 0.3357 | 0.941 | 0.000999 |
| MacAgPres | TCD4reg | C3 | CXCR4 | 1.5464 | 2.2962 | 1.9213 | 0.000999 |
| MacAgPres | TCD4reg | C3 | CXCR6 | 1.5464 | 0.3945 | 0.9705 | 0.000999 |
| MacAgPres | TCD4reg | C3 | ITGB2 | 1.5464 | 0.6238 | 1.0851 | 0.000999 |
| MacAgPres | TCD4reg | CALR | CD44 | 0.9 | 1.0896 | 0.9948 | 0.000999 |
| MacAgPres | TCD4reg | CALR | FAS | 0.9 | 0.2727 | 0.5863 | 0.000999 |
| MacAgPres | TCD4reg | CALR | IL7R | 0.9 | 0.9682 | 0.9341 | 0.000999 |
| MacAgPres | TCD4reg | CD86 | CD28 | 0.5864 | 0.313 | 0.4497 | 0.000999 |
| MacAgPres | TCD4reg | CD86 | CTLA4 | 0.5864 | 0.7909 | 0.6887 | 0.000999 |
| MacAgPres | TCD4reg | COL1A1 | CD44 | 1.0301 | 1.0896 | 1.0598 | 0.000999 |
| MacAgPres | TCD4reg | COL1A1 | ITGB1 | 1.0301 | 0.4001 | 0.7151 | 0.000999 |
| MacAgPres | TCD4reg | CXCL16 | CCR7 | 1.0017 | 0.3098 | 0.6558 | 0.000999 |
| MacAgPres | TCD4reg | CXCL16 | CXCR3 | 1.0017 | 0.3357 | 0.6687 | 0.000999 |
| MacAgPres | TCD4reg | CXCL16 | CXCR4 | 1.0017 | 2.2962 | 1.6489 | 0.000999 |
| MacAgPres | TCD4reg | CXCL16 | CXCR6 | 1.0017 | 0.3945 | 0.6981 | 0.000999 |
| MacAgPres | TCD4reg | CXCL16 | S1PR4 | 1.0017 | 0.2744 | 0.6381 | 0.000999 |
| MacAgPres | TCD4reg | CXCL2 | CCR7 | 0.5101 | 0.3098 | 0.41 | 0.000999 |

| | | | | | | | |
|---|---|---|---|---|---|---|---|
| MacAgPres | TCD4reg | CXCL2 | CXCR3 | 0.5101 | 0.3357 | 0.4229 | 0.000999 |
| MacAgPres | TCD4reg | CXCL2 | CXCR4 | 0.5101 | 2.2962 | 1.4031 | 0.000999 |
| MacAgPres | TCD4reg | CXCL2 | CXCR6 | 0.5101 | 0.3945 | 0.4523 | 0.000999 |
| MacAgPres | TCD4reg | CXCL2 | S1PR4 | 0.5101 | 0.2744 | 0.3923 | 0.000999 |
| MacAgPres | TCD4reg | CXCL3 | CCR7 | 0.427 | 0.3098 | 0.3684 | 0.000999 |
| MacAgPres | TCD4reg | CXCL3 | CXCR3 | 0.427 | 0.3357 | 0.3813 | 0.000999 |
| MacAgPres | TCD4reg | CXCL3 | CXCR4 | 0.427 | 2.2962 | 1.3616 | 0.000999 |
| MacAgPres | TCD4reg | CXCL3 | CXCR6 | 0.427 | 0.3945 | 0.4108 | 0.000999 |
| MacAgPres | TCD4reg | CXCL3 | S1PR4 | 0.427 | 0.2744 | 0.3507 | 0.000999 |
| MacAgPres | TCD4reg | FN1 | CD44 | 0.9125 | 1.0896 | 1.0011 | 0.000999 |
| MacAgPres | TCD4reg | FN1 | ITGB1 | 0.9125 | 0.4001 | 0.6563 | 0.000999 |
| MacAgPres | TCD4reg | FN1 | RPSA | 0.9125 | 2.4999 | 1.7062 | 0.000999 |
| MacAgPres | TCD4reg | GRN | TNFRSF1B | 1.7572 | 0.7982 | 1.2777 | 0.000999 |
| MacAgPres | TCD4reg | GSTP1 | CD44 | 1.4191 | 1.0896 | 1.2544 | 0.000999 |
| MacAgPres | TCD4reg | GSTP1 | FAS | 1.4191 | 0.2727 | 0.8459 | 0.000999 |
| MacAgPres | TCD4reg | GSTP1 | IL7R | 1.4191 | 0.9682 | 1.1936 | 0.000999 |
| MacAgPres | TCD4reg | HBEGF | CD44 | 0.4872 | 1.0896 | 0.7884 | 0.000999 |
| MacAgPres | TCD4reg | HLA-DMA | CD4 | 2.1626 | 0.429 | 1.2958 | 0.000999 |
| MacAgPres | TCD4reg | HLA-DRA | CD4 | 4.7477 | 0.429 | 2.5883 | 0.000999 |
| MacAgPres | TCD4reg | HLA-DRA | PTPRC | 4.7477 | 1.6603 | 3.204 | 0.000999 |
| MacAgPres | TCD4reg | HSP90B1 | ITGB1 | 0.7624 | 0.4001 | 0.5813 | 0.000999 |
| MacAgPres | TCD4reg | HSP90B1 | TNFRSF1B | 0.7624 | 0.7982 | 0.7803 | 0.000999 |
| MacAgPres | TCD4reg | LGALS3 | CD6 | 0.6864 | 0.2651 | 0.4757 | 0.000999 |
| MacAgPres | TCD4reg | LGALS3 | FAS | 0.6864 | 0.2727 | 0.4796 | 0.000999 |
| MacAgPres | TCD4reg | LGALS3 | ITGB1 | 0.6864 | 0.4001 | 0.5432 | 0.000999 |
| MacAgPres | TCD4reg | NAMPT | ANXA6 | 0.924 | 0.4557 | 0.6899 | 0.000999 |
| MacAgPres | TCD4reg | PECAM1 | CXCR4 | 0.2665 | 2.2962 | 1.2813 | 0.000999 |
| MacAgPres | TCD4reg | RTN4 | CD44 | 0.9772 | 1.0896 | 1.0334 | 0.000999 |
| MacAgPres | TCD4reg | RTN4 | CD99 | 0.9772 | 0.6603 | 0.8188 | 0.000999 |
| MacAgPres | TCD4reg | SPP1 | CD44 | 1.3063 | 1.0896 | 1.198 | 0.000999 |

| | | | | | | | |
|---|---|---|---|---|---|---|---|
| MacAgPres | TCD4reg | SPP1 | ITGB1 | 1.3063 | 0.4001 | 0.8532 | 0.000999 |
| MacAgPres | TCD4reg | TNFSF13B | ITGB1 | 0.8442 | 0.4001 | 0.6222 | 0.000999 |
| MacAgPres | TCD4reg | HMGB1 | FAS | 1.2957 | 0.2727 | 0.7842 | 0.004 |
| MacAgPres | TCD8em | A2M | TNFRSF14 | 0.9993 | 0.2915 | 0.6454 | 0.000999 |
| MacAgPres | TCD8em | ALCAM | CD6 | 0.2541 | 0.4204 | 0.3373 | 0.000999 |
| MacAgPres | TCD8em | C3 | CXCR3 | 1.5464 | 0.4189 | 0.9827 | 0.000999 |
| MacAgPres | TCD8em | C3 | CXCR4 | 1.5464 | 2.8389 | 2.1927 | 0.000999 |
| MacAgPres | TCD8em | C3 | CXCR6 | 1.5464 | 0.3399 | 0.9431 | 0.000999 |
| MacAgPres | TCD8em | C3 | ITGB2 | 1.5464 | 1.0018 | 1.2741 | 0.000999 |
| MacAgPres | TCD8em | CALR | CD44 | 0.9 | 1.119 | 1.0095 | 0.000999 |
| MacAgPres | TCD8em | CALR | IL7R | 0.9 | 0.7183 | 0.8091 | 0.000999 |
| MacAgPres | TCD8em | COL1A1 | CD44 | 1.0301 | 1.119 | 1.0746 | 0.000999 |
| MacAgPres | TCD8em | COL1A1 | ITGB1 | 1.0301 | 0.2557 | 0.6429 | 0.000999 |
| MacAgPres | TCD8em | CXCL16 | CXCR3 | 1.0017 | 0.4189 | 0.7103 | 0.000999 |
| MacAgPres | TCD8em | CXCL16 | CXCR4 | 1.0017 | 2.8389 | 1.9203 | 0.000999 |
| MacAgPres | TCD8em | CXCL16 | CXCR6 | 1.0017 | 0.3399 | 0.6708 | 0.000999 |
| MacAgPres | TCD8em | CXCL16 | S1PR4 | 1.0017 | 0.2677 | 0.6347 | 0.000999 |
| MacAgPres | TCD8em | CXCL2 | CXCR3 | 0.5101 | 0.4189 | 0.4645 | 0.000999 |
| MacAgPres | TCD8em | CXCL2 | CXCR4 | 0.5101 | 2.8389 | 1.6745 | 0.000999 |
| MacAgPres | TCD8em | CXCL2 | CXCR6 | 0.5101 | 0.3399 | 0.425 | 0.000999 |
| MacAgPres | TCD8em | CXCL2 | S1PR4 | 0.5101 | 0.2677 | 0.3889 | 0.000999 |
| MacAgPres | TCD8em | CXCL3 | CXCR3 | 0.427 | 0.4189 | 0.4229 | 0.000999 |
| MacAgPres | TCD8em | CXCL3 | CXCR4 | 0.427 | 2.8389 | 1.633 | 0.000999 |
| MacAgPres | TCD8em | CXCL3 | CXCR6 | 0.427 | 0.3399 | 0.3834 | 0.000999 |
| MacAgPres | TCD8em | CXCL3 | S1PR4 | 0.427 | 0.2677 | 0.3474 | 0.000999 |
| MacAgPres | TCD8em | FN1 | CD44 | 0.9125 | 1.119 | 1.0158 | 0.000999 |
| MacAgPres | TCD8em | FN1 | ITGA4 | 0.9125 | 0.2823 | 0.5974 | 0.000999 |
| MacAgPres | TCD8em | FN1 | ITGB1 | 0.9125 | 0.2557 | 0.5841 | 0.000999 |
| MacAgPres | TCD8em | FN1 | RPSA | 0.9125 | 2.3686 | 1.6406 | 0.000999 |
| MacAgPres | TCD8em | GRN | TNFRSF1B | 1.7572 | 0.3038 | 1.0305 | 0.000999 |

| | | | | | | | |
|---|---|---|---|---|---|---|---|
| MacAgPres | TCD8em | GSTP1 | CD44 | 1.4191 | 1.119 | 1.2691 | 0.000999 |
| MacAgPres | TCD8em | GSTP1 | IL7R | 1.4191 | 0.7183 | 1.0687 | 0.000999 |
| MacAgPres | TCD8em | GSTP1 | ITGA4 | 1.4191 | 0.2823 | 0.8507 | 0.000999 |
| MacAgPres | TCD8em | HBEGF | CD44 | 0.4872 | 1.119 | 0.8031 | 0.000999 |
| MacAgPres | TCD8em | HLA-DRA | PTPRC | 4.7477 | 1.7912 | 3.2694 | 0.000999 |
| MacAgPres | TCD8em | HSP90B1 | ITGB1 | 0.7624 | 0.2557 | 0.5091 | 0.000999 |
| MacAgPres | TCD8em | HSP90B1 | TNFRSF1B | 0.7624 | 0.3038 | 0.5331 | 0.000999 |
| MacAgPres | TCD8em | ITGB1 | CD47 | 0.3801 | 0.4276 | 0.4038 | 0.000999 |
| MacAgPres | TCD8em | ITGB1 | ITGA4 | 0.3801 | 0.2823 | 0.3312 | 0.000999 |
| MacAgPres | TCD8em | ITGB1 | RACK1 | 0.3801 | 2.4374 | 1.4087 | 0.000999 |
| MacAgPres | TCD8em | ITGB2 | ITGAL | 1.28 | 0.3162 | 0.7981 | 0.000999 |
| MacAgPres | TCD8em | LGALS3 | CD6 | 0.6864 | 0.4204 | 0.5534 | 0.000999 |
| MacAgPres | TCD8em | LGALS3 | ITGB1 | 0.6864 | 0.2557 | 0.4711 | 0.000999 |
| MacAgPres | TCD8em | LGALS3 | SPN | 0.6864 | 0.2603 | 0.4734 | 0.000999 |
| MacAgPres | TCD8em | NAMPT | ANXA6 | 0.924 | 0.552 | 0.738 | 0.000999 |
| MacAgPres | TCD8em | PECAM1 | CXCR4 | 0.2665 | 2.8389 | 1.5527 | 0.000999 |
| MacAgPres | TCD8em | RPS19 | RPSA | 3.5707 | 2.3686 | 2.9697 | 0.000999 |
| MacAgPres | TCD8em | RTN4 | CD44 | 0.9772 | 1.119 | 1.0481 | 0.000999 |
| MacAgPres | TCD8em | RTN4 | CD99 | 0.9772 | 1.1525 | 1.0649 | 0.000999 |
| MacAgPres | TCD8em | SPP1 | CD44 | 1.3063 | 1.119 | 1.2127 | 0.000999 |
| MacAgPres | TCD8em | SPP1 | ITGA4 | 1.3063 | 0.2823 | 0.7943 | 0.000999 |
| MacAgPres | TCD8em | SPP1 | ITGB1 | 1.3063 | 0.2557 | 0.781 | 0.000999 |
| MacAgPres | TCD8em | TNFSF13B | ITGB1 | 0.8442 | 0.2557 | 0.55 | 0.000999 |
| MacAgPres | TCD8ex | A2M | TNFRSF14 | 0.9993 | 0.3094 | 0.6544 | 0.000999 |
| MacAgPres | TCD8ex | ALCAM | CD6 | 0.2541 | 0.318 | 0.2861 | 0.000999 |
| MacAgPres | TCD8ex | ANXA1 | CD44 | 0.957 | 0.8219 | 0.8895 | 0.000999 |
| MacAgPres | TCD8ex | ANXA1 | CXCR3 | 0.957 | 0.3647 | 0.6609 | 0.000999 |
| MacAgPres | TCD8ex | ANXA1 | CXCR4 | 0.957 | 2.3472 | 1.6521 | 0.000999 |
| MacAgPres | TCD8ex | ANXA1 | CXCR6 | 0.957 | 0.6626 | 0.8098 | 0.000999 |
| MacAgPres | TCD8ex | ANXA1 | ITGA4 | 0.957 | 0.4432 | 0.7001 | 0.000999 |

| | | | | | | | |
|---|---|---|---|---|---|---|---|
| MacAgPres | TCD8ex | ANXA1 | VCAM1 | 0.957 | 0.4462 | 0.7016 | 0.000999 |
| MacAgPres | TCD8ex | APOE | VCAM1 | 2.1012 | 0.4462 | 1.2737 | 0.000999 |
| MacAgPres | TCD8ex | C3 | CXCR3 | 1.5464 | 0.3647 | 0.9555 | 0.000999 |
| MacAgPres | TCD8ex | C3 | CXCR4 | 1.5464 | 2.3472 | 1.9468 | 0.000999 |
| MacAgPres | TCD8ex | C3 | CXCR6 | 1.5464 | 0.6626 | 1.1045 | 0.000999 |
| MacAgPres | TCD8ex | C3 | ITGB2 | 1.5464 | 0.7635 | 1.1549 | 0.000999 |
| MacAgPres | TCD8ex | CD86 | CTLA4 | 0.5864 | 0.4883 | 0.5373 | 0.000999 |
| MacAgPres | TCD8ex | COL1A1 | CD44 | 1.0301 | 0.8219 | 0.926 | 0.000999 |
| MacAgPres | TCD8ex | COL1A1 | ITGB1 | 1.0301 | 0.2995 | 0.6648 | 0.000999 |
| MacAgPres | TCD8ex | CXCL16 | CXCR3 | 1.0017 | 0.3647 | 0.6832 | 0.000999 |
| MacAgPres | TCD8ex | CXCL16 | CXCR4 | 1.0017 | 2.3472 | 1.6745 | 0.000999 |
| MacAgPres | TCD8ex | CXCL16 | CXCR6 | 1.0017 | 0.6626 | 0.8321 | 0.000999 |
| MacAgPres | TCD8ex | CXCL2 | CXCR3 | 0.5101 | 0.3647 | 0.4374 | 0.000999 |
| MacAgPres | TCD8ex | CXCL2 | CXCR4 | 0.5101 | 2.3472 | 1.4287 | 0.000999 |
| MacAgPres | TCD8ex | CXCL2 | CXCR6 | 0.5101 | 0.6626 | 0.5863 | 0.000999 |
| MacAgPres | TCD8ex | CXCL3 | CXCR3 | 0.427 | 0.3647 | 0.3958 | 0.000999 |
| MacAgPres | TCD8ex | CXCL3 | CXCR4 | 0.427 | 2.3472 | 1.3871 | 0.000999 |
| MacAgPres | TCD8ex | CXCL3 | CXCR6 | 0.427 | 0.6626 | 0.5448 | 0.000999 |
| MacAgPres | TCD8ex | FN1 | CD44 | 0.9125 | 0.8219 | 0.8672 | 0.000999 |
| MacAgPres | TCD8ex | FN1 | ITGA4 | 0.9125 | 0.4432 | 0.6779 | 0.000999 |
| MacAgPres | TCD8ex | FN1 | ITGB1 | 0.9125 | 0.2995 | 0.606 | 0.000999 |
| MacAgPres | TCD8ex | FN1 | ITGB7 | 0.9125 | 0.3007 | 0.6066 | 0.000999 |
| MacAgPres | TCD8ex | FN1 | RPSA | 0.9125 | 2.2697 | 1.5911 | 0.000999 |
| MacAgPres | TCD8ex | GRN | TNFRSF1B | 1.7572 | 0.6436 | 1.2004 | 0.000999 |
| MacAgPres | TCD8ex | GSTP1 | CD44 | 1.4191 | 0.8219 | 1.1205 | 0.000999 |
| MacAgPres | TCD8ex | GSTP1 | ITGA4 | 1.4191 | 0.4432 | 0.9312 | 0.000999 |
| MacAgPres | TCD8ex | GSTP1 | VCAM1 | 1.4191 | 0.4462 | 0.9327 | 0.000999 |
| MacAgPres | TCD8ex | HBEGF | CD44 | 0.4872 | 0.8219 | 0.6546 | 0.000999 |
| MacAgPres | TCD8ex | HLA-DRA | PDCD1 | 4.7477 | 0.6109 | 2.6793 | 0.000999 |
| MacAgPres | TCD8ex | HLA-DRA | PTPRC | 4.7477 | 1.6762 | 3.2119 | 0.000999 |

| | | | | | | | |
|---|---|---|---|---|---|---|---|
| MacAgPres | TCD8ex | HSP90B1 | ITGB1 | 0.7624 | 0.2995 | 0.531 | 0.000999 |
| MacAgPres | TCD8ex | HSP90B1 | TNFRSF1B | 0.7624 | 0.6436 | 0.703 | 0.000999 |
| MacAgPres | TCD8ex | ITGB1 | CD47 | 0.3801 | 0.3952 | 0.3876 | 0.000999 |
| MacAgPres | TCD8ex | ITGB1 | ITGA4 | 0.3801 | 0.4432 | 0.4117 | 0.000999 |
| MacAgPres | TCD8ex | ITGB1 | VCAM1 | 0.3801 | 0.4462 | 0.4131 | 0.000999 |
| MacAgPres | TCD8ex | LGALS3 | CD6 | 0.6864 | 0.318 | 0.5022 | 0.000999 |
| MacAgPres | TCD8ex | LGALS3 | ITGB1 | 0.6864 | 0.2995 | 0.4929 | 0.000999 |
| MacAgPres | TCD8ex | LGALS3 | SPN | 0.6864 | 0.3486 | 0.5175 | 0.000999 |
| MacAgPres | TCD8ex | NAMPT | ANXA6 | 0.924 | 0.5678 | 0.7459 | 0.000999 |
| MacAgPres | TCD8ex | PECAM1 | CXCR4 | 0.2665 | 2.3472 | 1.3068 | 0.000999 |
| MacAgPres | TCD8ex | RTN4 | CD44 | 0.9772 | 0.8219 | 0.8996 | 0.000999 |
| MacAgPres | TCD8ex | RTN4 | CD99 | 0.9772 | 1.2049 | 1.091 | 0.000999 |
| MacAgPres | TCD8ex | SPP1 | CD44 | 1.3063 | 0.8219 | 1.0641 | 0.000999 |
| MacAgPres | TCD8ex | SPP1 | ITGA4 | 1.3063 | 0.4432 | 0.8748 | 0.000999 |
| MacAgPres | TCD8ex | SPP1 | ITGB1 | 1.3063 | 0.2995 | 0.8029 | 0.000999 |
| MacAgPres | TCD8ex | TNFSF13B | ITGB1 | 0.8442 | 0.2995 | 0.5719 | 0.000999 |
| MacAgPres | TCD8ex | ITGB1 | RACK1 | 0.3801 | 2.3353 | 1.3577 | 0.004 |
| MacAgPres | TGD | A2M | TNFRSF14 | 0.9993 | 0.4642 | 0.7317 | 0.000999 |
| MacAgPres | TGD | ALCAM | CD6 | 0.2541 | 0.4684 | 0.3613 | 0.000999 |
| MacAgPres | TGD | ANXA1 | CD44 | 0.957 | 1.1342 | 1.0456 | 0.000999 |
| MacAgPres | TGD | ANXA1 | CXCR3 | 0.957 | 1.1748 | 1.0659 | 0.000999 |
| MacAgPres | TGD | ANXA1 | ITGA4 | 0.957 | 0.3968 | 0.6769 | 0.000999 |
| MacAgPres | TGD | ARF1 | CD44 | 0.6151 | 1.1342 | 0.8746 | 0.000999 |
| MacAgPres | TGD | ARF1 | ITGA4 | 0.6151 | 0.3968 | 0.5059 | 0.000999 |
| MacAgPres | TGD | C3 | CXCR3 | 1.5464 | 1.1748 | 1.3606 | 0.000999 |
| MacAgPres | TGD | C3 | ITGB2 | 1.5464 | 1.4444 | 1.4954 | 0.000999 |
| MacAgPres | TGD | CALM1 | RPSA | 1.0626 | 2.8814 | 1.972 | 0.000999 |
| MacAgPres | TGD | CALR | CD44 | 0.9 | 1.1342 | 1.0171 | 0.000999 |
| MacAgPres | TGD | CALR | IL7R | 0.9 | 1.3862 | 1.1431 | 0.000999 |
| MacAgPres | TGD | CD99 | CD99 | 0.5386 | 1.7154 | 1.127 | 0.000999 |

| | | | | | | | |
|---|---|---|---|---|---|---|---|
| MacAgPres | TGD | COL1A1 | CD44 | 1.0301 | 1.1342 | 1.0821 | 0.000999 |
| MacAgPres | TGD | CXCL16 | CXCR3 | 1.0017 | 1.1748 | 1.0883 | 0.000999 |
| MacAgPres | TGD | CXCL2 | CXCR3 | 0.5101 | 1.1748 | 0.8424 | 0.000999 |
| MacAgPres | TGD | CXCL3 | CXCR3 | 0.427 | 1.1748 | 0.8009 | 0.000999 |
| MacAgPres | TGD | FN1 | CD44 | 0.9125 | 1.1342 | 1.0234 | 0.000999 |
| MacAgPres | TGD | FN1 | ITGA4 | 0.9125 | 0.3968 | 0.6547 | 0.000999 |
| MacAgPres | TGD | FN1 | ITGB7 | 0.9125 | 0.459 | 0.6858 | 0.000999 |
| MacAgPres | TGD | FN1 | RPSA | 0.9125 | 2.8814 | 1.8969 | 0.000999 |
| MacAgPres | TGD | GRN | TNFRSF1A | 1.7572 | 0.3099 | 1.0336 | 0.000999 |
| MacAgPres | TGD | GSTP1 | CD44 | 1.4191 | 1.1342 | 1.2767 | 0.000999 |
| MacAgPres | TGD | GSTP1 | IL7R | 1.4191 | 1.3862 | 1.4027 | 0.000999 |
| MacAgPres | TGD | GSTP1 | ITGA4 | 1.4191 | 0.3968 | 0.908 | 0.000999 |
| MacAgPres | TGD | HBEGF | CD44 | 0.4872 | 1.1342 | 0.8107 | 0.000999 |
| MacAgPres | TGD | HLA-A | CD8A | 2.9135 | 1.746 | 2.3298 | 0.000999 |
| MacAgPres | TGD | HLA-DRA | PTPRC | 4.7477 | 1.8653 | 3.3065 | 0.000999 |
| MacAgPres | TGD | HLA-E | CD8A | 1.9307 | 1.746 | 1.8384 | 0.000999 |
| MacAgPres | TGD | HLA-E | KLRC2 | 1.9307 | 0.8082 | 1.3695 | 0.000999 |
| MacAgPres | TGD | HLA-E | KLRD1 | 1.9307 | 1.3451 | 1.6379 | 0.000999 |
| MacAgPres | TGD | HLA-F | CD8A | 0.2712 | 1.746 | 1.0086 | 0.000999 |
| MacAgPres | TGD | HMGB1 | ITGA4 | 1.2957 | 0.3968 | 0.8463 | 0.000999 |
| MacAgPres | TGD | HMGB2 | CD44 | 0.4224 | 1.1342 | 0.7783 | 0.000999 |
| MacAgPres | TGD | ITGB1 | CD46 | 0.3801 | 0.3085 | 0.3443 | 0.000999 |
| MacAgPres | TGD | ITGB1 | ITGA4 | 0.3801 | 0.3968 | 0.3885 | 0.000999 |
| MacAgPres | TGD | ITGB1 | PTK2B | 0.3801 | 0.3668 | 0.3734 | 0.000999 |
| MacAgPres | TGD | ITGB1 | RACK1 | 0.3801 | 2.7287 | 1.5544 | 0.000999 |
| MacAgPres | TGD | ITGB2 | ITGAL | 1.28 | 0.3731 | 0.8266 | 0.000999 |
| MacAgPres | TGD | LGALS3 | CD6 | 0.6864 | 0.4684 | 0.5774 | 0.000999 |
| MacAgPres | TGD | LGALS3 | SPN | 0.6864 | 0.5644 | 0.6254 | 0.000999 |
| MacAgPres | TGD | LGALS3 | TGFBR2 | 0.6864 | 0.3677 | 0.527 | 0.000999 |
| MacAgPres | TGD | NAMPT | ANXA6 | 0.924 | 0.8553 | 0.8897 | 0.000999 |

| | | | | | | | |
|---|---|---|---|---|---|---|---|
| MacAgPres | TGD | PTPRC | CD2 | 0.8308 | 1.4522 | 1.1415 | 0.000999 |
| MacAgPres | TGD | PTPRC | CD8A | 0.8308 | 1.746 | 1.2884 | 0.000999 |
| MacAgPres | TGD | PTPRC | ITGAL | 0.8308 | 0.3731 | 0.602 | 0.000999 |
| MacAgPres | TGD | RPS19 | ITGA4 | 3.5707 | 0.3968 | 1.9838 | 0.000999 |
| MacAgPres | TGD | RPS19 | RPSA | 3.5707 | 2.8814 | 3.226 | 0.000999 |
| MacAgPres | TGD | RTN4 | CD44 | 0.9772 | 1.1342 | 1.0557 | 0.000999 |
| MacAgPres | TGD | RTN4 | CD99 | 0.9772 | 1.7154 | 1.3463 | 0.000999 |
| MacAgPres | TGD | SPP1 | CD44 | 1.3063 | 1.1342 | 1.2203 | 0.000999 |
| MacAgPres | TGD | SPP1 | ITGA4 | 1.3063 | 0.3968 | 0.8516 | 0.000999 |
| MacAgPres | TGD | SPP1 | ITGB1 | 1.3063 | 0.3657 | 0.836 | 0.000999 |
| MacAgPres | TGD | PTPRC | CD44 | 0.8308 | 1.1342 | 0.9825 | 0.003 |
| MacAgPres | TGD | MIF | ITGA4 | 0.4996 | 0.3968 | 0.4482 | 0.00599 |
| NKT | MacAgPres | ALOX5AP | ALOX5 | 0.5021 | 0.3892 | 0.4457 | 0.000999 |
| NKT | MacAgPres | ANXA1 | C3AR1 | 1.2259 | 0.7287 | 0.9773 | 0.000999 |
| NKT | MacAgPres | ANXA1 | C5AR1 | 1.2259 | 0.5708 | 0.8984 | 0.000999 |
| NKT | MacAgPres | ANXA1 | CD4 | 1.2259 | 0.5049 | 0.8654 | 0.000999 |
| NKT | MacAgPres | ANXA1 | CD44 | 1.2259 | 0.7335 | 0.9797 | 0.000999 |
| NKT | MacAgPres | ANXA1 | FPR1 | 1.2259 | 0.4477 | 0.8368 | 0.000999 |
| NKT | MacAgPres | ANXA1 | FPR3 | 1.2259 | 0.2875 | 0.7567 | 0.000999 |
| NKT | MacAgPres | ARF1 | CD4 | 0.7991 | 0.5049 | 0.652 | 0.000999 |
| NKT | MacAgPres | ARF1 | CD44 | 0.7991 | 0.7335 | 0.7663 | 0.000999 |
| NKT | MacAgPres | CALM1 | TNFRSF1A | 1.9335 | 0.3753 | 1.1544 | 0.000999 |
| NKT | MacAgPres | CALM1 | TNFRSF1B | 1.9335 | 0.4694 | 1.2015 | 0.000999 |
| NKT | MacAgPres | CALR | LRP1 | 0.9479 | 0.3779 | 0.6629 | 0.000999 |
| NKT | MacAgPres | CD2 | CD4 | 1.1465 | 0.5049 | 0.8257 | 0.000999 |
| NKT | MacAgPres | CD320 | TGFBR2 | 0.2545 | 0.2588 | 0.2567 | 0.000999 |
| NKT | MacAgPres | CD6 | ALCAM | 0.4881 | 0.2541 | 0.3711 | 0.000999 |
| NKT | MacAgPres | HMGB1 | LY96 | 1.195 | 0.726 | 0.9605 | 0.000999 |
| NKT | MacAgPres | HMGB1 | TLR2 | 1.195 | 0.327 | 0.761 | 0.000999 |
| NKT | MacAgPres | HMGB2 | CD44 | 0.6467 | 0.7335 | 0.6901 | 0.000999 |

| | | | | | | | |
|---|---|---|---|---|---|---|---|
| NKT | MacAgPres | HMGB2 | DAB2 | 0.6467 | 0.6335 | 0.6401 | 0.000999 |
| NKT | MacAgPres | HSP90B1 | LRP1 | 0.6138 | 0.3779 | 0.4958 | 0.000999 |
| NKT | MacAgPres | IL16 | CD4 | 0.3556 | 0.5049 | 0.4302 | 0.000999 |
| NKT | MacAgPres | IL16 | FCGR1A | 0.3556 | 0.769 | 0.5623 | 0.000999 |
| NKT | MacAgPres | ITGA4 | CD44 | 0.2946 | 0.7335 | 0.514 | 0.000999 |
| NKT | MacAgPres | ITGB1 | CD151 | 0.488 | 0.2751 | 0.3815 | 0.000999 |
| NKT | MacAgPres | ITGB1 | DAB2 | 0.488 | 0.6335 | 0.5607 | 0.000999 |
| NKT | MacAgPres | ITGB1 | PLAUR | 0.488 | 1.0849 | 0.7864 | 0.000999 |
| NKT | MacAgPres | ITGB2 | PLAUR | 1.6481 | 1.0849 | 1.3665 | 0.000999 |
| NKT | MacAgPres | LRPAP1 | LRP1 | 0.2807 | 0.3779 | 0.3293 | 0.000999 |
| NKT | MacAgPres | LRPAP1 | SORL1 | 0.2807 | 0.5617 | 0.4212 | 0.000999 |
| NKT | MacAgPres | LTB | TNFRSF1A | 0.2605 | 0.3753 | 0.3179 | 0.000999 |
| NKT | MacAgPres | MIF | CD74 | 0.6657 | 5.2676 | 2.9666 | 0.000999 |
| NKT | MacAgPres | PTPRC | CD4 | 1.8797 | 0.5049 | 1.1923 | 0.000999 |
| NKT | MacAgPres | PTPRC | CD44 | 1.8797 | 0.7335 | 1.3066 | 0.000999 |
| NKT | MacAgPres | RPS19 | C5AR1 | 3.5336 | 0.5708 | 2.0522 | 0.000999 |
| NKT | MacAgPres | RPS19 | CD4 | 3.5336 | 0.5049 | 2.0192 | 0.000999 |
| NKT | MacAgPres | TGFB1 | TGFBR1 | 0.5441 | 0.2605 | 0.4023 | 0.000999 |
| NKT | MacAgPres | TGFB1 | TGFBR2 | 0.5441 | 0.2588 | 0.4014 | 0.000999 |
| NKT | MacAgPres | TGFB1 | TLR2 | 0.5441 | 0.327 | 0.4355 | 0.000999 |
| NKT | MacAgPres | TIMP1 | LRP1 | 0.3005 | 0.3779 | 0.3392 | 0.000999 |
| NKT | MacAgPres | XCL2 | LPAR6 | 0.2774 | 0.6544 | 0.4659 | 0.000999 |
| NKT | MacAgPres | XCL2 | PTAFR | 0.2774 | 0.2882 | 0.2828 | 0.000999 |
| NKT | MacAgPres | CALR | CD44 | 0.9479 | 0.7335 | 0.8407 | 0.00599 |
| TCD4ex | MacAgPres | ALOX5AP | ALOX5 | 1.119 | 0.3892 | 0.7541 | 0.000999 |
| TCD4ex | MacAgPres | ANXA1 | C3AR1 | 0.5174 | 0.7287 | 0.6231 | 0.000999 |
| TCD4ex | MacAgPres | ANXA1 | C5AR1 | 0.5174 | 0.5708 | 0.5441 | 0.000999 |
| TCD4ex | MacAgPres | ANXA1 | FPR1 | 0.5174 | 0.4477 | 0.4826 | 0.000999 |
| TCD4ex | MacAgPres | ANXA1 | FPR3 | 0.5174 | 0.2875 | 0.4025 | 0.000999 |
| TCD4ex | MacAgPres | CALM1 | TNFRSF1A | 1.2957 | 0.3753 | 0.8355 | 0.000999 |

| | | | | | | | |
|---|---|---|---|---|---|---|---|
| TCD4ex | MacAgPres | CALM1 | TNFRSF1B | 1.2957 | 0.4694 | 0.8826 | 0.000999 |
| TCD4ex | MacAgPres | CALR | LRP1 | 0.748 | 0.3779 | 0.5629 | 0.000999 |
| TCD4ex | MacAgPres | CD2 | CD4 | 1.5731 | 0.5049 | 1.039 | 0.000999 |
| TCD4ex | MacAgPres | CD6 | ALCAM | 0.5338 | 0.2541 | 0.3939 | 0.000999 |
| TCD4ex | MacAgPres | CXCL13 | C3AR1 | 2.3367 | 0.7287 | 1.5327 | 0.000999 |
| TCD4ex | MacAgPres | CXCL13 | C5AR1 | 2.3367 | 0.5708 | 1.4537 | 0.000999 |
| TCD4ex | MacAgPres | CXCL13 | CXCR4 | 2.3367 | 1.7715 | 2.0541 | 0.000999 |
| TCD4ex | MacAgPres | CXCL13 | FPR1 | 2.3367 | 0.4477 | 1.3922 | 0.000999 |
| TCD4ex | MacAgPres | CXCL13 | P2RY13 | 2.3367 | 0.3022 | 1.3194 | 0.000999 |
| TCD4ex | MacAgPres | HMGB1 | LY96 | 1.3876 | 0.726 | 1.0568 | 0.000999 |
| TCD4ex | MacAgPres | HMGB1 | TLR2 | 1.3876 | 0.327 | 0.8573 | 0.000999 |
| TCD4ex | MacAgPres | HMGB2 | DAB2 | 1.0243 | 0.6335 | 0.8289 | 0.000999 |
| TCD4ex | MacAgPres | HSP90B1 | LRP1 | 0.3944 | 0.3779 | 0.3861 | 0.000999 |
| TCD4ex | MacAgPres | ICAM3 | ITGB2 | 0.573 | 1.28 | 0.9265 | 0.000999 |
| TCD4ex | MacAgPres | ITGB1 | CD151 | 0.4615 | 0.2751 | 0.3683 | 0.000999 |
| TCD4ex | MacAgPres | ITGB1 | DAB2 | 0.4615 | 0.6335 | 0.5475 | 0.000999 |
| TCD4ex | MacAgPres | ITGB1 | PLAUR | 0.4615 | 1.0849 | 0.7732 | 0.000999 |
| TCD4ex | MacAgPres | ITGB2 | PLAUR | 0.7871 | 1.0849 | 0.936 | 0.000999 |
| TCD4ex | MacAgPres | LRPAP1 | LRP1 | 0.2809 | 0.3779 | 0.3294 | 0.000999 |
| TCD4ex | MacAgPres | LRPAP1 | SORL1 | 0.2809 | 0.5617 | 0.4213 | 0.000999 |
| TCD4ex | MacAgPres | LTB | ITGB1 | 0.9655 | 0.3801 | 0.6728 | 0.000999 |
| TCD4ex | MacAgPres | LTB | TNFRSF1A | 0.9655 | 0.3753 | 0.6704 | 0.000999 |
| TCD4ex | MacAgPres | MIF | CD74 | 0.6999 | 5.2676 | 2.9837 | 0.000999 |
| TCD4ex | MacAgPres | NMB | GPR65 | 0.6391 | 0.3478 | 0.4935 | 0.000999 |
| TCD4ex | MacAgPres | NMB | LPAR6 | 0.6391 | 0.6544 | 0.6467 | 0.000999 |
| TCD4ex | MacAgPres | NMB | PTAFR | 0.6391 | 0.2882 | 0.4637 | 0.000999 |
| TCD4ex | MacAgPres | PTPRC | CD4 | 1.7493 | 0.5049 | 1.1271 | 0.000999 |
| TCD4ex | MacAgPres | RPS19 | C5AR1 | 3.6329 | 0.5708 | 2.1018 | 0.000999 |
| TCD4ex | MacAgPres | SELL | CD4 | 0.5618 | 0.5049 | 0.5333 | 0.000999 |
| TCD4ex | MacAgPres | SELL | SELPLG | 0.5618 | 0.2693 | 0.4155 | 0.000999 |

| | | | | | | | |
|---|---|---|---|---|---|---|---|
| TCD4ex | MacAgPres | PTPRC | CD44 | 1.7493 | 0.7335 | 1.2414 | 0.004 |
| TCD4ex | MacAgPres | ITGB1 | CD47 | 0.4615 | 0.3827 | 0.4221 | 0.00599 |
| TCD4reg | MacAgPres | ALOX5AP | ALOX5 | 0.4589 | 0.3892 | 0.4241 | 0.000999 |
| TCD4reg | MacAgPres | ARF1 | CD4 | 0.7185 | 0.5049 | 0.6117 | 0.000999 |
| TCD4reg | MacAgPres | CALM1 | TNFRSF1A | 1.534 | 0.3753 | 0.9546 | 0.000999 |
| TCD4reg | MacAgPres | CALR | LRP1 | 0.6835 | 0.3779 | 0.5307 | 0.000999 |
| TCD4reg | MacAgPres | CD2 | CD4 | 1.6566 | 0.5049 | 1.0808 | 0.000999 |
| TCD4reg | MacAgPres | CD28 | CD4 | 0.313 | 0.5049 | 0.4089 | 0.000999 |
| TCD4reg | MacAgPres | CD6 | ALCAM | 0.2651 | 0.2541 | 0.2596 | 0.000999 |
| TCD4reg | MacAgPres | HMGB1 | LY96 | 1.2547 | 0.726 | 0.9904 | 0.000999 |
| TCD4reg | MacAgPres | HMGB1 | TLR2 | 1.2547 | 0.327 | 0.7909 | 0.000999 |
| TCD4reg | MacAgPres | HMGB2 | DAB2 | 0.616 | 0.6335 | 0.6248 | 0.000999 |
| TCD4reg | MacAgPres | HSP90B1 | LRP1 | 0.4221 | 0.3779 | 0.4 | 0.000999 |
| TCD4reg | MacAgPres | ICAM3 | ITGB2 | 0.6663 | 1.28 | 0.9731 | 0.000999 |
| TCD4reg | MacAgPres | IL16 | CD4 | 0.2528 | 0.5049 | 0.3788 | 0.000999 |
| TCD4reg | MacAgPres | IL16 | FCGR1A | 0.2528 | 0.769 | 0.5109 | 0.000999 |
| TCD4reg | MacAgPres | ITGB1 | CD151 | 0.4001 | 0.2751 | 0.3376 | 0.000999 |
| TCD4reg | MacAgPres | ITGB1 | DAB2 | 0.4001 | 0.6335 | 0.5168 | 0.000999 |
| TCD4reg | MacAgPres | ITGB1 | PLAUR | 0.4001 | 1.0849 | 0.7425 | 0.000999 |
| TCD4reg | MacAgPres | ITGB2 | PLAUR | 0.6238 | 1.0849 | 0.8543 | 0.000999 |
| TCD4reg | MacAgPres | LGALS3 | CLEC7A | 0.4492 | 0.6841 | 0.5666 | 0.000999 |
| TCD4reg | MacAgPres | LGALS3 | FCGR2A | 0.4492 | 1.2286 | 0.8389 | 0.000999 |
| TCD4reg | MacAgPres | LGALS3 | FCGR2B | 0.4492 | 0.578 | 0.5136 | 0.000999 |
| TCD4reg | MacAgPres | LGALS3 | TGFBR2 | 0.4492 | 0.2588 | 0.354 | 0.000999 |
| TCD4reg | MacAgPres | LRPAP1 | LRP1 | 0.3101 | 0.3779 | 0.344 | 0.000999 |
| TCD4reg | MacAgPres | LRPAP1 | SORL1 | 0.3101 | 0.5617 | 0.4359 | 0.000999 |
| TCD4reg | MacAgPres | LTB | ITGB1 | 2.242 | 0.3801 | 1.3111 | 0.000999 |
| TCD4reg | MacAgPres | LTB | TNFRSF1A | 2.242 | 0.3753 | 1.3087 | 0.000999 |
| TCD4reg | MacAgPres | MIF | CD74 | 0.6295 | 5.2676 | 2.9485 | 0.000999 |
| TCD4reg | MacAgPres | PTPRC | CD4 | 1.6603 | 0.5049 | 1.0826 | 0.000999 |

| | | | | | | | |
|---|---|---|---|---|---|---|---|
| TCD4reg | MacAgPres | RPS19 | C5AR1 | 3.6897 | 0.5708 | 2.1302 | 0.000999 |
| TCD4reg | MacAgPres | RPS19 | CD4 | 3.6897 | 0.5049 | 2.0973 | 0.000999 |
| TCD4reg | MacAgPres | SELL | CD4 | 0.7062 | 0.5049 | 0.6055 | 0.000999 |
| TCD4reg | MacAgPres | TIMP1 | LRP1 | 0.4362 | 0.3779 | 0.407 | 0.000999 |
| TCD4reg | MacAgPres | SELL | SELPLG | 0.7062 | 0.2693 | 0.4877 | 0.002 |
| TCD8em | MacAgPres | ALOX5AP | ALOX5 | 0.7973 | 0.3892 | 0.5933 | 0.000999 |
| TCD8em | MacAgPres | ANXA1 | C3AR1 | 1.0693 | 0.7287 | 0.899 | 0.000999 |
| TCD8em | MacAgPres | ANXA1 | C5AR1 | 1.0693 | 0.5708 | 0.82 | 0.000999 |
| TCD8em | MacAgPres | ANXA1 | CD4 | 1.0693 | 0.5049 | 0.7871 | 0.000999 |
| TCD8em | MacAgPres | ANXA1 | FPR1 | 1.0693 | 0.4477 | 0.7585 | 0.000999 |
| TCD8em | MacAgPres | ANXA1 | FPR3 | 1.0693 | 0.2875 | 0.6784 | 0.000999 |
| TCD8em | MacAgPres | APOE | LRP1 | 0.2791 | 0.3779 | 0.3285 | 0.000999 |
| TCD8em | MacAgPres | APOE | SORL1 | 0.2791 | 0.5617 | 0.4204 | 0.000999 |
| TCD8em | MacAgPres | ARF1 | CD4 | 0.6684 | 0.5049 | 0.5866 | 0.000999 |
| TCD8em | MacAgPres | CALM1 | TNFRSF1A | 1.5262 | 0.3753 | 0.9507 | 0.000999 |
| TCD8em | MacAgPres | CALM1 | TNFRSF1B | 1.5262 | 0.4694 | 0.9978 | 0.000999 |
| TCD8em | MacAgPres | CALR | LRP1 | 0.7141 | 0.3779 | 0.546 | 0.000999 |
| TCD8em | MacAgPres | CD2 | CD4 | 1.5961 | 0.5049 | 1.0505 | 0.000999 |
| TCD8em | MacAgPres | CD6 | ALCAM | 0.4204 | 0.2541 | 0.3373 | 0.000999 |
| TCD8em | MacAgPres | HLA-DMA | CD4 | 0.2969 | 0.5049 | 0.4009 | 0.000999 |
| TCD8em | MacAgPres | HMGB1 | LY96 | 1.3636 | 0.726 | 1.0448 | 0.000999 |
| TCD8em | MacAgPres | HMGB1 | TLR2 | 1.3636 | 0.327 | 0.8453 | 0.000999 |
| TCD8em | MacAgPres | HMGB2 | DAB2 | 0.7212 | 0.6335 | 0.6773 | 0.000999 |
| TCD8em | MacAgPres | HSP90B1 | ITGB1 | 0.471 | 0.3801 | 0.4256 | 0.000999 |
| TCD8em | MacAgPres | HSP90B1 | LRP1 | 0.471 | 0.3779 | 0.4245 | 0.000999 |
| TCD8em | MacAgPres | HSP90B1 | TNFRSF1B | 0.471 | 0.4694 | 0.4702 | 0.000999 |
| TCD8em | MacAgPres | ICAM3 | ITGB2 | 0.5282 | 1.28 | 0.9041 | 0.000999 |
| TCD8em | MacAgPres | IFNG | IFNGR1 | 0.3967 | 0.7861 | 0.5914 | 0.000999 |
| TCD8em | MacAgPres | IFNG | IFNGR2 | 0.3967 | 0.5706 | 0.4836 | 0.000999 |
| TCD8em | MacAgPres | IL16 | CD4 | 0.4085 | 0.5049 | 0.4567 | 0.000999 |

| | | | | | | | |
|---|---|---|---|---|---|---|---|
| TCD8em | MacAgPres | IL16 | FCGR1A | 0.4085 | 0.769 | 0.5888 | 0.000999 |
| TCD8em | MacAgPres | ITGB1 | CD151 | 0.2557 | 0.2751 | 0.2654 | 0.000999 |
| TCD8em | MacAgPres | ITGB1 | DAB2 | 0.2557 | 0.6335 | 0.4446 | 0.000999 |
| TCD8em | MacAgPres | ITGB1 | PLAUR | 0.2557 | 1.0849 | 0.6703 | 0.000999 |
| TCD8em | MacAgPres | ITGB2 | PLAUR | 1.0018 | 1.0849 | 1.0433 | 0.000999 |
| TCD8em | MacAgPres | LGALS3 | CLEC7A | 0.2689 | 0.6841 | 0.4765 | 0.000999 |
| TCD8em | MacAgPres | LGALS3 | FCGR2A | 0.2689 | 1.2286 | 0.7488 | 0.000999 |
| TCD8em | MacAgPres | LGALS3 | FCGR2B | 0.2689 | 0.578 | 0.4235 | 0.000999 |
| TCD8em | MacAgPres | LRPAP1 | LRP1 | 0.2719 | 0.3779 | 0.3249 | 0.000999 |
| TCD8em | MacAgPres | LRPAP1 | SORL1 | 0.2719 | 0.5617 | 0.4168 | 0.000999 |
| TCD8em | MacAgPres | LTB | ITGB1 | 0.6352 | 0.3801 | 0.5076 | 0.000999 |
| TCD8em | MacAgPres | LTB | TNFRSF1A | 0.6352 | 0.3753 | 0.5052 | 0.000999 |
| TCD8em | MacAgPres | MIF | CD74 | 0.6063 | 5.2676 | 2.937 | 0.000999 |
| TCD8em | MacAgPres | PSAP | LRP1 | 0.5012 | 0.3779 | 0.4396 | 0.000999 |
| TCD8em | MacAgPres | PTPRC | CD4 | 1.7912 | 0.5049 | 1.148 | 0.000999 |
| TCD8em | MacAgPres | RPS19 | C5AR1 | 3.536 | 0.5708 | 2.0534 | 0.000999 |
| TCD8em | MacAgPres | RPS19 | CD4 | 3.536 | 0.5049 | 2.0204 | 0.000999 |
| TCD8em | MacAgPres | SELL | CD4 | 0.258 | 0.5049 | 0.3814 | 0.000999 |
| TCD8em | MacAgPres | TIMP1 | LRP1 | 0.2667 | 0.3779 | 0.3223 | 0.000999 |
| TCD8em | MacAgPres | TNF | TNFRSF1A | 0.2667 | 0.3753 | 0.321 | 0.000999 |
| TCD8em | MacAgPres | TNF | TNFRSF1B | 0.2667 | 0.4694 | 0.3681 | 0.000999 |
| TCD8em | MacAgPres | XCL2 | GPR65 | 0.295 | 0.3478 | 0.3214 | 0.000999 |
| TCD8em | MacAgPres | XCL2 | LPAR6 | 0.295 | 0.6544 | 0.4747 | 0.000999 |
| TCD8em | MacAgPres | XCL2 | PTAFR | 0.295 | 0.2882 | 0.2916 | 0.000999 |
| TCD8em | MacAgPres | IFNG | PTPN6 | 0.3967 | 0.2953 | 0.346 | 0.003 |
| TCD8em | MacAgPres | LGALS3 | ITGB1 | 0.2689 | 0.3801 | 0.3245 | 0.003 |
| TCD8ex | MacAgPres | AIMP1 | RACK1 | 0.2508 | 2.4781 | 1.3644 | 0.000999 |
| TCD8ex | MacAgPres | AIMP1 | RRBP1 | 0.2508 | 0.2634 | 0.2571 | 0.000999 |
| TCD8ex | MacAgPres | ALOX5AP | ALOX5 | 1.5024 | 0.3892 | 0.9458 | 0.000999 |
| TCD8ex | MacAgPres | ANXA1 | C3AR1 | 0.5645 | 0.7287 | 0.6466 | 0.000999 |

| | | | | | | | |
|---|---|---|---|---|---|---|---|
| TCD8ex | MacAgPres | ANXA1 | C5AR1 | 0.5645 | 0.5708 | 0.5676 | 0.000999 |
| TCD8ex | MacAgPres | ANXA1 | CD4 | 0.5645 | 0.5049 | 0.5347 | 0.000999 |
| TCD8ex | MacAgPres | ANXA1 | FPR1 | 0.5645 | 0.4477 | 0.5061 | 0.000999 |
| TCD8ex | MacAgPres | ANXA1 | FPR3 | 0.5645 | 0.2875 | 0.426 | 0.000999 |
| TCD8ex | MacAgPres | APOE | LRP1 | 0.2814 | 0.3779 | 0.3296 | 0.000999 |
| TCD8ex | MacAgPres | APOE | SORL1 | 0.2814 | 0.5617 | 0.4215 | 0.000999 |
| TCD8ex | MacAgPres | ARF1 | CD4 | 0.7835 | 0.5049 | 0.6442 | 0.000999 |
| TCD8ex | MacAgPres | CALM1 | TNFRSF1A | 1.4627 | 0.3753 | 0.919 | 0.000999 |
| TCD8ex | MacAgPres | CALR | LRP1 | 0.9634 | 0.3779 | 0.6706 | 0.000999 |
| TCD8ex | MacAgPres | CD2 | CD4 | 2.0502 | 0.5049 | 1.2775 | 0.000999 |
| TCD8ex | MacAgPres | CD6 | ALCAM | 0.318 | 0.2541 | 0.2861 | 0.000999 |
| TCD8ex | MacAgPres | CXCL13 | C3AR1 | 1.7749 | 0.7287 | 1.2518 | 0.000999 |
| TCD8ex | MacAgPres | CXCL13 | C5AR1 | 1.7749 | 0.5708 | 1.1728 | 0.000999 |
| TCD8ex | MacAgPres | CXCL13 | FPR1 | 1.7749 | 0.4477 | 1.1113 | 0.000999 |
| TCD8ex | MacAgPres | CXCL13 | P2RY13 | 1.7749 | 0.3022 | 1.0385 | 0.000999 |
| TCD8ex | MacAgPres | FASLG | TNFRSF1A | 0.2613 | 0.3753 | 0.3183 | 0.000999 |
| TCD8ex | MacAgPres | HLA-DMA | CD4 | 0.3991 | 0.5049 | 0.452 | 0.000999 |
| TCD8ex | MacAgPres | HMGB1 | LY96 | 1.5414 | 0.726 | 1.1337 | 0.000999 |
| TCD8ex | MacAgPres | HMGB1 | TLR2 | 1.5414 | 0.327 | 0.9342 | 0.000999 |
| TCD8ex | MacAgPres | HMGB2 | DAB2 | 0.85 | 0.6335 | 0.7418 | 0.000999 |
| TCD8ex | MacAgPres | HSP90B1 | LRP1 | 0.5666 | 0.3779 | 0.4723 | 0.000999 |
| TCD8ex | MacAgPres | ICAM3 | ITGB2 | 0.5405 | 1.28 | 0.9103 | 0.000999 |
| TCD8ex | MacAgPres | IFNG | IFNGR1 | 0.8575 | 0.7861 | 0.8218 | 0.000999 |
| TCD8ex | MacAgPres | IFNG | IFNGR2 | 0.8575 | 0.5706 | 0.7141 | 0.000999 |
| TCD8ex | MacAgPres | ITGB1 | CD151 | 0.2995 | 0.2751 | 0.2873 | 0.000999 |
| TCD8ex | MacAgPres | ITGB1 | DAB2 | 0.2995 | 0.6335 | 0.4665 | 0.000999 |
| TCD8ex | MacAgPres | ITGB1 | PLAUR | 0.2995 | 1.0849 | 0.6922 | 0.000999 |
| TCD8ex | MacAgPres | ITGB1 | RACK1 | 0.2995 | 2.4781 | 1.3888 | 0.000999 |
| TCD8ex | MacAgPres | ITGB2 | PLAUR | 0.7635 | 1.0849 | 0.9242 | 0.000999 |
| TCD8ex | MacAgPres | LGALS3 | CLEC7A | 0.3824 | 0.6841 | 0.5333 | 0.000999 |

| | | | | | | | |
|---|---|---|---|---|---|---|---|
| TCD8ex | MacAgPres | LGALS3 | FCGR2A | 0.3824 | 1.2286 | 0.8055 | 0.000999 |
| TCD8ex | MacAgPres | LGALS3 | FCGR2B | 0.3824 | 0.578 | 0.4802 | 0.000999 |
| TCD8ex | MacAgPres | LRPAP1 | LRP1 | 0.2971 | 0.3779 | 0.3375 | 0.000999 |
| TCD8ex | MacAgPres | LRPAP1 | SORL1 | 0.2971 | 0.5617 | 0.4294 | 0.000999 |
| TCD8ex | MacAgPres | LTB | ITGB1 | 0.4576 | 0.3801 | 0.4188 | 0.000999 |
| TCD8ex | MacAgPres | LTB | TNFRSF1A | 0.4576 | 0.3753 | 0.4165 | 0.000999 |
| TCD8ex | MacAgPres | MIF | CD74 | 0.749 | 5.2676 | 3.0083 | 0.000999 |
| TCD8ex | MacAgPres | PTPRC | CD4 | 1.6762 | 0.5049 | 1.0905 | 0.000999 |
| TCD8ex | MacAgPres | RPS19 | C5AR1 | 3.8353 | 0.5708 | 2.203 | 0.000999 |
| TCD8ex | MacAgPres | RPS19 | CD4 | 3.8353 | 0.5049 | 2.1701 | 0.000999 |
| TCD8ex | MacAgPres | VCAM1 | ITGB1 | 0.4462 | 0.3801 | 0.4131 | 0.000999 |
| TCD8ex | MacAgPres | VCAM1 | ITGB2 | 0.4462 | 1.28 | 0.8631 | 0.000999 |
| TCD8ex | MacAgPres | XCL2 | LPAR6 | 0.3157 | 0.6544 | 0.4851 | 0.000999 |
| TCD8ex | MacAgPres | XCL2 | PTAFR | 0.3157 | 0.2882 | 0.302 | 0.000999 |
| TGD | MacAgPres | ARF1 | CD4 | 0.9453 | 0.5049 | 0.7251 | 0.000999 |
| TGD | MacAgPres | ARF1 | CD44 | 0.9453 | 0.7335 | 0.8394 | 0.000999 |
| TGD | MacAgPres | CALM1 | RPSA | 1.7587 | 2.1141 | 1.9364 | 0.000999 |
| TGD | MacAgPres | CALM1 | TNFRSF1A | 1.7587 | 0.3753 | 1.067 | 0.000999 |
| TGD | MacAgPres | CALM1 | TNFRSF1B | 1.7587 | 0.4694 | 1.1141 | 0.000999 |
| TGD | MacAgPres | CD2 | CD4 | 1.4522 | 0.5049 | 0.9785 | 0.000999 |
| TGD | MacAgPres | CD2 | PTPRC | 1.4522 | 0.8308 | 1.1415 | 0.000999 |
| TGD | MacAgPres | CD6 | ALCAM | 0.4684 | 0.2541 | 0.3613 | 0.000999 |
| TGD | MacAgPres | CD99 | CD99 | 1.7154 | 0.5386 | 1.127 | 0.000999 |
| TGD | MacAgPres | HMGB1 | LY96 | 1.7525 | 0.726 | 1.2393 | 0.000999 |
| TGD | MacAgPres | HMGB1 | TLR2 | 1.7525 | 0.327 | 1.0398 | 0.000999 |
| TGD | MacAgPres | ICAM3 | ITGB2 | 0.8816 | 1.28 | 1.0808 | 0.000999 |
| TGD | MacAgPres | IL16 | CD4 | 0.9357 | 0.5049 | 0.7203 | 0.000999 |
| TGD | MacAgPres | IL16 | FCGR1A | 0.9357 | 0.769 | 0.8524 | 0.000999 |
| TGD | MacAgPres | ITGA4 | CD44 | 0.3968 | 0.7335 | 0.5652 | 0.000999 |
| TGD | MacAgPres | ITGA4 | CXCR4 | 0.3968 | 1.7715 | 1.0842 | 0.000999 |

| | | | | | | | |
|---|---|---|---|---|---|---|---|
| TGD | MacAgPres | ITGA4 | RPSA | 0.3968 | 2.1141 | 1.2555 | 0.000999 |
| TGD | MacAgPres | ITGB1 | PLAUR | 0.3657 | 1.0849 | 0.7253 | 0.000999 |
| TGD | MacAgPres | ITGB2 | PLAUR | 1.4444 | 1.0849 | 1.2646 | 0.000999 |
| TGD | MacAgPres | LTB | ITGB1 | 1.6829 | 0.3801 | 1.0315 | 0.000999 |
| TGD | MacAgPres | LTB | TNFRSF1A | 1.6829 | 0.3753 | 1.0291 | 0.000999 |
| TGD | MacAgPres | MIF | CD74 | 1.6335 | 5.2676 | 3.4505 | 0.000999 |
| TGD | MacAgPres | PTPRC | CD4 | 1.8653 | 0.5049 | 1.1851 | 0.000999 |
| TGD | MacAgPres | PTPRC | CD44 | 1.8653 | 0.7335 | 1.2994 | 0.000999 |
| TGD | MacAgPres | PTPRC | CXCR4 | 1.8653 | 1.7715 | 1.8184 | 0.000999 |
| TGD | MacAgPres | RPS19 | C5AR1 | 3.8902 | 0.5708 | 2.2305 | 0.000999 |
| TGD | MacAgPres | RPS19 | CD4 | 3.8902 | 0.5049 | 2.1975 | 0.000999 |
| TGD | MacAgPres | RPS19 | RPSA | 3.8902 | 2.1141 | 3.0022 | 0.000999 |
| TGD | MacAgPres | SELL | CD4 | 0.3555 | 0.5049 | 0.4302 | 0.000999 |
| TGD | MacAgPres | SELL | SELPLG | 0.3555 | 0.2693 | 0.3124 | 0.000999 |
| TGD | MacAgPres | SEMA4D | PTPRC | 0.5658 | 0.8308 | 0.6983 | 0.000999 |
| TGD | MacAgPres | TGFB1 | ITGB1 | 0.7703 | 0.3801 | 0.5752 | 0.000999 |
| TGD | MacAgPres | TGFB1 | TGFBR1 | 0.7703 | 0.2605 | 0.5154 | 0.000999 |
| TGD | MacAgPres | TGFB1 | TGFBR2 | 0.7703 | 0.2588 | 0.5146 | 0.000999 |
| TGD | MacAgPres | TGFB1 | TLR2 | 0.7703 | 0.327 | 0.5487 | 0.000999 |
| TGD | MacAgPres | XCL1 | GPR65 | 1.5822 | 0.3478 | 0.965 | 0.000999 |
| TGD | MacAgPres | XCL1 | LPAR6 | 1.5822 | 0.6544 | 1.1183 | 0.000999 |
| TGD | MacAgPres | XCL1 | PTAFR | 1.5822 | 0.2882 | 0.9352 | 0.000999 |
| TGD | MacAgPres | XCL2 | GPR65 | 0.4864 | 0.3478 | 0.4171 | 0.000999 |
| TGD | MacAgPres | XCL2 | LPAR6 | 0.4864 | 0.6544 | 0.5704 | 0.000999 |
| TGD | MacAgPres | XCL2 | PTAFR | 0.4864 | 0.2882 | 0.3873 | 0.000999 |
| TGD | MacAgPres | HSP90B1 | LRP1 | 0.9338 | 0.3779 | 0.6559 | 0.002 |
| TGD | MacAgPres | HSP90B1 | TNFRSF1B | 0.9338 | 0.4694 | 0.7016 | 0.003 |
| TGD | MacAgPres | HSP90B1 | ITGB1 | 0.9338 | 0.3801 | 0.657 | 0.00699 |

| Supplementary Table 8: Communication results between macrophages and lymphocytes subpopulations, and malignant cells | | | | | | | |
|---|---|---|---|---|---|---|---|
| **MacAgPres vs. malignant cells** | | | | | | | |
| celltype_01 | celltype_02 | ligand | receptor | ligand_mean_expression | receptor_mean_expression | ligand_receptor_mean_expression | pvalue |
| MacAgPres | breasttumor | ANXA1 | CD44 | 0.9542 | 0.3247 | 0.6395 | 0.000999 |
| MacAgPres | breasttumor | APOE | SCARB1 | 2.1025 | 0.2628 | 1.1826 | 0.000999 |
| MacAgPres | breasttumor | APOE | SDC2 | 2.1025 | 1.071 | 1.5868 | 0.000999 |
| MacAgPres | breasttumor | CALM1 | MYLK | 1.0629 | 0.4064 | 0.7346 | 0.000999 |
| MacAgPres | breasttumor | CALM1 | RPSA | 1.0629 | 2.358 | 1.7104 | 0.000999 |
| MacAgPres | breasttumor | CALM2 | MYLK | 1.5958 | 0.4064 | 1.0011 | 0.000999 |
| MacAgPres | breasttumor | CALM3 | MYLK | 0.4699 | 0.4064 | 0.4381 | 0.000999 |
| MacAgPres | breasttumor | CD99 | CD99 | 0.5375 | 0.3871 | 0.4623 | 0.000999 |
| MacAgPres | breasttumor | COL1A1 | CD44 | 1.0301 | 0.3247 | 0.6774 | 0.000999 |
| MacAgPres | breasttumor | COL1A1 | ITGB1 | 1.0301 | 0.2939 | 0.662 | 0.000999 |
| MacAgPres | breasttumor | FN1 | CD44 | 0.9127 | 0.3247 | 0.6187 | 0.000999 |
| MacAgPres | breasttumor | FN1 | ITGB1 | 0.9127 | 0.2939 | 0.6033 | 0.000999 |
| MacAgPres | breasttumor | FN1 | RPSA | 0.9127 | 2.358 | 1.6354 | 0.000999 |
| MacAgPres | breasttumor | FN1 | SDC2 | 0.9127 | 1.071 | 0.9919 | 0.000999 |
| MacAgPres | breasttumor | GRN | TNFRSF1A | 1.7556 | 0.2536 | 1.0046 | 0.000999 |
| MacAgPres | breasttumor | HLA-DRA | SCARB1 | 4.7481 | 0.2628 | 2.5054 | 0.000999 |
| MacAgPres | breasttumor | ITGB1 | CD151 | 0.3809 | 0.3569 | 0.3689 | 0.000999 |
| MacAgPres | breasttumor | ITGB1 | RACK1 | 0.3809 | 2.6334 | 1.5071 | 0.000999 |
| MacAgPres | breasttumor | PTPRC | CD44 | 0.8325 | 0.3247 | 0.5786 | 0.000999 |
| MacAgPres | breasttumor | RTN4 | CD44 | 0.9766 | 0.3247 | 0.6507 | 0.000999 |
| MacAgPres | breasttumor | RTN4 | CD99 | 0.9766 | 0.3871 | 0.6819 | 0.000999 |
| MacAgPres | breasttumor | SERPINA1 | SDC2 | 0.9564 | 1.071 | 1.0137 | 0.000999 |
| MacAgPres | breasttumor | SPP1 | CD44 | 1.3027 | 0.3247 | 0.8137 | 0.000999 |
| MacAgPres | breasttumor | SPP1 | ITGB1 | 1.3027 | 0.2939 | 0.7983 | 0.000999 |
| MacAgPres | breasttumor | TNFSF13B | ITGB1 | 0.8459 | 0.2939 | 0.5699 | 0.000999 |
| MacAgPres | breasttumor | HBEGF | CD44 | 0.485 | 0.3247 | 0.4048 | 0.005 |
| MacAgPres | breasttumor | RPS19 | RPSA | 3.5708 | 2.358 | 2.9644 | 0.00699 |

| | | | | | | | |
|---|---|---|---|---|---|---|---|
| MacAgPres | colorectaltumor | APOE | SDC1 | 2.1025 | 0.2575 | 1.18 | 0.000999 |
| MacAgPres | colorectaltumor | COL1A1 | ITGB1 | 1.0301 | 0.4214 | 0.7257 | 0.000999 |
| MacAgPres | colorectaltumor | COL1A1 | SDC1 | 1.0301 | 0.2575 | 0.6438 | 0.000999 |
| MacAgPres | colorectaltumor | FN1 | ITGB1 | 0.9127 | 0.4214 | 0.6671 | 0.000999 |
| MacAgPres | colorectaltumor | FN1 | RPSA | 0.9127 | 2.251 | 1.5819 | 0.000999 |
| MacAgPres | colorectaltumor | FN1 | SDC1 | 0.9127 | 0.2575 | 0.5851 | 0.000999 |
| MacAgPres | colorectaltumor | ITGB1 | CD151 | 0.3809 | 0.6439 | 0.5124 | 0.000999 |
| MacAgPres | colorectaltumor | ITGB1 | CD46 | 0.3809 | 0.3583 | 0.3696 | 0.000999 |
| MacAgPres | colorectaltumor | ITGB1 | CDH17 | 0.3809 | 0.254 | 0.3174 | 0.000999 |
| MacAgPres | colorectaltumor | ITGB1 | RACK1 | 0.3809 | 2.7146 | 1.5477 | 0.000999 |
| MacAgPres | colorectaltumor | ITGB1 | SDC1 | 0.3809 | 0.2575 | 0.3192 | 0.000999 |
| MacAgPres | colorectaltumor | LGALS3 | ITGB4 | 0.6866 | 0.2913 | 0.489 | 0.000999 |
| MacAgPres | colorectaltumor | MDK | SDC1 | 0.3952 | 0.2575 | 0.3263 | 0.000999 |
| MacAgPres | colorectaltumor | PECAM1 | SDC1 | 0.267 | 0.2575 | 0.2622 | 0.000999 |
| MacAgPres | colorectaltumor | SPP1 | ITGB1 | 1.3027 | 0.4214 | 0.862 | 0.000999 |
| MacAgPres | colorectaltumor | SPP1 | SDC1 | 1.3027 | 0.2575 | 0.7801 | 0.000999 |
| MacAgPres | colorectaltumor | TNFSF13B | ITGB1 | 0.8459 | 0.4214 | 0.6337 | 0.000999 |
| MacAgPres | livertumor | A2M | TNFRSF14 | 0.9998 | 0.2788 | 0.6393 | 0.000999 |
| MacAgPres | livertumor | ANXA1 | CD44 | 0.9542 | 0.8016 | 0.8779 | 0.000999 |
| MacAgPres | livertumor | ANXA1 | MET | 0.9542 | 0.2983 | 0.6262 | 0.000999 |
| MacAgPres | livertumor | ANXA1 | PTPRF | 0.9542 | 0.384 | 0.6691 | 0.000999 |
| MacAgPres | livertumor | APOE | SDC1 | 2.1025 | 0.5354 | 1.319 | 0.000999 |
| MacAgPres | livertumor | APOE | SDC4 | 2.1025 | 0.6068 | 1.3547 | 0.000999 |
| MacAgPres | livertumor | AREG | ERBB3 | 0.4587 | 0.2644 | 0.3615 | 0.000999 |
| MacAgPres | livertumor | ARF1 | INSR | 0.614 | 0.3039 | 0.459 | 0.000999 |
| MacAgPres | livertumor | CALM1 | INSR | 1.0629 | 0.3039 | 0.6834 | 0.000999 |
| MacAgPres | livertumor | CALM1 | RPSA | 1.0629 | 2.6317 | 1.8473 | 0.000999 |
| MacAgPres | livertumor | CALR | ITGA3 | 0.8981 | 0.3913 | 0.6447 | 0.000999 |
| MacAgPres | livertumor | CD99 | EPHB4 | 0.5375 | 0.4535 | 0.4955 | 0.000999 |
| MacAgPres | livertumor | CD99 | INSR | 0.5375 | 0.3039 | 0.4207 | 0.000999 |

| | | | | | | | |
|---|---|---|---|---|---|---|---|
| MacAgPres | livertumor | COL1A1 | CD44 | 1.0301 | 0.8016 | 0.9158 | 0.000999 |
| MacAgPres | livertumor | COL1A1 | ITGA2 | 1.0301 | 0.3983 | 0.7142 | 0.000999 |
| MacAgPres | livertumor | COL1A1 | ITGA3 | 1.0301 | 0.3913 | 0.7107 | 0.000999 |
| MacAgPres | livertumor | COL1A1 | ITGB1 | 1.0301 | 1.6044 | 1.3173 | 0.000999 |
| MacAgPres | livertumor | COL1A1 | SDC1 | 1.0301 | 0.5354 | 0.7828 | 0.000999 |
| MacAgPres | livertumor | FN1 | CD44 | 0.9127 | 0.8016 | 0.8572 | 0.000999 |
| MacAgPres | livertumor | FN1 | ITGA2 | 0.9127 | 0.3983 | 0.6555 | 0.000999 |
| MacAgPres | livertumor | FN1 | ITGA3 | 0.9127 | 0.3913 | 0.652 | 0.000999 |
| MacAgPres | livertumor | FN1 | ITGB1 | 0.9127 | 1.6044 | 1.2586 | 0.000999 |
| MacAgPres | livertumor | FN1 | RPSA | 0.9127 | 2.6317 | 1.7722 | 0.000999 |
| MacAgPres | livertumor | FN1 | SDC1 | 0.9127 | 0.5354 | 0.7241 | 0.000999 |
| MacAgPres | livertumor | FN1 | TFRC | 0.9127 | 0.2899 | 0.6013 | 0.000999 |
| MacAgPres | livertumor | HBEGF | CD44 | 0.485 | 0.8016 | 0.6433 | 0.000999 |
| MacAgPres | livertumor | HBEGF | ERBB3 | 0.485 | 0.2644 | 0.3747 | 0.000999 |
| MacAgPres | livertumor | HLA-A | ERBB3 | 2.913 | 0.2644 | 1.5887 | 0.000999 |
| MacAgPres | livertumor | HLA-DRA | EPHB4 | 4.7481 | 0.4535 | 2.6008 | 0.000999 |
| MacAgPres | livertumor | HLA-E | EPHB4 | 1.9296 | 0.4535 | 1.1915 | 0.000999 |
| MacAgPres | livertumor | HLA-E | ITGA6 | 1.9296 | 0.5211 | 1.2254 | 0.000999 |
| MacAgPres | livertumor | HLA-E | PLXNB2 | 1.9296 | 0.265 | 1.0973 | 0.000999 |
| MacAgPres | livertumor | HMGB2 | CDH1 | 0.4233 | 0.4896 | 0.4565 | 0.000999 |
| MacAgPres | livertumor | HMGB2 | DSG2 | 0.4233 | 0.7479 | 0.5856 | 0.000999 |
| MacAgPres | livertumor | HSP90B1 | ITGB1 | 0.7614 | 1.6044 | 1.1829 | 0.000999 |
| MacAgPres | livertumor | ITGB1 | CD151 | 0.3809 | 1.1085 | 0.7447 | 0.000999 |
| MacAgPres | livertumor | ITGB1 | ITGA2 | 0.3809 | 0.3983 | 0.3896 | 0.000999 |
| MacAgPres | livertumor | ITGB1 | ITGA6 | 0.3809 | 0.5211 | 0.451 | 0.000999 |
| MacAgPres | livertumor | ITGB1 | SDC1 | 0.3809 | 0.5354 | 0.4581 | 0.000999 |
| MacAgPres | livertumor | ITGB1 | VTN | 0.3809 | 0.7399 | 0.5604 | 0.000999 |
| MacAgPres | livertumor | LGALS3 | ITGA2 | 0.6866 | 0.3983 | 0.5425 | 0.000999 |
| MacAgPres | livertumor | LGALS3 | ITGB1 | 0.6866 | 1.6044 | 1.1455 | 0.000999 |
| MacAgPres | livertumor | LGALS3 | ITGB4 | 0.6866 | 0.5518 | 0.6192 | 0.000999 |

| | | | | | | | |
|---|---|---|---|---|---|---|---|
| MacAgPres | livertumor | MDK | SDC1 | 0.3952 | 0.5354 | 0.4653 | 0.000999 |
| MacAgPres | livertumor | MDK | SDC4 | 0.3952 | 0.6068 | 0.501 | 0.000999 |
| MacAgPres | livertumor | NAMPT | INSR | 0.9233 | 0.3039 | 0.6136 | 0.000999 |
| MacAgPres | livertumor | PECAM1 | SDC1 | 0.267 | 0.5354 | 0.4012 | 0.000999 |
| MacAgPres | livertumor | PECAM1 | SDC4 | 0.267 | 0.6068 | 0.4369 | 0.000999 |
| MacAgPres | livertumor | PTPRC | CD44 | 0.8325 | 0.8016 | 0.817 | 0.000999 |
| MacAgPres | livertumor | PTPRC | INSR | 0.8325 | 0.3039 | 0.5682 | 0.000999 |
| MacAgPres | livertumor | PTPRC | MET | 0.8325 | 0.2983 | 0.5654 | 0.000999 |
| MacAgPres | livertumor | RPS19 | RPSA | 3.5708 | 2.6317 | 3.1012 | 0.000999 |
| MacAgPres | livertumor | SPP1 | ITGB1 | 1.3027 | 1.6044 | 1.4536 | 0.000999 |
| MacAgPres | livertumor | SPP1 | SDC1 | 1.3027 | 0.5354 | 0.9191 | 0.000999 |
| MacAgPres | livertumor | TNFSF12 | TNFRSF12A | 0.3091 | 1.0484 | 0.6787 | 0.000999 |
| MacAgPres | livertumor | TNFSF13B | ITGB1 | 0.8459 | 1.6044 | 1.2252 | 0.000999 |
| MacAgPres | livertumor | SPP1 | CD44 | 1.3027 | 0.8016 | 1.0521 | 0.00699 |
| MacAgPres | livertumor | HMGB2 | CD44 | 0.4233 | 0.8016 | 0.6124 | 0.00899 |
| MacAgPres | lungtumor | APOE | SDC1 | 2.1025 | 0.2715 | 1.187 | 0.000999 |
| MacAgPres | lungtumor | APOE | SDC4 | 2.1025 | 0.3516 | 1.2271 | 0.000999 |
| MacAgPres | lungtumor | C3 | CXCR4 | 1.5464 | 0.2684 | 0.9074 | 0.000999 |
| MacAgPres | lungtumor | COL1A1 | CD44 | 1.0301 | 0.3802 | 0.7051 | 0.000999 |
| MacAgPres | lungtumor | COL1A1 | SDC1 | 1.0301 | 0.2715 | 0.6508 | 0.000999 |
| MacAgPres | lungtumor | CXCL16 | CXCR4 | 1.0026 | 0.2684 | 0.6355 | 0.000999 |
| MacAgPres | lungtumor | FN1 | CD44 | 0.9127 | 0.3802 | 0.6465 | 0.000999 |
| MacAgPres | lungtumor | FN1 | RPSA | 0.9127 | 1.6057 | 1.2592 | 0.000999 |
| MacAgPres | lungtumor | FN1 | SDC1 | 0.9127 | 0.2715 | 0.5921 | 0.000999 |
| MacAgPres | lungtumor | HBEGF | CD44 | 0.485 | 0.3802 | 0.4326 | 0.000999 |
| MacAgPres | lungtumor | ITGB1 | CD151 | 0.3809 | 0.6776 | 0.5292 | 0.000999 |
| MacAgPres | lungtumor | ITGB1 | CD46 | 0.3809 | 0.434 | 0.4074 | 0.000999 |
| MacAgPres | lungtumor | ITGB1 | CD47 | 0.3809 | 0.3255 | 0.3532 | 0.000999 |
| MacAgPres | lungtumor | ITGB1 | SDC1 | 0.3809 | 0.2715 | 0.3262 | 0.000999 |
| MacAgPres | lungtumor | PECAM1 | SDC1 | 0.267 | 0.2715 | 0.2693 | 0.000999 |

| | | | | | | | |
|---|---|---|---|---|---|---|---|
| MacAgPres | lungtumor | PECAM1 | SDC4 | 0.267 | 0.3516 | 0.3093 | 0.000999 |
| MacAgPres | lungtumor | PTPRC | CD44 | 0.8325 | 0.3802 | 0.6063 | 0.000999 |
| MacAgPres | lungtumor | PTPRC | CXCR4 | 0.8325 | 0.2684 | 0.5504 | 0.000999 |
| MacAgPres | lungtumor | RTN4 | CD44 | 0.9766 | 0.3802 | 0.6784 | 0.000999 |
| MacAgPres | lungtumor | RTN4 | CD99 | 0.9766 | 0.5938 | 0.7852 | 0.000999 |
| MacAgPres | lungtumor | TNFSF12 | TNFRSF12A | 0.3091 | 0.441 | 0.375 | 0.000999 |
| MacAgPres | melanomatumo | A2M | TNFRSF14 | 0.9998 | 0.5069 | 0.7533 | 0.000999 |
| MacAgPres | melanomatumo | ANXA1 | CD44 | 0.9542 | 1.2653 | 1.1098 | 0.000999 |
| MacAgPres | melanomatumo | ANXA1 | EDNRB | 0.9542 | 0.5666 | 0.7604 | 0.000999 |
| MacAgPres | melanomatumo | CALM1 | RPSA | 1.0629 | 3.7787 | 2.4208 | 0.000999 |
| MacAgPres | melanomatumo | CALR | CD44 | 0.8981 | 1.2653 | 1.0817 | 0.000999 |
| MacAgPres | melanomatumo | COL1A1 | CD44 | 1.0301 | 1.2653 | 1.1477 | 0.000999 |
| MacAgPres | melanomatumo | FN1 | CD44 | 0.9127 | 1.2653 | 1.089 | 0.000999 |
| MacAgPres | melanomatumo | FN1 | RPSA | 0.9127 | 3.7787 | 2.3457 | 0.000999 |
| MacAgPres | melanomatumo | HBEGF | CD44 | 0.485 | 1.2653 | 0.8752 | 0.000999 |
| MacAgPres | melanomatumo | HMGB2 | CD44 | 0.4233 | 1.2653 | 0.8443 | 0.000999 |
| MacAgPres | melanomatumo | ITGB1 | RACK1 | 0.3809 | 4.145 | 2.2629 | 0.000999 |
| MacAgPres | melanomatumo | PTPRC | CD44 | 0.8325 | 1.2653 | 1.0489 | 0.000999 |
| MacAgPres | melanomatumo | RPS19 | RPSA | 3.5708 | 3.7787 | 3.6747 | 0.000999 |
| MacAgPres | melanomatumo | RTN4 | CD44 | 0.9766 | 1.2653 | 1.121 | 0.000999 |
| MacAgPres | melanomatumo | SPP1 | CD44 | 1.3027 | 1.2653 | 1.284 | 0.000999 |
| MacAgPres | melanomatumo | ARF1 | CD44 | 0.614 | 1.2653 | 0.9397 | 0.002 |
| MacAgPres | melanomatumo | GSTP1 | CD44 | 1.4178 | 1.2653 | 1.3416 | 0.002 |
| MacAgPres | melanomatumo | NAMPT | ANXA6 | 0.9233 | 0.3625 | 0.6429 | 0.003 |
| MacAgPres | melanomatumo | SPP1 | NCSTN | 1.3027 | 0.2614 | 0.7821 | 0.00699 |
| MacAgPres | ovarytumor | AREG | FOLR1 | 0.4587 | 0.5619 | 0.5103 | 0.000999 |
| MacAgPres | ovarytumor | C3 | CXCR4 | 1.5464 | 0.3688 | 0.9576 | 0.000999 |
| MacAgPres | ovarytumor | CALM1 | RPSA | 1.0629 | 2.9542 | 2.0085 | 0.000999 |
| MacAgPres | ovarytumor | COL1A1 | ITGB1 | 1.0301 | 0.2783 | 0.6542 | 0.000999 |
| MacAgPres | ovarytumor | CXCL16 | CXCR4 | 1.0026 | 0.3688 | 0.6857 | 0.000999 |

| | | | | | | | |
|---|---|---|---|---|---|---|---|
| MacAgPres | ovarytumor | FN1 | ITGB1 | 0.9127 | 0.2783 | 0.5955 | 0.000999 |
| MacAgPres | ovarytumor | FN1 | RPSA | 0.9127 | 2.9542 | 1.9335 | 0.000999 |
| MacAgPres | ovarytumor | HSP90B1 | ITGB1 | 0.7614 | 0.2783 | 0.5198 | 0.000999 |
| MacAgPres | ovarytumor | ITGB1 | CD151 | 0.3809 | 0.4981 | 0.4395 | 0.000999 |
| MacAgPres | ovarytumor | ITGB1 | CD47 | 0.3809 | 0.4931 | 0.437 | 0.000999 |
| MacAgPres | ovarytumor | ITGB1 | RACK1 | 0.3809 | 2.9543 | 1.6676 | 0.000999 |
| MacAgPres | ovarytumor | LGALS3 | ITGB1 | 0.6866 | 0.2783 | 0.4825 | 0.000999 |
| MacAgPres | ovarytumor | PTPRC | CXCR4 | 0.8325 | 0.3688 | 0.6006 | 0.000999 |
| MacAgPres | ovarytumor | SPP1 | ITGB1 | 1.3027 | 0.2783 | 0.7905 | 0.000999 |
| MacAgPres | ovarytumor | TNFSF12 | TNFRSF12A | 0.3091 | 0.3901 | 0.3496 | 0.000999 |
| MacAgPres | ovarytumor | TNFSF13B | ITGB1 | 0.8459 | 0.2783 | 0.5621 | 0.000999 |
| breasttumor | MacAgPres | ANXA1 | C3AR1 | 0.4092 | 0.729 | 0.5691 | 0.000999 |
| breasttumor | MacAgPres | ANXA1 | CXCR4 | 0.4092 | 1.7737 | 1.0914 | 0.000999 |
| breasttumor | MacAgPres | APP | ALOX5 | 0.5409 | 0.39 | 0.4655 | 0.000999 |
| breasttumor | MacAgPres | APP | C3AR1 | 0.5409 | 0.729 | 0.635 | 0.000999 |
| breasttumor | MacAgPres | APP | C5AR1 | 0.5409 | 0.5707 | 0.5558 | 0.000999 |
| breasttumor | MacAgPres | APP | CXCR4 | 0.5409 | 1.7737 | 1.1573 | 0.000999 |
| breasttumor | MacAgPres | APP | FPR1 | 0.5409 | 0.4486 | 0.4948 | 0.000999 |
| breasttumor | MacAgPres | APP | LRP1 | 0.5409 | 0.3786 | 0.4598 | 0.000999 |
| breasttumor | MacAgPres | APP | SORL1 | 0.5409 | 0.5618 | 0.5514 | 0.000999 |
| breasttumor | MacAgPres | APP | TGFBR2 | 0.5409 | 0.2584 | 0.3997 | 0.000999 |
| breasttumor | MacAgPres | ARF1 | CD4 | 1.6219 | 0.5059 | 1.0639 | 0.000999 |
| breasttumor | MacAgPres | ARF1 | CD44 | 1.6219 | 0.734 | 1.178 | 0.000999 |
| breasttumor | MacAgPres | CALM1 | TNFRSF1B | 0.9015 | 0.4694 | 0.6855 | 0.000999 |
| breasttumor | MacAgPres | CALR | CD44 | 1.1835 | 0.734 | 0.9588 | 0.000999 |
| breasttumor | MacAgPres | CALR | LRP1 | 1.1835 | 0.3786 | 0.7811 | 0.000999 |
| breasttumor | MacAgPres | CD99 | CD99 | 0.3871 | 0.5375 | 0.4623 | 0.000999 |
| breasttumor | MacAgPres | COL2A1 | CD44 | 0.3935 | 0.734 | 0.5638 | 0.000999 |
| breasttumor | MacAgPres | COL2A1 | ITGB1 | 0.3935 | 0.3809 | 0.3872 | 0.000999 |
| breasttumor | MacAgPres | EFNA1 | TGFBR1 | 0.3824 | 0.2595 | 0.3209 | 0.000999 |

| | | | | | | | |
|---|---|---|---|---|---|---|---|
| breasttumor | MacAgPres | GSTP1 | CD44 | 2.5435 | 0.734 | 1.6388 | 0.000999 |
| breasttumor | MacAgPres | HEBP1 | C3AR1 | 0.4062 | 0.729 | 0.5676 | 0.000999 |
| breasttumor | MacAgPres | HEBP1 | C5AR1 | 0.4062 | 0.5707 | 0.4885 | 0.000999 |
| breasttumor | MacAgPres | HEBP1 | CXCR4 | 0.4062 | 1.7737 | 1.0899 | 0.000999 |
| breasttumor | MacAgPres | HEBP1 | FPR1 | 0.4062 | 0.4486 | 0.4274 | 0.000999 |
| breasttumor | MacAgPres | HEBP1 | FPR3 | 0.4062 | 0.2881 | 0.3471 | 0.000999 |
| breasttumor | MacAgPres | HMGB1 | LY96 | 1.1815 | 0.7263 | 0.9539 | 0.000999 |
| breasttumor | MacAgPres | HMGB1 | TLR2 | 1.1815 | 0.3254 | 0.7534 | 0.000999 |
| breasttumor | MacAgPres | HMGB2 | CD44 | 0.3139 | 0.734 | 0.524 | 0.000999 |
| breasttumor | MacAgPres | HMGB2 | DAB2 | 0.3139 | 0.6336 | 0.4738 | 0.000999 |
| breasttumor | MacAgPres | HSP90B1 | LRP1 | 0.7786 | 0.3786 | 0.5786 | 0.000999 |
| breasttumor | MacAgPres | HSP90B1 | TNFRSF1B | 0.7786 | 0.4694 | 0.624 | 0.000999 |
| breasttumor | MacAgPres | ITGB1 | CD47 | 0.2939 | 0.3834 | 0.3386 | 0.000999 |
| breasttumor | MacAgPres | ITGB1 | DAB2 | 0.2939 | 0.6336 | 0.4637 | 0.000999 |
| breasttumor | MacAgPres | ITGB1 | PLAUR | 0.2939 | 1.0857 | 0.6898 | 0.000999 |
| breasttumor | MacAgPres | LGALS3 | CLEC7A | 0.8391 | 0.683 | 0.761 | 0.000999 |
| breasttumor | MacAgPres | LGALS3 | FCGR2A | 0.8391 | 1.2301 | 1.0346 | 0.000999 |
| breasttumor | MacAgPres | LGALS3 | FCGR2B | 0.8391 | 0.5782 | 0.7086 | 0.000999 |
| breasttumor | MacAgPres | LGALS3 | ITGB1 | 0.8391 | 0.3809 | 0.61 | 0.000999 |
| breasttumor | MacAgPres | LGALS3 | TGFBR2 | 0.8391 | 0.2584 | 0.5487 | 0.000999 |
| breasttumor | MacAgPres | LRPAP1 | LRP1 | 0.2756 | 0.3786 | 0.3271 | 0.000999 |
| breasttumor | MacAgPres | LRPAP1 | SORL1 | 0.2756 | 0.5618 | 0.4187 | 0.000999 |
| breasttumor | MacAgPres | MDK | LRP1 | 0.592 | 0.3786 | 0.4853 | 0.000999 |
| breasttumor | MacAgPres | MIF | CD74 | 3.2716 | 5.2679 | 4.2698 | 0.000999 |
| breasttumor | MacAgPres | NPW | C3AR1 | 0.5478 | 0.729 | 0.6384 | 0.000999 |
| breasttumor | MacAgPres | NPW | C5AR1 | 0.5478 | 0.5707 | 0.5593 | 0.000999 |
| breasttumor | MacAgPres | NPW | CXCR4 | 0.5478 | 1.7737 | 1.1608 | 0.000999 |
| breasttumor | MacAgPres | NPW | FPR1 | 0.5478 | 0.4486 | 0.4982 | 0.000999 |
| breasttumor | MacAgPres | RPS19 | C5AR1 | 3.691 | 0.5707 | 2.1309 | 0.000999 |
| breasttumor | MacAgPres | RPS19 | CD4 | 3.691 | 0.5059 | 2.0984 | 0.000999 |

| | | | | | | | |
|---|---|---|---|---|---|---|---|
| breasttumor | MacAgPres | SERPINE2 | LRP1 | 0.6001 | 0.3786 | 0.4894 | 0.000999 |
| breasttumor | MacAgPres | TF | CD4 | 0.2688 | 0.5059 | 0.3873 | 0.000999 |
| colorectaltumor | MacAgPres | AIMP1 | RACK1 | 0.5265 | 2.4767 | 1.5016 | 0.000999 |
| colorectaltumor | MacAgPres | AIMP1 | RRBP1 | 0.5265 | 0.2629 | 0.3947 | 0.000999 |
| colorectaltumor | MacAgPres | APP | ALOX5 | 0.4966 | 0.39 | 0.4433 | 0.000999 |
| colorectaltumor | MacAgPres | APP | C3AR1 | 0.4966 | 0.729 | 0.6128 | 0.000999 |
| colorectaltumor | MacAgPres | APP | C5AR1 | 0.4966 | 0.5707 | 0.5337 | 0.000999 |
| colorectaltumor | MacAgPres | APP | CXCR4 | 0.4966 | 1.7737 | 1.1352 | 0.000999 |
| colorectaltumor | MacAgPres | APP | FPR1 | 0.4966 | 0.4486 | 0.4726 | 0.000999 |
| colorectaltumor | MacAgPres | APP | LRP1 | 0.4966 | 0.3786 | 0.4376 | 0.000999 |
| colorectaltumor | MacAgPres | APP | SORL1 | 0.4966 | 0.5618 | 0.5292 | 0.000999 |
| colorectaltumor | MacAgPres | APP | TGFBR2 | 0.4966 | 0.2584 | 0.3775 | 0.000999 |
| colorectaltumor | MacAgPres | GSTP1 | CD44 | 2.8039 | 0.734 | 1.769 | 0.000999 |
| colorectaltumor | MacAgPres | HMGB1 | LY96 | 2.155 | 0.7263 | 1.4406 | 0.000999 |
| colorectaltumor | MacAgPres | HMGB1 | TLR2 | 2.155 | 0.3254 | 1.2402 | 0.000999 |
| colorectaltumor | MacAgPres | HMGB2 | CD44 | 0.5347 | 0.734 | 0.6344 | 0.000999 |
| colorectaltumor | MacAgPres | HMGB2 | DAB2 | 0.5347 | 0.6336 | 0.5842 | 0.000999 |
| colorectaltumor | MacAgPres | ITGB1 | PLAUR | 0.4214 | 1.0857 | 0.7535 | 0.000999 |
| colorectaltumor | MacAgPres | LGALS3 | CLEC7A | 1.5563 | 0.683 | 1.1196 | 0.000999 |
| colorectaltumor | MacAgPres | LGALS3 | FCGR2A | 1.5563 | 1.2301 | 1.3932 | 0.000999 |
| colorectaltumor | MacAgPres | LGALS3 | FCGR2B | 1.5563 | 0.5782 | 1.0672 | 0.000999 |
| colorectaltumor | MacAgPres | LGALS3 | ITGB1 | 1.5563 | 0.3809 | 0.9686 | 0.000999 |
| colorectaltumor | MacAgPres | LGALS3 | TGFBR2 | 1.5563 | 0.2584 | 0.9073 | 0.000999 |
| colorectaltumor | MacAgPres | MDK | LRP1 | 0.7548 | 0.3786 | 0.5667 | 0.000999 |
| colorectaltumor | MacAgPres | MIF | CD74 | 1.6741 | 5.2679 | 3.471 | 0.000999 |
| colorectaltumor | MacAgPres | RPS19 | C5AR1 | 4.0956 | 0.5707 | 2.3331 | 0.000999 |
| colorectaltumor | MacAgPres | RPS19 | CD4 | 4.0956 | 0.5059 | 2.3007 | 0.000999 |
| colorectaltumor | MacAgPres | RPS19 | RPSA | 4.0956 | 2.1132 | 3.1044 | 0.000999 |
| colorectaltumor | MacAgPres | ITGB1 | DAB2 | 0.4214 | 0.6336 | 0.5275 | 0.002 |
| colorectaltumor | MacAgPres | CALM1 | TNFRSF1B | 1.1676 | 0.4694 | 0.8185 | 0.003 |

| | | | | | | | |
|---|---|---|---|---|---|---|---|
| colorectaltumor | MacAgPres | ARF1 | CD4 | 0.6489 | 0.5059 | 0.5774 | 0.00799 |
| livertumor | MacAgPres | AGRN | LRP1 | 0.3492 | 0.3786 | 0.3639 | 0.000999 |
| livertumor | MacAgPres | AGT | ATP6AP2 | 0.4701 | 0.777 | 0.6235 | 0.000999 |
| livertumor | MacAgPres | AGT | C3AR1 | 0.4701 | 0.729 | 0.5995 | 0.000999 |
| livertumor | MacAgPres | AGT | C5AR1 | 0.4701 | 0.5707 | 0.5204 | 0.000999 |
| livertumor | MacAgPres | AGT | CXCR4 | 0.4701 | 1.7737 | 1.1219 | 0.000999 |
| livertumor | MacAgPres | AGT | FPR1 | 0.4701 | 0.4486 | 0.4593 | 0.000999 |
| livertumor | MacAgPres | AGT | GPR65 | 0.4701 | 0.3485 | 0.4093 | 0.000999 |
| livertumor | MacAgPres | AGT | LPAR6 | 0.4701 | 0.6546 | 0.5623 | 0.000999 |
| livertumor | MacAgPres | AGT | P2RY13 | 0.4701 | 0.3012 | 0.3856 | 0.000999 |
| livertumor | MacAgPres | AGT | PTAFR | 0.4701 | 0.2888 | 0.3794 | 0.000999 |
| livertumor | MacAgPres | AIMP1 | RRBP1 | 0.3298 | 0.2629 | 0.2964 | 0.000999 |
| livertumor | MacAgPres | APOA1 | LRP1 | 0.2888 | 0.3786 | 0.3337 | 0.000999 |
| livertumor | MacAgPres | APOA1 | MSR1 | 0.2888 | 0.6006 | 0.4447 | 0.000999 |
| livertumor | MacAgPres | APP | ALOX5 | 0.5626 | 0.39 | 0.4763 | 0.000999 |
| livertumor | MacAgPres | APP | C3AR1 | 0.5626 | 0.729 | 0.6458 | 0.000999 |
| livertumor | MacAgPres | APP | C5AR1 | 0.5626 | 0.5707 | 0.5666 | 0.000999 |
| livertumor | MacAgPres | APP | CXCR4 | 0.5626 | 1.7737 | 1.1681 | 0.000999 |
| livertumor | MacAgPres | APP | FPR1 | 0.5626 | 0.4486 | 0.5056 | 0.000999 |
| livertumor | MacAgPres | APP | LRP1 | 0.5626 | 0.3786 | 0.4706 | 0.000999 |
| livertumor | MacAgPres | APP | SORL1 | 0.5626 | 0.5618 | 0.5622 | 0.000999 |
| livertumor | MacAgPres | APP | TGFBR2 | 0.5626 | 0.2584 | 0.4105 | 0.000999 |
| livertumor | MacAgPres | ARF1 | CD4 | 0.6936 | 0.5059 | 0.5997 | 0.000999 |
| livertumor | MacAgPres | CALR | LRP1 | 1.0416 | 0.3786 | 0.7101 | 0.000999 |
| livertumor | MacAgPres | CCL20 | C3AR1 | 1.0877 | 0.729 | 0.9084 | 0.000999 |
| livertumor | MacAgPres | CCL20 | C5AR1 | 1.0877 | 0.5707 | 0.8292 | 0.000999 |
| livertumor | MacAgPres | CCL20 | CXCR4 | 1.0877 | 1.7737 | 1.4307 | 0.000999 |
| livertumor | MacAgPres | CCL20 | FPR1 | 1.0877 | 0.4486 | 0.7681 | 0.000999 |
| livertumor | MacAgPres | CCL20 | P2RY13 | 1.0877 | 0.3012 | 0.6944 | 0.000999 |
| livertumor | MacAgPres | CDH1 | RACK1 | 0.4896 | 2.4767 | 1.4832 | 0.000999 |

| | | | | | | | |
|---|---|---|---|---|---|---|---|
| livertumor | MacAgPres | CXCL2 | CXCR4 | 0.3364 | 1.7737 | 1.055 | 0.000999 |
| livertumor | MacAgPres | CXCL3 | CXCR4 | 0.2543 | 1.7737 | 1.014 | 0.000999 |
| livertumor | MacAgPres | DSG2 | CD44 | 0.7479 | 0.734 | 0.741 | 0.000999 |
| livertumor | MacAgPres | DSG2 | CXCR4 | 0.7479 | 1.7737 | 1.2608 | 0.000999 |
| livertumor | MacAgPres | DSG2 | DAB2 | 0.7479 | 0.6336 | 0.6908 | 0.000999 |
| livertumor | MacAgPres | EFNA1 | TGFBR1 | 0.3633 | 0.2595 | 0.3114 | 0.000999 |
| livertumor | MacAgPres | FGA | ITGB2 | 0.2688 | 1.2797 | 0.7742 | 0.000999 |
| livertumor | MacAgPres | GSTP1 | CD44 | 1.7219 | 0.734 | 1.228 | 0.000999 |
| livertumor | MacAgPres | HMGB1 | LY96 | 1.4361 | 0.7263 | 1.0812 | 0.000999 |
| livertumor | MacAgPres | HMGB1 | TLR2 | 1.4361 | 0.3254 | 0.8807 | 0.000999 |
| livertumor | MacAgPres | HSP90B1 | LRP1 | 0.8951 | 0.3786 | 0.6369 | 0.000999 |
| livertumor | MacAgPres | HSP90B1 | TNFRSF1B | 0.8951 | 0.4694 | 0.6823 | 0.000999 |
| livertumor | MacAgPres | HSPG2 | LRP1 | 0.2746 | 0.3786 | 0.3266 | 0.000999 |
| livertumor | MacAgPres | ITGB1 | CD151 | 1.6044 | 0.2757 | 0.9401 | 0.000999 |
| livertumor | MacAgPres | ITGB1 | CD47 | 1.6044 | 0.3834 | 0.9939 | 0.000999 |
| livertumor | MacAgPres | ITGB1 | DAB2 | 1.6044 | 0.6336 | 1.119 | 0.000999 |
| livertumor | MacAgPres | ITGB1 | PLAUR | 1.6044 | 1.0857 | 1.3451 | 0.000999 |
| livertumor | MacAgPres | ITGB1 | RACK1 | 1.6044 | 2.4767 | 2.0406 | 0.000999 |
| livertumor | MacAgPres | LAMC2 | CD151 | 0.4847 | 0.2757 | 0.3802 | 0.000999 |
| livertumor | MacAgPres | LAMC2 | ITGB1 | 0.4847 | 0.3809 | 0.4328 | 0.000999 |
| livertumor | MacAgPres | LGALS3 | CLEC7A | 0.8172 | 0.683 | 0.7501 | 0.000999 |
| livertumor | MacAgPres | LGALS3 | FCGR2A | 0.8172 | 1.2301 | 1.0237 | 0.000999 |
| livertumor | MacAgPres | LGALS3 | FCGR2B | 0.8172 | 0.5782 | 0.6977 | 0.000999 |
| livertumor | MacAgPres | LGALS3 | TGFBR2 | 0.8172 | 0.2584 | 0.5378 | 0.000999 |
| livertumor | MacAgPres | MDK | LRP1 | 1.6101 | 0.3786 | 0.9944 | 0.000999 |
| livertumor | MacAgPres | MIF | CD74 | 1.7638 | 5.2679 | 3.5159 | 0.000999 |
| livertumor | MacAgPres | NTS | GPR65 | 0.3425 | 0.3485 | 0.3455 | 0.000999 |
| livertumor | MacAgPres | NTS | LPAR6 | 0.3425 | 0.6546 | 0.4985 | 0.000999 |
| livertumor | MacAgPres | NTS | PTAFR | 0.3425 | 0.2888 | 0.3156 | 0.000999 |
| livertumor | MacAgPres | OCLN | TGFBR1 | 0.3091 | 0.2595 | 0.2843 | 0.000999 |

| | | | | | | | |
|---|---|---|---|---|---|---|---|
| livertumor | MacAgPres | OCLN | TGFBR2 | 0.3091 | 0.2584 | 0.2837 | 0.000999 |
| livertumor | MacAgPres | PTPRF | RACK1 | 0.384 | 2.4767 | 1.4304 | 0.000999 |
| livertumor | MacAgPres | PTPRF | TGFBR2 | 0.384 | 0.2584 | 0.3212 | 0.000999 |
| livertumor | MacAgPres | RPS19 | C5AR1 | 4.0772 | 0.5707 | 2.324 | 0.000999 |
| livertumor | MacAgPres | RPS19 | CD4 | 4.0772 | 0.5059 | 2.2915 | 0.000999 |
| livertumor | MacAgPres | RPS19 | RPSA | 4.0772 | 2.1132 | 3.0952 | 0.000999 |
| livertumor | MacAgPres | SERPINA1 | LRP1 | 1.604 | 0.3786 | 0.9913 | 0.000999 |
| livertumor | MacAgPres | TF | CD4 | 0.523 | 0.5059 | 0.5144 | 0.000999 |
| livertumor | MacAgPres | TFPI | LRP1 | 0.5043 | 0.3786 | 0.4415 | 0.000999 |
| livertumor | MacAgPres | VEGFA | AXL | 0.5485 | 0.3964 | 0.4724 | 0.000999 |
| livertumor | MacAgPres | VTN | ITGB1 | 0.7399 | 0.3809 | 0.5604 | 0.000999 |
| livertumor | MacAgPres | VTN | PLAUR | 0.7399 | 1.0857 | 0.9128 | 0.000999 |
| lungtumor | MacAgPres | AIMP1 | RACK1 | 0.3345 | 2.4767 | 1.4056 | 0.000999 |
| lungtumor | MacAgPres | AIMP1 | RRBP1 | 0.3345 | 0.2629 | 0.2987 | 0.000999 |
| lungtumor | MacAgPres | ANXA1 | C3AR1 | 1.4026 | 0.729 | 1.0658 | 0.000999 |
| lungtumor | MacAgPres | ANXA1 | C5AR1 | 1.4026 | 0.5707 | 0.9867 | 0.000999 |
| lungtumor | MacAgPres | ANXA1 | CD4 | 1.4026 | 0.5059 | 0.9543 | 0.000999 |
| lungtumor | MacAgPres | ANXA1 | CD44 | 1.4026 | 0.734 | 1.0683 | 0.000999 |
| lungtumor | MacAgPres | ANXA1 | CXCR4 | 1.4026 | 1.7737 | 1.5882 | 0.000999 |
| lungtumor | MacAgPres | ANXA1 | FPR1 | 1.4026 | 0.4486 | 0.9256 | 0.000999 |
| lungtumor | MacAgPres | ANXA1 | FPR3 | 1.4026 | 0.2881 | 0.8454 | 0.000999 |
| lungtumor | MacAgPres | ARF1 | CD4 | 0.8791 | 0.5059 | 0.6925 | 0.000999 |
| lungtumor | MacAgPres | ARF1 | CD44 | 0.8791 | 0.734 | 0.8066 | 0.000999 |
| lungtumor | MacAgPres | CALM1 | RPSA | 0.8426 | 2.1132 | 1.4779 | 0.000999 |
| lungtumor | MacAgPres | CALM1 | TNFRSF1B | 0.8426 | 0.4694 | 0.656 | 0.000999 |
| lungtumor | MacAgPres | CALR | CD44 | 0.7795 | 0.734 | 0.7568 | 0.000999 |
| lungtumor | MacAgPres | CALR | LRP1 | 0.7795 | 0.3786 | 0.5791 | 0.000999 |
| lungtumor | MacAgPres | CCL20 | C3AR1 | 0.2885 | 0.729 | 0.5088 | 0.000999 |
| lungtumor | MacAgPres | CCL20 | C5AR1 | 0.2885 | 0.5707 | 0.4296 | 0.000999 |
| lungtumor | MacAgPres | CCL20 | CXCR4 | 0.2885 | 1.7737 | 1.0311 | 0.000999 |

| | | | | | | | |
|---|---|---|---|---|---|---|---|
| lungtumor | MacAgPres | CCL20 | FPR1 | 0.2885 | 0.4486 | 0.3686 | 0.000999 |
| lungtumor | MacAgPres | CCL20 | P2RY13 | 0.2885 | 0.3012 | 0.2949 | 0.000999 |
| lungtumor | MacAgPres | CXCL10 | C3AR1 | 0.3417 | 0.729 | 0.5354 | 0.000999 |
| lungtumor | MacAgPres | CXCL10 | C5AR1 | 0.3417 | 0.5707 | 0.4562 | 0.000999 |
| lungtumor | MacAgPres | CXCL10 | CXCR4 | 0.3417 | 1.7737 | 1.0577 | 0.000999 |
| lungtumor | MacAgPres | CXCL10 | FPR1 | 0.3417 | 0.4486 | 0.3951 | 0.000999 |
| lungtumor | MacAgPres | CXCL10 | P2RY13 | 0.3417 | 0.3012 | 0.3214 | 0.000999 |
| lungtumor | MacAgPres | EFNA1 | TGFBR1 | 0.3852 | 0.2595 | 0.3223 | 0.000999 |
| lungtumor | MacAgPres | FGA | ITGB1 | 0.4445 | 0.3809 | 0.4127 | 0.000999 |
| lungtumor | MacAgPres | FGA | ITGB2 | 0.4445 | 1.2797 | 0.8621 | 0.000999 |
| lungtumor | MacAgPres | GSTP1 | CD44 | 2.4093 | 0.734 | 1.5716 | 0.000999 |
| lungtumor | MacAgPres | HMGB1 | LY96 | 1.1943 | 0.7263 | 0.9603 | 0.000999 |
| lungtumor | MacAgPres | HMGB1 | TLR2 | 1.1943 | 0.3254 | 0.7598 | 0.000999 |
| lungtumor | MacAgPres | HMGB2 | CD44 | 0.5593 | 0.734 | 0.6467 | 0.000999 |
| lungtumor | MacAgPres | HMGB2 | DAB2 | 0.5593 | 0.6336 | 0.5965 | 0.000999 |
| lungtumor | MacAgPres | HSP90B1 | LRP1 | 0.6141 | 0.3786 | 0.4964 | 0.000999 |
| lungtumor | MacAgPres | HSP90B1 | TNFRSF1B | 0.6141 | 0.4694 | 0.5418 | 0.000999 |
| lungtumor | MacAgPres | LGALS3 | CLEC7A | 0.9915 | 0.683 | 0.8372 | 0.000999 |
| lungtumor | MacAgPres | LGALS3 | FCGR2A | 0.9915 | 1.2301 | 1.1108 | 0.000999 |
| lungtumor | MacAgPres | LGALS3 | FCGR2B | 0.9915 | 0.5782 | 0.7849 | 0.000999 |
| lungtumor | MacAgPres | LGALS3 | ITGB1 | 0.9915 | 0.3809 | 0.6862 | 0.000999 |
| lungtumor | MacAgPres | LGALS3 | TGFBR2 | 0.9915 | 0.2584 | 0.6249 | 0.000999 |
| lungtumor | MacAgPres | MDK | LRP1 | 1.2896 | 0.3786 | 0.8341 | 0.000999 |
| lungtumor | MacAgPres | MIF | CD74 | 1.4585 | 5.2679 | 3.3632 | 0.000999 |
| lungtumor | MacAgPres | RPS19 | C5AR1 | 3.5104 | 0.5707 | 2.0406 | 0.000999 |
| lungtumor | MacAgPres | RPS19 | CD4 | 3.5104 | 0.5059 | 2.0082 | 0.000999 |
| lungtumor | MacAgPres | RPS19 | RPSA | 3.5104 | 2.1132 | 2.8118 | 0.000999 |
| lungtumor | MacAgPres | RTN4 | CD44 | 0.4013 | 0.734 | 0.5677 | 0.000999 |
| lungtumor | MacAgPres | SAA1 | C3AR1 | 0.2539 | 0.729 | 0.4915 | 0.000999 |
| lungtumor | MacAgPres | SAA1 | C5AR1 | 0.2539 | 0.5707 | 0.4123 | 0.000999 |

| | | | | | | | |
|---|---|---|---|---|---|---|---|
| lungtumor | MacAgPres | SAA1 | CXCR4 | 0.2539 | 1.7737 | 1.0138 | 0.000999 |
| lungtumor | MacAgPres | SAA1 | FPR1 | 0.2539 | 0.4486 | 0.3512 | 0.000999 |
| lungtumor | MacAgPres | SAA1 | GPR65 | 0.2539 | 0.3485 | 0.3012 | 0.000999 |
| lungtumor | MacAgPres | SAA1 | LPAR6 | 0.2539 | 0.6546 | 0.4542 | 0.000999 |
| lungtumor | MacAgPres | SAA1 | P2RY13 | 0.2539 | 0.3012 | 0.2775 | 0.000999 |
| lungtumor | MacAgPres | SAA1 | PTAFR | 0.2539 | 0.2888 | 0.2713 | 0.000999 |
| lungtumor | MacAgPres | SAA1 | SELENOS | 0.2539 | 0.3478 | 0.3009 | 0.000999 |
| lungtumor | MacAgPres | SERPINA1 | LRP1 | 0.3624 | 0.3786 | 0.3705 | 0.000999 |
| lungtumor | MacAgPres | SPP1 | CD44 | 1.4874 | 0.734 | 1.1107 | 0.000999 |
| lungtumor | MacAgPres | SPP1 | ITGB1 | 1.4874 | 0.3809 | 0.9341 | 0.000999 |
| lungtumor | MacAgPres | TFPI | LRP1 | 0.3615 | 0.3786 | 0.3701 | 0.000999 |
| lungtumor | MacAgPres | TIMP1 | LRP1 | 0.6195 | 0.3786 | 0.4991 | 0.000999 |
| lungtumor | MacAgPres | HSP90B1 | ITGB1 | 0.6141 | 0.3809 | 0.4975 | 0.003 |
| lungtumor | MacAgPres | CALM1 | TNFRSF1A | 0.8426 | 0.3745 | 0.6085 | 0.004 |
| melanomatumo | MacAgPres | ARF1 | CD4 | 1.8545 | 0.5059 | 1.1802 | 0.000999 |
| melanomatumo | MacAgPres | ARF1 | CD44 | 1.8545 | 0.734 | 1.2943 | 0.000999 |
| melanomatumo | MacAgPres | GSTP1 | CD44 | 2.3341 | 0.734 | 1.5341 | 0.000999 |
| melanomatumo | MacAgPres | HAS2 | CD44 | 0.3078 | 0.734 | 0.5209 | 0.000999 |
| melanomatumo | MacAgPres | HEBP1 | CXCR4 | 0.3689 | 1.7737 | 1.0713 | 0.000999 |
| melanomatumo | MacAgPres | MIF | CD74 | 2.6939 | 5.2679 | 3.9809 | 0.000999 |
| melanomatumo | MacAgPres | SERPINE2 | LRP1 | 0.9323 | 0.3786 | 0.6555 | 0.000999 |
| melanomatumo | MacAgPres | CD320 | TGFBR2 | 0.2588 | 0.2584 | 0.2586 | 0.002 |
| melanomatumo | MacAgPres | HEBP1 | C3AR1 | 0.3689 | 0.729 | 0.5489 | 0.002 |
| melanomatumo | MacAgPres | HEBP1 | C5AR1 | 0.3689 | 0.5707 | 0.4698 | 0.003 |
| melanomatumo | MacAgPres | HEBP1 | FPR1 | 0.3689 | 0.4486 | 0.4087 | 0.003 |
| melanomatumo | MacAgPres | HMGB1 | LY96 | 1.6757 | 0.7263 | 1.201 | 0.004 |
| melanomatumo | MacAgPres | HMGB1 | TLR2 | 1.6757 | 0.3254 | 1.0005 | 0.004 |
| melanomatumo | MacAgPres | HEBP1 | FPR3 | 0.3689 | 0.2881 | 0.3285 | 0.00699 |
| ovarytumor | MacAgPres | ANXA1 | C3AR1 | 1.0314 | 0.729 | 0.8802 | 0.000999 |
| ovarytumor | MacAgPres | ANXA1 | C5AR1 | 1.0314 | 0.5707 | 0.8011 | 0.000999 |

| | | | | | | | |
|---|---|---|---|---|---|---|---|
| ovarytumor | MacAgPres | ANXA1 | CD4 | 1.0314 | 0.5059 | 0.7686 | 0.000999 |
| ovarytumor | MacAgPres | ANXA1 | CD44 | 1.0314 | 0.734 | 0.8827 | 0.000999 |
| ovarytumor | MacAgPres | ANXA1 | CXCR4 | 1.0314 | 1.7737 | 1.4025 | 0.000999 |
| ovarytumor | MacAgPres | ANXA1 | FPR1 | 1.0314 | 0.4486 | 0.74 | 0.000999 |
| ovarytumor | MacAgPres | ANXA1 | FPR3 | 1.0314 | 0.2881 | 0.6597 | 0.000999 |
| ovarytumor | MacAgPres | APOE | SORL1 | 0.837 | 0.5618 | 0.6994 | 0.000999 |
| ovarytumor | MacAgPres | APP | ALOX5 | 0.2907 | 0.39 | 0.3404 | 0.000999 |
| ovarytumor | MacAgPres | APP | C3AR1 | 0.2907 | 0.729 | 0.5099 | 0.000999 |
| ovarytumor | MacAgPres | APP | C5AR1 | 0.2907 | 0.5707 | 0.4307 | 0.000999 |
| ovarytumor | MacAgPres | APP | CXCR4 | 0.2907 | 1.7737 | 1.0322 | 0.000999 |
| ovarytumor | MacAgPres | APP | FPR1 | 0.2907 | 0.4486 | 0.3697 | 0.000999 |
| ovarytumor | MacAgPres | APP | LRP1 | 0.2907 | 0.3786 | 0.3347 | 0.000999 |
| ovarytumor | MacAgPres | APP | SORL1 | 0.2907 | 0.5618 | 0.4263 | 0.000999 |
| ovarytumor | MacAgPres | APP | TGFBR2 | 0.2907 | 0.2584 | 0.2745 | 0.000999 |
| ovarytumor | MacAgPres | ARF1 | CD4 | 0.7364 | 0.5059 | 0.6211 | 0.000999 |
| ovarytumor | MacAgPres | ARF1 | CD44 | 0.7364 | 0.734 | 0.7352 | 0.000999 |
| ovarytumor | MacAgPres | CALM1 | TNFRSF1A | 1.0087 | 0.3745 | 0.6916 | 0.000999 |
| ovarytumor | MacAgPres | CALM1 | TNFRSF1B | 1.0087 | 0.4694 | 0.739 | 0.000999 |
| ovarytumor | MacAgPres | CALR | CD44 | 0.7423 | 0.734 | 0.7381 | 0.000999 |
| ovarytumor | MacAgPres | CALR | LRP1 | 0.7423 | 0.3786 | 0.5604 | 0.000999 |
| ovarytumor | MacAgPres | COL1A1 | CD44 | 0.3504 | 0.734 | 0.5422 | 0.000999 |
| ovarytumor | MacAgPres | GRN | TNFRSF1B | 0.3896 | 0.4694 | 0.4295 | 0.000999 |
| ovarytumor | MacAgPres | GSTP1 | CD44 | 2.6185 | 0.734 | 1.6762 | 0.000999 |
| ovarytumor | MacAgPres | HMGB1 | LY96 | 1.7272 | 0.7263 | 1.2268 | 0.000999 |
| ovarytumor | MacAgPres | HMGB1 | TLR2 | 1.7272 | 0.3254 | 1.0263 | 0.000999 |
| ovarytumor | MacAgPres | HMGB2 | CD44 | 0.4974 | 0.734 | 0.6157 | 0.000999 |
| ovarytumor | MacAgPres | HMGB2 | DAB2 | 0.4974 | 0.6336 | 0.5655 | 0.000999 |
| ovarytumor | MacAgPres | HSP90B1 | ITGB1 | 0.6365 | 0.3809 | 0.5087 | 0.000999 |
| ovarytumor | MacAgPres | HSP90B1 | LRP1 | 0.6365 | 0.3786 | 0.5076 | 0.000999 |
| ovarytumor | MacAgPres | HSP90B1 | TNFRSF1B | 0.6365 | 0.4694 | 0.5529 | 0.000999 |

| | | | | | | | |
|---|---|---|---|---|---|---|---|
| ovarytumor | MacAgPres | ITGB1 | DAB2 | 0.2783 | 0.6336 | 0.4559 | 0.000999 |
| ovarytumor | MacAgPres | ITGB1 | PLAUR | 0.2783 | 1.0857 | 0.682 | 0.000999 |
| ovarytumor | MacAgPres | LGALS3 | CLEC7A | 0.5097 | 0.683 | 0.5963 | 0.000999 |
| ovarytumor | MacAgPres | LGALS3 | FCGR2A | 0.5097 | 1.2301 | 0.8699 | 0.000999 |
| ovarytumor | MacAgPres | LGALS3 | FCGR2B | 0.5097 | 0.5782 | 0.544 | 0.000999 |
| ovarytumor | MacAgPres | LGALS3 | TGFBR2 | 0.5097 | 0.2584 | 0.384 | 0.000999 |
| ovarytumor | MacAgPres | LRPAP1 | LRP1 | 0.3826 | 0.3786 | 0.3806 | 0.000999 |
| ovarytumor | MacAgPres | LRPAP1 | SORL1 | 0.3826 | 0.5618 | 0.4722 | 0.000999 |
| ovarytumor | MacAgPres | MDK | LRP1 | 1.5321 | 0.3786 | 0.9553 | 0.000999 |
| ovarytumor | MacAgPres | MIF | CD74 | 0.8565 | 5.2679 | 3.0622 | 0.000999 |
| ovarytumor | MacAgPres | RPS19 | C5AR1 | 3.9325 | 0.5707 | 2.2516 | 0.000999 |
| ovarytumor | MacAgPres | RPS19 | CD4 | 3.9325 | 0.5059 | 2.2192 | 0.000999 |
| ovarytumor | MacAgPres | RTN4 | CD44 | 0.4992 | 0.734 | 0.6166 | 0.000999 |
| ovarytumor | MacAgPres | RTN4 | CD99 | 0.4992 | 0.5375 | 0.5183 | 0.000999 |
| ovarytumor | MacAgPres | SERPING1 | LRP1 | 0.3678 | 0.3786 | 0.3732 | 0.000999 |
| ovarytumor | MacAgPres | SPP1 | CD44 | 0.3946 | 0.734 | 0.5643 | 0.000999 |
| ovarytumor | MacAgPres | TIMP1 | LRP1 | 0.7779 | 0.3786 | 0.5783 | 0.000999 |
| ovarytumor | MacAgPres | LGALS3 | ITGB1 | 0.5097 | 0.3809 | 0.4453 | 0.00599 |

| Supplementary Table 8: Communication results between macrophages and lymphocytes subpopulations, and malignant cells | | | | | | | |
|---|---|---|---|---|---|---|---|
| **RTM_IFN vs. lymphocytes** | | | | | | | |
| celltype_01 | celltype_02 | ligand | receptor | ligand_mean_expression | receptor_mean_expression | ligand_receptor_mean_expression | pvalue |
| RTMIFN | NKT | A2M | TNFRSF14 | 0.8393 | 0.2848 | 0.562 | 0.000999 |
| RTMIFN | NKT | ADM | PTGER2 | 0.3632 | 0.3627 | 0.3629 | 0.000999 |
| RTMIFN | NKT | APOE | SORL1 | 2.8059 | 0.4137 | 1.6098 | 0.000999 |
| RTMIFN | NKT | APP | CXCR4 | 0.2822 | 2.0373 | 1.1598 | 0.000999 |
| RTMIFN | NKT | APP | F2R | 0.2822 | 0.2978 | 0.29 | 0.000999 |
| RTMIFN | NKT | APP | SORL1 | 0.2822 | 0.4137 | 0.3479 | 0.000999 |
| RTMIFN | NKT | APP | TNFRSF14 | 0.2822 | 0.2848 | 0.2835 | 0.000999 |
| RTMIFN | NKT | C3 | CXCR4 | 0.6267 | 2.0373 | 1.332 | 0.000999 |
| RTMIFN | NKT | C3 | ITGB2 | 0.6267 | 1.6481 | 1.1374 | 0.000999 |
| RTMIFN | NKT | CCL20 | CXCR4 | 0.2626 | 2.0373 | 1.1499 | 0.000999 |
| RTMIFN | NKT | CCL20 | S1PR4 | 0.2626 | 0.4435 | 0.3531 | 0.000999 |
| RTMIFN | NKT | CCL20 | S1PR5 | 0.2626 | 0.508 | 0.3853 | 0.000999 |
| RTMIFN | NKT | CD58 | CD2 | 0.3142 | 1.1465 | 0.7304 | 0.000999 |
| RTMIFN | NKT | COL1A1 | CD44 | 0.6812 | 0.6724 | 0.6768 | 0.000999 |
| RTMIFN | NKT | COL1A1 | ITGB1 | 0.6812 | 0.488 | 0.5846 | 0.000999 |
| RTMIFN | NKT | CXCL10 | CXCR4 | 0.9959 | 2.0373 | 1.5166 | 0.000999 |
| RTMIFN | NKT | CXCL10 | S1PR4 | 0.9959 | 0.4435 | 0.7197 | 0.000999 |
| RTMIFN | NKT | CXCL10 | S1PR5 | 0.9959 | 0.508 | 0.752 | 0.000999 |
| RTMIFN | NKT | CXCL11 | CXCR4 | 0.3356 | 2.0373 | 1.1864 | 0.000999 |
| RTMIFN | NKT | CXCL11 | S1PR4 | 0.3356 | 0.4435 | 0.3896 | 0.000999 |
| RTMIFN | NKT | CXCL11 | S1PR5 | 0.3356 | 0.508 | 0.4218 | 0.000999 |
| RTMIFN | NKT | CXCL12 | CXCR4 | 0.2864 | 2.0373 | 1.1618 | 0.000999 |
| RTMIFN | NKT | CXCL12 | S1PR4 | 0.2864 | 0.4435 | 0.3649 | 0.000999 |
| RTMIFN | NKT | CXCL12 | S1PR5 | 0.2864 | 0.508 | 0.3972 | 0.000999 |
| RTMIFN | NKT | CXCL16 | CXCR4 | 0.9728 | 2.0373 | 1.505 | 0.000999 |
| RTMIFN | NKT | CXCL16 | S1PR4 | 0.9728 | 0.4435 | 0.7081 | 0.000999 |
| RTMIFN | NKT | CXCL16 | S1PR5 | 0.9728 | 0.508 | 0.7404 | 0.000999 |

| | | | | | | | |
|---|---|---|---|---|---|---|---|
| RTMIFN | NKT | CXCL2 | CXCR4 | 0.7506 | 2.0373 | 1.394 | 0.000999 |
| RTMIFN | NKT | CXCL2 | S1PR4 | 0.7506 | 0.4435 | 0.5971 | 0.000999 |
| RTMIFN | NKT | CXCL2 | S1PR5 | 0.7506 | 0.508 | 0.6293 | 0.000999 |
| RTMIFN | NKT | CXCL3 | CXCR4 | 0.4907 | 2.0373 | 1.264 | 0.000999 |
| RTMIFN | NKT | CXCL3 | S1PR4 | 0.4907 | 0.4435 | 0.4671 | 0.000999 |
| RTMIFN | NKT | CXCL3 | S1PR5 | 0.4907 | 0.508 | 0.4994 | 0.000999 |
| RTMIFN | NKT | CXCL9 | CXCR4 | 0.7322 | 2.0373 | 1.3847 | 0.000999 |
| RTMIFN | NKT | CXCL9 | S1PR4 | 0.7322 | 0.4435 | 0.5879 | 0.000999 |
| RTMIFN | NKT | CXCL9 | S1PR5 | 0.7322 | 0.508 | 0.6201 | 0.000999 |
| RTMIFN | NKT | FN1 | CD44 | 1.3064 | 0.6724 | 0.9894 | 0.000999 |
| RTMIFN | NKT | FN1 | ITGA4 | 1.3064 | 0.2946 | 0.8005 | 0.000999 |
| RTMIFN | NKT | FN1 | ITGB1 | 1.3064 | 0.488 | 0.8972 | 0.000999 |
| RTMIFN | NKT | FN1 | ITGB7 | 1.3064 | 0.4721 | 0.8893 | 0.000999 |
| RTMIFN | NKT | FN1 | RPSA | 1.3064 | 2.3639 | 1.8352 | 0.000999 |
| RTMIFN | NKT | GRN | TNFRSF1B | 1.8891 | 0.4949 | 1.192 | 0.000999 |
| RTMIFN | NKT | GSTP1 | CD44 | 1.6648 | 0.6724 | 1.1686 | 0.000999 |
| RTMIFN | NKT | GSTP1 | IL7R | 1.6648 | 0.5933 | 1.1291 | 0.000999 |
| RTMIFN | NKT | GSTP1 | ITGA4 | 1.6648 | 0.2946 | 0.9797 | 0.000999 |
| RTMIFN | NKT | HBEGF | CD44 | 0.3607 | 0.6724 | 0.5165 | 0.000999 |
| RTMIFN | NKT | HEBP1 | CXCR4 | 0.2764 | 2.0373 | 1.1568 | 0.000999 |
| RTMIFN | NKT | HLA-DRA | PTPRC | 4.1984 | 1.8797 | 3.039 | 0.000999 |
| RTMIFN | NKT | ICAM1 | IL2RG | 0.5523 | 1.1524 | 0.8523 | 0.000999 |
| RTMIFN | NKT | ICAM1 | ITGAL | 0.5523 | 0.6778 | 0.6151 | 0.000999 |
| RTMIFN | NKT | ICAM1 | ITGB2 | 0.5523 | 1.6481 | 1.1002 | 0.000999 |
| RTMIFN | NKT | ICAM1 | SPN | 0.5523 | 0.6398 | 0.596 | 0.000999 |
| RTMIFN | NKT | LGALS3 | CD6 | 1.3808 | 0.4881 | 0.9345 | 0.000999 |
| RTMIFN | NKT | LGALS3 | ITGB1 | 1.3808 | 0.488 | 0.9344 | 0.000999 |
| RTMIFN | NKT | LGALS3 | SPN | 1.3808 | 0.6398 | 1.0103 | 0.000999 |
| RTMIFN | NKT | LRPAP1 | SORL1 | 0.5174 | 0.4137 | 0.4655 | 0.000999 |
| RTMIFN | NKT | MMP9 | CD44 | 0.3709 | 0.6724 | 0.5217 | 0.000999 |

| | | | | | | | |
|---|---|---|---|---|---|---|---|
| RTMIFN | NKT | MMP9 | ITGB1 | 0.3709 | 0.488 | 0.4295 | 0.000999 |
| RTMIFN | NKT | NAMPT | ANXA6 | 1.0013 | 0.844 | 0.9226 | 0.000999 |
| RTMIFN | NKT | RTN4 | CD44 | 1.0112 | 0.6724 | 0.8418 | 0.000999 |
| RTMIFN | NKT | RTN4 | CD99 | 1.0112 | 1.4547 | 1.2329 | 0.000999 |
| RTMIFN | NKT | SPP1 | CD44 | 2.4585 | 0.6724 | 1.5655 | 0.000999 |
| RTMIFN | NKT | SPP1 | ITGA4 | 2.4585 | 0.2946 | 1.3765 | 0.000999 |
| RTMIFN | NKT | SPP1 | ITGB1 | 2.4585 | 0.488 | 1.4732 | 0.000999 |
| RTMIFN | NKT | TNFSF13B | ITGB1 | 1.0395 | 0.488 | 0.7637 | 0.000999 |
| RTMIFN | NKT | AIMP1 | RACK1 | 0.3209 | 2.4327 | 1.3768 | 0.00699 |
| RTMIFN | TCD4ex | A2M | TNFRSF14 | 0.8393 | 0.2806 | 0.5599 | 0.000999 |
| RTMIFN | TCD4ex | ADM | PTGER4 | 0.3632 | 0.4696 | 0.4164 | 0.000999 |
| RTMIFN | TCD4ex | ANXA1 | CCR7 | 1.0204 | 0.4844 | 0.7524 | 0.000999 |
| RTMIFN | TCD4ex | ANXA1 | CD4 | 1.0204 | 0.5864 | 0.8034 | 0.000999 |
| RTMIFN | TCD4ex | ANXA1 | CD44 | 1.0204 | 1.0244 | 1.0224 | 0.000999 |
| RTMIFN | TCD4ex | ANXA1 | CXCR4 | 1.0204 | 2.4201 | 1.7202 | 0.000999 |
| RTMIFN | TCD4ex | APP | CCR7 | 0.2822 | 0.4844 | 0.3833 | 0.000999 |
| RTMIFN | TCD4ex | APP | CXCR4 | 0.2822 | 2.4201 | 1.3512 | 0.000999 |
| RTMIFN | TCD4ex | C3 | CCR7 | 0.6267 | 0.4844 | 0.5555 | 0.000999 |
| RTMIFN | TCD4ex | C3 | CXCR4 | 0.6267 | 2.4201 | 1.5234 | 0.000999 |
| RTMIFN | TCD4ex | C3 | ITGB2 | 0.6267 | 0.7871 | 0.7069 | 0.000999 |
| RTMIFN | TCD4ex | CALR | IL7R | 1.0231 | 1.2353 | 1.1292 | 0.000999 |
| RTMIFN | TCD4ex | CCL20 | CCR7 | 0.2626 | 0.4844 | 0.3735 | 0.000999 |
| RTMIFN | TCD4ex | CCL20 | CXCR4 | 0.2626 | 2.4201 | 1.3413 | 0.000999 |
| RTMIFN | TCD4ex | CD40 | CD40LG | 0.4077 | 0.3948 | 0.4013 | 0.000999 |
| RTMIFN | TCD4ex | CD86 | CD28 | 0.516 | 0.3256 | 0.4208 | 0.000999 |
| RTMIFN | TCD4ex | CD86 | CTLA4 | 0.516 | 0.6294 | 0.5727 | 0.000999 |
| RTMIFN | TCD4ex | COL1A1 | CD44 | 0.6812 | 1.0244 | 0.8528 | 0.000999 |
| RTMIFN | TCD4ex | COL1A1 | ITGB1 | 0.6812 | 0.4615 | 0.5714 | 0.000999 |
| RTMIFN | TCD4ex | CXCL10 | CCR7 | 0.9959 | 0.4844 | 0.7402 | 0.000999 |
| RTMIFN | TCD4ex | CXCL10 | CXCR4 | 0.9959 | 2.4201 | 1.708 | 0.000999 |

| | | | | | | | |
|---|---|---|---|---|---|---|---|
| RTMIFN | TCD4ex | CXCL11 | CCR7 | 0.3356 | 0.4844 | 0.41 | 0.000999 |
| RTMIFN | TCD4ex | CXCL11 | CXCR4 | 0.3356 | 2.4201 | 1.3778 | 0.000999 |
| RTMIFN | TCD4ex | CXCL12 | CCR7 | 0.2864 | 0.4844 | 0.3854 | 0.000999 |
| RTMIFN | TCD4ex | CXCL12 | CD4 | 0.2864 | 0.5864 | 0.4364 | 0.000999 |
| RTMIFN | TCD4ex | CXCL12 | CXCR4 | 0.2864 | 2.4201 | 1.3532 | 0.000999 |
| RTMIFN | TCD4ex | CXCL16 | CCR7 | 0.9728 | 0.4844 | 0.7286 | 0.000999 |
| RTMIFN | TCD4ex | CXCL16 | CXCR4 | 0.9728 | 2.4201 | 1.6964 | 0.000999 |
| RTMIFN | TCD4ex | CXCL2 | CCR7 | 0.7506 | 0.4844 | 0.6175 | 0.000999 |
| RTMIFN | TCD4ex | CXCL2 | CXCR4 | 0.7506 | 2.4201 | 1.5853 | 0.000999 |
| RTMIFN | TCD4ex | CXCL3 | CCR7 | 0.4907 | 0.4844 | 0.4876 | 0.000999 |
| RTMIFN | TCD4ex | CXCL3 | CXCR4 | 0.4907 | 2.4201 | 1.4554 | 0.000999 |
| RTMIFN | TCD4ex | CXCL9 | CCR7 | 0.7322 | 0.4844 | 0.6083 | 0.000999 |
| RTMIFN | TCD4ex | CXCL9 | CXCR4 | 0.7322 | 2.4201 | 1.5761 | 0.000999 |
| RTMIFN | TCD4ex | FN1 | CD44 | 1.3064 | 1.0244 | 1.1654 | 0.000999 |
| RTMIFN | TCD4ex | FN1 | ITGB1 | 1.3064 | 0.4615 | 0.884 | 0.000999 |
| RTMIFN | TCD4ex | FN1 | RPSA | 1.3064 | 2.4895 | 1.898 | 0.000999 |
| RTMIFN | TCD4ex | GRN | TNFRSF1B | 1.8891 | 0.383 | 1.1361 | 0.000999 |
| RTMIFN | TCD4ex | GSTP1 | CD44 | 1.6648 | 1.0244 | 1.3446 | 0.000999 |
| RTMIFN | TCD4ex | GSTP1 | IL7R | 1.6648 | 1.2353 | 1.45 | 0.000999 |
| RTMIFN | TCD4ex | HBEGF | CD44 | 0.3607 | 1.0244 | 0.6925 | 0.000999 |
| RTMIFN | TCD4ex | HEBP1 | CCR7 | 0.2764 | 0.4844 | 0.3804 | 0.000999 |
| RTMIFN | TCD4ex | HEBP1 | CXCR4 | 0.2764 | 2.4201 | 1.3482 | 0.000999 |
| RTMIFN | TCD4ex | HLA-DMA | CD4 | 1.8152 | 0.5864 | 1.2008 | 0.000999 |
| RTMIFN | TCD4ex | HLA-DRA | CD4 | 4.1984 | 0.5864 | 2.3924 | 0.000999 |
| RTMIFN | TCD4ex | HLA-DRA | PDCD1 | 4.1984 | 0.457 | 2.3277 | 0.000999 |
| RTMIFN | TCD4ex | HLA-DRA | PTPRC | 4.1984 | 1.7493 | 2.9739 | 0.000999 |
| RTMIFN | TCD4ex | HSP90B1 | ITGB1 | 0.7956 | 0.4615 | 0.6285 | 0.000999 |
| RTMIFN | TCD4ex | HSP90B1 | TNFRSF1B | 0.7956 | 0.383 | 0.5893 | 0.000999 |
| RTMIFN | TCD4ex | ICAM1 | IL2RG | 0.5523 | 0.9695 | 0.7609 | 0.000999 |
| RTMIFN | TCD4ex | ICAM1 | ITGAL | 0.5523 | 0.2603 | 0.4063 | 0.000999 |

| | | | | | | | |
|---|---|---|---|---|---|---|---|
| RTMIFN | TCD4ex | ICAM1 | ITGB2 | 0.5523 | 0.7871 | 0.6697 | 0.000999 |
| RTMIFN | TCD4ex | ITGB2 | ITGAL | 1.456 | 0.2603 | 0.8581 | 0.000999 |
| RTMIFN | TCD4ex | LGALS3 | CD6 | 1.3808 | 0.5338 | 0.9573 | 0.000999 |
| RTMIFN | TCD4ex | LGALS3 | ITGB1 | 1.3808 | 0.4615 | 0.9212 | 0.000999 |
| RTMIFN | TCD4ex | MMP9 | CD44 | 0.3709 | 1.0244 | 0.6977 | 0.000999 |
| RTMIFN | TCD4ex | MMP9 | ITGB1 | 0.3709 | 0.4615 | 0.4162 | 0.000999 |
| RTMIFN | TCD4ex | NAMPT | ANXA6 | 1.0013 | 0.5386 | 0.7699 | 0.000999 |
| RTMIFN | TCD4ex | RTN4 | CD44 | 1.0112 | 1.0244 | 1.0178 | 0.000999 |
| RTMIFN | TCD4ex | RTN4 | CD99 | 1.0112 | 0.9165 | 0.9639 | 0.000999 |
| RTMIFN | TCD4ex | SPP1 | CD44 | 2.4585 | 1.0244 | 1.7415 | 0.000999 |
| RTMIFN | TCD4ex | SPP1 | ITGB1 | 2.4585 | 0.4615 | 1.46 | 0.000999 |
| RTMIFN | TCD4ex | TGFB1 | ITGB1 | 0.3271 | 0.4615 | 0.3943 | 0.000999 |
| RTMIFN | TCD4ex | TNFSF12 | TNFRSF25 | 0.3767 | 0.3259 | 0.3513 | 0.000999 |
| RTMIFN | TCD4ex | TNFSF13B | ITGB1 | 1.0395 | 0.4615 | 0.7505 | 0.000999 |
| RTMIFN | TCD4ex | CD58 | CD2 | 0.3142 | 1.5731 | 0.9436 | 0.002 |
| RTMIFN | TCD4ex | APP | TNFRSF14 | 0.2822 | 0.2806 | 0.2814 | 0.003 |
| RTMIFN | TCD4reg | A2M | TNFRSF14 | 0.8393 | 0.3843 | 0.6118 | 0.000999 |
| RTMIFN | TCD4reg | ANXA1 | CCR7 | 1.0204 | 0.3098 | 0.6651 | 0.000999 |
| RTMIFN | TCD4reg | ANXA1 | CD4 | 1.0204 | 0.429 | 0.7247 | 0.000999 |
| RTMIFN | TCD4reg | ANXA1 | CD44 | 1.0204 | 1.0896 | 1.055 | 0.000999 |
| RTMIFN | TCD4reg | ANXA1 | CXCR3 | 1.0204 | 0.3357 | 0.678 | 0.000999 |
| RTMIFN | TCD4reg | ANXA1 | CXCR4 | 1.0204 | 2.2962 | 1.6583 | 0.000999 |
| RTMIFN | TCD4reg | ANXA1 | CXCR6 | 1.0204 | 0.3945 | 0.7075 | 0.000999 |
| RTMIFN | TCD4reg | ANXA1 | FAS | 1.0204 | 0.2727 | 0.6466 | 0.000999 |
| RTMIFN | TCD4reg | APP | CCR7 | 0.2822 | 0.3098 | 0.296 | 0.000999 |
| RTMIFN | TCD4reg | APP | CXCR3 | 0.2822 | 0.3357 | 0.309 | 0.000999 |
| RTMIFN | TCD4reg | APP | CXCR4 | 0.2822 | 2.2962 | 1.2892 | 0.000999 |
| RTMIFN | TCD4reg | APP | CXCR6 | 0.2822 | 0.3945 | 0.3384 | 0.000999 |
| RTMIFN | TCD4reg | APP | TNFRSF14 | 0.2822 | 0.3843 | 0.3333 | 0.000999 |
| RTMIFN | TCD4reg | C3 | CCR7 | 0.6267 | 0.3098 | 0.4682 | 0.000999 |

| | | | | | | | |
|---|---|---|---|---|---|---|---|
| RTMIFN | TCD4reg | C3 | CXCR3 | 0.6267 | 0.3357 | 0.4812 | 0.000999 |
| RTMIFN | TCD4reg | C3 | CXCR4 | 0.6267 | 2.2962 | 1.4614 | 0.000999 |
| RTMIFN | TCD4reg | C3 | CXCR6 | 0.6267 | 0.3945 | 0.5106 | 0.000999 |
| RTMIFN | TCD4reg | C3 | ITGB2 | 0.6267 | 0.6238 | 0.6252 | 0.000999 |
| RTMIFN | TCD4reg | CALR | IL7R | 1.0231 | 0.9682 | 0.9956 | 0.000999 |
| RTMIFN | TCD4reg | CD86 | CD28 | 0.516 | 0.313 | 0.4145 | 0.000999 |
| RTMIFN | TCD4reg | CD86 | CTLA4 | 0.516 | 0.7909 | 0.6535 | 0.000999 |
| RTMIFN | TCD4reg | COL1A1 | CD44 | 0.6812 | 1.0896 | 0.8854 | 0.000999 |
| RTMIFN | TCD4reg | COL1A1 | ITGB1 | 0.6812 | 0.4001 | 0.5407 | 0.000999 |
| RTMIFN | TCD4reg | CXCL10 | CCR7 | 0.9959 | 0.3098 | 0.6529 | 0.000999 |
| RTMIFN | TCD4reg | CXCL10 | CXCR3 | 0.9959 | 0.3357 | 0.6658 | 0.000999 |
| RTMIFN | TCD4reg | CXCL10 | CXCR4 | 0.9959 | 2.2962 | 1.646 | 0.000999 |
| RTMIFN | TCD4reg | CXCL10 | CXCR6 | 0.9959 | 0.3945 | 0.6952 | 0.000999 |
| RTMIFN | TCD4reg | CXCL10 | S1PR4 | 0.9959 | 0.2744 | 0.6352 | 0.000999 |
| RTMIFN | TCD4reg | CXCL11 | CCR7 | 0.3356 | 0.3098 | 0.3227 | 0.000999 |
| RTMIFN | TCD4reg | CXCL11 | CXCR3 | 0.3356 | 0.3357 | 0.3356 | 0.000999 |
| RTMIFN | TCD4reg | CXCL11 | CXCR4 | 0.3356 | 2.2962 | 1.3159 | 0.000999 |
| RTMIFN | TCD4reg | CXCL11 | CXCR6 | 0.3356 | 0.3945 | 0.3651 | 0.000999 |
| RTMIFN | TCD4reg | CXCL11 | S1PR4 | 0.3356 | 0.2744 | 0.305 | 0.000999 |
| RTMIFN | TCD4reg | CXCL12 | CCR7 | 0.2864 | 0.3098 | 0.2981 | 0.000999 |
| RTMIFN | TCD4reg | CXCL12 | CD4 | 0.2864 | 0.429 | 0.3577 | 0.000999 |
| RTMIFN | TCD4reg | CXCL12 | CXCR3 | 0.2864 | 0.3357 | 0.311 | 0.000999 |
| RTMIFN | TCD4reg | CXCL12 | CXCR4 | 0.2864 | 2.2962 | 1.2913 | 0.000999 |
| RTMIFN | TCD4reg | CXCL12 | CXCR6 | 0.2864 | 0.3945 | 0.3404 | 0.000999 |
| RTMIFN | TCD4reg | CXCL12 | S1PR4 | 0.2864 | 0.2744 | 0.2804 | 0.000999 |
| RTMIFN | TCD4reg | CXCL16 | CCR7 | 0.9728 | 0.3098 | 0.6413 | 0.000999 |
| RTMIFN | TCD4reg | CXCL16 | CXCR3 | 0.9728 | 0.3357 | 0.6542 | 0.000999 |
| RTMIFN | TCD4reg | CXCL16 | CXCR4 | 0.9728 | 2.2962 | 1.6345 | 0.000999 |
| RTMIFN | TCD4reg | CXCL16 | CXCR6 | 0.9728 | 0.3945 | 0.6836 | 0.000999 |
| RTMIFN | TCD4reg | CXCL16 | S1PR4 | 0.9728 | 0.2744 | 0.6236 | 0.000999 |

| | | | | | | | |
|---|---|---|---|---|---|---|---|
| RTMIFN | TCD4reg | CXCL2 | CCR7 | 0.7506 | 0.3098 | 0.5302 | 0.000999 |
| RTMIFN | TCD4reg | CXCL2 | CXCR3 | 0.7506 | 0.3357 | 0.5431 | 0.000999 |
| RTMIFN | TCD4reg | CXCL2 | CXCR4 | 0.7506 | 2.2962 | 1.5234 | 0.000999 |
| RTMIFN | TCD4reg | CXCL2 | CXCR6 | 0.7506 | 0.3945 | 0.5726 | 0.000999 |
| RTMIFN | TCD4reg | CXCL2 | S1PR4 | 0.7506 | 0.2744 | 0.5125 | 0.000999 |
| RTMIFN | TCD4reg | CXCL3 | CCR7 | 0.4907 | 0.3098 | 0.4003 | 0.000999 |
| RTMIFN | TCD4reg | CXCL3 | CXCR3 | 0.4907 | 0.3357 | 0.4132 | 0.000999 |
| RTMIFN | TCD4reg | CXCL3 | CXCR4 | 0.4907 | 2.2962 | 1.3934 | 0.000999 |
| RTMIFN | TCD4reg | CXCL3 | CXCR6 | 0.4907 | 0.3945 | 0.4426 | 0.000999 |
| RTMIFN | TCD4reg | CXCL3 | S1PR4 | 0.4907 | 0.2744 | 0.3826 | 0.000999 |
| RTMIFN | TCD4reg | CXCL9 | CCR7 | 0.7322 | 0.3098 | 0.521 | 0.000999 |
| RTMIFN | TCD4reg | CXCL9 | CXCR3 | 0.7322 | 0.3357 | 0.5339 | 0.000999 |
| RTMIFN | TCD4reg | CXCL9 | CXCR4 | 0.7322 | 2.2962 | 1.5142 | 0.000999 |
| RTMIFN | TCD4reg | CXCL9 | CXCR6 | 0.7322 | 0.3945 | 0.5634 | 0.000999 |
| RTMIFN | TCD4reg | CXCL9 | S1PR4 | 0.7322 | 0.2744 | 0.5033 | 0.000999 |
| RTMIFN | TCD4reg | FN1 | CD44 | 1.3064 | 1.0896 | 1.198 | 0.000999 |
| RTMIFN | TCD4reg | FN1 | ITGB1 | 1.3064 | 0.4001 | 0.8533 | 0.000999 |
| RTMIFN | TCD4reg | FN1 | RPSA | 1.3064 | 2.4999 | 1.9032 | 0.000999 |
| RTMIFN | TCD4reg | GRN | TNFRSF1B | 1.8891 | 0.7982 | 1.3437 | 0.000999 |
| RTMIFN | TCD4reg | GSTP1 | CD44 | 1.6648 | 1.0896 | 1.3772 | 0.000999 |
| RTMIFN | TCD4reg | GSTP1 | FAS | 1.6648 | 0.2727 | 0.9687 | 0.000999 |
| RTMIFN | TCD4reg | GSTP1 | IL7R | 1.6648 | 0.9682 | 1.3165 | 0.000999 |
| RTMIFN | TCD4reg | HBEGF | CD44 | 0.3607 | 1.0896 | 0.7251 | 0.000999 |
| RTMIFN | TCD4reg | HEBP1 | CCR7 | 0.2764 | 0.3098 | 0.2931 | 0.000999 |
| RTMIFN | TCD4reg | HEBP1 | CXCR3 | 0.2764 | 0.3357 | 0.306 | 0.000999 |
| RTMIFN | TCD4reg | HEBP1 | CXCR4 | 0.2764 | 2.2962 | 1.2863 | 0.000999 |
| RTMIFN | TCD4reg | HEBP1 | CXCR6 | 0.2764 | 0.3945 | 0.3355 | 0.000999 |
| RTMIFN | TCD4reg | HLA-DMA | CD4 | 1.8152 | 0.429 | 1.1221 | 0.000999 |
| RTMIFN | TCD4reg | HLA-DRA | CD4 | 4.1984 | 0.429 | 2.3137 | 0.000999 |
| RTMIFN | TCD4reg | HLA-DRA | PTPRC | 4.1984 | 1.6603 | 2.9294 | 0.000999 |

| | | | | | | | |
|---|---|---|---|---|---|---|---|
| RTMIFN | TCD4reg | HSP90B1 | ITGB1 | 0.7956 | 0.4001 | 0.5978 | 0.000999 |
| RTMIFN | TCD4reg | HSP90B1 | TNFRSF1B | 0.7956 | 0.7982 | 0.7969 | 0.000999 |
| RTMIFN | TCD4reg | ICAM1 | IL2RA | 0.5523 | 0.7115 | 0.6319 | 0.000999 |
| RTMIFN | TCD4reg | ICAM1 | IL2RG | 0.5523 | 1.1601 | 0.8562 | 0.000999 |
| RTMIFN | TCD4reg | ICAM1 | ITGB2 | 0.5523 | 0.6238 | 0.588 | 0.000999 |
| RTMIFN | TCD4reg | LGALS3 | CD6 | 1.3808 | 0.2651 | 0.8229 | 0.000999 |
| RTMIFN | TCD4reg | LGALS3 | FAS | 1.3808 | 0.2727 | 0.8268 | 0.000999 |
| RTMIFN | TCD4reg | LGALS3 | ITGB1 | 1.3808 | 0.4001 | 0.8905 | 0.000999 |
| RTMIFN | TCD4reg | MMP9 | CD44 | 0.3709 | 1.0896 | 0.7303 | 0.000999 |
| RTMIFN | TCD4reg | MMP9 | ITGB1 | 0.3709 | 0.4001 | 0.3855 | 0.000999 |
| RTMIFN | TCD4reg | NAMPT | ANXA6 | 1.0013 | 0.4557 | 0.7285 | 0.000999 |
| RTMIFN | TCD4reg | RTN4 | CD44 | 1.0112 | 1.0896 | 1.0504 | 0.000999 |
| RTMIFN | TCD4reg | RTN4 | CD99 | 1.0112 | 0.6603 | 0.8357 | 0.000999 |
| RTMIFN | TCD4reg | SPP1 | CD44 | 2.4585 | 1.0896 | 1.7741 | 0.000999 |
| RTMIFN | TCD4reg | SPP1 | ITGB1 | 2.4585 | 0.4001 | 1.4293 | 0.000999 |
| RTMIFN | TCD4reg | TNFSF13B | ITGB1 | 1.0395 | 0.4001 | 0.7198 | 0.000999 |
| RTMIFN | TCD4reg | CALR | CD44 | 1.0231 | 1.0896 | 1.0564 | 0.002 |
| RTMIFN | TCD4reg | CALR | FAS | 1.0231 | 0.2727 | 0.6479 | 0.002 |
| RTMIFN | TCD4reg | TGFB1 | ITGB1 | 0.3271 | 0.4001 | 0.3636 | 0.002 |
| RTMIFN | TCD8em | A2M | TNFRSF14 | 0.8393 | 0.2915 | 0.5654 | 0.000999 |
| RTMIFN | TCD8em | ADM | PTGER4 | 0.3632 | 0.2847 | 0.324 | 0.000999 |
| RTMIFN | TCD8em | APP | CXCR3 | 0.2822 | 0.4189 | 0.3506 | 0.000999 |
| RTMIFN | TCD8em | APP | CXCR4 | 0.2822 | 2.8389 | 1.5606 | 0.000999 |
| RTMIFN | TCD8em | APP | CXCR6 | 0.2822 | 0.3399 | 0.3111 | 0.000999 |
| RTMIFN | TCD8em | APP | F2R | 0.2822 | 0.2695 | 0.2759 | 0.000999 |
| RTMIFN | TCD8em | APP | TNFRSF14 | 0.2822 | 0.2915 | 0.2869 | 0.000999 |
| RTMIFN | TCD8em | C3 | CXCR3 | 0.6267 | 0.4189 | 0.5228 | 0.000999 |
| RTMIFN | TCD8em | C3 | CXCR4 | 0.6267 | 2.8389 | 1.7328 | 0.000999 |
| RTMIFN | TCD8em | C3 | CXCR6 | 0.6267 | 0.3399 | 0.4833 | 0.000999 |
| RTMIFN | TCD8em | C3 | ITGB2 | 0.6267 | 1.0018 | 0.8142 | 0.000999 |

| | | | | | | | |
|---|---|---|---|---|---|---|---|
| RTMIFN | TCD8em | CCL20 | CXCR3 | 0.2626 | 0.4189 | 0.3407 | 0.000999 |
| RTMIFN | TCD8em | CCL20 | CXCR4 | 0.2626 | 2.8389 | 1.5507 | 0.000999 |
| RTMIFN | TCD8em | CCL20 | CXCR6 | 0.2626 | 0.3399 | 0.3012 | 0.000999 |
| RTMIFN | TCD8em | CCL20 | S1PR4 | 0.2626 | 0.2677 | 0.2652 | 0.000999 |
| RTMIFN | TCD8em | CD58 | CD2 | 0.3142 | 1.5961 | 0.9551 | 0.000999 |
| RTMIFN | TCD8em | COL1A1 | CD44 | 0.6812 | 1.119 | 0.9001 | 0.000999 |
| RTMIFN | TCD8em | COL1A1 | ITGB1 | 0.6812 | 0.2557 | 0.4685 | 0.000999 |
| RTMIFN | TCD8em | CXCL10 | CXCR3 | 0.9959 | 0.4189 | 0.7074 | 0.000999 |
| RTMIFN | TCD8em | CXCL10 | CXCR4 | 0.9959 | 2.8389 | 1.9174 | 0.000999 |
| RTMIFN | TCD8em | CXCL10 | CXCR6 | 0.9959 | 0.3399 | 0.6679 | 0.000999 |
| RTMIFN | TCD8em | CXCL10 | S1PR4 | 0.9959 | 0.2677 | 0.6318 | 0.000999 |
| RTMIFN | TCD8em | CXCL11 | CXCR3 | 0.3356 | 0.4189 | 0.3772 | 0.000999 |
| RTMIFN | TCD8em | CXCL11 | CXCR4 | 0.3356 | 2.8389 | 1.5873 | 0.000999 |
| RTMIFN | TCD8em | CXCL11 | CXCR6 | 0.3356 | 0.3399 | 0.3377 | 0.000999 |
| RTMIFN | TCD8em | CXCL11 | S1PR4 | 0.3356 | 0.2677 | 0.3017 | 0.000999 |
| RTMIFN | TCD8em | CXCL12 | CXCR3 | 0.2864 | 0.4189 | 0.3526 | 0.000999 |
| RTMIFN | TCD8em | CXCL12 | CXCR4 | 0.2864 | 2.8389 | 1.5626 | 0.000999 |
| RTMIFN | TCD8em | CXCL12 | CXCR6 | 0.2864 | 0.3399 | 0.3131 | 0.000999 |
| RTMIFN | TCD8em | CXCL12 | S1PR4 | 0.2864 | 0.2677 | 0.277 | 0.000999 |
| RTMIFN | TCD8em | CXCL16 | CXCR3 | 0.9728 | 0.4189 | 0.6958 | 0.000999 |
| RTMIFN | TCD8em | CXCL16 | CXCR4 | 0.9728 | 2.8389 | 1.9058 | 0.000999 |
| RTMIFN | TCD8em | CXCL16 | CXCR6 | 0.9728 | 0.3399 | 0.6563 | 0.000999 |
| RTMIFN | TCD8em | CXCL16 | S1PR4 | 0.9728 | 0.2677 | 0.6202 | 0.000999 |
| RTMIFN | TCD8em | CXCL2 | CXCR3 | 0.7506 | 0.4189 | 0.5847 | 0.000999 |
| RTMIFN | TCD8em | CXCL2 | CXCR4 | 0.7506 | 2.8389 | 1.7948 | 0.000999 |
| RTMIFN | TCD8em | CXCL2 | CXCR6 | 0.7506 | 0.3399 | 0.5452 | 0.000999 |
| RTMIFN | TCD8em | CXCL2 | S1PR4 | 0.7506 | 0.2677 | 0.5092 | 0.000999 |
| RTMIFN | TCD8em | CXCL3 | CXCR3 | 0.4907 | 0.4189 | 0.4548 | 0.000999 |
| RTMIFN | TCD8em | CXCL3 | CXCR4 | 0.4907 | 2.8389 | 1.6648 | 0.000999 |
| RTMIFN | TCD8em | CXCL3 | CXCR6 | 0.4907 | 0.3399 | 0.4153 | 0.000999 |

| | | | | | | | |
|---|---|---|---|---|---|---|---|
| RTMIFN | TCD8em | CXCL3 | S1PR4 | 0.4907 | 0.2677 | 0.3792 | 0.000999 |
| RTMIFN | TCD8em | CXCL9 | CXCR3 | 0.7322 | 0.4189 | 0.5755 | 0.000999 |
| RTMIFN | TCD8em | CXCL9 | CXCR4 | 0.7322 | 2.8389 | 1.7856 | 0.000999 |
| RTMIFN | TCD8em | CXCL9 | CXCR6 | 0.7322 | 0.3399 | 0.536 | 0.000999 |
| RTMIFN | TCD8em | CXCL9 | S1PR4 | 0.7322 | 0.2677 | 0.5 | 0.000999 |
| RTMIFN | TCD8em | FN1 | CD44 | 1.3064 | 1.119 | 1.2127 | 0.000999 |
| RTMIFN | TCD8em | FN1 | ITGA4 | 1.3064 | 0.2823 | 0.7944 | 0.000999 |
| RTMIFN | TCD8em | FN1 | ITGB1 | 1.3064 | 0.2557 | 0.7811 | 0.000999 |
| RTMIFN | TCD8em | FN1 | RPSA | 1.3064 | 2.3686 | 1.8375 | 0.000999 |
| RTMIFN | TCD8em | GRN | TNFRSF1B | 1.8891 | 0.3038 | 1.0964 | 0.000999 |
| RTMIFN | TCD8em | GSTP1 | CD44 | 1.6648 | 1.119 | 1.3919 | 0.000999 |
| RTMIFN | TCD8em | GSTP1 | IL7R | 1.6648 | 0.7183 | 1.1915 | 0.000999 |
| RTMIFN | TCD8em | GSTP1 | ITGA4 | 1.6648 | 0.2823 | 0.9736 | 0.000999 |
| RTMIFN | TCD8em | HBEGF | CD44 | 0.3607 | 1.119 | 0.7399 | 0.000999 |
| RTMIFN | TCD8em | HEBP1 | CXCR3 | 0.2764 | 0.4189 | 0.3476 | 0.000999 |
| RTMIFN | TCD8em | HEBP1 | CXCR4 | 0.2764 | 2.8389 | 1.5577 | 0.000999 |
| RTMIFN | TCD8em | HEBP1 | CXCR6 | 0.2764 | 0.3399 | 0.3081 | 0.000999 |
| RTMIFN | TCD8em | HLA-DRA | PTPRC | 4.1984 | 1.7912 | 2.9948 | 0.000999 |
| RTMIFN | TCD8em | HSP90B1 | ITGB1 | 0.7956 | 0.2557 | 0.5257 | 0.000999 |
| RTMIFN | TCD8em | HSP90B1 | TNFRSF1B | 0.7956 | 0.3038 | 0.5497 | 0.000999 |
| RTMIFN | TCD8em | ICAM1 | IL2RG | 0.5523 | 0.9501 | 0.7512 | 0.000999 |
| RTMIFN | TCD8em | ICAM1 | ITGAL | 0.5523 | 0.3162 | 0.4343 | 0.000999 |
| RTMIFN | TCD8em | ICAM1 | ITGB2 | 0.5523 | 1.0018 | 0.777 | 0.000999 |
| RTMIFN | TCD8em | ICAM1 | SPN | 0.5523 | 0.2603 | 0.4063 | 0.000999 |
| RTMIFN | TCD8em | ITGB2 | ITGAL | 1.456 | 0.3162 | 0.8861 | 0.000999 |
| RTMIFN | TCD8em | LGALS3 | CD6 | 1.3808 | 0.4204 | 0.9006 | 0.000999 |
| RTMIFN | TCD8em | LGALS3 | ITGB1 | 1.3808 | 0.2557 | 0.8183 | 0.000999 |
| RTMIFN | TCD8em | LGALS3 | SPN | 1.3808 | 0.2603 | 0.8206 | 0.000999 |
| RTMIFN | TCD8em | MMP9 | CD44 | 0.3709 | 1.119 | 0.745 | 0.000999 |
| RTMIFN | TCD8em | MMP9 | ITGB1 | 0.3709 | 0.2557 | 0.3133 | 0.000999 |

| | | | | | | | |
|---|---|---|---|---|---|---|---|
| RTMIFN | TCD8em | NAMPT | ANXA6 | 1.0013 | 0.552 | 0.7766 | 0.000999 |
| RTMIFN | TCD8em | RTN4 | CD44 | 1.0112 | 1.119 | 1.0651 | 0.000999 |
| RTMIFN | TCD8em | RTN4 | CD99 | 1.0112 | 1.1525 | 1.0819 | 0.000999 |
| RTMIFN | TCD8em | SPP1 | CD44 | 2.4585 | 1.119 | 1.7888 | 0.000999 |
| RTMIFN | TCD8em | SPP1 | ITGA4 | 2.4585 | 0.2823 | 1.3704 | 0.000999 |
| RTMIFN | TCD8em | SPP1 | ITGB1 | 2.4585 | 0.2557 | 1.3571 | 0.000999 |
| RTMIFN | TCD8em | TNFSF13B | ITGB1 | 1.0395 | 0.2557 | 0.6476 | 0.000999 |
| RTMIFN | TCD8em | ITGB1 | CD47 | 0.528 | 0.4276 | 0.4778 | 0.002 |
| RTMIFN | TCD8em | ITGB1 | ITGA4 | 0.528 | 0.2823 | 0.4052 | 0.002 |
| RTMIFN | TCD8em | ITGB1 | RACK1 | 0.528 | 2.4374 | 1.4827 | 0.002 |
| RTMIFN | TCD8em | CALR | CD44 | 1.0231 | 1.119 | 1.0711 | 0.004 |
| RTMIFN | TCD8em | CALR | IL7R | 1.0231 | 0.7183 | 0.8707 | 0.004 |
| RTMIFN | TCD8em | MIF | ITGA4 | 0.8578 | 0.2823 | 0.5701 | 0.00799 |
| RTMIFN | TCD8em | MIF | TNFRSF14 | 0.8578 | 0.2915 | 0.5746 | 0.00899 |
| RTMIFN | TCD8ex | A2M | TNFRSF14 | 0.8393 | 0.3094 | 0.5744 | 0.000999 |
| RTMIFN | TCD8ex | ADM | PTGER4 | 0.3632 | 0.3089 | 0.336 | 0.000999 |
| RTMIFN | TCD8ex | ANXA1 | CD44 | 1.0204 | 0.8219 | 0.9212 | 0.000999 |
| RTMIFN | TCD8ex | ANXA1 | CXCR3 | 1.0204 | 0.3647 | 0.6925 | 0.000999 |
| RTMIFN | TCD8ex | ANXA1 | CXCR4 | 1.0204 | 2.3472 | 1.6838 | 0.000999 |
| RTMIFN | TCD8ex | ANXA1 | CXCR6 | 1.0204 | 0.6626 | 0.8415 | 0.000999 |
| RTMIFN | TCD8ex | ANXA1 | ITGA4 | 1.0204 | 0.4432 | 0.7318 | 0.000999 |
| RTMIFN | TCD8ex | ANXA1 | VCAM1 | 1.0204 | 0.4462 | 0.7333 | 0.000999 |
| RTMIFN | TCD8ex | APOE | VCAM1 | 2.8059 | 0.4462 | 1.6261 | 0.000999 |
| RTMIFN | TCD8ex | APP | CXCR3 | 0.2822 | 0.3647 | 0.3235 | 0.000999 |
| RTMIFN | TCD8ex | APP | CXCR4 | 0.2822 | 2.3472 | 1.3147 | 0.000999 |
| RTMIFN | TCD8ex | APP | CXCR6 | 0.2822 | 0.6626 | 0.4724 | 0.000999 |
| RTMIFN | TCD8ex | APP | TNFRSF14 | 0.2822 | 0.3094 | 0.2958 | 0.000999 |
| RTMIFN | TCD8ex | C3 | CXCR3 | 0.6267 | 0.3647 | 0.4957 | 0.000999 |
| RTMIFN | TCD8ex | C3 | CXCR4 | 0.6267 | 2.3472 | 1.4869 | 0.000999 |
| RTMIFN | TCD8ex | C3 | CXCR6 | 0.6267 | 0.6626 | 0.6446 | 0.000999 |

| | | | | | | | |
|---|---|---|---|---|---|---|---|
| RTMIFN | TCD8ex | C3 | ITGB2 | 0.6267 | 0.7635 | 0.6951 | 0.000999 |
| RTMIFN | TCD8ex | CD86 | CTLA4 | 0.516 | 0.4883 | 0.5021 | 0.000999 |
| RTMIFN | TCD8ex | COL1A1 | CD44 | 0.6812 | 0.8219 | 0.7516 | 0.000999 |
| RTMIFN | TCD8ex | COL1A1 | ITGB1 | 0.6812 | 0.2995 | 0.4904 | 0.000999 |
| RTMIFN | TCD8ex | CXCL10 | CXCR3 | 0.9959 | 0.3647 | 0.6803 | 0.000999 |
| RTMIFN | TCD8ex | CXCL10 | CXCR4 | 0.9959 | 2.3472 | 1.6716 | 0.000999 |
| RTMIFN | TCD8ex | CXCL10 | CXCR6 | 0.9959 | 0.6626 | 0.8292 | 0.000999 |
| RTMIFN | TCD8ex | CXCL11 | CXCR3 | 0.3356 | 0.3647 | 0.3501 | 0.000999 |
| RTMIFN | TCD8ex | CXCL11 | CXCR4 | 0.3356 | 2.3472 | 1.3414 | 0.000999 |
| RTMIFN | TCD8ex | CXCL11 | CXCR6 | 0.3356 | 0.6626 | 0.4991 | 0.000999 |
| RTMIFN | TCD8ex | CXCL12 | CXCR3 | 0.2864 | 0.3647 | 0.3255 | 0.000999 |
| RTMIFN | TCD8ex | CXCL12 | CXCR4 | 0.2864 | 2.3472 | 1.3168 | 0.000999 |
| RTMIFN | TCD8ex | CXCL12 | CXCR6 | 0.2864 | 0.6626 | 0.4745 | 0.000999 |
| RTMIFN | TCD8ex | CXCL16 | CXCR3 | 0.9728 | 0.3647 | 0.6687 | 0.000999 |
| RTMIFN | TCD8ex | CXCL16 | CXCR4 | 0.9728 | 2.3472 | 1.66 | 0.000999 |
| RTMIFN | TCD8ex | CXCL16 | CXCR6 | 0.9728 | 0.6626 | 0.8177 | 0.000999 |
| RTMIFN | TCD8ex | CXCL2 | CXCR3 | 0.7506 | 0.3647 | 0.5576 | 0.000999 |
| RTMIFN | TCD8ex | CXCL2 | CXCR4 | 0.7506 | 2.3472 | 1.5489 | 0.000999 |
| RTMIFN | TCD8ex | CXCL2 | CXCR6 | 0.7506 | 0.6626 | 0.7066 | 0.000999 |
| RTMIFN | TCD8ex | CXCL3 | CXCR3 | 0.4907 | 0.3647 | 0.4277 | 0.000999 |
| RTMIFN | TCD8ex | CXCL3 | CXCR4 | 0.4907 | 2.3472 | 1.419 | 0.000999 |
| RTMIFN | TCD8ex | CXCL3 | CXCR6 | 0.4907 | 0.6626 | 0.5766 | 0.000999 |
| RTMIFN | TCD8ex | CXCL9 | CXCR3 | 0.7322 | 0.3647 | 0.5484 | 0.000999 |
| RTMIFN | TCD8ex | CXCL9 | CXCR4 | 0.7322 | 2.3472 | 1.5397 | 0.000999 |
| RTMIFN | TCD8ex | CXCL9 | CXCR6 | 0.7322 | 0.6626 | 0.6974 | 0.000999 |
| RTMIFN | TCD8ex | FN1 | CD44 | 1.3064 | 0.8219 | 1.0642 | 0.000999 |
| RTMIFN | TCD8ex | FN1 | ITGA4 | 1.3064 | 0.4432 | 0.8748 | 0.000999 |
| RTMIFN | TCD8ex | FN1 | ITGB1 | 1.3064 | 0.2995 | 0.803 | 0.000999 |
| RTMIFN | TCD8ex | FN1 | ITGB7 | 1.3064 | 0.3007 | 0.8036 | 0.000999 |
| RTMIFN | TCD8ex | FN1 | RPSA | 1.3064 | 2.2697 | 1.7881 | 0.000999 |

| | | | | | | | |
|---|---|---|---|---|---|---|---|
| RTMIFN | TCD8ex | GRN | TNFRSF1B | 1.8891 | 0.6436 | 1.2664 | 0.000999 |
| RTMIFN | TCD8ex | GSTP1 | CD44 | 1.6648 | 0.8219 | 1.2434 | 0.000999 |
| RTMIFN | TCD8ex | GSTP1 | ITGA4 | 1.6648 | 0.4432 | 1.054 | 0.000999 |
| RTMIFN | TCD8ex | GSTP1 | VCAM1 | 1.6648 | 0.4462 | 1.0555 | 0.000999 |
| RTMIFN | TCD8ex | HBEGF | CD44 | 0.3607 | 0.8219 | 0.5913 | 0.000999 |
| RTMIFN | TCD8ex | HEBP1 | CXCR3 | 0.2764 | 0.3647 | 0.3205 | 0.000999 |
| RTMIFN | TCD8ex | HEBP1 | CXCR4 | 0.2764 | 2.3472 | 1.3118 | 0.000999 |
| RTMIFN | TCD8ex | HEBP1 | CXCR6 | 0.2764 | 0.6626 | 0.4695 | 0.000999 |
| RTMIFN | TCD8ex | HLA-DRA | PDCD1 | 4.1984 | 0.6109 | 2.4046 | 0.000999 |
| RTMIFN | TCD8ex | HLA-DRA | PTPRC | 4.1984 | 1.6762 | 2.9373 | 0.000999 |
| RTMIFN | TCD8ex | ICAM1 | IL2RG | 0.5523 | 1.2126 | 0.8824 | 0.000999 |
| RTMIFN | TCD8ex | ICAM1 | ITGB2 | 0.5523 | 0.7635 | 0.6579 | 0.000999 |
| RTMIFN | TCD8ex | ICAM1 | SPN | 0.5523 | 0.3486 | 0.4505 | 0.000999 |
| RTMIFN | TCD8ex | LGALS3 | CD6 | 1.3808 | 0.318 | 0.8494 | 0.000999 |
| RTMIFN | TCD8ex | LGALS3 | ITGB1 | 1.3808 | 0.2995 | 0.8401 | 0.000999 |
| RTMIFN | TCD8ex | LGALS3 | SPN | 1.3808 | 0.3486 | 0.8647 | 0.000999 |
| RTMIFN | TCD8ex | MMP9 | CD44 | 0.3709 | 0.8219 | 0.5964 | 0.000999 |
| RTMIFN | TCD8ex | MMP9 | ITGB1 | 0.3709 | 0.2995 | 0.3352 | 0.000999 |
| RTMIFN | TCD8ex | NAMPT | ANXA6 | 1.0013 | 0.5678 | 0.7845 | 0.000999 |
| RTMIFN | TCD8ex | RTN4 | CD44 | 1.0112 | 0.8219 | 0.9166 | 0.000999 |
| RTMIFN | TCD8ex | RTN4 | CD99 | 1.0112 | 1.2049 | 1.108 | 0.000999 |
| RTMIFN | TCD8ex | SPP1 | CD44 | 2.4585 | 0.8219 | 1.6402 | 0.000999 |
| RTMIFN | TCD8ex | SPP1 | ITGA4 | 2.4585 | 0.4432 | 1.4509 | 0.000999 |
| RTMIFN | TCD8ex | SPP1 | ITGB1 | 2.4585 | 0.2995 | 1.379 | 0.000999 |
| RTMIFN | TCD8ex | TNFSF13B | ITGB1 | 1.0395 | 0.2995 | 0.6695 | 0.000999 |
| RTMIFN | TCD8ex | CD58 | CD2 | 0.3142 | 2.0502 | 1.1822 | 0.003 |
| RTMIFN | TCD8ex | CCL20 | CXCR4 | 0.2626 | 2.3472 | 1.3049 | 0.005 |
| RTMIFN | TCD8ex | ITGB1 | ITGA4 | 0.528 | 0.4432 | 0.4856 | 0.005 |
| RTMIFN | TCD8ex | ITGB1 | VCAM1 | 0.528 | 0.4462 | 0.4871 | 0.005 |
| RTMIFN | TCD8ex | ITGB1 | CD47 | 0.528 | 0.3952 | 0.4616 | 0.00599 |

| | | | | | | | |
|---|---|---|---|---|---|---|---|
| RTMIFN | TCD8ex | ITGB1 | RACK1 | 0.528 | 2.3353 | 1.4316 | 0.00599 |
| RTMIFN | TCD8ex | CCL20 | CXCR3 | 0.2626 | 0.3647 | 0.3136 | 0.00699 |
| RTMIFN | TCD8ex | CCL20 | CXCR6 | 0.2626 | 0.6626 | 0.4626 | 0.00699 |
| RTMIFN | TCD8ex | TGFB1 | ITGB1 | 0.3271 | 0.2995 | 0.3133 | 0.00799 |
| RTMIFN | TGD | A2M | TNFRSF14 | 0.8393 | 0.4642 | 0.6517 | 0.000999 |
| RTMIFN | TGD | ADM | PTGDR | 0.3632 | 0.7153 | 0.5392 | 0.000999 |
| RTMIFN | TGD | ANXA1 | CXCR3 | 1.0204 | 1.1748 | 1.0976 | 0.000999 |
| RTMIFN | TGD | ANXA1 | CXCR4 | 1.0204 | 1.4894 | 1.2549 | 0.000999 |
| RTMIFN | TGD | ANXA1 | ITGA4 | 1.0204 | 0.3968 | 0.7086 | 0.000999 |
| RTMIFN | TGD | APOE | SORL1 | 2.8059 | 0.3318 | 1.5689 | 0.000999 |
| RTMIFN | TGD | APP | CXCR3 | 0.2822 | 1.1748 | 0.7285 | 0.000999 |
| RTMIFN | TGD | APP | CXCR4 | 0.2822 | 1.4894 | 0.8858 | 0.000999 |
| RTMIFN | TGD | APP | SORL1 | 0.2822 | 0.3318 | 0.307 | 0.000999 |
| RTMIFN | TGD | APP | TGFBR2 | 0.2822 | 0.3677 | 0.325 | 0.000999 |
| RTMIFN | TGD | C3 | CXCR3 | 0.6267 | 1.1748 | 0.9007 | 0.000999 |
| RTMIFN | TGD | C3 | CXCR4 | 0.6267 | 1.4894 | 1.058 | 0.000999 |
| RTMIFN | TGD | C3 | ITGB2 | 0.6267 | 1.4444 | 1.0355 | 0.000999 |
| RTMIFN | TGD | CALR | IL7R | 1.0231 | 1.3862 | 1.2047 | 0.000999 |
| RTMIFN | TGD | CCL20 | CXCR3 | 0.2626 | 1.1748 | 0.7187 | 0.000999 |
| RTMIFN | TGD | CCL20 | CXCR4 | 0.2626 | 1.4894 | 0.876 | 0.000999 |
| RTMIFN | TGD | CD58 | CD2 | 0.3142 | 1.4522 | 0.8832 | 0.000999 |
| RTMIFN | TGD | CXCL10 | CXCR3 | 0.9959 | 1.1748 | 1.0853 | 0.000999 |
| RTMIFN | TGD | CXCL10 | CXCR4 | 0.9959 | 1.4894 | 1.2426 | 0.000999 |
| RTMIFN | TGD | CXCL11 | CXCR3 | 0.3356 | 1.1748 | 0.7552 | 0.000999 |
| RTMIFN | TGD | CXCL11 | CXCR4 | 0.3356 | 1.4894 | 0.9125 | 0.000999 |
| RTMIFN | TGD | CXCL12 | CXCR3 | 0.2864 | 1.1748 | 0.7306 | 0.000999 |
| RTMIFN | TGD | CXCL12 | CXCR4 | 0.2864 | 1.4894 | 0.8879 | 0.000999 |
| RTMIFN | TGD | CXCL16 | CXCR3 | 0.9728 | 1.1748 | 1.0738 | 0.000999 |
| RTMIFN | TGD | CXCL16 | CXCR4 | 0.9728 | 1.4894 | 1.2311 | 0.000999 |
| RTMIFN | TGD | CXCL2 | CXCR3 | 0.7506 | 1.1748 | 0.9627 | 0.000999 |

| | | | | | | | |
|---|---|---|---|---|---|---|---|
| RTMIFN | TGD | CXCL2 | CXCR4 | 0.7506 | 1.4894 | 1.12 | 0.000999 |
| RTMIFN | TGD | CXCL3 | CXCR3 | 0.4907 | 1.1748 | 0.8327 | 0.000999 |
| RTMIFN | TGD | CXCL3 | CXCR4 | 0.4907 | 1.4894 | 0.9901 | 0.000999 |
| RTMIFN | TGD | CXCL9 | CXCR3 | 0.7322 | 1.1748 | 0.9535 | 0.000999 |
| RTMIFN | TGD | CXCL9 | CXCR4 | 0.7322 | 1.4894 | 1.1108 | 0.000999 |
| RTMIFN | TGD | FN1 | CD44 | 1.3064 | 1.1342 | 1.2203 | 0.000999 |
| RTMIFN | TGD | FN1 | ITGA4 | 1.3064 | 0.3968 | 0.8516 | 0.000999 |
| RTMIFN | TGD | FN1 | ITGB1 | 1.3064 | 0.3657 | 0.8361 | 0.000999 |
| RTMIFN | TGD | FN1 | ITGB7 | 1.3064 | 0.459 | 0.8827 | 0.000999 |
| RTMIFN | TGD | FN1 | RPSA | 1.3064 | 2.8814 | 2.0939 | 0.000999 |
| RTMIFN | TGD | GRN | TNFRSF1A | 1.8891 | 0.3099 | 1.0995 | 0.000999 |
| RTMIFN | TGD | GSTP1 | CD44 | 1.6648 | 1.1342 | 1.3995 | 0.000999 |
| RTMIFN | TGD | GSTP1 | IL7R | 1.6648 | 1.3862 | 1.5255 | 0.000999 |
| RTMIFN | TGD | GSTP1 | ITGA4 | 1.6648 | 0.3968 | 1.0308 | 0.000999 |
| RTMIFN | TGD | HBEGF | CD44 | 0.3607 | 1.1342 | 0.7474 | 0.000999 |
| RTMIFN | TGD | HEBP1 | CXCR3 | 0.2764 | 1.1748 | 0.7256 | 0.000999 |
| RTMIFN | TGD | HEBP1 | CXCR4 | 0.2764 | 1.4894 | 0.8829 | 0.000999 |
| RTMIFN | TGD | HLA-DRA | PTPRC | 4.1984 | 1.8653 | 3.0319 | 0.000999 |
| RTMIFN | TGD | ICAM1 | IL2RG | 0.5523 | 1.561 | 1.0566 | 0.000999 |
| RTMIFN | TGD | ICAM1 | ITGAL | 0.5523 | 0.3731 | 0.4627 | 0.000999 |
| RTMIFN | TGD | ICAM1 | SPN | 0.5523 | 0.5644 | 0.5583 | 0.000999 |
| RTMIFN | TGD | ITGB1 | ITGA4 | 0.528 | 0.3968 | 0.4624 | 0.000999 |
| RTMIFN | TGD | ITGB1 | RACK1 | 0.528 | 2.7287 | 1.6283 | 0.000999 |
| RTMIFN | TGD | LGALS3 | CD6 | 1.3808 | 0.4684 | 0.9246 | 0.000999 |
| RTMIFN | TGD | LGALS3 | ITGB1 | 1.3808 | 0.3657 | 0.8732 | 0.000999 |
| RTMIFN | TGD | LGALS3 | SPN | 1.3808 | 0.5644 | 0.9726 | 0.000999 |
| RTMIFN | TGD | LGALS3 | TGFBR2 | 1.3808 | 0.3677 | 0.8742 | 0.000999 |
| RTMIFN | TGD | LRPAP1 | SORL1 | 0.5174 | 0.3318 | 0.4246 | 0.000999 |
| RTMIFN | TGD | MMP9 | CD44 | 0.3709 | 1.1342 | 0.7526 | 0.000999 |
| RTMIFN | TGD | MMP9 | ITGB1 | 0.3709 | 0.3657 | 0.3683 | 0.000999 |

| | | | | | | | |
|---|---|---|---|---|---|---|---|
| RTMIFN | TGD | NAMPT | ANXA6 | 1.0013 | 0.8553 | 0.9283 | 0.000999 |
| RTMIFN | TGD | RTN4 | CD44 | 1.0112 | 1.1342 | 1.0727 | 0.000999 |
| RTMIFN | TGD | RTN4 | CD99 | 1.0112 | 1.7154 | 1.3633 | 0.000999 |
| RTMIFN | TGD | SPP1 | CD44 | 2.4585 | 1.1342 | 1.7964 | 0.000999 |
| RTMIFN | TGD | SPP1 | ITGA4 | 2.4585 | 0.3968 | 1.4277 | 0.000999 |
| RTMIFN | TGD | SPP1 | ITGB1 | 2.4585 | 0.3657 | 1.4121 | 0.000999 |
| RTMIFN | TGD | TNFSF13B | ITGB1 | 1.0395 | 0.3657 | 0.7026 | 0.000999 |
| RTMIFN | TGD | VEGFA | PTK2B | 0.3864 | 0.3668 | 0.3766 | 0.000999 |
| RTMIFN | TGD | APP | TNFRSF14 | 0.2822 | 0.4642 | 0.3732 | 0.002 |
| RTMIFN | TGD | ICAM1 | ITGB2 | 0.5523 | 1.4444 | 0.9984 | 0.002 |
| RTMIFN | TGD | ITGB1 | PTK2B | 0.528 | 0.3668 | 0.4474 | 0.004 |
| RTMIFN | TGD | HLA-A | CD8A | 3.0435 | 1.746 | 2.3948 | 0.005 |
| NKT | RTMIFN | ALOX5AP | ALOX5 | 0.5021 | 0.4066 | 0.4544 | 0.000999 |
| NKT | RTMIFN | ANXA1 | C3AR1 | 1.2259 | 0.6481 | 0.937 | 0.000999 |
| NKT | RTMIFN | ANXA1 | C5AR1 | 1.2259 | 0.5634 | 0.8947 | 0.000999 |
| NKT | RTMIFN | ANXA1 | CD4 | 1.2259 | 0.7071 | 0.9665 | 0.000999 |
| NKT | RTMIFN | ANXA1 | CD44 | 1.2259 | 1.2543 | 1.2401 | 0.000999 |
| NKT | RTMIFN | ANXA1 | FPR1 | 1.2259 | 0.4203 | 0.8231 | 0.000999 |
| NKT | RTMIFN | ANXA1 | FPR3 | 1.2259 | 0.4976 | 0.8618 | 0.000999 |
| NKT | RTMIFN | ARF1 | CD4 | 0.7991 | 0.7071 | 0.7531 | 0.000999 |
| NKT | RTMIFN | ARF1 | CD44 | 0.7991 | 1.2543 | 1.0267 | 0.000999 |
| NKT | RTMIFN | CALM1 | TNFRSF1A | 1.9335 | 0.4908 | 1.2122 | 0.000999 |
| NKT | RTMIFN | CALR | CD40 | 0.9479 | 0.4077 | 0.6778 | 0.000999 |
| NKT | RTMIFN | CALR | CD44 | 0.9479 | 1.2543 | 1.1011 | 0.000999 |
| NKT | RTMIFN | CALR | LRP1 | 0.9479 | 0.5528 | 0.7504 | 0.000999 |
| NKT | RTMIFN | CD2 | CD4 | 1.1465 | 0.7071 | 0.9268 | 0.000999 |
| NKT | RTMIFN | GSTP1 | CD44 | 0.7909 | 1.2543 | 1.0226 | 0.000999 |
| NKT | RTMIFN | HLA-DRA | CD4 | 0.6626 | 0.7071 | 0.6848 | 0.000999 |
| NKT | RTMIFN | HMGB1 | LY96 | 1.195 | 1.1488 | 1.1719 | 0.000999 |
| NKT | RTMIFN | HMGB1 | TLR2 | 1.195 | 0.3529 | 0.7739 | 0.000999 |

| | | | | | | | |
|---|---|---|---|---|---|---|---|
| NKT | RTMIFN | HMGB2 | CD44 | 0.6467 | 1.2543 | 0.9505 | 0.000999 |
| NKT | RTMIFN | HMGB2 | DAB2 | 0.6467 | 0.5996 | 0.6231 | 0.000999 |
| NKT | RTMIFN | HSP90B1 | LRP1 | 0.6138 | 0.5528 | 0.5833 | 0.000999 |
| NKT | RTMIFN | IL16 | CD4 | 0.3556 | 0.7071 | 0.5313 | 0.000999 |
| NKT | RTMIFN | IL16 | FCGR1A | 0.3556 | 0.8743 | 0.6149 | 0.000999 |
| NKT | RTMIFN | ITGA4 | CD44 | 0.2946 | 1.2543 | 0.7744 | 0.000999 |
| NKT | RTMIFN | ITGA4 | TFRC | 0.2946 | 0.3853 | 0.3399 | 0.000999 |
| NKT | RTMIFN | ITGAL | ICAM1 | 0.6778 | 0.5523 | 0.6151 | 0.000999 |
| NKT | RTMIFN | ITGB1 | CD151 | 0.488 | 0.3579 | 0.4229 | 0.000999 |
| NKT | RTMIFN | ITGB1 | DAB2 | 0.488 | 0.5996 | 0.5438 | 0.000999 |
| NKT | RTMIFN | ITGB1 | ENG | 0.488 | 0.3505 | 0.4192 | 0.000999 |
| NKT | RTMIFN | ITGB1 | ITGAX | 0.488 | 0.2809 | 0.3845 | 0.000999 |
| NKT | RTMIFN | ITGB1 | PLAUR | 0.488 | 1.1076 | 0.7978 | 0.000999 |
| NKT | RTMIFN | ITGB2 | ICAM1 | 1.6481 | 0.5523 | 1.1002 | 0.000999 |
| NKT | RTMIFN | ITGB2 | ITGAX | 1.6481 | 0.2809 | 0.9645 | 0.000999 |
| NKT | RTMIFN | ITGB2 | PLAUR | 1.6481 | 1.1076 | 1.3779 | 0.000999 |
| NKT | RTMIFN | LRPAP1 | LRP1 | 0.2807 | 0.5528 | 0.4168 | 0.000999 |
| NKT | RTMIFN | LTB | CD40 | 0.2605 | 0.4077 | 0.3341 | 0.000999 |
| NKT | RTMIFN | LTB | TNFRSF1A | 0.2605 | 0.4908 | 0.3756 | 0.000999 |
| NKT | RTMIFN | MIF | CD74 | 0.6657 | 4.5905 | 2.6281 | 0.000999 |
| NKT | RTMIFN | PSAP | LRP1 | 0.6777 | 0.5528 | 0.6153 | 0.000999 |
| NKT | RTMIFN | PTPRC | CD4 | 1.8797 | 0.7071 | 1.2934 | 0.000999 |
| NKT | RTMIFN | PTPRC | CD44 | 1.8797 | 1.2543 | 1.567 | 0.000999 |
| NKT | RTMIFN | PTPRC | IFNAR1 | 1.8797 | 0.3883 | 1.134 | 0.000999 |
| NKT | RTMIFN | RPS19 | C5AR1 | 3.5336 | 0.5634 | 2.0485 | 0.000999 |
| NKT | RTMIFN | RPS19 | CD4 | 3.5336 | 0.7071 | 2.1203 | 0.000999 |
| NKT | RTMIFN | RPS19 | ICAM1 | 3.5336 | 0.5523 | 2.043 | 0.000999 |
| NKT | RTMIFN | RTN4 | CD44 | 0.4374 | 1.2543 | 0.8459 | 0.000999 |
| NKT | RTMIFN | TGFB1 | ENG | 0.5441 | 0.3505 | 0.4473 | 0.000999 |
| NKT | RTMIFN | TGFB1 | ITGB5 | 0.5441 | 0.2827 | 0.4134 | 0.000999 |

| | | | | | | | |
|---|---|---|---|---|---|---|---|
| NKT | RTMIFN | TGFB1 | TLR2 | 0.5441 | 0.3529 | 0.4485 | 0.000999 |
| NKT | RTMIFN | TIMP1 | LRP1 | 0.3005 | 0.5528 | 0.4266 | 0.000999 |
| NKT | RTMIFN | XCL2 | PTAFR | 0.2774 | 0.2739 | 0.2756 | 0.000999 |
| NKT | RTMIFN | XCL2 | LPAR6 | 0.2774 | 0.4248 | 0.3511 | 0.002 |
| TCD4ex | RTMIFN | ALOX5AP | ALOX5 | 1.119 | 0.4066 | 0.7628 | 0.000999 |
| TCD4ex | RTMIFN | ANXA1 | C3AR1 | 0.5174 | 0.6481 | 0.5828 | 0.000999 |
| TCD4ex | RTMIFN | ANXA1 | C5AR1 | 0.5174 | 0.5634 | 0.5404 | 0.000999 |
| TCD4ex | RTMIFN | ANXA1 | FPR1 | 0.5174 | 0.4203 | 0.4689 | 0.000999 |
| TCD4ex | RTMIFN | ANXA1 | FPR3 | 0.5174 | 0.4976 | 0.5075 | 0.000999 |
| TCD4ex | RTMIFN | APOE | LRP1 | 0.2557 | 0.5528 | 0.4042 | 0.000999 |
| TCD4ex | RTMIFN | AREG | ICAM1 | 0.3427 | 0.5523 | 0.4475 | 0.000999 |
| TCD4ex | RTMIFN | CALM1 | TNFRSF1A | 1.2957 | 0.4908 | 0.8932 | 0.000999 |
| TCD4ex | RTMIFN | CALR | CD40 | 0.748 | 0.4077 | 0.5778 | 0.000999 |
| TCD4ex | RTMIFN | CALR | LRP1 | 0.748 | 0.5528 | 0.6504 | 0.000999 |
| TCD4ex | RTMIFN | CD40LG | CD40 | 0.3948 | 0.4077 | 0.4013 | 0.000999 |
| TCD4ex | RTMIFN | CXCL13 | C3AR1 | 2.3367 | 0.6481 | 1.4924 | 0.000999 |
| TCD4ex | RTMIFN | CXCL13 | C5AR1 | 2.3367 | 0.5634 | 1.45 | 0.000999 |
| TCD4ex | RTMIFN | CXCL13 | FPR1 | 2.3367 | 0.4203 | 1.3785 | 0.000999 |
| TCD4ex | RTMIFN | HMGB1 | AGTRAP | 1.3876 | 0.38 | 0.8838 | 0.000999 |
| TCD4ex | RTMIFN | HMGB1 | LY96 | 1.3876 | 1.1488 | 1.2682 | 0.000999 |
| TCD4ex | RTMIFN | HMGB1 | TLR2 | 1.3876 | 0.3529 | 0.8703 | 0.000999 |
| TCD4ex | RTMIFN | HMGB2 | DAB2 | 1.0243 | 0.5996 | 0.8119 | 0.000999 |
| TCD4ex | RTMIFN | HSP90B1 | LRP1 | 0.3944 | 0.5528 | 0.4736 | 0.000999 |
| TCD4ex | RTMIFN | ICAM3 | ITGB2 | 0.573 | 1.456 | 1.0145 | 0.000999 |
| TCD4ex | RTMIFN | ITGAL | ICAM1 | 0.2603 | 0.5523 | 0.4063 | 0.000999 |
| TCD4ex | RTMIFN | ITGB1 | CD151 | 0.4615 | 0.3579 | 0.4097 | 0.000999 |
| TCD4ex | RTMIFN | ITGB1 | DAB2 | 0.4615 | 0.5996 | 0.5305 | 0.000999 |
| TCD4ex | RTMIFN | ITGB1 | ENG | 0.4615 | 0.3505 | 0.406 | 0.000999 |
| TCD4ex | RTMIFN | ITGB1 | ITGAX | 0.4615 | 0.2809 | 0.3712 | 0.000999 |
| TCD4ex | RTMIFN | ITGB1 | PLAUR | 0.4615 | 1.1076 | 0.7846 | 0.000999 |

| | | | | | | | |
|---|---|---|---|---|---|---|---|
| TCD4ex | RTMIFN | ITGB2 | ICAM1 | 0.7871 | 0.5523 | 0.6697 | 0.000999 |
| TCD4ex | RTMIFN | ITGB2 | ITGAM | 0.7871 | 0.2708 | 0.5289 | 0.000999 |
| TCD4ex | RTMIFN | ITGB2 | ITGAX | 0.7871 | 0.2809 | 0.534 | 0.000999 |
| TCD4ex | RTMIFN | ITGB2 | PLAUR | 0.7871 | 1.1076 | 0.9473 | 0.000999 |
| TCD4ex | RTMIFN | LRPAP1 | LRP1 | 0.2809 | 0.5528 | 0.4168 | 0.000999 |
| TCD4ex | RTMIFN | LTB | CD40 | 0.9655 | 0.4077 | 0.6866 | 0.000999 |
| TCD4ex | RTMIFN | LTB | TNFRSF1A | 0.9655 | 0.4908 | 0.7282 | 0.000999 |
| TCD4ex | RTMIFN | MIF | CD74 | 0.6999 | 4.5905 | 2.6452 | 0.000999 |
| TCD4ex | RTMIFN | NMB | LPAR6 | 0.6391 | 0.4248 | 0.5319 | 0.000999 |
| TCD4ex | RTMIFN | NMB | PTAFR | 0.6391 | 0.2739 | 0.4565 | 0.000999 |
| TCD4ex | RTMIFN | PSAP | LRP1 | 0.5291 | 0.5528 | 0.5409 | 0.000999 |
| TCD4ex | RTMIFN | PTPRC | IFNAR1 | 1.7493 | 0.3883 | 1.0688 | 0.000999 |
| TCD4ex | RTMIFN | RPS19 | C5AR1 | 3.6329 | 0.5634 | 2.0982 | 0.000999 |
| TCD4ex | RTMIFN | RPS19 | ICAM1 | 3.6329 | 0.5523 | 2.0926 | 0.000999 |
| TCD4ex | RTMIFN | SPP1 | ITGB5 | 0.3314 | 0.2827 | 0.3071 | 0.000999 |
| TCD4ex | RTMIFN | TIMP1 | LRP1 | 0.3374 | 0.5528 | 0.4451 | 0.000999 |
| TCD4ex | RTMIFN | NMB | GPR65 | 0.6391 | 0.3227 | 0.4809 | 0.00799 |
| TCD4reg | RTMIFN | ALOX5AP | ALOX5 | 0.4589 | 0.4066 | 0.4327 | 0.000999 |
| TCD4reg | RTMIFN | APOE | LRP1 | 0.2934 | 0.5528 | 0.4231 | 0.000999 |
| TCD4reg | RTMIFN | CALM1 | TNFRSF1A | 1.534 | 0.4908 | 1.0124 | 0.000999 |
| TCD4reg | RTMIFN | CALR | CD40 | 0.6835 | 0.4077 | 0.5456 | 0.000999 |
| TCD4reg | RTMIFN | CALR | LRP1 | 0.6835 | 0.5528 | 0.6182 | 0.000999 |
| TCD4reg | RTMIFN | HMGB1 | AGTRAP | 1.2547 | 0.38 | 0.8174 | 0.000999 |
| TCD4reg | RTMIFN | HMGB1 | LY96 | 1.2547 | 1.1488 | 1.2018 | 0.000999 |
| TCD4reg | RTMIFN | HMGB1 | TLR2 | 1.2547 | 0.3529 | 0.8038 | 0.000999 |
| TCD4reg | RTMIFN | HMGB2 | DAB2 | 0.616 | 0.5996 | 0.6078 | 0.000999 |
| TCD4reg | RTMIFN | HSP90B1 | LRP1 | 0.4221 | 0.5528 | 0.4875 | 0.000999 |
| TCD4reg | RTMIFN | ICAM3 | ITGB2 | 0.6663 | 1.456 | 1.0611 | 0.000999 |
| TCD4reg | RTMIFN | IL16 | FCGR1A | 0.2528 | 0.8743 | 0.5636 | 0.000999 |
| TCD4reg | RTMIFN | ITGB1 | CD151 | 0.4001 | 0.3579 | 0.379 | 0.000999 |

| | | | | | | | |
|---|---|---|---|---|---|---|---|
| TCD4reg | RTMIFN | ITGB1 | DAB2 | 0.4001 | 0.5996 | 0.4998 | 0.000999 |
| TCD4reg | RTMIFN | ITGB1 | ENG | 0.4001 | 0.3505 | 0.3753 | 0.000999 |
| TCD4reg | RTMIFN | ITGB1 | ITGAX | 0.4001 | 0.2809 | 0.3405 | 0.000999 |
| TCD4reg | RTMIFN | ITGB1 | PLAUR | 0.4001 | 1.1076 | 0.7539 | 0.000999 |
| TCD4reg | RTMIFN | ITGB2 | ICAM1 | 0.6238 | 0.5523 | 0.588 | 0.000999 |
| TCD4reg | RTMIFN | ITGB2 | ITGAM | 0.6238 | 0.2708 | 0.4473 | 0.000999 |
| TCD4reg | RTMIFN | ITGB2 | ITGAX | 0.6238 | 0.2809 | 0.4524 | 0.000999 |
| TCD4reg | RTMIFN | ITGB2 | PLAUR | 0.6238 | 1.1076 | 0.8657 | 0.000999 |
| TCD4reg | RTMIFN | LGALS3 | CLEC7A | 0.4492 | 0.6167 | 0.5329 | 0.000999 |
| TCD4reg | RTMIFN | LGALS3 | FCGR2A | 0.4492 | 1.0158 | 0.7325 | 0.000999 |
| TCD4reg | RTMIFN | LGALS3 | FCGR2B | 0.4492 | 0.7853 | 0.6172 | 0.000999 |
| TCD4reg | RTMIFN | LGALS3 | TFRC | 0.4492 | 0.3853 | 0.4172 | 0.000999 |
| TCD4reg | RTMIFN | LGALS3 | TGFBR2 | 0.4492 | 0.2932 | 0.3712 | 0.000999 |
| TCD4reg | RTMIFN | LRPAP1 | LRP1 | 0.3101 | 0.5528 | 0.4315 | 0.000999 |
| TCD4reg | RTMIFN | LTB | CD40 | 2.242 | 0.4077 | 1.3249 | 0.000999 |
| TCD4reg | RTMIFN | LTB | TNFRSF1A | 2.242 | 0.4908 | 1.3664 | 0.000999 |
| TCD4reg | RTMIFN | MIF | CD74 | 0.6295 | 4.5905 | 2.61 | 0.000999 |
| TCD4reg | RTMIFN | PSAP | LRP1 | 0.5938 | 0.5528 | 0.5733 | 0.000999 |
| TCD4reg | RTMIFN | PTPRC | IFNAR1 | 1.6603 | 0.3883 | 1.0243 | 0.000999 |
| TCD4reg | RTMIFN | RPS19 | C5AR1 | 3.6897 | 0.5634 | 2.1265 | 0.000999 |
| TCD4reg | RTMIFN | RPS19 | ICAM1 | 3.6897 | 0.5523 | 2.121 | 0.000999 |
| TCD4reg | RTMIFN | SPP1 | ITGB5 | 0.3125 | 0.2827 | 0.2976 | 0.000999 |
| TCD4reg | RTMIFN | TIMP1 | LRP1 | 0.4362 | 0.5528 | 0.4945 | 0.000999 |
| TCD4reg | RTMIFN | ARF1 | CD4 | 0.7185 | 0.7071 | 0.7128 | 0.002 |
| TCD4reg | RTMIFN | CD2 | CD4 | 1.6566 | 0.7071 | 1.1818 | 0.002 |
| TCD4reg | RTMIFN | CD28 | CD4 | 0.313 | 0.7071 | 0.51 | 0.002 |
| TCD4reg | RTMIFN | IL16 | CD4 | 0.2528 | 0.7071 | 0.4799 | 0.002 |
| TCD4reg | RTMIFN | PTPRC | CD4 | 1.6603 | 0.7071 | 1.1837 | 0.002 |
| TCD4reg | RTMIFN | RPS19 | CD4 | 3.6897 | 0.7071 | 2.1984 | 0.002 |
| TCD4reg | RTMIFN | SELL | CD4 | 0.7062 | 0.7071 | 0.7066 | 0.002 |

| | | | | | | | |
|---|---|---|---|---|---|---|---|
| TCD4reg | RTMIFN | HLA-DMA | CD4 | 0.278 | 0.7071 | 0.4925 | 0.003 |
| TCD4reg | RTMIFN | HLA-DRA | CD4 | 0.7704 | 0.7071 | 0.7387 | 0.00699 |
| TCD8em | RTMIFN | ALOX5AP | ALOX5 | 0.7973 | 0.4066 | 0.602 | 0.000999 |
| TCD8em | RTMIFN | ANXA1 | C3AR1 | 1.0693 | 0.6481 | 0.8587 | 0.000999 |
| TCD8em | RTMIFN | ANXA1 | C5AR1 | 1.0693 | 0.5634 | 0.8163 | 0.000999 |
| TCD8em | RTMIFN | ANXA1 | CD4 | 1.0693 | 0.7071 | 0.8882 | 0.000999 |
| TCD8em | RTMIFN | ANXA1 | FPR1 | 1.0693 | 0.4203 | 0.7448 | 0.000999 |
| TCD8em | RTMIFN | ANXA1 | FPR3 | 1.0693 | 0.4976 | 0.7834 | 0.000999 |
| TCD8em | RTMIFN | APOE | LRP1 | 0.2791 | 0.5528 | 0.416 | 0.000999 |
| TCD8em | RTMIFN | ARF1 | CD4 | 0.6684 | 0.7071 | 0.6877 | 0.000999 |
| TCD8em | RTMIFN | CALM1 | TNFRSF1A | 1.5262 | 0.4908 | 1.0085 | 0.000999 |
| TCD8em | RTMIFN | CALR | CD40 | 0.7141 | 0.4077 | 0.5609 | 0.000999 |
| TCD8em | RTMIFN | CALR | LRP1 | 0.7141 | 0.5528 | 0.6335 | 0.000999 |
| TCD8em | RTMIFN | CD2 | CD4 | 1.5961 | 0.7071 | 1.1516 | 0.000999 |
| TCD8em | RTMIFN | HLA-DMA | CD4 | 0.2969 | 0.7071 | 0.502 | 0.000999 |
| TCD8em | RTMIFN | HLA-DRA | CD4 | 0.952 | 0.7071 | 0.8295 | 0.000999 |
| TCD8em | RTMIFN | HMGB1 | AGTRAP | 1.3636 | 0.38 | 0.8718 | 0.000999 |
| TCD8em | RTMIFN | HMGB1 | LY96 | 1.3636 | 1.1488 | 1.2562 | 0.000999 |
| TCD8em | RTMIFN | HMGB1 | TLR2 | 1.3636 | 0.3529 | 0.8583 | 0.000999 |
| TCD8em | RTMIFN | HMGB2 | DAB2 | 0.7212 | 0.5996 | 0.6604 | 0.000999 |
| TCD8em | RTMIFN | HSP90B1 | LRP1 | 0.471 | 0.5528 | 0.5119 | 0.000999 |
| TCD8em | RTMIFN | ICAM3 | ITGB2 | 0.5282 | 1.456 | 0.9921 | 0.000999 |
| TCD8em | RTMIFN | IFNG | IFNGR1 | 0.3967 | 0.5594 | 0.4781 | 0.000999 |
| TCD8em | RTMIFN | IFNG | IFNGR2 | 0.3967 | 0.7809 | 0.5888 | 0.000999 |
| TCD8em | RTMIFN | IL16 | CD4 | 0.4085 | 0.7071 | 0.5578 | 0.000999 |
| TCD8em | RTMIFN | IL16 | FCGR1A | 0.4085 | 0.8743 | 0.6414 | 0.000999 |
| TCD8em | RTMIFN | ITGA4 | TFRC | 0.2823 | 0.3853 | 0.3338 | 0.000999 |
| TCD8em | RTMIFN | ITGAL | ICAM1 | 0.3162 | 0.5523 | 0.4343 | 0.000999 |
| TCD8em | RTMIFN | ITGB1 | CD151 | 0.2557 | 0.3579 | 0.3068 | 0.000999 |
| TCD8em | RTMIFN | ITGB1 | DAB2 | 0.2557 | 0.5996 | 0.4276 | 0.000999 |

| | | | | | | | |
|---|---|---|---|---|---|---|---|
| TCD8em | RTMIFN | ITGB1 | ENG | 0.2557 | 0.3505 | 0.3031 | 0.000999 |
| TCD8em | RTMIFN | ITGB1 | ITGAX | 0.2557 | 0.2809 | 0.2683 | 0.000999 |
| TCD8em | RTMIFN | ITGB1 | PLAUR | 0.2557 | 1.1076 | 0.6817 | 0.000999 |
| TCD8em | RTMIFN | ITGB2 | ICAM1 | 1.0018 | 0.5523 | 0.777 | 0.000999 |
| TCD8em | RTMIFN | ITGB2 | ITGAM | 1.0018 | 0.2708 | 0.6363 | 0.000999 |
| TCD8em | RTMIFN | ITGB2 | ITGAX | 1.0018 | 0.2809 | 0.6414 | 0.000999 |
| TCD8em | RTMIFN | ITGB2 | PLAUR | 1.0018 | 1.1076 | 1.0547 | 0.000999 |
| TCD8em | RTMIFN | LGALS3 | CLEC7A | 0.2689 | 0.6167 | 0.4428 | 0.000999 |
| TCD8em | RTMIFN | LGALS3 | FCGR2A | 0.2689 | 1.0158 | 0.6423 | 0.000999 |
| TCD8em | RTMIFN | LGALS3 | FCGR2B | 0.2689 | 0.7853 | 0.5271 | 0.000999 |
| TCD8em | RTMIFN | LGALS3 | TFRC | 0.2689 | 0.3853 | 0.3271 | 0.000999 |
| TCD8em | RTMIFN | LRPAP1 | LRP1 | 0.2719 | 0.5528 | 0.4123 | 0.000999 |
| TCD8em | RTMIFN | LTB | CD40 | 0.6352 | 0.4077 | 0.5214 | 0.000999 |
| TCD8em | RTMIFN | LTB | TNFRSF1A | 0.6352 | 0.4908 | 0.563 | 0.000999 |
| TCD8em | RTMIFN | MIF | CD74 | 0.6063 | 4.5905 | 2.5984 | 0.000999 |
| TCD8em | RTMIFN | PSAP | LRP1 | 0.5012 | 0.5528 | 0.527 | 0.000999 |
| TCD8em | RTMIFN | PTPRC | CD4 | 1.7912 | 0.7071 | 1.2491 | 0.000999 |
| TCD8em | RTMIFN | PTPRC | IFNAR1 | 1.7912 | 0.3883 | 1.0898 | 0.000999 |
| TCD8em | RTMIFN | RPS19 | C5AR1 | 3.536 | 0.5634 | 2.0497 | 0.000999 |
| TCD8em | RTMIFN | RPS19 | CD4 | 3.536 | 0.7071 | 2.1215 | 0.000999 |
| TCD8em | RTMIFN | RPS19 | ICAM1 | 3.536 | 0.5523 | 2.0441 | 0.000999 |
| TCD8em | RTMIFN | SELL | CD4 | 0.258 | 0.7071 | 0.4825 | 0.000999 |
| TCD8em | RTMIFN | SPP1 | ITGB5 | 0.3608 | 0.2827 | 0.3218 | 0.000999 |
| TCD8em | RTMIFN | TIMP1 | LRP1 | 0.2667 | 0.5528 | 0.4097 | 0.000999 |
| TCD8em | RTMIFN | TNF | TNFRSF1A | 0.2667 | 0.4908 | 0.3788 | 0.000999 |
| TCD8em | RTMIFN | XCL2 | LPAR6 | 0.295 | 0.4248 | 0.3599 | 0.000999 |
| TCD8em | RTMIFN | XCL2 | PTAFR | 0.295 | 0.2739 | 0.2844 | 0.000999 |
| TCD8em | RTMIFN | HSP90B1 | ITGB1 | 0.471 | 0.528 | 0.4995 | 0.002 |
| TCD8em | RTMIFN | LGALS3 | ITGB1 | 0.2689 | 0.528 | 0.3984 | 0.002 |
| TCD8em | RTMIFN | LTB | ITGB1 | 0.6352 | 0.528 | 0.5816 | 0.002 |

| | | | | | | | |
|---|---|---|---|---|---|---|---|
| TCD8em | RTMIFN | SPP1 | ITGB1 | 0.3608 | 0.528 | 0.4444 | 0.002 |
| TCD8ex | RTMIFN | ALOX5AP | ALOX5 | 1.5024 | 0.4066 | 0.9545 | 0.000999 |
| TCD8ex | RTMIFN | ANXA1 | C3AR1 | 0.5645 | 0.6481 | 0.6063 | 0.000999 |
| TCD8ex | RTMIFN | ANXA1 | C5AR1 | 0.5645 | 0.5634 | 0.5639 | 0.000999 |
| TCD8ex | RTMIFN | ANXA1 | CD4 | 0.5645 | 0.7071 | 0.6358 | 0.000999 |
| TCD8ex | RTMIFN | ANXA1 | CD44 | 0.5645 | 1.2543 | 0.9094 | 0.000999 |
| TCD8ex | RTMIFN | ANXA1 | FPR1 | 0.5645 | 0.4203 | 0.4924 | 0.000999 |
| TCD8ex | RTMIFN | ANXA1 | FPR3 | 0.5645 | 0.4976 | 0.531 | 0.000999 |
| TCD8ex | RTMIFN | APOE | LRP1 | 0.2814 | 0.5528 | 0.4171 | 0.000999 |
| TCD8ex | RTMIFN | AREG | ICAM1 | 0.2758 | 0.5523 | 0.414 | 0.000999 |
| TCD8ex | RTMIFN | ARF1 | CD4 | 0.7835 | 0.7071 | 0.7453 | 0.000999 |
| TCD8ex | RTMIFN | ARF1 | CD44 | 0.7835 | 1.2543 | 1.0189 | 0.000999 |
| TCD8ex | RTMIFN | CALM1 | TNFRSF1A | 1.4627 | 0.4908 | 0.9768 | 0.000999 |
| TCD8ex | RTMIFN | CALR | CD40 | 0.9634 | 0.4077 | 0.6855 | 0.000999 |
| TCD8ex | RTMIFN | CALR | CD44 | 0.9634 | 1.2543 | 1.1088 | 0.000999 |
| TCD8ex | RTMIFN | CALR | LRP1 | 0.9634 | 0.5528 | 0.7581 | 0.000999 |
| TCD8ex | RTMIFN | CD2 | CD4 | 2.0502 | 0.7071 | 1.3786 | 0.000999 |
| TCD8ex | RTMIFN | CXCL13 | C3AR1 | 1.7749 | 0.6481 | 1.2115 | 0.000999 |
| TCD8ex | RTMIFN | CXCL13 | C5AR1 | 1.7749 | 0.5634 | 1.1692 | 0.000999 |
| TCD8ex | RTMIFN | CXCL13 | FPR1 | 1.7749 | 0.4203 | 1.0976 | 0.000999 |
| TCD8ex | RTMIFN | FASLG | TNFRSF1A | 0.2613 | 0.4908 | 0.376 | 0.000999 |
| TCD8ex | RTMIFN | GSTP1 | CD44 | 0.8286 | 1.2543 | 1.0415 | 0.000999 |
| TCD8ex | RTMIFN | HLA-DMA | CD4 | 0.3991 | 0.7071 | 0.5531 | 0.000999 |
| TCD8ex | RTMIFN | HLA-DRA | CD4 | 1.247 | 0.7071 | 0.977 | 0.000999 |
| TCD8ex | RTMIFN | HMGB1 | AGTRAP | 1.5414 | 0.38 | 0.9607 | 0.000999 |
| TCD8ex | RTMIFN | HMGB1 | LY96 | 1.5414 | 1.1488 | 1.3451 | 0.000999 |
| TCD8ex | RTMIFN | HMGB1 | TLR2 | 1.5414 | 0.3529 | 0.9472 | 0.000999 |
| TCD8ex | RTMIFN | HMGB2 | CD44 | 0.85 | 1.2543 | 1.0522 | 0.000999 |
| TCD8ex | RTMIFN | HMGB2 | DAB2 | 0.85 | 0.5996 | 0.7248 | 0.000999 |
| TCD8ex | RTMIFN | HSP90B1 | LRP1 | 0.5666 | 0.5528 | 0.5597 | 0.000999 |

| | | | | | | | |
|---|---|---|---|---|---|---|---|
| TCD8ex | RTMIFN | ICAM3 | ITGB2 | 0.5405 | 1.456 | 0.9982 | 0.000999 |
| TCD8ex | RTMIFN | IFNG | IFNGR1 | 0.8575 | 0.5594 | 0.7085 | 0.000999 |
| TCD8ex | RTMIFN | IFNG | IFNGR2 | 0.8575 | 0.7809 | 0.8192 | 0.000999 |
| TCD8ex | RTMIFN | ITGA4 | CD44 | 0.4432 | 1.2543 | 0.8488 | 0.000999 |
| TCD8ex | RTMIFN | ITGA4 | TFRC | 0.4432 | 0.3853 | 0.4143 | 0.000999 |
| TCD8ex | RTMIFN | ITGB1 | DAB2 | 0.2995 | 0.5996 | 0.4495 | 0.000999 |
| TCD8ex | RTMIFN | ITGB1 | ENG | 0.2995 | 0.3505 | 0.325 | 0.000999 |
| TCD8ex | RTMIFN | ITGB1 | ITGAX | 0.2995 | 0.2809 | 0.2902 | 0.000999 |
| TCD8ex | RTMIFN | ITGB1 | PLAUR | 0.2995 | 1.1076 | 0.7036 | 0.000999 |
| TCD8ex | RTMIFN | ITGB2 | ICAM1 | 0.7635 | 0.5523 | 0.6579 | 0.000999 |
| TCD8ex | RTMIFN | ITGB2 | ITGAM | 0.7635 | 0.2708 | 0.5171 | 0.000999 |
| TCD8ex | RTMIFN | ITGB2 | ITGAX | 0.7635 | 0.2809 | 0.5222 | 0.000999 |
| TCD8ex | RTMIFN | ITGB2 | PLAUR | 0.7635 | 1.1076 | 0.9355 | 0.000999 |
| TCD8ex | RTMIFN | LGALS3 | CLEC7A | 0.3824 | 0.6167 | 0.4995 | 0.000999 |
| TCD8ex | RTMIFN | LGALS3 | FCGR2A | 0.3824 | 1.0158 | 0.6991 | 0.000999 |
| TCD8ex | RTMIFN | LGALS3 | FCGR2B | 0.3824 | 0.7853 | 0.5838 | 0.000999 |
| TCD8ex | RTMIFN | LGALS3 | TFRC | 0.3824 | 0.3853 | 0.3838 | 0.000999 |
| TCD8ex | RTMIFN | LRPAP1 | LRP1 | 0.2971 | 0.5528 | 0.4249 | 0.000999 |
| TCD8ex | RTMIFN | LTB | CD40 | 0.4576 | 0.4077 | 0.4326 | 0.000999 |
| TCD8ex | RTMIFN | LTB | TNFRSF1A | 0.4576 | 0.4908 | 0.4742 | 0.000999 |
| TCD8ex | RTMIFN | MIF | CD74 | 0.749 | 4.5905 | 2.6698 | 0.000999 |
| TCD8ex | RTMIFN | PSAP | LRP1 | 0.5016 | 0.5528 | 0.5272 | 0.000999 |
| TCD8ex | RTMIFN | PTPRC | CD4 | 1.6762 | 0.7071 | 1.1916 | 0.000999 |
| TCD8ex | RTMIFN | PTPRC | CD44 | 1.6762 | 1.2543 | 1.4653 | 0.000999 |
| TCD8ex | RTMIFN | PTPRC | IFNAR1 | 1.6762 | 0.3883 | 1.0323 | 0.000999 |
| TCD8ex | RTMIFN | RPS19 | C5AR1 | 3.8353 | 0.5634 | 2.1993 | 0.000999 |
| TCD8ex | RTMIFN | RPS19 | CD4 | 3.8353 | 0.7071 | 2.2712 | 0.000999 |
| TCD8ex | RTMIFN | RPS19 | ICAM1 | 3.8353 | 0.5523 | 2.1938 | 0.000999 |
| TCD8ex | RTMIFN | RTN4 | CD44 | 0.3033 | 1.2543 | 0.7788 | 0.000999 |
| TCD8ex | RTMIFN | SPP1 | CD44 | 0.2978 | 1.2543 | 0.7761 | 0.000999 |

| | | | | | | | |
|---|---|---|---|---|---|---|---|
| TCD8ex | RTMIFN | SPP1 | ITGB5 | 0.2978 | 0.2827 | 0.2903 | 0.000999 |
| TCD8ex | RTMIFN | VCAM1 | ITGB2 | 0.4462 | 1.456 | 0.9511 | 0.000999 |
| TCD8ex | RTMIFN | VCAM1 | TFRC | 0.4462 | 0.3853 | 0.4158 | 0.000999 |
| TCD8ex | RTMIFN | XCL2 | LPAR6 | 0.3157 | 0.4248 | 0.3702 | 0.000999 |
| TCD8ex | RTMIFN | XCL2 | PTAFR | 0.3157 | 0.2739 | 0.2948 | 0.000999 |
| TCD8ex | RTMIFN | ITGB1 | CD151 | 0.2995 | 0.3579 | 0.3287 | 0.002 |
| TCD8ex | RTMIFN | VCAM1 | ITGB1 | 0.4462 | 0.528 | 0.4871 | 0.005 |
| TCD8ex | RTMIFN | HSP90B1 | ITGB1 | 0.5666 | 0.528 | 0.5473 | 0.00599 |
| TCD8ex | RTMIFN | LTB | ITGB1 | 0.4576 | 0.528 | 0.4928 | 0.00599 |
| TCD8ex | RTMIFN | LGALS3 | ITGB1 | 0.3824 | 0.528 | 0.4552 | 0.00699 |
| TCD8ex | RTMIFN | SPP1 | ITGB1 | 0.2978 | 0.528 | 0.4129 | 0.00699 |
| TGD | RTMIFN | ANXA1 | C3AR1 | 0.7073 | 0.6481 | 0.6777 | 0.000999 |
| TGD | RTMIFN | ANXA1 | C5AR1 | 0.7073 | 0.5634 | 0.6353 | 0.000999 |
| TGD | RTMIFN | ANXA1 | CD4 | 0.7073 | 0.7071 | 0.7072 | 0.000999 |
| TGD | RTMIFN | ANXA1 | FPR1 | 0.7073 | 0.4203 | 0.5638 | 0.000999 |
| TGD | RTMIFN | ANXA1 | FPR3 | 0.7073 | 0.4976 | 0.6024 | 0.000999 |
| TGD | RTMIFN | AREG | ICAM1 | 0.4756 | 0.5523 | 0.5139 | 0.000999 |
| TGD | RTMIFN | ARF1 | CD4 | 0.9453 | 0.7071 | 0.8262 | 0.000999 |
| TGD | RTMIFN | CALM1 | TNFRSF1A | 1.7587 | 0.4908 | 1.1247 | 0.000999 |
| TGD | RTMIFN | CALM1 | TNFRSF1B | 1.7587 | 0.3537 | 1.0562 | 0.000999 |
| TGD | RTMIFN | CALR | CD40 | 0.9627 | 0.4077 | 0.6852 | 0.000999 |
| TGD | RTMIFN | CALR | LRP1 | 0.9627 | 0.5528 | 0.7577 | 0.000999 |
| TGD | RTMIFN | CD2 | CD4 | 1.4522 | 0.7071 | 1.0796 | 0.000999 |
| TGD | RTMIFN | HLA-G | LILRB1 | 0.3614 | 0.2805 | 0.321 | 0.000999 |
| TGD | RTMIFN | HMGB1 | AGTRAP | 1.7525 | 0.38 | 1.0663 | 0.000999 |
| TGD | RTMIFN | HMGB1 | LY96 | 1.7525 | 1.1488 | 1.4506 | 0.000999 |
| TGD | RTMIFN | HMGB1 | TLR2 | 1.7525 | 0.3529 | 1.0527 | 0.000999 |
| TGD | RTMIFN | HMGB2 | DAB2 | 0.3275 | 0.5996 | 0.4635 | 0.000999 |
| TGD | RTMIFN | HSP90B1 | LRP1 | 0.9338 | 0.5528 | 0.7433 | 0.000999 |
| TGD | RTMIFN | IL16 | CD4 | 0.9357 | 0.7071 | 0.8214 | 0.000999 |

| | | | | | | | |
|---|---|---|---|---|---|---|---|
| TGD | RTMIFN | IL16 | FCGR1A | 0.9357 | 0.8743 | 0.905 | 0.000999 |
| TGD | RTMIFN | ITGA4 | TFRC | 0.3968 | 0.3853 | 0.3911 | 0.000999 |
| TGD | RTMIFN | ITGAL | ICAM1 | 0.3731 | 0.5523 | 0.4627 | 0.000999 |
| TGD | RTMIFN | ITGB1 | DAB2 | 0.3657 | 0.5996 | 0.4826 | 0.000999 |
| TGD | RTMIFN | ITGB1 | ITGAX | 0.3657 | 0.2809 | 0.3233 | 0.000999 |
| TGD | RTMIFN | ITGB1 | PLAUR | 0.3657 | 1.1076 | 0.7366 | 0.000999 |
| TGD | RTMIFN | ITGB2 | ITGAX | 1.4444 | 0.2809 | 0.8627 | 0.000999 |
| TGD | RTMIFN | ITGB2 | PLAUR | 1.4444 | 1.1076 | 1.276 | 0.000999 |
| TGD | RTMIFN | LRPAP1 | LRP1 | 0.3161 | 0.5528 | 0.4344 | 0.000999 |
| TGD | RTMIFN | LTB | CD40 | 1.6829 | 0.4077 | 1.0453 | 0.000999 |
| TGD | RTMIFN | LTB | ITGB1 | 1.6829 | 0.528 | 1.1054 | 0.000999 |
| TGD | RTMIFN | LTB | TNFRSF1A | 1.6829 | 0.4908 | 1.0868 | 0.000999 |
| TGD | RTMIFN | MIF | CD74 | 1.6335 | 4.5905 | 3.112 | 0.000999 |
| TGD | RTMIFN | PTPRC | CD4 | 1.8653 | 0.7071 | 1.2862 | 0.000999 |
| TGD | RTMIFN | PTPRC | CD44 | 1.8653 | 1.2543 | 1.5598 | 0.000999 |
| TGD | RTMIFN | PTPRC | IFNAR1 | 1.8653 | 0.3883 | 1.1268 | 0.000999 |
| TGD | RTMIFN | RPS19 | C5AR1 | 3.8902 | 0.5634 | 2.2268 | 0.000999 |
| TGD | RTMIFN | RPS19 | CD4 | 3.8902 | 0.7071 | 2.2986 | 0.000999 |
| TGD | RTMIFN | RPS19 | ICAM1 | 3.8902 | 0.5523 | 2.2212 | 0.000999 |
| TGD | RTMIFN | SELL | CD4 | 0.3555 | 0.7071 | 0.5313 | 0.000999 |
| TGD | RTMIFN | TGFB1 | ENG | 0.7703 | 0.3505 | 0.5604 | 0.000999 |
| TGD | RTMIFN | TGFB1 | ITGB1 | 0.7703 | 0.528 | 0.6492 | 0.000999 |
| TGD | RTMIFN | TGFB1 | ITGB5 | 0.7703 | 0.2827 | 0.5265 | 0.000999 |
| TGD | RTMIFN | TGFB1 | TLR2 | 0.7703 | 0.3529 | 0.5616 | 0.000999 |
| TGD | RTMIFN | XCL1 | GPR65 | 1.5822 | 0.3227 | 0.9524 | 0.000999 |
| TGD | RTMIFN | XCL1 | LPAR6 | 1.5822 | 0.4248 | 1.0035 | 0.000999 |
| TGD | RTMIFN | XCL1 | PTAFR | 1.5822 | 0.2739 | 0.928 | 0.000999 |
| TGD | RTMIFN | XCL2 | LPAR6 | 0.4864 | 0.4248 | 0.4556 | 0.000999 |
| TGD | RTMIFN | XCL2 | PTAFR | 0.4864 | 0.2739 | 0.3802 | 0.000999 |
| TGD | RTMIFN | ITGB2 | ICAM1 | 1.4444 | 0.5523 | 0.9984 | 0.002 |

| | | | | | | | |
|---|---|---|---|---|---|---|---|
| TGD | RTMIFN | ICAM1 | ITGAX | 0.2915 | 0.2809 | 0.2862 | 0.003 |
| TGD | RTMIFN | ITGB1 | ENG | 0.3657 | 0.3505 | 0.3581 | 0.003 |
| TGD | RTMIFN | ICAM1 | IL2RA | 0.2915 | 0.2724 | 0.2819 | 0.004 |
| TGD | RTMIFN | ICAM3 | ITGB2 | 0.8816 | 1.456 | 1.1688 | 0.004 |
| TGD | RTMIFN | ALOX5AP | ALOX5 | 0.5564 | 0.4066 | 0.4815 | 0.00599 |
| TGD | RTMIFN | HSP90B1 | ITGB1 | 0.9338 | 0.528 | 0.7309 | 0.00599 |
| TGD | RTMIFN | MIF | TNFRSF14 | 1.6335 | 0.4163 | 1.0249 | 0.00599 |
| TGD | RTMIFN | HSP90B1 | TNFRSF1B | 0.9338 | 0.3537 | 0.6437 | 0.00999 |
| TGD | RTMIFN | ITGA4 | CD44 | 0.3968 | 1.2543 | 0.8256 | 0.00999 |

| **Supplementary Table 8: Communication results between macrophages and lymphocytes subpopulations, and malignant cells** | | | | | | | |
|---|---|---|---|---|---|---|---|
| **RTM_IFN vs. malignant cells** | | | | | | | |
| celltype_01 | celltype_02 | ligand | receptor | ligand_mean_expression | receptor_mean_expression | ligand_receptor_mean_expression | pvalue |
| RTMIFN | breasttumor | ANXA1 | CD44 | 1.0204 | 0.3247 | 0.6725 | 0.000999 |
| RTMIFN | breasttumor | APOE | SCARB1 | 2.8059 | 0.2628 | 1.5343 | 0.000999 |
| RTMIFN | breasttumor | APOE | SDC2 | 2.8059 | 1.071 | 1.9384 | 0.000999 |
| RTMIFN | breasttumor | CALM1 | MYLK | 1.3269 | 0.4064 | 0.8666 | 0.000999 |
| RTMIFN | breasttumor | CALM1 | RPSA | 1.3269 | 2.358 | 1.8424 | 0.000999 |
| RTMIFN | breasttumor | CALM1 | TNFRSF1A | 1.3269 | 0.2536 | 0.7902 | 0.000999 |
| RTMIFN | breasttumor | CALM2 | MYLK | 1.8628 | 0.4064 | 1.1346 | 0.000999 |
| RTMIFN | breasttumor | CALM3 | MYLK | 0.8659 | 0.4064 | 0.6361 | 0.000999 |
| RTMIFN | breasttumor | COL1A1 | CD44 | 0.6812 | 0.3247 | 0.503 | 0.000999 |
| RTMIFN | breasttumor | COL1A1 | ITGB1 | 0.6812 | 0.2939 | 0.4875 | 0.000999 |
| RTMIFN | breasttumor | FN1 | CD44 | 1.3064 | 0.3247 | 0.8156 | 0.000999 |
| RTMIFN | breasttumor | FN1 | ITGB1 | 1.3064 | 0.2939 | 0.8001 | 0.000999 |
| RTMIFN | breasttumor | FN1 | RPSA | 1.3064 | 2.358 | 1.8322 | 0.000999 |
| RTMIFN | breasttumor | FN1 | SDC2 | 1.3064 | 1.071 | 1.1887 | 0.000999 |
| RTMIFN | breasttumor | GRN | TNFRSF1A | 1.889 | 0.2536 | 1.0713 | 0.000999 |
| RTMIFN | breasttumor | HBEGF | CD44 | 0.3607 | 0.3247 | 0.3427 | 0.000999 |
| RTMIFN | breasttumor | HLA-DRA | SCARB1 | 4.1983 | 0.2628 | 2.2305 | 0.000999 |
| RTMIFN | breasttumor | LGALS3 | ITGB1 | 1.3808 | 0.2939 | 0.8373 | 0.000999 |
| RTMIFN | breasttumor | MMP9 | CD44 | 0.3709 | 0.3247 | 0.3478 | 0.000999 |
| RTMIFN | breasttumor | MMP9 | ITGB1 | 0.3709 | 0.2939 | 0.3324 | 0.000999 |
| RTMIFN | breasttumor | PTPRC | CD44 | 0.6543 | 0.3247 | 0.4895 | 0.000999 |
| RTMIFN | breasttumor | RTN4 | CD44 | 1.0111 | 0.3247 | 0.6679 | 0.000999 |
| RTMIFN | breasttumor | RTN4 | CD99 | 1.0111 | 0.3871 | 0.6991 | 0.000999 |
| RTMIFN | breasttumor | SERPINA1 | SDC2 | 1.1249 | 1.071 | 1.0979 | 0.000999 |
| RTMIFN | breasttumor | SPP1 | CD44 | 2.4585 | 0.3247 | 1.3916 | 0.000999 |
| RTMIFN | breasttumor | SPP1 | ITGB1 | 2.4585 | 0.2939 | 1.3762 | 0.000999 |
| RTMIFN | breasttumor | TGFB1 | ITGB1 | 0.3271 | 0.2939 | 0.3105 | 0.000999 |

| | | | | | | | |
|---|---|---|---|---|---|---|---|
| RTMIFN | breasttumor | TGFB1 | SDC2 | 0.3271 | 1.071 | 0.6991 | 0.000999 |
| RTMIFN | breasttumor | TNFSF13B | ITGB1 | 1.0395 | 0.2939 | 0.6667 | 0.000999 |
| RTMIFN | breasttumor | CD99 | CD99 | 0.6857 | 0.3871 | 0.5364 | 0.003 |
| RTMIFN | breasttumor | ITGB1 | CD151 | 0.528 | 0.3569 | 0.4424 | 0.005 |
| RTMIFN | breasttumor | ITGB1 | RACK1 | 0.528 | 2.6334 | 1.5807 | 0.005 |
| RTMIFN | colorectaltumor | APOE | SDC1 | 2.8059 | 0.2575 | 1.5317 | 0.000999 |
| RTMIFN | colorectaltumor | CD99 | CD99 | 0.6857 | 0.3443 | 0.515 | 0.000999 |
| RTMIFN | colorectaltumor | FN1 | CD44 | 1.3064 | 0.2512 | 0.7788 | 0.000999 |
| RTMIFN | colorectaltumor | FN1 | ITGB1 | 1.3064 | 0.4214 | 0.8639 | 0.000999 |
| RTMIFN | colorectaltumor | FN1 | RPSA | 1.3064 | 2.251 | 1.7787 | 0.000999 |
| RTMIFN | colorectaltumor | FN1 | SDC1 | 1.3064 | 0.2575 | 0.7819 | 0.000999 |
| RTMIFN | colorectaltumor | MMP9 | ITGB1 | 0.3709 | 0.4214 | 0.3962 | 0.000999 |
| RTMIFN | colorectaltumor | PTPRC | CD44 | 0.6543 | 0.2512 | 0.4527 | 0.000999 |
| RTMIFN | colorectaltumor | RTN4 | CD99 | 1.0111 | 0.3443 | 0.6777 | 0.000999 |
| RTMIFN | colorectaltumor | SPP1 | CD44 | 2.4585 | 0.2512 | 1.3548 | 0.000999 |
| RTMIFN | colorectaltumor | SPP1 | ITGB1 | 2.4585 | 0.4214 | 1.4399 | 0.000999 |
| RTMIFN | colorectaltumor | SPP1 | SDC1 | 2.4585 | 0.2575 | 1.358 | 0.000999 |
| RTMIFN | colorectaltumor | TGFB1 | ITGB1 | 0.3271 | 0.4214 | 0.3742 | 0.000999 |
| RTMIFN | colorectaltumor | TNFSF13B | ITGB1 | 1.0395 | 0.4214 | 0.7304 | 0.000999 |
| RTMIFN | colorectaltumor | RTN4 | CD44 | 1.0111 | 0.2512 | 0.6312 | 0.004 |
| RTMIFN | livertumor | A2M | TNFRSF14 | 0.8392 | 0.2788 | 0.559 | 0.000999 |
| RTMIFN | livertumor | ADM | RAMP1 | 0.3632 | 0.4499 | 0.4065 | 0.000999 |
| RTMIFN | livertumor | ANXA1 | CD44 | 1.0204 | 0.8016 | 0.911 | 0.000999 |
| RTMIFN | livertumor | ANXA1 | MET | 1.0204 | 0.2983 | 0.6593 | 0.000999 |
| RTMIFN | livertumor | ANXA1 | PTPRF | 1.0204 | 0.384 | 0.7022 | 0.000999 |
| RTMIFN | livertumor | APOE | SDC1 | 2.8059 | 0.5354 | 1.6706 | 0.000999 |
| RTMIFN | livertumor | APOE | SDC4 | 2.8059 | 0.6068 | 1.7063 | 0.000999 |
| RTMIFN | livertumor | CALM1 | INSR | 1.3269 | 0.3039 | 0.8154 | 0.000999 |
| RTMIFN | livertumor | CALM1 | RPSA | 1.3269 | 2.6317 | 1.9793 | 0.000999 |
| RTMIFN | livertumor | COL1A1 | CD44 | 0.6812 | 0.8016 | 0.7414 | 0.000999 |

| | | | | | | | |
|---|---|---|---|---|---|---|---|
| RTMIFN | livertumor | COL1A1 | ITGA2 | 0.6812 | 0.3983 | 0.5397 | 0.000999 |
| RTMIFN | livertumor | COL1A1 | ITGA3 | 0.6812 | 0.3913 | 0.5362 | 0.000999 |
| RTMIFN | livertumor | COL1A1 | ITGB1 | 0.6812 | 1.6044 | 1.1428 | 0.000999 |
| RTMIFN | livertumor | COL1A1 | SDC1 | 0.6812 | 0.5354 | 0.6083 | 0.000999 |
| RTMIFN | livertumor | FN1 | CD44 | 1.3064 | 0.8016 | 1.054 | 0.000999 |
| RTMIFN | livertumor | FN1 | ITGA2 | 1.3064 | 0.3983 | 0.8523 | 0.000999 |
| RTMIFN | livertumor | FN1 | ITGA3 | 1.3064 | 0.3913 | 0.8488 | 0.000999 |
| RTMIFN | livertumor | FN1 | ITGB1 | 1.3064 | 1.6044 | 1.4554 | 0.000999 |
| RTMIFN | livertumor | FN1 | RPSA | 1.3064 | 2.6317 | 1.969 | 0.000999 |
| RTMIFN | livertumor | FN1 | SDC1 | 1.3064 | 0.5354 | 0.9209 | 0.000999 |
| RTMIFN | livertumor | FN1 | TFRC | 1.3064 | 0.2899 | 0.7981 | 0.000999 |
| RTMIFN | livertumor | HAMP | SLC40A1 | 0.4003 | 0.4972 | 0.4487 | 0.000999 |
| RTMIFN | livertumor | HBEGF | ERBB3 | 0.3607 | 0.2644 | 0.3125 | 0.000999 |
| RTMIFN | livertumor | HLA-A | ERBB3 | 3.0434 | 0.2644 | 1.6539 | 0.000999 |
| RTMIFN | livertumor | HLA-DRA | EPHB4 | 4.1983 | 0.4535 | 2.3259 | 0.000999 |
| RTMIFN | livertumor | HLA-E | EPHB4 | 1.9163 | 0.4535 | 1.1849 | 0.000999 |
| RTMIFN | livertumor | HLA-E | ITGA6 | 1.9163 | 0.5211 | 1.2187 | 0.000999 |
| RTMIFN | livertumor | HLA-E | PLXNB2 | 1.9163 | 0.265 | 1.0906 | 0.000999 |
| RTMIFN | livertumor | HMGB2 | CDH1 | 0.4676 | 0.4896 | 0.4786 | 0.000999 |
| RTMIFN | livertumor | HMGB2 | DSG2 | 0.4676 | 0.7479 | 0.6078 | 0.000999 |
| RTMIFN | livertumor | LGALS3 | ITGA2 | 1.3808 | 0.3983 | 0.8895 | 0.000999 |
| RTMIFN | livertumor | LGALS3 | ITGB1 | 1.3808 | 1.6044 | 1.4926 | 0.000999 |
| RTMIFN | livertumor | LGALS3 | ITGB4 | 1.3808 | 0.5518 | 0.9663 | 0.000999 |
| RTMIFN | livertumor | LGALS3 | TFRC | 1.3808 | 0.2899 | 0.8353 | 0.000999 |
| RTMIFN | livertumor | MMP9 | CD44 | 0.3709 | 0.8016 | 0.5863 | 0.000999 |
| RTMIFN | livertumor | MMP9 | CDH1 | 0.3709 | 0.4896 | 0.4303 | 0.000999 |
| RTMIFN | livertumor | MMP9 | ITGB1 | 0.3709 | 1.6044 | 0.9877 | 0.000999 |
| RTMIFN | livertumor | NAMPT | INSR | 1.0012 | 0.3039 | 0.6526 | 0.000999 |
| RTMIFN | livertumor | PTPRC | CD44 | 0.6543 | 0.8016 | 0.7279 | 0.000999 |
| RTMIFN | livertumor | PTPRC | INSR | 0.6543 | 0.3039 | 0.4791 | 0.000999 |

| | | | | | | | |
|---|---|---|---|---|---|---|---|
| RTMIFN | livertumor | PTPRC | MET | 0.6543 | 0.2983 | 0.4763 | 0.000999 |
| RTMIFN | livertumor | SPP1 | CD44 | 2.4585 | 0.8016 | 1.63 | 0.000999 |
| RTMIFN | livertumor | SPP1 | ITGB1 | 2.4585 | 1.6044 | 2.0315 | 0.000999 |
| RTMIFN | livertumor | SPP1 | SDC1 | 2.4585 | 0.5354 | 1.497 | 0.000999 |
| RTMIFN | livertumor | TGFB1 | ITGB1 | 0.3271 | 1.6044 | 0.9658 | 0.000999 |
| RTMIFN | livertumor | TGFB1 | VTN | 0.3271 | 0.7399 | 0.5335 | 0.000999 |
| RTMIFN | livertumor | TNFSF12 | TNFRSF12A | 0.3767 | 1.0484 | 0.7125 | 0.000999 |
| RTMIFN | livertumor | TNFSF13B | ITGB1 | 1.0395 | 1.6044 | 1.322 | 0.000999 |
| RTMIFN | livertumor | HBEGF | CD44 | 0.3607 | 0.8016 | 0.5811 | 0.002 |
| RTMIFN | lungtumor | APOE | SDC1 | 2.8059 | 0.2715 | 1.5387 | 0.000999 |
| RTMIFN | lungtumor | APOE | SDC4 | 2.8059 | 0.3516 | 1.5787 | 0.000999 |
| RTMIFN | lungtumor | C3 | CXCR4 | 0.6266 | 0.2684 | 0.4475 | 0.000999 |
| RTMIFN | lungtumor | CALM1 | RPSA | 1.3269 | 1.6057 | 1.4663 | 0.000999 |
| RTMIFN | lungtumor | COL1A1 | CD44 | 0.6812 | 0.3802 | 0.5307 | 0.000999 |
| RTMIFN | lungtumor | COL1A1 | SDC1 | 0.6812 | 0.2715 | 0.4764 | 0.000999 |
| RTMIFN | lungtumor | CXCL10 | CXCR4 | 0.9959 | 0.2684 | 0.6321 | 0.000999 |
| RTMIFN | lungtumor | CXCL12 | CXCR4 | 0.2863 | 0.2684 | 0.2774 | 0.000999 |
| RTMIFN | lungtumor | CXCL16 | CXCR4 | 0.9727 | 0.2684 | 0.6206 | 0.000999 |
| RTMIFN | lungtumor | CXCL2 | CXCR4 | 0.7506 | 0.2684 | 0.5095 | 0.000999 |
| RTMIFN | lungtumor | CXCL3 | CXCR4 | 0.4907 | 0.2684 | 0.3796 | 0.000999 |
| RTMIFN | lungtumor | CXCL9 | CXCR4 | 0.7322 | 0.2684 | 0.5003 | 0.000999 |
| RTMIFN | lungtumor | FN1 | CD44 | 1.3064 | 0.3802 | 0.8433 | 0.000999 |
| RTMIFN | lungtumor | FN1 | RPSA | 1.3064 | 1.6057 | 1.4561 | 0.000999 |
| RTMIFN | lungtumor | FN1 | SDC1 | 1.3064 | 0.2715 | 0.7889 | 0.000999 |
| RTMIFN | lungtumor | HBEGF | CD44 | 0.3607 | 0.3802 | 0.3704 | 0.000999 |
| RTMIFN | lungtumor | MMP9 | CD44 | 0.3709 | 0.3802 | 0.3756 | 0.000999 |
| RTMIFN | lungtumor | PTPRC | CD44 | 0.6543 | 0.3802 | 0.5172 | 0.000999 |
| RTMIFN | lungtumor | PTPRC | CXCR4 | 0.6543 | 0.2684 | 0.4613 | 0.000999 |
| RTMIFN | lungtumor | RTN4 | CD44 | 1.0111 | 0.3802 | 0.6957 | 0.000999 |
| RTMIFN | lungtumor | RTN4 | CD99 | 1.0111 | 0.5938 | 0.8025 | 0.000999 |

| | | | | | | | |
|---|---|---|---|---|---|---|---|
| RTMIFN | lungtumor | SPP1 | CD44 | 2.4585 | 0.3802 | 1.4193 | 0.000999 |
| RTMIFN | lungtumor | SPP1 | SDC1 | 2.4585 | 0.2715 | 1.365 | 0.000999 |
| RTMIFN | lungtumor | TNFSF12 | TNFRSF12A | 0.3767 | 0.441 | 0.4088 | 0.000999 |
| RTMIFN | lungtumor | ITGB1 | CD151 | 0.528 | 0.6776 | 0.6028 | 0.002 |
| RTMIFN | lungtumor | ITGB1 | CD46 | 0.528 | 0.434 | 0.481 | 0.002 |
| RTMIFN | lungtumor | ITGB1 | CD47 | 0.528 | 0.3255 | 0.4267 | 0.002 |
| RTMIFN | lungtumor | ITGB1 | RACK1 | 0.528 | 2.2108 | 1.3694 | 0.002 |
| RTMIFN | lungtumor | ITGB1 | SDC1 | 0.528 | 0.2715 | 0.3997 | 0.002 |
| RTMIFN | lungtumor | CXCL11 | CXCR4 | 0.3356 | 0.2684 | 0.302 | 0.004 |
| RTMIFN | melanomatumo | AIMP1 | RACK1 | 0.3208 | 4.145 | 2.2329 | 0.000999 |
| RTMIFN | melanomatumo | ANXA1 | EDNRB | 1.0204 | 0.5666 | 0.7935 | 0.000999 |
| RTMIFN | melanomatumo | APP | EDNRB | 0.2822 | 0.5666 | 0.4244 | 0.000999 |
| RTMIFN | melanomatumo | CALM1 | RPSA | 1.3269 | 3.7787 | 2.5528 | 0.000999 |
| RTMIFN | melanomatumo | FN1 | RPSA | 1.3064 | 3.7787 | 2.5425 | 0.000999 |
| RTMIFN | melanomatumo | ITGB1 | RACK1 | 0.528 | 4.145 | 2.3365 | 0.000999 |
| RTMIFN | melanomatumo | NAMPT | ANXA6 | 1.0012 | 0.3625 | 0.6819 | 0.000999 |
| RTMIFN | melanomatumo | RPS19 | RPSA | 3.6736 | 3.7787 | 3.7261 | 0.000999 |
| RTMIFN | melanomatumo | RTN4 | CD99 | 1.0111 | 0.6845 | 0.8478 | 0.000999 |
| RTMIFN | melanomatumo | SPP1 | CD44 | 2.4585 | 1.2653 | 1.8619 | 0.000999 |
| RTMIFN | melanomatumo | SPP1 | ITGB1 | 2.4585 | 0.3547 | 1.4066 | 0.000999 |
| RTMIFN | melanomatumo | SPP1 | NCSTN | 2.4585 | 0.2614 | 1.36 | 0.000999 |
| RTMIFN | melanomatumo | CALR | NCSTN | 1.0231 | 0.2614 | 0.6422 | 0.002 |
| RTMIFN | melanomatumo | FN1 | CD44 | 1.3064 | 1.2653 | 1.2859 | 0.003 |
| RTMIFN | melanomatumo | FN1 | ITGB1 | 1.3064 | 0.3547 | 0.8306 | 0.005 |
| RTMIFN | melanomatumo | TNFSF13B | ITGB1 | 1.0395 | 0.3547 | 0.6971 | 0.005 |
| RTMIFN | melanomatumo | A2M | TNFRSF14 | 0.8392 | 0.5069 | 0.6731 | 0.00799 |
| RTMIFN | melanomatumo | APP | NCSTN | 0.2822 | 0.2614 | 0.2718 | 0.00799 |
| RTMIFN | ovarytumor | AREG | FOLR1 | 0.3003 | 0.5619 | 0.4311 | 0.000999 |
| RTMIFN | ovarytumor | C3 | CXCR4 | 0.6266 | 0.3688 | 0.4977 | 0.000999 |
| RTMIFN | ovarytumor | CCL20 | CXCR4 | 0.2626 | 0.3688 | 0.3157 | 0.000999 |

| | | | | | | | |
|---|---|---|---|---|---|---|---|
| RTMIFN | ovarytumor | COL1A1 | ITGB1 | 0.6812 | 0.2783 | 0.4797 | 0.000999 |
| RTMIFN | ovarytumor | CXCL10 | CXCR4 | 0.9959 | 0.3688 | 0.6823 | 0.000999 |
| RTMIFN | ovarytumor | CXCL11 | CXCR4 | 0.3356 | 0.3688 | 0.3522 | 0.000999 |
| RTMIFN | ovarytumor | CXCL12 | CXCR4 | 0.2863 | 0.3688 | 0.3276 | 0.000999 |
| RTMIFN | ovarytumor | CXCL16 | CXCR4 | 0.9727 | 0.3688 | 0.6708 | 0.000999 |
| RTMIFN | ovarytumor | CXCL2 | CXCR4 | 0.7506 | 0.3688 | 0.5597 | 0.000999 |
| RTMIFN | ovarytumor | CXCL3 | CXCR4 | 0.4907 | 0.3688 | 0.4297 | 0.000999 |
| RTMIFN | ovarytumor | CXCL9 | CXCR4 | 0.7322 | 0.3688 | 0.5505 | 0.000999 |
| RTMIFN | ovarytumor | FN1 | ITGB1 | 1.3064 | 0.2783 | 0.7923 | 0.000999 |
| RTMIFN | ovarytumor | FN1 | RPSA | 1.3064 | 2.9542 | 2.1303 | 0.000999 |
| RTMIFN | ovarytumor | LGALS3 | ITGB1 | 1.3808 | 0.2783 | 0.8295 | 0.000999 |
| RTMIFN | ovarytumor | MMP9 | ITGB1 | 0.3709 | 0.2783 | 0.3246 | 0.000999 |
| RTMIFN | ovarytumor | PTPRC | CXCR4 | 0.6543 | 0.3688 | 0.5115 | 0.000999 |
| RTMIFN | ovarytumor | SPP1 | ITGB1 | 2.4585 | 0.2783 | 1.3684 | 0.000999 |
| RTMIFN | ovarytumor | TGFB1 | ITGB1 | 0.3271 | 0.2783 | 0.3027 | 0.000999 |
| RTMIFN | ovarytumor | TNFSF12 | TNFRSF12A | 0.3767 | 0.3901 | 0.3834 | 0.000999 |
| RTMIFN | ovarytumor | TNFSF13B | ITGB1 | 1.0395 | 0.2783 | 0.6589 | 0.000999 |
| RTMIFN | ovarytumor | ITGB1 | CD151 | 0.528 | 0.4981 | 0.513 | 0.003 |
| RTMIFN | ovarytumor | ITGB1 | CD47 | 0.528 | 0.4931 | 0.5105 | 0.003 |
| RTMIFN | ovarytumor | ITGB1 | RACK1 | 0.528 | 2.9543 | 1.7411 | 0.003 |
| RTMIFN | ovarytumor | CALM1 | RPSA | 1.3269 | 2.9542 | 2.1406 | 0.004 |
| breasttumor | RTMIFN | A2M | LRP1 | 0.3129 | 0.5528 | 0.4329 | 0.000999 |
| breasttumor | RTMIFN | A2M | TNFRSF14 | 0.3129 | 0.4163 | 0.3646 | 0.000999 |
| breasttumor | RTMIFN | ANXA1 | C3AR1 | 0.4092 | 0.648 | 0.5286 | 0.000999 |
| breasttumor | RTMIFN | ANXA1 | C5AR1 | 0.4092 | 0.5634 | 0.4863 | 0.000999 |
| breasttumor | RTMIFN | ANXA1 | CD4 | 0.4092 | 0.707 | 0.5581 | 0.000999 |
| breasttumor | RTMIFN | ANXA1 | CD44 | 0.4092 | 1.2543 | 0.8317 | 0.000999 |
| breasttumor | RTMIFN | ANXA1 | CXCR4 | 0.4092 | 0.9336 | 0.6714 | 0.000999 |
| breasttumor | RTMIFN | ANXA1 | FPR1 | 0.4092 | 0.4203 | 0.4147 | 0.000999 |
| breasttumor | RTMIFN | ANXA1 | FPR3 | 0.4092 | 0.4975 | 0.4534 | 0.000999 |

| | | | | | | | |
|---|---|---|---|---|---|---|---|
| breasttumor | RTMIFN | APP | ALOX5 | 0.5409 | 0.4066 | 0.4738 | 0.000999 |
| breasttumor | RTMIFN | APP | C3AR1 | 0.5409 | 0.648 | 0.5945 | 0.000999 |
| breasttumor | RTMIFN | APP | C5AR1 | 0.5409 | 0.5634 | 0.5522 | 0.000999 |
| breasttumor | RTMIFN | APP | CXCR4 | 0.5409 | 0.9336 | 0.7373 | 0.000999 |
| breasttumor | RTMIFN | APP | FPR1 | 0.5409 | 0.4203 | 0.4806 | 0.000999 |
| breasttumor | RTMIFN | APP | LRP1 | 0.5409 | 0.5528 | 0.5469 | 0.000999 |
| breasttumor | RTMIFN | APP | TGFBR2 | 0.5409 | 0.2932 | 0.4171 | 0.000999 |
| breasttumor | RTMIFN | APP | TNFRSF14 | 0.5409 | 0.4163 | 0.4786 | 0.000999 |
| breasttumor | RTMIFN | ARF1 | CD4 | 1.6219 | 0.707 | 1.1645 | 0.000999 |
| breasttumor | RTMIFN | ARF1 | CD44 | 1.6219 | 1.2543 | 1.4381 | 0.000999 |
| breasttumor | RTMIFN | CALM1 | TNFRSF1B | 0.9015 | 0.3537 | 0.6276 | 0.000999 |
| breasttumor | RTMIFN | CALR | CD40 | 1.1835 | 0.4077 | 0.7956 | 0.000999 |
| breasttumor | RTMIFN | CALR | CD44 | 1.1835 | 1.2543 | 1.2189 | 0.000999 |
| breasttumor | RTMIFN | CALR | LRP1 | 1.1835 | 0.5528 | 0.8682 | 0.000999 |
| breasttumor | RTMIFN | COL2A1 | CD44 | 0.3935 | 1.2543 | 0.8239 | 0.000999 |
| breasttumor | RTMIFN | GSTP1 | CD44 | 2.5435 | 1.2543 | 1.8989 | 0.000999 |
| breasttumor | RTMIFN | HEBP1 | C3AR1 | 0.4062 | 0.648 | 0.5271 | 0.000999 |
| breasttumor | RTMIFN | HEBP1 | C5AR1 | 0.4062 | 0.5634 | 0.4848 | 0.000999 |
| breasttumor | RTMIFN | HEBP1 | CXCR4 | 0.4062 | 0.9336 | 0.6699 | 0.000999 |
| breasttumor | RTMIFN | HEBP1 | FPR1 | 0.4062 | 0.4203 | 0.4132 | 0.000999 |
| breasttumor | RTMIFN | HEBP1 | FPR3 | 0.4062 | 0.4975 | 0.4519 | 0.000999 |
| breasttumor | RTMIFN | HMGB1 | LY96 | 1.1815 | 1.1487 | 1.1651 | 0.000999 |
| breasttumor | RTMIFN | HMGB1 | TLR2 | 1.1815 | 0.3529 | 0.7672 | 0.000999 |
| breasttumor | RTMIFN | HMGB2 | CD44 | 0.3139 | 1.2543 | 0.7841 | 0.000999 |
| breasttumor | RTMIFN | HMGB2 | DAB2 | 0.3139 | 0.5995 | 0.4567 | 0.000999 |
| breasttumor | RTMIFN | HSP90B1 | LRP1 | 0.7786 | 0.5528 | 0.6657 | 0.000999 |
| breasttumor | RTMIFN | HSP90B1 | TNFRSF1B | 0.7786 | 0.3537 | 0.5661 | 0.000999 |
| breasttumor | RTMIFN | ITGB1 | CD47 | 0.2939 | 0.468 | 0.3809 | 0.000999 |
| breasttumor | RTMIFN | ITGB1 | DAB2 | 0.2939 | 0.5995 | 0.4467 | 0.000999 |
| breasttumor | RTMIFN | ITGB1 | ENG | 0.2939 | 0.3504 | 0.3222 | 0.000999 |

| | | | | | | | |
|---|---|---|---|---|---|---|---|
| breasttumor | RTMIFN | ITGB1 | ITGAX | 0.2939 | 0.2809 | 0.2874 | 0.000999 |
| breasttumor | RTMIFN | ITGB1 | PLAUR | 0.2939 | 1.1076 | 0.7007 | 0.000999 |
| breasttumor | RTMIFN | LGALS3 | CLEC7A | 0.8391 | 0.6166 | 0.7279 | 0.000999 |
| breasttumor | RTMIFN | LGALS3 | FCGR2A | 0.8391 | 1.0157 | 0.9274 | 0.000999 |
| breasttumor | RTMIFN | LGALS3 | FCGR2B | 0.8391 | 0.7852 | 0.8122 | 0.000999 |
| breasttumor | RTMIFN | LGALS3 | TFRC | 0.8391 | 0.3853 | 0.6122 | 0.000999 |
| breasttumor | RTMIFN | LGALS3 | TGFBR2 | 0.8391 | 0.2932 | 0.5661 | 0.000999 |
| breasttumor | RTMIFN | LRPAP1 | LRP1 | 0.2756 | 0.5528 | 0.4142 | 0.000999 |
| breasttumor | RTMIFN | MDK | LRP1 | 0.592 | 0.5528 | 0.5724 | 0.000999 |
| breasttumor | RTMIFN | MIF | CD74 | 3.2716 | 4.5904 | 3.931 | 0.000999 |
| breasttumor | RTMIFN | MIF | TNFRSF14 | 3.2716 | 0.4163 | 1.844 | 0.000999 |
| breasttumor | RTMIFN | NPW | C3AR1 | 0.5478 | 0.648 | 0.5979 | 0.000999 |
| breasttumor | RTMIFN | NPW | C5AR1 | 0.5478 | 0.5634 | 0.5556 | 0.000999 |
| breasttumor | RTMIFN | NPW | CXCR4 | 0.5478 | 0.9336 | 0.7407 | 0.000999 |
| breasttumor | RTMIFN | NPW | FPR1 | 0.5478 | 0.4203 | 0.484 | 0.000999 |
| breasttumor | RTMIFN | PSAP | LRP1 | 0.5972 | 0.5528 | 0.575 | 0.000999 |
| breasttumor | RTMIFN | RPS19 | C5AR1 | 3.691 | 0.5634 | 2.1272 | 0.000999 |
| breasttumor | RTMIFN | RPS19 | CD4 | 3.691 | 0.707 | 2.199 | 0.000999 |
| breasttumor | RTMIFN | RPS19 | ICAM1 | 3.691 | 0.5523 | 2.1216 | 0.000999 |
| breasttumor | RTMIFN | RTN4 | CD44 | 0.3318 | 1.2543 | 0.7931 | 0.000999 |
| breasttumor | RTMIFN | SERPINE2 | LRP1 | 0.6001 | 0.5528 | 0.5765 | 0.000999 |
| breasttumor | RTMIFN | TF | CD4 | 0.2688 | 0.707 | 0.4879 | 0.000999 |
| breasttumor | RTMIFN | TF | TFRC | 0.2688 | 0.3853 | 0.327 | 0.000999 |
| breasttumor | RTMIFN | TIMP1 | LRP1 | 0.7045 | 0.5528 | 0.6286 | 0.000999 |
| breasttumor | RTMIFN | CD99 | CD99 | 0.3871 | 0.6857 | 0.5364 | 0.003 |
| breasttumor | RTMIFN | COL2A1 | ITGB1 | 0.3935 | 0.528 | 0.4607 | 0.004 |
| breasttumor | RTMIFN | RTN4 | CD99 | 0.3318 | 0.6857 | 0.5088 | 0.004 |
| breasttumor | RTMIFN | HSP90B1 | ITGB1 | 0.7786 | 0.528 | 0.6533 | 0.005 |
| breasttumor | RTMIFN | CALM1 | TNFRSF1A | 0.9015 | 0.4908 | 0.6961 | 0.00699 |
| breasttumor | RTMIFN | LGALS3 | ITGB1 | 0.8391 | 0.528 | 0.6835 | 0.00899 |

| | | | | | | | |
|---|---|---|---|---|---|---|---|
| colorectaltumor | RTMIFN | ANXA1 | C3AR1 | 0.5916 | 0.648 | 0.6198 | 0.000999 |
| colorectaltumor | RTMIFN | ANXA1 | C5AR1 | 0.5916 | 0.5634 | 0.5775 | 0.000999 |
| colorectaltumor | RTMIFN | ANXA1 | CD4 | 0.5916 | 0.707 | 0.6493 | 0.000999 |
| colorectaltumor | RTMIFN | ANXA1 | CD44 | 0.5916 | 1.2543 | 0.923 | 0.000999 |
| colorectaltumor | RTMIFN | ANXA1 | CXCR4 | 0.5916 | 0.9336 | 0.7626 | 0.000999 |
| colorectaltumor | RTMIFN | ANXA1 | FPR1 | 0.5916 | 0.4203 | 0.5059 | 0.000999 |
| colorectaltumor | RTMIFN | ANXA1 | FPR3 | 0.5916 | 0.4975 | 0.5446 | 0.000999 |
| colorectaltumor | RTMIFN | APP | ALOX5 | 0.4966 | 0.4066 | 0.4516 | 0.000999 |
| colorectaltumor | RTMIFN | APP | C3AR1 | 0.4966 | 0.648 | 0.5723 | 0.000999 |
| colorectaltumor | RTMIFN | APP | C5AR1 | 0.4966 | 0.5634 | 0.53 | 0.000999 |
| colorectaltumor | RTMIFN | APP | CXCR4 | 0.4966 | 0.9336 | 0.7151 | 0.000999 |
| colorectaltumor | RTMIFN | APP | FPR1 | 0.4966 | 0.4203 | 0.4584 | 0.000999 |
| colorectaltumor | RTMIFN | APP | LRP1 | 0.4966 | 0.5528 | 0.5247 | 0.000999 |
| colorectaltumor | RTMIFN | APP | TGFBR2 | 0.4966 | 0.2932 | 0.3949 | 0.000999 |
| colorectaltumor | RTMIFN | APP | TNFRSF14 | 0.4966 | 0.4163 | 0.4565 | 0.000999 |
| colorectaltumor | RTMIFN | ARF1 | CD4 | 0.6489 | 0.707 | 0.678 | 0.000999 |
| colorectaltumor | RTMIFN | ARF1 | CD44 | 0.6489 | 1.2543 | 0.9516 | 0.000999 |
| colorectaltumor | RTMIFN | CALM1 | TNFRSF1A | 1.1676 | 0.4908 | 0.8292 | 0.000999 |
| colorectaltumor | RTMIFN | CALM1 | TNFRSF1B | 1.1676 | 0.3537 | 0.7606 | 0.000999 |
| colorectaltumor | RTMIFN | CALR | CD40 | 0.4934 | 0.4077 | 0.4505 | 0.000999 |
| colorectaltumor | RTMIFN | CALR | CD44 | 0.4934 | 1.2543 | 0.8738 | 0.000999 |
| colorectaltumor | RTMIFN | CALR | LRP1 | 0.4934 | 0.5528 | 0.5231 | 0.000999 |
| colorectaltumor | RTMIFN | CD99 | CD99 | 0.3443 | 0.6857 | 0.515 | 0.000999 |
| colorectaltumor | RTMIFN | COL1A1 | CD44 | 0.4875 | 1.2543 | 0.8709 | 0.000999 |
| colorectaltumor | RTMIFN | GSTP1 | CD44 | 2.8039 | 1.2543 | 2.0291 | 0.000999 |
| colorectaltumor | RTMIFN | HMGB1 | AGTRAP | 2.155 | 0.38 | 1.2675 | 0.000999 |
| colorectaltumor | RTMIFN | HMGB1 | LY96 | 2.155 | 1.1487 | 1.6518 | 0.000999 |
| colorectaltumor | RTMIFN | HMGB1 | TLR2 | 2.155 | 0.3529 | 1.254 | 0.000999 |
| colorectaltumor | RTMIFN | HMGB2 | CD44 | 0.5347 | 1.2543 | 0.8945 | 0.000999 |
| colorectaltumor | RTMIFN | HMGB2 | DAB2 | 0.5347 | 0.5995 | 0.5671 | 0.000999 |

| | | | | | | | |
|---|---|---|---|---|---|---|---|
| colorectaltumor | RTMIFN | HSP90B1 | LRP1 | 0.5887 | 0.5528 | 0.5707 | 0.000999 |
| colorectaltumor | RTMIFN | HSP90B1 | TNFRSF1B | 0.5887 | 0.3537 | 0.4712 | 0.000999 |
| colorectaltumor | RTMIFN | ITGB1 | DAB2 | 0.4214 | 0.5995 | 0.5105 | 0.000999 |
| colorectaltumor | RTMIFN | ITGB1 | ENG | 0.4214 | 0.3504 | 0.3859 | 0.000999 |
| colorectaltumor | RTMIFN | ITGB1 | ITGAX | 0.4214 | 0.2809 | 0.3512 | 0.000999 |
| colorectaltumor | RTMIFN | ITGB1 | PLAUR | 0.4214 | 1.1076 | 0.7645 | 0.000999 |
| colorectaltumor | RTMIFN | LGALS3 | CLEC7A | 1.5563 | 0.6166 | 1.0865 | 0.000999 |
| colorectaltumor | RTMIFN | LGALS3 | FCGR2A | 1.5563 | 1.0157 | 1.286 | 0.000999 |
| colorectaltumor | RTMIFN | LGALS3 | FCGR2B | 1.5563 | 0.7852 | 1.1707 | 0.000999 |
| colorectaltumor | RTMIFN | LGALS3 | TFRC | 1.5563 | 0.3853 | 0.9708 | 0.000999 |
| colorectaltumor | RTMIFN | LGALS3 | TGFBR2 | 1.5563 | 0.2932 | 0.9247 | 0.000999 |
| colorectaltumor | RTMIFN | LRPAP1 | LRP1 | 0.2533 | 0.5528 | 0.4031 | 0.000999 |
| colorectaltumor | RTMIFN | MDK | LRP1 | 0.7548 | 0.5528 | 0.6538 | 0.000999 |
| colorectaltumor | RTMIFN | MIF | CD74 | 1.6741 | 4.5904 | 3.1322 | 0.000999 |
| colorectaltumor | RTMIFN | MIF | TNFRSF14 | 1.6741 | 0.4163 | 1.0452 | 0.000999 |
| colorectaltumor | RTMIFN | RPS19 | C5AR1 | 4.0956 | 0.5634 | 2.3295 | 0.000999 |
| colorectaltumor | RTMIFN | RPS19 | CD4 | 4.0956 | 0.707 | 2.4013 | 0.000999 |
| colorectaltumor | RTMIFN | RPS19 | ICAM1 | 4.0956 | 0.5523 | 2.3239 | 0.000999 |
| colorectaltumor | RTMIFN | RTN4 | CD44 | 0.4148 | 1.2543 | 0.8346 | 0.000999 |
| colorectaltumor | RTMIFN | TIMP1 | LRP1 | 0.8931 | 0.5528 | 0.7229 | 0.000999 |
| livertumor | RTMIFN | AGRN | LRP1 | 0.3492 | 0.5528 | 0.451 | 0.000999 |
| livertumor | RTMIFN | AGT | ATP6AP2 | 0.4701 | 0.8552 | 0.6626 | 0.000999 |
| livertumor | RTMIFN | AGT | C3AR1 | 0.4701 | 0.648 | 0.559 | 0.000999 |
| livertumor | RTMIFN | AGT | C5AR1 | 0.4701 | 0.5634 | 0.5167 | 0.000999 |
| livertumor | RTMIFN | AGT | CXCR4 | 0.4701 | 0.9336 | 0.7018 | 0.000999 |
| livertumor | RTMIFN | AGT | FPR1 | 0.4701 | 0.4203 | 0.4452 | 0.000999 |
| livertumor | RTMIFN | AGT | GPR65 | 0.4701 | 0.3227 | 0.3964 | 0.000999 |
| livertumor | RTMIFN | AGT | LPAR6 | 0.4701 | 0.4247 | 0.4474 | 0.000999 |
| livertumor | RTMIFN | AGT | PTAFR | 0.4701 | 0.2739 | 0.372 | 0.000999 |
| livertumor | RTMIFN | ANXA1 | C3AR1 | 0.3041 | 0.648 | 0.4761 | 0.000999 |

| | | | | | | | |
|---|---|---|---|---|---|---|---|
| livertumor | RTMIFN | ANXA1 | C5AR1 | 0.3041 | 0.5634 | 0.4338 | 0.000999 |
| livertumor | RTMIFN | ANXA1 | CD4 | 0.3041 | 0.707 | 0.5056 | 0.000999 |
| livertumor | RTMIFN | ANXA1 | CXCR4 | 0.3041 | 0.9336 | 0.6189 | 0.000999 |
| livertumor | RTMIFN | ANXA1 | FPR1 | 0.3041 | 0.4203 | 0.3622 | 0.000999 |
| livertumor | RTMIFN | ANXA1 | FPR3 | 0.3041 | 0.4975 | 0.4008 | 0.000999 |
| livertumor | RTMIFN | APOA1 | ABCA1 | 0.2888 | 0.4259 | 0.3573 | 0.000999 |
| livertumor | RTMIFN | APOA1 | LRP1 | 0.2888 | 0.5528 | 0.4208 | 0.000999 |
| livertumor | RTMIFN | APOA1 | MSR1 | 0.2888 | 0.587 | 0.4379 | 0.000999 |
| livertumor | RTMIFN | APP | ALOX5 | 0.5626 | 0.4066 | 0.4846 | 0.000999 |
| livertumor | RTMIFN | APP | C3AR1 | 0.5626 | 0.648 | 0.6053 | 0.000999 |
| livertumor | RTMIFN | APP | C5AR1 | 0.5626 | 0.5634 | 0.563 | 0.000999 |
| livertumor | RTMIFN | APP | CXCR4 | 0.5626 | 0.9336 | 0.7481 | 0.000999 |
| livertumor | RTMIFN | APP | FPR1 | 0.5626 | 0.4203 | 0.4914 | 0.000999 |
| livertumor | RTMIFN | APP | LRP1 | 0.5626 | 0.5528 | 0.5577 | 0.000999 |
| livertumor | RTMIFN | AREG | ICAM1 | 0.448 | 0.5523 | 0.5001 | 0.000999 |
| livertumor | RTMIFN | ARF1 | CD4 | 0.6936 | 0.707 | 0.7003 | 0.000999 |
| livertumor | RTMIFN | ARF1 | CD44 | 0.6936 | 1.2543 | 0.9739 | 0.000999 |
| livertumor | RTMIFN | C3 | C3AR1 | 0.6584 | 0.648 | 0.6532 | 0.000999 |
| livertumor | RTMIFN | C3 | C5AR1 | 0.6584 | 0.5634 | 0.6109 | 0.000999 |
| livertumor | RTMIFN | C3 | CXCR4 | 0.6584 | 0.9336 | 0.796 | 0.000999 |
| livertumor | RTMIFN | C3 | FPR1 | 0.6584 | 0.4203 | 0.5394 | 0.000999 |
| livertumor | RTMIFN | C3 | ITGAM | 0.6584 | 0.2708 | 0.4646 | 0.000999 |
| livertumor | RTMIFN | C3 | ITGAX | 0.6584 | 0.2809 | 0.4697 | 0.000999 |
| livertumor | RTMIFN | C3 | ITGB2 | 0.6584 | 1.4559 | 1.0572 | 0.000999 |
| livertumor | RTMIFN | C3 | LRP1 | 0.6584 | 0.5528 | 0.6056 | 0.000999 |
| livertumor | RTMIFN | CALM1 | TNFRSF1B | 0.6596 | 0.3537 | 0.5066 | 0.000999 |
| livertumor | RTMIFN | CALR | CD40 | 1.0416 | 0.4077 | 0.7246 | 0.000999 |
| livertumor | RTMIFN | CALR | CD44 | 1.0416 | 1.2543 | 1.148 | 0.000999 |
| livertumor | RTMIFN | CALR | LRP1 | 1.0416 | 0.5528 | 0.7972 | 0.000999 |
| livertumor | RTMIFN | CCL20 | C3AR1 | 1.0877 | 0.648 | 0.8679 | 0.000999 |

| | | | | | | | |
|---|---|---|---|---|---|---|---|
| livertumor | RTMIFN | CCL20 | C5AR1 | 1.0877 | 0.5634 | 0.8255 | 0.000999 |
| livertumor | RTMIFN | CCL20 | CXCR4 | 1.0877 | 0.9336 | 1.0106 | 0.000999 |
| livertumor | RTMIFN | CCL20 | FPR1 | 1.0877 | 0.4203 | 0.754 | 0.000999 |
| livertumor | RTMIFN | CFH | ITGAM | 0.3373 | 0.2708 | 0.304 | 0.000999 |
| livertumor | RTMIFN | CXCL2 | C3AR1 | 0.3364 | 0.648 | 0.4922 | 0.000999 |
| livertumor | RTMIFN | CXCL2 | C5AR1 | 0.3364 | 0.5634 | 0.4499 | 0.000999 |
| livertumor | RTMIFN | CXCL2 | CXCR4 | 0.3364 | 0.9336 | 0.635 | 0.000999 |
| livertumor | RTMIFN | CXCL2 | FPR1 | 0.3364 | 0.4203 | 0.3783 | 0.000999 |
| livertumor | RTMIFN | CXCL3 | C3AR1 | 0.2543 | 0.648 | 0.4512 | 0.000999 |
| livertumor | RTMIFN | CXCL3 | C5AR1 | 0.2543 | 0.5634 | 0.4088 | 0.000999 |
| livertumor | RTMIFN | CXCL3 | CXCR4 | 0.2543 | 0.9336 | 0.5939 | 0.000999 |
| livertumor | RTMIFN | CXCL3 | FPR1 | 0.2543 | 0.4203 | 0.3373 | 0.000999 |
| livertumor | RTMIFN | DSG2 | CD44 | 0.7479 | 1.2543 | 1.0011 | 0.000999 |
| livertumor | RTMIFN | DSG2 | CXCR4 | 0.7479 | 0.9336 | 0.8408 | 0.000999 |
| livertumor | RTMIFN | DSG2 | DAB2 | 0.7479 | 0.5995 | 0.6737 | 0.000999 |
| livertumor | RTMIFN | FGA | ITGAX | 0.2688 | 0.2809 | 0.2748 | 0.000999 |
| livertumor | RTMIFN | FGA | ITGB2 | 0.2688 | 1.4559 | 0.8623 | 0.000999 |
| livertumor | RTMIFN | GSTP1 | CD44 | 1.7219 | 1.2543 | 1.4881 | 0.000999 |
| livertumor | RTMIFN | HMGB1 | AGTRAP | 1.4361 | 0.38 | 0.908 | 0.000999 |
| livertumor | RTMIFN | HMGB1 | LY96 | 1.4361 | 1.1487 | 1.2924 | 0.000999 |
| livertumor | RTMIFN | HMGB1 | TLR2 | 1.4361 | 0.3529 | 0.8945 | 0.000999 |
| livertumor | RTMIFN | HSP90B1 | LRP1 | 0.8951 | 0.5528 | 0.724 | 0.000999 |
| livertumor | RTMIFN | HSP90B1 | TNFRSF1B | 0.8951 | 0.3537 | 0.6244 | 0.000999 |
| livertumor | RTMIFN | HSPG2 | LRP1 | 0.2746 | 0.5528 | 0.4137 | 0.000999 |
| livertumor | RTMIFN | ITGB1 | DAB2 | 1.6044 | 0.5995 | 1.102 | 0.000999 |
| livertumor | RTMIFN | ITGB1 | ENG | 1.6044 | 0.3504 | 0.9774 | 0.000999 |
| livertumor | RTMIFN | ITGB1 | ITGAX | 1.6044 | 0.2809 | 0.9427 | 0.000999 |
| livertumor | RTMIFN | ITGB1 | PLAUR | 1.6044 | 1.1076 | 1.356 | 0.000999 |
| livertumor | RTMIFN | LGALS3 | CLEC7A | 0.8172 | 0.6166 | 0.7169 | 0.000999 |
| livertumor | RTMIFN | LGALS3 | FCGR2A | 0.8172 | 1.0157 | 0.9165 | 0.000999 |

| | | | | | | | |
|---|---|---|---|---|---|---|---|
| livertumor | RTMIFN | LGALS3 | FCGR2B | 0.8172 | 0.7852 | 0.8012 | 0.000999 |
| livertumor | RTMIFN | LRPAP1 | LRP1 | 0.273 | 0.5528 | 0.4129 | 0.000999 |
| livertumor | RTMIFN | MDK | LRP1 | 1.6101 | 0.5528 | 1.0814 | 0.000999 |
| livertumor | RTMIFN | MIF | CD74 | 1.7638 | 4.5904 | 3.1771 | 0.000999 |
| livertumor | RTMIFN | MIF | TNFRSF14 | 1.7638 | 0.4163 | 1.09 | 0.000999 |
| livertumor | RTMIFN | NTS | GPR65 | 0.3425 | 0.3227 | 0.3326 | 0.000999 |
| livertumor | RTMIFN | NTS | LPAR6 | 0.3425 | 0.4247 | 0.3836 | 0.000999 |
| livertumor | RTMIFN | NTS | PTAFR | 0.3425 | 0.2739 | 0.3082 | 0.000999 |
| livertumor | RTMIFN | RPS19 | C5AR1 | 4.0772 | 0.5634 | 2.3203 | 0.000999 |
| livertumor | RTMIFN | RPS19 | CD4 | 4.0772 | 0.707 | 2.3921 | 0.000999 |
| livertumor | RTMIFN | RPS19 | ICAM1 | 4.0772 | 0.5523 | 2.3147 | 0.000999 |
| livertumor | RTMIFN | RTN4 | CD44 | 1.0046 | 1.2543 | 1.1294 | 0.000999 |
| livertumor | RTMIFN | SEMA3C | NRP2 | 0.2576 | 0.3534 | 0.3055 | 0.000999 |
| livertumor | RTMIFN | SERPINA1 | LRP1 | 1.604 | 0.5528 | 1.0784 | 0.000999 |
| livertumor | RTMIFN | SERPING1 | LRP1 | 0.3498 | 0.5528 | 0.4513 | 0.000999 |
| livertumor | RTMIFN | TF | CD4 | 0.523 | 0.707 | 0.615 | 0.000999 |
| livertumor | RTMIFN | TF | TFRC | 0.523 | 0.3853 | 0.4541 | 0.000999 |
| livertumor | RTMIFN | TFPI | LRP1 | 0.5043 | 0.5528 | 0.5285 | 0.000999 |
| livertumor | RTMIFN | TIMP1 | LRP1 | 1.0257 | 0.5528 | 0.7892 | 0.000999 |
| livertumor | RTMIFN | VEGFA | NRP2 | 0.5485 | 0.3534 | 0.4509 | 0.000999 |
| livertumor | RTMIFN | VTN | ITGB5 | 0.7399 | 0.2827 | 0.5113 | 0.000999 |
| livertumor | RTMIFN | VTN | PLAUR | 0.7399 | 1.1076 | 0.9238 | 0.000999 |
| livertumor | RTMIFN | ANXA1 | CD44 | 0.3041 | 1.2543 | 0.7792 | 0.002 |
| livertumor | RTMIFN | APP | TNFRSF14 | 0.5626 | 0.4163 | 0.4894 | 0.002 |
| livertumor | RTMIFN | PTPRF | TGFBR2 | 0.384 | 0.2932 | 0.3386 | 0.002 |
| livertumor | RTMIFN | OCLN | TGFBR2 | 0.3091 | 0.2932 | 0.3011 | 0.00699 |
| livertumor | RTMIFN | APP | TGFBR2 | 0.5626 | 0.2932 | 0.4279 | 0.00799 |
| livertumor | RTMIFN | ITGB1 | CD47 | 1.6044 | 0.468 | 1.0362 | 0.00799 |
| lungtumor | RTMIFN | ANXA1 | C3AR1 | 1.4026 | 0.648 | 1.0253 | 0.000999 |
| lungtumor | RTMIFN | ANXA1 | C5AR1 | 1.4026 | 0.5634 | 0.983 | 0.000999 |

| | | | | | | | |
|---|---|---|---|---|---|---|---|
| lungtumor | RTMIFN | ANXA1 | CD4 | 1.4026 | 0.707 | 1.0548 | 0.000999 |
| lungtumor | RTMIFN | ANXA1 | CD44 | 1.4026 | 1.2543 | 1.3285 | 0.000999 |
| lungtumor | RTMIFN | ANXA1 | CXCR4 | 1.4026 | 0.9336 | 1.1681 | 0.000999 |
| lungtumor | RTMIFN | ANXA1 | FPR1 | 1.4026 | 0.4203 | 0.9115 | 0.000999 |
| lungtumor | RTMIFN | ANXA1 | FPR3 | 1.4026 | 0.4975 | 0.9501 | 0.000999 |
| lungtumor | RTMIFN | AREG | ICAM1 | 0.3537 | 0.5523 | 0.453 | 0.000999 |
| lungtumor | RTMIFN | ARF1 | CD4 | 0.8791 | 0.707 | 0.7931 | 0.000999 |
| lungtumor | RTMIFN | ARF1 | CD44 | 0.8791 | 1.2543 | 1.0667 | 0.000999 |
| lungtumor | RTMIFN | CALM1 | RPSA | 0.8426 | 2.1961 | 1.5193 | 0.000999 |
| lungtumor | RTMIFN | CALM1 | TNFRSF1A | 0.8426 | 0.4908 | 0.6667 | 0.000999 |
| lungtumor | RTMIFN | CALM1 | TNFRSF1B | 0.8426 | 0.3537 | 0.5981 | 0.000999 |
| lungtumor | RTMIFN | CALR | CD40 | 0.7795 | 0.4077 | 0.5936 | 0.000999 |
| lungtumor | RTMIFN | CALR | CD44 | 0.7795 | 1.2543 | 1.0169 | 0.000999 |
| lungtumor | RTMIFN | CALR | LRP1 | 0.7795 | 0.5528 | 0.6662 | 0.000999 |
| lungtumor | RTMIFN | CCL20 | C3AR1 | 0.2885 | 0.648 | 0.4683 | 0.000999 |
| lungtumor | RTMIFN | CCL20 | C5AR1 | 0.2885 | 0.5634 | 0.426 | 0.000999 |
| lungtumor | RTMIFN | CCL20 | CXCR4 | 0.2885 | 0.9336 | 0.6111 | 0.000999 |
| lungtumor | RTMIFN | CCL20 | FPR1 | 0.2885 | 0.4203 | 0.3544 | 0.000999 |
| lungtumor | RTMIFN | CXCL10 | C3AR1 | 0.3417 | 0.648 | 0.4949 | 0.000999 |
| lungtumor | RTMIFN | CXCL10 | C5AR1 | 0.3417 | 0.5634 | 0.4525 | 0.000999 |
| lungtumor | RTMIFN | CXCL10 | CXCR4 | 0.3417 | 0.9336 | 0.6376 | 0.000999 |
| lungtumor | RTMIFN | CXCL10 | FPR1 | 0.3417 | 0.4203 | 0.381 | 0.000999 |
| lungtumor | RTMIFN | FGA | ITGAX | 0.4445 | 0.2809 | 0.3627 | 0.000999 |
| lungtumor | RTMIFN | FGA | ITGB2 | 0.4445 | 1.4559 | 0.9502 | 0.000999 |
| lungtumor | RTMIFN | GRN | TNFRSF1A | 0.4026 | 0.4908 | 0.4467 | 0.000999 |
| lungtumor | RTMIFN | GRN | TNFRSF1B | 0.4026 | 0.3537 | 0.3781 | 0.000999 |
| lungtumor | RTMIFN | GSTP1 | CD44 | 2.4093 | 1.2543 | 1.8318 | 0.000999 |
| lungtumor | RTMIFN | HLA-DRA | CD4 | 0.9609 | 0.707 | 0.834 | 0.000999 |
| lungtumor | RTMIFN | HLA-DRA | PTPRC | 0.9609 | 0.6543 | 0.8076 | 0.000999 |
| lungtumor | RTMIFN | HMGB1 | LY96 | 1.1943 | 1.1487 | 1.1715 | 0.000999 |

| | | | | | | | |
|---|---|---|---|---|---|---|---|
| lungtumor | RTMIFN | HMGB1 | TLR2 | 1.1943 | 0.3529 | 0.7736 | 0.000999 |
| lungtumor | RTMIFN | HMGB2 | CD44 | 0.5593 | 1.2543 | 0.9068 | 0.000999 |
| lungtumor | RTMIFN | HMGB2 | DAB2 | 0.5593 | 0.5995 | 0.5794 | 0.000999 |
| lungtumor | RTMIFN | HSP90B1 | LRP1 | 0.6141 | 0.5528 | 0.5834 | 0.000999 |
| lungtumor | RTMIFN | HSP90B1 | TNFRSF1B | 0.6141 | 0.3537 | 0.4839 | 0.000999 |
| lungtumor | RTMIFN | LGALS3 | CLEC7A | 0.9915 | 0.6166 | 0.8041 | 0.000999 |
| lungtumor | RTMIFN | LGALS3 | FCGR2A | 0.9915 | 1.0157 | 1.0036 | 0.000999 |
| lungtumor | RTMIFN | LGALS3 | FCGR2B | 0.9915 | 0.7852 | 0.8884 | 0.000999 |
| lungtumor | RTMIFN | LGALS3 | TFRC | 0.9915 | 0.3853 | 0.6884 | 0.000999 |
| lungtumor | RTMIFN | LGALS3 | TGFBR2 | 0.9915 | 0.2932 | 0.6423 | 0.000999 |
| lungtumor | RTMIFN | MDK | LRP1 | 1.2896 | 0.5528 | 0.9212 | 0.000999 |
| lungtumor | RTMIFN | MIF | CD74 | 1.4585 | 4.5904 | 3.0245 | 0.000999 |
| lungtumor | RTMIFN | MIF | TNFRSF14 | 1.4585 | 0.4163 | 0.9374 | 0.000999 |
| lungtumor | RTMIFN | PSAP | LRP1 | 0.5457 | 0.5528 | 0.5492 | 0.000999 |
| lungtumor | RTMIFN | RPS19 | C5AR1 | 3.5104 | 0.5634 | 2.0369 | 0.000999 |
| lungtumor | RTMIFN | RPS19 | CD4 | 3.5104 | 0.707 | 2.1087 | 0.000999 |
| lungtumor | RTMIFN | RPS19 | ICAM1 | 3.5104 | 0.5523 | 2.0313 | 0.000999 |
| lungtumor | RTMIFN | RPS19 | RPSA | 3.5104 | 2.1961 | 2.8533 | 0.000999 |
| lungtumor | RTMIFN | RTN4 | CD44 | 0.4013 | 1.2543 | 0.8278 | 0.000999 |
| lungtumor | RTMIFN | SAA1 | C3AR1 | 0.2539 | 0.648 | 0.451 | 0.000999 |
| lungtumor | RTMIFN | SAA1 | C5AR1 | 0.2539 | 0.5634 | 0.4086 | 0.000999 |
| lungtumor | RTMIFN | SAA1 | CXCR4 | 0.2539 | 0.9336 | 0.5937 | 0.000999 |
| lungtumor | RTMIFN | SAA1 | FPR1 | 0.2539 | 0.4203 | 0.3371 | 0.000999 |
| lungtumor | RTMIFN | SAA1 | GPR65 | 0.2539 | 0.3227 | 0.2883 | 0.000999 |
| lungtumor | RTMIFN | SAA1 | LPAR6 | 0.2539 | 0.4247 | 0.3393 | 0.000999 |
| lungtumor | RTMIFN | SAA1 | PTAFR | 0.2539 | 0.2739 | 0.2639 | 0.000999 |
| lungtumor | RTMIFN | SERPINA1 | LRP1 | 0.3624 | 0.5528 | 0.4576 | 0.000999 |
| lungtumor | RTMIFN | SPP1 | CD44 | 1.4874 | 1.2543 | 1.3709 | 0.000999 |
| lungtumor | RTMIFN | SPP1 | ITGB5 | 1.4874 | 0.2827 | 0.8851 | 0.000999 |
| lungtumor | RTMIFN | TFPI | LRP1 | 0.3615 | 0.5528 | 0.4571 | 0.000999 |

| | | | | | | | |
|---|---|---|---|---|---|---|---|
| lungtumor | RTMIFN | TIMP1 | LRP1 | 0.6195 | 0.5528 | 0.5861 | 0.000999 |
| lungtumor | RTMIFN | FGA | ITGB1 | 0.4445 | 0.528 | 0.4862 | 0.002 |
| lungtumor | RTMIFN | HSP90B1 | ITGB1 | 0.6141 | 0.528 | 0.571 | 0.002 |
| lungtumor | RTMIFN | LGALS3 | ITGB1 | 0.9915 | 0.528 | 0.7597 | 0.002 |
| lungtumor | RTMIFN | SAA1 | SELENOS | 0.2539 | 0.628 | 0.441 | 0.002 |
| lungtumor | RTMIFN | SPP1 | ITGB1 | 1.4874 | 0.528 | 1.0077 | 0.002 |
| melanomatumo | RTMIFN | ARF1 | CD4 | 1.8545 | 0.707 | 1.2808 | 0.000999 |
| melanomatumo | RTMIFN | ARF1 | CD44 | 1.8545 | 1.2543 | 1.5544 | 0.000999 |
| melanomatumo | RTMIFN | CD320 | TGFBR2 | 0.2588 | 0.2932 | 0.276 | 0.000999 |
| melanomatumo | RTMIFN | HEBP1 | C3AR1 | 0.3689 | 0.648 | 0.5084 | 0.000999 |
| melanomatumo | RTMIFN | HEBP1 | C5AR1 | 0.3689 | 0.5634 | 0.4661 | 0.000999 |
| melanomatumo | RTMIFN | HEBP1 | CXCR4 | 0.3689 | 0.9336 | 0.6512 | 0.000999 |
| melanomatumo | RTMIFN | HMGB1 | LY96 | 1.6757 | 1.1487 | 1.4122 | 0.000999 |
| melanomatumo | RTMIFN | HMGB1 | TLR2 | 1.6757 | 0.3529 | 1.0143 | 0.000999 |
| melanomatumo | RTMIFN | MIF | CD74 | 2.6939 | 4.5904 | 3.6422 | 0.000999 |
| melanomatumo | RTMIFN | MIF | TNFRSF14 | 2.6939 | 0.4163 | 1.5551 | 0.000999 |
| melanomatumo | RTMIFN | SERPINE2 | LRP1 | 0.9323 | 0.5528 | 0.7425 | 0.000999 |
| melanomatumo | RTMIFN | GSTP1 | CD44 | 2.3341 | 1.2543 | 1.7942 | 0.002 |
| melanomatumo | RTMIFN | HMGB1 | AGTRAP | 1.6757 | 0.38 | 1.0279 | 0.002 |
| melanomatumo | RTMIFN | HEBP1 | FPR3 | 0.3689 | 0.4975 | 0.4332 | 0.003 |
| melanomatumo | RTMIFN | HAS2 | CD44 | 0.3078 | 1.2543 | 0.781 | 0.00799 |
| melanomatumo | RTMIFN | HEBP1 | FPR1 | 0.3689 | 0.4203 | 0.3946 | 0.00799 |
| melanomatumo | RTMIFN | LRPAP1 | LRP1 | 0.6123 | 0.5528 | 0.5826 | 0.00799 |
| ovarytumor | RTMIFN | ANXA1 | C3AR1 | 1.0314 | 0.648 | 0.8397 | 0.000999 |
| ovarytumor | RTMIFN | ANXA1 | C5AR1 | 1.0314 | 0.5634 | 0.7974 | 0.000999 |
| ovarytumor | RTMIFN | ANXA1 | CD4 | 1.0314 | 0.707 | 0.8692 | 0.000999 |
| ovarytumor | RTMIFN | ANXA1 | CD44 | 1.0314 | 1.2543 | 1.1428 | 0.000999 |
| ovarytumor | RTMIFN | ANXA1 | CXCR4 | 1.0314 | 0.9336 | 0.9825 | 0.000999 |
| ovarytumor | RTMIFN | ANXA1 | FPR1 | 1.0314 | 0.4203 | 0.7258 | 0.000999 |
| ovarytumor | RTMIFN | ANXA1 | FPR3 | 1.0314 | 0.4975 | 0.7645 | 0.000999 |

| | | | | | | | |
|---|---|---|---|---|---|---|---|
| ovarytumor | RTMIFN | APOE | LRP1 | 0.837 | 0.5528 | 0.6949 | 0.000999 |
| ovarytumor | RTMIFN | APP | ALOX5 | 0.2907 | 0.4066 | 0.3487 | 0.000999 |
| ovarytumor | RTMIFN | APP | C3AR1 | 0.2907 | 0.648 | 0.4694 | 0.000999 |
| ovarytumor | RTMIFN | APP | C5AR1 | 0.2907 | 0.5634 | 0.427 | 0.000999 |
| ovarytumor | RTMIFN | APP | CXCR4 | 0.2907 | 0.9336 | 0.6122 | 0.000999 |
| ovarytumor | RTMIFN | APP | FPR1 | 0.2907 | 0.4203 | 0.3555 | 0.000999 |
| ovarytumor | RTMIFN | APP | LRP1 | 0.2907 | 0.5528 | 0.4217 | 0.000999 |
| ovarytumor | RTMIFN | APP | TGFBR2 | 0.2907 | 0.2932 | 0.2919 | 0.000999 |
| ovarytumor | RTMIFN | APP | TNFRSF14 | 0.2907 | 0.4163 | 0.3535 | 0.000999 |
| ovarytumor | RTMIFN | ARF1 | CD4 | 0.7364 | 0.707 | 0.7217 | 0.000999 |
| ovarytumor | RTMIFN | ARF1 | CD44 | 0.7364 | 1.2543 | 0.9954 | 0.000999 |
| ovarytumor | RTMIFN | CALM1 | TNFRSF1A | 1.0087 | 0.4908 | 0.7497 | 0.000999 |
| ovarytumor | RTMIFN | CALM1 | TNFRSF1B | 1.0087 | 0.3537 | 0.6812 | 0.000999 |
| ovarytumor | RTMIFN | CALR | CD40 | 0.7423 | 0.4077 | 0.575 | 0.000999 |
| ovarytumor | RTMIFN | CALR | CD44 | 0.7423 | 1.2543 | 0.9983 | 0.000999 |
| ovarytumor | RTMIFN | CALR | LRP1 | 0.7423 | 0.5528 | 0.6475 | 0.000999 |
| ovarytumor | RTMIFN | COL1A1 | CD44 | 0.3504 | 1.2543 | 0.8023 | 0.000999 |
| ovarytumor | RTMIFN | GRN | TNFRSF1A | 0.3896 | 0.4908 | 0.4402 | 0.000999 |
| ovarytumor | RTMIFN | GRN | TNFRSF1B | 0.3896 | 0.3537 | 0.3716 | 0.000999 |
| ovarytumor | RTMIFN | GSTP1 | CD44 | 2.6185 | 1.2543 | 1.9364 | 0.000999 |
| ovarytumor | RTMIFN | HLA-DMA | CD4 | 0.2537 | 0.707 | 0.4804 | 0.000999 |
| ovarytumor | RTMIFN | HLA-DRA | CD4 | 1.2369 | 0.707 | 0.9719 | 0.000999 |
| ovarytumor | RTMIFN | HLA-DRA | PTPRC | 1.2369 | 0.6543 | 0.9456 | 0.000999 |
| ovarytumor | RTMIFN | HMGB1 | AGTRAP | 1.7272 | 0.38 | 1.0536 | 0.000999 |
| ovarytumor | RTMIFN | HMGB1 | LY96 | 1.7272 | 1.1487 | 1.438 | 0.000999 |
| ovarytumor | RTMIFN | HMGB1 | TLR2 | 1.7272 | 0.3529 | 1.0401 | 0.000999 |
| ovarytumor | RTMIFN | HMGB2 | CD44 | 0.4974 | 1.2543 | 0.8759 | 0.000999 |
| ovarytumor | RTMIFN | HMGB2 | DAB2 | 0.4974 | 0.5995 | 0.5485 | 0.000999 |
| ovarytumor | RTMIFN | HSP90B1 | LRP1 | 0.6365 | 0.5528 | 0.5946 | 0.000999 |
| ovarytumor | RTMIFN | HSP90B1 | TNFRSF1B | 0.6365 | 0.3537 | 0.4951 | 0.000999 |

| | | | | | | | |
|---|---|---|---|---|---|---|---|
| ovarytumor | RTMIFN | ITGB1 | DAB2 | 0.2783 | 0.5995 | 0.4389 | 0.000999 |
| ovarytumor | RTMIFN | ITGB1 | ENG | 0.2783 | 0.3504 | 0.3144 | 0.000999 |
| ovarytumor | RTMIFN | ITGB1 | ITGAX | 0.2783 | 0.2809 | 0.2796 | 0.000999 |
| ovarytumor | RTMIFN | ITGB1 | PLAUR | 0.2783 | 1.1076 | 0.6929 | 0.000999 |
| ovarytumor | RTMIFN | LGALS3 | CLEC7A | 0.5097 | 0.6166 | 0.5632 | 0.000999 |
| ovarytumor | RTMIFN | LGALS3 | FCGR2A | 0.5097 | 1.0157 | 0.7627 | 0.000999 |
| ovarytumor | RTMIFN | LGALS3 | FCGR2B | 0.5097 | 0.7852 | 0.6475 | 0.000999 |
| ovarytumor | RTMIFN | LGALS3 | TFRC | 0.5097 | 0.3853 | 0.4475 | 0.000999 |
| ovarytumor | RTMIFN | LGALS3 | TGFBR2 | 0.5097 | 0.2932 | 0.4015 | 0.000999 |
| ovarytumor | RTMIFN | LRPAP1 | LRP1 | 0.3826 | 0.5528 | 0.4677 | 0.000999 |
| ovarytumor | RTMIFN | MDK | LRP1 | 1.5321 | 0.5528 | 1.0424 | 0.000999 |
| ovarytumor | RTMIFN | MIF | CD74 | 0.8565 | 4.5904 | 2.7234 | 0.000999 |
| ovarytumor | RTMIFN | MIF | TNFRSF14 | 0.8565 | 0.4163 | 0.6364 | 0.000999 |
| ovarytumor | RTMIFN | PSAP | LRP1 | 0.6574 | 0.5528 | 0.6051 | 0.000999 |
| ovarytumor | RTMIFN | RPS19 | C5AR1 | 3.9325 | 0.5634 | 2.2479 | 0.000999 |
| ovarytumor | RTMIFN | RPS19 | CD4 | 3.9325 | 0.707 | 2.3198 | 0.000999 |
| ovarytumor | RTMIFN | RPS19 | ICAM1 | 3.9325 | 0.5523 | 2.2424 | 0.000999 |
| ovarytumor | RTMIFN | RTN4 | CD44 | 0.4992 | 1.2543 | 0.8767 | 0.000999 |
| ovarytumor | RTMIFN | RTN4 | CD99 | 0.4992 | 0.6857 | 0.5925 | 0.000999 |
| ovarytumor | RTMIFN | SERPING1 | LRP1 | 0.3678 | 0.5528 | 0.4603 | 0.000999 |
| ovarytumor | RTMIFN | SPP1 | CD44 | 0.3946 | 1.2543 | 0.8245 | 0.000999 |
| ovarytumor | RTMIFN | SPP1 | ITGB5 | 0.3946 | 0.2827 | 0.3387 | 0.000999 |
| ovarytumor | RTMIFN | TIMP1 | LRP1 | 0.7779 | 0.5528 | 0.6653 | 0.000999 |
| ovarytumor | RTMIFN | COL1A1 | ITGB1 | 0.3504 | 0.528 | 0.4392 | 0.003 |
| ovarytumor | RTMIFN | HSP90B1 | ITGB1 | 0.6365 | 0.528 | 0.5822 | 0.003 |
| ovarytumor | RTMIFN | ITGB1 | CD46 | 0.2783 | 0.2704 | 0.2743 | 0.003 |
| ovarytumor | RTMIFN | LGALS3 | ITGB1 | 0.5097 | 0.528 | 0.5188 | 0.004 |
| ovarytumor | RTMIFN | SPP1 | ITGB1 | 0.3946 | 0.528 | 0.4613 | 0.005 |

| **Supplementary Table 8: Communication results between macrophages and lymphocytes subpopulations, and malignant cells** | | | | | | | |
|---|---|---|---|---|---|---|---|
| **RTM_IM vs. lymphocytes** | | | | | | | |
| celltype_01 | celltype_02 | ligand | receptor | ligand_mean_expression | receptor_mean_expression | ligand_receptor_mean_expression | pvalue |
| RTMIM | NKT | A2M | TNFRSF14 | 1.094 | 0.2848 | 0.6894 | 0.000999 |
| RTMIM | NKT | ADM | PTGER2 | 0.2706 | 0.3627 | 0.3167 | 0.000999 |
| RTMIM | NKT | ANXA1 | CXCR4 | 1.1316 | 2.0373 | 1.5844 | 0.000999 |
| RTMIM | NKT | APOE | SORL1 | 1.1762 | 0.4137 | 0.7949 | 0.000999 |
| RTMIM | NKT | C3 | CXCR4 | 0.3607 | 2.0373 | 1.199 | 0.000999 |
| RTMIM | NKT | C3 | ITGB2 | 0.3607 | 1.6481 | 1.0044 | 0.000999 |
| RTMIM | NKT | COL1A1 | CD44 | 0.4864 | 0.6724 | 0.5794 | 0.000999 |
| RTMIM | NKT | COL1A1 | ITGB1 | 0.4864 | 0.488 | 0.4872 | 0.000999 |
| RTMIM | NKT | CXCL12 | CXCR4 | 0.3733 | 2.0373 | 1.2053 | 0.000999 |
| RTMIM | NKT | CXCL12 | S1PR4 | 0.3733 | 0.4435 | 0.4084 | 0.000999 |
| RTMIM | NKT | CXCL12 | S1PR5 | 0.3733 | 0.508 | 0.4407 | 0.000999 |
| RTMIM | NKT | CXCL16 | CXCR4 | 0.7537 | 2.0373 | 1.3955 | 0.000999 |
| RTMIM | NKT | CXCL16 | S1PR4 | 0.7537 | 0.4435 | 0.5986 | 0.000999 |
| RTMIM | NKT | CXCL16 | S1PR5 | 0.7537 | 0.508 | 0.6308 | 0.000999 |
| RTMIM | NKT | CXCL2 | CXCR4 | 1.4022 | 2.0373 | 1.7197 | 0.000999 |
| RTMIM | NKT | CXCL2 | S1PR4 | 1.4022 | 0.4435 | 0.9229 | 0.000999 |
| RTMIM | NKT | CXCL2 | S1PR5 | 1.4022 | 0.508 | 0.9551 | 0.000999 |
| RTMIM | NKT | CXCL3 | CXCR4 | 1.1602 | 2.0373 | 1.5988 | 0.000999 |
| RTMIM | NKT | CXCL3 | S1PR4 | 1.1602 | 0.4435 | 0.8019 | 0.000999 |
| RTMIM | NKT | CXCL3 | S1PR5 | 1.1602 | 0.508 | 0.8341 | 0.000999 |
| RTMIM | NKT | FN1 | CD44 | 0.3872 | 0.6724 | 0.5298 | 0.000999 |
| RTMIM | NKT | FN1 | ITGA4 | 0.3872 | 0.2946 | 0.3409 | 0.000999 |
| RTMIM | NKT | FN1 | ITGB1 | 0.3872 | 0.488 | 0.4376 | 0.000999 |
| RTMIM | NKT | FN1 | ITGB7 | 0.3872 | 0.4721 | 0.4297 | 0.000999 |
| RTMIM | NKT | FN1 | RPSA | 0.3872 | 2.3639 | 1.3755 | 0.000999 |
| RTMIM | NKT | GRN | TNFRSF1B | 1.4409 | 0.4949 | 0.9679 | 0.000999 |
| RTMIM | NKT | GSTP1 | CD44 | 1.3126 | 0.6724 | 0.9925 | 0.000999 |

| | | | | | | | |
|---|---|---|---|---|---|---|---|
| RTMIM | NKT | GSTP1 | IL7R | 1.3126 | 0.5933 | 0.953 | 0.000999 |
| RTMIM | NKT | GSTP1 | ITGA4 | 1.3126 | 0.2946 | 0.8036 | 0.000999 |
| RTMIM | NKT | HBEGF | CD44 | 0.6117 | 0.6724 | 0.6421 | 0.000999 |
| RTMIM | NKT | HLA-DRA | PTPRC | 3.9393 | 1.8797 | 2.9095 | 0.000999 |
| RTMIM | NKT | HSP90B1 | ITGB1 | 0.7626 | 0.488 | 0.6253 | 0.000999 |
| RTMIM | NKT | HSP90B1 | TNFRSF1B | 0.7626 | 0.4949 | 0.6288 | 0.000999 |
| RTMIM | NKT | ICAM1 | IL2RG | 0.6002 | 1.1524 | 0.8763 | 0.000999 |
| RTMIM | NKT | ICAM1 | ITGAL | 0.6002 | 0.6778 | 0.639 | 0.000999 |
| RTMIM | NKT | ICAM1 | ITGB2 | 0.6002 | 1.6481 | 1.1242 | 0.000999 |
| RTMIM | NKT | ICAM1 | SPN | 0.6002 | 0.6398 | 0.62 | 0.000999 |
| RTMIM | NKT | IGF1 | RACK1 | 0.4603 | 2.4327 | 1.4465 | 0.000999 |
| RTMIM | NKT | IL10 | IL10RA | 0.2565 | 0.5156 | 0.3861 | 0.000999 |
| RTMIM | NKT | LGALS3 | CD6 | 0.7 | 0.4881 | 0.5941 | 0.000999 |
| RTMIM | NKT | LGALS3 | ITGB1 | 0.7 | 0.488 | 0.594 | 0.000999 |
| RTMIM | NKT | LGALS3 | SPN | 0.7 | 0.6398 | 0.6699 | 0.000999 |
| RTMIM | NKT | LRPAP1 | SORL1 | 0.3277 | 0.4137 | 0.3707 | 0.000999 |
| RTMIM | NKT | NAMPT | ANXA6 | 1.2185 | 0.844 | 1.0312 | 0.000999 |
| RTMIM | NKT | PECAM1 | CXCR4 | 0.334 | 2.0373 | 1.1856 | 0.000999 |
| RTMIM | NKT | PLAU | CD44 | 0.3288 | 0.6724 | 0.5006 | 0.000999 |
| RTMIM | NKT | RTN4 | CD44 | 0.9278 | 0.6724 | 0.8001 | 0.000999 |
| RTMIM | NKT | RTN4 | CD99 | 0.9278 | 1.4547 | 1.1912 | 0.000999 |
| RTMIM | NKT | SPP1 | CD44 | 0.61 | 0.6724 | 0.6412 | 0.000999 |
| RTMIM | NKT | SPP1 | ITGA4 | 0.61 | 0.2946 | 0.4523 | 0.000999 |
| RTMIM | NKT | SPP1 | ITGB1 | 0.61 | 0.488 | 0.549 | 0.000999 |
| RTMIM | NKT | THBS1 | CD47 | 0.3964 | 0.5576 | 0.477 | 0.000999 |
| RTMIM | NKT | THBS1 | ITGA4 | 0.3964 | 0.2946 | 0.3455 | 0.000999 |
| RTMIM | NKT | THBS1 | ITGB1 | 0.3964 | 0.488 | 0.4422 | 0.000999 |
| RTMIM | NKT | TNF | CALM1 | 0.3679 | 1.9335 | 1.1507 | 0.000999 |
| RTMIM | NKT | TNF | RACK1 | 0.3679 | 2.4327 | 1.4003 | 0.000999 |
| RTMIM | NKT | TNF | RASA3 | 0.3679 | 0.2983 | 0.3331 | 0.000999 |

| | | | | | | | |
|---|---|---|---|---|---|---|---|
| RTMIM | NKT | TNF | TNFRSF1B | 0.3679 | 0.4949 | 0.4314 | 0.000999 |
| RTMIM | NKT | TNFSF13B | ITGB1 | 0.4492 | 0.488 | 0.4686 | 0.000999 |
| RTMIM | TCD4ex | A2M | TNFRSF14 | 1.094 | 0.2806 | 0.6873 | 0.000999 |
| RTMIM | TCD4ex | ADM | PTGER4 | 0.2706 | 0.4696 | 0.3701 | 0.000999 |
| RTMIM | TCD4ex | ANXA1 | CCR7 | 1.1316 | 0.4844 | 0.808 | 0.000999 |
| RTMIM | TCD4ex | ANXA1 | CD4 | 1.1316 | 0.5864 | 0.859 | 0.000999 |
| RTMIM | TCD4ex | ANXA1 | CD44 | 1.1316 | 1.0244 | 1.078 | 0.000999 |
| RTMIM | TCD4ex | ANXA1 | CXCR4 | 1.1316 | 2.4201 | 1.7758 | 0.000999 |
| RTMIM | TCD4ex | C3 | CCR7 | 0.3607 | 0.4844 | 0.4226 | 0.000999 |
| RTMIM | TCD4ex | C3 | CXCR4 | 0.3607 | 2.4201 | 1.3904 | 0.000999 |
| RTMIM | TCD4ex | C3 | ITGB2 | 0.3607 | 0.7871 | 0.5739 | 0.000999 |
| RTMIM | TCD4ex | CALR | IL7R | 0.7672 | 1.2353 | 1.0012 | 0.000999 |
| RTMIM | TCD4ex | CD86 | CD28 | 0.4042 | 0.3256 | 0.3649 | 0.000999 |
| RTMIM | TCD4ex | CD86 | CTLA4 | 0.4042 | 0.6294 | 0.5168 | 0.000999 |
| RTMIM | TCD4ex | COL1A1 | CD44 | 0.4864 | 1.0244 | 0.7554 | 0.000999 |
| RTMIM | TCD4ex | COL1A1 | ITGB1 | 0.4864 | 0.4615 | 0.4739 | 0.000999 |
| RTMIM | TCD4ex | CXCL12 | CCR7 | 0.3733 | 0.4844 | 0.4289 | 0.000999 |
| RTMIM | TCD4ex | CXCL12 | CD4 | 0.3733 | 0.5864 | 0.4799 | 0.000999 |
| RTMIM | TCD4ex | CXCL12 | CXCR4 | 0.3733 | 2.4201 | 1.3967 | 0.000999 |
| RTMIM | TCD4ex | CXCL16 | CCR7 | 0.7537 | 0.4844 | 0.619 | 0.000999 |
| RTMIM | TCD4ex | CXCL16 | CXCR4 | 0.7537 | 2.4201 | 1.5869 | 0.000999 |
| RTMIM | TCD4ex | CXCL2 | CCR7 | 1.4022 | 0.4844 | 0.9433 | 0.000999 |
| RTMIM | TCD4ex | CXCL2 | CXCR4 | 1.4022 | 2.4201 | 1.9111 | 0.000999 |
| RTMIM | TCD4ex | CXCL3 | CCR7 | 1.1602 | 0.4844 | 0.8223 | 0.000999 |
| RTMIM | TCD4ex | CXCL3 | CXCR4 | 1.1602 | 2.4201 | 1.7901 | 0.000999 |
| RTMIM | TCD4ex | FN1 | CD44 | 0.3872 | 1.0244 | 0.7058 | 0.000999 |
| RTMIM | TCD4ex | FN1 | ITGB1 | 0.3872 | 0.4615 | 0.4244 | 0.000999 |
| RTMIM | TCD4ex | FN1 | RPSA | 0.3872 | 2.4895 | 1.4384 | 0.000999 |
| RTMIM | TCD4ex | GRN | TNFRSF1B | 1.4409 | 0.383 | 0.912 | 0.000999 |
| RTMIM | TCD4ex | GSTP1 | CD44 | 1.3126 | 1.0244 | 1.1685 | 0.000999 |

| | | | | | | | |
|---|---|---|---|---|---|---|---|
| RTMIM | TCD4ex | GSTP1 | IL7R | 1.3126 | 1.2353 | 1.2739 | 0.000999 |
| RTMIM | TCD4ex | HBEGF | CD44 | 0.6117 | 1.0244 | 0.818 | 0.000999 |
| RTMIM | TCD4ex | HLA-DMA | CD4 | 1.7339 | 0.5864 | 1.1602 | 0.000999 |
| RTMIM | TCD4ex | HLA-DRA | CD4 | 3.9393 | 0.5864 | 2.2629 | 0.000999 |
| RTMIM | TCD4ex | HLA-DRA | PDCD1 | 3.9393 | 0.457 | 2.1981 | 0.000999 |
| RTMIM | TCD4ex | HLA-DRA | PTPRC | 3.9393 | 1.7493 | 2.8443 | 0.000999 |
| RTMIM | TCD4ex | HSP90B1 | ITGB1 | 0.7626 | 0.4615 | 0.6121 | 0.000999 |
| RTMIM | TCD4ex | HSP90B1 | TNFRSF1B | 0.7626 | 0.383 | 0.5728 | 0.000999 |
| RTMIM | TCD4ex | ICAM1 | IL2RG | 0.6002 | 0.9695 | 0.7848 | 0.000999 |
| RTMIM | TCD4ex | ICAM1 | ITGAL | 0.6002 | 0.2603 | 0.4303 | 0.000999 |
| RTMIM | TCD4ex | ICAM1 | ITGB2 | 0.6002 | 0.7871 | 0.6936 | 0.000999 |
| RTMIM | TCD4ex | IGF1 | RACK1 | 0.4603 | 2.4434 | 1.4519 | 0.000999 |
| RTMIM | TCD4ex | IL10 | CD28 | 0.2565 | 0.3256 | 0.2911 | 0.000999 |
| RTMIM | TCD4ex | ITGB2 | ITGAL | 0.862 | 0.2603 | 0.5611 | 0.000999 |
| RTMIM | TCD4ex | LGALS3 | CD6 | 0.7 | 0.5338 | 0.6169 | 0.000999 |
| RTMIM | TCD4ex | LGALS3 | ITGB1 | 0.7 | 0.4615 | 0.5808 | 0.000999 |
| RTMIM | TCD4ex | NAMPT | ANXA6 | 1.2185 | 0.5386 | 0.8785 | 0.000999 |
| RTMIM | TCD4ex | PECAM1 | CXCR4 | 0.334 | 2.4201 | 1.377 | 0.000999 |
| RTMIM | TCD4ex | PLAU | CD44 | 0.3288 | 1.0244 | 0.6766 | 0.000999 |
| RTMIM | TCD4ex | RTN4 | CD44 | 0.9278 | 1.0244 | 0.9761 | 0.000999 |
| RTMIM | TCD4ex | RTN4 | CD99 | 0.9278 | 0.9165 | 0.9221 | 0.000999 |
| RTMIM | TCD4ex | SPP1 | CD44 | 0.61 | 1.0244 | 0.8172 | 0.000999 |
| RTMIM | TCD4ex | SPP1 | ITGB1 | 0.61 | 0.4615 | 0.5357 | 0.000999 |
| RTMIM | TCD4ex | TGFB1 | ITGB1 | 0.4155 | 0.4615 | 0.4385 | 0.000999 |
| RTMIM | TCD4ex | THBS1 | CD47 | 0.3964 | 0.3456 | 0.371 | 0.000999 |
| RTMIM | TCD4ex | THBS1 | ITGB1 | 0.3964 | 0.4615 | 0.429 | 0.000999 |
| RTMIM | TCD4ex | TNF | CALM1 | 0.3679 | 1.2957 | 0.8318 | 0.000999 |
| RTMIM | TCD4ex | TNF | ICOS | 0.3679 | 0.532 | 0.4499 | 0.000999 |
| RTMIM | TCD4ex | TNF | RACK1 | 0.3679 | 2.4434 | 1.4056 | 0.000999 |
| RTMIM | TCD4ex | TNF | TNFRSF1B | 0.3679 | 0.383 | 0.3755 | 0.000999 |

| | | | | | | | |
|---|---|---|---|---|---|---|---|
| RTMIM | TCD4ex | TNFSF13B | ITGB1 | 0.4492 | 0.4615 | 0.4554 | 0.000999 |
| RTMIM | TCD4ex | CALR | CD44 | 0.7672 | 1.0244 | 0.8958 | 0.00999 |
| RTMIM | TCD4reg | A2M | TNFRSF14 | 1.094 | 0.3843 | 0.7392 | 0.000999 |
| RTMIM | TCD4reg | ANXA1 | CCR7 | 1.1316 | 0.3098 | 0.7207 | 0.000999 |
| RTMIM | TCD4reg | ANXA1 | CD4 | 1.1316 | 0.429 | 0.7803 | 0.000999 |
| RTMIM | TCD4reg | ANXA1 | CD44 | 1.1316 | 1.0896 | 1.1106 | 0.000999 |
| RTMIM | TCD4reg | ANXA1 | CXCR3 | 1.1316 | 0.3357 | 0.7336 | 0.000999 |
| RTMIM | TCD4reg | ANXA1 | CXCR4 | 1.1316 | 2.2962 | 1.7139 | 0.000999 |
| RTMIM | TCD4reg | ANXA1 | CXCR6 | 1.1316 | 0.3945 | 0.7631 | 0.000999 |
| RTMIM | TCD4reg | ANXA1 | FAS | 1.1316 | 0.2727 | 0.7022 | 0.000999 |
| RTMIM | TCD4reg | C3 | CCR7 | 0.3607 | 0.3098 | 0.3353 | 0.000999 |
| RTMIM | TCD4reg | C3 | CXCR3 | 0.3607 | 0.3357 | 0.3482 | 0.000999 |
| RTMIM | TCD4reg | C3 | CXCR4 | 0.3607 | 2.2962 | 1.3284 | 0.000999 |
| RTMIM | TCD4reg | C3 | CXCR6 | 0.3607 | 0.3945 | 0.3776 | 0.000999 |
| RTMIM | TCD4reg | C3 | ITGB2 | 0.3607 | 0.6238 | 0.4923 | 0.000999 |
| RTMIM | TCD4reg | CALR | CD44 | 0.7672 | 1.0896 | 0.9284 | 0.000999 |
| RTMIM | TCD4reg | CALR | FAS | 0.7672 | 0.2727 | 0.52 | 0.000999 |
| RTMIM | TCD4reg | CALR | IL7R | 0.7672 | 0.9682 | 0.8677 | 0.000999 |
| RTMIM | TCD4reg | CCL13 | CXCR3 | 0.3554 | 0.3357 | 0.3456 | 0.000999 |
| RTMIM | TCD4reg | CD86 | CD28 | 0.4042 | 0.313 | 0.3586 | 0.000999 |
| RTMIM | TCD4reg | CD86 | CTLA4 | 0.4042 | 0.7909 | 0.5976 | 0.000999 |
| RTMIM | TCD4reg | CD99 | CD99 | 0.8717 | 0.6603 | 0.766 | 0.000999 |
| RTMIM | TCD4reg | COL1A1 | CD44 | 0.4864 | 1.0896 | 0.788 | 0.000999 |
| RTMIM | TCD4reg | COL1A1 | ITGB1 | 0.4864 | 0.4001 | 0.4432 | 0.000999 |
| RTMIM | TCD4reg | CXCL12 | CCR7 | 0.3733 | 0.3098 | 0.3416 | 0.000999 |
| RTMIM | TCD4reg | CXCL12 | CD4 | 0.3733 | 0.429 | 0.4012 | 0.000999 |
| RTMIM | TCD4reg | CXCL12 | CXCR3 | 0.3733 | 0.3357 | 0.3545 | 0.000999 |
| RTMIM | TCD4reg | CXCL12 | CXCR4 | 0.3733 | 2.2962 | 1.3348 | 0.000999 |
| RTMIM | TCD4reg | CXCL12 | CXCR6 | 0.3733 | 0.3945 | 0.3839 | 0.000999 |
| RTMIM | TCD4reg | CXCL12 | S1PR4 | 0.3733 | 0.2744 | 0.3239 | 0.000999 |

| | | | | | | | |
|---|---|---|---|---|---|---|---|
| RTMIM | TCD4reg | CXCL16 | CCR7 | 0.7537 | 0.3098 | 0.5317 | 0.000999 |
| RTMIM | TCD4reg | CXCL16 | CXCR3 | 0.7537 | 0.3357 | 0.5447 | 0.000999 |
| RTMIM | TCD4reg | CXCL16 | CXCR4 | 0.7537 | 2.2962 | 1.5249 | 0.000999 |
| RTMIM | TCD4reg | CXCL16 | CXCR6 | 0.7537 | 0.3945 | 0.5741 | 0.000999 |
| RTMIM | TCD4reg | CXCL16 | S1PR4 | 0.7537 | 0.2744 | 0.514 | 0.000999 |
| RTMIM | TCD4reg | CXCL2 | CCR7 | 1.4022 | 0.3098 | 0.856 | 0.000999 |
| RTMIM | TCD4reg | CXCL2 | CXCR3 | 1.4022 | 0.3357 | 0.8689 | 0.000999 |
| RTMIM | TCD4reg | CXCL2 | CXCR4 | 1.4022 | 2.2962 | 1.8492 | 0.000999 |
| RTMIM | TCD4reg | CXCL2 | CXCR6 | 1.4022 | 0.3945 | 0.8983 | 0.000999 |
| RTMIM | TCD4reg | CXCL2 | S1PR4 | 1.4022 | 0.2744 | 0.8383 | 0.000999 |
| RTMIM | TCD4reg | CXCL3 | CCR7 | 1.1602 | 0.3098 | 0.735 | 0.000999 |
| RTMIM | TCD4reg | CXCL3 | CXCR3 | 1.1602 | 0.3357 | 0.7479 | 0.000999 |
| RTMIM | TCD4reg | CXCL3 | CXCR4 | 1.1602 | 2.2962 | 1.7282 | 0.000999 |
| RTMIM | TCD4reg | CXCL3 | CXCR6 | 1.1602 | 0.3945 | 0.7774 | 0.000999 |
| RTMIM | TCD4reg | CXCL3 | S1PR4 | 1.1602 | 0.2744 | 0.7173 | 0.000999 |
| RTMIM | TCD4reg | FN1 | CD44 | 0.3872 | 1.0896 | 0.7384 | 0.000999 |
| RTMIM | TCD4reg | FN1 | ITGB1 | 0.3872 | 0.4001 | 0.3937 | 0.000999 |
| RTMIM | TCD4reg | FN1 | RPSA | 0.3872 | 2.4999 | 1.4436 | 0.000999 |
| RTMIM | TCD4reg | GRN | TNFRSF1B | 1.4409 | 0.7982 | 1.1195 | 0.000999 |
| RTMIM | TCD4reg | GSTP1 | CD44 | 1.3126 | 1.0896 | 1.2011 | 0.000999 |
| RTMIM | TCD4reg | GSTP1 | FAS | 1.3126 | 0.2727 | 0.7927 | 0.000999 |
| RTMIM | TCD4reg | GSTP1 | IL7R | 1.3126 | 0.9682 | 1.1404 | 0.000999 |
| RTMIM | TCD4reg | HBEGF | CD44 | 0.6117 | 1.0896 | 0.8507 | 0.000999 |
| RTMIM | TCD4reg | HLA-DMA | CD4 | 1.7339 | 0.429 | 1.0814 | 0.000999 |
| RTMIM | TCD4reg | HLA-DRA | CD4 | 3.9393 | 0.429 | 2.1841 | 0.000999 |
| RTMIM | TCD4reg | HLA-DRA | PTPRC | 3.9393 | 1.6603 | 2.7998 | 0.000999 |
| RTMIM | TCD4reg | HSP90B1 | ITGB1 | 0.7626 | 0.4001 | 0.5814 | 0.000999 |
| RTMIM | TCD4reg | HSP90B1 | TNFRSF1B | 0.7626 | 0.7982 | 0.7804 | 0.000999 |
| RTMIM | TCD4reg | ICAM1 | IL2RA | 0.6002 | 0.7115 | 0.6558 | 0.000999 |
| RTMIM | TCD4reg | ICAM1 | IL2RG | 0.6002 | 1.1601 | 0.8802 | 0.000999 |

| | | | | | | | |
|---|---|---|---|---|---|---|---|
| RTMIM | TCD4reg | ICAM1 | ITGB2 | 0.6002 | 0.6238 | 0.612 | 0.000999 |
| RTMIM | TCD4reg | IGF1 | RACK1 | 0.4603 | 2.5298 | 1.495 | 0.000999 |
| RTMIM | TCD4reg | IL10 | CD28 | 0.2565 | 0.313 | 0.2848 | 0.000999 |
| RTMIM | TCD4reg | IL10 | IFNAR2 | 0.2565 | 0.2663 | 0.2614 | 0.000999 |
| RTMIM | TCD4reg | IL10 | IL10RA | 0.2565 | 0.4952 | 0.3759 | 0.000999 |
| RTMIM | TCD4reg | LGALS3 | CD6 | 0.7 | 0.2651 | 0.4826 | 0.000999 |
| RTMIM | TCD4reg | LGALS3 | FAS | 0.7 | 0.2727 | 0.4864 | 0.000999 |
| RTMIM | TCD4reg | LGALS3 | ITGB1 | 0.7 | 0.4001 | 0.5501 | 0.000999 |
| RTMIM | TCD4reg | NAMPT | ANXA6 | 1.2185 | 0.4557 | 0.8371 | 0.000999 |
| RTMIM | TCD4reg | PECAM1 | CXCR4 | 0.334 | 2.2962 | 1.3151 | 0.000999 |
| RTMIM | TCD4reg | PLAU | CD44 | 0.3288 | 1.0896 | 0.7092 | 0.000999 |
| RTMIM | TCD4reg | RTN4 | CD44 | 0.9278 | 1.0896 | 1.0087 | 0.000999 |
| RTMIM | TCD4reg | RTN4 | CD99 | 0.9278 | 0.6603 | 0.794 | 0.000999 |
| RTMIM | TCD4reg | SPP1 | CD44 | 0.61 | 1.0896 | 0.8498 | 0.000999 |
| RTMIM | TCD4reg | SPP1 | ITGB1 | 0.61 | 0.4001 | 0.505 | 0.000999 |
| RTMIM | TCD4reg | TGFB1 | ITGB1 | 0.4155 | 0.4001 | 0.4078 | 0.000999 |
| RTMIM | TCD4reg | THBS1 | CD47 | 0.3964 | 0.416 | 0.4062 | 0.000999 |
| RTMIM | TCD4reg | THBS1 | ITGB1 | 0.3964 | 0.4001 | 0.3983 | 0.000999 |
| RTMIM | TCD4reg | TNF | CALM1 | 0.3679 | 1.534 | 0.9509 | 0.000999 |
| RTMIM | TCD4reg | TNF | FAS | 0.3679 | 0.2727 | 0.3203 | 0.000999 |
| RTMIM | TCD4reg | TNF | ICOS | 0.3679 | 0.6754 | 0.5216 | 0.000999 |
| RTMIM | TCD4reg | TNF | RACK1 | 0.3679 | 2.5298 | 1.4488 | 0.000999 |
| RTMIM | TCD4reg | TNF | TNFRSF1B | 0.3679 | 0.7982 | 0.583 | 0.000999 |
| RTMIM | TCD4reg | TNFSF13B | ITGB1 | 0.4492 | 0.4001 | 0.4247 | 0.000999 |
| RTMIM | TCD4reg | ITGB1 | RACK1 | 0.4238 | 2.5298 | 1.4768 | 0.00799 |
| RTMIM | TCD8em | A2M | TNFRSF14 | 1.094 | 0.2915 | 0.6928 | 0.000999 |
| RTMIM | TCD8em | ADM | PTGER4 | 0.2706 | 0.2847 | 0.2777 | 0.000999 |
| RTMIM | TCD8em | ANXA1 | CXCR4 | 1.1316 | 2.8389 | 1.9853 | 0.000999 |
| RTMIM | TCD8em | C3 | CXCR3 | 0.3607 | 0.4189 | 0.3898 | 0.000999 |
| RTMIM | TCD8em | C3 | CXCR4 | 0.3607 | 2.8389 | 1.5998 | 0.000999 |

| | | | | | | | |
|---|---|---|---|---|---|---|---|
| RTMIM | TCD8em | C3 | CXCR6 | 0.3607 | 0.3399 | 0.3503 | 0.000999 |
| RTMIM | TCD8em | C3 | ITGB2 | 0.3607 | 1.0018 | 0.6813 | 0.000999 |
| RTMIM | TCD8em | CALR | IL7R | 0.7672 | 0.7183 | 0.7428 | 0.000999 |
| RTMIM | TCD8em | CCL13 | CXCR3 | 0.3554 | 0.4189 | 0.3872 | 0.000999 |
| RTMIM | TCD8em | COL1A1 | CD44 | 0.4864 | 1.119 | 0.8027 | 0.000999 |
| RTMIM | TCD8em | COL1A1 | ITGB1 | 0.4864 | 0.2557 | 0.3711 | 0.000999 |
| RTMIM | TCD8em | CXCL12 | CXCR3 | 0.3733 | 0.4189 | 0.3961 | 0.000999 |
| RTMIM | TCD8em | CXCL12 | CXCR4 | 0.3733 | 2.8389 | 1.6061 | 0.000999 |
| RTMIM | TCD8em | CXCL12 | CXCR6 | 0.3733 | 0.3399 | 0.3566 | 0.000999 |
| RTMIM | TCD8em | CXCL12 | S1PR4 | 0.3733 | 0.2677 | 0.3205 | 0.000999 |
| RTMIM | TCD8em | CXCL16 | CXCR3 | 0.7537 | 0.4189 | 0.5863 | 0.000999 |
| RTMIM | TCD8em | CXCL16 | CXCR4 | 0.7537 | 2.8389 | 1.7963 | 0.000999 |
| RTMIM | TCD8em | CXCL16 | CXCR6 | 0.7537 | 0.3399 | 0.5468 | 0.000999 |
| RTMIM | TCD8em | CXCL16 | S1PR4 | 0.7537 | 0.2677 | 0.5107 | 0.000999 |
| RTMIM | TCD8em | CXCL2 | CXCR3 | 1.4022 | 0.4189 | 0.9105 | 0.000999 |
| RTMIM | TCD8em | CXCL2 | CXCR4 | 1.4022 | 2.8389 | 2.1205 | 0.000999 |
| RTMIM | TCD8em | CXCL2 | CXCR6 | 1.4022 | 0.3399 | 0.871 | 0.000999 |
| RTMIM | TCD8em | CXCL2 | S1PR4 | 1.4022 | 0.2677 | 0.8349 | 0.000999 |
| RTMIM | TCD8em | CXCL3 | CXCR3 | 1.1602 | 0.4189 | 0.7896 | 0.000999 |
| RTMIM | TCD8em | CXCL3 | CXCR4 | 1.1602 | 2.8389 | 1.9996 | 0.000999 |
| RTMIM | TCD8em | CXCL3 | CXCR6 | 1.1602 | 0.3399 | 0.75 | 0.000999 |
| RTMIM | TCD8em | CXCL3 | S1PR4 | 1.1602 | 0.2677 | 0.714 | 0.000999 |
| RTMIM | TCD8em | FN1 | CD44 | 0.3872 | 1.119 | 0.7531 | 0.000999 |
| RTMIM | TCD8em | FN1 | ITGA4 | 0.3872 | 0.2823 | 0.3348 | 0.000999 |
| RTMIM | TCD8em | FN1 | ITGB1 | 0.3872 | 0.2557 | 0.3215 | 0.000999 |
| RTMIM | TCD8em | FN1 | RPSA | 0.3872 | 2.3686 | 1.3779 | 0.000999 |
| RTMIM | TCD8em | GRN | TNFRSF1B | 1.4409 | 0.3038 | 0.8723 | 0.000999 |
| RTMIM | TCD8em | GSTP1 | CD44 | 1.3126 | 1.119 | 1.2158 | 0.000999 |
| RTMIM | TCD8em | GSTP1 | IL7R | 1.3126 | 0.7183 | 1.0155 | 0.000999 |
| RTMIM | TCD8em | GSTP1 | ITGA4 | 1.3126 | 0.2823 | 0.7975 | 0.000999 |

| | | | | | | | |
|---|---|---|---|---|---|---|---|
| RTMIM | TCD8em | HBEGF | CD44 | 0.6117 | 1.119 | 0.8654 | 0.000999 |
| RTMIM | TCD8em | HLA-DRA | PTPRC | 3.9393 | 1.7912 | 2.8652 | 0.000999 |
| RTMIM | TCD8em | HSP90B1 | ITGB1 | 0.7626 | 0.2557 | 0.5092 | 0.000999 |
| RTMIM | TCD8em | HSP90B1 | TNFRSF1B | 0.7626 | 0.3038 | 0.5332 | 0.000999 |
| RTMIM | TCD8em | ICAM1 | IL2RG | 0.6002 | 0.9501 | 0.7752 | 0.000999 |
| RTMIM | TCD8em | ICAM1 | ITGAL | 0.6002 | 0.3162 | 0.4582 | 0.000999 |
| RTMIM | TCD8em | ICAM1 | ITGB2 | 0.6002 | 1.0018 | 0.801 | 0.000999 |
| RTMIM | TCD8em | ICAM1 | SPN | 0.6002 | 0.2603 | 0.4303 | 0.000999 |
| RTMIM | TCD8em | IGF1 | RACK1 | 0.4603 | 2.4374 | 1.4488 | 0.000999 |
| RTMIM | TCD8em | IL10 | IL10RA | 0.2565 | 0.4383 | 0.3474 | 0.000999 |
| RTMIM | TCD8em | ITGB1 | CD47 | 0.4238 | 0.4276 | 0.4257 | 0.000999 |
| RTMIM | TCD8em | ITGB1 | ITGA4 | 0.4238 | 0.2823 | 0.3531 | 0.000999 |
| RTMIM | TCD8em | ITGB1 | RACK1 | 0.4238 | 2.4374 | 1.4306 | 0.000999 |
| RTMIM | TCD8em | LGALS3 | CD6 | 0.7 | 0.4204 | 0.5602 | 0.000999 |
| RTMIM | TCD8em | LGALS3 | ITGB1 | 0.7 | 0.2557 | 0.4779 | 0.000999 |
| RTMIM | TCD8em | LGALS3 | SPN | 0.7 | 0.2603 | 0.4802 | 0.000999 |
| RTMIM | TCD8em | NAMPT | ANXA6 | 1.2185 | 0.552 | 0.8852 | 0.000999 |
| RTMIM | TCD8em | PECAM1 | CXCR4 | 0.334 | 2.8389 | 1.5864 | 0.000999 |
| RTMIM | TCD8em | PLAU | CD44 | 0.3288 | 1.119 | 0.7239 | 0.000999 |
| RTMIM | TCD8em | RTN4 | CD44 | 0.9278 | 1.119 | 1.0234 | 0.000999 |
| RTMIM | TCD8em | RTN4 | CD99 | 0.9278 | 1.1525 | 1.0401 | 0.000999 |
| RTMIM | TCD8em | SPP1 | CD44 | 0.61 | 1.119 | 0.8645 | 0.000999 |
| RTMIM | TCD8em | SPP1 | ITGA4 | 0.61 | 0.2823 | 0.4462 | 0.000999 |
| RTMIM | TCD8em | SPP1 | ITGB1 | 0.61 | 0.2557 | 0.4329 | 0.000999 |
| RTMIM | TCD8em | TGFB1 | ITGB1 | 0.4155 | 0.2557 | 0.3356 | 0.000999 |
| RTMIM | TCD8em | THBS1 | CD47 | 0.3964 | 0.4276 | 0.412 | 0.000999 |
| RTMIM | TCD8em | THBS1 | ITGA4 | 0.3964 | 0.2823 | 0.3394 | 0.000999 |
| RTMIM | TCD8em | THBS1 | ITGB1 | 0.3964 | 0.2557 | 0.3261 | 0.000999 |
| RTMIM | TCD8em | TNF | CALM1 | 0.3679 | 1.5262 | 0.947 | 0.000999 |
| RTMIM | TCD8em | TNF | RACK1 | 0.3679 | 2.4374 | 1.4026 | 0.000999 |

| | | | | | | | |
|---|---|---|---|---|---|---|---|
| RTMIM | TCD8em | TNFSF13B | ITGB1 | 0.4492 | 0.2557 | 0.3525 | 0.000999 |
| RTMIM | TCD8em | ANXA1 | CXCR3 | 1.1316 | 0.4189 | 0.7752 | 0.004 |
| RTMIM | TCD8em | TNF | TNFRSF1B | 0.3679 | 0.3038 | 0.3358 | 0.005 |
| RTMIM | TCD8em | CALR | CD44 | 0.7672 | 1.119 | 0.9431 | 0.00599 |
| RTMIM | TCD8em | ANXA1 | CXCR6 | 1.1316 | 0.3399 | 0.7357 | 0.00899 |
| RTMIM | TCD8ex | A2M | TNFRSF14 | 1.094 | 0.3094 | 0.7017 | 0.000999 |
| RTMIM | TCD8ex | ADM | PTGER4 | 0.2706 | 0.3089 | 0.2898 | 0.000999 |
| RTMIM | TCD8ex | ANXA1 | CD44 | 1.1316 | 0.8219 | 0.9768 | 0.000999 |
| RTMIM | TCD8ex | ANXA1 | CXCR3 | 1.1316 | 0.3647 | 0.7481 | 0.000999 |
| RTMIM | TCD8ex | ANXA1 | CXCR4 | 1.1316 | 2.3472 | 1.7394 | 0.000999 |
| RTMIM | TCD8ex | ANXA1 | CXCR6 | 1.1316 | 0.6626 | 0.8971 | 0.000999 |
| RTMIM | TCD8ex | ANXA1 | ITGA4 | 1.1316 | 0.4432 | 0.7874 | 0.000999 |
| RTMIM | TCD8ex | ANXA1 | VCAM1 | 1.1316 | 0.4462 | 0.7889 | 0.000999 |
| RTMIM | TCD8ex | APOE | VCAM1 | 1.1762 | 0.4462 | 0.8112 | 0.000999 |
| RTMIM | TCD8ex | C3 | CXCR3 | 0.3607 | 0.3647 | 0.3627 | 0.000999 |
| RTMIM | TCD8ex | C3 | CXCR4 | 0.3607 | 2.3472 | 1.354 | 0.000999 |
| RTMIM | TCD8ex | C3 | CXCR6 | 0.3607 | 0.6626 | 0.5116 | 0.000999 |
| RTMIM | TCD8ex | C3 | ITGB2 | 0.3607 | 0.7635 | 0.5621 | 0.000999 |
| RTMIM | TCD8ex | CCL13 | CXCR3 | 0.3554 | 0.3647 | 0.3601 | 0.000999 |
| RTMIM | TCD8ex | CD86 | CTLA4 | 0.4042 | 0.4883 | 0.4462 | 0.000999 |
| RTMIM | TCD8ex | COL1A1 | CD44 | 0.4864 | 0.8219 | 0.6542 | 0.000999 |
| RTMIM | TCD8ex | COL1A1 | ITGB1 | 0.4864 | 0.2995 | 0.3929 | 0.000999 |
| RTMIM | TCD8ex | CXCL12 | CXCR3 | 0.3733 | 0.3647 | 0.369 | 0.000999 |
| RTMIM | TCD8ex | CXCL12 | CXCR4 | 0.3733 | 2.3472 | 1.3603 | 0.000999 |
| RTMIM | TCD8ex | CXCL12 | CXCR6 | 0.3733 | 0.6626 | 0.518 | 0.000999 |
| RTMIM | TCD8ex | CXCL16 | CXCR3 | 0.7537 | 0.3647 | 0.5592 | 0.000999 |
| RTMIM | TCD8ex | CXCL16 | CXCR4 | 0.7537 | 2.3472 | 1.5504 | 0.000999 |
| RTMIM | TCD8ex | CXCL16 | CXCR6 | 0.7537 | 0.6626 | 0.7081 | 0.000999 |
| RTMIM | TCD8ex | CXCL2 | CXCR3 | 1.4022 | 0.3647 | 0.8834 | 0.000999 |
| RTMIM | TCD8ex | CXCL2 | CXCR4 | 1.4022 | 2.3472 | 1.8747 | 0.000999 |

| | | | | | | | |
|---|---|---|---|---|---|---|---|
| RTMIM | TCD8ex | CXCL2 | CXCR6 | 1.4022 | 0.6626 | 1.0324 | 0.000999 |
| RTMIM | TCD8ex | CXCL3 | CXCR3 | 1.1602 | 0.3647 | 0.7624 | 0.000999 |
| RTMIM | TCD8ex | CXCL3 | CXCR4 | 1.1602 | 2.3472 | 1.7537 | 0.000999 |
| RTMIM | TCD8ex | CXCL3 | CXCR6 | 1.1602 | 0.6626 | 0.9114 | 0.000999 |
| RTMIM | TCD8ex | FN1 | CD44 | 0.3872 | 0.8219 | 0.6046 | 0.000999 |
| RTMIM | TCD8ex | FN1 | ITGA4 | 0.3872 | 0.4432 | 0.4152 | 0.000999 |
| RTMIM | TCD8ex | FN1 | ITGB1 | 0.3872 | 0.2995 | 0.3434 | 0.000999 |
| RTMIM | TCD8ex | FN1 | ITGB7 | 0.3872 | 0.3007 | 0.344 | 0.000999 |
| RTMIM | TCD8ex | FN1 | RPSA | 0.3872 | 2.2697 | 1.3284 | 0.000999 |
| RTMIM | TCD8ex | GRN | TNFRSF1B | 1.4409 | 0.6436 | 1.0423 | 0.000999 |
| RTMIM | TCD8ex | GSTP1 | CD44 | 1.3126 | 0.8219 | 1.0673 | 0.000999 |
| RTMIM | TCD8ex | GSTP1 | ITGA4 | 1.3126 | 0.4432 | 0.8779 | 0.000999 |
| RTMIM | TCD8ex | GSTP1 | VCAM1 | 1.3126 | 0.4462 | 0.8794 | 0.000999 |
| RTMIM | TCD8ex | HBEGF | CD44 | 0.6117 | 0.8219 | 0.7168 | 0.000999 |
| RTMIM | TCD8ex | HLA-DRA | PDCD1 | 3.9393 | 0.6109 | 2.2751 | 0.000999 |
| RTMIM | TCD8ex | HLA-DRA | PTPRC | 3.9393 | 1.6762 | 2.8077 | 0.000999 |
| RTMIM | TCD8ex | HSP90B1 | ITGB1 | 0.7626 | 0.2995 | 0.5311 | 0.000999 |
| RTMIM | TCD8ex | HSP90B1 | TNFRSF1B | 0.7626 | 0.6436 | 0.7031 | 0.000999 |
| RTMIM | TCD8ex | ICAM1 | IL2RG | 0.6002 | 1.2126 | 0.9064 | 0.000999 |
| RTMIM | TCD8ex | ICAM1 | ITGB2 | 0.6002 | 0.7635 | 0.6818 | 0.000999 |
| RTMIM | TCD8ex | ICAM1 | SPN | 0.6002 | 0.3486 | 0.4744 | 0.000999 |
| RTMIM | TCD8ex | IGF1 | RACK1 | 0.4603 | 2.3353 | 1.3978 | 0.000999 |
| RTMIM | TCD8ex | IL10 | IL10RA | 0.2565 | 0.3816 | 0.3191 | 0.000999 |
| RTMIM | TCD8ex | ITGB1 | CD47 | 0.4238 | 0.3952 | 0.4095 | 0.000999 |
| RTMIM | TCD8ex | ITGB1 | ITGA4 | 0.4238 | 0.4432 | 0.4335 | 0.000999 |
| RTMIM | TCD8ex | ITGB1 | RACK1 | 0.4238 | 2.3353 | 1.3796 | 0.000999 |
| RTMIM | TCD8ex | ITGB1 | VCAM1 | 0.4238 | 0.4462 | 0.435 | 0.000999 |
| RTMIM | TCD8ex | LGALS3 | CD6 | 0.7 | 0.318 | 0.509 | 0.000999 |
| RTMIM | TCD8ex | LGALS3 | ITGB1 | 0.7 | 0.2995 | 0.4998 | 0.000999 |
| RTMIM | TCD8ex | LGALS3 | SPN | 0.7 | 0.3486 | 0.5243 | 0.000999 |

| | | | | | | | |
|---|---|---|---|---|---|---|---|
| RTMIM | TCD8ex | NAMPT | ANXA6 | 1.2185 | 0.5678 | 0.8931 | 0.000999 |
| RTMIM | TCD8ex | PECAM1 | CXCR4 | 0.334 | 2.3472 | 1.3406 | 0.000999 |
| RTMIM | TCD8ex | PLAU | CD44 | 0.3288 | 0.8219 | 0.5754 | 0.000999 |
| RTMIM | TCD8ex | RTN4 | CD44 | 0.9278 | 0.8219 | 0.8749 | 0.000999 |
| RTMIM | TCD8ex | RTN4 | CD99 | 0.9278 | 1.2049 | 1.0663 | 0.000999 |
| RTMIM | TCD8ex | SPP1 | CD44 | 0.61 | 0.8219 | 0.716 | 0.000999 |
| RTMIM | TCD8ex | SPP1 | ITGA4 | 0.61 | 0.4432 | 0.5266 | 0.000999 |
| RTMIM | TCD8ex | SPP1 | ITGB1 | 0.61 | 0.2995 | 0.4547 | 0.000999 |
| RTMIM | TCD8ex | TGFB1 | ITGB1 | 0.4155 | 0.2995 | 0.3575 | 0.000999 |
| RTMIM | TCD8ex | THBS1 | CD47 | 0.3964 | 0.3952 | 0.3958 | 0.000999 |
| RTMIM | TCD8ex | THBS1 | ITGA4 | 0.3964 | 0.4432 | 0.4198 | 0.000999 |
| RTMIM | TCD8ex | THBS1 | ITGB1 | 0.3964 | 0.2995 | 0.348 | 0.000999 |
| RTMIM | TCD8ex | TNF | CALM1 | 0.3679 | 1.4627 | 0.9153 | 0.000999 |
| RTMIM | TCD8ex | TNF | ICOS | 0.3679 | 0.3384 | 0.3531 | 0.000999 |
| RTMIM | TCD8ex | TNF | RACK1 | 0.3679 | 2.3353 | 1.3516 | 0.000999 |
| RTMIM | TCD8ex | TNF | TNFRSF1B | 0.3679 | 0.6436 | 0.5057 | 0.000999 |
| RTMIM | TCD8ex | TNFSF13B | ITGB1 | 0.4492 | 0.2995 | 0.3744 | 0.000999 |
| RTMIM | TGD | A2M | TNFRSF14 | 1.094 | 0.4642 | 0.7791 | 0.000999 |
| RTMIM | TGD | ADM | PTGDR | 0.2706 | 0.7153 | 0.493 | 0.000999 |
| RTMIM | TGD | ANXA1 | CD44 | 1.1316 | 1.1342 | 1.1329 | 0.000999 |
| RTMIM | TGD | ANXA1 | CXCR3 | 1.1316 | 1.1748 | 1.1532 | 0.000999 |
| RTMIM | TGD | ANXA1 | CXCR4 | 1.1316 | 1.4894 | 1.3105 | 0.000999 |
| RTMIM | TGD | ANXA1 | ITGA4 | 1.1316 | 0.3968 | 0.7642 | 0.000999 |
| RTMIM | TGD | APOE | SORL1 | 1.1762 | 0.3318 | 0.754 | 0.000999 |
| RTMIM | TGD | ARF1 | ITGA4 | 0.5727 | 0.3968 | 0.4848 | 0.000999 |
| RTMIM | TGD | C3 | CXCR3 | 0.3607 | 1.1748 | 0.7677 | 0.000999 |
| RTMIM | TGD | C3 | CXCR4 | 0.3607 | 1.4894 | 0.925 | 0.000999 |
| RTMIM | TGD | C3 | ITGB2 | 0.3607 | 1.4444 | 0.9026 | 0.000999 |
| RTMIM | TGD | CALM1 | RPSA | 1.0352 | 2.8814 | 1.9583 | 0.000999 |
| RTMIM | TGD | CALR | CD44 | 0.7672 | 1.1342 | 0.9507 | 0.000999 |

| | | | | | | | |
|---|---|---|---|---|---|---|---|
| RTMIM | TGD | CALR | IL7R | 0.7672 | 1.3862 | 1.0767 | 0.000999 |
| RTMIM | TGD | CCL13 | CXCR3 | 0.3554 | 1.1748 | 0.7651 | 0.000999 |
| RTMIM | TGD | CD99 | CD99 | 0.8717 | 1.7154 | 1.2936 | 0.000999 |
| RTMIM | TGD | COL1A1 | CD44 | 0.4864 | 1.1342 | 0.8103 | 0.000999 |
| RTMIM | TGD | CXCL12 | CXCR3 | 0.3733 | 1.1748 | 0.7741 | 0.000999 |
| RTMIM | TGD | CXCL12 | CXCR4 | 0.3733 | 1.4894 | 0.9314 | 0.000999 |
| RTMIM | TGD | CXCL16 | CXCR3 | 0.7537 | 1.1748 | 0.9642 | 0.000999 |
| RTMIM | TGD | CXCL16 | CXCR4 | 0.7537 | 1.4894 | 1.1215 | 0.000999 |
| RTMIM | TGD | CXCL2 | CXCR3 | 1.4022 | 1.1748 | 1.2885 | 0.000999 |
| RTMIM | TGD | CXCL2 | CXCR4 | 1.4022 | 1.4894 | 1.4458 | 0.000999 |
| RTMIM | TGD | CXCL3 | CXCR3 | 1.1602 | 1.1748 | 1.1675 | 0.000999 |
| RTMIM | TGD | CXCL3 | CXCR4 | 1.1602 | 1.4894 | 1.3248 | 0.000999 |
| RTMIM | TGD | FN1 | CD44 | 0.3872 | 1.1342 | 0.7607 | 0.000999 |
| RTMIM | TGD | FN1 | ITGA4 | 0.3872 | 0.3968 | 0.392 | 0.000999 |
| RTMIM | TGD | FN1 | ITGB7 | 0.3872 | 0.459 | 0.4231 | 0.000999 |
| RTMIM | TGD | FN1 | RPSA | 0.3872 | 2.8814 | 1.6343 | 0.000999 |
| RTMIM | TGD | GSTP1 | CD44 | 1.3126 | 1.1342 | 1.2234 | 0.000999 |
| RTMIM | TGD | GSTP1 | IL7R | 1.3126 | 1.3862 | 1.3494 | 0.000999 |
| RTMIM | TGD | GSTP1 | ITGA4 | 1.3126 | 0.3968 | 0.8547 | 0.000999 |
| RTMIM | TGD | HBEGF | CD44 | 0.6117 | 1.1342 | 0.8729 | 0.000999 |
| RTMIM | TGD | HLA-A | CD8A | 2.3127 | 1.746 | 2.0293 | 0.000999 |
| RTMIM | TGD | HLA-DRA | PTPRC | 3.9393 | 1.8653 | 2.9023 | 0.000999 |
| RTMIM | TGD | HLA-E | CD8A | 1.7652 | 1.746 | 1.7556 | 0.000999 |
| RTMIM | TGD | HLA-E | KLRC2 | 1.7652 | 0.8082 | 1.2867 | 0.000999 |
| RTMIM | TGD | HLA-E | KLRD1 | 1.7652 | 1.3451 | 1.5552 | 0.000999 |
| RTMIM | TGD | HMGB1 | ITGA4 | 1.2113 | 0.3968 | 0.8041 | 0.000999 |
| RTMIM | TGD | HMGB2 | CD44 | 0.3507 | 1.1342 | 0.7425 | 0.000999 |
| RTMIM | TGD | ICAM1 | IL2RG | 0.6002 | 1.561 | 1.0806 | 0.000999 |
| RTMIM | TGD | ICAM1 | ITGAL | 0.6002 | 0.3731 | 0.4867 | 0.000999 |
| RTMIM | TGD | ICAM1 | ITGB2 | 0.6002 | 1.4444 | 1.0223 | 0.000999 |

| | | | | | | | |
|---|---|---|---|---|---|---|---|
| RTMIM | TGD | ICAM1 | SPN | 0.6002 | 0.5644 | 0.5823 | 0.000999 |
| RTMIM | TGD | IGF1 | RACK1 | 0.4603 | 2.7287 | 1.5945 | 0.000999 |
| RTMIM | TGD | IL10 | IFNAR2 | 0.2565 | 0.4338 | 0.3452 | 0.000999 |
| RTMIM | TGD | IL10 | IL10RA | 0.2565 | 0.625 | 0.4407 | 0.000999 |
| RTMIM | TGD | ITGB1 | ITGA4 | 0.4238 | 0.3968 | 0.4103 | 0.000999 |
| RTMIM | TGD | ITGB1 | PTK2B | 0.4238 | 0.3668 | 0.3953 | 0.000999 |
| RTMIM | TGD | ITGB1 | RACK1 | 0.4238 | 2.7287 | 1.5763 | 0.000999 |
| RTMIM | TGD | ITGB2 | ITGAL | 0.862 | 0.3731 | 0.6176 | 0.000999 |
| RTMIM | TGD | LGALS3 | CD6 | 0.7 | 0.4684 | 0.5842 | 0.000999 |
| RTMIM | TGD | LGALS3 | SPN | 0.7 | 0.5644 | 0.6322 | 0.000999 |
| RTMIM | TGD | LRPAP1 | SORL1 | 0.3277 | 0.3318 | 0.3297 | 0.000999 |
| RTMIM | TGD | MIF | ITGA4 | 0.562 | 0.3968 | 0.4794 | 0.000999 |
| RTMIM | TGD | NAMPT | ANXA6 | 1.2185 | 0.8553 | 1.0369 | 0.000999 |
| RTMIM | TGD | PECAM1 | CXCR4 | 0.334 | 1.4894 | 0.9117 | 0.000999 |
| RTMIM | TGD | PLAU | CD44 | 0.3288 | 1.1342 | 0.7315 | 0.000999 |
| RTMIM | TGD | PTPRC | CD2 | 0.5365 | 1.4522 | 0.9943 | 0.000999 |
| RTMIM | TGD | PTPRC | CD8A | 0.5365 | 1.746 | 1.1413 | 0.000999 |
| RTMIM | TGD | PTPRC | CXCR4 | 0.5365 | 1.4894 | 1.0129 | 0.000999 |
| RTMIM | TGD | RPS19 | ITGA4 | 3.2584 | 0.3968 | 1.8276 | 0.000999 |
| RTMIM | TGD | RPS19 | RPSA | 3.2584 | 2.8814 | 3.0699 | 0.000999 |
| RTMIM | TGD | RTN4 | CD44 | 0.9278 | 1.1342 | 1.031 | 0.000999 |
| RTMIM | TGD | RTN4 | CD99 | 0.9278 | 1.7154 | 1.3216 | 0.000999 |
| RTMIM | TGD | SPP1 | CD44 | 0.61 | 1.1342 | 0.8721 | 0.000999 |
| RTMIM | TGD | SPP1 | ITGA4 | 0.61 | 0.3968 | 0.5034 | 0.000999 |
| RTMIM | TGD | THBS1 | CD47 | 0.3964 | 0.4765 | 0.4365 | 0.000999 |
| RTMIM | TGD | THBS1 | ITGA4 | 0.3964 | 0.3968 | 0.3966 | 0.000999 |
| RTMIM | TGD | TNF | CALM1 | 0.3679 | 1.7587 | 1.0633 | 0.000999 |
| RTMIM | TGD | TNF | RACK1 | 0.3679 | 2.7287 | 1.5483 | 0.000999 |
| RTMIM | TGD | VEGFA | PTK2B | 0.4127 | 0.3668 | 0.3897 | 0.000999 |
| RTMIM | TGD | ARF1 | CD44 | 0.5727 | 1.1342 | 0.8535 | 0.002 |

| | | | | | | | |
|---|---|---|---|---|---|---|---|
| RTMIM | TGD | ITGB1 | CD47 | 0.4238 | 0.4765 | 0.4502 | 0.004 |
| RTMIM | TGD | PTPRC | ITGAL | 0.5365 | 0.3731 | 0.4548 | 0.005 |
| NKT | RTMIM | ANXA1 | C3AR1 | 1.2259 | 0.6118 | 0.9189 | 0.000999 |
| NKT | RTMIM | ANXA1 | C5AR1 | 1.2259 | 0.8768 | 1.0514 | 0.000999 |
| NKT | RTMIM | ANXA1 | CCR1 | 1.2259 | 0.2693 | 0.7476 | 0.000999 |
| NKT | RTMIM | ANXA1 | CD4 | 1.2259 | 0.6545 | 0.9402 | 0.000999 |
| NKT | RTMIM | ANXA1 | CD44 | 1.2259 | 0.8187 | 1.0223 | 0.000999 |
| NKT | RTMIM | ANXA1 | FPR3 | 1.2259 | 0.511 | 0.8685 | 0.000999 |
| NKT | RTMIM | ARF1 | CD4 | 0.7991 | 0.6545 | 0.7268 | 0.000999 |
| NKT | RTMIM | ARF1 | CD44 | 0.7991 | 0.8187 | 0.8089 | 0.000999 |
| NKT | RTMIM | CALM1 | TNFRSF1A | 1.9335 | 0.5118 | 1.2226 | 0.000999 |
| NKT | RTMIM | CALM1 | TNFRSF1B | 1.9335 | 0.5749 | 1.2542 | 0.000999 |
| NKT | RTMIM | CALR | CD44 | 0.9479 | 0.8187 | 0.8833 | 0.000999 |
| NKT | RTMIM | CALR | LRP1 | 0.9479 | 0.5167 | 0.7323 | 0.000999 |
| NKT | RTMIM | CD2 | CD4 | 1.1465 | 0.6545 | 0.9005 | 0.000999 |
| NKT | RTMIM | CD320 | TGFBR2 | 0.2545 | 0.3722 | 0.3134 | 0.000999 |
| NKT | RTMIM | HMGB1 | LY96 | 1.195 | 0.9322 | 1.0636 | 0.000999 |
| NKT | RTMIM | HMGB1 | TLR2 | 1.195 | 0.2837 | 0.7393 | 0.000999 |
| NKT | RTMIM | HMGB2 | CD44 | 0.6467 | 0.8187 | 0.7327 | 0.000999 |
| NKT | RTMIM | HMGB2 | DAB2 | 0.6467 | 1.5391 | 1.0929 | 0.000999 |
| NKT | RTMIM | HSP90B1 | LRP1 | 0.6138 | 0.5167 | 0.5653 | 0.000999 |
| NKT | RTMIM | IL16 | CD4 | 0.3556 | 0.6545 | 0.505 | 0.000999 |
| NKT | RTMIM | IL16 | FCGR1A | 0.3556 | 0.3257 | 0.3406 | 0.000999 |
| NKT | RTMIM | ITGA4 | CD44 | 0.2946 | 0.8187 | 0.5566 | 0.000999 |
| NKT | RTMIM | ITGA4 | TFRC | 0.2946 | 0.6023 | 0.4484 | 0.000999 |
| NKT | RTMIM | ITGAL | ICAM1 | 0.6778 | 0.6002 | 0.639 | 0.000999 |
| NKT | RTMIM | ITGB1 | CD36 | 0.488 | 0.3859 | 0.4369 | 0.000999 |
| NKT | RTMIM | ITGB1 | CD46 | 0.488 | 0.2644 | 0.3762 | 0.000999 |
| NKT | RTMIM | ITGB1 | DAB2 | 0.488 | 1.5391 | 1.0135 | 0.000999 |
| NKT | RTMIM | ITGB1 | PLAUR | 0.488 | 1.1344 | 0.8112 | 0.000999 |

| | | | | | | | |
|---|---|---|---|---|---|---|---|
| NKT | RTMIM | ITGB2 | ICAM1 | 1.6481 | 0.6002 | 1.1242 | 0.000999 |
| NKT | RTMIM | ITGB2 | ITGAM | 1.6481 | 0.3355 | 0.9918 | 0.000999 |
| NKT | RTMIM | ITGB2 | PLAUR | 1.6481 | 1.1344 | 1.3912 | 0.000999 |
| NKT | RTMIM | LRPAP1 | LRP1 | 0.2807 | 0.5167 | 0.3987 | 0.000999 |
| NKT | RTMIM | LTB | TNFRSF1A | 0.2605 | 0.5118 | 0.3861 | 0.000999 |
| NKT | RTMIM | MIF | CD74 | 0.6657 | 4.5079 | 2.5868 | 0.000999 |
| NKT | RTMIM | PSAP | LRP1 | 0.6777 | 0.5167 | 0.5972 | 0.000999 |
| NKT | RTMIM | PTPRC | CD4 | 1.8797 | 0.6545 | 1.2671 | 0.000999 |
| NKT | RTMIM | PTPRC | CD44 | 1.8797 | 0.8187 | 1.3492 | 0.000999 |
| NKT | RTMIM | RPS19 | C5AR1 | 3.5336 | 0.8768 | 2.2052 | 0.000999 |
| NKT | RTMIM | RPS19 | CD4 | 3.5336 | 0.6545 | 2.0941 | 0.000999 |
| NKT | RTMIM | RPS19 | ICAM1 | 3.5336 | 0.6002 | 2.0669 | 0.000999 |
| NKT | RTMIM | SPN | SIGLEC1 | 0.6398 | 0.3312 | 0.4855 | 0.000999 |
| NKT | RTMIM | TGFB1 | TGFBR2 | 0.5441 | 0.3722 | 0.4581 | 0.000999 |
| NKT | RTMIM | TGFB1 | TLR2 | 0.5441 | 0.2837 | 0.4139 | 0.000999 |
| NKT | RTMIM | TIMP1 | LRP1 | 0.3005 | 0.5167 | 0.4086 | 0.000999 |
| NKT | RTMIM | XCL2 | LPAR6 | 0.2774 | 0.4942 | 0.3858 | 0.000999 |
| TCD4ex | RTMIM | ANXA1 | C3AR1 | 0.5174 | 0.6118 | 0.5646 | 0.000999 |
| TCD4ex | RTMIM | ANXA1 | C5AR1 | 0.5174 | 0.8768 | 0.6971 | 0.000999 |
| TCD4ex | RTMIM | ANXA1 | FPR3 | 0.5174 | 0.511 | 0.5142 | 0.000999 |
| TCD4ex | RTMIM | APOE | LRP1 | 0.2557 | 0.5167 | 0.3862 | 0.000999 |
| TCD4ex | RTMIM | AREG | ICAM1 | 0.3427 | 0.6002 | 0.4715 | 0.000999 |
| TCD4ex | RTMIM | BTLA | TNFRSF14 | 0.3493 | 0.2649 | 0.3071 | 0.000999 |
| TCD4ex | RTMIM | CALM1 | TNFRSF1A | 1.2957 | 0.5118 | 0.9037 | 0.000999 |
| TCD4ex | RTMIM | CALM1 | TNFRSF1B | 1.2957 | 0.5749 | 0.9353 | 0.000999 |
| TCD4ex | RTMIM | CALR | LRP1 | 0.748 | 0.5167 | 0.6324 | 0.000999 |
| TCD4ex | RTMIM | CD2 | CD4 | 1.5731 | 0.6545 | 1.1138 | 0.000999 |
| TCD4ex | RTMIM | CD28 | CD4 | 0.3256 | 0.6545 | 0.49 | 0.000999 |
| TCD4ex | RTMIM | CXCL13 | C3AR1 | 2.3367 | 0.6118 | 1.4743 | 0.000999 |
| TCD4ex | RTMIM | CXCL13 | C5AR1 | 2.3367 | 0.8768 | 1.6067 | 0.000999 |

| | | | | | | | |
|---|---|---|---|---|---|---|---|
| TCD4ex | RTMIM | CXCL13 | CCR1 | 2.3367 | 0.2693 | 1.303 | 0.000999 |
| TCD4ex | RTMIM | HMGB1 | LY96 | 1.3876 | 0.9322 | 1.1599 | 0.000999 |
| TCD4ex | RTMIM | HMGB1 | TLR2 | 1.3876 | 0.2837 | 0.8356 | 0.000999 |
| TCD4ex | RTMIM | HMGB2 | DAB2 | 1.0243 | 1.5391 | 1.2817 | 0.000999 |
| TCD4ex | RTMIM | HSP90B1 | LRP1 | 0.3944 | 0.5167 | 0.4556 | 0.000999 |
| TCD4ex | RTMIM | ICAM3 | ITGB2 | 0.573 | 0.862 | 0.7175 | 0.000999 |
| TCD4ex | RTMIM | ITGAL | ICAM1 | 0.2603 | 0.6002 | 0.4303 | 0.000999 |
| TCD4ex | RTMIM | ITGB1 | CD36 | 0.4615 | 0.3859 | 0.4237 | 0.000999 |
| TCD4ex | RTMIM | ITGB1 | DAB2 | 0.4615 | 1.5391 | 1.0003 | 0.000999 |
| TCD4ex | RTMIM | ITGB1 | PLAUR | 0.4615 | 1.1344 | 0.7979 | 0.000999 |
| TCD4ex | RTMIM | ITGB2 | ICAM1 | 0.7871 | 0.6002 | 0.6936 | 0.000999 |
| TCD4ex | RTMIM | ITGB2 | ITGAM | 0.7871 | 0.3355 | 0.5613 | 0.000999 |
| TCD4ex | RTMIM | ITGB2 | PLAUR | 0.7871 | 1.1344 | 0.9607 | 0.000999 |
| TCD4ex | RTMIM | LRPAP1 | LRP1 | 0.2809 | 0.5167 | 0.3988 | 0.000999 |
| TCD4ex | RTMIM | LTB | ITGB1 | 0.9655 | 0.4238 | 0.6947 | 0.000999 |
| TCD4ex | RTMIM | LTB | TNFRSF1A | 0.9655 | 0.5118 | 0.7386 | 0.000999 |
| TCD4ex | RTMIM | MIF | CD74 | 0.6999 | 4.5079 | 2.6039 | 0.000999 |
| TCD4ex | RTMIM | NMB | LPAR6 | 0.6391 | 0.4942 | 0.5667 | 0.000999 |
| TCD4ex | RTMIM | PTPRC | CD4 | 1.7493 | 0.6545 | 1.2019 | 0.000999 |
| TCD4ex | RTMIM | PTPRC | CD44 | 1.7493 | 0.8187 | 1.284 | 0.000999 |
| TCD4ex | RTMIM | RPS19 | C5AR1 | 3.6329 | 0.8768 | 2.2548 | 0.000999 |
| TCD4ex | RTMIM | RPS19 | CD4 | 3.6329 | 0.6545 | 2.1437 | 0.000999 |
| TCD4ex | RTMIM | RPS19 | ICAM1 | 3.6329 | 0.6002 | 2.1166 | 0.000999 |
| TCD4ex | RTMIM | SELL | CD4 | 0.5618 | 0.6545 | 0.6081 | 0.000999 |
| TCD4ex | RTMIM | TIMP1 | LRP1 | 0.3374 | 0.5167 | 0.4271 | 0.000999 |
| TCD4ex | RTMIM | ARF1 | CD4 | 0.6827 | 0.6545 | 0.6686 | 0.003 |
| TCD4ex | RTMIM | ANXA1 | CCR1 | 0.5174 | 0.2693 | 0.3934 | 0.00799 |
| TCD4reg | RTMIM | APOE | LRP1 | 0.2934 | 0.5167 | 0.4051 | 0.000999 |
| TCD4reg | RTMIM | APOE | SDC3 | 0.2934 | 0.286 | 0.2897 | 0.000999 |
| TCD4reg | RTMIM | ARF1 | CD4 | 0.7185 | 0.6545 | 0.6865 | 0.000999 |

| | | | | | | | |
|---|---|---|---|---|---|---|---|
| TCD4reg | RTMIM | CALM1 | TNFRSF1A | 1.534 | 0.5118 | 1.0229 | 0.000999 |
| TCD4reg | RTMIM | CALR | LRP1 | 0.6835 | 0.5167 | 0.6001 | 0.000999 |
| TCD4reg | RTMIM | CD2 | CD4 | 1.6566 | 0.6545 | 1.1556 | 0.000999 |
| TCD4reg | RTMIM | CD28 | CD4 | 0.313 | 0.6545 | 0.4837 | 0.000999 |
| TCD4reg | RTMIM | CD99 | CD99 | 0.6603 | 0.8717 | 0.766 | 0.000999 |
| TCD4reg | RTMIM | HMGB1 | LY96 | 1.2547 | 0.9322 | 1.0935 | 0.000999 |
| TCD4reg | RTMIM | HMGB1 | TLR2 | 1.2547 | 0.2837 | 0.7692 | 0.000999 |
| TCD4reg | RTMIM | HMGB2 | DAB2 | 0.616 | 1.5391 | 1.0776 | 0.000999 |
| TCD4reg | RTMIM | HSP90B1 | LRP1 | 0.4221 | 0.5167 | 0.4694 | 0.000999 |
| TCD4reg | RTMIM | ICAM3 | ITGB2 | 0.6663 | 0.862 | 0.7641 | 0.000999 |
| TCD4reg | RTMIM | IL16 | CD4 | 0.2528 | 0.6545 | 0.4536 | 0.000999 |
| TCD4reg | RTMIM | IL16 | FCGR1A | 0.2528 | 0.3257 | 0.2892 | 0.000999 |
| TCD4reg | RTMIM | ITGB1 | CD36 | 0.4001 | 0.3859 | 0.393 | 0.000999 |
| TCD4reg | RTMIM | ITGB1 | DAB2 | 0.4001 | 1.5391 | 0.9696 | 0.000999 |
| TCD4reg | RTMIM | ITGB1 | PLAUR | 0.4001 | 1.1344 | 0.7672 | 0.000999 |
| TCD4reg | RTMIM | ITGB2 | ICAM1 | 0.6238 | 0.6002 | 0.612 | 0.000999 |
| TCD4reg | RTMIM | ITGB2 | ITGAM | 0.6238 | 0.3355 | 0.4797 | 0.000999 |
| TCD4reg | RTMIM | ITGB2 | PLAUR | 0.6238 | 1.1344 | 0.8791 | 0.000999 |
| TCD4reg | RTMIM | LGALS3 | CLEC7A | 0.4492 | 0.4812 | 0.4652 | 0.000999 |
| TCD4reg | RTMIM | LGALS3 | COLEC12 | 0.4492 | 0.3221 | 0.3856 | 0.000999 |
| TCD4reg | RTMIM | LGALS3 | FCGR2A | 0.4492 | 1.172 | 0.8106 | 0.000999 |
| TCD4reg | RTMIM | LGALS3 | FCGR2B | 0.4492 | 0.6872 | 0.5682 | 0.000999 |
| TCD4reg | RTMIM | LGALS3 | TFRC | 0.4492 | 0.6023 | 0.5257 | 0.000999 |
| TCD4reg | RTMIM | LGALS3 | TGFBR2 | 0.4492 | 0.3722 | 0.4107 | 0.000999 |
| TCD4reg | RTMIM | LRPAP1 | LRP1 | 0.3101 | 0.5167 | 0.4134 | 0.000999 |
| TCD4reg | RTMIM | LTB | ITGB1 | 2.242 | 0.4238 | 1.3329 | 0.000999 |
| TCD4reg | RTMIM | LTB | TNFRSF1A | 2.242 | 0.5118 | 1.3769 | 0.000999 |
| TCD4reg | RTMIM | MIF | CD74 | 0.6295 | 4.5079 | 2.5687 | 0.000999 |
| TCD4reg | RTMIM | PSAP | LRP1 | 0.5938 | 0.5167 | 0.5553 | 0.000999 |
| TCD4reg | RTMIM | PTPRC | CD4 | 1.6603 | 0.6545 | 1.1574 | 0.000999 |

| | | | | | | | |
|---|---|---|---|---|---|---|---|
| TCD4reg | RTMIM | RPS19 | C5AR1 | 3.6897 | 0.8768 | 2.2832 | 0.000999 |
| TCD4reg | RTMIM | RPS19 | CD4 | 3.6897 | 0.6545 | 2.1721 | 0.000999 |
| TCD4reg | RTMIM | RPS19 | ICAM1 | 3.6897 | 0.6002 | 2.1449 | 0.000999 |
| TCD4reg | RTMIM | RTN4 | CD99 | 0.3528 | 0.8717 | 0.6122 | 0.000999 |
| TCD4reg | RTMIM | SELL | CD4 | 0.7062 | 0.6545 | 0.6803 | 0.000999 |
| TCD4reg | RTMIM | TIMP1 | LRP1 | 0.4362 | 0.5167 | 0.4765 | 0.000999 |
| TCD4reg | RTMIM | ITGB1 | CD46 | 0.4001 | 0.2644 | 0.3323 | 0.002 |
| TCD8em | RTMIM | ANXA1 | C3AR1 | 1.0693 | 0.6118 | 0.8406 | 0.000999 |
| TCD8em | RTMIM | ANXA1 | C5AR1 | 1.0693 | 0.8768 | 0.973 | 0.000999 |
| TCD8em | RTMIM | ANXA1 | CCR1 | 1.0693 | 0.2693 | 0.6693 | 0.000999 |
| TCD8em | RTMIM | ANXA1 | CD4 | 1.0693 | 0.6545 | 0.8619 | 0.000999 |
| TCD8em | RTMIM | ANXA1 | FPR3 | 1.0693 | 0.511 | 0.7901 | 0.000999 |
| TCD8em | RTMIM | APOE | LRP1 | 0.2791 | 0.5167 | 0.3979 | 0.000999 |
| TCD8em | RTMIM | APOE | SDC3 | 0.2791 | 0.286 | 0.2826 | 0.000999 |
| TCD8em | RTMIM | ARF1 | CD4 | 0.6684 | 0.6545 | 0.6614 | 0.000999 |
| TCD8em | RTMIM | CALM1 | TNFRSF1A | 1.5262 | 0.5118 | 1.019 | 0.000999 |
| TCD8em | RTMIM | CALM1 | TNFRSF1B | 1.5262 | 0.5749 | 1.0505 | 0.000999 |
| TCD8em | RTMIM | CALR | LRP1 | 0.7141 | 0.5167 | 0.6154 | 0.000999 |
| TCD8em | RTMIM | CD2 | CD4 | 1.5961 | 0.6545 | 1.1253 | 0.000999 |
| TCD8em | RTMIM | HLA-DMA | CD4 | 0.2969 | 0.6545 | 0.4757 | 0.000999 |
| TCD8em | RTMIM | HLA-DRA | CD4 | 0.952 | 0.6545 | 0.8033 | 0.000999 |
| TCD8em | RTMIM | HMGB1 | LY96 | 1.3636 | 0.9322 | 1.1479 | 0.000999 |
| TCD8em | RTMIM | HMGB1 | TLR2 | 1.3636 | 0.2837 | 0.8236 | 0.000999 |
| TCD8em | RTMIM | HMGB2 | DAB2 | 0.7212 | 1.5391 | 1.1301 | 0.000999 |
| TCD8em | RTMIM | HSP90B1 | ITGB1 | 0.471 | 0.4238 | 0.4474 | 0.000999 |
| TCD8em | RTMIM | HSP90B1 | LRP1 | 0.471 | 0.5167 | 0.4939 | 0.000999 |
| TCD8em | RTMIM | HSP90B1 | TNFRSF1B | 0.471 | 0.5749 | 0.523 | 0.000999 |
| TCD8em | RTMIM | IFNG | IFNGR1 | 0.3967 | 0.428 | 0.4123 | 0.000999 |
| TCD8em | RTMIM | IFNG | IFNGR2 | 0.3967 | 0.5349 | 0.4658 | 0.000999 |
| TCD8em | RTMIM | IL16 | CD4 | 0.4085 | 0.6545 | 0.5315 | 0.000999 |

| | | | | | | | |
|---|---|---|---|---|---|---|---|
| TCD8em | RTMIM | IL16 | FCGR1A | 0.4085 | 0.3257 | 0.3671 | 0.000999 |
| TCD8em | RTMIM | ITGA4 | TFRC | 0.2823 | 0.6023 | 0.4423 | 0.000999 |
| TCD8em | RTMIM | ITGAL | ICAM1 | 0.3162 | 0.6002 | 0.4582 | 0.000999 |
| TCD8em | RTMIM | ITGB1 | CD36 | 0.2557 | 0.3859 | 0.3208 | 0.000999 |
| TCD8em | RTMIM | ITGB1 | CD46 | 0.2557 | 0.2644 | 0.2601 | 0.000999 |
| TCD8em | RTMIM | ITGB1 | DAB2 | 0.2557 | 1.5391 | 0.8974 | 0.000999 |
| TCD8em | RTMIM | ITGB1 | PLAUR | 0.2557 | 1.1344 | 0.695 | 0.000999 |
| TCD8em | RTMIM | ITGB2 | ICAM1 | 1.0018 | 0.6002 | 0.801 | 0.000999 |
| TCD8em | RTMIM | ITGB2 | ITGAM | 1.0018 | 0.3355 | 0.6687 | 0.000999 |
| TCD8em | RTMIM | ITGB2 | PLAUR | 1.0018 | 1.1344 | 1.0681 | 0.000999 |
| TCD8em | RTMIM | LGALS3 | CLEC7A | 0.2689 | 0.4812 | 0.3751 | 0.000999 |
| TCD8em | RTMIM | LGALS3 | COLEC12 | 0.2689 | 0.3221 | 0.2955 | 0.000999 |
| TCD8em | RTMIM | LGALS3 | FCGR2A | 0.2689 | 1.172 | 0.7205 | 0.000999 |
| TCD8em | RTMIM | LGALS3 | FCGR2B | 0.2689 | 0.6872 | 0.4781 | 0.000999 |
| TCD8em | RTMIM | LGALS3 | ITGB1 | 0.2689 | 0.4238 | 0.3464 | 0.000999 |
| TCD8em | RTMIM | LGALS3 | TFRC | 0.2689 | 0.6023 | 0.4356 | 0.000999 |
| TCD8em | RTMIM | LGALS3 | TGFBR2 | 0.2689 | 0.3722 | 0.3205 | 0.000999 |
| TCD8em | RTMIM | LRPAP1 | LRP1 | 0.2719 | 0.5167 | 0.3943 | 0.000999 |
| TCD8em | RTMIM | LTB | ITGB1 | 0.6352 | 0.4238 | 0.5295 | 0.000999 |
| TCD8em | RTMIM | LTB | TNFRSF1A | 0.6352 | 0.5118 | 0.5735 | 0.000999 |
| TCD8em | RTMIM | MIF | CD74 | 0.6063 | 4.5079 | 2.5571 | 0.000999 |
| TCD8em | RTMIM | PSAP | LRP1 | 0.5012 | 0.5167 | 0.509 | 0.000999 |
| TCD8em | RTMIM | PTPRC | CD4 | 1.7912 | 0.6545 | 1.2228 | 0.000999 |
| TCD8em | RTMIM | RPS19 | C5AR1 | 3.536 | 0.8768 | 2.2064 | 0.000999 |
| TCD8em | RTMIM | RPS19 | CD4 | 3.536 | 0.6545 | 2.0952 | 0.000999 |
| TCD8em | RTMIM | RPS19 | ICAM1 | 3.536 | 0.6002 | 2.0681 | 0.000999 |
| TCD8em | RTMIM | SELL | CD4 | 0.258 | 0.6545 | 0.4562 | 0.000999 |
| TCD8em | RTMIM | SPN | SIGLEC1 | 0.2603 | 0.3312 | 0.2958 | 0.000999 |
| TCD8em | RTMIM | SPP1 | ITGB1 | 0.3608 | 0.4238 | 0.3923 | 0.000999 |
| TCD8em | RTMIM | TIMP1 | LRP1 | 0.2667 | 0.5167 | 0.3917 | 0.000999 |

| | | | | | | | |
|---|---|---|---|---|---|---|---|
| TCD8em | RTMIM | TNF | NRP1 | 0.2667 | 0.4884 | 0.3776 | 0.000999 |
| TCD8em | RTMIM | TNF | TNFRSF1A | 0.2667 | 0.5118 | 0.3892 | 0.000999 |
| TCD8em | RTMIM | TNF | TNFRSF1B | 0.2667 | 0.5749 | 0.4208 | 0.000999 |
| TCD8em | RTMIM | XCL2 | LPAR6 | 0.295 | 0.4942 | 0.3946 | 0.000999 |
| TCD8em | RTMIM | IFNG | PTPN6 | 0.3967 | 0.3038 | 0.3502 | 0.002 |
| TCD8ex | RTMIM | AIMP1 | RRBP1 | 0.2508 | 0.2584 | 0.2546 | 0.000999 |
| TCD8ex | RTMIM | ANXA1 | C3AR1 | 0.5645 | 0.6118 | 0.5882 | 0.000999 |
| TCD8ex | RTMIM | ANXA1 | C5AR1 | 0.5645 | 0.8768 | 0.7206 | 0.000999 |
| TCD8ex | RTMIM | ANXA1 | CCR1 | 0.5645 | 0.2693 | 0.4169 | 0.000999 |
| TCD8ex | RTMIM | ANXA1 | CD4 | 0.5645 | 0.6545 | 0.6095 | 0.000999 |
| TCD8ex | RTMIM | ANXA1 | FPR3 | 0.5645 | 0.511 | 0.5377 | 0.000999 |
| TCD8ex | RTMIM | APOE | LRP1 | 0.2814 | 0.5167 | 0.399 | 0.000999 |
| TCD8ex | RTMIM | APOE | SDC3 | 0.2814 | 0.286 | 0.2837 | 0.000999 |
| TCD8ex | RTMIM | AREG | ICAM1 | 0.2758 | 0.6002 | 0.438 | 0.000999 |
| TCD8ex | RTMIM | ARF1 | CD4 | 0.7835 | 0.6545 | 0.719 | 0.000999 |
| TCD8ex | RTMIM | CALM1 | TNFRSF1A | 1.4627 | 0.5118 | 0.9872 | 0.000999 |
| TCD8ex | RTMIM | CALR | LRP1 | 0.9634 | 0.5167 | 0.7401 | 0.000999 |
| TCD8ex | RTMIM | CD2 | CD4 | 2.0502 | 0.6545 | 1.3523 | 0.000999 |
| TCD8ex | RTMIM | CXCL13 | C3AR1 | 1.7749 | 0.6118 | 1.1934 | 0.000999 |
| TCD8ex | RTMIM | CXCL13 | C5AR1 | 1.7749 | 0.8768 | 1.3258 | 0.000999 |
| TCD8ex | RTMIM | CXCL13 | CCR1 | 1.7749 | 0.2693 | 1.0221 | 0.000999 |
| TCD8ex | RTMIM | FASLG | TNFRSF1A | 0.2613 | 0.5118 | 0.3865 | 0.000999 |
| TCD8ex | RTMIM | HLA-DMA | CD4 | 0.3991 | 0.6545 | 0.5268 | 0.000999 |
| TCD8ex | RTMIM | HLA-DRA | CD4 | 1.247 | 0.6545 | 0.9507 | 0.000999 |
| TCD8ex | RTMIM | HMGB1 | LY96 | 1.5414 | 0.9322 | 1.2368 | 0.000999 |
| TCD8ex | RTMIM | HMGB1 | TLR2 | 1.5414 | 0.2837 | 0.9126 | 0.000999 |
| TCD8ex | RTMIM | HMGB2 | DAB2 | 0.85 | 1.5391 | 1.1946 | 0.000999 |
| TCD8ex | RTMIM | HSP90B1 | ITGB1 | 0.5666 | 0.4238 | 0.4952 | 0.000999 |
| TCD8ex | RTMIM | HSP90B1 | LRP1 | 0.5666 | 0.5167 | 0.5417 | 0.000999 |
| TCD8ex | RTMIM | ICAM3 | ITGB2 | 0.5405 | 0.862 | 0.7012 | 0.000999 |

| | | | | | | | |
|---|---|---|---|---|---|---|---|
| TCD8ex | RTMIM | IFNG | IFNGR1 | 0.8575 | 0.428 | 0.6428 | 0.000999 |
| TCD8ex | RTMIM | IFNG | IFNGR2 | 0.8575 | 0.5349 | 0.6962 | 0.000999 |
| TCD8ex | RTMIM | ITGA4 | TFRC | 0.4432 | 0.6023 | 0.5228 | 0.000999 |
| TCD8ex | RTMIM | ITGB1 | CD36 | 0.2995 | 0.3859 | 0.3427 | 0.000999 |
| TCD8ex | RTMIM | ITGB1 | DAB2 | 0.2995 | 1.5391 | 0.9193 | 0.000999 |
| TCD8ex | RTMIM | ITGB1 | PLAUR | 0.2995 | 1.1344 | 0.7169 | 0.000999 |
| TCD8ex | RTMIM | ITGB2 | ICAM1 | 0.7635 | 0.6002 | 0.6818 | 0.000999 |
| TCD8ex | RTMIM | ITGB2 | ITGAM | 0.7635 | 0.3355 | 0.5495 | 0.000999 |
| TCD8ex | RTMIM | ITGB2 | PLAUR | 0.7635 | 1.1344 | 0.9489 | 0.000999 |
| TCD8ex | RTMIM | LGALS3 | CLEC7A | 0.3824 | 0.4812 | 0.4318 | 0.000999 |
| TCD8ex | RTMIM | LGALS3 | COLEC12 | 0.3824 | 0.3221 | 0.3522 | 0.000999 |
| TCD8ex | RTMIM | LGALS3 | FCGR2A | 0.3824 | 1.172 | 0.7772 | 0.000999 |
| TCD8ex | RTMIM | LGALS3 | FCGR2B | 0.3824 | 0.6872 | 0.5348 | 0.000999 |
| TCD8ex | RTMIM | LGALS3 | ITGB1 | 0.3824 | 0.4238 | 0.4031 | 0.000999 |
| TCD8ex | RTMIM | LGALS3 | TFRC | 0.3824 | 0.6023 | 0.4923 | 0.000999 |
| TCD8ex | RTMIM | LGALS3 | TGFBR2 | 0.3824 | 0.3722 | 0.3773 | 0.000999 |
| TCD8ex | RTMIM | LRPAP1 | LRP1 | 0.2971 | 0.5167 | 0.4069 | 0.000999 |
| TCD8ex | RTMIM | LTB | ITGB1 | 0.4576 | 0.4238 | 0.4407 | 0.000999 |
| TCD8ex | RTMIM | LTB | TNFRSF1A | 0.4576 | 0.5118 | 0.4847 | 0.000999 |
| TCD8ex | RTMIM | MIF | CD74 | 0.749 | 4.5079 | 2.6285 | 0.000999 |
| TCD8ex | RTMIM | PSAP | LRP1 | 0.5016 | 0.5167 | 0.5092 | 0.000999 |
| TCD8ex | RTMIM | PTPRC | CD4 | 1.6762 | 0.6545 | 1.1653 | 0.000999 |
| TCD8ex | RTMIM | PTPRC | CD44 | 1.6762 | 0.8187 | 1.2475 | 0.000999 |
| TCD8ex | RTMIM | RPS19 | C5AR1 | 3.8353 | 0.8768 | 2.356 | 0.000999 |
| TCD8ex | RTMIM | RPS19 | CD4 | 3.8353 | 0.6545 | 2.2449 | 0.000999 |
| TCD8ex | RTMIM | RPS19 | ICAM1 | 3.8353 | 0.6002 | 2.2177 | 0.000999 |
| TCD8ex | RTMIM | SPN | SIGLEC1 | 0.3486 | 0.3312 | 0.3399 | 0.000999 |
| TCD8ex | RTMIM | SPP1 | ITGB1 | 0.2978 | 0.4238 | 0.3608 | 0.000999 |
| TCD8ex | RTMIM | VCAM1 | ITGB1 | 0.4462 | 0.4238 | 0.435 | 0.000999 |
| TCD8ex | RTMIM | VCAM1 | ITGB2 | 0.4462 | 0.862 | 0.6541 | 0.000999 |

| | | | | | | | |
|---|---|---|---|---|---|---|---|
| TCD8ex | RTMIM | VCAM1 | TFRC | 0.4462 | 0.6023 | 0.5243 | 0.000999 |
| TCD8ex | RTMIM | XCL2 | LPAR6 | 0.3157 | 0.4942 | 0.405 | 0.000999 |
| TGD | RTMIM | AREG | ICAM1 | 0.4756 | 0.6002 | 0.5379 | 0.000999 |
| TGD | RTMIM | ARF1 | CD4 | 0.9453 | 0.6545 | 0.7999 | 0.000999 |
| TGD | RTMIM | ARF1 | CD44 | 0.9453 | 0.8187 | 0.882 | 0.000999 |
| TGD | RTMIM | CALM1 | RPSA | 1.7587 | 1.7137 | 1.7362 | 0.000999 |
| TGD | RTMIM | CALM1 | TNFRSF1A | 1.7587 | 0.5118 | 1.1352 | 0.000999 |
| TGD | RTMIM | CALM1 | TNFRSF1B | 1.7587 | 0.5749 | 1.1668 | 0.000999 |
| TGD | RTMIM | CALR | LRP1 | 0.9627 | 0.5167 | 0.7397 | 0.000999 |
| TGD | RTMIM | CD2 | CD4 | 1.4522 | 0.6545 | 1.0533 | 0.000999 |
| TGD | RTMIM | CD2 | PTPRC | 1.4522 | 0.5365 | 0.9943 | 0.000999 |
| TGD | RTMIM | CD99 | CD99 | 1.7154 | 0.8717 | 1.2936 | 0.000999 |
| TGD | RTMIM | HMGB1 | LY96 | 1.7525 | 0.9322 | 1.3423 | 0.000999 |
| TGD | RTMIM | HMGB1 | TLR2 | 1.7525 | 0.2837 | 1.0181 | 0.000999 |
| TGD | RTMIM | HMGB2 | DAB2 | 0.3275 | 1.5391 | 0.9333 | 0.000999 |
| TGD | RTMIM | HSP90B1 | LRP1 | 0.9338 | 0.5167 | 0.7253 | 0.000999 |
| TGD | RTMIM | ICAM3 | ITGB2 | 0.8816 | 0.862 | 0.8718 | 0.000999 |
| TGD | RTMIM | IL16 | CD4 | 0.9357 | 0.6545 | 0.7951 | 0.000999 |
| TGD | RTMIM | IL16 | FCGR1A | 0.9357 | 0.3257 | 0.6307 | 0.000999 |
| TGD | RTMIM | ITGA4 | CD44 | 0.3968 | 0.8187 | 0.6078 | 0.000999 |
| TGD | RTMIM | ITGA4 | CXCR4 | 0.3968 | 0.8828 | 0.6398 | 0.000999 |
| TGD | RTMIM | ITGA4 | TFRC | 0.3968 | 0.6023 | 0.4996 | 0.000999 |
| TGD | RTMIM | ITGAL | ICAM1 | 0.3731 | 0.6002 | 0.4867 | 0.000999 |
| TGD | RTMIM | ITGB1 | DAB2 | 0.3657 | 1.5391 | 0.9524 | 0.000999 |
| TGD | RTMIM | ITGB2 | ICAM1 | 1.4444 | 0.6002 | 1.0223 | 0.000999 |
| TGD | RTMIM | ITGB2 | ITGAM | 1.4444 | 0.3355 | 0.89 | 0.000999 |
| TGD | RTMIM | ITGB2 | PLAUR | 1.4444 | 1.1344 | 1.2894 | 0.000999 |
| TGD | RTMIM | LTB | ITGB1 | 1.6829 | 0.4238 | 1.0534 | 0.000999 |
| TGD | RTMIM | LTB | TNFRSF1A | 1.6829 | 0.5118 | 1.0973 | 0.000999 |
| TGD | RTMIM | MIF | CD74 | 1.6335 | 4.5079 | 3.0707 | 0.000999 |

| | | | | | | | |
|---|---|---|---|---|---|---|---|
| TGD | RTMIM | MIF | TNFRSF14 | 1.6335 | 0.2649 | 0.9492 | 0.000999 |
| TGD | RTMIM | PTPRC | CD4 | 1.8653 | 0.6545 | 1.2599 | 0.000999 |
| TGD | RTMIM | PTPRC | CD44 | 1.8653 | 0.8187 | 1.342 | 0.000999 |
| TGD | RTMIM | PTPRC | CXCR4 | 1.8653 | 0.8828 | 1.3741 | 0.000999 |
| TGD | RTMIM | RPS19 | C5AR1 | 3.8902 | 0.8768 | 2.3835 | 0.000999 |
| TGD | RTMIM | RPS19 | CD4 | 3.8902 | 0.6545 | 2.2723 | 0.000999 |
| TGD | RTMIM | RPS19 | ICAM1 | 3.8902 | 0.6002 | 2.2452 | 0.000999 |
| TGD | RTMIM | RPS19 | RPSA | 3.8902 | 1.7137 | 2.8019 | 0.000999 |
| TGD | RTMIM | SELL | CD4 | 0.3555 | 0.6545 | 0.505 | 0.000999 |
| TGD | RTMIM | SEMA4D | PTPRC | 0.5658 | 0.5365 | 0.5511 | 0.000999 |
| TGD | RTMIM | SPN | SIGLEC1 | 0.5644 | 0.3312 | 0.4478 | 0.000999 |
| TGD | RTMIM | TGFB1 | ITGB1 | 0.7703 | 0.4238 | 0.5971 | 0.000999 |
| TGD | RTMIM | TGFB1 | TGFBR2 | 0.7703 | 0.3722 | 0.5713 | 0.000999 |
| TGD | RTMIM | TGFB1 | TLR2 | 0.7703 | 0.2837 | 0.527 | 0.000999 |
| TGD | RTMIM | XCL1 | LPAR6 | 1.5822 | 0.4942 | 1.0382 | 0.000999 |
| TGD | RTMIM | XCL2 | LPAR6 | 0.4864 | 0.4942 | 0.4903 | 0.000999 |
| TGD | RTMIM | HSP90B1 | TNFRSF1B | 0.9338 | 0.5749 | 0.7544 | 0.002 |
| TGD | RTMIM | ITGB1 | PLAUR | 0.3657 | 1.1344 | 0.75 | 0.002 |
| TGD | RTMIM | ITGA4 | RPSA | 0.3968 | 1.7137 | 1.0553 | 0.005 |
| TGD | RTMIM | HSP90B1 | ITGB1 | 0.9338 | 0.4238 | 0.6788 | 0.00799 |
| TGD | RTMIM | AIMP1 | RRBP1 | 0.2684 | 0.2584 | 0.2634 | 0.00999 |

**Supplementary Table 8: Communication results between macrophages and lymphocytes subpopulations, and malignant cells**

**RTM_IM vs. malignant cells**

| celltype_01 | celltype_02 | ligand | receptor | ligand_mean_expression | receptor_mean_expression | ligand_receptor_mean_expression | pvalue |
|---|---|---|---|---|---|---|---|
| RTMIM | breasttumor | ANXA1 | CD44 | 1.1309 | 0.3247 | 0.7278 | 0.000999 |
| RTMIM | breasttumor | APOE | SCARB1 | 1.1772 | 0.2628 | 0.72 | 0.000999 |
| RTMIM | breasttumor | APOE | SDC2 | 1.1772 | 1.071 | 1.1241 | 0.000999 |
| RTMIM | breasttumor | CALM1 | MYLK | 1.036 | 0.4064 | 0.7212 | 0.000999 |
| RTMIM | breasttumor | CALM1 | RPSA | 1.036 | 2.358 | 1.697 | 0.000999 |
| RTMIM | breasttumor | CALM2 | MYLK | 1.9125 | 0.4064 | 1.1594 | 0.000999 |
| RTMIM | breasttumor | CALM3 | MYLK | 0.6155 | 0.4064 | 0.5109 | 0.000999 |
| RTMIM | breasttumor | CD99 | CD99 | 0.87 | 0.3871 | 0.6285 | 0.000999 |
| RTMIM | breasttumor | COL1A1 | ITGB1 | 0.4843 | 0.2939 | 0.3891 | 0.000999 |
| RTMIM | breasttumor | FN1 | ITGB1 | 0.3859 | 0.2939 | 0.3399 | 0.000999 |
| RTMIM | breasttumor | FN1 | RPSA | 0.3859 | 2.358 | 1.372 | 0.000999 |
| RTMIM | breasttumor | FN1 | SDC2 | 0.3859 | 1.071 | 0.7285 | 0.000999 |
| RTMIM | breasttumor | GRN | TNFRSF1A | 1.4421 | 0.2536 | 0.8478 | 0.000999 |
| RTMIM | breasttumor | HBEGF | CD44 | 0.6122 | 0.3247 | 0.4685 | 0.000999 |
| RTMIM | breasttumor | HLA-DRA | SCARB1 | 3.9396 | 0.2628 | 2.1012 | 0.000999 |
| RTMIM | breasttumor | IGF1 | RACK1 | 0.4606 | 2.6334 | 1.547 | 0.000999 |
| RTMIM | breasttumor | ITGB1 | CD151 | 0.4226 | 0.3569 | 0.3897 | 0.000999 |
| RTMIM | breasttumor | ITGB1 | RACK1 | 0.4226 | 2.6334 | 1.528 | 0.000999 |
| RTMIM | breasttumor | PTPRC | CD44 | 0.5369 | 0.3247 | 0.4308 | 0.000999 |
| RTMIM | breasttumor | RTN4 | CD44 | 0.9264 | 0.3247 | 0.6256 | 0.000999 |
| RTMIM | breasttumor | RTN4 | CD99 | 0.9264 | 0.3871 | 0.6567 | 0.000999 |
| RTMIM | breasttumor | SERPINA1 | SDC2 | 0.3379 | 1.071 | 0.7045 | 0.000999 |
| RTMIM | breasttumor | SPP1 | CD44 | 0.6105 | 0.3247 | 0.4676 | 0.000999 |
| RTMIM | breasttumor | SPP1 | ITGB1 | 0.6105 | 0.2939 | 0.4522 | 0.000999 |
| RTMIM | breasttumor | TGFB1 | ITGB1 | 0.4158 | 0.2939 | 0.3549 | 0.000999 |
| RTMIM | breasttumor | TGFB1 | SDC2 | 0.4158 | 1.071 | 0.7434 | 0.000999 |
| RTMIM | breasttumor | THBS1 | ITGB1 | 0.3968 | 0.2939 | 0.3453 | 0.000999 |

| | | | | | | | |
|---|---|---|---|---|---|---|---|
| RTMIM | breasttumor | TNF | CALM1 | 0.3681 | 0.9015 | 0.6348 | 0.000999 |
| RTMIM | breasttumor | TNF | LTBR | 0.3681 | 0.2857 | 0.3269 | 0.000999 |
| RTMIM | breasttumor | TNF | RACK1 | 0.3681 | 2.6334 | 1.5008 | 0.000999 |
| RTMIM | breasttumor | TNF | TNFRSF1A | 0.3681 | 0.2536 | 0.3109 | 0.000999 |
| RTMIM | breasttumor | TNFSF13B | ITGB1 | 0.4496 | 0.2939 | 0.3717 | 0.000999 |
| RTMIM | colorectaltumor | APOE | SDC1 | 1.1772 | 0.2575 | 0.7173 | 0.000999 |
| RTMIM | colorectaltumor | CALM1 | RPSA | 1.036 | 2.251 | 1.6435 | 0.000999 |
| RTMIM | colorectaltumor | COL1A1 | SDC1 | 0.4843 | 0.2575 | 0.3709 | 0.000999 |
| RTMIM | colorectaltumor | FN1 | RPSA | 0.3859 | 2.251 | 1.3185 | 0.000999 |
| RTMIM | colorectaltumor | FN1 | SDC1 | 0.3859 | 0.2575 | 0.3217 | 0.000999 |
| RTMIM | colorectaltumor | IGF1 | RACK1 | 0.4606 | 2.7146 | 1.5876 | 0.000999 |
| RTMIM | colorectaltumor | ITGB1 | CD151 | 0.4226 | 0.6439 | 0.5333 | 0.000999 |
| RTMIM | colorectaltumor | ITGB1 | CDH17 | 0.4226 | 0.254 | 0.3383 | 0.000999 |
| RTMIM | colorectaltumor | ITGB1 | RACK1 | 0.4226 | 2.7146 | 1.5686 | 0.000999 |
| RTMIM | colorectaltumor | ITGB1 | SDC1 | 0.4226 | 0.2575 | 0.34 | 0.000999 |
| RTMIM | colorectaltumor | PECAM1 | SDC1 | 0.3342 | 0.2575 | 0.2958 | 0.000999 |
| RTMIM | colorectaltumor | RPS19 | RPSA | 3.2576 | 2.251 | 2.7543 | 0.000999 |
| RTMIM | colorectaltumor | SPP1 | SDC1 | 0.6105 | 0.2575 | 0.434 | 0.000999 |
| RTMIM | colorectaltumor | THBS1 | SDC1 | 0.3968 | 0.2575 | 0.3271 | 0.000999 |
| RTMIM | colorectaltumor | TNF | CALM1 | 0.3681 | 1.1676 | 0.7679 | 0.000999 |
| RTMIM | colorectaltumor | TNF | RACK1 | 0.3681 | 2.7146 | 1.5414 | 0.000999 |
| RTMIM | livertumor | A2M | TNFRSF14 | 1.0949 | 0.2788 | 0.6869 | 0.000999 |
| RTMIM | livertumor | ADM | RAMP1 | 0.2709 | 0.4499 | 0.3604 | 0.000999 |
| RTMIM | livertumor | ANXA1 | CD44 | 1.1309 | 0.8016 | 0.9663 | 0.000999 |
| RTMIM | livertumor | ANXA1 | MET | 1.1309 | 0.2983 | 0.7146 | 0.000999 |
| RTMIM | livertumor | ANXA1 | PTPRF | 1.1309 | 0.384 | 0.7575 | 0.000999 |
| RTMIM | livertumor | APOE | SDC1 | 1.1772 | 0.5354 | 0.8563 | 0.000999 |
| RTMIM | livertumor | APOE | SDC4 | 1.1772 | 0.6068 | 0.892 | 0.000999 |
| RTMIM | livertumor | ARF1 | INSR | 0.5732 | 0.3039 | 0.4386 | 0.000999 |
| RTMIM | livertumor | CALM1 | INSR | 1.036 | 0.3039 | 0.67 | 0.000999 |

| | | | | | | | |
|---|---|---|---|---|---|---|---|
| RTMIM | livertumor | CALM1 | RPSA | 1.036 | 2.6317 | 1.8339 | 0.000999 |
| RTMIM | livertumor | CALR | ITGA3 | 0.7678 | 0.3913 | 0.5796 | 0.000999 |
| RTMIM | livertumor | CD99 | EPHB4 | 0.87 | 0.4535 | 0.6617 | 0.000999 |
| RTMIM | livertumor | CD99 | INSR | 0.87 | 0.3039 | 0.587 | 0.000999 |
| RTMIM | livertumor | COL1A1 | ITGA2 | 0.4843 | 0.3983 | 0.4413 | 0.000999 |
| RTMIM | livertumor | COL1A1 | ITGA3 | 0.4843 | 0.3913 | 0.4378 | 0.000999 |
| RTMIM | livertumor | COL1A1 | ITGB1 | 0.4843 | 1.6044 | 1.0444 | 0.000999 |
| RTMIM | livertumor | COL1A1 | SDC1 | 0.4843 | 0.5354 | 0.5099 | 0.000999 |
| RTMIM | livertumor | FN1 | ITGA2 | 0.3859 | 0.3983 | 0.3921 | 0.000999 |
| RTMIM | livertumor | FN1 | ITGA3 | 0.3859 | 0.3913 | 0.3886 | 0.000999 |
| RTMIM | livertumor | FN1 | ITGB1 | 0.3859 | 1.6044 | 0.9952 | 0.000999 |
| RTMIM | livertumor | FN1 | RPSA | 0.3859 | 2.6317 | 1.5088 | 0.000999 |
| RTMIM | livertumor | FN1 | SDC1 | 0.3859 | 0.5354 | 0.4607 | 0.000999 |
| RTMIM | livertumor | HBEGF | ERBB3 | 0.6122 | 0.2644 | 0.4383 | 0.000999 |
| RTMIM | livertumor | HLA-A | ERBB3 | 2.313 | 0.2644 | 1.2887 | 0.000999 |
| RTMIM | livertumor | HLA-DRA | EPHB4 | 3.9396 | 0.4535 | 2.1965 | 0.000999 |
| RTMIM | livertumor | HLA-E | EPHB4 | 1.7651 | 0.4535 | 1.1093 | 0.000999 |
| RTMIM | livertumor | HLA-E | ITGA6 | 1.7651 | 0.5211 | 1.1431 | 0.000999 |
| RTMIM | livertumor | HLA-E | PLXNB2 | 1.7651 | 0.265 | 1.015 | 0.000999 |
| RTMIM | livertumor | HMGB2 | CDH1 | 0.351 | 0.4896 | 0.4203 | 0.000999 |
| RTMIM | livertumor | HMGB2 | DSG2 | 0.351 | 0.7479 | 0.5495 | 0.000999 |
| RTMIM | livertumor | HSP90B1 | ITGB1 | 0.7617 | 1.6044 | 1.183 | 0.000999 |
| RTMIM | livertumor | IGF1 | INSR | 0.4606 | 0.3039 | 0.3823 | 0.000999 |
| RTMIM | livertumor | IGF1 | RACK1 | 0.4606 | 2.7655 | 1.6131 | 0.000999 |
| RTMIM | livertumor | ITGB1 | CD151 | 0.4226 | 1.1085 | 0.7655 | 0.000999 |
| RTMIM | livertumor | ITGB1 | ITGA6 | 0.4226 | 0.5211 | 0.4718 | 0.000999 |
| RTMIM | livertumor | ITGB1 | SDC1 | 0.4226 | 0.5354 | 0.479 | 0.000999 |
| RTMIM | livertumor | ITGB1 | VTN | 0.4226 | 0.7399 | 0.5812 | 0.000999 |
| RTMIM | livertumor | LGALS3 | ITGA2 | 0.7006 | 0.3983 | 0.5494 | 0.000999 |
| RTMIM | livertumor | LGALS3 | ITGB1 | 0.7006 | 1.6044 | 1.1525 | 0.000999 |

| | | | | | | | |
|---|---|---|---|---|---|---|---|
| RTMIM | livertumor | LGALS3 | ITGB4 | 0.7006 | 0.5518 | 0.6262 | 0.000999 |
| RTMIM | livertumor | NAMPT | INSR | 1.2179 | 0.3039 | 0.7609 | 0.000999 |
| RTMIM | livertumor | PECAM1 | SDC1 | 0.3342 | 0.5354 | 0.4348 | 0.000999 |
| RTMIM | livertumor | PECAM1 | SDC4 | 0.3342 | 0.6068 | 0.4705 | 0.000999 |
| RTMIM | livertumor | PTPRC | INSR | 0.5369 | 0.3039 | 0.4204 | 0.000999 |
| RTMIM | livertumor | PTPRC | MET | 0.5369 | 0.2983 | 0.4176 | 0.000999 |
| RTMIM | livertumor | RPS19 | RPSA | 3.2576 | 2.6317 | 2.9446 | 0.000999 |
| RTMIM | livertumor | SPP1 | ITGB1 | 0.6105 | 1.6044 | 1.1075 | 0.000999 |
| RTMIM | livertumor | SPP1 | SDC1 | 0.6105 | 0.5354 | 0.573 | 0.000999 |
| RTMIM | livertumor | TGFB1 | ITGB1 | 0.4158 | 1.6044 | 1.0101 | 0.000999 |
| RTMIM | livertumor | TGFB1 | VTN | 0.4158 | 0.7399 | 0.5779 | 0.000999 |
| RTMIM | livertumor | THBS1 | CD47 | 0.3968 | 0.5289 | 0.4628 | 0.000999 |
| RTMIM | livertumor | THBS1 | ITGA3 | 0.3968 | 0.3913 | 0.394 | 0.000999 |
| RTMIM | livertumor | THBS1 | ITGB1 | 0.3968 | 1.6044 | 1.0006 | 0.000999 |
| RTMIM | livertumor | THBS1 | SDC1 | 0.3968 | 0.5354 | 0.4661 | 0.000999 |
| RTMIM | livertumor | TNF | LTBR | 0.3681 | 0.2777 | 0.3229 | 0.000999 |
| RTMIM | livertumor | TNF | RACK1 | 0.3681 | 2.7655 | 1.5668 | 0.000999 |
| RTMIM | livertumor | TNFSF13B | ITGB1 | 0.4496 | 1.6044 | 1.027 | 0.000999 |
| RTMIM | livertumor | VEGFA | VTN | 0.4114 | 0.7399 | 0.5757 | 0.000999 |
| RTMIM | livertumor | HBEGF | CD44 | 0.6122 | 0.8016 | 0.7069 | 0.00699 |
| RTMIM | livertumor | PTPRC | CD44 | 0.5369 | 0.8016 | 0.6692 | 0.00699 |
| RTMIM | lungtumor | APOE | SDC1 | 1.1772 | 0.2715 | 0.7243 | 0.000999 |
| RTMIM | lungtumor | APOE | SDC4 | 1.1772 | 0.3516 | 0.7644 | 0.000999 |
| RTMIM | lungtumor | CALM1 | RPSA | 1.036 | 1.6057 | 1.3209 | 0.000999 |
| RTMIM | lungtumor | CD99 | CD99 | 0.87 | 0.5938 | 0.7319 | 0.000999 |
| RTMIM | lungtumor | COL1A1 | CD44 | 0.4843 | 0.3802 | 0.4322 | 0.000999 |
| RTMIM | lungtumor | COL1A1 | SDC1 | 0.4843 | 0.2715 | 0.3779 | 0.000999 |
| RTMIM | lungtumor | CXCL12 | CXCR4 | 0.3736 | 0.2684 | 0.321 | 0.000999 |
| RTMIM | lungtumor | CXCL16 | CXCR4 | 0.7543 | 0.2684 | 0.5113 | 0.000999 |
| RTMIM | lungtumor | CXCL2 | CXCR4 | 1.4017 | 0.2684 | 0.8351 | 0.000999 |

| | | | | | | | |
|---|---|---|---|---|---|---|---|
| RTMIM | lungtumor | CXCL3 | CXCR4 | 1.1581 | 0.2684 | 0.7133 | 0.000999 |
| RTMIM | lungtumor | FN1 | CD44 | 0.3859 | 0.3802 | 0.3831 | 0.000999 |
| RTMIM | lungtumor | FN1 | RPSA | 0.3859 | 1.6057 | 0.9958 | 0.000999 |
| RTMIM | lungtumor | FN1 | SDC1 | 0.3859 | 0.2715 | 0.3287 | 0.000999 |
| RTMIM | lungtumor | HBEGF | CD44 | 0.6122 | 0.3802 | 0.4962 | 0.000999 |
| RTMIM | lungtumor | IGF1 | RACK1 | 0.4606 | 2.2108 | 1.3357 | 0.000999 |
| RTMIM | lungtumor | ITGB1 | CD151 | 0.4226 | 0.6776 | 0.5501 | 0.000999 |
| RTMIM | lungtumor | ITGB1 | CD46 | 0.4226 | 0.434 | 0.4283 | 0.000999 |
| RTMIM | lungtumor | ITGB1 | CD47 | 0.4226 | 0.3255 | 0.374 | 0.000999 |
| RTMIM | lungtumor | ITGB1 | RACK1 | 0.4226 | 2.2108 | 1.3167 | 0.000999 |
| RTMIM | lungtumor | ITGB1 | SDC1 | 0.4226 | 0.2715 | 0.347 | 0.000999 |
| RTMIM | lungtumor | PECAM1 | CXCR4 | 0.3342 | 0.2684 | 0.3013 | 0.000999 |
| RTMIM | lungtumor | PECAM1 | SDC1 | 0.3342 | 0.2715 | 0.3029 | 0.000999 |
| RTMIM | lungtumor | PECAM1 | SDC4 | 0.3342 | 0.3516 | 0.3429 | 0.000999 |
| RTMIM | lungtumor | PLAU | CD44 | 0.3274 | 0.3802 | 0.3538 | 0.000999 |
| RTMIM | lungtumor | PTPRC | CD44 | 0.5369 | 0.3802 | 0.4585 | 0.000999 |
| RTMIM | lungtumor | PTPRC | CXCR4 | 0.5369 | 0.2684 | 0.4026 | 0.000999 |
| RTMIM | lungtumor | RTN4 | CD44 | 0.9264 | 0.3802 | 0.6533 | 0.000999 |
| RTMIM | lungtumor | RTN4 | CD99 | 0.9264 | 0.5938 | 0.7601 | 0.000999 |
| RTMIM | lungtumor | THBS1 | CD47 | 0.3968 | 0.3255 | 0.3611 | 0.000999 |
| RTMIM | lungtumor | THBS1 | SDC1 | 0.3968 | 0.2715 | 0.3341 | 0.000999 |
| RTMIM | lungtumor | TNF | CALM1 | 0.3681 | 0.8426 | 0.6054 | 0.000999 |
| RTMIM | lungtumor | TNF | RACK1 | 0.3681 | 2.2108 | 1.2895 | 0.000999 |
| RTMIM | melanomatumo | ANXA1 | CD44 | 1.1309 | 1.2653 | 1.1981 | 0.000999 |
| RTMIM | melanomatumo | ANXA1 | EDNRB | 1.1309 | 0.5666 | 0.8488 | 0.000999 |
| RTMIM | melanomatumo | CALM1 | RPSA | 1.036 | 3.7787 | 2.4073 | 0.000999 |
| RTMIM | melanomatumo | CALR | CD44 | 0.7678 | 1.2653 | 1.0166 | 0.000999 |
| RTMIM | melanomatumo | COL1A1 | CD44 | 0.4843 | 1.2653 | 0.8748 | 0.000999 |
| RTMIM | melanomatumo | FN1 | CD44 | 0.3859 | 1.2653 | 0.8256 | 0.000999 |
| RTMIM | melanomatumo | FN1 | RPSA | 0.3859 | 3.7787 | 2.0823 | 0.000999 |

| | | | | | | | |
|---|---|---|---|---|---|---|---|
| RTMIM | melanomatumo | HBEGF | CD44 | 0.6122 | 1.2653 | 0.9388 | 0.000999 |
| RTMIM | melanomatumo | HMGB2 | CD44 | 0.351 | 1.2653 | 0.8082 | 0.000999 |
| RTMIM | melanomatumo | IGF1 | RACK1 | 0.4606 | 4.145 | 2.3028 | 0.000999 |
| RTMIM | melanomatumo | ITGB1 | RACK1 | 0.4226 | 4.145 | 2.2838 | 0.000999 |
| RTMIM | melanomatumo | PLAU | CD44 | 0.3274 | 1.2653 | 0.7964 | 0.000999 |
| RTMIM | melanomatumo | PTPRC | CD44 | 0.5369 | 1.2653 | 0.9011 | 0.000999 |
| RTMIM | melanomatumo | RPS19 | RPSA | 3.2576 | 3.7787 | 3.5181 | 0.000999 |
| RTMIM | melanomatumo | RTN4 | CD44 | 0.9264 | 1.2653 | 1.0959 | 0.000999 |
| RTMIM | melanomatumo | SPP1 | CD44 | 0.6105 | 1.2653 | 0.9379 | 0.000999 |
| RTMIM | melanomatumo | TNF | RACK1 | 0.3681 | 4.145 | 2.2566 | 0.000999 |
| RTMIM | melanomatumo | NAMPT | ANXA6 | 1.2179 | 0.3625 | 0.7902 | 0.002 |
| RTMIM | melanomatumo | GSTP1 | CD44 | 1.3137 | 1.2653 | 1.2895 | 0.003 |
| RTMIM | melanomatumo | ARF1 | CD44 | 0.5732 | 1.2653 | 0.9193 | 0.004 |
| RTMIM | melanomatumo | A2M | TNFRSF14 | 1.0949 | 0.5069 | 0.8009 | 0.005 |
| RTMIM | ovarytumor | C3 | CXCR4 | 0.3594 | 0.3688 | 0.3641 | 0.000999 |
| RTMIM | ovarytumor | CALM1 | RPSA | 1.036 | 2.9542 | 1.9951 | 0.000999 |
| RTMIM | ovarytumor | COL1A1 | ITGB1 | 0.4843 | 0.2783 | 0.3813 | 0.000999 |
| RTMIM | ovarytumor | CXCL12 | CXCR4 | 0.3736 | 0.3688 | 0.3712 | 0.000999 |
| RTMIM | ovarytumor | CXCL16 | CXCR4 | 0.7543 | 0.3688 | 0.5615 | 0.000999 |
| RTMIM | ovarytumor | CXCL2 | CXCR4 | 1.4017 | 0.3688 | 0.8853 | 0.000999 |
| RTMIM | ovarytumor | CXCL3 | CXCR4 | 1.1581 | 0.3688 | 0.7635 | 0.000999 |
| RTMIM | ovarytumor | FN1 | ITGB1 | 0.3859 | 0.2783 | 0.3321 | 0.000999 |
| RTMIM | ovarytumor | FN1 | RPSA | 0.3859 | 2.9542 | 1.6701 | 0.000999 |
| RTMIM | ovarytumor | IGF1 | RACK1 | 0.4606 | 2.9543 | 1.7075 | 0.000999 |
| RTMIM | ovarytumor | ITGB1 | CD151 | 0.4226 | 0.4981 | 0.4603 | 0.000999 |
| RTMIM | ovarytumor | ITGB1 | CD47 | 0.4226 | 0.4931 | 0.4578 | 0.000999 |
| RTMIM | ovarytumor | ITGB1 | RACK1 | 0.4226 | 2.9543 | 1.6884 | 0.000999 |
| RTMIM | ovarytumor | LGALS3 | ITGB1 | 0.7006 | 0.2783 | 0.4894 | 0.000999 |
| RTMIM | ovarytumor | PECAM1 | CXCR4 | 0.3342 | 0.3688 | 0.3515 | 0.000999 |
| RTMIM | ovarytumor | PTPRC | CXCR4 | 0.5369 | 0.3688 | 0.4528 | 0.000999 |

| | | | | | | | |
|---|---|---|---|---|---|---|---|
| RTMIM | ovarytumor | SPP1 | ITGB1 | 0.6105 | 0.2783 | 0.4444 | 0.000999 |
| RTMIM | ovarytumor | TGFB1 | ITGB1 | 0.4158 | 0.2783 | 0.3471 | 0.000999 |
| RTMIM | ovarytumor | THBS1 | CD47 | 0.3968 | 0.4931 | 0.4449 | 0.000999 |
| RTMIM | ovarytumor | THBS1 | ITGB1 | 0.3968 | 0.2783 | 0.3375 | 0.000999 |
| RTMIM | ovarytumor | TNF | CALM1 | 0.3681 | 1.0087 | 0.6884 | 0.000999 |
| RTMIM | ovarytumor | TNF | RACK1 | 0.3681 | 2.9543 | 1.6612 | 0.000999 |
| RTMIM | ovarytumor | TNFSF13B | ITGB1 | 0.4496 | 0.2783 | 0.3639 | 0.000999 |
| RTMIM | ovarytumor | HSP90B1 | ITGB1 | 0.7617 | 0.2783 | 0.52 | 0.003 |
| breasttumor | RTMIM | ANXA1 | C5AR1 | 0.4092 | 0.8753 | 0.6422 | 0.000999 |
| breasttumor | RTMIM | ANXA1 | CXCR4 | 0.4092 | 0.8835 | 0.6463 | 0.000999 |
| breasttumor | RTMIM | APP | C3AR1 | 0.5409 | 0.6107 | 0.5758 | 0.000999 |
| breasttumor | RTMIM | APP | C5AR1 | 0.5409 | 0.8753 | 0.7081 | 0.000999 |
| breasttumor | RTMIM | APP | CCR1 | 0.5409 | 0.2695 | 0.4052 | 0.000999 |
| breasttumor | RTMIM | APP | CD36 | 0.5409 | 0.3862 | 0.4636 | 0.000999 |
| breasttumor | RTMIM | APP | CXCR4 | 0.5409 | 0.8835 | 0.7122 | 0.000999 |
| breasttumor | RTMIM | APP | LRP1 | 0.5409 | 0.5172 | 0.5291 | 0.000999 |
| breasttumor | RTMIM | APP | TGFBR2 | 0.5409 | 0.3725 | 0.4567 | 0.000999 |
| breasttumor | RTMIM | APP | TNFRSF14 | 0.5409 | 0.2651 | 0.403 | 0.000999 |
| breasttumor | RTMIM | ARF1 | CD4 | 1.6219 | 0.655 | 1.1384 | 0.000999 |
| breasttumor | RTMIM | ARF1 | CD44 | 1.6219 | 0.8194 | 1.2207 | 0.000999 |
| breasttumor | RTMIM | CALM1 | TNFRSF1A | 0.9015 | 0.5106 | 0.706 | 0.000999 |
| breasttumor | RTMIM | CALM1 | TNFRSF1B | 0.9015 | 0.5754 | 0.7384 | 0.000999 |
| breasttumor | RTMIM | CALR | CD44 | 1.1835 | 0.8194 | 1.0015 | 0.000999 |
| breasttumor | RTMIM | CALR | LRP1 | 1.1835 | 0.5172 | 0.8503 | 0.000999 |
| breasttumor | RTMIM | CD99 | CD99 | 0.3871 | 0.87 | 0.6285 | 0.000999 |
| breasttumor | RTMIM | COL2A1 | CD44 | 0.3935 | 0.8194 | 0.6065 | 0.000999 |
| breasttumor | RTMIM | COL2A1 | ITGB1 | 0.3935 | 0.4226 | 0.408 | 0.000999 |
| breasttumor | RTMIM | GSTP1 | CD44 | 2.5435 | 0.8194 | 1.6815 | 0.000999 |
| breasttumor | RTMIM | HEBP1 | C3AR1 | 0.4062 | 0.6107 | 0.5085 | 0.000999 |
| breasttumor | RTMIM | HEBP1 | C5AR1 | 0.4062 | 0.8753 | 0.6408 | 0.000999 |

| | | | | | | | |
|---|---|---|---|---|---|---|---|
| breasttumor | RTMIM | HEBP1 | CCR1 | 0.4062 | 0.2695 | 0.3379 | 0.000999 |
| breasttumor | RTMIM | HEBP1 | CXCR4 | 0.4062 | 0.8835 | 0.6448 | 0.000999 |
| breasttumor | RTMIM | HEBP1 | FPR3 | 0.4062 | 0.5114 | 0.4588 | 0.000999 |
| breasttumor | RTMIM | HMGB1 | LY96 | 1.1815 | 0.9314 | 1.0564 | 0.000999 |
| breasttumor | RTMIM | HMGB1 | TLR2 | 1.1815 | 0.2839 | 0.7327 | 0.000999 |
| breasttumor | RTMIM | HMGB2 | CD44 | 0.3139 | 0.8194 | 0.5667 | 0.000999 |
| breasttumor | RTMIM | HMGB2 | DAB2 | 0.3139 | 1.5382 | 0.9261 | 0.000999 |
| breasttumor | RTMIM | HSP90B1 | ITGB1 | 0.7786 | 0.4226 | 0.6006 | 0.000999 |
| breasttumor | RTMIM | HSP90B1 | LRP1 | 0.7786 | 0.5172 | 0.6479 | 0.000999 |
| breasttumor | RTMIM | HSP90B1 | TNFRSF1B | 0.7786 | 0.5754 | 0.677 | 0.000999 |
| breasttumor | RTMIM | ITGB1 | CD36 | 0.2939 | 0.3862 | 0.34 | 0.000999 |
| breasttumor | RTMIM | ITGB1 | CD47 | 0.2939 | 0.3411 | 0.3175 | 0.000999 |
| breasttumor | RTMIM | ITGB1 | DAB2 | 0.2939 | 1.5382 | 0.916 | 0.000999 |
| breasttumor | RTMIM | ITGB1 | PLAUR | 0.2939 | 1.1353 | 0.7146 | 0.000999 |
| breasttumor | RTMIM | LGALS3 | CLEC7A | 0.8391 | 0.4816 | 0.6604 | 0.000999 |
| breasttumor | RTMIM | LGALS3 | COLEC12 | 0.8391 | 0.3224 | 0.5807 | 0.000999 |
| breasttumor | RTMIM | LGALS3 | FCGR2A | 0.8391 | 1.1714 | 1.0052 | 0.000999 |
| breasttumor | RTMIM | LGALS3 | FCGR2B | 0.8391 | 0.6878 | 0.7634 | 0.000999 |
| breasttumor | RTMIM | LGALS3 | ITGB1 | 0.8391 | 0.4226 | 0.6308 | 0.000999 |
| breasttumor | RTMIM | LGALS3 | TFRC | 0.8391 | 0.6028 | 0.7209 | 0.000999 |
| breasttumor | RTMIM | LGALS3 | TGFBR2 | 0.8391 | 0.3725 | 0.6058 | 0.000999 |
| breasttumor | RTMIM | LRPAP1 | LRP1 | 0.2756 | 0.5172 | 0.3964 | 0.000999 |
| breasttumor | RTMIM | MDK | LRP1 | 0.592 | 0.5172 | 0.5546 | 0.000999 |
| breasttumor | RTMIM | MDK | SDC3 | 0.592 | 0.2862 | 0.4391 | 0.000999 |
| breasttumor | RTMIM | MIF | CD74 | 3.2716 | 4.5086 | 3.8901 | 0.000999 |
| breasttumor | RTMIM | MIF | TNFRSF14 | 3.2716 | 0.2651 | 1.7683 | 0.000999 |
| breasttumor | RTMIM | NPW | C3AR1 | 0.5478 | 0.6107 | 0.5793 | 0.000999 |
| breasttumor | RTMIM | NPW | C5AR1 | 0.5478 | 0.8753 | 0.7116 | 0.000999 |
| breasttumor | RTMIM | NPW | CCR1 | 0.5478 | 0.2695 | 0.4087 | 0.000999 |
| breasttumor | RTMIM | NPW | CXCR4 | 0.5478 | 0.8835 | 0.7157 | 0.000999 |

| | | | | | | | |
|---|---|---|---|---|---|---|---|
| breasttumor | RTMIM | RPS19 | C5AR1 | 3.691 | 0.8753 | 2.2832 | 0.000999 |
| breasttumor | RTMIM | RPS19 | CD4 | 3.691 | 0.655 | 2.173 | 0.000999 |
| breasttumor | RTMIM | RPS19 | ICAM1 | 3.691 | 0.5982 | 2.1446 | 0.000999 |
| breasttumor | RTMIM | SERPINE2 | LRP1 | 0.6001 | 0.5172 | 0.5587 | 0.000999 |
| breasttumor | RTMIM | TF | CD4 | 0.2688 | 0.655 | 0.4619 | 0.000999 |
| breasttumor | RTMIM | TF | TFRC | 0.2688 | 0.6028 | 0.4358 | 0.000999 |
| breasttumor | RTMIM | TIMP1 | LRP1 | 0.7045 | 0.5172 | 0.6108 | 0.000999 |
| breasttumor | RTMIM | ANXA1 | CD4 | 0.4092 | 0.655 | 0.5321 | 0.003 |
| colorectaltumor | RTMIM | AIMP1 | RACK1 | 0.5265 | 2.2204 | 1.3735 | 0.000999 |
| colorectaltumor | RTMIM | AIMP1 | RRBP1 | 0.5265 | 0.2586 | 0.3926 | 0.000999 |
| colorectaltumor | RTMIM | APP | C3AR1 | 0.4966 | 0.6107 | 0.5537 | 0.000999 |
| colorectaltumor | RTMIM | APP | C5AR1 | 0.4966 | 0.8753 | 0.686 | 0.000999 |
| colorectaltumor | RTMIM | APP | CCR1 | 0.4966 | 0.2695 | 0.3831 | 0.000999 |
| colorectaltumor | RTMIM | APP | CD36 | 0.4966 | 0.3862 | 0.4414 | 0.000999 |
| colorectaltumor | RTMIM | APP | CXCR4 | 0.4966 | 0.8835 | 0.6901 | 0.000999 |
| colorectaltumor | RTMIM | APP | LRP1 | 0.4966 | 0.5172 | 0.5069 | 0.000999 |
| colorectaltumor | RTMIM | APP | TGFBR2 | 0.4966 | 0.3725 | 0.4345 | 0.000999 |
| colorectaltumor | RTMIM | APP | TNFRSF14 | 0.4966 | 0.2651 | 0.3809 | 0.000999 |
| colorectaltumor | RTMIM | ARF1 | CD4 | 0.6489 | 0.655 | 0.6519 | 0.000999 |
| colorectaltumor | RTMIM | ARF1 | CD44 | 0.6489 | 0.8194 | 0.7342 | 0.000999 |
| colorectaltumor | RTMIM | CALM1 | TNFRSF1B | 1.1676 | 0.5754 | 0.8715 | 0.000999 |
| colorectaltumor | RTMIM | GSTP1 | CD44 | 2.8039 | 0.8194 | 1.8117 | 0.000999 |
| colorectaltumor | RTMIM | HMGB1 | LY96 | 2.155 | 0.9314 | 1.5432 | 0.000999 |
| colorectaltumor | RTMIM | HMGB1 | TLR2 | 2.155 | 0.2839 | 1.2194 | 0.000999 |
| colorectaltumor | RTMIM | HMGB2 | CD44 | 0.5347 | 0.8194 | 0.6771 | 0.000999 |
| colorectaltumor | RTMIM | HMGB2 | DAB2 | 0.5347 | 1.5382 | 1.0365 | 0.000999 |
| colorectaltumor | RTMIM | ITGB1 | DAB2 | 0.4214 | 1.5382 | 0.9798 | 0.000999 |
| colorectaltumor | RTMIM | ITGB1 | PLAUR | 0.4214 | 1.1353 | 0.7783 | 0.000999 |
| colorectaltumor | RTMIM | LGALS3 | CLEC7A | 1.5563 | 0.4816 | 1.0189 | 0.000999 |
| colorectaltumor | RTMIM | LGALS3 | COLEC12 | 1.5563 | 0.3224 | 0.9393 | 0.000999 |

| | | | | | | | |
|---|---|---|---|---|---|---|---|
| colorectaltumor | RTMIM | LGALS3 | FCGR2A | 1.5563 | 1.1714 | 1.3638 | 0.000999 |
| colorectaltumor | RTMIM | LGALS3 | FCGR2B | 1.5563 | 0.6878 | 1.122 | 0.000999 |
| colorectaltumor | RTMIM | LGALS3 | ITGB1 | 1.5563 | 0.4226 | 0.9894 | 0.000999 |
| colorectaltumor | RTMIM | LGALS3 | TFRC | 1.5563 | 0.6028 | 1.0795 | 0.000999 |
| colorectaltumor | RTMIM | LGALS3 | TGFBR2 | 1.5563 | 0.3725 | 0.9644 | 0.000999 |
| colorectaltumor | RTMIM | MDK | LRP1 | 0.7548 | 0.5172 | 0.636 | 0.000999 |
| colorectaltumor | RTMIM | MDK | SDC3 | 0.7548 | 0.2862 | 0.5205 | 0.000999 |
| colorectaltumor | RTMIM | MIF | CD74 | 1.6741 | 4.5086 | 3.0913 | 0.000999 |
| colorectaltumor | RTMIM | MIF | TNFRSF14 | 1.6741 | 0.2651 | 0.9696 | 0.000999 |
| colorectaltumor | RTMIM | RPS19 | C5AR1 | 4.0956 | 0.8753 | 2.4854 | 0.000999 |
| colorectaltumor | RTMIM | RPS19 | CD4 | 4.0956 | 0.655 | 2.3753 | 0.000999 |
| colorectaltumor | RTMIM | RPS19 | ICAM1 | 4.0956 | 0.5982 | 2.3469 | 0.000999 |
| colorectaltumor | RTMIM | RPS19 | RPSA | 4.0956 | 1.7151 | 2.9053 | 0.000999 |
| colorectaltumor | RTMIM | CALM1 | TNFRSF1A | 1.1676 | 0.5106 | 0.8391 | 0.002 |
| colorectaltumor | RTMIM | COL1A1 | CD44 | 0.4875 | 0.8194 | 0.6535 | 0.00599 |
| livertumor | RTMIM | AGRN | LRP1 | 0.3492 | 0.5172 | 0.4332 | 0.000999 |
| livertumor | RTMIM | AGT | ATP6AP2 | 0.4701 | 0.6081 | 0.5391 | 0.000999 |
| livertumor | RTMIM | AGT | C3AR1 | 0.4701 | 0.6107 | 0.5404 | 0.000999 |
| livertumor | RTMIM | AGT | C5AR1 | 0.4701 | 0.8753 | 0.6727 | 0.000999 |
| livertumor | RTMIM | AGT | CCR1 | 0.4701 | 0.2695 | 0.3698 | 0.000999 |
| livertumor | RTMIM | AGT | CXCR4 | 0.4701 | 0.8835 | 0.6768 | 0.000999 |
| livertumor | RTMIM | AGT | LPAR6 | 0.4701 | 0.4946 | 0.4824 | 0.000999 |
| livertumor | RTMIM | APOA1 | ABCA1 | 0.2888 | 0.4351 | 0.362 | 0.000999 |
| livertumor | RTMIM | APOA1 | CD36 | 0.2888 | 0.3862 | 0.3375 | 0.000999 |
| livertumor | RTMIM | APOA1 | LRP1 | 0.2888 | 0.5172 | 0.403 | 0.000999 |
| livertumor | RTMIM | APOA1 | MSR1 | 0.2888 | 0.7036 | 0.4962 | 0.000999 |
| livertumor | RTMIM | APP | C3AR1 | 0.5626 | 0.6107 | 0.5866 | 0.000999 |
| livertumor | RTMIM | APP | C5AR1 | 0.5626 | 0.8753 | 0.7189 | 0.000999 |
| livertumor | RTMIM | APP | CCR1 | 0.5626 | 0.2695 | 0.416 | 0.000999 |
| livertumor | RTMIM | APP | CD36 | 0.5626 | 0.3862 | 0.4744 | 0.000999 |

| | | | | | | | |
|---|---|---|---|---|---|---|---|
| livertumor | RTMIM | APP | CXCR4 | 0.5626 | 0.8835 | 0.723 | 0.000999 |
| livertumor | RTMIM | APP | LRP1 | 0.5626 | 0.5172 | 0.5399 | 0.000999 |
| livertumor | RTMIM | APP | TGFBR2 | 0.5626 | 0.3725 | 0.4675 | 0.000999 |
| livertumor | RTMIM | APP | TNFRSF14 | 0.5626 | 0.2651 | 0.4138 | 0.000999 |
| livertumor | RTMIM | AREG | ICAM1 | 0.448 | 0.5982 | 0.5231 | 0.000999 |
| livertumor | RTMIM | ARF1 | CD4 | 0.6936 | 0.655 | 0.6743 | 0.000999 |
| livertumor | RTMIM | C3 | C3AR1 | 0.6584 | 0.6107 | 0.6346 | 0.000999 |
| livertumor | RTMIM | C3 | C5AR1 | 0.6584 | 0.8753 | 0.7669 | 0.000999 |
| livertumor | RTMIM | C3 | CCR1 | 0.6584 | 0.2695 | 0.464 | 0.000999 |
| livertumor | RTMIM | C3 | CXCR4 | 0.6584 | 0.8835 | 0.771 | 0.000999 |
| livertumor | RTMIM | C3 | ITGAM | 0.6584 | 0.3358 | 0.4971 | 0.000999 |
| livertumor | RTMIM | C3 | ITGB2 | 0.6584 | 0.8627 | 0.7606 | 0.000999 |
| livertumor | RTMIM | C3 | LRP1 | 0.6584 | 0.5172 | 0.5878 | 0.000999 |
| livertumor | RTMIM | CALR | CD44 | 1.0416 | 0.8194 | 0.9305 | 0.000999 |
| livertumor | RTMIM | CALR | LRP1 | 1.0416 | 0.5172 | 0.7794 | 0.000999 |
| livertumor | RTMIM | CCL20 | C3AR1 | 1.0877 | 0.6107 | 0.8492 | 0.000999 |
| livertumor | RTMIM | CCL20 | C5AR1 | 1.0877 | 0.8753 | 0.9815 | 0.000999 |
| livertumor | RTMIM | CCL20 | CCR1 | 1.0877 | 0.2695 | 0.6786 | 0.000999 |
| livertumor | RTMIM | CCL20 | CXCR4 | 1.0877 | 0.8835 | 0.9856 | 0.000999 |
| livertumor | RTMIM | CDH1 | RACK1 | 0.4896 | 2.2204 | 1.355 | 0.000999 |
| livertumor | RTMIM | CFH | ITGAM | 0.3373 | 0.3358 | 0.3366 | 0.000999 |
| livertumor | RTMIM | DSG2 | CD44 | 0.7479 | 0.8194 | 0.7837 | 0.000999 |
| livertumor | RTMIM | DSG2 | CXCR4 | 0.7479 | 0.8835 | 0.8157 | 0.000999 |
| livertumor | RTMIM | DSG2 | DAB2 | 0.7479 | 1.5382 | 1.1431 | 0.000999 |
| livertumor | RTMIM | FGA | ITGB2 | 0.2688 | 0.8627 | 0.5657 | 0.000999 |
| livertumor | RTMIM | GSTP1 | CD44 | 1.7219 | 0.8194 | 1.2706 | 0.000999 |
| livertumor | RTMIM | HMGB1 | LY96 | 1.4361 | 0.9314 | 1.1837 | 0.000999 |
| livertumor | RTMIM | HMGB1 | TLR2 | 1.4361 | 0.2839 | 0.86 | 0.000999 |
| livertumor | RTMIM | HSP90B1 | LRP1 | 0.8951 | 0.5172 | 0.7061 | 0.000999 |
| livertumor | RTMIM | HSP90B1 | TNFRSF1B | 0.8951 | 0.5754 | 0.7352 | 0.000999 |

| | | | | | | | |
|---|---|---|---|---|---|---|---|
| livertumor | RTMIM | HSPG2 | LRP1 | 0.2746 | 0.5172 | 0.3959 | 0.000999 |
| livertumor | RTMIM | ITGB1 | CD36 | 1.6044 | 0.3862 | 0.9953 | 0.000999 |
| livertumor | RTMIM | ITGB1 | CD46 | 1.6044 | 0.2646 | 0.9345 | 0.000999 |
| livertumor | RTMIM | ITGB1 | CD47 | 1.6044 | 0.3411 | 0.9728 | 0.000999 |
| livertumor | RTMIM | ITGB1 | DAB2 | 1.6044 | 1.5382 | 1.5713 | 0.000999 |
| livertumor | RTMIM | ITGB1 | PLAUR | 1.6044 | 1.1353 | 1.3699 | 0.000999 |
| livertumor | RTMIM | ITGB1 | RACK1 | 1.6044 | 2.2204 | 1.9124 | 0.000999 |
| livertumor | RTMIM | LAMC2 | ITGB1 | 0.4847 | 0.4226 | 0.4536 | 0.000999 |
| livertumor | RTMIM | LGALS3 | CLEC7A | 0.8172 | 0.4816 | 0.6494 | 0.000999 |
| livertumor | RTMIM | LGALS3 | COLEC12 | 0.8172 | 0.3224 | 0.5698 | 0.000999 |
| livertumor | RTMIM | LGALS3 | FCGR2A | 0.8172 | 1.1714 | 0.9943 | 0.000999 |
| livertumor | RTMIM | LGALS3 | FCGR2B | 0.8172 | 0.6878 | 0.7525 | 0.000999 |
| livertumor | RTMIM | LGALS3 | TFRC | 0.8172 | 0.6028 | 0.71 | 0.000999 |
| livertumor | RTMIM | LGALS3 | TGFBR2 | 0.8172 | 0.3725 | 0.5948 | 0.000999 |
| livertumor | RTMIM | MDK | LRP1 | 1.6101 | 0.5172 | 1.0636 | 0.000999 |
| livertumor | RTMIM | MDK | SDC3 | 1.6101 | 0.2862 | 0.9482 | 0.000999 |
| livertumor | RTMIM | MIF | CD74 | 1.7638 | 4.5086 | 3.1362 | 0.000999 |
| livertumor | RTMIM | MIF | TNFRSF14 | 1.7638 | 0.2651 | 1.0144 | 0.000999 |
| livertumor | RTMIM | NTS | LPAR6 | 0.3425 | 0.4946 | 0.4186 | 0.000999 |
| livertumor | RTMIM | OCLN | TGFBR2 | 0.3091 | 0.3725 | 0.3408 | 0.000999 |
| livertumor | RTMIM | PTPRF | RACK1 | 0.384 | 2.2204 | 1.3022 | 0.000999 |
| livertumor | RTMIM | PTPRF | TGFBR2 | 0.384 | 0.3725 | 0.3783 | 0.000999 |
| livertumor | RTMIM | RPS19 | C5AR1 | 4.0772 | 0.8753 | 2.4763 | 0.000999 |
| livertumor | RTMIM | RPS19 | CD4 | 4.0772 | 0.655 | 2.3661 | 0.000999 |
| livertumor | RTMIM | RPS19 | ICAM1 | 4.0772 | 0.5982 | 2.3377 | 0.000999 |
| livertumor | RTMIM | RPS19 | RPSA | 4.0772 | 1.7151 | 2.8961 | 0.000999 |
| livertumor | RTMIM | SEMA3C | NRP1 | 0.2576 | 0.4888 | 0.3732 | 0.000999 |
| livertumor | RTMIM | SEMA3C | NRP2 | 0.2576 | 0.3047 | 0.2812 | 0.000999 |
| livertumor | RTMIM | SERPINA1 | LRP1 | 1.604 | 0.5172 | 1.0606 | 0.000999 |
| livertumor | RTMIM | SPP1 | CD44 | 1.0048 | 0.8194 | 0.9121 | 0.000999 |

| | | | | | | | |
|---|---|---|---|---|---|---|---|
| livertumor | RTMIM | TF | CD4 | 0.523 | 0.655 | 0.589 | 0.000999 |
| livertumor | RTMIM | TF | TFRC | 0.523 | 0.6028 | 0.5629 | 0.000999 |
| livertumor | RTMIM | TFPI | LRP1 | 0.5043 | 0.5172 | 0.5107 | 0.000999 |
| livertumor | RTMIM | VEGFA | LYVE1 | 0.5485 | 1.0598 | 0.8041 | 0.000999 |
| livertumor | RTMIM | VEGFA | NRP1 | 0.5485 | 0.4888 | 0.5186 | 0.000999 |
| livertumor | RTMIM | VEGFA | NRP2 | 0.5485 | 0.3047 | 0.4266 | 0.000999 |
| livertumor | RTMIM | VTN | ITGB1 | 0.7399 | 0.4226 | 0.5812 | 0.000999 |
| livertumor | RTMIM | VTN | PLAUR | 0.7399 | 1.1353 | 0.9376 | 0.000999 |
| livertumor | RTMIM | AIMP1 | RRBP1 | 0.3298 | 0.2586 | 0.2942 | 0.002 |
| livertumor | RTMIM | VEGFA | AXL | 0.5485 | 0.3057 | 0.4271 | 0.00599 |
| livertumor | RTMIM | RTN4 | CD99 | 1.0046 | 0.87 | 0.9373 | 0.00899 |
| lungtumor | RTMIM | AIMP1 | RRBP1 | 0.3345 | 0.2586 | 0.2966 | 0.000999 |
| lungtumor | RTMIM | ANXA1 | C3AR1 | 1.4026 | 0.6107 | 1.0067 | 0.000999 |
| lungtumor | RTMIM | ANXA1 | C5AR1 | 1.4026 | 0.8753 | 1.139 | 0.000999 |
| lungtumor | RTMIM | ANXA1 | CCR1 | 1.4026 | 0.2695 | 0.8361 | 0.000999 |
| lungtumor | RTMIM | ANXA1 | CD4 | 1.4026 | 0.655 | 1.0288 | 0.000999 |
| lungtumor | RTMIM | ANXA1 | CD44 | 1.4026 | 0.8194 | 1.111 | 0.000999 |
| lungtumor | RTMIM | ANXA1 | CXCR4 | 1.4026 | 0.8835 | 1.1431 | 0.000999 |
| lungtumor | RTMIM | ANXA1 | FPR3 | 1.4026 | 0.5114 | 0.957 | 0.000999 |
| lungtumor | RTMIM | AREG | ICAM1 | 0.3537 | 0.5982 | 0.476 | 0.000999 |
| lungtumor | RTMIM | ARF1 | CD4 | 0.8791 | 0.655 | 0.7671 | 0.000999 |
| lungtumor | RTMIM | ARF1 | CD44 | 0.8791 | 0.8194 | 0.8493 | 0.000999 |
| lungtumor | RTMIM | CALM1 | TNFRSF1A | 0.8426 | 0.5106 | 0.6766 | 0.000999 |
| lungtumor | RTMIM | CALM1 | TNFRSF1B | 0.8426 | 0.5754 | 0.709 | 0.000999 |
| lungtumor | RTMIM | CALR | CD44 | 0.7795 | 0.8194 | 0.7995 | 0.000999 |
| lungtumor | RTMIM | CALR | LRP1 | 0.7795 | 0.5172 | 0.6484 | 0.000999 |
| lungtumor | RTMIM | CCL20 | C3AR1 | 0.2885 | 0.6107 | 0.4496 | 0.000999 |
| lungtumor | RTMIM | CCL20 | C5AR1 | 0.2885 | 0.8753 | 0.5819 | 0.000999 |
| lungtumor | RTMIM | CCL20 | CCR1 | 0.2885 | 0.2695 | 0.279 | 0.000999 |
| lungtumor | RTMIM | CCL20 | CXCR4 | 0.2885 | 0.8835 | 0.586 | 0.000999 |

| | | | | | | | |
|---|---|---|---|---|---|---|---|
| lungtumor | RTMIM | CD99 | CD99 | 0.5938 | 0.87 | 0.7319 | 0.000999 |
| lungtumor | RTMIM | CXCL10 | C3AR1 | 0.3417 | 0.6107 | 0.4762 | 0.000999 |
| lungtumor | RTMIM | CXCL10 | C5AR1 | 0.3417 | 0.8753 | 0.6085 | 0.000999 |
| lungtumor | RTMIM | CXCL10 | CCR1 | 0.3417 | 0.2695 | 0.3056 | 0.000999 |
| lungtumor | RTMIM | CXCL10 | CXCR4 | 0.3417 | 0.8835 | 0.6126 | 0.000999 |
| lungtumor | RTMIM | FGA | ITGB1 | 0.4445 | 0.4226 | 0.4335 | 0.000999 |
| lungtumor | RTMIM | FGA | ITGB2 | 0.4445 | 0.8627 | 0.6536 | 0.000999 |
| lungtumor | RTMIM | GRN | TNFRSF1B | 0.4026 | 0.5754 | 0.489 | 0.000999 |
| lungtumor | RTMIM | GSTP1 | CD44 | 2.4093 | 0.8194 | 1.6143 | 0.000999 |
| lungtumor | RTMIM | HMGB1 | LY96 | 1.1943 | 0.9314 | 1.0628 | 0.000999 |
| lungtumor | RTMIM | HMGB1 | TLR2 | 1.1943 | 0.2839 | 0.7391 | 0.000999 |
| lungtumor | RTMIM | HMGB2 | CD44 | 0.5593 | 0.8194 | 0.6894 | 0.000999 |
| lungtumor | RTMIM | HMGB2 | DAB2 | 0.5593 | 1.5382 | 1.0488 | 0.000999 |
| lungtumor | RTMIM | HSP90B1 | ITGB1 | 0.6141 | 0.4226 | 0.5183 | 0.000999 |
| lungtumor | RTMIM | HSP90B1 | LRP1 | 0.6141 | 0.5172 | 0.5656 | 0.000999 |
| lungtumor | RTMIM | HSP90B1 | TNFRSF1B | 0.6141 | 0.5754 | 0.5947 | 0.000999 |
| lungtumor | RTMIM | LGALS3 | CLEC7A | 0.9915 | 0.4816 | 0.7366 | 0.000999 |
| lungtumor | RTMIM | LGALS3 | COLEC12 | 0.9915 | 0.3224 | 0.6569 | 0.000999 |
| lungtumor | RTMIM | LGALS3 | FCGR2A | 0.9915 | 1.1714 | 1.0814 | 0.000999 |
| lungtumor | RTMIM | LGALS3 | FCGR2B | 0.9915 | 0.6878 | 0.8397 | 0.000999 |
| lungtumor | RTMIM | LGALS3 | ITGB1 | 0.9915 | 0.4226 | 0.707 | 0.000999 |
| lungtumor | RTMIM | LGALS3 | TFRC | 0.9915 | 0.6028 | 0.7972 | 0.000999 |
| lungtumor | RTMIM | LGALS3 | TGFBR2 | 0.9915 | 0.3725 | 0.682 | 0.000999 |
| lungtumor | RTMIM | MDK | LRP1 | 1.2896 | 0.5172 | 0.9034 | 0.000999 |
| lungtumor | RTMIM | MDK | SDC3 | 1.2896 | 0.2862 | 0.7879 | 0.000999 |
| lungtumor | RTMIM | MIF | CD74 | 1.4585 | 4.5086 | 2.9836 | 0.000999 |
| lungtumor | RTMIM | MIF | TNFRSF14 | 1.4585 | 0.2651 | 0.8618 | 0.000999 |
| lungtumor | RTMIM | RPS19 | C5AR1 | 3.5104 | 0.8753 | 2.1929 | 0.000999 |
| lungtumor | RTMIM | RPS19 | CD4 | 3.5104 | 0.655 | 2.0827 | 0.000999 |
| lungtumor | RTMIM | RPS19 | ICAM1 | 3.5104 | 0.5982 | 2.0543 | 0.000999 |

| | | | | | | | |
|---|---|---|---|---|---|---|---|
| lungtumor | RTMIM | RPS19 | RPSA | 3.5104 | 1.7151 | 2.6128 | 0.000999 |
| lungtumor | RTMIM | RTN4 | CD44 | 0.4013 | 0.8194 | 0.6103 | 0.000999 |
| lungtumor | RTMIM | RTN4 | CD99 | 0.4013 | 0.87 | 0.6356 | 0.000999 |
| lungtumor | RTMIM | SAA1 | C3AR1 | 0.2539 | 0.6107 | 0.4323 | 0.000999 |
| lungtumor | RTMIM | SAA1 | C5AR1 | 0.2539 | 0.8753 | 0.5646 | 0.000999 |
| lungtumor | RTMIM | SAA1 | CCR1 | 0.2539 | 0.2695 | 0.2617 | 0.000999 |
| lungtumor | RTMIM | SAA1 | CXCR4 | 0.2539 | 0.8835 | 0.5687 | 0.000999 |
| lungtumor | RTMIM | SAA1 | LPAR6 | 0.2539 | 0.4946 | 0.3743 | 0.000999 |
| lungtumor | RTMIM | SERPINA1 | LRP1 | 0.3624 | 0.5172 | 0.4398 | 0.000999 |
| lungtumor | RTMIM | SPP1 | CD44 | 1.4874 | 0.8194 | 1.1534 | 0.000999 |
| lungtumor | RTMIM | SPP1 | ITGB1 | 1.4874 | 0.4226 | 0.955 | 0.000999 |
| lungtumor | RTMIM | TFPI | LRP1 | 0.3615 | 0.5172 | 0.4393 | 0.000999 |
| lungtumor | RTMIM | TIMP1 | LRP1 | 0.6195 | 0.5172 | 0.5683 | 0.000999 |
| melanomatumo | RTMIM | ARF1 | CD4 | 1.8545 | 0.655 | 1.2547 | 0.000999 |
| melanomatumo | RTMIM | ARF1 | CD44 | 1.8545 | 0.8194 | 1.3369 | 0.000999 |
| melanomatumo | RTMIM | CD320 | TGFBR2 | 0.2588 | 0.3725 | 0.3156 | 0.000999 |
| melanomatumo | RTMIM | GSTP1 | CD44 | 2.3341 | 0.8194 | 1.5767 | 0.000999 |
| melanomatumo | RTMIM | HAS2 | CD44 | 0.3078 | 0.8194 | 0.5636 | 0.000999 |
| melanomatumo | RTMIM | MIF | CD74 | 2.6939 | 4.5086 | 3.6013 | 0.000999 |
| melanomatumo | RTMIM | MIF | TNFRSF14 | 2.6939 | 0.2651 | 1.4795 | 0.000999 |
| melanomatumo | RTMIM | SERPINE2 | LRP1 | 0.9323 | 0.5172 | 0.7247 | 0.000999 |
| melanomatumo | RTMIM | HMGB1 | LY96 | 1.6757 | 0.9314 | 1.3035 | 0.003 |
| melanomatumo | RTMIM | HMGB1 | TLR2 | 1.6757 | 0.2839 | 0.9798 | 0.003 |
| melanomatumo | RTMIM | LRPAP1 | LRP1 | 0.6123 | 0.5172 | 0.5648 | 0.005 |
| ovarytumor | RTMIM | ANXA1 | C3AR1 | 1.0314 | 0.6107 | 0.8211 | 0.000999 |
| ovarytumor | RTMIM | ANXA1 | C5AR1 | 1.0314 | 0.8753 | 0.9534 | 0.000999 |
| ovarytumor | RTMIM | ANXA1 | CCR1 | 1.0314 | 0.2695 | 0.6505 | 0.000999 |
| ovarytumor | RTMIM | ANXA1 | CD4 | 1.0314 | 0.655 | 0.8432 | 0.000999 |
| ovarytumor | RTMIM | ANXA1 | CD44 | 1.0314 | 0.8194 | 0.9254 | 0.000999 |
| ovarytumor | RTMIM | ANXA1 | CXCR4 | 1.0314 | 0.8835 | 0.9574 | 0.000999 |

| | | | | | | | |
|---|---|---|---|---|---|---|---|
| ovarytumor | RTMIM | ANXA1 | FPR3 | 1.0314 | 0.5114 | 0.7714 | 0.000999 |
| ovarytumor | RTMIM | APOE | LRP1 | 0.837 | 0.5172 | 0.6771 | 0.000999 |
| ovarytumor | RTMIM | APOE | SDC3 | 0.837 | 0.2862 | 0.5616 | 0.000999 |
| ovarytumor | RTMIM | APP | C3AR1 | 0.2907 | 0.6107 | 0.4507 | 0.000999 |
| ovarytumor | RTMIM | APP | C5AR1 | 0.2907 | 0.8753 | 0.583 | 0.000999 |
| ovarytumor | RTMIM | APP | CCR1 | 0.2907 | 0.2695 | 0.2801 | 0.000999 |
| ovarytumor | RTMIM | APP | CD36 | 0.2907 | 0.3862 | 0.3384 | 0.000999 |
| ovarytumor | RTMIM | APP | CXCR4 | 0.2907 | 0.8835 | 0.5871 | 0.000999 |
| ovarytumor | RTMIM | APP | LRP1 | 0.2907 | 0.5172 | 0.4039 | 0.000999 |
| ovarytumor | RTMIM | APP | TGFBR2 | 0.2907 | 0.3725 | 0.3316 | 0.000999 |
| ovarytumor | RTMIM | APP | TNFRSF14 | 0.2907 | 0.2651 | 0.2779 | 0.000999 |
| ovarytumor | RTMIM | ARF1 | CD4 | 0.7364 | 0.655 | 0.6957 | 0.000999 |
| ovarytumor | RTMIM | ARF1 | CD44 | 0.7364 | 0.8194 | 0.7779 | 0.000999 |
| ovarytumor | RTMIM | CALM1 | TNFRSF1A | 1.0087 | 0.5106 | 0.7596 | 0.000999 |
| ovarytumor | RTMIM | CALM1 | TNFRSF1B | 1.0087 | 0.5754 | 0.792 | 0.000999 |
| ovarytumor | RTMIM | CALR | CD44 | 0.7423 | 0.8194 | 0.7808 | 0.000999 |
| ovarytumor | RTMIM | CALR | LRP1 | 0.7423 | 0.5172 | 0.6297 | 0.000999 |
| ovarytumor | RTMIM | CCL2 | CCR1 | 0.3526 | 0.2695 | 0.3111 | 0.000999 |
| ovarytumor | RTMIM | COL1A1 | CD36 | 0.3504 | 0.3862 | 0.3683 | 0.000999 |
| ovarytumor | RTMIM | COL1A1 | CD44 | 0.3504 | 0.8194 | 0.5849 | 0.000999 |
| ovarytumor | RTMIM | COL1A1 | ITGB1 | 0.3504 | 0.4226 | 0.3865 | 0.000999 |
| ovarytumor | RTMIM | GRN | TNFRSF1A | 0.3896 | 0.5106 | 0.4501 | 0.000999 |
| ovarytumor | RTMIM | GRN | TNFRSF1B | 0.3896 | 0.5754 | 0.4825 | 0.000999 |
| ovarytumor | RTMIM | GSTP1 | CD44 | 2.6185 | 0.8194 | 1.7189 | 0.000999 |
| ovarytumor | RTMIM | HLA-DMA | CD4 | 0.2537 | 0.655 | 0.4543 | 0.000999 |
| ovarytumor | RTMIM | HLA-DRA | CD4 | 1.2369 | 0.655 | 0.9459 | 0.000999 |
| ovarytumor | RTMIM | HMGB1 | LY96 | 1.7272 | 0.9314 | 1.3293 | 0.000999 |
| ovarytumor | RTMIM | HMGB1 | TLR2 | 1.7272 | 0.2839 | 1.0056 | 0.000999 |
| ovarytumor | RTMIM | HMGB2 | CD44 | 0.4974 | 0.8194 | 0.6584 | 0.000999 |
| ovarytumor | RTMIM | HMGB2 | DAB2 | 0.4974 | 1.5382 | 1.0178 | 0.000999 |

| | | | | | | | |
|---|---|---|---|---|---|---|---|
| ovarytumor | RTMIM | HSP90B1 | ITGB1 | 0.6365 | 0.4226 | 0.5295 | 0.000999 |
| ovarytumor | RTMIM | HSP90B1 | LRP1 | 0.6365 | 0.5172 | 0.5768 | 0.000999 |
| ovarytumor | RTMIM | HSP90B1 | TNFRSF1B | 0.6365 | 0.5754 | 0.6059 | 0.000999 |
| ovarytumor | RTMIM | ITGB1 | CD36 | 0.2783 | 0.3862 | 0.3322 | 0.000999 |
| ovarytumor | RTMIM | ITGB1 | CD46 | 0.2783 | 0.2646 | 0.2715 | 0.000999 |
| ovarytumor | RTMIM | ITGB1 | DAB2 | 0.2783 | 1.5382 | 0.9082 | 0.000999 |
| ovarytumor | RTMIM | ITGB1 | PLAUR | 0.2783 | 1.1353 | 0.7068 | 0.000999 |
| ovarytumor | RTMIM | LGALS3 | CLEC7A | 0.5097 | 0.4816 | 0.4957 | 0.000999 |
| ovarytumor | RTMIM | LGALS3 | COLEC12 | 0.5097 | 0.3224 | 0.4161 | 0.000999 |
| ovarytumor | RTMIM | LGALS3 | FCGR2A | 0.5097 | 1.1714 | 0.8405 | 0.000999 |
| ovarytumor | RTMIM | LGALS3 | FCGR2B | 0.5097 | 0.6878 | 0.5988 | 0.000999 |
| ovarytumor | RTMIM | LGALS3 | ITGB1 | 0.5097 | 0.4226 | 0.4661 | 0.000999 |
| ovarytumor | RTMIM | LGALS3 | TFRC | 0.5097 | 0.6028 | 0.5563 | 0.000999 |
| ovarytumor | RTMIM | LGALS3 | TGFBR2 | 0.5097 | 0.3725 | 0.4411 | 0.000999 |
| ovarytumor | RTMIM | LRPAP1 | LRP1 | 0.3826 | 0.5172 | 0.4499 | 0.000999 |
| ovarytumor | RTMIM | MDK | LRP1 | 1.5321 | 0.5172 | 1.0246 | 0.000999 |
| ovarytumor | RTMIM | MDK | SDC3 | 1.5321 | 0.2862 | 0.9091 | 0.000999 |
| ovarytumor | RTMIM | MIF | CD74 | 0.8565 | 4.5086 | 2.6825 | 0.000999 |
| ovarytumor | RTMIM | MIF | TNFRSF14 | 0.8565 | 0.2651 | 0.5608 | 0.000999 |
| ovarytumor | RTMIM | PSAP | LRP1 | 0.6574 | 0.5172 | 0.5873 | 0.000999 |
| ovarytumor | RTMIM | RPS19 | C5AR1 | 3.9325 | 0.8753 | 2.4039 | 0.000999 |
| ovarytumor | RTMIM | RPS19 | CD4 | 3.9325 | 0.655 | 2.2938 | 0.000999 |
| ovarytumor | RTMIM | RPS19 | ICAM1 | 3.9325 | 0.5982 | 2.2654 | 0.000999 |
| ovarytumor | RTMIM | RTN4 | CD44 | 0.4992 | 0.8194 | 0.6593 | 0.000999 |
| ovarytumor | RTMIM | RTN4 | CD99 | 0.4992 | 0.87 | 0.6846 | 0.000999 |
| ovarytumor | RTMIM | SERPING1 | LRP1 | 0.3678 | 0.5172 | 0.4425 | 0.000999 |
| ovarytumor | RTMIM | SPP1 | CD44 | 0.3946 | 0.8194 | 0.607 | 0.000999 |
| ovarytumor | RTMIM | SPP1 | ITGB1 | 0.3946 | 0.4226 | 0.4086 | 0.000999 |
| ovarytumor | RTMIM | TIMP1 | LRP1 | 0.7779 | 0.5172 | 0.6475 | 0.000999 |

| Supplementary Table 8: Communication results between macrophages and lymphocytes subpopulations, and malignant cells | | | | | | | |
|---|---|---|---|---|---|---|---|
| **RTM_LA vs. lymphocytes** | | | | | | | |
| celltype_01 | celltype_02 | ligand | receptor | ligand_mean_expression | receptor_mean_expression | ligand_receptor_mean_expression | pvalue |
| RTMLA | NKT | A2M | TNFRSF14 | 0.7883 | 0.2848 | 0.5366 | 0.000999 |
| RTMLA | NKT | ADAM9 | ITGB1 | 0.2698 | 0.488 | 0.3789 | 0.000999 |
| RTMLA | NKT | ADM | PTGER2 | 0.2603 | 0.3627 | 0.3115 | 0.000999 |
| RTMLA | NKT | APOE | SORL1 | 3.8365 | 0.4137 | 2.1251 | 0.000999 |
| RTMLA | NKT | C3 | CXCR4 | 0.253 | 2.0373 | 1.1451 | 0.000999 |
| RTMLA | NKT | C3 | ITGB2 | 0.253 | 1.6481 | 0.9505 | 0.000999 |
| RTMLA | NKT | COL1A1 | CD44 | 0.3314 | 0.6724 | 0.5019 | 0.000999 |
| RTMLA | NKT | COL1A1 | ITGB1 | 0.3314 | 0.488 | 0.4097 | 0.000999 |
| RTMLA | NKT | CXCL16 | CXCR4 | 1.0236 | 2.0373 | 1.5305 | 0.000999 |
| RTMLA | NKT | CXCL16 | S1PR4 | 1.0236 | 0.4435 | 0.7336 | 0.000999 |
| RTMLA | NKT | CXCL16 | S1PR5 | 1.0236 | 0.508 | 0.7658 | 0.000999 |
| RTMLA | NKT | CXCL2 | CXCR4 | 0.4557 | 2.0373 | 1.2465 | 0.000999 |
| RTMLA | NKT | CXCL2 | S1PR4 | 0.4557 | 0.4435 | 0.4496 | 0.000999 |
| RTMLA | NKT | CXCL2 | S1PR5 | 0.4557 | 0.508 | 0.4818 | 0.000999 |
| RTMLA | NKT | CXCL3 | CXCR4 | 0.2688 | 2.0373 | 1.1531 | 0.000999 |
| RTMLA | NKT | CXCL3 | S1PR4 | 0.2688 | 0.4435 | 0.3562 | 0.000999 |
| RTMLA | NKT | CXCL3 | S1PR5 | 0.2688 | 0.508 | 0.3884 | 0.000999 |
| RTMLA | NKT | FN1 | CD44 | 1.1804 | 0.6724 | 0.9264 | 0.000999 |
| RTMLA | NKT | FN1 | ITGA4 | 1.1804 | 0.2946 | 0.7375 | 0.000999 |
| RTMLA | NKT | FN1 | ITGB1 | 1.1804 | 0.488 | 0.8342 | 0.000999 |
| RTMLA | NKT | FN1 | ITGB7 | 1.1804 | 0.4721 | 0.8262 | 0.000999 |
| RTMLA | NKT | FN1 | RPSA | 1.1804 | 2.3639 | 1.7721 | 0.000999 |
| RTMLA | NKT | GRN | TNFRSF1B | 2.0692 | 0.4949 | 1.2821 | 0.000999 |
| RTMLA | NKT | GSTP1 | CD44 | 1.8059 | 0.6724 | 1.2392 | 0.000999 |
| RTMLA | NKT | GSTP1 | IL7R | 1.8059 | 0.5933 | 1.1996 | 0.000999 |
| RTMLA | NKT | GSTP1 | ITGA4 | 1.8059 | 0.2946 | 1.0502 | 0.000999 |
| RTMLA | NKT | HEBP1 | CXCR4 | 0.2648 | 2.0373 | 1.1511 | 0.000999 |

| | | | | | | | |
|---|---|---|---|---|---|---|---|
| RTMLA | NKT | HLA-DRA | PTPRC | 4.3591 | 1.8797 | 3.1194 | 0.000999 |
| RTMLA | NKT | HSP90B1 | ITGB1 | 0.8143 | 0.488 | 0.6511 | 0.000999 |
| RTMLA | NKT | HSP90B1 | TNFRSF1B | 0.8143 | 0.4949 | 0.6546 | 0.000999 |
| RTMLA | NKT | LGALS3 | CD6 | 1.9146 | 0.4881 | 1.2014 | 0.000999 |
| RTMLA | NKT | LGALS3 | ITGB1 | 1.9146 | 0.488 | 1.2013 | 0.000999 |
| RTMLA | NKT | LGALS3 | SPN | 1.9146 | 0.6398 | 1.2772 | 0.000999 |
| RTMLA | NKT | LRPAP1 | SORL1 | 0.6395 | 0.4137 | 0.5266 | 0.000999 |
| RTMLA | NKT | MIF | ITGA4 | 0.8715 | 0.2946 | 0.583 | 0.000999 |
| RTMLA | NKT | MIF | TNFRSF14 | 0.8715 | 0.2848 | 0.5781 | 0.000999 |
| RTMLA | NKT | NAMPT | ANXA6 | 0.7571 | 0.844 | 0.8005 | 0.000999 |
| RTMLA | NKT | PECAM1 | CXCR4 | 0.3607 | 2.0373 | 1.199 | 0.000999 |
| RTMLA | NKT | RTN4 | CD44 | 0.9644 | 0.6724 | 0.8184 | 0.000999 |
| RTMLA | NKT | RTN4 | CD99 | 0.9644 | 1.4547 | 1.2095 | 0.000999 |
| RTMLA | NKT | SPP1 | CD44 | 1.3361 | 0.6724 | 1.0042 | 0.000999 |
| RTMLA | NKT | SPP1 | ITGA4 | 1.3361 | 0.2946 | 0.8153 | 0.000999 |
| RTMLA | NKT | SPP1 | ITGB1 | 1.3361 | 0.488 | 0.912 | 0.000999 |
| RTMLA | NKT | THBS1 | CD47 | 0.2853 | 0.5576 | 0.4214 | 0.000999 |
| RTMLA | NKT | THBS1 | ITGA4 | 0.2853 | 0.2946 | 0.2899 | 0.000999 |
| RTMLA | NKT | THBS1 | ITGB1 | 0.2853 | 0.488 | 0.3866 | 0.000999 |
| RTMLA | NKT | TNFSF13B | ITGB1 | 0.6721 | 0.488 | 0.58 | 0.000999 |
| RTMLA | NKT | VEGFB | ITGB1 | 0.2653 | 0.488 | 0.3766 | 0.000999 |
| RTMLA | NKT | AIMP1 | RACK1 | 0.2613 | 2.4327 | 1.347 | 0.004 |
| RTMLA | TCD4ex | A2M | TNFRSF14 | 0.7883 | 0.2806 | 0.5345 | 0.000999 |
| RTMLA | TCD4ex | ADAM9 | ITGB1 | 0.2698 | 0.4615 | 0.3657 | 0.000999 |
| RTMLA | TCD4ex | ADM | PTGER4 | 0.2603 | 0.4696 | 0.3649 | 0.000999 |
| RTMLA | TCD4ex | ANXA1 | CCR7 | 1.1107 | 0.4844 | 0.7976 | 0.000999 |
| RTMLA | TCD4ex | ANXA1 | CD4 | 1.1107 | 0.5864 | 0.8486 | 0.000999 |
| RTMLA | TCD4ex | ANXA1 | CD44 | 1.1107 | 1.0244 | 1.0675 | 0.000999 |
| RTMLA | TCD4ex | ANXA1 | CXCR4 | 1.1107 | 2.4201 | 1.7654 | 0.000999 |
| RTMLA | TCD4ex | C3 | CCR7 | 0.253 | 0.4844 | 0.3687 | 0.000999 |

| | | | | | | | |
|---|---|---|---|---|---|---|---|
| RTMLA | TCD4ex | C3 | CXCR4 | 0.253 | 2.4201 | 1.3365 | 0.000999 |
| RTMLA | TCD4ex | C3 | ITGB2 | 0.253 | 0.7871 | 0.52 | 0.000999 |
| RTMLA | TCD4ex | CALR | IL7R | 0.8929 | 1.2353 | 1.0641 | 0.000999 |
| RTMLA | TCD4ex | CD86 | CD28 | 0.47 | 0.3256 | 0.3978 | 0.000999 |
| RTMLA | TCD4ex | CD86 | CTLA4 | 0.47 | 0.6294 | 0.5497 | 0.000999 |
| RTMLA | TCD4ex | COL1A1 | CD44 | 0.3314 | 1.0244 | 0.6779 | 0.000999 |
| RTMLA | TCD4ex | COL1A1 | ITGB1 | 0.3314 | 0.4615 | 0.3965 | 0.000999 |
| RTMLA | TCD4ex | CXCL16 | CCR7 | 1.0236 | 0.4844 | 0.754 | 0.000999 |
| RTMLA | TCD4ex | CXCL16 | CXCR4 | 1.0236 | 2.4201 | 1.7218 | 0.000999 |
| RTMLA | TCD4ex | CXCL2 | CCR7 | 0.4557 | 0.4844 | 0.47 | 0.000999 |
| RTMLA | TCD4ex | CXCL2 | CXCR4 | 0.4557 | 2.4201 | 1.4379 | 0.000999 |
| RTMLA | TCD4ex | CXCL3 | CCR7 | 0.2688 | 0.4844 | 0.3766 | 0.000999 |
| RTMLA | TCD4ex | CXCL3 | CXCR4 | 0.2688 | 2.4201 | 1.3444 | 0.000999 |
| RTMLA | TCD4ex | FN1 | CD44 | 1.1804 | 1.0244 | 1.1024 | 0.000999 |
| RTMLA | TCD4ex | FN1 | ITGB1 | 1.1804 | 0.4615 | 0.8209 | 0.000999 |
| RTMLA | TCD4ex | FN1 | RPSA | 1.1804 | 2.4895 | 1.8349 | 0.000999 |
| RTMLA | TCD4ex | GRN | TNFRSF1B | 2.0692 | 0.383 | 1.2261 | 0.000999 |
| RTMLA | TCD4ex | GSTP1 | CD44 | 1.8059 | 1.0244 | 1.4152 | 0.000999 |
| RTMLA | TCD4ex | GSTP1 | IL7R | 1.8059 | 1.2353 | 1.5206 | 0.000999 |
| RTMLA | TCD4ex | HEBP1 | CCR7 | 0.2648 | 0.4844 | 0.3746 | 0.000999 |
| RTMLA | TCD4ex | HEBP1 | CXCR4 | 0.2648 | 2.4201 | 1.3425 | 0.000999 |
| RTMLA | TCD4ex | HLA-DMA | CD4 | 1.9759 | 0.5864 | 1.2811 | 0.000999 |
| RTMLA | TCD4ex | HLA-DRA | CD4 | 4.3591 | 0.5864 | 2.4727 | 0.000999 |
| RTMLA | TCD4ex | HLA-DRA | PDCD1 | 4.3591 | 0.457 | 2.408 | 0.000999 |
| RTMLA | TCD4ex | HLA-DRA | PTPRC | 4.3591 | 1.7493 | 3.0542 | 0.000999 |
| RTMLA | TCD4ex | HSP90B1 | ITGB1 | 0.8143 | 0.4615 | 0.6379 | 0.000999 |
| RTMLA | TCD4ex | HSP90B1 | TNFRSF1B | 0.8143 | 0.383 | 0.5987 | 0.000999 |
| RTMLA | TCD4ex | ITGB2 | ITGAL | 1.2598 | 0.2603 | 0.76 | 0.000999 |
| RTMLA | TCD4ex | LGALS3 | CD6 | 1.9146 | 0.5338 | 1.2242 | 0.000999 |
| RTMLA | TCD4ex | LGALS3 | ITGB1 | 1.9146 | 0.4615 | 1.188 | 0.000999 |

| | | | | | | | |
|---|---|---|---|---|---|---|---|
| RTMLA | TCD4ex | NAMPT | ANXA6 | 0.7571 | 0.5386 | 0.6479 | 0.000999 |
| RTMLA | TCD4ex | PECAM1 | CXCR4 | 0.3607 | 2.4201 | 1.3904 | 0.000999 |
| RTMLA | TCD4ex | RTN4 | CD44 | 0.9644 | 1.0244 | 0.9944 | 0.000999 |
| RTMLA | TCD4ex | RTN4 | CD99 | 0.9644 | 0.9165 | 0.9405 | 0.000999 |
| RTMLA | TCD4ex | SPP1 | CD44 | 1.3361 | 1.0244 | 1.1802 | 0.000999 |
| RTMLA | TCD4ex | SPP1 | ITGB1 | 1.3361 | 0.4615 | 0.8988 | 0.000999 |
| RTMLA | TCD4ex | THBS1 | CD47 | 0.2853 | 0.3456 | 0.3154 | 0.000999 |
| RTMLA | TCD4ex | THBS1 | ITGB1 | 0.2853 | 0.4615 | 0.3734 | 0.000999 |
| RTMLA | TCD4ex | TNFSF13B | ITGB1 | 0.6721 | 0.4615 | 0.5668 | 0.000999 |
| RTMLA | TCD4ex | VEGFB | ITGB1 | 0.2653 | 0.4615 | 0.3634 | 0.000999 |
| RTMLA | TCD4ex | MIF | TNFRSF14 | 0.8715 | 0.2806 | 0.5761 | 0.003 |
| RTMLA | TCD4reg | A2M | TNFRSF14 | 0.7883 | 0.3843 | 0.5863 | 0.000999 |
| RTMLA | TCD4reg | ADAM9 | ITGB1 | 0.2698 | 0.4001 | 0.335 | 0.000999 |
| RTMLA | TCD4reg | ANXA1 | CCR7 | 1.1107 | 0.3098 | 0.7103 | 0.000999 |
| RTMLA | TCD4reg | ANXA1 | CD4 | 1.1107 | 0.429 | 0.7698 | 0.000999 |
| RTMLA | TCD4reg | ANXA1 | CD44 | 1.1107 | 1.0896 | 1.1002 | 0.000999 |
| RTMLA | TCD4reg | ANXA1 | CXCR3 | 1.1107 | 0.3357 | 0.7232 | 0.000999 |
| RTMLA | TCD4reg | ANXA1 | CXCR4 | 1.1107 | 2.2962 | 1.7034 | 0.000999 |
| RTMLA | TCD4reg | ANXA1 | CXCR6 | 1.1107 | 0.3945 | 0.7526 | 0.000999 |
| RTMLA | TCD4reg | ANXA1 | FAS | 1.1107 | 0.2727 | 0.6917 | 0.000999 |
| RTMLA | TCD4reg | C3 | CCR7 | 0.253 | 0.3098 | 0.2814 | 0.000999 |
| RTMLA | TCD4reg | C3 | CXCR3 | 0.253 | 0.3357 | 0.2943 | 0.000999 |
| RTMLA | TCD4reg | C3 | CXCR4 | 0.253 | 2.2962 | 1.2746 | 0.000999 |
| RTMLA | TCD4reg | C3 | CXCR6 | 0.253 | 0.3945 | 0.3238 | 0.000999 |
| RTMLA | TCD4reg | C3 | ITGB2 | 0.253 | 0.6238 | 0.4384 | 0.000999 |
| RTMLA | TCD4reg | CALR | CD44 | 0.8929 | 1.0896 | 0.9913 | 0.000999 |
| RTMLA | TCD4reg | CALR | FAS | 0.8929 | 0.2727 | 0.5828 | 0.000999 |
| RTMLA | TCD4reg | CALR | IL7R | 0.8929 | 0.9682 | 0.9306 | 0.000999 |
| RTMLA | TCD4reg | CCL13 | CXCR3 | 0.3688 | 0.3357 | 0.3522 | 0.000999 |
| RTMLA | TCD4reg | CD86 | CD28 | 0.47 | 0.313 | 0.3915 | 0.000999 |

| | | | | | | | |
|---|---|---|---|---|---|---|---|
| RTMLA | TCD4reg | CD86 | CTLA4 | 0.47 | 0.7909 | 0.6305 | 0.000999 |
| RTMLA | TCD4reg | COL1A1 | CD44 | 0.3314 | 1.0896 | 0.7105 | 0.000999 |
| RTMLA | TCD4reg | COL1A1 | ITGB1 | 0.3314 | 0.4001 | 0.3657 | 0.000999 |
| RTMLA | TCD4reg | CXCL16 | CCR7 | 1.0236 | 0.3098 | 0.6667 | 0.000999 |
| RTMLA | TCD4reg | CXCL16 | CXCR3 | 1.0236 | 0.3357 | 0.6797 | 0.000999 |
| RTMLA | TCD4reg | CXCL16 | CXCR4 | 1.0236 | 2.2962 | 1.6599 | 0.000999 |
| RTMLA | TCD4reg | CXCL16 | CXCR6 | 1.0236 | 0.3945 | 0.7091 | 0.000999 |
| RTMLA | TCD4reg | CXCL16 | S1PR4 | 1.0236 | 0.2744 | 0.649 | 0.000999 |
| RTMLA | TCD4reg | CXCL2 | CCR7 | 0.4557 | 0.3098 | 0.3827 | 0.000999 |
| RTMLA | TCD4reg | CXCL2 | CXCR3 | 0.4557 | 0.3357 | 0.3957 | 0.000999 |
| RTMLA | TCD4reg | CXCL2 | CXCR4 | 0.4557 | 2.2962 | 1.3759 | 0.000999 |
| RTMLA | TCD4reg | CXCL2 | CXCR6 | 0.4557 | 0.3945 | 0.4251 | 0.000999 |
| RTMLA | TCD4reg | CXCL2 | S1PR4 | 0.4557 | 0.2744 | 0.365 | 0.000999 |
| RTMLA | TCD4reg | CXCL3 | CCR7 | 0.2688 | 0.3098 | 0.2893 | 0.000999 |
| RTMLA | TCD4reg | CXCL3 | CXCR3 | 0.2688 | 0.3357 | 0.3022 | 0.000999 |
| RTMLA | TCD4reg | CXCL3 | CXCR4 | 0.2688 | 2.2962 | 1.2825 | 0.000999 |
| RTMLA | TCD4reg | CXCL3 | CXCR6 | 0.2688 | 0.3945 | 0.3317 | 0.000999 |
| RTMLA | TCD4reg | CXCL3 | S1PR4 | 0.2688 | 0.2744 | 0.2716 | 0.000999 |
| RTMLA | TCD4reg | FN1 | CD44 | 1.1804 | 1.0896 | 1.135 | 0.000999 |
| RTMLA | TCD4reg | FN1 | ITGB1 | 1.1804 | 0.4001 | 0.7902 | 0.000999 |
| RTMLA | TCD4reg | FN1 | RPSA | 1.1804 | 2.4999 | 1.8402 | 0.000999 |
| RTMLA | TCD4reg | GRN | TNFRSF1B | 2.0692 | 0.7982 | 1.4337 | 0.000999 |
| RTMLA | TCD4reg | GSTP1 | CD44 | 1.8059 | 1.0896 | 1.4478 | 0.000999 |
| RTMLA | TCD4reg | GSTP1 | FAS | 1.8059 | 0.2727 | 1.0393 | 0.000999 |
| RTMLA | TCD4reg | GSTP1 | IL7R | 1.8059 | 0.9682 | 1.3871 | 0.000999 |
| RTMLA | TCD4reg | HEBP1 | CCR7 | 0.2648 | 0.3098 | 0.2873 | 0.000999 |
| RTMLA | TCD4reg | HEBP1 | CXCR3 | 0.2648 | 0.3357 | 0.3003 | 0.000999 |
| RTMLA | TCD4reg | HEBP1 | CXCR4 | 0.2648 | 2.2962 | 1.2805 | 0.000999 |
| RTMLA | TCD4reg | HEBP1 | CXCR6 | 0.2648 | 0.3945 | 0.3297 | 0.000999 |
| RTMLA | TCD4reg | HLA-DMA | CD4 | 1.9759 | 0.429 | 1.2024 | 0.000999 |

| | | | | | | | |
|---|---|---|---|---|---|---|---|
| RTMLA | TCD4reg | HLA-DRA | CD4 | 4.3591 | 0.429 | 2.394 | 0.000999 |
| RTMLA | TCD4reg | HLA-DRA | PTPRC | 4.3591 | 1.6603 | 3.0097 | 0.000999 |
| RTMLA | TCD4reg | HSP90B1 | ITGB1 | 0.8143 | 0.4001 | 0.6072 | 0.000999 |
| RTMLA | TCD4reg | HSP90B1 | TNFRSF1B | 0.8143 | 0.7982 | 0.8062 | 0.000999 |
| RTMLA | TCD4reg | LGALS3 | CD6 | 1.9146 | 0.2651 | 1.0898 | 0.000999 |
| RTMLA | TCD4reg | LGALS3 | FAS | 1.9146 | 0.2727 | 1.0937 | 0.000999 |
| RTMLA | TCD4reg | LGALS3 | ITGB1 | 1.9146 | 0.4001 | 1.1573 | 0.000999 |
| RTMLA | TCD4reg | MIF | TNFRSF14 | 0.8715 | 0.3843 | 0.6279 | 0.000999 |
| RTMLA | TCD4reg | NAMPT | ANXA6 | 0.7571 | 0.4557 | 0.6064 | 0.000999 |
| RTMLA | TCD4reg | PECAM1 | CXCR4 | 0.3607 | 2.2962 | 1.3284 | 0.000999 |
| RTMLA | TCD4reg | RTN4 | CD44 | 0.9644 | 1.0896 | 1.027 | 0.000999 |
| RTMLA | TCD4reg | RTN4 | CD99 | 0.9644 | 0.6603 | 0.8123 | 0.000999 |
| RTMLA | TCD4reg | SPP1 | CD44 | 1.3361 | 1.0896 | 1.2128 | 0.000999 |
| RTMLA | TCD4reg | SPP1 | ITGB1 | 1.3361 | 0.4001 | 0.8681 | 0.000999 |
| RTMLA | TCD4reg | THBS1 | CD47 | 0.2853 | 0.416 | 0.3506 | 0.000999 |
| RTMLA | TCD4reg | THBS1 | ITGB1 | 0.2853 | 0.4001 | 0.3427 | 0.000999 |
| RTMLA | TCD4reg | TNFSF13B | ITGB1 | 0.6721 | 0.4001 | 0.5361 | 0.000999 |
| RTMLA | TCD4reg | VEGFB | ITGB1 | 0.2653 | 0.4001 | 0.3327 | 0.000999 |
| RTMLA | TCD4reg | CD99 | CD99 | 0.805 | 0.6603 | 0.7327 | 0.00599 |
| RTMLA | TCD4reg | MIF | CD74 | 0.8715 | 2.0716 | 1.4716 | 0.00599 |
| RTMLA | TCD8em | A2M | TNFRSF14 | 0.7883 | 0.2915 | 0.5399 | 0.000999 |
| RTMLA | TCD8em | ADAM9 | ITGB1 | 0.2698 | 0.2557 | 0.2628 | 0.000999 |
| RTMLA | TCD8em | ADM | PTGER4 | 0.2603 | 0.2847 | 0.2725 | 0.000999 |
| RTMLA | TCD8em | C3 | CXCR3 | 0.253 | 0.4189 | 0.3359 | 0.000999 |
| RTMLA | TCD8em | C3 | CXCR4 | 0.253 | 2.8389 | 1.5459 | 0.000999 |
| RTMLA | TCD8em | C3 | CXCR6 | 0.253 | 0.3399 | 0.2964 | 0.000999 |
| RTMLA | TCD8em | C3 | ITGB2 | 0.253 | 1.0018 | 0.6274 | 0.000999 |
| RTMLA | TCD8em | CALR | IL7R | 0.8929 | 0.7183 | 0.8056 | 0.000999 |
| RTMLA | TCD8em | CCL13 | CXCR3 | 0.3688 | 0.4189 | 0.3938 | 0.000999 |
| RTMLA | TCD8em | CXCL16 | CXCR3 | 1.0236 | 0.4189 | 0.7213 | 0.000999 |

| | | | | | | | |
|---|---|---|---|---|---|---|---|
| RTMLA | TCD8em | CXCL16 | CXCR4 | 1.0236 | 2.8389 | 1.9313 | 0.000999 |
| RTMLA | TCD8em | CXCL16 | CXCR6 | 1.0236 | 0.3399 | 0.6817 | 0.000999 |
| RTMLA | TCD8em | CXCL16 | S1PR4 | 1.0236 | 0.2677 | 0.6457 | 0.000999 |
| RTMLA | TCD8em | CXCL2 | CXCR3 | 0.4557 | 0.4189 | 0.4373 | 0.000999 |
| RTMLA | TCD8em | CXCL2 | CXCR4 | 0.4557 | 2.8389 | 1.6473 | 0.000999 |
| RTMLA | TCD8em | CXCL2 | CXCR6 | 0.4557 | 0.3399 | 0.3978 | 0.000999 |
| RTMLA | TCD8em | CXCL2 | S1PR4 | 0.4557 | 0.2677 | 0.3617 | 0.000999 |
| RTMLA | TCD8em | CXCL3 | CXCR3 | 0.2688 | 0.4189 | 0.3439 | 0.000999 |
| RTMLA | TCD8em | CXCL3 | CXCR4 | 0.2688 | 2.8389 | 1.5539 | 0.000999 |
| RTMLA | TCD8em | CXCL3 | CXCR6 | 0.2688 | 0.3399 | 0.3043 | 0.000999 |
| RTMLA | TCD8em | CXCL3 | S1PR4 | 0.2688 | 0.2677 | 0.2683 | 0.000999 |
| RTMLA | TCD8em | FN1 | CD44 | 1.1804 | 1.119 | 1.1497 | 0.000999 |
| RTMLA | TCD8em | FN1 | ITGA4 | 1.1804 | 0.2823 | 0.7314 | 0.000999 |
| RTMLA | TCD8em | FN1 | ITGB1 | 1.1804 | 0.2557 | 0.7181 | 0.000999 |
| RTMLA | TCD8em | FN1 | RPSA | 1.1804 | 2.3686 | 1.7745 | 0.000999 |
| RTMLA | TCD8em | GRN | TNFRSF1B | 2.0692 | 0.3038 | 1.1865 | 0.000999 |
| RTMLA | TCD8em | GSTP1 | CD44 | 1.8059 | 1.119 | 1.4625 | 0.000999 |
| RTMLA | TCD8em | GSTP1 | IL7R | 1.8059 | 0.7183 | 1.2621 | 0.000999 |
| RTMLA | TCD8em | GSTP1 | ITGA4 | 1.8059 | 0.2823 | 1.0441 | 0.000999 |
| RTMLA | TCD8em | HEBP1 | CXCR3 | 0.2648 | 0.4189 | 0.3419 | 0.000999 |
| RTMLA | TCD8em | HEBP1 | CXCR4 | 0.2648 | 2.8389 | 1.5519 | 0.000999 |
| RTMLA | TCD8em | HEBP1 | CXCR6 | 0.2648 | 0.3399 | 0.3024 | 0.000999 |
| RTMLA | TCD8em | HLA-DRA | PTPRC | 4.3591 | 1.7912 | 3.0751 | 0.000999 |
| RTMLA | TCD8em | HSP90B1 | ITGB1 | 0.8143 | 0.2557 | 0.535 | 0.000999 |
| RTMLA | TCD8em | HSP90B1 | TNFRSF1B | 0.8143 | 0.3038 | 0.559 | 0.000999 |
| RTMLA | TCD8em | ITGB2 | ITGAL | 1.2598 | 0.3162 | 0.788 | 0.000999 |
| RTMLA | TCD8em | LGALS3 | CD6 | 1.9146 | 0.4204 | 1.1675 | 0.000999 |
| RTMLA | TCD8em | LGALS3 | ITGB1 | 1.9146 | 0.2557 | 1.0852 | 0.000999 |
| RTMLA | TCD8em | LGALS3 | SPN | 1.9146 | 0.2603 | 1.0875 | 0.000999 |
| RTMLA | TCD8em | MIF | ITGA4 | 0.8715 | 0.2823 | 0.5769 | 0.000999 |

| | | | | | | | |
|---|---|---|---|---|---|---|---|
| RTMLA | TCD8em | MIF | TNFRSF14 | 0.8715 | 0.2915 | 0.5815 | 0.000999 |
| RTMLA | TCD8em | NAMPT | ANXA6 | 0.7571 | 0.552 | 0.6546 | 0.000999 |
| RTMLA | TCD8em | PECAM1 | CXCR4 | 0.3607 | 2.8389 | 1.5998 | 0.000999 |
| RTMLA | TCD8em | RTN4 | CD44 | 0.9644 | 1.119 | 1.0417 | 0.000999 |
| RTMLA | TCD8em | RTN4 | CD99 | 0.9644 | 1.1525 | 1.0584 | 0.000999 |
| RTMLA | TCD8em | SPP1 | CD44 | 1.3361 | 1.119 | 1.2276 | 0.000999 |
| RTMLA | TCD8em | SPP1 | ITGA4 | 1.3361 | 0.2823 | 0.8092 | 0.000999 |
| RTMLA | TCD8em | SPP1 | ITGB1 | 1.3361 | 0.2557 | 0.7959 | 0.000999 |
| RTMLA | TCD8em | THBS1 | CD47 | 0.2853 | 0.4276 | 0.3564 | 0.000999 |
| RTMLA | TCD8em | THBS1 | ITGA4 | 0.2853 | 0.2823 | 0.2838 | 0.000999 |
| RTMLA | TCD8em | THBS1 | ITGB1 | 0.2853 | 0.2557 | 0.2705 | 0.000999 |
| RTMLA | TCD8em | TNFSF13B | ITGB1 | 0.6721 | 0.2557 | 0.4639 | 0.000999 |
| RTMLA | TCD8em | VEGFB | ITGB1 | 0.2653 | 0.2557 | 0.2605 | 0.000999 |
| RTMLA | TCD8em | CALR | CD44 | 0.8929 | 1.119 | 1.006 | 0.003 |
| RTMLA | TCD8em | MIF | CD74 | 0.8715 | 2.4538 | 1.6626 | 0.003 |
| RTMLA | TCD8ex | A2M | TNFRSF14 | 0.7883 | 0.3094 | 0.5489 | 0.000999 |
| RTMLA | TCD8ex | ADAM9 | ITGB1 | 0.2698 | 0.2995 | 0.2846 | 0.000999 |
| RTMLA | TCD8ex | ADM | PTGER4 | 0.2603 | 0.3089 | 0.2846 | 0.000999 |
| RTMLA | TCD8ex | ANXA1 | CD44 | 1.1107 | 0.8219 | 0.9663 | 0.000999 |
| RTMLA | TCD8ex | ANXA1 | CXCR3 | 1.1107 | 0.3647 | 0.7377 | 0.000999 |
| RTMLA | TCD8ex | ANXA1 | CXCR4 | 1.1107 | 2.3472 | 1.729 | 0.000999 |
| RTMLA | TCD8ex | ANXA1 | CXCR6 | 1.1107 | 0.6626 | 0.8866 | 0.000999 |
| RTMLA | TCD8ex | ANXA1 | ITGA4 | 1.1107 | 0.4432 | 0.777 | 0.000999 |
| RTMLA | TCD8ex | ANXA1 | VCAM1 | 1.1107 | 0.4462 | 0.7785 | 0.000999 |
| RTMLA | TCD8ex | APOE | VCAM1 | 3.8365 | 0.4462 | 2.1413 | 0.000999 |
| RTMLA | TCD8ex | C3 | CXCR3 | 0.253 | 0.3647 | 0.3088 | 0.000999 |
| RTMLA | TCD8ex | C3 | CXCR4 | 0.253 | 2.3472 | 1.3001 | 0.000999 |
| RTMLA | TCD8ex | C3 | CXCR6 | 0.253 | 0.6626 | 0.4578 | 0.000999 |
| RTMLA | TCD8ex | C3 | ITGB2 | 0.253 | 0.7635 | 0.5082 | 0.000999 |
| RTMLA | TCD8ex | CCL13 | CXCR3 | 0.3688 | 0.3647 | 0.3667 | 0.000999 |

| | | | | | | | |
|---|---|---|---|---|---|---|---|
| RTMLA | TCD8ex | CD86 | CTLA4 | 0.47 | 0.4883 | 0.4791 | 0.000999 |
| RTMLA | TCD8ex | COL1A1 | CD44 | 0.3314 | 0.8219 | 0.5767 | 0.000999 |
| RTMLA | TCD8ex | COL1A1 | ITGB1 | 0.3314 | 0.2995 | 0.3154 | 0.000999 |
| RTMLA | TCD8ex | CXCL16 | CXCR3 | 1.0236 | 0.3647 | 0.6941 | 0.000999 |
| RTMLA | TCD8ex | CXCL16 | CXCR4 | 1.0236 | 2.3472 | 1.6854 | 0.000999 |
| RTMLA | TCD8ex | CXCL16 | CXCR6 | 1.0236 | 0.6626 | 0.8431 | 0.000999 |
| RTMLA | TCD8ex | CXCL2 | CXCR3 | 0.4557 | 0.3647 | 0.4102 | 0.000999 |
| RTMLA | TCD8ex | CXCL2 | CXCR4 | 0.4557 | 2.3472 | 1.4014 | 0.000999 |
| RTMLA | TCD8ex | CXCL2 | CXCR6 | 0.4557 | 0.6626 | 0.5591 | 0.000999 |
| RTMLA | TCD8ex | CXCL3 | CXCR3 | 0.2688 | 0.3647 | 0.3167 | 0.000999 |
| RTMLA | TCD8ex | CXCL3 | CXCR4 | 0.2688 | 2.3472 | 1.308 | 0.000999 |
| RTMLA | TCD8ex | CXCL3 | CXCR6 | 0.2688 | 0.6626 | 0.4657 | 0.000999 |
| RTMLA | TCD8ex | FN1 | CD44 | 1.1804 | 0.8219 | 1.0012 | 0.000999 |
| RTMLA | TCD8ex | FN1 | ITGA4 | 1.1804 | 0.4432 | 0.8118 | 0.000999 |
| RTMLA | TCD8ex | FN1 | ITGB1 | 1.1804 | 0.2995 | 0.7399 | 0.000999 |
| RTMLA | TCD8ex | FN1 | ITGB7 | 1.1804 | 0.3007 | 0.7406 | 0.000999 |
| RTMLA | TCD8ex | FN1 | RPSA | 1.1804 | 2.2697 | 1.725 | 0.000999 |
| RTMLA | TCD8ex | GRN | TNFRSF1B | 2.0692 | 0.6436 | 1.3564 | 0.000999 |
| RTMLA | TCD8ex | GSTP1 | CD44 | 1.8059 | 0.8219 | 1.3139 | 0.000999 |
| RTMLA | TCD8ex | GSTP1 | ITGA4 | 1.8059 | 0.4432 | 1.1246 | 0.000999 |
| RTMLA | TCD8ex | GSTP1 | VCAM1 | 1.8059 | 0.4462 | 1.1261 | 0.000999 |
| RTMLA | TCD8ex | HEBP1 | CXCR3 | 0.2648 | 0.3647 | 0.3148 | 0.000999 |
| RTMLA | TCD8ex | HEBP1 | CXCR4 | 0.2648 | 2.3472 | 1.306 | 0.000999 |
| RTMLA | TCD8ex | HEBP1 | CXCR6 | 0.2648 | 0.6626 | 0.4637 | 0.000999 |
| RTMLA | TCD8ex | HLA-DRA | PDCD1 | 4.3591 | 0.6109 | 2.485 | 0.000999 |
| RTMLA | TCD8ex | HLA-DRA | PTPRC | 4.3591 | 1.6762 | 3.0176 | 0.000999 |
| RTMLA | TCD8ex | HSP90B1 | ITGB1 | 0.8143 | 0.2995 | 0.5569 | 0.000999 |
| RTMLA | TCD8ex | HSP90B1 | TNFRSF1B | 0.8143 | 0.6436 | 0.729 | 0.000999 |
| RTMLA | TCD8ex | LGALS3 | CD6 | 1.9146 | 0.318 | 1.1163 | 0.000999 |
| RTMLA | TCD8ex | LGALS3 | ITGB1 | 1.9146 | 0.2995 | 1.107 | 0.000999 |

| | | | | | | | |
|---|---|---|---|---|---|---|---|
| RTMLA | TCD8ex | LGALS3 | SPN | 1.9146 | 0.3486 | 1.1316 | 0.000999 |
| RTMLA | TCD8ex | NAMPT | ANXA6 | 0.7571 | 0.5678 | 0.6625 | 0.000999 |
| RTMLA | TCD8ex | PECAM1 | CXCR4 | 0.3607 | 2.3472 | 1.354 | 0.000999 |
| RTMLA | TCD8ex | RTN4 | CD44 | 0.9644 | 0.8219 | 0.8932 | 0.000999 |
| RTMLA | TCD8ex | RTN4 | CD99 | 0.9644 | 1.2049 | 1.0846 | 0.000999 |
| RTMLA | TCD8ex | SPP1 | CD44 | 1.3361 | 0.8219 | 1.079 | 0.000999 |
| RTMLA | TCD8ex | SPP1 | ITGA4 | 1.3361 | 0.4432 | 0.8897 | 0.000999 |
| RTMLA | TCD8ex | SPP1 | ITGB1 | 1.3361 | 0.2995 | 0.8178 | 0.000999 |
| RTMLA | TCD8ex | THBS1 | CD47 | 0.2853 | 0.3952 | 0.3402 | 0.000999 |
| RTMLA | TCD8ex | THBS1 | ITGA4 | 0.2853 | 0.4432 | 0.3642 | 0.000999 |
| RTMLA | TCD8ex | THBS1 | ITGB1 | 0.2853 | 0.2995 | 0.2924 | 0.000999 |
| RTMLA | TCD8ex | TNFSF13B | ITGB1 | 0.6721 | 0.2995 | 0.4858 | 0.000999 |
| RTMLA | TCD8ex | VEGFB | ITGB1 | 0.2653 | 0.2995 | 0.2824 | 0.000999 |
| RTMLA | TGD | A2M | TNFRSF14 | 0.7883 | 0.4642 | 0.6263 | 0.000999 |
| RTMLA | TGD | ADAM9 | ITGB1 | 0.2698 | 0.3657 | 0.3177 | 0.000999 |
| RTMLA | TGD | ADM | PTGDR | 0.2603 | 0.7153 | 0.4878 | 0.000999 |
| RTMLA | TGD | AIMP1 | RACK1 | 0.2613 | 2.7287 | 1.495 | 0.000999 |
| RTMLA | TGD | ANXA1 | CD44 | 1.1107 | 1.1342 | 1.1225 | 0.000999 |
| RTMLA | TGD | ANXA1 | CXCR3 | 1.1107 | 1.1748 | 1.1427 | 0.000999 |
| RTMLA | TGD | ANXA1 | CXCR4 | 1.1107 | 1.4894 | 1.3 | 0.000999 |
| RTMLA | TGD | ANXA1 | ITGA4 | 1.1107 | 0.3968 | 0.7538 | 0.000999 |
| RTMLA | TGD | APOE | SORL1 | 3.8365 | 0.3318 | 2.0841 | 0.000999 |
| RTMLA | TGD | C3 | CXCR3 | 0.253 | 1.1748 | 0.7139 | 0.000999 |
| RTMLA | TGD | C3 | CXCR4 | 0.253 | 1.4894 | 0.8712 | 0.000999 |
| RTMLA | TGD | C3 | ITGB2 | 0.253 | 1.4444 | 0.8487 | 0.000999 |
| RTMLA | TGD | CALM1 | RPSA | 1.1468 | 2.8814 | 2.0141 | 0.000999 |
| RTMLA | TGD | CALR | IL7R | 0.8929 | 1.3862 | 1.1396 | 0.000999 |
| RTMLA | TGD | CCL13 | CXCR3 | 0.3688 | 1.1748 | 0.7718 | 0.000999 |
| RTMLA | TGD | CD99 | CD99 | 0.805 | 1.7154 | 1.2602 | 0.000999 |
| RTMLA | TGD | CXCL16 | CXCR3 | 1.0236 | 1.1748 | 1.0992 | 0.000999 |

| | | | | | | | |
|---|---|---|---|---|---|---|---|
| RTMLA | TGD | CXCL16 | CXCR4 | 1.0236 | 1.4894 | 1.2565 | 0.000999 |
| RTMLA | TGD | CXCL2 | CXCR3 | 0.4557 | 1.1748 | 0.8152 | 0.000999 |
| RTMLA | TGD | CXCL2 | CXCR4 | 0.4557 | 1.4894 | 0.9725 | 0.000999 |
| RTMLA | TGD | CXCL3 | CXCR3 | 0.2688 | 1.1748 | 0.7218 | 0.000999 |
| RTMLA | TGD | CXCL3 | CXCR4 | 0.2688 | 1.4894 | 0.8791 | 0.000999 |
| RTMLA | TGD | FN1 | CD44 | 1.1804 | 1.1342 | 1.1573 | 0.000999 |
| RTMLA | TGD | FN1 | ITGA4 | 1.1804 | 0.3968 | 0.7886 | 0.000999 |
| RTMLA | TGD | FN1 | ITGB1 | 1.1804 | 0.3657 | 0.773 | 0.000999 |
| RTMLA | TGD | FN1 | ITGB7 | 1.1804 | 0.459 | 0.8197 | 0.000999 |
| RTMLA | TGD | FN1 | RPSA | 1.1804 | 2.8814 | 2.0309 | 0.000999 |
| RTMLA | TGD | GRN | TNFRSF1A | 2.0692 | 0.3099 | 1.1895 | 0.000999 |
| RTMLA | TGD | GSTP1 | CD44 | 1.8059 | 1.1342 | 1.4701 | 0.000999 |
| RTMLA | TGD | GSTP1 | IL7R | 1.8059 | 1.3862 | 1.5961 | 0.000999 |
| RTMLA | TGD | GSTP1 | ITGA4 | 1.8059 | 0.3968 | 1.1014 | 0.000999 |
| RTMLA | TGD | HEBP1 | CXCR3 | 0.2648 | 1.1748 | 0.7198 | 0.000999 |
| RTMLA | TGD | HEBP1 | CXCR4 | 0.2648 | 1.4894 | 0.8771 | 0.000999 |
| RTMLA | TGD | HLA-A | CD8A | 2.8931 | 1.746 | 2.3196 | 0.000999 |
| RTMLA | TGD | HLA-DRA | PTPRC | 4.3591 | 1.8653 | 3.1122 | 0.000999 |
| RTMLA | TGD | HLA-E | CD8A | 1.6584 | 1.746 | 1.7022 | 0.000999 |
| RTMLA | TGD | HLA-E | KLRD1 | 1.6584 | 1.3451 | 1.5017 | 0.000999 |
| RTMLA | TGD | HLA-F | CD8A | 0.2798 | 1.746 | 1.0129 | 0.000999 |
| RTMLA | TGD | ITGAM | ICAM1 | 0.2768 | 0.2915 | 0.2841 | 0.000999 |
| RTMLA | TGD | ITGB1 | ITGA4 | 0.3114 | 0.3968 | 0.3541 | 0.000999 |
| RTMLA | TGD | ITGB1 | RACK1 | 0.3114 | 2.7287 | 1.52 | 0.000999 |
| RTMLA | TGD | LGALS3 | CD6 | 1.9146 | 0.4684 | 1.1915 | 0.000999 |
| RTMLA | TGD | LGALS3 | ITGB1 | 1.9146 | 0.3657 | 1.1401 | 0.000999 |
| RTMLA | TGD | LGALS3 | SPN | 1.9146 | 0.5644 | 1.2395 | 0.000999 |
| RTMLA | TGD | LGALS3 | TGFBR2 | 1.9146 | 0.3677 | 1.1411 | 0.000999 |
| RTMLA | TGD | LRPAP1 | SORL1 | 0.6395 | 0.3318 | 0.4856 | 0.000999 |
| RTMLA | TGD | NAMPT | ANXA6 | 0.7571 | 0.8553 | 0.8062 | 0.000999 |

| | | | | | | | |
|---|---|---|---|---|---|---|---|
| RTMLA | TGD | PECAM1 | CXCR4 | 0.3607 | 1.4894 | 0.9251 | 0.000999 |
| RTMLA | TGD | PTPRC | CD2 | 0.5032 | 1.4522 | 0.9777 | 0.000999 |
| RTMLA | TGD | PTPRC | CD8A | 0.5032 | 1.746 | 1.1246 | 0.000999 |
| RTMLA | TGD | RPS19 | RPSA | 3.4668 | 2.8814 | 3.1741 | 0.000999 |
| RTMLA | TGD | RTN4 | CD44 | 0.9644 | 1.1342 | 1.0493 | 0.000999 |
| RTMLA | TGD | RTN4 | CD99 | 0.9644 | 1.7154 | 1.3399 | 0.000999 |
| RTMLA | TGD | SPP1 | CD44 | 1.3361 | 1.1342 | 1.2351 | 0.000999 |
| RTMLA | TGD | SPP1 | ITGA4 | 1.3361 | 0.3968 | 0.8665 | 0.000999 |
| RTMLA | TGD | SPP1 | ITGB1 | 1.3361 | 0.3657 | 0.8509 | 0.000999 |
| RTMLA | TGD | THBS1 | ITGA4 | 0.2853 | 0.3968 | 0.341 | 0.000999 |
| RTMLA | TGD | THBS1 | ITGB1 | 0.2853 | 0.3657 | 0.3255 | 0.000999 |
| RTMLA | TGD | TNFSF13B | ITGB1 | 0.6721 | 0.3657 | 0.5189 | 0.000999 |
| RTMLA | TGD | VEGFB | ITGB1 | 0.2653 | 0.3657 | 0.3155 | 0.000999 |
| RTMLA | TGD | ITGB1 | PTK2B | 0.3114 | 0.3668 | 0.3391 | 0.002 |
| RTMLA | TGD | ITGB2 | ITGAL | 1.2598 | 0.3731 | 0.8165 | 0.002 |
| RTMLA | TGD | ARF1 | ITGA4 | 0.6482 | 0.3968 | 0.5225 | 0.004 |
| NKT | RTMLA | ALOX5AP | ALOX5 | 0.5021 | 0.3103 | 0.4062 | 0.000999 |
| NKT | RTMLA | ANXA1 | C3AR1 | 1.2259 | 0.501 | 0.8635 | 0.000999 |
| NKT | RTMLA | ANXA1 | C5AR1 | 1.2259 | 0.6059 | 0.9159 | 0.000999 |
| NKT | RTMLA | ANXA1 | CCR1 | 1.2259 | 0.3215 | 0.7737 | 0.000999 |
| NKT | RTMLA | ANXA1 | CD4 | 1.2259 | 0.6388 | 0.9324 | 0.000999 |
| NKT | RTMLA | ANXA1 | CD44 | 1.2259 | 1.1046 | 1.1653 | 0.000999 |
| NKT | RTMLA | ANXA1 | FPR1 | 1.2259 | 0.3438 | 0.7849 | 0.000999 |
| NKT | RTMLA | ANXA1 | FPR3 | 1.2259 | 0.6513 | 0.9386 | 0.000999 |
| NKT | RTMLA | ARF1 | CD4 | 0.7991 | 0.6388 | 0.719 | 0.000999 |
| NKT | RTMLA | ARF1 | CD44 | 0.7991 | 1.1046 | 0.9519 | 0.000999 |
| NKT | RTMLA | CALM1 | TNFRSF1A | 1.9335 | 0.3864 | 1.16 | 0.000999 |
| NKT | RTMLA | CALR | CD44 | 0.9479 | 1.1046 | 1.0263 | 0.000999 |
| NKT | RTMLA | CALR | LRP1 | 0.9479 | 0.5181 | 0.733 | 0.000999 |
| NKT | RTMLA | CD2 | CD4 | 1.1465 | 0.6388 | 0.8927 | 0.000999 |

| | | | | | | | |
|---|---|---|---|---|---|---|---|
| NKT | RTMLA | GSTP1 | CD44 | 0.7909 | 1.1046 | 0.9477 | 0.000999 |
| NKT | RTMLA | HLA-DRA | CD4 | 0.6626 | 0.6388 | 0.6507 | 0.000999 |
| NKT | RTMLA | HMGB1 | LY96 | 1.195 | 1.2708 | 1.2329 | 0.000999 |
| NKT | RTMLA | HMGB2 | CD44 | 0.6467 | 1.1046 | 0.8757 | 0.000999 |
| NKT | RTMLA | HMGB2 | DAB2 | 0.6467 | 1.1614 | 0.9041 | 0.000999 |
| NKT | RTMLA | HSP90B1 | LRP1 | 0.6138 | 0.5181 | 0.566 | 0.000999 |
| NKT | RTMLA | IL16 | CD4 | 0.3556 | 0.6388 | 0.4972 | 0.000999 |
| NKT | RTMLA | IL16 | FCGR1A | 0.3556 | 0.4902 | 0.4229 | 0.000999 |
| NKT | RTMLA | ITGA4 | CD44 | 0.2946 | 1.1046 | 0.6996 | 0.000999 |
| NKT | RTMLA | ITGA4 | TFRC | 0.2946 | 0.5963 | 0.4454 | 0.000999 |
| NKT | RTMLA | ITGB1 | CD36 | 0.488 | 0.3707 | 0.4293 | 0.000999 |
| NKT | RTMLA | ITGB1 | DAB2 | 0.488 | 1.1614 | 0.8247 | 0.000999 |
| NKT | RTMLA | ITGB1 | ENG | 0.488 | 0.3636 | 0.4258 | 0.000999 |
| NKT | RTMLA | ITGB1 | ITGAX | 0.488 | 0.257 | 0.3725 | 0.000999 |
| NKT | RTMLA | ITGB1 | PLAUR | 0.488 | 0.7331 | 0.6106 | 0.000999 |
| NKT | RTMLA | ITGB2 | ITGAX | 1.6481 | 0.257 | 0.9525 | 0.000999 |
| NKT | RTMLA | ITGB2 | PLAUR | 1.6481 | 0.7331 | 1.1906 | 0.000999 |
| NKT | RTMLA | LRPAP1 | LRP1 | 0.2807 | 0.5181 | 0.3994 | 0.000999 |
| NKT | RTMLA | LTB | TNFRSF1A | 0.2605 | 0.3864 | 0.3234 | 0.000999 |
| NKT | RTMLA | MIF | CD74 | 0.6657 | 4.4485 | 2.5571 | 0.000999 |
| NKT | RTMLA | PSAP | LRP1 | 0.6777 | 0.5181 | 0.5979 | 0.000999 |
| NKT | RTMLA | PTPRC | CD4 | 1.8797 | 0.6388 | 1.2592 | 0.000999 |
| NKT | RTMLA | PTPRC | CD44 | 1.8797 | 1.1046 | 1.4921 | 0.000999 |
| NKT | RTMLA | RPS19 | C5AR1 | 3.5336 | 0.6059 | 2.0698 | 0.000999 |
| NKT | RTMLA | RPS19 | CD4 | 3.5336 | 0.6388 | 2.0862 | 0.000999 |
| NKT | RTMLA | RTN4 | CD44 | 0.4374 | 1.1046 | 0.771 | 0.000999 |
| NKT | RTMLA | TGFB1 | ENG | 0.5441 | 0.3636 | 0.4538 | 0.000999 |
| NKT | RTMLA | TIMP1 | LRP1 | 0.3005 | 0.5181 | 0.4093 | 0.000999 |
| NKT | RTMLA | XCL2 | PTAFR | 0.2774 | 0.2762 | 0.2768 | 0.000999 |
| NKT | RTMLA | ITGB1 | CD46 | 0.488 | 0.3134 | 0.4007 | 0.002 |

| | | | | | | | |
|---|---|---|---|---|---|---|---|
| NKT | RTMLA | TGFB1 | TGFBR2 | 0.5441 | 0.3296 | 0.4369 | 0.002 |
| NKT | RTMLA | CD320 | TGFBR2 | 0.2545 | 0.3296 | 0.2921 | 0.003 |
| NKT | RTMLA | ITGB2 | ITGAM | 1.6481 | 0.2768 | 0.9625 | 0.00999 |
| TCD4ex | RTMLA | ALOX5AP | ALOX5 | 1.119 | 0.3103 | 0.7146 | 0.000999 |
| TCD4ex | RTMLA | ANXA1 | C3AR1 | 0.5174 | 0.501 | 0.5092 | 0.000999 |
| TCD4ex | RTMLA | ANXA1 | C5AR1 | 0.5174 | 0.6059 | 0.5617 | 0.000999 |
| TCD4ex | RTMLA | ANXA1 | CCR1 | 0.5174 | 0.3215 | 0.4195 | 0.000999 |
| TCD4ex | RTMLA | ANXA1 | FPR1 | 0.5174 | 0.3438 | 0.4306 | 0.000999 |
| TCD4ex | RTMLA | ANXA1 | FPR3 | 0.5174 | 0.6513 | 0.5843 | 0.000999 |
| TCD4ex | RTMLA | APOE | LRP1 | 0.2557 | 0.5181 | 0.3869 | 0.000999 |
| TCD4ex | RTMLA | CALM1 | TNFRSF1A | 1.2957 | 0.3864 | 0.8411 | 0.000999 |
| TCD4ex | RTMLA | CALR | LRP1 | 0.748 | 0.5181 | 0.633 | 0.000999 |
| TCD4ex | RTMLA | CD2 | CD4 | 1.5731 | 0.6388 | 1.106 | 0.000999 |
| TCD4ex | RTMLA | CXCL13 | C3AR1 | 2.3367 | 0.501 | 1.4189 | 0.000999 |
| TCD4ex | RTMLA | CXCL13 | C5AR1 | 2.3367 | 0.6059 | 1.4713 | 0.000999 |
| TCD4ex | RTMLA | CXCL13 | CCR1 | 2.3367 | 0.3215 | 1.3291 | 0.000999 |
| TCD4ex | RTMLA | CXCL13 | FPR1 | 2.3367 | 0.3438 | 1.3402 | 0.000999 |
| TCD4ex | RTMLA | HMGB1 | LY96 | 1.3876 | 1.2708 | 1.3292 | 0.000999 |
| TCD4ex | RTMLA | HMGB2 | DAB2 | 1.0243 | 1.1614 | 1.0929 | 0.000999 |
| TCD4ex | RTMLA | HSP90B1 | LRP1 | 0.3944 | 0.5181 | 0.4562 | 0.000999 |
| TCD4ex | RTMLA | ICAM3 | ITGB2 | 0.573 | 1.2598 | 0.9164 | 0.000999 |
| TCD4ex | RTMLA | ITGB1 | CD36 | 0.4615 | 0.3707 | 0.4161 | 0.000999 |
| TCD4ex | RTMLA | ITGB1 | CD47 | 0.4615 | 0.532 | 0.4968 | 0.000999 |
| TCD4ex | RTMLA | ITGB1 | DAB2 | 0.4615 | 1.1614 | 0.8114 | 0.000999 |
| TCD4ex | RTMLA | ITGB1 | ENG | 0.4615 | 0.3636 | 0.4125 | 0.000999 |
| TCD4ex | RTMLA | ITGB1 | ITGAX | 0.4615 | 0.257 | 0.3592 | 0.000999 |
| TCD4ex | RTMLA | ITGB1 | PLAUR | 0.4615 | 0.7331 | 0.5973 | 0.000999 |
| TCD4ex | RTMLA | ITGB2 | ITGAM | 0.7871 | 0.2768 | 0.5319 | 0.000999 |
| TCD4ex | RTMLA | ITGB2 | ITGAX | 0.7871 | 0.257 | 0.522 | 0.000999 |
| TCD4ex | RTMLA | ITGB2 | PLAUR | 0.7871 | 0.7331 | 0.7601 | 0.000999 |

| | | | | | | | |
|---|---|---|---|---|---|---|---|
| TCD4ex | RTMLA | LRPAP1 | LRP1 | 0.2809 | 0.5181 | 0.3995 | 0.000999 |
| TCD4ex | RTMLA | LTB | TNFRSF1A | 0.9655 | 0.3864 | 0.676 | 0.000999 |
| TCD4ex | RTMLA | MIF | CD74 | 0.6999 | 4.4485 | 2.5742 | 0.000999 |
| TCD4ex | RTMLA | NMB | PTAFR | 0.6391 | 0.2762 | 0.4576 | 0.000999 |
| TCD4ex | RTMLA | PSAP | LRP1 | 0.5291 | 0.5181 | 0.5236 | 0.000999 |
| TCD4ex | RTMLA | PTPRC | CD4 | 1.7493 | 0.6388 | 1.1941 | 0.000999 |
| TCD4ex | RTMLA | RPS19 | C5AR1 | 3.6329 | 0.6059 | 2.1194 | 0.000999 |
| TCD4ex | RTMLA | TIMP1 | LRP1 | 0.3374 | 0.5181 | 0.4278 | 0.000999 |
| TCD4ex | RTMLA | PTPRC | CD44 | 1.7493 | 1.1046 | 1.427 | 0.002 |
| TCD4ex | RTMLA | ITGB1 | CD46 | 0.4615 | 0.3134 | 0.3875 | 0.00999 |
| TCD4reg | RTMLA | ALOX5AP | ALOX5 | 0.4589 | 0.3103 | 0.3846 | 0.000999 |
| TCD4reg | RTMLA | APOE | LRP1 | 0.2934 | 0.5181 | 0.4058 | 0.000999 |
| TCD4reg | RTMLA | ARF1 | CD4 | 0.7185 | 0.6388 | 0.6787 | 0.000999 |
| TCD4reg | RTMLA | CALM1 | TNFRSF1A | 1.534 | 0.3864 | 0.9602 | 0.000999 |
| TCD4reg | RTMLA | CALR | LRP1 | 0.6835 | 0.5181 | 0.6008 | 0.000999 |
| TCD4reg | RTMLA | CD2 | CD4 | 1.6566 | 0.6388 | 1.1477 | 0.000999 |
| TCD4reg | RTMLA | CD28 | CD4 | 0.313 | 0.6388 | 0.4759 | 0.000999 |
| TCD4reg | RTMLA | HMGB1 | LY96 | 1.2547 | 1.2708 | 1.2628 | 0.000999 |
| TCD4reg | RTMLA | HMGB2 | DAB2 | 0.616 | 1.1614 | 0.8887 | 0.000999 |
| TCD4reg | RTMLA | HSP90B1 | LRP1 | 0.4221 | 0.5181 | 0.4701 | 0.000999 |
| TCD4reg | RTMLA | ICAM3 | ITGB2 | 0.6663 | 1.2598 | 0.963 | 0.000999 |
| TCD4reg | RTMLA | IL16 | CD4 | 0.2528 | 0.6388 | 0.4458 | 0.000999 |
| TCD4reg | RTMLA | IL16 | FCGR1A | 0.2528 | 0.4902 | 0.3715 | 0.000999 |
| TCD4reg | RTMLA | ITGB1 | CD36 | 0.4001 | 0.3707 | 0.3854 | 0.000999 |
| TCD4reg | RTMLA | ITGB1 | DAB2 | 0.4001 | 1.1614 | 0.7807 | 0.000999 |
| TCD4reg | RTMLA | ITGB1 | ENG | 0.4001 | 0.3636 | 0.3818 | 0.000999 |
| TCD4reg | RTMLA | ITGB1 | ITGAX | 0.4001 | 0.257 | 0.3285 | 0.000999 |
| TCD4reg | RTMLA | ITGB1 | PLAUR | 0.4001 | 0.7331 | 0.5666 | 0.000999 |
| TCD4reg | RTMLA | ITGB2 | ITGAM | 0.6238 | 0.2768 | 0.4503 | 0.000999 |
| TCD4reg | RTMLA | ITGB2 | ITGAX | 0.6238 | 0.257 | 0.4404 | 0.000999 |

| | | | | | | | |
|---|---|---|---|---|---|---|---|
| TCD4reg | RTMLA | ITGB2 | PLAUR | 0.6238 | 0.7331 | 0.6785 | 0.000999 |
| TCD4reg | RTMLA | LGALS3 | CLEC7A | 0.4492 | 0.4847 | 0.4669 | 0.000999 |
| TCD4reg | RTMLA | LGALS3 | FCGR2A | 0.4492 | 1.0354 | 0.7423 | 0.000999 |
| TCD4reg | RTMLA | LGALS3 | FCGR2B | 0.4492 | 0.7235 | 0.5863 | 0.000999 |
| TCD4reg | RTMLA | LGALS3 | TFRC | 0.4492 | 0.5963 | 0.5227 | 0.000999 |
| TCD4reg | RTMLA | LGALS3 | TGFBR2 | 0.4492 | 0.3296 | 0.3894 | 0.000999 |
| TCD4reg | RTMLA | LRPAP1 | LRP1 | 0.3101 | 0.5181 | 0.4141 | 0.000999 |
| TCD4reg | RTMLA | LTB | TNFRSF1A | 2.242 | 0.3864 | 1.3142 | 0.000999 |
| TCD4reg | RTMLA | MIF | CD74 | 0.6295 | 4.4485 | 2.539 | 0.000999 |
| TCD4reg | RTMLA | PSAP | LRP1 | 0.5938 | 0.5181 | 0.556 | 0.000999 |
| TCD4reg | RTMLA | PTPRC | CD4 | 1.6603 | 0.6388 | 1.1496 | 0.000999 |
| TCD4reg | RTMLA | RPS19 | C5AR1 | 3.6897 | 0.6059 | 2.1478 | 0.000999 |
| TCD4reg | RTMLA | RPS19 | CD4 | 3.6897 | 0.6388 | 2.1643 | 0.000999 |
| TCD4reg | RTMLA | SELL | CD4 | 0.7062 | 0.6388 | 0.6725 | 0.000999 |
| TCD4reg | RTMLA | TIMP1 | LRP1 | 0.4362 | 0.5181 | 0.4771 | 0.000999 |
| TCD4reg | RTMLA | ITGB1 | CD46 | 0.4001 | 0.3134 | 0.3568 | 0.002 |
| TCD4reg | RTMLA | HLA-DMA | CD4 | 0.278 | 0.6388 | 0.4584 | 0.003 |
| TCD4reg | RTMLA | CD99 | CD99 | 0.6603 | 0.805 | 0.7327 | 0.00599 |
| TCD4reg | RTMLA | ITGB1 | CD47 | 0.4001 | 0.532 | 0.4661 | 0.00799 |
| TCD8em | RTMLA | ALOX5AP | ALOX5 | 0.7973 | 0.3103 | 0.5538 | 0.000999 |
| TCD8em | RTMLA | ANXA1 | C3AR1 | 1.0693 | 0.501 | 0.7851 | 0.000999 |
| TCD8em | RTMLA | ANXA1 | C5AR1 | 1.0693 | 0.6059 | 0.8376 | 0.000999 |
| TCD8em | RTMLA | ANXA1 | CCR1 | 1.0693 | 0.3215 | 0.6954 | 0.000999 |
| TCD8em | RTMLA | ANXA1 | CD4 | 1.0693 | 0.6388 | 0.854 | 0.000999 |
| TCD8em | RTMLA | ANXA1 | FPR1 | 1.0693 | 0.3438 | 0.7065 | 0.000999 |
| TCD8em | RTMLA | ANXA1 | FPR3 | 1.0693 | 0.6513 | 0.8603 | 0.000999 |
| TCD8em | RTMLA | APOE | LRP1 | 0.2791 | 0.5181 | 0.3986 | 0.000999 |
| TCD8em | RTMLA | ARF1 | CD4 | 0.6684 | 0.6388 | 0.6536 | 0.000999 |
| TCD8em | RTMLA | CALM1 | TNFRSF1A | 1.5262 | 0.3864 | 0.9563 | 0.000999 |
| TCD8em | RTMLA | CALR | LRP1 | 0.7141 | 0.5181 | 0.6161 | 0.000999 |

| | | | | | | | |
|---|---|---|---|---|---|---|---|
| TCD8em | RTMLA | CD2 | CD4 | 1.5961 | 0.6388 | 1.1175 | 0.000999 |
| TCD8em | RTMLA | HLA-DMA | CD4 | 0.2969 | 0.6388 | 0.4679 | 0.000999 |
| TCD8em | RTMLA | HLA-DRA | CD4 | 0.952 | 0.6388 | 0.7954 | 0.000999 |
| TCD8em | RTMLA | HMGB1 | LY96 | 1.3636 | 1.2708 | 1.3172 | 0.000999 |
| TCD8em | RTMLA | HMGB2 | DAB2 | 0.7212 | 1.1614 | 0.9413 | 0.000999 |
| TCD8em | RTMLA | HSP90B1 | LRP1 | 0.471 | 0.5181 | 0.4946 | 0.000999 |
| TCD8em | RTMLA | ICAM3 | ITGB2 | 0.5282 | 1.2598 | 0.894 | 0.000999 |
| TCD8em | RTMLA | IFNG | IFNGR1 | 0.3967 | 0.611 | 0.5039 | 0.000999 |
| TCD8em | RTMLA | IFNG | IFNGR2 | 0.3967 | 0.5888 | 0.4928 | 0.000999 |
| TCD8em | RTMLA | IL16 | CD4 | 0.4085 | 0.6388 | 0.5236 | 0.000999 |
| TCD8em | RTMLA | IL16 | FCGR1A | 0.4085 | 0.4902 | 0.4493 | 0.000999 |
| TCD8em | RTMLA | ITGA4 | TFRC | 0.2823 | 0.5963 | 0.4393 | 0.000999 |
| TCD8em | RTMLA | ITGB1 | CD36 | 0.2557 | 0.3707 | 0.3132 | 0.000999 |
| TCD8em | RTMLA | ITGB1 | CD46 | 0.2557 | 0.3134 | 0.2846 | 0.000999 |
| TCD8em | RTMLA | ITGB1 | DAB2 | 0.2557 | 1.1614 | 0.7086 | 0.000999 |
| TCD8em | RTMLA | ITGB1 | ENG | 0.2557 | 0.3636 | 0.3097 | 0.000999 |
| TCD8em | RTMLA | ITGB1 | ITGAX | 0.2557 | 0.257 | 0.2564 | 0.000999 |
| TCD8em | RTMLA | ITGB1 | PLAUR | 0.2557 | 0.7331 | 0.4944 | 0.000999 |
| TCD8em | RTMLA | ITGB2 | ITGAM | 1.0018 | 0.2768 | 0.6393 | 0.000999 |
| TCD8em | RTMLA | ITGB2 | ITGAX | 1.0018 | 0.257 | 0.6294 | 0.000999 |
| TCD8em | RTMLA | ITGB2 | PLAUR | 1.0018 | 0.7331 | 0.8675 | 0.000999 |
| TCD8em | RTMLA | LGALS3 | CLEC7A | 0.2689 | 0.4847 | 0.3768 | 0.000999 |
| TCD8em | RTMLA | LGALS3 | FCGR2A | 0.2689 | 1.0354 | 0.6521 | 0.000999 |
| TCD8em | RTMLA | LGALS3 | FCGR2B | 0.2689 | 0.7235 | 0.4962 | 0.000999 |
| TCD8em | RTMLA | LGALS3 | TFRC | 0.2689 | 0.5963 | 0.4326 | 0.000999 |
| TCD8em | RTMLA | LRPAP1 | LRP1 | 0.2719 | 0.5181 | 0.395 | 0.000999 |
| TCD8em | RTMLA | LTB | TNFRSF1A | 0.6352 | 0.3864 | 0.5108 | 0.000999 |
| TCD8em | RTMLA | MIF | CD74 | 0.6063 | 4.4485 | 2.5274 | 0.000999 |
| TCD8em | RTMLA | PSAP | LRP1 | 0.5012 | 0.5181 | 0.5097 | 0.000999 |
| TCD8em | RTMLA | PTPRC | CD4 | 1.7912 | 0.6388 | 1.215 | 0.000999 |

| | | | | | | | |
|---|---|---|---|---|---|---|---|
| TCD8em | RTMLA | RPS19 | C5AR1 | 3.536 | 0.6059 | 2.071 | 0.000999 |
| TCD8em | RTMLA | RPS19 | CD4 | 3.536 | 0.6388 | 2.0874 | 0.000999 |
| TCD8em | RTMLA | SELL | CD4 | 0.258 | 0.6388 | 0.4484 | 0.000999 |
| TCD8em | RTMLA | TIMP1 | LRP1 | 0.2667 | 0.5181 | 0.3924 | 0.000999 |
| TCD8em | RTMLA | TNF | TNFRSF1A | 0.2667 | 0.3864 | 0.3266 | 0.000999 |
| TCD8em | RTMLA | XCL2 | PTAFR | 0.295 | 0.2762 | 0.2856 | 0.000999 |
| TCD8ex | RTMLA | ALOX5AP | ALOX5 | 1.5024 | 0.3103 | 0.9063 | 0.000999 |
| TCD8ex | RTMLA | ANXA1 | C3AR1 | 0.5645 | 0.501 | 0.5328 | 0.000999 |
| TCD8ex | RTMLA | ANXA1 | C5AR1 | 0.5645 | 0.6059 | 0.5852 | 0.000999 |
| TCD8ex | RTMLA | ANXA1 | CCR1 | 0.5645 | 0.3215 | 0.443 | 0.000999 |
| TCD8ex | RTMLA | ANXA1 | CD4 | 0.5645 | 0.6388 | 0.6017 | 0.000999 |
| TCD8ex | RTMLA | ANXA1 | CD44 | 0.5645 | 1.1046 | 0.8345 | 0.000999 |
| TCD8ex | RTMLA | ANXA1 | FPR1 | 0.5645 | 0.3438 | 0.4542 | 0.000999 |
| TCD8ex | RTMLA | ANXA1 | FPR3 | 0.5645 | 0.6513 | 0.6079 | 0.000999 |
| TCD8ex | RTMLA | APOE | LRP1 | 0.2814 | 0.5181 | 0.3997 | 0.000999 |
| TCD8ex | RTMLA | ARF1 | CD4 | 0.7835 | 0.6388 | 0.7112 | 0.000999 |
| TCD8ex | RTMLA | ARF1 | CD44 | 0.7835 | 1.1046 | 0.9441 | 0.000999 |
| TCD8ex | RTMLA | CALM1 | TNFRSF1A | 1.4627 | 0.3864 | 0.9246 | 0.000999 |
| TCD8ex | RTMLA | CALR | CD44 | 0.9634 | 1.1046 | 1.034 | 0.000999 |
| TCD8ex | RTMLA | CALR | LRP1 | 0.9634 | 0.5181 | 0.7407 | 0.000999 |
| TCD8ex | RTMLA | CD2 | CD4 | 2.0502 | 0.6388 | 1.3445 | 0.000999 |
| TCD8ex | RTMLA | CXCL13 | C3AR1 | 1.7749 | 0.501 | 1.138 | 0.000999 |
| TCD8ex | RTMLA | CXCL13 | C5AR1 | 1.7749 | 0.6059 | 1.1904 | 0.000999 |
| TCD8ex | RTMLA | CXCL13 | CCR1 | 1.7749 | 0.3215 | 1.0482 | 0.000999 |
| TCD8ex | RTMLA | CXCL13 | FPR1 | 1.7749 | 0.3438 | 1.0594 | 0.000999 |
| TCD8ex | RTMLA | FASLG | TNFRSF1A | 0.2613 | 0.3864 | 0.3238 | 0.000999 |
| TCD8ex | RTMLA | GSTP1 | CD44 | 0.8286 | 1.1046 | 0.9666 | 0.000999 |
| TCD8ex | RTMLA | HLA-DMA | CD4 | 0.3991 | 0.6388 | 0.519 | 0.000999 |
| TCD8ex | RTMLA | HLA-DRA | CD4 | 1.247 | 0.6388 | 0.9429 | 0.000999 |
| TCD8ex | RTMLA | HMGB1 | LY96 | 1.5414 | 1.2708 | 1.4061 | 0.000999 |

| | | | | | | | |
|---|---|---|---|---|---|---|---|
| TCD8ex | RTMLA | HMGB2 | CD44 | 0.85 | 1.1046 | 0.9773 | 0.000999 |
| TCD8ex | RTMLA | HMGB2 | DAB2 | 0.85 | 1.1614 | 1.0057 | 0.000999 |
| TCD8ex | RTMLA | HSP90B1 | LRP1 | 0.5666 | 0.5181 | 0.5424 | 0.000999 |
| TCD8ex | RTMLA | ICAM3 | ITGB2 | 0.5405 | 1.2598 | 0.9001 | 0.000999 |
| TCD8ex | RTMLA | IFNG | IFNGR1 | 0.8575 | 0.611 | 0.7343 | 0.000999 |
| TCD8ex | RTMLA | IFNG | IFNGR2 | 0.8575 | 0.5888 | 0.7232 | 0.000999 |
| TCD8ex | RTMLA | ITGA4 | CD44 | 0.4432 | 1.1046 | 0.7739 | 0.000999 |
| TCD8ex | RTMLA | ITGA4 | TFRC | 0.4432 | 0.5963 | 0.5197 | 0.000999 |
| TCD8ex | RTMLA | ITGB1 | CD36 | 0.2995 | 0.3707 | 0.3351 | 0.000999 |
| TCD8ex | RTMLA | ITGB1 | DAB2 | 0.2995 | 1.1614 | 0.7304 | 0.000999 |
| TCD8ex | RTMLA | ITGB1 | ENG | 0.2995 | 0.3636 | 0.3315 | 0.000999 |
| TCD8ex | RTMLA | ITGB1 | ITGAX | 0.2995 | 0.257 | 0.2782 | 0.000999 |
| TCD8ex | RTMLA | ITGB1 | PLAUR | 0.2995 | 0.7331 | 0.5163 | 0.000999 |
| TCD8ex | RTMLA | ITGB2 | ITGAM | 0.7635 | 0.2768 | 0.5201 | 0.000999 |
| TCD8ex | RTMLA | ITGB2 | ITGAX | 0.7635 | 0.257 | 0.5102 | 0.000999 |
| TCD8ex | RTMLA | ITGB2 | PLAUR | 0.7635 | 0.7331 | 0.7483 | 0.000999 |
| TCD8ex | RTMLA | LGALS3 | CLEC7A | 0.3824 | 0.4847 | 0.4335 | 0.000999 |
| TCD8ex | RTMLA | LGALS3 | FCGR2A | 0.3824 | 1.0354 | 0.7089 | 0.000999 |
| TCD8ex | RTMLA | LGALS3 | FCGR2B | 0.3824 | 0.7235 | 0.553 | 0.000999 |
| TCD8ex | RTMLA | LGALS3 | TFRC | 0.3824 | 0.5963 | 0.4893 | 0.000999 |
| TCD8ex | RTMLA | LRPAP1 | LRP1 | 0.2971 | 0.5181 | 0.4076 | 0.000999 |
| TCD8ex | RTMLA | LTB | TNFRSF1A | 0.4576 | 0.3864 | 0.422 | 0.000999 |
| TCD8ex | RTMLA | MIF | CD74 | 0.749 | 4.4485 | 2.5987 | 0.000999 |
| TCD8ex | RTMLA | PSAP | LRP1 | 0.5016 | 0.5181 | 0.5099 | 0.000999 |
| TCD8ex | RTMLA | PTPRC | CD4 | 1.6762 | 0.6388 | 1.1575 | 0.000999 |
| TCD8ex | RTMLA | PTPRC | CD44 | 1.6762 | 1.1046 | 1.3904 | 0.000999 |
| TCD8ex | RTMLA | RPS19 | C5AR1 | 3.8353 | 0.6059 | 2.2206 | 0.000999 |
| TCD8ex | RTMLA | RPS19 | CD4 | 3.8353 | 0.6388 | 2.2371 | 0.000999 |
| TCD8ex | RTMLA | RTN4 | CD44 | 0.3033 | 1.1046 | 0.7039 | 0.000999 |
| TCD8ex | RTMLA | SPP1 | CD44 | 0.2978 | 1.1046 | 0.7012 | 0.000999 |

| | | | | | | | |
|---|---|---|---|---|---|---|---|
| TCD8ex | RTMLA | VCAM1 | ITGB2 | 0.4462 | 1.2598 | 0.853 | 0.000999 |
| TCD8ex | RTMLA | VCAM1 | TFRC | 0.4462 | 0.5963 | 0.5212 | 0.000999 |
| TCD8ex | RTMLA | XCL2 | PTAFR | 0.3157 | 0.2762 | 0.2959 | 0.000999 |
| TCD8ex | RTMLA | ITGB1 | CD46 | 0.2995 | 0.3134 | 0.3065 | 0.002 |
| TCD8ex | RTMLA | ITGB1 | CD47 | 0.2995 | 0.532 | 0.4158 | 0.002 |
| TCD8ex | RTMLA | LGALS3 | TGFBR2 | 0.3824 | 0.3296 | 0.356 | 0.005 |
| TGD | RTMLA | ARF1 | CD4 | 0.9453 | 0.6388 | 0.792 | 0.000999 |
| TGD | RTMLA | ARF1 | CD44 | 0.9453 | 1.1046 | 1.0249 | 0.000999 |
| TGD | RTMLA | CALM1 | TNFRSF1A | 1.7587 | 0.3864 | 1.0725 | 0.000999 |
| TGD | RTMLA | CALM1 | TNFRSF1B | 1.7587 | 0.3314 | 1.045 | 0.000999 |
| TGD | RTMLA | CALR | LRP1 | 0.9627 | 0.5181 | 0.7404 | 0.000999 |
| TGD | RTMLA | CD2 | CD4 | 1.4522 | 0.6388 | 1.0455 | 0.000999 |
| TGD | RTMLA | CD2 | PTPRC | 1.4522 | 0.5032 | 0.9777 | 0.000999 |
| TGD | RTMLA | CD99 | CD99 | 1.7154 | 0.805 | 1.2602 | 0.000999 |
| TGD | RTMLA | COL1A1 | CD36 | 0.4518 | 0.3707 | 0.4113 | 0.000999 |
| TGD | RTMLA | HMGB1 | LY96 | 1.7525 | 1.2708 | 1.5116 | 0.000999 |
| TGD | RTMLA | HMGB2 | DAB2 | 0.3275 | 1.1614 | 0.7444 | 0.000999 |
| TGD | RTMLA | HSP90B1 | LRP1 | 0.9338 | 0.5181 | 0.726 | 0.000999 |
| TGD | RTMLA | ICAM1 | ITGAM | 0.2915 | 0.2768 | 0.2841 | 0.000999 |
| TGD | RTMLA | ICAM1 | ITGAX | 0.2915 | 0.257 | 0.2742 | 0.000999 |
| TGD | RTMLA | ICAM3 | ITGB2 | 0.8816 | 1.2598 | 1.0707 | 0.000999 |
| TGD | RTMLA | IL16 | CD4 | 0.9357 | 0.6388 | 0.7873 | 0.000999 |
| TGD | RTMLA | IL16 | FCGR1A | 0.9357 | 0.4902 | 0.7129 | 0.000999 |
| TGD | RTMLA | ITGA4 | CD44 | 0.3968 | 1.1046 | 0.7507 | 0.000999 |
| TGD | RTMLA | ITGA4 | TFRC | 0.3968 | 0.5963 | 0.4965 | 0.000999 |
| TGD | RTMLA | ITGB1 | CD36 | 0.3657 | 0.3707 | 0.3682 | 0.000999 |
| TGD | RTMLA | ITGB1 | DAB2 | 0.3657 | 1.1614 | 0.7635 | 0.000999 |
| TGD | RTMLA | ITGB1 | ENG | 0.3657 | 0.3636 | 0.3646 | 0.000999 |
| TGD | RTMLA | ITGB1 | PLAUR | 0.3657 | 0.7331 | 0.5494 | 0.000999 |
| TGD | RTMLA | ITGB2 | ITGAM | 1.4444 | 0.2768 | 0.8606 | 0.000999 |

| | | | | | | | |
|---|---|---|---|---|---|---|---|
| TGD | RTMLA | ITGB2 | ITGAX | 1.4444 | 0.257 | 0.8507 | 0.000999 |
| TGD | RTMLA | ITGB2 | PLAUR | 1.4444 | 0.7331 | 1.0888 | 0.000999 |
| TGD | RTMLA | LTB | ITGB1 | 1.6829 | 0.3114 | 0.9971 | 0.000999 |
| TGD | RTMLA | LTB | TNFRSF1A | 1.6829 | 0.3864 | 1.0347 | 0.000999 |
| TGD | RTMLA | MIF | CD74 | 1.6335 | 4.4485 | 3.041 | 0.000999 |
| TGD | RTMLA | MIF | TNFRSF14 | 1.6335 | 0.31 | 0.9717 | 0.000999 |
| TGD | RTMLA | PTPRC | CD4 | 1.8653 | 0.6388 | 1.2521 | 0.000999 |
| TGD | RTMLA | PTPRC | CD44 | 1.8653 | 1.1046 | 1.485 | 0.000999 |
| TGD | RTMLA | PTPRC | CXCR4 | 1.8653 | 1.0274 | 1.4464 | 0.000999 |
| TGD | RTMLA | RPS19 | C5AR1 | 3.8902 | 0.6059 | 2.2481 | 0.000999 |
| TGD | RTMLA | RPS19 | CD4 | 3.8902 | 0.6388 | 2.2645 | 0.000999 |
| TGD | RTMLA | SELL | CD4 | 0.3555 | 0.6388 | 0.4972 | 0.000999 |
| TGD | RTMLA | TGFB1 | ENG | 0.7703 | 0.3636 | 0.567 | 0.000999 |
| TGD | RTMLA | TGFB1 | ITGB1 | 0.7703 | 0.3114 | 0.5409 | 0.000999 |
| TGD | RTMLA | TGFB1 | TGFBR2 | 0.7703 | 0.3296 | 0.55 | 0.000999 |
| TGD | RTMLA | XCL1 | PTAFR | 1.5822 | 0.2762 | 0.9292 | 0.000999 |
| TGD | RTMLA | XCL2 | PTAFR | 0.4864 | 0.2762 | 0.3813 | 0.000999 |
| TGD | RTMLA | ITGB1 | ITGAX | 0.3657 | 0.257 | 0.3113 | 0.002 |
| TGD | RTMLA | HSP90B1 | TNFRSF1B | 0.9338 | 0.3314 | 0.6326 | 0.00699 |

| Supplementary Table 8: Communication results between macrophages and lymphocytes subpopulations, and malignant cells | | | | | | | |
|---|---|---|---|---|---|---|---|
| **RTM_LA vs. malignant cells** | | | | | | | |
| celltype_01 | celltype_02 | ligand | receptor | ligand_mean_expression | receptor_mean_expression | ligand_receptor_mean_expression | pvalue |
| RTMLA | breasttumor | ADAM9 | ITGB1 | 0.2698 | 0.2939 | 0.2818 | 0.000999 |
| RTMLA | breasttumor | AIMP1 | RACK1 | 0.2613 | 2.6334 | 1.4473 | 0.000999 |
| RTMLA | breasttumor | ANXA1 | CD44 | 1.1107 | 0.3247 | 0.7177 | 0.000999 |
| RTMLA | breasttumor | APOE | SCARB1 | 3.8364 | 0.2628 | 2.0496 | 0.000999 |
| RTMLA | breasttumor | APOE | SDC2 | 3.8364 | 1.071 | 2.4537 | 0.000999 |
| RTMLA | breasttumor | CALM1 | MYLK | 1.1468 | 0.4064 | 0.7766 | 0.000999 |
| RTMLA | breasttumor | CALM1 | RPSA | 1.1468 | 2.358 | 1.7524 | 0.000999 |
| RTMLA | breasttumor | CALM1 | TNFRSF1A | 1.1468 | 0.2536 | 0.7002 | 0.000999 |
| RTMLA | breasttumor | CALM2 | MYLK | 1.7861 | 0.4064 | 1.0962 | 0.000999 |
| RTMLA | breasttumor | CALM3 | MYLK | 0.9246 | 0.4064 | 0.6655 | 0.000999 |
| RTMLA | breasttumor | CD99 | CD99 | 0.805 | 0.3871 | 0.596 | 0.000999 |
| RTMLA | breasttumor | COL1A1 | ITGB1 | 0.3314 | 0.2939 | 0.3126 | 0.000999 |
| RTMLA | breasttumor | FN1 | CD44 | 1.1803 | 0.3247 | 0.7525 | 0.000999 |
| RTMLA | breasttumor | FN1 | ITGB1 | 1.1803 | 0.2939 | 0.7371 | 0.000999 |
| RTMLA | breasttumor | FN1 | RPSA | 1.1803 | 2.358 | 1.7692 | 0.000999 |
| RTMLA | breasttumor | FN1 | SDC2 | 1.1803 | 1.071 | 1.1257 | 0.000999 |
| RTMLA | breasttumor | GRN | TNFRSF1A | 2.0692 | 0.2536 | 1.1614 | 0.000999 |
| RTMLA | breasttumor | HLA-DRA | SCARB1 | 4.359 | 0.2628 | 2.3109 | 0.000999 |
| RTMLA | breasttumor | LGALS3 | ITGB1 | 1.9145 | 0.2939 | 1.1042 | 0.000999 |
| RTMLA | breasttumor | PTPRC | CD44 | 0.5031 | 0.3247 | 0.4139 | 0.000999 |
| RTMLA | breasttumor | RTN4 | CD44 | 0.9643 | 0.3247 | 0.6445 | 0.000999 |
| RTMLA | breasttumor | RTN4 | CD99 | 0.9643 | 0.3871 | 0.6757 | 0.000999 |
| RTMLA | breasttumor | SERPINA1 | SDC2 | 1.0253 | 1.071 | 1.0482 | 0.000999 |
| RTMLA | breasttumor | SPP1 | CD44 | 1.336 | 0.3247 | 0.8304 | 0.000999 |
| RTMLA | breasttumor | SPP1 | ITGB1 | 1.336 | 0.2939 | 0.815 | 0.000999 |
| RTMLA | breasttumor | THBS1 | ITGB1 | 0.2852 | 0.2939 | 0.2896 | 0.000999 |
| RTMLA | breasttumor | TNFSF13B | ITGB1 | 0.6721 | 0.2939 | 0.483 | 0.000999 |

| | | | | | | | |
|---|---|---|---|---|---|---|---|
| RTMLA | breasttumor | VEGFB | ITGB1 | 0.2653 | 0.2939 | 0.2796 | 0.000999 |
| RTMLA | colorectaltumor | ADAM9 | ITGB1 | 0.2698 | 0.4214 | 0.3456 | 0.000999 |
| RTMLA | colorectaltumor | APOE | SDC1 | 3.8364 | 0.2575 | 2.0469 | 0.000999 |
| RTMLA | colorectaltumor | CALM1 | RPSA | 1.1468 | 2.251 | 1.6989 | 0.000999 |
| RTMLA | colorectaltumor | FN1 | ITGB1 | 1.1803 | 0.4214 | 0.8009 | 0.000999 |
| RTMLA | colorectaltumor | FN1 | RPSA | 1.1803 | 2.251 | 1.7157 | 0.000999 |
| RTMLA | colorectaltumor | FN1 | SDC1 | 1.1803 | 0.2575 | 0.7189 | 0.000999 |
| RTMLA | colorectaltumor | HSP90B1 | ITGB1 | 0.8143 | 0.4214 | 0.6178 | 0.000999 |
| RTMLA | colorectaltumor | ITGB1 | CD151 | 0.3114 | 0.6439 | 0.4777 | 0.000999 |
| RTMLA | colorectaltumor | ITGB1 | RACK1 | 0.3114 | 2.7146 | 1.513 | 0.000999 |
| RTMLA | colorectaltumor | LGALS3 | ITGB1 | 1.9145 | 0.4214 | 1.168 | 0.000999 |
| RTMLA | colorectaltumor | LGALS3 | ITGB4 | 1.9145 | 0.2913 | 1.1029 | 0.000999 |
| RTMLA | colorectaltumor | PECAM1 | SDC1 | 0.3607 | 0.2575 | 0.3091 | 0.000999 |
| RTMLA | colorectaltumor | SPP1 | ITGB1 | 1.336 | 0.4214 | 0.8787 | 0.000999 |
| RTMLA | colorectaltumor | SPP1 | SDC1 | 1.336 | 0.2575 | 0.7967 | 0.000999 |
| RTMLA | colorectaltumor | THBS1 | ITGB1 | 0.2852 | 0.4214 | 0.3533 | 0.000999 |
| RTMLA | colorectaltumor | THBS1 | SDC1 | 0.2852 | 0.2575 | 0.2713 | 0.000999 |
| RTMLA | colorectaltumor | TNFSF13B | ITGB1 | 0.6721 | 0.4214 | 0.5467 | 0.000999 |
| RTMLA | colorectaltumor | VEGFB | ITGB1 | 0.2653 | 0.4214 | 0.3433 | 0.000999 |
| RTMLA | colorectaltumor | ITGB1 | CDH17 | 0.3114 | 0.254 | 0.2827 | 0.003 |
| RTMLA | colorectaltumor | ITGB1 | SDC1 | 0.3114 | 0.2575 | 0.2844 | 0.00799 |
| RTMLA | livertumor | A2M | TNFRSF14 | 0.7883 | 0.2788 | 0.5336 | 0.000999 |
| RTMLA | livertumor | ADAM9 | CDH1 | 0.2698 | 0.4896 | 0.3797 | 0.000999 |
| RTMLA | livertumor | ADAM9 | ITGA3 | 0.2698 | 0.3913 | 0.3305 | 0.000999 |
| RTMLA | livertumor | ADAM9 | ITGA6 | 0.2698 | 0.5211 | 0.3955 | 0.000999 |
| RTMLA | livertumor | ADAM9 | ITGB1 | 0.2698 | 1.6044 | 0.9371 | 0.000999 |
| RTMLA | livertumor | ADM | RAMP1 | 0.2603 | 0.4499 | 0.3551 | 0.000999 |
| RTMLA | livertumor | AIMP1 | RACK1 | 0.2613 | 2.7655 | 1.5134 | 0.000999 |
| RTMLA | livertumor | ANXA1 | CD44 | 1.1107 | 0.8016 | 0.9561 | 0.000999 |
| RTMLA | livertumor | ANXA1 | MET | 1.1107 | 0.2983 | 0.7045 | 0.000999 |

| | | | | | | | |
|---|---|---|---|---|---|---|---|
| RTMLA | livertumor | ANXA1 | PTPRF | 1.1107 | 0.384 | 0.7474 | 0.000999 |
| RTMLA | livertumor | APOE | SDC1 | 3.8364 | 0.5354 | 2.1859 | 0.000999 |
| RTMLA | livertumor | APOE | SDC4 | 3.8364 | 0.6068 | 2.2216 | 0.000999 |
| RTMLA | livertumor | CALM1 | INSR | 1.1468 | 0.3039 | 0.7254 | 0.000999 |
| RTMLA | livertumor | CALM1 | RPSA | 1.1468 | 2.6317 | 1.8892 | 0.000999 |
| RTMLA | livertumor | CD99 | EPHB4 | 0.805 | 0.4535 | 0.6292 | 0.000999 |
| RTMLA | livertumor | CD99 | INSR | 0.805 | 0.3039 | 0.5545 | 0.000999 |
| RTMLA | livertumor | COL1A1 | ITGA2 | 0.3314 | 0.3983 | 0.3648 | 0.000999 |
| RTMLA | livertumor | COL1A1 | ITGA3 | 0.3314 | 0.3913 | 0.3613 | 0.000999 |
| RTMLA | livertumor | COL1A1 | ITGB1 | 0.3314 | 1.6044 | 0.9679 | 0.000999 |
| RTMLA | livertumor | COL1A1 | SDC1 | 0.3314 | 0.5354 | 0.4334 | 0.000999 |
| RTMLA | livertumor | FN1 | CD44 | 1.1803 | 0.8016 | 0.991 | 0.000999 |
| RTMLA | livertumor | FN1 | ITGA2 | 1.1803 | 0.3983 | 0.7893 | 0.000999 |
| RTMLA | livertumor | FN1 | ITGA3 | 1.1803 | 0.3913 | 0.7858 | 0.000999 |
| RTMLA | livertumor | FN1 | ITGB1 | 1.1803 | 1.6044 | 1.3924 | 0.000999 |
| RTMLA | livertumor | FN1 | RPSA | 1.1803 | 2.6317 | 1.906 | 0.000999 |
| RTMLA | livertumor | FN1 | SDC1 | 1.1803 | 0.5354 | 0.8579 | 0.000999 |
| RTMLA | livertumor | FN1 | TFRC | 1.1803 | 0.2899 | 0.7351 | 0.000999 |
| RTMLA | livertumor | HLA-A | ERBB3 | 2.8931 | 0.2644 | 1.5787 | 0.000999 |
| RTMLA | livertumor | HLA-DRA | EPHB4 | 4.359 | 0.4535 | 2.4062 | 0.000999 |
| RTMLA | livertumor | HLA-E | EPHB4 | 1.6583 | 0.4535 | 1.0559 | 0.000999 |
| RTMLA | livertumor | HLA-E | ITGA6 | 1.6583 | 0.5211 | 1.0897 | 0.000999 |
| RTMLA | livertumor | HLA-E | PLXNB2 | 1.6583 | 0.265 | 0.9616 | 0.000999 |
| RTMLA | livertumor | HMGB2 | CDH1 | 0.3931 | 0.4896 | 0.4414 | 0.000999 |
| RTMLA | livertumor | HMGB2 | DSG2 | 0.3931 | 0.7479 | 0.5705 | 0.000999 |
| RTMLA | livertumor | HSP90B1 | ITGB1 | 0.8143 | 1.6044 | 1.2094 | 0.000999 |
| RTMLA | livertumor | LGALS3 | ITGA2 | 1.9145 | 0.3983 | 1.1564 | 0.000999 |
| RTMLA | livertumor | LGALS3 | ITGB1 | 1.9145 | 1.6044 | 1.7595 | 0.000999 |
| RTMLA | livertumor | LGALS3 | ITGB4 | 1.9145 | 0.5518 | 1.2332 | 0.000999 |
| RTMLA | livertumor | LGALS3 | TFRC | 1.9145 | 0.2899 | 1.1022 | 0.000999 |

| | | | | | | | |
|---|---|---|---|---|---|---|---|
| RTMLA | livertumor | NAMPT | INSR | 0.7571 | 0.3039 | 0.5305 | 0.000999 |
| RTMLA | livertumor | PECAM1 | SDC1 | 0.3607 | 0.5354 | 0.4481 | 0.000999 |
| RTMLA | livertumor | PECAM1 | SDC4 | 0.3607 | 0.6068 | 0.4838 | 0.000999 |
| RTMLA | livertumor | PTPRC | INSR | 0.5031 | 0.3039 | 0.4035 | 0.000999 |
| RTMLA | livertumor | PTPRC | MET | 0.5031 | 0.2983 | 0.4007 | 0.000999 |
| RTMLA | livertumor | SPP1 | ITGB1 | 1.336 | 1.6044 | 1.4702 | 0.000999 |
| RTMLA | livertumor | SPP1 | SDC1 | 1.336 | 0.5354 | 0.9357 | 0.000999 |
| RTMLA | livertumor | THBS1 | ITGA3 | 0.2852 | 0.3913 | 0.3383 | 0.000999 |
| RTMLA | livertumor | THBS1 | ITGB1 | 0.2852 | 1.6044 | 0.9448 | 0.000999 |
| RTMLA | livertumor | THBS1 | SDC1 | 0.2852 | 0.5354 | 0.4103 | 0.000999 |
| RTMLA | livertumor | TNFSF13B | ITGB1 | 0.6721 | 1.6044 | 1.1383 | 0.000999 |
| RTMLA | livertumor | VEGFB | ITGB1 | 0.2653 | 1.6044 | 0.9349 | 0.000999 |
| RTMLA | livertumor | CD99 | CD99 | 0.805 | 0.5906 | 0.6978 | 0.002 |
| RTMLA | livertumor | HAMP | SLC40A1 | 0.303 | 0.4972 | 0.4001 | 0.002 |
| RTMLA | livertumor | PTPRC | CD44 | 0.5031 | 0.8016 | 0.6524 | 0.002 |
| RTMLA | livertumor | THBS1 | CD47 | 0.2852 | 0.5289 | 0.4071 | 0.002 |
| RTMLA | livertumor | RPS19 | RPSA | 3.4667 | 2.6317 | 3.0492 | 0.00599 |
| RTMLA | lungtumor | APOE | SDC1 | 3.8364 | 0.2715 | 2.0539 | 0.000999 |
| RTMLA | lungtumor | APOE | SDC4 | 3.8364 | 0.3516 | 2.094 | 0.000999 |
| RTMLA | lungtumor | CALM1 | RPSA | 1.1468 | 1.6057 | 1.3763 | 0.000999 |
| RTMLA | lungtumor | CD99 | CD99 | 0.805 | 0.5938 | 0.6994 | 0.000999 |
| RTMLA | lungtumor | COL1A1 | CD44 | 0.3314 | 0.3802 | 0.3558 | 0.000999 |
| RTMLA | lungtumor | COL1A1 | SDC1 | 0.3314 | 0.2715 | 0.3014 | 0.000999 |
| RTMLA | lungtumor | CXCL16 | CXCR4 | 1.0236 | 0.2684 | 0.646 | 0.000999 |
| RTMLA | lungtumor | CXCL2 | CXCR4 | 0.4556 | 0.2684 | 0.362 | 0.000999 |
| RTMLA | lungtumor | CXCL3 | CXCR4 | 0.2688 | 0.2684 | 0.2686 | 0.000999 |
| RTMLA | lungtumor | FN1 | CD44 | 1.1803 | 0.3802 | 0.7803 | 0.000999 |
| RTMLA | lungtumor | FN1 | RPSA | 1.1803 | 1.6057 | 1.393 | 0.000999 |
| RTMLA | lungtumor | FN1 | SDC1 | 1.1803 | 0.2715 | 0.7259 | 0.000999 |
| RTMLA | lungtumor | PECAM1 | CXCR4 | 0.3607 | 0.2684 | 0.3146 | 0.000999 |

| | | | | | | | |
|---|---|---|---|---|---|---|---|
| RTMLA | lungtumor | PECAM1 | SDC1 | 0.3607 | 0.2715 | 0.3161 | 0.000999 |
| RTMLA | lungtumor | PECAM1 | SDC4 | 0.3607 | 0.3516 | 0.3562 | 0.000999 |
| RTMLA | lungtumor | PTPRC | CD44 | 0.5031 | 0.3802 | 0.4417 | 0.000999 |
| RTMLA | lungtumor | PTPRC | CXCR4 | 0.5031 | 0.2684 | 0.3858 | 0.000999 |
| RTMLA | lungtumor | RTN4 | CD44 | 0.9643 | 0.3802 | 0.6723 | 0.000999 |
| RTMLA | lungtumor | RTN4 | CD99 | 0.9643 | 0.5938 | 0.7791 | 0.000999 |
| RTMLA | lungtumor | THBS1 | CD47 | 0.2852 | 0.3255 | 0.3054 | 0.000999 |
| RTMLA | lungtumor | THBS1 | SDC1 | 0.2852 | 0.2715 | 0.2784 | 0.000999 |
| RTMLA | lungtumor | ITGB1 | CD151 | 0.3114 | 0.6776 | 0.4945 | 0.00999 |
| RTMLA | melanomatumo | AIMP1 | RACK1 | 0.2613 | 4.145 | 2.2032 | 0.000999 |
| RTMLA | melanomatumo | ANXA1 | EDNRB | 1.1107 | 0.5666 | 0.8387 | 0.000999 |
| RTMLA | melanomatumo | CALM1 | RPSA | 1.1468 | 3.7787 | 2.4627 | 0.000999 |
| RTMLA | melanomatumo | FN1 | RPSA | 1.1803 | 3.7787 | 2.4795 | 0.000999 |
| RTMLA | melanomatumo | ITGB1 | RACK1 | 0.3114 | 4.145 | 2.2282 | 0.000999 |
| RTMLA | melanomatumo | RPS19 | RPSA | 3.4667 | 3.7787 | 3.6227 | 0.000999 |
| RTMLA | melanomatumo | NAMPT | ANXA6 | 0.7571 | 0.3625 | 0.5598 | 0.002 |
| RTMLA | melanomatumo | SPP1 | NCSTN | 1.336 | 0.2614 | 0.7987 | 0.00699 |
| RTMLA | melanomatumo | A2M | TNFRSF14 | 0.7883 | 0.5069 | 0.6476 | 0.00799 |
| RTMLA | ovarytumor | ADAM9 | ITGB1 | 0.2698 | 0.2783 | 0.274 | 0.000999 |
| RTMLA | ovarytumor | AREG | FOLR1 | 0.3085 | 0.5619 | 0.4352 | 0.000999 |
| RTMLA | ovarytumor | CXCL16 | CXCR4 | 1.0236 | 0.3688 | 0.6962 | 0.000999 |
| RTMLA | ovarytumor | CXCL2 | CXCR4 | 0.4556 | 0.3688 | 0.4122 | 0.000999 |
| RTMLA | ovarytumor | CXCL3 | CXCR4 | 0.2688 | 0.3688 | 0.3188 | 0.000999 |
| RTMLA | ovarytumor | FN1 | ITGB1 | 1.1803 | 0.2783 | 0.7293 | 0.000999 |
| RTMLA | ovarytumor | FN1 | RPSA | 1.1803 | 2.9542 | 2.0673 | 0.000999 |
| RTMLA | ovarytumor | LGALS3 | ITGB1 | 1.9145 | 0.2783 | 1.0964 | 0.000999 |
| RTMLA | ovarytumor | PECAM1 | CXCR4 | 0.3607 | 0.3688 | 0.3647 | 0.000999 |
| RTMLA | ovarytumor | PTPRC | CXCR4 | 0.5031 | 0.3688 | 0.436 | 0.000999 |
| RTMLA | ovarytumor | SPP1 | ITGB1 | 1.336 | 0.2783 | 0.8072 | 0.000999 |
| RTMLA | ovarytumor | THBS1 | CD47 | 0.2852 | 0.4931 | 0.3892 | 0.000999 |

| | | | | | | | |
|---|---|---|---|---|---|---|---|
| RTMLA | ovarytumor | THBS1 | ITGB1 | 0.2852 | 0.2783 | 0.2818 | 0.000999 |
| RTMLA | ovarytumor | TNFSF13B | ITGB1 | 0.6721 | 0.2783 | 0.4752 | 0.000999 |
| RTMLA | ovarytumor | VEGFB | ITGB1 | 0.2653 | 0.2783 | 0.2718 | 0.000999 |
| RTMLA | ovarytumor | HSP90B1 | ITGB1 | 0.8143 | 0.2783 | 0.5463 | 0.002 |
| RTMLA | ovarytumor | CALM1 | RPSA | 1.1468 | 2.9542 | 2.0505 | 0.003 |
| breasttumor | RTMLA | A2M | LRP1 | 0.3129 | 0.5181 | 0.4155 | 0.000999 |
| breasttumor | RTMLA | A2M | TNFRSF14 | 0.3129 | 0.31 | 0.3115 | 0.000999 |
| breasttumor | RTMLA | ANXA1 | C3AR1 | 0.4092 | 0.501 | 0.4551 | 0.000999 |
| breasttumor | RTMLA | ANXA1 | C5AR1 | 0.4092 | 0.6059 | 0.5075 | 0.000999 |
| breasttumor | RTMLA | ANXA1 | CCR1 | 0.4092 | 0.3215 | 0.3653 | 0.000999 |
| breasttumor | RTMLA | ANXA1 | CD4 | 0.4092 | 0.6388 | 0.524 | 0.000999 |
| breasttumor | RTMLA | ANXA1 | CD44 | 0.4092 | 1.1045 | 0.7569 | 0.000999 |
| breasttumor | RTMLA | ANXA1 | CXCR4 | 0.4092 | 1.0274 | 0.7183 | 0.000999 |
| breasttumor | RTMLA | ANXA1 | FPR1 | 0.4092 | 0.3438 | 0.3765 | 0.000999 |
| breasttumor | RTMLA | ANXA1 | FPR3 | 0.4092 | 0.6512 | 0.5302 | 0.000999 |
| breasttumor | RTMLA | APP | ALOX5 | 0.5409 | 0.3103 | 0.4256 | 0.000999 |
| breasttumor | RTMLA | APP | C3AR1 | 0.5409 | 0.501 | 0.521 | 0.000999 |
| breasttumor | RTMLA | APP | C5AR1 | 0.5409 | 0.6059 | 0.5734 | 0.000999 |
| breasttumor | RTMLA | APP | CCR1 | 0.5409 | 0.3215 | 0.4312 | 0.000999 |
| breasttumor | RTMLA | APP | CD36 | 0.5409 | 0.3707 | 0.4558 | 0.000999 |
| breasttumor | RTMLA | APP | CXCR4 | 0.5409 | 1.0274 | 0.7842 | 0.000999 |
| breasttumor | RTMLA | APP | FPR1 | 0.5409 | 0.3438 | 0.4424 | 0.000999 |
| breasttumor | RTMLA | APP | LRP1 | 0.5409 | 0.5181 | 0.5295 | 0.000999 |
| breasttumor | RTMLA | APP | TGFBR2 | 0.5409 | 0.3296 | 0.4353 | 0.000999 |
| breasttumor | RTMLA | APP | TNFRSF14 | 0.5409 | 0.31 | 0.4255 | 0.000999 |
| breasttumor | RTMLA | ARF1 | CD4 | 1.6219 | 0.6388 | 1.1303 | 0.000999 |
| breasttumor | RTMLA | ARF1 | CD44 | 1.6219 | 1.1045 | 1.3632 | 0.000999 |
| breasttumor | RTMLA | CALM1 | TNFRSF1B | 0.9015 | 0.3313 | 0.6164 | 0.000999 |
| breasttumor | RTMLA | CALR | CD44 | 1.1835 | 1.1045 | 1.144 | 0.000999 |
| breasttumor | RTMLA | CALR | LRP1 | 1.1835 | 0.5181 | 0.8508 | 0.000999 |

| | | | | | | | |
|---|---|---|---|---|---|---|---|
| breasttumor | RTMLA | CD99 | CD99 | 0.3871 | 0.805 | 0.596 | 0.000999 |
| breasttumor | RTMLA | COL2A1 | CD44 | 0.3935 | 1.1045 | 0.749 | 0.000999 |
| breasttumor | RTMLA | GSTP1 | CD44 | 2.5435 | 1.1045 | 1.824 | 0.000999 |
| breasttumor | RTMLA | HEBP1 | C3AR1 | 0.4062 | 0.501 | 0.4536 | 0.000999 |
| breasttumor | RTMLA | HEBP1 | C5AR1 | 0.4062 | 0.6059 | 0.506 | 0.000999 |
| breasttumor | RTMLA | HEBP1 | CCR1 | 0.4062 | 0.3215 | 0.3639 | 0.000999 |
| breasttumor | RTMLA | HEBP1 | CXCR4 | 0.4062 | 1.0274 | 0.7168 | 0.000999 |
| breasttumor | RTMLA | HEBP1 | FPR1 | 0.4062 | 0.3438 | 0.375 | 0.000999 |
| breasttumor | RTMLA | HEBP1 | FPR3 | 0.4062 | 0.6512 | 0.5287 | 0.000999 |
| breasttumor | RTMLA | HMGB1 | LY96 | 1.1815 | 1.2707 | 1.2261 | 0.000999 |
| breasttumor | RTMLA | HMGB2 | CD44 | 0.3139 | 1.1045 | 0.7092 | 0.000999 |
| breasttumor | RTMLA | HMGB2 | DAB2 | 0.3139 | 1.1613 | 0.7376 | 0.000999 |
| breasttumor | RTMLA | HSP90B1 | LRP1 | 0.7786 | 0.5181 | 0.6483 | 0.000999 |
| breasttumor | RTMLA | HSP90B1 | TNFRSF1B | 0.7786 | 0.3313 | 0.555 | 0.000999 |
| breasttumor | RTMLA | ITGB1 | CD36 | 0.2939 | 0.3707 | 0.3323 | 0.000999 |
| breasttumor | RTMLA | ITGB1 | CD46 | 0.2939 | 0.3134 | 0.3037 | 0.000999 |
| breasttumor | RTMLA | ITGB1 | CD47 | 0.2939 | 0.532 | 0.4129 | 0.000999 |
| breasttumor | RTMLA | ITGB1 | DAB2 | 0.2939 | 1.1613 | 0.7276 | 0.000999 |
| breasttumor | RTMLA | ITGB1 | ENG | 0.2939 | 0.3636 | 0.3287 | 0.000999 |
| breasttumor | RTMLA | ITGB1 | ITGAX | 0.2939 | 0.257 | 0.2754 | 0.000999 |
| breasttumor | RTMLA | ITGB1 | PLAUR | 0.2939 | 0.7331 | 0.5135 | 0.000999 |
| breasttumor | RTMLA | LGALS3 | CLEC7A | 0.8391 | 0.4847 | 0.6619 | 0.000999 |
| breasttumor | RTMLA | LGALS3 | FCGR2A | 0.8391 | 1.0353 | 0.9372 | 0.000999 |
| breasttumor | RTMLA | LGALS3 | FCGR2B | 0.8391 | 0.7235 | 0.7813 | 0.000999 |
| breasttumor | RTMLA | LGALS3 | TFRC | 0.8391 | 0.5962 | 0.7177 | 0.000999 |
| breasttumor | RTMLA | LRPAP1 | LRP1 | 0.2756 | 0.5181 | 0.3968 | 0.000999 |
| breasttumor | RTMLA | MDK | LRP1 | 0.592 | 0.5181 | 0.555 | 0.000999 |
| breasttumor | RTMLA | MIF | CD74 | 3.2716 | 4.4484 | 3.86 | 0.000999 |
| breasttumor | RTMLA | MIF | TNFRSF14 | 3.2716 | 0.31 | 1.7908 | 0.000999 |
| breasttumor | RTMLA | NPW | C3AR1 | 0.5478 | 0.501 | 0.5244 | 0.000999 |

| | | | | | | | |
|---|---|---|---|---|---|---|---|
| breasttumor | RTMLA | NPW | C5AR1 | 0.5478 | 0.6059 | 0.5769 | 0.000999 |
| breasttumor | RTMLA | NPW | CCR1 | 0.5478 | 0.3215 | 0.4347 | 0.000999 |
| breasttumor | RTMLA | NPW | CXCR4 | 0.5478 | 1.0274 | 0.7876 | 0.000999 |
| breasttumor | RTMLA | NPW | FPR1 | 0.5478 | 0.3438 | 0.4458 | 0.000999 |
| breasttumor | RTMLA | RPS19 | C5AR1 | 3.691 | 0.6059 | 2.1485 | 0.000999 |
| breasttumor | RTMLA | RPS19 | CD4 | 3.691 | 0.6388 | 2.1649 | 0.000999 |
| breasttumor | RTMLA | RTN4 | CD44 | 0.3318 | 1.1045 | 0.7182 | 0.000999 |
| breasttumor | RTMLA | RTN4 | CD99 | 0.3318 | 0.805 | 0.5684 | 0.000999 |
| breasttumor | RTMLA | SERPINE2 | LRP1 | 0.6001 | 0.5181 | 0.5591 | 0.000999 |
| breasttumor | RTMLA | TF | CD4 | 0.2688 | 0.6388 | 0.4538 | 0.000999 |
| breasttumor | RTMLA | TF | TFRC | 0.2688 | 0.5962 | 0.4325 | 0.000999 |
| breasttumor | RTMLA | TIMP1 | LRP1 | 0.7045 | 0.5181 | 0.6113 | 0.000999 |
| breasttumor | RTMLA | LGALS3 | TGFBR2 | 0.8391 | 0.3296 | 0.5844 | 0.003 |
| colorectaltumor | RTMLA | ANXA1 | CXCR4 | 0.5916 | 1.0274 | 0.8095 | 0.000999 |
| colorectaltumor | RTMLA | APP | ALOX5 | 0.4966 | 0.3103 | 0.4035 | 0.000999 |
| colorectaltumor | RTMLA | APP | C3AR1 | 0.4966 | 0.501 | 0.4988 | 0.000999 |
| colorectaltumor | RTMLA | APP | C5AR1 | 0.4966 | 0.6059 | 0.5513 | 0.000999 |
| colorectaltumor | RTMLA | APP | CCR1 | 0.4966 | 0.3215 | 0.4091 | 0.000999 |
| colorectaltumor | RTMLA | APP | CD36 | 0.4966 | 0.3707 | 0.4337 | 0.000999 |
| colorectaltumor | RTMLA | APP | CXCR4 | 0.4966 | 1.0274 | 0.762 | 0.000999 |
| colorectaltumor | RTMLA | APP | FPR1 | 0.4966 | 0.3438 | 0.4202 | 0.000999 |
| colorectaltumor | RTMLA | APP | LRP1 | 0.4966 | 0.5181 | 0.5074 | 0.000999 |
| colorectaltumor | RTMLA | APP | TGFBR2 | 0.4966 | 0.3296 | 0.4131 | 0.000999 |
| colorectaltumor | RTMLA | APP | TNFRSF14 | 0.4966 | 0.31 | 0.4033 | 0.000999 |
| colorectaltumor | RTMLA | ARF1 | CD4 | 0.6489 | 0.6388 | 0.6438 | 0.000999 |
| colorectaltumor | RTMLA | ARF1 | CD44 | 0.6489 | 1.1045 | 0.8767 | 0.000999 |
| colorectaltumor | RTMLA | CALM1 | TNFRSF1B | 1.1676 | 0.3313 | 0.7495 | 0.000999 |
| colorectaltumor | RTMLA | CALR | CD44 | 0.4934 | 1.1045 | 0.799 | 0.000999 |
| colorectaltumor | RTMLA | COL1A1 | CD36 | 0.4875 | 0.3707 | 0.4291 | 0.000999 |
| colorectaltumor | RTMLA | COL1A1 | CD44 | 0.4875 | 1.1045 | 0.796 | 0.000999 |

| | | | | | | | |
|---|---|---|---|---|---|---|---|
| colorectaltumor | RTMLA | GSTP1 | CD44 | 2.8039 | 1.1045 | 1.9542 | 0.000999 |
| colorectaltumor | RTMLA | HMGB1 | LY96 | 2.155 | 1.2707 | 1.7129 | 0.000999 |
| colorectaltumor | RTMLA | HMGB2 | CD44 | 0.5347 | 1.1045 | 0.8196 | 0.000999 |
| colorectaltumor | RTMLA | HMGB2 | DAB2 | 0.5347 | 1.1613 | 0.848 | 0.000999 |
| colorectaltumor | RTMLA | ITGB1 | CD36 | 0.4214 | 0.3707 | 0.3961 | 0.000999 |
| colorectaltumor | RTMLA | ITGB1 | CD47 | 0.4214 | 0.532 | 0.4767 | 0.000999 |
| colorectaltumor | RTMLA | ITGB1 | DAB2 | 0.4214 | 1.1613 | 0.7914 | 0.000999 |
| colorectaltumor | RTMLA | ITGB1 | ENG | 0.4214 | 0.3636 | 0.3925 | 0.000999 |
| colorectaltumor | RTMLA | ITGB1 | ITGAX | 0.4214 | 0.257 | 0.3392 | 0.000999 |
| colorectaltumor | RTMLA | ITGB1 | PLAUR | 0.4214 | 0.7331 | 0.5773 | 0.000999 |
| colorectaltumor | RTMLA | LGALS3 | FCGR2A | 1.5563 | 1.0353 | 1.2958 | 0.000999 |
| colorectaltumor | RTMLA | MDK | LRP1 | 0.7548 | 0.5181 | 0.6365 | 0.000999 |
| colorectaltumor | RTMLA | MIF | CD74 | 1.6741 | 4.4484 | 3.0612 | 0.000999 |
| colorectaltumor | RTMLA | MIF | TNFRSF14 | 1.6741 | 0.31 | 0.992 | 0.000999 |
| colorectaltumor | RTMLA | RPS19 | C5AR1 | 4.0956 | 0.6059 | 2.3507 | 0.000999 |
| colorectaltumor | RTMLA | RPS19 | CD4 | 4.0956 | 0.6388 | 2.3672 | 0.000999 |
| colorectaltumor | RTMLA | RPS19 | RPSA | 4.0956 | 1.7725 | 2.9341 | 0.004 |
| colorectaltumor | RTMLA | LGALS3 | FCGR2B | 1.5563 | 0.7235 | 1.1399 | 0.00599 |
| colorectaltumor | RTMLA | CALM1 | TNFRSF1A | 1.1676 | 0.3864 | 0.777 | 0.00799 |
| livertumor | RTMLA | AGRN | LRP1 | 0.3492 | 0.5181 | 0.4337 | 0.000999 |
| livertumor | RTMLA | AGT | ATP6AP2 | 0.4701 | 0.8973 | 0.6837 | 0.000999 |
| livertumor | RTMLA | AGT | C3AR1 | 0.4701 | 0.501 | 0.4855 | 0.000999 |
| livertumor | RTMLA | AGT | C5AR1 | 0.4701 | 0.6059 | 0.538 | 0.000999 |
| livertumor | RTMLA | AGT | CCR1 | 0.4701 | 0.3215 | 0.3958 | 0.000999 |
| livertumor | RTMLA | AGT | CXCR4 | 0.4701 | 1.0274 | 0.7487 | 0.000999 |
| livertumor | RTMLA | AGT | FPR1 | 0.4701 | 0.3438 | 0.4069 | 0.000999 |
| livertumor | RTMLA | AGT | PTAFR | 0.4701 | 0.2762 | 0.3731 | 0.000999 |
| livertumor | RTMLA | ANXA1 | CXCR4 | 0.3041 | 1.0274 | 0.6658 | 0.000999 |
| livertumor | RTMLA | APOA1 | ABCA1 | 0.2888 | 0.4677 | 0.3783 | 0.000999 |
| livertumor | RTMLA | APOA1 | CD36 | 0.2888 | 0.3707 | 0.3297 | 0.000999 |

| | | | | | | | |
|---|---|---|---|---|---|---|---|
| livertumor | RTMLA | APOA1 | LRP1 | 0.2888 | 0.5181 | 0.4034 | 0.000999 |
| livertumor | RTMLA | APOA1 | MSR1 | 0.2888 | 0.868 | 0.5784 | 0.000999 |
| livertumor | RTMLA | APP | ALOX5 | 0.5626 | 0.3103 | 0.4364 | 0.000999 |
| livertumor | RTMLA | APP | C3AR1 | 0.5626 | 0.501 | 0.5318 | 0.000999 |
| livertumor | RTMLA | APP | C5AR1 | 0.5626 | 0.6059 | 0.5842 | 0.000999 |
| livertumor | RTMLA | APP | CCR1 | 0.5626 | 0.3215 | 0.442 | 0.000999 |
| livertumor | RTMLA | APP | CD36 | 0.5626 | 0.3707 | 0.4666 | 0.000999 |
| livertumor | RTMLA | APP | CXCR4 | 0.5626 | 1.0274 | 0.795 | 0.000999 |
| livertumor | RTMLA | APP | FPR1 | 0.5626 | 0.3438 | 0.4532 | 0.000999 |
| livertumor | RTMLA | APP | LRP1 | 0.5626 | 0.5181 | 0.5403 | 0.000999 |
| livertumor | RTMLA | APP | TGFBR2 | 0.5626 | 0.3296 | 0.4461 | 0.000999 |
| livertumor | RTMLA | APP | TNFRSF14 | 0.5626 | 0.31 | 0.4363 | 0.000999 |
| livertumor | RTMLA | ARF1 | CD4 | 0.6936 | 0.6388 | 0.6662 | 0.000999 |
| livertumor | RTMLA | ARF1 | CD44 | 0.6936 | 1.1045 | 0.8991 | 0.000999 |
| livertumor | RTMLA | C3 | C3AR1 | 0.6584 | 0.501 | 0.5797 | 0.000999 |
| livertumor | RTMLA | C3 | C5AR1 | 0.6584 | 0.6059 | 0.6322 | 0.000999 |
| livertumor | RTMLA | C3 | CCR1 | 0.6584 | 0.3215 | 0.49 | 0.000999 |
| livertumor | RTMLA | C3 | CXCR4 | 0.6584 | 1.0274 | 0.8429 | 0.000999 |
| livertumor | RTMLA | C3 | FPR1 | 0.6584 | 0.3438 | 0.5011 | 0.000999 |
| livertumor | RTMLA | C3 | ITGAM | 0.6584 | 0.2768 | 0.4676 | 0.000999 |
| livertumor | RTMLA | C3 | ITGAX | 0.6584 | 0.257 | 0.4577 | 0.000999 |
| livertumor | RTMLA | C3 | ITGB2 | 0.6584 | 1.2597 | 0.9591 | 0.000999 |
| livertumor | RTMLA | C3 | LRP1 | 0.6584 | 0.5181 | 0.5883 | 0.000999 |
| livertumor | RTMLA | CALR | CD44 | 1.0416 | 1.1045 | 1.0731 | 0.000999 |
| livertumor | RTMLA | CALR | LRP1 | 1.0416 | 0.5181 | 0.7799 | 0.000999 |
| livertumor | RTMLA | CCL20 | C3AR1 | 1.0877 | 0.501 | 0.7943 | 0.000999 |
| livertumor | RTMLA | CCL20 | C5AR1 | 1.0877 | 0.6059 | 0.8468 | 0.000999 |
| livertumor | RTMLA | CCL20 | CCR1 | 1.0877 | 0.3215 | 0.7046 | 0.000999 |
| livertumor | RTMLA | CCL20 | CXCR4 | 1.0877 | 1.0274 | 1.0575 | 0.000999 |
| livertumor | RTMLA | CCL20 | FPR1 | 1.0877 | 0.3438 | 0.7157 | 0.000999 |

| | | | | | | | |
|---|---|---|---|---|---|---|---|
| livertumor | RTMLA | CFH | ITGAM | 0.3373 | 0.2768 | 0.3071 | 0.000999 |
| livertumor | RTMLA | CXCL2 | C3AR1 | 0.3364 | 0.501 | 0.4187 | 0.000999 |
| livertumor | RTMLA | CXCL2 | C5AR1 | 0.3364 | 0.6059 | 0.4711 | 0.000999 |
| livertumor | RTMLA | CXCL2 | CCR1 | 0.3364 | 0.3215 | 0.3289 | 0.000999 |
| livertumor | RTMLA | CXCL2 | CXCR4 | 0.3364 | 1.0274 | 0.6819 | 0.000999 |
| livertumor | RTMLA | CXCL2 | FPR1 | 0.3364 | 0.3438 | 0.3401 | 0.000999 |
| livertumor | RTMLA | CXCL3 | C3AR1 | 0.2543 | 0.501 | 0.3776 | 0.000999 |
| livertumor | RTMLA | CXCL3 | C5AR1 | 0.2543 | 0.6059 | 0.4301 | 0.000999 |
| livertumor | RTMLA | CXCL3 | CCR1 | 0.2543 | 0.3215 | 0.2879 | 0.000999 |
| livertumor | RTMLA | CXCL3 | CXCR4 | 0.2543 | 1.0274 | 0.6408 | 0.000999 |
| livertumor | RTMLA | CXCL3 | FPR1 | 0.2543 | 0.3438 | 0.299 | 0.000999 |
| livertumor | RTMLA | DSG2 | CD44 | 0.7479 | 1.1045 | 0.9262 | 0.000999 |
| livertumor | RTMLA | DSG2 | CXCR4 | 0.7479 | 1.0274 | 0.8877 | 0.000999 |
| livertumor | RTMLA | DSG2 | DAB2 | 0.7479 | 1.1613 | 0.9546 | 0.000999 |
| livertumor | RTMLA | FGA | ITGAX | 0.2688 | 0.257 | 0.2629 | 0.000999 |
| livertumor | RTMLA | FGA | ITGB2 | 0.2688 | 1.2597 | 0.7643 | 0.000999 |
| livertumor | RTMLA | HMGB1 | LY96 | 1.4361 | 1.2707 | 1.3534 | 0.000999 |
| livertumor | RTMLA | HSP90B1 | LRP1 | 0.8951 | 0.5181 | 0.7066 | 0.000999 |
| livertumor | RTMLA | HSP90B1 | TNFRSF1B | 0.8951 | 0.3313 | 0.6132 | 0.000999 |
| livertumor | RTMLA | HSPG2 | LRP1 | 0.2746 | 0.5181 | 0.3964 | 0.000999 |
| livertumor | RTMLA | ITGB1 | CD36 | 1.6044 | 0.3707 | 0.9876 | 0.000999 |
| livertumor | RTMLA | ITGB1 | CD46 | 1.6044 | 0.3134 | 0.9589 | 0.000999 |
| livertumor | RTMLA | ITGB1 | CD47 | 1.6044 | 0.532 | 1.0682 | 0.000999 |
| livertumor | RTMLA | ITGB1 | DAB2 | 1.6044 | 1.1613 | 1.3829 | 0.000999 |
| livertumor | RTMLA | ITGB1 | ENG | 1.6044 | 0.3636 | 0.984 | 0.000999 |
| livertumor | RTMLA | ITGB1 | ITGAX | 1.6044 | 0.257 | 0.9307 | 0.000999 |
| livertumor | RTMLA | ITGB1 | PLAUR | 1.6044 | 0.7331 | 1.1688 | 0.000999 |
| livertumor | RTMLA | ITGB1 | RACK1 | 1.6044 | 2.3077 | 1.9561 | 0.000999 |
| livertumor | RTMLA | MDK | LRP1 | 1.6101 | 0.5181 | 1.0641 | 0.000999 |
| livertumor | RTMLA | MIF | CD74 | 1.7638 | 4.4484 | 3.1061 | 0.000999 |

| | | | | | | | |
|---|---|---|---|---|---|---|---|
| livertumor | RTMLA | MIF | TNFRSF14 | 1.7638 | 0.31 | 1.0369 | 0.000999 |
| livertumor | RTMLA | NTS | PTAFR | 0.3425 | 0.2762 | 0.3093 | 0.000999 |
| livertumor | RTMLA | OCLN | TGFBR2 | 0.3091 | 0.3296 | 0.3194 | 0.000999 |
| livertumor | RTMLA | PTPRF | TGFBR2 | 0.384 | 0.3296 | 0.3568 | 0.000999 |
| livertumor | RTMLA | RPS19 | C5AR1 | 4.0772 | 0.6059 | 2.3416 | 0.000999 |
| livertumor | RTMLA | RPS19 | CD4 | 4.0772 | 0.6388 | 2.358 | 0.000999 |
| livertumor | RTMLA | RTN4 | CD44 | 1.0046 | 1.1045 | 1.0546 | 0.000999 |
| livertumor | RTMLA | RTN4 | CD99 | 1.0046 | 0.805 | 0.9048 | 0.000999 |
| livertumor | RTMLA | SEMA3C | NRP2 | 0.2576 | 0.3367 | 0.2972 | 0.000999 |
| livertumor | RTMLA | SERPINA1 | LRP1 | 1.604 | 0.5181 | 1.0611 | 0.000999 |
| livertumor | RTMLA | SERPING1 | LRP1 | 0.3498 | 0.5181 | 0.4339 | 0.000999 |
| livertumor | RTMLA | TF | CD4 | 0.523 | 0.6388 | 0.5809 | 0.000999 |
| livertumor | RTMLA | TF | TFRC | 0.523 | 0.5962 | 0.5596 | 0.000999 |
| livertumor | RTMLA | TFPI | LRP1 | 0.5043 | 0.5181 | 0.5112 | 0.000999 |
| livertumor | RTMLA | VEGFA | NRP2 | 0.5485 | 0.3367 | 0.4426 | 0.000999 |
| livertumor | RTMLA | VTN | PLAUR | 0.7399 | 0.7331 | 0.7365 | 0.000999 |
| livertumor | RTMLA | CD99 | CD99 | 0.5906 | 0.805 | 0.6978 | 0.002 |
| livertumor | RTMLA | LRPAP1 | LRP1 | 0.273 | 0.5181 | 0.3956 | 0.00599 |
| livertumor | RTMLA | GSTP1 | CD44 | 1.7219 | 1.1045 | 1.4132 | 0.00699 |
| lungtumor | RTMLA | ANXA1 | C3AR1 | 1.4026 | 0.501 | 0.9518 | 0.000999 |
| lungtumor | RTMLA | ANXA1 | C5AR1 | 1.4026 | 0.6059 | 1.0043 | 0.000999 |
| lungtumor | RTMLA | ANXA1 | CCR1 | 1.4026 | 0.3215 | 0.8621 | 0.000999 |
| lungtumor | RTMLA | ANXA1 | CD4 | 1.4026 | 0.6388 | 1.0207 | 0.000999 |
| lungtumor | RTMLA | ANXA1 | CD44 | 1.4026 | 1.1045 | 1.2536 | 0.000999 |
| lungtumor | RTMLA | ANXA1 | CXCR4 | 1.4026 | 1.0274 | 1.215 | 0.000999 |
| lungtumor | RTMLA | ANXA1 | FPR1 | 1.4026 | 0.3438 | 0.8732 | 0.000999 |
| lungtumor | RTMLA | ANXA1 | FPR3 | 1.4026 | 0.6512 | 1.0269 | 0.000999 |
| lungtumor | RTMLA | ARF1 | CD4 | 0.8791 | 0.6388 | 0.759 | 0.000999 |
| lungtumor | RTMLA | ARF1 | CD44 | 0.8791 | 1.1045 | 0.9918 | 0.000999 |
| lungtumor | RTMLA | CALM1 | TNFRSF1A | 0.8426 | 0.3864 | 0.6145 | 0.000999 |

| | | | | | | | |
|---|---|---|---|---|---|---|---|
| lungtumor | RTMLA | CALM1 | TNFRSF1B | 0.8426 | 0.3313 | 0.587 | 0.000999 |
| lungtumor | RTMLA | CALR | CD44 | 0.7795 | 1.1045 | 0.942 | 0.000999 |
| lungtumor | RTMLA | CALR | LRP1 | 0.7795 | 0.5181 | 0.6488 | 0.000999 |
| lungtumor | RTMLA | CCL20 | C3AR1 | 0.2885 | 0.501 | 0.3948 | 0.000999 |
| lungtumor | RTMLA | CCL20 | C5AR1 | 0.2885 | 0.6059 | 0.4472 | 0.000999 |
| lungtumor | RTMLA | CCL20 | CCR1 | 0.2885 | 0.3215 | 0.305 | 0.000999 |
| lungtumor | RTMLA | CCL20 | CXCR4 | 0.2885 | 1.0274 | 0.658 | 0.000999 |
| lungtumor | RTMLA | CCL20 | FPR1 | 0.2885 | 0.3438 | 0.3162 | 0.000999 |
| lungtumor | RTMLA | CD99 | CD99 | 0.5938 | 0.805 | 0.6994 | 0.000999 |
| lungtumor | RTMLA | CXCL10 | C3AR1 | 0.3417 | 0.501 | 0.4213 | 0.000999 |
| lungtumor | RTMLA | CXCL10 | C5AR1 | 0.3417 | 0.6059 | 0.4738 | 0.000999 |
| lungtumor | RTMLA | CXCL10 | CCR1 | 0.3417 | 0.3215 | 0.3316 | 0.000999 |
| lungtumor | RTMLA | CXCL10 | CXCR4 | 0.3417 | 1.0274 | 0.6845 | 0.000999 |
| lungtumor | RTMLA | CXCL10 | FPR1 | 0.3417 | 0.3438 | 0.3427 | 0.000999 |
| lungtumor | RTMLA | FGA | ITGAX | 0.4445 | 0.257 | 0.3507 | 0.000999 |
| lungtumor | RTMLA | FGA | ITGB2 | 0.4445 | 1.2597 | 0.8521 | 0.000999 |
| lungtumor | RTMLA | GRN | TNFRSF1B | 0.4026 | 0.3313 | 0.367 | 0.000999 |
| lungtumor | RTMLA | GSTP1 | CD44 | 2.4093 | 1.1045 | 1.7569 | 0.000999 |
| lungtumor | RTMLA | HLA-DRA | CD4 | 0.9609 | 0.6388 | 0.7999 | 0.000999 |
| lungtumor | RTMLA | HLA-DRA | PTPRC | 0.9609 | 0.5031 | 0.732 | 0.000999 |
| lungtumor | RTMLA | HMGB1 | LY96 | 1.1943 | 1.2707 | 1.2325 | 0.000999 |
| lungtumor | RTMLA | HMGB2 | CD44 | 0.5593 | 1.1045 | 0.8319 | 0.000999 |
| lungtumor | RTMLA | HMGB2 | DAB2 | 0.5593 | 1.1613 | 0.8603 | 0.000999 |
| lungtumor | RTMLA | HSP90B1 | LRP1 | 0.6141 | 0.5181 | 0.5661 | 0.000999 |
| lungtumor | RTMLA | HSP90B1 | TNFRSF1B | 0.6141 | 0.3313 | 0.4727 | 0.000999 |
| lungtumor | RTMLA | LGALS3 | CLEC7A | 0.9915 | 0.4847 | 0.7381 | 0.000999 |
| lungtumor | RTMLA | LGALS3 | FCGR2A | 0.9915 | 1.0353 | 1.0134 | 0.000999 |
| lungtumor | RTMLA | LGALS3 | FCGR2B | 0.9915 | 0.7235 | 0.8575 | 0.000999 |
| lungtumor | RTMLA | LGALS3 | TFRC | 0.9915 | 0.5962 | 0.7939 | 0.000999 |
| lungtumor | RTMLA | LGALS3 | TGFBR2 | 0.9915 | 0.3296 | 0.6606 | 0.000999 |

| | | | | | | | |
|---|---|---|---|---|---|---|---|
| lungtumor | RTMLA | MDK | LRP1 | 1.2896 | 0.5181 | 0.9038 | 0.000999 |
| lungtumor | RTMLA | MIF | CD74 | 1.4585 | 4.4484 | 2.9535 | 0.000999 |
| lungtumor | RTMLA | MIF | TNFRSF14 | 1.4585 | 0.31 | 0.8843 | 0.000999 |
| lungtumor | RTMLA | PSAP | LRP1 | 0.5457 | 0.5181 | 0.5319 | 0.000999 |
| lungtumor | RTMLA | RPS19 | C5AR1 | 3.5104 | 0.6059 | 2.0582 | 0.000999 |
| lungtumor | RTMLA | RPS19 | CD4 | 3.5104 | 0.6388 | 2.0746 | 0.000999 |
| lungtumor | RTMLA | RTN4 | CD44 | 0.4013 | 1.1045 | 0.7529 | 0.000999 |
| lungtumor | RTMLA | SAA1 | C3AR1 | 0.2539 | 0.501 | 0.3774 | 0.000999 |
| lungtumor | RTMLA | SAA1 | C5AR1 | 0.2539 | 0.6059 | 0.4299 | 0.000999 |
| lungtumor | RTMLA | SAA1 | CCR1 | 0.2539 | 0.3215 | 0.2877 | 0.000999 |
| lungtumor | RTMLA | SAA1 | CXCR4 | 0.2539 | 1.0274 | 0.6406 | 0.000999 |
| lungtumor | RTMLA | SAA1 | FPR1 | 0.2539 | 0.3438 | 0.2988 | 0.000999 |
| lungtumor | RTMLA | SAA1 | PTAFR | 0.2539 | 0.2762 | 0.265 | 0.000999 |
| lungtumor | RTMLA | SERPINA1 | LRP1 | 0.3624 | 0.5181 | 0.4402 | 0.000999 |
| lungtumor | RTMLA | SPP1 | CD44 | 1.4874 | 1.1045 | 1.296 | 0.000999 |
| lungtumor | RTMLA | TFPI | LRP1 | 0.3615 | 0.5181 | 0.4398 | 0.000999 |
| lungtumor | RTMLA | TIMP1 | LRP1 | 0.6195 | 0.5181 | 0.5688 | 0.000999 |
| lungtumor | RTMLA | RTN4 | CD99 | 0.4013 | 0.805 | 0.6031 | 0.00599 |
| melanomatumo | RTMLA | ARF1 | CD4 | 1.8545 | 0.6388 | 1.2466 | 0.000999 |
| melanomatumo | RTMLA | ARF1 | CD44 | 1.8545 | 1.1045 | 1.4795 | 0.000999 |
| melanomatumo | RTMLA | CD320 | TGFBR2 | 0.2588 | 0.3296 | 0.2942 | 0.000999 |
| melanomatumo | RTMLA | GSTP1 | CD44 | 2.3341 | 1.1045 | 1.7193 | 0.000999 |
| melanomatumo | RTMLA | HAS2 | CD44 | 0.3078 | 1.1045 | 0.7062 | 0.000999 |
| melanomatumo | RTMLA | HMGB1 | LY96 | 1.6757 | 1.2707 | 1.4732 | 0.000999 |
| melanomatumo | RTMLA | MIF | CD74 | 2.6939 | 4.4484 | 3.5712 | 0.000999 |
| melanomatumo | RTMLA | MIF | TNFRSF14 | 2.6939 | 0.31 | 1.502 | 0.000999 |
| melanomatumo | RTMLA | SERPINE2 | LRP1 | 0.9323 | 0.5181 | 0.7252 | 0.000999 |
| melanomatumo | RTMLA | HEBP1 | CXCR4 | 0.3689 | 1.0274 | 0.6981 | 0.004 |
| ovarytumor | RTMLA | ANXA1 | C3AR1 | 1.0314 | 0.501 | 0.7662 | 0.000999 |
| ovarytumor | RTMLA | ANXA1 | C5AR1 | 1.0314 | 0.6059 | 0.8186 | 0.000999 |

| | | | | | | | |
|---|---|---|---|---|---|---|---|
| ovarytumor | RTMLA | ANXA1 | CCR1 | 1.0314 | 0.3215 | 0.6765 | 0.000999 |
| ovarytumor | RTMLA | ANXA1 | CD4 | 1.0314 | 0.6388 | 0.8351 | 0.000999 |
| ovarytumor | RTMLA | ANXA1 | CD44 | 1.0314 | 1.1045 | 1.068 | 0.000999 |
| ovarytumor | RTMLA | ANXA1 | CXCR4 | 1.0314 | 1.0274 | 1.0294 | 0.000999 |
| ovarytumor | RTMLA | ANXA1 | FPR1 | 1.0314 | 0.3438 | 0.6876 | 0.000999 |
| ovarytumor | RTMLA | ANXA1 | FPR3 | 1.0314 | 0.6512 | 0.8413 | 0.000999 |
| ovarytumor | RTMLA | APOE | LRP1 | 0.837 | 0.5181 | 0.6776 | 0.000999 |
| ovarytumor | RTMLA | APP | ALOX5 | 0.2907 | 0.3103 | 0.3005 | 0.000999 |
| ovarytumor | RTMLA | APP | C3AR1 | 0.2907 | 0.501 | 0.3959 | 0.000999 |
| ovarytumor | RTMLA | APP | C5AR1 | 0.2907 | 0.6059 | 0.4483 | 0.000999 |
| ovarytumor | RTMLA | APP | CCR1 | 0.2907 | 0.3215 | 0.3061 | 0.000999 |
| ovarytumor | RTMLA | APP | CD36 | 0.2907 | 0.3707 | 0.3307 | 0.000999 |
| ovarytumor | RTMLA | APP | CXCR4 | 0.2907 | 1.0274 | 0.6591 | 0.000999 |
| ovarytumor | RTMLA | APP | FPR1 | 0.2907 | 0.3438 | 0.3173 | 0.000999 |
| ovarytumor | RTMLA | APP | LRP1 | 0.2907 | 0.5181 | 0.4044 | 0.000999 |
| ovarytumor | RTMLA | APP | TGFBR2 | 0.2907 | 0.3296 | 0.3102 | 0.000999 |
| ovarytumor | RTMLA | APP | TNFRSF14 | 0.2907 | 0.31 | 0.3004 | 0.000999 |
| ovarytumor | RTMLA | ARF1 | CD4 | 0.7364 | 0.6388 | 0.6876 | 0.000999 |
| ovarytumor | RTMLA | ARF1 | CD44 | 0.7364 | 1.1045 | 0.9205 | 0.000999 |
| ovarytumor | RTMLA | CALM1 | TNFRSF1A | 1.0087 | 0.3864 | 0.6975 | 0.000999 |
| ovarytumor | RTMLA | CALM1 | TNFRSF1B | 1.0087 | 0.3313 | 0.67 | 0.000999 |
| ovarytumor | RTMLA | CALR | CD44 | 0.7423 | 1.1045 | 0.9234 | 0.000999 |
| ovarytumor | RTMLA | CALR | LRP1 | 0.7423 | 0.5181 | 0.6302 | 0.000999 |
| ovarytumor | RTMLA | CCL2 | CCR1 | 0.3526 | 0.3215 | 0.3371 | 0.000999 |
| ovarytumor | RTMLA | COL1A1 | CD36 | 0.3504 | 0.3707 | 0.3606 | 0.000999 |
| ovarytumor | RTMLA | COL1A1 | CD44 | 0.3504 | 1.1045 | 0.7275 | 0.000999 |
| ovarytumor | RTMLA | GRN | TNFRSF1A | 0.3896 | 0.3864 | 0.388 | 0.000999 |
| ovarytumor | RTMLA | GRN | TNFRSF1B | 0.3896 | 0.3313 | 0.3604 | 0.000999 |
| ovarytumor | RTMLA | GSTP1 | CD44 | 2.6185 | 1.1045 | 1.8615 | 0.000999 |
| ovarytumor | RTMLA | HLA-DMA | CD4 | 0.2537 | 0.6388 | 0.4462 | 0.000999 |

| | | | | | | | |
|---|---|---|---|---|---|---|---|
| ovarytumor | RTMLA | HLA-DRA | CD4 | 1.2369 | 0.6388 | 0.9378 | 0.000999 |
| ovarytumor | RTMLA | HLA-DRA | PTPRC | 1.2369 | 0.5031 | 0.87 | 0.000999 |
| ovarytumor | RTMLA | HMGB1 | LY96 | 1.7272 | 1.2707 | 1.499 | 0.000999 |
| ovarytumor | RTMLA | HMGB2 | CD44 | 0.4974 | 1.1045 | 0.801 | 0.000999 |
| ovarytumor | RTMLA | HMGB2 | DAB2 | 0.4974 | 1.1613 | 0.8294 | 0.000999 |
| ovarytumor | RTMLA | HSP90B1 | LRP1 | 0.6365 | 0.5181 | 0.5773 | 0.000999 |
| ovarytumor | RTMLA | HSP90B1 | TNFRSF1B | 0.6365 | 0.3313 | 0.4839 | 0.000999 |
| ovarytumor | RTMLA | ITGB1 | CD36 | 0.2783 | 0.3707 | 0.3245 | 0.000999 |
| ovarytumor | RTMLA | ITGB1 | CD46 | 0.2783 | 0.3134 | 0.2959 | 0.000999 |
| ovarytumor | RTMLA | ITGB1 | DAB2 | 0.2783 | 1.1613 | 0.7198 | 0.000999 |
| ovarytumor | RTMLA | ITGB1 | ENG | 0.2783 | 0.3636 | 0.3209 | 0.000999 |
| ovarytumor | RTMLA | ITGB1 | ITGAX | 0.2783 | 0.257 | 0.2676 | 0.000999 |
| ovarytumor | RTMLA | ITGB1 | PLAUR | 0.2783 | 0.7331 | 0.5057 | 0.000999 |
| ovarytumor | RTMLA | LGALS3 | CLEC7A | 0.5097 | 0.4847 | 0.4972 | 0.000999 |
| ovarytumor | RTMLA | LGALS3 | FCGR2A | 0.5097 | 1.0353 | 0.7725 | 0.000999 |
| ovarytumor | RTMLA | LGALS3 | FCGR2B | 0.5097 | 0.7235 | 0.6166 | 0.000999 |
| ovarytumor | RTMLA | LGALS3 | TFRC | 0.5097 | 0.5962 | 0.553 | 0.000999 |
| ovarytumor | RTMLA | LGALS3 | TGFBR2 | 0.5097 | 0.3296 | 0.4197 | 0.000999 |
| ovarytumor | RTMLA | LRPAP1 | LRP1 | 0.3826 | 0.5181 | 0.4504 | 0.000999 |
| ovarytumor | RTMLA | MDK | LRP1 | 1.5321 | 0.5181 | 1.0251 | 0.000999 |
| ovarytumor | RTMLA | MIF | CD74 | 0.8565 | 4.4484 | 2.6524 | 0.000999 |
| ovarytumor | RTMLA | MIF | TNFRSF14 | 0.8565 | 0.31 | 0.5832 | 0.000999 |
| ovarytumor | RTMLA | PSAP | LRP1 | 0.6574 | 0.5181 | 0.5878 | 0.000999 |
| ovarytumor | RTMLA | RPS19 | C5AR1 | 3.9325 | 0.6059 | 2.2692 | 0.000999 |
| ovarytumor | RTMLA | RPS19 | CD4 | 3.9325 | 0.6388 | 2.2857 | 0.000999 |
| ovarytumor | RTMLA | RTN4 | CD44 | 0.4992 | 1.1045 | 0.8019 | 0.000999 |
| ovarytumor | RTMLA | RTN4 | CD99 | 0.4992 | 0.805 | 0.6521 | 0.000999 |
| ovarytumor | RTMLA | SERPING1 | LRP1 | 0.3678 | 0.5181 | 0.443 | 0.000999 |
| ovarytumor | RTMLA | SPP1 | CD44 | 0.3946 | 1.1045 | 0.7496 | 0.000999 |
| ovarytumor | RTMLA | TIMP1 | LRP1 | 0.7779 | 0.5181 | 0.648 | 0.000999 |

| Supplementary Table 8: Communication results between macrophages and lymphocytes subpopulations, and malignant cells | | | | | | | |
|---|---|---|---|---|---|---|---|
| **RTM-likeMT vs. lymphocytes** | | | | | | | |
| celltype_01 | celltype_02 | ligand | receptor | ligand_mean_expression | receptor_mean_expression | ligand_receptor_mean_expression | pvalue |
| RTM-likeMT | NKT | A2M | TNFRSF14 | 0.7163 | 0.2848 | 0.5005 | 0.000999 |
| RTM-likeMT | NKT | ADM | PTGER2 | 0.4876 | 0.3627 | 0.4252 | 0.000999 |
| RTM-likeMT | NKT | APOE | SORL1 | 3.1184 | 0.4137 | 1.766 | 0.000999 |
| RTM-likeMT | NKT | CCL20 | CXCR4 | 0.9244 | 2.0373 | 1.4808 | 0.000999 |
| RTM-likeMT | NKT | CCL20 | S1PR4 | 0.9244 | 0.4435 | 0.6839 | 0.000999 |
| RTM-likeMT | NKT | CCL20 | S1PR5 | 0.9244 | 0.508 | 0.7162 | 0.000999 |
| RTM-likeMT | NKT | COL1A1 | CD44 | 0.2826 | 0.6724 | 0.4775 | 0.000999 |
| RTM-likeMT | NKT | COL1A1 | ITGB1 | 0.2826 | 0.488 | 0.3853 | 0.000999 |
| RTM-likeMT | NKT | CXCL16 | CXCR4 | 1.0911 | 2.0373 | 1.5642 | 0.000999 |
| RTM-likeMT | NKT | CXCL16 | S1PR4 | 1.0911 | 0.4435 | 0.7673 | 0.000999 |
| RTM-likeMT | NKT | CXCL16 | S1PR5 | 1.0911 | 0.508 | 0.7996 | 0.000999 |
| RTM-likeMT | NKT | CXCL2 | CXCR4 | 1.9342 | 2.0373 | 1.9857 | 0.000999 |
| RTM-likeMT | NKT | CXCL2 | S1PR4 | 1.9342 | 0.4435 | 1.1889 | 0.000999 |
| RTM-likeMT | NKT | CXCL2 | S1PR5 | 1.9342 | 0.508 | 1.2211 | 0.000999 |
| RTM-likeMT | NKT | CXCL3 | CXCR4 | 1.7116 | 2.0373 | 1.8744 | 0.000999 |
| RTM-likeMT | NKT | CXCL3 | S1PR4 | 1.7116 | 0.4435 | 1.0776 | 0.000999 |
| RTM-likeMT | NKT | CXCL3 | S1PR5 | 1.7116 | 0.508 | 1.1098 | 0.000999 |
| RTM-likeMT | NKT | FN1 | CD44 | 2.2884 | 0.6724 | 1.4804 | 0.000999 |
| RTM-likeMT | NKT | FN1 | ITGA4 | 2.2884 | 0.2946 | 1.2915 | 0.000999 |
| RTM-likeMT | NKT | FN1 | ITGB1 | 2.2884 | 0.488 | 1.3882 | 0.000999 |
| RTM-likeMT | NKT | FN1 | ITGB7 | 2.2884 | 0.4721 | 1.3803 | 0.000999 |
| RTM-likeMT | NKT | FN1 | RPSA | 2.2884 | 2.3639 | 2.3262 | 0.000999 |
| RTM-likeMT | NKT | GRN | TNFRSF1B | 1.6859 | 0.4949 | 1.0904 | 0.000999 |
| RTM-likeMT | NKT | GSTP1 | CD44 | 1.5481 | 0.6724 | 1.1103 | 0.000999 |
| RTM-likeMT | NKT | GSTP1 | IL7R | 1.5481 | 0.5933 | 1.0707 | 0.000999 |
| RTM-likeMT | NKT | GSTP1 | ITGA4 | 1.5481 | 0.2946 | 0.9213 | 0.000999 |
| RTM-likeMT | NKT | HBEGF | CD44 | 0.4141 | 0.6724 | 0.5433 | 0.000999 |

| | | | | | | | |
|---|---|---|---|---|---|---|---|
| RTM-likeMT | NKT | HLA-DRA | PTPRC | 3.9047 | 1.8797 | 2.8922 | 0.000999 |
| RTM-likeMT | NKT | ICAM1 | IL2RG | 0.3448 | 1.1524 | 0.7486 | 0.000999 |
| RTM-likeMT | NKT | ICAM1 | ITGAL | 0.3448 | 0.6778 | 0.5113 | 0.000999 |
| RTM-likeMT | NKT | ICAM1 | ITGB2 | 0.3448 | 1.6481 | 0.9964 | 0.000999 |
| RTM-likeMT | NKT | ICAM1 | SPN | 0.3448 | 0.6398 | 0.4923 | 0.000999 |
| RTM-likeMT | NKT | IL10 | IL10RA | 0.2855 | 0.5156 | 0.4006 | 0.000999 |
| RTM-likeMT | NKT | LGALS3 | CD6 | 0.8673 | 0.4881 | 0.6777 | 0.000999 |
| RTM-likeMT | NKT | LGALS3 | ITGB1 | 0.8673 | 0.488 | 0.6776 | 0.000999 |
| RTM-likeMT | NKT | LGALS3 | SPN | 0.8673 | 0.6398 | 0.7535 | 0.000999 |
| RTM-likeMT | NKT | LRPAP1 | SORL1 | 0.4493 | 0.4137 | 0.4315 | 0.000999 |
| RTM-likeMT | NKT | MIF | CD74 | 1.0074 | 1.6792 | 1.3433 | 0.000999 |
| RTM-likeMT | NKT | MIF | ITGA4 | 1.0074 | 0.2946 | 0.651 | 0.000999 |
| RTM-likeMT | NKT | MIF | TNFRSF14 | 1.0074 | 0.2848 | 0.6461 | 0.000999 |
| RTM-likeMT | NKT | NAMPT | ANXA6 | 1.5359 | 0.844 | 1.1899 | 0.000999 |
| RTM-likeMT | NKT | PECAM1 | CXCR4 | 0.3422 | 2.0373 | 1.1898 | 0.000999 |
| RTM-likeMT | NKT | RTN4 | CD44 | 0.9166 | 0.6724 | 0.7945 | 0.000999 |
| RTM-likeMT | NKT | RTN4 | CD99 | 0.9166 | 1.4547 | 1.1857 | 0.000999 |
| RTM-likeMT | NKT | SPP1 | CD44 | 0.7979 | 0.6724 | 0.7352 | 0.000999 |
| RTM-likeMT | NKT | SPP1 | ITGA4 | 0.7979 | 0.2946 | 0.5462 | 0.000999 |
| RTM-likeMT | NKT | SPP1 | ITGB1 | 0.7979 | 0.488 | 0.6429 | 0.000999 |
| RTM-likeMT | NKT | THBS1 | CD47 | 0.7512 | 0.5576 | 0.6544 | 0.000999 |
| RTM-likeMT | NKT | THBS1 | ITGA4 | 0.7512 | 0.2946 | 0.5229 | 0.000999 |
| RTM-likeMT | NKT | THBS1 | ITGB1 | 0.7512 | 0.488 | 0.6196 | 0.000999 |
| RTM-likeMT | NKT | TNFSF13B | ITGB1 | 0.7616 | 0.488 | 0.6248 | 0.000999 |
| RTM-likeMT | TCD4ex | A2M | TNFRSF14 | 0.7163 | 0.2806 | 0.4985 | 0.000999 |
| RTM-likeMT | TCD4ex | ADM | PTGER4 | 0.4876 | 0.4696 | 0.4786 | 0.000999 |
| RTM-likeMT | TCD4ex | ANXA1 | CCR7 | 1.1172 | 0.4844 | 0.8008 | 0.000999 |
| RTM-likeMT | TCD4ex | ANXA1 | CD4 | 1.1172 | 0.5864 | 0.8518 | 0.000999 |
| RTM-likeMT | TCD4ex | ANXA1 | CD44 | 1.1172 | 1.0244 | 1.0708 | 0.000999 |
| RTM-likeMT | TCD4ex | ANXA1 | CXCR4 | 1.1172 | 2.4201 | 1.7686 | 0.000999 |

| | | | | | | | |
|---|---|---|---|---|---|---|---|
| RTM-likeMT | TCD4ex | CCL20 | CCR7 | 0.9244 | 0.4844 | 0.7044 | 0.000999 |
| RTM-likeMT | TCD4ex | CCL20 | CXCR4 | 0.9244 | 2.4201 | 1.6722 | 0.000999 |
| RTM-likeMT | TCD4ex | CD86 | CD28 | 0.5683 | 0.3256 | 0.4469 | 0.000999 |
| RTM-likeMT | TCD4ex | CD86 | CTLA4 | 0.5683 | 0.6294 | 0.5989 | 0.000999 |
| RTM-likeMT | TCD4ex | COL1A1 | CD44 | 0.2826 | 1.0244 | 0.6535 | 0.000999 |
| RTM-likeMT | TCD4ex | COL1A1 | ITGB1 | 0.2826 | 0.4615 | 0.372 | 0.000999 |
| RTM-likeMT | TCD4ex | CXCL16 | CCR7 | 1.0911 | 0.4844 | 0.7878 | 0.000999 |
| RTM-likeMT | TCD4ex | CXCL16 | CXCR4 | 1.0911 | 2.4201 | 1.7556 | 0.000999 |
| RTM-likeMT | TCD4ex | CXCL2 | CCR7 | 1.9342 | 0.4844 | 1.2093 | 0.000999 |
| RTM-likeMT | TCD4ex | CXCL2 | CXCR4 | 1.9342 | 2.4201 | 2.1771 | 0.000999 |
| RTM-likeMT | TCD4ex | CXCL3 | CCR7 | 1.7116 | 0.4844 | 1.098 | 0.000999 |
| RTM-likeMT | TCD4ex | CXCL3 | CXCR4 | 1.7116 | 2.4201 | 2.0658 | 0.000999 |
| RTM-likeMT | TCD4ex | FN1 | CD44 | 2.2884 | 1.0244 | 1.6564 | 0.000999 |
| RTM-likeMT | TCD4ex | FN1 | ITGB1 | 2.2884 | 0.4615 | 1.375 | 0.000999 |
| RTM-likeMT | TCD4ex | FN1 | RPSA | 2.2884 | 2.4895 | 2.389 | 0.000999 |
| RTM-likeMT | TCD4ex | GRN | TNFRSF1B | 1.6859 | 0.383 | 1.0345 | 0.000999 |
| RTM-likeMT | TCD4ex | GSTP1 | CD44 | 1.5481 | 1.0244 | 1.2863 | 0.000999 |
| RTM-likeMT | TCD4ex | GSTP1 | IL7R | 1.5481 | 1.2353 | 1.3917 | 0.000999 |
| RTM-likeMT | TCD4ex | HBEGF | CD44 | 0.4141 | 1.0244 | 0.7193 | 0.000999 |
| RTM-likeMT | TCD4ex | HLA-DMA | CD4 | 1.7556 | 0.5864 | 1.171 | 0.000999 |
| RTM-likeMT | TCD4ex | HLA-DRA | CD4 | 3.9047 | 0.5864 | 2.2456 | 0.000999 |
| RTM-likeMT | TCD4ex | HLA-DRA | PDCD1 | 3.9047 | 0.457 | 2.1808 | 0.000999 |
| RTM-likeMT | TCD4ex | HLA-DRA | PTPRC | 3.9047 | 1.7493 | 2.827 | 0.000999 |
| RTM-likeMT | TCD4ex | HSP90B1 | ITGB1 | 0.5443 | 0.4615 | 0.5029 | 0.000999 |
| RTM-likeMT | TCD4ex | ICAM1 | IL2RG | 0.3448 | 0.9695 | 0.6571 | 0.000999 |
| RTM-likeMT | TCD4ex | ICAM1 | ITGAL | 0.3448 | 0.2603 | 0.3025 | 0.000999 |
| RTM-likeMT | TCD4ex | ICAM1 | ITGB2 | 0.3448 | 0.7871 | 0.5659 | 0.000999 |
| RTM-likeMT | TCD4ex | IL10 | CD28 | 0.2855 | 0.3256 | 0.3055 | 0.000999 |
| RTM-likeMT | TCD4ex | ITGB2 | ITGAL | 1.4781 | 0.2603 | 0.8692 | 0.000999 |
| RTM-likeMT | TCD4ex | LGALS3 | CD6 | 0.8673 | 0.5338 | 0.7005 | 0.000999 |

| | | | | | | | |
|---|---|---|---|---|---|---|---|
| RTM-likeMT | TCD4ex | LGALS3 | ITGB1 | 0.8673 | 0.4615 | 0.6644 | 0.000999 |
| RTM-likeMT | TCD4ex | MIF | TNFRSF14 | 1.0074 | 0.2806 | 0.644 | 0.000999 |
| RTM-likeMT | TCD4ex | NAMPT | ANXA6 | 1.5359 | 0.5386 | 1.0373 | 0.000999 |
| RTM-likeMT | TCD4ex | PECAM1 | CXCR4 | 0.3422 | 2.4201 | 1.3811 | 0.000999 |
| RTM-likeMT | TCD4ex | RTN4 | CD44 | 0.9166 | 1.0244 | 0.9705 | 0.000999 |
| RTM-likeMT | TCD4ex | RTN4 | CD99 | 0.9166 | 0.9165 | 0.9166 | 0.000999 |
| RTM-likeMT | TCD4ex | SPP1 | CD44 | 0.7979 | 1.0244 | 0.9111 | 0.000999 |
| RTM-likeMT | TCD4ex | SPP1 | ITGB1 | 0.7979 | 0.4615 | 0.6297 | 0.000999 |
| RTM-likeMT | TCD4ex | THBS1 | CD47 | 0.7512 | 0.3456 | 0.5484 | 0.000999 |
| RTM-likeMT | TCD4ex | THBS1 | ITGB1 | 0.7512 | 0.4615 | 0.6064 | 0.000999 |
| RTM-likeMT | TCD4ex | TNFSF13B | ITGB1 | 0.7616 | 0.4615 | 0.6116 | 0.000999 |
| RTM-likeMT | TCD4ex | HSP90B1 | TNFRSF1B | 0.5443 | 0.383 | 0.4637 | 0.00899 |
| RTM-likeMT | TCD4reg | A2M | TNFRSF14 | 0.7163 | 0.3843 | 0.5503 | 0.000999 |
| RTM-likeMT | TCD4reg | ANXA1 | CCR7 | 1.1172 | 0.3098 | 0.7135 | 0.000999 |
| RTM-likeMT | TCD4reg | ANXA1 | CD4 | 1.1172 | 0.429 | 0.7731 | 0.000999 |
| RTM-likeMT | TCD4reg | ANXA1 | CD44 | 1.1172 | 1.0896 | 1.1034 | 0.000999 |
| RTM-likeMT | TCD4reg | ANXA1 | CXCR3 | 1.1172 | 0.3357 | 0.7264 | 0.000999 |
| RTM-likeMT | TCD4reg | ANXA1 | CXCR4 | 1.1172 | 2.2962 | 1.7067 | 0.000999 |
| RTM-likeMT | TCD4reg | ANXA1 | CXCR6 | 1.1172 | 0.3945 | 0.7559 | 0.000999 |
| RTM-likeMT | TCD4reg | ANXA1 | FAS | 1.1172 | 0.2727 | 0.695 | 0.000999 |
| RTM-likeMT | TCD4reg | CCL20 | CCR7 | 0.9244 | 0.3098 | 0.6171 | 0.000999 |
| RTM-likeMT | TCD4reg | CCL20 | CXCR3 | 0.9244 | 0.3357 | 0.63 | 0.000999 |
| RTM-likeMT | TCD4reg | CCL20 | CXCR4 | 0.9244 | 2.2962 | 1.6103 | 0.000999 |
| RTM-likeMT | TCD4reg | CCL20 | CXCR6 | 0.9244 | 0.3945 | 0.6594 | 0.000999 |
| RTM-likeMT | TCD4reg | CCL20 | S1PR4 | 0.9244 | 0.2744 | 0.5994 | 0.000999 |
| RTM-likeMT | TCD4reg | CD86 | CD28 | 0.5683 | 0.313 | 0.4407 | 0.000999 |
| RTM-likeMT | TCD4reg | CD86 | CTLA4 | 0.5683 | 0.7909 | 0.6796 | 0.000999 |
| RTM-likeMT | TCD4reg | CD99 | CD99 | 0.8632 | 0.6603 | 0.7618 | 0.000999 |
| RTM-likeMT | TCD4reg | CXCL16 | CCR7 | 1.0911 | 0.3098 | 0.7005 | 0.000999 |
| RTM-likeMT | TCD4reg | CXCL16 | CXCR3 | 1.0911 | 0.3357 | 0.7134 | 0.000999 |

| | | | | | | | |
|---|---|---|---|---|---|---|---|
| RTM-likeMT | TCD4reg | CXCL16 | CXCR4 | 1.0911 | 2.2962 | 1.6937 | 0.000999 |
| RTM-likeMT | TCD4reg | CXCL16 | CXCR6 | 1.0911 | 0.3945 | 0.7428 | 0.000999 |
| RTM-likeMT | TCD4reg | CXCL16 | S1PR4 | 1.0911 | 0.2744 | 0.6828 | 0.000999 |
| RTM-likeMT | TCD4reg | CXCL2 | CCR7 | 1.9342 | 0.3098 | 1.122 | 0.000999 |
| RTM-likeMT | TCD4reg | CXCL2 | CXCR3 | 1.9342 | 0.3357 | 1.1349 | 0.000999 |
| RTM-likeMT | TCD4reg | CXCL2 | CXCR4 | 1.9342 | 2.2962 | 2.1152 | 0.000999 |
| RTM-likeMT | TCD4reg | CXCL2 | CXCR6 | 1.9342 | 0.3945 | 1.1644 | 0.000999 |
| RTM-likeMT | TCD4reg | CXCL2 | S1PR4 | 1.9342 | 0.2744 | 1.1043 | 0.000999 |
| RTM-likeMT | TCD4reg | CXCL3 | CCR7 | 1.7116 | 0.3098 | 1.0107 | 0.000999 |
| RTM-likeMT | TCD4reg | CXCL3 | CXCR3 | 1.7116 | 0.3357 | 1.0236 | 0.000999 |
| RTM-likeMT | TCD4reg | CXCL3 | CXCR4 | 1.7116 | 2.2962 | 2.0039 | 0.000999 |
| RTM-likeMT | TCD4reg | CXCL3 | CXCR6 | 1.7116 | 0.3945 | 1.053 | 0.000999 |
| RTM-likeMT | TCD4reg | CXCL3 | S1PR4 | 1.7116 | 0.2744 | 0.993 | 0.000999 |
| RTM-likeMT | TCD4reg | FN1 | CD44 | 2.2884 | 1.0896 | 1.689 | 0.000999 |
| RTM-likeMT | TCD4reg | FN1 | ITGB1 | 2.2884 | 0.4001 | 1.3443 | 0.000999 |
| RTM-likeMT | TCD4reg | FN1 | RPSA | 2.2884 | 2.4999 | 2.3942 | 0.000999 |
| RTM-likeMT | TCD4reg | GRN | TNFRSF1B | 1.6859 | 0.7982 | 1.2421 | 0.000999 |
| RTM-likeMT | TCD4reg | GSTP1 | CD44 | 1.5481 | 1.0896 | 1.3189 | 0.000999 |
| RTM-likeMT | TCD4reg | GSTP1 | FAS | 1.5481 | 0.2727 | 0.9104 | 0.000999 |
| RTM-likeMT | TCD4reg | GSTP1 | IL7R | 1.5481 | 0.9682 | 1.2582 | 0.000999 |
| RTM-likeMT | TCD4reg | HBEGF | CD44 | 0.4141 | 1.0896 | 0.7519 | 0.000999 |
| RTM-likeMT | TCD4reg | HLA-DMA | CD4 | 1.7556 | 0.429 | 1.0923 | 0.000999 |
| RTM-likeMT | TCD4reg | HLA-DRA | CD4 | 3.9047 | 0.429 | 2.1668 | 0.000999 |
| RTM-likeMT | TCD4reg | HLA-DRA | PTPRC | 3.9047 | 1.6603 | 2.7825 | 0.000999 |
| RTM-likeMT | TCD4reg | ICAM1 | IL2RA | 0.3448 | 0.7115 | 0.5281 | 0.000999 |
| RTM-likeMT | TCD4reg | ICAM1 | IL2RG | 0.3448 | 1.1601 | 0.7524 | 0.000999 |
| RTM-likeMT | TCD4reg | ICAM1 | ITGB2 | 0.3448 | 0.6238 | 0.4843 | 0.000999 |
| RTM-likeMT | TCD4reg | IL10 | CD28 | 0.2855 | 0.313 | 0.2993 | 0.000999 |
| RTM-likeMT | TCD4reg | IL10 | IFNAR2 | 0.2855 | 0.2663 | 0.2759 | 0.000999 |
| RTM-likeMT | TCD4reg | IL10 | IL10RA | 0.2855 | 0.4952 | 0.3903 | 0.000999 |

| | | | | | | | |
|---|---|---|---|---|---|---|---|
| RTM-likeMT | TCD4reg | LGALS3 | CD6 | 0.8673 | 0.2651 | 0.5662 | 0.000999 |
| RTM-likeMT | TCD4reg | LGALS3 | FAS | 0.8673 | 0.2727 | 0.57 | 0.000999 |
| RTM-likeMT | TCD4reg | LGALS3 | ITGB1 | 0.8673 | 0.4001 | 0.6337 | 0.000999 |
| RTM-likeMT | TCD4reg | MIF | CD74 | 1.0074 | 2.0716 | 1.5395 | 0.000999 |
| RTM-likeMT | TCD4reg | MIF | TNFRSF14 | 1.0074 | 0.3843 | 0.6959 | 0.000999 |
| RTM-likeMT | TCD4reg | NAMPT | ANXA6 | 1.5359 | 0.4557 | 0.9958 | 0.000999 |
| RTM-likeMT | TCD4reg | PECAM1 | CXCR4 | 0.3422 | 2.2962 | 1.3192 | 0.000999 |
| RTM-likeMT | TCD4reg | RTN4 | CD44 | 0.9166 | 1.0896 | 1.0031 | 0.000999 |
| RTM-likeMT | TCD4reg | RTN4 | CD99 | 0.9166 | 0.6603 | 0.7885 | 0.000999 |
| RTM-likeMT | TCD4reg | SPP1 | CD44 | 0.7979 | 1.0896 | 0.9438 | 0.000999 |
| RTM-likeMT | TCD4reg | SPP1 | ITGB1 | 0.7979 | 0.4001 | 0.599 | 0.000999 |
| RTM-likeMT | TCD4reg | THBS1 | CD47 | 0.7512 | 0.416 | 0.5836 | 0.000999 |
| RTM-likeMT | TCD4reg | THBS1 | ITGB1 | 0.7512 | 0.4001 | 0.5757 | 0.000999 |
| RTM-likeMT | TCD4reg | TNFSF13B | ITGB1 | 0.7616 | 0.4001 | 0.5809 | 0.000999 |
| RTM-likeMT | TCD4reg | COL1A1 | ITGB1 | 0.2826 | 0.4001 | 0.3413 | 0.002 |
| RTM-likeMT | TCD4reg | HSP90B1 | ITGB1 | 0.5443 | 0.4001 | 0.4722 | 0.002 |
| RTM-likeMT | TCD4reg | HSP90B1 | TNFRSF1B | 0.5443 | 0.7982 | 0.6713 | 0.002 |
| RTM-likeMT | TCD8em | A2M | TNFRSF14 | 0.7163 | 0.2915 | 0.5039 | 0.000999 |
| RTM-likeMT | TCD8em | ADM | PTGER4 | 0.4876 | 0.2847 | 0.3862 | 0.000999 |
| RTM-likeMT | TCD8em | CCL20 | CXCR3 | 0.9244 | 0.4189 | 0.6716 | 0.000999 |
| RTM-likeMT | TCD8em | CCL20 | CXCR4 | 0.9244 | 2.8389 | 1.8816 | 0.000999 |
| RTM-likeMT | TCD8em | CCL20 | CXCR6 | 0.9244 | 0.3399 | 0.6321 | 0.000999 |
| RTM-likeMT | TCD8em | CCL20 | S1PR4 | 0.9244 | 0.2677 | 0.596 | 0.000999 |
| RTM-likeMT | TCD8em | CXCL16 | CXCR3 | 1.0911 | 0.4189 | 0.755 | 0.000999 |
| RTM-likeMT | TCD8em | CXCL16 | CXCR4 | 1.0911 | 2.8389 | 1.965 | 0.000999 |
| RTM-likeMT | TCD8em | CXCL16 | CXCR6 | 1.0911 | 0.3399 | 0.7155 | 0.000999 |
| RTM-likeMT | TCD8em | CXCL16 | S1PR4 | 1.0911 | 0.2677 | 0.6794 | 0.000999 |
| RTM-likeMT | TCD8em | CXCL2 | CXCR3 | 1.9342 | 0.4189 | 1.1765 | 0.000999 |
| RTM-likeMT | TCD8em | CXCL2 | CXCR4 | 1.9342 | 2.8389 | 2.3866 | 0.000999 |
| RTM-likeMT | TCD8em | CXCL2 | CXCR6 | 1.9342 | 0.3399 | 1.137 | 0.000999 |

| | | | | | | | |
|---|---|---|---|---|---|---|---|
| RTM-likeMT | TCD8em | CXCL2 | S1PR4 | 1.9342 | 0.2677 | 1.101 | 0.000999 |
| RTM-likeMT | TCD8em | CXCL3 | CXCR3 | 1.7116 | 0.4189 | 1.0652 | 0.000999 |
| RTM-likeMT | TCD8em | CXCL3 | CXCR4 | 1.7116 | 2.8389 | 2.2752 | 0.000999 |
| RTM-likeMT | TCD8em | CXCL3 | CXCR6 | 1.7116 | 0.3399 | 1.0257 | 0.000999 |
| RTM-likeMT | TCD8em | CXCL3 | S1PR4 | 1.7116 | 0.2677 | 0.9896 | 0.000999 |
| RTM-likeMT | TCD8em | FN1 | CD44 | 2.2884 | 1.119 | 1.7037 | 0.000999 |
| RTM-likeMT | TCD8em | FN1 | ITGA4 | 2.2884 | 0.2823 | 1.2854 | 0.000999 |
| RTM-likeMT | TCD8em | FN1 | ITGB1 | 2.2884 | 0.2557 | 1.2721 | 0.000999 |
| RTM-likeMT | TCD8em | FN1 | RPSA | 2.2884 | 2.3686 | 2.3285 | 0.000999 |
| RTM-likeMT | TCD8em | GRN | TNFRSF1B | 1.6859 | 0.3038 | 0.9948 | 0.000999 |
| RTM-likeMT | TCD8em | GSTP1 | CD44 | 1.5481 | 1.119 | 1.3336 | 0.000999 |
| RTM-likeMT | TCD8em | GSTP1 | IL7R | 1.5481 | 0.7183 | 1.1332 | 0.000999 |
| RTM-likeMT | TCD8em | GSTP1 | ITGA4 | 1.5481 | 0.2823 | 0.9152 | 0.000999 |
| RTM-likeMT | TCD8em | HBEGF | CD44 | 0.4141 | 1.119 | 0.7666 | 0.000999 |
| RTM-likeMT | TCD8em | HLA-DRA | PTPRC | 3.9047 | 1.7912 | 2.8479 | 0.000999 |
| RTM-likeMT | TCD8em | ICAM1 | IL2RG | 0.3448 | 0.9501 | 0.6474 | 0.000999 |
| RTM-likeMT | TCD8em | ICAM1 | ITGAL | 0.3448 | 0.3162 | 0.3305 | 0.000999 |
| RTM-likeMT | TCD8em | ICAM1 | ITGB2 | 0.3448 | 1.0018 | 0.6733 | 0.000999 |
| RTM-likeMT | TCD8em | ICAM1 | SPN | 0.3448 | 0.2603 | 0.3026 | 0.000999 |
| RTM-likeMT | TCD8em | IL10 | IL10RA | 0.2855 | 0.4383 | 0.3619 | 0.000999 |
| RTM-likeMT | TCD8em | ITGB2 | ITGAL | 1.4781 | 0.3162 | 0.8972 | 0.000999 |
| RTM-likeMT | TCD8em | LGALS3 | CD6 | 0.8673 | 0.4204 | 0.6439 | 0.000999 |
| RTM-likeMT | TCD8em | LGALS3 | ITGB1 | 0.8673 | 0.2557 | 0.5615 | 0.000999 |
| RTM-likeMT | TCD8em | LGALS3 | SPN | 0.8673 | 0.2603 | 0.5638 | 0.000999 |
| RTM-likeMT | TCD8em | MIF | CD74 | 1.0074 | 2.4538 | 1.7306 | 0.000999 |
| RTM-likeMT | TCD8em | MIF | ITGA4 | 1.0074 | 0.2823 | 0.6449 | 0.000999 |
| RTM-likeMT | TCD8em | MIF | TNFRSF14 | 1.0074 | 0.2915 | 0.6495 | 0.000999 |
| RTM-likeMT | TCD8em | NAMPT | ANXA6 | 1.5359 | 0.552 | 1.044 | 0.000999 |
| RTM-likeMT | TCD8em | PECAM1 | CXCR4 | 0.3422 | 2.8389 | 1.5906 | 0.000999 |
| RTM-likeMT | TCD8em | RTN4 | CD44 | 0.9166 | 1.119 | 1.0178 | 0.000999 |

| | | | | | | | |
|---|---|---|---|---|---|---|---|
| RTM-likeMT | TCD8em | RTN4 | CD99 | 0.9166 | 1.1525 | 1.0346 | 0.000999 |
| RTM-likeMT | TCD8em | SPP1 | CD44 | 0.7979 | 1.119 | 0.9585 | 0.000999 |
| RTM-likeMT | TCD8em | SPP1 | ITGA4 | 0.7979 | 0.2823 | 0.5401 | 0.000999 |
| RTM-likeMT | TCD8em | SPP1 | ITGB1 | 0.7979 | 0.2557 | 0.5268 | 0.000999 |
| RTM-likeMT | TCD8em | THBS1 | CD47 | 0.7512 | 0.4276 | 0.5894 | 0.000999 |
| RTM-likeMT | TCD8em | THBS1 | ITGA4 | 0.7512 | 0.2823 | 0.5168 | 0.000999 |
| RTM-likeMT | TCD8em | THBS1 | ITGB1 | 0.7512 | 0.2557 | 0.5035 | 0.000999 |
| RTM-likeMT | TCD8em | TNFSF13B | ITGB1 | 0.7616 | 0.2557 | 0.5087 | 0.000999 |
| RTM-likeMT | TCD8ex | A2M | TNFRSF14 | 0.7163 | 0.3094 | 0.5129 | 0.000999 |
| RTM-likeMT | TCD8ex | ADM | PTGER4 | 0.4876 | 0.3089 | 0.3983 | 0.000999 |
| RTM-likeMT | TCD8ex | ANXA1 | CD44 | 1.1172 | 0.8219 | 0.9696 | 0.000999 |
| RTM-likeMT | TCD8ex | ANXA1 | CXCR3 | 1.1172 | 0.3647 | 0.7409 | 0.000999 |
| RTM-likeMT | TCD8ex | ANXA1 | CXCR4 | 1.1172 | 2.3472 | 1.7322 | 0.000999 |
| RTM-likeMT | TCD8ex | ANXA1 | CXCR6 | 1.1172 | 0.6626 | 0.8899 | 0.000999 |
| RTM-likeMT | TCD8ex | ANXA1 | ITGA4 | 1.1172 | 0.4432 | 0.7802 | 0.000999 |
| RTM-likeMT | TCD8ex | ANXA1 | VCAM1 | 1.1172 | 0.4462 | 0.7817 | 0.000999 |
| RTM-likeMT | TCD8ex | APOE | VCAM1 | 3.1184 | 0.4462 | 1.7823 | 0.000999 |
| RTM-likeMT | TCD8ex | CCL20 | CXCR3 | 0.9244 | 0.3647 | 0.6445 | 0.000999 |
| RTM-likeMT | TCD8ex | CCL20 | CXCR4 | 0.9244 | 2.3472 | 1.6358 | 0.000999 |
| RTM-likeMT | TCD8ex | CCL20 | CXCR6 | 0.9244 | 0.6626 | 0.7935 | 0.000999 |
| RTM-likeMT | TCD8ex | CD86 | CTLA4 | 0.5683 | 0.4883 | 0.5283 | 0.000999 |
| RTM-likeMT | TCD8ex | COL1A1 | CD44 | 0.2826 | 0.8219 | 0.5522 | 0.000999 |
| RTM-likeMT | TCD8ex | COL1A1 | ITGB1 | 0.2826 | 0.2995 | 0.291 | 0.000999 |
| RTM-likeMT | TCD8ex | CXCL16 | CXCR3 | 1.0911 | 0.3647 | 0.7279 | 0.000999 |
| RTM-likeMT | TCD8ex | CXCL16 | CXCR4 | 1.0911 | 2.3472 | 1.7192 | 0.000999 |
| RTM-likeMT | TCD8ex | CXCL16 | CXCR6 | 1.0911 | 0.6626 | 0.8768 | 0.000999 |
| RTM-likeMT | TCD8ex | CXCL2 | CXCR3 | 1.9342 | 0.3647 | 1.1494 | 0.000999 |
| RTM-likeMT | TCD8ex | CXCL2 | CXCR4 | 1.9342 | 2.3472 | 2.1407 | 0.000999 |
| RTM-likeMT | TCD8ex | CXCL2 | CXCR6 | 1.9342 | 0.6626 | 1.2984 | 0.000999 |
| RTM-likeMT | TCD8ex | CXCL3 | CXCR3 | 1.7116 | 0.3647 | 1.0381 | 0.000999 |

| | | | | | | | |
|---|---|---|---|---|---|---|---|
| RTM-likeMT | TCD8ex | CXCL3 | CXCR4 | 1.7116 | 2.3472 | 2.0294 | 0.000999 |
| RTM-likeMT | TCD8ex | CXCL3 | CXCR6 | 1.7116 | 0.6626 | 1.1871 | 0.000999 |
| RTM-likeMT | TCD8ex | FN1 | CD44 | 2.2884 | 0.8219 | 1.5552 | 0.000999 |
| RTM-likeMT | TCD8ex | FN1 | ITGA4 | 2.2884 | 0.4432 | 1.3658 | 0.000999 |
| RTM-likeMT | TCD8ex | FN1 | ITGB1 | 2.2884 | 0.2995 | 1.294 | 0.000999 |
| RTM-likeMT | TCD8ex | FN1 | ITGB7 | 2.2884 | 0.3007 | 1.2946 | 0.000999 |
| RTM-likeMT | TCD8ex | FN1 | RPSA | 2.2884 | 2.2697 | 2.2791 | 0.000999 |
| RTM-likeMT | TCD8ex | GRN | TNFRSF1B | 1.6859 | 0.6436 | 1.1648 | 0.000999 |
| RTM-likeMT | TCD8ex | GSTP1 | CD44 | 1.5481 | 0.8219 | 1.185 | 0.000999 |
| RTM-likeMT | TCD8ex | GSTP1 | ITGA4 | 1.5481 | 0.4432 | 0.9957 | 0.000999 |
| RTM-likeMT | TCD8ex | GSTP1 | VCAM1 | 1.5481 | 0.4462 | 0.9972 | 0.000999 |
| RTM-likeMT | TCD8ex | HBEGF | CD44 | 0.4141 | 0.8219 | 0.618 | 0.000999 |
| RTM-likeMT | TCD8ex | HLA-DRA | PDCD1 | 3.9047 | 0.6109 | 2.2578 | 0.000999 |
| RTM-likeMT | TCD8ex | HLA-DRA | PTPRC | 3.9047 | 1.6762 | 2.7904 | 0.000999 |
| RTM-likeMT | TCD8ex | ICAM1 | IL2RG | 0.3448 | 1.2126 | 0.7787 | 0.000999 |
| RTM-likeMT | TCD8ex | ICAM1 | ITGB2 | 0.3448 | 0.7635 | 0.5541 | 0.000999 |
| RTM-likeMT | TCD8ex | ICAM1 | SPN | 0.3448 | 0.3486 | 0.3467 | 0.000999 |
| RTM-likeMT | TCD8ex | IL10 | IL10RA | 0.2855 | 0.3816 | 0.3335 | 0.000999 |
| RTM-likeMT | TCD8ex | LGALS3 | CD6 | 0.8673 | 0.318 | 0.5927 | 0.000999 |
| RTM-likeMT | TCD8ex | LGALS3 | ITGB1 | 0.8673 | 0.2995 | 0.5834 | 0.000999 |
| RTM-likeMT | TCD8ex | LGALS3 | SPN | 0.8673 | 0.3486 | 0.608 | 0.000999 |
| RTM-likeMT | TCD8ex | MIF | CD74 | 1.0074 | 2.5058 | 1.7566 | 0.000999 |
| RTM-likeMT | TCD8ex | MIF | ITGA4 | 1.0074 | 0.4432 | 0.7253 | 0.000999 |
| RTM-likeMT | TCD8ex | MIF | TNFRSF14 | 1.0074 | 0.3094 | 0.6584 | 0.000999 |
| RTM-likeMT | TCD8ex | NAMPT | ANXA6 | 1.5359 | 0.5678 | 1.0519 | 0.000999 |
| RTM-likeMT | TCD8ex | PECAM1 | CXCR4 | 0.3422 | 2.3472 | 1.3447 | 0.000999 |
| RTM-likeMT | TCD8ex | RTN4 | CD44 | 0.9166 | 0.8219 | 0.8693 | 0.000999 |
| RTM-likeMT | TCD8ex | RTN4 | CD99 | 0.9166 | 1.2049 | 1.0608 | 0.000999 |
| RTM-likeMT | TCD8ex | SPP1 | CD44 | 0.7979 | 0.8219 | 0.8099 | 0.000999 |
| RTM-likeMT | TCD8ex | SPP1 | ITGA4 | 0.7979 | 0.4432 | 0.6206 | 0.000999 |

| | | | | | | | |
|---|---|---|---|---|---|---|---|
| RTM-likeMT | TCD8ex | SPP1 | ITGB1 | 0.7979 | 0.2995 | 0.5487 | 0.000999 |
| RTM-likeMT | TCD8ex | THBS1 | CD47 | 0.7512 | 0.3952 | 0.5732 | 0.000999 |
| RTM-likeMT | TCD8ex | THBS1 | ITGA4 | 0.7512 | 0.4432 | 0.5972 | 0.000999 |
| RTM-likeMT | TCD8ex | THBS1 | ITGB1 | 0.7512 | 0.2995 | 0.5254 | 0.000999 |
| RTM-likeMT | TCD8ex | TNFSF13B | ITGB1 | 0.7616 | 0.2995 | 0.5305 | 0.000999 |
| RTM-likeMT | TGD | A2M | TNFRSF14 | 0.7163 | 0.4642 | 0.5902 | 0.000999 |
| RTM-likeMT | TGD | ADM | PTGDR | 0.4876 | 0.7153 | 0.6015 | 0.000999 |
| RTM-likeMT | TGD | ANXA1 | CXCR3 | 1.1172 | 1.1748 | 1.146 | 0.000999 |
| RTM-likeMT | TGD | ANXA1 | CXCR4 | 1.1172 | 1.4894 | 1.3033 | 0.000999 |
| RTM-likeMT | TGD | ANXA1 | ITGA4 | 1.1172 | 0.3968 | 0.757 | 0.000999 |
| RTM-likeMT | TGD | APOE | SORL1 | 3.1184 | 0.3318 | 1.7251 | 0.000999 |
| RTM-likeMT | TGD | CALM1 | RPSA | 0.8272 | 2.8814 | 1.8543 | 0.000999 |
| RTM-likeMT | TGD | CALR | IL7R | 0.7092 | 1.3862 | 1.0477 | 0.000999 |
| RTM-likeMT | TGD | CCL20 | CXCR3 | 0.9244 | 1.1748 | 1.0496 | 0.000999 |
| RTM-likeMT | TGD | CCL20 | CXCR4 | 0.9244 | 1.4894 | 1.2069 | 0.000999 |
| RTM-likeMT | TGD | CD99 | CD99 | 0.8632 | 1.7154 | 1.2893 | 0.000999 |
| RTM-likeMT | TGD | CXCL16 | CXCR3 | 1.0911 | 1.1748 | 1.133 | 0.000999 |
| RTM-likeMT | TGD | CXCL16 | CXCR4 | 1.0911 | 1.4894 | 1.2903 | 0.000999 |
| RTM-likeMT | TGD | CXCL2 | CXCR3 | 1.9342 | 1.1748 | 1.5545 | 0.000999 |
| RTM-likeMT | TGD | CXCL2 | CXCR4 | 1.9342 | 1.4894 | 1.7118 | 0.000999 |
| RTM-likeMT | TGD | CXCL3 | CXCR3 | 1.7116 | 1.1748 | 1.4432 | 0.000999 |
| RTM-likeMT | TGD | CXCL3 | CXCR4 | 1.7116 | 1.4894 | 1.6005 | 0.000999 |
| RTM-likeMT | TGD | FN1 | CD44 | 2.2884 | 1.1342 | 1.7113 | 0.000999 |
| RTM-likeMT | TGD | FN1 | ITGA4 | 2.2884 | 0.3968 | 1.3426 | 0.000999 |
| RTM-likeMT | TGD | FN1 | ITGB1 | 2.2884 | 0.3657 | 1.3271 | 0.000999 |
| RTM-likeMT | TGD | FN1 | ITGB7 | 2.2884 | 0.459 | 1.3737 | 0.000999 |
| RTM-likeMT | TGD | FN1 | RPSA | 2.2884 | 2.8814 | 2.5849 | 0.000999 |
| RTM-likeMT | TGD | GRN | TNFRSF1A | 1.6859 | 0.3099 | 0.9979 | 0.000999 |
| RTM-likeMT | TGD | GSTP1 | IL7R | 1.5481 | 1.3862 | 1.4672 | 0.000999 |
| RTM-likeMT | TGD | GSTP1 | ITGA4 | 1.5481 | 0.3968 | 0.9725 | 0.000999 |

| | | | | | | | |
|---|---|---|---|---|---|---|---|
| RTM-likeMT | TGD | HLA-A | CD8A | 2.8234 | 1.746 | 2.2847 | 0.000999 |
| RTM-likeMT | TGD | HLA-DRA | PTPRC | 3.9047 | 1.8653 | 2.885 | 0.000999 |
| RTM-likeMT | TGD | HLA-E | CD8A | 1.7632 | 1.746 | 1.7546 | 0.000999 |
| RTM-likeMT | TGD | HLA-E | KLRC2 | 1.7632 | 0.8082 | 1.2857 | 0.000999 |
| RTM-likeMT | TGD | HLA-E | KLRD1 | 1.7632 | 1.3451 | 1.5541 | 0.000999 |
| RTM-likeMT | TGD | HLA-F | CD8A | 0.3364 | 1.746 | 1.0412 | 0.000999 |
| RTM-likeMT | TGD | ICAM1 | IL2RG | 0.3448 | 1.561 | 0.9529 | 0.000999 |
| RTM-likeMT | TGD | ICAM1 | ITGAL | 0.3448 | 0.3731 | 0.359 | 0.000999 |
| RTM-likeMT | TGD | ICAM1 | SPN | 0.3448 | 0.5644 | 0.4546 | 0.000999 |
| RTM-likeMT | TGD | IL10 | IFNAR2 | 0.2855 | 0.4338 | 0.3596 | 0.000999 |
| RTM-likeMT | TGD | IL10 | IL10RA | 0.2855 | 0.625 | 0.4552 | 0.000999 |
| RTM-likeMT | TGD | ITGB1 | ITGA4 | 0.3078 | 0.3968 | 0.3523 | 0.000999 |
| RTM-likeMT | TGD | ITGB1 | RACK1 | 0.3078 | 2.7287 | 1.5182 | 0.000999 |
| RTM-likeMT | TGD | ITGB2 | ITGAL | 1.4781 | 0.3731 | 0.9256 | 0.000999 |
| RTM-likeMT | TGD | LGALS3 | CD6 | 0.8673 | 0.4684 | 0.6679 | 0.000999 |
| RTM-likeMT | TGD | LGALS3 | ITGB1 | 0.8673 | 0.3657 | 0.6165 | 0.000999 |
| RTM-likeMT | TGD | LGALS3 | SPN | 0.8673 | 0.5644 | 0.7158 | 0.000999 |
| RTM-likeMT | TGD | LGALS3 | TGFBR2 | 0.8673 | 0.3677 | 0.6175 | 0.000999 |
| RTM-likeMT | TGD | NAMPT | ANXA6 | 1.5359 | 0.8553 | 1.1956 | 0.000999 |
| RTM-likeMT | TGD | PECAM1 | CXCR4 | 0.3422 | 1.4894 | 0.9158 | 0.000999 |
| RTM-likeMT | TGD | PTPRC | CD2 | 0.5973 | 1.4522 | 1.0247 | 0.000999 |
| RTM-likeMT | TGD | PTPRC | CD8A | 0.5973 | 1.746 | 1.1716 | 0.000999 |
| RTM-likeMT | TGD | RPS19 | ITGA4 | 3.5378 | 0.3968 | 1.9673 | 0.000999 |
| RTM-likeMT | TGD | RPS19 | RPSA | 3.5378 | 2.8814 | 3.2096 | 0.000999 |
| RTM-likeMT | TGD | RTN4 | CD99 | 0.9166 | 1.7154 | 1.316 | 0.000999 |
| RTM-likeMT | TGD | SPP1 | ITGA4 | 0.7979 | 0.3968 | 0.5974 | 0.000999 |
| RTM-likeMT | TGD | SPP1 | ITGB1 | 0.7979 | 0.3657 | 0.5818 | 0.000999 |
| RTM-likeMT | TGD | THBS1 | CD47 | 0.7512 | 0.4765 | 0.6139 | 0.000999 |
| RTM-likeMT | TGD | THBS1 | ITGA4 | 0.7512 | 0.3968 | 0.574 | 0.000999 |
| RTM-likeMT | TGD | THBS1 | ITGB1 | 0.7512 | 0.3657 | 0.5584 | 0.000999 |

| | | | | | | | |
|---|---|---|---|---|---|---|---|
| RTM-likeMT | TGD | TNFSF13B | ITGB1 | 0.7616 | 0.3657 | 0.5636 | 0.000999 |
| RTM-likeMT | TGD | VEGFA | PTK2B | 0.512 | 0.3668 | 0.4394 | 0.000999 |
| RTM-likeMT | TGD | ARF1 | ITGA4 | 0.5429 | 0.3968 | 0.4699 | 0.002 |
| RTM-likeMT | TGD | ITGAM | ICAM1 | 0.4553 | 0.2915 | 0.3734 | 0.00699 |
| NKT | RTM-likeMT | ALOX5AP | ALOX5 | 0.5021 | 0.4191 | 0.4606 | 0.000999 |
| NKT | RTM-likeMT | ANXA1 | C3AR1 | 1.2259 | 0.4884 | 0.8572 | 0.000999 |
| NKT | RTM-likeMT | ANXA1 | C5AR1 | 1.2259 | 1.271 | 1.2485 | 0.000999 |
| NKT | RTM-likeMT | ANXA1 | CCR1 | 1.2259 | 0.2778 | 0.7519 | 0.000999 |
| NKT | RTM-likeMT | ANXA1 | CD4 | 1.2259 | 0.6983 | 0.9621 | 0.000999 |
| NKT | RTM-likeMT | ANXA1 | CD44 | 1.2259 | 1.4501 | 1.338 | 0.000999 |
| NKT | RTM-likeMT | ANXA1 | FPR1 | 1.2259 | 0.5 | 0.863 | 0.000999 |
| NKT | RTM-likeMT | ANXA1 | FPR3 | 1.2259 | 0.3995 | 0.8127 | 0.000999 |
| NKT | RTM-likeMT | ARF1 | CD4 | 0.7991 | 0.6983 | 0.7487 | 0.000999 |
| NKT | RTM-likeMT | ARF1 | CD44 | 0.7991 | 1.4501 | 1.1246 | 0.000999 |
| NKT | RTM-likeMT | CALM1 | TNFRSF1A | 1.9335 | 0.283 | 1.1083 | 0.000999 |
| NKT | RTM-likeMT | CALM1 | TNFRSF1B | 1.9335 | 0.7608 | 1.3472 | 0.000999 |
| NKT | RTM-likeMT | CALR | CD44 | 0.9479 | 1.4501 | 1.199 | 0.000999 |
| NKT | RTM-likeMT | CALR | LRP1 | 0.9479 | 0.6199 | 0.7839 | 0.000999 |
| NKT | RTM-likeMT | CD2 | CD4 | 1.1465 | 0.6983 | 0.9224 | 0.000999 |
| NKT | RTM-likeMT | GSTP1 | CD44 | 0.7909 | 1.4501 | 1.1205 | 0.000999 |
| NKT | RTM-likeMT | GZMB | NOTCH2 | 1.8361 | 0.3052 | 1.0706 | 0.000999 |
| NKT | RTM-likeMT | HLA-DRA | CD4 | 0.6626 | 0.6983 | 0.6805 | 0.000999 |
| NKT | RTM-likeMT | HMGB1 | AGTRAP | 1.195 | 0.3997 | 0.7973 | 0.000999 |
| NKT | RTM-likeMT | HMGB1 | LY96 | 1.195 | 1.4354 | 1.3152 | 0.000999 |
| NKT | RTM-likeMT | HMGB1 | TLR2 | 1.195 | 0.3069 | 0.7509 | 0.000999 |
| NKT | RTM-likeMT | HMGB2 | CD44 | 0.6467 | 1.4501 | 1.0484 | 0.000999 |
| NKT | RTM-likeMT | HMGB2 | DAB2 | 0.6467 | 0.7068 | 0.6768 | 0.000999 |
| NKT | RTM-likeMT | HSP90B1 | LRP1 | 0.6138 | 0.6199 | 0.6169 | 0.000999 |
| NKT | RTM-likeMT | HSP90B1 | TNFRSF1B | 0.6138 | 0.7608 | 0.6873 | 0.000999 |
| NKT | RTM-likeMT | IL16 | CD4 | 0.3556 | 0.6983 | 0.5269 | 0.000999 |

| | | | | | | | |
|---|---|---|---|---|---|---|---|
| NKT | RTM-likeMT | IL16 | FCGR1A | 0.3556 | 0.4653 | 0.4104 | 0.000999 |
| NKT | RTM-likeMT | ITGA4 | CD44 | 0.2946 | 1.4501 | 0.8723 | 0.000999 |
| NKT | RTM-likeMT | ITGA4 | TFRC | 0.2946 | 0.9121 | 0.6033 | 0.000999 |
| NKT | RTM-likeMT | ITGAL | ICAM1 | 0.6778 | 0.3448 | 0.5113 | 0.000999 |
| NKT | RTM-likeMT | ITGB1 | CD46 | 0.488 | 0.3174 | 0.4027 | 0.000999 |
| NKT | RTM-likeMT | ITGB1 | DAB2 | 0.488 | 0.7068 | 0.5974 | 0.000999 |
| NKT | RTM-likeMT | ITGB1 | ENG | 0.488 | 0.3413 | 0.4146 | 0.000999 |
| NKT | RTM-likeMT | ITGB1 | PLAUR | 0.488 | 1.2013 | 0.8446 | 0.000999 |
| NKT | RTM-likeMT | ITGB1 | PTK2B | 0.488 | 0.3072 | 0.3976 | 0.000999 |
| NKT | RTM-likeMT | ITGB2 | ICAM1 | 1.6481 | 0.3448 | 0.9964 | 0.000999 |
| NKT | RTM-likeMT | ITGB2 | ITGAM | 1.6481 | 0.4553 | 1.0517 | 0.000999 |
| NKT | RTM-likeMT | ITGB2 | PLAUR | 1.6481 | 1.2013 | 1.4247 | 0.000999 |
| NKT | RTM-likeMT | LRPAP1 | LRP1 | 0.2807 | 0.6199 | 0.4503 | 0.000999 |
| NKT | RTM-likeMT | LTB | TNFRSF1A | 0.2605 | 0.283 | 0.2717 | 0.000999 |
| NKT | RTM-likeMT | MIF | CD74 | 0.6657 | 4.0179 | 2.3418 | 0.000999 |
| NKT | RTM-likeMT | PSAP | LRP1 | 0.6777 | 0.6199 | 0.6488 | 0.000999 |
| NKT | RTM-likeMT | PTPRC | CD4 | 1.8797 | 0.6983 | 1.289 | 0.000999 |
| NKT | RTM-likeMT | PTPRC | CD44 | 1.8797 | 1.4501 | 1.6649 | 0.000999 |
| NKT | RTM-likeMT | PTPRC | IFNAR1 | 1.8797 | 0.3337 | 1.1067 | 0.000999 |
| NKT | RTM-likeMT | RPS19 | C5AR1 | 3.5336 | 1.271 | 2.4023 | 0.000999 |
| NKT | RTM-likeMT | RPS19 | CD4 | 3.5336 | 0.6983 | 2.116 | 0.000999 |
| NKT | RTM-likeMT | RPS19 | ICAM1 | 3.5336 | 0.3448 | 1.9392 | 0.000999 |
| NKT | RTM-likeMT | RTN4 | CD44 | 0.4374 | 1.4501 | 0.9438 | 0.000999 |
| NKT | RTM-likeMT | SPN | SIGLEC1 | 0.6398 | 0.2533 | 0.4465 | 0.000999 |
| NKT | RTM-likeMT | TGFB1 | ENG | 0.5441 | 0.3413 | 0.4427 | 0.000999 |
| NKT | RTM-likeMT | TGFB1 | FCN1 | 0.5441 | 1.1076 | 0.8259 | 0.000999 |
| NKT | RTM-likeMT | TGFB1 | TGFBR2 | 0.5441 | 0.3477 | 0.4459 | 0.000999 |
| NKT | RTM-likeMT | TGFB1 | TLR2 | 0.5441 | 0.3069 | 0.4255 | 0.000999 |
| NKT | RTM-likeMT | TIMP1 | LRP1 | 0.3005 | 0.6199 | 0.4602 | 0.000999 |
| NKT | RTM-likeMT | CD320 | TGFBR2 | 0.2545 | 0.3477 | 0.3011 | 0.002 |

| | | | | | | | |
|---|---|---|---|---|---|---|---|
| TCD4ex | RTM-likeMT | ALOX5AP | ALOX5 | 1.119 | 0.4191 | 0.769 | 0.000999 |
| TCD4ex | RTM-likeMT | ANXA1 | C3AR1 | 0.5174 | 0.4884 | 0.5029 | 0.000999 |
| TCD4ex | RTM-likeMT | ANXA1 | C5AR1 | 0.5174 | 1.271 | 0.8942 | 0.000999 |
| TCD4ex | RTM-likeMT | ANXA1 | CCR1 | 0.5174 | 0.2778 | 0.3976 | 0.000999 |
| TCD4ex | RTM-likeMT | ANXA1 | CD44 | 0.5174 | 1.4501 | 0.9838 | 0.000999 |
| TCD4ex | RTM-likeMT | ANXA1 | FPR1 | 0.5174 | 0.5 | 0.5087 | 0.000999 |
| TCD4ex | RTM-likeMT | ANXA1 | FPR3 | 0.5174 | 0.3995 | 0.4585 | 0.000999 |
| TCD4ex | RTM-likeMT | APOE | LRP1 | 0.2557 | 0.6199 | 0.4378 | 0.000999 |
| TCD4ex | RTM-likeMT | AREG | ICAM1 | 0.3427 | 0.3448 | 0.3438 | 0.000999 |
| TCD4ex | RTM-likeMT | ARF1 | CD44 | 0.6827 | 1.4501 | 1.0664 | 0.000999 |
| TCD4ex | RTM-likeMT | BTLA | TNFRSF14 | 0.3493 | 0.3616 | 0.3554 | 0.000999 |
| TCD4ex | RTM-likeMT | CALM1 | TNFRSF1A | 1.2957 | 0.283 | 0.7894 | 0.000999 |
| TCD4ex | RTM-likeMT | CALM1 | TNFRSF1B | 1.2957 | 0.7608 | 1.0283 | 0.000999 |
| TCD4ex | RTM-likeMT | CALR | CD44 | 0.748 | 1.4501 | 1.099 | 0.000999 |
| TCD4ex | RTM-likeMT | CALR | LRP1 | 0.748 | 0.6199 | 0.6839 | 0.000999 |
| TCD4ex | RTM-likeMT | CD2 | CD4 | 1.5731 | 0.6983 | 1.1357 | 0.000999 |
| TCD4ex | RTM-likeMT | CXCL13 | C3AR1 | 2.3367 | 0.4884 | 1.4126 | 0.000999 |
| TCD4ex | RTM-likeMT | CXCL13 | C5AR1 | 2.3367 | 1.271 | 1.8038 | 0.000999 |
| TCD4ex | RTM-likeMT | CXCL13 | CCR1 | 2.3367 | 0.2778 | 1.3072 | 0.000999 |
| TCD4ex | RTM-likeMT | CXCL13 | FPR1 | 2.3367 | 0.5 | 1.4184 | 0.000999 |
| TCD4ex | RTM-likeMT | GSTP1 | CD44 | 0.4481 | 1.4501 | 0.9491 | 0.000999 |
| TCD4ex | RTM-likeMT | HMGB1 | AGTRAP | 1.3876 | 0.3997 | 0.8936 | 0.000999 |
| TCD4ex | RTM-likeMT | HMGB1 | LY96 | 1.3876 | 1.4354 | 1.4115 | 0.000999 |
| TCD4ex | RTM-likeMT | HMGB1 | TLR2 | 1.3876 | 0.3069 | 0.8472 | 0.000999 |
| TCD4ex | RTM-likeMT | HMGB2 | CD44 | 1.0243 | 1.4501 | 1.2372 | 0.000999 |
| TCD4ex | RTM-likeMT | HMGB2 | DAB2 | 1.0243 | 0.7068 | 0.8656 | 0.000999 |
| TCD4ex | RTM-likeMT | HSP90B1 | LRP1 | 0.3944 | 0.6199 | 0.5071 | 0.000999 |
| TCD4ex | RTM-likeMT | HSP90B1 | TNFRSF1B | 0.3944 | 0.7608 | 0.5776 | 0.000999 |
| TCD4ex | RTM-likeMT | ICAM3 | ITGB2 | 0.573 | 1.4781 | 1.0256 | 0.000999 |
| TCD4ex | RTM-likeMT | ITGAL | ICAM1 | 0.2603 | 0.3448 | 0.3025 | 0.000999 |

| | | | | | | | |
|---|---|---|---|---|---|---|---|
| TCD4ex | RTM-likeMT | ITGB1 | DAB2 | 0.4615 | 0.7068 | 0.5842 | 0.000999 |
| TCD4ex | RTM-likeMT | ITGB1 | ENG | 0.4615 | 0.3413 | 0.4014 | 0.000999 |
| TCD4ex | RTM-likeMT | ITGB1 | PLAUR | 0.4615 | 1.2013 | 0.8314 | 0.000999 |
| TCD4ex | RTM-likeMT | ITGB1 | PTK2B | 0.4615 | 0.3072 | 0.3844 | 0.000999 |
| TCD4ex | RTM-likeMT | ITGB2 | ICAM1 | 0.7871 | 0.3448 | 0.5659 | 0.000999 |
| TCD4ex | RTM-likeMT | ITGB2 | ITGAM | 0.7871 | 0.4553 | 0.6212 | 0.000999 |
| TCD4ex | RTM-likeMT | ITGB2 | PLAUR | 0.7871 | 1.2013 | 0.9942 | 0.000999 |
| TCD4ex | RTM-likeMT | LRPAP1 | LRP1 | 0.2809 | 0.6199 | 0.4504 | 0.000999 |
| TCD4ex | RTM-likeMT | LTB | TNFRSF1A | 0.9655 | 0.283 | 0.6243 | 0.000999 |
| TCD4ex | RTM-likeMT | MIF | CD74 | 0.6999 | 4.0179 | 2.3589 | 0.000999 |
| TCD4ex | RTM-likeMT | NMB | GPR65 | 0.6391 | 0.3775 | 0.5083 | 0.000999 |
| TCD4ex | RTM-likeMT | PSAP | LRP1 | 0.5291 | 0.6199 | 0.5745 | 0.000999 |
| TCD4ex | RTM-likeMT | PTPRC | CD4 | 1.7493 | 0.6983 | 1.2238 | 0.000999 |
| TCD4ex | RTM-likeMT | PTPRC | CD44 | 1.7493 | 1.4501 | 1.5997 | 0.000999 |
| TCD4ex | RTM-likeMT | PTPRC | IFNAR1 | 1.7493 | 0.3337 | 1.0415 | 0.000999 |
| TCD4ex | RTM-likeMT | RPS19 | C5AR1 | 3.6329 | 1.271 | 2.452 | 0.000999 |
| TCD4ex | RTM-likeMT | RPS19 | ICAM1 | 3.6329 | 0.3448 | 1.9888 | 0.000999 |
| TCD4ex | RTM-likeMT | RTN4 | CD44 | 0.355 | 1.4501 | 0.9026 | 0.000999 |
| TCD4ex | RTM-likeMT | SELL | CD4 | 0.5618 | 0.6983 | 0.63 | 0.000999 |
| TCD4ex | RTM-likeMT | SPP1 | CD44 | 0.3314 | 1.4501 | 0.8907 | 0.000999 |
| TCD4ex | RTM-likeMT | TIMP1 | LRP1 | 0.3374 | 0.6199 | 0.4787 | 0.000999 |
| TCD4ex | RTM-likeMT | ITGB1 | CD46 | 0.4615 | 0.3174 | 0.3895 | 0.005 |
| TCD4ex | RTM-likeMT | CD28 | CD4 | 0.3256 | 0.6983 | 0.5119 | 0.00599 |
| TCD4ex | RTM-likeMT | ARF1 | CD4 | 0.6827 | 0.6983 | 0.6905 | 0.00899 |
| TCD4reg | RTM-likeMT | ALOX5AP | ALOX5 | 0.4589 | 0.4191 | 0.439 | 0.000999 |
| TCD4reg | RTM-likeMT | APOE | LRP1 | 0.2934 | 0.6199 | 0.4567 | 0.000999 |
| TCD4reg | RTM-likeMT | ARF1 | CD4 | 0.7185 | 0.6983 | 0.7084 | 0.000999 |
| TCD4reg | RTM-likeMT | ARF1 | CD44 | 0.7185 | 1.4501 | 1.0843 | 0.000999 |
| TCD4reg | RTM-likeMT | CALM1 | TNFRSF1A | 1.534 | 0.283 | 0.9085 | 0.000999 |
| TCD4reg | RTM-likeMT | CALR | CD44 | 0.6835 | 1.4501 | 1.0668 | 0.000999 |

| | | | | | | | |
|---|---|---|---|---|---|---|---|
| TCD4reg | RTM-likeMT | CALR | LRP1 | 0.6835 | 0.6199 | 0.6517 | 0.000999 |
| TCD4reg | RTM-likeMT | CD2 | CD4 | 1.6566 | 0.6983 | 1.1775 | 0.000999 |
| TCD4reg | RTM-likeMT | CD28 | CD4 | 0.313 | 0.6983 | 0.5057 | 0.000999 |
| TCD4reg | RTM-likeMT | CD99 | CD99 | 0.6603 | 0.8632 | 0.7618 | 0.000999 |
| TCD4reg | RTM-likeMT | GSTP1 | CD44 | 0.4917 | 1.4501 | 0.9709 | 0.000999 |
| TCD4reg | RTM-likeMT | HLA-DMA | CD4 | 0.278 | 0.6983 | 0.4881 | 0.000999 |
| TCD4reg | RTM-likeMT | HMGB1 | AGTRAP | 1.2547 | 0.3997 | 0.8272 | 0.000999 |
| TCD4reg | RTM-likeMT | HMGB1 | LY96 | 1.2547 | 1.4354 | 1.345 | 0.000999 |
| TCD4reg | RTM-likeMT | HMGB1 | TLR2 | 1.2547 | 0.3069 | 0.7808 | 0.000999 |
| TCD4reg | RTM-likeMT | HMGB2 | CD44 | 0.616 | 1.4501 | 1.0331 | 0.000999 |
| TCD4reg | RTM-likeMT | HMGB2 | DAB2 | 0.616 | 0.7068 | 0.6614 | 0.000999 |
| TCD4reg | RTM-likeMT | HSP90B1 | LRP1 | 0.4221 | 0.6199 | 0.521 | 0.000999 |
| TCD4reg | RTM-likeMT | ICAM3 | ITGB2 | 0.6663 | 1.4781 | 1.0722 | 0.000999 |
| TCD4reg | RTM-likeMT | IL16 | CD4 | 0.2528 | 0.6983 | 0.4756 | 0.000999 |
| TCD4reg | RTM-likeMT | IL16 | FCGR1A | 0.2528 | 0.4653 | 0.3591 | 0.000999 |
| TCD4reg | RTM-likeMT | ITGB1 | CD46 | 0.4001 | 0.3174 | 0.3588 | 0.000999 |
| TCD4reg | RTM-likeMT | ITGB1 | DAB2 | 0.4001 | 0.7068 | 0.5534 | 0.000999 |
| TCD4reg | RTM-likeMT | ITGB1 | ENG | 0.4001 | 0.3413 | 0.3707 | 0.000999 |
| TCD4reg | RTM-likeMT | ITGB1 | PLAUR | 0.4001 | 1.2013 | 0.8007 | 0.000999 |
| TCD4reg | RTM-likeMT | ITGB1 | PTK2B | 0.4001 | 0.3072 | 0.3537 | 0.000999 |
| TCD4reg | RTM-likeMT | ITGB2 | ICAM1 | 0.6238 | 0.3448 | 0.4843 | 0.000999 |
| TCD4reg | RTM-likeMT | ITGB2 | ITGAM | 0.6238 | 0.4553 | 0.5396 | 0.000999 |
| TCD4reg | RTM-likeMT | ITGB2 | PLAUR | 0.6238 | 1.2013 | 0.9126 | 0.000999 |
| TCD4reg | RTM-likeMT | LGALS3 | CLEC7A | 0.4492 | 0.7806 | 0.6149 | 0.000999 |
| TCD4reg | RTM-likeMT | LGALS3 | FCGR2A | 0.4492 | 1.0278 | 0.7385 | 0.000999 |
| TCD4reg | RTM-likeMT | LGALS3 | FCGR2B | 0.4492 | 0.898 | 0.6736 | 0.000999 |
| TCD4reg | RTM-likeMT | LGALS3 | TFRC | 0.4492 | 0.9121 | 0.6806 | 0.000999 |
| TCD4reg | RTM-likeMT | LGALS3 | TGFBR2 | 0.4492 | 0.3477 | 0.3984 | 0.000999 |
| TCD4reg | RTM-likeMT | LRPAP1 | LRP1 | 0.3101 | 0.6199 | 0.465 | 0.000999 |
| TCD4reg | RTM-likeMT | LTB | TNFRSF1A | 2.242 | 0.283 | 1.2625 | 0.000999 |

| | | | | | | | |
|---|---|---|---|---|---|---|---|
| TCD4reg | RTM-likeMT | MIF | CD74 | 0.6295 | 4.0179 | 2.3237 | 0.000999 |
| TCD4reg | RTM-likeMT | PSAP | LRP1 | 0.5938 | 0.6199 | 0.6069 | 0.000999 |
| TCD4reg | RTM-likeMT | PTPRC | CD4 | 1.6603 | 0.6983 | 1.1793 | 0.000999 |
| TCD4reg | RTM-likeMT | PTPRC | CD44 | 1.6603 | 1.4501 | 1.5552 | 0.000999 |
| TCD4reg | RTM-likeMT | PTPRC | IFNAR1 | 1.6603 | 0.3337 | 0.997 | 0.000999 |
| TCD4reg | RTM-likeMT | RPS19 | C5AR1 | 3.6897 | 1.271 | 2.4803 | 0.000999 |
| TCD4reg | RTM-likeMT | RPS19 | CD4 | 3.6897 | 0.6983 | 2.194 | 0.000999 |
| TCD4reg | RTM-likeMT | RPS19 | ICAM1 | 3.6897 | 0.3448 | 2.0172 | 0.000999 |
| TCD4reg | RTM-likeMT | RTN4 | CD44 | 0.3528 | 1.4501 | 0.9014 | 0.000999 |
| TCD4reg | RTM-likeMT | RTN4 | CD99 | 0.3528 | 0.8632 | 0.608 | 0.000999 |
| TCD4reg | RTM-likeMT | SELL | CD4 | 0.7062 | 0.6983 | 0.7022 | 0.000999 |
| TCD4reg | RTM-likeMT | SPP1 | CD44 | 0.3125 | 1.4501 | 0.8813 | 0.000999 |
| TCD4reg | RTM-likeMT | TIMP1 | LRP1 | 0.4362 | 0.6199 | 0.5281 | 0.000999 |
| TCD8em | RTM-likeMT | ALOX5AP | ALOX5 | 0.7973 | 0.4191 | 0.6082 | 0.000999 |
| TCD8em | RTM-likeMT | ANXA1 | C3AR1 | 1.0693 | 0.4884 | 0.7788 | 0.000999 |
| TCD8em | RTM-likeMT | ANXA1 | C5AR1 | 1.0693 | 1.271 | 1.1701 | 0.000999 |
| TCD8em | RTM-likeMT | ANXA1 | CCR1 | 1.0693 | 0.2778 | 0.6735 | 0.000999 |
| TCD8em | RTM-likeMT | ANXA1 | CD4 | 1.0693 | 0.6983 | 0.8838 | 0.000999 |
| TCD8em | RTM-likeMT | ANXA1 | CD44 | 1.0693 | 1.4501 | 1.2597 | 0.000999 |
| TCD8em | RTM-likeMT | ANXA1 | FPR1 | 1.0693 | 0.5 | 0.7847 | 0.000999 |
| TCD8em | RTM-likeMT | ANXA1 | FPR3 | 1.0693 | 0.3995 | 0.7344 | 0.000999 |
| TCD8em | RTM-likeMT | APOE | LRP1 | 0.2791 | 0.6199 | 0.4495 | 0.000999 |
| TCD8em | RTM-likeMT | ARF1 | CD4 | 0.6684 | 0.6983 | 0.6833 | 0.000999 |
| TCD8em | RTM-likeMT | ARF1 | CD44 | 0.6684 | 1.4501 | 1.0592 | 0.000999 |
| TCD8em | RTM-likeMT | CALM1 | TNFRSF1A | 1.5262 | 0.283 | 0.9046 | 0.000999 |
| TCD8em | RTM-likeMT | CALM1 | TNFRSF1B | 1.5262 | 0.7608 | 1.1435 | 0.000999 |
| TCD8em | RTM-likeMT | CALR | CD44 | 0.7141 | 1.4501 | 1.0821 | 0.000999 |
| TCD8em | RTM-likeMT | CALR | LRP1 | 0.7141 | 0.6199 | 0.667 | 0.000999 |
| TCD8em | RTM-likeMT | CD2 | CD4 | 1.5961 | 0.6983 | 1.1472 | 0.000999 |
| TCD8em | RTM-likeMT | GSTP1 | CD44 | 0.767 | 1.4501 | 1.1086 | 0.000999 |

| | | | | | | | |
|---|---|---|---|---|---|---|---|
| TCD8em | RTM-likeMT | GZMB | NOTCH2 | 0.4781 | 0.3052 | 0.3916 | 0.000999 |
| TCD8em | RTM-likeMT | HLA-DMA | CD4 | 0.2969 | 0.6983 | 0.4976 | 0.000999 |
| TCD8em | RTM-likeMT | HLA-DRA | CD4 | 0.952 | 0.6983 | 0.8252 | 0.000999 |
| TCD8em | RTM-likeMT | HMGB1 | AGTRAP | 1.3636 | 0.3997 | 0.8816 | 0.000999 |
| TCD8em | RTM-likeMT | HMGB1 | LY96 | 1.3636 | 1.4354 | 1.3995 | 0.000999 |
| TCD8em | RTM-likeMT | HMGB1 | TLR2 | 1.3636 | 0.3069 | 0.8352 | 0.000999 |
| TCD8em | RTM-likeMT | HMGB2 | CD44 | 0.7212 | 1.4501 | 1.0856 | 0.000999 |
| TCD8em | RTM-likeMT | HMGB2 | DAB2 | 0.7212 | 0.7068 | 0.714 | 0.000999 |
| TCD8em | RTM-likeMT | HSP90B1 | LRP1 | 0.471 | 0.6199 | 0.5455 | 0.000999 |
| TCD8em | RTM-likeMT | HSP90B1 | TNFRSF1B | 0.471 | 0.7608 | 0.6159 | 0.000999 |
| TCD8em | RTM-likeMT | ICAM3 | ITGB2 | 0.5282 | 1.4781 | 1.0032 | 0.000999 |
| TCD8em | RTM-likeMT | IFNG | IFNGR1 | 0.3967 | 0.5267 | 0.4617 | 0.000999 |
| TCD8em | RTM-likeMT | IFNG | IFNGR2 | 0.3967 | 0.7344 | 0.5655 | 0.000999 |
| TCD8em | RTM-likeMT | IL16 | CD4 | 0.4085 | 0.6983 | 0.5534 | 0.000999 |
| TCD8em | RTM-likeMT | IL16 | FCGR1A | 0.4085 | 0.4653 | 0.4369 | 0.000999 |
| TCD8em | RTM-likeMT | ITGA4 | CD44 | 0.2823 | 1.4501 | 0.8662 | 0.000999 |
| TCD8em | RTM-likeMT | ITGA4 | TFRC | 0.2823 | 0.9121 | 0.5972 | 0.000999 |
| TCD8em | RTM-likeMT | ITGAL | ICAM1 | 0.3162 | 0.3448 | 0.3305 | 0.000999 |
| TCD8em | RTM-likeMT | ITGB1 | CD46 | 0.2557 | 0.3174 | 0.2866 | 0.000999 |
| TCD8em | RTM-likeMT | ITGB1 | DAB2 | 0.2557 | 0.7068 | 0.4813 | 0.000999 |
| TCD8em | RTM-likeMT | ITGB1 | ENG | 0.2557 | 0.3413 | 0.2985 | 0.000999 |
| TCD8em | RTM-likeMT | ITGB1 | PLAUR | 0.2557 | 1.2013 | 0.7285 | 0.000999 |
| TCD8em | RTM-likeMT | ITGB1 | PTK2B | 0.2557 | 0.3072 | 0.2815 | 0.000999 |
| TCD8em | RTM-likeMT | ITGB2 | ICAM1 | 1.0018 | 0.3448 | 0.6733 | 0.000999 |
| TCD8em | RTM-likeMT | ITGB2 | ITGAM | 1.0018 | 0.4553 | 0.7285 | 0.000999 |
| TCD8em | RTM-likeMT | ITGB2 | PLAUR | 1.0018 | 1.2013 | 1.1016 | 0.000999 |
| TCD8em | RTM-likeMT | LGALS3 | CLEC7A | 0.2689 | 0.7806 | 0.5248 | 0.000999 |
| TCD8em | RTM-likeMT | LGALS3 | FCGR2A | 0.2689 | 1.0278 | 0.6484 | 0.000999 |
| TCD8em | RTM-likeMT | LGALS3 | FCGR2B | 0.2689 | 0.898 | 0.5834 | 0.000999 |
| TCD8em | RTM-likeMT | LGALS3 | TFRC | 0.2689 | 0.9121 | 0.5905 | 0.000999 |

| | | | | | | | |
|---|---|---|---|---|---|---|---|
| TCD8em | RTM-likeMT | LGALS3 | TGFBR2 | 0.2689 | 0.3477 | 0.3083 | 0.000999 |
| TCD8em | RTM-likeMT | LRPAP1 | LRP1 | 0.2719 | 0.6199 | 0.4459 | 0.000999 |
| TCD8em | RTM-likeMT | LRPAP1 | SORL1 | 0.2719 | 0.3023 | 0.2871 | 0.000999 |
| TCD8em | RTM-likeMT | LTB | TNFRSF1A | 0.6352 | 0.283 | 0.4591 | 0.000999 |
| TCD8em | RTM-likeMT | MIF | CD74 | 0.6063 | 4.0179 | 2.3121 | 0.000999 |
| TCD8em | RTM-likeMT | PSAP | LRP1 | 0.5012 | 0.6199 | 0.5606 | 0.000999 |
| TCD8em | RTM-likeMT | PTPRC | CD4 | 1.7912 | 0.6983 | 1.2447 | 0.000999 |
| TCD8em | RTM-likeMT | PTPRC | CD44 | 1.7912 | 1.4501 | 1.6207 | 0.000999 |
| TCD8em | RTM-likeMT | PTPRC | IFNAR1 | 1.7912 | 0.3337 | 1.0624 | 0.000999 |
| TCD8em | RTM-likeMT | RPS19 | C5AR1 | 3.536 | 1.271 | 2.4035 | 0.000999 |
| TCD8em | RTM-likeMT | RPS19 | CD4 | 3.536 | 0.6983 | 2.1172 | 0.000999 |
| TCD8em | RTM-likeMT | RPS19 | ICAM1 | 3.536 | 0.3448 | 1.9404 | 0.000999 |
| TCD8em | RTM-likeMT | RTN4 | CD44 | 0.3632 | 1.4501 | 0.9066 | 0.000999 |
| TCD8em | RTM-likeMT | SELL | CD4 | 0.258 | 0.6983 | 0.4781 | 0.000999 |
| TCD8em | RTM-likeMT | SPN | SIGLEC1 | 0.2603 | 0.2533 | 0.2568 | 0.000999 |
| TCD8em | RTM-likeMT | SPP1 | CD44 | 0.3608 | 1.4501 | 0.9054 | 0.000999 |
| TCD8em | RTM-likeMT | TIMP1 | LRP1 | 0.2667 | 0.6199 | 0.4433 | 0.000999 |
| TCD8em | RTM-likeMT | TNF | TNFRSF1A | 0.2667 | 0.283 | 0.2749 | 0.000999 |
| TCD8em | RTM-likeMT | TNF | TNFRSF1B | 0.2667 | 0.7608 | 0.5138 | 0.000999 |
| TCD8em | RTM-likeMT | XCL2 | GPR65 | 0.295 | 0.3775 | 0.3362 | 0.000999 |
| TCD8ex | RTM-likeMT | ALOX5AP | ALOX5 | 1.5024 | 0.4191 | 0.9607 | 0.000999 |
| TCD8ex | RTM-likeMT | ANXA1 | C3AR1 | 0.5645 | 0.4884 | 0.5265 | 0.000999 |
| TCD8ex | RTM-likeMT | ANXA1 | C5AR1 | 0.5645 | 1.271 | 0.9177 | 0.000999 |
| TCD8ex | RTM-likeMT | ANXA1 | CCR1 | 0.5645 | 0.2778 | 0.4211 | 0.000999 |
| TCD8ex | RTM-likeMT | ANXA1 | CD4 | 0.5645 | 0.6983 | 0.6314 | 0.000999 |
| TCD8ex | RTM-likeMT | ANXA1 | CD44 | 0.5645 | 1.4501 | 1.0073 | 0.000999 |
| TCD8ex | RTM-likeMT | ANXA1 | FPR1 | 0.5645 | 0.5 | 0.5323 | 0.000999 |
| TCD8ex | RTM-likeMT | ANXA1 | FPR3 | 0.5645 | 0.3995 | 0.482 | 0.000999 |
| TCD8ex | RTM-likeMT | APOE | LRP1 | 0.2814 | 0.6199 | 0.4506 | 0.000999 |
| TCD8ex | RTM-likeMT | APOE | SORL1 | 0.2814 | 0.3023 | 0.2918 | 0.000999 |

| | | | | | | | |
|---|---|---|---|---|---|---|---|
| TCD8ex | RTM-likeMT | AREG | ICAM1 | 0.2758 | 0.3448 | 0.3103 | 0.000999 |
| TCD8ex | RTM-likeMT | ARF1 | CD4 | 0.7835 | 0.6983 | 0.7409 | 0.000999 |
| TCD8ex | RTM-likeMT | ARF1 | CD44 | 0.7835 | 1.4501 | 1.1168 | 0.000999 |
| TCD8ex | RTM-likeMT | CALM1 | TNFRSF1A | 1.4627 | 0.283 | 0.8729 | 0.000999 |
| TCD8ex | RTM-likeMT | CALM1 | TNFRSF1B | 1.4627 | 0.7608 | 1.1118 | 0.000999 |
| TCD8ex | RTM-likeMT | CALR | CD44 | 0.9634 | 1.4501 | 1.2067 | 0.000999 |
| TCD8ex | RTM-likeMT | CALR | LRP1 | 0.9634 | 0.6199 | 0.7916 | 0.000999 |
| TCD8ex | RTM-likeMT | CD2 | CD4 | 2.0502 | 0.6983 | 1.3742 | 0.000999 |
| TCD8ex | RTM-likeMT | CXCL13 | C3AR1 | 1.7749 | 0.4884 | 1.1317 | 0.000999 |
| TCD8ex | RTM-likeMT | CXCL13 | C5AR1 | 1.7749 | 1.271 | 1.523 | 0.000999 |
| TCD8ex | RTM-likeMT | CXCL13 | CCR1 | 1.7749 | 0.2778 | 1.0264 | 0.000999 |
| TCD8ex | RTM-likeMT | CXCL13 | FPR1 | 1.7749 | 0.5 | 1.1375 | 0.000999 |
| TCD8ex | RTM-likeMT | FASLG | TNFRSF1A | 0.2613 | 0.283 | 0.2721 | 0.000999 |
| TCD8ex | RTM-likeMT | GSTP1 | CD44 | 0.8286 | 1.4501 | 1.1394 | 0.000999 |
| TCD8ex | RTM-likeMT | GZMB | NOTCH2 | 1.9794 | 0.3052 | 1.1423 | 0.000999 |
| TCD8ex | RTM-likeMT | HLA-DMA | CD4 | 0.3991 | 0.6983 | 0.5487 | 0.000999 |
| TCD8ex | RTM-likeMT | HLA-DRA | CD4 | 1.247 | 0.6983 | 0.9726 | 0.000999 |
| TCD8ex | RTM-likeMT | HMGB1 | AGTRAP | 1.5414 | 0.3997 | 0.9706 | 0.000999 |
| TCD8ex | RTM-likeMT | HMGB1 | LY96 | 1.5414 | 1.4354 | 1.4884 | 0.000999 |
| TCD8ex | RTM-likeMT | HMGB1 | TLR2 | 1.5414 | 0.3069 | 0.9242 | 0.000999 |
| TCD8ex | RTM-likeMT | HMGB2 | CD44 | 0.85 | 1.4501 | 1.1501 | 0.000999 |
| TCD8ex | RTM-likeMT | HMGB2 | DAB2 | 0.85 | 0.7068 | 0.7784 | 0.000999 |
| TCD8ex | RTM-likeMT | HSP90B1 | LRP1 | 0.5666 | 0.6199 | 0.5933 | 0.000999 |
| TCD8ex | RTM-likeMT | ICAM3 | ITGB2 | 0.5405 | 1.4781 | 1.0093 | 0.000999 |
| TCD8ex | RTM-likeMT | IFNG | IFNGR1 | 0.8575 | 0.5267 | 0.6921 | 0.000999 |
| TCD8ex | RTM-likeMT | IFNG | IFNGR2 | 0.8575 | 0.7344 | 0.7959 | 0.000999 |
| TCD8ex | RTM-likeMT | ITGA4 | CD44 | 0.4432 | 1.4501 | 0.9467 | 0.000999 |
| TCD8ex | RTM-likeMT | ITGA4 | TFRC | 0.4432 | 0.9121 | 0.6777 | 0.000999 |
| TCD8ex | RTM-likeMT | ITGB1 | DAB2 | 0.2995 | 0.7068 | 0.5031 | 0.000999 |
| TCD8ex | RTM-likeMT | ITGB1 | ENG | 0.2995 | 0.3413 | 0.3204 | 0.000999 |

| | | | | | | | |
|---|---|---|---|---|---|---|---|
| TCD8ex | RTM-likeMT | ITGB1 | PLAUR | 0.2995 | 1.2013 | 0.7504 | 0.000999 |
| TCD8ex | RTM-likeMT | ITGB1 | PTK2B | 0.2995 | 0.3072 | 0.3034 | 0.000999 |
| TCD8ex | RTM-likeMT | ITGB2 | ICAM1 | 0.7635 | 0.3448 | 0.5541 | 0.000999 |
| TCD8ex | RTM-likeMT | ITGB2 | ITGAM | 0.7635 | 0.4553 | 0.6094 | 0.000999 |
| TCD8ex | RTM-likeMT | ITGB2 | PLAUR | 0.7635 | 1.2013 | 0.9824 | 0.000999 |
| TCD8ex | RTM-likeMT | LGALS3 | CLEC7A | 0.3824 | 0.7806 | 0.5815 | 0.000999 |
| TCD8ex | RTM-likeMT | LGALS3 | FCGR2A | 0.3824 | 1.0278 | 0.7051 | 0.000999 |
| TCD8ex | RTM-likeMT | LGALS3 | FCGR2B | 0.3824 | 0.898 | 0.6402 | 0.000999 |
| TCD8ex | RTM-likeMT | LGALS3 | TFRC | 0.3824 | 0.9121 | 0.6472 | 0.000999 |
| TCD8ex | RTM-likeMT | LGALS3 | TGFBR2 | 0.3824 | 0.3477 | 0.365 | 0.000999 |
| TCD8ex | RTM-likeMT | LRPAP1 | LRP1 | 0.2971 | 0.6199 | 0.4585 | 0.000999 |
| TCD8ex | RTM-likeMT | LRPAP1 | SORL1 | 0.2971 | 0.3023 | 0.2997 | 0.000999 |
| TCD8ex | RTM-likeMT | LTB | TNFRSF1A | 0.4576 | 0.283 | 0.3703 | 0.000999 |
| TCD8ex | RTM-likeMT | MIF | CD74 | 0.749 | 4.0179 | 2.3835 | 0.000999 |
| TCD8ex | RTM-likeMT | PSAP | LRP1 | 0.5016 | 0.6199 | 0.5608 | 0.000999 |
| TCD8ex | RTM-likeMT | PTPRC | CD4 | 1.6762 | 0.6983 | 1.1872 | 0.000999 |
| TCD8ex | RTM-likeMT | PTPRC | CD44 | 1.6762 | 1.4501 | 1.5631 | 0.000999 |
| TCD8ex | RTM-likeMT | PTPRC | IFNAR1 | 1.6762 | 0.3337 | 1.0049 | 0.000999 |
| TCD8ex | RTM-likeMT | RPS19 | C5AR1 | 3.8353 | 1.271 | 2.5531 | 0.000999 |
| TCD8ex | RTM-likeMT | RPS19 | CD4 | 3.8353 | 0.6983 | 2.2668 | 0.000999 |
| TCD8ex | RTM-likeMT | RPS19 | ICAM1 | 3.8353 | 0.3448 | 2.09 | 0.000999 |
| TCD8ex | RTM-likeMT | RTN4 | CD44 | 0.3033 | 1.4501 | 0.8767 | 0.000999 |
| TCD8ex | RTM-likeMT | SPN | SIGLEC1 | 0.3486 | 0.2533 | 0.301 | 0.000999 |
| TCD8ex | RTM-likeMT | SPP1 | CD44 | 0.2978 | 1.4501 | 0.874 | 0.000999 |
| TCD8ex | RTM-likeMT | VCAM1 | ITGB2 | 0.4462 | 1.4781 | 0.9622 | 0.000999 |
| TCD8ex | RTM-likeMT | VCAM1 | TFRC | 0.4462 | 0.9121 | 0.6791 | 0.000999 |
| TCD8ex | RTM-likeMT | ITGB1 | CD46 | 0.2995 | 0.3174 | 0.3085 | 0.002 |
| TCD8ex | RTM-likeMT | HSP90B1 | TNFRSF1B | 0.5666 | 0.7608 | 0.6637 | 0.00799 |
| TGD | RTM-likeMT | ALOX5AP | ALOX5 | 0.5564 | 0.4191 | 0.4878 | 0.000999 |
| TGD | RTM-likeMT | ANXA1 | C5AR1 | 0.7073 | 1.271 | 0.9891 | 0.000999 |

| | | | | | | | |
|---|---|---|---|---|---|---|---|
| TGD | RTM-likeMT | ARF1 | CD4 | 0.9453 | 0.6983 | 0.8218 | 0.000999 |
| TGD | RTM-likeMT | ARF1 | CD44 | 0.9453 | 1.4501 | 1.1977 | 0.000999 |
| TGD | RTM-likeMT | CALM1 | RPSA | 1.7587 | 1.7822 | 1.7704 | 0.000999 |
| TGD | RTM-likeMT | CALM1 | TNFRSF1A | 1.7587 | 0.283 | 1.0209 | 0.000999 |
| TGD | RTM-likeMT | CALM1 | TNFRSF1B | 1.7587 | 0.7608 | 1.2598 | 0.000999 |
| TGD | RTM-likeMT | CALR | CD44 | 0.9627 | 1.4501 | 1.2064 | 0.000999 |
| TGD | RTM-likeMT | CALR | LRP1 | 0.9627 | 0.6199 | 0.7913 | 0.000999 |
| TGD | RTM-likeMT | CD2 | CD4 | 1.4522 | 0.6983 | 1.0753 | 0.000999 |
| TGD | RTM-likeMT | CD2 | PTPRC | 1.4522 | 0.5973 | 1.0247 | 0.000999 |
| TGD | RTM-likeMT | CD99 | CD99 | 1.7154 | 0.8632 | 1.2893 | 0.000999 |
| TGD | RTM-likeMT | COL1A1 | CD44 | 0.4518 | 1.4501 | 0.951 | 0.000999 |
| TGD | RTM-likeMT | HMGB1 | AGTRAP | 1.7525 | 0.3997 | 1.0761 | 0.000999 |
| TGD | RTM-likeMT | HMGB1 | LY96 | 1.7525 | 1.4354 | 1.5939 | 0.000999 |
| TGD | RTM-likeMT | HMGB1 | TLR2 | 1.7525 | 0.3069 | 1.0297 | 0.000999 |
| TGD | RTM-likeMT | HSP90B1 | ITGB1 | 0.9338 | 0.3078 | 0.6208 | 0.000999 |
| TGD | RTM-likeMT | HSP90B1 | LRP1 | 0.9338 | 0.6199 | 0.7769 | 0.000999 |
| TGD | RTM-likeMT | HSP90B1 | TNFRSF1B | 0.9338 | 0.7608 | 0.8473 | 0.000999 |
| TGD | RTM-likeMT | ICAM3 | ITGB2 | 0.8816 | 1.4781 | 1.1799 | 0.000999 |
| TGD | RTM-likeMT | IL16 | CD4 | 0.9357 | 0.6983 | 0.817 | 0.000999 |
| TGD | RTM-likeMT | IL16 | FCGR1A | 0.9357 | 0.4653 | 0.7005 | 0.000999 |
| TGD | RTM-likeMT | ITGA4 | CD44 | 0.3968 | 1.4501 | 0.9235 | 0.000999 |
| TGD | RTM-likeMT | ITGA4 | TFRC | 0.3968 | 0.9121 | 0.6545 | 0.000999 |
| TGD | RTM-likeMT | ITGAL | ICAM1 | 0.3731 | 0.3448 | 0.359 | 0.000999 |
| TGD | RTM-likeMT | ITGB1 | DAB2 | 0.3657 | 0.7068 | 0.5362 | 0.000999 |
| TGD | RTM-likeMT | ITGB1 | ENG | 0.3657 | 0.3413 | 0.3535 | 0.000999 |
| TGD | RTM-likeMT | ITGB1 | PLAUR | 0.3657 | 1.2013 | 0.7835 | 0.000999 |
| TGD | RTM-likeMT | ITGB2 | PLAUR | 1.4444 | 1.2013 | 1.3229 | 0.000999 |
| TGD | RTM-likeMT | LRPAP1 | LRP1 | 0.3161 | 0.6199 | 0.468 | 0.000999 |
| TGD | RTM-likeMT | LTB | ITGB1 | 1.6829 | 0.3078 | 0.9953 | 0.000999 |
| TGD | RTM-likeMT | LTB | TNFRSF1A | 1.6829 | 0.283 | 0.983 | 0.000999 |

| | | | | | | | |
|---|---|---|---|---|---|---|---|
| TGD | RTM-likeMT | MIF | CD74 | 1.6335 | 4.0179 | 2.8257 | 0.000999 |
| TGD | RTM-likeMT | MIF | TNFRSF14 | 1.6335 | 0.3616 | 0.9975 | 0.000999 |
| TGD | RTM-likeMT | PTPRC | CD4 | 1.8653 | 0.6983 | 1.2818 | 0.000999 |
| TGD | RTM-likeMT | PTPRC | CD44 | 1.8653 | 1.4501 | 1.6577 | 0.000999 |
| TGD | RTM-likeMT | PTPRC | CXCR4 | 1.8653 | 0.9973 | 1.4313 | 0.000999 |
| TGD | RTM-likeMT | PTPRC | IFNAR1 | 1.8653 | 0.3337 | 1.0995 | 0.000999 |
| TGD | RTM-likeMT | RPS19 | C5AR1 | 3.8902 | 1.271 | 2.5806 | 0.000999 |
| TGD | RTM-likeMT | RPS19 | CD4 | 3.8902 | 0.6983 | 2.2942 | 0.000999 |
| TGD | RTM-likeMT | RPS19 | ICAM1 | 3.8902 | 0.3448 | 2.1175 | 0.000999 |
| TGD | RTM-likeMT | SELL | CD4 | 0.3555 | 0.6983 | 0.5269 | 0.000999 |
| TGD | RTM-likeMT | SPN | SIGLEC1 | 0.5644 | 0.2533 | 0.4088 | 0.000999 |
| TGD | RTM-likeMT | TGFB1 | ENG | 0.7703 | 0.3413 | 0.5558 | 0.000999 |
| TGD | RTM-likeMT | TGFB1 | FCN1 | 0.7703 | 1.1076 | 0.939 | 0.000999 |
| TGD | RTM-likeMT | TGFB1 | ITGB1 | 0.7703 | 0.3078 | 0.5391 | 0.000999 |
| TGD | RTM-likeMT | TGFB1 | TGFBR2 | 0.7703 | 0.3477 | 0.559 | 0.000999 |
| TGD | RTM-likeMT | TGFB1 | TLR2 | 0.7703 | 0.3069 | 0.5386 | 0.000999 |
| TGD | RTM-likeMT | XCL1 | GPR65 | 1.5822 | 0.3775 | 0.9798 | 0.000999 |
| TGD | RTM-likeMT | XCL2 | GPR65 | 0.4864 | 0.3775 | 0.4319 | 0.000999 |
| TGD | RTM-likeMT | HMGB2 | DAB2 | 0.3275 | 0.7068 | 0.5171 | 0.002 |
| TGD | RTM-likeMT | ICAM1 | ITGAM | 0.2915 | 0.4553 | 0.3734 | 0.00699 |

| Supplementary Table 8: Communication results between macrophages and lymphocytes subpopulations, and malignant cells | | | | | | | |
|---|---|---|---|---|---|---|---|
| **RTM-likeMT vs. malignant cells** | | | | | | | |
| celltype_01 | celltype_02 | ligand | receptor | ligand_mean_expression | receptor_mean_expression | ligand_receptor_mean_expression | pvalue |
| RTM-likeMT | breasttumor | ANXA1 | CD44 | 1.1151 | 0.3247 | 0.7199 | 0.000999 |
| RTM-likeMT | breasttumor | APOE | SCARB1 | 3.1194 | 0.2628 | 1.6911 | 0.000999 |
| RTM-likeMT | breasttumor | APOE | SDC2 | 3.1194 | 1.071 | 2.0952 | 0.000999 |
| RTM-likeMT | breasttumor | CALM2 | MYLK | 1.6298 | 0.4064 | 1.0181 | 0.000999 |
| RTM-likeMT | breasttumor | CALM3 | MYLK | 1.0656 | 0.4064 | 0.736 | 0.000999 |
| RTM-likeMT | breasttumor | CD99 | CD99 | 0.8652 | 0.3871 | 0.6261 | 0.000999 |
| RTM-likeMT | breasttumor | COL1A1 | ITGB1 | 0.2832 | 0.2939 | 0.2885 | 0.000999 |
| RTM-likeMT | breasttumor | FN1 | CD44 | 2.2847 | 0.3247 | 1.3047 | 0.000999 |
| RTM-likeMT | breasttumor | FN1 | ITGB1 | 2.2847 | 0.2939 | 1.2893 | 0.000999 |
| RTM-likeMT | breasttumor | FN1 | RPSA | 2.2847 | 2.358 | 2.3214 | 0.000999 |
| RTM-likeMT | breasttumor | FN1 | SDC2 | 2.2847 | 1.071 | 1.6779 | 0.000999 |
| RTM-likeMT | breasttumor | GRN | TNFRSF1A | 1.6831 | 0.2536 | 0.9684 | 0.000999 |
| RTM-likeMT | breasttumor | HBEGF | CD44 | 0.4151 | 0.3247 | 0.3699 | 0.000999 |
| RTM-likeMT | breasttumor | HLA-DRA | SCARB1 | 3.9038 | 0.2628 | 2.0833 | 0.000999 |
| RTM-likeMT | breasttumor | PTPRC | CD44 | 0.5986 | 0.3247 | 0.4617 | 0.000999 |
| RTM-likeMT | breasttumor | RTN4 | CD44 | 0.9153 | 0.3247 | 0.62 | 0.000999 |
| RTM-likeMT | breasttumor | RTN4 | CD99 | 0.9153 | 0.3871 | 0.6512 | 0.000999 |
| RTM-likeMT | breasttumor | SERPINA1 | SDC2 | 1.0071 | 1.071 | 1.0391 | 0.000999 |
| RTM-likeMT | breasttumor | SPP1 | CD44 | 0.7998 | 0.3247 | 0.5622 | 0.000999 |
| RTM-likeMT | breasttumor | SPP1 | ITGB1 | 0.7998 | 0.2939 | 0.5468 | 0.000999 |
| RTM-likeMT | breasttumor | THBS1 | ITGB1 | 0.753 | 0.2939 | 0.5234 | 0.000999 |
| RTM-likeMT | breasttumor | TNFSF13B | ITGB1 | 0.76 | 0.2939 | 0.5269 | 0.000999 |
| RTM-likeMT | colorectaltumor | APOE | SDC1 | 3.1194 | 0.2575 | 1.6884 | 0.000999 |
| RTM-likeMT | colorectaltumor | FN1 | ITGB1 | 2.2847 | 0.4214 | 1.3531 | 0.000999 |
| RTM-likeMT | colorectaltumor | FN1 | RPSA | 2.2847 | 2.251 | 2.2679 | 0.000999 |
| RTM-likeMT | colorectaltumor | FN1 | SDC1 | 2.2847 | 0.2575 | 1.2711 | 0.000999 |
| RTM-likeMT | colorectaltumor | ITGB1 | CD151 | 0.3085 | 0.6439 | 0.4762 | 0.000999 |

| | | | | | | | |
|---|---|---|---|---|---|---|---|
| RTM-likeMT | colorectaltumor | ITGB1 | CDH17 | 0.3085 | 0.254 | 0.2812 | 0.000999 |
| RTM-likeMT | colorectaltumor | ITGB1 | RACK1 | 0.3085 | 2.7146 | 1.5115 | 0.000999 |
| RTM-likeMT | colorectaltumor | ITGB1 | SDC1 | 0.3085 | 0.2575 | 0.283 | 0.000999 |
| RTM-likeMT | colorectaltumor | PECAM1 | SDC1 | 0.343 | 0.2575 | 0.3002 | 0.000999 |
| RTM-likeMT | colorectaltumor | SPP1 | ITGB1 | 0.7998 | 0.4214 | 0.6106 | 0.000999 |
| RTM-likeMT | colorectaltumor | SPP1 | SDC1 | 0.7998 | 0.2575 | 0.5286 | 0.000999 |
| RTM-likeMT | colorectaltumor | THBS1 | CD47 | 0.753 | 0.2608 | 0.5069 | 0.000999 |
| RTM-likeMT | colorectaltumor | THBS1 | ITGB1 | 0.753 | 0.4214 | 0.5872 | 0.000999 |
| RTM-likeMT | colorectaltumor | THBS1 | SDC1 | 0.753 | 0.2575 | 0.5052 | 0.000999 |
| RTM-likeMT | colorectaltumor | TNFSF13B | ITGB1 | 0.76 | 0.4214 | 0.5907 | 0.000999 |
| RTM-likeMT | colorectaltumor | FN1 | CD44 | 2.2847 | 0.2512 | 1.2679 | 0.005 |
| RTM-likeMT | colorectaltumor | CALM1 | RPSA | 0.8292 | 2.251 | 1.5401 | 0.00799 |
| RTM-likeMT | livertumor | A2M | TNFRSF14 | 0.7132 | 0.2788 | 0.496 | 0.000999 |
| RTM-likeMT | livertumor | ADM | RAMP1 | 0.4888 | 0.4499 | 0.4694 | 0.000999 |
| RTM-likeMT | livertumor | ANXA1 | MET | 1.1151 | 0.2983 | 0.7067 | 0.000999 |
| RTM-likeMT | livertumor | ANXA1 | PTPRF | 1.1151 | 0.384 | 0.7496 | 0.000999 |
| RTM-likeMT | livertumor | APOE | SDC1 | 3.1194 | 0.5354 | 1.8274 | 0.000999 |
| RTM-likeMT | livertumor | APOE | SDC4 | 3.1194 | 0.6068 | 1.8631 | 0.000999 |
| RTM-likeMT | livertumor | AREG | ERBB3 | 0.8359 | 0.2644 | 0.5502 | 0.000999 |
| RTM-likeMT | livertumor | CALM1 | INSR | 0.8292 | 0.3039 | 0.5665 | 0.000999 |
| RTM-likeMT | livertumor | CALM1 | RPSA | 0.8292 | 2.6317 | 1.7304 | 0.000999 |
| RTM-likeMT | livertumor | CD99 | EPHB4 | 0.8652 | 0.4535 | 0.6593 | 0.000999 |
| RTM-likeMT | livertumor | CD99 | INSR | 0.8652 | 0.3039 | 0.5846 | 0.000999 |
| RTM-likeMT | livertumor | COL1A1 | ITGA2 | 0.2832 | 0.3983 | 0.3407 | 0.000999 |
| RTM-likeMT | livertumor | COL1A1 | ITGA3 | 0.2832 | 0.3913 | 0.3372 | 0.000999 |
| RTM-likeMT | livertumor | COL1A1 | ITGB1 | 0.2832 | 1.6044 | 0.9438 | 0.000999 |
| RTM-likeMT | livertumor | COL1A1 | SDC1 | 0.2832 | 0.5354 | 0.4093 | 0.000999 |
| RTM-likeMT | livertumor | EREG | ERBB3 | 0.7435 | 0.2644 | 0.5039 | 0.000999 |
| RTM-likeMT | livertumor | FN1 | CD44 | 2.2847 | 0.8016 | 1.5432 | 0.000999 |
| RTM-likeMT | livertumor | FN1 | ITGA2 | 2.2847 | 0.3983 | 1.3415 | 0.000999 |

| | | | | | | | |
|---|---|---|---|---|---|---|---|
| RTM-likeMT | livertumor | FN1 | ITGA3 | 2.2847 | 0.3913 | 1.338 | 0.000999 |
| RTM-likeMT | livertumor | FN1 | ITGB1 | 2.2847 | 1.6044 | 1.9446 | 0.000999 |
| RTM-likeMT | livertumor | FN1 | RPSA | 2.2847 | 2.6317 | 2.4582 | 0.000999 |
| RTM-likeMT | livertumor | FN1 | SDC1 | 2.2847 | 0.5354 | 1.4101 | 0.000999 |
| RTM-likeMT | livertumor | FN1 | TFRC | 2.2847 | 0.2899 | 1.2873 | 0.000999 |
| RTM-likeMT | livertumor | HAMP | SLC40A1 | 0.9966 | 0.4972 | 0.7469 | 0.000999 |
| RTM-likeMT | livertumor | HBEGF | ERBB3 | 0.4151 | 0.2644 | 0.3397 | 0.000999 |
| RTM-likeMT | livertumor | HLA-A | ERBB3 | 2.8219 | 0.2644 | 1.5431 | 0.000999 |
| RTM-likeMT | livertumor | HLA-DRA | EPHB4 | 3.9038 | 0.4535 | 2.1786 | 0.000999 |
| RTM-likeMT | livertumor | HLA-E | EPHB4 | 1.7672 | 0.4535 | 1.1103 | 0.000999 |
| RTM-likeMT | livertumor | HLA-E | ITGA6 | 1.7672 | 0.5211 | 1.1442 | 0.000999 |
| RTM-likeMT | livertumor | HLA-E | PLXNB2 | 1.7672 | 0.265 | 1.0161 | 0.000999 |
| RTM-likeMT | livertumor | HMGB2 | CDH1 | 0.5216 | 0.4896 | 0.5056 | 0.000999 |
| RTM-likeMT | livertumor | HMGB2 | DSG2 | 0.5216 | 0.7479 | 0.6347 | 0.000999 |
| RTM-likeMT | livertumor | HSP90B1 | ITGB1 | 0.5456 | 1.6044 | 1.075 | 0.000999 |
| RTM-likeMT | livertumor | LGALS3 | ITGA2 | 0.8659 | 0.3983 | 0.6321 | 0.000999 |
| RTM-likeMT | livertumor | LGALS3 | ITGB1 | 0.8659 | 1.6044 | 1.2352 | 0.000999 |
| RTM-likeMT | livertumor | LGALS3 | ITGB4 | 0.8659 | 0.5518 | 0.7089 | 0.000999 |
| RTM-likeMT | livertumor | NAMPT | INSR | 1.5328 | 0.3039 | 0.9184 | 0.000999 |
| RTM-likeMT | livertumor | PECAM1 | SDC1 | 0.343 | 0.5354 | 0.4392 | 0.000999 |
| RTM-likeMT | livertumor | PECAM1 | SDC4 | 0.343 | 0.6068 | 0.4749 | 0.000999 |
| RTM-likeMT | livertumor | PTPRC | INSR | 0.5986 | 0.3039 | 0.4513 | 0.000999 |
| RTM-likeMT | livertumor | PTPRC | MET | 0.5986 | 0.2983 | 0.4485 | 0.000999 |
| RTM-likeMT | livertumor | RPS19 | RPSA | 3.537 | 2.6317 | 3.0844 | 0.000999 |
| RTM-likeMT | livertumor | SPP1 | ITGB1 | 0.7998 | 1.6044 | 1.2021 | 0.000999 |
| RTM-likeMT | livertumor | SPP1 | SDC1 | 0.7998 | 0.5354 | 0.6676 | 0.000999 |
| RTM-likeMT | livertumor | THBS1 | CD47 | 0.753 | 0.5289 | 0.6409 | 0.000999 |
| RTM-likeMT | livertumor | THBS1 | ITGA3 | 0.753 | 0.3913 | 0.5721 | 0.000999 |
| RTM-likeMT | livertumor | THBS1 | ITGB1 | 0.753 | 1.6044 | 1.1787 | 0.000999 |
| RTM-likeMT | livertumor | THBS1 | SDC1 | 0.753 | 0.5354 | 0.6442 | 0.000999 |

| | | | | | | | |
|---|---|---|---|---|---|---|---|
| RTM-likeMT | livertumor | TNFSF13B | ITGB1 | 0.76 | 1.6044 | 1.1822 | 0.000999 |
| RTM-likeMT | livertumor | VEGFA | VTN | 0.5132 | 0.7399 | 0.6266 | 0.000999 |
| RTM-likeMT | livertumor | VEGFA | AXL | 0.5132 | 0.3099 | 0.4116 | 0.003 |
| RTM-likeMT | lungtumor | APOE | SDC1 | 3.1194 | 0.2715 | 1.6955 | 0.000999 |
| RTM-likeMT | lungtumor | APOE | SDC4 | 3.1194 | 0.3516 | 1.7355 | 0.000999 |
| RTM-likeMT | lungtumor | CCL20 | CXCR4 | 0.9187 | 0.2684 | 0.5936 | 0.000999 |
| RTM-likeMT | lungtumor | CD99 | CD99 | 0.8652 | 0.5938 | 0.7295 | 0.000999 |
| RTM-likeMT | lungtumor | COL1A1 | SDC1 | 0.2832 | 0.2715 | 0.2773 | 0.000999 |
| RTM-likeMT | lungtumor | CXCL16 | CXCR4 | 1.0903 | 0.2684 | 0.6793 | 0.000999 |
| RTM-likeMT | lungtumor | CXCL2 | CXCR4 | 1.9307 | 0.2684 | 1.0995 | 0.000999 |
| RTM-likeMT | lungtumor | CXCL3 | CXCR4 | 1.7093 | 0.2684 | 0.9889 | 0.000999 |
| RTM-likeMT | lungtumor | FN1 | CD44 | 2.2847 | 0.3802 | 1.3325 | 0.000999 |
| RTM-likeMT | lungtumor | FN1 | RPSA | 2.2847 | 1.6057 | 1.9452 | 0.000999 |
| RTM-likeMT | lungtumor | FN1 | SDC1 | 2.2847 | 0.2715 | 1.2781 | 0.000999 |
| RTM-likeMT | lungtumor | HBEGF | CD44 | 0.4151 | 0.3802 | 0.3976 | 0.000999 |
| RTM-likeMT | lungtumor | PECAM1 | CXCR4 | 0.343 | 0.2684 | 0.3057 | 0.000999 |
| RTM-likeMT | lungtumor | PECAM1 | SDC1 | 0.343 | 0.2715 | 0.3073 | 0.000999 |
| RTM-likeMT | lungtumor | PECAM1 | SDC4 | 0.343 | 0.3516 | 0.3473 | 0.000999 |
| RTM-likeMT | lungtumor | PTPRC | CD44 | 0.5986 | 0.3802 | 0.4894 | 0.000999 |
| RTM-likeMT | lungtumor | PTPRC | CXCR4 | 0.5986 | 0.2684 | 0.4335 | 0.000999 |
| RTM-likeMT | lungtumor | RTN4 | CD44 | 0.9153 | 0.3802 | 0.6478 | 0.000999 |
| RTM-likeMT | lungtumor | RTN4 | CD99 | 0.9153 | 0.5938 | 0.7546 | 0.000999 |
| RTM-likeMT | lungtumor | THBS1 | CD47 | 0.753 | 0.3255 | 0.5392 | 0.000999 |
| RTM-likeMT | lungtumor | THBS1 | SDC1 | 0.753 | 0.2715 | 0.5122 | 0.000999 |
| RTM-likeMT | lungtumor | ITGB1 | CD151 | 0.3085 | 0.6776 | 0.493 | 0.003 |
| RTM-likeMT | lungtumor | COL1A1 | CD44 | 0.2832 | 0.3802 | 0.3317 | 0.00599 |
| RTM-likeMT | lungtumor | ITGB1 | SDC1 | 0.3085 | 0.2715 | 0.29 | 0.00599 |
| RTM-likeMT | melanomatumo | ANXA1 | EDNRB | 1.1151 | 0.5666 | 0.8409 | 0.000999 |
| RTM-likeMT | melanomatumo | CALM1 | RPSA | 0.8292 | 3.7787 | 2.3039 | 0.000999 |
| RTM-likeMT | melanomatumo | FN1 | RPSA | 2.2847 | 3.7787 | 3.0317 | 0.000999 |

| | | | | | | | |
|---|---|---|---|---|---|---|---|
| RTM-likeMT | melanomatumo | ITGB1 | RACK1 | 0.3085 | 4.145 | 2.2268 | 0.000999 |
| RTM-likeMT | melanomatumo | NAMPT | ANXA6 | 1.5328 | 0.3625 | 0.9476 | 0.000999 |
| RTM-likeMT | melanomatumo | RPS19 | RPSA | 3.537 | 3.7787 | 3.6578 | 0.000999 |
| RTM-likeMT | melanomatumo | FN1 | ITGB1 | 2.2847 | 0.3547 | 1.3197 | 0.005 |
| RTM-likeMT | ovarytumor | AREG | FOLR1 | 0.8359 | 0.5619 | 0.6989 | 0.000999 |
| RTM-likeMT | ovarytumor | CCL20 | CXCR4 | 0.9187 | 0.3688 | 0.6438 | 0.000999 |
| RTM-likeMT | ovarytumor | CXCL16 | CXCR4 | 1.0903 | 0.3688 | 0.7295 | 0.000999 |
| RTM-likeMT | ovarytumor | CXCL2 | CXCR4 | 1.9307 | 0.3688 | 1.1497 | 0.000999 |
| RTM-likeMT | ovarytumor | CXCL3 | CXCR4 | 1.7093 | 0.3688 | 1.0391 | 0.000999 |
| RTM-likeMT | ovarytumor | FN1 | ITGB1 | 2.2847 | 0.2783 | 1.2815 | 0.000999 |
| RTM-likeMT | ovarytumor | FN1 | RPSA | 2.2847 | 2.9542 | 2.6195 | 0.000999 |
| RTM-likeMT | ovarytumor | LGALS3 | ITGB1 | 0.8659 | 0.2783 | 0.5721 | 0.000999 |
| RTM-likeMT | ovarytumor | PECAM1 | CXCR4 | 0.343 | 0.3688 | 0.3559 | 0.000999 |
| RTM-likeMT | ovarytumor | PTPRC | CXCR4 | 0.5986 | 0.3688 | 0.4837 | 0.000999 |
| RTM-likeMT | ovarytumor | SPP1 | ITGB1 | 0.7998 | 0.2783 | 0.539 | 0.000999 |
| RTM-likeMT | ovarytumor | THBS1 | CD47 | 0.753 | 0.4931 | 0.623 | 0.000999 |
| RTM-likeMT | ovarytumor | THBS1 | ITGB1 | 0.753 | 0.2783 | 0.5156 | 0.000999 |
| RTM-likeMT | ovarytumor | TNFSF13B | ITGB1 | 0.76 | 0.2783 | 0.5191 | 0.000999 |
| breasttumor | RTM-likeMT | A2M | LRP1 | 0.3129 | 0.6166 | 0.4648 | 0.000999 |
| breasttumor | RTM-likeMT | A2M | TNFRSF14 | 0.3129 | 0.3624 | 0.3377 | 0.000999 |
| breasttumor | RTM-likeMT | ANXA1 | C3AR1 | 0.4092 | 0.4896 | 0.4494 | 0.000999 |
| breasttumor | RTM-likeMT | ANXA1 | C5AR1 | 0.4092 | 1.2739 | 0.8416 | 0.000999 |
| breasttumor | RTM-likeMT | ANXA1 | CCR1 | 0.4092 | 0.2784 | 0.3438 | 0.000999 |
| breasttumor | RTM-likeMT | ANXA1 | CD4 | 0.4092 | 0.6999 | 0.5545 | 0.000999 |
| breasttumor | RTM-likeMT | ANXA1 | CD44 | 0.4092 | 1.4501 | 0.9296 | 0.000999 |
| breasttumor | RTM-likeMT | ANXA1 | CXCR4 | 0.4092 | 0.994 | 0.7016 | 0.000999 |
| breasttumor | RTM-likeMT | ANXA1 | FPR1 | 0.4092 | 0.4978 | 0.4535 | 0.000999 |
| breasttumor | RTM-likeMT | ANXA1 | FPR3 | 0.4092 | 0.4004 | 0.4048 | 0.000999 |
| breasttumor | RTM-likeMT | APP | ALOX5 | 0.5409 | 0.4201 | 0.4805 | 0.000999 |
| breasttumor | RTM-likeMT | APP | C3AR1 | 0.5409 | 0.4896 | 0.5153 | 0.000999 |

| | | | | | | | |
|---|---|---|---|---|---|---|---|
| breasttumor | RTM-likeMT | APP | C5AR1 | 0.5409 | 1.2739 | 0.9074 | 0.000999 |
| breasttumor | RTM-likeMT | APP | CCR1 | 0.5409 | 0.2784 | 0.4097 | 0.000999 |
| breasttumor | RTM-likeMT | APP | CXCR4 | 0.5409 | 0.994 | 0.7675 | 0.000999 |
| breasttumor | RTM-likeMT | APP | FPR1 | 0.5409 | 0.4978 | 0.5194 | 0.000999 |
| breasttumor | RTM-likeMT | APP | LRP1 | 0.5409 | 0.6166 | 0.5788 | 0.000999 |
| breasttumor | RTM-likeMT | APP | SORL1 | 0.5409 | 0.303 | 0.422 | 0.000999 |
| breasttumor | RTM-likeMT | APP | TGFBR2 | 0.5409 | 0.3485 | 0.4447 | 0.000999 |
| breasttumor | RTM-likeMT | APP | TNFRSF14 | 0.5409 | 0.3624 | 0.4517 | 0.000999 |
| breasttumor | RTM-likeMT | ARF1 | CD4 | 1.6219 | 0.6999 | 1.1609 | 0.000999 |
| breasttumor | RTM-likeMT | ARF1 | CD44 | 1.6219 | 1.4501 | 1.536 | 0.000999 |
| breasttumor | RTM-likeMT | CALM1 | TNFRSF1B | 0.9015 | 0.7592 | 0.8304 | 0.000999 |
| breasttumor | RTM-likeMT | CALR | CD44 | 1.1835 | 1.4501 | 1.3168 | 0.000999 |
| breasttumor | RTM-likeMT | CALR | LRP1 | 1.1835 | 0.6166 | 0.9001 | 0.000999 |
| breasttumor | RTM-likeMT | CD99 | CD99 | 0.3871 | 0.8652 | 0.6261 | 0.000999 |
| breasttumor | RTM-likeMT | COL2A1 | CD44 | 0.3935 | 1.4501 | 0.9218 | 0.000999 |
| breasttumor | RTM-likeMT | GSTP1 | CD44 | 2.5435 | 1.4501 | 1.9968 | 0.000999 |
| breasttumor | RTM-likeMT | HEBP1 | C3AR1 | 0.4062 | 0.4896 | 0.4479 | 0.000999 |
| breasttumor | RTM-likeMT | HEBP1 | C5AR1 | 0.4062 | 1.2739 | 0.8401 | 0.000999 |
| breasttumor | RTM-likeMT | HEBP1 | CCR1 | 0.4062 | 0.2784 | 0.3423 | 0.000999 |
| breasttumor | RTM-likeMT | HEBP1 | CXCR4 | 0.4062 | 0.994 | 0.7001 | 0.000999 |
| breasttumor | RTM-likeMT | HEBP1 | FPR1 | 0.4062 | 0.4978 | 0.452 | 0.000999 |
| breasttumor | RTM-likeMT | HEBP1 | FPR3 | 0.4062 | 0.4004 | 0.4033 | 0.000999 |
| breasttumor | RTM-likeMT | HMGB1 | AGTRAP | 1.1815 | 0.3972 | 0.7893 | 0.000999 |
| breasttumor | RTM-likeMT | HMGB1 | LY96 | 1.1815 | 1.4353 | 1.3084 | 0.000999 |
| breasttumor | RTM-likeMT | HMGB1 | TLR2 | 1.1815 | 0.3076 | 0.7445 | 0.000999 |
| breasttumor | RTM-likeMT | HMGB2 | CD44 | 0.3139 | 1.4501 | 0.882 | 0.000999 |
| breasttumor | RTM-likeMT | HMGB2 | DAB2 | 0.3139 | 0.7084 | 0.5112 | 0.000999 |
| breasttumor | RTM-likeMT | HSP90B1 | LRP1 | 0.7786 | 0.6166 | 0.6976 | 0.000999 |
| breasttumor | RTM-likeMT | HSP90B1 | TNFRSF1B | 0.7786 | 0.7592 | 0.7689 | 0.000999 |
| breasttumor | RTM-likeMT | ITGB1 | CD46 | 0.2939 | 0.3148 | 0.3043 | 0.000999 |

| | | | | | | | |
|---|---|---|---|---|---|---|---|
| breasttumor | RTM-likeMT | ITGB1 | CD47 | 0.2939 | 0.3748 | 0.3343 | 0.000999 |
| breasttumor | RTM-likeMT | ITGB1 | DAB2 | 0.2939 | 0.7084 | 0.5012 | 0.000999 |
| breasttumor | RTM-likeMT | ITGB1 | ENG | 0.2939 | 0.3421 | 0.318 | 0.000999 |
| breasttumor | RTM-likeMT | ITGB1 | PLAUR | 0.2939 | 1.2007 | 0.7473 | 0.000999 |
| breasttumor | RTM-likeMT | ITGB1 | PTK2B | 0.2939 | 0.3079 | 0.3009 | 0.000999 |
| breasttumor | RTM-likeMT | LGALS3 | CLEC7A | 0.8391 | 0.7777 | 0.8084 | 0.000999 |
| breasttumor | RTM-likeMT | LGALS3 | FCGR2A | 0.8391 | 1.0268 | 0.933 | 0.000999 |
| breasttumor | RTM-likeMT | LGALS3 | FCGR2B | 0.8391 | 0.8966 | 0.8679 | 0.000999 |
| breasttumor | RTM-likeMT | LGALS3 | TFRC | 0.8391 | 0.9108 | 0.8749 | 0.000999 |
| breasttumor | RTM-likeMT | LGALS3 | TGFBR2 | 0.8391 | 0.3485 | 0.5938 | 0.000999 |
| breasttumor | RTM-likeMT | LRPAP1 | LRP1 | 0.2756 | 0.6166 | 0.4461 | 0.000999 |
| breasttumor | RTM-likeMT | LRPAP1 | SORL1 | 0.2756 | 0.303 | 0.2893 | 0.000999 |
| breasttumor | RTM-likeMT | MDK | LRP1 | 0.592 | 0.6166 | 0.6043 | 0.000999 |
| breasttumor | RTM-likeMT | MDK | NOTCH2 | 0.592 | 0.3059 | 0.4489 | 0.000999 |
| breasttumor | RTM-likeMT | MIF | CD74 | 3.2716 | 4.0176 | 3.6446 | 0.000999 |
| breasttumor | RTM-likeMT | MIF | TNFRSF14 | 3.2716 | 0.3624 | 1.817 | 0.000999 |
| breasttumor | RTM-likeMT | NPW | C3AR1 | 0.5478 | 0.4896 | 0.5187 | 0.000999 |
| breasttumor | RTM-likeMT | NPW | C5AR1 | 0.5478 | 1.2739 | 0.9109 | 0.000999 |
| breasttumor | RTM-likeMT | NPW | CCR1 | 0.5478 | 0.2784 | 0.4131 | 0.000999 |
| breasttumor | RTM-likeMT | NPW | CXCR4 | 0.5478 | 0.994 | 0.7709 | 0.000999 |
| breasttumor | RTM-likeMT | NPW | FPR1 | 0.5478 | 0.4978 | 0.5228 | 0.000999 |
| breasttumor | RTM-likeMT | RPS19 | C5AR1 | 3.691 | 1.2739 | 2.4825 | 0.000999 |
| breasttumor | RTM-likeMT | RPS19 | CD4 | 3.691 | 0.6999 | 2.1955 | 0.000999 |
| breasttumor | RTM-likeMT | RPS19 | ICAM1 | 3.691 | 0.3422 | 2.0166 | 0.000999 |
| breasttumor | RTM-likeMT | RTN4 | CD44 | 0.3318 | 1.4501 | 0.8909 | 0.000999 |
| breasttumor | RTM-likeMT | RTN4 | CD99 | 0.3318 | 0.8652 | 0.5985 | 0.000999 |
| breasttumor | RTM-likeMT | SERPINE2 | LRP1 | 0.6001 | 0.6166 | 0.6084 | 0.000999 |
| breasttumor | RTM-likeMT | TF | CD4 | 0.2688 | 0.6999 | 0.4843 | 0.000999 |
| breasttumor | RTM-likeMT | TF | TFRC | 0.2688 | 0.9108 | 0.5898 | 0.000999 |
| breasttumor | RTM-likeMT | COL2A1 | ITGB1 | 0.3935 | 0.3085 | 0.351 | 0.003 |

| | | | | | | | |
|---|---|---|---|---|---|---|---|
| colorectaltumor | RTM-likeMT | ANXA1 | C5AR1 | 0.5916 | 1.2739 | 0.9328 | 0.000999 |
| colorectaltumor | RTM-likeMT | ANXA1 | CD44 | 0.5916 | 1.4501 | 1.0208 | 0.000999 |
| colorectaltumor | RTM-likeMT | APP | ALOX5 | 0.4966 | 0.4201 | 0.4584 | 0.000999 |
| colorectaltumor | RTM-likeMT | APP | C3AR1 | 0.4966 | 0.4896 | 0.4931 | 0.000999 |
| colorectaltumor | RTM-likeMT | APP | C5AR1 | 0.4966 | 1.2739 | 0.8853 | 0.000999 |
| colorectaltumor | RTM-likeMT | APP | CCR1 | 0.4966 | 0.2784 | 0.3875 | 0.000999 |
| colorectaltumor | RTM-likeMT | APP | CXCR4 | 0.4966 | 0.994 | 0.7453 | 0.000999 |
| colorectaltumor | RTM-likeMT | APP | FPR1 | 0.4966 | 0.4978 | 0.4972 | 0.000999 |
| colorectaltumor | RTM-likeMT | APP | LRP1 | 0.4966 | 0.6166 | 0.5566 | 0.000999 |
| colorectaltumor | RTM-likeMT | APP | SORL1 | 0.4966 | 0.303 | 0.3998 | 0.000999 |
| colorectaltumor | RTM-likeMT | APP | TGFBR2 | 0.4966 | 0.3485 | 0.4226 | 0.000999 |
| colorectaltumor | RTM-likeMT | APP | TNFRSF14 | 0.4966 | 0.3624 | 0.4295 | 0.000999 |
| colorectaltumor | RTM-likeMT | ARF1 | CD4 | 0.6489 | 0.6999 | 0.6744 | 0.000999 |
| colorectaltumor | RTM-likeMT | ARF1 | CD44 | 0.6489 | 1.4501 | 1.0495 | 0.000999 |
| colorectaltumor | RTM-likeMT | CALM1 | TNFRSF1A | 1.1676 | 0.2803 | 0.7239 | 0.000999 |
| colorectaltumor | RTM-likeMT | CALM1 | TNFRSF1B | 1.1676 | 0.7592 | 0.9634 | 0.000999 |
| colorectaltumor | RTM-likeMT | CALR | CD44 | 0.4934 | 1.4501 | 0.9717 | 0.000999 |
| colorectaltumor | RTM-likeMT | COL1A1 | CD44 | 0.4875 | 1.4501 | 0.9688 | 0.000999 |
| colorectaltumor | RTM-likeMT | GSTP1 | CD44 | 2.8039 | 1.4501 | 2.127 | 0.000999 |
| colorectaltumor | RTM-likeMT | HMGB1 | AGTRAP | 2.155 | 0.3972 | 1.2761 | 0.000999 |
| colorectaltumor | RTM-likeMT | HMGB1 | LY96 | 2.155 | 1.4353 | 1.7951 | 0.000999 |
| colorectaltumor | RTM-likeMT | HMGB1 | TLR2 | 2.155 | 0.3076 | 1.2313 | 0.000999 |
| colorectaltumor | RTM-likeMT | HMGB2 | CD44 | 0.5347 | 1.4501 | 0.9924 | 0.000999 |
| colorectaltumor | RTM-likeMT | HMGB2 | DAB2 | 0.5347 | 0.7084 | 0.6216 | 0.000999 |
| colorectaltumor | RTM-likeMT | HSP90B1 | LRP1 | 0.5887 | 0.6166 | 0.6027 | 0.000999 |
| colorectaltumor | RTM-likeMT | HSP90B1 | TNFRSF1B | 0.5887 | 0.7592 | 0.674 | 0.000999 |
| colorectaltumor | RTM-likeMT | ITGB1 | DAB2 | 0.4214 | 0.7084 | 0.5649 | 0.000999 |
| colorectaltumor | RTM-likeMT | ITGB1 | ENG | 0.4214 | 0.3421 | 0.3818 | 0.000999 |
| colorectaltumor | RTM-likeMT | ITGB1 | PLAUR | 0.4214 | 1.2007 | 0.811 | 0.000999 |
| colorectaltumor | RTM-likeMT | ITGB1 | PTK2B | 0.4214 | 0.3079 | 0.3647 | 0.000999 |

| | | | | | | | |
|---|---|---|---|---|---|---|---|
| colorectaltumor | RTM-likeMT | LGALS3 | CLEC7A | 1.5563 | 0.7777 | 1.167 | 0.000999 |
| colorectaltumor | RTM-likeMT | LGALS3 | FCGR2A | 1.5563 | 1.0268 | 1.2915 | 0.000999 |
| colorectaltumor | RTM-likeMT | LGALS3 | FCGR2B | 1.5563 | 0.8966 | 1.2264 | 0.000999 |
| colorectaltumor | RTM-likeMT | LGALS3 | ITGB1 | 1.5563 | 0.3085 | 0.9324 | 0.000999 |
| colorectaltumor | RTM-likeMT | LGALS3 | TFRC | 1.5563 | 0.9108 | 1.2335 | 0.000999 |
| colorectaltumor | RTM-likeMT | LGALS3 | TGFBR2 | 1.5563 | 0.3485 | 0.9524 | 0.000999 |
| colorectaltumor | RTM-likeMT | MDK | LRP1 | 0.7548 | 0.6166 | 0.6857 | 0.000999 |
| colorectaltumor | RTM-likeMT | MDK | NOTCH2 | 0.7548 | 0.3059 | 0.5304 | 0.000999 |
| colorectaltumor | RTM-likeMT | MIF | CD74 | 1.6741 | 4.0176 | 2.8458 | 0.000999 |
| colorectaltumor | RTM-likeMT | MIF | TNFRSF14 | 1.6741 | 0.3624 | 1.0182 | 0.000999 |
| colorectaltumor | RTM-likeMT | RPS19 | C5AR1 | 4.0956 | 1.2739 | 2.6848 | 0.000999 |
| colorectaltumor | RTM-likeMT | RPS19 | CD4 | 4.0956 | 0.6999 | 2.3977 | 0.000999 |
| colorectaltumor | RTM-likeMT | RPS19 | ICAM1 | 4.0956 | 0.3422 | 2.2189 | 0.000999 |
| colorectaltumor | RTM-likeMT | RPS19 | RPSA | 4.0956 | 1.7816 | 2.9386 | 0.000999 |
| colorectaltumor | RTM-likeMT | RTN4 | CD44 | 0.4148 | 1.4501 | 0.9325 | 0.000999 |
| colorectaltumor | RTM-likeMT | LRPAP1 | LRP1 | 0.2533 | 0.6166 | 0.435 | 0.002 |
| colorectaltumor | RTM-likeMT | ITGB1 | CD47 | 0.4214 | 0.3748 | 0.3981 | 0.003 |
| livertumor | RTM-likeMT | AGRN | LRP1 | 0.3492 | 0.6166 | 0.4829 | 0.000999 |
| livertumor | RTM-likeMT | AGT | ATP6AP2 | 0.4701 | 0.6631 | 0.5666 | 0.000999 |
| livertumor | RTM-likeMT | AGT | C3AR1 | 0.4701 | 0.4896 | 0.4798 | 0.000999 |
| livertumor | RTM-likeMT | AGT | C5AR1 | 0.4701 | 1.2739 | 0.872 | 0.000999 |
| livertumor | RTM-likeMT | AGT | CCR1 | 0.4701 | 0.2784 | 0.3742 | 0.000999 |
| livertumor | RTM-likeMT | AGT | CXCR4 | 0.4701 | 0.994 | 0.732 | 0.000999 |
| livertumor | RTM-likeMT | AGT | FPR1 | 0.4701 | 0.4978 | 0.4839 | 0.000999 |
| livertumor | RTM-likeMT | AGT | GPR65 | 0.4701 | 0.3783 | 0.4242 | 0.000999 |
| livertumor | RTM-likeMT | ANXA1 | C5AR1 | 0.3041 | 1.2739 | 0.789 | 0.000999 |
| livertumor | RTM-likeMT | APOA1 | ABCA1 | 0.2888 | 0.3999 | 0.3443 | 0.000999 |
| livertumor | RTM-likeMT | APOA1 | LRP1 | 0.2888 | 0.6166 | 0.4527 | 0.000999 |
| livertumor | RTM-likeMT | APOA1 | MSR1 | 0.2888 | 0.6732 | 0.481 | 0.000999 |
| livertumor | RTM-likeMT | APP | ALOX5 | 0.5626 | 0.4201 | 0.4913 | 0.000999 |

| | | | | | | | |
|---|---|---|---|---|---|---|---|
| livertumor | RTM-likeMT | APP | C3AR1 | 0.5626 | 0.4896 | 0.5261 | 0.000999 |
| livertumor | RTM-likeMT | APP | C5AR1 | 0.5626 | 1.2739 | 0.9182 | 0.000999 |
| livertumor | RTM-likeMT | APP | CCR1 | 0.5626 | 0.2784 | 0.4205 | 0.000999 |
| livertumor | RTM-likeMT | APP | CXCR4 | 0.5626 | 0.994 | 0.7783 | 0.000999 |
| livertumor | RTM-likeMT | APP | FPR1 | 0.5626 | 0.4978 | 0.5302 | 0.000999 |
| livertumor | RTM-likeMT | APP | LRP1 | 0.5626 | 0.6166 | 0.5896 | 0.000999 |
| livertumor | RTM-likeMT | APP | SORL1 | 0.5626 | 0.303 | 0.4328 | 0.000999 |
| livertumor | RTM-likeMT | APP | TGFBR2 | 0.5626 | 0.3485 | 0.4555 | 0.000999 |
| livertumor | RTM-likeMT | APP | TNFRSF14 | 0.5626 | 0.3624 | 0.4625 | 0.000999 |
| livertumor | RTM-likeMT | ARF1 | CD4 | 0.6936 | 0.6999 | 0.6967 | 0.000999 |
| livertumor | RTM-likeMT | ARF1 | CD44 | 0.6936 | 1.4501 | 1.0718 | 0.000999 |
| livertumor | RTM-likeMT | C3 | C3AR1 | 0.6584 | 0.4896 | 0.574 | 0.000999 |
| livertumor | RTM-likeMT | C3 | C5AR1 | 0.6584 | 1.2739 | 0.9662 | 0.000999 |
| livertumor | RTM-likeMT | C3 | C5AR2 | 0.6584 | 0.2839 | 0.4712 | 0.000999 |
| livertumor | RTM-likeMT | C3 | CCR1 | 0.6584 | 0.2784 | 0.4684 | 0.000999 |
| livertumor | RTM-likeMT | C3 | CXCR4 | 0.6584 | 0.994 | 0.8262 | 0.000999 |
| livertumor | RTM-likeMT | C3 | FPR1 | 0.6584 | 0.4978 | 0.5781 | 0.000999 |
| livertumor | RTM-likeMT | C3 | ITGAM | 0.6584 | 0.4529 | 0.5557 | 0.000999 |
| livertumor | RTM-likeMT | C3 | ITGB2 | 0.6584 | 1.4753 | 1.0669 | 0.000999 |
| livertumor | RTM-likeMT | C3 | LRP1 | 0.6584 | 0.6166 | 0.6375 | 0.000999 |
| livertumor | RTM-likeMT | CALM1 | TNFRSF1B | 0.6596 | 0.7592 | 0.7094 | 0.000999 |
| livertumor | RTM-likeMT | CALR | CD44 | 1.0416 | 1.4501 | 1.2459 | 0.000999 |
| livertumor | RTM-likeMT | CALR | LRP1 | 1.0416 | 0.6166 | 0.8291 | 0.000999 |
| livertumor | RTM-likeMT | CCL20 | C3AR1 | 1.0877 | 0.4896 | 0.7886 | 0.000999 |
| livertumor | RTM-likeMT | CCL20 | C5AR1 | 1.0877 | 1.2739 | 1.1808 | 0.000999 |
| livertumor | RTM-likeMT | CCL20 | CCR1 | 1.0877 | 0.2784 | 0.683 | 0.000999 |
| livertumor | RTM-likeMT | CCL20 | CXCR4 | 1.0877 | 0.994 | 1.0408 | 0.000999 |
| livertumor | RTM-likeMT | CCL20 | FPR1 | 1.0877 | 0.4978 | 0.7927 | 0.000999 |
| livertumor | RTM-likeMT | CFH | ITGAM | 0.3373 | 0.4529 | 0.3951 | 0.000999 |
| livertumor | RTM-likeMT | DSG2 | CD44 | 0.7479 | 1.4501 | 1.099 | 0.000999 |

| | | | | | | | |
|---|---|---|---|---|---|---|---|
| livertumor | RTM-likeMT | DSG2 | CXCR4 | 0.7479 | 0.994 | 0.871 | 0.000999 |
| livertumor | RTM-likeMT | DSG2 | DAB2 | 0.7479 | 0.7084 | 0.7282 | 0.000999 |
| livertumor | RTM-likeMT | FGA | ITGB2 | 0.2688 | 1.4753 | 0.8721 | 0.000999 |
| livertumor | RTM-likeMT | GSTP1 | CD44 | 1.7219 | 1.4501 | 1.586 | 0.000999 |
| livertumor | RTM-likeMT | HMGB1 | AGTRAP | 1.4361 | 0.3972 | 0.9166 | 0.000999 |
| livertumor | RTM-likeMT | HMGB1 | LY96 | 1.4361 | 1.4353 | 1.4357 | 0.000999 |
| livertumor | RTM-likeMT | HMGB1 | TLR2 | 1.4361 | 0.3076 | 0.8718 | 0.000999 |
| livertumor | RTM-likeMT | HSP90B1 | LRP1 | 0.8951 | 0.6166 | 0.7559 | 0.000999 |
| livertumor | RTM-likeMT | HSP90B1 | TNFRSF1B | 0.8951 | 0.7592 | 0.8272 | 0.000999 |
| livertumor | RTM-likeMT | HSPG2 | LRP1 | 0.2746 | 0.6166 | 0.4456 | 0.000999 |
| livertumor | RTM-likeMT | ITGB1 | CD46 | 1.6044 | 0.3148 | 0.9596 | 0.000999 |
| livertumor | RTM-likeMT | ITGB1 | CD47 | 1.6044 | 0.3748 | 0.9896 | 0.000999 |
| livertumor | RTM-likeMT | ITGB1 | DAB2 | 1.6044 | 0.7084 | 1.1564 | 0.000999 |
| livertumor | RTM-likeMT | ITGB1 | ENG | 1.6044 | 0.3421 | 0.9733 | 0.000999 |
| livertumor | RTM-likeMT | ITGB1 | PLAUR | 1.6044 | 1.2007 | 1.4026 | 0.000999 |
| livertumor | RTM-likeMT | ITGB1 | PTK2B | 1.6044 | 0.3079 | 0.9562 | 0.000999 |
| livertumor | RTM-likeMT | ITGB1 | RACK1 | 1.6044 | 2.3206 | 1.9625 | 0.000999 |
| livertumor | RTM-likeMT | LGALS3 | CLEC7A | 0.8172 | 0.7777 | 0.7975 | 0.000999 |
| livertumor | RTM-likeMT | LGALS3 | FCGR2A | 0.8172 | 1.0268 | 0.922 | 0.000999 |
| livertumor | RTM-likeMT | LGALS3 | FCGR2B | 0.8172 | 0.8966 | 0.8569 | 0.000999 |
| livertumor | RTM-likeMT | LGALS3 | TFRC | 0.8172 | 0.9108 | 0.864 | 0.000999 |
| livertumor | RTM-likeMT | LRPAP1 | LRP1 | 0.273 | 0.6166 | 0.4448 | 0.000999 |
| livertumor | RTM-likeMT | MDK | LRP1 | 1.6101 | 0.6166 | 1.1134 | 0.000999 |
| livertumor | RTM-likeMT | MDK | NOTCH2 | 1.6101 | 0.3059 | 0.958 | 0.000999 |
| livertumor | RTM-likeMT | MIF | CD74 | 1.7638 | 4.0176 | 2.8907 | 0.000999 |
| livertumor | RTM-likeMT | MIF | TNFRSF14 | 1.7638 | 0.3624 | 1.0631 | 0.000999 |
| livertumor | RTM-likeMT | NTS | GPR65 | 0.3425 | 0.3783 | 0.3604 | 0.000999 |
| livertumor | RTM-likeMT | OCLN | TGFBR2 | 0.3091 | 0.3485 | 0.3288 | 0.000999 |
| livertumor | RTM-likeMT | PTPRF | TGFBR2 | 0.384 | 0.3485 | 0.3663 | 0.000999 |
| livertumor | RTM-likeMT | RPS19 | C5AR1 | 4.0772 | 1.2739 | 2.6756 | 0.000999 |

| | | | | | | | |
|---|---|---|---|---|---|---|---|
| livertumor | RTM-likeMT | RPS19 | CD4 | 4.0772 | 0.6999 | 2.3886 | 0.000999 |
| livertumor | RTM-likeMT | RPS19 | ICAM1 | 4.0772 | 0.3422 | 2.2097 | 0.000999 |
| livertumor | RTM-likeMT | RTN4 | CD44 | 1.0046 | 1.4501 | 1.2273 | 0.000999 |
| livertumor | RTM-likeMT | RTN4 | CD99 | 1.0046 | 0.8652 | 0.9349 | 0.000999 |
| livertumor | RTM-likeMT | SERPINA1 | LRP1 | 1.604 | 0.6166 | 1.1103 | 0.000999 |
| livertumor | RTM-likeMT | SERPING1 | LRP1 | 0.3498 | 0.6166 | 0.4832 | 0.000999 |
| livertumor | RTM-likeMT | SPP1 | CD44 | 1.0048 | 1.4501 | 1.2274 | 0.000999 |
| livertumor | RTM-likeMT | TF | CD4 | 0.523 | 0.6999 | 0.6115 | 0.000999 |
| livertumor | RTM-likeMT | TF | TFRC | 0.523 | 0.9108 | 0.7169 | 0.000999 |
| livertumor | RTM-likeMT | TFPI | LRP1 | 0.5043 | 0.6166 | 0.5604 | 0.000999 |
| livertumor | RTM-likeMT | VEGFA | LYVE1 | 0.5485 | 0.7793 | 0.6639 | 0.000999 |
| livertumor | RTM-likeMT | VEGFA | PTK2B | 0.5485 | 0.3079 | 0.4282 | 0.000999 |
| livertumor | RTM-likeMT | VTN | PLAUR | 0.7399 | 1.2007 | 0.9703 | 0.000999 |
| lungtumor | RTM-likeMT | ANXA1 | C3AR1 | 1.4026 | 0.4896 | 0.9461 | 0.000999 |
| lungtumor | RTM-likeMT | ANXA1 | C5AR1 | 1.4026 | 1.2739 | 1.3383 | 0.000999 |
| lungtumor | RTM-likeMT | ANXA1 | CCR1 | 1.4026 | 0.2784 | 0.8405 | 0.000999 |
| lungtumor | RTM-likeMT | ANXA1 | CD4 | 1.4026 | 0.6999 | 1.0513 | 0.000999 |
| lungtumor | RTM-likeMT | ANXA1 | CD44 | 1.4026 | 1.4501 | 1.4264 | 0.000999 |
| lungtumor | RTM-likeMT | ANXA1 | CXCR4 | 1.4026 | 0.994 | 1.1983 | 0.000999 |
| lungtumor | RTM-likeMT | ANXA1 | FPR1 | 1.4026 | 0.4978 | 0.9502 | 0.000999 |
| lungtumor | RTM-likeMT | ANXA1 | FPR3 | 1.4026 | 0.4004 | 0.9015 | 0.000999 |
| lungtumor | RTM-likeMT | AREG | ICAM1 | 0.3537 | 0.3422 | 0.348 | 0.000999 |
| lungtumor | RTM-likeMT | ARF1 | CD4 | 0.8791 | 0.6999 | 0.7895 | 0.000999 |
| lungtumor | RTM-likeMT | ARF1 | CD44 | 0.8791 | 1.4501 | 1.1646 | 0.000999 |
| lungtumor | RTM-likeMT | CALM1 | TNFRSF1B | 0.8426 | 0.7592 | 0.8009 | 0.000999 |
| lungtumor | RTM-likeMT | CALR | CD44 | 0.7795 | 1.4501 | 1.1148 | 0.000999 |
| lungtumor | RTM-likeMT | CALR | LRP1 | 0.7795 | 0.6166 | 0.6981 | 0.000999 |
| lungtumor | RTM-likeMT | CCL20 | C3AR1 | 0.2885 | 0.4896 | 0.389 | 0.000999 |
| lungtumor | RTM-likeMT | CCL20 | C5AR1 | 0.2885 | 1.2739 | 0.7812 | 0.000999 |
| lungtumor | RTM-likeMT | CCL20 | CCR1 | 0.2885 | 0.2784 | 0.2835 | 0.000999 |

| | | | | | | | |
|---|---|---|---|---|---|---|---|
| lungtumor | RTM-likeMT | CCL20 | CXCR4 | 0.2885 | 0.994 | 0.6413 | 0.000999 |
| lungtumor | RTM-likeMT | CCL20 | FPR1 | 0.2885 | 0.4978 | 0.3932 | 0.000999 |
| lungtumor | RTM-likeMT | CD99 | CD99 | 0.5938 | 0.8652 | 0.7295 | 0.000999 |
| lungtumor | RTM-likeMT | CXCL10 | C3AR1 | 0.3417 | 0.4896 | 0.4156 | 0.000999 |
| lungtumor | RTM-likeMT | CXCL10 | C5AR1 | 0.3417 | 1.2739 | 0.8078 | 0.000999 |
| lungtumor | RTM-likeMT | CXCL10 | CCR1 | 0.3417 | 0.2784 | 0.31 | 0.000999 |
| lungtumor | RTM-likeMT | CXCL10 | CXCR4 | 0.3417 | 0.994 | 0.6678 | 0.000999 |
| lungtumor | RTM-likeMT | CXCL10 | FPR1 | 0.3417 | 0.4978 | 0.4197 | 0.000999 |
| lungtumor | RTM-likeMT | FGA | ITGB2 | 0.4445 | 1.4753 | 0.9599 | 0.000999 |
| lungtumor | RTM-likeMT | GRN | TNFRSF1B | 0.4026 | 0.7592 | 0.5809 | 0.000999 |
| lungtumor | RTM-likeMT | GSTP1 | CD44 | 2.4093 | 1.4501 | 1.9297 | 0.000999 |
| lungtumor | RTM-likeMT | HLA-DRA | CD4 | 0.9609 | 0.6999 | 0.8304 | 0.000999 |
| lungtumor | RTM-likeMT | HLA-DRA | PTPRC | 0.9609 | 0.5986 | 0.7798 | 0.000999 |
| lungtumor | RTM-likeMT | HMGB1 | AGTRAP | 1.1943 | 0.3972 | 0.7958 | 0.000999 |
| lungtumor | RTM-likeMT | HMGB1 | LY96 | 1.1943 | 1.4353 | 1.3148 | 0.000999 |
| lungtumor | RTM-likeMT | HMGB1 | TLR2 | 1.1943 | 0.3076 | 0.7509 | 0.000999 |
| lungtumor | RTM-likeMT | HMGB2 | CD44 | 0.5593 | 1.4501 | 1.0047 | 0.000999 |
| lungtumor | RTM-likeMT | HMGB2 | DAB2 | 0.5593 | 0.7084 | 0.6339 | 0.000999 |
| lungtumor | RTM-likeMT | HSP90B1 | LRP1 | 0.6141 | 0.6166 | 0.6154 | 0.000999 |
| lungtumor | RTM-likeMT | HSP90B1 | TNFRSF1B | 0.6141 | 0.7592 | 0.6867 | 0.000999 |
| lungtumor | RTM-likeMT | LGALS3 | CLEC7A | 0.9915 | 0.7777 | 0.8846 | 0.000999 |
| lungtumor | RTM-likeMT | LGALS3 | FCGR2A | 0.9915 | 1.0268 | 1.0092 | 0.000999 |
| lungtumor | RTM-likeMT | LGALS3 | FCGR2B | 0.9915 | 0.8966 | 0.9441 | 0.000999 |
| lungtumor | RTM-likeMT | LGALS3 | TFRC | 0.9915 | 0.9108 | 0.9512 | 0.000999 |
| lungtumor | RTM-likeMT | LGALS3 | TGFBR2 | 0.9915 | 0.3485 | 0.67 | 0.000999 |
| lungtumor | RTM-likeMT | MDK | LRP1 | 1.2896 | 0.6166 | 0.9531 | 0.000999 |
| lungtumor | RTM-likeMT | MDK | NOTCH2 | 1.2896 | 0.3059 | 0.7977 | 0.000999 |
| lungtumor | RTM-likeMT | MIF | CD74 | 1.4585 | 4.0176 | 2.7381 | 0.000999 |
| lungtumor | RTM-likeMT | MIF | TNFRSF14 | 1.4585 | 0.3624 | 0.9105 | 0.000999 |
| lungtumor | RTM-likeMT | PSAP | LRP1 | 0.5457 | 0.6166 | 0.5812 | 0.000999 |

| | | | | | | | |
|---|---|---|---|---|---|---|---|
| lungtumor | RTM-likeMT | RPS19 | C5AR1 | 3.5104 | 1.2739 | 2.3922 | 0.000999 |
| lungtumor | RTM-likeMT | RPS19 | CD4 | 3.5104 | 0.6999 | 2.1052 | 0.000999 |
| lungtumor | RTM-likeMT | RPS19 | ICAM1 | 3.5104 | 0.3422 | 1.9263 | 0.000999 |
| lungtumor | RTM-likeMT | RTN4 | CD44 | 0.4013 | 1.4501 | 0.9257 | 0.000999 |
| lungtumor | RTM-likeMT | RTN4 | CD99 | 0.4013 | 0.8652 | 0.6332 | 0.000999 |
| lungtumor | RTM-likeMT | SAA1 | C3AR1 | 0.2539 | 0.4896 | 0.3717 | 0.000999 |
| lungtumor | RTM-likeMT | SAA1 | C5AR1 | 0.2539 | 1.2739 | 0.7639 | 0.000999 |
| lungtumor | RTM-likeMT | SAA1 | CCR1 | 0.2539 | 0.2784 | 0.2661 | 0.000999 |
| lungtumor | RTM-likeMT | SAA1 | CXCR4 | 0.2539 | 0.994 | 0.6239 | 0.000999 |
| lungtumor | RTM-likeMT | SAA1 | FPR1 | 0.2539 | 0.4978 | 0.3758 | 0.000999 |
| lungtumor | RTM-likeMT | SAA1 | GPR65 | 0.2539 | 0.3783 | 0.3161 | 0.000999 |
| lungtumor | RTM-likeMT | SERPINA1 | LRP1 | 0.3624 | 0.6166 | 0.4895 | 0.000999 |
| lungtumor | RTM-likeMT | SPP1 | CD44 | 1.4874 | 1.4501 | 1.4687 | 0.000999 |
| lungtumor | RTM-likeMT | SPP1 | ITGB1 | 1.4874 | 0.3085 | 0.8979 | 0.000999 |
| lungtumor | RTM-likeMT | TFPI | LRP1 | 0.3615 | 0.6166 | 0.489 | 0.000999 |
| lungtumor | RTM-likeMT | TIMP1 | LRP1 | 0.6195 | 0.6166 | 0.6181 | 0.000999 |
| lungtumor | RTM-likeMT | FGA | ITGB1 | 0.4445 | 0.3085 | 0.3765 | 0.003 |
| lungtumor | RTM-likeMT | CALM1 | RPSA | 0.8426 | 1.7816 | 1.3121 | 0.005 |
| lungtumor | RTM-likeMT | RPS19 | RPSA | 3.5104 | 1.7816 | 2.646 | 0.005 |
| lungtumor | RTM-likeMT | SAA1 | SELENOS | 0.2539 | 0.3901 | 0.322 | 0.00699 |
| melanomatumo | RTM-likeMT | ARF1 | CD4 | 1.8545 | 0.6999 | 1.2772 | 0.000999 |
| melanomatumo | RTM-likeMT | ARF1 | CD44 | 1.8545 | 1.4501 | 1.6523 | 0.000999 |
| melanomatumo | RTM-likeMT | CD320 | TGFBR2 | 0.2588 | 0.3485 | 0.3037 | 0.000999 |
| melanomatumo | RTM-likeMT | GSTP1 | CD44 | 2.3341 | 1.4501 | 1.8921 | 0.000999 |
| melanomatumo | RTM-likeMT | HAS2 | CD44 | 0.3078 | 1.4501 | 0.8789 | 0.000999 |
| melanomatumo | RTM-likeMT | HEBP1 | C5AR1 | 0.3689 | 1.2739 | 0.8214 | 0.000999 |
| melanomatumo | RTM-likeMT | HMGB1 | AGTRAP | 1.6757 | 0.3972 | 1.0365 | 0.000999 |
| melanomatumo | RTM-likeMT | HMGB1 | LY96 | 1.6757 | 1.4353 | 1.5555 | 0.000999 |
| melanomatumo | RTM-likeMT | HMGB1 | TLR2 | 1.6757 | 0.3076 | 0.9916 | 0.000999 |
| melanomatumo | RTM-likeMT | MIF | CD74 | 2.6939 | 4.0176 | 3.3558 | 0.000999 |

| | | | | | | | |
|---|---|---|---|---|---|---|---|
| melanomatumo | RTM-likeMT | MIF | TNFRSF14 | 2.6939 | 0.3624 | 1.5282 | 0.000999 |
| melanomatumo | RTM-likeMT | SERPINE2 | LRP1 | 0.9323 | 0.6166 | 0.7744 | 0.000999 |
| melanomatumo | RTM-likeMT | HEBP1 | CXCR4 | 0.3689 | 0.994 | 0.6814 | 0.002 |
| melanomatumo | RTM-likeMT | HEBP1 | FPR3 | 0.3689 | 0.4004 | 0.3846 | 0.003 |
| melanomatumo | RTM-likeMT | HEBP1 | C3AR1 | 0.3689 | 0.4896 | 0.4292 | 0.004 |
| melanomatumo | RTM-likeMT | HEBP1 | FPR1 | 0.3689 | 0.4978 | 0.4333 | 0.004 |
| melanomatumo | RTM-likeMT | COPA | NOTCH2 | 0.4147 | 0.3059 | 0.3603 | 0.005 |
| melanomatumo | RTM-likeMT | HEBP1 | CCR1 | 0.3689 | 0.2784 | 0.3236 | 0.005 |
| melanomatumo | RTM-likeMT | CALM1 | TNFRSF1B | 1.0928 | 0.7592 | 0.926 | 0.00599 |
| ovarytumor | RTM-likeMT | ANXA1 | C3AR1 | 1.0314 | 0.4896 | 0.7605 | 0.000999 |
| ovarytumor | RTM-likeMT | ANXA1 | C5AR1 | 1.0314 | 1.2739 | 1.1527 | 0.000999 |
| ovarytumor | RTM-likeMT | ANXA1 | CCR1 | 1.0314 | 0.2784 | 0.6549 | 0.000999 |
| ovarytumor | RTM-likeMT | ANXA1 | CD4 | 1.0314 | 0.6999 | 0.8656 | 0.000999 |
| ovarytumor | RTM-likeMT | ANXA1 | CD44 | 1.0314 | 1.4501 | 1.2407 | 0.000999 |
| ovarytumor | RTM-likeMT | ANXA1 | CXCR4 | 1.0314 | 0.994 | 1.0127 | 0.000999 |
| ovarytumor | RTM-likeMT | ANXA1 | FPR1 | 1.0314 | 0.4978 | 0.7646 | 0.000999 |
| ovarytumor | RTM-likeMT | ANXA1 | FPR3 | 1.0314 | 0.4004 | 0.7159 | 0.000999 |
| ovarytumor | RTM-likeMT | APOE | LRP1 | 0.837 | 0.6166 | 0.7268 | 0.000999 |
| ovarytumor | RTM-likeMT | APOE | SORL1 | 0.837 | 0.303 | 0.57 | 0.000999 |
| ovarytumor | RTM-likeMT | APP | ALOX5 | 0.2907 | 0.4201 | 0.3554 | 0.000999 |
| ovarytumor | RTM-likeMT | APP | C3AR1 | 0.2907 | 0.4896 | 0.3901 | 0.000999 |
| ovarytumor | RTM-likeMT | APP | C5AR1 | 0.2907 | 1.2739 | 0.7823 | 0.000999 |
| ovarytumor | RTM-likeMT | APP | CCR1 | 0.2907 | 0.2784 | 0.2846 | 0.000999 |
| ovarytumor | RTM-likeMT | APP | CXCR4 | 0.2907 | 0.994 | 0.6424 | 0.000999 |
| ovarytumor | RTM-likeMT | APP | FPR1 | 0.2907 | 0.4978 | 0.3943 | 0.000999 |
| ovarytumor | RTM-likeMT | APP | LRP1 | 0.2907 | 0.6166 | 0.4537 | 0.000999 |
| ovarytumor | RTM-likeMT | APP | SORL1 | 0.2907 | 0.303 | 0.2968 | 0.000999 |
| ovarytumor | RTM-likeMT | APP | TGFBR2 | 0.2907 | 0.3485 | 0.3196 | 0.000999 |
| ovarytumor | RTM-likeMT | APP | TNFRSF14 | 0.2907 | 0.3624 | 0.3266 | 0.000999 |
| ovarytumor | RTM-likeMT | ARF1 | CD4 | 0.7364 | 0.6999 | 0.7182 | 0.000999 |

| | | | | | | | |
|---|---|---|---|---|---|---|---|
| ovarytumor | RTM-likeMT | ARF1 | CD44 | 0.7364 | 1.4501 | 1.0932 | 0.000999 |
| ovarytumor | RTM-likeMT | CALM1 | TNFRSF1B | 1.0087 | 0.7592 | 0.8839 | 0.000999 |
| ovarytumor | RTM-likeMT | CALR | CD44 | 0.7423 | 1.4501 | 1.0962 | 0.000999 |
| ovarytumor | RTM-likeMT | CALR | LRP1 | 0.7423 | 0.6166 | 0.6794 | 0.000999 |
| ovarytumor | RTM-likeMT | CCL2 | CCR1 | 0.3526 | 0.2784 | 0.3155 | 0.000999 |
| ovarytumor | RTM-likeMT | COL1A1 | CD44 | 0.3504 | 1.4501 | 0.9002 | 0.000999 |
| ovarytumor | RTM-likeMT | GRN | TNFRSF1B | 0.3896 | 0.7592 | 0.5744 | 0.000999 |
| ovarytumor | RTM-likeMT | GSTP1 | CD44 | 2.6185 | 1.4501 | 2.0343 | 0.000999 |
| ovarytumor | RTM-likeMT | HLA-DMA | CD4 | 0.2537 | 0.6999 | 0.4768 | 0.000999 |
| ovarytumor | RTM-likeMT | HLA-DRA | CD4 | 1.2369 | 0.6999 | 0.9684 | 0.000999 |
| ovarytumor | RTM-likeMT | HLA-DRA | PTPRC | 1.2369 | 0.5986 | 0.9178 | 0.000999 |
| ovarytumor | RTM-likeMT | HMGB1 | AGTRAP | 1.7272 | 0.3972 | 1.0622 | 0.000999 |
| ovarytumor | RTM-likeMT | HMGB1 | LY96 | 1.7272 | 1.4353 | 1.5812 | 0.000999 |
| ovarytumor | RTM-likeMT | HMGB1 | TLR2 | 1.7272 | 0.3076 | 1.0174 | 0.000999 |
| ovarytumor | RTM-likeMT | HMGB2 | CD44 | 0.4974 | 1.4501 | 0.9737 | 0.000999 |
| ovarytumor | RTM-likeMT | HMGB2 | DAB2 | 0.4974 | 0.7084 | 0.6029 | 0.000999 |
| ovarytumor | RTM-likeMT | HSP90B1 | LRP1 | 0.6365 | 0.6166 | 0.6265 | 0.000999 |
| ovarytumor | RTM-likeMT | HSP90B1 | TNFRSF1B | 0.6365 | 0.7592 | 0.6978 | 0.000999 |
| ovarytumor | RTM-likeMT | ITGB1 | CD46 | 0.2783 | 0.3148 | 0.2965 | 0.000999 |
| ovarytumor | RTM-likeMT | ITGB1 | DAB2 | 0.2783 | 0.7084 | 0.4934 | 0.000999 |
| ovarytumor | RTM-likeMT | ITGB1 | ENG | 0.2783 | 0.3421 | 0.3102 | 0.000999 |
| ovarytumor | RTM-likeMT | ITGB1 | PLAUR | 0.2783 | 1.2007 | 0.7395 | 0.000999 |
| ovarytumor | RTM-likeMT | ITGB1 | PTK2B | 0.2783 | 0.3079 | 0.2931 | 0.000999 |
| ovarytumor | RTM-likeMT | LGALS3 | CLEC7A | 0.5097 | 0.7777 | 0.6437 | 0.000999 |
| ovarytumor | RTM-likeMT | LGALS3 | FCGR2A | 0.5097 | 1.0268 | 0.7683 | 0.000999 |
| ovarytumor | RTM-likeMT | LGALS3 | FCGR2B | 0.5097 | 0.8966 | 0.7032 | 0.000999 |
| ovarytumor | RTM-likeMT | LGALS3 | TFRC | 0.5097 | 0.9108 | 0.7103 | 0.000999 |
| ovarytumor | RTM-likeMT | LGALS3 | TGFBR2 | 0.5097 | 0.3485 | 0.4291 | 0.000999 |
| ovarytumor | RTM-likeMT | LRPAP1 | LRP1 | 0.3826 | 0.6166 | 0.4996 | 0.000999 |
| ovarytumor | RTM-likeMT | LRPAP1 | SORL1 | 0.3826 | 0.303 | 0.3428 | 0.000999 |

| | | | | | | | |
|---|---|---|---|---|---|---|---|
| ovarytumor | RTM-likeMT | MDK | LRP1 | 1.5321 | 0.6166 | 1.0743 | 0.000999 |
| ovarytumor | RTM-likeMT | MDK | NOTCH2 | 1.5321 | 0.3059 | 0.919 | 0.000999 |
| ovarytumor | RTM-likeMT | MIF | CD74 | 0.8565 | 4.0176 | 2.437 | 0.000999 |
| ovarytumor | RTM-likeMT | MIF | TNFRSF14 | 0.8565 | 0.3624 | 0.6094 | 0.000999 |
| ovarytumor | RTM-likeMT | PSAP | LRP1 | 0.6574 | 0.6166 | 0.637 | 0.000999 |
| ovarytumor | RTM-likeMT | RPS19 | C5AR1 | 3.9325 | 1.2739 | 2.6032 | 0.000999 |
| ovarytumor | RTM-likeMT | RPS19 | CD4 | 3.9325 | 0.6999 | 2.3162 | 0.000999 |
| ovarytumor | RTM-likeMT | RPS19 | ICAM1 | 3.9325 | 0.3422 | 2.1373 | 0.000999 |
| ovarytumor | RTM-likeMT | RTN4 | CD44 | 0.4992 | 1.4501 | 0.9746 | 0.000999 |
| ovarytumor | RTM-likeMT | RTN4 | CD99 | 0.4992 | 0.8652 | 0.6822 | 0.000999 |
| ovarytumor | RTM-likeMT | SERPING1 | LRP1 | 0.3678 | 0.6166 | 0.4922 | 0.000999 |
| ovarytumor | RTM-likeMT | SPP1 | CD44 | 0.3946 | 1.4501 | 0.9224 | 0.000999 |
| ovarytumor | RTM-likeMT | TIMP1 | LRP1 | 0.7779 | 0.6166 | 0.6973 | 0.000999 |
| ovarytumor | RTM-likeMT | CALM1 | TNFRSF1A | 1.0087 | 0.2803 | 0.6445 | 0.002 |
