## Supplementary Table 9 for "Identification of novel myeloid-derived cell states with implication in cancer outcome"

**Supplementary Table 9: Univariate Cox Regression analysis results**

| TCGA-OV | | |
|---|---|---|
| **Cell States** | **HR (95% CI for HR)** | **P.value** |
| Mac_Angio | 1.1 (0.93-1.2) | 0.42 |
| Mac_Reg | 1 (0.92-1.2) | 0.5 |
| **RTM_IM** | **1.1 (1-1.3)** | **0.035** |
| Malignant Cells | 0.93 (0.82-1.1) | 0.28 |
| Mac_Rec | 0.95 (0.83-1.1) | 0.43 |
| Fibroblasts | 1 (0.91-1.2) | 0.63 |
| **TCD4_ex** | **0.87 (0.77-0.99)** | **0.04** |
| Endothelial | 1 (0.88-1.1) | 0.98 |
| TCD8_naive | 0.93 (0.82-1.1) | 0.26 |
| pDC | 0.94 (0.83-1.1) | 0.37 |
| B_cells | 0.91 (0.81-1) | 0.16 |
| NK_rest | 0.93 (0.82-1.1) | 0.25 |
| TCD4_reg | 0.9 (0.79-1) | 0.1 |
| RTM_LA | 1 (0.91-1.2) | 0.69 |
| cDC2B_FCER1A | 0.97 (0.85-1.1) | 0.61 |
| cDC2A_AREG | 0.98 (0.86-1.1) | 0.78 |
| TCD4_em | 0.96 (0.84-1.1) | 0.54 |
| cDC2_CD14 | 0.96 (0.85-1.1) | 0.58 |
| Mac_Hypo | 1 (0.92-1.2) | 0.49 |
| TCD4_naive | 0.95 (0.83-1.1) | 0.4 |
| TCD8_em | 0.92 (0.81-1) | 0.21 |
| Mac_IFN | 1 (0.91-1.2) | 0.64 |
| **TCD8_ex** | **0.86 (0.76-0.98)** | **0.02** |
| **MonoInter_CXCL10** | **0.86 (0.76-0.98)** | **0.024** |
| Epithelial Cells | 0.93 (0.82-1.1) | 0.24 |
| NK_cyto | 0.96 (0.84-1.1) | 0.51 |
| RTM_IFN | 0.97 (0.85-1.1) | 0.58 |

| TCGA-BRCA: Basal | | |
|---|---|---|
| **Cell States** | **HR (95% CI for HR)** | **P.value** |
| TCD4_em | 1.1 (0.75-1.6) | 0.66 |
| TCD4_naive | 1 (0.7-1.5) | 0.93 |
| TCD8_naive | 0.97 (0.67-1.4) | 0.88 |
| TCD8_em | 0.88 (0.61-1.3) | 0.5 |
| B_cells | 0.77 (0.54-1.1) | 0.15 |
| TCD4_ex | 0.94 (0.64-1.4) | 0.77 |
| NK_rest | 0.88 (0.62-1.3) | 0.48 |
| TCD4_reg | 0.87 (0.59-1.3) | 0.47 |
| NK_cyto | 0.67 (0.44-1) | 0.069 |
| Mac_Reg | 0.98 (0.67-1.5) | 0.93 |
| cDC1_CLEC9A | 0.91 (0.64-1.3) | 0.58 |
| cDC2A_AREG | 0.8 (0.51-1.3) | 0.36 |
| cDC2B_FCER1A | 0.87 (0.53-1.4) | 0.56 |
| RTM_IM | 1.5 (1-2.1) | 0.04 |
| **Mono_FCG3RA** | **1.8 (1.2-2.6)** | **0.0032** |
| Mac_Rec | 1.1 (0.74-1.7) | 0.61 |
| Neutrophil_MMP9 | 1.1 (0.74-1.5) | 0.73 |
| Mast_cells | 1.4 (0.93-2.2) | 0.1 |
| **NKT** | **0.63 (0.42-0.94)** | **0.024** |
| RTM_LA | 0.84 (0.59-1.2) | 0.32 |
| TCD8_ex | 0.97 (0.68-1.4) | 0.87 |
| Mono_IL1B | 0.8 (0.56-1.2) | 0.24 |
| Mono_CD14_FOS+ | 0.84 (0.57-1.2) | 0.38 |
| MonoInter_CLEC10A | 1.1 (0.78-1.6) | 0.53 |
| Mac_Angio | 1.3 (0.85-1.8) | 0.25 |
| cDC2_CD14 | 0.66 (0.42-1) | 0.071 |
| MonoInter_CXCL10 | 0.79 (0.55-1.1) | 0.2 |

| | | |
|---|---|---|
| RTM_like_MT | 0.97 (0.86-1.1) | 0.69 |
| Mac_LA | 1 (0.89-1.2) | 0.83 |
| Mast_cells | 0.94 (0.83-1.1) | 0.3 |
| MonoInter_CLEC10A | 1 (0.89-1.1) | 0.86 |
| Mono_IL1B | 1.1 (0.93-1.2) | 0.41 |
| cDC2_CD207 | 0.9 (0.79-1) | 0.091 |
| NKT | 0.92 (0.81-1) | 0.2 |
| cDC1_CLEC9A | 0.94 (0.83-1.1) | 0.33 |
| cDC2_FCGR3A | 0.9 (0.79-1) | 0.1 |
| Mono_FCG3RA | 0.97 (0.85-1.1) | 0.62 |
| Mono_CD14_FOS+ | 1 (0.92-1.2) | 0.48 |
| Age | 1 (1-1) | **0.0011** |
| Stage | -1.6 (0.011-0.85) | 0.1421 |
| Subtype_mRNA | 0.26 (0.45-1.1) | 0.0675 |

| **TCGA-BRCA: HER2** | | |
|---|---|---|
| **Cell States** | **HR (95% CI for HR)** | **P.value** |
| TCD4_em | 1.3 (0.83-2) | 0.25 |
| TCD4_naive | 1.1 (0.69-1.7) | 0.71 |
| TCD8_naive | 0.79 (0.49-1.3) | 0.34 |
| TCD8_em | 0.88 (0.55-1.4) | 0.59 |
| B_cells | 0.8 (0.5-1.3) | 0.37 |
| TCD4_ex | 0.75 (0.44-1.3) | 0.27 |
| NK_rest | 1.1 (0.67-1.7) | 0.78 |
| TCD4_reg | 0.97 (0.59-1.6) | 0.91 |
| NK_cyto | 0.71 (0.39-1.3) | 0.27 |
| Mac_Reg | 1.1 (0.62-1.9) | 0.77 |
| cDC1_CLEC9A | 0.78 (0.49-1.3) | 0.31 |
| cDC2A_AREG | 1.1 (0.63-1.8) | 0.82 |
| cDC2B_FCER1A | 1.2 (0.74-2.1) | 0.42 |

| | | |
|---|---|---|
| Endothelial | 1 (0.69-1.5) | 0.95 |
| Fibroblasts | 1.3 (0.85-2) | 0.23 |
| pDC | 1.1 (0.73-1.6) | 0.74 |
| Neutrophil_TAGLN2 | 1.2 (0.85-1.8) | 0.26 |
| Mac_LA | 1.3 (0.86-1.9) | 0.22 |
| Neutrophil_CXCL8 | 1.3 (0.88-1.9) | 0.2 |
| Mac_IFN | 0.89 (0.61-1.3) | 0.56 |
| Mac_Hypo | 1.3 (0.9-1.9) | 0.16 |
| Mac_Alv_like | 0.83 (0.57-1.2) | 0.32 |
| RTM_IFN | 0.91 (0.66-1.3) | 0.58 |
| **cDC2_FCGR3A** | **0.63 (0.44-0.91)** | **0.013** |
| Epithelial Cells | 0.92 (0.57-1.5) | 0.76 |
| Malignant Cells | 0.99 (0.67-1.5) | 0.94 |
| TGD | 0.98 (0.69-1.4) | 0.89 |
| Age | 1 (0.97-1) | 0.84 |
| **Stage** | **20 (0-Inf)** | **8.925e-05** |

| **TCGA-BRCA: Luminal A** | | |
|---|---|---|
| **Cell States** | **HR (95% CI for HR)** | **P.value** |
| TCD4_em | 1 (0.76-1.4) | 0.83 |
| TCD4_naive | 0.78 (0.57-1.1) | 0.1 |
| TCD8_naive | 0.81 (0.6-1.1) | 0.17 |
| TCD8_em | 0.86 (0.64-1.2) | 0.34 |
| B_cells | 0.78 (0.57-1.1) | 0.13 |
| **TCD4_ex** | **0.7 (0.5-0.98)** | **0.039** |
| NK_rest | 0.85 (0.63-1.1) | 0.29 |
| TCD4_reg | 0.93 (0.68-1.3) | 0.65 |
| NK_cyto | 1.2 (0.85-1.6) | 0.31 |
| Mac_Reg | 0.86 (0.63-1.2) | 0.31 |
| **cDC1_CLEC9A** | **0.69 (0.5-0.97)** | **0.034** |

| | | |
|---|---|---|
| RTM_IM | 0.96 (0.53-1.7) | 0.88 |
| Mono_FCG3RA | 1.5 (0.93-2.5) | 0.093 |
| Mac_Rec | 0.82 (0.52-1.3) | 0.41 |
| Neutrophil_MMP9 | 0.9 (0.59-1.4) | 0.63 |
| Mast_cells | 0.67 (0.35-1.3) | 0.22 |
| NKT | 0.71 (0.43-1.2) | 0.19 |
| RTM_LA | 0.76 (0.48-1.2) | 0.25 |
| TCD8_ex | 0.79 (0.5-1.2) | 0.31 |
| Mono_IL1B | 0.92 (0.57-1.5) | 0.73 |
| Mono_CD14_FOS+ | 1 (0.62-1.8) | 0.86 |
| MonoInter_CLEC10A | 1.1 (0.68-1.9) | 0.64 |
| Mac_Angio | 0.85 (0.54-1.3) | 0.47 |
| cDC2_CD14 | 0.84 (0.52-1.4) | 0.47 |
| MonoInter_CXCL10 | 0.93 (0.59-1.5) | 0.74 |
| Endothelial | 1.3 (0.8-2.3) | 0.27 |
| Fibroblasts | 0.76 (0.46-1.3) | 0.28 |
| pDC | 0.71 (0.41-1.2) | 0.2 |
| Neutrophil_TAGLN2 | 0.74 (0.41-1.3) | 0.32 |
| Mac_LA | 0.91 (0.55-1.5) | 0.72 |
| Neutrophil_CXCL8 | 0.78 (0.44-1.4) | 0.39 |
| Mac_IFN | 0.86 (0.51-1.4) | 0.55 |
| Mac_Hypo | 1 (0.61-1.7) | 0.95 |
| Mac_Alv_like | 0.92 (0.57-1.5) | 0.72 |
| RTM_IFN | 0.93 (0.58-1.5) | 0.78 |
| cDC2_FCGR3A | 0.8 (0.47-1.3) | 0.4 |
| Epithelial Cells | 0.98 (0.64-1.5) | 0.93 |
| **Malignant Cells** | **1.9 (1-3.4)** | **0.045** |
| TGD | 0.86 (0.55-1.3) | 0.51 |
| **Age** | **1.1 (1-1.1)** | **0.0021** |
| **Stage** | **1.1 (0.084-6.8)** | **0.0025** |

| | | |
|---|---|---|
| cDC2A_AREG | 0.76 (0.57-1) | 0.066 |
| cDC2B_FCER1A | 0.83 (0.61-1.1) | 0.25 |
| RTM_IM | 1 (0.77-1.4) | 0.76 |
| **Mono_FCG3RA** | **0.64 (0.47-0.89)** | **0.0075** |
| Mac_Rec | 0.74 (0.53-1) | 0.083 |
| Neutrophil_MMP9 | 1.1 (0.81-1.5) | 0.51 |
| Mast_cells | 0.87 (0.64-1.2) | 0.4 |
| NKT | 1.1 (0.79-1.5) | 0.64 |
| RTM_LA | 0.84 (0.61-1.1) | 0.28 |
| TCD8_ex | 0.83 (0.61-1.1) | 0.25 |
| Mono_IL1B | 0.84 (0.61-1.1) | 0.27 |
| Mono_CD14_FOS+ | 0.96 (0.7-1.3) | 0.8 |
| MonoInter_CLEC10A | 0.88 (0.65-1.2) | 0.39 |
| Mac_Angio | 0.84 (0.62-1.1) | 0.25 |
| **cDC2_CD14** | **0.72 (0.53-0.96)** | **0.028** |
| MonoInter_CXCL10 | 0.82 (0.59-1.1) | 0.24 |
| Endothelial | 1.1 (0.8-1.5) | 0.55 |
| Fibroblasts | 0.89 (0.67-1.2) | 0.42 |
| **pDC** | **0.71 (0.53-0.97)** | **0.029** |
| Neutrophil_TAGLN2 | 0.93 (0.68-1.3) | 0.65 |
| Mac_LA | 0.92 (0.67-1.3) | 0.59 |
| Neutrophil_CXCL8 | 0.93 (0.69-1.3) | 0.66 |
| Mac_IFN | 0.83 (0.6-1.1) | 0.24 |
| Mac_Hypo | 0.79 (0.58-1.1) | 0.13 |
| Mac_Alv_like | 0.85 (0.63-1.2) | 0.31 |
| RTM_IFN | 0.8 (0.57-1.1) | 0.18 |
| **cDC2_FCGR3A** | **0.72 (0.53-0.99)** | **0.041** |
| Epithelial Cells | 0.99 (0.73-1.4) | 0.97 |
| Malignant Cells | 1.2 (0.92-1.6) | 0.17 |
| TGD | 0.78 (0.58-1.1) | 0.12 |

| TCGA-BRCA: Luminal B | | |
|---|---|---|
| **Cell States** | **HR (95% CI for HR)** | **P.value** |
| TCD4_em | 0.56 (0.39-0.83) | 0.0034 |
| TCD4_naive | 0.87 (0.59-1.3) | 0.48 |
| TCD8_naive | 0.91 (0.63-1.3) | 0.6 |
| TCD8_em | 0.95 (0.66-1.4) | 0.8 |
| B_cells | 0.67 (0.44-1) | 0.056 |
| TCD4_ex | 1.1 (0.72-1.6) | 0.75 |
| NK_rest | 0.68 (0.47-0.99) | 0.043 |
| TCD4_reg | 0.94 (0.65-1.4) | 0.76 |
| NK_cyto | 1.1 (0.75-1.6) | 0.63 |
| Mac_Reg | 0.97 (0.66-1.4) | 0.88 |
| cDC1_CLEC9A | 0.95 (0.65-1.4) | 0.78 |
| cDC2A_AREG | 0.84 (0.51-1.4) | 0.5 |
| cDC2B_FCER1A | 0.91 (0.59-1.4) | 0.68 |
| RTM_IM | 1.1 (0.72-1.7) | 0.66 |
| Mono_FCG3RA | 1.1 (0.78-1.6) | 0.55 |
| Mac_Rec | 1.3 (0.87-1.8) | 0.23 |
| Neutrophil_MMP9 | 1 (0.7-1.5) | 0.97 |
| Mast_cells | 1.3 (0.9-2) | 0.15 |
| NKT | 1.4 (0.93-2) | 0.11 |
| RTM_LA | 1.2 (0.82-1.7) | 0.37 |
| TCD8_ex | 0.99 (0.69-1.4) | 0.95 |
| Mono_IL1B | 1.3 (0.89-1.8) | 0.2 |
| Mono_CD14_FOS+ | 0.98 (0.68-1.4) | 0.93 |
| MonoInter_CLEC10A | 1.2 (0.83-1.7) | 0.37 |
| **Mac_Angio** | **1.5 (1-2.2)** | **0.048** |

| | | |
|---|---|---|
| **Age** | **1.1 (1-1.1)** | **2e-06** |
| **Stage** | **1.1 (0.64-5.7)** | **0.02362** |

| TCGA-LUAD | | |
|---|---|---|
| **Cell States** | **HR (95% CI for HR)** | **P.value** |
| TCD8_ex | 0.97 (0.85-1.1) | 0.7 |
| TCD4_ex | 0.95 (0.83-1.1) | 0.44 |
| TCD4_em | 0.93 (0.81-1.1) | 0.31 |
| TCD4_reg | 0.96 (0.84-1.1) | 0.51 |
| Mac_IFN | 1 (0.9-1.2) | 0.65 |
| TCD8_em | 0.89 (0.78-1) | 0.095 |
| RTM_IFN | 1 (0.87-1.1) | 0.94 |
| NK_cyto | 0.95 (0.83-1.1) | 0.45 |
| MonoInter_CXCL10 | 1 (0.88-1.1) | 0.99 |
| TCD8_naive | 0.96 (0.84-1.1) | 0.51 |
| **Malignant Cells** | **1.3 (1.1-1.4)** | **0.00088** |
| Mac_Rec | 1 (0.89-1.2) | 0.83 |
| **B_cells** | **0.82 (0.72-0.94)** | **0.0034** |
| cDC2A_AREG | 0.95 (0.84-1.1) | 0.47 |
| Mac_Alv_like | 0.92 (0.81-1.1) | 0.24 |
| NKT | 0.9 (0.78-1) | 0.12 |
| RTM_LA | 0.9 (0.78-1) | 0.1 |
| TCD4_naive | 0.95 (0.82-1.1) | 0.42 |
| **Mast_cells** | **0.8 (0.7-0.92)** | **0.0014** |
| Mono_FCG3RA | 1 (0.9-1.2) | 0.65 |
| Endothelial | 0.95 (0.83-1.1) | 0.46 |
| **Mac_Angio** | **1.2 (1.1-1.4)** | **0.0059** |
| RTM_like_MT | 0.98 (0.86-1.1) | 0.78 |
| NK_rest | 0.96 (0.83-1.1) | 0.52 |
| Epithelial Cells | 0.92 (0.81-1) | 0.2 |

| | | |
|---|---|---|
| cDC2_CD14 | 1.1 (0.75-1.6) | 0.67 |
| MonoInter_CXCL10 | 1.1 (0.77-1.6) | 0.56 |
| Endothelial | 0.66 (0.4-1.1) | 0.12 |
| **Fibroblasts** | **1.6 (1.1-2.4)** | **0.011** |
| pDC | 0.66 (0.42-1) | 0.065 |
| Neutrophil_TAGLN2 | 1.1 (0.78-1.7) | 0.49 |
| Mac_LA | 1.5 (0.99-2.2) | 0.055 |
| Neutrophil_CXCL8 | 1.1 (0.79-1.7) | 0.49 |
| Mac_IFN | 0.96 (0.67-1.4) | 0.83 |
| Mac_Hypo | 1.4 (0.93-2) | 0.11 |
| Mac_Alv_like | 1.1 (0.74-1.6) | 0.7 |
| RTM_IFN | 0.93 (0.65-1.3) | 0.69 |
| cDC2_FCGR3A | 0.8 (0.55-1.2) | 0.23 |
| Epithelial Cells | 0.82 (0.53-1.3) | 0.38 |
| Malignant Cells | 0.81 (0.55-1.2) | 0.3 |
| TGD | 0.83 (0.56-1.2) | 0.37 |
| Age | 1 (0.99-1.1) | 0.1 |
| **Stage** | **1.6 (0.28-18)** | **0.01858** |

| **TCGA-LUSC** | | |
|---|---|---|
| **Cell States** | **HR (95% CI for HR)** | **P.value** |
| Mac_Rec | 0.98 (0.86-1.1) | 0.74 |
| Endothelial | 1.1 (0.99-1.3) | 0.06 |
| TCD4_ex | 0.95 (0.83-1.1) | 0.42 |
| Mono_CD14_FOS- | 1 (0.88-1.1) | 0.96 |
| TCD4_reg | 0.95 (0.83-1.1) | 0.46 |
| Fibroblasts | 1.1 (0.97-1.2) | 0.15 |
| Malignant Cells | 0.9 (0.79-1) | 0.1 |
| Mast_cells | 0.98 (0.86-1.1) | 0.74 |
| RTM_LA | 0.91 (0.8-1) | 0.14 |
| Fibroblasts | 1 (0.87-1.1) | 0.95 |
| MonoInter_CLEC10A | 1.1 (0.95-1.2) | 0.2 |
| pDC | 0.99 (0.87-1.1) | 0.94 |
| **RTM_IM** | **0.86 (0.75-0.98)** | **0.025** |
| **Mono_IL1B** | **1.2 (1.1-1.4)** | **0.0022** |
| cDC2_FCGR3A | 1 (0.9-1.2) | 0.67 |
| Mac_Reg | 0.92 (0.81-1.1) | 0.24 |
| Mac_Hypo | 1.1 (0.97-1.3) | 0.12 |
| Mono_CD14_FOS- | 1 (0.89-1.2) | 0.76 |
| cDC1_CLEC9A | 0.85 (0.74-0.96) | 0.013 |
| **cDC2_CD207** | **0.84 (0.74-0.95)** | **0.0078** |
| **cDC2B_FCER1A** | **0.85 (0.75-0.97)** | **0.016** |
| **cDC2_CXCL8** | **1.2 (1-1.4)** | **0.011** |
| cDC2_CD14 | 0.91 (0.79-1) | 0.15 |
| Mac_LA | 0.96 (0.84-1.1) | 0.57 |
| Neutrophil_TAGLN2 | 1.1 (0.98-1.3) | 0.11 |
| **Neutrophil_MMP9** | **1.2 (1.1-1.4)** | **0.0017** |
| **Neutrophil_CXCL8** | **1.1 (1-1.3)** | **0.043** |
| Age | 1 (0.99-1) | 0.51 |
| gender | 1.1 (0.79-1.4) | 0.67 |
| **Stage** | **1.3 (1.6-3.5)** | **1.822e-11** |

| **TCGA-LIHC** | | |
|---|---|---|
| **Cell States** | **HR (95% CI for HR)** | **P.value** |
| NK_rest | 0.99 (0.84-1.2) | 0.89 |
| Endothelial | 1 (0.87-1.2) | 0.75 |
| Fibroblasts | 1.1 (0.9-1.2) | 0.49 |
| TCD4_naive | 0.91 (0.77-1.1) | 0.22 |
| TCD8_naive | 0.86 (0.73-1) | 0.055 |
| Epithelial Cells | 1.2 (0.99-1.4) | 0.074 |

| | | | | | |
|---|---|---|---|---|---|
| Epithelial Cells | 1 (0.87-1.1) | 0.96 | Malignant Cells | 0.86 (0.73-1) | 0.078 |
| **Mac_Angio** | **1.2 (1-1.4)** | **0.0094** | TCD8_em | 0.91 (0.77-1.1) | 0.22 |
| TCD4_em | 1 (0.9-1.2) | 0.67 | TCD4_em | 0.94 (0.8-1.1) | 0.44 |
| TCD8_ex | 0.96 (0.84-1.1) | 0.53 | NKT | 0.87 (0.74-1) | 0.081 |
| TCD4_naive | 0.89 (0.78-1) | 0.093 | TCD8_ex | 0.94 (0.8-1.1) | 0.44 |
| RTM_IM | 1.1 (0.93-1.2) | 0.37 | TCD4_reg | 0.98 (0.83-1.1) | 0.77 |
| TCD8_em | 0.96 (0.84-1.1) | 0.53 | B_cells | 0.93 (0.79-1.1) | 0.35 |
| Mac_IFN | 0.99 (0.86-1.1) | 0.82 | NK_cyto | 0.94 (0.8-1.1) | 0.42 |
| cDC2B_FCER1A | 0.91 (0.8-1) | 0.16 | pDC | 1 (0.86-1.2) | 0.88 |
| RTM_IFN | 1.1 (0.99-1.3) | 0.072 | **cDC2_CXCL8** | **0.81 (0.69-0.96)** | **0.012** |
| NKT | 0.92 (0.81-1) | 0.19 | RTM_IM | 0.86 (0.73-1) | 0.059 |
| Mac_LA | 1.1 (0.95-1.2) | 0.22 | Mono_FCG3RA | 0.95 (0.81-1.1) | 0.51 |
| **NK_cyto** | **0.85 (0.75-0.96)** | **0.012** | cDC1_CLEC9A | 0.94 (0.8-1.1) | 0.46 |
| B_cells | 0.99 (0.87-1.1) | 0.86 | Mac_Rec | 0.91 (0.77-1.1) | 0.24 |
| Mac_Hypo | 1.1 (0.97-1.3) | 0.15 | **Mac_Reg** | **0.82 (0.69-0.97)** | **0.017** |
| TCD8_naive | 0.93 (0.82-1.1) | 0.27 | MonoInter_CLEC10A | 0.95 (0.81-1.1) | 0.57 |
| **Mono_IL1B** | **1.2 (1-1.3)** | **0.02** | cDC2A_AREG | 0.91 (0.78-1.1) | 0.26 |
| cDC2A_AREG | 0.91 (0.8-1) | 0.17 | cDC2_CD14 | 0.88 (0.75-1) | 0.12 |
| NK_rest | 0.97 (0.85-1.1) | 0.68 | Mono_IL1B | 1 (0.88-1.2) | 0.73 |
| Mono_FCG3RA | 0.95 (0.84-1.1) | 0.46 | RTM_IFN | 1.1 (0.95-1.3) | 0.18 |
| pDC | 1 (0.89-1.2) | 0.86 | **Mac_IFN** | **1.3 (1.1-1.5)** | **0.0014** |
| cDC2_CD14 | 0.91 (0.8-1) | 0.15 | cDC_LAMP3 | 0.91 (0.78-1.1) | 0.27 |
| RTM_like_MT | 0.93 (0.82-1.1) | 0.24 | TCD4_ex | 1 (0.85-1.2) | 0.99 |
| Mac_Alv_like | 0.93 (0.82-1.1) | 0.26 | **RTM_LA** | **1.3 (1.1-1.5)** | **0.0013** |
| cDC2_CXCL8 | 1.1 (0.94-1.2) | 0.32 | MonoInter_CXCL10 | 0.89 (0.75-1) | 0.15 |
| MonoInter_CLEC10A | 1 (0.88-1.1) | 0.97 | cDC2_FCGR3A | 0.86 (0.73-1) | 0.058 |
| cDC1_CLEC9A | 0.98 (0.86-1.1) | 0.8 | Mac_Angio | 1 (0.88-1.2) | 0.64 |
| MonoInter_CXCL10 | 0.98 (0.86-1.1) | 0.72 | cDC2_CD207 | 0.91 (0.77-1.1) | 0.23 |
| cDC2_CD207 | 0.92 (0.81-1.1) | 0.22 | Mac_Hypo | 0.98 (0.84-1.2) | 0.82 |
| Mac_Reg | 0.99 (0.87-1.1) | 0.93 | Mac_Alv_like | 1 (0.89-1.2) | 0.63 |

| | | |
|---|---|---|
| Neutrophil_CXCL8 | 1 (0.9-1.2) | 0.69 |
| Neutrophil_TAGLN2 | 1.1 (0.94-1.2) | 0.32 |
| Neutrophil_MMP9 | 1.1 (0.96-1.3) | 0.18 |
| cDC2_FCGR3A | 1 (0.91-1.2) | 0.55 |
| **Stage** | **0.56 (0.92-1.8)** | **0.007** |
| Age | 1 (0.99-1) | 0.28 |
| gender | 1.3 (0.9-1.8) | 0.17 |

| **TCGA-COAD** | | |
|---|---|---|
| **Cell States** | **HR (95% CI for HR)** | **P.value** |
| Endothelial | 1.1 (0.92-1.3) | 0.27 |
| TCD8_em | 0.95 (0.8-1.1) | 0.62 |
| Epithelial Cells | 0.98 (0.82-1.2) | 0.82 |
| TCD4_ex | 0.95 (0.78-1.1) | 0.55 |
| cDC2_CD14 | 0.85 (0.71-1) | 0.09 |
| TCD4_naive | 0.99 (0.82-1.2) | 0.95 |
| TCD4_reg | 0.9 (0.75-1.1) | 0.27 |
| NK_rest | 0.91 (0.76-1.1) | 0.3 |
| B_cells | 0.94 (0.78-1.1) | 0.51 |
| TCD8_ex | 1 (0.85-1.2) | 0.82 |
| Mast_cells | 0.92 (0.76-1.1) | 0.36 |
| RTM_LA | 1 (0.84-1.2) | 0.93 |
| Mac_IFN | 1 (0.86-1.2) | 0.75 |
| Fibroblasts | 1.1 (0.91-1.3) | 0.32 |
| Mono_IL1B | 0.95 (0.79-1.1) | 0.57 |
| TCD4_em | 0.97 (0.8-1.2) | 0.74 |
| RTM_IM | 1.1 (0.91-1.3) | 0.37 |
| Mac_Hypo | 1 (0.84-1.2) | 0.98 |
| Mac_Reg | 0.94 (0.78-1.1) | 0.48 |
| Malignant Cells | 1 (0.84-1.2) | 0.93 |

| | | |
|---|---|---|
| Age | 1 (1-1) | 0.14 |
| gender | 0.89 (0.62-1.3) | 0.56 |
| **Stage** | **1 (0.85-2.4)** | **3.139e-05** |

| **TCGA-READ** | | |
|---|---|---|
| **Cell States** | **HR (95% CI for HR)** | **P.value** |
| Endothelial | 0.92 (0.64-1.3) | 0.67 |
| TCD8_em | 0.98 (0.69-1.4) | 0.92 |
| Epithelial Cells | 1 (0.69-1.4) | 0.99 |
| TCD4_ex | 0.81 (0.56-1.2) | 0.24 |
| cDC2_CD14 | 0.83 (0.57-1.2) | 0.3 |
| **TCD4_naive** | **0.68 (0.48-0.96)** | **0.028** |
| TCD4_reg | 0.9 (0.63-1.3) | 0.53 |
| NK_rest | 0.88 (0.62-1.3) | 0.48 |
| B_cells | 0.88 (0.6-1.3) | 0.5 |
| TCD8_ex | 0.87 (0.61-1.2) | 0.43 |
| Mast_cells | 1.1 (0.74-1.5) | 0.77 |
| RTM_LA | 1.3 (0.91-1.9) | 0.14 |
| Mac_IFN | 1.1 (0.79-1.6) | 0.51 |
| Fibroblasts | 1.1 (0.77-1.6) | 0.56 |
| Mono_IL1B | 0.87 (0.62-1.2) | 0.41 |
| TCD4_em | 0.93 (0.66-1.3) | 0.68 |
| RTM_IM | 1.3 (0.94-1.9) | 0.11 |
| Mac_Hypo | 1.2 (0.85-1.6) | 0.33 |
| Mac_Reg | 0.93 (0.66-1.3) | 0.69 |
| Malignant Cells | 0.94 (0.66-1.3) | 0.74 |
| RTM_IFN | 1.2 (0.85-1.7) | 0.3 |
| pDC | 0.78 (0.55-1.1) | 0.17 |

| | | |
|---|---|---|
| RTM_IFN | 1.1 (0.89-1.3) | 0.47 |
| pDC | 0.93 (0.78-1.1) | 0.46 |
| NK_cyto | 0.84 (0.69-1) | 0.067 |
| cDC2A_AREG | 0.85 (0.7-1) | 0.074 |
| cDC2B_FCER1A | 0.95 (0.79-1.1) | 0.55 |
| cDC_LAMP3 | 0.89 (0.74-1.1) | 0.2 |
| TCD8_naive | 0.87 (0.72-1) | 0.14 |
| MonoInter_CLEC10A | 0.92 (0.76-1.1) | 0.35 |
| Mac_Rec | 0.97 (0.8-1.2) | 0.71 |
| Mac_LA | 1 (0.87-1.3) | 0.63 |
| Mono_CD14_FOS+ | 0.96 (0.8-1.2) | 0.67 |
| Mac_Angio | 1 (0.86-1.2) | 0.69 |
| NKT | 0.98 (0.81-1.2) | 0.8 |
| Mono_CD14_FOS- | 1 (0.85-1.2) | 0.78 |
| Neutrophil_TAGLN2 | 0.91 (0.76-1.1) | 0.34 |
| MonoInter_CXCL10 | 0.96 (0.8-1.2) | 0.7 |
| Neutrophil_CXCL8 | 0.93 (0.78-1.1) | 0.44 |
| Neutrophil_MMP9 | 0.88 (0.73-1.1) | 0.2 |
| Mac_Alv_like | 1 (0.87-1.3) | 0.65 |
| **Age** | **1 (1-1)** | **0.013** |
| gender | 1 (0.66-1.5) | 0.99 |
| **Stage** | **1.3 (0.57-4.1)** | **2.324e-09** |

| **TCGA-SKCM : Metastasis** | | |
|---|---|---|
| **Cell States** | **HR (95% CI for HR)** | **P.value** |
| **TCD4_reg** | **0.62 (0.54-0.71)** | **1.1e-12** |
| **TCD8_ex** | **0.67 (0.59-0.77)** | **2.1e-09** |
| **TCD4_naive** | **0.86 (0.76-0.98)** | **0.027** |
| **B_cells** | **0.68 (0.6-0.77)** | **1.1e-09** |
| **NK_cyto** | **0.76 (0.67-0.87)** | **3.8e-05** |

| | | |
|---|---|---|
| NK_cyto | 0.99 (0.71-1.4) | 0.97 |
| cDC2A_AREG | 0.9 (0.65-1.3) | 0.55 |
| cDC2B_FCER1A | 0.96 (0.68-1.4) | 0.84 |
| cDC_LAMP3 | 0.93 (0.65-1.3) | 0.71 |
| TCD8_naive | 0.83 (0.59-1.2) | 0.28 |
| MonoInter_CLEC10A | 1 (0.73-1.4) | 0.95 |
| Mac_Rec | 1.1 (0.82-1.6) | 0.42 |
| Mac_LA | 1.3 (0.92-1.9) | 0.13 |
| Mono_CD14_FOS+ | 0.93 (0.67-1.3) | 0.65 |
| Mac_Angio | 1.4 (0.96-2) | 0.086 |
| NKT | 1.2 (0.83-1.6) | 0.38 |
| Mono_CD14_FOS- | 1 (0.72-1.5) | 0.9 |
| Neutrophil_TAGLN2 | 0.89 (0.63-1.3) | 0.51 |
| MonoInter_CXCL10 | 1.1 (0.77-1.5) | 0.62 |
| Neutrophil_CXCL8 | 1 (0.72-1.4) | 0.92 |
| Neutrophil_MMP9 | 0.87 (0.62-1.2) | 0.44 |
| Mac_Alv_like | 1.4 (0.97-2) | 0.078 |
| **Age** | **1.1 (1-1.1)** | **0.0017** |
| gender | 0.82 (0.38-1.8) | 0.61 |
| **Stage** | **0.88 (0.19-5.6)** | **0.007** |

| **TCGA-SKCM : Primary** | | |
|---|---|---|
| **Cell States** | **HR (95% CI for HR)** | **P.value** |
| TCD4_reg | 0.73 (0.53-1) | 0.058 |
| TCD8_ex | 0.75 (0.54-1) | 0.083 |
| TCD4_naive | 0.99 (0.71-1.4) | 0.97 |
| B_cells | 0.79 (0.59-1.1) | 0.14 |
| NK_cyto | 0.84 (0.61-1.2) | 0.28 |
| **TCD8_em** | **0.65 (0.46-0.92)** | **0.016** |
| TCD8_naive | 0.87 (0.62-1.2) | 0.45 |

| | | |
|---|---|---|
| **TCD8_em** | **0.72 (0.63-0.81)** | **3e-07** |
| **TCD8_naive** | **0.78 (0.68-0.88)** | **0.00011** |
| **pDC** | **0.66 (0.58-0.76)** | **1.9e-09** |
| **TCD4_em** | **0.81 (0.71-0.93)** | **0.0019** |
| Mono_IL1B | 0.95 (0.84-1.1) | 0.43 |
| **Malignant Cells** | **1.3 (1.1-1.4)** | **0.00018** |
| **Epithelial Cells** | **0.86 (0.76-0.98)** | **0.021** |
| Mono_CD14_FOS- | 0.9 (0.79-1) | 0.087 |
| **TCD4_ex** | **0.87 (0.77-0.99)** | **0.038** |
| Fibroblasts | 1.1 (0.94-1.2) | 0.33 |
| **Endothelial** | **0.84 (0.74-0.96)** | **0.0079** |
| **MonoInter_CXCL10** | **0.77 (0.68-0.88)** | **6.8e-05** |
| **NK_rest** | **0.8 (0.7-0.9)** | **0.00043** |
| **NKT** | **0.76 (0.67-0.87)** | **3.2e-05** |
| **RTM_IFN** | **0.73 (0.64-0.83)** | **1.5e-06** |
| gender | 1.1 (0.82-1.5) | 0.4832 |
| **Age** | **1 (1-1)** | **0.0017** |

| TCGA-UVM | | |
|---|---|---|
| **Cell States** | **HR (95% CI for HR)** | **P.value** |
| **Epithelial Cells** | **0.36 (0.22-0.58)** | **3.5e-05** |
| RTM_IM | 0.85 (0.6-1.2) | 0.39 |
| **Malignant Cells** | **2.8 (1.7-4.6)** | **3.1e-05** |
| NK_rest | 0.97 (0.69-1.4) | 0.87 |
| Endothelial | 1 (0.71-1.4) | 0.98 |
| Mono_IL1B | 0.91 (0.64-1.3) | 0.6 |
| Mac_Reg | 1.1 (0.75-1.5) | 0.69 |
| TCD4_naive | 0.88 (0.62-1.3) | 0.48 |
| Fibroblasts | 1.2 (0.84-1.7) | 0.32 |
| TCD4_reg | 1.3 (0.92-2) | 0.12 |

| | | |
|---|---|---|
| pDC | 0.94 (0.69-1.3) | 0.72 |
| TCD4_em | 0.91 (0.66-1.2) | 0.54 |
| Mono_IL1B | 1.2 (0.86-1.7) | 0.29 |
| Malignant Cells | 1 (0.73-1.5) | 0.86 |
| Epithelial Cells | 0.97 (0.7-1.4) | 0.88 |
| Mono_CD14_FOS- | 1.1 (0.78-1.5) | 0.61 |
| TCD4_ex | 0.87 (0.61-1.2) | 0.43 |
| Fibroblasts | 1.2 (0.9-1.7) | 0.18 |
| Endothelial | 1.1 (0.79-1.6) | 0.52 |
| MonoInter_CXCL10 | 0.71 (0.5-1) | 0.055 |
| NK_rest | 0.84 (0.6-1.2) | 0.28 |
| NKT | 0.86 (0.62-1.2) | 0.35 |
| **RTM_IFN** | **0.63 (0.44-0.91)** | **0.013** |
| Age | 0.99 (0.97-1) | 0.68 |

| | | |
|---|---|---|
| TCD8_naive | 1.2 (0.85-1.7) | 0.29 |
| NK_cyto | 1.5 (1-2.2) | 0.052 |
| **RTM_IFN** | **1.7 (1.2-2.5)** | **0.0056** |
| **TCD8_ex** | **1.7 (1.1-2.5)** | **0.011** |
| **NKT** | **1.8 (1.2-2.7)** | **0.0045** |
| B_cells | 0.83 (0.55-1.2) | 0.37 |
| TCD4_em | 1 (0.72-1.4) | 0.97 |
| TCD4_ex | 1.1 (0.74-1.5) | 0.74 |
| TCD8_em | 1.4 (0.98-2.1) | 0.065 |
| pDC | 1 (0.73-1.4) | 0.86 |
| Mac_Rec | 1.1 (0.75-1.5) | 0.21 |
| **Age** | **1 (1-1.1)** | **0.0008332** |
| gender | 1.7 (0.72-4.1) | 0.22 |
| Stage | 4.3 (0.51-3.1) | 13 |
