## Supplementary Table 10 for "Identification of novel myeloid-derived cell states with implication in cancer outcome"

**Supplementary Table 10: Multivariate Cox Regression analysis results**

**TCGA-OV**

| | p. value | HR | HR low | HR high |
|---|---|---|---|---|
| RTM_IM | 0.0049 | 1.2 | 1.1 | 1.4 |
| TCD4_ex | 0.9 | 0.98 | 0.78 | 1.2 |
| TCD8_ex | 0.58 | 0.94 | 0.74 | 1.2 |
| MonoInter_CXCL10 | 0.16 | 0.86 | 0.7 | 1.1 |
| Age | 0.0014 | 1 | 1 | 1 |

**TCGA-BRCA: Basal**

| | p. value | HR | HR low | HR high |
|---|---|---|---|---|
| RTM_IM | 0.18 | 1.3 | 0.87 | 2 |
| Mono_FCG3RA | 0.0081 | 1.9 | 1.2 | 3 |
| NKT | 0.27 | 0.74 | 0.44 | 1.3 |
| cDC2_FCGR3A | 0.28 | 0.75 | 0.45 | 1.3 |
| Stage2 | 1 | 1.4e+08 | 0 | Inf |
| Stage3 | 1 | 5.5e+08 | 0 | Inf |
| Stage4 | 1 | 1.9e+09 | 0 | Inf |

**TCGA-BRCA: HER2**

| | p. value | HR | HR low | HR high |
|---|---|---|---|---|
| breast_tumor | 0.94 | 1 | 0.53 | 2 |
| Age | 0.0098 | 1.1 | 1 | 1.2 |
| Stage2 | 0.44 | 0.4 | 0.039 | 4.1 |
| Stage3 | 0.79 | 1.4 | 0.14 | 14 |
| Stage4 | 0.013 | 21 | 1.9 | 230 |

**TCGA-BRCA: Luminal A**

| | p. value | HR | HR low | HR high |
|---|---|---|---|---|
| TCD4_ex | 0.14 | 0.74 | 0.5 | 1.1 |
| cDC1_CLEC9A | 0.62 | 0.88 | 0.54 | 1.4 |
| Mono_FCG3RA | 0.029 | 0.66 | 0.45 | 0.96 |
| cDC2_CD14 | 0.46 | 0.85 | 0.55 | 1.3 |
| pDC | 0.65 | 0.92 | 0.64 | 1.3 |
| cDC2_FCGR3A | 0.89 | 1 | 0.63 | 1.7 |
| Age | 1.3e-06 | 1.1 | 1 | 1.1 |
| Stage2 | 0.081 | 2.7 | 0.89 | 8.3 |
| Stage3 | 0.0079 | 5.1 | 1.5 | 17 |
| Stage4 | 3e-04 | 19 | 3.9 | 95 |

**TCGA-BRCA: Luminal B**

| | p. value | HR | HR low | HR high |
|---|---|---|---|---|
| TCD4_em | 0.06 | 0.64 | 0.41 | 1 |
| NK_rest | 0.15 | 0.74 | 0.49 | 1.1 |
| Mac_Angio | 0.052 | 1.5 | 1 | 2.3 |
| Fibroblasts | 0.22 | 1.3 | 0.84 | 2.1 |
| Stage2 | 0.36 | 2.7 | 0.33 | 22 |
| Stage3 | 0.14 | 4.9 | 0.6 | 40 |
| Stage4 | 0.0081 | 24 | 2.3 | 250 |

**TCGA-LIHC**

| | p. value | HR | HR low | HR high |
|---|---|---|---|---|
| cDC2_CXCL8 | 0.018 | 0.8 | 0.66 | 0.96 |
| Mac_Reg | 0.38 | 0.92 | 0.76 | 1.1 |
| Mac_IFN | 0.0062 | 1.3 | 1.1 | 1.6 |
| RTM_LA | 0.086 | 1.2 | 0.98 | 1.4 |
| Stage2 | 0.26 | 1.3 | 0.8 | 2.2 |
| Stage3 | 3e-04 | 2.3 | 1.5 | 3.6 |
| Stage4 | 0.0033 | 6.3 | 1.8 | 22 |

**TCGA-READ**

| | p. value | HR | HR low | HR high |
|---|---|---|---|---|
| TCD4_naive | 0.027 | 0.64 | 0.42 | 0.95 |
| Age | 0.002 | 1.1 | 1 | 1.1 |
| Stage2 | 1 | 1 | 0.18 | 5.5 |
| Stage3 | 0.41 | 1.9 | 0.4 | 9.6 |
| Stage4 | 0.011 | 7.7 | 1.6 | 37 |

**TCGA-SKCM : Primary**

| | p. value | HR | HR low | HR high |
|---|---|---|---|---|
| TCD8_em | 0.25 | 0.78 | 0.51 | 1.2 |
| RTM_IFN | 0.17 | 0.73 | 0.47 | 1.1 |

**TCGA-UVM**

| | p. value | HR | HR low | HR high |
|---|---|---|---|---|
| Epithelial Cells | 0.2 | 0.53 | 0.2 | 1.4 |
| Malignant Cells | 0.25 | 1.7 | 0.7 | 4.1 |
| RTM_IFN | 0.41 | 0.71 | 0.31 | 1.6 |
| TCD8_ex | 0.55 | 1.3 | 0.52 | 3.4 |
| NKT | 0.38 | 1.4 | 0.69 | 2.7 |
| Age | 0.08 | 1 | 1 | 1.1 |
| Stage3 | 0.74 | 1.2 | 0.44 | 3.2 |
| Stage4 | 0.0023 | 41 | 3.8 | 450 |

**TCGA-SKCM : Metastasis**

| | p. value | HR | HR low | HR high |
|---|---|---|---|---|
| TCD4_reg | 0.023 | 0.77 | 0.62 | 0.96 |
| TCD8_ex | 0.15 | 0.83 | 0.65 | 1.1 |
| TCD4_naive | 0.13 | 1.2 | 0.96 | 1.4 |
| B_cells | 0.017 | 0.78 | 0.64 | 0.96 |
| NK_cyto | 0.37 | 1.1 | 0.9 | 1.3 |
| TCD8_em | 0.7 | 1.1 | 0.82 | 1.3 |
| TCD8_naive | 0.74 | 0.97 | 0.8 | 1.2 |
| pDC | 0.053 | 0.82 | 0.67 | 1 |
| TCD4_em | 0.44 | 0.94 | 0.8 | 1.1 |
| Malignant Cells | 0.14 | 0.85 | 0.7 | 1.1 |
| Epithelial Cells | 0.032 | 0.85 | 0.73 | 0.99 |
| TCD4_ex | 0.11 | 1.1 | 0.97 | 1.4 |
| Endothelial | 0.24 | 0.89 | 0.73 | 1.1 |
| MonoInter_CXCI | 0.37 | 0.93 | 0.8 | 1.1 |
| NK_rest | 0.25 | 0.91 | 0.77 | 1.1 |
| NKT | 0.4 | 0.93 | 0.78 | 1.1 |
| RTM_IFN | 0.28 | 1.1 | 0.9 | 1.4 |
| Age | 0.019 | 1 | 1 | 1 |


**TCGA-BRCA: Luminal B**

| | p. value | HR | HR low | HR high |
|---|---|---|---|---|
| TCD4_em | 0.06 | 0.64 | 0.41 | 1 |
| NK_rest | 0.15 | 0.74 | 0.49 | 1.1 |
| Mac_Angio | 0.052 | 1.5 | 1 | 2.3 |
| Fibroblasts | 0.22 | 1.3 | 0.84 | 2.1 |
| Stage2 | 0.36 | 2.7 | 0.33 | 22 |
| Stage3 | 0.14 | 4.9 | 0.6 | 40 |
| Stage4 | 0.0081 | 24 | 2.3 | 250 |


**TCGA-LUAD**

| | p. value | HR | HR low | HR high |
|---|---|---|---|---|
| lung_tumor | 0.92 | 1 | 0.84 | 1.2 |
| B_cells | 0.12 | 0.88 | 0.74 | 1 |
| Mast_cells | 0.033 | 0.83 | 0.7 | 0.98 |
| Mac_Angio | 0.098 | 1.2 | 0.97 | 1.4 |
| RTM_IM | 0.62 | 0.96 | 0.82 | 1.1 |
| Mono_IL1B | 0.86 | 1 | 0.82 | 1.3 |
| cDC1_CLEC9A | 0.97 | 1 | 0.78 | 1.3 |
| cDC2_CD207 | 0.069 | 0.79 | 0.61 | 1 |
| cDC2B_FCER1A | 0.54 | 1.1 | 0.84 | 1.4 |
| cDC2_CXCL8 | 0.045 | 1.3 | 1 | 1.6 |
| Neutrophil_MMP9 | 0.4 | 1.1 | 0.89 | 1.4 |
| Neutrophil_CXCL8 | 0.27 | 0.89 | 0.72 | 1.1 |
| Stage2 | 7.5e-05 | 2.2 | 1.5 | 3.2 |
| Stage3 | 3.1e-08 | 3.1 | 2.1 | 4.6 |
| Stage4 | 3.9e-05 | 3.4 | 1.9 | 6.2 |

**TCGA-LUSC**

| | p. value | HR | HR low | HR high |
|---|---|---|---|---|
| Mac_Angio | 0.0064 | 1.2 | 1.1 | 1.4 |
| NK_cyto | 0.0021 | 0.81 | 0.71 | 0.93 |
| Mono_IL1B | 0.1 | 1.1 | 0.98 | 1.3 |
| Stage2 | 0.13 | 1.3 | 0.93 | 1.9 |
| Stage3 | 0.0061 | 1.7 | 1.2 | 2.5 |
| Stage4 | 0.042 | 2.6 | 1 | 6.5 |

* only $p < 0.05$ were considered significant.
For TCGA-COAD no significant findings were found.
