## Supplementary Table 11 for "Identification of novel myeloid-derived cell states with implication in cancer outcome"

**Supplementary Table 11: Description of INCA cohorts TNBC and HGSOC**

| | INCA - TNBC | | | | | |
|---|---|---|---|---|---|---|
| **Characteristic** | **High TREM2**, N = 58[1] | **Low TREM2**, N = 52[1] | **p-value**[2] | **High FOLR2**, N = 29[1] | **Low FOLR2**, N = 83[1] | **p-value**[2] |
| **Staging** | | | 0.076 | | | 0.018 |
| I | 14 (24%) | 22 (42%) | | 4 (14%) | 34 (41%) | |
| II | 23 (40%) | 12 (23%) | | 10 (34%) | 25 (30%) | |
| III | 21 (36%) | 18 (35%) | | 15 (52%) | 24 (29%) | |
| **Neoadjuvant_QT** | | | | | | |
| Yes | 58 (100%) | 52 (100%) | | 29 (100%) | 83 (100%) | |
| **Status Vital** | | | 0.009 | | | 0.026 |
| Alive | 31 (53%) | 15 (29%) | | 17 (59%) | 29 (35%) | |
| Dead | 27 (47%) | 37 (71%) | | 12 (41%) | 54 (65%) | |
| **Recurrence** | | | 0.059 | | | 0.024 |
| 0 | 23 (40%) | 30 (58%) | | 9 (31%) | 46 (55%) | |
| 1 | 35 (60%) | 22 (42%) | | 20 (69%) | 37 (45%) | |
| **OS (years)** | 3.17 (1.52) | 3.57 (1.57) | 0.3 | 2.83 (1.62) | 3.57 (1.48) | 0.034 |
| **RFS (years)** | 31 (21) | 39 (21) | 0.2 | 30 (21) | 37 (21) | 0.2 |
| **FOLR2_group** | | | 0.004 | | | |
| High FOLR2 | 22 (38%) | 7 (13%) | | | | |
| Low FOLR2 | 36 (62%) | 45 (87%) | | | | |
| **CD8** | | | 0.3 | | | 0.7 |
| High CD8 | 11 (34%) | 17 (47%) | | 9 (43%) | 19 (39%) | |
| Low CD8 | 21 (66%) | 19 (53%) | | 12 (57%) | 30 (61%) | |
| Unknown | 26 | 16 | | 8 | 34 | |
| **PDL1_CPS** | | | 0.077 | | | 0.4 |
| High PDL1 | 19 (48%) | 11 (28%) | | 6 (29%) | 24 (40%) | |
| Low PDL1 | 21 (53%) | 28 (72%) | | 15 (71%) | 36 (60%) | |
| Unknown | 18 | 13 | | 8 | 23 | |
| **PD1** | | | 0.2 | | | 0.13 |
| High PD1 | 4 (13%) | 1 (2.7%) | | 3 (15%) | 2 (3.9%) | |
| Low PD1 | 28 (88%) | 36 (97%) | | 17 (85%) | 49 (96%) | |

| | | | | | |
|---|---|---|---|---|---|
| Low PD1 | 28 (88%) | 36 (97%) | 17 (85%) | 49 (96%) | |
| Unknown | 26 | 15 | 9 | 32 | |
| **TREM2_group** | | | | | 0.004 |
| High TREM2 | | | 22 (76%) | 36 (44%) | |
| Low TREM2 | | | 7 (24%) | 45 (56%) | |
| Unknown | | | 0 | 2 | |

[1] n (%); Mean (SD)

[2] Pearson’s Chi-squared test; Wilcoxon rank sum test; Fisher’s exact test

**Supplementary Table 11: Brazilian cohort - INCA - HGSOC**

| | INCA - HGSOC | | | | | |
|---|---|---|---|---|---|---|
| **Characteristic** | **High TREM2**, N = 32[1] | **Low TREM2**, N = 116[1] | **p-value**[2] | **High FOLR2**, N = 20[1] | **Low FOLR2**, N = 91[1] | **p-value**[2] |
| **Diagnosis_age** | 58 (36 - 84) | 58 (23 - 81) | >0.9 | 60 (36 - 76) | 57 (29 - 81) | 0.2 |
| **Staging** | | | 0.6 | | | 0.8 |
| I | 3 (9.4%) | 5 (4.3%) | | 2 (10%) | 4 (4.4%) | |
| II | 4 (13%) | 10 (8.6%) | | 1 (5.0%) | 8 (8.8%) | |
| III | 19 (59%) | 77 (66%) | | 13 (65%) | 61 (67%) | |
| IV | 6 (19%) | 24 (21%) | | 4 (20%) | 18 (20%) | |
| **Neoadjuvant_QT** | | | >0.9 | | | 0.3 |
| N | 19 (59%) | 66 (59%) | | 9 (47%) | 52 (60%) | |
| S | 13 (41%) | 45 (41%) | | 10 (53%) | 35 (40%) | |
| Unknown | 0 | 5 | | 1 | 4 | |
| **Adjuvant_QT** | | | 0.4 | | | >0.9 |
| N | 2 (6.7%) | 15 (14%) | | 2 (11%) | 10 (13%) | |
| S | 28 (93%) | 89 (86%) | | 16 (89%) | 69 (87%) | |
| Unknown | 2 | 12 | | 2 | 12 | |
| **Recurrence** | | | >0.9 | | | >0.9 |
| N | 17 (61%) | 58 (61%) | | 11 (61%) | 46 (61%) | |
| S | 11 (39%) | 37 (39%) | | 7 (39%) | 30 (39%) | |
| Unknown | 4 | 21 | | 2 | 15 | |
| **Persistence** | | | 0.6 | | | 0.3 |
| N | 15 (54%) | 56 (60%) | | 8 (44%) | 44 (58%) | |
| S | 13 (46%) | 38 (40%) | | 10 (56%) | 32 (42%) | |
| Unknown | 4 | 22 | | 2 | 15 | |
| **Status Vital** | | | 0.2 | | | 0.7 |
| Alive | 2 (6.3%) | 18 (16%) | | 2 (10%) | 15 (17%) | |
| Dead | 30 (94%) | 94 (84%) | | 18 (90%) | 73 (83%) | |
| Unknown | 0 | 4 | | 0 | 3 | |
| **PFS (years)** | 1.49 (1.12) | 1.99 (1.68) | 0.5 | 1.10 (1.05) | 2.02 (1.63) | 0.007 |
| Unknown | 2 | 7 | | 1 | 5 | |
| **OS (years)** | 2.83 (1.40) | 3.04 (1.82) | 0.6 | 2.82 (1.66) | 3.03 (1.71) | 0.6 |

| | | | | | | |
|---|---|---|---|---|---|---|
| **OS (years)** | 2.83 (1.40) | 3.04 (1.82) | 0.6 | 2.82 (1.66) | 3.03 (1.71) | 0.6 |
| Unknown | 2 | 10 | | 1 | 8 | |
| **CD68_intratumoral** | | | <0.001 | | | 0.030 |
| High CD68 | 11 (50%) | 3 (2.9%) | | 4 (27%) | 4 (5.7%) | |
| Low CD68 | 11 (50%) | 102 (97%) | | 11 (73%) | 66 (94%) | |
| Unknown | 10 | 11 | | 5 | 21 | |
| **CD8_Intratumoral** | | | <0.001 | | | 0.6 |
| High CD8_Intratumoral | 26 (81%) | 34 (32%) | | 9 (50%) | 37 (44%) | |
| Low CD8_Intratumoral | 6 (19%) | 73 (68%) | | 9 (50%) | 47 (56%) | |
| Unknown | 0 | 9 | | 2 | 7 | |
| **PD1_Intratumoral** | | | 0.11 | | | 0.005 |
| High PD1_Intratumoral | 12 (38%) | 25 (23%) | | 10 (56%) | 19 (23%) | |
| Low PD1_Intratumoral | 20 (63%) | 82 (77%) | | 8 (44%) | 65 (77%) | |
| Unknown | 0 | 9 | | 2 | 7 | |
| **FOLR2_intratumoral** | | | 0.2 | | | |
| High FOLR2 | 6 (32%) | 12 (17%) | | | | |
| Low FOLR2 | 13 (68%) | 57 (83%) | | | | |
| Unknown | 13 | 47 | | | | |
| **KI67** | | | 0.5 | | | 0.003 |
| High KI67 | 10 (31%) | 39 (37%) | | 12 (71%) | 26 (32%) | |
| Low KI67 | 22 (69%) | 66 (63%) | | 5 (29%) | 55 (68%) | |
| Unknown | 0 | 11 | | 3 | 10 | |
| **TREM2_CD68_intratumoral** | | | | | | 0.2 |
| High TREM2 | | | | 6 (33%) | 13 (19%) | |
| Low TREM2 | | | | 12 (67%) | 57 (81%) | |
| Unknown | | | | 2 | 21 | |

[1] Mean (Range); n (%); Mean (SD)

[2] Wilcoxon rank sum test; Fisher's exact test; Pearson's Chi-squared test
